# Unifying multimodal single-cell data with a mixture-of-experts *β*-variational autoencoder framework

**DOI:** 10.1101/2025.02.28.640429

**Authors:** Andrew J. Ashford, Trevor Enright, Julia Somers, Olga Nikolova, Emek Demir

## Abstract

Multimodal single-cell assays profile complementary layers of cell state, but integration is complicated by modality mismatch, sparsity, and uneven cohort coverage. We present UniVI (**Uni**fied **V**ariational **I**nference), a scalable mixture-of-experts *β*-variational autoencoder that learns a shared latent space while preserving modality-specific structure. UniVI couples modality-specific encoders/de-coders with a shared latent prior and a symmetric cross-modal alignment objective, enabling consistent integration of paired measurements without curated feature-link graphs or pre-annotated reference atlases; optional supervised heads can be added when labels are available. Across paired RNA–protein (CITE-seq) and RNA–chromatin (10x Multiome, SHARE-seq) data spanning human PBMCs and mouse back skin—a non-hematopoietic tissue with continuous differentiation hierarchies—UniVI produces coherent embeddings, improves label transfer, and enables cross-modal reconstruction and denoising. Extending to tri-modal measurements, UniVI maintains robust three-way alignment among RNA, chromatin accessibility, and surface proteins (TEA-seq), and accommodates DNA methylation in a paired scNMT-seq mouse gastrulation proof-of-concept under beta-binomial likelihoods. Performance degrades gracefully under severe cell-type imbalance and in the presence of modality-exclusive populations. In an acute myeloid leukemia mosaic design, a paired RNA–protein bridge anchors independent RNA-only and protein+genotype cohorts, revealing genotype-associated neighborhoods that sharpen with mutation-aware fine-tuning. UniVI thus provides a flexible, interpretable framework for multimodal integration across paired, tri-modal, and mosaic study designs and supports practical reference-to-query projection in partially observed studies.

## Introduction

Multimodal single-cell assays now routinely measure complementary layers of cell state within the same experiment, enabling joint views of phenotype, transcription, and regulation (Baysoy et al. 2023). Cellular indexing of transcriptomes and epitopes by sequencing (CITE-seq) couples RNA with surface epitopes (Stoeckius et al. 2017); single-cell combinatorial indexing chromatin accessibility and mRNA (SHARE-seq) jointly profiles chromatin accessibility and gene expression (Ma et al. 2020); and assay for transposase-accessible chromatin with select antigen profiling by sequencing (ASAP-seq) integrates accessibility with protein readouts (Mimitou et al. 2021). More recently, *tri*-modal protocols such as transcriptome, epitopes, and chromatin accessibility by sequencing (TEA-seq) connect three layers in the same cells (Swanson et al. 2021). Together these assays resolve cell states and regulatory programs that are difficult to recover from any single modality alone (Mimitou et al. 2019).

Despite rapid experimental progress, multimodal data are still generated unevenly across tissues, conditions, and cohorts (Argelaguet et al. 2021; Baysoy et al. 2023; Heumos et al. 2023). Many studies pair a modest multimodal “anchor” subset with much larger unimodal collections, with substantial shifts in cell-type composition, sequencing depth, and technical noise (Heumos et al. 2023; Ashuach et al. 2023). Modalities themselves differ in fundamental ways: RNA measurements are overdispersed and zero-rich (Eraslan et al. 2019; Svensson 2020); chromatin accessibility (ATAC) is extremely sparse and often treated as near-binary across large feature spaces (Buenrostro et al. 2015; Satpathy et al. 2019); antibody-derived tags (ADTs) are lower-dimensional but phenotypically specific (Stoeckius et al. 2017); and DNA methylation and genomic alterations introduce additional nonlinearities and coverage variability (Angermueller et al. 2016; Chaligne et al. 2021). Integration in this setting is therefore not merely batch correction: when cross-modal evidence is weak, methods that enforce uniform correspondence can *over-align* modalities, obscuring modality-specific biology and producing spurious matches (Liu et al. 2025; Fu et al. 2025). These challenges intensify in large mosaic and disease-focused designs, where transcriptomic, protein, and genotype readouts may exist in separate cohorts that only partially overlap in modality and biology, and require methods that (i) learn from a limited paired “bridge”, (ii) support parameter-frozen projection of additional cohorts, and (iii) avoid or flag over-confident correspondence.

Computational strategies for multimodal integration differ primarily in how they define cross-modality correspondence and how strongly they enforce it. *Shared-projection* methods learn a common subspace via canonical correlation analysis (CCA) (Hotelling 1936; Hardoon et al. 2004) or anchor-based workflows such as Seurat (Butler et al. 2018; Stuart et al. 2019), with extensions like MaxFuse augmenting CCA-style co-embedding with within-modality graph smoothing for weakly linked settings (Chen et al. 2024). *Neighborhood-based* approaches integrate modality-specific graphs (Seurat WNN) (Hao et al. 2021), align mutual nearest neighbors across datasets (Haghverdi et al. 2018), or iteratively remove batch effects in a shared low-dimensional space (Harmony) (Korsunsky et al. 2019). *Factorization* models—including SVD, NMF, and MOFA-family extensions for mosaic designs—decompose measurements into shared and modality- or dataset-specific factors (Welch et al. 2019; Argelaguet et al. 2018; Argelaguet et al. 2020; C Gao et al. 2021). A unifying view for many of these methods is *manifold alignment* (MA): learning a shared embedding that preserves within-modality structure while bringing matched states across modalities into correspondence (Wang and Mahadevan 2009; Singh et al. 2020). *Optimal transport* (OT) provides a closely related formulation natural for perturbational settings with genuine population shift (Peyré and Cuturi 2019; Bunne et al. 2023); for same-population designs, MA can be interpreted as an approximately zero-shift special case that is more resistant to over-aligning modality-specific variance (K Cao et al. 2022). OT- and matching-based formulations have also been applied broadly to single-cell, spatial, and mosaic-integration tasks (Stark et al. 2020; Jain et al. 2021; Moriel et al. 2021; Demetci et al. 2022; Huizing et al. 2022; He et al. 2024).

Integration methods further differ in their use of *prior information*. *Feature-level priors* encode hypothesized cross-modality correspondences such as peak–gene guidance graphs to orient alignment (Pliner et al. 2018; Granja et al. 2021; Stuart et al. 2021; ZJ Cao and G Gao 2022), while *reference priors* project new datasets onto a pre-annotated multimodal atlas via anchor-based mapping (Stuart et al. 2019; Hao et al. 2023). Both can be unreliable in practice: feature-link graphs may be incomplete or context-dependent, and reference atlases degrade under composition shift, rare states, or disease-driven regulatory rewiring—concerns that intensify for atypical modality combinations such as RNA with methylation or genomic alterations. Deep generative models extend this landscape by coupling probabilistic encoders/decoders with modality-aware likelihoods (Stahlschmidt et al. 2022): scVI provides a principled framework for scRNA-seq, while to-talVI and MultiVI generalize to CITE-seq and RNA–ATAC, respectively (Lopez et al. 2018; Gayoso et al. 2021; Ashuach et al. 2023), and mixture-of-experts (MoE) formulations have been proposed for multimodal variational autoencoder (VAE) models (Shi et al. 2019; Minoura et al. 2021).

We developed UniVI to address these heterogeneous regimes while explicitly separating shared from modality-specific structure. UniVI is a MoE *β*-VAE (Jordan and Jacobs 1994; Kingma and Welling 2013; Higgins et al. 2017; Shazeer et al. 2017; Minoura et al. 2021) that learns a unified latent representation across modalities using modality-specific encoders and decoders coupled by a shared latent prior and a symmetric cross-modal regularization objective. UniVI is *prior-light*: it learns cross-modality correspondence directly from paired cells, avoiding reliance on curated feature-link graphs or pre-annotated reference atlases that may be unavailable outside canonical modality pairs. Beyond per-modality reconstruction error, UniVI applies an explicit *symmetric* divergence penalty between modality-specific posteriors for the same paired cell, directly coupling modalities at the level of (*µ, σ*) and reducing spurious alignment when one modality is noisier, sparser, or lower-dimensional. UniVI occupies a similar design space to deep generative models such as totalVI/MultiVI (Gayoso et al. 2021; Ashuach et al. 2023) and product-of-experts (PoE) VAEs (Shi et al. 2019; Minoura et al. 2021), but is specifically optimized for “mosaic” regimes with heterogeneous overlaps, extreme composition imbalances, or large unimodal subsets, and fuses modalities through MoE aggregation rather than feature concatenation (which can amplify modality scale differences) or graph-level fusion alone (e.g., WNN) (Hao et al. 2021). MoE aggregation reweights modalities across the manifold so that informative views dominate locally when others are noisy, sparse, or missing. A single configuration interface decouples architecture from training pipeline, enabling reproducible swaps of modality-specific likelihoods and encoder/decoder pairs; optional supervised heads can be attached as auxiliary signals but are not required for alignment.

Recent benchmarking studies emphasize that integration performance is highly contingent on the experimental regime—the degree of cell-state overlap, paired versus mosaic data, and disparities in modality-specific sparsity (Xiao et al. 2024; Liu et al. 2025; Fu et al. 2025; Zhou et al. 2025). These factors introduce ambiguous correspondences or over-confident alignments distributed heterogeneously along the manifold, yet aggregate performance metrics rarely surface where integration is locally reliable. UniVI accordingly includes a comprehensive diagnostic suite, pairing standard alignment and label-transfer metrics with neighborhood- and reconstruction-based scores that flag regions where modality-specific structure should be interpreted cautiously.

In this work, we evaluate UniVI across progressively more demanding multimodal study designs. We first establish performance on fully paired bimodal benchmarks: CITE-seq PBMCs (RNA–ADT) (Hao et al. 2021) and 10x Multiome PBMCs (RNA–ATAC) (10x Genomics 2021a), assessing single-cell correspondence, bidirectional label transfer, modality mixing, and cross-modal reconstruction. To test generalization beyond hematopoietic biology, we extend to paired SHARE-seq RNA–ATAC measurements of late-anagen mouse back skin (Ma et al. 2020), a non-hematopoietic tissue with continuous differentiation hierarchies and substantially lower per-cell ATAC complexity. As a proof-of-concept on a modality class with substantially different measurement statistics, we apply UniVI to paired scNMT-seq mouse gastrulation data (Argelaguet et al. 2019), jointly integrating RNA, CpG methylation, and GpC accessibility via beta-binomial likelihoods. We then evaluate *parameter-frozen* reference-to-query transfer: a UniVI model trained on a paired Multiome reference (10x Genomics 2021b) embeds independent RNA-only (Ding et al. 2020) and ATAC-only (Satpathy et al. 2019) PBMC cohorts by encoder inference alone, with optional lightweight supervised refinement applied only after projection. We extend to fully paired tri-modal alignment in TEA-seq (Swanson et al. 2021) under sample holdout. We then study a disease-focused mosaic acute myeloid leukemia (AML) setting, training a paired RNA–ADT bridge on AML CITE-seq (Knorr et al. 2023) and projecting RNA+genotype (Galen et al. 2019) and ADT+genotype (Demaree et al. 2021) cohorts to examine genotype-associated organization with optional mutation-aware refinement. Finally, we benchmark UniVI against widely used integration methods under a unified cross-validation runner, perform regime-focused robustness and ablation analyses (overlap sweeps, cell-type-specific modality dropout), and stress-test computational scaling across CUDA and Apple Metal (MPS) backends.

## Results

### Overview of UniVI and evaluation roadmap

UniVI is a multimodal MoE *β*-VAE that learns a shared latent manifold across heterogeneous single-cell modalities while preserving modality-specific structure (Fig. 1). The model uses modality-specific encoders and decoders with modality-appropriate likelihoods, coupled through a shared latent prior (Fig. 1A). For paired cells, UniVI yields modality-specific Gaussian posteriors (e.g., *Z*_RNA_, *Z*_ADT_, *Z*_ATAC_) and also provides a fused representation via MoE aggregation for visualization and neighborhood analyses (Fig. 1A).

**Figure 1.**
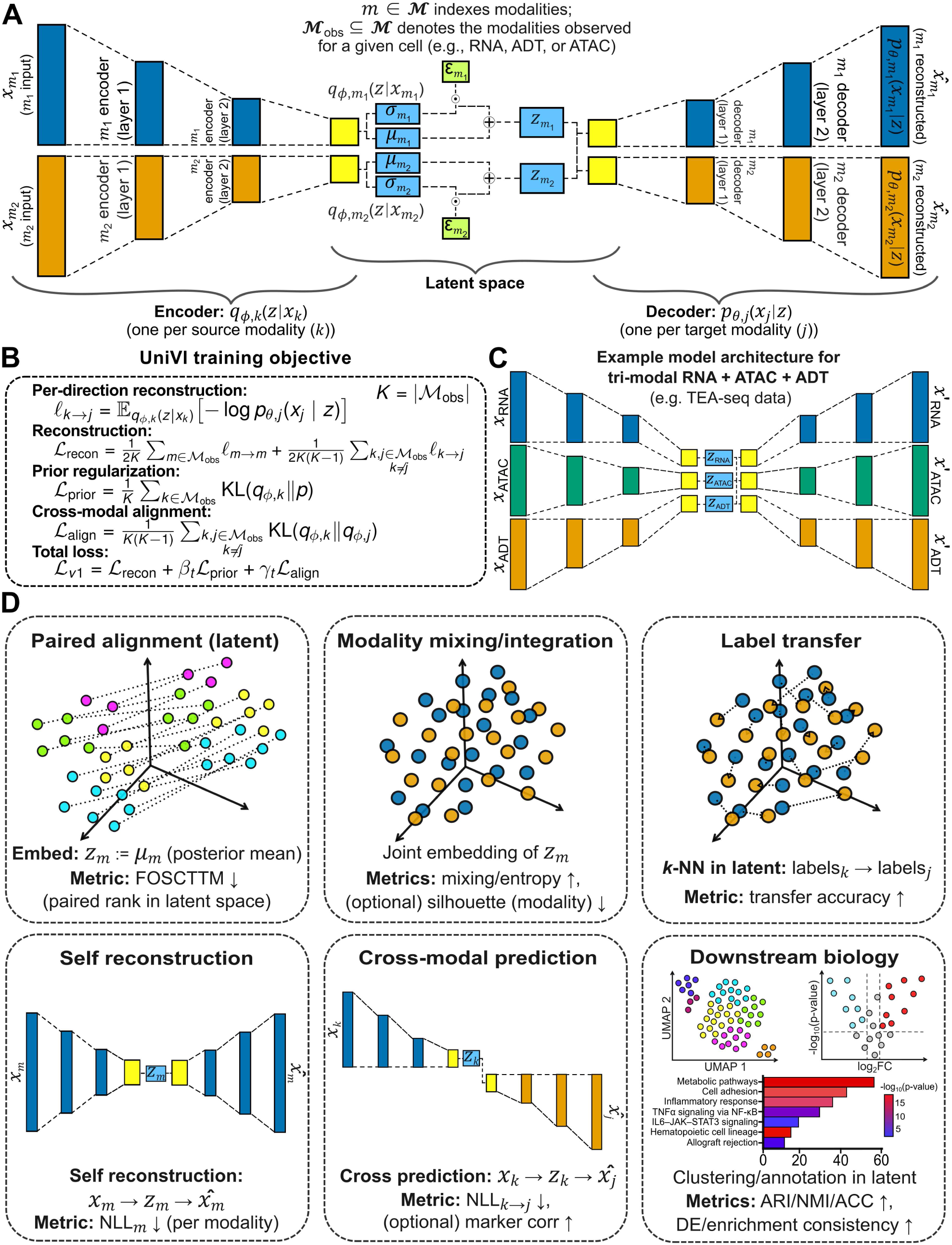
UniVI overview and evaluation roadmap. (A) UniVI (`loss_mode=”v1”`) with modality-specific encoders/decoders, a shared latent prior, and MoE aggregation. (B) v1 training objective: reconstruction, KL to the shared prior, and symmetric paired posterior alignment. (C) Evaluated study designs spanning paired, tri-modal, bridge, and mosaic regimes. Example tri-modal RNA-ATAC-ADT architecture shown to highlight UniVI’s built-in modularity. (D) Metrics and diagnostics used across regimes (paired correspondence, modality mixing, label transfer, cross-modal reconstruction, biological consistency).

In our analyses, UniVI is trained with a paired objective (`loss_mode=”v1”`) that combines modality-specific reconstruction and Kullback–Leibler (KL) regularization with an explicit *symmetric* alignment term encouraging agreement between modality-specific posteriors for the same jointly-measured cell (Fig. 1B). This posterior-level coupling supports single-cell correspondence without relying on curated feature-link graphs or reference atlases.

Following the order outlined above, the Results progress through four study-design regimes: paired bimodal assays (Fig. 1A), fully paired tri-modal integration (Fig. 1C), parameter-frozen reference-to-query bridging, and disease-focused mosaic integration. Across all regimes we report a common set of quantitative metrics and marker/neighborhood diagnostics—paired correspondence, modality mixing, bidirectional label transfer, cross-modal reconstruction, and biological consistency in the fused space (Fig. 1D)—to distinguish well-supported alignment from regions where modality-specific structure should be interpreted cautiously. Several main-text panels are complemented by expanded supplemental views (Supp. Figs. S1–S11; Supp. Tables S1–S8) providing additional marker validations, alternative visualizations, and sensitivity analyses.

### UniVI integrates paired CITE-seq RNA and protein profiles at scale

We first evaluated UniVI on the Hao et al. (2021) CITE-seq PBMC dataset, a large paired RNA–ADT benchmark in which transcription and surface epitopes provide complementary views of immune identity. On a held-out paired test set of *n* = 112,359 cells (Methods; Supp. Methods), a shared UMAP (McInnes et al. 2018) constructed from stacked RNA- and ADT-derived embeddings formed coherent immune lineages with minimal residual separation by modality (Fig. 2A,B). Single-cell correspondence was strong: FOSCTTM (Singh et al. 2020) was 0.0209 ± 0.00022 on a 20,000-cell subsample of paired test embeddings (mean ± SEM), indicating that true cross-modality partners typically rank among the nearest neighbors. In the fused latent space, a *k*-NN-based modality mixing score (*k* = 20; Methods) was 0.487 (close to the 0.5 expected for two well-mixed modalities), versus 0.216 on modality-specific embeddings—consistent with UniVI retaining modality-specific structure in *Z*_RNA_ and *Z*_ADT_ while yielding a well-aligned fused manifold. Using `celltype.l2` annotations and *k*-NN label transfer (*k* = 15; Methods; Supp. Methods), UniVI achieved high bidirectional performance: RNA→ADT accuracy 0.958 (macro-F1 0.733) and ADT→RNA accuracy 0.962 (macro-F1 0.763). Confusion matrices show that errors concentrate among closely related immune states rather than unrelated lineages (Fig. 2C,D), indicating that cross-modality neighborhoods preserve meaningful semantic structure across both abundant and rare populations.

**Figure 2.**
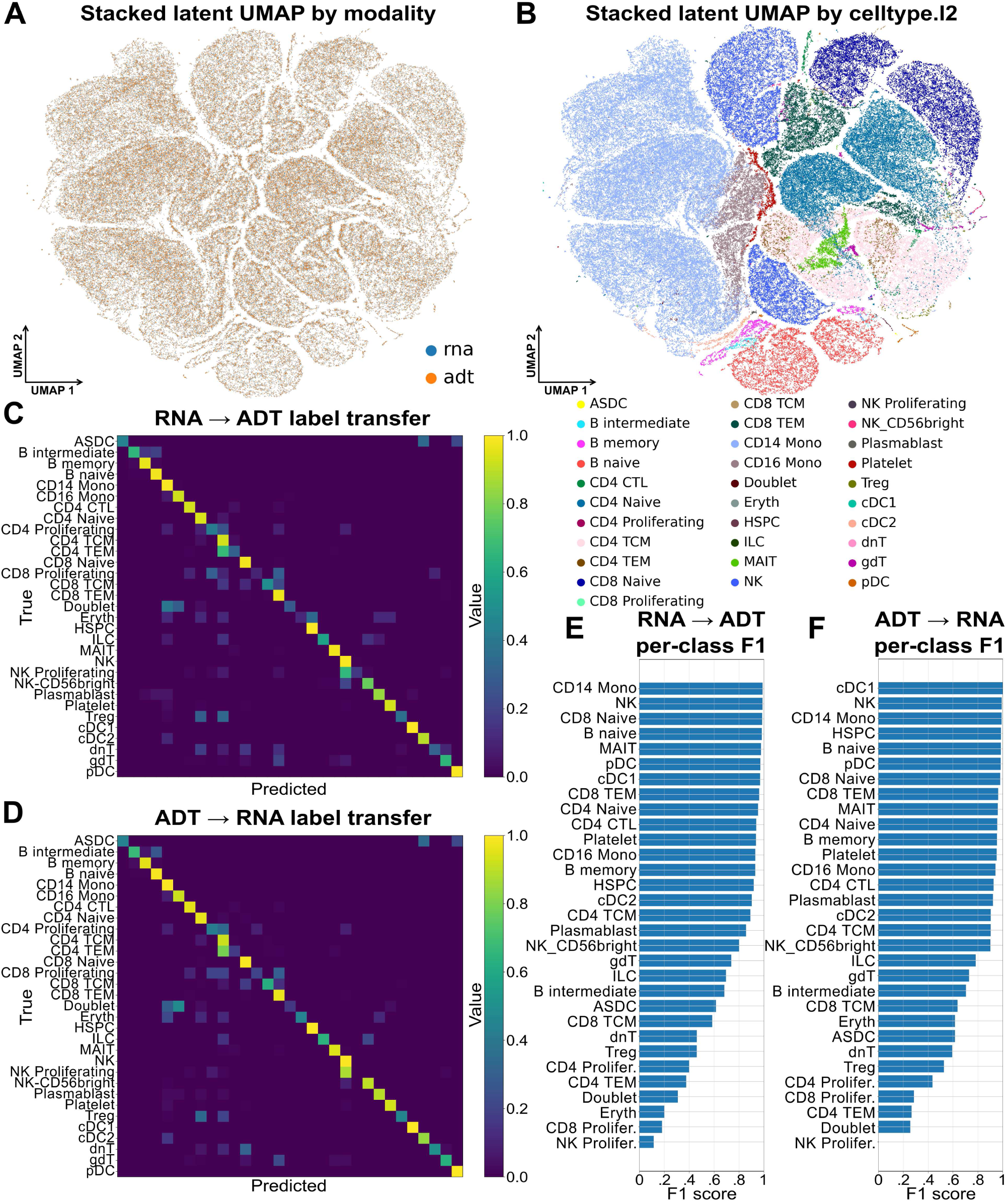
UniVI integrates paired CITE-seq RNA and protein profiles. (A) Stacked latent UMAP of held-out test embeddings colored by modality (RNA vs ADT). (B) Same UMAP colored by `celltype.l2`. (C,D) Row-normalized confusion matrices for RNA→ADT (C) and ADT→RNA (D) *k*-NN label transfer (*k* = 15). (E,F) Per-class F1 scores for RNA→ADT (E) and ADT→RNA (F). Summary (test set): RNA→ADT ACC 0.958 (macro-F1 0.733); ADT→RNA ACC 0.962 (macro-F1 0.763). FOSCTTM on a 20,000-cell subsample: 0.0209 ± 0.00022 (SEM).

### UniVI provides cross-modal denoising and reconstruction in CITE-seq

Beyond joint embedding, UniVI supports cross-modal reconstruction by encoding one modality and decoding another (Methods). In the held-out CITE-seq test set, cross-modal reconstructions preserved lineage-level marker structure across both directions (Fig. 3). A stacked latent UMAP colored by reference coarse-level cell types provides the shared coordinate frame for these comparisons and shows clear separation of major immune compartments (Fig. 3C). When conditioned on ADT-derived latents, reconstructed RNA profiles preserved canonical lineage markers while producing smoother within-lineage gradients, recapitulating B cell (*CD79A*), myeloid (*LYZ*), cytotoxic/NK (*NKG7*), and T cell (*TRAC*) compartments in the same latent coordinate system used for the observed data (Fig. 3D,F,H,J). Conversely, RNA→ADT reconstructions captured the expected compartment-level structure for matched markers, including CD19 in B cells, CD14 in monocytes, CD56-1 in cytotoxic/NK cells, and CD3-1 in T cells (Fig. 3E,G,I,K). Reconstructed features in both directions maintained strong between-lineage contrasts consistent with observed profiles (Fig. 3A,B), demonstrating that UniVI learns an aligned manifold supporting biologically faithful cross-modal prediction; concordance persists over an expanded marker set and finer immune subtypes (Supp. Fig. S1).

**Figure 3.**
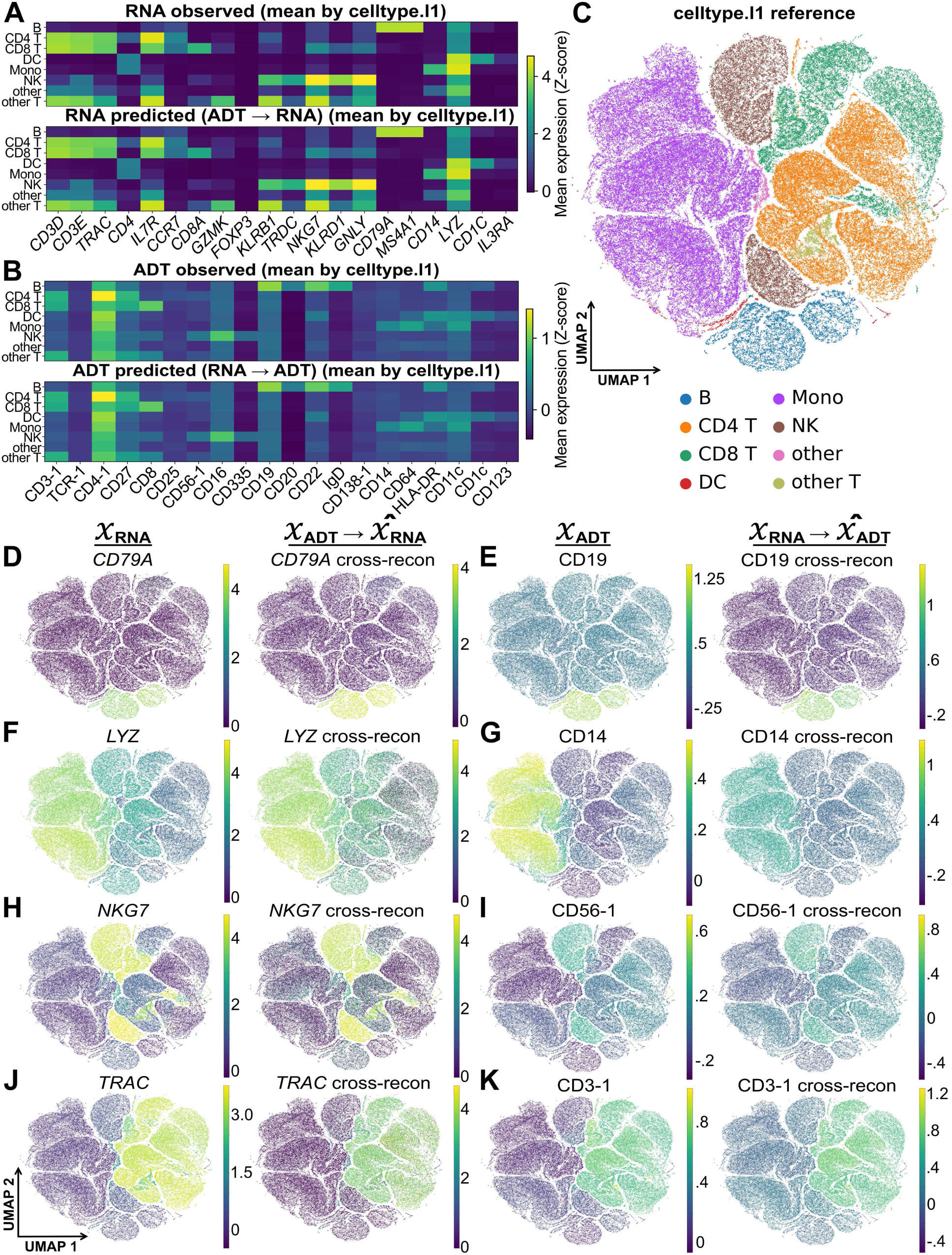
UniVI supports cross-modal reconstruction and denoising in CITE-seq. (A,B) Cell-type–aggregated marker means for RNA (A) and ADT (B), shown for observed and cross-modally reconstructed profiles (*Z*-scored per feature across cell types). (C) Stacked test-set latent UMAP colored by reference coarse-level cell types. (D–K) Observed (left) vs cross-modal reconstructions (right) for representative markers: B cells—*CD79A* (RNA, D), CD19 (ADT, E); mono/mac—*LYZ* (RNA, F), CD14 (ADT, G); cytotoxic/NK—*NKG7* (RNA, H), CD56-1 (ADT, I); T cells—*TRAC* (RNA, J), CD3-1 (ADT, K). Directions are ADT→RNA (D,F,H,J) and RNA→ADT (E,G,I,K).

### UniVI aligns paired RNA and ATAC profiles in 10x Multiome PBMCs

We next evaluated UniVI on paired 10x Multiome PBMC data (10x Genomics 2021a), a challenging regime because chromatin accessibility is extremely sparse and indirectly coupled to gene expression. UniVI was evaluated on a held-out paired test set of *n* = 3,137 cells spanning 19 immune populations (Methods; Supp. Methods; Fig. 4). In a shared latent UMAP constructed from stacked *Z*_RNA_ and *Z*_ATAC_, major PBMC lineages form coherent regions and paired cells co-localize at single-cell resolution, with broad modality interleaving rather than assay-specific separation (Fig. 4A,B). Residual modality structure is most visible within the naïve T cell compartment, where RNA- and ATAC-derived points occupy partially offset lobes, though paired points remain closely linked; FOSCTTM on the full test set is 0.0479 ± 0.00107 (SEM), indicating strong pairing fidelity despite this local distortion. 3D visualization of the same held-out latent coordinates (Supp. Methods; Supp. Fig. S2) confirms that the alignment is not an artifact of 2D UMAP projection and that local distortions are spatially limited.

**Figure 4.**
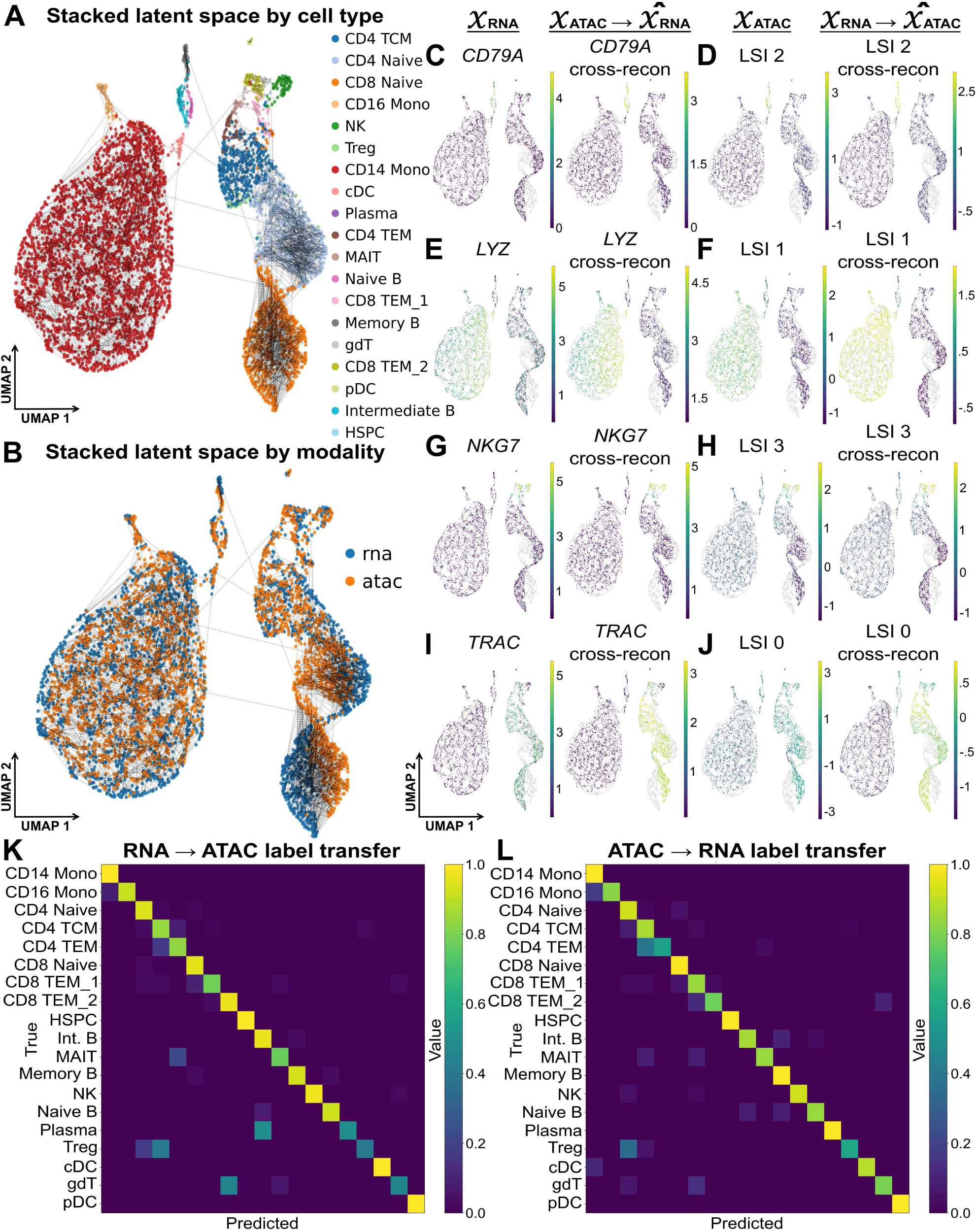
UniVI integrates paired RNA and ATAC profiles in 10x Multiome PBMCs. (A) Stacked latent UMAP of the held-out test set (*n* = 3,137) from modality-specific posterior means (*Z*_RNA_, *Z*_ATAC_); dashed segments connect paired embeddings; colored by `cell_type`. (B) Same UMAP colored by modality (RNA vs ATAC). (C–J) Observed (left) vs cross-modal reconstructions (right): ATAC→RNA overlays for lineage markers (*CD79A*, *LYZ*, *NKG7*, *TRAC*) and RNA→ATAC reconstructions shown in the ATAC LSI space (example components labeled per panel). (K,L) Row-normalized confusion matrices for *k*-NN label transfer (*k* = 3) RNA→ATAC and ATAC→RNA. Summary: FOSCTTM 0.0479 ± 0.00107 (SEM); RNA→ATAC ACC 0.960 (macro-F1 0.858); ATAC→RNA ACC 0.961 (macro-F1 0.892).

Under ATAC→RNA decoding, reconstructed RNA recapitulated canonical lineage markers in the expected regions of the latent space (*CD79A*, *LYZ*, *NKG7*, *TRAC* across B, myeloid, cytotoxic/NK, and T cell compartments; Fig. 4C,E,G,I). Conversely, RNA→ATAC decoding produced structured variation in the TF–IDF+LSI representation commonly used for scATAC-seq (Fig. 4D,F,H,J); because individual LSI dimensions are not directly interpretable at peak resolution, we interpret these reconstructions at the level of lineage- and program-scale geometry. A *k*-NN modality mixing score on modality-specific embeddings (*k* = 20) was 0.230, consistent with retention of modality-specific information despite global alignment. Using *k*-NN label transfer (*k* = 3 due to the presence of extreme minority classes in the test set; Methods), RNA→ATAC achieved accuracy 0.960 (macro-F1 0.858) and ATAC→RNA accuracy 0.961 (macro-F1 0.892) (Fig. 4K,L); errors concentrated among closely related T cell states rather than spanning unrelated lineages, establishing strong paired RNA–ATAC correspondence and biologically coherent cross-modal reconstruction in Multiome PBMCs.

### UniVI aligns paired SHARE-seq mouse skin data across developmentally diverse cell types

To assess generalization beyond hematopoietic biology, we evaluated UniVI on SHARE-seq paired RNA and ATAC measurements of late-anagen mouse back skin (Ma et al. 2020), spanning 22 cell types across epidermal, hair-follicle differentiation, dermal mesenchymal, neural-crest-derived, vascular, and immune lineages with continuous differentiation trajectories. SHARE-seq is also a more demanding paired RNA–ATAC regime than 10x Multiome, with substantially lower per-cell ATAC complexity (median 3,220 fragments per cell vs ≥10,000 typical for 10x Multiome; details in Supp. Methods). After QC, UniVI was evaluated on a held-out paired test set of *n* = 3,139 cells (Supp. Fig. S3).

Stacked *Z*_RNA_ and *Z*_ATAC_ embeddings formed coherent regions for each annotated cell type and broadly interleaved modalities across the manifold (Supp. Fig. S3A,B), with FOSCTTM 0.0546 ± 0.00199 (mean ± SEM)—comparable to Multiome PBMCs (0.0479 ± 0.00107) despite the increased heterogeneity and reduced ATAC depth—and bidirectional Recall@10 of 0.168 (RNA→ATAC) and 0.162 (ATAC→RNA). The *k*-NN modality mixing score on stacked embeddings (*k* = 30) was 0.326, with residual local separation concentrated within continuous differentiation trajectories rather than at lineage boundaries. Cross-modal reconstructions recovered lineage-scale structure across diverse skin cell types: ATAC→RNA decoding yielded mean per-feature Pearson *r* = 0.132 (MSE 0.994 on *Z*-scored expression; Supp. Fig. S3D), and RNA→ATAC decoding into the TF–IDF+LSI representation yielded *r* = 0.184 (MSE 0.196; Supp. Fig. S3C). These values are lower than for paired CITE-seq RNA–ADT, consistent with both the indirect coupling of chromatin accessibility to gene expression and SHARE-seq’s reduced per-cell ATAC depth; we therefore interpret cross-reconstructions in this regime at the level of lineage- and program-scale geometry (Discussion). *k*-NN label transfer (*k* = 15) for ATAC→RNA achieved accuracy 0.774 with macro-F1 0.666 across all 22 populations; the gap reflects degradation on rare populations (melanocytes *n* = 43, Schwann cells *n* = 40, sebaceous gland *n* = 47), where small anchor counts limit both cross-modal learning and *k*-NN retrieval. Errors concentrated among biologically related states along the hair-shaft differentiation trajectory (TAC-1, TAC-2, IRS, medulla, cuticle/cortex) rather than spanning unrelated lineages (Supp. Fig. S3B), demonstrating that UniVI’s paired RNA–ATAC alignment extends to a non-hematopoietic tissue with substantially greater lineage and developmental-origin diversity than the PBMC datasets, while making the limits of cross-modal reconstruction in low-complexity ATAC settings explicit.

### UniVI accommodates DNA methylation in tri-modal scNMT-seq mouse gastrulation

To evaluate whether UniVI extends to modality classes with substantially different measurement statistics and weak feature correspondence—specifically DNA methylation—we trained UniVI on the scNMT-seq mouse gastrulation dataset (Argelaguet et al. 2019), which jointly profiles RNA, CpG methylation, and GpC chromatin accessibility in the same cells across the E4.5–E7.5 window of mouse gastrulation. Methylation and accessibility modalities used beta-binomial likelihoods modeling per-feature (methylated, total) counts directly; RNA used a Gaussian likelihood on log-normalized expression (Methods; Supp. Methods, Supp. Table S8). Cells were assigned to an 85/5/10 train/validation/test split, and we report both an inductive evaluation on the held-out paired test set (*n* = 114; Supp. Fig. S4A–E) and a transductive evaluation on the full paired dataset (*n* = 1,140; Supp. Fig. S4F–J), reflecting the modest cohort size that limits sample-level holdout statistics.

On the inductive held-out set, the stacked latent UMAP from modality-specific posterior means showed broad interleaving of RNA, CpG, and GpC points across the manifold (Supp. Fig. S4A), with developmental stage recovered in canonical order (E4.5 → E5.5 → E6.5 → E7.5; Supp. Fig. S4B) and embryo-of-origin not dominating the geometry (Supp. Fig. S4C). Pijuan-Sala atlas-transferred `lineage10x` annotations (Pijuan-Sala et al. 2019) formed coherent regions consistent with gastrulating lineage compartments (Supp. Fig. S4D). MoE gating weights averaged within each `lineage10x` group remained balanced across the three modalities (per-lineage range ∼ 0.20–0.46; Supp. Fig. S4E), with mature mesoderm preferentially weighting CpG/GpC and primitive endoderm preferentially weighting RNA—indicating that the framework leverages methylation information rather than collapsing onto RNA. Inductive FOSCTTM was 0.192 ± 0.024 for RNA–CpG correspondence (Recall@1 = 0.070, Recall@10 = 0.526).

Applying the same checkpoint transductively to the full paired dataset resolved finer lineage structure, including ExE_ectoderm, ExE_mesoderm, Parietal/Pharyngeal/Intermediate mesoderm, Rostral neuroectoderm, and Surface_ectoderm (Supp. Fig. S4F–I), with sharper per-lineage MoE contrast than at inductive scale (e.g., Surface_ectoderm strongly weighting GpC; ExE_mesoderm strongly weighting CpG; Supp. Fig. S4J). On the full dataset, FOSCTTM dropped to 0.029 ± 0.003 (Recall@1 = 0.250, Recall@10 = 0.691), with mean MoE gating remaining balanced across modalities (CpG 0.31, GpC 0.40, RNA 0.29). Together, these results show that UniVI can integrate single-cell DNA methylation alongside RNA and chromatin accessibility under modality-appropriate likelihoods, recovering both global developmental ordering and lineage-specific modality weighting without enforcing artificial feature correspondence between assays with fundamentally different measurement statistics.

### A Multiome reference bridges independent unimodal RNA and ATAC cohorts

We next evaluated UniVI in a reference-to-query bridge design in which a paired Multiome dataset serves as a transferable reference for independent unimodal cohorts. A UniVI model trained on paired 10x Multiome PBMC cells (10x Genomics 2021b) embedded Ding et al. (2020) (RNA-only) and Satpathy et al. (2019) (ATAC-only) cells into the same latent geometry by encoder inference without updating trained generative parameters (Methods; Supp. Methods; Fig. 5). In a shared UMAP across Multiome, Ding, and Satpathy, projected unimodal cells co-localized with the Multiome reference primarily by coarse immune identity (Fig. 5A,B). Residual structure associated with technology or platform remained visible (Fig. 5C), consistent with realistic cohort shift, but was secondary to lineage-scale organization—indicating that a paired Multiome-trained model provides a usable geometric bridge across independent unimodal datasets.

**Figure 5.**
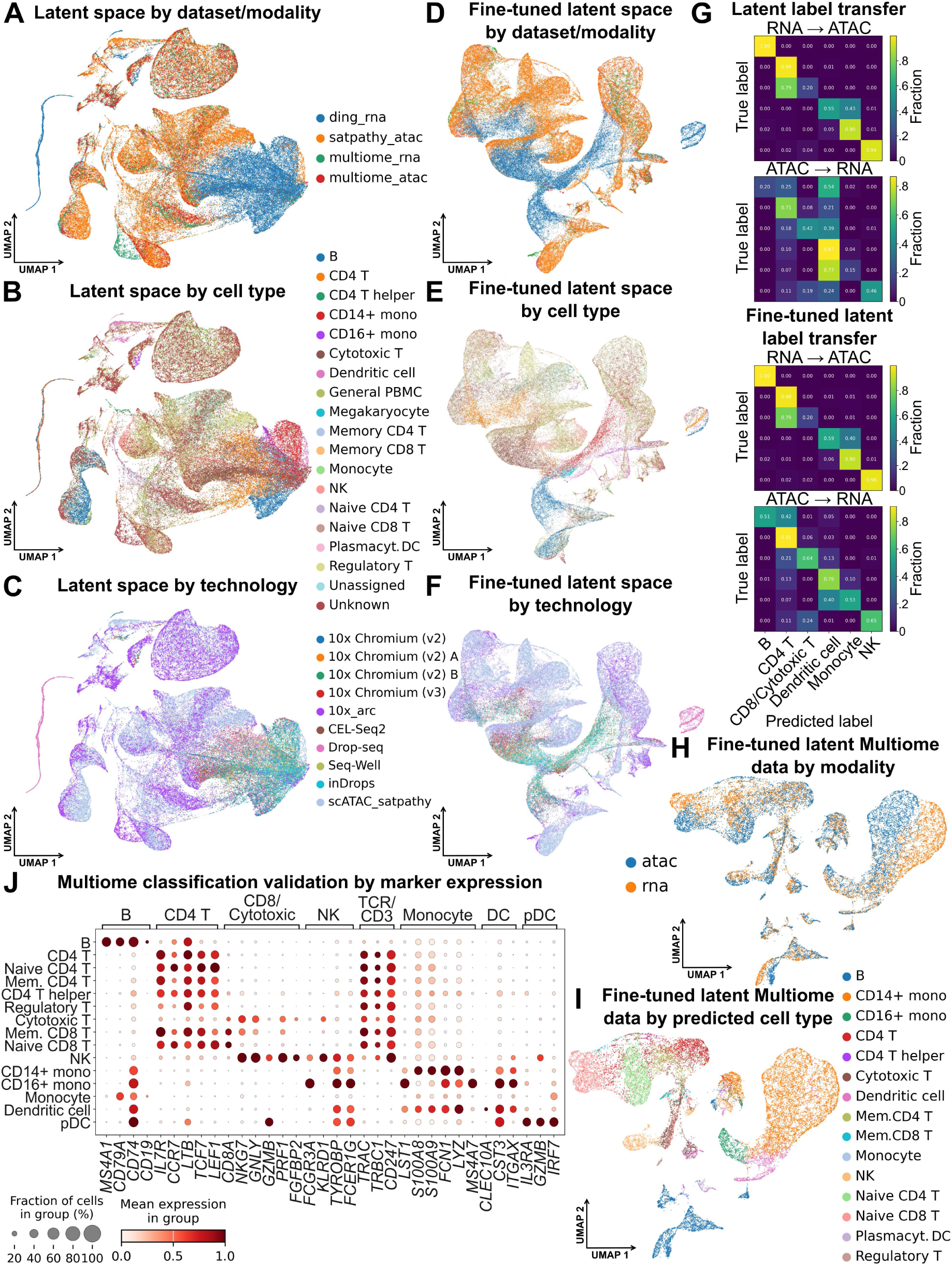
A Multiome reference bridges independent unimodal RNA-only and ATAC-only PBMC cohorts. (A–C) Joint latent UMAP of Multiome reference (RNA+ATAC) with projected Ding (RNA-only) and Satpathy (ATAC-only) cells, colored by (A) dataset/modality, (B) harmonized coarse PBMC label, and (C) technology/platform. (D–F) Same views after optional supervised refinement (classification head; decoders frozen), showing increased within-lineage cross-cohort co-localization. (G) Cross-cohort *k*-NN label transfer (*k* = 15) before (top) and after (bottom) refinement in both directions (Ding↔Satpathy). (H) Multiome bridge cells only, colored by modality. (I) Multiome bridge cells colored by predicted coarse label from the refinement head. (J) Marker-based validation in Multiome RNA (dot plot: fraction expressing and mean expression per predicted group).

Starting from the Multiome-trained checkpoint, optional supervised refinement after projection (Methods) using harmonized coarse PBMC labels preserved the global PBMC layout while increasing within-lineage cross-cohort co-localization and reducing technology-driven stratification (Fig. 5D–F). Cross-cohort consistency was quantified via bidirectional *k*-NN label transfer using a harmonized coarse PBMC vocabulary: in the parameter-frozen latent space, Ding→Satpathy reached accuracy 0.824 (macro-F1 0.768) while Satpathy→Ding was weaker (accuracy 0.408, macro-F1 0.410), consistent with asymmetric domain shift; after refinement, Ding→Satpathy remained high (0.825 / 0.774) and Satpathy→Ding improved to 0.665 / 0.623 (Fig. 5G). The Multiome bridge dataset itself lacked curated labels and was not used as a source of supervision; applying the refined classifier yielded coherent predicted coarse PBMC labels (Fig. 5H,I), and marker-based validation in Multiome RNA showed enrichment of canonical lineage markers in predicted groups (Fig. 5J; expanded marker view in Supp. Fig. S5). Together, these results show that a paired Multiome reference can support parameter-frozen projection of independent RNA-only and ATAC-only cohorts into a shared latent geometry, and that optional light supervision can improve cross-cohort semantic consistency without re-learning the underlying generative mapping.

### Tri-modal TEA-seq integration is robust and preserves concordant biology across all modalities

We evaluated UniVI on tri-modal TEA-seq PBMCs from Swanson et al. (2021) using a simple well holdout: models trained on wells 3–4 and 6 were evaluated on held-out well 5 (Supp. Methods). Because wells correspond to within-run capture/library partitions rather than independent biological replicates, we treat this as a modest robustness control; nonetheless, UniVI maintained strong three-way alignment on the held-out sample (Fig. 6). On well 5 cells, pairwise FOSCTTM was low and broadly comparable across modality pairs (Fig. 6A): RNA vs ADT 0.0747 ± 0.0011, RNA vs ATAC 0.0598 ± 0.0010, ADT vs ATAC 0.0593 ± 0.0009 (mean ± SEM), indicating that performance is not driven by a single dominant pairing. Neighborhood composition in the stacked latent space showed substantial cross-modality mixing (mean different-modality neighbor fraction 0.5605; local modality entropy mean 0.9332, median 0.9833; modality silhouette (Rousseeuw 1987) −0.0033), distances to same- vs different-modality neighbors overlapped strongly, and modality-colored UMAPs showed clear interleaving among RNA, ADT, and ATAC (Fig. 6B–D).

**Figure 6.**
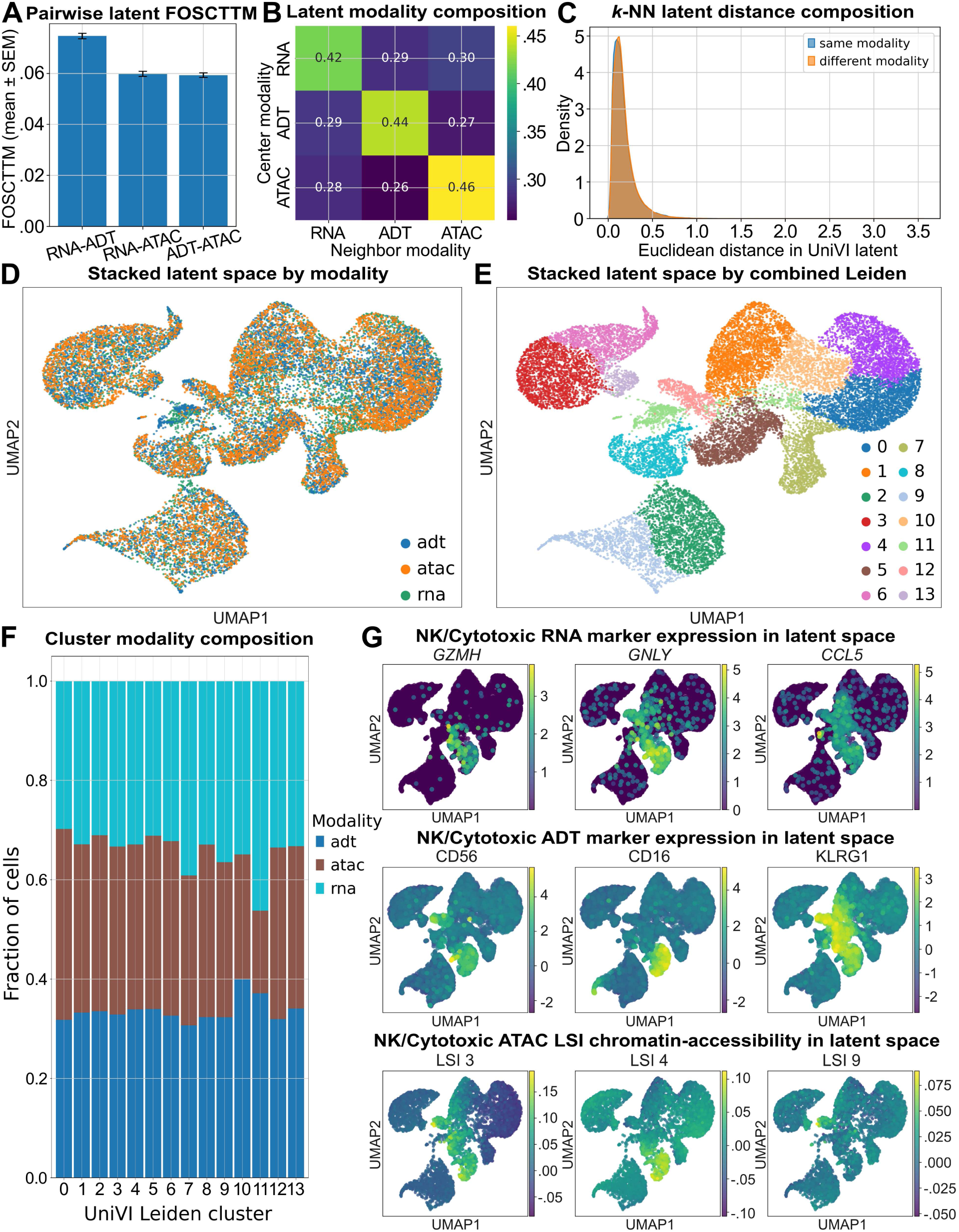
UniVI integrates tri-modal TEA-seq PBMCs and is stable under a held-out partition. Model trained on wells 3–4 and 6; evaluated on held-out well 5 (all panels). (A) Pairwise FOSCTTM (mean ± SEM) on the held-out well 5 sample for RNA–ADT, RNA–ATAC, and ADT–ATAC. (B) *k*-NN neighbor modality composition (*k* = 30) in the stacked latent space for well 5 cells. (C) Distances to same- vs different-modality neighbors (*k* = 30) in the stacked latent space. (D) Stacked latent UMAP colored by modality (RNA, ADT, ATAC). (E) Same UMAP colored by Leiden clusters on the stacked *k*-NN graph (14 clusters). (F) Modality composition per Leiden cluster. (G) Marker overlays illustrating cross-modality concordance in a cytotoxic lymphocyte region: RNA (*GZMH*, *GNLY*, *CCL5*), ADT (CD56, CD16, KLRG1), and representative ATAC LSI dimensions (LSI 3, LSI 4, LSI 9).

Leiden clustering (Traag et al. 2019) on the stacked latent representation yielded coherent groups with broadly balanced cluster-level modality composition (Fig. 6E,F), and modality-specific neighborhood graphs produced broadly concordant partitions in the same joint coordinates (Supp. Fig. S6A). Marker overlays revealed expected immune compartments with cross-modality agreement (Fig. 6G): a cytotoxic lymphocyte region exhibits elevated RNA expression of *GZMH*, *GNLY*, and *CCL5*, higher ADT signal for CD56, CD16, and KLRG1, and structured variation in representative ATAC LSI dimensions. Expanded lineage marker panels further support RNA–protein concordance for B cells, CD4 T cells, and myeloid/DC compartments (Supp. Fig. S6B–G), indicating that the learned tri-modal manifold is stable under a within-run partition holdout while retaining biologically interpretable structure shared across RNA, protein, and chromatin accessibility.

### Generalization beyond healthy PBMCs: AML mosaic integration bridges RNA, protein, and genotype

We next evaluated UniVI in a disease-focused mosaic AML setting where no single dataset contains all modalities. A model trained on paired patient-derived AML CITE-seq RNA+ADT cells (Knorr et al. 2023) was used to project unimodal AML scRNA-seq (RNA+genotype) (Galen et al. 2019) and DNA-barcoded antibody sequencing (DAb-seq) (protein+genotype) (Demaree et al. 2021) into a shared latent space by parameter-frozen encoder inference (Methods; Supp. Methods; Fig. 7). In the projected latent space, the paired CITE-seq bridge cohort defines the backbone geometry while van Galen and DAb-seq cells map into partially overlapping subregions; van Galen cell-state structure remains coherent after projection, and DAb-seq cells populate heterogeneous neighborhoods spanning progenitor-like and more differentiated regions (Fig. 7A,D)—an expected pattern in mosaic designs where cohorts differ in platform, feature construction, and sampled biology. Even without mutation supervision during bridge training, genotype signal was spatially structured in the shared manifold: DAb-seq *NPM1* status concentrated in specific regions, and neighborhood-based transfer propagated non-uniform *NPM1* probabilities across cohorts in both directions; sparse *NPM1* labels in the RNA-only cohort showed compatible localization patterns, though limited by missingness and occasional annotation conflicts (Fig. 7B,C,E,F).

**Figure 7.**
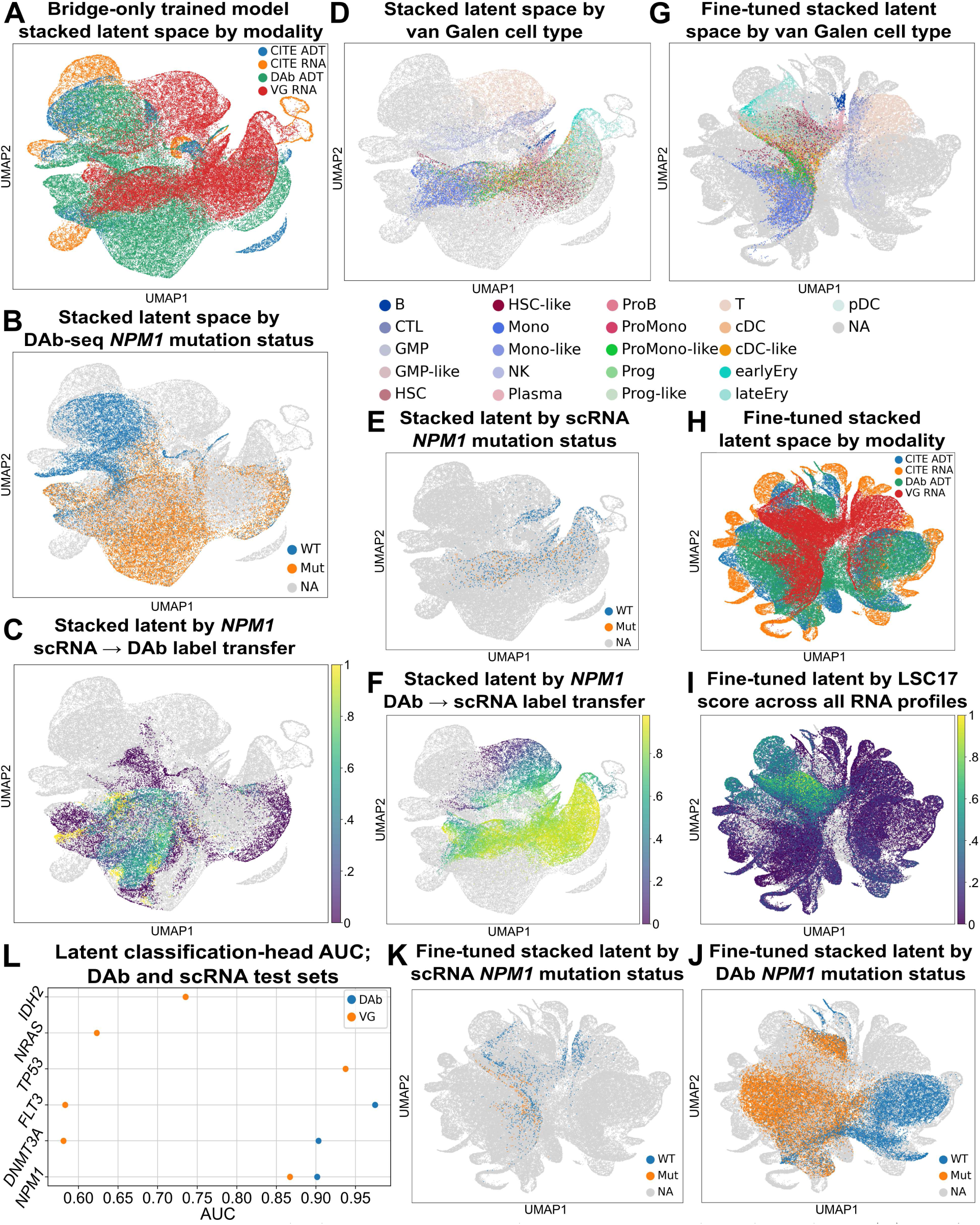
AML mosaic integration bridges RNA, protein, and genotype across independent cohorts. (A) Joint latent UMAP colored by dataset/modality (AML CITE RNA/ADT, van Galen RNA, DAb-seq ADT). (B) Joint UMAP highlighting DAb-seq *NPM1* status (WT/MUT; unlabeled gray). (C) *k*-NN probability transfer of *NPM1* from van Galen→DAb-seq (*k* = 60), shown on DAb-seq cells. (D) Joint UMAP colored by van Galen cell-state labels. (E) Joint UMAP colored by van Galen *NPM1* status for labeled cells. (F) *k*-NN probability transfer of *NPM1* from DAb-seq→van Galen (*k* = 50), shown on van Galen cells. (G) Refined UMAP colored by van Galen cell-state labels. (H) Joint UMAP after mutation-head refinement (encoders fine-tuned; decoders frozen), colored by dataset/modality. (I) Refined UMAP colored by LSC17 stemness score. (J) Refined UMAP colored by DAb-seq *NPM1* status. (K) Refined UMAP colored by van Galen *NPM1* status (labeled cells only). (L) Mutation-head test performance (AUC) across recurrent AML genes in DAb-seq and van Galen (genes with insufficient labeled test cells omitted/NA).

Optional post hoc refinement with mutation-prediction heads (encoders fine-tuned, decoders frozen; Methods; Supp. Methods) increased cross-cohort interleaving (Fig. 7H) and strengthened separation of DAb-seq *NPM1* status along a major manifold axis (Fig. 7J), and preserved the dominant van Galen state organization (Fig. 7G). Projecting the LSC17 stemness score (Ng et al. 2016) (Supp. Methods) onto the refined latent space revealed a broad differentiation gradient with higher scores concentrated in progenitor-like regions (Fig. 7I); the van Galen cohort contributes a prominent high-LSC17 cluster localizing to LSC-enriched and other immature progenitor-like compartments, consistent with its sampling and the expected AML hierarchy, while the DAb-seq cohort—spanning both peripheral blood and bone marrow—shows weaker enrichment for immature AML neighborhoods. These patterns indicate that the refined manifold captures a biologically interpretable stemness continuum coherent across cohorts and not explained by genotype structure alone.

Mutation-head performance tracks the availability of direct genotype supervision (Fig. 7L). In DAb-seq, where cell-level genotypes are observed for thousands of cells per target, heads achieve strong discrimination on densely labeled drivers (*NPM1* AUC = 0.90, *DNMT3A* AUC = 0.90, *FLT3* AUC = 0.97; *n* = 2,745–6,036 labeled test cells). The van Galen cohort contains substantially fewer labeled cells per gene (often tens to hundreds, with pervasive missingness; Supp. Table S4), yielding strong *NPM1* performance (AUC = 0.87, AP = 0.63; *n* = 295) but near-chance *DNMT3A* and *FLT3* (AUC ≈ 0.58). Targeted mutation calling from scRNA-seq has limited sensitivity, so wild-type calls in van Galen frequently reflect *no detected mutant transcript* rather than confidently genotyped WT, further constraining achievable performance; sporadic high-AUC targets (e.g., *TP53*, *n* = 12) rest on very small labeled test sets. Overall, auxiliary supervision is most informative where genotypes are directly observed at scale.

#### Supplemental AML mosaic context and expanded genotype landscapes

To interpret where paired anchors constrain the shared geometry, the bridge cohort’s clinical/molecular context (Supp. Table S1) and patient distribution across the fine-tuned manifold (Supp. Fig. S7A–C) indicate that bridge patients occupy multiple regions rather than a single localized cluster. Beyond *NPM1*, structured gene-specific probability landscapes for additional AML drivers are visible across projected cohorts (Supp. Fig. S7D–I). Modality composition and label availability differ sharply across datasets (Supp. Tables S1–S5): DAb-seq provides dense cell-level genotype labels supporting both evaluation and refinement, whereas van Galen provides sparse targeted readouts that are informative when present but incomplete—contextualizing why genotype grounding is strongest where direct labels are dense (DAb-seq), while still enabling structured genotype-associated patterns across the full mosaic embedding.

### Benchmarking against commonly used multimodal integration methods

We benchmarked UniVI against a broad set of widely used multimodal integration baselines spanning deep generative models, manifold alignment, graph/neighbor fusion, and factorization-based approaches (Fig. 8). All methods were run through a unified evaluation runner with consistent folds, preprocessing, and metric definitions; approaches whose standard workflows incorporate held-out cells during fitting were treated as transductive and explicitly flagged (Methods; Supp. Methods; Supp. Table S6). We distinguish **inductive** methods, whose parameters (or linear transforms) are learned on the training split and applied unchanged to validation/test cells, from **transductive** workflows, where the end-to-end procedure constructs a joint graph, factorization, or harmonized embedding over *all cells provided* and so the evaluated cells may influence the representation during fitting. This labeling matters because transductive workflows can appear advantaged on objectives computed on the same cells used during fitting, whereas inductive evaluations more directly probe *generalization*: can a model trained on one cohort produce stable embeddings for held-out cells by forward inference alone? In many applications (reference mapping, atlas extension, longitudinal deployment), the inductive setting is the relevant operating regime, so we report all methods under their standard workflows and flag transductive procedures.

**Figure 8.**
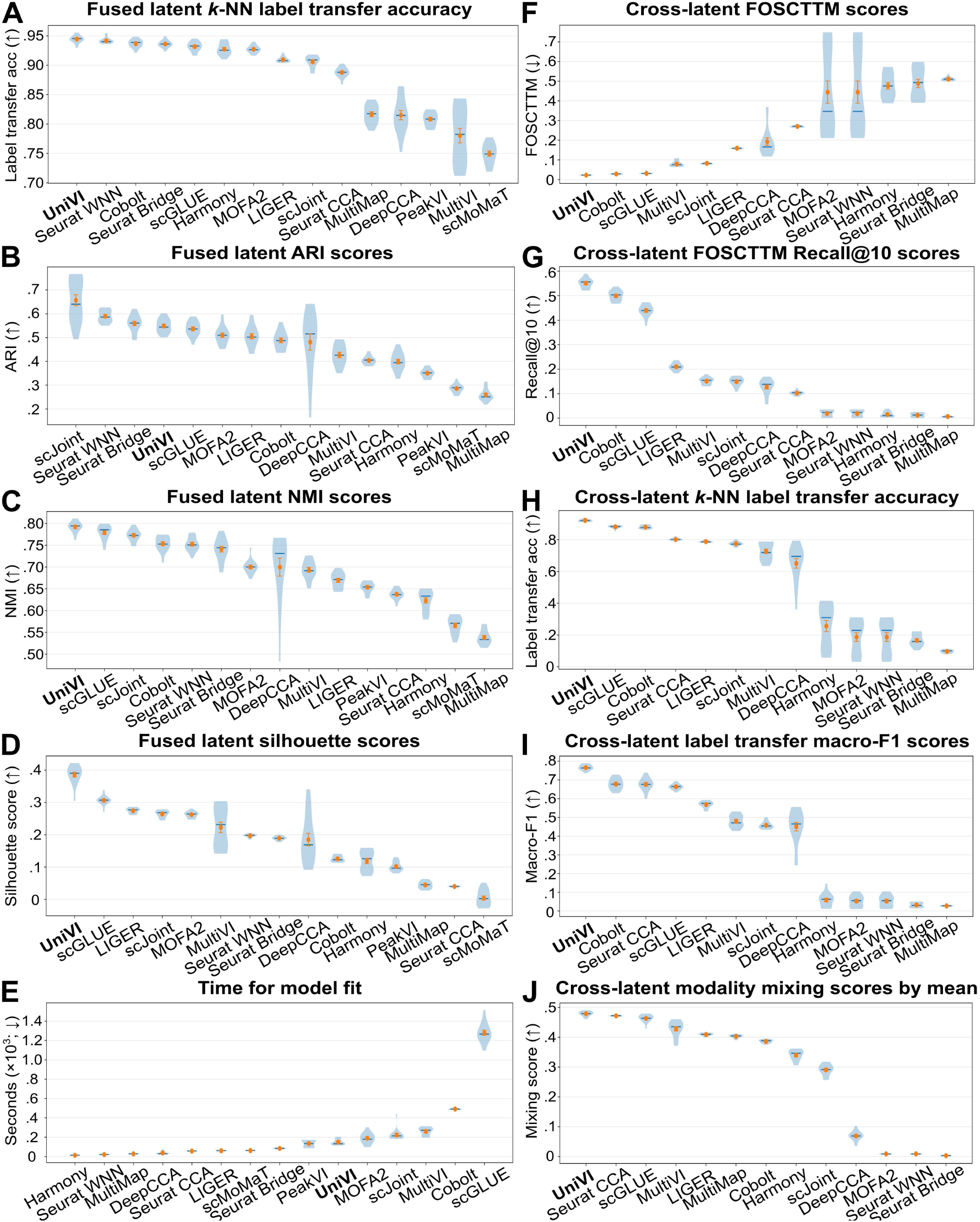
Benchmark comparison across multimodal integration methods using fused-space and cross-latent metrics. Violin plots summarize cross-validation folds and random seeds (orange markers: mean). (A) Fused-latent *k*-NN label transfer accuracy (*k* = 15). (B) Fused-latent *k*-means ARI (*k* set to the number of evaluated label classes). (C) Fused-latent *k*-means NMI (*k* set to the number of evaluated label classes). (D) Fused-latent silhouette score by ground-truth labels. (E) Wall-clock fit time (seconds). (F) Cross-latent FOSCTTM between modality-specific embeddings. (G) Cross-latent Recall@10. (H) Cross-latent *k*-NN label transfer accuracy (*k* = 15). (I) Cross-latent *k*-NN label-transfer macro-F1 (*k* = 15). (J) Modality mixing on stacked modality-specific embeddings (*k* = 30).

On metrics that reflect how well a *single* integrated embedding supports downstream analyses, UniVI is consistently among the strongest performers: it achieves the highest (or near-highest) *k*-NN label-transfer accuracy in the fused latent space (Fig. 8A) and leads on *k*-means ARI (Hubert and Arabie 1985) and NMI (Kvålseth 2017) (Fig. 8B,C). These gains do not come from over-mixing: UniVI also shows the best silhouette scores by ground-truth labels (Fig. 8D), indicating well-separated biological populations alongside cross-modality integration. Across baselines, we observe characteristic specialization: methods that prioritize cross-modal matching can yield strong correspondence on some metrics but sacrifice global organization or label separability (lower ARI/NMI/silhouette), while methods that preserve broad biological structure in one modality can under-align the other, leading to weaker cross-latent transfer and reduced modality interleaving (Fig. 8A–D,F–J).

Evaluated on modality-specific embeddings, UniVI also performs strongly on single-cell correspondence: low cross-latent FOSCTTM (Fig. 8F), the highest Recall@10 (Fig. 8G), and best-in-class cross-latent *k*-NN label-transfer accuracy and macro-F1 (Fig. 8H,I), supporting robust transfer across both common and rarer populations. UniVI further exhibits high modality mixing on stacked modality-specific embeddings (Fig. 8J), consistent with the qualitative co-localization patterns shown in earlier paired analyses (Figs. 2–4). Several baselines show a familiar failure mode—weak interleaving, or improved mixing accompanied by degraded label separation—which UniVI avoids. Wall-clock fit times vary substantially across methods under identical runner settings (Fig. 8E), reflecting differing optimization costs; the core pattern is that UniVI occupies a favorable regime where strong fused-space utility and cross-modality correspondence are achieved together rather than requiring a choice between the two.

### Sensitivity analyses reveal a broad stable operating region

To verify that the reported results reflect a robust operating point rather than a narrow optimum, we performed targeted sensitivity analyses in paired 10x Multiome PBMCs spanning objective weights, architectural regularization, and data scale. Sweeping the KL weight *β* and cross-modal coupling weight *γ* revealed a broad interior region in which paired correspondence (FOSCTTM, Recall@10), bidirectional label transfer macro-F1, and fused-space NMI/ARI were jointly strong (Supp. Fig. S8); metric-specific optima occurred at different (*β, γ*) settings, consistent with an expected trade-oH between strict paired retrieval, semantic stability, and global structure rather than a single tuned corner. The dominant failure pattern was confined to extreme regularization: very large *β* degraded correspondence and label transfer (underfitting), while removing coupling (*γ* = 0) weakened cross-modal retrieval. Sweeps over latent dimensionality and encoder/decoder dropout produced smooth, non-catastrophic trade-offs rather than brittle optima (Supp. Figs. S9–S10). Profiling training as a function of effective paired training size showed diminishing returns beyond moderate dataset sizes, with predictable wall-clock time and memory growth (Supp. Fig. S11). Together these analyses indicate that UniVI supports robust defaults for common multimodal study designs without extensive dataset-specific hyperparameter search.

### Robustness under reduced cross-modality overlap reveals an anchor threshold

Many multimodal studies deviate from fully paired designs: only a subset of cells may have both assays, while the remainder are effectively unimodal due to capture failures, assay-specific filtering, or cohort-specific missingness. These conditions reduce the number of *paired anchors* that directly constrain cross-modality geometry and can induce failure modes ranging from under-alignment to over-confident matching in weakly supported regions. To quantify this dependence, we evaluated UniVI under a controlled *overlap sweep* in paired 10x Multiome PBMCs, progressively reducing the fraction of cells retaining both RNA and ATAC while preserving leakage-free preprocessing and inductive evaluation within each cross-validation replicate (Supp. Methods; Fig. 9). We considered two missingness patterns—dropping ATAC (*Drop ATAC*) or dropping RNA (*Drop RNA*)—to test whether robustness depends on which modality is sparsified.

**Figure 9.**
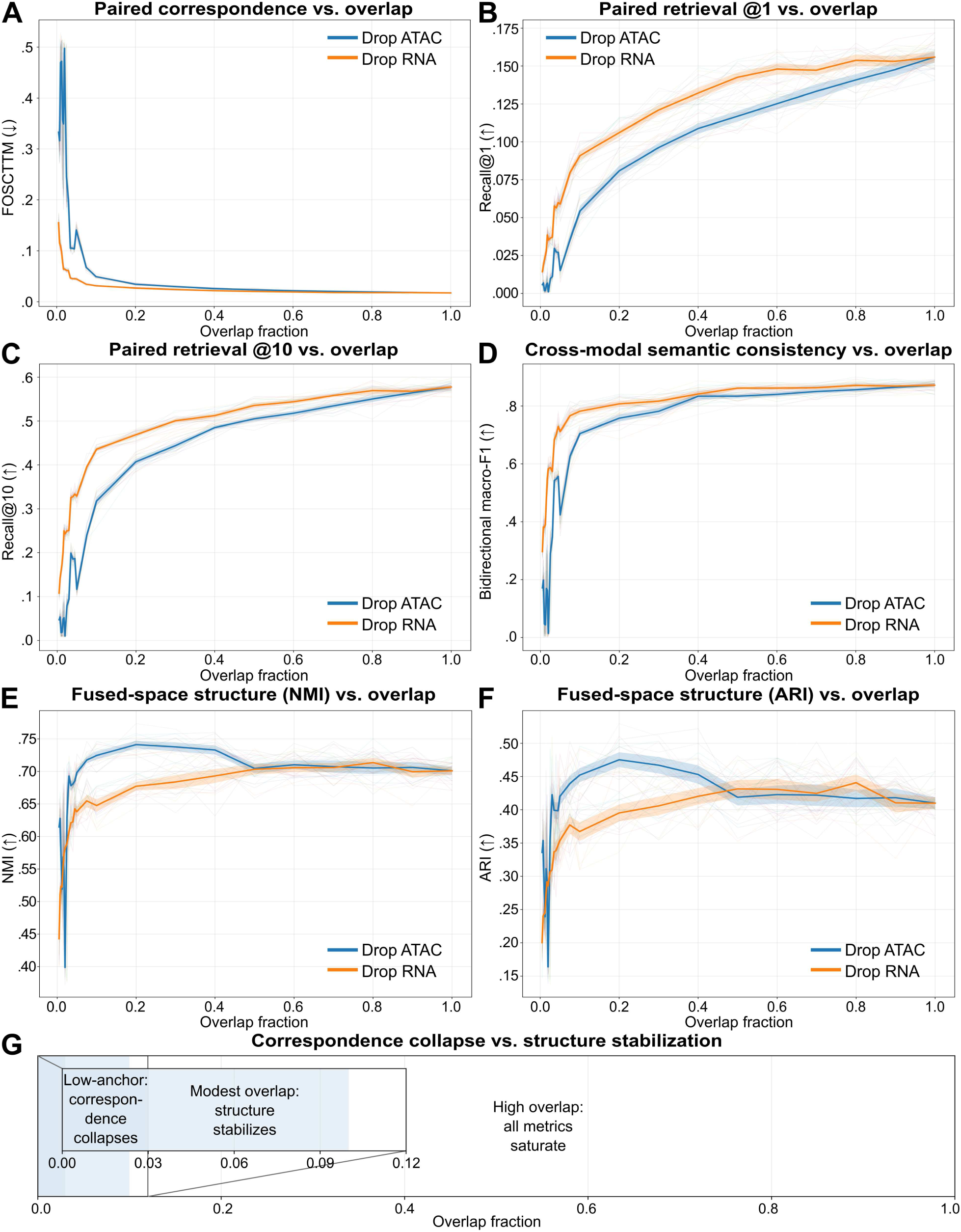
Reduced paired overlap reveals an anchor threshold. Paired 10x Multiome PBMCs were evaluated under an overlap sweep that reduced the fraction of paired RNA–ATAC anchors (Supp. Methods), either by dropping ATAC (*Drop ATAC*, blue) or dropping RNA (*Drop RNA*, orange). Lines show means across cross-validation replicates with uncertainty bands. (A) Paired correspondence (FOSCTTM; lower is better). (B–C) Paired retrieval (Recall@1, Recall@10; higher is better). (D) Cross-modality semantic consistency (bidirectional *k*-NN label transfer macro-F1; higher is better). (E–F) Fused-space biological structure (*k*-means NMI, ARI; higher is better). (G) Schematic summary of low-anchor collapse, modest-overlap stabilization, and high-overlap saturation.

Strict correspondence degrades nonlinearly as overlap decreases (Fig. 9A–C). At extremely low overlap, FOSCTTM increases sharply and paired retrieval drops toward chance, consistent with an under-constrained geometry when too few paired cells exist to reliably link modalities. At 3% overlap, Recall@10 is ≈ 0.09 for *Drop ATAC* on average, whereas *Drop RNA* remains substantially higher at the same fraction, highlighting an asymmetry in robustness depending on which modality is missing. As overlap increases beyond this low-anchor regime, correspondence improves rapidly and then continues to increase more gradually with additional anchors. In contrast, cross-modality semantic consistency measured by bidirectional *k*-NN label transfer (macro-F1) degrades more gracefully and becomes stable once modest overlap is available (Fig. 9D); by 10% overlap, bidirectional macro-F1 reaches ≈ 0.70 for *Drop ATAC*, even though strict paired retrieval continues to improve at higher overlap fractions. This separation indicates that, once a sufficient anchor subset exists, UniVI preserves cross-modal *semantic neighborhoods* even when exact one-to-one pairing fidelity remains partially underdetermined. Fused-space structure metrics improve rapidly once overlap exits the low-anchor regime and remain stable across intermediateto-high overlap fractions (Fig. 9E,F; fused-space NMI ≈ 0.72 by 10% overlap for *Drop ATAC*). Together, these results support a practical design principle for partially paired multimodal studies: UniVI benefits most from ensuring that each major population is represented by a modest paired anchor subset linking modalities, rather than requiring fully paired coverage everywhere (Fig. 9G).

### Localized missingness stress tests show graceful degradation and interpretable mixture-of-experts behavior

Missingness in real multimodal studies is often *localized* rather than global: specific lineages can fail one assay more frequently, and mosaic cohorts may contribute cells primarily from one modality. To probe UniVI’s behavior under structured missingness, we performed a cell-type-specific modality dropout stress test in paired 10x Multiome PBMCs: for each annotated population, we masked one modality (RNA or ATAC) *only for that population* during training while leaving all other populations fully paired, with evaluation on unchanged paired validation/test splits (Supp. Methods; Fig. 10). Across targeted ablations, the overall fused latent geometry remained stable: unperturbed populations retained coherent neighborhoods and relative positions, while deviations were concentrated within the ablated group (Fig. 10A). The stress test also exposes a readable mechanism: when one modality is removed for a specific population, MoE gating shifts locally toward the remaining modality while gating patterns elsewhere remain qualitatively unchanged (Fig. 10B). We interpret the signed gating preference (e.g., *w*_RNA_ − *w*_ATAC_) as a *support map* indicating which modality drives the fused representation across the manifold and which regions should be interpreted cautiously when one modality is weak or absent.

**Figure 10.**
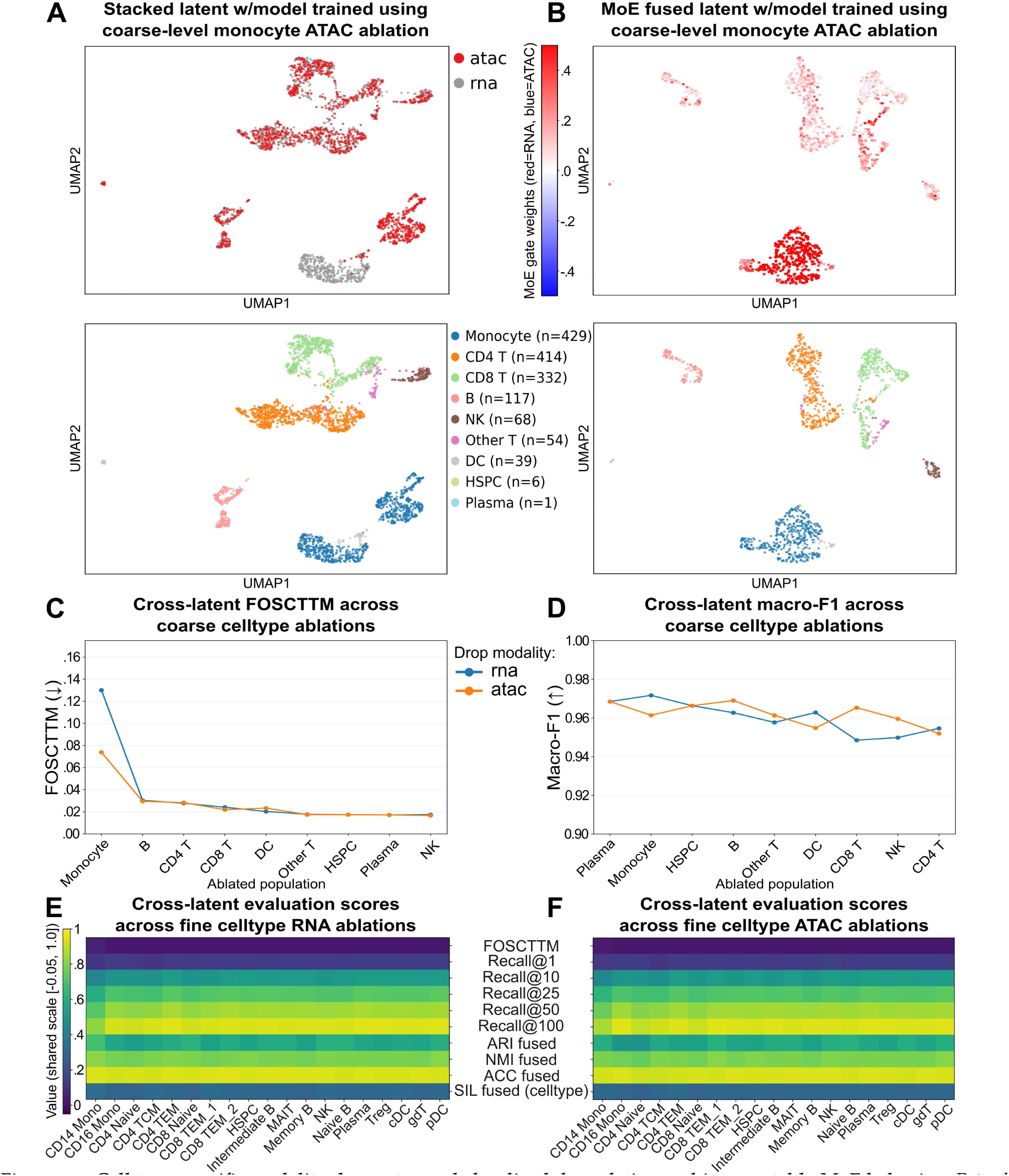
Cell-type-specific modality dropout reveals localized degradation and interpretable MoE behavior. Paired 10x Multiome PBMCs were subjected to targeted modality ablation in which one modality (RNA or ATAC) was masked *only for a single annotated population* during training; evaluation was performed on unchanged paired validation/test splits (Supp. Methods). (A) Stacked latent UMAP colored by modality for an illustrative ablation condition, showing that the perturbation is localized while global geometry is preserved. (B) MoE fused-latent UMAP colored by signed gating preference (e.g., *w*_RNA_ − *w*_ATAC_), highlighting a localized shift toward the remaining modality within the ablated region. (C) Coarse-label summary of correspondence degradation under targeted dropout, quantified on paired test embeddings (e.g., FOSCTTM; lower is better), comparing RNA-drop vs ATAC-drop conditions across ablated populations. (D) Coarse-label summary of semantic stability under targeted dropout (cross-modality label transfer macro-F1; higher is better). (E,F) Fine cell-type heatmaps summarizing correspondence and clustering diagnostics across all evaluated populations for (E) RNA-drop and (F) ATAC-drop conditions (metrics indicated per row).

Correspondence losses were largely confined to the perturbed population rather than propagating across the manifold (Fig. 10C), with fine-grained heatmaps revealing within-lineage heterogeneity under RNA- and ATAC-masked conditions (Fig. 10E,F). Despite reduced cross-modal correspondence in ablated regions, cross-modality semantic consistency remained high overall (label-transfer macro-F1; Fig. 10D), consistent with a regime where the model maintains global label structure even when one region becomes effectively unimodal. Overall, UniVI degrades gracefully under localized missingness: global organization is preserved, performance impacts are concentrated in the ablated region, and MoE gating provides an interpretable indicator of modality support, aligning with practical study designs where missingness is structured rather than uniform.

## Discussion

Multimodal single-cell integration is increasingly asked to do more than co-embed two well-matched assays from a single experiment. Modalities differ sharply in noise model, sparsity, and dynamic range, and studies often include only a modest paired “anchor” subset alongside much larger unimodal cohorts with composition shift and structured missingness. In these regimes, visually mixed embeddings can be misleading: apparent alignment can reflect aggressive coupling rather than well-supported cross-modal evidence. UniVI was designed around this gap. It couples modality-specific encoders and decoders through a shared latent prior, explicitly penalizes disagreement between modality-specific posteriors for paired cells, and provides an interpretable MoE fused representation. Together, these choices aim to preserve shared structure where evidence exists while avoiding uniform correspondence when it is weak.

Across fully paired bimodal benchmarks, UniVI produced aligned latent geometries supporting both single-cell correspondence and downstream biological utility. In CITE-seq PBMCs, RNA- and ADT-derived embeddings co-localize with minimal modality separation, and cross-modal reconstructions (RNA↔ADT) recover lineage and subtype marker patterns in both directions, providing an orthogonal check that the latent representation captures biologically meaningful cross-modal signal (Figs. 2–3; Supp. Fig. S1). In paired RNA–ATAC Multiome PBMCs, UniVI maintains strong correspondence and label transfer despite the extreme sparsity and indirect coupling of accessibility to expression (Fig. 4; Supp. Fig. S2). The mouse skin SHARE-seq evaluation then extends paired RNA–ATAC alignment to a non-hematopoietic tissue with continuous differentiation trajectories and substantially lower per-cell ATAC complexity, where lineage-scale structure is preserved across the embedding and cross-modal reconstructions recover program-level biology (Supp. Fig. S3). Together these results support posterior-level coupling as a practical mechanism for stabilizing alignment across modalities with mismatched geometry and information content, in both hematopoietic and non-hematopoietic settings.

A more stringent test of modality-agnostic generalization is whether UniVI extends to assays whose measurement statistics differ substantially from counts and surface intensities—most notably DNA methylation. The scNMT-seq mouse gastrulation analysis (Results; Supp. Fig. S4) addresses this directly: under beta-binomial likelihoods on per-feature (methylated, total) pairs, UniVI produces a coherent tri-modal latent space in which MoE gating remains balanced across RNA, CpG methylation, and GpC accessibility rather than collapsing onto RNA. We view this as a proof-of-concept rather than a comprehensive evaluation. Paired single-cell datasets combining RNA with DNA methylation remain scarce, the available cohort here (*n* = 1,140 paired cells) is modest in size, and broader characterization on methylation-containing modality pairs—across additional tissue contexts, aggregation regimes (e.g., CpG-island vs. genebody), and methylation-only or methylation+accessibility study designs—will require additional paired cohorts as they become available.

A core contribution beyond paired bimodal evaluation is the partially paired, bridge, and mosaic regimes. In the Multiome bridge regime, a paired-reference model projects independent RNA-only and ATAC-only PBMC cohorts into a shared geometry without updating generative parameters (Fig. 5; Supp. Fig. S5), with optional lightweight supervised refinement improving cross-cohort semantic consistency without refitting the generative bridge. UniVI extends cleanly to tri-modal measurements (TEA-seq; Fig. 6; Supp. Fig. S6) and to the disease-focused mosaic AML regime that most motivates “prior-light” integration, where a paired RNA–protein bridge co-organizes independent RNA+genotype and protein+genotype cohorts (Fig. 7; Supp. Fig. S7); mutation-head refinement sharpens genotype grounding and reveals a stemness continuum aligned with LSC17, with discrimination strongest where dense cell-level genotypes exist (DAb-seq) and constrained where labels are sparse (van Galen). Together, these settings illustrate that UniVI’s reach extends to clinically relevant study designs where prior-heavy integration assumptions break down.

Benchmarking against widely used methods under a unified runner places UniVI at or near the top across fused-space metrics (label transfer, ARI/NMI, silhouette) while maintaining strong cross-latent correspondence and stacked-embedding modality mixing (Fig. 8); the inductive-versus-transductive distinction matters here because UniVI is explicitly designed for forward-inference embedding of held-out cells, the operating regime relevant to reference mapping and atlas extension. Beyond point performance, the overlap sweep reveals a low-anchor threshold where strict correspondence collapses before manifold-level structure and semantic consistency stabilize (Fig. 9), and the cell-type-specific dropout stress test shows degradation concentrated in the perturbed population while global structure remains stable, with MoE gating providing an interpretable “support map” over where evidence is present (Fig. 10). These results motivate a regime-first evaluation: reporting correspondence, semantic stability, mixing, and reconstruction together rather than treating visual mixing as sufficient.

Hyperparameter and scaling analyses further support these conclusions. Joint sweeps over the KL weight *β* and the cross-modal coupling weight *γ* identify a broad interior region in which paired correspondence, bidirectional label transfer, and fused-space structure are jointly strong, with metric-specific optima distributed across the grid rather than concentrated at a single tuned corner (Supp. Fig. S8); the dominant failure pattern is confined to extreme regularization, where very large *β* underfits correspondence and label transfer and *γ* = 0 weakens cross-modal retrieval. Sweeps over latent dimensionality and encoder/decoder dropout likewise produce smooth, non-catastrophic trade-offs rather than brittle optima (Supp. Figs. S9–S10). Profiling training as a function of effective paired training size shows diminishing returns beyond moderate dataset sizes with predictable wall-clock and memory growth, and execution on an Apple Metal (MPS) backend documents laptop-class feasibility alongside the CUDA cluster runs that produced the main-text figures (Supp. Fig. S11). Together these analyses indicate that UniVI’s reported operating point reflects a robust default rather than a narrow optimum, supporting use across common multimodal study designs without extensive dataset-specific hyperparameter search.

Several limitations follow from these results. Although we evaluated UniVI in a non-hematopoietic tissue (mouse skin SHARE-seq) and on a methylation-containing tri-modal assay (scNMT-seq), the bulk of our benchmarking remains within hematopoietic biology (PBMC and AML cohorts), reflecting the current availability of richly annotated paired multimodal datasets; broader characterization across solid tumors, brain, and developmental atlases will be required to fully establish cross-tissue generality. While MoE fusion improves robustness to modality dominance and missingness, more explicit uncertainty calibration—including flagging regions where correspondence is weakly supported—remains an important methodological direction. Bridge and mosaic settings necessarily involve cohort shift; parameter-frozen projection is valuable for reuse and comparability, but projected “islands” should be interpreted using the diagnostics emphasized here rather than assumed to imply one-to-one correspondence. Mutation-head evaluation in the AML setting used cell-level splits due to metadata constraints; stronger patient-level generalization claims will require cohorts with harmonized patient identifiers and dense genotype labels supporting patient-holdout evaluation. Finally, reconstruction-based diagnostics for extremely sparse modalities (notably ATAC) depend on the chosen representation, and ATAC cross-reconstructions are best interpreted as capturing lineage- and program-scale geometry rather than peak-level biology, motivating future extensions that incorporate richer ATAC targets.

That last direction—peak-resolved inference—we are already actively pursuing. By training with count-aware likelihoods (negative binomial for RNA, Poisson or Bernoulli for ATAC) and coupling cross-modal generation to a promoter- and enhancer-derived peak-to-gene map, UniVI supports targeted in-silico peak perturbations whose predicted transcriptomic effects can be evaluated against chromosome- and width-matched null peak sets. Initial analyses (reference implementation provided in Supplemental Notebook S1; see Code availability) suggest that linked-peak perturbations elicit gene-specific responses that are not recapitulated by matched null peaks. A full characterization—including dose-response behavior, cell-type-resolved effects, and benchmarking against established peak-to-gene methods—will be reported separately.

Taken together, our results support UniVI as a prior-light, modality-agnostic generative framework that performs strongly in fully paired benchmarks while being explicitly evaluated—and behaviorally constrained—in the partially paired, bridge, and mosaic regimes that increasingly define real multimodal studies. As multimodal datasets continue to expand in scale and heterogeneity, methods will be judged not only by point performance on canonical paired tasks but by how gracefully they degrade under weak cross-modal evidence and how clearly they surface where integration is reliable.

## Methods

### UniVI model architecture and objective

UniVI is a multimodal *β*-VAE with modality-specific encoders and decoders coupled through a shared latent prior. For each modality *m* ∈ *M*, the encoder defines a diagonal-Gaussian posterior

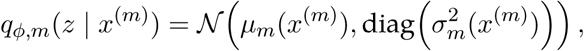

and the corresponding decoder defines a modality-appropriate likelihood *p_θ,m_*(*x*^(*m*)^ | *z*) under the shared prior *p*(*z*) = *N* (0*, I*). Latent samples are obtained using the reparameterization

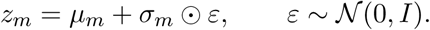

Encoders and decoders were implemented as multilayer perceptrons with normalization and dropout, with dataset-specific widths and depths reported in Supp. Table S8.

#### Training objective used throughout this study

All models were trained using `loss_mode=”v1”`, `v1_recon=”avg”`, and `normalize_v1_terms=True`. For cell *i*, let *M_i_* ⊆ *M* denote the set of observed modalities and let *K_i_* = |*M_i_*|. For source modality *k* and reconstruction target *j*, define the directed target reconstruction loss

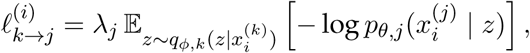

where *λ_j_*denotes the effective target-specific reconstruction scale, including any configured modality reconstruction weight and, when enabled, feature-dimension normalization.

For *K_i_* ≥ 2, v1_recon=”avg” assigns one half of the total reconstruction weight to self-reconstruction and one half to directed cross-modal reconstruction:

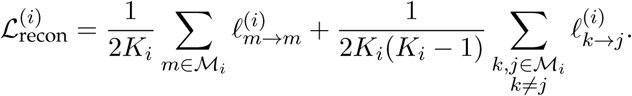

For *K_i_*= 1, the reconstruction term reduces to the available modality’s self-reconstruction loss.

Because normalize_v1_terms=True, KL regularization is averaged across observed modality-specific posteriors:

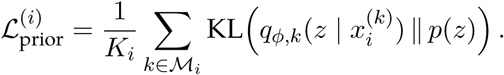

For *K_i_*≥ 2, cross-modal posterior alignment is the mean over all directed modality pairs:

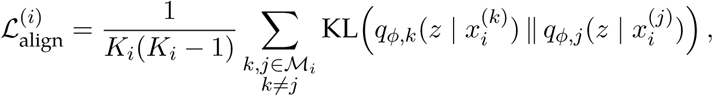

and 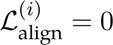 for *K_i_* = 1. Equivalently, the two directed terms for each unordered modality pair form a symmetric KL divergence with equal directional weighting.

The per-cell generative objective is

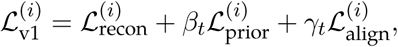

where *β_t_* and *γ_t_* are epoch-dependent weights annealed to the configured maxima. Per-cell losses are averaged across each minibatch. Auxiliary supervised losses, when used, were added separately during the refinement stages described below.

#### Closed-form Gaussian KL

For 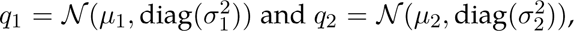 each directed KL term is

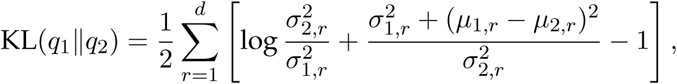

where *d* is the latent dimension. This couples modalities directly at the level of posterior means and variances for the same cell.

#### Precision-weighted MoE fused representation

The reconstruction objective above is evaluated from the modality-specific posterior samples and is distinct from fused-latent construction. For analyses requiring a single representation per cell, UniVI combines the available modality-specific Gaussian experts using their element-wise posterior precisions. Let

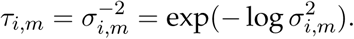

When learned MoE gating is enabled, a router produces probabilities *α_i,m_* by a softmax across available modalities, and the effective precision is

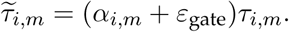

When learned gating is disabled, *τ_i,m_* = *τ_i,m_*. The fused diagonal-Gaussian posterior is

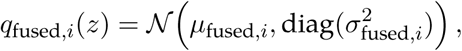

with

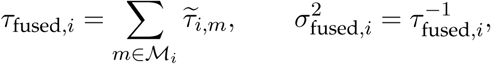

and

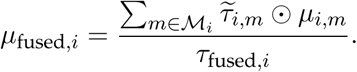

All operations are element-wise across latent dimensions. Learned MoE gating was enabled for selected fused-latent analyses; otherwise, fusion used posterior precision alone. In all cases, fused-latent construction did not replace the balanced self/cross reconstruction objective used for training.

### Optimization and training schedules

UniVI was trained in PyTorch using AdamW (Adam with decoupled weight decay) (Loshchilov and Hutter 2019; Paszke et al. 2019) with mini-batches, gradient clipping, and early stopping on validation objectives. Unless otherwise stated, learning rates were in the range 10^—4^–10^—3^ with weight decay 10^—5^–10^—4^ and batch sizes 128–256 depending on dataset and hardware constraints. KL regularization and cross-modality alignment terms were introduced gradually via epoch-based annealing schedules. All stochasticity (initialization, minibatch order, and any data subsampling) was controlled by fixed run-level seeds. Complete per-experiment configurations (widths, depths, learning rates, annealing schedules, and seeds) are reported in Supp. Table S8.

### Latent representations used for evaluation

Modality-specific embeddings refer to encoder posterior means derived from a single modality (e.g., *Z*_RNA_, *Z*_ADT_, and *Z*_ATAC_). The fused embedding *Z*_fused_ is the precision-weighted posterior mean *µ*_fused_ defined above and was used for selected fused-space visualizations, neighborhood analyses, and supervised heads. When learned gating was enabled, router probabilities modulated the posterior precisions before fusion; otherwise, the fused embedding used precision-only aggregation. For analyses assessing modality interleaving beyond paired retrieval, we used a *stacked* embedding containing one point per cell per modality in the shared latent coordinate system.

### Optional supervised refinement after projection

When labels were available and harmonizable across projected cohorts, we optionally refined latent representations by attaching a lightweight prediction head to the latent posterior mean and optimizing an auxiliary supervised loss. This refinement was used *only when indicated* (e.g., to improve cross-cohort semantic consistency in PBMC bridging or to strengthen genotype grounding in AML mosaics) and was not required for generative integration. UniVI decoders were kept frozen throughout supervised refinement. Refinement followed a two-stage schedule (head warmup with encoders/decoders frozen, then optional encoder unfreezing at a reduced learning rate); missing labels were masked and excluded from supervised loss terms and evaluation. Detailed head architectures, loss functions, and stage-specific learning rates are reported in Supp. Methods and Supp. Table S8.

### Datasets

We evaluated UniVI across the study designs enumerated in the Introduction; dataset-specific splits, preprocessing, and inductive/transductive treatment are summarized in Supp. Table S7 and Supp. Methods. Below we list each dataset and its accession.

#### PBMC CITE-seq (paired RNA–ADT)

We analyzed PBMC CITE-seq data with paired RNA and ADT measurements and immune annotations (Stoeckius et al. 2017; Hao et al. 2021) (GEO: GSE164378).

#### PBMC 10x Multiome (paired RNA–ATAC)

We used PBMC 10x Multiome data in which each cell has matched RNA and ATAC profiles (10x Genomics 2021a).

#### Mouse skin SHARE-seq (paired RNA–ATAC)

SHARE-seq paired RNA and ATAC from late-anagen mouse back skin (Ma et al. 2020) (GEO: GSE140203). Cell-type annotations from the original publication were used as ground truth. Dataset-specific QC, feature construction, and split details are reported in Supp. Methods.

#### Mouse gastrulation scNMT-seq (tri-modal RNA + DNA methylation + chromatin accessibility)

scNMT-seq mouse gastrulation data (Argelaguet et al. 2019) (GEO: GSE121708) jointly profiling RNA, CpG methylation, and GpC accessibility in the same cells. Methylation and accessibility modalities used a beta-binomial likelihood operating on per-feature (successes, coverage) pairs at the genebody level; RNA used a Gaussian likelihood on log-normalized expression. Split definitions and hyperparameters are reported in Supp. Methods and Supp. Table S8.

#### Bridge regime (paired Multiome reference + external RNA-only and ATAC-only cohorts)

For reference-to-query bridging, UniVI was trained exclusively on a paired Multiome PBMC reference (10x Genomics 2021b). We treated Ding et al. (2020) (GEO: GSE132044) as an RNA-only cohort and Satpathy et al. (2019) (GEO: GSE129785) as an ATAC-only cohort; both were embedded into the learned latent space by forward inference through the appropriate modality encoder, without updating trained generative parameters.

#### PBMC TEA-seq (paired RNA+ADT+ATAC)

We analyzed TEA-seq PBMCs (Swanson et al. 2021) (GEO: GSE158013), which jointly profile RNA, ADT, and ATAC in the same cells and include multiple replicate wells for across-well holdout evaluation.

#### AML mosaic regime (paired RNA–ADT bridge + RNA-only + protein+genotype)

We trained UniVI as a paired RNA–protein bridge on AML CITE-seq (Knorr et al. 2023) (GEO: GSE220474), then projected (i) AML scRNA-seq with targeted genotyping from Galen et al. (2019) (GEO: GSE116256) and (ii) DAb-seq (protein+targeted genotype) from Demaree et al. (2021) (NCBI BioProject: PRJNA602320) using the corresponding modality encoder. Where mutation labels were available, we optionally performed supervised refinement using per-gene mutation heads. Additional cohort context, mutation readout totals, and label availability are provided in Supp. Tables S1–S5.

### Preprocessing and feature construction

Raw counts were stored in `.layers[”counts”]` and model inputs were stored in `.X` (or a specified `X_key`). Preprocessing steps that learn parameters were fit on training data only and applied unchanged thereafter, including for projected cohorts in bridge/mosaic settings (see Supp. Table S7).

#### RNA

RNA inputs were library-size normalized to a fixed depth (target sum 10^4^ counts per cell) followed by log(1+*x*). Highly variable genes (HVGs) were selected on training cells only using SCANPY with `}avor=”seurat_v3”`; the resulting gene set was reused unchanged for validation/test. When per-gene scaling was used (e.g., for Gaussian decoders), *Z*-scoring statistics were fit on training cells and applied unchanged thereafter. Dataset-specific HVG counts, gene-family exclusions, and harmonization choices (e.g., reuse of reference HVGs in bridging; gene intersections in AML mosaics) are reported in Supp. Methods.

#### ADT

ADT features were CLR-normalized per cell. Low-information/control antibodies (e.g., isotype controls) were excluded during panel harmonization when applicable. When Gaussian decoders were used, ADT features were optionally *Z*-scored per marker using training statistics and applied unchanged thereafter. Cross-cohort marker harmonization for AML mosaic projection is detailed in Supp. Methods.

#### ATAC

For scATAC-seq datasets, we started from author-provided peak-by-cell (or tile-by-cell) matrices when available. Peak/tile matrices were transformed using TF–IDF followed by truncated SVD/LSI: TF–IDF was fit on training cells (scikit-learn `T|dfTransformer` with `use_idf=True`, `smooth_idf=True`, L2 normalization), applied unchanged thereafter, then LSI was computed with `TruncatedSVD`. This TF–IDF+LSI representation is widely used in scATAC-seq workflows (Cusanovich et al. 2015; Granja et al. 2021; Stuart et al. 2021). When scaling was enabled, LSI components were standardized using training statistics and applied unchanged thereafter. Dataset-specific tile/peak construction, depth-confounded component handling, and fragment thresholds are reported in Supp. Methods.

#### Mutation labels (AML)

Mutation calls were represented as per-gene binary targets for recurrent AML genes where available. Labels were stored as a multi-label target matrix (one column per gene), and missing calls were explicitly masked so they were excluded from supervised loss terms and per-gene evaluation (Supp. Tables S2–S4).

### Splits, leakage control, and inductive evaluation

All splits were defined as barcode-indexed maps (AnnData `.obs_names`) and persisted to disk for exact reproducibility. For paired datasets, pairing was enforced by strict barcode intersection (1:1 pairing) prior to split assignment. Unless otherwise stated, model parameters were learned on training splits (with early stopping on validation where applicable), and all validation/test embeddings were obtained by forward inference without updating fitted parameters. Preprocessing transforms that learn parameters (HVG selection, scaling, TF–IDF, SVD/LSI, standardization) were fit using training cells only and applied unchanged to validation/test and to projected cohorts (Supp. Table S7). Dataset-specific stratification labels, per-class caps, well/sample holdout definitions, and patient-level considerations are detailed in Supp. Methods.

### Evaluation metrics

We quantified multimodal integration using complementary metrics assessing (i) paired correspondence at single-cell resolution, (ii) semantic consistency via cross-modality label transfer, (iii) modality interleaving in shared latent neighborhoods, and (iv) label separation in fused space. Unless otherwise stated, distances were computed in latent space using squared Euclidean distance. Neighbor sizes *k* are reported in figure captions. UMAP visualizations were computed for qualitative assessment of latent geometry.

#### Paired alignment: FOSCTTM and Recall@K

For paired datasets, we computed FOSCTTM (Singh et al. 2020; Liu et al. 2025) using squared Euclidean distances between modality-specific embeddings. For each cell in modality A, we ranked all cells in modality B by distance and computed the fraction closer than the true paired partner; lower values indicate stronger correspondence. We additionally report Recall@K, defined as the fraction of cells whose true paired partner is among the top-*K* nearest cross-modality neighbors (as measured by *k*-NN).

#### *k*-NN label transfer: accuracy and macro-F1

We performed bidirectional *k*-NN label transfer (Wolf et al. 2018; Stuart et al. 2019). In each direction (A→B and B→A), target labels were assigned by majority vote from the *k* nearest neighbors in the source embedding space and evaluated against ground truth. We report accuracy (ACC) and macro-F1; confusion matrices were row-normalized.

#### Modality mixing on stacked embeddings

We assessed modality interleaving on the stacked embedding (Luecken et al. 2022) by building a latent *k*-NN graph and summarizing interleaving as the fraction of neighbors drawn from a different modality. When reported, complementary diagnostics (e.g., modality entropy or silhouette by modality) were computed on the same graph.

#### Fused-space clustering and label separation: ARI, NMI, and silhouette

We ran *k*-means clustering on the fused embedding with *k* set to the number of evaluated label classes and compared assignments to ground-truth labels using ARI and NMI. We also computed a label-based silhouette score on the fused embedding.

#### Cross-modal reconstruction diagnostics

We evaluated cross-modal reconstruction by encoding one modality and decoding another (e.g., RNA→ADT and ADT→RNA; ATAC→RNA and RNA→ATAC) (Gayoso et al. 2021; Ashuach et al. 2023). Reconstructions were used as diagnostics to confirm that latent structure captures cross-modal signal rather than only geometric alignment.

### Benchmarking protocol and baselines

We benchmarked UniVI against representative integration approaches spanning CCA/anchor workflows and graph fusion (Seurat CCA, WNN, Seurat Bridge) (Butler et al. 2018; Stuart et al. 2019; Hao et al. 2021; Hao et al. 2023), factorization/latent factor models (LIGER, MOFA2) (Welch et al. 2019; Argelaguet et al. 2018; Argelaguet et al. 2020), shared-embedding correction (Harmony) (Korsunsky et al. 2019), MA (MultiMAP) (Jain et al. 2021), deep generative models (PeakVI, MultiVI) (Ashuach et al. 2022; Ashuach et al. 2023), deep alignment (DeepCCA) (Andrew et al. 2013), and additional multimodal frameworks (CoBOLT, scGLUE, scJoint, scMoMaT) (Gong et al. 2021; ZJ Cao and G Gao 2022; Lin et al. 2022; Zhang et al. 2023). Methods were executed through a unified benchmarking runner where possible, with split maps, preprocessing, and metric computation applied consistently across methods.

#### Inductive/transductive categorization and embedding eligibility

Within each replicate run (random seed × fold), preprocessing was fit on training cells only and applied unchanged to validation/test, and persisted fold maps were reused across methods. Methods that support clean out-of-sample embedding of held-out cells without refitting parameters were evaluated inductively; methods whose standard workflows incorporate held-out cells during fitting were treated as transductive and flagged. The resulting categorization, runner-specific notes for each baseline, and per-method embedding/metric eligibility are summarized in Supp. Table S6 and Supp. Methods.

#### Repeated runs and aggregation

We evaluated each method over five random seeds and 3-fold cross-validation per seed (5 × 3 = 15 runs per method per dataset). Benchmark panels report the distribution across replicate runs.

### Implementation and reproducibility

All analyses were performed in Python using the AnnData ecosystem (SCANPY/Muon) (Wolf et al. 2018; Virshup et al. 2021; Muon Team 2022). UniVI was implemented in PyTorch (Paszke et al. 2019). Random seeds were controlled at the runner level, and train/validation/test split maps were persisted as barcode-indexed files and reused across reruns and modality-specific objects. When methods required separate software stacks (e.g., Python vs R), we executed them in separate environments while reusing the same fold maps and seed sets. The principal training and evaluation workloads were executed on the Oregon Health & Science University (OHSU) Advanced Research Computing (ARC) cluster, which provided the GPU resources used for all main-text figures (see Supp. Table S8 for specific resource used per-figure). Selected hyperparameter sweep/scaling/resource profiling experiments were additionally executed on an Apple MacBook Pro (M1 Max) to document laptop-class feasibility (Supp. Fig. S11; Supp. Table S8).

## Code availability

The UniVI implementation, configuration files, scripts, and analysis notebooks used to generate the main-text and supplemental figures are openly available as:

- **GitHub repository:** https://github.com/Ashford-A/UniVI
- **PyPI package:** https://pypi.org/project/univi
- **Conda package (conda-forge):** https://anaconda.org/conda-forge/univi
- **Supplemental Code archive:** Supplemental_Code.zip—a snapshot of the GitHub repository pinned to release v0.4.7, containing the univi/ package source, para meter_|les/, scripts/, envs/, notebooks/, Supplemental_Notebook_ S1.ipynb, and MANIFEST.txt.

This manuscript was prepared against release `v0.4.7`, with matching tags on GitHub, PyPI, and conda-forge; we recommend pinning to this release when reproducing specific numerical results. The canonical, versioned source is the GitHub repository, which also hosts any post-publication updates.

### Python package

The `univi` package (`import univi`) provides the model architecture, objectives, training utilities, evaluation metrics, and plotting helpers used throughout this manuscript: construction and training of UniVI models with modality-specific encoders/decoders and selectable objectives (e.g., `loss_mode=”v1”`); encoding of single-cell AnnData objects into modality-specific and fused latent embeddings; cross-modal reconstruction and prediction (e.g., RNA→ADT, ADT→RNA, ATAC→RNA), with outputs ready for attachment to AnnData `.layers` and `.obsm`; and evaluation routines for correspondence (FOSCTTM, Recall@*k*), *k*-NN label transfer (accuracy and macro-F1), mixing diagnostics, and feature-level reconstruction summaries.

### Per-figure reproducibility

All analysis notebooks reside under `notebooks/GR_manu script_reproducibility/` with figure-keyed names (e.g., `UniVI_manuscript_GR-`Figure 2 `CITE_paired.ipynb` reproduces Fig. 2, and analogously through Fig. 10). Supplemental figures are co-located with their corresponding main-text notebook: Supp. Fig. S1 alongside Fig. 3 in `UniVI_manuscript_GR-`Figure 3 `CITE_paired_biological_latent.ipynb` (which also covers Fig. 2); Supp. Fig. S2 in the Fig. 4 notebook; Supp. Fig. S5 in the Fig. 5 notebook; Supp. Fig. S6 in the Fig. 6 notebook; and Supp. Fig. S7 in the Fig. 7 notebook. Supp. Figs. S3 and S4 are produced by dedicated standalone notebooks (`UniVI_manuscript_GR-Supple mouse_skin_SHARE-seq_integration.ipynb` and `UniVI_manuscript_GR-Supple scNMT-seq_mouse_gastrulation_data.ipynb`, respectively), and Supp. Figs. S8–S11 by the grid-sweep notebook (`UniVI_manuscript_GR-Supple grid-sweep.ipynb`) and its `_compile_plots_from_results_df` companion. Figures whose panels aggregate multiple sweeps (Figs. 8–10) likewise have a `_compile_plots_from_results_df` companion notebook that reassembles published panels from persisted results tables.

Dataset-specific preprocessing, model hyperparameters, and training schedules are pinned in matching configurations under `parameter_|les/` (e.g., `params_citeseq_pbmc_GR_|g2_3.json`, `params_multiome_pbmc_GR_|g4.json`, `params_aml_citeseq_ GR_|g7.json`; also listed in Supp. Tables S7–S8). Reproducible entry-point scripts live under `scripts/`, with a master driver `scripts/revision_reproduce_all.sh` that orchestrates the full pipeline.

### Supplemental Notebook S1

Supplemental_Notebook_S1.ipynb (mirrored in no tebooks/GR_manuscript_reproducibility/) provides the reference implementation for the peak-resolved inference experiment discussed in the Discussion: count-aware likelihoods (negative binomial RNA, Poisson or Bernoulli ATAC), the promoter- and enhancer-derived peak-to-gene linkage map, the in-silico peak-perturbation procedure (modes: o{/on/ set/scale/add), and the chromosome- and width-matched null-peak specificity test.

### Environments

Example environment specifications under envs/—notably envs/univi_ v0.4.7_env.yml, matching release v0.4.7—together with pyproject.toml and conda.recipe/, support installation and rerunning the notebooks and scripts under consistent dependencies.

## Acknowledgments

We thank Hyeyoung Cho for foundational contributions that helped shape the early direction of this project. We also acknowledge the Oregon Health & Science University (OHSU) Advanced Computing Center (ACC) for providing computational infrastructure support, including Exa-cloud and the ACC research cluster (ARC). The research reported in this publication utilized computational infrastructure supported by the Office of Research Infrastructure Programs, Office of the Director, National Institutes of Health, under Award Number S10OD034224. Analyses were conducted using a combination of institutional computing resources and local workstation hardware for profiling experiments. Finally, we thank the investigators who generated the publicly available datasets analyzed in this study, as well as the maintainers of public repositories that enable open data sharing and open-source software.

## Author contributions

Conceptualization: AJA, ED. Methodology: AJA, ED, TE, JS. Investigation: AJA, TE. Software: AJA, TE. Formal analysis: AJA. Visualization: AJA. Writing–original draft: AJA, ED. Writing–review & editing: AJA, ED, TE; ON contributed editorial input. Supervision: ED.

