## Supplemental Material for "Unifying multimodal single-cell data with a mixture-of-experts *β*-variational autoencoder framework"

This document provides supplemental figures, tables, and methods supporting the main text, "Unifying multimodal single-cell data with a mixture-of-experts $\beta$-variational autoencoder framework". Figure and table numbering uses the prefix "S" (e.g., Fig. S1, Table S1).

#### Abbreviations

AML, acute myeloid leukemia; AP, average precision; ARI, adjusted Rand index; AUC, area under the (ROC) curve; BM, bone marrow; BN, batch normalization; CCA, canonical correlation analysis; CITE-seq, cellular indexing of transcriptomes and epitopes by sequencing; CLR, centered log-ratio; CV, cross-validation; ELN, European LeukemiaNet; ES, early stopping; ExE, extra-embryonic; FOSCTTM, fraction of samples closer than the true match; GEO, Gene Expression Omnibus; HLA, human leukocyte antigen; HVG, highly variable genes; iNMF, integrative non-negative matrix factorization; IRS, inner root sheath; ITD, internal tandem duplication; KD, kinase domain; KL, Kullback–Leibler (divergence); $k$-NN, $k$-nearest neighbors; LayerNorm, layer normalization; LR, learning rate; LSC17, leukemic stem cell 17-gene signature; LSI, latent semantic indexing; LSN, library-size normalization; MLP, multilayer perceptron; MoE, mixture-of-experts; MSE, mean squared error; Mut, mutant; NMI, normalized mutual information; PBMC, peripheral blood mononuclear cell; PCR, polymerase chain reaction; qPCR, quantitative PCR; R@$k$, Recall at $k$; ROC, receiver operating characteristic; RSS, resident set size; scATAC-seq, single-cell assay for transposase-accessible chromatin using sequencing; scRNA-seq, single-cell RNA sequencing; SEM, standard error of the mean; SVD, singular value decomposition; TAC, transit-amplifying cell; TF–IDF, term frequency–inverse document frequency; Tx, treatment; VAE, variational autoencoder; VAF, variant allele frequency; WD, weight decay; WHO, World Health Organization; WNN, weighted nearest neighbors; WT, wild-type.

### Supplemental Figures

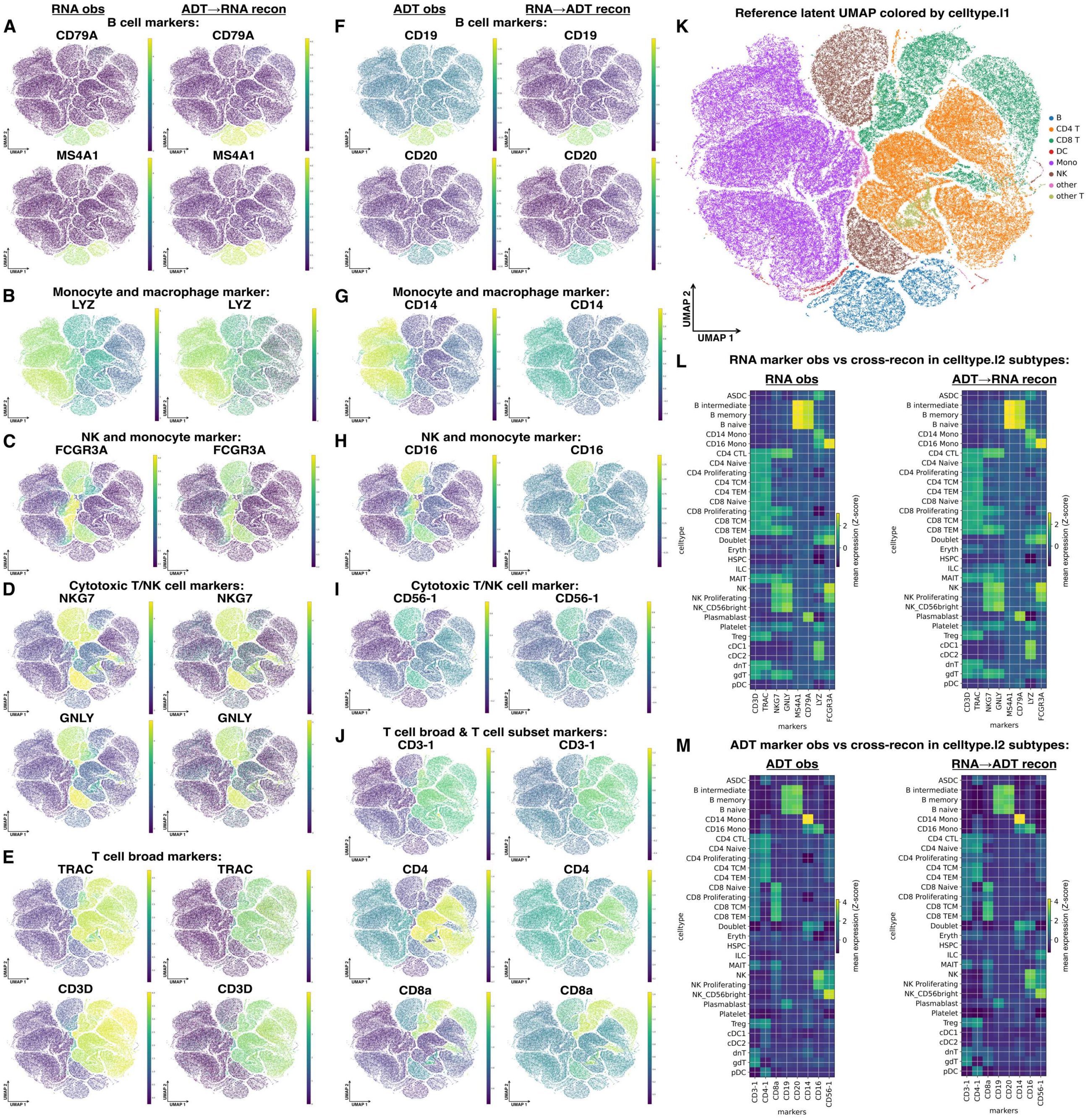


**Figure S1. Expanded version of Fig. 3 demonstrating that UniVI preserves CITE-seq biological structure and enables accurate RNA↔ADT cross-modality reconstruction across an extended marker set and finer immune subtypes.** This supplemental figure mirrors Fig. 3 but expands the marker panels shown on the UMAP and evaluates marker recovery at higher-resolution annotations (`celltype.l2`). In all UMAP panels, the horizontal axis is UMAP1 and the vertical axis is UMAP2. For marker overlays, the color scale denotes *Z-scored normalized expression/abundance* values computed *per marker across cells* (after the modality-specific normalization used in this study), consistent with the labeling used in the corresponding main-figure panels. (A–E) RNA marker overlays on the reference UMAP comparing observed RNA ($X_{\mathrm{RNA}}$) to RNA predicted from ADT cross-reconstruction ($X_{\mathrm{ADT}} \rightarrow \hat{X}_{\mathrm{RNA}}$), shown for representative B cell (*CD79A*, *MS4A1*), monocyte/macrophage (*LYZ*), NK/monocyte (*FCGR3A*), cytotoxic T/NK (*NKG7*, *GNLY*), and T cell (*TRAC*, *CD3D*) markers. (F–J) ADT marker overlays comparing observed ADT ($X_{\mathrm{ADT}}$) to ADT predicted from RNA cross-reconstruction ($X_{\mathrm{RNA}} \rightarrow \hat{X}_{\mathrm{ADT}}$), including B cell (CD19, CD20), monocyte/macrophage (CD14), NK/monocyte (CD16), cytotoxic T/NK (CD56-1), and broad/subset T cell markers (CD3-1, CD4, CD8a). (K) Reference UMAP colored by coarse `celltype.l1`. (L) Mean RNA marker expression by `celltype.l2` subtypes for observed RNA (left) versus ADT→RNA cross-reconstruction (right), demonstrating subtype-specific marker patterns are retained after cross-modality prediction. (M) Mean ADT marker abundance by `celltype.l2` subtypes for observed ADT (left) versus RNA→ADT cross-reconstruction (right), showing concordant recovery of subtype-enriched surface markers.

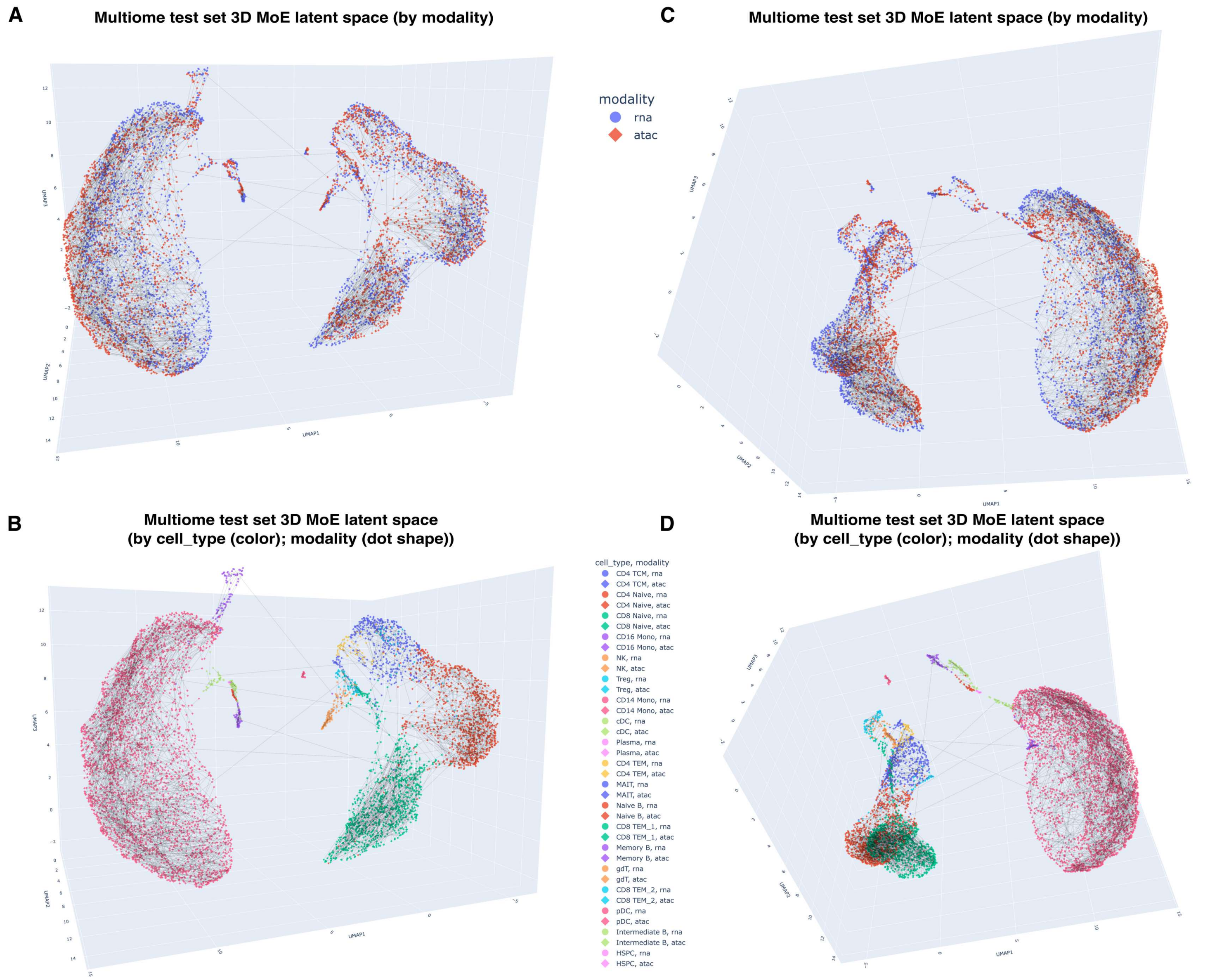


**Figure S2. 3D visualization of the held-out 10x Multiome test set in the UniVI latent space confirms cross-modality co-localization beyond 2D projection effects.** Using the same UniVI model trained for Fig. 4, we encoded the held-out paired test cells and plotted modality-specific posterior means in 3D latent coordinates. (A,C) Points colored by modality (RNA vs ATAC) show broad interleaving across the manifold. (B,D) Points colored by `cell_type` (with modality indicated by marker shape) show that major immune populations form coherent regions while maintaining close RNA–ATAC co-localization. Panels (A,B) and (C,D) show two camera views of the same embedding to reduce reliance on any single projection angle.

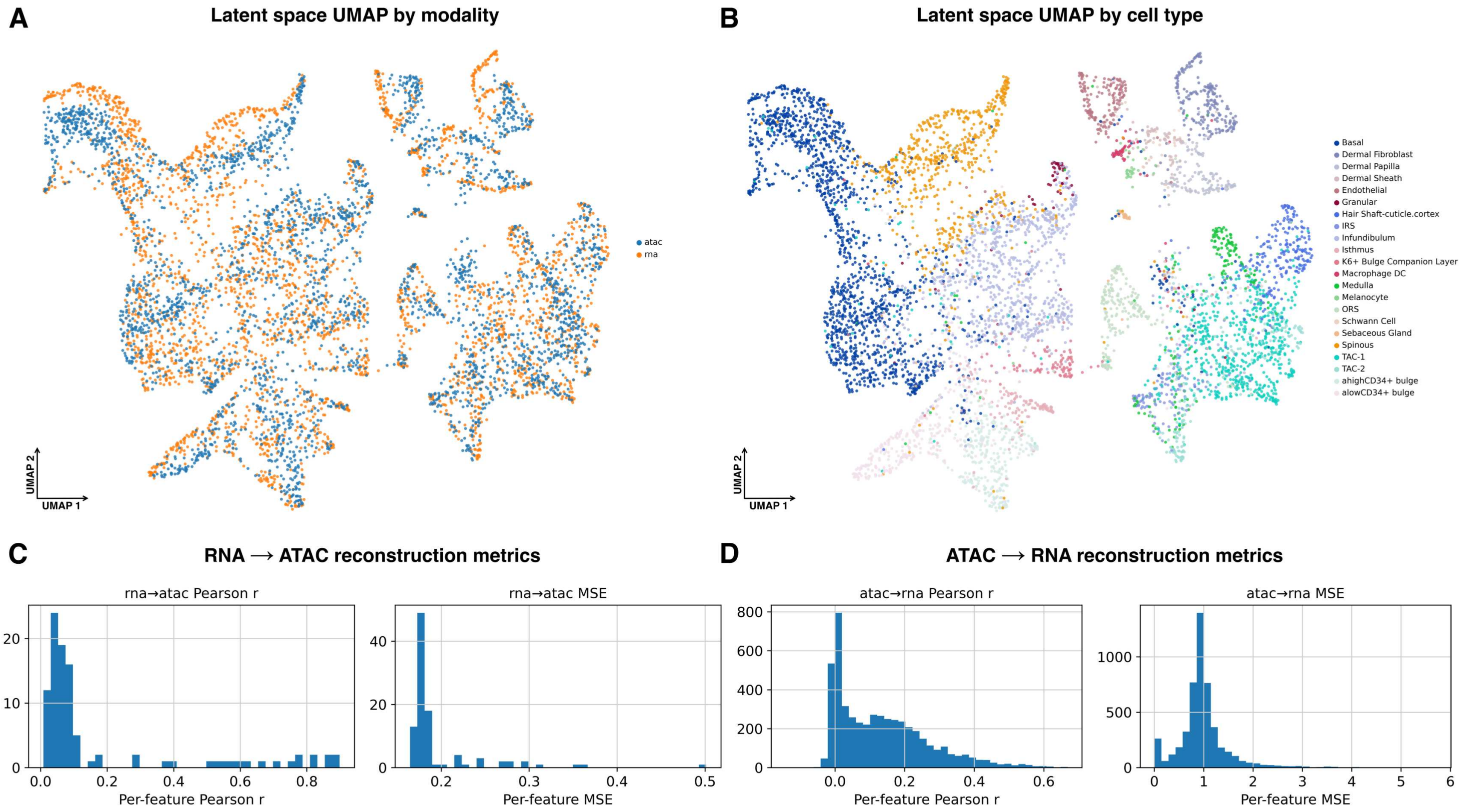


**Figure S3. UniVI extends to non-hematopoietic tissue and continuous differentiation hierarchies in paired SHARE-seq mouse skin.** Paired SHARE-seq RNA and ATAC measurements of late-anagen mouse back skin (Ma et al. 2020) were integrated with UniVI; held-out paired test cells ($n = 3{,}139$) span 22 cell types from epidermal, hair-follicle differentiation, dermal mesenchymal, neural-crest-derived, vascular, and immune lineages (Methods). (A) Stacked latent UMAP from modality-specific posterior means ($Z_{\mathrm{RNA}}$, $Z_{\mathrm{ATAC}}$) colored by modality (RNA vs ATAC), showing broad cross-modal interleaving despite the substantially lower per-cell ATAC complexity of SHARE-seq compared with 10x Multiome PBMCs. (B) Same UMAP colored by original `cell_type` annotation; coherent regions form per lineage, with adjacent compartments corresponding to known biological transitions along the hair-follicle differentiation trajectory (e.g., TAC-1 → TAC-2 → IRS / medulla / hair shaft cuticle). (C) Cross-modal RNA → ATAC reconstruction metrics on the held-out paired test set in the ATAC TF–IDF + LSI representation: per-feature Pearson $r$ (left) and per-feature MSE (right) histograms. (D) Cross-modal ATAC → RNA reconstruction metrics on $Z$-scored log-normalized expression: per-feature Pearson $r$ (left) and per-feature MSE (right) histograms. Summary: FOSCTTM $0.0546 \pm 0.00199$ (SEM); bidirectional Recall@10 0.168 (RNA→ATAC) and 0.162 (ATAC→RNA); $k$-NN modality mixing 0.326 ($k = 30$); ATAC→RNA $k$-NN label-transfer accuracy 0.774 (macro-F1 0.666); mean per-feature Pearson $r$ 0.184 (RNA→ATAC) and 0.132 (ATAC→RNA).

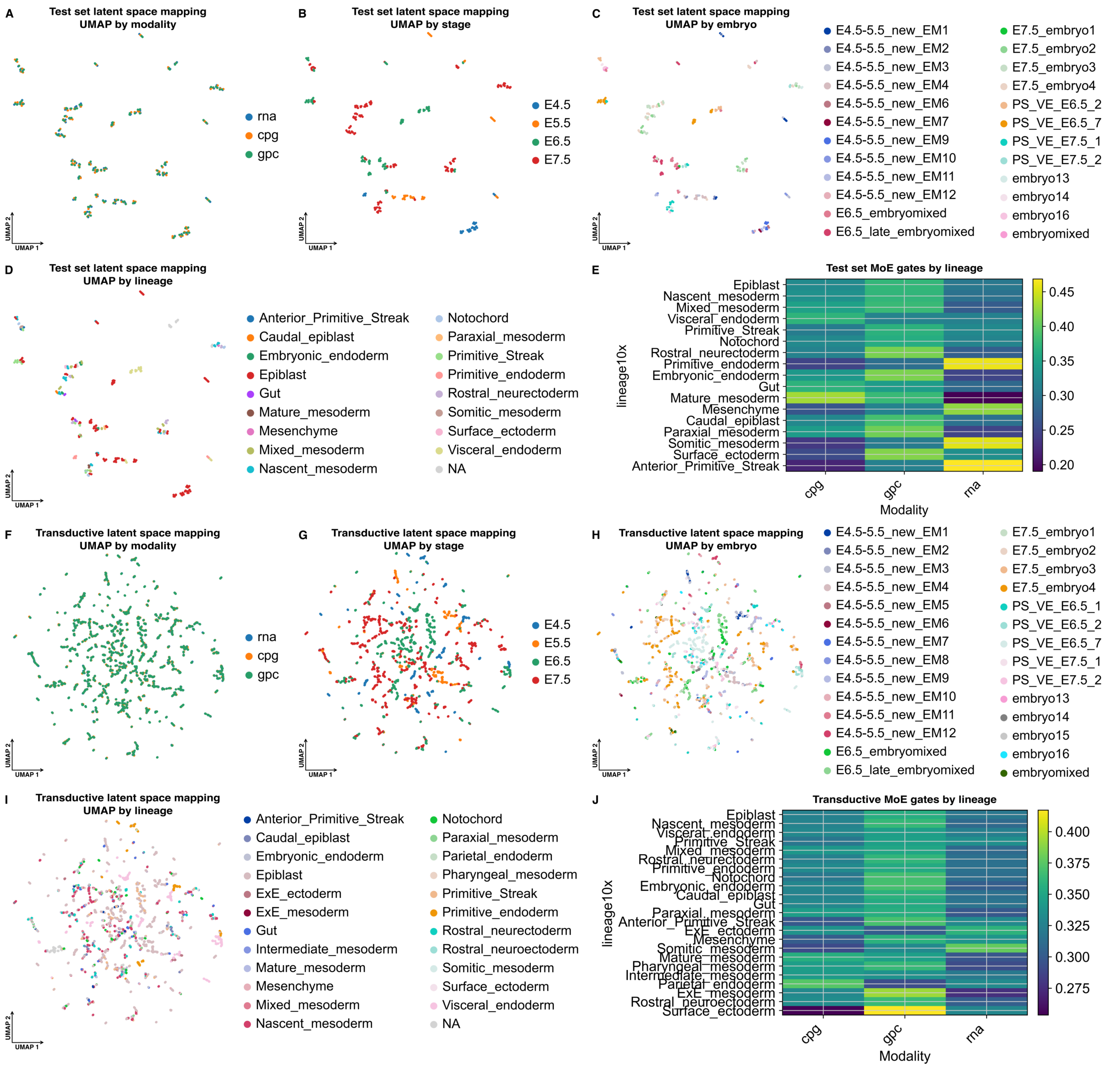


**Figure S4. UniVI accommodates paired single-cell DNA methylation in a tri-modal scNMT-seq mouse gastrulation proof-of-concept.** Paired scNMT-seq mouse gastrulation data (Argelaguet et al. 2019) jointly profiling RNA, CpG methylation, and GpC accessibility were integrated under tri-modal training; methylation and accessibility used beta-binomial likelihoods on per-feature (successes, coverage) pairs, while RNA used a Gaussian likelihood on log-normalized expression (Methods). **Top rows (A–E):** *inductive* test set ($n = 114$; 85%/5%/10% split, seed 0). (A) Stacked latent UMAP from modality-specific posterior means, colored by modality (RNA, CpG, GpC). (B) Same UMAP colored by developmental stage (E4.5–E7.5), recapitulating stage order despite modest test size. (C) Same UMAP colored by embryo of origin; embryo-level batch effects are not dominant. (D) Same UMAP colored by `lineage10x` (Pijuan-Sala atlas transfer (Pijuan-Sala et al. 2019)), showing lineage-coherent regions. (E) Mixture-of-experts (MoE) gating weights averaged within each `lineage10x` group; gating is balanced across modalities (per-lineage range ~0.20–0.46), with mature mesoderm preferentially weighting CpG/GpC and primitive endoderm preferentially weighting RNA. **Bottom rows (F–J):** *transductive* evaluation on the full paired dataset ($n = 1{,}140$; same checkpoint, forward inference). (F–I) As in (A–D), with additional lineages resolved (ExE_ectoderm, ExE_mesoderm, Parietal/Pharyngeal/Intermediate mesoderm, Rostral neuroectoderm). (J) MoE gating as in (E), with sharper per-lineage contrast (Surface_ectoderm: strong GpC; ExE_mesoderm: strong CpG). Summary: inductive FOSCTTM $0.192 \pm 0.024$ (R@1 0.070, R@10 0.526); transductive FOSCTTM $0.029 \pm 0.003$ (R@1 0.250, R@10 0.691). Mean MoE gating: CpG 0.31, GpC 0.40, RNA 0.29.

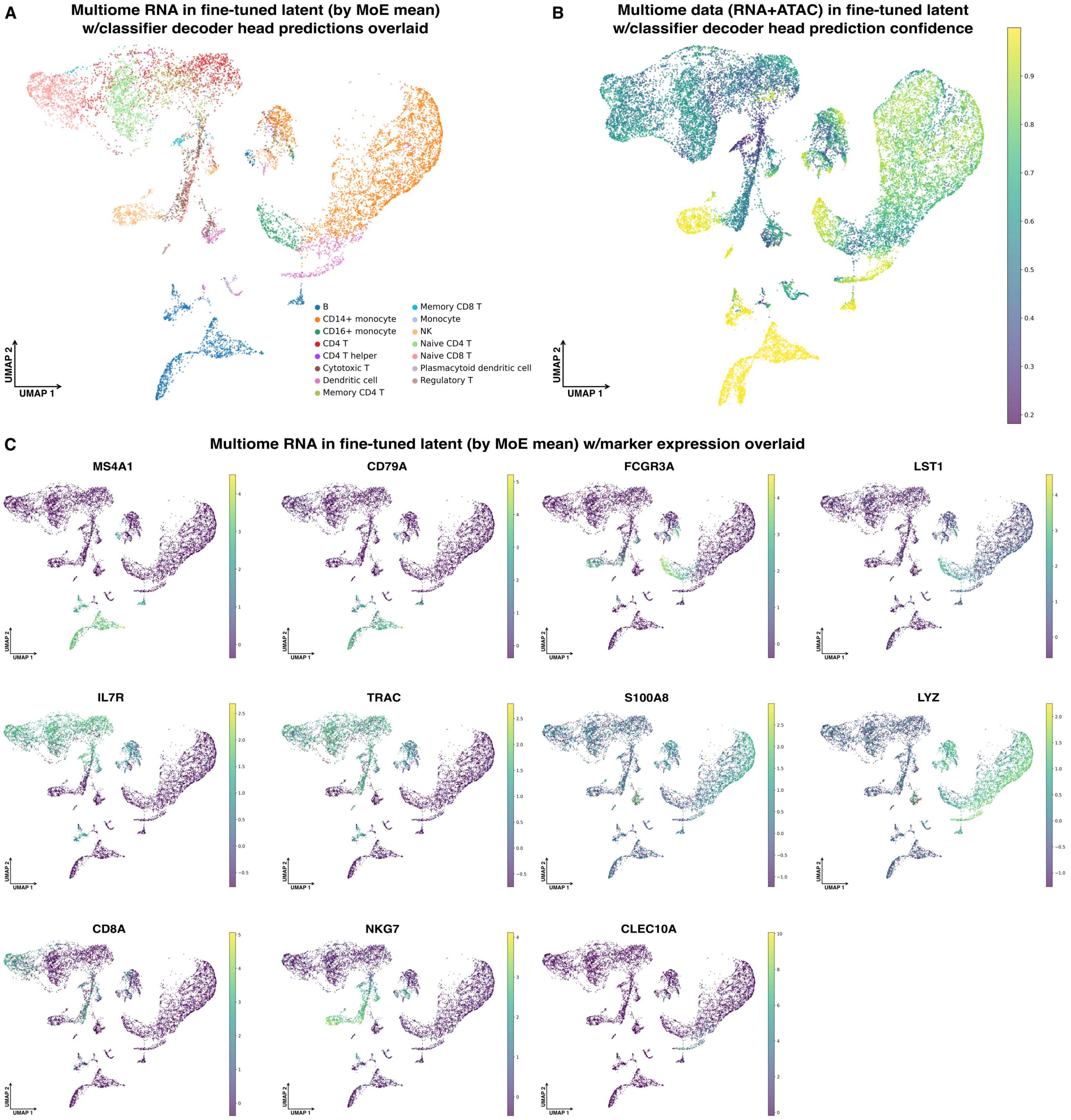


**Figure S5. Expanded marker-based validation of transferred coarse PBMC labels on unlabeled Multiome bridge cells after supervised refinement (expands Fig. 5H–J).** Starting from the Multiome-trained bridge checkpoint, we applied the lightweight refinement classifier head (decoders frozen; Methods) and then evaluated predictions on the Multiome bridge cells, which were not used as a source of supervision. In all UMAP panels, the horizontal axis is UMAP1 and the vertical axis is UMAP2; marker overlay intensities are shown as *Z-scored normalized expression* values computed *per marker across cells* (after modality-specific normalization), consistent with the corresponding main-figure panels. (A) Multiome RNA cells embedded in the refined latent space (MoE mean) with classifier-predicted coarse PBMC labels overlaid, showing coherent lineage-scale regions. (B) The same embedding colored by prediction confidence, highlighting that high-confidence calls concentrate within well-separated lineages while lower-confidence predictions occur near transitional boundaries. (C) Expanded RNA marker overlays in Multiome (e.g., *MS4A1*/*CD79A* for B cells; *FCGR3A*/*LST1* for myeloid; *IL7R*/*TRAC*/*CD8A*/*NKG7* for T/NK; *S100A8*/*LYZ* for inflammatory myeloid; and *CLEC10A* for dendritic subsets) confirm that predicted label structure agrees with canonical lineage marker expression across the refined embedding.

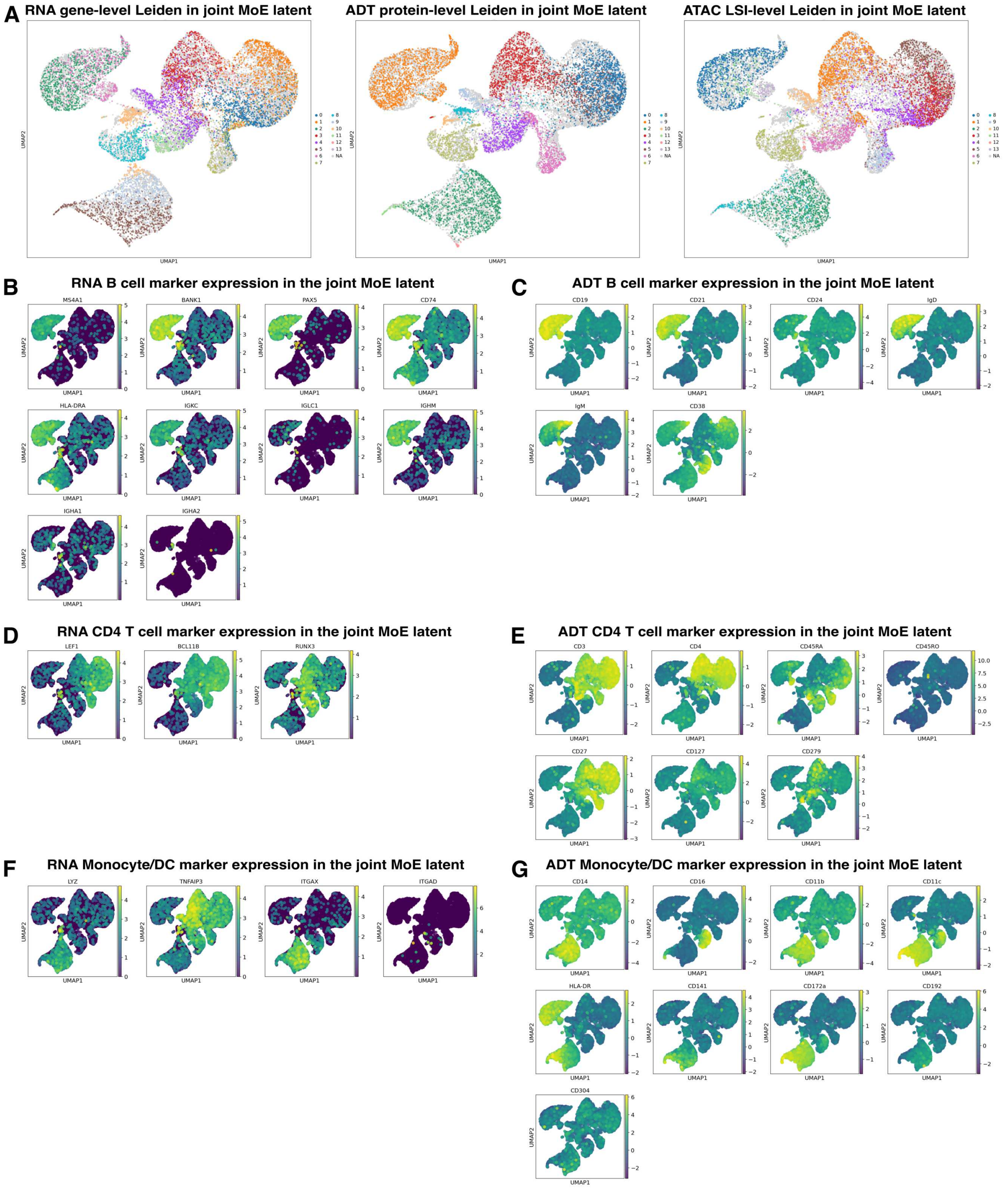


**Figure S6. Expanded view of unsupervised structure and cross-modality marker concordance in the TEA-seq well 5 sample (expands Fig. 6D–G).** UniVI was trained on TEA-seq wells 3–4 and 6 and evaluated on well 5 (Methods); wells correspond to within-run capture/library partitions and are used here as a lightweight robustness control. In all UMAP panels, the horizontal axis is UMAP1 and the vertical axis is UMAP2. Marker overlays (RNA or ADT) are displayed as *Z-scored normalized expression/abundance* values computed *per marker across cells* (after modality-specific normalization), consistent with the corresponding main-figure panels. (A) Latent UMAP colored by unsupervised Leiden clusters computed separately from RNA gene-level, ADT protein-level, and ATAC LSI-level neighborhood graphs in the shared MoE latent space, illustrating that major partitions are broadly consistent across modalities. (B,C) B cell lineage signals in the joint latent space, shown as RNA expression overlays (*MS4A1*, *BANK1*, *PAX5*, *CD74*, *HLA-DRA*, *IGKC*, *IGLC1*, *IGHM*, *IGHA1*, *IGHA2*) alongside corresponding ADT protein overlays (CD19, CD21, CD24, IgD, IgM, CD38). (D,E) CD4 T cell lineage signals, shown as RNA overlays (*LEF1*, *BCL11B*, *RUNX3*) and ADT overlays (CD3, CD4, CD45RA, CD45RO, CD27, CD127, CD279). (F,G) Monocyte/dendritic compartments, shown as RNA overlays (*LYZ*, *TNFAIP3*, *ITGAX*, *ITGAD*) and ADT overlays (CD14, CD16, CD11b, CD11c, HLA-DR, CD141, CD172a, CD192, CD304). Together, these panels provide an expanded qualitative view that the shared latent geometry supports coherent immune-lineage structure and concordant marker patterns across RNA, protein, and chromatin accessibility in this held-out well.

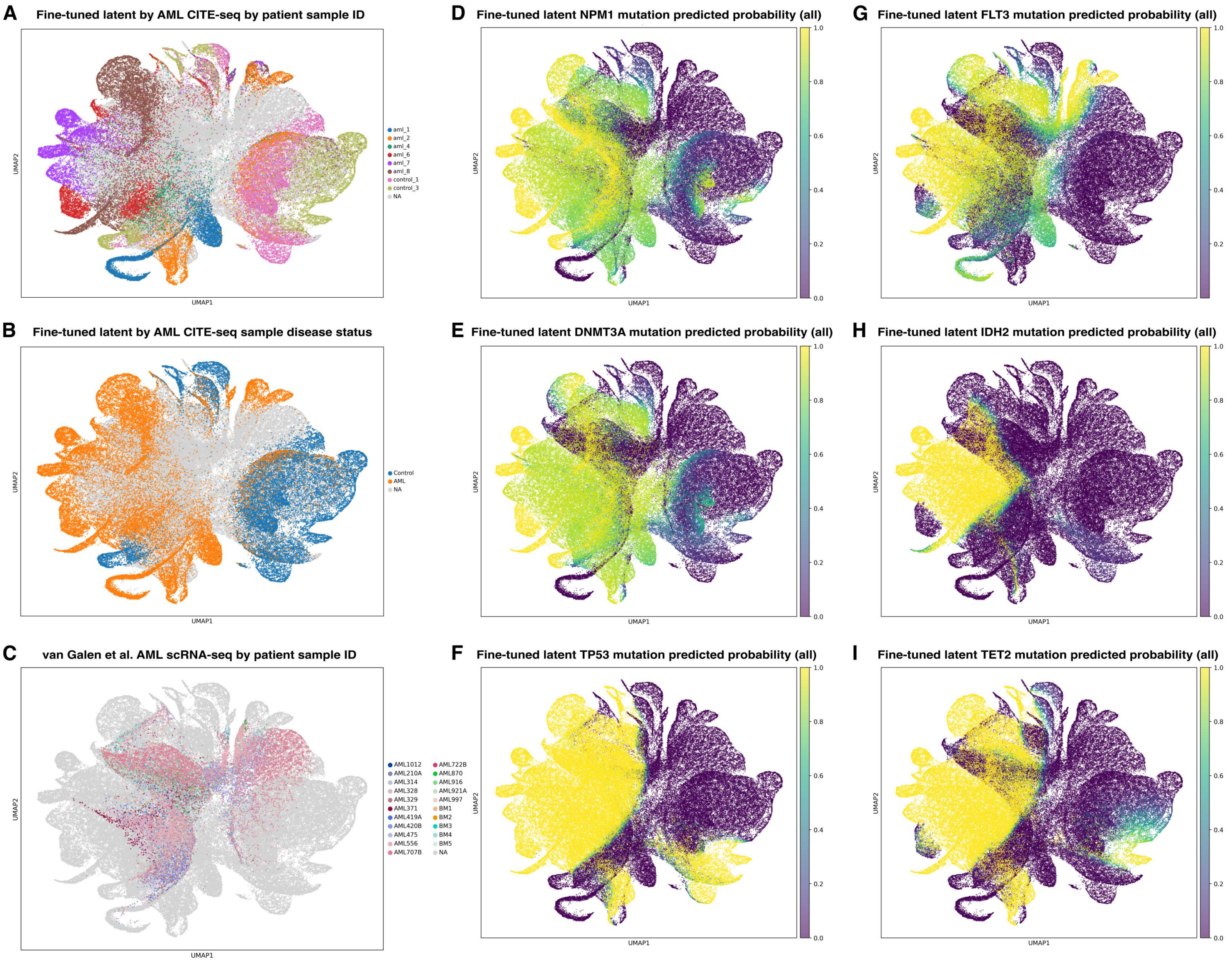


**Figure S7. Bridge cohort context and mutation-head grounding in the fine-tuned AML mosaic latent space (expands Fig. 7).** UniVI was trained on paired AML CITE-seq RNA+ADT cells as the bridge cohort and then optionally fine-tuned with auxiliary mutation-classification heads, yielding a joint latent space in which genotype-associated structure can be visualized across cohorts. (A) Fine-tuned latent UMAP with AML CITE-seq bridge patients colored by `CITE_sample_id` (other cohorts in light gray) to show where paired, mutation-supervised anchors lie in the manifold. (B) Same embedding colored by CITE-seq disease status (AML vs control) to contextualize bridge composition. (C) van Galen scRNA-seq cohort colored by patient/sample ID (other cohorts in light gray), highlighting patient-level heterogeneity in the unimodal RNA cohort relative to the bridge-anchored geometry. (D–I) Predicted mutation probabilities from the fine-tuned latent mutation heads over all embedded cells for representative genes spanning common AML drivers and bridge-supervision targets: (D) *NPM1*, (E) *DNMT3A*, (F) *TP53*, (G) *FLT3*, (H) *IDH2*, and (I) *TET2*. High-probability regions indicate latent neighborhoods whose shared multimodal context supports consistent genotype predictions, demonstrating that mutation-aware refinement can ground portions of the shared manifold in clinically relevant genetic programs without requiring that all cohorts share genotype measurements.

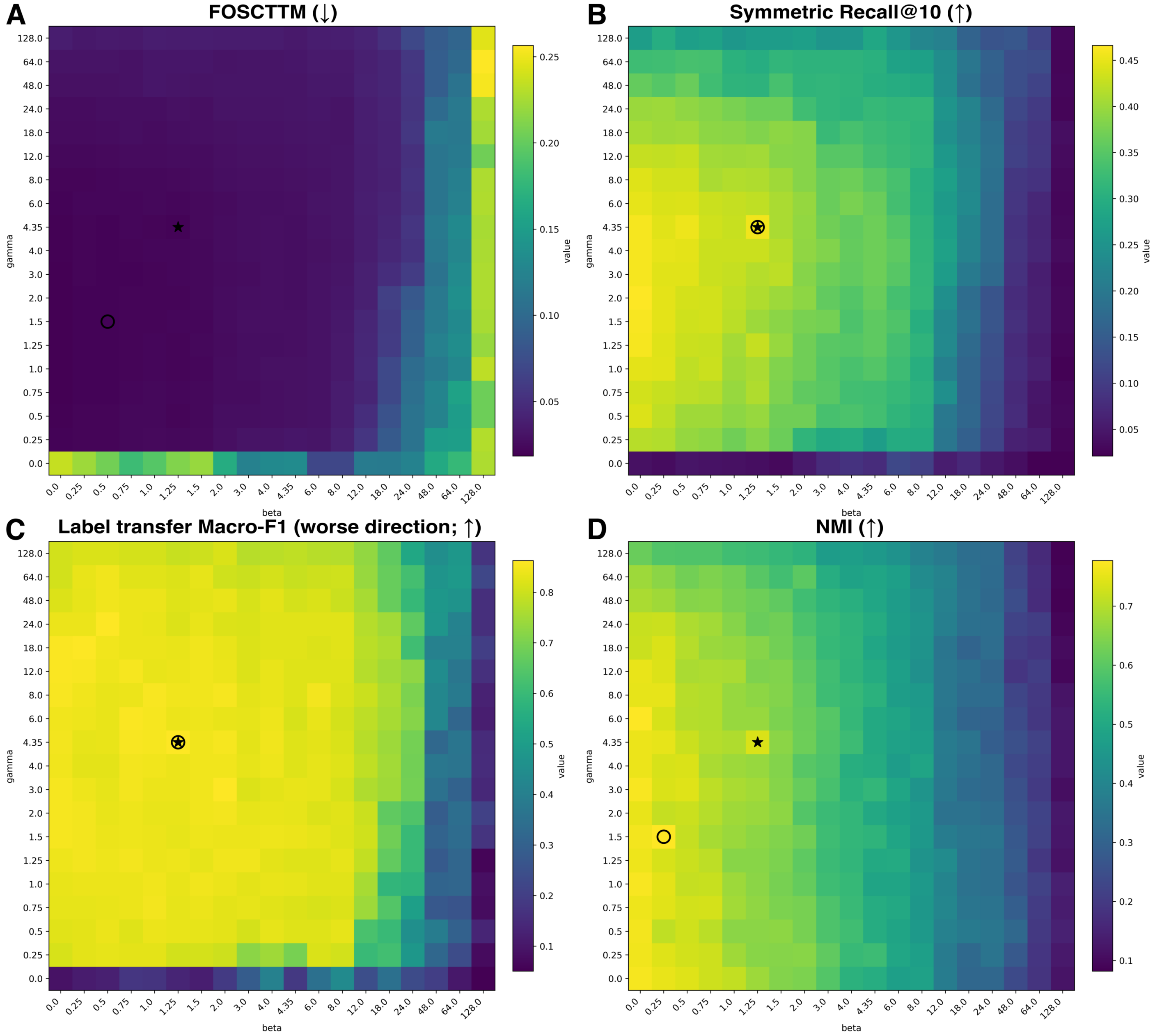


**Figure S8. Loss-weight sensitivity over $(\beta, \gamma)$ reveals a broad stable operating region with metric-specific optima.** Paired 10x Multiome PBMCs were used to sweep the KL weight $\beta$ and the cross-modal posterior coupling weight $\gamma$ in UniVI (`loss_mode="v1"`; Methods). Each panel summarizes a different evaluation objective computed on held-out paired test embeddings: (A) FOSCTTM (lower is better), (B) symmetric Recall@10 (higher is better), (C) bidirectional $k$-NN label-transfer macro-F1 (higher is better; worst-direction summary), and (D) fused-space NMI (higher is better). Circles indicate the best-scoring non-degenerate configuration for the corresponding metric (excluding $\beta = 0$ and $\gamma = 0$), highlighting expected trade-offs between strict paired retrieval, semantic consistency, and global structure rather than a single tuned corner. Stars mark the manuscript default configuration (for Multiome data).

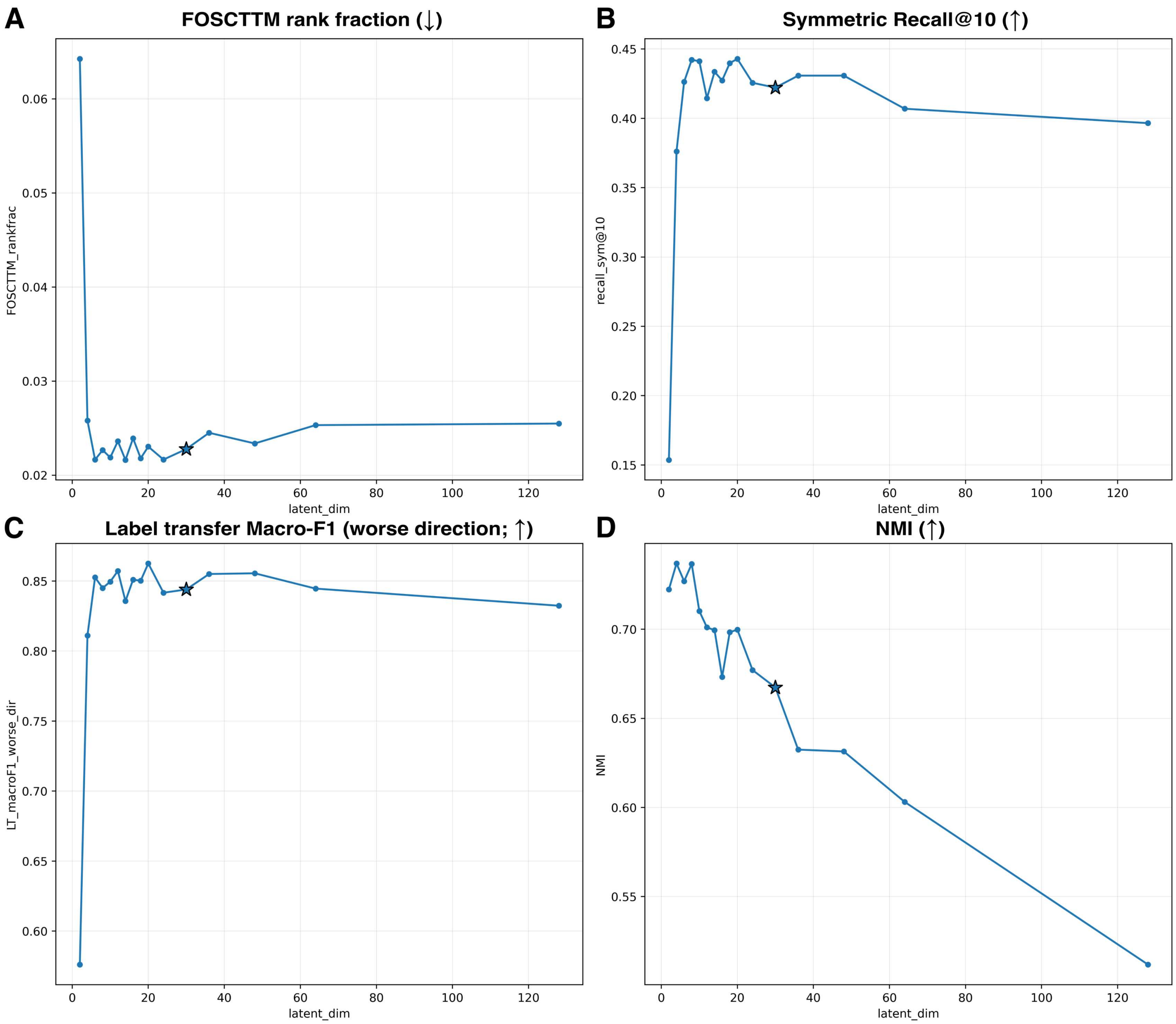


**Figure S9. Latent dimensionality sensitivity in paired Multiome PBMCs.** We swept the UniVI latent dimensionality while holding all other preprocessing and training settings fixed, and evaluated performance on held-out paired test embeddings. Panels report complementary objectives: (A) paired correspondence via FOSCTTM rank fraction (lower is better), (B) strict paired retrieval via symmetric Recall@10 (higher is better), (C) cross-modality semantic consistency via label-transfer macro-F1 in the worse transfer direction (higher is better), and (D) fused-space biological structure via $k$-means NMI against curated labels (higher is better). The star marks the manuscript default latent dimension (for Multiome data).

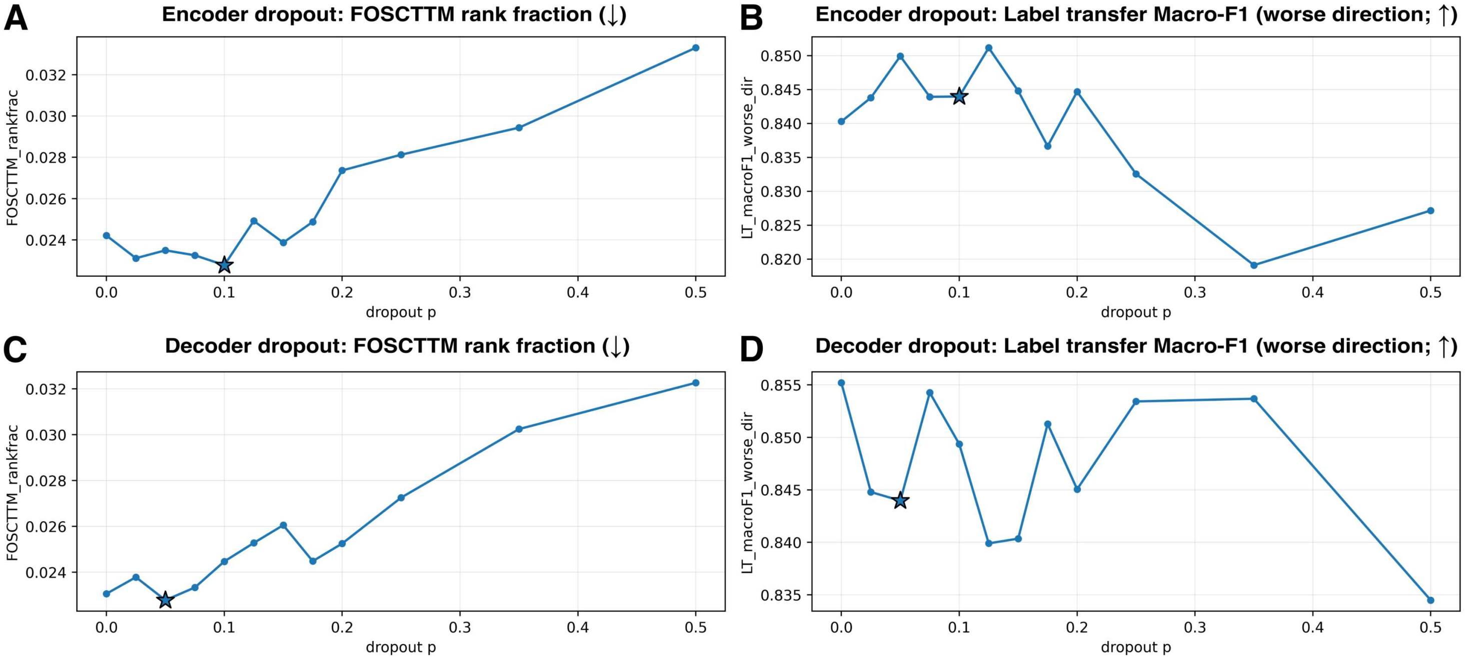


**Figure S10. Encoder/decoder dropout sensitivity reveals smooth correspondence–semantics trade-offs.** We swept dropout probability $p$ in the encoder (top row) or decoder (bottom row) while holding all other settings fixed, and evaluated on held-out paired test embeddings. (A) Encoder dropout: correspondence measured by FOSCTTM rank fraction (lower is better). (B) Encoder dropout: semantic consistency measured by cross-modality label-transfer macro-F1 in the worse transfer direction (higher is better). (C) Decoder dropout: correspondence measured by FOSCTTM rank fraction (lower is better). (D) Decoder dropout: semantic consistency measured by cross-modality label-transfer macro-F1 in the worse transfer direction (higher is better). Stars denote the manuscript default dropout settings (encoder $p = 0.10$ in A–B; decoder $p = 0.05$ in C–D).

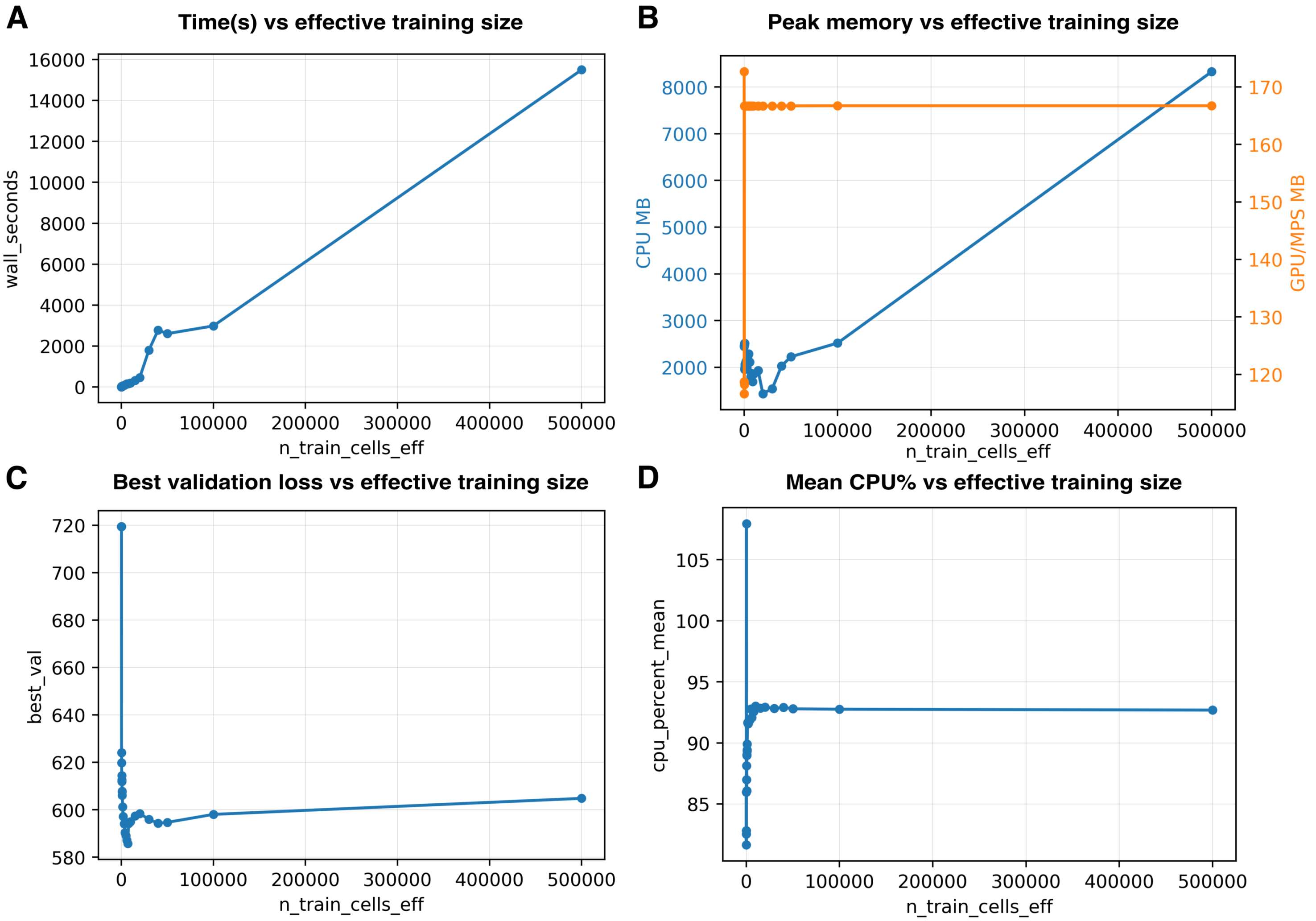


**Figure S11. Computational scaling with effective training set size.** We profiled UniVI training as a function of effective training size by resampling from a fixed underlying paired Multiome training set (Methods), isolating computational scaling from gains due to additional unique observations. Panels report (A) wall-clock training time, (B) peak memory usage (CPU RSS and accelerator peak when available), (C) best validation loss achieved during early stopping, and (D) mean CPU utilization.

### Supplemental Tables

**Table S1. AML CITE-seq bridge cohort (paired RNA+ADT) used to anchor the AML mosaic integration.** Samples provide paired RNA–ADT anchors for bridge training/projection (Methods). Mutation-head fine-tuning uses DAb-seq and van Galen labels; this table is included to interpret patient-local structure in the fine-tuned latent space. ELN: F=favorable, I=intermediate, A=adverse.

| Sample | Age | Sex | ELN | Karyotype | Tx | Key genes* |
|---|---|---|---|---|---|---|
| aml1 | 64 | F | – | Normal | 7+3 | – |
| aml2 | 60 | M | A | Complex | decitabine | *SRSF2*, *TET2*, *TP53*, *RAD21*, *GATA2* |
| aml3 | 75 | M | A | Complex | 7+3 | *TET2*, *ASXL1* |
| aml4 | 73 | M | I | Unclassified | 7+3 | *KRAS*, *NPM1* |
| aml5 | 64 | M | I | Unclassified | 7+7 + midostaurin | *DNMT3A*, *NPM1*, *FLT3*, *TET2* |
| aml6 | 44 | F | F | +8 | 7+3 + crenolanib | *DNMT3A*, *NPM1* |
| aml7 | 54 | M | I | Normal | 7+3 | *TET2* |
| aml8 | 83 | M | F | inv(16), -7q | decitabine | *KIT*, *FLT3*, *DNMT3A* |

*Gene-level alterations shown only for drivers/head targets referenced in Results/figures. Variant strings omitted for readability. *Source:* condensed from Knorr et al. supplementary patient summary.

**Table S2. Demaree et al. DAb-seq AML cohort summary (protein+genotype).** DAb-seq provides dense cell-level genotype labels used for mutation overlays/evaluation in Fig. 7.

| Patient | Clinical | Karyotype / WHO | ELN | Genotype notes (targets/labels) |
|---|---|---|---|---|
| Patient #1 | Gemtuzumab treated | Normal; acute mono/ monocytic leukemia | Intermediate (presumed) | *NPM1* mut; *FLT3*-ITD by PCR; *CEBPA* PCR negative; no panel reported. |
| Patient #2 | Pediatric | 47,XY; *MLL* rearrangement (11q23) | – | qPCR positive *FLT3* KD (D835/I836); no panel reported. |
| Patient #3 | FLT3 inhibitor (gilteritinib) | 46,XY; AML with mutated *NPM1* | Intermediate | Panel at diagnosis: *NPM1* (38.7%), *DNMT3A* (41.5%), *IDH2* (44.9%) VAF. |

*Source:* condensed from Demaree et al. supplementary patient/treatment summary.

**Table S3. van Galen AML scRNA-seq cohort (collapsed by sample; timepoints removed).** Key alterations list unique gene symbols parsed from the original annotations (variant strings omitted). "–" denotes not reported in the original supplement.

| Sample | Tissue | Sex | Age | Blasts | Key genes | Cytogenetics | Tx | Cells |
|---|---|---|---|---|---|---|---|---|
| BM1 | BM | M | 52 | – | – | – | – | 108 |
| BM2 | BM | M | 21 | – | – | – | – | 188 |
| BM3 | BM | M | 56 | – | – | – | – | 643 |
| BM4 | BM | M | 23 | – | – | – | – | 3738 |
| BM5 CD34+ | BM | M | 45 | – | – | – | – | 1431 |
| BM5 CD34+CD38- | BM | M | 45 | – | – | – | – | 1590 |
| AML1012 | BM | F | 32 | 65% | *KRAS, NRAS, NOTCH2, SF3A1* | inv(16); +8/+21 | *CBFB-MYH11* | 1136 |
| AML210A | BM | M | 67 | 83% | *DNMT3A, NPM1, TET2, FLT3* | 46,XY | – | 748 |
| AML419A | BM | F | 54 | 60% | *CEBPA, DNMT3A, NPM1, FLT3, JAK3* | 46,XX | – | 1189 |
| AML916 | BM | F | 57 | 75% | *TP53* | 46,XX | – | 933 |
| AML921A | BM | M | 42 | 70% | *DNMT3A, RUNX1, SETD2* | 46,XY | – | 3813 |
| AML314 | BM | M | 54 | 28% | *BCOR, RUNX1* | 46,XY | – | 162 |
| AML371 | BM | M | 51 | 16% | *NRAS, WT1* | der(16)/t(16;18) | – | 756 |
| AML475 | BM | M | 70 | 1% | *DNMT3A, BCOR, BCORL1* | 46,XY | – | 423 |
| AML722B | BM | F | 52 | 84% | *IDH2, BCORL1, ASXL1, PHF6, PTPN11* | i(7); +8 | – | 79 |
| AML870 | BM | M | 32 | 89% | *ZRSR2* | t(9;11) | *MLL-** | 345 |
| AML997 | BM | M | 62 | 16% | *DNMT3A, NPM1, CEBPA, FLT3* | 46,XY | – | 83 |
| AML329 | BM | F | 73 | 37% | *NPM1, FLT3, NOTCH1, SMC3* | 46,XX | – | 525 |
| AML420B | BM | M | 58 | 29% | *IDH2, TP53, SH2B3* | add(1p36) mosaic | – | 485 |
| AML556 | BM | M | 70 | 79% | *DNMT3A, NPM1, NRAS, TET2, ATM* | 46,XY | – | 2328 |
| AML328 | BM | F | 74 | 55% | *DNMT3A, TP53, FLT3* | inv(3)/-7 complex | – | 1094 |
| AML707B | BM | M | 26 | 76% | *BRCC3, KIT, RAD21* | t(8;21) | *RUNX1-RUNX1T1* | 1586 |
| OCI-AML3 | Cell line | M | 57 | – | *DNMT3A, NPM1, NRAS, RAD21, SMC3, BCORL1, ATM, SETD2* | hyperdiploid | – | 1178 |
| MUTZ3 | Cell line | M | 29 | – | *SF3B1, KRAS, ASXL1, GATA2, IKZF1* | near-diploid | – | 1502 |

*Source:* condensed from van Galen et al. supplementary patient mutation summary.

**Table S4. Targeted mutation detection summary (collapsed by sample; timepoints removed).** Rows are collapsed to unique (*Sample, Mutation, sequencing method*) combinations across all assayed timepoints. Reported totals sum transcript counts and genotyped cells over timepoints for the same sample and mutation. **Total_cells** denotes the summed number of profiled cells across timepoints for that sample in this targeted assay panel. Primer names are listed as unique identifiers used for each target.

| Sample | Mutation | Method | WT | Mut | Genotyped | Total_cells | Frac. | Primer(s) |
|---|---|---|---|---|---|---|---|---|
| AML1012 | *NRAS*.G13D | Illumina | 13 | 9 | 22 | 1136 | 1.9% | PvG1067-Next_NRAS_11 |
| AML210A | *DNMT3A*.R882C | Illumina | 6 | 4 | 9 | 748 | 1.2% | PvG1060-Next_DNMT3A_2623 |
| AML210A | *FLT3*.ITD | Illumina | 0 | 0 | 0 | 748 | 0.0% | PvG1073-Next_FLT3_1761 |
| AML210A | *NPM1*.W288fs | Illumina | 71 | 45 | 92 | 748 | 12.3% | PvG1066-Next_NPM1_833 |
| AML210A | *TET2*.S358G | Illumina | 0 | 0 | 0 | 748 | 0.0% | Cannot amplify: 8,221 bp from polyA site |
| AML328 | *DNMT3A*.L637Q | Illumina | 8 | 7 | 15 | 6405 | 0.2% | PvG1071-Next_DNMT3A_1885 |
| AML328 | *FLT3*.ITD | Illumina | 0 | 0 | 0 | 6405 | 0.0% | PvG1072-Next_FLT3_1711 |
| AML328 | *FLT3*.ITD | Oxford nanopore | 20 | 13 | 33 | 1094 | 3.0% | PvG1072-Next_FLT3_1711 |
| AML328 | *TP53*.Q144P/P152R | Illumina | 7 | 28 | 35 | 6405 | 0.5% | PvG1082-Next_TP53_405 |
| AML328 | *TP53*.Q144P/P152R | Oxford nanopore | 24 | 73 | 97 | 6405 | 1.5% | PvG1082-Next_TP53_405 |
| AML329 | *FLT3*.ITD | Illumina | 24 | 12 | 36 | 1702 | 2.1% | PvG1061-Next_FLT3_1740 |
| AML329 | *NPM1*.W288fs | Illumina | 416 | 206 | 295 | 1702 | 17.3% | PvG1066-Next_NPM1_833 |
| AML371 | *NRAS*.Q61K | Illumina | 8 | 6 | 14 | 960 | 1.5% | PvG1068-Next_NRAS_151 |
| AML371 | *WT1*.T309fs | Illumina | 0 | 0 | 0 | 960 | 0.0% | PvG1125-Next_WT1_1093 |
| AML419A | *DNMT3A*.R882C | Illumina | 22 | 18 | 38 | 1189 | 3.2% | PvG1060-Next_DNMT3A_2623 |
| AML419A | *FLT3*.A680V | Illumina | 56 | 15 | 67 | 1189 | 5.6% | PvG1075-Next_FLT3_2019 |
| AML419A | *FLT3*.A680V/ N841K/ITD | Oxford nanopore | 15 | 10 | 25 | 1189 | 2.1% | PvG1168-Next_FLT3_1322 |
| AML419A | *FLT3*.ITD | Illumina | 17 | 7 | 24 | 1189 | 2.0% | PvG1073-Next_FLT3_1761 |
| AML419A | *FLT3*.N841K | Illumina | 52 | 14 | 64 | 1189 | 5.4% | PvG1062-Next_FLT3_2482 |
| AML419A | *NPM1*.W288fs | Illumina | 148 | 116 | 210 | 1189 | 17.7% | PvG1066-Next_NPM1_833 |
| AML420B | *IDH2*.R140Q | Illumina | 49 | 3 | 51 | 2510 | 2.0% | PvG1064-Next_IDH2_392 |
| AML420B | *TP53*.R273L | Illumina | 20 | 3 | 22 | 2510 | 0.9% | PvG1114-Next_TP53_794 |
| AML475 | *DNMT3A*.R882H | Illumina | 9 | 6 | 13 | 525 | 2.5% | PvG1060-Next_DNMT3A_2623 |
| AML556 | *DNMT3A*.R882C | Illumina | 96 | 22 | 102 | 4982 | 2.0% | PvG1060-Next_DNMT3A_2623 |
| AML556 | *NPM1*.W288fs | Illumina | 1862 | 382 | 1411 | 4982 | 28.3% | PvG1066-Next_NPM1_833 |
| AML556 | *NRAS*.G12D | Illumina | 98 | 22 | 117 | 4982 | 2.3% | PvG1067-Next_NRAS_11 |
| AML556 | *NRAS*.Q61H | Illumina | 98 | 22 | 117 | 4982 | 2.3% | PvG1068-Next_NRAS_151 |
| AML556 | *TET2*.L1804fs | Illumina | 10 | 6 | 16 | 4982 | 0.3% | PvG1112-Next_TET2_5363 |
| AML556 | *TET2*.S1059stp | Illumina | 22 | 13 | 35 | 4982 | 0.7% | PvG1111-Next_TET2_3139 |
| AML722B | *IDH2*.R172K | Illumina | 2 | 0 | 2 | 152 | 1.3% | PvG1065-Next_IDH2_491 |
| AML916 | *TP53*.C238Y | Illumina | 0 | 22 | 21 | 933 | 2.3% | PvG1091-Next_TP53_683 |
| AML921A | *DNMT3A*.R882H | Illumina | 135 | 105 | 210 | 3813 | 5.5% | PvG1060-Next_DNMT3A_2623 |
| AML997 | *DNMT3A*.R882H | Illumina | 3 | 0 | 3 | 270 | 1.1% | PvG1060-Next_DNMT3A_2623 |
| AML997 | *FLT3*.ITD | Illumina | 0 | 0 | 0 | 270 | 0.0% | Cannot design primer: insufficient information |
| AML997 | *NPM1*.W288fs | Illumina | 63 | 7 | 50 | 270 | 18.5% | PvG1066-Next_NPM1_833 |

*Source:* condensed from van Galen et al. supplementary patient mutation summary.

**Table S5. AML mosaic cohorts, modalities, and mutation-label availability.** Bridge training uses paired AML CITE-seq (RNA+ADT); van Galen (RNA-only) and DAb-seq (ADT+genotype) are projected via encoder inference (Methods).

| Cohort | Modalities | Bridge | Proj. | Dense geno | Notes / typical label use |
|---|---|---|---|---|---|
| Knorr et al. (2023) AML CITE-seq | RNA+ADT | ✓ | – | – | Paired anchors; patient context in Supp. Table S1. |
| Demaree et al. (2021) DAb-seq | ADT+ genotype | – | ✓ | ✓ | Primary source of mutation labels for overlays and head evaluation; patient context in Supp. Table S2. |
| Galen et al. (2019) scRNA-seq | RNA+ genotype | – | ✓ | – | Sparse/incomplete mutation annotations; used when labels exist and for neighborhood transfer; patient context in Supp. Tables S3 and S4. |

**Table S6. Benchmark methods evaluated in Fig. 8, with method family and evaluation regime in our unified runner. Inductive:** fit on train split; apply to val/test by forward inference or a fixed transform (no refit). **Transductive:** standard workflow fits jointly on all cells supplied to the runner; held-out cells can influence the learned representation. Regime labels reflect *our runner defaults* (not all possible out-of-sample extensions). **Prior information:** external side inputs beyond assay matrices (e.g., labels, batch covariates, feature correspondences, graphs, or paired cells).

| Method | Reference | Method family | Runner | Regime | Notes (as evaluated in our pipeline) |
|---|---|---|---|---|---|
| UniVI | This work | Deep generative (MoE $\beta$-VAE; multimodal latent) | PyTorch | Inductive | Trained on train split only; val/test embedded by parameter-frozen encoder inference (no parameter updates). **Prior information:** none required beyond assay matrices. |
| scGLUE | Cao and Gao (2022) | Graph-linked latent alignment / manifold coupling | PyTorch | Inductive | Fit on train split; held-out cells embedded via learned encoders (no retraining / no re-graphing on test). **Prior information:** guidance (regulatory) graph linking features across modalities (e.g., peak$\longrightarrow$gene / TF-target edges). |
| scJoint | Lin et al. (2022) | Neural alignment / supervised alignment | PyTorch | Inductive | Fit on train split; held-out cells embedded by forward pass only (no adaptation). **Prior information:** supervised labels (typically cell type labels on a reference modality) used to guide alignment. |
| CoBOLT | Gong et al. (2021) | Deep probabilistic latent model (multi-omics coupling) | PyTorch | Transductive | As run here, CoBOLT was fit jointly over the full evaluation set supplied to the runner (no strict train-only fit with test-only transform in the default workflow). **Prior information:** none required beyond assay labels. |
| MultiVI | Ashuach et al. (2023) | Deep generative (scVI family; RNA+ATAC) | PyTorch | Inductive | Fit on train split; held-out cells embedded by amortized inference (no refit / no joint optimization over test cells). **Prior information:** none required beyond assay matrices. |
| PeakVI | Ashuach et al. (2022) | Deep generative (scVI family; ATAC) | PyTorch | Inductive | ATAC-only VAE. Fit on train split; held-out cells embedded by amortized inference (no refit). **Prior information:** none required beyond assay matrices. |
| DeepCCA | Andrew et al. (2013) | Deep representation learning (CCA objective) | PyTorch | Inductive | Neural encoders trained on train split to maximize cross-view correlation; held-out cells embedded by forward pass only. **Prior information:** paired observations across modalities during training. |
| MultiMAP | Jain et al. (2021) | Manifold alignment (nearest-neighbor / graph-based) | PyTorch | Transductive | As run here, constructs a joint graph/embedding over all cells supplied for integration; we therefore provided the full evaluation set to the method (no strict train-only fit with test-only transform in the default workflow). **Prior information:** none external; relies on kNN graphs computed from the supplied representations (dependent on chosen features / PCs / LSI). |
| scMoMaT | R Zhang et al. (2023) | Matrix tri-factorization / latent factor fusion | PyTorch | Transductive | Joint factorization is optimized over all embedded cells in the standard workflow; our runner supplies all cells being evaluated (no default out-of-sample mapping used). **Prior information:** none required beyond assay matrices. |
| Harmony | Korsunsky et al. (2019) | Batch correction / embedding harmonization | R | Transductive | No standard “fit on train, transform test” interface; harmonization is performed on the embedding of all supplied cells, so our runner supplies the full evaluation set. **Prior information:** batch labels (or grouping variable) to define nuisance structure. |
| Seurat WNN | Hao et al. (2021) | Graph / neighbor fusion (weighted nearest neighbors) | R | Transductive | WNN graph construction and smoothing are computed on all supplied cells; our runner supplies the full evaluation set (no strict held-out inference in the default workflow). **Prior information:** modality-specific reduced spaces as inputs (e.g., PCA for RNA/ADT; LSI for ATAC). |
| Seurat CCA | Stuart et al. (2019) | Anchor-based integration / CCA alignment | R | Transductive | Anchors/CCA are fit over the cells supplied for integration in the default workflow; our runner supplies the full evaluation set to match standard usage. **Prior information:** a shared feature space or defined feature correspondences used for CCA/anchors. |
| Seurat Bridge | Hao et al. (2023) | Reference mapping / anchor transfer (bridge integration) | R | Inductive | Implemented as Seurat reference mapping: train split defines the reference; held-out cells are mapped to the fixed reference (no recomputation of anchors on test). **Prior information:** a pre-defined reference (and bridge if used) plus consistent feature mapping into the reference space. |
| LIGER | Welch et al. (2019) | Matrix factorization (iNMF) | R | Transductive | Standard iNMF is fit jointly across supplied cells; our runner supplies the full evaluation set (no default out-of-sample transform used). **Prior information:** matched features across datasets/modalities (or a defined shared feature subset). |
| MOFA2 | Argelaguet et al. (2020) | Factor analysis (multi-view latent factors) | R | Transductive | Factors are learned from the supplied dataset in our runner; we fit MOFA2 on all evaluated cells (no out-of-sample factor inference used). **Prior information:** none required beyond assay matrices. |

**Runner caveat:** Transductive methods are run with the full evaluation set supplied to the workflow; inductive methods apply train-fit parameters unchanged to val/test (Methods).

**Table S7. Datasets, splits, and preprocessing used across main + supplemental analyses in this manuscript.** Parameter-learning transforms (HVG selection, scaling, TF–IDF/LSI, standardization) are fit on train and applied unchanged to val/test and projected cohorts.

| Col | Scope | Dataset | Accession / source | Modalities | Train/Val/Test | Stratify / caps | Preprocessing | Drop LSI1 |
|---|---|---|---|---|---|---|---|---|
| **A** | Figs. 2–3 + Supp. Fig. S1 | Hao et al. (2021) (PBMC CITE-seq) | GEO: `GSE164378` | RNA+ADT | 43,927/5,478/ 112,359 | `celltype.l3` cap 1200/150 otherwise 80/10/10 +capped extras | **RNA:** HVG 2000; LSN $10^4$+log1p; $z$(train-fit)<br>**ADT:** 228 ADTs; CLR(cell); $z$(train-fit) | – |
| **B** | Fig. 4 + Supp. Fig. S2 | 10x Genomics (2021a) (PBMC Multiome) | 10k Human PBMCs, Multiome v1.0, Chromium X | RNA+ATAC | 5,777/717/3,137 | `cell_type` cap 800/100 otherwise 80/10/10 +capped extras | **RNA:** HVG 2000; LSN $10^4$+log1p; $z$(train-fit)<br>**ATAC:** 107,194 peaks; TF–IDF; LSI(101)*; (train-fit) | No |
| **C** | Fig. 5 + Supp. Fig. S5 | 10x Genomics (2021b) (PBMC Multiome) | PBMC from a Healthy Donor – No Cell Sorting (10k) | RNA+ATAC | 10,810/1,201/– | 90/10/0 | **RNA:** HVG 2000 (ref); LSN $10^4$+log1p; $z$(train-fit)<br>**ATAC:** 85,485 peaks; TF–IDF; LSI(101)*; (train-fit) | No |
| **D** | Fig. 5 + Supp. Fig. S5 | Ding et al. (2020) (PBMC scRNA-seq) | GEO: `GSE132044` | RNA | –/–/30,495 | 0/0/100 (query projection) | **RNA:** overlap genes (Multiome∩Ding); HVGs in overlap; LSN $10^4$+log1p; $z$(train-fit) | – |
| **E** | Fig. 5 + Supp. Fig. S5 | Satpathy et al. (2019) (PBMC scATAC-seq) | GEO: `GSE129785` | ATAC | –/–/47,148 | 0/0/100 (query projection) | **ATAC:** 85,485 peaks; TF–IDF; LSI(101)*; $z$(train-fit) | No |
| **F** | Fig. 6 + Supp. Fig. S6 | Swanson et al. (2021) (PBMC TEA-seq) | GEO: `GSE158013` | RNA+ADT+ ATAC | 17,978/2,247/ 9,669 | 80/10/10+well holdout (train wells 3–4, 6; eval well 5) | **RNA:** HVG 2000; LSN $10^4$+log1p; $z$(train-fit)<br>**ADT:** 47 ADTs; CLR(cell); $z$(train-fit)<br>**ATAC:** 155,930 tiles; TF–IDF; LSI(101)*; $z$(train-fit) | No |
| **G** | Fig. 7 + Supp. Fig. S7 | Knorr et al. (2023) (AML CITE-seq) | GEO: `GSE220474` | RNA+ADT | 28,414/4,652/ 10,113 | `sample_id` group holdout (8/1/2 samples) | **RNA:** 21,782 genes; LSN $10^4$+log1p; $z$(train-fit)<br>**ADT:** 18 shared markers; CLR(cell); $z$(train-fit); clip $[-10, 10]$ | – |
| **H** | Fig. 7 + Supp. Fig. S7 | Galen et al. (2019) (AML scRNA-seq) | GEO: `GSE116256` | RNA+geno | –/–/37,627 | 0/0/100 (query projection) | **RNA:** apply bridge RNA transform | – |
| **I** | Fig. 7 + Supp. Fig. S7 | Demaree et al. (2021) (DAb-seq) | NCBI BioProject: `PRJNA602320` | ADT+geno | –/–/40,239 | 0/0/100 (query projection) | **ADT:** apply bridge ADT transform | – |
| **J** | Figs. 8–9 | 10x Genomics (2021a) (PBMC Multiome) | 10k Human PBMCs, Multiome v1.0, Chromium X | RNA+ATAC | 7,704/963/964† | 80/10/10; repeated CV† | **RNA:** HVG 2000; LSN $10^4$+log1p; $z$(train-fit)<br>**ATAC:** 107,194 peaks; TF–IDF; LSI(101)*; (train-fit)‡ | Yes‡ |
| **K** | Fig. 10 + Supp. Figs. S8–S11 | 10x Genomics (2021a) (PBMC Multiome) | 10k Human PBMCs, Multiome v1.0, Chromium X | RNA+ATAC | 7,223/963/ 1,445† | 75/10/15; repeated CV† | **RNA:** HVG 2000; LSN $10^4$+log1p; $z$(train-fit)<br>**ATAC:** 107,194 peaks; TF–IDF; LSI(101)*; (train-fit)‡ | Yes‡ |
| **L** | Supp. Fig. S3 | Ma et al. (2020) (Mouse skin SHARE-seq) | GEO: `GSE140203`; sample `GSM4156597` | RNA+ATAC | 24,961/3,120/ 3,139 | `cell_type` 80/10/10 stratified; `Mix` excluded | **RNA:** HVG 5000 (`seurat_v3`; *mt-*/*Rps*/*Rpl*/*Mrps*/*Mrpl*/*Rik*/-*ps* excluded); LSN $10^4$+log1p; $z$(train-fit)<br>**ATAC:** 233,589 peaks; ENCODE mm10 blacklist; log-TF–IDF; LSI(101)*; $z$(train-fit) | Yes |
| **M** | Supp. Fig. S4 | Argelaguet et al. (2019) (Mouse gastrulation scNMT-seq) | GEO: `GSE121708`§ | RNA+CpG+ GpC | 969/57/114 | random 85/5/10; inductive + transductive metrics reported | **RNA:** LSN $10^4$+log1p (no HVG; no $z$-scoring)<br>**CpG:** genebody; beta-binomial on (successes, coverage); parsed-bundle defaults<br>**GpC:** genebody; beta-binomial on (successes, coverage); parsed-bundle defaults | No |

**Abbreviations:** LSN = library-size normalization (target sum $10^4$ counts/cell); HVG = highly variable genes (SCANPY `flavor="seurat_v3"` where applicable); CLR = centered-log-ratio normalization; $z$(train-fit) = $z$-scoring with mean/std fit on training cells and applied unchanged to validation/test; LSI = latent semantic indexing (truncated SVD of TF–IDF representation).

**Footnotes:**

* For ATAC TF–IDF + LSI pipelines, LSI component 0 (depth-confounded) is dropped from downstream model input; the value in parentheses indicates the number of LSI components fit ($n$) before this drop, leaving $n - 1$ components used as input.

† Counts shown are per-fold averages; benchmarking and ablation analyses used repeated cross-validation (Methods).

‡ Preprocessing transforms refit per fold; cross-validation enabled for these analyses.

§ The raw data are deposited at `GSE121708`; analyses used the parsed feature-level bundle distributed via the EBI FTP server (`ftp://ftp.ebi.ac.uk/pub/databases/scnmt_gastrulation/scnmt_gastrulation.tar.gz`).

**Table S8. UniVI training + fine-tuning hyperparameters used in this manuscript.** ES = early stopping on validation loss; schedules are epoch ranges: KL(a–b), Align(a–b). All models used `loss_mode="v1"`, `v1_recon="avg"`, and `normalize_v1_terms=True`. Learned MoE gating was enabled only for selected fused-latent analyses and did not alter the reconstruction objective.

| Col | Scope | Modalities | Batch | Latent dim | Symm hidden dims | $\beta/\gamma$ | Drop (enc/dec) | Epochs | LR / WD | Device | Schedules / ES |
|---|---|---|---|---|---|---|---|---|---|---|---|
| **A** | Figs. 2–3 + Supp. Fig. S1 | rna+adt | 256 | 30 | rna: [512,256,128]<br>adt: [128,64] | 1.5/5.0 | 0.10/0.00 | 3000 | $10^{-3}/10^{-4}$ | cuda | loss=v1; BN enc yes/dec no; KL(0–0), Align(0–0); ES pat=50/100 |
| **B** | Fig. 4 + Supp. Fig. S2 | rna+atac | 128 | 30 | rna: [512,256,128]<br>atac:[128,64] | 1.25/4.35 | 0.10/0.05 | 5000 | $10^{-3}/10^{-4}$ | cuda | loss=v1; BN enc yes/dec no;KL(50–85), Align(75–110); ES pat=300 |
| **C** | Fig. 5 + Supp. Fig. S5 | rna+atac | 256 | 30 | rna: [512,256,128]<br>atac:[128,64] | 1.0/5.0 | 0.25/0.05 | 5000 | $10^{-3}/10^{-4}$ | cuda | `head_then_joint`;decoders frozen;warmup=300, n=3000; LR(bb)=$10^{-5}$;LR(head)=$3 \times 10^{-4}$ |
| **D** | Fig. 6 + Supp. Fig. S6 | rna+adt+atac | 256 | 30 | rna: [512,256,128]<br>adt: [128,64]<br>atac:[128,64] | 1.15/1.35 | 0.10/0.00 | 3000 | $10^{-4}/10^{-4}$ | cuda | loss=v1;KL(0–50), Align(25–75);ES pat=200 |
| **E** | Fig. 7 + Supp. Fig. S7 | rna+adt | 256 | 30 | rna: [1024,512,256,128,64]<br>adt: [128,64,32] | 1.15/1.75 | 0.10/0.05 | 3000 | $10^{-4}/10^{-5}$ | cuda | 7 gene heads; warmup$\rightarrow$ unfreeze; decoders frozen; $\lambda$ MSE$(z_s, z_t)$ |
| **F** | Fig. 8 | rna+atac | 256 | 30 | rna: [512,256,128]<br>atac:[256,128,64] | 1.35/3.75 | 0.10/0.05 | 200 | $10^{-3}/10^{-4}$ | cuda | benchmark runner; KL(25–75), Align(45–95); ES pat=50; multi-seed |
| **G** | Fig. 9 | rna+atac | 256 | 30 | rna: [512,256,128]<br>atac:[128,64] | 1.25/4.35 | 0.10/0.05 | 5000 | $10^{-3}/10^{-4}$ | mps | overlap sweep;KL(0–60), Align(25–85);ES pat=100 |
| **H** | Fig. 10 | rna+atac | 128 | 30 | rna: [512,256,128]<br>atac:[128,64] | 1.25/4.35 | 0.10/0.05 | 5000 | $10^{-3}/10^{-4}$ | mps | ablations/sweeps;KL(50–100), Align(75–125);ES pat=50 |
| **I** | Supp. Fig. S3 | rna+atac | 128 | 30 | rna: [1024,512,256,128]<br>atac:[256,128,64] | 1.25/6.35 | 0.10/0.00 | 5000 | $10^{-3}/10^{-4}$ | mps | KL(50–85), Align(75–110);ES pat=200 |
| **J** | Supp. Fig. S4 | rna+cpg+gpc | 24 | 20 | rna: [512,256,128,64]<br>cpg: [256,128,64]<br>gpc: [256,128,64] | 1.35/6.25 | 0.010/0.005 | 5000 | $10^{-3}/10^{-4}$ | mps | KL(30–60), Align(40–70);ES pat=300 |
| **K** | Supp. Figs. S8– S11 | rna+atac | 128 | 30 | rna: [512,256,128]<br>atac:[128,64] | 1.25/4.35 | 0.10/0.05 | 5000 | $10^{-3}/10^{-4}$ | mps | ablations/sweeps;KL(50–100), Align(75–125);ES pat=50 |

### Supplemental Methods

This document records implementation details, dataset-specific preprocessing protocols, regime-specific deviations from the default pipeline, hyperparameter selection procedures, and stress-test configurations referenced from the main Methods. Complete numerical settings (per-experiment widths, depths, learning rates, annealing schedules, and seeds) are provided in Supp. Tables S7–S8.

**Shared training-objective configuration.** All models reported in this study were trained using `loss_mode="v1"`, `v1_recon="avg"`, and `normalize_v1_terms=True`. Thus, training used the balanced self/cross reconstruction, normalized KL-to-prior, and normalized directed posterior-alignment terms defined in the main Methods. MoE fusion was used separately to construct selected fused latent spaces. Where learned MoE gating was enabled, router probabilities modulated the modality posterior precisions before analytic Gaussian fusion; otherwise, fusion used posterior precision alone. No reported model used fused-posterior reconstruction during training.

#### Training schedule details

**Warmup and annealing.** KL regularization ($\beta$) and cross-modality alignment ($\gamma$) terms were introduced gradually via epoch-based annealing schedules to stabilize the early phase of training and avoid posterior collapse. Per-experiment annealing schedules (start epoch, end epoch, schedule shape) are recorded in Supp. Table S8.

#### Supervised refinement: head architectures and schedules

**Losses and heads.** Prediction heads were trained with early stopping on supervised validation performance (Supp. Table S8). Unless otherwise noted, heads were lightweight MLPs with `dropout=0.1` and LayerNorm applied to hidden activations. In the Multiome bridge analysis (Fig. 5; Supp. Fig. S5), we used a single multi-class classification head to predict harmonized coarse PBMC labels (cross-entropy loss) with three hidden layers `[64, 64, 32]`. In the AML mosaic analysis (Fig. 7; Supp. Fig. S7), we used one binary mutation head per gene (binary cross-entropy), masking missing labels per gene so unlabeled cells do not contribute to that gene's supervised loss. Mutation heads used two hidden layers `[64, 32]`.

**Training schedule (head warmup $\rightarrow$ optional small-LR encoder fine-tuning).** Supervised refinement used two stages: (i) head warmup, training only head parameters with UniVI encoders/decoders frozen; then (ii) optional encoder unfreezing with a reduced learning rate, updating only the relevant encoder(s) together with the head. Missing labels were masked and excluded from supervised loss terms and evaluation.

#### Dataset-specific preprocessing details

**RNA — dataset-specific HVG and harmonization choices.** For bridging, HVGs (and any RNA scaling) were defined on the paired Multiome reference and reused unchanged for Ding RNA projection. For AML mosaics, RNA inputs were restricted to genes shared across AML CITE-seq and van Galen scRNA-seq. Dataset-specific HVG counts and gene-family exclusions for the SHARE-seq analysis are reported below.

**ADT — cross-cohort marker harmonization for AML mosaics.** For AML mosaic projection, the CLR/scaling transformation learned on AML CITE-seq training cells was applied unchanged to DAb-seq proteins; ADT inputs were harmonized by canonical marker names and restricted to shared markers.

**ATAC — TEA-seq tile construction.** For TEA-seq, ATAC inputs were constructed from a tile-by-cell matrix (SnapATAC2) (K Zhang et al. 2024); a fragments cutoff (ATAC `n_fragments` $\geq 1500$) was applied in the configuration used for Fig. 6.

#### Splits and stratification details

**Paired PBMC benchmarks (stratification and caps).** To stabilize optimization under class imbalance while preserving realistic evaluation, we constructed stratified train/validation splits by curated labels and applied per-class caps in train and validation. Paired cells not selected due to caps were assigned to the paired test set (`unused_to_test=True`), yielding a long-tailed held-out test set that reflects the natural label distribution. For the Hao et al. (2021) CITE-seq PBMC benchmark (Figs. 2–3), splits were stratified using `celltype.l3`, with caps of 1,200 cells per label for training and 150 per label for validation; remaining paired cells were assigned to test. For reporting, we evaluate label-transfer and structure metrics primarily on coarser label vocabularies (`celltype.l2` and, when indicated, `celltype.l1`). For the initial 10x Multiome PBMC analysis (Fig. 4), splits were stratified by `cell_type` with caps of 800 cells per label for training and 100 per label for validation; remaining paired cells were assigned to the paired test set. For all other 10x Multiome analyses, we used uncapped stratified train/validation/test splits over all paired cells, with proportions set to either *80%/10%/10%* or *75%/10%/15%* (Supp. Table S7).

**TEA-seq well-held-out split.** Models were trained on wells 3–4 and 6 and evaluated on held-out well 5 (prefix `X066-MP0C1W5`); preprocessing (RNA/ADT scaling; ATAC TF–IDF/LSI) was fit on training wells and applied unchanged to the held-out well.

**AML mutation-head evaluation split.** Where we trained mutation-prediction heads, evaluation used stratified cell-level splits due to metadata constraints; reported AUC/AP therefore reflect within-cohort generalization under cell-level holdout rather than strict patient-level generalization. Labels were handled per gene with masking of missing calls.

#### Regime-specific protocols

The main study-design regimes are defined in Methods (*Study design and evaluation regimes*). Here we record implementation details and deviations from the default pipeline (complete numerical settings in Supp. Tables S7–S8).

**3D latent visualization of the Multiome test set.** We generated a 3D visualization of the held-out 10x Multiome test set using the UniVI checkpoint trained for Fig. 4 (`loss_mode="v1"`). We encoded paired test cells by forward inference and plotted modality-specific posterior means ($Z_{\mathrm{RNA}}$ and $Z_{\mathrm{ATAC}}$) under two camera rotations; points were colored by modality or by `cell_type` (with modality indicated by marker shape) (Supp. Fig. S2).

**Reference-to-query bridging (parameter-frozen projection).** Bridge experiments treated the paired multimodal dataset as a reference used to learn a shared latent geometry and modality mappings, then embedded unimodal query cohorts by forward inference through the appropriate modality encoder without refitting parameters (Lotfollahi et al. 2022). This "reference-to-query mapping" setup is closely related in spirit to anchor-based reference mapping workflows (e.g., Seurat) (Stuart et al. 2019; Hao et al. 2023). All preprocessing transforms were fit on the reference training split and applied unchanged to projected cohorts. When indicated, we applied auxiliary latent-space supervision after projection (see Methods: *Optional supervised refinement after projection*; Supp. Table S8).

**TEA-seq well-held-out evaluation (stacked latent; Leiden and unimodal cluster concordance; visualization/diagnosis only).** All TEA-seq evaluation was performed on held-out well 5 using latent representations learned from training wells (3–4 and 6). We built a $k$-NN graph on the stacked well 5 latent points (Methods: *Latent representations used for evaluation*), symmetrized it, and applied Leiden community detection (Traag et al. 2019); the resolution was chosen for an interpretable granularity (14 clusters in Fig. 6E). Cluster modality composition was summarized over stacked points (Fig. 6F). These clusters were used for visualization/diagnosis only. As an orthogonal check, we computed unimodal Leiden clusters in native feature spaces (RNA expression, ADT features, and ATAC TF–IDF/LSI), mapped these labels onto the shared MoE latent coordinates, and compared modality concordance (Supp. Fig. S6A), alongside expanded marker overlays (Supp. Fig. S6B–G).

**AML mosaic integration via a paired RNA–protein bridge.** For AML mosaics, UniVI was trained on paired AML CITE-seq RNA+ADT as a bridge, then used to project van Galen scRNA-seq (RNA+genotype) and DAb-seq (protein+genotype) into the learned latent space using the corresponding modality encoders.

**LSC17 stemness score calculation in AML RNA cohorts.** To quantify leukemia stemness programs in all AML RNA feature sets, we computed the LSC17 signature score per cell using the published 17-gene panel (*DNMT3B*, *ZBTB46*, *NYNRIN*, *ARHGAP22*, *LAPTM4B*, *MMRN1*, *DPYSL3*, *FAM30A*, *CDK6*, *CPXM1*, *SOCS2*, *SMIM24*, *EMP1*, *BEX3*, *CD34*, *AKR1C3*, *ADGRG1*; LSC17). (Ng et al. 2016) For each AnnData object, gene symbols were matched to `adata.var_names` to ensure robust matching across cohorts. We extracted the expression matrix for the matched LSC17 genes from the specified normalized expression layer (here, `layer='log1p'`; otherwise `adata.X`) and computed the per-cell LSC17 score as the arithmetic mean across the available LSC17 genes:

$$\mathrm{LSC17\_score}_i = \frac{1}{|\mathcal{G}_i|} \sum_{g \in \mathcal{G}_i} x_{ig},$$

where $\mathcal{G}_i$ denotes the set of LSC17 genes present in the dataset and $x_{ig}$ is the normalized expression for cell $i$ and gene $g$. If one or more genes were absent from a dataset, scores were computed using the subset of present genes, and the number of matched genes was recorded.

To facilitate comparisons within each dataset, we also computed a dataset-specific standardized score ($Z$-score) across all cells in that AnnData object:

$$\text{LSC17\_z}_i = \frac{\text{LSC17\_score}_i - \mu}{\sigma + 10^{-8}},$$

where $\mu$ and $\sigma$ are the mean and standard deviation of `LSC17_score` across cells in that object. Scores were stored in `adata.obs` as `LSC17_score`, `LSC17_z`, and `LSC17_n_genes` (the number of LSC17 genes used for scoring).

**Mouse skin SHARE-seq paired RNA–ATAC integration.** For the SHARE-seq mouse skin analysis (Supp. Fig. S3), we applied dataset-specific QC and preprocessing reflecting SHARE-seq's higher sparsity and per-cell complexity profile compared with 10x Multiome. Cells annotated as "Mix" (an ambiguous-state label assigned by the original authors) were excluded prior to analysis, consistent with prior benchmarking on this dataset (Lee et al. 2023). After QC, $n = 31{,}220$ paired cells from 22 cell types were retained (75%/10%/15% stratified train/validation/test split). The held-out test set used in Supp. Fig. S3 contained $n = 3{,}139$ paired cells.

RNA QC: cells were retained with total counts in $[300, 30{,}000]$, $\geq 200$ detected genes, and mitochondrial fraction $\leq 5\%$. ATAC QC: cells were retained with peak counts in $[250, 12{,}500]$. After paired-barcode intersection, we excluded cells annotated as "Mix" as above. Peaks overlapping the ENCODE mm10 blacklist (Boyle Lab v2) (Amemiya et al. 2019) were removed prior to feature construction. RNA HVGs ($n = 5{,}000$) were selected on training cells only using SCANPY with `flavor="seurat_v3"`, with mitochondrial (*mt-*), ribosomal (*Rps*, *Rpl*, *Mrps*, *Mrpl*), predicted/uncharacterized (*Rik*), and pseudogene (*-ps*) families excluded from the HVG candidate pool prior to ranking. ATAC peaks were retained if accessible in 0.25%–80% of training cells; TF–IDF used the Signac convention $\log(\text{TF} \cdot \text{IDF} \cdot 10^4 + 1)$ with L2 row normalization, followed by TruncatedSVD to 101 components with LSI component 0 (depth-confounded) dropped from downstream model input. All preprocessing parameters (HVG selection, RNA $Z$-scoring, TF–IDF, LSI, LSI standardization) were fit on training cells and applied unchanged to validation/test splits. Model hyperparameters are recorded in Supp. Table S8.

**Mouse gastrulation scNMT-seq tri-modal integration.** For the scNMT-seq mouse gastrulation analysis (Supp. Fig. S4), we used the parsed EBI gastrulation bundle, loading RNA, CpG methylation (genebody-level), and GpC accessibility (genebody-level) into a paired AnnData triplet. Both methylation modalities used a beta-binomial likelihood operating on per-feature (successes, coverage) pairs, while RNA used a Gaussian likelihood on log-normalized expression. Reflecting the modest dataset size, we did not enforce the original publication's joint `pass_rnaQC` / `pass_metQC` / `pass_accQC` filter and did not apply additional cell- or feature-level filtering beyond the parsed bundle's defaults; methylation and accessibility features were retained as supplied. RNA inputs were library-size normalized to a target sum of $10^4$ counts per cell followed by $\log(1+x)$; methylation and accessibility were modeled directly from per-feature (successes, coverage) pairs via beta-binomial likelihoods, with the supplied fractional values in `.X` reserved for diagnostic visualization. Paired cells were assigned to 85%/5%/10% train/validation/test splits with seed 0. We report both an inductive evaluation on the held-out paired test set ($n = 114$;

Supp. Fig. S4A–E) and a transductive evaluation in which the same checkpoint is applied by forward inference to the full paired dataset ($n = 1{,}140$; Supp. Fig. S4F–J), reflecting the modest cohort size that limits sample-level holdout statistics. Model hyperparameters are recorded in Supp. Table S8.

#### Stress tests, hyperparameter sweeps, and scaling analyses

**Cell-type-specific modality dropout stress test (local missingness).** In paired 10x Multiome PBMCs, we retrained UniVI with one modality (RNA or ATAC) masked for training cells in a single annotated population $\ell$, while leaving all other training populations fully paired; evaluation used unchanged, fully paired validation/test splits by forward inference only. We reported per-ablated-label paired correspondence (FOSCTTM/Recall@K) and fused-space structure metrics (e.g., label transfer and ARI/NMI; Methods: *Evaluation metrics*). For diagnosis, we inspected stacked-embedding UMAP overlap and MoE gating preference summaries (e.g., $w_{\mathrm{RNA}} - w_{\mathrm{ATAC}}$).

**Global overlap stress test (controlled reduction of paired anchors).** We performed an overlap sweep in paired 10x Multiome PBMCs by retaining pairing for an $f$ fraction of cells and treating the remainder as unimodal (RNA-only or ATAC-only) according to the configuration, for $f \in \{0.0025, 0.005, 0.0075, 0.01, 0.015, 0.02, 0.025, 0.03, 0.035, 0.04, 0.05, 0.075, 0.10, 0.20, \ldots, 1.0\}$.

**Cross-validation protocol and metrics for overlap sweeps.** Overlap-sweep results were aggregated over 20 replicate runs defined by repeated splits of the paired Multiome dataset (fixed 75%/10%/15% train/validation/test per replicate, persisted as barcode-indexed maps). Preprocessing was refit on the training split within each replicate and applied unchanged thereafter. We reported mean and uncertainty across runs for: (i) paired correspondence (FOSCTTM; Recall@$K$ for $K \in \{1, 10, 25, 50, 100\}$), (ii) bidirectional $k$-NN label transfer (accuracy and macro-F1), and (iii) fused-space structure metrics (ARI/NMI and silhouette).

**Selection of $\beta$ and $\gamma$ defaults.** We treated $\beta$ (KL strength) and $\gamma$ (paired alignment strength) as multi-objective knobs and performed a $(\beta, \gamma)$ grid sweep in paired Multiome PBMCs, assessing fused-space structure (ARI), paired correspondence (FOSCTTM), cross-modality semantic consistency (label-transfer macro-F1), and strict paired retrieval (Recall@10). We selected the manuscript default from a broad region performing consistently well across objectives (Supp. Fig. S8).

**Computational scaling and resource profiling.** We measured computational scaling by increasing effective training set sizes via resampling (with replacement) from a fixed underlying set of paired Multiome training cells. For each effective size, we recorded wall-clock training time, peak memory (CPU RSS and accelerator peak when available), best validation loss under early stopping, and mean CPU utilization. Because resampling reuses the same observations, this experiment characterizes scaling behavior rather than statistical gains from additional unique cells (Supp. Fig. S11).

#### Benchmarking: per-method embedding and metric eligibility

For each baseline method, we defined a fused embedding for structure metrics and modality-specific embeddings when available. Metrics requiring modality-specific embeddings (paired retrieval/FOSCTTM, cross-modality label transfer, stacked modality mixing) were computed only when both modality embeddings were available from that method (or could be constructed without violating intended usage). Per-method categorization (inductive vs. transductive) and runner-specific notes are summarized in Supp. Table S6.

### References

10x Genomics 2021a. 10k human PBMCs, multiome v1.0, Chromium X, single-cell multiome ATAC + gene expression dataset generated by a Chromium X series instrument, analyzed using Cell Ranger ARC 2.0.0.

10x Genomics 2021b. PBMCs from a healthy donor – no cell sorting (10k), single-cell multiome ATAC + gene expression dataset generated by a Chromium Controller instrument and analyzed by Cell Ranger ARC 2.0.0.

Amemiya HM, Kundaje A, and Boyle AP. 2019. The ENCODE Blacklist: Identification of Problematic Regions of the Genome. *Scientific Reports*. **9**: 9354.

Andrew G, Arora R, Bilmes J, and Livescu K 2013. Deep Canonical Correlation Analysis. In: *Proceedings of the 30th International Conference on Machine Learning*. Ed. by S Dasgupta and D McAllester. Vol. 28. 3. Atlanta, Georgia, USA: PMLR, pp. 1247–1255.

Argelaguet R, Arnol D, Bredikhin D, Deloro Y, Velten B, Marioni JC, and Stegle O. 2020. MOFA+: a statistical framework for comprehensive integration of multi-modal single-cell data. *Genome Biology*. **21**: 111.

Argelaguet R et al. 2019. Multi-omics profiling of mouse gastrulation at single-cell resolution. *Nature*. **576**: 487–491.

Ashuach T, Gabitto MI, Koodli RV, Saldi GA, Jordan MI, and Yosef N. 2023. MultiVI: deep generative model for the integration of multimodal data. *Nature Methods*. **20**: 1222–1231.

Ashuach T, Reidenbach DA, Gayoso A, and Yosef N. 2022. PeakVI: A deep generative model for single cell chromatin accessibility analysis. *Cell Reports Methods*. **2**: 100570.

Cao ZJ and Gao G. 2022. Multi-omics single-cell data integration and regulatory inference with graph-linked embedding. *Nature Biotechnology*. **40**: 1458–1466.

Demaree B, Delley CL, Vasudevan HN, Peretz CAC, Ruff D, Smith CC, and Abate AR. 2021. Joint profiling of DNA and proteins in single cells to dissect genotype–phenotype associations in leukemia. *Nature Communications*. **12**: 1583.

Ding J et al. 2020. Systematic comparison of single-cell and single-nucleus RNA-sequencing methods. *Nature Biotechnology*. **38**: 737–746.

Galen P van et al. 2019. Single-Cell RNA-Seq Reveals AML Hierarchies Relevant to Disease Progression and Immunity. *Cell*. **176**: 1265–1281.e24.

Gong B, Y. Z, and E. P. 2021. CoBOLT: integrative analysis of multimodal single-cell sequencing data. *Genome Biology*. **22**: 351.

Hao Y et al. 2021. Integrated analysis of multimodal single-cell data. *Cell*. **184**: 3573–3587.e29.

Hao Y et al. 2023. Dictionary learning for integrative, multimodal and scalable single-cell analysis. *Nature Biotechnology*. **42**: 293–304.

Jain MS et al. 2021. MultiMAP: Dimensionality reduction and integration of multimodal data. *Genome Biology*. **22**: 346.

Knorr K et al. 2023. Systematic evaluation of AML-associated antigens identifies anti-U5 SNRNP200 therapeutic antibodies for the treatment of acute myeloid leukemia. *Nature Cancer*. **4**: 1675–1692.

Korsunsky I, Millard N, Fan J, Slowikowski K, Zhang F, Wei K, Baglaenko Y, Brenner M, Loh PR, and Raychaudhuri S. 2019. Fast, sensitive and accurate integration of single-cell data with Harmony. *Nature Methods*. **16**: 1289–1296.

Lee MYY, Kaestner KH, and Li M. 2023. Benchmarking algorithms for joint integration of unpaired and paired single-cell RNA-seq and ATAC-seq data. *Genome Biology*. **24**: 244.

Lin Y, Wu TY, Wan S, Yang JYH, Wong WH, and Wang YXR. 2022. scJoint integrates atlas-scale single-cell RNA-seq and ATAC-seq data with transfer learning. *Nature Biotechnology*. **40**: 703–710.

Lotfollahi M et al. 2022. Mapping single-cell data to reference atlases by transfer learning. *Nature Biotechnology*. **40**: 121–130.

Ma S et al. 2020. Chromatin potential identified by shared single-cell profiling of RNA and chromatin. *Cell*. **183**: 1103–1116.e20.

Ng SWK et al. 2016. A 17-gene stemness score for rapid determination of risk in acute leukaemia. *Nature*. **540**: 433–437.

Pijuan-Sala B et al. 2019. A single-cell molecular map of mouse gastrulation and early organogenesis. *Nature*. **566**: 490–495.

Satpathy AT et al. 2019. Massively parallel single-cell chromatin landscapes of human immune cell development and intratumoral T cell exhaustion. *Nature Biotechnology*. **37**: 925–936.

Stuart T, Butler A, Hoffman P, Hafemeister C, Papalexi E, Mauck WMI, Hao Y, Stoeckius M, Smibert P, and Satija R. 2019. Comprehensive integration of single-cell data. *Cell*. **177**: 1888–1902.e21.

Swanson E et al. 2021. Simultaneous trimodal single-cell measurement of transcripts, epitopes, and chromatin accessibility using TEA-seq. *eLife*. **10**: e63632.

Traag VA, Waltman L, and Eck NJ van. 2019. From Louvain to Leiden: guaranteeing well-connected communities. *Scientific Reports*. **9**: 5233.

Welch JD, Kozareva V, Ferreira A, Vanderburg C, Martin C, and Macosko EZ. 2019. Single-cell multi-omic integration compares and contrasts features of brain cell identity. *Cell*. **177**: 1873–1887.e17.

Zhang K, Zemke NR, Armand EJ, and Ren B. 2024. A fast, scalable and versatile tool for analysis of single-cell omics data. *Nature Methods*. **21**: 217–227.

Zhang R, Zhou T, Zhong W, Xie D, Wu F, and Wang K. 2023. scMoMaT jointly performs single cell mosaic integration and multi-modal bio-marker detection. *Nature Communications*. **14**: 384.
