## Supplemental Code for "Unifying multimodal single-cell data with a mixture-of-experts *β*-variational autoencoder framework": meta.yaml

```
{% set name = "univi" %}
{% set version = "0.4.7" %}

package:
  name: {{ name }}
  version: {{ version }}

source:
  # Use the PyPI sdist for released versions
  url: https://pypi.io/packages/source/u/{{ name }}/{{ name }}-{{ version }}.tar.gz
  # optional but recommended once you look it up:
  # sha256: f49634d0e4a76b73464031048950e80a281434e8a41610003d1d1da2186f0c34

build:
  noarch: python
  # IMPORTANT: no-build-isolation so pip uses the conda host env (with setuptools/wheel)
  script: {{ PYTHON }} -m pip install . --no-deps --no-build-isolation -vv

requirements:
  host:
    - python >=3.10
    - pip
    - setuptools
    - wheel

  run:
    - python >=3.10

    # Core numerics / data
    - numpy >=1.26
    - scipy >=1.11
    - pandas >=2.1

    # Single-cell ecosystem
    - anndata >=0.10
    - scanpy >=1.11

    # DL / ML
    - pytorch >=2.2
    - scikit-learn >=1.3

    # I/O and stats helpers
    - h5py >=3.10
    - pyyaml >=6.0

    # plotting / viz
    - matplotlib >=3.8
    - seaborn >=0.13

    # clustering / graphs
    - igraph >=0.11
    - leidenalg >=0.10

    # misc
    - tqdm >=4.66

test:
  imports:
    - univi
  commands:
    - python -c "import univi; print(getattr(univi, '__version__', 'no_version_attr'))"

about:
  home: https://github.com/Ashford-A/UniVI
  summary: "UniVI: a scalable multi-modal variational autoencoder toolkit for seamless integration and analysis of multimodal single-cell data."
  license: MIT
  # LICENSE is at repo root (../ from conda.recipe/)
  license_file: ../LICENSE

extra:
  recipe-maintainers:
    - Ashford-A
```
