## Supplemental Code for "Unifying multimodal single-cell data with a mixture-of-experts *β*-variational autoencoder framework": univi_env.yml

```
name: univi_env
channels:
  - conda-forge
  - bioconda
  - defaults
dependencies:
  - _libgcc_mutex=0.1=conda_forge
  - _openmp_mutex=4.5=2_gnu
  - bzip2=1.0.8=hda65f42_8
  - ca-certificates=2025.11.12=hbd8a1cb_0
  - ld_impl_linux-64=2.45=h1aa0949_0
  - libexpat=2.7.1=hecca717_0
  - libffi=3.5.2=h9ec8514_0
  - libgcc=15.2.0=h767d61c_7
  - libgcc-ng=15.2.0=h69a702a_7
  - libgomp=15.2.0=h767d61c_7
  - liblzma=5.8.1=hb9d3cd8_2
  - libnsl=2.0.1=hb9d3cd8_1
  - libsqlite=3.51.0=h0c1763c_0
  - libstdcxx=15.2.0=h8f9b012_7
  - libuuid=2.41.2=he9a06e4_0
  - libxcrypt=4.4.36=hd590300_1
  - libzlib=1.3.1=hb9d3cd8_2
  - ncurses=6.5=h2d0b736_3
  - openssl=3.6.0=h26f9b46_0
  - pip=25.3=pyh8b19718_0
  - python=3.10.19=h3c07f61_2_cpython
  - readline=8.2=h8c095d6_2
  - setuptools=80.9.0=pyhff2d567_0
  - tk=8.6.13=noxft_ha0e22de_103
  - wheel=0.45.1=pyhd8ed1ab_1
  - zstd=1.5.7=hb8e6e7a_2
  - pip:
      - anndata==0.10.9
      - anyio==4.11.0
      - argon2-cffi==25.1.0
      - argon2-cffi-bindings==25.1.0
      - array-api-compat==1.12.0
      - arrow==1.4.0
      - asttokens==3.0.1
      - async-lru==2.0.5
      - attrs==25.4.0
      - babel==2.17.0
      - beautifulsoup4==4.14.2
      - bleach==6.3.0
      - certifi==2025.11.12
      - cffi==2.0.0
      - charset-normalizer==3.4.4
      - choreographer==1.2.1
      - comm==0.2.3
      - contourpy==1.3.2
      - cycler==0.12.1
      - cykhash==2.0.1
      - debugpy==1.8.17
      - decorator==5.2.1
      - defusedxml==0.7.1
      - dill==0.4.0
      - exceptiongroup==1.3.0
      - executing==2.2.1
      - fastjsonschema==2.21.2
      - filelock==3.20.0
      - fonttools==4.60.1
      - fqdn==1.5.1
      - fsspec==2025.10.0
      - h11==0.16.0
      - h5py==3.15.1
      - hmmlearn==0.3.3
      - httpcore==1.0.9
      - httpx==0.28.1
      - idna==3.11
      - igraph==1.0.0
      - importlib-metadata==8.7.0
      - iniconfig==2.3.0
      - ipykernel==7.1.0
      - ipython==8.37.0
      - ipywidgets==8.1.8
      - isoduration==20.11.0
      - jedi==0.19.2
      - jinja2==3.1.6
      - joblib==1.5.2
      - json5==0.12.1
      - jsonpointer==3.0.0
      - jsonschema==4.25.1
      - jsonschema-specifications==2025.9.1
      - jupyter-client==8.6.3
      - jupyter-core==5.9.1
      - jupyter-events==0.12.0
      - jupyter-lsp==2.3.0
      - jupyter-server==2.17.0
      - jupyter-server-terminals==0.5.3
      - jupyterlab==4.4.10
      - jupyterlab-pygments==0.3.0
      - jupyterlab-server==2.28.0
      - jupyterlab-widgets==3.0.16
      - kaleido==1.2.0
      - kiwisolver==1.4.9
      - lark==1.3.1
      - legacy-api-wrap==1.5
      - leidenalg==0.11.0
      - llvmlite==0.45.1
      - logistro==2.0.1
      - macs3==3.0.3
      - markupsafe==3.0.3
      - matplotlib==3.10.7
      - matplotlib-inline==0.2.1
      - mistune==3.1.4
      - mpmath==1.3.0
      - multiprocess==0.70.18
      - natsort==8.4.0
      - nbclient==0.10.2
      - nbconvert==7.16.6
      - nbformat==5.10.4
      - nest-asyncio==1.6.0
      - networkx==3.4.2
      - notebook==7.4.7
      - notebook-shim==0.2.4
      - numba==0.62.1
      - numpy==1.26.4
      - nvidia-cublas-cu12==12.8.4.1
      - nvidia-cuda-cupti-cu12==12.8.90
      - nvidia-cuda-nvrtc-cu12==12.8.93
      - nvidia-cuda-runtime-cu12==12.8.90
      - nvidia-cudnn-cu12==9.10.2.21
      - nvidia-cufft-cu12==11.3.3.83
      - nvidia-cufile-cu12==1.13.1.3
      - nvidia-curand-cu12==10.3.9.90
      - nvidia-cusolver-cu12==11.7.3.90
      - nvidia-cusparse-cu12==12.5.8.93
      - nvidia-cusparselt-cu12==0.7.1
      - nvidia-nccl-cu12==2.27.5
      - nvidia-nvjitlink-cu12==12.8.93
      - nvidia-nvshmem-cu12==3.3.20
      - nvidia-nvtx-cu12==12.8.90
      - orjson==3.11.4
      - overrides==7.7.0
      - packaging==25.0
      - pandas==2.1.1
      - pandocfilters==1.5.1
      - parso==0.8.5
      - patsy==1.0.2
      - pexpect==4.9.0
      - pillow==12.0.0
      - platformdirs==4.5.0
      - plotly==5.24.1
      - pluggy==1.6.0
      - polars==1.35.2
      - polars-runtime-32==1.35.2
      - pooch==1.8.2
      - prometheus-client==0.23.1
      - prompt-toolkit==3.0.52
      - psutil==7.1.3
      - ptyprocess==0.7.0
      - pure-eval==0.2.3
      - pyarrow==22.0.0
      - pycparser==2.23
      - pyfaidx==0.7.2.2
      - pygments==2.19.2
      - pynndescent==0.5.13
      - pyparsing==3.2.5
      - pytest==9.0.1
      - pytest-timeout==2.4.0
      - python-dateutil==2.9.0.post0
      - python-json-logger==4.0.0
      - pytz==2025.2
      - pyyaml==6.0.3
      - pyzmq==27.1.0
      - referencing==0.37.0
      - requests==2.32.5
      - rfc3339-validator==0.1.4
      - rfc3986-validator==0.1.1
      - rfc3987-syntax==1.1.0
      - rpds-py==0.29.0
      - rustworkx==0.17.1
      - scanpy==1.11.5
      - scikit-learn==1.7.2
      - scipy==1.13.1
      - seaborn==0.13.2
      - send2trash==1.8.3
      - session-info2==0.2.3
      - simplejson==3.20.2
      - six==1.17.0
      - snapatac2==2.8.0
      - sniffio==1.3.1
      - soupsieve==2.8
      - stack-data==0.6.3
      - statsmodels==0.14.5
      - sympy==1.14.0
      - tenacity==9.1.2
      - terminado==0.18.1
      - texttable==1.7.0
      - threadpoolctl==3.6.0
      - tinycss2==1.4.0
      - tomli==2.3.0
      - torch==2.9.1
      - tornado==6.5.2
      - tqdm==4.67.1
      - traitlets==5.14.3
      - triton==3.5.1
      - typeguard==4.4.4
      - typing-extensions==4.15.0
      - tzdata==2025.2
      - umap-learn==0.5.9.post2
      - uri-template==1.3.0
      - urllib3==2.5.0
      - wcwidth==0.2.14
      - webcolors==25.10.0
      - webencodings==0.5.1
      - websocket-client==1.9.0
      - widgetsnbextension==4.0.15
      - zipp==3.23.0
```
