## Supplemental Code for "Unifying multimodal single-cell data with a mixture-of-experts *β*-variational autoencoder framework": univi_v0.4.7_env.yml

```
name: univi_v0.4.7
channels:
  - conda-forge
dependencies:
  - python=3.10
  - ipywidgets
  - jupyterlab
  - jupyterlab_widgets
  - notebook
  - widgetsnbextension
  - pip:
    - anndata==0.11.4
    - array-api-compat==1.13.0
    - backports.tarfile==1.2.0
    - biothings_client==0.4.1
    - build==1.4.0
    - choreographer==1.2.1
    - contourpy==1.3.2
    - cycler==0.12.1
    - cykhash==2.0.1
    - dill==0.4.1
    - docutils==0.22.4
    - et_xmlfile==2.0.0
    - filelock==3.24.3
    - fonttools==4.61.1
    - fsspec==2026.2.0
    - h5py==3.15.1
    - hmmlearn==0.3.3
    - igraph==1.0.0
    - iniconfig==2.3.0
    - ipympl==0.10.0
    - ipython==8.38.0
    - joblib==1.5.3
    - kaleido==1.2.0
    - kiwisolver==1.4.9
    - legacy-api-wrap==1.5
    - leidenalg==0.11.0
    - llvmlite==0.46.0
    - logistro==2.0.1
    - matplotlib==3.10.8
    - mpmath==1.3.0
    - multiprocess==0.70.19
    - mygene==3.2.2
    - natsort==8.4.0
    - ncls==0.0.70
    - networkx==3.4.2
    - numba==0.64.0
    - numpy==2.2.6
    - openpyxl==3.1.5
    - orjson==3.11.7
    - pandas==2.3.3
    - pandocfilters==1.5.1
    - patsy==1.0.2
    - pillow==12.1.1
    - plotly==5.24.1
    - polars==1.38.1
    - polars-runtime-32==1.38.1
    - pooch==1.9.0
    - pyarrow==23.0.1
    - pycparser==3.0
    - pyfaidx==0.7.2.2
    - pynndescent==0.6.0
    - pyparsing==3.3.2
    - pyranges==0.1.4
    - pytest==9.0.2
    - pytest-timeout==2.4.0
    - python-json-logger==4.0.0
    - rustworkx==0.17.1
    - scanpy==1.11.5
    - scikit-learn==1.7.2
    - scikit-misc==0.5.2
    - scipy==1.15.3
    - seaborn==0.13.2
    - session-info2==0.4
    - simplejson==3.20.2
    - snapatac2==2.8.0
    - sorted_nearest==0.0.41
    - statsmodels==0.14.6
    - sympy==1.14.0
    - tabulate==0.9.0
    - tenacity==9.1.4
    - texttable==1.7.0
    - threadpoolctl==3.6.0
    - tinycss2==1.4.0
    - torch==2.10.0
    - torchaudio==2.10.0
    - torchvision==0.25.0
    - typeguard==4.5.0
    - umap-learn==0.5.11
    - univi==0.4.7
    - uv==0.10.3

prefix: "you/path/to/envs/univi_v0.4.7"
```
