## Supplemental Code for "Unifying multimodal single-cell data with a mixture-of-experts *β*-variational autoencoder framework": UniVI_working_environment_v2_minimal.yml

```
name: UniVI_working_environment_v2
channels:
  - conda-forge
  - bioconda
  - defaults
dependencies:
  - nose=1.3.7
  - ujson=5.7.0
  - python-jsonrpc-server=0.4.0
  - click=8.1.3
  - libuuid=2.32.1
  - mkl_fft=1.3.1
  - liblapacke=3.9.0
  - conda-pack=0.7.0
  - snappy=1.1.9
  - wcwidth=0.2.6
  - three-merge=0.1.1
  - conda-build=3.23.3
  - clyent=1.2.2
  - pyzmq=25.0.0
  - mock=5.0.1
  - jbig=2.1
  - r-base=3.6.3
  - mypy_extensions=1.0.0
  - cloudpickle=2.2.1
  - alsa-lib=1.2.3.2
  - libdeflate=1.14
  - markupsafe=2.1.2
  - gfortran_impl_linux-64=9.3.0
  - bottleneck=1.3.6
  - brotlipy=0.7.0
  - libcblas=3.9.0
  - gfortran_linux-64=9.3.0
  - pyrsistent=0.19.3
  - lcms2=2.14
  - cftime=1.6.2
  - pickleshare=0.7.5
  - libopus=1.3.1
  - babel=2.12.1
  - harfbuzz=2.8.1
  - zstd=1.5.2
  - defusedxml=0.7.1
  - numexpr=2.7.3
  - toml=0.10.2
  - msgpack-python=1.0.5
  - spyder-kernels=2.0.4
  - blosc=1.21.3
  - multipledispatch=0.6.0
  - text-unidecode=1.3
  - pyflakes=2.2.0
  - libgfortran5=9.3.0
  - fsspec=2023.3.0
  - contextlib2=21.6.0
  - docutils=0.19
  - python-language-server=0.36.2
  - python-tzdata=2022.7
  - greenlet=2.0.2
  - pysocks=1.7.1
  - xorg-renderproto=0.11.1
  - libclang=11.1.0
  - libzip=1.9.2
  - pathlib2=2.3.7.post1
  - backports.shutil_get_terminal_size=1.0.0
  - hdf4=4.2.15
  - _r-mutex=1.0.1
  - tomli=2.0.1
  - cached-property=1.5.2
  - zope=1.0
  - zeromq=4.3.4
  - krb5=1.19.3
  - bzip2=1.0.8
  - ptyprocess=0.7.0
  - comm=0.1.2
  - pkginfo=1.9.6
  - unicodedata2=15.0.0
  - stdlib-list=0.8.0
  - libpng=1.6.39
  - zipp=3.15.0
  - libstdcxx-devel_linux-64=9.3.0
  - mkl=2022.2.1
  - arrow=1.2.3
  - requests=2.28.2
  - tzdata=2022g
  - six=1.16.0
  - jupyterlab=3.5.0
  - cycler=0.11.0
  - libopenblas=0.3.12
  - bwidget=1.9.14
  - testpath=0.6.0
  - libogg=1.3.4
  - nspr=4.35
  - ipykernel=6.21.2
  - gcc_linux-64=9.3.0
  - path=16.6.0
  - pycparser=2.21
  - xorg-xextproto=7.3.0
  - mccabe=0.6.1
  - mpmath=1.3.0
  - ply=3.11
  - asn1crypto=1.5.1
  - locket=1.0.0
  - gsl=2.6
  - pyhamcrest=2.0.4
  - mpfr=4.1.0
  - patch=2.7.6
  - r-matrix=1.2_17
  - tornado=6.2
  - websocket-client=1.5.1
  - nss=3.88
  - patsy=0.5.3
  - snowballstemmer=2.2.0
  - textdistance=4.5.0
  - zict=2.2.0
  - python_abi=3.8
  - libxcb=1.13
  - libglib=2.68.3
  - qtconsole-base=5.4.0
  - libwebp-base=1.2.4
  - path.py=12.5.0
  - pytz-deprecation-shim=0.1.0.post0
  - isort=5.12.0
  - bokeh=2.4.3
  - nbclassic=0.5.3
  - conda-verify=3.1.1
  - argon2-cffi-bindings=21.2.0
  - jupyter_server=2.4.0
  - kiwisolver=1.4.4
  - graphite2=1.3.13
  - gmpy2=2.1.2
  - pyqt=5.12.3
  - sleef=3.5.1
  - fonts-conda-forge=1
  - imagecodecs-lite=2019.12.3
  - unicodecsv=0.14.1
  - liblief=0.12.3
  - libbrotlicommon=1.0.9
  - zstandard=0.19.0
  - _libgcc_mutex=0.1
  - cytoolz=0.12.0
  - contourpy=1.0.7
  - libgcc-devel_linux-64=9.3.0
  - fastcache=1.1.0
  - brotli-bin=1.0.9
  - gmp=6.2.1
  - webencodings=0.5.1
  - jedi=0.17.2
  - libgomp=12.2.0
  - spyder=5.0.5
  - xorg-libsm=1.2.3
  - ipython_genutils=0.2.0
  - xorg-libxau=1.0.9
  - keyutils=1.6.1
  - pylint=2.17.0
  - pixman=0.40.0
  - openssl=1.1.1t
  - iniconfig=2.0.0
  - cached_property=1.5.2
  - rtree=1.0.1
  - tomlkit=0.11.6
  - bleach=6.0.0
  - openpyxl=3.1.1
  - conda-package-handling=2.0.2
  - pycosat=0.6.4
  - libcurl=7.77.0
  - jupyter_console=6.6.3
  - astroid=2.15.0
  - tktable=2.10
  - rope=1.7.0
  - conda-content-trust=0.1.3
  - sphinxcontrib-devhelp=1.0.2
  - certifi=2022.12.7
  - libevent=2.1.10
  - tqdm=4.65.0
  - libedit=3.1.20191231
  - libbrotlienc=1.0.9
  - ncurses=6.2
  - anaconda=custom
  - freetype=2.12.1
  - sphinx=6.1.3
  - zope.event=4.6
  - fbpca=1.0
  - libtool=2.4.7
  - sphinxcontrib=1.0
  - notebook=6.5.3
  - xlsxwriter=3.0.8
  - libllvm9=9.0.1
  - pyerfa=2.0.0.1
  - xorg-libxrender=0.9.10
  - jinja2=3.1.2
  - libnghttp2=1.51.0
  - matplotlib=3.7.1
  - pip
  - matplotlib-base=3.7.1
  - networkx=3.0
  - jinja2-time=0.2.0
  - python-fastjsonschema=2.16.3
  - binaryornot=0.4.4
  - pygments=2.14.0
  - ripgrep=13.0.0
  - jupyterlab_pygments=0.2.2
  - sphinxcontrib-qthelp=1.0.3
  - pango=1.48.6
  - anaconda-client=1.11.1
  - entrypoints=0.4
  - intervaltree=3.1.0
  - fontconfig=2.14.2
  - lz4=4.3.2
  - more-itertools=9.1.0
  - terminado=0.17.1
  - patchelf=0.17.2
  - expat=2.5.0
  - scipy=1.6.3
  - ld_impl_linux-64=2.35.1
  - libev=4.33
  - libsodium=1.0.18
  - binutils_impl_linux-64=2.35.1
  - imagesize=1.4.1
  - prompt_toolkit=3.0.38
  - libgcc-ng=12.2.0
  - libxkbcommon=1.0.3
  - json5=0.9.5
  - liblapack=3.9.0
  - packaging=23.0
  - pathspec=0.11.0
  - dunamai=1.16.0
  - unidecode=1.3.6
  - attrs=22.2.0
  - flake8=3.8.4
  - werkzeug=2.2.3
  - zope.interface=5.5.2
  - libnetcdf=4.8.0
  - sysroot_linux-64=2.12
  - xorg-xproto=7.0.31
  - sniffio=1.3.0
  - jupyter_events=0.6.3
  - backcall=0.2.0
  - qstylizer=0.2.2
  - distributed=2023.3.0
  - gstreamer=1.18.4
  - pywavelets=1.4.1
  - scikit-image=0.19.3
  - backports.zoneinfo=0.2.1
  - flask=2.2.3
  - et_xmlfile=1.1.0
  - libsqlite=3.40.0
  - tk=8.6.12
  - xorg-kbproto=1.0.7
  - curl=7.77.0
  - xorg-libxdmcp=1.1.3
  - pyjwt=2.6.0
  - pthread-stubs=0.4
  - pytables=3.6.1
  - atomicwrites=1.4.1
  - cookiecutter=2.1.1
  - xlwt=1.3.0
  - rpy2=3.4.5
  - soupsieve=2.3.2.post1
  - libxml2=2.9.12
  - joblib=1.2.0
  - nbformat=5.7.3
  - rfc3339-validator=0.1.4
  - anyio=3.6.2
  - mysql-common=8.0.32
  - qtconsole=5.4.0
  - libbrotlidec=1.0.9
  - libllvm11=11.1.0
  - widgetsnbextension=4.0.5
  - tblib=1.7.0
  - traitlets=5.9.0
  - nbconvert-core=7.2.9
  - psutil=5.9.4
  - sortedcontainers=2.4.0
  - py=1.11.0
  - watchdog=2.3.1
  - pycurl=7.43.0.6
  - netcdf4=1.5.7
  - lazy-object-proxy=1.9.0
  - sphinxcontrib-applehelp=1.0.4
  - pytz=2022.7.1
  - conda-env=2.6.0
  - intel-openmp=2022.1.0
  - _ipyw_jlab_nb_ext_conf=0.1.0
  - glib=2.68.3
  - conda-repo-cli=1.0.27
  - qt=5.12.9
  - pyopenssl=23.0.0
  - gxx_linux-64=9.3.0
  - pkgutil-resolve-name=1.3.10
  - pydocstyle=6.3.0
  - c-ares=1.18.1
  - jupyterlab_widgets=3.0.5
  - argon2-cffi=21.3.0
  - tzlocal=4.2
  - gettext=0.21.1
  - wurlitzer=3.0.3
  - sphinxcontrib-websupport=1.2.4
  - _openmp_mutex=4.5
  - statsmodels=0.13.5
  - astropy=5.2.1
  - lzo=2.10
  - libspatialindex=1.9.3
  - nbclient=0.7.2
  - libvorbis=1.3.7
  - pytest=7.2.2
  - ruamel_yaml=0.15.80
  - cffi=1.14.5
  - kernel-headers_linux-64=2.6.32
  - importlib_metadata=6.0.0
  - nest-asyncio=1.5.6
  - sphinxcontrib-jsmath=1.0.1
  - mysql-libs=8.0.32
  - parso=0.7.0
  - gevent=22.10.2
  - h5py=3.3.0
  - jpeg=9e
  - pyls-spyder=0.3.2
  - python-slugify=8.0.1
  - anaconda-navigator=2.3.2
  - python=3.8.10
  - make=4.3
  - pyodbc=4.0.35
  - gcc_impl_linux-64=9.3.0
  - beautifulsoup4=4.11.2
  - sphinxcontrib-htmlhelp=2.0.1
  - libarchive=3.5.2
  - r-lattice=0.20_38
  - zlib=1.2.13
  - colorama=0.4.6
  - unixodbc=2.3.10
  - rfc3986-validator=0.1.1
  - toolz=0.12.0
  - pandas=1.5.3
  - anndata2ri=1.1
  - ipywidgets=8.0.4
  - jupyter_server_terminals=0.4.4
  - glib-tools=2.68.3
  - get_terminal_size=1.0.0
  - libffi=3.3
  - conda=23.1.0
  - importlib_resources=5.12.0
  - mkl-service=2.4.0
  - python-json-logger=2.0.7
  - cairo=1.16.0
  - pyls-black=0.4.6
  - binutils_linux-64=2.35.1
  - tifffile=2019.7.26.2
  - diff-match-patch=20200713
  - fonttools=4.39.0
  - libprotobuf=3.21.12
  - stack_data=0.6.2
  - seaborn=0.12.2
  - font-ttf-source-code-pro=2.038
  - glob2=0.7
  - pillow=9.2.0
  - threadpoolctl=3.1.0
  - sip=6.7.7
  - heapdict=1.0.1
  - tbb=2021.7.0
  - llvm-openmp=15.0.7
  - libuv=1.44.2
  - numpydoc=1.5.0
  - platformdirs=3.1.0
  - pcre=8.45
  - geosketch=1.2
  - pyparsing=3.0.9
  - alabaster=0.7.13
  - ruamel.yaml=0.17.21
  - prompt-toolkit=3.0.38
  - sphinxcontrib-serializinghtml=1.1.5
  - sed=4.8
  - session-info=1.0.0
  - jupyter_client=8.0.3
  - dbus=1.13.6
  - backports.functools_lru_cache=1.6.4
  - mistune=2.0.5
  - singledispatch=3.6.1
  - readline=8.1
  - fonts-conda-ecosystem=1
  - pathtools=0.1.2
  - idna=3.4
  - brotli=1.0.9
  - pep8=1.7.1
  - libtiff=4.4.0
  - pexpect=4.8.0
  - enum34=1.1.10
  - sympy=1.11.1
  - font-ttf-dejavu-sans-mono=2.37
  - anaconda-project=0.11.1
  - pyxdg=0.28
  - python-annoy=1.17.1
  - arviz=0.11.2
  - pystan=2.19.1.1
  - plotly=5.13.0
  - cryptography=38.0.4
  - nbconvert-pandoc=7.2.9
  - numba=0.56.4
  - libstdcxx-ng=12.2.0
  - autopep8=1.5.5
  - xorg-libx11=1.8.4
  - libpq=13.3
  - libzlib=1.2.13
  - nltk=3.8.1
  - icu=68.2
  - lz4-c=1.9.3
  - flit-core=3.8.0
  - _anaconda_depends=2020.07
  - font-ttf-inconsolata=3.000
  - sortedcollections=2.1.0
  - scanorama=1.7.3
  - qdarkstyle=3.0.2
  - chardet=5.1.0
  - regex=2022.10.31
  - html5lib=1.1
  - pluggy=1.0.0
  - yapf=0.32.0
  - libiconv=1.16
  - matplotlib-inline=0.1.6
  - xz=5.2.6
  - exceptiongroup=1.1.0
  - libblas=3.9.0
  - partd=1.3.0
  - lerc=4.0.0
  - bitarray=2.7.3
  - black=23.1.0
  - dask=2023.3.0
  - yaml=0.2.5
  - munkres=1.1.4
  - pandoc=2.19.2
  - scikit-learn=1.1.1
  - pycodestyle=2.6.0
  - dask-core=2023.3.0
  - future=0.18.3
  - numpy=1.21.5
  - font-ttf-ubuntu=0.83
  - tinycss2=1.2.1
  - backports=1.0
  - imageio=2.26.0
  - libssh2=1.10.0
  - ca-certificates=2022.12.7
  - jdcal=1.4.1
  - get_version=3.5.1
  - libgfortran-ng=9.3.0
  - pandocfilters=1.5.0
  - pytoolconfig=1.2.5
  - mpc=1.3.1
  - jeepney=0.8.0
  - llvmlite=0.39.1
  - typing=3.10.0.0
  - notebook-shim=0.2.2
  - gst-plugins-base=1.18.4
  - bkcharts=0.2
  - cython=0.29.33
  - importlib-metadata=6.0.0
  - nbconvert=7.2.9
  - send2trash=1.8.0
  - simplegeneric=0.8.1
  - tenacity=8.2.2
  - hdf5=1.10.6
  - decorator=5.1.1
  - mkl_random=1.2.2
  - ruamel.yaml.clib=0.2.7
  - lxml=4.6.3
  - itsdangerous=2.1.2
  - filelock=3.9.0
  - py-lief=0.12.3
  - xorg-libice=1.0.10
  - wheel=0.38.4
  - xarray=2022.11.0
  - jsonschema=4.17.3
  - jupyter=1.0.0
  - jupyterlab_server=2.20.0
  - pathlib=1.0.1
  - gxx_impl_linux-64=9.3.0
  - jaraco.classes=3.2.3
  - libxslt=1.1.33
  - argh=0.26.2
  - openjpeg=2.5.0
  - fribidi=1.0.10
  - python-libarchive-c=4.0
  - jupyter_core=5.2.0
  - olefile=0.46
  - wrapt=1.15.0
  - dill=0.3.6
  - prometheus_client=0.16.0
  - pyyaml=6.0
  - debugpy=1.6.6
  - boto=2.49.0
  - qtpy=2.3.0
  - sqlite=3.36.0
  - pure_eval=0.2.2
  - urllib3=1.26.14
  - qtawesome=1.2.2
  - inflection=0.5.1
  - conda-token=0.4.0
  - python-dateutil=2.8.2
  - navigator-updater=0.3.0
  - conda-package-streaming=0.7.0
  - blas=2.106
  - xorg-libxext=1.3.4
  - sqlalchemy=2.0.5.post1
```
