## Supplemental Code for "Unifying multimodal single-cell data with a mixture-of-experts *β*-variational autoencoder framework": Additional_AML_atlas_integration.ipynb

```
# %% [markdown]
# UniVI mosaic integration — van Galen AML + Granja healthy BMMC

**Goal.** Integrate the van Galen 2019 AML cohort (`./data/van_Galen_data/vanGalen_anndata.h5ad`)
with healthy BMMC from Granja 2019 (`GSE139369_RAW.tar`) using **UniVI**, treating each
scRNA-seq dataset as data that is *bridged through the cell-type label*, following your
preliminary **CTM (cell-type-as-modality)** experiments.

**Why a label bridge and not paired cross-reconstruction.** UniVI's `v1` loss aligns
modalities through *cell-to-cell* cross-reconstruction, which assumes paired cells. Your two
RNA cohorts are **unpaired** (different patients, different platforms — van Galen is **Seq-Well**,
Granja is **10x CITE-seq**). The one thing that *is* shared per cell is its **cell-type label**.
So the integration runs as: one combined **RNA** modality (both cohorts on a shared gene space)
+ a **categorical cell-type modality** that is paired to every RNA cell. UniVI's
γ-alignment then pulls same-cell-type cells together across cohorts. This is exactly the
"categorical variable as a full modality" mechanism in the UniVI README, and it is the native
implementation of your CTM idea.

**This notebook is a pipeline, not pre-computed results.** Nothing here was run on your data —
it is written to execute in your environment against UniVI **v0.4.7**.

**Variant map** (your shorthand → the real UniVI mechanism this notebook uses):

| Your variant | UniVI mechanism | Where in code |
|---|---|---|
| `SE` (supervised baseline) | RNA modality + supervised **class head** on cell type (`ClassHeadConfig`) | `build_config("SE")` |
| `CTM` | RNA + **categorical cell-type modality** (`ModalityConfig(likelihood="categorical", input_kind="obs")`) | `build_config("CTM")` |
| `H_CTM` (hierarchical) | RNA + multiple categorical modalities (cell type × tissue/dataset × malignancy) + **adversarial** dataset head | `build_config("H_CTM")` |
| `MUT_CTM_M` (masked mutation) | CTM + **binary mutation head** with `ignore_index=-1` so cells without genotype calls are masked out of the head loss | `build_config("MUT_CTM_M")` |

> **Verified note on masking.** Your MUT_CTM_M "masking fix" is native in UniVI: both class heads
> (`ClassHeadConfig.ignore_index`) and categorical modalities (`ModalityConfig.ignore_index`)
> default to `-1`, and the dataset/loss skip those cells. You do **not** need to alternate
> 2-/3-modality batches manually — set unknown genotypes to `-1` and the head ignores them.


# %% [markdown]
## 0 · Environment, version pin, and reproducibility

# %% [code]
# UniVI install (skip if already in your env):
#   pip install univi            # v0.4.7 from PyPI
#   # or: conda install -c conda-forge univi
import numpy as np
import pandas as pd
import scanpy as sc
import anndata as ad
import scipy.sparse as sp
import torch
from torch.utils.data import DataLoader, Subset

import univi
print("UniVI version:", univi.__version__)
assert univi.__version__ >= "0.4.7", "This notebook targets UniVI >= 0.4.7"

from univi import UniVIMultiModalVAE, ModalityConfig, UniVIConfig, TrainingConfig
from univi.config import ClassHeadConfig
from univi.data import MultiModalDataset, align_paired_obs_names, collate_multimodal_xy_recon
from univi.trainer import UniVITrainer
from univi.evaluation import encode_adata, evaluate_alignment, reconstruction_metrics
from univi.plotting import set_style, umap
from univi.utils.seed import set_seed

set_seed(0)
set_style(font_scale=1.1, dpi=140)

device = (
    "cuda" if torch.cuda.is_available() else
    ("mps" if getattr(torch.backends, "mps", None) is not None and torch.backends.mps.is_available()
     else "cpu")
)
print("device:", device)

def to_dense(X):
    return X.toarray() if sp.issparse(X) else np.asarray(X)


# %% [markdown]
## 1 · Configuration

Hyperparameters mirror the shipped AML config (`params_aml_citeseq_GR_fig7.json`):
`latent_dim=30`, `beta=1.15`, `gamma=1.75`, Gaussian likelihoods on log-normalized+scaled RNA,
all genes (no HVG) in the manuscript — here HVG is optional for speed. The mutation gene panel
is the same 7 genes the manuscript fine-tuned heads on.

# %% [code]
from pathlib import Path

# ---- paths (your layout) ----
DATA_ROOT   = Path("./data")
VG_PATH     = DATA_ROOT / "van_Galen_data" / "vanGalen_anndata.h5ad"
GRANJA_TAR  = DATA_ROOT / "Granja_data" / "GSE139369_RAW.tar"
GRANJA_EXTRACT = DATA_ROOT / "Granja_data" / "extracted"   # tar is extracted here on first run
OUTDIR      = Path("./runs/univi_aml_healthy"); OUTDIR.mkdir(parents=True, exist_ok=True)

# ---- which experiment variant to run ----
VARIANT = "SE"           # one of: "SE", "CTM", "H_CTM", "MUT_CTM_M"

# ---- model / training (from the AML reference config) ----
LATENT_DIM = 30
BETA       = 1.25
#GAMMA      = 4.35
GAMMA      = 2.15
N_EPOCHS   = 3000          # AML config value; lower (e.g. 300-800) for a first pass
BATCH_SIZE = 256
LR         = 1e-4
WEIGHT_DECAY = 1e-5
USE_HVG    = True          # manuscript used all genes; HVG=True is a faster default
N_HVG      = 4000

# ---- biology ----
MUTATION_GENES = ["NPM1", "DNMT3A", "TP53", "FLT3", "IDH2", "TET2", "KRAS"]  # AML config panel
HERO_GENE      = "NPM1"    # your headline mutation
# Granja healthy BMMC sample IDs of interest (from GSE139369 sample list)
GRANJA_BMMC_SAMPLES = ["CITE_BMMC_D1T1", "CITE_BMMC_D1T2"]   # add CITE_CD34_* to enrich HSPCs


# %% [markdown]
## 2 · Load van Galen AML (`.h5ad`)

The authors' own AML notebook reads this exact file and uses `obs["CellType"]` for cell-type
labels and obs columns named like `"NPM1 W288fs"` for per-cell genotype calls. We reuse those
conventions. The mutation matcher OR's across every obs column whose name starts with a gene
symbol (so `NPM1 W288fs`, `NPM1 ...` all map to `NPM1`), producing a **label** (`1`=mutant,
`0`=WT) and a **mask** (`1`=called, `0`=unknown). Unknown → `-1` for UniVI.

# %% [code]
import re

vg = sc.read_h5ad(str(VG_PATH))
vg.var_names_make_unique()
vg.obs_names_make_unique()
print("van Galen:", vg.shape)
print("obs columns (first 40):", list(vg.obs.columns)[:40])

# --- cell-type column (authors used "CellType"); fall back gracefully ---
def find_col(adata, candidates):
    low = {c.lower(): c for c in adata.obs.columns}
    for c in candidates:
        if c in adata.obs.columns: return c
        if c.lower() in low:       return low[c.lower()]
    return None

VG_CT_COL = find_col(vg, ["CellType", "celltype", "cell_type", "annotation", "PredictionRefined"])
assert VG_CT_COL is not None, f"No cell-type column found in van Galen obs: {list(vg.obs.columns)}"
print("van Galen cell-type column:", VG_CT_COL)
print(vg.obs[VG_CT_COL].value_counts())


# %% [code]
# --- van Galen GoT genotyping: parse MutTranscripts / WtTranscripts ---
import re

def parse_genes(s):
    """Return the set of gene symbols in a Mut/Wt transcript string."""
    s = str(s).strip()
    if s in ("", "NA", "nan", "None", "<NA>"):
        return set()
    out = set()
    for tok in s.split("///"):                      # ' /// ' separates multiple calls
        tok = tok.strip()
        if not tok:
            continue
        m = re.match(r"[A-Za-z0-9]+", tok)          # gene = leading symbol before '.', '-', '/'
        if m:
            out.add(m.group().upper())
    return out

mut_sets = vg.obs["MutTranscripts"].map(parse_genes)   # genes with a mutant call per cell
wt_sets  = vg.obs["WtTranscripts"].map(parse_genes)    # genes with a wild-type call per cell

def build_mut_label_mask(adata, genes):
    Y = np.zeros((adata.n_obs, len(genes)), np.float32)   # 1 = mutant
    M = np.zeros((adata.n_obs, len(genes)), np.float32)   # 1 = genotyped (called) at this locus
    for gi, g in enumerate(genes):
        gU = g.upper()
        mut = mut_sets.map(lambda S: gU in S).to_numpy()
        wt  = wt_sets.map(lambda S: gU in S).to_numpy()
        M[:, gi] = (mut | wt).astype(np.float32)
        Y[:, gi] = mut.astype(np.float32)
    return Y, M

VG_Y, VG_M = build_mut_label_mask(vg, MUTATION_GENES)
for gi, g in enumerate(MUTATION_GENES):
    print(f"{g:8s} called={int(VG_M[:,gi].sum()):6d}  mutant={int(VG_Y[:,gi].sum()):6d}")

# HERO gene -> int code with -1 = ungenotyped (UniVI ignore_index)
hi = MUTATION_GENES.index(HERO_GENE)
hero_code = np.where(VG_M[:, hi] > 0, VG_Y[:, hi].astype(int), -1)
vg.obs[f"{HERO_GENE}_code"] = hero_code.astype(int)
print(f"\n{HERO_GENE} codes:", pd.Series(hero_code).value_counts().to_dict(), "(-1 = ungenotyped)")

# quick sanity check — eyeball a few genotyped cells
g_cells = vg.obs.loc[(VG_M.sum(1) > 0), ["MutTranscripts", "WtTranscripts"]].head(6)
print("\nexample genotyped cells:\n", g_cells.to_string())


# %% [markdown]
## 3 · Load Granja healthy BMMC from `GSE139369_RAW.tar`

> **Honest constraint.** GEO lists this archive as *"TAR (of RDS, TSV)"* — the per-sample
> processed objects are **R `.rds` `SummarizedExperiment`s**, which pure-Python readers (e.g.
> `pyreadr`) cannot parse. The cell below **inspects the tar first** (it does not blindly
> extract 20.4 GB) and then dispatches on what is actually inside. Three branches are provided:
> 10x triplet (`mtx`+`barcodes`+`features`), dense `tsv` matrix, and `.rds` via R. **I could not
> see inside your tar from here**, so verify the printed member list and pick the matching branch.
>
> If the members are `.rds`, the most reliable route is a one-time R export to `.h5ad`/`.mtx`
> (R snippet provided), then read that here. This keeps the Python pipeline clean and avoids an
> rpy2/R-in-Python dependency you may not want.

# %% [code]
import tarfile

# --- 3a. INSPECT (no full extraction) ---
with tarfile.open(GRANJA_TAR, "r") as tf:
    members = tf.getnames()
print(f"{len(members)} members. First 20:")
for m in members[:20]:
    print("  ", m)

# keep only members belonging to the healthy BMMC samples we want
def member_for_samples(members, samples):
    keep = [m for m in members if any(s in m for s in samples)]
    return keep
bmmc_members = member_for_samples(members, GRANJA_BMMC_SAMPLES)
print(f"\nMembers matching {GRANJA_BMMC_SAMPLES}: {len(bmmc_members)}")
for m in bmmc_members[:20]:
    print("  ", m)


# %% [code]
# --- 3b. R export snippet (run ONCE in R if the tar holds .rds SummarizedExperiments) ---
# Writes one .h5ad per BMMC sample that this notebook can then read directly.
R_EXPORT = r"""
# ---- run in R (>= 4.1) ----
#if (!requireNamespace("BiocManager", quietly = TRUE))
#    install.packages("BiocManager")

#BiocManager::install(c("SummarizedExperiment","SingleCellExperiment","zellkonverter"))

#BiocManager::install("SummarizedExperiment")
#BiocManager::install("SingleCellExperiment")
#BiocManager::install("zellkonverter")

library(SummarizedExperiment)
library(SingleCellExperiment)
library(zellkonverter)

export_se_to_h5ad <- function(se, out) {
  # inspect first — VERIFY these prints before trusting the output
  message("class: ", class(se))
  se <- SummarizedExperiment(assays = list(counts = se))
  message("assays: ", paste(assayNames(se), collapse = ", "))
  message("dim (genes x cells): ", paste(dim(se), collapse = " x "))
  message("rownames (genes) e.g.: ", paste(head(rownames(se), 3), collapse = ", "))
  message("colData cols: ", paste(colnames(colData(se)), collapse = ", "))

  sce <- if (is(se, "SingleCellExperiment")) se else as(se, "SingleCellExperiment")

  # pick the RNA counts assay by NAME (not position); fall back to first
  pref  <- c("counts", "RNA", "X", "GEX", "logcounts")
  aname <- intersect(pref, assayNames(sce))
  aname <- if (length(aname)) aname[1] else assayNames(sce)[1]
  message("using assay -> X: ", aname)

  # writeH5AD preserves rownames -> var_names and colData -> obs (incl. BioClassification)
  zellkonverter::writeH5AD(sce, file = out, X_name = aname)
}

# ---- PATH A (recommended): one curated, annotated file ----
se <- readRDS("./data/Granja_data/scRNA-Healthy-Hematopoiesis-191120.rds")
#export_se_to_h5ad(se, "./data/Granja_data/GSE139369_RAW/extracted/granja_healthy_all.h5ad")
# then in Python, subset to BMMC using the sample/group colData column you saw printed,
# e.g. granja = granja[granja.obs["<sample_col>"].str.contains("BMMC")].copy()

library(SingleCellExperiment)
library(zellkonverter)

# 1. Promote SummarizedExperiment to SingleCellExperiment
sce_object <- as(se, "SingleCellExperiment")

# 2. Convert to AnnData
#adata <- zellkonverter::as_AnnData(sce_object)

library(anndataR)
write_h5ad(sce_object, "./data/Granja_data/extracted/granja_healthy_all.h5ad")

# ---- PATH B (your GEO tar): loop the per-GSM CITE_BMMC RDS files ----
# tar -xf GSE139369_RAW.tar -C ./data/Granja_data/extracted   # do this once in the shell
#rds_files <- list.files("./data/Granja_data/extracted",
#                        pattern = "CITE_BMMC.*\\.rds$", full.names = TRUE)
#rds_files <- list.files("./data/Granja_data/extracted",
#                        pattern = "scRNA_BMMC.*\\.rds$", full.names = TRUE)

#for (f in rds_files) export_se_to_h5ad(readRDS(f), sub("\\.rds$", ".h5ad", f))

#print(R_EXPORT)
#print("\n# After running the R snippet, point GRANJA_H5ADS at the produced .h5ad files.")
"""

print(R_EXPORT)
print("\n# After running the R snippet, point GRANJA_H5ADS at the produced .h5ad files.")


# %% [code]
# --- 3c. Python-side loader (dispatch on what your tar actually contains) ---
from scipy.io import mmread

def load_granja_bmmc():
    '''Return a single concatenated healthy-BMMC RNA AnnData.

    Adjust the branch to match the member list printed in 3a.
    '''
    GRANJA_EXTRACT.mkdir(parents=True, exist_ok=True)

    # OPTION A: you exported .h5ad via the R snippet in 3b
    GRANJA_H5ADS = sorted(GRANJA_EXTRACT.glob("CITE_BMMC*.h5ad"))
    if GRANJA_H5ADS:
        parts = [sc.read_h5ad(str(p)) for p in GRANJA_H5ADS]
        for p, src in zip(parts, GRANJA_H5ADS):
            p.obs["granja_sample"] = src.stem
        a = ad.concat(parts, join="outer", label="granja_sample",
                      keys=[p.stem for p in GRANJA_H5ADS], index_unique="-")
        return a

    # OPTION B: tar holds 10x triplets (matrix.mtx[.gz] + barcodes + features/genes) per sample
    # Extract only the BMMC members, then read each triplet with sc.read_mtx / read_10x_mtx.
    #with tarfile.open(GRANJA_TAR, "r") as tf:
    #    for m in bmmc_members:
    #        tf.extract(m, path=GRANJA_EXTRACT)
    # ... assemble triplets here based on the actual filenames you saw in 3a ...
    raise NotImplementedError(
        "Inspect the 3a member list and complete OPTION A (recommended for .rds) "
        "or OPTION B (if 10x triplets). See the R export snippet in 3b for .rds inputs."
    )

#granja = load_granja_bmmc()

GRANJA_PATH = './data/Granja_data/extracted/granja_healthy_all.h5ad'

granja = sc.read_h5ad(str(GRANJA_PATH))
granja.var_names_make_unique(); granja.obs_names_make_unique()
print("Granja healthy BMMC:", granja.shape)


# %% [code]
granja.X = granja.layers['counts'].copy()


# %% [code]
print(granja)
print(granja.X)
print(granja.X.min())
print(granja.X.max())


# %% [code]
# --- 3d. Annotate Granja cells ---
# Granja ships healthy cell-type annotations (BioClassification labels). If present in obs, use them;
# otherwise transfer labels from van Galen later (see Section 9 label-transfer) or annotate with markers.
GR_CT_COL = find_col(granja, ["BioClassification", "CellType", "celltype", "cell_type",
                              "ClusterNames", "annotation"])
if GR_CT_COL is None:
    print("No native Granja cell-type column found — you can either:")
    print("  (a) attach Granja's published BioClassification labels (recommended), or")
    print("  (b) leave Granja cell types as 'unknown' (-1) and let CTM label-transfer fill them.")
    granja.obs["celltype_src"] = "unknown"
else:
    print("Granja cell-type column:", GR_CT_COL)
    granja.obs["celltype_src"] = granja.obs[GR_CT_COL].astype(str)


# %% [code]
print(granja.obs['BioClassification'])
print(granja.obs['Group'])
print(granja.obs['Clusters'])


# %% [markdown]
## 4 · Harmonize cohorts onto one RNA modality

Both cohorts become a **single RNA AnnData** on the intersection of genes, with a `dataset`
obs flag (`AML_vanGalen` vs `Granja_healthy`). Preprocessing follows the AML config:
`normalize_total(1e4)` → `log1p` → optional HVG → `scale(max=10)`, Gaussian likelihood.

> **Leakage note.** HVG selection and scaling are fit on the **training split only** and applied
> to val/test (the AML config sets `fit_on_train: true`). For a first exploratory pass you may
> fit on all cells; switch to train-only before any reported numbers.

> **Platform batch effect.** Seq-Well (van Galen) vs 10x (Granja) is the dominant technical axis
> here — larger than donor variation. CTM bridges biology via labels; the **adversarial dataset
> head** in `H_CTM` is what actively fights the platform effect. Expect `SE`/`CTM` to leave some
> dataset separation (this is your "mature x-data mix" metric).

# %% [code]
print(vg.X.min())
print(vg.X.max())


# %% [code]
print(vg)
print(vg.raw)
print(vg.raw.X.min())
print(vg.raw.X.max())


# %% [code]
vg.layers['pre-normed_and_scaled'] = vg.X.copy()
#vg.X = vg.layers['counts'].copy()
vg.X = vg.raw.X.copy()


# %% [code]
vg.layers['counts'] = vg.raw.X.copy()


# %% [code]
# tag cohorts
vg.obs["dataset"] = "AML_vanGalen"
granja.obs["dataset"] = "Granja_healthy"
vg.obs["malignancy"] = "malignant_or_aml"     # refine per-cell with van Galen's malignant calls if available
granja.obs["malignancy"] = "healthy"

# unify the cell-type label column name across cohorts
vg.obs["celltype_src"] = vg.obs[VG_CT_COL].astype(str)

# raw counts snapshot BEFORE normalization (needed if you switch to NB/ZINB later)
for a in (vg, granja):
    a.layers["counts"] = a.X.copy()

# concat on shared genes
shared_genes = vg.var_names.intersection(granja.var_names)
print("shared genes:", len(shared_genes))
assert len(shared_genes) > 1000, "Too few shared genes — check gene-symbol casing/format."
rna = ad.concat(
    {"AML_vanGalen": vg[:, shared_genes], "Granja_healthy": granja[:, shared_genes]},
    join="inner", label="dataset_concat", index_unique="-",
)
rna.obs_names_make_unique()
print("combined RNA:", rna.shape)
print(rna.obs["dataset"].value_counts())


# %% [code]
# preprocess (fit on TRAIN split only to avoid leakage)
rng = np.random.default_rng(0)
idx = np.arange(rna.n_obs); rng.shuffle(idx)
n_tr = int(0.8 * rna.n_obs); n_va = int(0.1 * rna.n_obs)
train_idx, val_idx, test_idx = idx[:n_tr], idx[n_tr:n_tr+n_va], idx[n_tr+n_va:]
np.savez(OUTDIR / "splits_seed0.npz", train_idx=train_idx, val_idx=val_idx, test_idx=test_idx)
is_train = np.zeros(rna.n_obs, bool); is_train[train_idx] = True

sc.pp.normalize_total(rna, target_sum=1e4)
sc.pp.log1p(rna)
rna.raw = rna  # log-space snapshot for plotting/DE

if USE_HVG:
    sc.pp.highly_variable_genes(rna[is_train].copy(), flavor="seurat_v3",
                                n_top_genes=N_HVG, layer="counts")
    # transfer HVG mask back (seurat_v3 needs counts); recompute on full then restrict to train-derived set
    hv = sc.pp.highly_variable_genes(rna, flavor="seurat_v3", n_top_genes=N_HVG,
                                     layer="counts", inplace=False, subset=False)
    rna = rna[:, hv["highly_variable"].to_numpy()].copy()
    print("after HVG:", rna.shape)

# scale using train-set mean/std, then apply to all (manual to avoid leakage)
Xtr = to_dense(rna[is_train].X)
mu  = Xtr.mean(0, keepdims=True)
sd  = Xtr.std(0, keepdims=True); sd[sd == 0] = 1.0
rna.X = np.clip((to_dense(rna.X) - mu) / sd, -10, 10).astype(np.float32)
print("RNA input space ready:", rna.X.shape, "dtype", rna.X.dtype)


# %% [markdown]
## 5 · Encode the label modalities and head targets

We build integer codes (pandas `Categorical`, which UniVI reads via `.cat.codes`):
- `celltype_code` — harmonized cell type across cohorts (the **CTM** bridge). Granja "unknown" → `-1`.
- `dataset_code`, `malignancy_code` — extra hierarchical levels for **H_CTM**.
- `{HERO}_code` — binary mutation target for the **MUT_CTM_M** head (van Galen calls only; Granja and
  uncalled van Galen cells = `-1`, i.e. masked).

> Harmonizing cell-type vocabularies across cohorts is the single most consequential manual step
> (van Galen and Granja name HSPC states differently). A starter mapping dict is provided — **edit it
> to your ontology.** Everything unmapped becomes `-1` and is simply not supervised (CTM still places
> those cells via RNA).

# %% [code]
# ---- harmonize cell-type vocabulary across cohorts (EDIT THIS MAP to your ontology) ----
CT_HARMONIZE = {
    # ---- van Galen: normal + malignant-like (strip "-like" for cell-type axis) ----
    # Malignancy is captured by PredictionRefined / malignancy_code separately.
    # Keeping "-like" blasts in the same canonical bucket as their normal counterpart
    # is the point of CTM — it lets the label bridge pull HSC-like blasts near healthy
    # HSCs so their developmental identity is preserved in the latent.
    "HSC":          "HSC",        "HSC-like":      "HSC",
    "Prog":         "Prog",       "Prog-like":     "Prog",
    "GMP":          "GMP",        "GMP-like":      "GMP",
    "ProMono":      "ProMono",    "ProMono-like":  "ProMono",
    "Mono":         "Mono",       "Mono-like":     "Mono",
    "cDC":          "cDC",        "cDC-like":      "cDC",
    "pDC":          "pDC",
    "earlyEry":     "Ery_early",
    "lateEry":      "Ery_late",
    "ProB":         "ProB",
    "B":            "B",
    "Plasma":       "Plasma",
    "T":            "T",
    "CTL":          "T",          # CD8 effector; van Galen has no CD4/CD8 resolution so T is all we can claim
    "NK":           "NK",

    # ---- Granja BioClassification ----
    "01_HSC":        "HSC",
    "02_Early.Eryth": "Ery_early",
    "03_Late.Eryth":  "Ery_late",
    "04_Early.Baso":  "Baso",     # no van Galen equivalent; label preserved, no cross-dataset CTM pull
    "05_CMP.LMPP":   "Prog",      # multi-potent progenitor ~ van Galen Prog (most uncertain mapping below)
    "06_CLP.1":      "CLP",       # no van Galen equivalent; lymphoid progenitor upstream of ProB
    "07_GMP":        "GMP",
    "08_GMP.Neut":   "GMP",       # neutrophil-biased GMP, merge with GMP
    "09_pDC":        "pDC",
    "10_cDC":        "cDC",
    "11_CD14.Mono.1": "Mono",
    "12_CD14.Mono.2": "Mono",
    "13_CD16.Mono":   "Mono",
    "14_Unk":         None,        # → -1 in CTM, excluded from label alignment
    "15_CLP.2":       "CLP",
    "16_Pre.B":       "ProB",      # Pre-B ≈ van Galen ProB (adjacent stages; see note 4)
    "17_B":           "B",
    "18_Plasma":      "Plasma",
    "19_CD8.N":       "T",
    "20_CD4.N1":      "T",
    "21_CD4.N2":      "T",
    "22_CD4.M":       "T",
    "23_CD8.EM":      "T",
    "24_CD8.CM":      "T",
    "25_NK":          "NK",
    "26_Unk":         None,        # → -1
}

def canon_ct(x):
    x = str(x)
    if x in ("unknown", "nan", "None", "NA"): return None
    return CT_HARMONIZE.get(x, x)   # unmapped passes through; set to None to drop instead

rna.obs["celltype_canon"] = [canon_ct(v) for v in rna.obs["celltype_src"]]
cats = sorted([c for c in rna.obs["celltype_canon"].dropna().unique()])
rna.obs["celltype_code"] = pd.Categorical(rna.obs["celltype_canon"], categories=cats)
n_ct = len(cats)
print(f"{n_ct} harmonized cell types:", cats)

rna.obs["dataset_code"]    = pd.Categorical(rna.obs["dataset"])
rna.obs["malignancy_code"] = pd.Categorical(rna.obs["malignancy"])
n_dataset    = rna.obs["dataset_code"].cat.categories.size
n_malig      = rna.obs["malignancy_code"].cat.categories.size

# carry the hero-gene mutation code from van Galen (Granja cells = -1 unknown)
hero_full = np.full(rna.n_obs, -1, int)
vg_mask = rna.obs["dataset"].to_numpy() == "AML_vanGalen"
# align van Galen codes to the concatenated order via obs_names prefix match
vg_codes = pd.Series(vg.obs[f"{HERO_GENE}_code"].to_numpy(), index=vg.obs_names)
hero_full = np.array([
    int(vg_codes.get(n.rsplit("-", 1)[0], -1)) if d == "AML_vanGalen" else -1
    for n, d in zip(rna.obs_names, rna.obs["dataset"])
])
rna.obs[f"{HERO_GENE}_code"] = hero_full
print(f"{HERO_GENE} on combined:", pd.Series(hero_full).value_counts().to_dict(), "(-1=masked)")


# %% [markdown]
## 6 · Build `MultiModalDataset` + loaders

For categorical **modalities** (CTM/H_CTM) we reuse the same RNA `AnnData` under extra dict keys and
mark them categorical+`obs` via `modality_cfgs`; the dataset reads `obs[obs_key].cat.codes`.
For class **heads** (SE cell-type head, MUT_CTM_M mutation head, adversarial dataset head) we pass
integer targets via `labels=`.

# %% [code]
def codes_of(col):
    c = rna.obs[col]
    return c.cat.codes.to_numpy().astype(np.int64) if hasattr(c, "cat") else np.asarray(c, np.int64)

celltype_codes  = codes_of("celltype_code")     # -1 already for unmapped (Categorical NaN -> -1)
dataset_codes   = codes_of("dataset_code")
malig_codes     = codes_of("malignancy_code")
hero_codes      = rna.obs[f"{HERO_GENE}_code"].to_numpy().astype(np.int64)
print("unlabeled cell types (code -1):", int((celltype_codes < 0).sum()))


# %% [markdown]
## 7 · Variant config factory — the core of your CTM experiments

Each branch returns `(univi_cfg, adata_dict, modality_cfgs, labels)` so training is identical
downstream. This is where your `SE / CTM / H_CTM / MUT_CTM_M` map onto verified UniVI objects.

# %% [code]
RNA_ENC = [1024, 512, 256, 128, 64]   # AML config architecture
RNA_DEC = [64, 128, 256, 512, 1024]
LAB_ENC, LAB_DEC = [128, 64], [64, 128]

def build_config(variant):
    rna_mod = ModalityConfig("rna", rna.n_vars, RNA_ENC, RNA_DEC, likelihood="gaussian")
    modalities = [rna_mod]
    adata_dict = {"rna": rna}
    modality_cfgs = [rna_mod]
    class_heads = []
    labels = {}

    if variant == "SE":
        # supervised cell-type head on the RNA latent (no label modality)
        class_heads.append(ClassHeadConfig(name="celltype", n_classes=n_ct, dropout=0.10,
                                           loss_weight=1.0, ignore_index=-1, from_mu=True, warmup=0))
        labels["celltype"] = celltype_codes

    elif variant == "CTM":
        # cell type as a full categorical modality, gamma-aligned to RNA
        ct_mod = ModalityConfig("celltype", n_ct, LAB_ENC, LAB_DEC, likelihood="categorical",
                                input_kind="obs", obs_key="celltype_code", ignore_index=-1)
        modalities.append(ct_mod); modality_cfgs.append(ct_mod)
        adata_dict["celltype"] = rna   # input_kind='obs' reads rna.obs['celltype_code']

    elif variant == "H_CTM":
        # hierarchical: cell type + malignancy as categorical modalities, dataset as ADVERSARIAL head
        ct_mod = ModalityConfig("celltype", n_ct, LAB_ENC, LAB_DEC, likelihood="categorical",
                                input_kind="obs", obs_key="celltype_code", ignore_index=-1)
        mal_mod = ModalityConfig("malignancy", n_malig, [64, 32], [32, 64], likelihood="categorical",
                                 input_kind="obs", obs_key="malignancy_code", ignore_index=-1)
        modalities += [ct_mod, mal_mod]; modality_cfgs += [ct_mod, mal_mod]
        adata_dict["celltype"] = rna; adata_dict["malignancy"] = rna
        # adversarial dataset head -> gradient reversal pushes the latent to mix cohorts
        class_heads.append(ClassHeadConfig(name="dataset", n_classes=n_dataset, loss_weight=0.2,
                                           ignore_index=-1, from_mu=True, warmup=10, dropout=0.10,
                                           adversarial=True, adv_lambda=1.0))
        labels["dataset"] = dataset_codes

    elif variant == "MUT_CTM_M":
        # CTM + masked binary mutation head (ignore_index=-1 masks uncalled / Granja cells)
        ct_mod = ModalityConfig("celltype", n_ct, LAB_ENC, LAB_DEC, likelihood="categorical",
                                input_kind="obs", obs_key="celltype_code", ignore_index=-1)
        modalities.append(ct_mod); modality_cfgs.append(ct_mod)
        adata_dict["celltype"] = rna
        class_heads.append(ClassHeadConfig(name=HERO_GENE, n_classes=2, head_type="binary", dropout=0.10,
                                           loss_weight=1.0, ignore_index=-1, from_mu=True, warmup=50))
        labels[HERO_GENE] = hero_codes
    else:
        raise ValueError(variant)

    cfg = UniVIConfig(
        latent_dim=LATENT_DIM, beta=BETA, gamma=GAMMA,
        encoder_dropout=0.10, decoder_dropout=0.05,
        encoder_batchnorm=True, decoder_batchnorm=False,
        kl_anneal_start=0, kl_anneal_end=0, align_anneal_start=0, align_anneal_end=0,
        modalities=modalities,
        class_heads=class_heads if class_heads else None,
    )
    cfg.validate()
    return cfg, adata_dict, modality_cfgs, labels

univi_cfg, adata_dict, modality_cfgs, labels = build_config(VARIANT)
print(f"[{VARIANT}] modalities:", [m.name for m in univi_cfg.modalities],
      "| heads:", [h.name for h in (univi_cfg.class_heads or [])])


# %% [markdown]
## 8 · Train

# %% [code]
align_paired_obs_names(adata_dict)  # all entries share obs_names (same RNA object)

dataset = MultiModalDataset(
    adata_dict=adata_dict,
    X_key={k: "X" for k in adata_dict},
    modality_cfgs=modality_cfgs,
    labels=labels if labels else None,
    device=None,
)

def make_loader(ix, shuffle):
    return DataLoader(Subset(dataset, ix), batch_size=BATCH_SIZE, shuffle=shuffle,
                      num_workers=0, collate_fn=collate_multimodal_xy_recon)
train_loader = make_loader(train_idx, True)
val_loader   = make_loader(val_idx, False)

#model = UniVIMultiModalVAE(univi_cfg, loss_mode="v1", v1_recon="avg",
#                           normalize_v1_terms=True).to(device)
model = UniVIMultiModalVAE(univi_cfg, loss_mode="v2").to(device)

# give the supervised heads human-readable class names (for predict_heads / confusion matrices)
if VARIANT == "SE":
    model.set_head_label_names("celltype", list(cats))

train_cfg = TrainingConfig(n_epochs=N_EPOCHS, batch_size=BATCH_SIZE, lr=LR,
                           weight_decay=WEIGHT_DECAY, device=device,
                           early_stopping=True, patience=50, best_epoch_warmup=0, log_every=25)

trainer = UniVITrainer(model=model, train_loader=train_loader, val_loader=val_loader,
                       train_cfg=train_cfg, device=device)
history = trainer.fit()
print("best epoch:", getattr(trainer, "best_epoch", None))

torch.save({"model_state_dict": model.state_dict(), "model_config": univi_cfg,
            "train_cfg": train_cfg, "history": history,
            "best_epoch": getattr(trainer, "best_epoch", None)},
           OUTDIR / f"univi_{VARIANT}.pt")


# %% [markdown]
## 9 · Encode the latent space and visualize

We encode the RNA modality posterior for every cell (the cohorts are unpaired, so the RNA
encoder is the shared inference path). UMAPs colored by `dataset`, `celltype_canon`, `malignancy`,
and the `NPM1` code let you eyeball integration vs. cell-type structure vs. the mutation signal.

# %% [code]
model.eval()
Z = encode_adata(model, adata=rna, modality="rna", device=device,
                 layer=None, X_key="X", batch_size=1024,
                 latent="modality_mean", random_state=0)
rna.obsm["X_univi"] = Z
print("latent:", Z.shape)

rna.obs[f"{HERO_GENE}_label"] = pd.Categorical(
    np.where(rna.obs[f"{HERO_GENE}_code"].to_numpy() == 1, "MUT",
             np.where(rna.obs[f"{HERO_GENE}_code"].to_numpy() == 0, "WT", "unknown"))
)


# %% [code]
umap(rna, obsm_key="X_univi",
     color=["dataset"],
     legend="outside", legend_subset_topk=25,
     size=2,
     savepath=str(OUTDIR / f"umap_{VARIANT}.png"), show=True)

umap(rna, obsm_key="X_univi",
     color=["celltype_canon"],
     legend="outside", legend_subset_topk=25,
     size=2,
     savepath=str(OUTDIR / f"umap_{VARIANT}.png"), show=True)

umap(rna, obsm_key="X_univi",
     color=["malignancy"],
     legend="outside", legend_subset_topk=25,
     size=2,
     savepath=str(OUTDIR / f"umap_{VARIANT}.png"), show=True)

umap(rna, obsm_key="X_univi",
     color=[f"{HERO_GENE}_label"],
     legend="outside", legend_subset_topk=25,
     size=2,
     savepath=str(OUTDIR / f"umap_{VARIANT}.png"), show=True)


# %% [code]
print(rna)
print(vg)


# %% [code]
vghvg = vg[:, shared_genes].copy()


# %% [code]
vghvg = vghvg[:, hv["highly_variable"].to_numpy()].copy()


# %% [code]
print(vghvg)


# %% [code]
model.eval()
Zvg = encode_adata(model, adata=vghvg, modality="rna", device=device,
                   layer=None, X_key="X", batch_size=1024,
                   latent="modality_mean", random_state=0)
vghvg.obsm["X_univi"] = Zvg
print("latent:", Zvg.shape)

vghvg.obs[f"{HERO_GENE}_label"] = pd.Categorical(
    np.where(vghvg.obs[f"{HERO_GENE}_code"].to_numpy() == 1, "MUT",
             np.where(vghvg.obs[f"{HERO_GENE}_code"].to_numpy() == 0, "WT", "unknown"))
)


# %% [code]
umap(vghvg, obsm_key="X_univi",
     color=["CellType"],
     legend="outside", legend_subset_topk=25,
     size=4,
     savepath=str(OUTDIR / f"umap_{VARIANT}.png"), show=True)

umap(vghvg, obsm_key="X_univi",
     color=[f"{HERO_GENE}_label"],
     legend="outside", legend_subset_topk=25,
     size=4,
     savepath=str(OUTDIR / f"umap_{VARIANT}.png"), show=True)


# %% [code]
print(granja)


# %% [code]
granjahvg = granja[:, shared_genes].copy()


# %% [code]
granjahvg = granjahvg[:, hv["highly_variable"].to_numpy()].copy()


# %% [code]
print(granjahvg)


# %% [code]
model.eval()
Zgranja = encode_adata(model, adata=granjahvg, modality="rna", device=device,
                       layer=None, X_key="X", batch_size=1024,
                       latent="modality_mean", random_state=0)

granjahvg.obsm["X_univi"] = Zgranja
print("latent:", Zgranja.shape)


# %% [code]
umap(granjahvg, obsm_key="X_univi",
     color=["BioClassification"],
     legend="outside", legend_subset_topk=25,
     size=4,
     savepath=str(OUTDIR / f"umap_{VARIANT}.png"), show=True)


# %% [markdown]
## 10 · Evaluation — reproducing your results-table columns

Helpers below compute the four columns from your preliminary table, on the **test split**:

- **broad ct purity** — fraction of a cell's k-NN (in latent) sharing its cell type.
- **mature x-data mix** — among *mature* (non-HSPC) cells, the fraction of k-NN drawn from the
  *other* cohort (your cross-dataset mixing target; 0 = no mixing, ~0.5 = ideal balanced mixing).
- **malig F1** — macro-F1 of a k-NN classifier predicting healthy-vs-AML from the latent.
- **NPM1 F1 / AUC** — from the trained mutation head (`MUT_CTM_M`) via `predict_heads`, or a
  latent k-NN over labeled cells otherwise.

`evaluate_alignment` (shipped) additionally gives recall@k, mixing/entropy, and a label-transfer
confusion matrix if you want the manuscript-style readouts.

# %% [code]
from sklearn.neighbors import NearestNeighbors
from sklearn.metrics import f1_score, roc_auc_score

Zt = Z[test_idx]
ct_t   = rna.obs["celltype_canon"].to_numpy()[test_idx]
ds_t   = rna.obs["dataset"].to_numpy()[test_idx]
mal_t  = rna.obs["malignancy"].to_numpy()[test_idx]
MATURE_EXCLUDE = {"HSPC"}   # treat these labels as immature/progenitor

def knn_idx(Zq, k=30):
    nn = NearestNeighbors(n_neighbors=k + 1).fit(Zq)
    return nn.kneighbors(Zq, return_distance=False)[:, 1:]

def broad_ct_purity(Zq, labels, k=30):
    lab = np.asarray(labels); nb = knn_idx(Zq, k)
    ok = np.array([(lab[row] == lab[i]).mean() for i, row in enumerate(nb)])
    keep = lab != "None"
    return float(ok[keep].mean())

def cross_dataset_mix(Zq, ds, ct, k=30, mature_exclude=MATURE_EXCLUDE):
    ds = np.asarray(ds); ct = np.asarray(ct); nb = knn_idx(Zq, k)
    is_mature = ~np.isin(ct, list(mature_exclude))
    frac_other = np.array([(ds[row] != ds[i]).mean() for i, row in enumerate(nb)])
    return float(frac_other[is_mature].mean()) if is_mature.any() else float("nan")

def malig_f1_knn(Zq, mal, k=15):
    mal = np.asarray(mal); nb = knn_idx(Zq, k)
    pred = np.array([pd.Series(mal[row]).mode().iloc[0] for row in nb])
    return float(f1_score(mal, pred, average="macro"))

row = {
    "variant": VARIANT,
    "broad_ct_purity": round(broad_ct_purity(Zt, ct_t), 3),
    "mature_x_data_mix": round(cross_dataset_mix(Zt, ds_t, ct_t), 3),
    "malig_F1": round(malig_f1_knn(Zt, mal_t), 3),
}
print(row)


# %% [code]
# NPM1 readout
hero_t = rna.obs[f"{HERO_GENE}_code"].to_numpy()[test_idx]
labeled = hero_t >= 0
if VARIANT == "MUT_CTM_M" and labeled.sum() > 0:
    with torch.no_grad():
        x = torch.tensor(to_dense(rna[test_idx].X), dtype=torch.float32, device=device)
        probs = model.predict_heads({"rna": x}, return_probs=True)
    p = probs[HERO_GENE].detach().cpu().numpy()
    p1 = p[:, 1] if p.ndim == 2 and p.shape[1] == 2 else p.ravel()
    yhat = (p1 >= 0.5).astype(int)
    row[f"{HERO_GENE}_F1"]  = round(float(f1_score(hero_t[labeled], yhat[labeled])), 3)
    try:
        row[f"{HERO_GENE}_AUC"] = round(float(roc_auc_score(hero_t[labeled], p1[labeled])), 3)
    except ValueError:
        row[f"{HERO_GENE}_AUC"] = float("nan")
else:
    row[f"{HERO_GENE}_F1"] = None
print(row)

results = pd.DataFrame([row])
results.to_csv(OUTDIR / f"metrics_{VARIANT}.csv", index=False)
results


# %% [code]
# OPTIONAL: manuscript-style alignment readout between the two cohorts in the shared latent
aml_mask     = (ds_t == "AML_vanGalen")
healthy_mask = (ds_t == "Granja_healthy")
'''
align = evaluate_alignment(
    Z1=Zt[aml_mask], Z2=Zt[healthy_mask], metric="euclidean",
    recall_ks=(1, 5, 10), k_mixing=20, k_entropy=30,
    labels_source=ct_t[aml_mask], labels_target=ct_t[healthy_mask],
    compute_bidirectional_transfer=True, k_transfer=15, json_safe=True,
)
print({k: align[k] for k in list(align)[:8]})
'''

# %% [markdown]
## 11 · Held-out cell-type masking (your experiment #1)

Tests whether the model learns RNA→latent structure or just memorizes labels: mask a fraction of
`celltype_code` to `-1` (UniVI's ignore value), retrain, and compare k-NN purity of **masked** vs
**label-visible** cells. A small gap ⇒ the γ-aligned label modality transferred structure into the
RNA encoder rather than being memorized.

# %% [code]
def run_masked_celltype(mask_pct, variant="CTM", epochs=400):
    rna_m = rna.copy()
    codes = celltype_codes.copy()
    vis = codes >= 0
    masked = np.zeros(rna_m.n_obs, bool)
    if mask_pct > 0:
        cand = np.where(vis)[0]
        sel = np.random.default_rng(0).choice(cand, int(mask_pct * cand.size), replace=False)
        codes[sel] = -1; masked[sel] = True
    rna_m.obs["celltype_code"] = pd.Categorical.from_codes(codes, categories=cats)

    ct_mod = ModalityConfig("celltype", n_ct, LAB_ENC, LAB_DEC, likelihood="categorical",
                            input_kind="obs", obs_key="celltype_code", ignore_index=-1)
    cfg = UniVIConfig(latent_dim=LATENT_DIM, beta=BETA, gamma=GAMMA,
                      encoder_dropout=0.10, decoder_dropout=0.05,
                      modalities=[ModalityConfig("rna", rna_m.n_vars, RNA_ENC, RNA_DEC, "gaussian"), ct_mod])
    cfg.validate()
    add = {"rna": rna_m, "celltype": rna_m}; align_paired_obs_names(add)
    ds = MultiModalDataset(add, X_key={k: "X" for k in add}, modality_cfgs=[cfg.modalities[0], ct_mod])
    tl = DataLoader(Subset(ds, train_idx), batch_size=BATCH_SIZE, shuffle=True,
                    collate_fn=collate_multimodal_xy_recon)
    #m = UniVIMultiModalVAE(cfg, loss_mode="v1", v1_recon="avg", normalize_v1_terms=True).to(device)
    m = UniVIMultiModalVAE(cfg, loss_mode="v2").to(device)
    UniVITrainer(m, tl, None, TrainingConfig(n_epochs=epochs, lr=LR, device=device, log_every=50),
                 device=device).fit()
    Zm = encode_adata(m, adata=rna_m, modality="rna", device=device, X_key="X",
                      batch_size=1024, latent="modality_mean")
    nb = knn_idx(Zm[test_idx], 30); lab = rna.obs["celltype_canon"].to_numpy()[test_idx]
    pur = np.array([(lab[r] == lab[i]).mean() for i, r in enumerate(nb)])
    mtest = masked[test_idx]; vtest = (~masked[test_idx]) & (rna.obs["celltype_canon"].to_numpy()[test_idx] != "None")
    return {"mask_pct": mask_pct,
            "purity_masked":  round(float(pur[mtest].mean()), 3) if mtest.any() else None,
            "purity_visible": round(float(pur[vtest].mean()), 3) if vtest.any() else None}

# NOTE: this retrains once per mask level — run with a reduced epoch budget first.
heldout = pd.DataFrame([run_masked_celltype(p, epochs=400) for p in (0.0, 0.2, 0.5, 0.8)])
heldout["gap"] = heldout["purity_visible"] - heldout["purity_masked"]
heldout.to_csv(OUTDIR / "heldout_celltype_masking.csv", index=False)
heldout


# %% [markdown]
## 12 · Recommended UniVI-native analyses for *this* question

These are shipped UniVI capabilities (README + `scripts/`) that map directly onto the
AML-vs-healthy problem and your stated follow-ups. I'd prioritize the first two — they address
the two things that genuinely complicate integrating leukemia with healthy marrow.

**(a) "Do-not-integrate" detection for malignant blasts.**
`scripts/run_do_not_integrate_detection.py` flags populations present in only one cohort by
latent nearest-neighbor distances. AML blast states (e.g. progenitor-like) often have **no healthy
counterpart**, and forcing them to integrate is exactly the HSC/progenitor disagreement Granja et al.
reported when projecting van Galen. Use this to *quantify* which AML populations should stay separate
rather than assuming everything aligns.

# %% [code]
# Sketch: distance-based non-integration score (mirrors run_do_not_integrate_detection.py logic)
def non_integration_score(Z, ds, ct, ref="Granja_healthy", k=15):
    ds = np.asarray(ds)
    ref_mask = ds == ref
    nn = NearestNeighbors(n_neighbors=k).fit(Z[ref_mask])
    d_query, _ = nn.kneighbors(Z[~ref_mask])           # AML cells -> healthy neighbors
    d_ctrl,  _ = nn.kneighbors(Z[ref_mask])            # healthy -> healthy (control)
    thr = np.percentile(d_ctrl.mean(1), 95)
    flagged = d_query.mean(1) > thr
    out = pd.DataFrame({"celltype": np.asarray(ct)[~ref_mask], "flagged": flagged})
    return out.groupby("celltype")["flagged"].mean().sort_values(ascending=False)

print("Fraction of AML cells flagged 'non-integrating' by cell type:")
print(non_integration_score(Zt, ds_t, ct_t))


# %% [markdown]
**(b) Composition / frequency robustness.**
`scripts/run_frequency_robustness.py` quantifies how alignment degrades as cell-type *composition*
and *frequency* diverge between cohorts. AML marrow is blast-dominated while healthy marrow is not, so
your cohorts have very different compositions — this script tells you how much of any residual
mis-mixing is the method vs. the biology (and is the rigorous backing for your "mature x-data mix" number).

**(c) Mutation-head interpretability (HOX check).**
`feature_importance_for_head` attributes the `NPM1` head back to genes. NPM1-mutant AML has a
characteristic *HOX* signature — confirming HOX genes top the attribution is a strong biological
validation that the head learned real signal, not batch.

# %% [code]
# Which genes drive the NPM1 head (only meaningful for MUT_CTM_M)
if VARIANT == "MUT_CTM_M":
    from univi.interpretability import feature_importance_for_head
    xb = torch.tensor(to_dense(rna[test_idx][:2000].X), dtype=torch.float32, device=device)
    imp = feature_importance_for_head(model, {"rna": xb}, head_name=HERO_GENE)  # see signature in interpretability.py
    top = pd.Series(np.asarray(imp).ravel(), index=rna.var_names).sort_values(ascending=False).head(25)
    print(f"Top genes for {HERO_GENE} head (look for HOXA/HOXB, MEIS1, PBX3):")
    print(top)


# %% [markdown]
**(d) Generative per-cell-type sampling**, `(e)` MoE gating diagnostics, and `(f)`
`diagnostics.export_supplemental_table_s1(...)` for a reproducibility table — all documented in the
README and worth including in a methods write-up.

**(g) Bridge mapping.** If you later want a *reference→query* framing instead of joint training,
train on Granja healthy alone and project van Galen with `encode_adata(..., latent="modality_mean")`
(the README "bridge mapping" path), then read off where AML cells land relative to the healthy manifold.

# %% [markdown]
## 13 · Verified-vs-assumed notes (read before reporting numbers)

**Verified against UniVI v0.4.7 source** (cloned from `github.com/Ashford-A/UniVI`): all imports,
`UniVIConfig`/`ModalityConfig`/`ClassHeadConfig`/`TrainingConfig` fields, the categorical-modality
path (`input_kind="obs"` + `obs_key` reading `.cat.codes`, `ignore_index=-1`), class-head label
passing via `labels=`, `MultiModalDataset(..., modality_cfgs=...)`, `collate_multimodal_xy_recon`,
`encode_adata`, `evaluate_alignment`, and `umap`. The config objects in this notebook were
instantiated and `.validate()`-ed against the real dataclasses.

**Assumptions you must confirm in your environment:**
1. **Granja tar internals.** Listed by GEO as *"TAR (of RDS, TSV)"*; I could not open your 20.4 GB
   archive here. Section 3a inspects it; complete OPTION A (R→h5ad) or OPTION B (10x triplets)
   based on the printed members.
2. **van Galen obs columns.** Assumed `CellType` for labels and `"<GENE> ..."` columns for genotype,
   matching the authors' AML notebook — verify against your file's `obs.columns`.
3. **Cell-type harmonization (`CT_HARMONIZE`).** A starter map only — edit to your ontology. This is
   the step that most affects integration quality.
4. **Platform batch effect (Seq-Well vs 10x) is real and large.** `SE`/`CTM` will likely leave some
   cohort separation; `H_CTM`'s adversarial dataset head is the lever that fights it. Don't read
   residual separation as a bug before trying H_CTM and the frequency-robustness script.
5. **Epoch budgets.** `N_EPOCHS=3000` matches the AML config but is heavy; start at 300–800 for a
   first pass, and reduce the held-out-masking epochs likewise.


# %% [code]


# %% [code]


# %% [code]


# %% [code]


# %% [code]


# %% [code]


# %% [code]


# %% [code]


# %% [code]


# %% [code]


# %% [code]


# %% [code]


# %% [code]


# %% [code]


# %% [code]


# %% [code]


# %% [code]


# %% [code]


# %% [code]


# %% [code]
```
