## Supplemental Code for "Unifying multimodal single-cell data with a mixture-of-experts *β*-variational autoencoder framework": Fig02_CITE_paired.ipynb

```
# %% [markdown]
# UniVI manuscript - Figure 2 generation reproducible workflow
### CITE-seq embeddings before/after integration; modality mixing statistics; label-transfer confusion matrices and per-class performance.

Andrew Ashford, Pathways + Omics Group, Oregon Health & Science University, Portland, OR - 12/6/2025

This Jupyter Notebook will house the end-to-end workflow to generate the panels in Figure 2 of our manuscript, "Unifying multimodal single-cell data with a mixture-of-experts β-variational autoencoder framework" which is currently being revised for Genome Research and is available currently on bioRxiv at the following link: https://www.biorxiv.org/content/10.1101/2025.02.28.640429v1.full


# %% [markdown]
### Import modules

# %% [code]
# Import non-UniVI modules
import os
import sys
from pathlib import Path

import numpy as np
import pandas as pd
import scanpy as sc
import anndata as ad
import torch

from torch.utils.data import DataLoader, Subset


# %% [code]
# Import required UniVI modules
from univi import (
    ModalityConfig,
    UniVIConfig,
    TrainingConfig,
    UniVIMultiModalVAE,
    matching,
    UniVITrainer,
    write_univi_latent,
    MultiModalDataset,
)

import univi as uv
import univi.evaluation as ue
import univi.plotting as up


# %% [code]
# Double check UniVI module version
print("Installed version is univi v" + str(uv.__version__))


# %% [markdown]
### Specify device to use for model

Set "device" - preferably device should be "cuda" for speedier model implementation/training (NOTE: can use "mps" on MacBook M1 chips to use their GPU). Requires GPU and the correct packages/versions.


# %% [code]
print("torch:", torch.__version__)
print("torch.cuda.is_available():", torch.cuda.is_available())
device = "cuda" if torch.cuda.is_available() else "cpu"
print("Using device:", device)


# %% [markdown]
### Specify file paths

Where data lives.

# %% [code]
DATA_ROOT = Path("/home/groups/precepts/ashforda/UniVI_v2/UniVI_older-non_git/data/Hao_CITE-seq_data")

RNA_PATH = DATA_ROOT / "Hao_RNA_data.h5ad"
ADT_PATH = DATA_ROOT / "Hao_ADT_data.h5ad"

print("RNA file:", RNA_PATH)
print("ADT file:", ADT_PATH)


# %% [markdown]
### Read in data

Read data into AnnData objects using the paths in the code chunk above.

# %% [code]
rna = sc.read_h5ad(RNA_PATH)
adt = sc.read_h5ad(ADT_PATH)

print(rna)
print(adt)

print("RNA obs names head:", rna.obs_names[:5].tolist())
print("ADT obs names head:", adt.obs_names[:5].tolist())


# %% [markdown]
### Align cells between RNA and ADT

Make sure the cell indices are aligned so UniVI knows which samples are paired.

# %% [code]
# Intersect barcodes
common_cells = rna.obs_names.intersection(adt.obs_names)
print("Common cells:", len(common_cells))

rna = rna[common_cells].copy()
adt = adt[common_cells].copy()

# Make sure order matches
adt = adt[rna.obs_names].copy()

assert np.array_equal(rna.obs_names.values, adt.obs_names.values)
print("obs_names aligned between RNA and ADT.")


# %% [markdown]
### Stratify the data by celltype so that we train a balanced/generalizeable model

Also specifying all the data preprocessing functions below, will be used in their own respective sections.

# %% [code]
# -----------------------------
# Stratified split with cap
# -----------------------------
def stratified_split(
    idx,
    labels,
    frac_train=0.8,
    frac_val=0.1,
    seed=0,
    max_per_label=None,
    unused_to_test=True,
):
    """
    For each label:
      - shuffle its indices
      - keep up to max_per_label as "used"
      - split used -> train/val/test by fractions
      - leftover beyond max_per_label -> unused
    If unused_to_test=True, unused is appended into test and not returned separately.
    """
    rng = np.random.default_rng(seed)
    idx = np.asarray(idx)
    labels = np.asarray(labels)

    train_idx, val_idx, test_idx, unused_idx = [], [], [], []

    # stable label order (preserves category order if categorical, else sorted unique)
    uniq = pd.unique(labels)

    for lab in uniq:
        m = idx[labels == lab]
        rng.shuffle(m)

        if max_per_label is not None:
            used = m[:max_per_label]
            leftover = m[max_per_label:]
            if leftover.size:
                unused_idx.append(leftover)
        else:
            used = m

        n = used.size
        if n == 0:
            continue

        n_train = int(frac_train * n)
        n_val   = int(frac_val * n)
        # remainder -> test
        train_idx.append(used[:n_train])
        val_idx.append(used[n_train:n_train + n_val])
        test_idx.append(used[n_train + n_val:])

    train_idx = np.concatenate(train_idx) if train_idx else np.array([], dtype=int)
    val_idx   = np.concatenate(val_idx)   if val_idx   else np.array([], dtype=int)
    test_idx  = np.concatenate(test_idx)  if test_idx  else np.array([], dtype=int)
    unused_idx = np.concatenate(unused_idx) if unused_idx else np.array([], dtype=int)

    if unused_to_test and unused_idx.size:
        test_idx = np.concatenate([test_idx, unused_idx])
        unused_idx = np.array([], dtype=int)

    return train_idx, val_idx, test_idx, unused_idx


# %% [code]
# -----------------------------
# Small helper functions
# -----------------------------
def _ensure_counts_layer(adata, layer_name="counts"):
    if layer_name in adata.layers:
        return
    # Fall back to X as counts if needed (only if X is actually raw counts!)
    adata.layers[layer_name] = adata.X.copy()

def _subset_by_idx_pair(rna, adt, idx):
    # Assumes rna/adt are already paired in the same order
    return rna[idx].copy(), adt[idx].copy()

def _clr_normalize_dense(X, eps=1e-8):
    """
    CLR per cell: log1p(x) - mean(log1p(x)) across features.
    Works for dense or sparse; returns dense float32 array.
    """
    if sp.issparse(X):
        X = X.toarray()
    X = X.astype(np.float32, copy=False)
    X = np.log1p(X)
    X = X - X.mean(axis=1, keepdims=True)
    return X

def preprocess_citeseq_splits(
    rna_train, adt_train,
    rna_val, adt_val,
    rna_test, adt_test,
    *,
    rna_counts_layer="counts",
    adt_counts_layer="counts",
    n_hvg=2000,
    target_sum=1e4,
    rna_make_log1p=True,
    adt_make_clr=True,
):
    """
    Train-fit, then transform val/test the same way.

    RNA:
      - ensures counts layer
      - HVGs learned on train only (Seurat v3 if possible)
      - normalize_total + log1p (applied to all splits)
      - subsets to HVGs for all splits

    ADT:
      - ensures counts layer
      - CLR per cell (applied to all splits) -> stored in .X (dense float32)
    """
    # ensure counts layers exist
    for a in (rna_train, rna_val, rna_test):
        _ensure_counts_layer(a, rna_counts_layer)
    for a in (adt_train, adt_val, adt_test):
        _ensure_counts_layer(a, adt_counts_layer)

    # ---- RNA: pick HVGs on TRAIN only
    rna_train_tmp = rna_train.copy()
    rna_train_tmp.X = rna_train_tmp.layers[rna_counts_layer]

    try:
        sc.pp.highly_variable_genes(
            rna_train_tmp,
            n_top_genes=int(n_hvg),
            flavor="seurat_v3",
            layer=None,   # we set .X already
        )
    except Exception:
        sc.pp.highly_variable_genes(
            rna_train_tmp,
            n_top_genes=int(n_hvg),
            flavor="seurat",
            layer=None,
        )

    hvg_mask = rna_train_tmp.var["highly_variable"].to_numpy()
    hvg_genes = rna_train_tmp.var_names[hvg_mask].tolist()

    # ---- RNA: normalize/log1p + subset to HVGs (train/val/test)
    def _rna_transform(adata):
        # subset genes first to minimize memory
        ad = adata[:, hvg_genes].copy()
        X = ad.layers[rna_counts_layer]

        # normalize_total on counts
        if sp.issparse(X):
            cell_sums = np.asarray(X.sum(axis=1)).ravel()
            scale = (float(target_sum) / np.maximum(cell_sums, 1e-12)).astype(np.float32)
            Xn = X.multiply(scale[:, None])
        else:
            cell_sums = X.sum(axis=1, keepdims=True)
            Xn = (X / np.maximum(cell_sums, 1e-12)) * float(target_sum)

        if rna_make_log1p:
            if sp.issparse(Xn):
                Xn = Xn.tocsr(copy=True)
                Xn.data = np.log1p(Xn.data).astype(np.float32, copy=False)
            else:
                Xn = np.log1p(Xn).astype(np.float32, copy=False)

        ad.X = Xn
        # keep original counts too (still subsetted)
        ad.layers[rna_counts_layer] = ad.layers[rna_counts_layer]
        return ad

    rna_train_pp = _rna_transform(rna_train)
    rna_val_pp   = _rna_transform(rna_val)
    rna_test_pp  = _rna_transform(rna_test)

    # ---- ADT: CLR into .X (train/val/test)
    def _adt_transform(adata):
        ad = adata.copy()
        Xc = ad.layers[adt_counts_layer]
        if adt_make_clr:
            ad.X = _clr_normalize_dense(Xc)
        else:
            # keep counts as X (dense float32)
            ad.X = Xc.toarray().astype(np.float32) if sp.issparse(Xc) else Xc.astype(np.float32)
        return ad

    adt_train_pp = _adt_transform(adt_train)
    adt_val_pp   = _adt_transform(adt_val)
    adt_test_pp  = _adt_transform(adt_test)

    return (rna_train_pp, adt_train_pp,
            rna_val_pp,   adt_val_pp,
            rna_test_pp,  adt_test_pp,
            hvg_genes)


# %% [code]
# Check the counts of each celltype.l1 in the data
print(rna.obs["celltype.l1"].value_counts())


# %% [code]
# Check the counts of each celltype.l2 in the data
print(rna.obs["celltype.l2"].value_counts())


# %% [code]
# Check the counts of each celltype.l3 in the data
print(rna.obs["celltype.l3"].value_counts())


# %% [code]
# -----------------------------
# Use it on Hao CITE-seq
# -----------------------------

labels = rna.obs["celltype.l1"].astype(str).to_numpy()
idx = np.arange(rna.n_obs)

train_idx, val_idx, test_idx, unused_idx = stratified_split(
    idx, labels,
    frac_train=0.8, frac_val=0.1, seed=0,
    max_per_label=2000,
    unused_to_test=True,      # set True if you want unused merged into test
)

# paired splits
rna_train, adt_train = _subset_by_idx_pair(rna, adt, train_idx)
rna_val,   adt_val   = _subset_by_idx_pair(rna, adt, val_idx)
rna_test,  adt_test  = _subset_by_idx_pair(rna, adt, test_idx)

if unused_idx.size:
    rna_unused, adt_unused = _subset_by_idx_pair(rna, adt, unused_idx)
else:
    rna_unused, adt_unused = None, None


# %% [markdown]
### Data preprocessing

* RNA preprocessing (log1p + HVG + scale → Gaussian decoder)

* ADT preprocessing (CLR + scale → Gaussian decoder)

Running the preprocessing functions specified above. Of note, we're performing the preprocessing per-train/val/test split.

# %% [code]
import scipy.sparse as sp

# preprocessing (fit on train, apply to val/test)
(rna_train_pp, adt_train_pp,
 rna_val_pp,   adt_val_pp,
 rna_test_pp,  adt_test_pp,
 hvg_genes) = preprocess_citeseq_splits(
    rna_train, adt_train,
    rna_val,   adt_val,
    rna_test,  adt_test,
    n_hvg=2000,
    target_sum=1e4,
    rna_make_log1p=True,
    adt_make_clr=True,
)


# %% [code]
'''
# optionally preprocess "unused" with same HVGs + same transforms (no refit)
if rna_unused is not None:
    # reuse the same transform logic by treating unused as "test"
    (rna_unused_pp, adt_unused_pp,
     _, _, _, _, _) = preprocess_citeseq_splits(
        rna_train, adt_train,     # only used to learn HVGs (already learned here, but fine)
        rna_unused, adt_unused,
        rna_unused, adt_unused,
        n_hvg=len(hvg_genes),
        target_sum=1e4,
        rna_make_log1p=True,
        adt_make_clr=True,
    )
    # the function returns in (train,val,test) slots; we passed unused into val/test
    # so grab one of them:
    # rna_unused_pp == rna_train_pp from that call; use the "val" one instead:
    # Instead, do a tiny manual transform with the already-computed hvg_genes:
    # (If you want this cleaned up, I can give you a one-liner helper.)
'''

# %% [code]
# Sanity check above preprocessed data objects - start with RNA
print(rna_train_pp)
print(rna_val_pp)
print(rna_test_pp)


# %% [code]
# Now sanity check ADT data objects
print(adt_train_pp)
print(adt_val_pp)
print(adt_test_pp)


# %% [markdown]
### Wrap into MultiModalDataset & DataLoaders

# %% [code]
from univi.data import align_paired_obs_names

pin_memory = (device == "cuda")

# Make sure each split is paired and ordered the same across modalities
# This was erroring out due to a function bug, fixed it and should be fixed by manuscript publication
#train_dict = align_paired_obs_names({"rna": rna_train_pp, "adt": adt_train_pp})
#val_dict   = align_paired_obs_names({"rna": rna_val_pp,   "adt": adt_val_pp})
#test_dict  = align_paired_obs_names({"rna": rna_test_pp,  "adt": adt_test_pp})

# Using this instead for now since we know the data are already paired from above code
assert (rna_train_pp.obs_names == adt_train_pp.obs_names).all()
assert (rna_val_pp.obs_names   == adt_val_pp.obs_names).all()
assert (rna_test_pp.obs_names  == adt_test_pp.obs_names).all()

train_dict = {"rna": rna_train_pp, "adt": adt_train_pp}
val_dict   = {"rna": rna_val_pp,   "adt": adt_val_pp}
test_dict  = {"rna": rna_test_pp,  "adt": adt_test_pp}

# Build datasets (CPU tensors; trainer/model moves to GPU)
train_ds = MultiModalDataset(adata_dict=train_dict, X_key="X", device=None)
val_ds   = MultiModalDataset(adata_dict=val_dict,   X_key="X", device=None)
test_ds  = MultiModalDataset(adata_dict=test_dict,  X_key="X", device=None)

batch_size = 256
num_workers = 0

train_loader = DataLoader(
    train_ds,
    batch_size=batch_size,
    shuffle=True,
    num_workers=num_workers,
    pin_memory=pin_memory,
    drop_last=False,
)

val_loader = DataLoader(
    val_ds,
    batch_size=batch_size,
    shuffle=False,
    num_workers=num_workers,
    pin_memory=pin_memory,
    drop_last=False,
)

test_loader = DataLoader(
    test_ds,
    batch_size=batch_size,
    shuffle=False,
    num_workers=num_workers,
    pin_memory=pin_memory,
    drop_last=False,
)

print("n_train / n_val / n_test:", train_ds.n_cells, val_ds.n_cells, test_ds.n_cells)
print("batches:", len(train_loader), len(val_loader), len(test_loader))

# sanity check one batch
x = next(iter(train_loader))
print({k: v.shape for k, v in x.items()})


# %% [markdown]
### UniVI configs

# %% [code]
univi_cfg = UniVIConfig(
    latent_dim=30,
    beta=1.5,
    gamma=5.0,
    encoder_dropout=0.1,
    decoder_dropout=0.0,
    encoder_batchnorm=True,
    decoder_batchnorm=False,
    kl_anneal_start=0,
    kl_anneal_end=0,
    align_anneal_start=0,
    align_anneal_end=0,
    modalities=[
        ModalityConfig(
            name="rna",
            input_dim=rna_train_pp.n_vars,
            encoder_hidden=[512, 256, 128],
            decoder_hidden=[128, 256, 512],
            likelihood="gaussian",
        ),
        ModalityConfig(
            name="adt",
            input_dim=adt_train_pp.n_vars,
            encoder_hidden=[128, 64],
            decoder_hidden=[64, 128],
            likelihood="gaussian",
        ),
    ],
)


# %% [markdown]
If you want raw-count NB/ZINB instead, you’d:
* Put raw counts in .layers["counts"]
* Set X_key="counts" in the dataset
* Use likelihood="nb" or "zinb" above

But for this "Figure 2" notebook, the scaled Gaussian setup is nicely stable and good for the best-integrated latent space.


# %% [markdown]
### Instantiate model and model objective

# %% [code]
model = UniVIMultiModalVAE(
    univi_cfg,
    loss_mode="v1",      # cross-recon + cross-posterior alignment - "lite"/"v2" just use single joint latent and L2 norm latent means for alignment,
                         # leads to weaker alignment but depends less on paired feature reconstruction in training.
    #v1_recon="cross",   # full k→j cross-recon
    v1_recon="avg",      # average of cross-and self- recon
    #v1_recon_mix=0.5,
    normalize_v1_terms=True,
).to(device)


# %% [markdown]
### Instantiate TrainingConfig & trainer

# %% [code]
train_cfg = TrainingConfig(
    n_epochs=3000,
    batch_size=batch_size,
    lr=1e-3,
    weight_decay=1e-4,
    device=device,
    log_every=20,
    grad_clip=5.0,
    num_workers=0,
    seed=42,
    early_stopping=True,
    patience=50,
    min_delta=0.0,
)

print(train_cfg)
print(univi_cfg)


# %% [code]
trainer = UniVITrainer(
    model=model,
    train_loader=train_loader,
    val_loader=val_loader,
    train_cfg=train_cfg,
    device=device,
)

print(trainer)


# %% [markdown]
### Fit the model

# %% [code]
history = trainer.fit()

# history is a dict with keys like "train_loss", "val_loss", "beta", "gamma"
print("Training finished.")
print("Best val loss:", np.min(history["val_loss"]))


# %% [code]
import matplotlib.pyplot as plt

plt.figure()
plt.plot(history["train_loss"], label="train")
plt.plot(history["val_loss"], label="val")
plt.xlabel("Epoch")
plt.ylabel("Loss")
plt.legend()
plt.title("UniVI training (CITE-seq)")
plt.show()


# %% [markdown]
### Write latent z back to AnnData

# %% [code]
print("Test batches:", len(test_loader))


# %% [code]
# Make sure adata_dict has the *same ordering* as during training
adata_dict = {
    "rna": rna_test_pp,
    "adt": adt_test_pp,
}

Z = write_univi_latent(
    model,
    adata_dict,
    obsm_key="X_univi",    # will be added to each AnnData in adata_dict
    batch_size=512,
    device=device,
    use_mean=True,         # deterministic: use encoder means instead of noisy samples
    #use_mean=False,       # Stochastic
)

print("UniVI latent shape:",  Z.shape)
print("rna.obsm keys:",       rna_test_pp.obsm.keys())
print("adt.obsm keys:",       adt_test_pp.obsm.keys())


# %% [markdown]
Now both rna and adt have a shared latent:

* rna_test_pp.obsm["X_univi"]

* adt_test_pp.obsm["X_univi"]


# %% [markdown]
### Quick UMAP + visualization

You can now treat X_univi like any other embedding.

# %% [code]
# Check out many of each cell type are in the test set for level 2 cell annotations
print(rna_test_pp.obs["celltype.l2"].value_counts())


# %% [code]
# Use RNA AnnData just to get a UMAP; ADT has matching .obsm["X_univi"]
rna_univi = rna_test_pp.copy()
rna_univi.obsm["X_univi"] = rna_test_pp.obsm["X_univi"]

# wipe any stale neighbors artifacts
rna_univi.uns.pop("neighbors", None)
rna_univi.obsp.pop("connectivities", None)
rna_univi.obsp.pop("distances", None)


# %% [code]
sc.pp.neighbors(rna_univi, use_rep="X_univi", n_neighbors=15, key_added="univi")
sc.tl.umap(rna_univi, neighbors_key="univi")


# %% [code]
# Copy back UMAP to ADT so they share coords
adt_test_pp.obsm["X_univi_umap"] = rna_univi.obsm["X_umap"].copy()


# %% [code]
# Scanpy defaults (affects sc.pl.*)
sc.set_figure_params(
    figsize=(10, 8),   # bigger canvas
    dpi=100,            # on-screen sharpness
    dpi_save=300,       # saved file sharpness
    fontsize=10,
    frameon=False,
)

# Matplotlib defaults (affects plt.*)
plt.rcParams.update({
    "figure.figsize": (10, 8),
    "figure.dpi": 100,
    "savefig.dpi": 300,
    "savefig.bbox": "tight",
    "savefig.pad_inches": 0.1,
    "axes.titlesize": 12,
    "axes.labelsize": 8,
    "legend.fontsize": 10,
})


# %% [code]
# Example: color by cell type and modality (if you have those obs fields)
sc.pl.umap(
    rna_univi,
    color=["celltype.l1"],
    frameon=False,
    size=3.0,
    wspace=0.4,
)


# %% [code]
# Make sure X_univi exists in both
assert "X_univi" in rna_test_pp.obsm and "X_univi" in adt_test_pp.obsm
assert rna_test_pp.obsm["X_univi"].shape == adt_test_pp.obsm["X_univi"].shape  # paired case

rna_u = rna_test_pp.copy()
adt_u = adt_test_pp.copy()
rna_u.obs["modality"] = "rna"
adt_u.obs["modality"] = "adt"

# concatenate (keeps obs, stacks cells)
combo = ad.concat([rna_u, adt_u], join="outer", label="modality", keys=["rna","adt"], index_unique="-")

# IMPORTANT: carry over the latent explicitly (concat won't always merge obsm how you want)
combo.obsm["X_univi"] = np.vstack([rna_u.obsm["X_univi"], adt_u.obsm["X_univi"]])


# %% [code]
# neighbors on the latent
sc.pp.neighbors(combo, use_rep="X_univi", n_neighbors=15)


# %% [code]
# umap on the latent using the neighbors
sc.tl.umap(combo)


# %% [code]
sc.pl.umap(
    combo,
    color=["modality"],
    frameon=False,
    size=3.0,
    wspace=0.4,
)


# %% [code]
sc.pl.umap(
    combo,
    color=["celltype.l1"],
    frameon=False,
    size=3.0,
    wspace=0.4,
)


# %% [code]
sc.pl.umap(
    combo,
    color=["celltype.l2"],
    frameon=False,
    size=3.0,
    wspace=0.4,
)


# %% [code]
sc.pl.umap(
    combo,
    color=["celltype.l3"],
    frameon=False,
    size=3.0,
    wspace=0.4,
)


# %% [markdown]
### All evaluation metrics beyond just the UMAP embeddings of the shared test set latent

# %% [code]
import json

from sklearn.neighbors import NearestNeighbors
from sklearn.metrics import confusion_matrix, accuracy_score, f1_score

# ----------------------------
# User settings
# ----------------------------
outdir = "figures/Figure2_CITEseq_metrics_reproducibility"
os.makedirs(outdir, exist_ok=True)

celltype_key = "celltype.l2"   # change to celltype.l1 / celltype.l3 as needed
k_mix = 20                     # k for modality mixing
k_lt  = 15                     # k for label transfer kNN
seed = 42


# %% [code]
from sklearn.neighbors import NearestNeighbors
from sklearn.metrics import confusion_matrix, accuracy_score, f1_score

# ----------------------------
# Metrics helpers
# ----------------------------
def foscttm_chunked(Z1, Z2, block=512):
    """
    Exact FOSCTTM computed in blocks to avoid NxN memory blowups.
    Assumes 1:1 pairing between rows i in Z1 and Z2.
    """
    Z1 = np.asarray(Z1, dtype=np.float32)
    Z2 = np.asarray(Z2, dtype=np.float32)
    assert Z1.shape == Z2.shape
    n = Z1.shape[0]

    Z2_T = Z2.T
    n2 = np.sum(Z2 * Z2, axis=1)  # (n,)

    fos = np.empty(n, dtype=np.float32)

    for i0 in range(0, n, block):
        i1 = min(i0 + block, n)
        A = Z1[i0:i1]  # (b, d)
        n1 = np.sum(A * A, axis=1)[:, None]  # (b,1)

        d2 = n1 + n2[None, :] - 2.0 * (A @ Z2_T)  # (b,n)

        true = d2[np.arange(i1 - i0), np.arange(i0, i1)]
        fos[i0:i1] = (d2 < true[:, None]).sum(axis=1) / (n - 1)

    return float(fos.mean()), float(fos.std(ddof=1) / np.sqrt(n))


def modality_mixing_score(Z, modality_labels, k=20, metric="euclidean"):
    """
    Vanilla modality mixing:
    Mean fraction of kNN neighbors that differ in modality.
    Use this for *non-duplicated* sets (e.g., concatenated modality-specific embeddings).
    """
    Z = np.asarray(Z, dtype=np.float32)
    modality_labels = np.asarray(modality_labels)

    n = Z.shape[0]
    if n <= 1:
        return 0.0

    k_eff = int(min(max(int(k), 1), n - 1))
    nn = NearestNeighbors(n_neighbors=k_eff + 1, metric=metric)
    nn.fit(Z)
    nbrs = nn.kneighbors(Z, return_distance=False)[:, 1:]  # drop self

    same = (modality_labels[nbrs] == modality_labels[:, None])
    return float((~same).mean())


def modality_mixing_score_excluding_pairs(Z, modality_labels, cell_ids, k=20, metric="euclidean"):
    """
    Pair-aware modality mixing for 'combo' style stacked data (same cell appears twice: RNA + ADT),
    where the fused embedding may be identical (or extremely close) for the paired duplicates.

    Computes: for each row, the fraction of neighbors from the other modality,
    AFTER removing the paired duplicate (same cell_id) from its neighbor list.

    cell_ids must map each row to a shared cell identifier (same for RNA+ADT copy).
    """
    Z = np.asarray(Z, dtype=np.float32)
    modality_labels = np.asarray(modality_labels)
    cell_ids = np.asarray(cell_ids).astype(str)

    n = Z.shape[0]
    if n <= 2:
        return 0.0

    # Need enough neighbors so we can drop self + paired duplicate and still have k
    k_eff = int(min(max(int(k), 1), n - 2))
    nn = NearestNeighbors(n_neighbors=k_eff + 2, metric=metric)
    nn.fit(Z)
    nbrs = nn.kneighbors(Z, return_distance=False)[:, 1:]  # drop self

    # Build "pair index": for each row i, pair[i] is the index of the other modality copy
    first = {}
    pair = np.full(n, -1, dtype=np.int64)
    for i, cid in enumerate(cell_ids):
        if cid in first:
            j = first[cid]
            pair[i] = j
            pair[j] = i
        else:
            first[cid] = i

    frac_other = np.empty(n, dtype=np.float32)
    for i in range(n):
        neigh = nbrs[i]

        # remove paired duplicate if present
        pj = pair[i]
        if pj != -1:
            neigh = neigh[neigh != pj]

        neigh = neigh[:k_eff]
        frac_other[i] = (modality_labels[neigh] != modality_labels[i]).mean()

    return float(frac_other.mean())


def knn_label_transfer(Z_source, y_source, Z_target, y_target, k=15, metric="euclidean"):
    """
    kNN label transfer: predict y_target from neighbors in Z_source.
    Returns predictions, accuracy, macro-F1, confusion matrix.
    """
    Z_source = np.asarray(Z_source, dtype=np.float32)
    Z_target = np.asarray(Z_target, dtype=np.float32)
    y_source = np.asarray(y_source, dtype=str)
    y_target = np.asarray(y_target, dtype=str)

    nn = NearestNeighbors(n_neighbors=k, metric=metric)
    nn.fit(Z_source)
    nbrs = nn.kneighbors(Z_target, return_distance=False)

    preds = []
    for inds in nbrs:
        votes = y_source[inds]
        vals, cnts = np.unique(votes, return_counts=True)
        preds.append(vals[np.argmax(cnts)])
    preds = np.asarray(preds, dtype=str)

    acc = float(accuracy_score(y_target, preds))
    macro_f1 = float(f1_score(y_target, preds, average="macro"))
    classes = np.unique(np.concatenate([y_source, y_target]))
    cm = confusion_matrix(y_target, preds, labels=classes)
    return preds, acc, macro_f1, cm, classes


# %% [code]
from univi.evaluation import encode_adata

Z_rna = encode_adata(model, rna_test_pp, modality="rna",
                     latent="modality_mean", device=device, batch_size=1024)
Z_adt = encode_adata(model, adt_test_pp, modality="adt",
                     latent="modality_mean", device=device, batch_size=1024)

key_rna = "encode_adata(modality_mean)"
key_adt = "encode_adata(modality_mean)"

rna_test_pp.obsm["X_univi_rna"] = Z_rna
adt_test_pp.obsm["X_univi_adt"] = Z_adt


# %% [code]
print("max|Z_rna-Z_adt|:", float(np.max(np.abs(Z_rna - Z_adt))))
print("mean L2(Z_rna-Z_adt):", float(np.mean(np.linalg.norm(Z_rna - Z_adt, axis=1))))


# %% [code]
# ---------- PRE-FLIGHT CHECK + FIX (run once right before metrics/plots) ----------
def _preflight_align_for_metrics(
    rna_test_pp, adt_test_pp, Z_rna, Z_adt, celltype_key="celltype.l2"
):
    import numpy as np

    # 1) basic shape checks
    print("rna_test_pp n_obs:", rna_test_pp.n_obs)
    print("adt_test_pp n_obs:", adt_test_pp.n_obs)
    print("Z_rna shape:", np.asarray(Z_rna).shape)
    print("Z_adt shape:", np.asarray(Z_adt).shape)

    # 2) enforce pairing by obs_names intersection (preserves RNA order)
    common = rna_test_pp.obs_names.intersection(adt_test_pp.obs_names)
    if len(common) != rna_test_pp.n_obs or len(common) != adt_test_pp.n_obs:
        print(f"[preflight] restricting to common paired cells: {len(common)}")

    rna_al = rna_test_pp[common].copy()
    adt_al = adt_test_pp[common].copy()
    adt_al = adt_al[rna_al.obs_names].copy()  # force identical order

    # 3) slice Z arrays to match aligned AnnData order (assumes Z were computed in the same order)
    # If your Z were computed from rna_test_pp/adt_test_pp directly, this is safe.
    # If not, you *must* recompute Z after alignment.
    if np.asarray(Z_rna).shape[0] != rna_al.n_obs or np.asarray(Z_adt).shape[0] != adt_al.n_obs:
        raise ValueError(
            "[preflight] Z arrays do not match aligned AnnData n_obs. "
            "Recompute Z_rna/Z_adt from rna_al/adt_al."
        )

    # 4) labels must come from the SAME objects you predict on
    labels_rna = rna_al.obs[celltype_key].astype(str).to_numpy()
    labels_adt = adt_al.obs[celltype_key].astype(str).to_numpy()

    # 5) final asserts (this prevents your exact error)
    assert rna_al.n_obs == adt_al.n_obs
    assert (rna_al.obs_names == adt_al.obs_names).all()
    assert len(labels_rna) == np.asarray(Z_rna).shape[0]
    assert len(labels_adt) == np.asarray(Z_adt).shape[0]

    print("[preflight] OK: paired, aligned, and lengths match.")
    return rna_al, adt_al, labels_rna, labels_adt


# run it:
rna_test_pp, adt_test_pp, labels_rna, labels_adt = _preflight_align_for_metrics(
    rna_test_pp, adt_test_pp, Z_rna, Z_adt, celltype_key=celltype_key
)


# %% [code]
# ----------------------------
# Compute metrics (Figure 2)
# ----------------------------

# 0) Fused latent (if you truly want it)
Z_fused = np.asarray(combo.obsm["X_univi"])
mods_fused = combo.obs["modality"].to_numpy()

# derive cell_ids that are identical for RNA+ADT copies
# works with index_unique="-" that produces "<orig>-rna" / "<orig>-adt"
cell_ids = combo.obs_names.to_series().str.rsplit("-", n=1).str[0].to_numpy()

mix_fused = modality_mixing_score_excluding_pairs(Z_fused, mods_fused, cell_ids, k=k_mix)

# 1) Modality-specific latents (Z_rna, Z_adt) already computed above

# 2) FOSCTTM on modality-specific latents (subsample ok)
n_total = int(rna_test_pp.n_obs)
n_fos = int(min(20000, n_total))     # subsample size for FOSCTTM eval
rng = np.random.default_rng(seed)
sub = rng.choice(n_total, size=n_fos, replace=False)

fos_rna_adt, fos_sem = foscttm_chunked(Z_rna[sub], Z_adt[sub], block=512)

# 3) Modality mixing on modality-specific latents
Z_concat = np.vstack([Z_rna, Z_adt])
mods = np.array(["rna"] * Z_rna.shape[0] + ["adt"] * Z_adt.shape[0], dtype=object)
mix_modality_specific = modality_mixing_score(Z_concat, mods, k=k_mix)

# 4) Label transfer (modality-specific)
# Make labels that match each embedding 1:1
labels_rna = rna_test_pp.obs[celltype_key].astype(str).to_numpy()
labels_adt = adt_test_pp.obs[celltype_key].astype(str).to_numpy()

assert Z_rna.shape[0] == labels_rna.shape[0], (Z_rna.shape, labels_rna.shape)
assert Z_adt.shape[0] == labels_adt.shape[0], (Z_adt.shape, labels_adt.shape)

pred_adt, acc_r2a, f1_r2a, cm_r2a, classes = knn_label_transfer(
    Z_source=Z_rna, y_source=labels_rna,
    Z_target=Z_adt, y_target=labels_adt,
    k=k_lt
)
pred_rna, acc_a2r, f1_a2r, cm_a2r, _ = knn_label_transfer(
    Z_source=Z_adt, y_source=labels_adt,
    Z_target=Z_rna, y_target=labels_rna,
    k=k_lt
)

metrics = {
    "embedding_keys": {"rna": key_rna, "adt": key_adt, "fused": "combo.obsm['X_univi']"},
    "n_test": n_total,
    "celltype_key": celltype_key,

    "FOSCTTM_metric": "euclidean_sq",
    "FOSCTTM_subsample_n": n_fos,
    "FOSCTTM_rna_vs_adt_mean": float(fos_rna_adt),
    "FOSCTTM_rna_vs_adt_sem": float(fos_sem),

    "modality_mixing_k": int(k_mix),
    "modality_mixing_fused": float(mix_fused),
    "modality_mixing_modality_specific": float(mix_modality_specific),

    "label_transfer_k": int(k_lt),
    "label_transfer_rna_to_adt_acc": float(acc_r2a),
    "label_transfer_rna_to_adt_macroF1": float(f1_r2a),
    "label_transfer_adt_to_rna_acc": float(acc_a2r),
    "label_transfer_adt_to_rna_macroF1": float(f1_a2r),
}

print(json.dumps(metrics, indent=2))
with open(os.path.join(outdir, "figure2_reproducibility_metrics.json"), "w") as f:
    json.dump(metrics, f, indent=2)
pd.DataFrame([metrics]).to_csv(os.path.join(outdir, "figure2_reproducibility_metrics.csv"), index=False)


# %% [code]
from sklearn.neighbors import NearestNeighbors
from sklearn.metrics import precision_recall_fscore_support

# If plots ever don't show in classic notebook, run once:
# %matplotlib inline

# -------------------------
# show-or-save helper
# -------------------------
def _finish(fig=None, outpath=None, dpi=200, show=True, close=True):
    if outpath is not None:
        plt.savefig(outpath, dpi=dpi, bbox_inches="tight")
    if show:
        plt.show()
    if close:
        plt.close(fig if fig is not None else plt.gcf())

# -------------------------
# plots
# -------------------------
def plot_metrics_bar(
    metrics,
    keys,
    title="Figure 2 reproducibility summary metrics",
    outpath=None,
    err_keys=None,          # dict: metric_key -> error_metric_key
    default_err=np.nan,     # np.nan = no error bar drawn; 0.0 = zero-length
    capsize=4,
):
    vals = np.array([float(metrics[k]) for k in keys], dtype=float)

    if err_keys is None:
        yerr = np.full_like(vals, default_err, dtype=float)
    else:
        yerr = np.array(
            [float(metrics[err_keys[k]]) if k in err_keys and err_keys[k] in metrics else default_err
             for k in keys],
            dtype=float
        )

    fig = plt.figure(figsize=(10, 6))
    plt.bar(keys, vals, yerr=yerr, capsize=capsize)
    plt.xticks(rotation=45, ha="right")
    plt.title(title)
    plt.tight_layout()

    # inline “finish” behavior
    if outpath is not None:
        plt.savefig(outpath, dpi=300, bbox_inches="tight")
    plt.show()
    plt.close(fig)


def plot_confusion(cm, classes, title, normalize="true", outpath=None, cmap="viridis"):
    import numpy as np
    import matplotlib.pyplot as plt

    cm = np.asarray(cm, dtype=float)
    if normalize == "true":
        cm = cm / (cm.sum(axis=1, keepdims=True) + 1e-12)
        subtitle = "Row-normalized"
    elif normalize == "pred":
        cm = cm / (cm.sum(axis=0, keepdims=True) + 1e-12)
        subtitle = "Col-normalized"
    elif normalize == "all":
        cm = cm / (cm.sum() + 1e-12)
        subtitle = "Global-normalized"
    else:
        subtitle = "Counts"

    classes = np.asarray(classes, dtype=str)
    n = len(classes)

    fig, ax = plt.subplots(figsize=(10, 8))
    im = ax.imshow(cm, interpolation="nearest", cmap=cmap, aspect="auto")

    # kill any gridlines (including ones from styles)
    ax.grid(False)
    ax.minorticks_off()

    ax.set_title(f"{title}\n({subtitle})")
    ax.set_xlabel("Predicted")
    ax.set_ylabel("True")

    ax.set_xticks(np.arange(n))
    ax.set_yticks(np.arange(n))
    ax.set_xticklabels(classes, rotation=90)
    ax.set_yticklabels(classes)

    cbar = fig.colorbar(im, ax=ax)
    cbar.set_label("value")

    fig.tight_layout()

    if outpath is not None:
        fig.savefig(outpath, dpi=300, bbox_inches="tight")

    plt.show()
    plt.close(fig)


def plot_per_class_f1(y_true, y_pred, title="Per-class F1", outpath=None, top_n=None):
    y_true = np.asarray(y_true).astype(str)
    y_pred = np.asarray(y_pred).astype(str)
    classes = np.unique(np.concatenate([y_true, y_pred]))

    _, _, f1, support = precision_recall_fscore_support(
        y_true, y_pred, labels=classes, zero_division=0
    )

    df = pd.DataFrame({"class": classes, "f1": f1, "support": support})
    df = df.sort_values(["f1", "support"], ascending=[True, False])

    if top_n is not None:
        df = df.head(int(top_n))

    fig = plt.figure(figsize=(10, 0.35 * len(df) + 2.0))
    plt.barh(df["class"], df["f1"])
    plt.xlabel("F1")
    plt.title(title)
    plt.xlim(0, 1)
    plt.tight_layout()
    _finish(fig, outpath=outpath, show=True, close=True)
    return df

def plot_modality_mixing_hist(Z, mods, k=20, metric="euclidean", title="Modality mixing", outpath=None):
    Z = np.asarray(Z, dtype=np.float32)
    mods = np.asarray(mods)

    n = Z.shape[0]
    k_eff = int(min(max(int(k), 1), n - 1))

    nn = NearestNeighbors(n_neighbors=k_eff + 1, metric=metric)
    nn.fit(Z)
    nbrs = nn.kneighbors(Z, return_distance=False)[:, 1:]
    frac_other = (mods[nbrs] != mods[:, None]).mean(axis=1)

    fig = plt.figure(figsize=(7, 4.5))
    plt.hist(frac_other, bins=60)
    plt.xlabel("Fraction of kNN from other modality")
    plt.ylabel("# cells")
    plt.title(title)
    plt.tight_layout()
    _finish(fig, outpath=outpath, show=True, close=True)
    return frac_other

def plot_foscttm_sanity(Z1, Z2, idx, title="FOSCTTM sanity", outpath=None):
    Z1s = np.asarray(Z1[idx], dtype=np.float32)
    Z2s = np.asarray(Z2[idx], dtype=np.float32)

    d_true = np.sum((Z1s - Z2s) ** 2, axis=1)

    nn = NearestNeighbors(n_neighbors=51, metric="euclidean")
    nn.fit(np.asarray(Z2, dtype=np.float32))
    dist, _ = nn.kneighbors(Z1s)
    d_min = dist[:, 0] ** 2

    fig = plt.figure(figsize=(5.5, 5.5))
    plt.scatter(d_true, d_min, s=8, alpha=0.4)
    mx = np.percentile(np.concatenate([d_true, d_min]), 99)
    plt.plot([0, mx], [0, mx], linewidth=1)
    plt.xlim(0, mx); plt.ylim(0, mx)
    plt.xlabel("d(true match)^2")
    plt.ylabel("d(nearest neighbor)^2")
    plt.title(title)
    plt.tight_layout()
    _finish(fig, outpath=outpath, show=True, close=True)

def plot_paired_distance_hist(Z_rna, Z_adt, idx, title="Paired latent distance (subset)", outpath=None):
    d_pair = np.sqrt(np.sum((Z_rna[idx] - Z_adt[idx]) ** 2, axis=1))
    fig = plt.figure(figsize=(7, 4.5))
    plt.hist(d_pair, bins=80)
    plt.xlabel("||z_rna - z_adt||")
    plt.ylabel("# cells")
    plt.title(title)
    plt.tight_layout()
    _finish(fig, outpath=outpath, show=True, close=True)


# %% [code]
# 1) Summary metric bar
plot_metrics_bar(
    metrics,
    keys=[
        "FOSCTTM_rna_vs_adt_mean",
        "modality_mixing_fused",
        "modality_mixing_modality_specific",
        "label_transfer_rna_to_adt_acc",
        "label_transfer_rna_to_adt_macroF1",
    ],
    err_keys={
        #"FOSCTTM_rna_vs_adt_mean": "FOSCTTM_rna_vs_adt_sem",
        # add more here if you compute SEM/CI for them later
    },
    default_err=np.nan,   # no error bars for the others
    outpath=None,
)


# %% [code]
# 2) Confusion matrices (row-normalized)
plot_confusion(cm_r2a, classes, title=f"Label transfer (RNA → ADT), k={k_lt}", normalize="true", outpath=None)
plot_confusion(cm_a2r, classes, title=f"Label transfer (ADT → RNA), k={k_lt}", normalize="true", outpath=None)


# %% [code]
# 3) Per-class F1
_ = plot_per_class_f1(labels_adt, pred_adt, title="RNA → ADT: per-class F1", top_n=100, outpath=None)
_ = plot_per_class_f1(labels_rna, pred_rna, title="ADT → RNA: per-class F1", top_n=100, outpath=None)


# %% [code]
# 4) Modality-mixing distributions
_ = plot_modality_mixing_hist(Z_fused, mods_fused, k=k_mix, title=f"Mixing (fused latent), k={k_mix}", outpath=None)

Z_concat = np.vstack([Z_rna, Z_adt])
mods_concat = np.array(["rna"] * Z_rna.shape[0] + ["adt"] * Z_adt.shape[0], dtype=object)
_ = plot_modality_mixing_hist(Z_concat, mods_concat, k=k_mix, title=f"Mixing (modality-specific latents), k={k_mix}", outpath=None)


# %% [code]
# 5) FOSCTTM sanity + paired distance hist
sub_sanity = np.asarray(sub)[:min(5000, len(sub))]
plot_foscttm_sanity(Z_rna, Z_adt, sub_sanity, outpath=None)

sub_dist = np.asarray(sub)[:min(50000, len(sub))]
plot_paired_distance_hist(Z_rna, Z_adt, sub_dist, outpath=None)


# %% [code]
# 6) UMAPs (Scanpy already shows inline)
# (these are heavy at 294k cells; feel free to set size=1.5)
#sc.pl.umap(combo, color=["modality"], frameon=False, size=2.0)
#sc.pl.umap(combo, color=[celltype_key], frameon=False, size=2.0)


# %% [markdown]
#### Interactive 3-D UMAPs

# %% [code]
import plotly.express as px
import plotly.graph_objects as go

def plot_umap3d_interactive(
    combo,
    umap_key="X_umap_3d",         # <- default for your CITE-seq object
    color_by="celltype.l2",
    symbol_by="modality",
    hover_cols=("cell_id", "modality"),

    point_size=3,
    opacity=0.85,

    # paired lines between modalities (optional)
    draw_pair_lines=True,         # <- default False for typical CITE-seq (often not stacked)
    id_key="cell_id",             # will auto-create from obs_names if missing
    modality_key="modality",
    mod_a=None,                   # if None, infer from available modalities
    mod_b=None,
    line_sample=None,             # None = all
    random_state=0,
    line_width=2.5,
    line_opacity=0.35,
    line_color="rgba(0,0,0,0.25)",

    width=900,
    height=700,
):
    # --- embedding checks ---
    assert umap_key in combo.obsm, f"combo.obsm['{umap_key}'] missing."
    assert combo.obsm[umap_key].shape[1] == 3, f"combo.obsm['{umap_key}'] must be 3D."

    coords = combo.obsm[umap_key]
    df = combo.obs.copy()

    # --- ensure an ID column exists (use obs_names if not) ---
    if id_key not in df.columns:
        df[id_key] = combo.obs_names.astype(str)

    # coords
    df = df.assign(u1=coords[:, 0], u2=coords[:, 1], u3=coords[:, 2])

    # --- validations ---
    if color_by is not None and color_by not in df.columns:
        raise KeyError(f"color_by='{color_by}' not found in combo.obs.")
    if symbol_by is not None and symbol_by not in df.columns:
        raise KeyError(f"symbol_by='{symbol_by}' not found in combo.obs.")
    if draw_pair_lines and modality_key not in df.columns:
        raise KeyError(f"Need '{modality_key}' in combo.obs to draw pair lines.")

    hover_data = {c: True for c in hover_cols if c in df.columns}
    if color_by is not None:
        hover_data[color_by] = True

    fig = px.scatter_3d(
        df,
        x="u1", y="u2", z="u3",
        color=color_by if color_by is not None else None,
        symbol=symbol_by if symbol_by is not None else None,
        hover_data=hover_data,
        opacity=opacity,
    )
    fig.update_traces(marker=dict(size=point_size))

    # --- paired lines (only works if you have stacked modalities: same cell_id appears multiple times) ---
    if draw_pair_lines:
        vc = df[id_key].value_counts()
        has_pairs = (vc >= 2).any()
        if not has_pairs:
            # Nothing to connect; skip gracefully
            draw_pair_lines = False
        else:
            d2 = df[[id_key, modality_key, "u1", "u2", "u3"]].copy()

            # infer modalities to link if not provided
            mods = list(pd.unique(d2[modality_key].astype(str)))
            if mod_a is None or mod_b is None:
                if len(mods) < 2:
                    raise ValueError(f"Not enough modalities in '{modality_key}' to link: {mods}")
                mod_a, mod_b = mods[0], mods[1]

            wide = d2.pivot(index=id_key, columns=modality_key, values=["u1", "u2", "u3"])
            needed = [(("u1", mod_a), ("u1", mod_b)),
                      (("u2", mod_a), ("u2", mod_b)),
                      (("u3", mod_a), ("u3", mod_b))]
            if not all(a in wide.columns and b in wide.columns for a, b in needed):
                raise KeyError(
                    f"Could not find both modalities '{mod_a}' and '{mod_b}' for paired links. "
                    f"Available modalities: {mods}"
                )

            wide = wide.dropna(subset=[(c, m) for c in ("u1", "u2", "u3") for m in (mod_a, mod_b)], how="any")

            if line_sample is not None and wide.shape[0] > line_sample:
                wide = wide.sample(n=line_sample, random_state=random_state)

            x = np.empty(wide.shape[0] * 3)
            y = np.empty(wide.shape[0] * 3)
            z = np.empty(wide.shape[0] * 3)

            x[0::3] = wide[("u1", mod_a)].to_numpy()
            y[0::3] = wide[("u2", mod_a)].to_numpy()
            z[0::3] = wide[("u3", mod_a)].to_numpy()

            x[1::3] = wide[("u1", mod_b)].to_numpy()
            y[1::3] = wide[("u2", mod_b)].to_numpy()
            z[1::3] = wide[("u3", mod_b)].to_numpy()

            x[2::3] = np.nan
            y[2::3] = np.nan
            z[2::3] = np.nan

            fig.add_trace(
                go.Scatter3d(
                    x=x, y=y, z=z,
                    mode="lines",
                    line=dict(width=line_width, color=line_color),
                    opacity=line_opacity,
                    hoverinfo="skip",
                    showlegend=False,
                    name=f"paired links ({mod_a}↔{mod_b})",
                )
            )

    fig.update_layout(
        width=width,
        height=height,
        margin=dict(l=10, r=10, t=40, b=10),
        scene=dict(xaxis_title="UMAP1", yaxis_title="UMAP2", zaxis_title="UMAP3"),
        title=f"Interactive 3D UMAP (key={umap_key}, color={color_by}, symbol={symbol_by})",
    )

    return fig


# %% [code]
# Scanpy writes to combo.obsm["X_umap"] (now 3D). If you want to preserve 2D:
# Compute 2D first (done), copy it
combo.obsm["X_umap_2d"] = combo.obsm["X_umap"].copy()


# %% [code]
# Then compute 3D UMAP - might take a bit since we're using, like, 150k cells for the test set for the figures
sc.tl.umap(combo, n_components=3, random_state=42)


# %% [code]
# Then add 2d back to .obsm['X_umap'] and 3d to .obsm['X_umap_3d']
combo.obsm["X_umap_3d"] = combo.obsm["X_umap"].copy()
combo.obsm["X_umap"] = combo.obsm["X_umap_2d"]  # restore default 2D


# %% [code]
import plotly.io as pio
#pio.renderers.default

# Classic Jupyter Notebook:
pio.renderers.default = "notebook_connected"

# JupyterLab:
#pio.renderers.default = "jupyterlab"

# VS Code notebooks:
#pio.renderers.default = "vscode"


# %% [code]
# After you’ve created your 3D UMAP via function and added it to combo:
threedee_umap_fig = plot_umap3d_interactive(
    combo,
    color_by=label_key,
    symbol_by="modality",
    draw_pair_lines=True,
)


# %% [code]
threedee_umap_fig.show()


# %% [code]


# %% [code]


# %% [code]


# %% [code]


# %% [code]


# %% [code]


# %% [code]


# %% [code]


# %% [code]


# %% [code]


# %% [code]


# %% [code]


# %% [code]


# %% [code]


# %% [code]


# %% [code]
```
