## Supplemental Code for "Unifying multimodal single-cell data with a mixture-of-experts *β*-variational autoencoder framework": Fig03_CITE_biological_latent.ipynb

````
# %% [markdown]
# UniVI manuscript - Figure 3 generation reproducible workflow
### CITE-seq cross-modal prediction and data generation, validated with marker expression

Andrew Ashford, Pathways + Omics Group, Oregon Health & Science University, Portland, OR - 12/6/2025

This Jupyter Notebook will house the end-to-end workflow to generate the panels in Figure 3 of our manuscript, "Unifying multimodal single-cell data with a mixture-of-experts β-variational autoencoder framework" which is currently being revised for Genome Research and is available currently on bioRxiv at the following link: https://www.biorxiv.org/content/10.1101/2025.02.28.640429v1.full

GitHub for the project can be found at: https://github.com/Ashford-A/UniVI

Package is pip installable via the command: 
```bash
pip install univi
```


# %% [markdown]
### Import modules

# %% [code]
# Import non-UniVI modules
import os
import sys
from pathlib import Path

import numpy as np
import pandas as pd
import scanpy as sc
import anndata as ad
import torch

from torch.utils.data import DataLoader, Subset


# %% [code]
# Import required UniVI modules
from univi import (
    ModalityConfig,
    UniVIConfig,
    TrainingConfig,
    UniVIMultiModalVAE,
    matching,
    UniVITrainer,
    write_univi_latent,
    MultiModalDataset,
)

import univi as uv
import univi.evaluation as ue
import univi.plotting as up


# %% [code]
# Double check UniVI module version
print("Installed version is univi v" + str(uv.__version__))


# %% [markdown]
### Specify device to use for model

Set "device" - preferably device should be "cuda" for speedier model implementation/training. Requires GPU and the correct packages/versions.


# %% [code]
print("torch:", torch.__version__)
print("torch.cuda.is_available():", torch.cuda.is_available())
device = "cuda" if torch.cuda.is_available() else "cpu"
print("Using device:", device)


# %% [markdown]
### Specify file paths

Where data lives.

# %% [code]
DATA_ROOT = Path("/home/groups/precepts/ashforda/UniVI_v2/UniVI_older-non_git/data/Hao_CITE-seq_data")

RNA_PATH = DATA_ROOT / "Hao_RNA_data.h5ad"
ADT_PATH = DATA_ROOT / "Hao_ADT_data.h5ad"

print("RNA file:", RNA_PATH)
print("ADT file:", ADT_PATH)


# %% [markdown]
### Read in data

Read data into AnnData objects using the paths in the code chunk above.

# %% [code]
rna = sc.read_h5ad(RNA_PATH)

print(rna)
print("RNA obs names head:", rna.obs_names[:5].tolist())


# %% [code]
adt = sc.read_h5ad(ADT_PATH)

print(adt)
print("ADT obs names head:", adt.obs_names[:5].tolist())


# %% [markdown]
### Align cells between RNA and ADT

Make sure the cell indices are aligned so UniVI knows which samples are paired.

# %% [code]
# Intersect barcodes
common_cells = rna.obs_names.intersection(adt.obs_names)
print("Common cells:", len(common_cells))

rna = rna[common_cells].copy()
adt = adt[common_cells].copy()

# Make sure order matches
adt = adt[rna.obs_names].copy()

assert np.array_equal(rna.obs_names.values, adt.obs_names.values)
print("obs_names aligned between RNA and ADT.")


# %% [markdown]
### Stratify the data by celltype so that we train a balanced/generalizeable model

Also specifying all the data preprocessing functions below, will be used in their own respective sections.

# %% [code]
# -----------------------------
# Stratified split with cap
# -----------------------------
def stratified_split(
    idx,
    labels,
    frac_train=0.8,
    frac_val=0.1,
    seed=0,
    max_per_label=None,
    unused_to_test=True,
):
    """
    For each label:
      - shuffle its indices
      - keep up to max_per_label as "used"
      - split used -> train/val/test by fractions
      - leftover beyond max_per_label -> unused
    If unused_to_test=True, unused is appended into test and not returned separately.
    """
    rng = np.random.default_rng(seed)
    idx = np.asarray(idx)
    labels = np.asarray(labels)

    train_idx, val_idx, test_idx, unused_idx = [], [], [], []

    # stable label order (preserves category order if categorical, else sorted unique)
    uniq = pd.unique(labels)

    for lab in uniq:
        m = idx[labels == lab]
        rng.shuffle(m)

        if max_per_label is not None:
            used = m[:max_per_label]
            leftover = m[max_per_label:]
            if leftover.size:
                unused_idx.append(leftover)
        else:
            used = m

        n = used.size
        if n == 0:
            continue

        n_train = int(frac_train * n)
        n_val   = int(frac_val * n)
        # remainder -> test
        train_idx.append(used[:n_train])
        val_idx.append(used[n_train:n_train + n_val])
        test_idx.append(used[n_train + n_val:])

    train_idx = np.concatenate(train_idx) if train_idx else np.array([], dtype=int)
    val_idx   = np.concatenate(val_idx)   if val_idx   else np.array([], dtype=int)
    test_idx  = np.concatenate(test_idx)  if test_idx  else np.array([], dtype=int)
    unused_idx = np.concatenate(unused_idx) if unused_idx else np.array([], dtype=int)

    if unused_to_test and unused_idx.size:
        test_idx = np.concatenate([test_idx, unused_idx])
        unused_idx = np.array([], dtype=int)

    return train_idx, val_idx, test_idx, unused_idx


# %% [code]
# -----------------------------
# Small helper functions
# -----------------------------
def _ensure_counts_layer(adata, layer_name="counts"):
    if layer_name in adata.layers:
        return
    # Fall back to X as counts if needed (only if X is actually raw counts!)
    adata.layers[layer_name] = adata.X.copy()

def _subset_by_idx_pair(rna, adt, idx):
    # Assumes rna/adt are already paired in the same order
    return rna[idx].copy(), adt[idx].copy()

def _clr_normalize_dense(X, eps=1e-8):
    """
    CLR per cell: log1p(x) - mean(log1p(x)) across features.
    Works for dense or sparse; returns dense float32 array.
    """
    if sp.issparse(X):
        X = X.toarray()
    X = X.astype(np.float32, copy=False)
    X = np.log1p(X)
    X = X - X.mean(axis=1, keepdims=True)
    return X

def preprocess_citeseq_splits(
    rna_train, adt_train,
    rna_val, adt_val,
    rna_test, adt_test,
    *,
    rna_counts_layer="counts",
    adt_counts_layer="counts",
    n_hvg=2000,
    target_sum=1e4,
    rna_make_log1p=True,
    adt_make_clr=True,
):
    """
    Train-fit, then transform val/test the same way.

    RNA:
      - ensures counts layer
      - HVGs learned on train only (Seurat v3 if possible)
      - normalize_total + log1p (applied to all splits)
      - subsets to HVGs for all splits

    ADT:
      - ensures counts layer
      - CLR per cell (applied to all splits) -> stored in .X (dense float32)
    """
    # ensure counts layers exist
    for a in (rna_train, rna_val, rna_test):
        _ensure_counts_layer(a, rna_counts_layer)
    for a in (adt_train, adt_val, adt_test):
        _ensure_counts_layer(a, adt_counts_layer)

    # ---- RNA: pick HVGs on TRAIN only
    rna_train_tmp = rna_train.copy()
    rna_train_tmp.X = rna_train_tmp.layers[rna_counts_layer]

    try:
        sc.pp.highly_variable_genes(
            rna_train_tmp,
            n_top_genes=int(n_hvg),
            flavor="seurat_v3",
            layer=None,   # we set .X already
        )
    except Exception:
        sc.pp.highly_variable_genes(
            rna_train_tmp,
            n_top_genes=int(n_hvg),
            flavor="seurat",
            layer=None,
        )

    hvg_mask = rna_train_tmp.var["highly_variable"].to_numpy()
    hvg_genes = rna_train_tmp.var_names[hvg_mask].tolist()

    # ---- RNA: normalize/log1p + subset to HVGs (train/val/test)
    def _rna_transform(adata):
        # subset genes first to minimize memory
        ad = adata[:, hvg_genes].copy()
        X = ad.layers[rna_counts_layer]

        # normalize_total on counts
        if sp.issparse(X):
            cell_sums = np.asarray(X.sum(axis=1)).ravel()
            scale = (float(target_sum) / np.maximum(cell_sums, 1e-12)).astype(np.float32)
            Xn = X.multiply(scale[:, None])
        else:
            cell_sums = X.sum(axis=1, keepdims=True)
            Xn = (X / np.maximum(cell_sums, 1e-12)) * float(target_sum)

        if rna_make_log1p:
            if sp.issparse(Xn):
                Xn = Xn.tocsr(copy=True)
                Xn.data = np.log1p(Xn.data).astype(np.float32, copy=False)
            else:
                Xn = np.log1p(Xn).astype(np.float32, copy=False)

        ad.X = Xn
        # keep original counts too (still subsetted)
        ad.layers[rna_counts_layer] = ad.layers[rna_counts_layer]
        return ad

    rna_train_pp = _rna_transform(rna_train)
    rna_val_pp   = _rna_transform(rna_val)
    rna_test_pp  = _rna_transform(rna_test)

    # ---- ADT: CLR into .X (train/val/test)
    def _adt_transform(adata):
        ad = adata.copy()
        Xc = ad.layers[adt_counts_layer]
        if adt_make_clr:
            ad.X = _clr_normalize_dense(Xc)
        else:
            # keep counts as X (dense float32)
            ad.X = Xc.toarray().astype(np.float32) if sp.issparse(Xc) else Xc.astype(np.float32)
        return ad

    adt_train_pp = _adt_transform(adt_train)
    adt_val_pp   = _adt_transform(adt_val)
    adt_test_pp  = _adt_transform(adt_test)

    return (rna_train_pp, adt_train_pp,
            rna_val_pp,   adt_val_pp,
            rna_test_pp,  adt_test_pp,
            hvg_genes)


# %% [code]
# Check the counts of each celltype.l1 in the data
print(rna.obs["celltype.l1"].value_counts())


# %% [code]
# Check the counts of each celltype.l2 in the data
print(rna.obs["celltype.l2"].value_counts())


# %% [code]
# Check the counts of each celltype.l3 in the data
print(rna.obs["celltype.l3"].value_counts())


# %% [code]
# -----------------------------
# Use it on Hao CITE-seq
# -----------------------------
labels = rna.obs["celltype.l3"].astype(str).to_numpy()
idx = np.arange(rna.n_obs)

train_idx, val_idx, test_idx, unused_idx = stratified_split(
    idx, labels,
    frac_train=0.8, frac_val=0.1, seed=42,
    max_per_label=1200,
    unused_to_test=True,      # set True if you want unused merged into test
)

# paired splits
rna_train, adt_train = _subset_by_idx_pair(rna, adt, train_idx)
rna_val,   adt_val   = _subset_by_idx_pair(rna, adt, val_idx)
rna_test,  adt_test  = _subset_by_idx_pair(rna, adt, test_idx)

if unused_idx.size:
    rna_unused, adt_unused = _subset_by_idx_pair(rna, adt, unused_idx)
else:
    rna_unused, adt_unused = None, None


# %% [markdown]
### Data preprocessing

* RNA preprocessing (log1p + HVG + scale → Gaussian decoder)

* ADT preprocessing (CLR + scale → Gaussian decoder)

Running the preprocessing functions specified above. Of note, we're performing the preprocessing per-train/val/test split.

# %% [code]
import scipy.sparse as sp

# preprocessing (fit on train, apply to val/test)
(rna_train_pp, adt_train_pp,
 rna_val_pp,   adt_val_pp,
 rna_test_pp,  adt_test_pp,
 hvg_genes) = preprocess_citeseq_splits(
    rna_train, adt_train,
    rna_val,   adt_val,
    rna_test,  adt_test,
    n_hvg=2000,
    target_sum=1e4,
    rna_make_log1p=True,
    adt_make_clr=True,
)


# %% [code]
'''
# optionally preprocess "unused" with same HVGs + same transforms (no refit)
if rna_unused is not None:
    # reuse the same transform logic by treating unused as "test"
    (rna_unused_pp, adt_unused_pp,
     _, _, _, _, _) = preprocess_citeseq_splits(
        rna_train, adt_train,     # only used to learn HVGs (already learned here, but fine)
        rna_unused, adt_unused,
        rna_unused, adt_unused,
        n_hvg=len(hvg_genes),
        target_sum=1e4,
        rna_make_log1p=True,
        adt_make_clr=True,
    )
    # the function returns in (train,val,test) slots; we passed unused into val/test
    # so grab one of them:
    # rna_unused_pp == rna_train_pp from that call; use the "val" one instead:
    # Instead, do a tiny manual transform with the already-computed hvg_genes:
    # (If you want this cleaned up, I can give you a one-liner helper.)
'''

# %% [code]
# Sanity check above preprocessed data objects - start with RNA
print(rna_train_pp)
print(rna_val_pp)
print(rna_test_pp)


# %% [code]
print(rna_train_pp.obs['celltype.l3'].value_counts())
print(rna_val_pp.obs['celltype.l3'].value_counts())
print(rna_test_pp.obs['celltype.l3'].value_counts())


# %% [code]
# Now sanity check ADT data objects
print(adt_train_pp)
print(adt_val_pp)
print(adt_test_pp)


# %% [markdown]
### Wrap into MultiModalDataset & DataLoaders

# %% [code]
from univi.data import align_paired_obs_names

pin_memory = (device == "cuda")

# Make sure each split is paired and ordered the same across modalities
# This was erroring out due to a function bug, fixed it and should be fixed by manuscript publication
#train_dict = align_paired_obs_names({"rna": rna_train_pp, "adt": adt_train_pp})
#val_dict   = align_paired_obs_names({"rna": rna_val_pp,   "adt": adt_val_pp})
#test_dict  = align_paired_obs_names({"rna": rna_test_pp,  "adt": adt_test_pp})

# Using this instead for now since we know the data are already paired from above code
assert (rna_train_pp.obs_names == adt_train_pp.obs_names).all()
assert (rna_val_pp.obs_names   == adt_val_pp.obs_names).all()
assert (rna_test_pp.obs_names  == adt_test_pp.obs_names).all()

train_dict = {"rna": rna_train_pp, "adt": adt_train_pp}
val_dict   = {"rna": rna_val_pp,   "adt": adt_val_pp}
test_dict  = {"rna": rna_test_pp,  "adt": adt_test_pp}

# Build datasets (CPU tensors; trainer/model moves to GPU)
train_ds = MultiModalDataset(adata_dict=train_dict, X_key="X", device=None)
val_ds   = MultiModalDataset(adata_dict=val_dict,   X_key="X", device=None)
test_ds  = MultiModalDataset(adata_dict=test_dict,  X_key="X", device=None)

batch_size = 256
num_workers = 0

train_loader = DataLoader(
    train_ds,
    batch_size=batch_size,
    shuffle=True,
    num_workers=num_workers,
    pin_memory=pin_memory,
    drop_last=False,
)

val_loader = DataLoader(
    val_ds,
    batch_size=batch_size,
    shuffle=False,
    num_workers=num_workers,
    pin_memory=pin_memory,
    drop_last=False,
)

test_loader = DataLoader(
    test_ds,
    batch_size=batch_size,
    shuffle=False,
    num_workers=num_workers,
    pin_memory=pin_memory,
    drop_last=False,
)

print("n_train / n_val / n_test:", train_ds.n_cells, val_ds.n_cells, test_ds.n_cells)
print("batches:", len(train_loader), len(val_loader), len(test_loader))

# sanity check one batch
x = next(iter(train_loader))
print({k: v.shape for k, v in x.items()})


# %% [markdown]
### UniVI configs

# %% [code]
univi_cfg = UniVIConfig(
    latent_dim=30,
    beta=1.5,
    gamma=5.0,
    encoder_dropout=0.1,
    decoder_dropout=0.0,
    encoder_batchnorm=True,
    decoder_batchnorm=False,
    kl_anneal_start=0,
    kl_anneal_end=0,
    align_anneal_start=0,
    align_anneal_end=0,
    modalities=[
        ModalityConfig(
            name="rna",
            input_dim=rna_train_pp.n_vars,
            encoder_hidden=[512, 256, 128],
            decoder_hidden=[128, 256, 512],
            likelihood="gaussian",
        ),
        ModalityConfig(
            name="adt",
            input_dim=adt_train_pp.n_vars,
            encoder_hidden=[128, 64],
            decoder_hidden=[64, 128],
            likelihood="gaussian",
        ),
    ],
)


# %% [markdown]
If you want raw-count NB/ZINB instead, you’d:
* Put raw counts in .layers["counts"]
* Set X_key="counts" in the dataset
* Use likelihood="nb" or "zinb" above

But for this "Figure 2" notebook, the scaled Gaussian setup is nicely stable and good for the best-integrated latent space.


# %% [markdown]
### Instantiate model and model objective

# %% [code]
model = UniVIMultiModalVAE(
    univi_cfg,
    loss_mode="v1",      # cross-recon + cross-posterior alignment
    #v1_recon="cross",   # full k→j cross-recon
    v1_recon="avg",
    #v1_recon_mix=0.5,
    normalize_v1_terms=True,
).to(device)


# %% [markdown]
### Instantiate TrainingConfig & trainer

# %% [code]
train_cfg = TrainingConfig(
    n_epochs=3000,
    batch_size=batch_size,
    lr=1e-3,
    weight_decay=1e-4,
    device=device,
    log_every=20,
    grad_clip=5.0,
    num_workers=0,
    seed=42,
    early_stopping=True,
    patience=100,
    min_delta=0.0,
)

print(train_cfg)
print(univi_cfg)


# %% [code]
trainer = UniVITrainer(
    model=model,
    train_loader=train_loader,
    val_loader=val_loader,
    train_cfg=train_cfg,
    device=device,
)

print(trainer)


# %% [markdown]
### Fit the model

# %% [code]
history = trainer.fit()

# history is a dict with keys like "train_loss", "val_loss", "beta", "gamma"
print("Training finished.")
print("Best val loss:", np.min(history["val_loss"]))


# %% [code]
import matplotlib.pyplot as plt

plt.figure()
plt.plot(history["train_loss"], label="train")
plt.plot(history["val_loss"], label="val")
plt.xlabel("Epoch")
plt.ylabel("Loss")
plt.legend()
plt.title("UniVI training (CITE-seq)")
plt.show()


# %% [markdown]
### Quick UMAP + visualization

You can now treat X_univi like any other embedding.

# %% [code]
# Check out many of each cell type are in the test set for level 2 cell annotations
print(rna_test_pp.obs["celltype.l2"].value_counts())


# %% [code]
# Scanpy defaults (affects sc.pl.*)
sc.set_figure_params(
    figsize=(10, 8),   # bigger canvas
    dpi=100,            # on-screen sharpness
    dpi_save=300,       # saved file sharpness
    fontsize=10,
    frameon=False,
)

# Matplotlib defaults (affects plt.*)
plt.rcParams.update({
    "figure.figsize": (10, 8),
    "figure.dpi": 100,
    "savefig.dpi": 300,
    "savefig.bbox": "tight",
    "savefig.pad_inches": 0.1,
    "axes.titlesize": 12,
    "axes.labelsize": 10,
    "legend.fontsize": 10,
})


# %% [code]
# Make sure adata_dict has the *same ordering* as during training
adata_dict = {
    "rna": rna_test_pp,
    "adt": adt_test_pp,
}

Z = write_univi_latent(
    model,
    adata_dict,
    obsm_key="X_univi",   # will be added to each AnnData in adata_dict
    batch_size=512,
    device=device,
    use_mean=True,       # deterministic: use encoder means instead of noisy samples
    #use_mean=False,       # Stochastic
)

print("UniVI latent shape:",  Z.shape)
print("rna.obsm keys:",       rna_test_pp.obsm.keys())
print("adt.obsm keys:",       adt_test_pp.obsm.keys())


# %% [code]
# Make sure X_univi exists in both
assert "X_univi" in rna_test_pp.obsm and "X_univi" in adt_test_pp.obsm
assert rna_test_pp.obsm["X_univi"].shape == adt_test_pp.obsm["X_univi"].shape  # paired case

rna_u = rna_test_pp.copy()
adt_u = adt_test_pp.copy()
rna_u.obs["modality"] = "rna"
adt_u.obs["modality"] = "adt"

# concatenate (keeps obs, stacks cells)
combo = ad.concat([rna_u, adt_u], join="outer", label="modality", keys=["rna","adt"], index_unique="-")

# IMPORTANT: carry over the latent explicitly (concat won't always merge obsm how you want)
combo.obsm["X_univi"] = np.vstack([rna_u.obsm["X_univi"], adt_u.obsm["X_univi"]])


# %% [code]
# neighbors on the latent
sc.pp.neighbors(combo, use_rep="X_univi", n_neighbors=15)


# %% [code]
# umap on the latent using the neighbors
sc.tl.umap(combo)


# %% [code]
sc.pl.umap(
    combo,
    color=["modality"],
    frameon=False,
    size=3,
    wspace=0.4,
    alpha=0.5
)


# %% [code]
sc.pl.umap(
    combo,
    color=["celltype.l1"],
    frameon=False,
    size=3,
    wspace=0.4,
)


# %% [code]
sc.pl.umap(
    combo,
    color=["celltype.l2"],
    frameon=False,
    size=3,
    wspace=0.4,
)


# %% [code]
sc.pl.umap(
    combo,
    color=["celltype.l3"],
    frameon=False,
    size=3,
    wspace=0.4,
)


# %% [markdown]
### Evaluation metrics for the model prior to Figure 3 analyses (Figure 2 metrics)

# %% [code]
import json

from sklearn.neighbors import NearestNeighbors
from sklearn.metrics import confusion_matrix, accuracy_score, f1_score

# ----------------------------
# User settings
# ----------------------------
outdir = "figures/Figure3_CITEseq_cross_recon_reproducibility"
os.makedirs(outdir, exist_ok=True)

celltype_key = "celltype.l2"   # change to celltype.l1 / celltype.l3 as needed
k_mix = 10                     # k for modality mixing
k_lt  = 5                      # k for label transfer kNN
seed = 42


# %% [code]
from sklearn.neighbors import NearestNeighbors
from sklearn.metrics import confusion_matrix, accuracy_score, f1_score

# ----------------------------
# Metrics helpers
# ----------------------------
def foscttm_chunked(Z1, Z2, block=512):
    """
    Exact FOSCTTM computed in blocks to avoid NxN memory blowups.
    Assumes 1:1 pairing between rows i in Z1 and Z2.
    """
    Z1 = np.asarray(Z1, dtype=np.float32)
    Z2 = np.asarray(Z2, dtype=np.float32)
    assert Z1.shape == Z2.shape
    n = Z1.shape[0]

    Z2_T = Z2.T
    n2 = np.sum(Z2 * Z2, axis=1)  # (n,)

    fos = np.empty(n, dtype=np.float32)

    for i0 in range(0, n, block):
        i1 = min(i0 + block, n)
        A = Z1[i0:i1]  # (b, d)
        n1 = np.sum(A * A, axis=1)[:, None]  # (b,1)

        d2 = n1 + n2[None, :] - 2.0 * (A @ Z2_T)  # (b,n)

        true = d2[np.arange(i1 - i0), np.arange(i0, i1)]
        fos[i0:i1] = (d2 < true[:, None]).sum(axis=1) / (n - 1)

    return float(fos.mean()), float(fos.std(ddof=1) / np.sqrt(n))


def modality_mixing_score(Z, modality_labels, k=20, metric="euclidean"):
    """
    Vanilla modality mixing:
    Mean fraction of kNN neighbors that differ in modality.
    Use this for *non-duplicated* sets (e.g., concatenated modality-specific embeddings).
    """
    Z = np.asarray(Z, dtype=np.float32)
    modality_labels = np.asarray(modality_labels)

    n = Z.shape[0]
    if n <= 1:
        return 0.0

    k_eff = int(min(max(int(k), 1), n - 1))
    nn = NearestNeighbors(n_neighbors=k_eff + 1, metric=metric)
    nn.fit(Z)
    nbrs = nn.kneighbors(Z, return_distance=False)[:, 1:]  # drop self

    same = (modality_labels[nbrs] == modality_labels[:, None])
    return float((~same).mean())


def modality_mixing_score_excluding_pairs(Z, modality_labels, cell_ids, k=20, metric="euclidean"):
    """
    Pair-aware modality mixing for 'combo' style stacked data (same cell appears twice: RNA + ADT),
    where the fused embedding may be identical (or extremely close) for the paired duplicates.

    Computes: for each row, the fraction of neighbors from the other modality,
    AFTER removing the paired duplicate (same cell_id) from its neighbor list.

    cell_ids must map each row to a shared cell identifier (same for RNA+ADT copy).
    """
    Z = np.asarray(Z, dtype=np.float32)
    modality_labels = np.asarray(modality_labels)
    cell_ids = np.asarray(cell_ids).astype(str)

    n = Z.shape[0]
    if n <= 2:
        return 0.0

    # Need enough neighbors so we can drop self + paired duplicate and still have k
    k_eff = int(min(max(int(k), 1), n - 2))
    nn = NearestNeighbors(n_neighbors=k_eff + 2, metric=metric)
    nn.fit(Z)
    nbrs = nn.kneighbors(Z, return_distance=False)[:, 1:]  # drop self

    # Build "pair index": for each row i, pair[i] is the index of the other modality copy
    first = {}
    pair = np.full(n, -1, dtype=np.int64)
    for i, cid in enumerate(cell_ids):
        if cid in first:
            j = first[cid]
            pair[i] = j
            pair[j] = i
        else:
            first[cid] = i

    frac_other = np.empty(n, dtype=np.float32)
    for i in range(n):
        neigh = nbrs[i]

        # remove paired duplicate if present
        pj = pair[i]
        if pj != -1:
            neigh = neigh[neigh != pj]

        neigh = neigh[:k_eff]
        frac_other[i] = (modality_labels[neigh] != modality_labels[i]).mean()

    return float(frac_other.mean())


def knn_label_transfer(Z_source, y_source, Z_target, y_target, k=15, metric="euclidean"):
    """
    kNN label transfer: predict y_target from neighbors in Z_source.
    Returns predictions, accuracy, macro-F1, confusion matrix.
    """
    Z_source = np.asarray(Z_source, dtype=np.float32)
    Z_target = np.asarray(Z_target, dtype=np.float32)
    y_source = np.asarray(y_source, dtype=str)
    y_target = np.asarray(y_target, dtype=str)

    nn = NearestNeighbors(n_neighbors=k, metric=metric)
    nn.fit(Z_source)
    nbrs = nn.kneighbors(Z_target, return_distance=False)

    preds = []
    for inds in nbrs:
        votes = y_source[inds]
        vals, cnts = np.unique(votes, return_counts=True)
        preds.append(vals[np.argmax(cnts)])
    preds = np.asarray(preds, dtype=str)

    acc = float(accuracy_score(y_target, preds))
    macro_f1 = float(f1_score(y_target, preds, average="macro"))
    classes = np.unique(np.concatenate([y_source, y_target]))
    cm = confusion_matrix(y_target, preds, labels=classes)
    return preds, acc, macro_f1, cm, classes


# %% [code]
from univi.evaluation import encode_adata

Z_rna = encode_adata(model, rna_test_pp, modality="rna",
                     latent="modality_mean", device=device, batch_size=1024)
Z_adt = encode_adata(model, adt_test_pp, modality="adt",
                     latent="modality_mean", device=device, batch_size=1024)

key_rna = "encode_adata(modality_mean)"
key_adt = "encode_adata(modality_mean)"

rna_test_pp.obsm["X_univi_rna"] = Z_rna
adt_test_pp.obsm["X_univi_adt"] = Z_adt


# %% [code]
print("max|Z_rna-Z_adt|:", float(np.max(np.abs(Z_rna - Z_adt))))
print("mean L2(Z_rna-Z_adt):", float(np.mean(np.linalg.norm(Z_rna - Z_adt, axis=1))))


# %% [code]
# ---------- PRE-FLIGHT CHECK + FIX (run once right before metrics/plots) ----------
def _preflight_align_for_metrics(
    rna_test_pp, adt_test_pp, Z_rna, Z_adt, celltype_key="celltype.l2"
):
    import numpy as np

    # 1) basic shape checks
    print("rna_test_pp n_obs:", rna_test_pp.n_obs)
    print("adt_test_pp n_obs:", adt_test_pp.n_obs)
    print("Z_rna shape:", np.asarray(Z_rna).shape)
    print("Z_adt shape:", np.asarray(Z_adt).shape)

    # 2) enforce pairing by obs_names intersection (preserves RNA order)
    common = rna_test_pp.obs_names.intersection(adt_test_pp.obs_names)
    if len(common) != rna_test_pp.n_obs or len(common) != adt_test_pp.n_obs:
        print(f"[preflight] restricting to common paired cells: {len(common)}")

    rna_al = rna_test_pp[common].copy()
    adt_al = adt_test_pp[common].copy()
    adt_al = adt_al[rna_al.obs_names].copy()  # force identical order

    # 3) slice Z arrays to match aligned AnnData order (assumes Z were computed in the same order)
    # If your Z were computed from rna_test_pp/adt_test_pp directly, this is safe.
    # If not, you *must* recompute Z after alignment.
    if np.asarray(Z_rna).shape[0] != rna_al.n_obs or np.asarray(Z_adt).shape[0] != adt_al.n_obs:
        raise ValueError(
            "[preflight] Z arrays do not match aligned AnnData n_obs. "
            "Recompute Z_rna/Z_adt from rna_al/adt_al."
        )

    # 4) labels must come from the SAME objects you predict on
    labels_rna = rna_al.obs[celltype_key].astype(str).to_numpy()
    labels_adt = adt_al.obs[celltype_key].astype(str).to_numpy()

    # 5) final asserts (this prevents your exact error)
    assert rna_al.n_obs == adt_al.n_obs
    assert (rna_al.obs_names == adt_al.obs_names).all()
    assert len(labels_rna) == np.asarray(Z_rna).shape[0]
    assert len(labels_adt) == np.asarray(Z_adt).shape[0]

    print("[preflight] OK: paired, aligned, and lengths match.")
    return rna_al, adt_al, labels_rna, labels_adt


# run it:
rna_test_pp, adt_test_pp, labels_rna, labels_adt = _preflight_align_for_metrics(
    rna_test_pp, adt_test_pp, Z_rna, Z_adt, celltype_key=celltype_key
)


# %% [code]
# ----------------------------
# Compute metrics (Figure 3)
# ----------------------------

# 0) Fused latent (if you truly want it)
Z_fused = np.asarray(combo.obsm["X_univi"])
mods_fused = combo.obs["modality"].to_numpy()

# derive cell_ids that are identical for RNA+ADT copies
# works with index_unique="-" that produces "<orig>-rna" / "<orig>-adt"
cell_ids = combo.obs_names.to_series().str.rsplit("-", n=1).str[0].to_numpy()

mix_fused = modality_mixing_score_excluding_pairs(Z_fused, mods_fused, cell_ids, k=k_mix)

# 1) Modality-specific latents (Z_rna, Z_adt) already computed above

# 2) FOSCTTM on modality-specific latents (subsample ok)
n_total = int(rna_test_pp.n_obs)
n_fos = int(min(20000, n_total))
rng = np.random.default_rng(seed)
sub = rng.choice(n_total, size=n_fos, replace=False)

fos_rna_adt, fos_sem = foscttm_chunked(Z_rna[sub], Z_adt[sub], block=512)

# 3) Modality mixing on modality-specific latents
Z_concat = np.vstack([Z_rna, Z_adt])
mods = np.array(["rna"] * Z_rna.shape[0] + ["adt"] * Z_adt.shape[0], dtype=object)
mix_modality_specific = modality_mixing_score(Z_concat, mods, k=k_mix)

# 4) Label transfer (modality-specific)
# Make labels that match each embedding 1:1
labels_rna = rna_test_pp.obs[celltype_key].astype(str).to_numpy()
labels_adt = adt_test_pp.obs[celltype_key].astype(str).to_numpy()

assert Z_rna.shape[0] == labels_rna.shape[0], (Z_rna.shape, labels_rna.shape)
assert Z_adt.shape[0] == labels_adt.shape[0], (Z_adt.shape, labels_adt.shape)

pred_adt, acc_r2a, f1_r2a, cm_r2a, classes = knn_label_transfer(
    Z_source=Z_rna, y_source=labels_rna,
    Z_target=Z_adt, y_target=labels_adt,
    k=k_lt
)
pred_rna, acc_a2r, f1_a2r, cm_a2r, _ = knn_label_transfer(
    Z_source=Z_adt, y_source=labels_adt,
    Z_target=Z_rna, y_target=labels_rna,
    k=k_lt
)

metrics = {
    "embedding_keys": {"rna": key_rna, "adt": key_adt, "fused": "combo.obsm['X_univi']"},
    "n_test": n_total,
    "celltype_key": celltype_key,

    "FOSCTTM_metric": "euclidean_sq",
    "FOSCTTM_subsample_n": n_fos,
    "FOSCTTM_rna_vs_adt_mean": float(fos_rna_adt),
    "FOSCTTM_rna_vs_adt_sem": float(fos_sem),

    "modality_mixing_k": int(k_mix),
    "modality_mixing_fused": float(mix_fused),
    "modality_mixing_modality_specific": float(mix_modality_specific),

    "label_transfer_k": int(k_lt),
    "label_transfer_rna_to_adt_acc": float(acc_r2a),
    "label_transfer_rna_to_adt_macroF1": float(f1_r2a),
    "label_transfer_adt_to_rna_acc": float(acc_a2r),
    "label_transfer_adt_to_rna_macroF1": float(f1_a2r),
}

print(json.dumps(metrics, indent=2))
with open(os.path.join(outdir, "figure3_metrics.json"), "w") as f:
    json.dump(metrics, f, indent=2)
pd.DataFrame([metrics]).to_csv(os.path.join(outdir, "figure3_metrics.csv"), index=False)


# %% [code]
from sklearn.neighbors import NearestNeighbors
from sklearn.metrics import precision_recall_fscore_support

# If plots ever don't show in classic notebook, run once:
# %matplotlib inline

# -------------------------
# show-or-save helper
# -------------------------
def _finish(fig=None, outpath=None, dpi=300, show=True, close=True):
    if outpath is not None:
        plt.savefig(outpath, dpi=dpi, bbox_inches="tight")
    if show:
        plt.show()
    if close:
        plt.close(fig if fig is not None else plt.gcf())

# -------------------------
# plots
# -------------------------
def plot_metrics_bar(
    metrics,
    keys,
    title="Figure 2 summary metrics",
    outpath=None,
    err_keys=None,          # dict: metric_key -> error_metric_key
    default_err=np.nan,     # np.nan = no error bar drawn; 0.0 = zero-length
    capsize=4,
):
    vals = np.array([float(metrics[k]) for k in keys], dtype=float)

    if err_keys is None:
        yerr = np.full_like(vals, default_err, dtype=float)
    else:
        yerr = np.array(
            [float(metrics[err_keys[k]]) if k in err_keys and err_keys[k] in metrics else default_err
             for k in keys],
            dtype=float
        )

    fig = plt.figure(figsize=(10, 6))
    plt.bar(keys, vals, yerr=yerr, capsize=capsize)
    plt.xticks(rotation=45, ha="right")
    plt.title(title)
    plt.tight_layout()

    # inline “finish” behavior
    if outpath is not None:
        plt.savefig(outpath, dpi=300, bbox_inches="tight")
    plt.show()
    plt.close(fig)


def plot_confusion(cm, classes, title, normalize="true", outpath=None, cmap="viridis"):
    import numpy as np
    import matplotlib.pyplot as plt

    cm = np.asarray(cm, dtype=float)
    if normalize == "true":
        cm = cm / (cm.sum(axis=1, keepdims=True) + 1e-12)
        subtitle = "Row-normalized"
    elif normalize == "pred":
        cm = cm / (cm.sum(axis=0, keepdims=True) + 1e-12)
        subtitle = "Col-normalized"
    elif normalize == "all":
        cm = cm / (cm.sum() + 1e-12)
        subtitle = "Global-normalized"
    else:
        subtitle = "Counts"

    classes = np.asarray(classes, dtype=str)
    n = len(classes)

    fig, ax = plt.subplots(figsize=(14, 13))
    im = ax.imshow(cm, interpolation="nearest", cmap=cmap, aspect="auto")

    # kill any gridlines (including ones from styles)
    ax.grid(False)
    ax.minorticks_off()

    ax.set_title(f"{title}\n({subtitle})")
    ax.set_xlabel("Predicted")
    ax.set_ylabel("True")

    ax.set_xticks(np.arange(n))
    ax.set_yticks(np.arange(n))
    ax.set_xticklabels(classes, rotation=90)
    ax.set_yticklabels(classes)

    cbar = fig.colorbar(im, ax=ax)
    cbar.set_label("value")

    fig.tight_layout()

    if outpath is not None:
        fig.savefig(outpath, dpi=300, bbox_inches="tight")

    plt.show()
    plt.close(fig)


def plot_per_class_f1(y_true, y_pred, title="Per-class F1", outpath=None, top_n=None):
    y_true = np.asarray(y_true).astype(str)
    y_pred = np.asarray(y_pred).astype(str)
    classes = np.unique(np.concatenate([y_true, y_pred]))

    _, _, f1, support = precision_recall_fscore_support(
        y_true, y_pred, labels=classes, zero_division=0
    )

    df = pd.DataFrame({"class": classes, "f1": f1, "support": support})
    df = df.sort_values(["f1", "support"], ascending=[True, False])

    if top_n is not None:
        df = df.head(int(top_n))

    fig = plt.figure(figsize=(10, 0.35 * len(df) + 2.0))
    plt.barh(df["class"], df["f1"])
    plt.xlabel("F1")
    plt.title(title)
    plt.xlim(0, 1)
    plt.tight_layout()
    _finish(fig, outpath=outpath, show=True, close=True)
    return df

def plot_modality_mixing_hist(Z, mods, k=20, metric="euclidean", title="Modality mixing", outpath=None):
    Z = np.asarray(Z, dtype=np.float32)
    mods = np.asarray(mods)

    n = Z.shape[0]
    k_eff = int(min(max(int(k), 1), n - 1))

    nn = NearestNeighbors(n_neighbors=k_eff + 1, metric=metric)
    nn.fit(Z)
    nbrs = nn.kneighbors(Z, return_distance=False)[:, 1:]
    frac_other = (mods[nbrs] != mods[:, None]).mean(axis=1)

    fig = plt.figure(figsize=(7, 4.5))
    plt.hist(frac_other, bins=60)
    plt.xlabel("Fraction of kNN from other modality")
    plt.ylabel("# cells")
    plt.title(title)
    plt.tight_layout()
    _finish(fig, outpath=outpath, show=True, close=True)
    return frac_other

def plot_foscttm_sanity(Z1, Z2, idx, title="FOSCTTM sanity", outpath=None):
    Z1s = np.asarray(Z1[idx], dtype=np.float32)
    Z2s = np.asarray(Z2[idx], dtype=np.float32)

    d_true = np.sum((Z1s - Z2s) ** 2, axis=1)

    nn = NearestNeighbors(n_neighbors=51, metric="euclidean")
    nn.fit(np.asarray(Z2, dtype=np.float32))
    dist, _ = nn.kneighbors(Z1s)
    d_min = dist[:, 0] ** 2

    fig = plt.figure(figsize=(5.5, 5.5))
    plt.scatter(d_true, d_min, s=8, alpha=0.4)
    mx = np.percentile(np.concatenate([d_true, d_min]), 99)
    plt.plot([0, mx], [0, mx], linewidth=1)
    plt.xlim(0, mx); plt.ylim(0, mx)
    plt.xlabel("d(true match)^2")
    plt.ylabel("d(nearest neighbor)^2")
    plt.title(title)
    plt.tight_layout()
    _finish(fig, outpath=outpath, show=True, close=True)

def plot_paired_distance_hist(Z_rna, Z_adt, idx, title="Paired latent distance (subset)", outpath=None):
    d_pair = np.sqrt(np.sum((Z_rna[idx] - Z_adt[idx]) ** 2, axis=1))
    fig = plt.figure(figsize=(7, 4.5))
    plt.hist(d_pair, bins=80)
    plt.xlabel("||z_rna - z_adt||")
    plt.ylabel("# cells")
    plt.title(title)
    plt.tight_layout()
    _finish(fig, outpath=outpath, show=True, close=True)


# %% [code]
# 1) Summary metric bar
plot_metrics_bar(
    metrics,
    keys=[
        "FOSCTTM_rna_vs_adt_mean",
        "modality_mixing_fused",
        "modality_mixing_modality_specific",
        "label_transfer_rna_to_adt_acc",
        "label_transfer_rna_to_adt_macroF1",
    ],
    err_keys={
        #"FOSCTTM_rna_vs_adt_mean": "FOSCTTM_rna_vs_adt_sem",
        # add more here if you compute SEM/CI for them later
    },
    default_err=np.nan,   # no error bars for the others
    outpath=None,
)


# %% [code]
# 2) Confusion matrices (row-normalized)
plot_confusion(cm_r2a, classes, title=f"Label transfer (RNA → ADT), k={k_lt}", normalize="true", outpath=None)
plot_confusion(cm_a2r, classes, title=f"Label transfer (ADT → RNA), k={k_lt}", normalize="true", outpath=None)


# %% [code]
# 3) Per-class F1
_ = plot_per_class_f1(labels_adt, pred_adt, title="RNA → ADT: per-class F1", top_n=100, outpath=None)
_ = plot_per_class_f1(labels_rna, pred_rna, title="ADT → RNA: per-class F1", top_n=100, outpath=None)


# %% [code]
# 4) Modality-mixing distributions
_ = plot_modality_mixing_hist(Z_fused, mods_fused, k=k_mix, title=f"Mixing (fused latent), k={k_mix}", outpath=None)

Z_concat = np.vstack([Z_rna, Z_adt])
mods_concat = np.array(["rna"] * Z_rna.shape[0] + ["adt"] * Z_adt.shape[0], dtype=object)
_ = plot_modality_mixing_hist(Z_concat, mods_concat, k=k_mix, title=f"Mixing (modality-specific latents), k={k_mix}", outpath=None)


# %% [code]
# 5) FOSCTTM sanity + paired distance hist
sub_sanity = np.asarray(sub)[:min(5000, len(sub))]
plot_foscttm_sanity(Z_rna, Z_adt, sub_sanity, outpath=None)

sub_dist = np.asarray(sub)[:min(50000, len(sub))]
plot_paired_distance_hist(Z_rna, Z_adt, sub_dist, outpath=None)


# %% [markdown]
### Perform Figure 3 cross-reconstruction experiments using the trained model and the test set

# %% [code]
import scipy.sparse as sp
from univi.evaluation import encode_adata

# -----------------------------
# 0) markers
# -----------------------------
def pick_existing(var_names, candidates):
    s = set(var_names)
    return [x for x in candidates if x in s]

# ---- ADT marker proteins ----
adt_markers_wanted = [
    "CD3-1","CD3-2","TCR-1","TCR-2",
    "CD4-1","CD4-2","CD8","CD8a",
    "CD45RA","CD45RO","CD27","CD28",
    "CD69","CD25","TIGIT","CD279",
    "CD56-1","CD56-2","CD16","CD335","CD314","CD244","CD57",
    "CD19","CD20","CD22","CD21",
    "CD79a","CD79b","IgD","IgM",
    "CD38-1","CD38-2","CD138-1","CD138-2","CD319","CD269",
    "CD14","CD16",
    "CD11b-1","CD11b-2","CD11c","CD13","CD15","CD33",
    "CD64","CD68","CD163",
    "HLA-DR","CD86","CD80",
    "CD1c","CD141","CD11c","HLA-DR",
    "CD123","CD303","CD304",
    "CD34","CD117","CD135","CD133-1","CD133-2",
]
adt_markers = pick_existing(adt_test_pp.var_names, adt_markers_wanted)
print("ADT markers used:", adt_markers)

# ---- RNA marker genes ----
rna_markers_wanted = [
    "PTPRC",
    "TRAC","TRBC1","CD3D","CD3E",
    "CD4","IL7R","CCR7","LTB","TCF7","LEF1",
    "CD8A","CD8B",
    "GZMK","GZMB","PRF1","NKG7","GNLY","FGFBP2",
    "KLRD1","TRAT1","LCK",
    "FOXP3","IL2RA","CTLA4","IKZF2","TIGIT",
    "KLRB1","TRDC","TRGC1","TRGC2",
    "MS4A1","CD79A","CD79B","CD74",
    "CD37","CD22","BANK1","TCL1A",
    "MZB1","JCHAIN","XBP1","TNFRSF17",
    "LYZ","LST1","TYROBP","S100A8","S100A9","FCN1","VCAN",
    "FCGR3A","MS4A7","LGALS3","CTSS","CX3CR1",
    "FCER1A","CD1C","CLEC10A",
    "CLEC4C","IL3RA","TCF4","GZMB",
    "CD34","KIT","MKI67","TOP2A",
    "PPBP","PF4",
]
rna_markers = pick_existing(rna_test_pp.var_names, rna_markers_wanted)
print("RNA markers used:", rna_markers)

# indices in RNA space for marker subset
rna_idx = np.array([rna_test_pp.var_names.get_loc(g) for g in rna_markers], dtype=int)


# %% [code]
# -----------------------------
# 1) paired sanity
# -----------------------------
assert rna_test_pp.n_obs == adt_test_pp.n_obs
assert (rna_test_pp.obs_names == adt_test_pp.obs_names).all()


# %% [code]
# -----------------------------
# 2) obs matrices (what the model saw)
# -----------------------------
def _dense_X(adata):
    X = adata.X
    if sp.issparse(X):
        X = X.toarray()
    return np.asarray(X, dtype=np.float32)

X_adt_obs = _dense_X(adt_test_pp)                 # CLR (dense)
X_rna_obs = _dense_X(rna_test_pp)                 # log1p HVG space
X_rna_obs_sub = X_rna_obs[:, rna_idx]


# %% [code]
# -----------------------------
# 3) cross-modal predictions (FULL)
# -----------------------------
@torch.no_grad()
def cross_modal_predict_all(model, adata_src, src_mod, tgt_mod, device="cpu", batch_size=512, use_moe=True):
    from univi.data import _get_matrix
    model.eval()

    X = _get_matrix(adata_src, layer=None, X_key="X")
    if sp.issparse(X):
        X = X.toarray()
    X = np.asarray(X, dtype=np.float32)

    dev = torch.device(device)
    out = []
    for start in range(0, X.shape[0], int(batch_size)):
        end = min(start + int(batch_size), X.shape[0])
        xb = torch.as_tensor(X[start:end], dtype=torch.float32, device=dev)

        mu_dict, logvar_dict = model.encode_modalities({src_mod: xb})
        if use_moe and hasattr(model, "mixture_of_experts"):
            mu_z, _ = model.mixture_of_experts(mu_dict, logvar_dict)
        else:
            mu_z = mu_dict[src_mod]

        xhat = model.decode_modalities(mu_z)[tgt_mod]  # (B, n_tgt)
        out.append(xhat.detach().cpu().numpy())

    return np.vstack(out).astype(np.float32, copy=False)

# RNA -> ADT (predict all proteins)
X_adt_hat = cross_modal_predict_all(model, rna_test_pp, "rna", "adt", device=device, batch_size=512)

# ADT -> RNA (predict full RNA HVG space, then subset markers)
X_rna_hat_full = cross_modal_predict_all(model, adt_test_pp, "adt", "rna", device=device, batch_size=512)
X_rna_hat_sub  = X_rna_hat_full[:, rna_idx]


# %% [code]
# -----------------------------
# 4) ONE fixed 2D embedding from observed RNA latent
# -----------------------------
def compute_fixed_umap_from_rep(adata, rep_key, out_umap_key="X_fig3_umap",
                               neighbors_key="fig3_neighbors",
                               n_neighbors=15, min_dist=0.5):
    a = adata.copy()

    # ---- hard reset any stale neighbor/umap state ----
    a.uns.pop("neighbors", None)
    a.uns.pop("umap", None)
    for k in ("connectivities", "distances"):
        if k in a.obsp:
            del a.obsp[k]
    # if scanpy stored keyed neighbors previously, remove them too
    if "neighbors" in a.uns and isinstance(a.uns["neighbors"], dict):
        a.uns["neighbors"].pop(neighbors_key, None)

    # ---- compute neighbors+umap from your rep ----
    sc.pp.neighbors(a, use_rep=rep_key, n_neighbors=n_neighbors, key_added=neighbors_key)
    sc.tl.umap(a, neighbors_key=neighbors_key, min_dist=min_dist)

    # write back just the 2D coords to your original object
    adata.obsm[out_umap_key] = a.obsm["X_umap"].copy()
    return adata


# %% [code]
# observed RNA latent (encoder mean)
Z_rna_obs = encode_adata(
    model,
    rna_test_pp,
    modality="rna",
    latent="modality_mean",
    device=device,
    batch_size=1024,
)
print("Z_rna_obs:", Z_rna_obs.shape)


# %% [code]
# usage
_tmp = rna_test_pp.copy()
_tmp.obsm["X_fig3_latent"] = Z_rna_obs

compute_fixed_umap_from_rep(
    _tmp,
    rep_key="X_fig3_latent",
    out_umap_key="X_fig3_umap",
    neighbors_key="fig3_neighbors",
    n_neighbors=15,
    min_dist=0.5,
)

# copy coords back (or just do it directly on rna_test_pp if you prefer)
rna_test_pp.obsm["X_fig3_umap"] = _tmp.obsm["X_fig3_umap"].copy()


# %% [code]
# -----------------------------
# 5) add obs/hat/delta onto rna_test_pp for plotting on X_fig3_umap
# -----------------------------
def add_obs(adata, key, values):
    adata.obs[key] = np.asarray(values, dtype=np.float32)

for p in adt_markers:
    j = adt_test_pp.var_names.get_loc(p)
    obs = X_adt_obs[:, j]
    hat = X_adt_hat[:, j]
    add_obs(rna_test_pp, f"ADT_obs_{p}", obs)
    add_obs(rna_test_pp, f"ADT_hat_{p}", hat)
    add_obs(rna_test_pp, f"ADT_delta_{p}", hat - obs)
    add_obs(rna_test_pp, f"ADT_absdelta_{p}", np.abs(hat - obs))

for t, g in enumerate(rna_markers):
    obs = X_rna_obs_sub[:, t]
    hat = X_rna_hat_sub[:, t]
    add_obs(rna_test_pp, f"RNA_obs_{g}", obs)
    add_obs(rna_test_pp, f"RNA_hat_{g}", hat)
    add_obs(rna_test_pp, f"RNA_delta_{g}", hat - obs)
    add_obs(rna_test_pp, f"RNA_absdelta_{g}", np.abs(hat - obs))

# Optional “activation score” per cell (mean absolute delta across your marker panels)
rna_test_pp.obs["ADT_activation_score"] = rna_test_pp.obs[[f"ADT_absdelta_{p}" for p in adt_markers]].mean(axis=1).astype(np.float32)
rna_test_pp.obs["RNA_activation_score"] = rna_test_pp.obs[[f"RNA_absdelta_{g}" for g in rna_markers]].mean(axis=1).astype(np.float32)


# %% [code]
# -----------------------------
# 6) quick plots (same coords for everything)
# -----------------------------
sc.pl.embedding(rna_test_pp, basis="X_fig3_umap", color=["celltype.l3"], size=4, frameon=False)

sc.pl.embedding(
    rna_test_pp,
    basis="X_fig3_umap",
    color=["ADT_activation_score", "RNA_activation_score"],
    ncols=2, size=4, frameon=False
)

# Example: plot first 6 proteins and first 6 genes as (obs | hat | delta)
def plot_triplet(prefix, feat, q=0.99, size=4):
    k_obs = f"{prefix}_obs_{feat}"
    k_hat = f"{prefix}_hat_{feat}"
    k_del = f"{prefix}_delta_{feat}"
    d = rna_test_pp.obs[k_del].to_numpy()
    lim = float(np.quantile(np.abs(d), q))
    sc.pl.embedding(rna_test_pp, basis="X_fig3_umap", color=[k_obs, k_hat], ncols=2, size=size, frameon=False)
    sc.pl.embedding(rna_test_pp, basis="X_fig3_umap", color=[k_del], size=size, frameon=False, vmin=-lim, vmax=lim)

#for p in adt_markers[:len(adt_markers)]:
for p in adt_markers[:5]:
    plot_triplet("ADT", p)

#for g in rna_markers[:len(rna_markers)]:
for g in rna_markers[:5]:
    plot_triplet("RNA", g)


# %% [markdown]
### Additional analysis plots to interrogate the above results

# %% [code]
print(combo)
rna_x_univi = combo[combo.obs['modality'] == 'rna']
adt_x_univi = combo[combo.obs['modality'] == 'adt']


# %% [code]
sc.pl.embedding(
    rna_x_univi,
    basis='X_umap',
    color=[
        "modality",
        "celltype.l3",
    ],
    ncols=2,
    size=3,
    frameon=False,
    wspace=0.35,
)


# %% [code]
rna_test_pp.obsm['X_fig3_umap'] = rna_x_univi.obsm['X_umap']
adt_test_pp.obsm['X_fig3_umap'] = adt_x_univi.obsm['X_umap']


# %% [code]
#print(set(rna_test_pp.var_names))
#print(set(adt_test_pp.var_names))


# %% [code]
X_rna_hat = X_rna_hat_full.copy()


# %% [code]
# Assumes already have
# - rna_test_pp, adt_test_pp (paired + aligned)
# - X_rna_obs, X_rna_hat (n_cells x n_rna_features) in SAME var order as rna_test_pp.var_names
# - X_adt_obs, X_adt_hat (n_cells x n_adt_features) in SAME var order as adt_test_pp.var_names
# - rna_markers (list of genes) and adt_markers (list of proteins) that exist in var_names
# - a fixed UMAP basis on rna_test_pp: rna_test_pp.obsm["X_umap_fig3"]
#
# If your basis key differs, set:
UMAP_BASIS = "X_fig3_umap"

import os
import numpy as np
import pandas as pd
import scipy.sparse as sp
import matplotlib.pyplot as plt

from sklearn.metrics import r2_score
from sklearn.neighbors import NearestNeighbors
from scipy.stats import spearmanr


# -----------------------------
# 0) output / plotting defaults
# -----------------------------
outdir_analysis = os.path.join(outdir, "additional_analysis")
os.makedirs(outdir_analysis, exist_ok=True)

def _finish(fig=None, outpath=None, dpi=400, show=True, close=True):
    if outpath is not None:
        plt.savefig(outpath, dpi=dpi, bbox_inches="tight")
    if show:
        plt.show()
    if close:
        plt.close(fig if fig is not None else plt.gcf())


def _as_float32(X):
    return np.asarray(X, dtype=np.float32)

def _check_shapes():
    assert X_rna_obs.shape == X_rna_hat.shape, (X_rna_obs.shape, X_rna_hat.shape)
    assert X_adt_obs.shape == X_adt_hat.shape, (X_adt_obs.shape, X_adt_hat.shape)
    assert X_rna_obs.shape[0] == rna_test_pp.n_obs
    assert X_adt_obs.shape[0] == adt_test_pp.n_obs
    assert X_rna_obs.shape[1] == rna_test_pp.n_vars
    assert X_adt_obs.shape[1] == adt_test_pp.n_vars

_check_shapes()

# robust paired checks (important for per-cell summaries)
assert rna_test_pp.n_obs == adt_test_pp.n_obs
assert (rna_test_pp.obs_names == adt_test_pp.obs_names).all()


# %% [code]
# -----------------------------
# 1) attach per-feature / per-cell summaries
# -----------------------------
def per_feature_pearson(X, Y):
    X = _as_float32(X); Y = _as_float32(Y)
    Xc = X - X.mean(axis=0, keepdims=True)
    Yc = Y - Y.mean(axis=0, keepdims=True)
    num = (Xc * Yc).sum(axis=0)
    den = np.sqrt((Xc**2).sum(axis=0) * (Yc**2).sum(axis=0)) + 1e-8
    return num / den

def per_feature_spearman(X, Y):
    # fast-ish: rank by argsort-of-argsort per feature
    Xr = np.argsort(np.argsort(X, axis=0), axis=0).astype(np.float32)
    Yr = np.argsort(np.argsort(Y, axis=0), axis=0).astype(np.float32)
    return per_feature_pearson(Xr, Yr)

def per_feature_rmse(X, Y):
    X = _as_float32(X); Y = _as_float32(Y)
    return np.sqrt(np.mean((X - Y) ** 2, axis=0))

def per_feature_mae(X, Y):
    X = _as_float32(X); Y = _as_float32(Y)
    return np.mean(np.abs(X - Y), axis=0)

# Per-feature stats (ALL features)
adt_stats_all = pd.DataFrame({
    "feature": adt_test_pp.var_names.to_numpy(),
    "pearson_r": per_feature_pearson(X_adt_obs, X_adt_hat),
    "spearman_r": per_feature_spearman(X_adt_obs, X_adt_hat),
    "rmse": per_feature_rmse(X_adt_obs, X_adt_hat),
    "mae": per_feature_mae(X_adt_obs, X_adt_hat),
})
rna_stats_all = pd.DataFrame({
    "feature": rna_test_pp.var_names.to_numpy(),
    "pearson_r": per_feature_pearson(X_rna_obs, X_rna_hat),
    "spearman_r": per_feature_spearman(X_rna_obs, X_rna_hat),
    "rmse": per_feature_rmse(X_rna_obs, X_rna_hat),
    "mae": per_feature_mae(X_rna_obs, X_rna_hat),
})

# marker-only views
adt_stats_markers = adt_stats_all[adt_stats_all["feature"].isin(list(adt_markers))].copy()
rna_stats_markers = rna_stats_all[rna_stats_all["feature"].isin(list(rna_markers))].copy()

# save tables
adt_stats_all.to_csv(os.path.join(outdir_analysis, "adt_feature_stats_all.csv"), index=False)
rna_stats_all.to_csv(os.path.join(outdir_analysis, "rna_feature_stats_all.csv"), index=False)
adt_stats_markers.to_csv(os.path.join(outdir_analysis, "adt_feature_stats_markers.csv"), index=False)
rna_stats_markers.to_csv(os.path.join(outdir_analysis, "rna_feature_stats_markers.csv"), index=False)


# Per-cell summaries (all features and marker-only)
def per_cell_scores(X_obs, X_hat, feat_idx=None):
    Xo = _as_float32(X_obs)
    Xh = _as_float32(X_hat)
    if feat_idx is not None:
        Xo = Xo[:, feat_idx]
        Xh = Xh[:, feat_idx]
    resid = Xh - Xo
    mae = np.mean(np.abs(resid), axis=1)
    rmse = np.sqrt(np.mean(resid**2, axis=1))
    # per-cell corr across features (guard against constant vectors)
    xo = Xo - Xo.mean(axis=1, keepdims=True)
    xh = Xh - Xh.mean(axis=1, keepdims=True)
    num = np.sum(xo * xh, axis=1)
    den = np.sqrt(np.sum(xo**2, axis=1) * np.sum(xh**2, axis=1)) + 1e-8
    pr = num / den
    return mae.astype(np.float32), rmse.astype(np.float32), pr.astype(np.float32)

# indices for markers (safe even if markers empty)
rna_marker_idx = np.array([rna_test_pp.var_names.get_loc(g) for g in rna_markers], dtype=int) if len(rna_markers) else None
adt_marker_idx = np.array([adt_test_pp.var_names.get_loc(p) for p in adt_markers], dtype=int) if len(adt_markers) else None

rna_mae_all, rna_rmse_all, rna_pr_all = per_cell_scores(X_rna_obs, X_rna_hat, feat_idx=None)
adt_mae_all, adt_rmse_all, adt_pr_all = per_cell_scores(X_adt_obs, X_adt_hat, feat_idx=None)

if rna_marker_idx is not None and rna_marker_idx.size:
    rna_mae_mk, rna_rmse_mk, rna_pr_mk = per_cell_scores(X_rna_obs, X_rna_hat, feat_idx=rna_marker_idx)
else:
    rna_mae_mk = rna_rmse_mk = rna_pr_mk = np.full(rna_test_pp.n_obs, np.nan, dtype=np.float32)

if adt_marker_idx is not None and adt_marker_idx.size:
    adt_mae_mk, adt_rmse_mk, adt_pr_mk = per_cell_scores(X_adt_obs, X_adt_hat, feat_idx=adt_marker_idx)
else:
    adt_mae_mk = adt_rmse_mk = adt_pr_mk = np.full(adt_test_pp.n_obs, np.nan, dtype=np.float32)

# attach to rna_test_pp for easy embedding plots on a single basis
rna_test_pp.obs["RNA_xrec_mae_all"] = rna_mae_all
rna_test_pp.obs["RNA_xrec_rmse_all"] = rna_rmse_all
rna_test_pp.obs["RNA_xrec_pearson_all"] = rna_pr_all

rna_test_pp.obs["ADT_xrec_mae_all"] = adt_mae_all
rna_test_pp.obs["ADT_xrec_rmse_all"] = adt_rmse_all
rna_test_pp.obs["ADT_xrec_pearson_all"] = adt_pr_all

rna_test_pp.obs["RNA_xrec_mae_markers"] = rna_mae_mk
rna_test_pp.obs["RNA_xrec_rmse_markers"] = rna_rmse_mk
rna_test_pp.obs["RNA_xrec_pearson_markers"] = rna_pr_mk

rna_test_pp.obs["ADT_xrec_mae_markers"] = adt_mae_mk
rna_test_pp.obs["ADT_xrec_rmse_markers"] = adt_rmse_mk
rna_test_pp.obs["ADT_xrec_pearson_markers"] = adt_pr_mk


# %% [code]
# -----------------------------
# 2) UMAP diagnostics (per-cell error / agreement)
# -----------------------------
# These are great “where does it fail?” maps.
import scanpy as sc

sc.pl.embedding(
    rna_test_pp,
    basis=UMAP_BASIS,
    color=[
        "RNA_xrec_mae_markers",
        "ADT_xrec_mae_markers",
        "RNA_xrec_pearson_markers",
        "ADT_xrec_pearson_markers",
    ],
    ncols=2,
    size=3,
    frameon=False,
    wspace=0.35,
)


# %% [code]
# -----------------------------
# 3) feature performance plots (markers + global extremes)
# -----------------------------
def plot_feature_perf(df, title, top=20, bottom=20, outpath=None):
    d0 = df.sort_values("pearson_r", ascending=False).copy()
    top = min(int(top), len(d0))
    bottom = min(int(bottom), len(d0))
    d = pd.concat([d0.head(top), d0.tail(bottom)], axis=0)

    fig, ax = plt.subplots(figsize=(7.5, 0.22 * len(d) + 1.0))
    ax.barh(d["feature"], d["pearson_r"])
    ax.invert_yaxis()

    ax.set_xlim(-1, 1)
    ax.set_xlabel("Pearson r (obs vs pred)")
    ax.set_title(title, pad=4)
    ax.grid(False)

    # ---- tighten top/bottom whitespace ----
    ax.margins(y=0)                    # removes default 5% y padding
    ax.set_ylim(len(d) - 0.5, -0.5)    # tight bounds around bars (works with invert_yaxis too)

    fig.tight_layout(pad=0.2)
    _finish(fig, outpath=outpath)

'''
plot_feature_perf(
    adt_stats_all, "ADT (RNA→ADT): best/worst proteins (all)",
    top=25, bottom=25,
    outpath=os.path.join(outdir_analysis, "adt_feature_perf_best_worst.png"),
)
plot_feature_perf(
    rna_stats_all, "RNA (ADT→RNA): best/worst genes (all)",
    top=25, bottom=25,
    outpath=os.path.join(outdir_analysis, "rna_feature_perf_best_worst.png"),
)
'''

if len(adt_stats_markers):
    plot_feature_perf(
        adt_stats_markers, "ADT (RNA→ADT): marker proteins",
        top=len(adt_stats_markers), bottom=0,
        outpath=os.path.join(outdir_analysis, "adt_marker_perf_reproducibility.png"),
    )

if len(rna_stats_markers):
    plot_feature_perf(
        rna_stats_markers, "RNA (ADT→RNA): marker genes",
        top=len(rna_stats_markers), bottom=0,
        outpath=os.path.join(outdir_analysis, "rna_marker_perf_reproducibility.png"),
    )


# %% [code]
# -----------------------------
# 4) obs-vs-pred scatter/hexbin + calibration for markers
# -----------------------------
def calibration_bins(x_obs, x_pred, n_bins=30):
    x_obs = np.asarray(x_obs).ravel()
    x_pred = np.asarray(x_pred).ravel()
    qs = np.quantile(x_pred, np.linspace(0, 1, n_bins + 1))
    qs = np.unique(qs)
    if qs.size < 3:
        return None
    bin_id = np.digitize(x_pred, qs[1:-1], right=True)
    mu_obs  = np.array([x_obs[bin_id == b].mean() if np.any(bin_id == b) else np.nan for b in range(qs.size - 1)])
    mu_pred = np.array([x_pred[bin_id == b].mean() if np.any(bin_id == b) else np.nan for b in range(qs.size - 1)])
    return mu_pred, mu_obs

def plot_calibration(x_obs, x_pred, title, n_bins=30, outpath=None):
    out = calibration_bins(x_obs, x_pred, n_bins=n_bins)
    if out is None:
        print(f"[calibration] skipped: {title} (not enough unique values)")
        return
    mu_pred, mu_obs = out
    fig, ax = plt.subplots(figsize=(5.3, 4.8))
    ax.plot(mu_pred, mu_obs, marker="o", linewidth=1)
    lo = np.nanmin([mu_pred.min(), mu_obs.min()])
    hi = np.nanmax([mu_pred.max(), mu_obs.max()])
    ax.plot([lo, hi], [lo, hi], linewidth=1)
    ax.set_title(title)
    ax.set_xlabel("Mean predicted (bin)")
    ax.set_ylabel("Mean observed (bin)")
    ax.grid(False)
    fig.tight_layout()
    _finish(fig, outpath=outpath)

def hexbin_obs_pred(x_obs, x_pred, title, gridsize=80, outpath=None):
    x_obs = np.asarray(x_obs).ravel()
    x_pred = np.asarray(x_pred).ravel()
    fig, ax = plt.subplots(figsize=(5.5, 5.0))
    hb = ax.hexbin(x_obs, x_pred, gridsize=gridsize, mincnt=1)
    ax.set_title(title)
    ax.set_xlabel("Observed")
    ax.set_ylabel("Predicted")
    lo = np.nanpercentile(np.concatenate([x_obs, x_pred]), 1)
    hi = np.nanpercentile(np.concatenate([x_obs, x_pred]), 99)
    ax.plot([lo, hi], [lo, hi], linewidth=1)
    ax.grid(False)
    fig.colorbar(hb, ax=ax, label="count")
    fig.tight_layout()
    _finish(fig, outpath=outpath)

# ADT marker examples
for p in list(adt_markers)[:6]:
    j = adt_test_pp.var_names.get_loc(p)
    hexbin_obs_pred(
        X_adt_obs[:, j], X_adt_hat[:, j],
        title=f"ADT (RNA→ADT): {p}",
        outpath=os.path.join(outdir_analysis, f"hexbin_ADT_{p}.png"),
    )
    plot_calibration(
        X_adt_obs[:, j], X_adt_hat[:, j],
        title=f"Calibration (RNA→ADT): {p}",
        n_bins=40,
        outpath=os.path.join(outdir_analysis, f"calib_ADT_{p}.png"),
    )

# RNA marker examples
for g in list(rna_markers)[:6]:
    j = rna_test_pp.var_names.get_loc(g)
    hexbin_obs_pred(
        X_rna_obs[:, j], X_rna_hat[:, j],
        title=f"RNA (ADT→RNA): {g}",
        outpath=os.path.join(outdir_analysis, f"hexbin_RNA_{g}.png"),
    )
    plot_calibration(
        X_rna_obs[:, j], X_rna_hat[:, j],
        title=f"Calibration (ADT→RNA): {g}",
        n_bins=40,
        outpath=os.path.join(outdir_analysis, f"calib_RNA_{g}.png"),
    )


# %% [code]
# -----------------------------
# 5) kNN “where is error concentrated?” using the UMAP neighbor graph
# -----------------------------
# This uses neighbors computed on the fixed UMAP rep (or at least same cells).
# If absent, compute a fresh graph and avoid Scanpy's stale/broken .uns["neighbors"].
def ensure_neighbors_for_basis(adata, basis_key, neighbors_key="analysis_neighbors", n_neighbors=30, metric="euclidean"):
    # Already computed?
    if neighbors_key in adata.uns and f"{neighbors_key}_connectivities" in adata.obsp:
        return

    # ---- HARD RESET: clear broken/default neighbors state that can crash Scanpy ----
    adata.uns.pop("neighbors", None)
    adata.uns.pop("umap", None)
    for k in ("connectivities", "distances"):
        if k in adata.obsp:
            del adata.obsp[k]

    # Clear any partial keyed neighbors too
    adata.uns.pop(neighbors_key, None)
    for k in (f"{neighbors_key}_connectivities", f"{neighbors_key}_distances"):
        if k in adata.obsp:
            del adata.obsp[k]

    # Use the embedding in .obsm as the representation
    if basis_key not in adata.obsm:
        raise KeyError(f"{basis_key} not in adata.obsm (available: {list(adata.obsm.keys())})")

    sc.pp.neighbors(
        adata,
        use_rep=basis_key,          # yes, neighbors can be computed on a 2D embedding
        n_neighbors=int(n_neighbors),
        metric=metric,
        key_added=neighbors_key,
    )

def local_smooth(values, conn):
    # conn: sparse connectivities matrix (n x n)
    v = _as_float32(values).reshape(-1, 1)
    w = conn @ v
    d = np.asarray(conn.sum(axis=1)).ravel().astype(np.float32) + 1e-8
    return (w.ravel() / d).astype(np.float32)

# build neighbor graph on the 2D embedding just for smoothing/regions
ensure_neighbors_for_basis(rna_test_pp, UMAP_BASIS, neighbors_key="analysis_neighbors", n_neighbors=30)
conn = rna_test_pp.obsp["analysis_neighbors_connectivities"]

rna_test_pp.obs["RNA_xrec_mae_markers_smoothed"] = local_smooth(
    rna_test_pp.obs["RNA_xrec_mae_markers"].to_numpy(), conn
)
rna_test_pp.obs["ADT_xrec_mae_markers_smoothed"] = local_smooth(
    rna_test_pp.obs["ADT_xrec_mae_markers"].to_numpy(), conn
)

sc.pl.embedding(
    rna_test_pp,
    basis=UMAP_BASIS,
    color=["RNA_xrec_mae_markers_smoothed", "ADT_xrec_mae_markers_smoothed"],
    ncols=2, size=3, frameon=False, wspace=0.35
)


# %% [code]
# -----------------------------
# 6) “hard vs easy” cells: top error quantiles + marker panels
# -----------------------------
def label_hard_cells(adata, key, q=0.95, out_key=None):
    x = adata.obs[key].to_numpy()
    thr = float(np.nanquantile(x, q))
    out_key = out_key or f"{key}_hard_q{int(q*100)}"
    adata.obs[out_key] = (x >= thr)
    print(f"[hard-cells] {out_key}: threshold={thr:.4g}, n_hard={(adata.obs[out_key]).sum()}/{adata.n_obs}")
    return out_key

k_hard_rna = label_hard_cells(rna_test_pp, "RNA_xrec_mae_markers", q=0.95)
k_hard_adt = label_hard_cells(rna_test_pp, "ADT_xrec_mae_markers", q=0.95)

sc.pl.embedding(
    rna_test_pp,
    basis=UMAP_BASIS,
    color=[k_hard_rna, k_hard_adt, "celltype.l2"],
    ncols=3, size=3, frameon=False, wspace=0.35
)


# %% [code]
# Optional: compare marker expression between hard/easy for one or two markers
def violin_hard_vs_easy(adata, value_key, hard_key, title=None, outpath=None):
    hard = adata.obs[hard_key].to_numpy().astype(bool)
    x1 = adata.obs[value_key].to_numpy()[~hard]
    x2 = adata.obs[value_key].to_numpy()[hard]
    fig, ax = plt.subplots(figsize=(5.2, 4.0))
    ax.violinplot([x1, x2], showmeans=True, showextrema=False)
    ax.set_xticks([1, 2])
    ax.set_xticklabels(["easy", "hard"])
    ax.set_ylabel(value_key)
    ax.set_title(title or f"{value_key} (easy vs hard)")
    ax.grid(False)
    fig.tight_layout()
    _finish(fig, outpath=outpath)

# If you already created obs/hat/delta columns earlier, you can point these at them:
# e.g., violin_hard_vs_easy(rna_test_pp, "ADT_absdelta_CD3-1", k_hard_adt)
# Here we just show the per-cell MAE as an example:
violin_hard_vs_easy(
    rna_test_pp,
    "RNA_xrec_mae_markers",
    k_hard_rna,
    title="RNA cross-recon error (markers): easy vs hard",
    outpath=os.path.join(outdir_analysis, "violin_RNA_mae_markers_easy_vs_hard_reproducibility.png"),
)
violin_hard_vs_easy(
    rna_test_pp,
    "ADT_xrec_mae_markers",
    k_hard_adt,
    title="ADT cross-recon error (markers): easy vs hard",
    outpath=os.path.join(outdir_analysis, "violin_ADT_mae_markers_easy_vs_hard_reproducibility.png"),
)


# %% [code]
# -----------------------------
# 7) report a short summary block
# -----------------------------
def summarize_df(df, name, features=None):
    if features is not None:
        df = df[df["feature"].isin(features)].copy()
    print(f"\n=== {name} ===")
    if len(df) == 0:
        print("(empty)")
        return
    print("n_features:", len(df))
    print("pearson_r median / mean:", float(df["pearson_r"].median()), float(df["pearson_r"].mean()))
    print("pearson_r 10th/90th:", float(df["pearson_r"].quantile(0.1)), float(df["pearson_r"].quantile(0.9)))
    print("worst 5 (pearson_r):")
    display(df.sort_values("pearson_r").head(5))
    print("best 5 (pearson_r):")
    display(df.sort_values("pearson_r", ascending=False).head(5))

summarize_df(adt_stats_all, "ADT (RNA→ADT) all proteins")
summarize_df(rna_stats_all, "RNA (ADT→RNA) all genes")
summarize_df(adt_stats_markers, "ADT (RNA→ADT) marker proteins", features=adt_markers)
summarize_df(rna_stats_markers, "RNA (ADT→RNA) marker genes", features=rna_markers)

print(f"\nSaved additional analysis outputs to: {outdir_analysis}")


# %% [code]
from scipy.cluster.hierarchy import linkage, leaves_list
from scipy.spatial.distance import pdist
import numpy as np
import matplotlib.pyplot as plt

def cluster_order_df(df, axis=0, metric="correlation", method="average"):
    M = df.to_numpy(dtype=np.float32)
    if axis == 1:
        M = M.T
        labels = np.array(df.columns)
    else:
        labels = np.array(df.index)

    # correlation distance works well for marker-pattern similarity
    D = pdist(M, metric=metric)
    Z = linkage(D, method=method)
    ord_idx = leaves_list(Z)
    return labels[ord_idx]

def plot_heatmap_ordered(df, title, row_order=None, col_order=None, vmin=None, vmax=None, cmap="viridis"):
    d = df.copy()
    if row_order is not None:
        d = d.reindex(row_order)
    if col_order is not None:
        d = d.loc[:, col_order]

    M = d.to_numpy(dtype=np.float32)

    fig_h = max(4.0, 0.28 * d.shape[0] + 1.5)
    fig_w = max(6.0, 0.35 * d.shape[1] + 2.5)
    fig, ax = plt.subplots(figsize=(fig_w, fig_h))
    im = ax.imshow(M, aspect="auto", interpolation="nearest", cmap=cmap, vmin=vmin, vmax=vmax)

    ax.set_title(title, pad=10)
    ax.set_xlabel("markers")
    ax.set_ylabel("celltype")
    ax.set_xticks(np.arange(d.shape[1]))
    ax.set_xticklabels(d.columns, rotation=90)
    ax.set_yticks(np.arange(d.shape[0]))
    ax.set_yticklabels(d.index)

    cbar = fig.colorbar(im, ax=ax, fraction=0.046, pad=0.04)
    cbar.set_label("mean expression (Z-score)")
    ax.grid(False)
    fig.tight_layout()
    plt.show()
    plt.close(fig)


# %% [code]
import numpy as np
import pandas as pd
import scipy.sparse as sp

def mean_by_celltype(
    X,
    adata,
    features,
    celltype_key="celltype.l2",
    use="var_names",          # "var_names" (genes/proteins) OR "obs_names" if X is in cell-order only
    agg="mean",               # "mean" or "median"
    fill_missing=np.nan,
    sort_celltypes=True,
):
    
    if celltype_key not in adata.obs:
        raise KeyError(f"{celltype_key!r} not in adata.obs")

    # ---- ensure dense float32 (safe + consistent)
    if sp.issparse(X):
        X = X.toarray()
    X = np.asarray(X, dtype=np.float32)

    # ---- map features -> column indices (keep requested order)
    feats = list(features)
    if use != "var_names":
        raise ValueError("Only use='var_names' is supported for now.")

    name_to_idx = {k: i for i, k in enumerate(adata.var_names.to_numpy())}
    idx = [name_to_idx.get(f, -1) for f in feats]

    # subset existing cols; remember where to re-insert missing
    keep_pos = [j for j, k in enumerate(idx) if k >= 0]
    keep_idx = [k for k in idx if k >= 0]

    if len(keep_idx) == 0:
        # nothing matched: return empty-ish frame with correct shape
        celltypes = adata.obs[celltype_key].astype(str)
        ct_order = sorted(celltypes.unique()) if sort_celltypes else celltypes.unique().tolist()
        return pd.DataFrame(fill_missing, index=ct_order, columns=feats, dtype=np.float32)

    Xsub = X[:, keep_idx]  # (n_cells, n_keep_features)

    # ---- group by celltype
    celltypes = adata.obs[celltype_key].astype(str).to_numpy()
    ct_vals = pd.Categorical(celltypes)  # preserves observed order; we can sort later
    ct_categories = ct_vals.categories.tolist()
    if sort_celltypes:
        ct_categories = sorted(ct_categories)

    # build output for kept features first
    out_kept = {}
    for ct in ct_categories:
        mask = (celltypes == ct)
        if not np.any(mask):
            continue
        block = Xsub[mask]
        if agg == "mean":
            out_kept[ct] = block.mean(axis=0)
        elif agg == "median":
            out_kept[ct] = np.median(block, axis=0)
        else:
            raise ValueError("agg must be 'mean' or 'median'")

    df_kept = pd.DataFrame.from_dict(out_kept, orient="index")
    df_kept.index.name = celltype_key
    df_kept.columns = [feats[j] for j in keep_pos]

    # ---- reinsert missing features as fill_missing, preserving requested column order
    df = pd.DataFrame(index=df_kept.index, columns=feats, dtype=np.float32)
    df.loc[:, :] = fill_missing
    df.loc[:, df_kept.columns] = df_kept

    return df


# %% [code]
# ----------------------------
# Build celltype means
# ----------------------------
# RNA: observed vs ADT→RNA predicted
rna_obs_ct = mean_by_celltype(X_rna_obs, rna_test_pp, rna_markers, celltype_key)
rna_hat_ct = mean_by_celltype(X_rna_hat, rna_test_pp, rna_markers, celltype_key)

# ADT: observed vs RNA→ADT predicted
adt_obs_ct = mean_by_celltype(X_adt_obs, adt_test_pp, adt_markers, celltype_key)
adt_hat_ct = mean_by_celltype(X_adt_hat, adt_test_pp, adt_markers, celltype_key)

# Align row order across obs/hat (same celltype ordering)
rna_hat_ct = rna_hat_ct.reindex(rna_obs_ct.index)
adt_hat_ct = adt_hat_ct.reindex(adt_obs_ct.index)


# %% [code]
# Compute ONE row order from observed (or from delta), then reuse for all panels
#row_order = cluster_order_df(rna_obs_ct, axis=0, metric="correlation", method="average")
row_order = cluster_order_df(adt_obs_ct, axis=0, metric="correlation", method="average")

# Keep columns in your existing marker order
rna_col_order = list(rna_obs_ct.columns)


# %% [code]
# Scanpy defaults (affects sc.pl.*)
sc.set_figure_params(
    figsize=(10, 8),    # bigger canvas
    dpi=100,            # on-screen sharpness
    dpi_save=300,       # saved file sharpness
    fontsize=10,
    frameon=False,
)

# Matplotlib defaults (affects plt.*)
plt.rcParams.update({
    "figure.figsize": (10, 8),
    "figure.dpi": 100,
    "savefig.dpi": 300,
    "savefig.bbox": "tight",
    "savefig.pad_inches": 0.1,
    "axes.titlesize": 12,
    "axes.labelsize": 10,
    "legend.fontsize": 10,
})


# %% [code]
# ----------------------------
# Plot for RNA
# ----------------------------
plot_heatmap_ordered(rna_obs_ct, "RNA observed (mean by celltype.l3)", row_order=row_order, col_order=rna_col_order, vmin=-0.5, vmax=5)
plot_heatmap_ordered(rna_hat_ct, "RNA predicted (ADT→RNA) (mean by celltype.l3)", row_order=row_order, col_order=rna_col_order, vmin=-0.5, vmax=5)

# Delta is super informative on a symmetric scale
rna_delta_ct = (rna_hat_ct - rna_obs_ct).reindex(rna_obs_ct.index).loc[:, rna_col_order]
plot_heatmap_ordered(rna_delta_ct, "RNA delta (pred - obs)", row_order=row_order, col_order=rna_col_order, vmin=-1.25, vmax=1.25)


# %% [code]
# Keep columns in your existing marker order
adt_col_order = list(adt_obs_ct.columns)


# %% [code]
# ----------------------------
# Plot for ADT
# ----------------------------
plot_heatmap_ordered(adt_obs_ct, "ADT observed (mean by celltype.l3)", row_order=row_order, col_order=adt_col_order, vmin=-0.75, vmax=1.5)
plot_heatmap_ordered(adt_hat_ct, "ADT predicted (RNA→ADT) (mean by celltype.l3)", row_order=row_order, col_order=adt_col_order, vmin=-0.75, vmax=1.5)

# Delta is super informative on a symmetric scale
adt_delta_ct = (adt_hat_ct - adt_obs_ct).reindex(adt_obs_ct.index).loc[:, adt_col_order]
plot_heatmap_ordered(adt_delta_ct, "ADT delta (pred - obs)", row_order=row_order, col_order=adt_col_order, vmin=-0.25, vmax=0.25)


# %% [code]


# %% [markdown]
### Main Figure 3 stuff (actually this time)

# %% [code]
import os
import numpy as np
import pandas as pd
import scipy.sparse as sp
import matplotlib.pyplot as plt
import scanpy as sc

# -----------------------------
# 0) small utils
# -----------------------------
def _ensure_dir(p):
    os.makedirs(p, exist_ok=True)
    return p

def _as_float32(X):
    if sp.issparse(X):
        X = X.toarray()
    return np.asarray(X, dtype=np.float32)

def _basis_name(basis):
    # scanpy expects basis="umap" for obsm["X_umap"]
    # if you pass "X_umap" it will often look for "X_X_umap" -> bad
    return basis[2:] if isinstance(basis, str) and basis.startswith("X_") else basis

def _scanpy_fig_from_plot():
    # scanpy plots into the current matplotlib figure in many versions
    return plt.gcf()

# -----------------------------
# 1) your metrics (as-is)
# -----------------------------
def per_feature_pearson(X, Y):
    X = _as_float32(X); Y = _as_float32(Y)
    Xc = X - X.mean(axis=0, keepdims=True)
    Yc = Y - Y.mean(axis=0, keepdims=True)
    num = (Xc * Yc).sum(axis=0)
    den = np.sqrt((Xc**2).sum(axis=0) * (Yc**2).sum(axis=0)) + 1e-8
    return num / den

def per_feature_spearman(X, Y):
    Xr = np.argsort(np.argsort(_as_float32(X), axis=0), axis=0).astype(np.float32)
    Yr = np.argsort(np.argsort(_as_float32(Y), axis=0), axis=0).astype(np.float32)
    return per_feature_pearson(Xr, Yr)

def per_feature_rmse(X, Y):
    X = _as_float32(X); Y = _as_float32(Y)
    return np.sqrt(np.mean((X - Y) ** 2, axis=0))

def per_feature_mae(X, Y):
    X = _as_float32(X); Y = _as_float32(Y)
    return np.mean(np.abs(X - Y), axis=0)

def per_cell_scores(X_obs, X_hat, feat_idx=None):
    Xo = _as_float32(X_obs)
    Xh = _as_float32(X_hat)
    if feat_idx is not None:
        Xo = Xo[:, feat_idx]
        Xh = Xh[:, feat_idx]
    resid = Xh - Xo
    mae = np.mean(np.abs(resid), axis=1)
    rmse = np.sqrt(np.mean(resid**2, axis=1))
    xo = Xo - Xo.mean(axis=1, keepdims=True)
    xh = Xh - Xh.mean(axis=1, keepdims=True)
    num = np.sum(xo * xh, axis=1)
    den = np.sqrt(np.sum(xo**2, axis=1) * np.sum(xh**2, axis=1)) + 1e-8
    pr = num / den
    return mae.astype(np.float32), rmse.astype(np.float32), pr.astype(np.float32)

# -----------------------------
# 2) plotting helpers
# -----------------------------
def plot_feature_perf(df, title, metric_col="pearson_r", top=25, bottom=25,
                      outpath=None, xlim=(-1, 1), show=True, close=False):
    d0 = df.sort_values(metric_col, ascending=False).copy()
    top = min(int(top), len(d0))
    bottom = min(int(bottom), len(d0))
    d = pd.concat([d0.head(top), d0.tail(bottom)], axis=0)

    fig, ax = plt.subplots(figsize=(7.5, 0.22 * len(d) + 1.2))
    ax.barh(d["feature"], d[metric_col])
    ax.invert_yaxis()
    ax.set_xlim(*xlim)
    ax.set_xlabel(metric_col)
    ax.set_title(title, pad=6)
    ax.grid(False)

    # ---- tighten top/bottom whitespace ----
    ax.margins(y=0)                    # removes default 5% y padding
    ax.set_ylim(len(d) - 0.5, -0.5)    # tightbounds around bars (works with invert_yaxis too)

    fig.tight_layout(pad=0.2)

    if outpath is not None:
        _ensure_dir(os.path.dirname(outpath))
        fig.savefig(outpath, dpi=300, bbox_inches="tight")
    if show:
        plt.show()
    if close:
        plt.close(fig)
    return fig

def add_obs_hat_to_adata_obs(adata, X_obs, X_hat, feat_names, feature_axis_names, prefix="RNA_"):
    """
    Adds columns like:
      f"{prefix}obs_{FEAT}" and f"{prefix}hat_{FEAT}"
    into adata.obs.
    """
    Xo = _as_float32(X_obs)
    Xh = _as_float32(X_hat)

    feat_names = list(feat_names)
    axis = pd.Index(feature_axis_names)

    kept = []
    for f in feat_names:
        if f not in axis:
            continue
        j = axis.get_loc(f)
        adata.obs[f"{prefix}obs_{f}"] = Xo[:, j]
        adata.obs[f"{prefix}hat_{f}"] = Xh[:, j]
        kept.append(f)

    missing = sorted(set(feat_names) - set(kept))
    if missing:
        print(f"[warn] {prefix} missing {len(missing)} features (skipped). Example: {missing[:5]}")
    return kept

def plot_umap_obs_vs_hat(adata, basis, feats, prefix, outdir,
                         ncols=2, size=10, show_first_n=0, close=True):
    """
    For each feat, saves a 2-panel obs vs hat UMAP.
    show_first_n: display the first N figures in notebook (to avoid spam).
    """
    basis = _basis_name(basis)
    outdir = _ensure_dir(outdir)
    shown = 0

    for f in feats:
        cols = [f"{prefix}obs_{f}", f"{prefix}hat_{f}"]
        cols = [c for c in cols if c in adata.obs.columns]
        if len(cols) != 2:
            continue

        sc.pl.embedding(
            adata,
            basis=basis,
            color=cols,
            ncols=ncols,
            size=size,
            frameon=False,
            wspace=0.35,
            show=(shown < int(show_first_n)),
        )
        fig = _scanpy_fig_from_plot()
        fig.savefig(os.path.join(outdir, f"{prefix}{f}_obs_vs_hat_umap.png"),
                    dpi=300, bbox_inches="tight")
        if close or (shown >= int(show_first_n)):
            plt.close(fig)
        shown += 1

def _group_means(X, groups):
    # groups: (N,) array-like of strings
    X = _as_float32(X)
    g = pd.Categorical(groups)
    levels = list(g.categories)
    out = np.zeros((len(levels), X.shape[1]), dtype=np.float32)
    for i, lab in enumerate(levels):
        idx = np.where(g == lab)[0]
        if idx.size:
            out[i] = X[idx].mean(axis=0)
        else:
            out[i] = np.nan
    return levels, out

def _zscore_cols(M):
    M = np.asarray(M, dtype=np.float32)
    mu = np.nanmean(M, axis=0, keepdims=True)
    sd = np.nanstd(M, axis=0, keepdims=True) + 1e-8
    return (M - mu) / sd

def plot_obs_pred_delta_heatmaps(X_obs, X_hat, feature_names, groups,
                                 title_prefix, outdir,
                                 zscore="col", clip=5.0, show=True, close=False):
    outdir = _ensure_dir(outdir)

    levels, Mo = _group_means(X_obs, groups)
    _,      Mh = _group_means(X_hat, groups)
    Md = Mh - Mo

    if zscore == "col":
        Mo = _zscore_cols(Mo)
        Mh = _zscore_cols(Mh)
        Md = _zscore_cols(Md)

    if clip is not None:
        Mo = np.clip(Mo, -clip, clip)
        Mh = np.clip(Mh, -clip, clip)
        Md = np.clip(Md, -clip, clip)

    def _plot_one(M, ttl, path):
        fig, ax = plt.subplots(figsize=(0.22*len(feature_names)+3.5, 0.26*len(levels)+2.0))
        im = ax.imshow(M, aspect="auto", interpolation="nearest")
        ax.set_title(ttl)
        ax.set_xlabel("markers")
        ax.set_ylabel("celltype")
        ax.set_xticks(np.arange(len(feature_names)))
        ax.set_xticklabels(feature_names, rotation=90)
        ax.set_yticks(np.arange(len(levels)))
        ax.set_yticklabels(levels)
        plt.colorbar(im, ax=ax, fraction=0.03, pad=0.02)
        fig.tight_layout()
        fig.savefig(path, dpi=300, bbox_inches="tight")
        if show:
            plt.show()
        if close:
            plt.close(fig)

    _plot_one(Mo, f"{title_prefix} | observed (mean by {len(levels)} groups)",
              os.path.join(outdir, f"{title_prefix}__obs.png"))
    _plot_one(Mh, f"{title_prefix} | predicted (mean by {len(levels)} groups)",
              os.path.join(outdir, f"{title_prefix}__hat.png"))
    _plot_one(Md, f"{title_prefix} | delta (pred - obs)",
              os.path.join(outdir, f"{title_prefix}__delta.png"))

def plot_violin_by_celltype(adata, keys, groupby, outpath=None, show=True):
    sc.pl.violin(adata, keys=keys, groupby=groupby, rotation=90, show=show)
    fig = _scanpy_fig_from_plot()
    if outpath is not None:
        _ensure_dir(os.path.dirname(outpath))
        fig.savefig(outpath, dpi=300, bbox_inches="tight")
    if not show:
        plt.close(fig)

# -----------------------------
# 3) stats + fig3 builder (fixed)
# -----------------------------
def compute_and_save_all_stats(
    rna_test_pp, adt_test_pp,
    X_rna_obs, X_rna_hat,
    X_adt_obs, X_adt_hat,
    rna_markers, adt_markers,
    outdir_analysis
):
    _ensure_dir(outdir_analysis)

    rna_stats_all = pd.DataFrame({
        "feature": rna_test_pp.var_names.to_numpy(),
        "pearson_r":  per_feature_pearson(X_rna_obs, X_rna_hat),
        "spearman_r": per_feature_spearman(X_rna_obs, X_rna_hat),
        "rmse":       per_feature_rmse(X_rna_obs, X_rna_hat),
        "mae":        per_feature_mae(X_rna_obs, X_rna_hat),
    })
    adt_stats_all = pd.DataFrame({
        "feature": adt_test_pp.var_names.to_numpy(),
        "pearson_r":  per_feature_pearson(X_adt_obs, X_adt_hat),
        "spearman_r": per_feature_spearman(X_adt_obs, X_adt_hat),
        "rmse":       per_feature_rmse(X_adt_obs, X_adt_hat),
        "mae":        per_feature_mae(X_adt_obs, X_adt_hat),
    })

    rna_markers = [g for g in list(rna_markers) if g in rna_test_pp.var_names]
    adt_markers = [p for p in list(adt_markers) if p in adt_test_pp.var_names]

    rna_stats_markers = rna_stats_all[rna_stats_all["feature"].isin(rna_markers)].copy()
    adt_stats_markers = adt_stats_all[adt_stats_all["feature"].isin(adt_markers)].copy()

    rna_stats_all.to_csv(os.path.join(outdir_analysis, "rna_feature_stats_all.csv"), index=False)
    adt_stats_all.to_csv(os.path.join(outdir_analysis, "adt_feature_stats_all.csv"), index=False)
    rna_stats_markers.to_csv(os.path.join(outdir_analysis, "rna_feature_stats_markers.csv"), index=False)
    adt_stats_markers.to_csv(os.path.join(outdir_analysis, "adt_feature_stats_markers.csv"), index=False)

    # per-cell (all + markers)
    rna_marker_idx = np.array([rna_test_pp.var_names.get_loc(g) for g in rna_markers], dtype=int) if len(rna_markers) else None
    adt_marker_idx = np.array([adt_test_pp.var_names.get_loc(p) for p in adt_markers], dtype=int) if len(adt_markers) else None

    rna_mae_all, rna_rmse_all, rna_pr_all = per_cell_scores(X_rna_obs, X_rna_hat, feat_idx=None)
    adt_mae_all, adt_rmse_all, adt_pr_all = per_cell_scores(X_adt_obs, X_adt_hat, feat_idx=None)

    if rna_marker_idx is not None and rna_marker_idx.size:
        rna_mae_mk, rna_rmse_mk, rna_pr_mk = per_cell_scores(X_rna_obs, X_rna_hat, feat_idx=rna_marker_idx)
    else:
        rna_mae_mk = rna_rmse_mk = rna_pr_mk = np.full(rna_test_pp.n_obs, np.nan, dtype=np.float32)

    if adt_marker_idx is not None and adt_marker_idx.size:
        adt_mae_mk, adt_rmse_mk, adt_pr_mk = per_cell_scores(X_adt_obs, X_adt_hat, feat_idx=adt_marker_idx)
    else:
        adt_mae_mk = adt_rmse_mk = adt_pr_mk = np.full(rna_test_pp.n_obs, np.nan, dtype=np.float32)

    # attach to rna_test_pp (one basis)
    rna_test_pp.obs["RNA_xrec_mae_all"] = rna_mae_all
    rna_test_pp.obs["RNA_xrec_rmse_all"] = rna_rmse_all
    rna_test_pp.obs["RNA_xrec_pearson_all"] = rna_pr_all

    rna_test_pp.obs["ADT_xrec_mae_all"] = adt_mae_all
    rna_test_pp.obs["ADT_xrec_rmse_all"] = adt_rmse_all
    rna_test_pp.obs["ADT_xrec_pearson_all"] = adt_pr_all

    rna_test_pp.obs["RNA_xrec_mae_markers"] = rna_mae_mk
    rna_test_pp.obs["RNA_xrec_rmse_markers"] = rna_rmse_mk
    rna_test_pp.obs["RNA_xrec_pearson_markers"] = rna_pr_mk

    rna_test_pp.obs["ADT_xrec_mae_markers"] = adt_mae_mk
    rna_test_pp.obs["ADT_xrec_rmse_markers"] = adt_rmse_mk
    rna_test_pp.obs["ADT_xrec_pearson_markers"] = adt_pr_mk

    return rna_stats_all, adt_stats_all, rna_stats_markers, adt_stats_markers

def make_fig3_recon_panels(
    rna_test_pp,
    adt_test_pp,
    X_rna_obs, X_rna_hat,
    X_adt_obs, X_adt_hat,
    rna_markers, adt_markers,
    group_key="celltype.l2",
    umap_basis="umap",     # <-- pass "umap" for obsm["X_umap"], or "fig3_umap" for obsm["X_fig3_umap"]
    outdir="fig3_outputs",
    top=25, bottom=25,
    show=True,             # <-- controls notebook display
    show_first_n_marker_umaps=6,   # avoid 200 figures popping up
):
    outdir = _ensure_dir(outdir)
    outdir_tables = _ensure_dir(os.path.join(outdir, "tables"))
    outdir_bars   = _ensure_dir(os.path.join(outdir, "barplots"))
    outdir_umap   = _ensure_dir(os.path.join(outdir, "umap_markers"))
    outdir_heat   = _ensure_dir(os.path.join(outdir, "heatmaps"))
    outdir_diag   = _ensure_dir(os.path.join(outdir, "umap_diagnostics"))

    basis = _basis_name(umap_basis)

    # tables + per-cell diagnostics
    rna_stats_all, adt_stats_all, rna_stats_mk, adt_stats_mk = compute_and_save_all_stats(
        rna_test_pp, adt_test_pp,
        X_rna_obs, X_rna_hat,
        X_adt_obs, X_adt_hat,
        rna_markers, adt_markers,
        outdir_tables
    )

    # barplots
    plot_feature_perf(
        adt_stats_mk, title="ADT (RNA→ADT): marker proteins", metric_col="pearson_r",
        top=top, bottom=bottom,
        outpath=os.path.join(outdir_bars, "adt_markers_pearson_top_bottom.png"),
        xlim=(-1, 1),
        show=show, close=False
    )
    plot_feature_perf(
        rna_stats_mk, title="RNA (ADT→RNA): marker genes", metric_col="pearson_r",
        top=top, bottom=bottom,
        outpath=os.path.join(outdir_bars, "rna_markers_pearson_top_bottom.png"),
        xlim=(-1, 1),
        show=show, close=False
    )

    # diagnostics on the SAME embedding basis
    sc.pl.embedding(
        rna_test_pp,
        basis=basis,
        color=[
            "RNA_xrec_mae_markers", "ADT_xrec_mae_markers",
            "RNA_xrec_pearson_markers", "ADT_xrec_pearson_markers",
        ],
        ncols=2,
        size=12,
        frameon=False,
        wspace=0.35,
        show=show
    )
    fig = _scanpy_fig_from_plot()
    fig.savefig(os.path.join(outdir_diag, "per_cell_diagnostics_markers.png"), dpi=300, bbox_inches="tight")
    if not show:
        plt.close(fig)

    # marker UMAPs: add obs/hat values onto rna_test_pp.obs, then plot
    kept_rna = add_obs_hat_to_adata_obs(
        rna_test_pp, X_rna_obs, X_rna_hat,
        feat_names=list(rna_markers),
        feature_axis_names=rna_test_pp.var_names,
        prefix="RNA_"
    )
    plot_umap_obs_vs_hat(
        rna_test_pp, basis=basis,
        feats=kept_rna, prefix="RNA_",
        outdir=os.path.join(outdir_umap, "rna_markers"),
        ncols=2, size=12,
        show_first_n=(show_first_n_marker_umaps if show else 0),
        close=True
    )

    kept_adt = add_obs_hat_to_adata_obs(
        rna_test_pp, X_adt_obs, X_adt_hat,
        feat_names=list(adt_markers),
        feature_axis_names=adt_test_pp.var_names,
        prefix="ADT_"
    )
    plot_umap_obs_vs_hat(
        rna_test_pp, basis=basis,
        feats=kept_adt, prefix="ADT_",
        outdir=os.path.join(outdir_umap, "adt_markers"),
        ncols=2, size=12,
        show_first_n=(show_first_n_marker_umaps if show else 0),
        close=True
    )

    # mean-by-celltype heatmaps (markers only)
    groups = rna_test_pp.obs[group_key].astype(str).to_numpy()

    if kept_rna:
        rna_idx = [rna_test_pp.var_names.get_loc(g) for g in kept_rna]
        plot_obs_pred_delta_heatmaps(
            X_rna_obs[:, rna_idx],
            X_rna_hat[:, rna_idx],
            feature_names=kept_rna,
            groups=groups,
            title_prefix="RNA_ADTtoRNA_markers",
            outdir=os.path.join(outdir_heat, "rna_markers"),
            zscore="col",
            clip=5.0,
            show=show,
            close=False
        )

    if kept_adt:
        adt_idx = [adt_test_pp.var_names.get_loc(p) for p in kept_adt]
        plot_obs_pred_delta_heatmaps(
            X_adt_obs[:, adt_idx],
            X_adt_hat[:, adt_idx],
            feature_names=kept_adt,
            groups=groups,
            title_prefix="ADT_RNAtoADT_markers",
            outdir=os.path.join(outdir_heat, "adt_markers"),
            zscore="col",
            clip=5.0,
            show=show,
            close=False
        )

    print(f"[done] Fig3 outputs written to: {outdir}")
    return {
        "rna_stats_all": rna_stats_all,
        "adt_stats_all": adt_stats_all,
        "rna_stats_markers": rna_stats_mk,
        "adt_stats_markers": adt_stats_mk,
        "outdir": outdir
    }


# %% [code]
# RNA markers and ADT markers lists to use for expression UMAPs for Figure 3
# Canonical, ordered marker panels for PBMC CITE-seq
'''
CANON_RNA = [
    # T
    "CD3D", "TRAC", "IL7R",
    # Cytotoxic / NK
    "NKG7", "GNLY", "PRF1",
    # B / plasma
    "MS4A1", "CD79A", "MZB1", "JCHAIN",
    # Myeloid
    "LYZ", "FCGR3A",
]
'''

CANON_RNA = [
    # T
    #"CD3D", "TRAC",
    # Cytotoxic / NK
    #"NKG7", "GNLY",
    # B / plasma
    #"MS4A1", "CD79A", #"MZB1", "JCHAIN",
    # Myeloid
    #"LYZ", "FCGR3A",
    "CD8"
]

'''
CANON_ADT = [
    # T
    "CD3-1", "CD4-1", "CD8a",
    # naive/memory
    "CD45RA", "CD45RO",
    # B
    "CD19", "CD20",
    # myeloid / DC
    "CD14", "CD16", "CD11c",
    # APC / NK
    "HLA-DR", "CD56-1",
]
'''

def pick_first_present(options, varnames):
    for x in options:
        if x in varnames:
            return x
    return None

CD3 = pick_first_present(["CD3-1","CD3-2"], adt_test_pp.var_names)
CD4 = pick_first_present(["CD4-1","CD4-2"], adt_test_pp.var_names)
CD56 = pick_first_present(["CD56-1","CD56-2"], adt_test_pp.var_names)
'''
CANON_ADT = [x for x in [
    CD3, CD4, "CD8a", 
    "CD45RA", "CD45RO",
    "CD19", "CD20",
    "CD14", "CD16",
    "HLA-DR", CD56
] if x is not None]
'''
CANON_ADT = [x for x in [
    #CD3, CD4, 
    #"CD8a", "CD19", 
    #"CD19", "CD20", #"CD14", 
    #"CD16", CD56
    "GNLY"
] if x is not None]

# keep only those present (preserve order!)
canon_rna = [g for g in CANON_RNA if g in rna_test_pp.var_names]
canon_adt = [p for p in CANON_ADT if p in adt_test_pp.var_names]

print("RNA canonical panel:", canon_rna)
print("ADT canonical panel:", canon_adt)


# %% [code]
UMAP_BASIS = "fig3_umap"    # X_fig3_umap
GROUP_KEY  = "celltype.l2"

results = make_fig3_recon_panels(
    rna_test_pp=rna_test_pp,
    adt_test_pp=adt_test_pp,
    X_rna_obs=X_rna_obs,
    X_rna_hat=X_rna_hat,
    X_adt_obs=X_adt_obs,
    X_adt_hat=X_adt_hat,
    rna_markers=canon_rna,
    adt_markers=canon_adt,
    group_key=GROUP_KEY,
    umap_basis=UMAP_BASIS,
    outdir=os.path.join(outdir_analysis, "fig3"),
    top=25,
    bottom=0,
    show=True,                      # <-- shows inline
    show_first_n_marker_umaps=6     # <-- avoids 200 inline plots
)


# %% [code]
print(set(rna_test_pp.var_names))
print(set(adt_test_pp.var_names))


# %% [code]


# %% [code]


# %% [code]


# %% [code]


# %% [code]


# %% [code]


# %% [code]


# %% [code]


# %% [code]


# %% [code]
````
