## Supplemental Code for "Unifying multimodal single-cell data with a mixture-of-experts *β*-variational autoencoder framework": Fig04_Multiome_paired.ipynb

````
# %% [markdown]
# UniVI manuscript - Figure 4 generation reproducible workflow
### Multiome RNA + ATAC latent embedding by cell type and modality; examples of predicted accessibility programs from RNA; predicted gene programs from aTAC; alignment and reconstruction metrics

Andrew Ashford, Pathways + Omics Group, Oregon Health & Science University, Portland, OR - 12/7/2025

This Jupyter Notebook will house the end-to-end workflow to generate the panels in Figure 4 of our manuscript, "Unifying multimodal single-cell data with a mixture-of-experts β-variational autoencoder framework" which is currently being revised for Genome Research and is available currently on bioRxiv at the following link: https://www.biorxiv.org/content/10.1101/2025.02.28.640429v1.full

GitHub for the project - including a Quickstart guide - can be found at: https://github.com/Ashford-A/UniVI

Package is pip installable via the command: 
```bash
pip install univi
```


# %% [markdown]
### Import modules

# %% [code]
# Import non-UniVI modules
import os
import sys
from pathlib import Path

import numpy as np
import pandas as pd
import scanpy as sc
import anndata as ad
import torch

from torch.utils.data import DataLoader, Subset


# %% [code]
import scipy.sparse as sp
from sklearn.feature_extraction.text import TfidfTransformer
from sklearn.decomposition import TruncatedSVD
from sklearn.preprocessing import StandardScaler

from sklearn.neighbors import NearestNeighbors
from sklearn.metrics import confusion_matrix, accuracy_score, f1_score


# %% [code]
# Import required UniVI modules
from univi import (
    ModalityConfig,
    UniVIConfig,
    TrainingConfig,
    UniVIMultiModalVAE,
    matching,
    UniVITrainer,
    write_univi_latent,
    MultiModalDataset,
)

import univi as uv
import univi.evaluation as ue
import univi.plotting as up


# %% [code]
# Double check UniVI module version
print("Installed version is univi v" + str(uv.__version__))


# %% [markdown]
### Specify device to use for model

Set "device" - preferably device should be "cuda" for speedier model implementation/training. Requires GPU and the correct packages/versions.


# %% [code]
print("torch:", torch.__version__)
print("torch.cuda.is_available():", torch.cuda.is_available())
device = "cuda" if torch.cuda.is_available() else "cpu"
print("Using device:", device)


# %% [markdown]
### Specify file paths

Where data lives.

# %% [code]
DATA_ROOT = Path("/home/groups/precepts/ashforda/UniVI_v2/UniVI_older-non_git/data/PBMC_10x_Multiome_data/10x_Genomics_Multiome_data")

RNA_PATH = DATA_ROOT / "10x-Multiome-Pbmc10k-RNA.h5ad"
ATAC_PATH = DATA_ROOT / "10x-Multiome-Pbmc10k-ATAC.h5ad"

print("RNA file:", RNA_PATH)
print("ATAC file:", ATAC_PATH)


# %% [markdown]
### Read in data

Read data into AnnData objects using the paths in the code chunk above.

# %% [code]
rna = sc.read_h5ad(RNA_PATH)
atac = sc.read_h5ad(ATAC_PATH)

print(rna)
print(atac)

print("RNA obs names head:", rna.obs_names[:5].tolist())
print("ATAC obs names head:", atac.obs_names[:5].tolist())


# %% [markdown]
### Align cells between RNA and ATAC

Make sure the cell indices are aligned so UniVI knows which samples are paired.

# %% [code]
# Intersect barcodes
common_cells = rna.obs_names.intersection(atac.obs_names)
print("Common cells:", len(common_cells))

rna = rna[common_cells].copy()
atac = atac[common_cells].copy()

# Make sure order matches
atac = atac[rna.obs_names].copy()

assert np.array_equal(rna.obs_names.values, atac.obs_names.values)
print("obs_names aligned between RNA and ATAC.")


# %% [markdown]
### Stratify the data by celltype so that we train a balanced/generalizeable model

Also specifying all the data preprocessing functions below, will be used in their own respective sections.

# %% [code]
# -----------------------------
# Stratified split with cap
# -----------------------------
def stratified_split(
    idx,
    labels,
    frac_train=0.8,
    frac_val=0.1,
    seed=0,
    max_per_label=None,
    unused_to_test=True,
):
    """
    For each label:
      - shuffle its indices
      - keep up to max_per_label as "used"
      - split used -> train/val/test by fractions
      - leftover beyond max_per_label -> unused
    If unused_to_test=True, unused is appended into test and not returned separately.
    """
    rng = np.random.default_rng(seed)
    idx = np.asarray(idx)
    labels = np.asarray(labels)

    train_idx, val_idx, test_idx, unused_idx = [], [], [], []

    # stable label order (preserves category order if categorical, else sorted unique)
    uniq = pd.unique(labels)

    for lab in uniq:
        m = idx[labels == lab]
        rng.shuffle(m)

        if max_per_label is not None:
            used = m[:max_per_label]
            leftover = m[max_per_label:]
            if leftover.size:
                unused_idx.append(leftover)
        else:
            used = m

        n = used.size
        if n == 0:
            continue

        n_train = int(frac_train * n)
        n_val   = int(frac_val * n)
        # remainder -> test
        train_idx.append(used[:n_train])
        val_idx.append(used[n_train:n_train + n_val])
        test_idx.append(used[n_train + n_val:])

    train_idx = np.concatenate(train_idx) if train_idx else np.array([], dtype=int)
    val_idx   = np.concatenate(val_idx)   if val_idx   else np.array([], dtype=int)
    test_idx  = np.concatenate(test_idx)  if test_idx  else np.array([], dtype=int)
    unused_idx = np.concatenate(unused_idx) if unused_idx else np.array([], dtype=int)

    if unused_to_test and unused_idx.size:
        test_idx = np.concatenate([test_idx, unused_idx])
        unused_idx = np.array([], dtype=int)

    return train_idx, val_idx, test_idx, unused_idx


# %% [code]
# -----------------------------
# Small helper functions
# -----------------------------
def _ensure_counts_layer(adata, layer_name="counts"):
    if layer_name in adata.layers:
        return
    # Fall back to X as counts if needed (only if X is actually raw counts!)
    adata.layers[layer_name] = adata.X.copy()

def _subset_by_idx_pair(rna, adt, idx):
    # Assumes rna/adt are already paired in the same order
    return rna[idx].copy(), adt[idx].copy()

def preprocess_multiome_splits(
    rna_train, atac_train,
    rna_val,   atac_val,
    rna_test,  atac_test,
    *,
    rna_counts_layer="counts",
    atac_counts_layer="counts",
    n_hvg=2000,
    target_sum=1e4,
    n_lsi=50,
    seed=0,
):
    # --- ensure counts layers ---
    for a in (rna_train, rna_val, rna_test):
        if rna_counts_layer not in a.layers:
            a.layers[rna_counts_layer] = a.X.copy()
    for a in (atac_train, atac_val, atac_test):
        if atac_counts_layer not in a.layers:
            a.layers[atac_counts_layer] = a.X.copy()

    # =========================
    # RNA: HVG on TRAIN only
    # =========================
    rna_train_tmp = rna_train.copy()
    rna_train_tmp.X = rna_train_tmp.layers[rna_counts_layer]
    try:
        sc.pp.highly_variable_genes(rna_train_tmp, n_top_genes=int(n_hvg), flavor="seurat_v3")
    except Exception:
        sc.pp.highly_variable_genes(rna_train_tmp, n_top_genes=int(n_hvg), flavor="seurat")

    hvg = rna_train_tmp.var_names[rna_train_tmp.var["highly_variable"].to_numpy()].tolist()

    def _rna_transform(a):
        ad = a[:, hvg].copy()
        X = ad.layers[rna_counts_layer]
        sc.pp.normalize_total(ad, target_sum=float(target_sum), layer=rna_counts_layer)
        sc.pp.log1p(ad, layer=rna_counts_layer)
        ad.X = ad.layers[rna_counts_layer].copy()
        return ad

    rna_train_pp = _rna_transform(rna_train)
    rna_val_pp   = _rna_transform(rna_val)
    rna_test_pp  = _rna_transform(rna_test)

    # =========================
    # ATAC: TFIDF + SVD (LSI) fit on TRAIN only
    # =========================
    Xtr = atac_train.layers[atac_counts_layer]
    Xva = atac_val.layers[atac_counts_layer]
    Xte = atac_test.layers[atac_counts_layer]

    if not sp.issparse(Xtr):
        Xtr = sp.csr_matrix(Xtr)
    if not sp.issparse(Xva):
        Xva = sp.csr_matrix(Xva)
    if not sp.issparse(Xte):
        Xte = sp.csr_matrix(Xte)

    tfidf = TfidfTransformer(norm="l2", use_idf=True, smooth_idf=True, sublinear_tf=False)
    Xtr_t = tfidf.fit_transform(Xtr)
    Xva_t = tfidf.transform(Xva)
    Xte_t = tfidf.transform(Xte)

    svd = TruncatedSVD(n_components=int(n_lsi), random_state=int(seed))
    Ztr = svd.fit_transform(Xtr_t)
    Zva = svd.transform(Xva_t)
    Zte = svd.transform(Xte_t)

    # scale using TRAIN only (good for Gaussian decoder stability)
    scaler = StandardScaler(with_mean=True, with_std=True)
    Ztr = scaler.fit_transform(Ztr).astype(np.float32, copy=False)
    Zva = scaler.transform(Zva).astype(np.float32, copy=False)
    Zte = scaler.transform(Zte).astype(np.float32, copy=False)

    # Build ATAC-LSI AnnDatas (keeps pairing/obs; var are "LSI_0..")
    atac_train_lsi = ad.AnnData(X=Ztr, obs=atac_train.obs.copy(),
                               var=pd.DataFrame(index=[f"LSI_{i}" for i in range(Ztr.shape[1])]))
    atac_val_lsi   = ad.AnnData(X=Zva, obs=atac_val.obs.copy(),
                               var=pd.DataFrame(index=[f"LSI_{i}" for i in range(Zva.shape[1])]))
    atac_test_lsi  = ad.AnnData(X=Zte, obs=atac_test.obs.copy(),
                               var=pd.DataFrame(index=[f"LSI_{i}" for i in range(Zte.shape[1])]))

    # keep raw peaks around for biology (optional; you already have atac_* originals)
    return (rna_train_pp, atac_train_lsi,
            rna_val_pp,   atac_val_lsi,
            rna_test_pp,  atac_test_lsi,
            hvg, tfidf, svd, scaler)

import numpy as np
import pandas as pd
import anndata as ad
import scanpy as sc
import scipy.sparse as sp

def transform_rna_with_hvg(rna, *, hvg, counts_layer="counts", target_sum=1e4):
    a = rna[:, hvg].copy()

    # make sure counts exist
    if counts_layer not in a.layers:
        a.layers[counts_layer] = a.X.copy()

    # IMPORTANT: don't overwrite your raw counts layer; work in a fresh layer
    a.layers["log1p"] = a.layers[counts_layer].copy()
    sc.pp.normalize_total(a, target_sum=float(target_sum), layer="log1p")
    sc.pp.log1p(a, layer="log1p")
    a.X = a.layers["log1p"]
    return a

def transform_atac_with_lsi(atac, *, counts_layer="counts", tfidf=None, svd=None, scaler=None):
    if tfidf is None or svd is None or scaler is None:
        raise ValueError("Need tfidf, svd, scaler objects from preprocess_multiome_splits().")

    a = atac.copy()
    if counts_layer not in a.layers:
        a.layers[counts_layer] = a.X.copy()

    X = a.layers[counts_layer]
    if not sp.issparse(X):
        X = sp.csr_matrix(X)

    Xt = tfidf.transform(X)
    Z  = svd.transform(Xt)
    Z  = scaler.transform(Z).astype(np.float32, copy=False)

    atac_lsi = ad.AnnData(
        X=Z,
        obs=a.obs.copy(),
        var=pd.DataFrame(index=[f"LSI_{i}" for i in range(Z.shape[1])]),
    )
    return atac_lsi


# %% [code]
# Check the counts of each celltype.l1 in the data
print(rna.obs["cell_type"].value_counts())


# %% [code]
# --------------------------------------------------
# Use it on 10k PBMC 10x Genomic Multiome RNA + ATAC
# --------------------------------------------------
labels = rna.obs["cell_type"].astype(str).to_numpy()
idx = np.arange(rna.n_obs)

train_idx, val_idx, test_idx, unused_idx = stratified_split(
    idx, labels,
    frac_train=0.8, frac_val=0.1, seed=0,
    max_per_label=5000,
    unused_to_test=True,      # set True if you want unused merged into test
)

# paired splits
rna_train, atac_train = _subset_by_idx_pair(rna, atac, train_idx)
rna_val,   atac_val   = _subset_by_idx_pair(rna, atac, val_idx)
rna_test,  atac_test  = _subset_by_idx_pair(rna, atac, test_idx)

if unused_idx.size:
    rna_unused, atac_unused = _subset_by_idx_pair(rna, atac, unused_idx)
else:
    rna_unused, atac_unused = None, None


# %% [code]
print(rna_train.obs['cell_type'].value_counts())


# %% [code]
def upsample_train_to_target(train_idx, labels, target_per_label=1000, seed=0, shuffle=True):
    rng = np.random.default_rng(seed)
    train_idx = np.asarray(train_idx, dtype=int)
    labels = np.asarray(labels)

    y = labels[train_idx]
    out = []

    # stable-ish label order
    for lab in pd.unique(y):
        m = train_idx[y == lab]
        if m.size == 0:
            continue

        replace = (m.size < int(target_per_label))
        out.append(rng.choice(m, size=int(target_per_label), replace=replace))

    out = np.concatenate(out).astype(int, copy=False)

    if shuffle:
        rng.shuffle(out)
    return out

# ---- usage (paired adatas) ----
# make sure rna/atac aligned first
'''
assert (rna.obs_names == atac.obs_names).all()

labels = rna.obs["cell_type"].astype(str).to_numpy()

train_idx_bal = upsample_train_to_target(
    train_idx,
    labels,
    target_per_label=1500,
    seed=0,
    shuffle=True,
)

rna_train_bal  = rna[train_idx_bal].copy()
atac_train_bal = atac[train_idx_bal].copy()
'''

# %% [markdown]
### Data preprocessing

* RNA preprocessing (log1p + HVG + scale → Gaussian decoder)

* ATAC preprocessing (TF-IDF + LSI + scale → Gaussian decoder)

Running the preprocessing functions specified above. Of note, we're performing the preprocessing per-train/val/test split.

# %% [code]
import scipy.sparse as sp

# preprocessing (fit on train, apply to val/test)
(rna_train_pp, atac_train_lsi,
 rna_val_pp,   atac_val_lsi,
 rna_test_pp,  atac_test_lsi,
 hvg_genes, tfidf, svd, scaler) = preprocess_multiome_splits(
    rna_train, atac_train,
    rna_val,   atac_val,
    rna_test,  atac_test,
    n_hvg=2000, n_lsi=100, seed=42
)


# %% [code]
'''
# 1) Fit preprocessing on UNIQUE train (your existing call)
(rna_train_pp_u, atac_train_lsi_u,
 rna_val_pp,     atac_val_lsi,
 rna_test_pp,    atac_test_lsi,
 hvg, tfidf, svd, scaler) = preprocess_multiome_splits(
    rna_train, atac_train,
    rna_val,   atac_val,
    rna_test,  atac_test,
    n_hvg=5000, n_lsi=200, seed=42
)

# 2) Upsample indices INSIDE the train split
train_idx_u = np.arange(rna_train.n_obs)
train_idx_bal = upsample_train_to_target(
    train_idx_u,
    rna_train.obs["cell_type"].to_numpy(),
    target_per_label=1500,
    seed=0,
)

# 3) Build upsampled paired train adatas
rna_train_bal  = rna_train[train_idx_bal].copy()
atac_train_bal = atac_train[train_idx_bal].copy()

# (optional) if duplicates annoy scanpy later:
# rna_train_bal.obs_names_make_unique()
# atac_train_bal.obs_names_make_unique()

# 4) Transform upsampled train using the FIT objects
rna_train_pp  = transform_rna_with_hvg(rna_train_bal, hvg=hvg, counts_layer="counts", target_sum=1e4)
atac_train_lsi = transform_atac_with_lsi(atac_train_bal, counts_layer="counts", tfidf=tfidf, svd=svd, scaler=scaler)
'''

# %% [code]
# Sanity check above preprocessed data objects - start with RNA
print(rna_train_pp)
print(rna_val_pp)
print(rna_test_pp)


# %% [code]
# Now sanity check ATAC LSI data objects
print(atac_train_lsi)
print(atac_val_lsi)
print(atac_test_lsi)


# %% [markdown]
### Wrap into MultiModalDataset & DataLoaders

# %% [code]
from univi.data import align_paired_obs_names

pin_memory = (device == "cuda")

# Make sure each split is paired and ordered the same across modalities
# This was erroring out due to a function bug, fixed it and should be fixed by manuscript publication
#train_dict = align_paired_obs_names({"rna": rna_train_pp, "adt": atac_train_lsi})
#val_dict   = align_paired_obs_names({"rna": rna_val_pp,   "adt": atac_val_lsi})
#test_dict  = align_paired_obs_names({"rna": rna_test_pp,  "adt": atac_test_lsi})

# Using this instead for now since we know the data are already paired from above code
assert (rna_train_pp.obs_names == atac_train_lsi.obs_names).all()
assert (rna_val_pp.obs_names   == atac_val_lsi.obs_names).all()
assert (rna_test_pp.obs_names  == atac_test_lsi.obs_names).all()

train_dict = {"rna": rna_train_pp, "atac": atac_train_lsi}
val_dict   = {"rna": rna_val_pp,   "atac": atac_val_lsi}
test_dict  = {"rna": rna_test_pp,  "atac": atac_test_lsi}

# Build datasets (CPU tensors; trainer/model moves to GPU)
train_ds = MultiModalDataset(adata_dict=train_dict, X_key="X", device=None)
val_ds   = MultiModalDataset(adata_dict=val_dict,   X_key="X", device=None)
test_ds  = MultiModalDataset(adata_dict=test_dict,  X_key="X", device=None)

batch_size = 128
num_workers = 0


train_loader = DataLoader(
    train_ds,
    batch_size=batch_size,
    shuffle=True,
    num_workers=num_workers,
    pin_memory=pin_memory,
    drop_last=False,
)

'''
from torch.utils.data import DataLoader, WeightedRandomSampler

# labels for the TRAIN split (same order as train_ds cells)
y = rna_train_pp.obs["cell_type"].astype(str).to_numpy()

# inverse frequency weights
vals, counts = np.unique(y, return_counts=True)
freq = dict(zip(vals, counts))
w = np.array([1.0 / freq[c] for c in y], dtype=np.float64)
w = w / w.sum()

sampler = WeightedRandomSampler(
    weights=torch.as_tensor(w, dtype=torch.double),
    num_samples=len(w),   # one "epoch" worth of draws
    replacement=True
)

train_loader = DataLoader(
    train_ds,
    batch_size=batch_size,
    sampler=sampler,      # <-- instead of shuffle=True
    shuffle=False,
    num_workers=num_workers,
    pin_memory=pin_memory,
    drop_last=False,
)
'''

val_loader = DataLoader(
    val_ds,
    batch_size=batch_size,
    shuffle=False,
    num_workers=num_workers,
    pin_memory=pin_memory,
    drop_last=False,
)

test_loader = DataLoader(
    test_ds,
    batch_size=batch_size,
    shuffle=False,
    num_workers=num_workers,
    pin_memory=pin_memory,
    drop_last=False,
)

print("n_train / n_val / n_test:", train_ds.n_cells, val_ds.n_cells, test_ds.n_cells)
print("batches:", len(train_loader), len(val_loader), len(test_loader))

# sanity check one batch
x = next(iter(train_loader))
print({k: v.shape for k, v in x.items()})


# %% [markdown]
### UniVI configs

# %% [code]
y_codes = rna.obs["cell_type"].astype("category").cat.codes.to_numpy()
n_classes = int(y_codes.max() + 1)


# %% [code]
univi_cfg = UniVIConfig(
    latent_dim=30,
    beta=1.25,
    gamma=4.35,
    encoder_dropout=0.1,
    decoder_dropout=0.05,
    encoder_batchnorm=True,
    decoder_batchnorm=False,
    kl_anneal_start=50,
    kl_anneal_end=85,
    align_anneal_start=75,
    align_anneal_end=110,
    modalities=[
        ModalityConfig(
            name="rna",
            input_dim=rna_train_pp.n_vars,
            encoder_hidden=[512, 256, 128],
            decoder_hidden=[128, 256, 512],
            likelihood="gaussian",
        ),
        ModalityConfig(
            name="atac",
            input_dim=atac_train_lsi.n_vars,
            encoder_hidden=[128, 64],
            decoder_hidden=[64, 128],
            likelihood="gaussian",
        ),
    ],
)


# %% [markdown]
If you want raw-count NB/ZINB instead, you’d:
* Put raw counts in .layers["counts"]
* Set X_key="counts" in the dataset
* Use likelihood="nb" or "zinb" above

But for this "Figure 4" notebook, the scaled Gaussian setup is nicely stable and good for the best-integrated latent space.


# %% [markdown]
### Instantiate model and model objective

# %% [code]

model = UniVIMultiModalVAE(
    univi_cfg,
    loss_mode="v1",      # cross-recon + cross-posterior alignment
    #loss_mode="lite"
    #v1_recon="cross",   # full k→j cross-recon
    v1_recon="avg",
    #v1_recon_mix=0.5,
    normalize_v1_terms=True,
).to(device)

'''
model = UniVIMultiModalVAE(
    univi_cfg,
    loss_mode="v1",      # cross-recon + cross-posterior alignment
    #loss_mode="lite"
    #v1_recon="cross",   # full k→j cross-recon
    v1_recon="avg",
    #v1_recon_mix=0.5,
    normalize_v1_terms=True,
    n_label_classes=n_classes,
    label_loss_weight=5.0,
    label_ignore_index=-1,
    classify_from_mu=True,
).to(device)
'''
'''
model = UniVIMultiModalVAE(
    univi_cfg,
    loss_mode="lite",

    # Optional: keep the decoder-side classification head too
    n_label_classes=n_classes,
    label_loss_weight=2.0,

    # Encoder-side label expert injected into fusion
    use_label_encoder=True,
    label_moe_weight=3.5,      # >1 => labels influence fusion more
    unlabeled_logvar=20.0,     # very high => tiny precision => ignored in fusion
    label_encoder_warmup=5,    # wait N epochs before injecting labels into fusion
    label_ignore_index=-1,
).to("cuda")
'''

# %% [markdown]
### Instantiate TrainingConfig & trainer

# %% [code]
train_cfg = TrainingConfig(
    n_epochs=5000,
    batch_size=batch_size,
    lr=1e-3,
    weight_decay=1e-4,
    device=device,
    log_every=100,
    grad_clip=5.0,
    num_workers=0,
    seed=42,
    early_stopping=True,
    patience=300,         # Setting kind of high patience since training takes ~1s/iteration because of small-ish dataset
    min_delta=0.0,
)

print(train_cfg)
print(univi_cfg)


# %% [code]
trainer = UniVITrainer(
    model=model,
    train_loader=train_loader,
    val_loader=val_loader,
    train_cfg=train_cfg,
    device=device,
)

print(trainer)


# %% [markdown]
### Fit the model

# %% [code]
history = trainer.fit()

# history is a dict with keys like "train_loss", "val_loss", "beta", "gamma"
print("Training finished.")
print("Best val loss:", np.min(history["val_loss"]))


# %% [code]
import matplotlib.pyplot as plt

plt.figure()
plt.plot(history["train_loss"], label="train")
plt.plot(history["val_loss"], label="val")
plt.xlabel("Epoch")
plt.ylabel("Loss")
plt.legend()
plt.title("UniVI training (Multiome)")
plt.show()


# %% [markdown]
### Write latent z back to AnnData

# %% [code]
print("Test batches:", len(test_loader))


# %% [code]
# Make sure adata_dict has the *same ordering* as during training
adata_dict = {
    "rna": rna_test_pp,
    "atac": atac_test_lsi,
}

Z = write_univi_latent(
    model,
    adata_dict,
    obsm_key="X_univi",   # will be added to each AnnData in adata_dict
    batch_size=512,
    device=device,
    use_mean=True,       # deterministic: use encoder means instead of noisy samples
    #use_mean=False,       # Stochastic
)

print("UniVI latent shape:",  Z.shape)
print("rna.obsm keys:",       rna_test_pp.obsm.keys())
print("adt.obsm keys:",       atac_test_lsi.obsm.keys())


# %% [markdown]
Now both rna and atac have a shared latent:

* rna_test_pp.obsm["X_univi"]

* atac_test_pp.obsm["X_univi"]


# %% [markdown]
### Code to save/load modal if so inclined

# %% [code]
beta_used = "1.25"
gamma_used = "4.35"
latent_dims_used = "30"


# %% [code]
output_dir = f'./results/univi_TEA-seq_Figure_4_Multiome_beta-{beta_used}_gamma-{gamma_used}_latent_dims-{latent_dims_used}_gaussian_all_reproducibility/'
out_file = f"trained_model_beta-{beta_used}_gamma-{gamma_used}_latent_dims-{latent_dims_used}_gaussian_both_reproducibility.pt"


# %% [code]
from dataclasses import asdict

os.makedirs(output_dir, exist_ok=True)


ckpt_path = output_dir + out_file


rna = rna_test_pp.copy()
atac = atac_test_lsi.copy()


# %% [code]
n_hvg = len(rna.var_names)
target_sum = 1e4
n_lsi = len(atac.var_names)
rna_hvg_names = rna.var_names


# after training
#history = trainer.fit()

import sys, platform, hashlib, numpy as np, torch

def _hash_list(x):
    h = hashlib.sha256()
    for s in x:
        h.update(str(s).encode())
        h.update(b"\n")
    return h.hexdigest()

ckpt = {
    # core
    "state_dict": trainer.model.state_dict(),
    "univi_cfg": asdict(univi_cfg),

    # best model info
    "best_epoch": trainer.best_epoch,
    "best_val_loss": float(trainer.best_val_loss),

    # splits (STORE THESE!)
    "splits": {
        "train": np.asarray(train_idx, dtype=np.int64),
        "val":   np.asarray(val_idx, dtype=np.int64),
        "test":  np.asarray(test_idx, dtype=np.int64),
    },

    # preprocessing + feature sets (examples; include what you actually use)
    "preproc": {
        "rna":  {"layer": "counts", "n_hvg": n_hvg, "target_sum": target_sum, "log1p": True, "zscore": True},
        "adt":  {"layer": "counts", "clr": True, "zscore": True},
        "atac": {"layer": "counts", "tfidf": True, "n_lsi": n_lsi, "lsi_drop_first": True, "zscore": True},
    },
    "features": {
        "rna_hvgs": list(rna_hvg_names),          # list[str]
        "atac_peaks": list(atac.var_names),       # list[str] (or whatever LSI used)
    },

    # provenance checks
    "data_fingerprint": {
        "rna_obs_hash": _hash_list(rna.obs_names),
        "rna_var_hash": _hash_list(rna.var_names),
        "atac_obs_hash": _hash_list(atac.obs_names),
        "atac_var_hash": _hash_list(atac.var_names),
    },

    # environment versions
    "versions": {
        "python": sys.version,
        "platform": platform.platform(),
        "torch": torch.__version__,
        "numpy": np.__version__,
        "univi": uv.__version__,
        "anndata": ad.__version__,
        "scanpy": sc.__version__,
    },

    # history (optional)
    "history": history,
}

torch.save(ckpt, ckpt_path)
print("Saved best model to:", ckpt_path)


# %% [code]
# Later to reload model
import torch
from univi.config import UniVIConfig, ModalityConfig
from univi.models.univi import UniVIMultiModalVAE

print("torch:", torch.__version__)
print("torch.cuda.is_available():", torch.cuda.is_available())
device = "cuda" if torch.cuda.is_available() else "cpu"
print("Using device:", device)

#device = "cuda"  # or "cuda" if available


ckpt = torch.load(
    #output_dir + out_file,
    ckpt_path,
    map_location=device,
    weights_only=False,
)


# ---- Rebuild UniVIConfig, making sure modalities are ModalityConfig objects ----
cfg_dict = ckpt["univi_cfg"]

# If this is an OmegaConf object or similar, make sure it's a plain dict
try:
    from omegaconf import DictConfig, OmegaConf
    if isinstance(cfg_dict, DictConfig):
        cfg_dict = OmegaConf.to_container(cfg_dict, resolve=True)
except ImportError:
    pass


# Now rehydrate each modality
modalities = [ModalityConfig(**m) for m in cfg_dict["modalities"]]
cfg_dict = {**cfg_dict, "modalities": modalities}

univi_cfg_loaded = UniVIConfig(**cfg_dict)

# ---- Rebuild model + load weights ----
model_loaded = UniVIMultiModalVAE(univi_cfg_loaded).to(device)
model_loaded.load_state_dict(ckpt["state_dict"])

print("Best epoch was:", ckpt.get("best_epoch"), "val loss =", ckpt.get("best_val_loss"))


print(sorted(ckpt.keys()))
#print(ckpt['splits'])


# %% [markdown]
### Figure 4A–D: UMAP (modality + celltype levels)

# %% [code]
from univi.evaluation import encode_adata

Z_rna = encode_adata(model, rna_test_pp, modality="rna",
                     latent="modality_mean", device=device, batch_size=1024)
Z_atac = encode_adata(model, atac_test_lsi, modality="atac",
                     latent="modality_mean", device=device, batch_size=1024)

key_rna = "encode_adata(modality_mean)"
key_atac = "encode_adata(modality_mean)"

rna_test_pp.obsm["X_univi_rna"] = Z_rna
atac_test_lsi.obsm["X_univi_atac"] = Z_atac


# %% [code]
rna_u = rna_test_pp.copy()
atac_u = atac_test_lsi.copy()

rna_u.obs["modality"] = "rna"
atac_u.obs["modality"] = "atac"

combo = ad.concat([rna_u, atac_u], join="outer", label="modality", keys=["rna","atac"], index_unique="-")

# IMPORTANT: put the correct latent for each half
combo.obsm["X_univi_sep"] = np.vstack([Z_rna, Z_atac]).astype(np.float32)


# %% [code]
# neighbors/umap on the *separate* stacked latent
sc.pp.neighbors(combo, use_rep="X_univi_sep", n_neighbors=30)
sc.tl.umap(combo)


# %% [code]
'''
# Scanpy defaults (affects sc.pl.*)
sc.set_figure_params(
    figsize=(14, 12),   # bigger canvas
    dpi=200,            # on-screen sharpness
    dpi_save=600,       # saved file sharpness
    fontsize=14,
    frameon=False,
)

# Matplotlib defaults (affects plt.*)
plt.rcParams.update({
    "figure.figsize": (14, 12),
    "figure.dpi": 200,
    "savefig.dpi": 600,
    "savefig.bbox": "tight",
    "savefig.pad_inches": 0.1,
    "axes.titlesize": 16,
    "axes.labelsize": 14,
    "legend.fontsize": 12,
})
'''

# %% [code]
celltype_key = 'cell_type'


# %% [code]
# plot UMAPs
#sc.pl.umap(combo, color=["modality"], frameon=False, size=25.0)
#sc.pl.umap(combo, color=[celltype_key], frameon=False, size=25.0)


# %% [code]
# Check out many of each cell type are in the test set for level 2 cell annotations
print(rna_test_pp.obs["cell_type"].value_counts())


# %% [markdown]
### Figure 4E–F: alignment metrics (FOSCTTM + label transfer + mixing)

# %% [code]
from sklearn.neighbors import NearestNeighbors
from sklearn.metrics import precision_recall_fscore_support

# If plots ever don't show in classic notebook, run once:
# %matplotlib inline

# -------------------------
# show-or-save helper
# -------------------------
def _finish(fig=None, outpath=None, dpi=100, show=True, close=True):
    if outpath is not None:
        plt.savefig(outpath, dpi=dpi, bbox_inches="tight")
    if show:
        plt.show()
    if close:
        plt.close(fig if fig is not None else plt.gcf())

# -------------------------
# plots
# -------------------------
def plot_metrics_bar(
    metrics,
    keys,
    title="Figure 4 summary metrics",
    outpath=None,
    err_keys=None,          # dict: metric_key -> error_metric_key
    default_err=np.nan,     # np.nan = no error bar drawn; 0.0 = zero-length
    capsize=4,
):
    vals = np.array([float(metrics[k]) for k in keys], dtype=float)

    if err_keys is None:
        yerr = np.full_like(vals, default_err, dtype=float)
    else:
        yerr = np.array(
            [float(metrics[err_keys[k]]) if k in err_keys and err_keys[k] in metrics else default_err
             for k in keys],
            dtype=float
        )

    fig = plt.figure(figsize=(10, 6))
    plt.bar(keys, vals, yerr=yerr, capsize=capsize)
    plt.xticks(rotation=45, ha="right")
    plt.title(title)
    plt.tight_layout()

    # inline “finish” behavior
    if outpath is not None:
        plt.savefig(outpath, dpi=300, bbox_inches="tight")
    plt.show()
    plt.close(fig)


def plot_confusion(cm, classes, title, normalize="true", outpath=None, cmap="viridis"):
    import numpy as np
    import matplotlib.pyplot as plt

    cm = np.asarray(cm, dtype=float)
    if normalize == "true":
        cm = cm / (cm.sum(axis=1, keepdims=True) + 1e-12)
        subtitle = "Row-normalized"
    elif normalize == "pred":
        cm = cm / (cm.sum(axis=0, keepdims=True) + 1e-12)
        subtitle = "Col-normalized"
    elif normalize == "all":
        cm = cm / (cm.sum() + 1e-12)
        subtitle = "Global-normalized"
    else:
        subtitle = "Counts"

    classes = np.asarray(classes, dtype=str)
    n = len(classes)

    fig, ax = plt.subplots(figsize=(14, 13))
    im = ax.imshow(cm, interpolation="nearest", cmap=cmap, aspect="auto")

    # kill any gridlines (including ones from styles)
    ax.grid(False)
    ax.minorticks_off()

    ax.set_title(f"{title}\n({subtitle})")
    ax.set_xlabel("Predicted")
    ax.set_ylabel("True")

    ax.set_xticks(np.arange(n))
    ax.set_yticks(np.arange(n))
    ax.set_xticklabels(classes, rotation=90)
    ax.set_yticklabels(classes)

    cbar = fig.colorbar(im, ax=ax)
    cbar.set_label("value")

    fig.tight_layout()

    if outpath is not None:
        fig.savefig(outpath, dpi=300, bbox_inches="tight")

    plt.show()
    plt.close(fig)


def plot_per_class_f1(y_true, y_pred, title="Per-class F1", outpath=None, top_n=None):
    y_true = np.asarray(y_true).astype(str)
    y_pred = np.asarray(y_pred).astype(str)
    classes = np.unique(np.concatenate([y_true, y_pred]))

    _, _, f1, support = precision_recall_fscore_support(
        y_true, y_pred, labels=classes, zero_division=0
    )

    df = pd.DataFrame({"class": classes, "f1": f1, "support": support})
    df = df.sort_values(["f1", "support"], ascending=[True, False])

    if top_n is not None:
        df = df.head(int(top_n))

    fig = plt.figure(figsize=(10, 0.35 * len(df) + 2.0))
    plt.barh(df["class"], df["f1"])
    plt.xlabel("F1")
    plt.title(title)
    plt.xlim(0, 1)
    plt.tight_layout()
    _finish(fig, outpath=outpath, show=True, close=True)
    return df

def plot_modality_mixing_hist(Z, mods, k=20, metric="euclidean", title="Modality mixing", outpath=None):
    Z = np.asarray(Z, dtype=np.float32)
    mods = np.asarray(mods)

    n = Z.shape[0]
    k_eff = int(min(max(int(k), 1), n - 1))

    nn = NearestNeighbors(n_neighbors=k_eff + 1, metric=metric)
    nn.fit(Z)
    nbrs = nn.kneighbors(Z, return_distance=False)[:, 1:]
    frac_other = (mods[nbrs] != mods[:, None]).mean(axis=1)

    fig = plt.figure(figsize=(7, 4.5))
    plt.hist(frac_other, bins=60)
    plt.xlabel("Fraction of kNN from other modality")
    plt.ylabel("# cells")
    plt.title(title)
    plt.tight_layout()
    _finish(fig, outpath=outpath, show=True, close=True)
    return frac_other

def plot_foscttm_sanity(Z1, Z2, idx, title="FOSCTTM sanity", outpath=None):
    Z1s = np.asarray(Z1[idx], dtype=np.float32)
    Z2s = np.asarray(Z2[idx], dtype=np.float32)

    d_true = np.sum((Z1s - Z2s) ** 2, axis=1)

    nn = NearestNeighbors(n_neighbors=51, metric="euclidean")
    nn.fit(np.asarray(Z2, dtype=np.float32))
    dist, _ = nn.kneighbors(Z1s)
    d_min = dist[:, 0] ** 2

    fig = plt.figure(figsize=(5.5, 5.5))
    plt.scatter(d_true, d_min, s=8, alpha=0.4)
    mx = np.percentile(np.concatenate([d_true, d_min]), 99)
    plt.plot([0, mx], [0, mx], linewidth=1)
    plt.xlim(0, mx); plt.ylim(0, mx)
    plt.xlabel("d(true match)^2")
    plt.ylabel("d(nearest neighbor)^2")
    plt.title(title)
    plt.tight_layout()
    _finish(fig, outpath=outpath, show=True, close=True)

def plot_paired_distance_hist(Z_rna, Z_adt, idx, title="Paired latent distance (subset)", outpath=None):
    d_pair = np.sqrt(np.sum((Z_rna[idx] - Z_adt[idx]) ** 2, axis=1))
    fig = plt.figure(figsize=(7, 4.5))
    plt.hist(d_pair, bins=80)
    plt.xlabel("||z_rna - z_adt||")
    plt.ylabel("# cells")
    plt.title(title)
    plt.tight_layout()
    _finish(fig, outpath=outpath, show=True, close=True)


# %% [code]
from sklearn.neighbors import NearestNeighbors
from sklearn.metrics import confusion_matrix, accuracy_score, f1_score

# ----------------------------
# Metrics helpers
# ----------------------------
def foscttm_chunked(Z1, Z2, block=512):
    """
    Exact FOSCTTM computed in blocks to avoid NxN memory blowups.
    Assumes 1:1 pairing between rows i in Z1 and Z2.
    """
    Z1 = np.asarray(Z1, dtype=np.float32)
    Z2 = np.asarray(Z2, dtype=np.float32)
    assert Z1.shape == Z2.shape
    n = Z1.shape[0]

    Z2_T = Z2.T
    n2 = np.sum(Z2 * Z2, axis=1)  # (n,)

    fos = np.empty(n, dtype=np.float32)

    for i0 in range(0, n, block):
        i1 = min(i0 + block, n)
        A = Z1[i0:i1]  # (b, d)
        n1 = np.sum(A * A, axis=1)[:, None]  # (b,1)

        d2 = n1 + n2[None, :] - 2.0 * (A @ Z2_T)  # (b,n)

        true = d2[np.arange(i1 - i0), np.arange(i0, i1)]
        fos[i0:i1] = (d2 < true[:, None]).sum(axis=1) / (n - 1)

    return float(fos.mean()), float(fos.std(ddof=1) / np.sqrt(n))


def modality_mixing_score(Z, modality_labels, k=20, metric="euclidean"):
    """
    Vanilla modality mixing:
    Mean fraction of kNN neighbors that differ in modality.
    Use this for *non-duplicated* sets (e.g., concatenated modality-specific embeddings).
    """
    Z = np.asarray(Z, dtype=np.float32)
    modality_labels = np.asarray(modality_labels)

    n = Z.shape[0]
    if n <= 1:
        return 0.0

    k_eff = int(min(max(int(k), 1), n - 1))
    nn = NearestNeighbors(n_neighbors=k_eff + 1, metric=metric)
    nn.fit(Z)
    nbrs = nn.kneighbors(Z, return_distance=False)[:, 1:]  # drop self

    same = (modality_labels[nbrs] == modality_labels[:, None])
    return float((~same).mean())


def modality_mixing_score_excluding_pairs(Z, modality_labels, cell_ids, k=20, metric="euclidean"):
    """
    Pair-aware modality mixing for 'combo' style stacked data (same cell appears twice: RNA + ADT),
    where the fused embedding may be identical (or extremely close) for the paired duplicates.

    Computes: for each row, the fraction of neighbors from the other modality,
    AFTER removing the paired duplicate (same cell_id) from its neighbor list.

    cell_ids must map each row to a shared cell identifier (same for RNA+ADT copy).
    """
    Z = np.asarray(Z, dtype=np.float32)
    modality_labels = np.asarray(modality_labels)
    cell_ids = np.asarray(cell_ids).astype(str)

    n = Z.shape[0]
    if n <= 2:
        return 0.0

    # Need enough neighbors so we can drop self + paired duplicate and still have k
    k_eff = int(min(max(int(k), 1), n - 2))
    nn = NearestNeighbors(n_neighbors=k_eff + 2, metric=metric)
    nn.fit(Z)
    nbrs = nn.kneighbors(Z, return_distance=False)[:, 1:]  # drop self

    # Build "pair index": for each row i, pair[i] is the index of the other modality copy
    first = {}
    pair = np.full(n, -1, dtype=np.int64)
    for i, cid in enumerate(cell_ids):
        if cid in first:
            j = first[cid]
            pair[i] = j
            pair[j] = i
        else:
            first[cid] = i

    frac_other = np.empty(n, dtype=np.float32)
    for i in range(n):
        neigh = nbrs[i]

        # remove paired duplicate if present
        pj = pair[i]
        if pj != -1:
            neigh = neigh[neigh != pj]

        neigh = neigh[:k_eff]
        frac_other[i] = (modality_labels[neigh] != modality_labels[i]).mean()

    return float(frac_other.mean())


def knn_label_transfer(Z_source, y_source, Z_target, y_target, k=15, metric="euclidean"):
    """
    kNN label transfer: predict y_target from neighbors in Z_source.
    Returns predictions, accuracy, macro-F1, confusion matrix.
    """
    Z_source = np.asarray(Z_source, dtype=np.float32)
    Z_target = np.asarray(Z_target, dtype=np.float32)
    y_source = np.asarray(y_source, dtype=str)
    y_target = np.asarray(y_target, dtype=str)

    nn = NearestNeighbors(n_neighbors=k, metric=metric)
    nn.fit(Z_source)
    nbrs = nn.kneighbors(Z_target, return_distance=False)

    preds = []
    for inds in nbrs:
        votes = y_source[inds]
        vals, cnts = np.unique(votes, return_counts=True)
        preds.append(vals[np.argmax(cnts)])
    preds = np.asarray(preds, dtype=str)

    acc = float(accuracy_score(y_target, preds))
    macro_f1 = float(f1_score(y_target, preds, average="macro"))
    classes = np.unique(np.concatenate([y_source, y_target]))
    cm = confusion_matrix(y_target, preds, labels=classes)
    return preds, acc, macro_f1, cm, classes


# %% [code]
from univi.evaluation import encode_adata

Z_rna  = encode_adata(model, rna_test_pp,      modality="rna",  latent="modality_mean", device=device, batch_size=1024)
Z_atac = encode_adata(model, atac_test_lsi,    modality="atac", latent="modality_mean", device=device, batch_size=1024)

labels_rna  = rna_test_pp.obs["cell_type"].astype(str).to_numpy()
labels_atac = atac_test_lsi.obs["cell_type"].astype(str).to_numpy()
assert Z_rna.shape[0] == labels_rna.shape[0]
assert Z_atac.shape[0] == labels_atac.shape[0]

# FOSCTTM (subsample)
n_total = rna_test_pp.n_obs
sub = np.random.default_rng(42).choice(n_total, size=min(20000, n_total), replace=False)
fos_mean, fos_sem = foscttm_chunked(Z_rna[sub], Z_atac[sub], block=512)

# modality mixing on stacked embeddings (NOT fused duplicates)
Z_concat = np.vstack([Z_rna, Z_atac])
mods = np.array(["rna"]*Z_rna.shape[0] + ["atac"]*Z_atac.shape[0], dtype=object)
mix = modality_mixing_score(Z_concat, mods, k=30)

# label transfer
pred_atac, acc_r2a, f1_r2a, cm_r2a, classes = knn_label_transfer(Z_rna,  labels_rna,  Z_atac, labels_atac, k=3)
pred_rna,  acc_a2r, f1_a2r, cm_a2r, _       = knn_label_transfer(Z_atac, labels_atac, Z_rna,  labels_rna,  k=3)

print("FOSCTTM:", fos_mean, "±", fos_sem)
print("mixing:", mix)
print("RNA→ATAC acc/F1:", acc_r2a, f1_r2a)
print("ATAC→RNA acc/F1:", acc_a2r, f1_a2r)

plot_confusion(cm_r2a, classes, title="Label transfer (RNA→ATAC)", normalize="true")
plot_confusion(cm_a2r, classes, title="Label transfer (ATAC→RNA)", normalize="true")


# %% [code]
print(atac_train_lsi.obs['cell_type'].value_counts())


# %% [code]
print(atac_val_lsi.obs['cell_type'].value_counts())


# %% [code]
print(atac_test_lsi.obs['cell_type'].value_counts())


# %% [code]
#rna_u = rna_test_pp.copy()
#atac_u = atac_test_lsi.copy()

#rna_u.obs["modality"] = "rna"
#atac_u.obs["modality"] = "atac"

#combo = ad.concat([rna_u, atac_u], join="outer", label="modality", keys=["rna","atac"], index_unique="-")

# IMPORTANT: put the correct latent for each half
#combo.obsm["X_univi_sep"] = np.vstack([Z_rna, Z_atac]).astype(np.float32)


# %% [code]
# neighbors/umap on the *separate* stacked latent
#sc.pp.neighbors(combo, use_rep="X_univi_sep", n_neighbors=15)
#sc.tl.umap(combo)


# %% [code]
# 6) UMAPs
#sc.pl.umap(combo, color=["modality"], frameon=False, size=25.0)
#sc.pl.umap(combo, color=[celltype_key], frameon=False, size=25.0)


# %% [markdown]
### Figure 4G–H: cross-modal prediction panels (interpretable)

# %% [code]
# 1) Predict ATAC-LSI from RNA, and RNA from ATAC-LSI
# If ue.cross_modal_predict exists in your version, use it; otherwise use your helper.
try:
    X_atac_hat = ue.cross_modal_predict(model, rna_test_pp, "rna", "atac", device=device, batch_size=1024)
    X_rna_hat  = ue.cross_modal_predict(model, atac_test_lsi, "atac", "rna", device=device, batch_size=1024)
except Exception:
    # fallback: use your cross_modal_predict_subset and request all features
    X_atac_hat = cross_modal_predict_subset(model, rna_test_pp, "rna", "atac",
                                           feat_idx=np.arange(atac_test_lsi.n_vars),
                                           device=device, batch_size=1024)
    X_rna_hat  = cross_modal_predict_subset(model, atac_test_lsi, "atac", "rna",
                                           feat_idx=np.arange(rna_test_pp.n_vars),
                                           device=device, batch_size=1024)

X_atac_obs = np.asarray(atac_test_lsi.X, dtype=np.float32)
X_rna_obs  = np.asarray(rna_test_pp.X.toarray() if sp.issparse(rna_test_pp.X) else rna_test_pp.X, dtype=np.float32)

# 2) Pick a few LSI dims to show as "accessibility programs"
lsi_dims = [0, 1, 2, 3]  # choose ones that look biological; you can rank by variance too

for d in lsi_dims:
    rna_test_pp.obs[f"ATAC_LSI{d}_obs"] = X_atac_obs[:, d]
    rna_test_pp.obs[f"ATAC_LSI{d}_hat"] = X_atac_hat[:, d]

# ensure you have a UMAP basis to plot on (use the shared latent UMAP for consistency)
if "X_umap" not in rna_test_pp.obsm:
    tmp = rna_test_pp.copy()
    sc.pp.neighbors(tmp, use_rep="X_univi", n_neighbors=15)
    sc.tl.umap(tmp)
    rna_test_pp.obsm["X_umap"] = tmp.obsm["X_umap"].copy()

# plot obs vs predicted for LSI programs (nice 2×4 grid)
keys = []
for d in lsi_dims:
    keys += [f"ATAC_LSI{d}_obs", f"ATAC_LSI{d}_hat"]


# %% [code]
# Scanpy defaults (affects sc.pl.*)
sc.set_figure_params(
    figsize=(10, 8),    # bigger canvas
    dpi=100,            # on-screen sharpness
    dpi_save=300,       # saved file sharpness
    fontsize=10,
    frameon=False,
)

# Matplotlib defaults (affects plt.*)
plt.rcParams.update({
    "figure.figsize": (10, 8),
    "figure.dpi": 100,
    "savefig.dpi": 300,
    "savefig.bbox": "tight",
    "savefig.pad_inches": 0.1,
    "axes.titlesize": 12,
    "axes.labelsize": 10,
    "legend.fontsize": 10,
})


# %% [code]
'''
rna_u = rna_test_pp.copy()
atac_u = atac_test_lsi.copy()

rna_u.obs["modality"] = "rna"
atac_u.obs["modality"] = "atac"

combo = ad.concat([rna_u, atac_u], join="outer", label="modality", keys=["rna","atac"], index_unique="-")

# IMPORTANT: put the correct latent for each half
combo.obsm["X_univi_sep"] = np.vstack([Z_rna, Z_atac]).astype(np.float32)

# neighbors/umap on the *separate* stacked latent
sc.pp.neighbors(combo, use_rep="X_univi_sep", n_neighbors=15)
sc.tl.umap(combo)

# plot UMAPs
sc.pl.umap(combo, color=["modality"], frameon=False, size=25.0)
sc.pl.umap(combo, color=[celltype_key], frameon=False, size=25.0)
'''

# %% [code]
print(rna_test_pp)


# %% [code]
#sc.pl.umap(combo, color=keys, ncols=4, frameon=False, size=75, wspace=0.25)
#sc.pl.embedding(rna_test_pp, color='cell_type', ncols=4, frameon=False, basis='X_umap', size=75, wspace=0.25)


# %% [code]
rna_markers = ["IL7R","CCR7","NKG7","GNLY","MS4A1","CD74","LYZ","S100A9"]
rna_markers = [g for g in rna_markers if g in rna_test_pp.var_names]

for g in rna_markers:
    j = rna_test_pp.var_names.get_loc(g)
    rna_test_pp.obs[f"RNA_{g}_obs"] = X_rna_obs[:, j]
    rna_test_pp.obs[f"RNA_{g}_hat"] = X_rna_hat[:, j]

keys = []
for g in rna_markers[:6]:
    keys += [f"RNA_{g}_obs", f"RNA_{g}_hat"]


# %% [code]
#sc.pl.umap(rna_test_pp, color=keys, ncols=4, frameon=False, size=75, wspace=0.25)


# %% [code]
def pearson_per_feature(X, Y):
    X = np.asarray(X, dtype=np.float32); Y = np.asarray(Y, dtype=np.float32)
    Xc = X - X.mean(axis=0, keepdims=True)
    Yc = Y - Y.mean(axis=0, keepdims=True)
    num = (Xc * Yc).sum(axis=0)
    den = np.sqrt((Xc**2).sum(axis=0) * (Yc**2).sum(axis=0)) + 1e-8
    return num / den

r_lsi = pearson_per_feature(X_atac_obs, X_atac_hat)
r_rna = pearson_per_feature(X_rna_obs,  X_rna_hat)

plt.figure(figsize=(6,4))
plt.hist(r_lsi, bins=50)
plt.title("RNA→ATAC (LSI) per-dimension Pearson r")
plt.xlabel("r"); plt.ylabel("# dims")
plt.tight_layout(); plt.show()

plt.figure(figsize=(6,4))
plt.hist(r_rna, bins=50)
plt.title("ATAC→RNA per-gene Pearson r")
plt.xlabel("r"); plt.ylabel("# genes")
plt.tight_layout(); plt.show()


# %% [code]
# Check out many of each cell type are in the test set for level 2 cell annotations
print(rna_test_pp.obs["cell_type"].value_counts())


# %% [markdown]
### All evaluation metrics beyond just the UMAP embeddings of the shared test set latent

# %% [code]
import json

from sklearn.neighbors import NearestNeighbors
from sklearn.metrics import confusion_matrix, accuracy_score, f1_score

# ----------------------------
# User settings
# ----------------------------
outdir = "figures/Figure4_Multiome_eval_reproducibility"
os.makedirs(outdir, exist_ok=True)

celltype_key = "cell_type"   # change to celltype.l1 / celltype.l3 as needed
k_mix = 20                     # k for modality mixing
k_lt  = 3                      # k for label transfer kNN
seed = 42


# %% [code]
# ---------- PRE-FLIGHT CHECK + FIX (run once right before metrics/plots) ----------
def _preflight_align_for_metrics(
    rna_test_pp, adt_test_pp, Z_rna, Z_adt, celltype_key="cell_type"
):
    import numpy as np

    # 1) basic shape checks
    print("rna_test_pp n_obs:", rna_test_pp.n_obs)
    print("adt_test_pp n_obs:", adt_test_pp.n_obs)
    print("Z_rna shape:", np.asarray(Z_rna).shape)
    print("Z_adt shape:", np.asarray(Z_adt).shape)

    # 2) enforce pairing by obs_names intersection (preserves RNA order)
    common = rna_test_pp.obs_names.intersection(adt_test_pp.obs_names)
    if len(common) != rna_test_pp.n_obs or len(common) != adt_test_pp.n_obs:
        print(f"[preflight] restricting to common paired cells: {len(common)}")

    rna_al = rna_test_pp[common].copy()
    adt_al = adt_test_pp[common].copy()
    adt_al = adt_al[rna_al.obs_names].copy()  # force identical order

    # 3) slice Z arrays to match aligned AnnData order (assumes Z were computed in the same order)
    # If your Z were computed from rna_test_pp/adt_test_pp directly, this is safe.
    # If not, you *must* recompute Z after alignment.
    if np.asarray(Z_rna).shape[0] != rna_al.n_obs or np.asarray(Z_adt).shape[0] != adt_al.n_obs:
        raise ValueError(
            "[preflight] Z arrays do not match aligned AnnData n_obs. "
            "Recompute Z_rna/Z_adt from rna_al/adt_al."
        )

    # 4) labels must come from the SAME objects you predict on
    labels_rna = rna_al.obs[celltype_key].astype(str).to_numpy()
    labels_adt = adt_al.obs[celltype_key].astype(str).to_numpy()

    # 5) final asserts (this prevents your exact error)
    assert rna_al.n_obs == adt_al.n_obs
    assert (rna_al.obs_names == adt_al.obs_names).all()
    assert len(labels_rna) == np.asarray(Z_rna).shape[0]
    assert len(labels_adt) == np.asarray(Z_adt).shape[0]

    print("[preflight] OK: paired, aligned, and lengths match.")
    return rna_al, adt_al, labels_rna, labels_adt


# run it:
rna_test_pp, atac_test_lsi, labels_rna, labels_atac = _preflight_align_for_metrics(
    rna_test_pp, atac_test_lsi, Z_rna, Z_atac, celltype_key=celltype_key
)


# %% [code]
'''
from univi.evaluation import encode_adata

Z_rna = encode_adata(model, rna_test_pp, modality="rna",
                     latent="modality_mean", device=device, batch_size=1024)
Z_atac = encode_adata(model, atac_test_lsi, modality="atac",
                     latent="modality_mean", device=device, batch_size=1024)

key_rna = "encode_adata(modality_mean)"
key_atac = "encode_adata(modality_mean)"

rna_test_pp.obsm["X_univi_rna"] = Z_rna
atac_test_lsi.obsm["X_univi_atac"] = Z_atac


print("max|Z_rna-Z_atac|:", float(np.max(np.abs(Z_rna - Z_atac))))
print("mean L2(Z_rna-Z_atac):", float(np.mean(np.linalg.norm(Z_rna - Z_atac, axis=1))))
'''

# %% [code]
# ----------------------------
# Compute metrics (test model performance)
# ----------------------------

# 0) Fused latent (if you truly want it)
#Z_fused = np.asarray(combo.obsm["X_univi"])
#mods_fused = combo.obs["modality"].to_numpy()
Z_fused = np.asarray(combo.obsm["X_univi"])
mods_fused = combo.obs["modality"].to_numpy()

# derive cell_ids that are identical for RNA+ADT copies
# works with index_unique="-" that produces "<orig>-rna" / "<orig>-adt"
cell_ids = combo.obs_names.to_series().str.rsplit("-", n=1).str[0].to_numpy()

mix_fused = modality_mixing_score_excluding_pairs(Z_fused, mods_fused, cell_ids, k=k_mix)

# 1) Modality-specific latents (Z_rna, Z_atac) already computed above


# 2) FOSCTTM on modality-specific latents (subsample ok)
n_total = int(rna_test_pp.n_obs)
n_fos = int(min(20000, n_total))
rng = np.random.default_rng(seed)
sub = rng.choice(n_total, size=n_fos, replace=False)

fos_rna_atac, fos_sem = foscttm_chunked(Z_rna[sub], Z_atac[sub], block=512)

# 3) Modality mixing on modality-specific latents
Z_concat = np.vstack([Z_rna, Z_atac])
mods = np.array(["rna"] * Z_rna.shape[0] + ["atac"] * Z_atac.shape[0], dtype=object)
mix_modality_specific = modality_mixing_score(Z_concat, mods, k=k_mix)

# 4) Label transfer (modality-specific)
# Make labels that match each embedding 1:1
labels_rna = rna_test_pp.obs[celltype_key].astype(str).to_numpy()
labels_atac = atac_test_lsi.obs[celltype_key].astype(str).to_numpy()

assert Z_rna.shape[0] == labels_rna.shape[0], (Z_rna.shape, labels_rna.shape)
assert Z_atac.shape[0] == labels_atac.shape[0], (Z_atac.shape, labels_atac.shape)

pred_atac, acc_r2a, f1_r2a, cm_r2a, classes = knn_label_transfer(
    Z_source=Z_rna, y_source=labels_rna,
    Z_target=Z_atac, y_target=labels_atac,
    k=k_lt
)
pred_rna, acc_a2r, f1_a2r, cm_a2r, _ = knn_label_transfer(
    Z_source=Z_atac, y_source=labels_atac,
    Z_target=Z_rna, y_target=labels_rna,
    k=k_lt
)

metrics = {
    "embedding_keys": {"rna": key_rna, "atac": key_atac, "fused": "combo.obsm['X_univi']"},
    "n_test": n_total,
    "celltype_key": celltype_key,

    "FOSCTTM_metric": "euclidean_sq",
    "FOSCTTM_subsample_n": n_fos,
    "FOSCTTM_rna_vs_atac_mean": float(fos_rna_atac),
    "FOSCTTM_rna_vs_atac_sem": float(fos_sem),

    "modality_mixing_k": int(k_mix),
    "modality_mixing_fused": float(mix_fused),
    "modality_mixing_modality_specific": float(mix_modality_specific),

    "label_transfer_k": int(k_lt),
    "label_transfer_rna_to_atac_acc": float(acc_r2a),
    "label_transfer_rna_to_atac_macroF1": float(f1_r2a),
    "label_transfer_atac_to_rna_acc": float(acc_a2r),
    "label_transfer_atac_to_rna_macroF1": float(f1_a2r),
}

print(json.dumps(metrics, indent=2))
with open(os.path.join(outdir, "figure4_metrics.json"), "w") as f:
    json.dump(metrics, f, indent=2)
pd.DataFrame([metrics]).to_csv(os.path.join(outdir, "figure4_metrics.csv"), index=False)


# %% [markdown]
### Peaks → Gene Activity (promoter-based) code

# %% [code]
import re

def build_promoters_from_gtf(
    gtf_path,
    *,
    upstream=2000,
    downstream=500,
    use_feature="gene",
    gene_name_col="gene_name",
    gene_id_col="gene_id",
):
    """
    Load GTF via PyRanges and construct promoter intervals around TSS.
    Uses '+' strand TSS=Start, '-' strand TSS=End.
    Returns DataFrame with Chromosome, Start, End, Strand, gene_name, gene_id.
    """
    try:
        import pyranges as pr
    except ImportError as e:
        raise ImportError("Please `pip install pyranges` to use gene activity.") from e

    gr = pr.read_gtf(gtf_path)
    df = gr.df

    if "Feature" not in df.columns:
        raise ValueError("Unexpected GTF columns; expected a 'Feature' column.")
    df = df[df["Feature"] == use_feature].copy()

    # keep only standard chromosomes if you want (optional)
    # df = df[df["Chromosome"].astype(str).str.match(r"^chr(\d+|X|Y|M)$")]

    if gene_name_col not in df.columns:
        # fall back to gene_id as name if needed
        df[gene_name_col] = df.get(gene_id_col, df["gene_id"] if "gene_id" in df.columns else "gene")

    # Compute TSS and promoter bounds
    tss = np.where(df["Strand"].values == "+", df["Start"].values, df["End"].values)
    pstart = (tss - int(upstream)).astype(np.int64)
    pend   = (tss + int(downstream)).astype(np.int64)
    pstart[pstart < 0] = 0

    out = pd.DataFrame({
        "Chromosome": df["Chromosome"].astype(str).values,
        "Start": pstart,
        "End": pend,
        "Strand": df["Strand"].astype(str).values,
        "gene_name": df[gene_name_col].astype(str).values,
        "gene_id": df[gene_id_col].astype(str).values if gene_id_col in df.columns else df[gene_name_col].astype(str).values,
    })

    # drop duplicate gene_name promoters (keep first) to avoid double counting
    out = out.drop_duplicates(subset=["gene_name"], keep="first").reset_index(drop=True)
    return out

def make_peak_to_gene_map(
    atac_adata,
    promoters_df,
    *,
    prefer_nearest_tss=True,
):
    """
    Build a sparse peaks->genes mapping matrix M (n_peaks x n_genes).
    If prefer_nearest_tss=True, assign each peak to the nearest overlapping promoter (by distance to TSS proxy),
    otherwise a peak can contribute to multiple genes (not usually desired).
    Returns: (M_csr, gene_names)
    """
    try:
        import pyranges as pr
    except ImportError as e:
        raise ImportError("Please `pip install pyranges` to use gene activity.") from e

    peaks_df = _parse_peak_varnames(atac_adata.var_names)
    peaks_df["peak_idx"] = np.arange(peaks_df.shape[0], dtype=np.int64)

    prom = promoters_df.copy()
    prom["gene_idx"] = pd.Index(prom["gene_name"]).factorize()[0]
    # factorize above is stable but we want explicit ordering:
    gene_names = prom["gene_name"].to_numpy()
    gene_to_idx = {g:i for i,g in enumerate(gene_names)}
    prom["gene_idx"] = prom["gene_name"].map(gene_to_idx).astype(np.int64)

    gr_peaks = pr.PyRanges(peaks_df[["Chromosome","Start","End","peak_idx"]])
    gr_prom  = pr.PyRanges(prom[["Chromosome","Start","End","Strand","gene_name","gene_idx"]])

    ov = gr_peaks.join(gr_prom).df
    if ov.empty:
        raise ValueError("No peak↔promoter overlaps found. Check genome build / chr naming / promoter window sizes.")

    # If desired: pick a single gene per peak among overlaps (nearest promoter midpoint as proxy)
    if prefer_nearest_tss:
        # promoter midpoint proxy
        ov["prom_mid"] = ((ov["Start_b"].values + ov["End_b"].values) / 2.0)
        peak_mid = ((ov["Start"].values + ov["End"].values) / 2.0)
        ov["dist"] = np.abs(peak_mid - ov["prom_mid"])
        ov = ov.sort_values(["peak_idx","dist"]).drop_duplicates("peak_idx", keep="first")

    n_peaks = atac_adata.n_vars
    n_genes = len(gene_names)

    rows = ov["peak_idx"].to_numpy(dtype=np.int64)
    cols = ov["gene_idx"].to_numpy(dtype=np.int64)
    data = np.ones(rows.shape[0], dtype=np.float32)

    M = sp.csr_matrix((data, (rows, cols)), shape=(n_peaks, n_genes))
    return M, gene_names

def compute_gene_activity_adata(
    atac_adata,
    peak_to_gene_map,
    gene_names,
    *,
    counts_layer="counts",
    binarize=False,
):
    """
    Compute gene activity matrix (cells x genes) using sparse matmul:
      GA = ATAC_counts (cells x peaks) @ M (peaks x genes)

    Returns AnnData with X = GA (sparse), var_names = gene_names, obs copied.
    """
    Xp = atac_adata.layers[counts_layer] if counts_layer in atac_adata.layers else atac_adata.X
    if not sp.issparse(Xp):
        Xp = sp.csr_matrix(Xp)

    if binarize:
        Xp = Xp.copy()
        Xp.data[:] = 1.0

    GA = Xp @ peak_to_gene_map  # (cells x genes) sparse
    GA = GA.tocsr()

    ga = ad.AnnData(
        X=GA,
        obs=atac_adata.obs.copy(),
        var=pd.DataFrame(index=pd.Index(gene_names, name="gene")),
    )
    return ga


# %% [code]
def _parse_peak_varnames(varnames):
    """
    Parse ATAC peak names into (Chromosome, Start, End).

    Expected *core* pattern somewhere in each name:
        'chrX:123-456'

    If a name can't be parsed, we assign dummy coordinates
    'chrUn:0-1' so it won't overlap any promoters, but we keep
    array length == n_peaks (so downstream matrices still align).

    Returns
    -------
    peaks_df : DataFrame with columns ['Chromosome','Start','End']
               length == len(varnames)
    """
    chrom = []
    start = []
    end   = []
    bad   = []

    # regex that finds 'chr...:start-end' anywhere
    pat = re.compile(r"(chr[0-9XYM]+):(\d+)-(\d+)")

    for v in varnames:
        m = pat.search(v)
        if m:
            c, s, e = m.groups()
            chrom.append(c)
            start.append(int(s))
            end.append(int(e))
        else:
            bad.append(v)
            # dummy coord that won't intersect any promoter
            chrom.append("chrUn")
            start.append(0)
            end.append(1)

    if bad:
        print(f"[parse peaks] WARNING: could not parse {len(bad)} peaks; "
              "assigned dummy coords 'chrUn:0-1'. Examples:")
        for b in bad[:10]:
            print("   ", b)

    return pd.DataFrame(
        {"Chromosome": chrom, "Start": start, "End": end},
        index=pd.Index(varnames, name="peak_id"),
    )


# %% [code]
GTF_PATH = "/home/groups/precepts/ashforda/scOPE_github_stuff/data/reference/Homo_sapiens_GRCh38.p13.gencode.annotation.gtf"

# 1) build promoters from GTF
promoters = build_promoters_from_gtf(
    GTF_PATH,
    upstream=2000,
    downstream=500,
    use_feature="gene",   # typical
)

# 2) build peak→gene mapping using TRAIN peaks (assumes peaks identical across splits)
M_peak_gene, gene_names = make_peak_to_gene_map(atac_train, promoters, prefer_nearest_tss=True)

# 3) compute gene activity matrices for each split
atac_train_ga = compute_gene_activity_adata(atac_train, M_peak_gene, gene_names, counts_layer="counts", binarize=False)
atac_val_ga   = compute_gene_activity_adata(atac_val,   M_peak_gene, gene_names, counts_layer="counts", binarize=False)
atac_test_ga  = compute_gene_activity_adata(atac_test,  M_peak_gene, gene_names, counts_layer="counts", binarize=False)

# pairing sanity
assert (rna_train.obs_names == atac_train_ga.obs_names).all()
assert (rna_val.obs_names   == atac_val_ga.obs_names).all()
assert (rna_test.obs_names  == atac_test_ga.obs_names).all()

print(atac_train_ga, atac_val_ga, atac_test_ga)


# %% [code]
def preprocess_gene_activity_gaussian(ga_adata, target_sum=1e4, log1p=True):
    ga = ga_adata.copy()
    # normalize_total works on .X if it's sparse
    sc.pp.normalize_total(ga, target_sum=float(target_sum))
    if log1p:
        sc.pp.log1p(ga)
    return ga

atac_train_ga_pp = preprocess_gene_activity_gaussian(atac_train_ga, target_sum=1e4, log1p=True)
atac_val_ga_pp   = preprocess_gene_activity_gaussian(atac_val_ga,   target_sum=1e4, log1p=True)
atac_test_ga_pp  = preprocess_gene_activity_gaussian(atac_test_ga,  target_sum=1e4, log1p=True)


# %% [code]
# If dimensionality is too large, you can select top variable genes on train
# and subset all splits (same as HVG):

def subset_hvg_like(train_ga, val_ga, test_ga, n_top=5000):
    tmp = train_ga.copy()
    sc.pp.highly_variable_genes(tmp, n_top_genes=int(n_top), flavor="seurat")
    hv = tmp.var_names[tmp.var["highly_variable"].to_numpy()]
    return train_ga[:, hv].copy(), val_ga[:, hv].copy(), test_ga[:, hv].copy()

atac_train_ga_pp, atac_val_ga_pp, atac_test_ga_pp = subset_hvg_like(
    atac_train_ga_pp, atac_val_ga_pp, atac_test_ga_pp, n_top=5000
)


# %% [code]
train_dict = {"rna": rna_train_pp, "atac": atac_train_ga_pp}
val_dict   = {"rna": rna_val_pp,   "atac": atac_val_ga_pp}
test_dict  = {"rna": rna_test_pp,  "atac": atac_test_ga_pp}


# %% [code]
# Scanpy defaults (affects sc.pl.*)
sc.set_figure_params(
    figsize=(14, 12),   # bigger canvas
    dpi=200,            # on-screen sharpness
    dpi_save=600,       # saved file sharpness
    fontsize=14,
    frameon=False,
)

# Matplotlib defaults (affects plt.*)
plt.rcParams.update({
    "figure.figsize": (14, 12),
    "figure.dpi": 200,
    "savefig.dpi": 600,
    "savefig.bbox": "tight",
    "savefig.pad_inches": 0.1,
    "axes.titlesize": 16,
    "axes.labelsize": 14,
    "legend.fontsize": 12,
})


# %% [code]
# pick the split you want to visualize
ga = atac_test_ga_pp.copy()

# bring over labels from RNA if GA doesn't already have them
# (they should match by obs_names because you asserted pairing)
if "cell_type" in rna_test.obs and "cell_type" not in ga.obs:
    ga.obs["cell_type"] = rna_test.obs["cell_type"].astype(str).values

# neighbors/umap on gene-activity
ga.uns.pop("neighbors", None)
ga.uns.pop("umap", None)
for k in ("connectivities", "distances"):
    if k in ga.obsp:
        del ga.obsp[k]

sc.pp.pca(ga, n_comps=50)
sc.pp.neighbors(ga, use_rep="X_pca", n_neighbors=15)
sc.tl.umap(ga)

sc.pl.umap(ga, color=["cell_type"], frameon=False, size=75)


# %% [code]
# PBMC-ish marker set (edit freely)
ga_markers = [
    "MS4A1", "CD79A", "CD74",     # B
    "NKG7", "GNLY", "PRF1",       # NK / cytotoxic
    "LYZ", "S100A8", "S100A9",    # mono
    "IL7R", "CCR7", "LTB",        # CD4 naive/TCM-ish
    "FCGR3A", "LST1",             # CD16 mono-ish
]

# keep only genes present
ga_markers = [g for g in ga_markers if g in ga.var_names]
print("GA markers present:", ga_markers)
'''
sc.pl.umap(
    ga,
    color=ga_markers,
    frameon=False,
    ncols=4,
    wspace=0.25,
    size=75,
)
'''

# %% [code]
# compute mean gene activity per cell type (on your selected markers or top HVGs)
genes_for_heatmap = ga_markers  # or use ga.var_names[:200] / etc.

X = ga[:, genes_for_heatmap].X
if not hasattr(X, "toarray"):
    X_dense = np.asarray(X)
else:
    X_dense = X.toarray()

df = pd.DataFrame(X_dense, index=ga.obs_names, columns=genes_for_heatmap)
df["cell_type"] = ga.obs["cell_type"].astype(str).values

pb = df.groupby("cell_type")[genes_for_heatmap].mean()

# z-score per gene for visualization
pb_z = (pb - pb.mean(axis=0)) / (pb.std(axis=0) + 1e-8)

plt.figure(figsize=(0.5 * len(genes_for_heatmap) + 4, 0.35 * pb_z.shape[0] + 3))
plt.imshow(pb_z.values, aspect="auto", interpolation="nearest")
plt.yticks(np.arange(pb_z.shape[0]), pb_z.index)
plt.xticks(np.arange(pb_z.shape[1]), pb_z.columns, rotation=90)
plt.colorbar(label="z-scored mean gene activity")
plt.title("Gene activity pseudo-bulk by cell type")
plt.tight_layout()
plt.show()


# %% [code]
# Use paired test splits
ga = atac_test_ga_pp
rna = rna_test.copy()

# make sure same order
common = rna.obs_names.intersection(ga.obs_names)
rna = rna[common].copy()
ga  = ga[common].copy()
ga  = ga[rna.obs_names].copy()

# choose shared genes
shared = np.intersect1d(rna.var_names.astype(str), ga.var_names.astype(str))
shared = shared[:2000]  # keep it light; or pick markers
print("n shared genes:", len(shared))

# pull matrices
Xr = rna[:, shared].X
Xg = ga[:, shared].X
if hasattr(Xr, "toarray"): Xr = Xr.toarray()
if hasattr(Xg, "toarray"): Xg = Xg.toarray()
Xr = np.asarray(Xr, dtype=np.float32)
Xg = np.asarray(Xg, dtype=np.float32)

# per-gene Pearson (obs vs GA)
Xr_c = Xr - Xr.mean(axis=0, keepdims=True)
Xg_c = Xg - Xg.mean(axis=0, keepdims=True)
num = (Xr_c * Xg_c).sum(axis=0)
den = np.sqrt((Xr_c**2).sum(axis=0) * (Xg_c**2).sum(axis=0)) + 1e-8
r = num / den

plt.figure(figsize=(6,4))
plt.hist(r, bins=60)
plt.xlabel("Pearson r (RNA vs gene activity), per gene")
plt.ylabel("# genes")
plt.title("RNA↔Gene-activity agreement (paired test)")
plt.tight_layout()
plt.show()

# show top genes
top = np.argsort(r)[-20:][::-1]
print(pd.DataFrame({"gene": shared[top], "pearson_r": r[top]}))


# %% [code]
# Example: use gene activity as atac modality adata in your UniVI pipeline
atac_train_use = atac_train_ga_pp
atac_val_use   = atac_val_ga_pp
atac_test_use  = atac_test_ga_pp

print(atac_train_use, atac_val_use, atac_test_use)


# %% [code]


# %% [markdown]
### Other Figure 4/supplemental figures stuff for Multiome analysis

# %% [code]
import numpy as np
import anndata as ad
import scanpy as sc

# ------------------------------------------------------
# 1) modality-specific latents (separate encoders)
# ------------------------------------------------------
Z_rna = encode_adata(
    model, rna_test_pp,
    modality="rna",
    latent="modality_mean",   # <- RNA encoder mean
    device=device,
    batch_size=1024,
)

Z_atac = encode_adata(
    model, atac_test_lsi,
    modality="atac",
    latent="modality_mean",   # <- ATAC encoder mean
    device=device,
    batch_size=1024,
)

assert Z_rna.shape == Z_atac.shape
assert (rna_test_pp.obs_names == atac_test_lsi.obs_names).all()

# (optional sanity: they should NOT be identical if truly modality-specific)
print("mean L2(Z_rna - Z_adt):", float(np.mean(np.linalg.norm(Z_rna - Z_atac, axis=1))))

# ------------------------------------------------------
# 2) build combo and overlay the *two different* embeddings
# ------------------------------------------------------
rna_u = rna_test_pp.copy()
atac_u = atac_test_lsi.copy()
rna_u.obs["modality"] = "rna"
atac_u.obs["modality"] = "atac"

combo = ad.concat([rna_u, atac_u], join="outer", index_unique="-")

# IMPORTANT: this is the embedding you’ll use for UMAP/plots
combo.obsm["X_latent"] = np.vstack([Z_rna, Z_atac])

# ------------------------------------------------------
# 3) one UMAP on the stacked embedding
# ------------------------------------------------------
# clear stale graphs if re-running
for key in ("neighbors", "umap"):
    combo.uns.pop(key, None)
for k in ("connectivities", "distances"):
    if k in combo.obsp:
        del combo.obsp[k]

sc.pp.neighbors(combo, use_rep="X_latent", n_neighbors=15)
sc.tl.umap(combo, min_dist=0.5, random_state=0)


# %% [code]
# quick check: do the two modalities overlay?
sc.pl.umap(combo, color=["modality"], frameon=False, size=50)


# %% [code]
# quick check: do the two modalities overlay?
sc.pl.umap(combo, color=["cell_type"], frameon=False, size=50)


# %% [code]
#import anndata as ad
#import numpy as np
#import scanpy as sc

rna_u  = rna_test_pp.copy()
atac_u = atac_test_lsi.copy()
rna_u.obs["modality"]  = "rna"
atac_u.obs["modality"] = "atac"

combo = ad.concat([rna_u, atac_u], join="outer", index_unique="-")
combo.obsm["X_latent"] = np.vstack([Z_rna, Z_atac]).astype(np.float32)

sc.pp.neighbors(combo, use_rep="X_latent", n_neighbors=15)
sc.tl.umap(combo, min_dist=0.5, random_state=0)

# same embedding, different colorings:
sc.pl.umap(combo, color="modality", ncols=2, wspace=0.35, frameon=False, size=50)
sc.pl.umap(combo, color="cell_type", ncols=2, wspace=0.35, frameon=False, size=50)

# RNA expression: plot only RNA rows (cleanest)
#sc.pl.umap(combo[combo.obs["modality"]=="rna"], color=["NKG7","MS4A1"], frameon=False, size=50)

# ATAC programs/LSI: plot only ATAC rows (cleanest)
#sc.pl.umap(combo[combo.obs["modality"]=="atac"], color=["LSI_0","LSI_1"], frameon=False, size=50)


# %% [code]
import numpy as np
import scipy.sparse as sp
import torch

@torch.no_grad()
def cross_modal_predict_all(
    model,
    adata_src,
    src_mod,
    tgt_mod,
    *,
    device="cpu",
    batch_size=512,
    X_key="X",
    layer=None,
    use_moe=True,
):
    """
    Predict target-modality features for ALL cells in adata_src using UniVI.
    Returns a (n_cells, n_tgt_features) float32 numpy array.
    """
    from univi.data import _get_matrix

    model.eval()
    dev = torch.device(device)

    X = _get_matrix(adata_src, layer=layer, X_key=X_key)
    if sp.issparse(X):
        X = X.toarray()
    X = np.asarray(X, dtype=np.float32)

    outs = []
    for start in range(0, X.shape[0], int(batch_size)):
        end = min(start + int(batch_size), X.shape[0])
        xb = torch.as_tensor(X[start:end], dtype=torch.float32, device=dev)

        mu_dict, logvar_dict = model.encode_modalities({src_mod: xb})

        if use_moe and hasattr(model, "mixture_of_experts"):
            mu_z, _ = model.mixture_of_experts(mu_dict, logvar_dict)
        else:
            mu_z = mu_dict[src_mod]

        xhat = model.decode_modalities(mu_z)[tgt_mod]
        outs.append(xhat.detach().cpu().numpy())

    return np.vstack(outs).astype(np.float32, copy=False)


# %% [code]
import numpy as np, scipy.sparse as sp
import anndata as ad
import scanpy as sc

# 0) assume paired + aligned
assert (rna_test_pp.obs_names == atac_test_lsi.obs_names).all()

# 1) make combo + embedding (use your existing Z_rna/Z_atac)
rna_u = rna_test_pp.copy();  rna_u.obs["modality"]  = "rna"
atac_u = atac_test_lsi.copy(); atac_u.obs["modality"] = "atac"

combo = ad.concat([rna_u, atac_u], join="outer", index_unique="-")
combo.obsm["X_latent"] = np.vstack([Z_rna, Z_atac]).astype(np.float32)

# compute 2D UMAP once (stored in combo.obsm["X_umap"])
sc.pp.neighbors(combo, use_rep="X_latent", n_neighbors=15)
sc.tl.umap(combo, min_dist=0.5, random_state=0)

# 2) observed ATAC (LSI space)
A_obs = atac_test_lsi.X
if sp.issparse(A_obs): A_obs = A_obs.toarray()
A_obs = np.asarray(A_obs, dtype=np.float32)

# 3) predicted ATAC from RNA (same LSI dim count)
A_hat = cross_modal_predict_all(
    model, rna_test_pp, src_mod="rna", tgt_mod="atac",
    device=device, batch_size=512
).astype(np.float32)

assert A_hat.shape == A_obs.shape

# 4) write obs/hat scalars onto the correct modality rows in combo
mods = combo.obs["modality"].astype(str).to_numpy()
is_rna  = (mods == "rna")
is_atac = (mods == "atac")

dims = [0, 1, 2, 3, 4, 5, 6, 7, 8, 9]
#dims = [36, 37, 38, 39, 40, 41, 42, 43, 44, 45]

for j in dims:
    combo.obs[f"ATAC_LSI_{j}_obs"] = np.nan
    combo.obs[f"ATAC_LSI_{j}_hat"] = np.nan
    combo.obs.loc[combo.obs_names[is_atac], f"ATAC_LSI_{j}_obs"] = A_obs[:, j]
    combo.obs.loc[combo.obs_names[is_rna],  f"ATAC_LSI_{j}_hat"] = A_hat[:, j]

# 5) plot on SAME embedding
for j in dims:
    sc.pl.umap(
        combo,
        color=[f"ATAC_LSI_{j}_obs", f"ATAC_LSI_{j}_hat"],
        ncols=2, wspace=0.35, size=50, frameon=False
    )

# also show the embedding itself
sc.pl.umap(combo, color=["modality", "cell_type"], ncols=2, wspace=0.35, size=50, frameon=False)

# per-cell RMSE across chosen dims (on RNA rows)
J = np.array(dims, dtype=int)
rmse = np.sqrt(((A_hat[:, J] - A_obs[:, J])**2).mean(axis=1)).astype(np.float32)

combo.obs["ATAC_LSI_rmse"] = np.nan
combo.obs.loc[combo.obs_names[is_rna], "ATAC_LSI_rmse"] = rmse

sc.pl.umap(combo, color=["ATAC_LSI_rmse"], frameon=False, size=50)


# %% [code]


# %% [code]
import numpy as np, scipy.sparse as sp
import anndata as ad
import scanpy as sc

# 0) assume paired + aligned
assert (rna_test_pp.obs_names == atac_test_lsi.obs_names).all()

# 1) make combo + embedding (use your existing Z_rna/Z_atac)
rna_u = rna_test_pp.copy();   rna_u.obs["modality"]  = "rna"
atac_u = atac_test_lsi.copy(); atac_u.obs["modality"] = "atac"

combo = ad.concat([rna_u, atac_u], join="outer", index_unique="-")
combo.obsm["X_latent"] = np.vstack([Z_rna, Z_atac]).astype(np.float32)

# 2D UMAP once (stored in combo.obsm["X_umap"])
sc.pp.neighbors(combo, use_rep="X_latent", n_neighbors=15)
sc.tl.umap(combo, min_dist=0.5, random_state=0)

mods = combo.obs["modality"].astype(str).to_numpy()
is_rna  = (mods == "rna")
is_atac = (mods == "atac")

# 2) observed RNA (per cell x gene)
R_obs = rna_test_pp.X
if sp.issparse(R_obs): R_obs = R_obs.toarray()
R_obs = np.asarray(R_obs, dtype=np.float32)

# 3) predicted RNA from ATAC (same gene dim)
R_hat = cross_modal_predict_all(
    model, atac_test_lsi, src_mod="atac", tgt_mod="rna",
    device=device, batch_size=512
).astype(np.float32)

assert R_hat.shape == R_obs.shape

# 4) choose marker genes
#marker_genes = ["IL7R","CCR7","NKG7","GNLY","MS4A1","CD74","LYZ","S100A9"]
marker_genes = ["CD79A", "TRAC"]
marker_genes = [g for g in marker_genes if g in rna_test_pp.var_names]

gene_idx = [rna_test_pp.var_names.get_loc(g) for g in marker_genes]

# 5) write obs/hat scalars onto correct modality rows (obs on RNA rows, hat on ATAC rows)
for g, j in zip(marker_genes, gene_idx):
    combo.obs[f"RNA_{g}_obs"] = np.nan
    combo.obs[f"RNA_{g}_hat"] = np.nan
    combo.obs.loc[combo.obs_names[is_rna],  f"RNA_{g}_obs"] = R_obs[:, j]
    combo.obs.loc[combo.obs_names[is_atac], f"RNA_{g}_hat"] = R_hat[:, j]

# 6) plot on SAME embedding
for g in marker_genes:
    sc.pl.umap(
        combo,
        color=[f"RNA_{g}_obs", f"RNA_{g}_hat"],
        ncols=2, wspace=0.35, size=50, frameon=False
    )

# also show embedding itself
sc.pl.umap(combo, color=["modality", "cell_type"], ncols=2, wspace=0.35, size=50, frameon=False)

# 7) per-cell RMSE across marker genes (attach to ATAC rows, since hat lives there)
J = np.array(gene_idx, dtype=int)
rmse = np.sqrt(((R_hat[:, J] - R_obs[:, J])**2).mean(axis=1)).astype(np.float32)

combo.obs["RNA_marker_rmse"] = np.nan
combo.obs.loc[combo.obs_names[is_atac], "RNA_marker_rmse"] = rmse

sc.pl.umap(combo, color=["RNA_marker_rmse"], frameon=False, size=50)


# %% [code]
#import numpy as np
#import pandas as pd
#import anndata as ad
#import scanpy as sc
#import matplotlib.pyplot as plt
from matplotlib.collections import LineCollection

def overlay_umap_with_pair_lines(
    rna_adata,
    z_rna_key="X_univi_rna_enc",
    z_atac_key="X_univi_atac_enc",
    label_key=None,
    n_neighbors=15,
    random_state=0,
    point_size=8,
    line_sample=300,        # None = all pairs (can get messy)
    line_alpha=0.25,
    line_lw=1.5,
    line_color="k",         # or "gray", "tab:blue", etc.
):
    assert z_rna_key in rna_adata.obsm, f"Missing {z_rna_key}"
    assert z_atac_key in rna_adata.obsm, f"Missing {z_atac_key}"
    assert rna_adata.obsm[z_rna_key].shape == rna_adata.obsm[z_atac_key].shape

    # Two copies of same cells
    a = rna_adata.copy()
    b = rna_adata.copy()
    a.obs = a.obs.copy()
    b.obs = b.obs.copy()

    cell_ids = rna_adata.obs_names.astype(str)
    a.obs["cell_id"] = cell_ids
    b.obs["cell_id"] = cell_ids
    a.obs["modality"] = "rna_enc"
    b.obs["modality"] = "atac_enc"

    combo = ad.concat([a, b], join="outer", axis=0, index_unique="-")
    combo.obsm["X_overlay"] = np.vstack([rna_adata.obsm[z_rna_key], rna_adata.obsm[z_atac_key]])

    # Fresh neighbors/umap state
    combo.uns.pop("neighbors", None)
    combo.uns.pop("umap", None)
    for k in ("connectivities", "distances"):
        if k in combo.obsp:
            del combo.obsp[k]

    sc.pp.neighbors(combo, use_rep="X_overlay", n_neighbors=n_neighbors, random_state=random_state)
    sc.tl.umap(combo, random_state=random_state)

    # Table of coords
    um = pd.DataFrame(combo.obsm["X_umap"], columns=["umap1", "umap2"], index=combo.obs_names)
    df = combo.obs[["cell_id", "modality"]].join(um)

    # One row per cell_id with both modalities present
    wide = df.pivot(index="cell_id", columns="modality", values=["umap1", "umap2"]).dropna()

    # Optional subsample lines
    if line_sample is not None and wide.shape[0] > line_sample:
        wide = wide.sample(n=line_sample, random_state=random_state)

    x1 = wide[("umap1", "rna_enc")].to_numpy()
    y1 = wide[("umap2", "rna_enc")].to_numpy()
    x2 = wide[("umap1", "atac_enc")].to_numpy()
    y2 = wide[("umap2", "atac_enc")].to_numpy()

    # Build line segments for LineCollection: (n_lines, 2 points, 2 coords)
    segs = np.stack(
        [np.column_stack([x1, y1]), np.column_stack([x2, y2])],
        axis=1
    )

    # Plot
    fig, ax = plt.subplots(figsize=(8, 8), dpi=200)

    for mod in ["rna_enc", "atac_enc"]:
        sub = df[df["modality"] == mod]
        ax.scatter(sub["umap1"], sub["umap2"], s=point_size, alpha=0.7, label=mod)

    lc = LineCollection(
        segs,
        colors=line_color,
        linestyles=":",
        linewidths=line_lw,
        alpha=line_alpha,
    )
    ax.add_collection(lc)

    ax.set_xticks([]); ax.set_yticks([])
    ax.set_xlabel("UMAP1"); ax.set_ylabel("UMAP2")
    ax.legend(frameon=False, loc="best")
    ax.set_title("Overlay UMAP with paired-cell dotted links")
    plt.tight_layout()
    plt.show()

    # Optional extra scanpy plots (no pair lines)
    if label_key is not None and label_key in combo.obs:
        sc.pl.umap(combo, color=[label_key, "modality"], frameon=False, wspace=0.4)

    return combo


# %% [code]
label_key = 'cell_type'


# %% [code]
print(combo)


# %% [code]
import numpy as np
import anndata as ad
import scanpy as sc

def build_combo_for_umaps(rna_test_pp, atac_test_lsi, Z_rna, Z_atac, celltype_key="cell_type"):
    # --- pairing sanity ---
    assert (rna_test_pp.obs_names == atac_test_lsi.obs_names).all()
    assert Z_rna.shape == Z_atac.shape
    assert Z_rna.shape[0] == rna_test_pp.n_obs

    # --- copies with clear metadata ---
    r = rna_test_pp.copy()
    a = atac_test_lsi.copy()
    r.obs = r.obs.copy(); a.obs = a.obs.copy()

    r.obs["modality"] = "rna"
    a.obs["modality"] = "atac"
    r.obs["cell_id"]  = r.obs_names.astype(str)
    a.obs["cell_id"]  = a.obs_names.astype(str)

    # --- concat (2N rows) ---
    combo = ad.concat([r, a], join="outer", axis=0, index_unique="-")

    # --- ONE clean rep for neighbors/UMAP (no NaNs) ---
    combo.obsm["X_latent"] = np.vstack([Z_rna, Z_atac]).astype(np.float32, copy=False)

    # optional: stash fused/shared latent too, if you have it
    if ("X_univi" in rna_test_pp.obsm) and ("X_univi" in atac_test_lsi.obsm):
        combo.obsm["X_fused"] = np.vstack([rna_test_pp.obsm["X_univi"], atac_test_lsi.obsm["X_univi"]]).astype(np.float32)

    # final sanity: rep must be finite
    X = combo.obsm["X_latent"]
    if not np.isfinite(X).all():
        good = np.isfinite(X).all(axis=1)
        combo = combo[good].copy()
        combo.obsm["X_latent"] = np.asarray(combo.obsm["X_latent"], dtype=np.float32)

    return combo


def compute_umap_2d_and_3d(combo, rep_key="X_latent", n_neighbors=15, random_state=0, min_dist=0.5):
    # wipe stale state (safe if rerun)
    for key in ("neighbors", "umap"):
        combo.uns.pop(key, None)
    for k in ("connectivities", "distances"):
        if k in combo.obsp:
            del combo.obsp[k]

    # neighbors once
    sc.pp.neighbors(combo, use_rep=rep_key, n_neighbors=n_neighbors, random_state=random_state)

    # 2D
    sc.tl.umap(combo, n_components=2, min_dist=min_dist, random_state=random_state)
    combo.obsm["X_umap2"] = combo.obsm["X_umap"].copy()

    # 3D (reuses same neighbor graph)
    sc.tl.umap(combo, n_components=3, min_dist=min_dist, random_state=random_state)
    combo.obsm["X_umap3"] = combo.obsm["X_umap"].copy()

    # default back to 2D for scanpy plotting
    combo.obsm["X_umap"] = combo.obsm["X_umap2"].copy()
    return combo


# %% [code]
combo = build_combo_for_umaps(rna_test_pp, atac_test_lsi, Z_rna, Z_atac, celltype_key="cell_type")
combo = compute_umap_2d_and_3d(combo, rep_key="X_latent", n_neighbors=15, random_state=0)

# scanpy 2D plots
#sc.pl.umap(combo, color=["modality", "cell_type"], frameon=False, size=50)


# %% [code]
import numpy as np
import pandas as pd
import matplotlib.pyplot as plt
from matplotlib.collections import LineCollection

def overlay_pair_lines_2d(
    combo,
    *,
    basis_key="X_umap2",
    id_key="cell_id",
    modality_key="modality",
    mod_a="rna",
    mod_b="atac",

    color_by="cell_type",
    cmap="tab20",
    show_legend=True,
    legend_max=50,

    line_sample=5000,
    line_alpha=0.50,
    line_lw=1.5,
    line_color="k",

    point_size=75,
    point_alpha=0.85,

    figsize=(16, 14),
    dpi=400,
):
    coords = np.asarray(combo.obsm[basis_key])
    if coords.shape[1] != 2:
        raise ValueError(f"{basis_key} must be 2D for matplotlib overlay (got shape {coords.shape}).")

    df = combo.obs[[id_key, modality_key]].copy()
    df["u1"] = coords[:, 0]
    df["u2"] = coords[:, 1]

    # -----------------------------
    # Coloring
    # -----------------------------
    color_map = None
    if color_by is None:
        df["_color"] = "C0"

    else:
        if color_by not in combo.obs.columns:
            raise KeyError(f"color_by='{color_by}' not found in combo.obs.")
        df[color_by] = combo.obs[color_by].astype(str).values

        if color_by == modality_key or color_by == "modality":
            # Force classic blue/orange
            color_map = {str(mod_a): "C0", str(mod_b): "C1"}
            df["_color"] = df[color_by].map(color_map).fillna("0.7")
        else:
            cats = pd.unique(df[color_by])
            cmap_obj = plt.get_cmap(cmap, len(cats))
            color_map = {c: cmap_obj(i) for i, c in enumerate(cats)}
            df["_color"] = df[color_by].map(color_map)

    # -----------------------------
    # Pair lines
    # -----------------------------
    wide = df.pivot(index=id_key, columns=modality_key, values=["u1", "u2"]).dropna()
    if line_sample is not None and wide.shape[0] > int(line_sample):
        wide = wide.sample(n=int(line_sample), random_state=0)

    x1 = wide[("u1", mod_a)].to_numpy()
    y1 = wide[("u2", mod_a)].to_numpy()
    x2 = wide[("u1", mod_b)].to_numpy()
    y2 = wide[("u2", mod_b)].to_numpy()
    segs = np.stack([np.c_[x1, y1], np.c_[x2, y2]], axis=1)

    fig, ax = plt.subplots(figsize=figsize, dpi=dpi)

    for mod in [mod_a, mod_b]:
        sub = df[df[modality_key].astype(str) == str(mod)]
        ax.scatter(
            sub["u1"], sub["u2"],
            s=point_size,
            c=sub["_color"].tolist(),
            alpha=point_alpha,
            linewidths=0,
            label=str(mod),
        )

    ax.add_collection(LineCollection(
        segs, colors=line_color, linestyles=":", linewidths=line_lw, alpha=line_alpha
    ))

    ax.set_xticks([]); ax.set_yticks([])
    ax.set_xlabel("UMAP1"); ax.set_ylabel("UMAP2")
    ax.set_title(f"UMAP overlay + paired links (colored by {color_by})")

    # -----------------------------
    # Legend
    # -----------------------------
    if show_legend and color_by is not None and color_map is not None:
        cats = list(pd.unique(df[color_by]))
        cats_show = cats[:int(legend_max)]
        handles = [
            plt.Line2D([0], [0], marker="o", color="w",
                       markerfacecolor=color_map.get(c, "0.7"), markersize=7, label=c)
            for c in cats_show
        ]
        ax.legend(
            handles=handles,
            title=color_by,
            frameon=False,
            loc="center left",
            bbox_to_anchor=(1.02, 0.5),
            borderaxespad=0.0,
        )
        fig.subplots_adjust(right=0.72)
    else:
        ax.legend(frameon=False, loc="best")

    plt.tight_layout()
    
    #plt.show()
    
    return fig


# %% [code]
celltype_2d_umap = overlay_pair_lines_2d(combo, basis_key="X_umap2", mod_a="rna", mod_b="atac", color_by="cell_type", line_sample=5000)


# %% [code]
#celltype_2d_umap.show()


# %% [code]
modality_2d_umap = overlay_pair_lines_2d(combo, basis_key="X_umap2", mod_a="rna", mod_b="atac", color_by="modality", line_sample=5000)


# %% [code]
#modality_2d_umap.show()


# %% [code]
import numpy as np
import pandas as pd
import plotly.express as px
import plotly.graph_objects as go


def plot_umap3d_interactive(
    adata,
    *,
    rep_key="X_latent",           # representation to build neighbors/UMAP from (e.g., "X_latent")
    umap_key="X_umap3",           # where to store/read 3D coords
    n_neighbors=15,
    min_dist=0.5,
    random_state=0,

    color_by=None,                # obs key (categorical/continuous). None -> no color
    symbol_by=None,               # obs key (e.g., "modality")
    hover_cols=("cell_id", "modality", "cell_type"),

    point_size=3,
    opacity=0.85,

    draw_pair_lines=True,
    id_key="cell_id",
    modality_key="modality",
    mod_a="rna",
    mod_b="atac",
    line_sample=None,             # None=all
    line_width=2.5,
    line_opacity=0.45,
    line_color="rgba(0,0,0,0.25)",

    width=950,
    height=750,
    title=None,
):
    """
    Compute (if needed) a 3D UMAP in `adata.obsm[umap_key]` and return a Plotly 3D scatter.
    Optionally draws faint paired lines between modality copies using `id_key` and `modality_key`.

    Requirements:
      - `rep_key` exists in `adata.obsm` and is finite (no NaN/Inf)
      - If draw_pair_lines=True: `id_key` and `modality_key` exist in `adata.obs`,
        and each `id_key` appears twice (once per mod_a, once per mod_b).
    """
    import scanpy as sc

    # -----------------------
    # 0) Validate rep
    # -----------------------
    if rep_key not in adata.obsm:
        raise KeyError(f"rep_key='{rep_key}' not found in adata.obsm. Available: {list(adata.obsm.keys())}")

    X = np.asarray(adata.obsm[rep_key], dtype=np.float32)
    if X.ndim != 2:
        raise ValueError(f"adata.obsm['{rep_key}'] must be 2D; got shape {X.shape}")
    if not np.isfinite(X).all():
        bad = (~np.isfinite(X).all(axis=1)).sum()
        raise ValueError(f"Input contains NaN/Inf in '{rep_key}'. Bad rows: {bad}/{X.shape[0]}")

    # -----------------------
    # 1) Compute 3D UMAP if missing/wrong
    # -----------------------
    need_umap = (umap_key not in adata.obsm) or (adata.obsm[umap_key].shape[1] != 3)
    if need_umap:
        # Clear stale scanpy state safely
        for key in ("neighbors", "umap"):
            adata.uns.pop(key, None)
        for k in ("connectivities", "distances"):
            if k in adata.obsp:
                del adata.obsp[k]

        sc.pp.neighbors(adata, use_rep=rep_key, n_neighbors=int(n_neighbors), random_state=int(random_state))
        sc.tl.umap(adata, n_components=3, min_dist=float(min_dist), random_state=int(random_state))
        adata.obsm[umap_key] = adata.obsm["X_umap"].copy()

    coords = np.asarray(adata.obsm[umap_key], dtype=np.float32)
    if coords.shape[1] != 3:
        raise ValueError(f"{umap_key} must be 3D; got shape {coords.shape}")

    # -----------------------
    # 2) Build plotting DF
    # -----------------------
    df = adata.obs.copy()
    df = df.assign(u1=coords[:, 0], u2=coords[:, 1], u3=coords[:, 2])

    if color_by is not None and color_by not in df.columns:
        raise KeyError(f"color_by='{color_by}' not in adata.obs.")
    if symbol_by is not None and symbol_by not in df.columns:
        raise KeyError(f"symbol_by='{symbol_by}' not in adata.obs.")

    hover_data = {c: True for c in hover_cols if c in df.columns}
    if color_by is not None:
        hover_data[color_by] = True
    if symbol_by is not None:
        hover_data[symbol_by] = True

    fig = px.scatter_3d(
        df,
        x="u1", y="u2", z="u3",
        color=color_by if color_by is not None else None,
        symbol=symbol_by if symbol_by is not None else None,
        hover_data=hover_data,
        opacity=float(opacity),
    )
    fig.update_traces(marker=dict(size=float(point_size)))

    # -----------------------
    # 3) Optional paired lines
    # -----------------------
    if draw_pair_lines:
        if id_key not in df.columns or modality_key not in df.columns:
            raise KeyError(f"Need '{id_key}' and '{modality_key}' in adata.obs to draw pair lines.")

        d2 = df[[id_key, modality_key, "u1", "u2", "u3"]].copy()
        wide = d2.pivot(index=id_key, columns=modality_key, values=["u1", "u2", "u3"]).dropna()

        # ensure both modalities exist
        need_cols = [("u1", mod_a), ("u2", mod_a), ("u3", mod_a),
                    ("u1", mod_b), ("u2", mod_b), ("u3", mod_b)]
        missing = [c for c in need_cols if c not in wide.columns]
        if missing:
            raise KeyError(
                f"Missing required pivot columns for mod_a='{mod_a}', mod_b='{mod_b}'. "
                f"Check values in adata.obs['{modality_key}']."
            )

        if line_sample is not None and wide.shape[0] > int(line_sample):
            wide = wide.sample(n=int(line_sample), random_state=int(random_state))

        # Build segments separated by NaNs (Plotly trick)
        n = wide.shape[0]
        x = np.empty(n * 3, dtype=float)
        y = np.empty(n * 3, dtype=float)
        z = np.empty(n * 3, dtype=float)

        x[0::3] = wide[("u1", mod_a)].to_numpy()
        y[0::3] = wide[("u2", mod_a)].to_numpy()
        z[0::3] = wide[("u3", mod_a)].to_numpy()

        x[1::3] = wide[("u1", mod_b)].to_numpy()
        y[1::3] = wide[("u2", mod_b)].to_numpy()
        z[1::3] = wide[("u3", mod_b)].to_numpy()

        x[2::3] = np.nan
        y[2::3] = np.nan
        z[2::3] = np.nan

        fig.add_trace(
            go.Scatter3d(
                x=x, y=y, z=z,
                mode="lines",
                line=dict(width=float(line_width), color=line_color),
                opacity=float(line_opacity),
                hoverinfo="skip",
                showlegend=False,
                name=f"paired links ({mod_a}↔{mod_b})",
            )
        )

    # -----------------------
    # 4) Layout polish
    # -----------------------
    if title is None:
        if color_by is None and symbol_by is None:
            title = "Interactive 3D UMAP"
        else:
            title = f"Interactive 3D UMAP (color={color_by}, symbol={symbol_by})"

    fig.update_layout(
        width=int(width),
        height=int(height),
        margin=dict(l=10, r=10, t=50, b=10),
        title=title,
        scene=dict(
            xaxis_title="UMAP1",
            yaxis_title="UMAP2",
            zaxis_title="UMAP3",
        ),
        legend=dict(itemsizing="constant"),
    )
    return fig


# %% [code]
import plotly.io as pio
pio.renderers.default

# Classic Jupyter Notebook:
#pio.renderers.default = "notebook_connected"

# JupyterLab:
#pio.renderers.default = "jupyterlab"

# VS Code notebooks:
#pio.renderers.default = "vscode"


# %% [code]
celltype_fig = plot_umap3d_interactive(
    combo,
    rep_key="X_latent",
    color_by="cell_type",
    symbol_by="modality",
    draw_pair_lines=True,
    id_key="cell_id",
    modality_key="modality",
    mod_a="rna",
    mod_b="atac",
    line_sample=5000,
)


# %% [code]
config = {
    "displayModeBar": True,      # <- force modebar to always show
    "displaylogo": False,
    "toImageButtonOptions": {    # <- controls what the button exports
        "format": "png",
        "filename": "umap3d",
        "width": 2000,
        "height": 1600,
        "scale": 9,              # higher = higher-res PNG from the button
    },
}


# %% [code]
celltype_fig.show(config=config)


# %% [code]
modality_fig = plot_umap3d_interactive(
    combo,
    rep_key="X_latent",
    color_by="modality",
    symbol_by="modality",
    draw_pair_lines=True,
    id_key="cell_id",
    modality_key="modality",
    mod_a="rna",
    mod_b="atac",
    line_sample=5000,
)


# %% [code]
modality_fig.show(config=config)


# %% [code]


# %% [code]


# %% [code]


# %% [code]


# %% [code]


# %% [code]


# %% [code]


# %% [code]


# %% [code]


# %% [code]


# %% [code]


# %% [code]


# %% [code]


# %% [code]
````
