## Supplemental Code for "Unifying multimodal single-cell data with a mixture-of-experts *β*-variational autoencoder framework": Fig05_Multiome_bridge_finetuning.ipynb

````
# %% [markdown]
# UniVI manuscript - Figure 5 generation reproducible workflow
### Multiome RNA + ATAC as a bridge to integrate separate unimodal datasets; use Multiome data to train a model and then use the trained model on outside unimodal scRNA and scATAC data

Andrew Ashford, Pathways + Omics Group, Oregon Health & Science University, Portland, OR - 12/10/2025

This Jupyter Notebook will house the end-to-end workflow to generate the panels in Figure 5 of our manuscript, "Unifying multimodal single-cell data with a mixture-of-experts β-variational autoencoder framework" which is currently being revised for Genome Research and is available currently on bioRxiv at the following link: https://www.biorxiv.org/content/10.1101/2025.02.28.640429v1.full

GitHub for the project - including a Quickstart guide - can be found at: https://github.com/Ashford-A/UniVI

Package is pip installable via the command: 
```bash
pip install univi
```


# %% [markdown]
### Import modules

# %% [code]
# Import non-UniVI modules
import os
import sys
from pathlib import Path

import numpy as np
import pandas as pd
import scanpy as sc
import anndata as ad
import torch

from torch.utils.data import DataLoader, Subset


# %% [code]
import scipy.sparse as sp
from sklearn.feature_extraction.text import TfidfTransformer
from sklearn.decomposition import TruncatedSVD
from sklearn.preprocessing import StandardScaler

from sklearn.neighbors import NearestNeighbors
from sklearn.metrics import confusion_matrix, accuracy_score, f1_score


# %% [code]
# Import required UniVI modules
from univi import (
    ModalityConfig,
    UniVIConfig,
    TrainingConfig,
    UniVIMultiModalVAE,
    matching,
    UniVITrainer,
    write_univi_latent,
    MultiModalDataset,
)

import univi as uv
import univi.evaluation as ue
import univi.plotting as up


# %% [code]
# Double check UniVI module version
print("Installed version is univi v" + str(uv.__version__))


# %% [markdown]
### Specify device to use for model

Set "device" - preferably device should be "cuda" for speedier model implementation/training. Requires GPU and the correct packages/versions.


# %% [code]
print("torch:", torch.__version__)
print("torch.cuda.is_available():", torch.cuda.is_available())
device = "cuda" if torch.cuda.is_available() else "cpu"
print("Using device:", device)


# %% [markdown]
### Specify file paths

Where data lives.

# %% [code]
DATA_ROOT = Path("/home/groups/precepts/ashforda/UniVI_v2/UniVI_older-non_git/data/PBMC_10x_Multiome_data/MultiVI_combined_data")

MULTI_PATH = DATA_ROOT / "10x_Genomics_Multiome_split.h5ad"
UNI_RNA_PATH = DATA_ROOT / "Ding_scRNA_data_split.h5ad"
UNI_ATAC_PATH = DATA_ROOT / "Satpathy_scATAC_data_split.h5ad"

print("Combined Multiome file: ", MULTI_PATH)
print("Unimodal RNA file:", UNI_RNA_PATH)
print("Unimodal ATAC file:", UNI_ATAC_PATH)


# %% [markdown]
### Read in data

Read data into AnnData objects using the paths in the code chunk above.

# %% [code]
multi = sc.read_h5ad(MULTI_PATH)

print(multi)


# %% [code]
# Split into rna and atac adata objects for downstream use using the .var['modality'] variable
print(multi.var['modality'])


# %% [code]
rna_mask = multi.var['modality'] == 'Gene Expression'
atac_mask = multi.var['modality'] == 'Peaks'

print(rna_mask.value_counts())   # sanity check
print(atac_mask.value_counts())  # sanity check

rna = multi[:, rna_mask].copy()
atac = multi[:, atac_mask].copy()

print(rna)
print(atac)

print("RNA obs names head:", rna.obs_names[:5].tolist())
print("ATAC obs names head:", atac.obs_names[:5].tolist())


# %% [code]
# Read in unimodal data objects
uni_rna = sc.read_h5ad(UNI_RNA_PATH)
uni_atac = sc.read_h5ad(UNI_ATAC_PATH)

print(uni_rna)
print(uni_atac)

print("RNA obs names head:", uni_rna.obs_names[:5].tolist())
print("ATAC obs names head:", uni_atac.obs_names[:5].tolist())


# %% [markdown]
### Align cells between RNA and ATAC

Make sure the cell indices are aligned so UniVI knows which samples are paired.

# %% [code]
# Intersect barcodes
common_cells = rna.obs_names.intersection(atac.obs_names)
print("Common cells:", len(common_cells))

rna = rna[common_cells].copy()
atac = atac[common_cells].copy()

# Make sure order matches
atac = atac[rna.obs_names].copy()

assert np.array_equal(rna.obs_names.values, atac.obs_names.values)
print("obs_names aligned between RNA and ATAC.")


# %% [markdown]
### Set up Multiome RNA + ATAC bridge data train/val/test splits

# %% [code]
rng = np.random.default_rng(42)
n = rna.n_obs
idx = np.arange(n)
rng.shuffle(idx)

n_train = int(0.90 * n)
n_val   = int(0.10 * n)
n_test  = n - n_train - n_val

idx_train = idx[:n_train]
idx_val   = idx[n_train:n_train+n_val]
idx_test  = idx[n_train+n_val:]

rna_train  = rna[idx_train].copy()
rna_val    = rna[idx_val].copy()
rna_test   = rna[idx_test].copy()

atac_train = atac[idx_train].copy()
atac_val   = atac[idx_val].copy()
atac_test  = atac[idx_test].copy()


# %% [markdown]
### Subset each dataset by shared features and preprocess individually

Also specifying all the data preprocessing functions below, will be used in their own respective sections.

* RNA preprocessing (log1p + HVG + scale → Gaussian decoder)

* ATAC preprocessing (TF-IDF + LSI + scale → Gaussian decoder)


# %% [code]
import numpy as np
import pandas as pd
import scipy.sparse as sp
from sklearn.feature_extraction.text import TfidfTransformer
from sklearn.decomposition import TruncatedSVD
from sklearn.preprocessing import StandardScaler

import anndata as ad
import scanpy as sc


# =============================================================================
# 1) Multiome preprocessing (train/val/test) with shared statistics
# =============================================================================

def preprocess_multiome_splits(
    rna_train, atac_train,
    rna_val,   atac_val,
    rna_test,  atac_test,
    *,
    rna_counts_layer="counts",
    atac_counts_layer="counts",
    n_hvg=2000,
    hvg_list=None,          # <--- NEW
    target_sum=1e4,
    n_lsi=50,
    seed=0,
):
    """
    Preprocess paired RNA/ATAC Multiome splits.

    RNA:
      - define HVGs on TRAIN only
      - normalize_total + log1p on train/val/test
      - StandardScaler fit on TRAIN, applied to val/test

    ATAC:
      - TF-IDF + SVD (LSI) fit on TRAIN only
      - StandardScaler in LSI space fit on TRAIN, applied to val/test

    Returns:
      rna_train_pp, atac_train_lsi,
      rna_val_pp,   atac_val_lsi,
      rna_test_pp,  atac_test_lsi,
      hvg,
      tfidf, svd, atac_scaler,
      rna_scaler
    """

    # --- ensure counts layers exist ---
    for a in (rna_train, rna_val, rna_test):
        if rna_counts_layer not in a.layers:
            a.layers[rna_counts_layer] = a.X.copy()
    for a in (atac_train, atac_val, atac_test):
        if atac_counts_layer not in a.layers:
            a.layers[atac_counts_layer] = a.X.copy()

    # =========================
    # RNA: choose HVGs
    # =========================
    if hvg_list is None:
        # define HVGs *from the bridge / train split* (current behavior)
        rna_train_tmp = rna_train.copy()
        rna_train_tmp.X = rna_train_tmp.layers[rna_counts_layer]

        try:
            sc.pp.highly_variable_genes(
                rna_train_tmp,
                n_top_genes=int(n_hvg),
                flavor="seurat_v3",
            )
        except Exception:
            sc.pp.highly_variable_genes(
                rna_train_tmp,
                n_top_genes=int(n_hvg),
                flavor="seurat",
            )

        hvg = rna_train_tmp.var_names[
            rna_train_tmp.var["highly_variable"].to_numpy()
        ].tolist()
    else:
        # use user-supplied HVG list (e.g. from Ding / joint)
        # enforce intersection + order w.r.t. rna_train
        hvg = [g for g in hvg_list if g in rna_train.var_names]
        if len(hvg) == 0:
            raise ValueError(
                "Provided hvg_list has no overlap with rna_train.var_names."
            )
        # reindex to training order
        hvg = list(rna_train.var_names.intersection(hvg))
        if len(hvg) == 0:
            raise ValueError("After intersection, no HVGs remain in rna_train.")

    def _rna_lognorm_to_X(a):
        """Subset to HVGs, normalize_total + log1p into X (no scaling yet)."""
        adata = a[:, hvg].copy()

        if rna_counts_layer not in adata.layers:
            adata.layers[rna_counts_layer] = adata.X.copy()

        # if no cells, just propagate counts
        if adata.n_obs == 0:
            X = adata.layers[rna_counts_layer]
            if sp.issparse(X):
                adata.X = X.copy()
            else:
                adata.X = np.asarray(X, dtype=np.float32)
            return adata

        adata.layers["log1p"] = adata.layers[rna_counts_layer].copy()
        sc.pp.normalize_total(adata, target_sum=float(target_sum), layer="log1p")
        sc.pp.log1p(adata, layer="log1p")

        X = adata.layers["log1p"]
        if sp.issparse(X):
            X = X.toarray()
        adata.X = np.asarray(X, dtype=np.float32)
        return adata

    rna_train_pp = _rna_lognorm_to_X(rna_train)
    rna_val_pp   = _rna_lognorm_to_X(rna_val)
    rna_test_pp  = _rna_lognorm_to_X(rna_test)

    # ---- RNA scaling (feature-wise z-score across cells; fit on TRAIN only) ----
    rna_scaler = StandardScaler(with_mean=True, with_std=True)

    def _scale_rna_inplace(adata, scaler, fit: bool):
        if adata.n_obs == 0:
            return
        X = adata.X
        if sp.issparse(X):
            X = X.toarray()
        if fit:
            Xs = scaler.fit_transform(X)
        else:
            # sanity check: same #features
            if scaler.n_features_in_ != X.shape[1]:
                raise ValueError(
                    f"[RNA] StandardScaler expects {scaler.n_features_in_} features, "
                    f"but got {X.shape[1]}. Check that HVG list matches training."
                )
            Xs = scaler.transform(X)
        adata.X = Xs.astype(np.float32, copy=False)

    _scale_rna_inplace(rna_train_pp, rna_scaler, fit=True)
    _scale_rna_inplace(rna_val_pp,   rna_scaler, fit=False)
    _scale_rna_inplace(rna_test_pp,  rna_scaler, fit=False)

    # =========================
    # ATAC: TFIDF + SVD (LSI) fit on TRAIN only
    # =========================
    Xtr = atac_train.layers[atac_counts_layer]
    Xva = atac_val.layers[atac_counts_layer]
    Xte = atac_test.layers[atac_counts_layer]

    if not sp.issparse(Xtr):
        Xtr = sp.csr_matrix(Xtr)
    if not sp.issparse(Xva):
        Xva = sp.csr_matrix(Xva)
    if not sp.issparse(Xte):
        Xte = sp.csr_matrix(Xte)

    tfidf = TfidfTransformer(norm="l2", use_idf=True, smooth_idf=True, sublinear_tf=False)
    Xtr_t = tfidf.fit_transform(Xtr)
    Xva_t = tfidf.transform(Xva)
    Xte_t = tfidf.transform(Xte)

    svd = TruncatedSVD(n_components=int(n_lsi), random_state=int(seed))
    Ztr = svd.fit_transform(Xtr_t)
    Zva = svd.transform(Xva_t)
    Zte = svd.transform(Xte_t)

    atac_scaler = StandardScaler(with_mean=True, with_std=True)
    Ztr = atac_scaler.fit_transform(Ztr).astype(np.float32, copy=False)
    # sanity: LSI dim is fixed by svd, so this should always match
    Zva = atac_scaler.transform(Zva).astype(np.float32, copy=False)
    Zte = atac_scaler.transform(Zte).astype(np.float32, copy=False)

    atac_train_lsi = ad.AnnData(
        X=Ztr,
        obs=atac_train.obs.copy(),
        var=pd.DataFrame(index=[f"LSI_{i}" for i in range(Ztr.shape[1])]),
    )
    atac_val_lsi = ad.AnnData(
        X=Zva,
        obs=atac_val.obs.copy(),
        var=pd.DataFrame(index=[f"LSI_{i}" for i in range(Zva.shape[1])]),
    )
    atac_test_lsi = ad.AnnData(
        X=Zte,
        obs=atac_test.obs.copy(),
        var=pd.DataFrame(index=[f"LSI_{i}" for i in range(Zte.shape[1])]),
    )

    return (
        rna_train_pp,
        atac_train_lsi,
        rna_val_pp,
        atac_val_lsi,
        rna_test_pp,
        atac_test_lsi,
        hvg,
        tfidf,
        svd,
        atac_scaler,  # ATAC LSI scaler
        rna_scaler,   # RNA scaler
    )


# =============================================================================
# 2) Transform NEW / UNIMODAL data with the SAME pipeline
# =============================================================================

def transform_rna_with_hvg(
    rna,
    *,
    hvg,
    counts_layer="counts",
    target_sum=1e4,
    rna_scaler: StandardScaler | None = None,
):
    """
    Apply the same RNA preprocessing to a new AnnData:
      - subset to given HVGs (MUST match training HVGs/order)
      - normalize_total + log1p
      - optional z-score with `rna_scaler` fit on the train split
    """
    # sanity: make sure all HVGs are present
    missing = [g for g in hvg if g not in rna.var_names]
    if len(missing) > 0:
        raise ValueError(
            f"transform_rna_with_hvg: {len(missing)} HVGs are missing from the input "
            f"AnnData (e.g. {missing[:5]} ...). You must use the same gene set as training."
        )

    a = rna[:, hvg].copy()

    if counts_layer not in a.layers:
        a.layers[counts_layer] = a.X.copy()

    a.layers["log1p"] = a.layers[counts_layer].copy()
    sc.pp.normalize_total(a, target_sum=float(target_sum), layer="log1p")
    sc.pp.log1p(a, layer="log1p")

    X = a.layers["log1p"]
    if sp.issparse(X):
        X = X.toarray()
    X = np.asarray(X, dtype=np.float32)

    if rna_scaler is not None:
        if rna_scaler.n_features_in_ != X.shape[1]:
            raise ValueError(
                f"[RNA external] StandardScaler expects {rna_scaler.n_features_in_} features, "
                f"but got {X.shape[1]}. Make sure HVG list and order match training."
            )
        X = rna_scaler.transform(X)

    a.X = X.astype(np.float32, copy=False)
    return a


def transform_atac_with_lsi(
    atac,
    *,
    counts_layer="counts",
    tfidf=None,
    svd=None,
    atac_scaler=None,
):
    """
    Apply the same ATAC preprocessing to new AnnData:
      - ensure counts layer
      - TF-IDF using the fitted `tfidf`
      - project to LSI using the fitted `svd`
      - z-score in LSI space with `atac_scaler`
    """
    if tfidf is None or svd is None or atac_scaler is None:
        raise ValueError(
            "Need tfidf, svd, atac_scaler objects from preprocess_multiome_splits()."
        )

    a = atac.copy()
    if counts_layer not in a.layers:
        a.layers[counts_layer] = a.X.copy()

    X = a.layers[counts_layer]
    if not sp.issparse(X):
        X = sp.csr_matrix(X)

    Xt = tfidf.transform(X)
    Z  = svd.transform(Xt)
    Z  = atac_scaler.transform(Z).astype(np.float32, copy=False)

    atac_lsi = ad.AnnData(
        X=Z,
        obs=a.obs.copy(),
        var=pd.DataFrame(index=[f"LSI_{i}" for i in range(Z.shape[1])]),
    )
    return atac_lsi


def compute_hvgs_from_reference(
    ref_rna,
    bridge_rna=None,
    *,
    counts_layer="counts",
    n_hvg=5000,
) -> list[str]:
    """
    Compute HVGs from a reference RNA dataset (e.g. Ding),
    optionally restricted to genes shared with the bridge RNA.
    """
    ref = ref_rna.copy()

    if counts_layer not in ref.layers:
        ref.layers[counts_layer] = ref.X.copy()

    # restrict to shared genes if a bridge AnnData is provided
    if bridge_rna is not None:
        shared = ref.var_names.intersection(bridge_rna.var_names)
        ref = ref[:, shared].copy()

    ref.X = ref.layers[counts_layer]

    try:
        sc.pp.highly_variable_genes(
            ref,
            n_top_genes=int(n_hvg),
            flavor="seurat_v3",
        )
    except Exception:
        sc.pp.highly_variable_genes(
            ref,
            n_top_genes=int(n_hvg),
            flavor="seurat",
        )

    hvgs = ref.var_names[ref.var["highly_variable"].to_numpy()].tolist()
    return hvgs


# %% [code]
# Pseudocode idea
combined = ad.concat([uni_rna, rna], join="inner", label="dataset")

sc.pp.highly_variable_genes(
    combined,
    n_top_genes=2000,
    flavor="seurat_v3",
    batch_key="dataset",  # important: HVGs robust across datasets
)

hvg = combined.var_names[combined.var["highly_variable"]].tolist()


# %% [code]
# say you have ding_rna and multiome_rna
#hvg_ding = compute_hvgs_from_reference(
#    ref_rna=uni_rna,
#    bridge_rna=rna_train,      # multiome RNA train split
#    counts_layer="counts",
#    n_hvg=5000,
#)


# %% [code]
# 1) Fit on multiome train/val/test
(
    rna_train_pp,
    atac_train_lsi,
    rna_val_pp,
    atac_val_lsi,
    rna_test_pp,
    atac_test_lsi,
    hvg_used,
    tfidf,
    svd,
    atac_scaler,
    rna_scaler,
) = preprocess_multiome_splits(
    rna_train, atac_train,
    rna_val,   atac_val,
    rna_test,  atac_test,
    rna_counts_layer="counts",
    atac_counts_layer="counts",
    n_hvg=2000,          # ignored when hvg_list is provided
    #hvg_list=hvg_ding,
    hvg_list=hvg,
    target_sum=1e4,
    n_lsi=100,
    seed=42,
)

print("Multiome is using HVGs:", len(hvg_used))
print("RNA train mean/std:", float(rna_train_pp.X.mean()), float(rna_train_pp.X.std()))
print("ATAC train mean/std:", float(atac_train_lsi.X.mean()), float(atac_train_lsi.X.std()))


# %% [code]
# 2) Transform unimodal RNA & ATAC using the SAME HVGs / LSI
#    (this is the “bridge” part)
# RNA: keep only bridge HVGs that exist in the unimodal dataset

uni_rna_pp = transform_rna_with_hvg(
    uni_rna,
    hvg=hvg,
    counts_layer="counts",
    target_sum=1e4,
    rna_scaler=rna_scaler,
)


# %% [code]
# ATAC: need overlapping peaks; align first
common_peaks = atac.var_names.intersection(uni_atac.var_names)
print("Shared ATAC peaks (bridge vs unimodal):", len(common_peaks))

atac_bridge_aligned = atac[:, common_peaks].copy()
uni_atac_aligned    = uni_atac[:, common_peaks].copy()

uni_atac_lsi = transform_atac_with_lsi(
    uni_atac_aligned,
    counts_layer="counts",
    tfidf=tfidf,
    svd=svd,
    atac_scaler=atac_scaler,
)

# Now you have:
# - rna_train_pp / rna_val_pp / rna_test_pp 
# - atac_train_lsi / atac_val_lsi / atac_test_lsi   
#   (for UniVI training + eval)
# - uni_rna_pp 
# - uni_atac_lsi   
#   (unimodal datasets mapped into the same feature spaces) 


# %% [code]
print(rna_train_pp)
print(rna_val_pp)
print(rna_test_pp)


# %% [code]
print(rna_train_pp.X.min())
print(rna_train_pp.X.max())


# %% [code]
print(atac_train_lsi)
print(atac_val_lsi)
print(atac_test_lsi)


# %% [code]
print(atac_train_lsi.X.min())
print(atac_train_lsi.X.max())


# %% [code]
print(uni_rna_pp)


# %% [code]
print(uni_atac_lsi)


# %% [markdown]
### Wrap into MultiModalDataset & DataLoaders

# %% [code]
from univi.data import align_paired_obs_names

pin_memory = (device == "cuda")

# Make sure each split is paired and ordered the same across modalities
# This was erroring out due to a function bug, fixed it and should be fixed by manuscript publication
#train_dict = align_paired_obs_names({"rna": rna_train_pp, "adt": atac_train_lsi})
#val_dict   = align_paired_obs_names({"rna": rna_val_pp,   "adt": atac_val_lsi})
#test_dict  = align_paired_obs_names({"rna": rna_test_pp,  "adt": atac_test_lsi})

# Using this instead for now since we know the data are already paired from above code
assert (rna_train_pp.obs_names == atac_train_lsi.obs_names).all()
assert (rna_val_pp.obs_names   == atac_val_lsi.obs_names).all()
assert (rna_test_pp.obs_names  == atac_test_lsi.obs_names).all()

train_dict = {"rna": rna_train_pp, "atac": atac_train_lsi}
val_dict   = {"rna": rna_val_pp,   "atac": atac_val_lsi}
test_dict  = {"rna": rna_test_pp,  "atac": atac_test_lsi}

# Build datasets (CPU tensors; trainer/model moves to GPU)
train_ds = MultiModalDataset(adata_dict=train_dict, X_key="X", device=None)
val_ds   = MultiModalDataset(adata_dict=val_dict,   X_key="X", device=None)
test_ds  = MultiModalDataset(adata_dict=test_dict,  X_key="X", device=None)

batch_size = 256
num_workers = 0

train_loader = DataLoader(
    train_ds,
    batch_size=batch_size,
    shuffle=True,
    num_workers=num_workers,
    pin_memory=pin_memory,
    drop_last=False,
)

'''
from torch.utils.data import DataLoader, WeightedRandomSampler

# labels for the TRAIN split (same order as train_ds cells)
y = rna_train_pp.obs["cell_type"].astype(str).to_numpy()

# inverse frequency weights
vals, counts = np.unique(y, return_counts=True)
freq = dict(zip(vals, counts))
w = np.array([1.0 / freq[c] for c in y], dtype=np.float64)
w = w / w.sum()

sampler = WeightedRandomSampler(
    weights=torch.as_tensor(w, dtype=torch.double),
    num_samples=len(w),   # one "epoch" worth of draws
    replacement=True
)

train_loader = DataLoader(
    train_ds,
    batch_size=batch_size,
    sampler=sampler,      # <-- instead of shuffle=True
    shuffle=False,
    num_workers=num_workers,
    pin_memory=pin_memory,
    drop_last=False,
)
'''

val_loader = DataLoader(
    val_ds,
    batch_size=batch_size,
    shuffle=False,
    num_workers=num_workers,
    pin_memory=pin_memory,
    drop_last=False,
)

test_loader = DataLoader(
    test_ds,
    batch_size=batch_size,
    shuffle=False,
    num_workers=num_workers,
    pin_memory=pin_memory,
    drop_last=False,
)

print("n_train / n_val / n_test:", train_ds.n_cells, val_ds.n_cells, test_ds.n_cells)
print("batches:", len(train_loader), len(val_loader), len(test_loader))

# sanity check one batch
x = next(iter(train_loader))
print({k: v.shape for k, v in x.items()})


# %% [markdown]
### UniVI configs

# %% [code]
#y_codes = rna.obs["cell_type"].astype("category").cat.codes.to_numpy()
#n_classes = int(y_codes.max() + 1)


# %% [code]
univi_cfg = UniVIConfig(
    latent_dim=30,
    beta=1.0,
    gamma=5.0,
    encoder_dropout=0.25,
    decoder_dropout=0.05,
    encoder_batchnorm=True,
    decoder_batchnorm=False,
    kl_anneal_start=0,
    kl_anneal_end=25,
    align_anneal_start=15,
    align_anneal_end=40,
    modalities=[
        ModalityConfig(
            name="rna",
            input_dim=rna_train_pp.n_vars,
            encoder_hidden=[512, 256, 128],
            decoder_hidden=[128, 256, 512],
            likelihood="gaussian",
        ),
        ModalityConfig(
            name="atac",
            input_dim=atac_train_lsi.n_vars,
            encoder_hidden=[128, 64],
            decoder_hidden=[64, 128],
            likelihood="gaussian",
        ),
    ],
)


# %% [markdown]
If you want raw-count NB/ZINB instead, you’d:
* Put raw counts in .layers["counts"]
* Set X_key="counts" in the dataset
* Use likelihood="nb" or "zinb" above

But for this "Figure 5" notebook, the scaled Gaussian setup is nicely stable and good for the best-integrated latent space.


# %% [markdown]
### Instantiate model and model objective

# %% [code]

model = UniVIMultiModalVAE(
    univi_cfg,
    loss_mode="v1",      # cross-recon + cross-posterior alignment
    #loss_mode="lite"
    #v1_recon="cross",   # full k→j cross-recon
    v1_recon="avg",
    #v1_recon_mix=0.5,
    normalize_v1_terms=True,
).to(device)

'''
model = UniVIMultiModalVAE(
    univi_cfg,
    loss_mode="v1",      # cross-recon + cross-posterior alignment
    #loss_mode="lite"
    #v1_recon="cross",   # full k→j cross-recon
    v1_recon="avg",
    #v1_recon_mix=0.5,
    normalize_v1_terms=True,
    n_label_classes=n_classes,
    label_loss_weight=5.0,
    label_ignore_index=-1,
    classify_from_mu=True,
).to(device)
'''
'''
model = UniVIMultiModalVAE(
    univi_cfg,
    loss_mode="lite",

    # Optional: keep the decoder-side classification head too
    n_label_classes=n_classes,
    label_loss_weight=2.0,

    # Encoder-side label expert injected into fusion
    use_label_encoder=True,
    label_moe_weight=3.5,      # >1 => labels influence fusion more
    unlabeled_logvar=20.0,     # very high => tiny precision => ignored in fusion
    label_encoder_warmup=5,    # wait N epochs before injecting labels into fusion
    label_ignore_index=-1,
).to("cuda")
'''

# %% [markdown]
### Instantiate TrainingConfig & trainer

# %% [code]
train_cfg = TrainingConfig(
    n_epochs=5000,
    batch_size=batch_size,
    lr=1e-3,
    weight_decay=1e-4,
    device=device,
    log_every=100,
    grad_clip=5.0,
    num_workers=0,
    seed=42,
    early_stopping=True,
    patience=150,         # Setting kind of high patience since training takes ~1s/iteration because of small-ish dataset
    min_delta=0.0,
)

print(train_cfg)
print(univi_cfg)


# %% [code]
trainer = UniVITrainer(
    model=model,
    train_loader=train_loader,
    val_loader=val_loader,
    train_cfg=train_cfg,
    device=device,
)

print(trainer)


# %% [markdown]
### Fit the model

# %% [code]
history = trainer.fit()

# history is a dict with keys like "train_loss", "val_loss", "beta", "gamma"
print("Training finished.")
print("Best val loss:", np.min(history["val_loss"]))


# %% [code]
import matplotlib.pyplot as plt

plt.figure()
plt.plot(history["train_loss"], label="train")
plt.plot(history["val_loss"], label="val")
plt.xlabel("Epoch")
plt.ylabel("Loss")
plt.legend()
plt.title("UniVI training (Multiome)")
plt.show()


# %% [markdown]
### Write latent z back to AnnData

# %% [code]
'''
# If I want to later use existing model weights to instantiate a model and change objectives.
# For example

import torch
from pathlib import Path
import univi as uv

# -----------------------
# 1) Load checkpoint
# -----------------------
ckpt_path = Path("checkpoints/pbmc_bridge_stage1_best.pt")
ckpt = torch.load(ckpt_path, map_location=device)

model_config_stage1 = ckpt.get("model_config")
train_config_stage1 = ckpt.get("train_config")

# -----------------------
# 2) Rebuild the model
# -----------------------
# However you normally do this:
model = uv.models.UniVI(**model_config_stage1["model_kwargs"])
model.to(device)

# Load weights
model.load_state_dict(ckpt["model_state"])
print("Loaded stage-1 weights from", ckpt_path)

# -----------------------
# 3) New training config (change objective)
# -----------------------
# You can start from the old config dict and tweak:
train_config_stage2 = train_config_stage1.copy()
train_config_stage2.update(
    {
        "mode": "lite",        # or "v1" -> "lite", or whatever flag you’re using
        "max_epochs": 30,      # shorter fine-tuning
        "lr": 1e-4,            # usually smaller LR for fine-tuning
        # optionally change loss weights here:
        # "lambda_align": 0.5,
        # "lambda_recon": 1.0,
        # "lambda_cls": 1.0,
    }
)

optimizer = torch.optim.AdamW(model.parameters(), lr=train_config_stage2["lr"])

# -----------------------
# 4) New trainer for stage 2
# -----------------------
trainer2 = uv.UniVITrainer(
    model=model,
    optimizer=optimizer,
    train_loader=train_loader_stage2,
    val_loader=val_loader_stage2,
    config=train_config_stage2,
    device=device,
)

trainer2.train()

'''


# %% [code]
from univi.evaluation import encode_adata

# Unimodal RNA → UniVI latent
Z_uni_rna = encode_adata(
    model,
    uni_rna_pp,
    modality="rna",
    latent="modality_mean",   # or "moe_mean" if you want fused-ish
    device=device,
    batch_size=1024,
)
uni_rna_pp.obsm["X_univi_rna"] = Z_uni_rna

# Unimodal ATAC → UniVI latent
Z_uni_atac = encode_adata(
    model,
    uni_atac_lsi,
    modality="atac",
    latent="modality_mean",
    device=device,
    batch_size=1024,
)
uni_atac_lsi.obsm["X_univi_atac"] = Z_uni_atac

print(Z_uni_rna.shape, Z_uni_atac.shape)


# %% [markdown]
Now both rna and atac have a shared latent:

* rna_test_pp.obsm["X_univi"]

* atac_test_pp.obsm["X_univi"]


# %% [markdown]
### Code to save/load modal if so inclined

# %% [code]
'''
A) You already have rna_train_pp / rna_val_pp / rna_test_pp (no resplitting)
from univi.utils.io import save_anndata_splits

save_anndata_splits(
    outdir=output_dir,
    prefix="rna",
    splits={"train": rna_train_pp, "val": rna_val_pp, "test": rna_test_pp},
    copy=False,
)

B) You want to split from the original using obs_names / indices / masks
save_anndata_splits(
    adata=rna,
    outdir=output_dir,
    prefix="rna",
    split_map={
        "train": rna_train_pp.obs_names.tolist(),
        "val":   rna_val_pp.obs_names.tolist(),
        "test":  rna_test_pp.obs_names.tolist(),
    },
    copy=False,
)

C) Only save the JSON split map (no huge .h5ad writes)
save_anndata_splits(
    adata=rna,
    outdir=output_dir,
    prefix="rna",
    split_map={
        "train": rna_train_pp.obs_names.tolist(),
        "val":   rna_val_pp.obs_names.tolist(),
        "test":  rna_test_pp.obs_names.tolist(),
    },
    save_h5ad=False,
    save_split_map=True,
)
'''


# %% [code]
beta_used = "1.0"
gamma_used = "5.0"
latent_dims_used = "30"


# %% [code]
output_dir = f'./results/univi_bridging_unimodal_data_pbmc_beta-{beta_used}_gamma-{gamma_used}_latent_dims-{latent_dims_used}_gaussian_both_reproducibility/'


# %% [code]
out_file = f"trained_multiome_model_beta-{beta_used}_gamma-{gamma_used}_latent_dims-{latent_dims_used}_gaussian_both_reproducibility.pt"


# %% [code]
from dataclasses import asdict

os.makedirs(output_dir, exist_ok=True)

# after training
#history = trainer.fit()

# trainer.model already has the best weights (because we restored best_state_dict)
ckpt_path = output_dir + out_file
torch.save(
    {
        "state_dict": trainer.model.state_dict(),
        "univi_cfg": asdict(univi_cfg),
        "best_epoch": trainer.best_epoch,
        "best_val_loss": trainer.best_val_loss,
    },
    ckpt_path,
)
print("Saved best model to:", ckpt_path)


# %% [code]
# Later to reload model
import torch
from univi.config import UniVIConfig, ModalityConfig
from univi.models.univi import UniVIMultiModalVAE

print("torch:", torch.__version__)
print("torch.cuda.is_available():", torch.cuda.is_available())
device = "cuda" if torch.cuda.is_available() else "cpu"
print("Using device:", device)

#device = "cuda"  # or "cuda" if available

ckpt = torch.load(
    output_dir + out_file,
    map_location=device,
)


# %% [code]
# ---- Rebuild UniVIConfig, making sure modalities are ModalityConfig objects ----
cfg_dict = ckpt["univi_cfg"]

# If this is an OmegaConf object or similar, make sure it's a plain dict
try:
    from omegaconf import DictConfig, OmegaConf
    if isinstance(cfg_dict, DictConfig):
        cfg_dict = OmegaConf.to_container(cfg_dict, resolve=True)
except ImportError:
    pass


# %% [code]
# Now rehydrate each modality
modalities = [ModalityConfig(**m) for m in cfg_dict["modalities"]]
cfg_dict = {**cfg_dict, "modalities": modalities}

univi_cfg_loaded = UniVIConfig(**cfg_dict)

# ---- Rebuild model + load weights ----
model_loaded = UniVIMultiModalVAE(univi_cfg_loaded).to(device)
model_loaded.load_state_dict(ckpt["state_dict"])

print("Best epoch was:", ckpt.get("best_epoch"), "val loss =", ckpt.get("best_val_loss"))


# %% [markdown]
### Figure 5 stuff

# %% [code]
from sklearn.neighbors import NearestNeighbors
from sklearn.metrics import confusion_matrix, accuracy_score, f1_score

# ----------------------------
# Metrics helpers
# ----------------------------
def foscttm_chunked(Z1, Z2, block=512):
    """
    Exact FOSCTTM computed in blocks to avoid NxN memory blowups.
    Assumes 1:1 pairing between rows i in Z1 and Z2.
    """
    Z1 = np.asarray(Z1, dtype=np.float32)
    Z2 = np.asarray(Z2, dtype=np.float32)
    assert Z1.shape == Z2.shape
    n = Z1.shape[0]

    Z2_T = Z2.T
    n2 = np.sum(Z2 * Z2, axis=1)  # (n,)

    fos = np.empty(n, dtype=np.float32)

    for i0 in range(0, n, block):
        i1 = min(i0 + block, n)
        A = Z1[i0:i1]  # (b, d)
        n1 = np.sum(A * A, axis=1)[:, None]  # (b,1)

        d2 = n1 + n2[None, :] - 2.0 * (A @ Z2_T)  # (b,n)

        true = d2[np.arange(i1 - i0), np.arange(i0, i1)]
        fos[i0:i1] = (d2 < true[:, None]).sum(axis=1) / (n - 1)

    return float(fos.mean()), float(fos.std(ddof=1) / np.sqrt(n))


def modality_mixing_score(Z, modality_labels, k=20, metric="euclidean"):
    """
    Vanilla modality mixing:
    Mean fraction of kNN neighbors that differ in modality.
    Use this for *non-duplicated* sets (e.g., concatenated modality-specific embeddings).
    """
    Z = np.asarray(Z, dtype=np.float32)
    modality_labels = np.asarray(modality_labels)

    n = Z.shape[0]
    if n <= 1:
        return 0.0

    k_eff = int(min(max(int(k), 1), n - 1))
    nn = NearestNeighbors(n_neighbors=k_eff + 1, metric=metric)
    nn.fit(Z)
    nbrs = nn.kneighbors(Z, return_distance=False)[:, 1:]  # drop self

    same = (modality_labels[nbrs] == modality_labels[:, None])
    return float((~same).mean())


def modality_mixing_score_excluding_pairs(Z, modality_labels, cell_ids, k=20, metric="euclidean"):
    """
    Pair-aware modality mixing for 'combo' style stacked data (same cell appears twice: RNA + ADT),
    where the fused embedding may be identical (or extremely close) for the paired duplicates.

    Computes: for each row, the fraction of neighbors from the other modality,
    AFTER removing the paired duplicate (same cell_id) from its neighbor list.

    cell_ids must map each row to a shared cell identifier (same for RNA+ADT copy).
    """
    Z = np.asarray(Z, dtype=np.float32)
    modality_labels = np.asarray(modality_labels)
    cell_ids = np.asarray(cell_ids).astype(str)

    n = Z.shape[0]
    if n <= 2:
        return 0.0

    # Need enough neighbors so we can drop self + paired duplicate and still have k
    k_eff = int(min(max(int(k), 1), n - 2))
    nn = NearestNeighbors(n_neighbors=k_eff + 2, metric=metric)
    nn.fit(Z)
    nbrs = nn.kneighbors(Z, return_distance=False)[:, 1:]  # drop self

    # Build "pair index": for each row i, pair[i] is the index of the other modality copy
    first = {}
    pair = np.full(n, -1, dtype=np.int64)
    for i, cid in enumerate(cell_ids):
        if cid in first:
            j = first[cid]
            pair[i] = j
            pair[j] = i
        else:
            first[cid] = i

    frac_other = np.empty(n, dtype=np.float32)
    for i in range(n):
        neigh = nbrs[i]

        # remove paired duplicate if present
        pj = pair[i]
        if pj != -1:
            neigh = neigh[neigh != pj]

        neigh = neigh[:k_eff]
        frac_other[i] = (modality_labels[neigh] != modality_labels[i]).mean()

    return float(frac_other.mean())


def knn_label_transfer(Z_source, y_source, Z_target, y_target, k=15, metric="euclidean"):
    """
    kNN label transfer: predict y_target from neighbors in Z_source.
    Returns predictions, accuracy, macro-F1, confusion matrix.
    """
    Z_source = np.asarray(Z_source, dtype=np.float32)
    Z_target = np.asarray(Z_target, dtype=np.float32)
    y_source = np.asarray(y_source, dtype=str)
    y_target = np.asarray(y_target, dtype=str)

    nn = NearestNeighbors(n_neighbors=k, metric=metric)
    nn.fit(Z_source)
    nbrs = nn.kneighbors(Z_target, return_distance=False)

    preds = []
    for inds in nbrs:
        votes = y_source[inds]
        vals, cnts = np.unique(votes, return_counts=True)
        preds.append(vals[np.argmax(cnts)])
    preds = np.asarray(preds, dtype=str)

    acc = float(accuracy_score(y_target, preds))
    macro_f1 = float(f1_score(y_target, preds, average="macro"))
    classes = np.unique(np.concatenate([y_source, y_target]))
    cm = confusion_matrix(y_target, preds, labels=classes)
    return preds, acc, macro_f1, cm, classes


from sklearn.neighbors import NearestNeighbors
from sklearn.metrics import precision_recall_fscore_support

# If plots ever don't show in classic notebook, run once:
# %matplotlib inline

# -------------------------
# show-or-save helper
# -------------------------
def _finish(fig=None, outpath=None, dpi=300, show=True, close=True):
    if outpath is not None:
        plt.savefig(outpath, dpi=dpi, bbox_inches="tight")
    if show:
        plt.show()
    if close:
        plt.close(fig if fig is not None else plt.gcf())

# -------------------------
# plots
# -------------------------
def plot_metrics_bar(
    metrics,
    keys,
    title="Figure 2 summary metrics",
    outpath=None,
    err_keys=None,          # dict: metric_key -> error_metric_key
    default_err=np.nan,     # np.nan = no error bar drawn; 0.0 = zero-length
    capsize=4,
):
    vals = np.array([float(metrics[k]) for k in keys], dtype=float)

    if err_keys is None:
        yerr = np.full_like(vals, default_err, dtype=float)
    else:
        yerr = np.array(
            [float(metrics[err_keys[k]]) if k in err_keys and err_keys[k] in metrics else default_err
             for k in keys],
            dtype=float
        )

    fig = plt.figure(figsize=(10, 6))
    plt.bar(keys, vals, yerr=yerr, capsize=capsize)
    plt.xticks(rotation=45, ha="right")
    plt.title(title)
    plt.tight_layout()

    # inline “finish” behavior
    if outpath is not None:
        plt.savefig(outpath, dpi=300, bbox_inches="tight")
    plt.show()
    plt.close(fig)


def plot_confusion(cm, classes, title, normalize="true", outpath=None, cmap="viridis"):
    import numpy as np
    import matplotlib.pyplot as plt

    cm = np.asarray(cm, dtype=float)
    if normalize == "true":
        cm = cm / (cm.sum(axis=1, keepdims=True) + 1e-12)
        subtitle = "Row-normalized"
    elif normalize == "pred":
        cm = cm / (cm.sum(axis=0, keepdims=True) + 1e-12)
        subtitle = "Col-normalized"
    elif normalize == "all":
        cm = cm / (cm.sum() + 1e-12)
        subtitle = "Global-normalized"
    else:
        subtitle = "Counts"

    classes = np.asarray(classes, dtype=str)
    n = len(classes)

    fig, ax = plt.subplots(figsize=(14, 13))
    im = ax.imshow(cm, interpolation="nearest", cmap=cmap, aspect="auto")

    # kill any gridlines (including ones from styles)
    ax.grid(False)
    ax.minorticks_off()

    ax.set_title(f"{title}\n({subtitle})")
    ax.set_xlabel("Predicted")
    ax.set_ylabel("True")

    ax.set_xticks(np.arange(n))
    ax.set_yticks(np.arange(n))
    ax.set_xticklabels(classes, rotation=90)
    ax.set_yticklabels(classes)

    cbar = fig.colorbar(im, ax=ax)
    cbar.set_label("value")

    fig.tight_layout()

    if outpath is not None:
        fig.savefig(outpath, dpi=300, bbox_inches="tight")

    plt.show()
    plt.close(fig)


def plot_per_class_f1(y_true, y_pred, title="Per-class F1", outpath=None, top_n=None):
    y_true = np.asarray(y_true).astype(str)
    y_pred = np.asarray(y_pred).astype(str)
    classes = np.unique(np.concatenate([y_true, y_pred]))

    _, _, f1, support = precision_recall_fscore_support(
        y_true, y_pred, labels=classes, zero_division=0
    )

    df = pd.DataFrame({"class": classes, "f1": f1, "support": support})
    df = df.sort_values(["f1", "support"], ascending=[True, False])

    if top_n is not None:
        df = df.head(int(top_n))

    fig = plt.figure(figsize=(10, 0.35 * len(df) + 2.0))
    plt.barh(df["class"], df["f1"])
    plt.xlabel("F1")
    plt.title(title)
    plt.xlim(0, 1)
    plt.tight_layout()
    _finish(fig, outpath=outpath, show=True, close=True)
    return df

def plot_modality_mixing_hist(Z, mods, k=20, metric="euclidean", title="Modality mixing", outpath=None):
    Z = np.asarray(Z, dtype=np.float32)
    mods = np.asarray(mods)

    n = Z.shape[0]
    k_eff = int(min(max(int(k), 1), n - 1))

    nn = NearestNeighbors(n_neighbors=k_eff + 1, metric=metric)
    nn.fit(Z)
    nbrs = nn.kneighbors(Z, return_distance=False)[:, 1:]
    frac_other = (mods[nbrs] != mods[:, None]).mean(axis=1)

    fig = plt.figure(figsize=(7, 4.5))
    plt.hist(frac_other, bins=60)
    plt.xlabel("Fraction of kNN from other modality")
    plt.ylabel("# cells")
    plt.title(title)
    plt.tight_layout()
    _finish(fig, outpath=outpath, show=True, close=True)
    return frac_other

def plot_foscttm_sanity(Z1, Z2, idx, title="FOSCTTM sanity", outpath=None):
    Z1s = np.asarray(Z1[idx], dtype=np.float32)
    Z2s = np.asarray(Z2[idx], dtype=np.float32)

    d_true = np.sum((Z1s - Z2s) ** 2, axis=1)

    nn = NearestNeighbors(n_neighbors=51, metric="euclidean")
    nn.fit(np.asarray(Z2, dtype=np.float32))
    dist, _ = nn.kneighbors(Z1s)
    d_min = dist[:, 0] ** 2

    fig = plt.figure(figsize=(5.5, 5.5))
    plt.scatter(d_true, d_min, s=8, alpha=0.4)
    mx = np.percentile(np.concatenate([d_true, d_min]), 99)
    plt.plot([0, mx], [0, mx], linewidth=1)
    plt.xlim(0, mx); plt.ylim(0, mx)
    plt.xlabel("d(true match)^2")
    plt.ylabel("d(nearest neighbor)^2")
    plt.title(title)
    plt.tight_layout()
    _finish(fig, outpath=outpath, show=True, close=True)

def plot_paired_distance_hist(Z_rna, Z_adt, idx, title="Paired latent distance (subset)", outpath=None):
    d_pair = np.sqrt(np.sum((Z_rna[idx] - Z_adt[idx]) ** 2, axis=1))
    fig = plt.figure(figsize=(7, 4.5))
    plt.hist(d_pair, bins=80)
    plt.xlabel("||z_rna - z_adt||")
    plt.ylabel("# cells")
    plt.title(title)
    plt.tight_layout()
    _finish(fig, outpath=outpath, show=True, close=True)


# %% [code]
celltype_key = "celltype"


# %% [code]
print(uni_rna_pp)
print(uni_atac_lsi)


# %% [code]
import numpy as np
import pandas as pd
import scanpy as sc
from sklearn.neighbors import NearestNeighbors
import matplotlib.pyplot as plt

# ------------------------------------------------
# Helper: find a reasonable raw celltype column
# ------------------------------------------------
def _get_celltype_series(
    adata,
    candidates=("celltype", "cell_type", "celltype.l2", "cell_type_l2", "annot", "leiden")
):
    """Return a string Series of celltype labels if any candidate is present, else None."""
    for key in candidates:
        if key in adata.obs:
            return adata.obs[key].astype(str), key
    return None, None


# %% [code]
print(set(uni_rna_pp.obs['celltype']))


# %% [code]
print(set(uni_atac_lsi.obs['celltype']))


# %% [code]
# Canonical, "figure-style" labels
CANON = {
    "B":                           "B",
    "Memory B":                    "Memory B",
    "CD14+ monocyte":              "CD14+ monocyte",
    "CD16+ monocyte":              "CD16+ monocyte",
    "Monocyte":                    "Monocyte",
    "NK":                          "NK",
    "CD4 T":                       "CD4 T",
    "CD4 T Helper":                "CD4 T helper",
    "Memory CD4 T":                "Memory CD4 T",
    "Naive CD4 T":                 "Naive CD4 T",
    "Memory CD8 T":                "Memory CD8 T",
    "Naive CD8 T":                 "Naive CD8 T",
    "Cytotoxic T":                 "Cytotoxic T",
    "Regulatory T":                "Regulatory T",
    "Dendritic Cell":              "Dendritic cell",
    "Plasmacytoid dendritic cell": "Plasmacytoid dendritic cell",
    "Conventional dendritic cell": "Conventional dendritic cell",
    "Megakaryocyte":               "Megakaryocyte",
    "General PBMC":                "General PBMC",
    "Unassigned":                  "Unassigned",
    "Unknown":                     "Unknown",
}

HARMONIZATION_MAP = {
    # 'Plasmacytoid dendritic cell', 'Natural killer cell', 'CD4+ T cell', 
    # 'Megakaryocyte', 'Cytotoxic T cell', 'CD16+ monocyte', 'B cell', 
    # 'Unassigned', 'CD14+ monocyte', 'Dendritic cell'
    "ding_rna": {
        # ---- T cells ----
        "Naive CD4 T":          "Naive CD4 T",
        "CD4 Naive":            "Naive CD4 T",

        "CD4 TCM":              "Memory CD4 T",
        "CD4 TEM":              "Memory CD4 T",
        "CD4 Memory":           "Memory CD4 T",

        "Naive CD8 T":          "Naive CD8 T",
        "CD8 Naive":            "Naive CD8 T",

        "CD8 TCM":              "Memory CD8 T",
        "CD8 TEM":              "Memory CD8 T",
        "CD8 TEMRA":            "Cytotoxic T",
        "CD8 Effector":         "Cytotoxic T",
        "CD8 Cytotoxic":        "Cytotoxic T",
        "Cytotoxic T cell":     "Cytotoxic T",

        "CD4 T cell":           "CD4 T",
        "CD4+ T cell":          "CD4 T",
        "CD4 Helper T cell":    "CD4 T helper",
        "Treg":                 "Regulatory T",
        "Regulatory T":         "Regulatory T",

        # ---- NK / NKT ----
        "NK":                   "NK",
        "NK cell":              "NK",
        "Natural killer cell":  "NK",

        # ---- B cells ----
        "B":                    "B",
        "B cell":               "B",
        "B cells":              "B",
        "Memory B":             "Memory B",
        "Memory B cell":        "Memory B",
        "Memory B cells":       "Memory B",

        # ---- Monocytes ----
        "CD14+ Monocyte":       "CD14+ monocyte",
        "CD14+ monocyte":       "CD14+ monocyte",
        "CD14 Monocyte":        "CD14+ monocyte",
        "Mono CD14":            "CD14+ monocyte",

        "CD16+ Monocyte":       "CD16+ monocyte",
        "CD16+ monocyte":       "CD16+ monocyte",
        "CD16 Monocyte":        "CD16+ monocyte",
        "Mono CD16":            "CD16+ monocyte",

        "Monocyte":             "Monocyte",
        "Monocytes":            "Monocyte",

        # ---- DC / pDC ----
        "Dendritic cell":       "Dendritic cell",
        "Dendritic cells":      "Dendritic cell",
        "DC":                   "Dendritic cell",
        "cDC":                  "Dendritic cell",

        "pDC":                  "Plasmacytoid dendritic cell",
        "BM pDC":               "Plasmacytoid dendritic cell",  # Maybe leave as BM?

        # ---- Progenitors / others ----
        "HSPC":                 "HSPC",
        "Progenitor":           "HSPC",
        "Megakaryocyte":        "Megakaryocyte",
        "PBMC":                 "General PBMC",

        # ---- catch-alls ----
        "Unassigned":           "Unassigned",
        "Unknown":              "Unknown",
    },

    # 'Dendritic_Cells', 'Naive_CD4_T_Cells', 'Monocytes', 'Memory_CD4_T_Cells', 
    # 'Memory_CD8_T_Cells', 'Regulatory_T_Cells', 'B_Cells', 'PBMC', 'NK_Cells', 
    # 'Naive_CD8_T_Cells', 'Bone_Marrow', 'BM_pDC', 'CD4_HelperT'
    "satpathy_atac": {
        # ---- T cells ----
        "Naive_CD4_T_Cells":    "Naive CD4 T",
        "CD4 T naive":          "Naive CD4 T",

        "Memory_CD4_T_Cells":   "Memory CD4 T",
        "CD4 T memory":         "Memory CD4 T",
        
        "CD4_HelperT":          "CD4 T helper",

        "Naive_CD8_T_Cells":    "Naive CD8 T",
        "CD8 Naive":            "Naive CD8 T",
        "CD8 T naive":          "Naive CD8 T",

        "CD8 Effector":         "Memory CD8 T",
        "Memory_CD8_T_Cells":   "Memory CD8 T",
        "CD8 Cytotoxic":        "Cytotoxic T",

        "Regulatory_T_Cells":   "Regulatory T",
        "Regulatory T":         "Regulatory T",
        "TREG":                 "Regulatory T",

        # ---- NK ----
        "NK":                   "NK",
        "NK cell":              "NK",
        "NK_Cells":             "NK",
        "Natural killer cell":  "NK",

        # ---- B cells ----
        "B":                    "B",
        "B cell":               "B",
        "B cells":              "B",
        "B_Cells":              "B",
        "Memory B":             "Memory B",
        "Memory B cell":        "Memory B",
        "Memory B cells":       "Memory B",

        # ---- Monocytes ----
        "CD14 Monocyte":        "CD14+ monocyte",
        "Mono CD14":            "CD14+ monocyte",

        "CD16 Monocyte":        "CD16+ monocyte",
        "Mono CD16":            "CD16+ monocyte",

        "Monocyte":             "Monocyte",
        "Monocytes":            "Monocyte",

        # ---- DC / pDC ----
        "Dendritic cell":       "Dendritic cell",
        "Dendritic_Cells":      "Dendritic cell",
        "Conventional DC":      "Dendritic cell",
        "cDC":                  "Conventional dendritic cell",

        "BM_pDC":               "Plasmacytoid dendritic cell",
        "pDC":                  "Plasmacytoid dendritic cell",

        # ---- Progenitors / others ----
        "HSPC":                 "HSPC",
        "Progenitor":           "HSPC",
        "PBMC":                 "General PBMC",
        "Bone_Marrow":          "General PBMC",
        "Megakaryocyte":        "Megakaryocyte",

        # ---- catch-alls ----
        "Unassigned":           "Unassigned",
        "Unknown":              "Unknown",
    },

    # Multiome still unlabeled for now
    "multiome_rna":  {},
    "multiome_atac": {},
}


# %% [code]
def add_harmonized_labels(adata, dataset_name):
    raw, key = _get_celltype_series(adata)
    if raw is None:
        adata.obs["celltype_raw"] = pd.Series(
            [f"unlabeled_{dataset_name}"] * adata.n_obs,
            index=adata.obs_names,
            dtype="object",
        )
        adata.obs["celltype_harmonized"] = adata.obs["celltype_raw"].copy()
        return adata

    adata.obs["celltype_raw"] = raw.astype(str)
    mapping = HARMONIZATION_MAP.get(dataset_name, {})

    # first-pass: dataset-specific mapping
    harm = adata.obs["celltype_raw"].map(mapping)

    # fall back to raw where unmapped
    mask_missing = harm.isna()
    harm[mask_missing] = adata.obs.loc[mask_missing, "celltype_raw"]

    # snap to canonical spelling if possible
    harm = harm.map(lambda x: CANON.get(x, x))

    adata.obs["celltype_harmonized"] = harm.astype("category")
    return adata


# %% [code]
from univi.evaluation import encode_adata

# ------------------------------------------------
# 2.1 Build a "bridge" reference from all Multiome splits
# ------------------------------------------------
bridge_rna = ad.concat(
    {"rna_train": rna_train_pp, "rna_val": rna_val_pp, "rna_test": rna_test_pp},
    axis=0,
    join="outer",
    label="split",
    index_unique=None,
)
bridge_atac = ad.concat(
    {"atac_train": atac_train_lsi, "atac_val": atac_val_lsi, "atac_test": atac_test_lsi},
    axis=0,
    join="outer",
    label="split",
    index_unique=None,
)

print("Bridge RNA:", bridge_rna.shape)
print("Bridge ATAC:", bridge_atac.shape)


# %% [code]
# ------------------------------------------------
# 2.2 Encode bridge and unimodal into UniVI latent
# ------------------------------------------------
# Bridge RNA
Z_bridge_rna = encode_adata(
    model,
    bridge_rna,
    modality="rna",
    latent="modality_mean",
    device=device,
    batch_size=1024,
)
bridge_rna.obsm["X_univi"] = Z_bridge_rna

# Bridge ATAC
Z_bridge_atac = encode_adata(
    model,
    bridge_atac,
    modality="atac",
    latent="modality_mean",
    device=device,
    batch_size=1024,
)
bridge_atac.obsm["X_univi"] = Z_bridge_atac

# Unimodal (already encoded earlier)
uni_rna_pp.obsm["X_univi"] = uni_rna_pp.obsm["X_univi_rna"]
uni_atac_lsi.obsm["X_univi"] = uni_atac_lsi.obsm["X_univi_atac"]


# %% [code]
# ------------------------------------------------
# 2.3 Annotate dataset + modality + reference flags
# ------------------------------------------------
bridge_rna.obs["dataset"] = "multiome_rna"
bridge_rna.obs["modality"] = "rna"
bridge_rna.obs["is_reference"] = True

bridge_atac.obs["dataset"] = "multiome_atac"
bridge_atac.obs["modality"] = "atac"
bridge_atac.obs["is_reference"] = True

uni_rna_pp.obs["dataset"] = "ding_rna"
uni_rna_pp.obs["modality"] = "rna"
uni_rna_pp.obs["is_reference"] = False

uni_atac_lsi.obs["dataset"] = "satpathy_atac"
uni_atac_lsi.obs["modality"] = "atac"
uni_atac_lsi.obs["is_reference"] = False


# %% [code]
# ------------------------------------------------
# 2.4 Add harmonized labels where possible
#     (bridge gets 'unlabeled_multiome_*' if no celltypes)
# ------------------------------------------------
bridge_rna = add_harmonized_labels(bridge_rna, "multiome")
bridge_atac = add_harmonized_labels(bridge_atac, "multiome")
uni_rna_pp  = add_harmonized_labels(uni_rna_pp,  "ding_rna")
uni_atac_lsi = add_harmonized_labels(uni_atac_lsi, "satpathy_atac")


# %% [code]
COARSE_MAP = {
    # B lineage
    "B": "B",
    "B cell": "B",
    "Memory B": "B",
    "Naive B": "B",

    # Monocytes
    "CD14+ monocyte": "Monocyte",
    "CD16+ monocyte": "Monocyte",
    "Monocyte": "Monocyte",

    # CD4 T lineage
    "CD4 T": "CD4 T",
    "CD4 T helper": "CD4 T",
    "Naive CD4 T": "CD4 T",
    "Memory CD4 T": "CD4 T",
    "Regulatory T": "CD4 T",
    "Treg": "CD4 T",

    # CD8 / cytotoxic T lineage
    "CD8 T": "CD8 / cytotoxic T",
    "Cytotoxic T": "CD8 / cytotoxic T",
    "Naive CD8 T": "CD8 / cytotoxic T",
    "Memory CD8 T": "CD8 / cytotoxic T",

    # NK
    "NK": "NK",
    "NK cell": "NK",

    # DCs
    "Dendritic cell": "Dendritic cell",
    "Plasmacytoid dendritic cell": "Dendritic cell",
    "pDC": "Dendritic cell",

    # Others
    "Megakaryocyte": "Megakaryocyte",
    "HSPC": "HSPC / progenitor",
    "General PBMC": "Other",
    "Unknown": "Other",
}
    

# %% [code]
for name, adata in {
    "ding_rna": uni_rna_pp,
    "satpathy_atac": uni_atac_lsi,
    "multiome_rna": bridge_rna,
    "multiome_atac": bridge_atac,
}.items():
    fine = adata.obs["celltype_harmonized"]  # whatever column you’re using now
    adata.obs["celltype_harmonized_coarse"] = fine.map(COARSE_MAP).fillna("Other")
    

# %% [code]
combo = ad.concat(
    {
        "ding_rna":      uni_rna_pp,
        "satpathy_atac": uni_atac_lsi,
        "multiome_rna":  bridge_rna,
        "multiome_atac": bridge_atac,
    },
    axis=0,
    join="outer",
    label="dataset",
    index_unique=None,
)

# sanity check: unique labels per dataset
for ds in ["ding_rna", "satpathy_atac"]:
    print("\nDataset:", ds)
    mask = combo.obs["dataset"] == ds
    print("raw:", combo.obs.loc[mask, "celltype_raw"].unique())
    print("harm:", combo.obs.loc[mask, "celltype_harmonized"].unique())


# %% [code]
# Check which raw labels got mapped to what, per dataset
for ds in ["ding_rna", "satpathy_atac"]:
    print("\n=== Dataset:", ds, "===")
    mask = combo.obs["dataset"] == ds
    tab = pd.crosstab(
        combo.obs.loc[mask, "celltype_raw"],
        combo.obs.loc[mask, "celltype_harmonized"]
    )
    print(tab)

# Dataset composition per harmonized label
print(
    pd.crosstab(
        combo.obs["celltype_harmonized"],
        combo.obs["dataset"],
        normalize="index"
    )
)


# %% [code]
# Check which raw labels got mapped to what, per dataset
for ds in ["ding_rna", "satpathy_atac"]:
    print("\n=== Dataset:", ds, "===")
    mask = combo.obs["dataset"] == ds
    tab = pd.crosstab(
        combo.obs.loc[mask, "celltype_raw"],
        combo.obs.loc[mask, "celltype_harmonized_coarse"]
    )
    print(tab)

# Dataset composition per harmonized label
print(
    pd.crosstab(
        combo.obs["celltype_harmonized_coarse"],
        combo.obs["dataset"],
        normalize="index"
    )
)


# %% [code]
# ------------------------------------------------
# 2.5 Concatenate all cells into one AnnData with shared latent
# ------------------------------------------------
'''
combo = ad.concat(
    {
        "multiome_rna": bridge_rna,
        "multiome_atac": bridge_atac,
        "ding_rna": uni_rna_pp,
        "satpathy_atac": uni_atac_lsi,
    },
    join="outer",
    index_unique=None,
    label="source",
)

print("Combined latent AnnData:", combo.shape)
assert "X_univi" in combo.obsm, "Missing X_univi latent in combined AnnData"
'''

# %% [code]
print(combo)
print(combo.obs['modality'])
print(combo.obs['celltype_harmonized'])
print(combo.obs['celltype_harmonized_coarse'])
print(combo.obs['dataset'])


# %% [code]
# ------------------------------------------------
# 3.1 Set plotting defaults
# ------------------------------------------------

sc.set_figure_params(
    figsize=(10, 8),
    dpi=100,
    dpi_save=300,
    fontsize=10,
    frameon=False,
)
plt.rcParams.update({
    "figure.figsize": (10, 8),
    "figure.dpi": 100,
    "savefig.dpi": 300,
    "savefig.bbox": "tight",
    "savefig.pad_inches": 0.1,
})


# %% [code]
# ------------------------------------------------
# 3.2 Compute neighbors/UMAP on UniVI latent
# ------------------------------------------------
sc.pp.neighbors(combo, n_neighbors=20, use_rep="X_univi", metric="euclidean")
#sc.pp.neighbors(combo, n_neighbors=30, use_rep="X_univi", metric="cosine")


# %% [code]
sc.tl.umap(combo, min_dist=0.3, spread=1.0)


# %% [code]
# ------------------------------------------------
# 3.3 UMAP views
# ------------------------------------------------
sc.pl.umap(
    combo,
    color=["dataset"],
    wspace=0.3,
    frameon=False,
    size=20,
    alpha=0.65,
    title=["Dataset"],
)


# %% [code]
sc.pl.umap(
    combo,
    color=["modality"],
    wspace=0.3,
    frameon=False,
    size=20,
    alpha=0.65,
    title=["Modality"],
)


# %% [code]
sc.pl.umap(
    combo,
    color=["tech"],
    wspace=0.3,
    frameon=False,
    size=20,
    alpha=0.65,
    title=["Sequencing technology"],
)
    

# %% [code]
# If you want to only show labels where they exist, subset:
has_labels = ~combo.obs["celltype_harmonized"].str.startswith("unlabeled_")
if has_labels.sum() > 0:
    sc.pl.umap(
        combo[has_labels],
        color=["celltype_harmonized"],
        wspace=0.3,
        frameon=False,
        size=20,
        alpha=0.65,
        title=["Harmonized cell types (unimodal)"],
    )
    
    sc.pl.umap(
        combo[has_labels],
        color=["celltype_harmonized_coarse"],
        wspace=0.3,
        frameon=False,
        size=20,
        alpha=0.65,
        title=["Harmonized coarse cell types (unimodal)"],
    )
    
    sc.pl.umap(
        combo[has_labels],
        color=["dataset"],
        wspace=0.3,
        frameon=False,
        size=20,
        alpha=0.65,
        title=["Dataset"],
    )
    

# %% [code]
# Show where the reference lives versus unimodal projections
sc.pl.umap(
    combo,
    color=["is_reference"],
    wspace=0.3,
    frameon=False,
    size=20,
    alpha=0.65,
    title=["Reference vs unimodal"],
)


# %% [code]
def add_reference_neighbor_fraction(
    adata,
    use_rep="X_univi",
    ref_key="is_reference",
    k=30,
    suffix="k30",
):
    """
    For each cell, compute the fraction of its k nearest neighbors that are from the reference.
    Adds:
      .obs[f'ref_neighbor_frac_{suffix}']
    """
    Z = np.asarray(adata.obsm[use_rep], dtype=np.float32)
    ref = adata.obs[ref_key].values.astype(bool)

    k_eff = int(min(max(int(k), 1), Z.shape[0] - 1))
    nn = NearestNeighbors(n_neighbors=k_eff + 1, metric="euclidean")
    nn.fit(Z)
    nbrs = nn.kneighbors(Z, return_distance=False)[:, 1:]  # drop self

    frac = ref[nbrs].mean(axis=1)
    col = f"ref_neighbor_frac_{suffix}"
    adata.obs[col] = frac
    return col


# %% [code]
# Compute and store
ref_frac_col = add_reference_neighbor_fraction(combo, use_rep="X_univi", k=30, suffix="k30")

# Look at unimodal only
unimodal_mask = ~combo.obs["is_reference"].values
print("Unimodal cells:", unimodal_mask.sum())
print("Reference-neighbor fraction (unimodal):")
print(combo.obs.loc[unimodal_mask, ref_frac_col].describe())


# %% [code]
def plot_ref_neighbor_fraction_by_label(
    combo,
    frac_col,
    label_col="celltype_harmonized_coarse",
    outpath=None,
):
    df = combo.obs.loc[~combo.obs["is_reference"], [frac_col, label_col, "dataset"]].copy()
    df = df[df[label_col].notna()]

    # order labels by median ref-neighbor fraction
    med = df.groupby(label_col)[frac_col].median().sort_values(ascending=False)
    order = med.index.tolist()
    plt.figure(figsize=(0.35 * len(order) + 4, 4.5))
    plt.boxplot(
        [df.loc[df[label_col] == lab, frac_col].values for lab in order],
        labels=order,
        showfliers=False,
    )
    plt.xticks(rotation=90)
    plt.ylabel("Reference-neighbor fraction")
    plt.title("Multiome reference support per harmonized coarse label")
    plt.tight_layout()
    if outpath is not None:
        plt.savefig(outpath, dpi=300, bbox_inches="tight")
    plt.show()

plot_ref_neighbor_fraction_by_label(combo, ref_frac_col)


# %% [code]
from sklearn.metrics import accuracy_score, f1_score, confusion_matrix

def knn_label_transfer_unpaired(Z_source, y_source, Z_target, y_target, k=15, metric="euclidean"):
    nn = NearestNeighbors(n_neighbors=k, metric=metric)
    nn.fit(Z_source)
    nbrs = nn.kneighbors(Z_target, return_distance=False)

    preds = []
    for inds in nbrs:
        votes = y_source[inds]
        vals, cnts = np.unique(votes, return_counts=True)
        preds.append(vals[np.argmax(cnts)])
    preds = np.asarray(preds, dtype=str)

    acc = float(accuracy_score(y_target, preds))
    macro_f1 = float(f1_score(y_target, preds, average="macro"))
    classes = np.unique(np.concatenate([y_source, y_target]))
    cm = confusion_matrix(y_target, preds, labels=classes)
    return preds, acc, macro_f1, cm, classes


# %% [code]
# Extract unimodal latents + labels
Z_uni_rna = uni_rna_pp.obsm["X_univi"]
y_uni_rna = uni_rna_pp.obs["celltype_harmonized_coarse"].astype(str).to_numpy()

Z_uni_atac = uni_atac_lsi.obsm["X_univi"]
y_uni_atac = uni_atac_lsi.obs["celltype_harmonized_coarse"].astype(str).to_numpy()


# %% [code]
# RNA -> ATAC label transfer
pred_atac_from_rna, acc_r2a, f1_r2a, cm_r2a, classes_r2a = knn_label_transfer_unpaired(
    Z_source=Z_uni_rna,
    y_source=y_uni_rna,
    Z_target=Z_uni_atac,
    y_target=y_uni_atac,
    k=10,
)
print("Unimodal RNA -> ATAC label transfer acc / macroF1:", acc_r2a, f1_r2a)


# %% [code]
# ATAC -> RNA label transfer
pred_rna_from_atac, acc_a2r, f1_a2r, cm_a2r, classes_a2r = knn_label_transfer_unpaired(
    Z_source=Z_uni_atac,
    y_source=y_uni_atac,
    Z_target=Z_uni_rna,
    y_target=y_uni_rna,
    k=10,
)
print("Unimodal ATAC -> RNA label transfer acc / macroF1:", acc_a2r, f1_a2r)


# %% [code]
import numpy as np
import pandas as pd
from sklearn.neighbors import NearestNeighbors
import matplotlib.pyplot as plt

def compute_ref_neighbor_fraction(
    adata,
    latent_key="X_univi",
    dataset_key="dataset",
    label_key="harmonized_label",
    ref_datasets=("multiome_rna", "multiome_atac"),
    uni_datasets=("ding_rna", "satpathy_atac"),
    k=30,
):
    """
    For each unimodal cell, compute the fraction of its k nearest neighbors
    (in latent space) that come from the reference (Multiome) datasets.
    """
    Z = np.asarray(adata.obsm[latent_key], dtype=np.float32)

    ds = adata.obs[dataset_key].astype(str)
    is_ref = ds.isin(ref_datasets).to_numpy()
    is_uni = ds.isin(uni_datasets).to_numpy()

    if is_ref.sum() == 0:
        raise ValueError("No reference cells found – check `ref_datasets`/`dataset_key`.")
    if is_uni.sum() == 0:
        raise ValueError("No unimodal cells found – check `uni_datasets`/`dataset_key`.")

    # fit neighbors on *all* cells
    nn = NearestNeighbors(n_neighbors=k + 1).fit(Z)
    dists, idx = nn.kneighbors(Z[is_uni])

    # drop self (first neighbor)
    idx_neighbors = idx[:, 1:]
    neigh_is_ref = is_ref[idx_neighbors]   # shape (n_uni, k)

    frac_ref = neigh_is_ref.mean(axis=1)

    df = pd.DataFrame(
        {
            "ref_neighbor_frac_k{}".format(k): frac_ref,
            dataset_key: ds.to_numpy()[is_uni],
            label_key: adata.obs[label_key].astype(str).to_numpy()[is_uni],
            "is_reference": False,
        },
        index=adata.obs_names[is_uni],
    )
    return df

# ---- run it ----
df_uni = compute_ref_neighbor_fraction(
    combo,
    latent_key="X_univi",
    dataset_key="dataset",
    label_key="celltype_harmonized_coarse",  # change if your column name differs
    ref_datasets=("multiome_rna", "multiome_atac"),
    uni_datasets=("ding_rna", "satpathy_atac"),
    k=30,
)

print("Unimodal cells:", df_uni.shape[0])
print(df_uni["ref_neighbor_frac_k30"].describe())


# %% [code]
# order labels by median reference support
med = df_uni.groupby("celltype_harmonized_coarse")["ref_neighbor_frac_k30"].median().sort_values(ascending=False)
order = med.index.tolist()

plt.figure(figsize=(12, 3.5))
df_uni.boxplot(
    column="ref_neighbor_frac_k30",
    by="celltype_harmonized_coarse",
    positions=range(len(order)),
    grid=False,
)
# the default pandas boxplot ignores `order`, so do it manually:
plt.clf()
plt.figure(figsize=(12, 3.5))
data = [df_uni.loc[df_uni["celltype_harmonized_coarse"] == lab, "ref_neighbor_frac_k30"] for lab in order]
plt.boxplot(data, showfliers=False)
plt.xticks(range(1, len(order) + 1), order, rotation=90)
plt.ylabel("Reference-neighbor fraction (k=30)")
plt.title("Multiome reference support per harmonized label")
plt.tight_layout()
plt.show()


# %% [code]
def plot_dataset_composition_by_label(
    adata,
    label_key="harmonized_label",
    dataset_key="dataset",
    min_cells=50,
):
    obs = adata.obs[[label_key, dataset_key]].copy()
    obs[label_key] = obs[label_key].astype(str)
    obs[dataset_key] = obs[dataset_key].astype(str)

    counts = (
        obs.groupby([label_key, dataset_key])
        .size()
        .unstack(fill_value=0)
    )

    # optionally drop very tiny labels
    counts = counts[counts.sum(axis=1) >= min_cells]

    frac = counts.div(counts.sum(axis=1), axis=0)

    plt.figure(figsize=(10, 4))
    bottom = np.zeros(frac.shape[0])
    x = np.arange(frac.shape[0])

    for ds in frac.columns:
        plt.bar(x, frac[ds].values, bottom=bottom, label=ds)
        bottom += frac[ds].values

    plt.xticks(x, frac.index, rotation=90)
    plt.ylabel("Fraction of cells")
    plt.title("Dataset composition per harmonized label")
    plt.legend(bbox_to_anchor=(1.05, 1), loc="upper left")
    plt.tight_layout()
    plt.show()

# ---- run it ----
plot_dataset_composition_by_label(
    combo,
    label_key="celltype_harmonized_coarse",
    dataset_key="dataset",
    min_cells=100,
)


# %% [code]
def compute_ref_vs_nn_ratio(
    adata,
    latent_key="X_univi",
    dataset_key="dataset",
    ref_datasets=("multiome_rna", "multiome_atac"),
    uni_datasets=("ding_rna", "satpathy_atac"),
):
    Z = np.asarray(adata.obsm[latent_key], dtype=np.float32)

    ds = adata.obs[dataset_key].astype(str)
    is_ref = ds.isin(ref_datasets).to_numpy()
    is_uni = ds.isin(uni_datasets).to_numpy()

    Z_ref = Z[is_ref]
    Z_uni = Z[is_uni]

    if Z_ref.shape[0] == 0 or Z_uni.shape[0] == 0:
        raise ValueError("Need both reference and unimodal cells.")

    # nearest reference for each unimodal cell
    nn_ref = NearestNeighbors(n_neighbors=1).fit(Z_ref)
    d_ref, _ = nn_ref.kneighbors(Z_uni)
    d_ref = d_ref[:, 0]

    # nearest overall neighbor (excluding self within unimodal pool)
    nn_uni = NearestNeighbors(n_neighbors=2).fit(Z_uni)
    d_all, idx_all = nn_uni.kneighbors(Z_uni)
    d_nn = d_all[:, 1]  # skip self

    ratio = d_ref / d_nn
    s = pd.Series(ratio, index=adata.obs_names[is_uni], name="ref_vs_nn_ratio")
    return s

# ---- run it ----
ratio = compute_ref_vs_nn_ratio(
    combo,
    latent_key="X_univi",
    dataset_key="dataset",
    ref_datasets=("multiome_rna", "multiome_atac"),
    uni_datasets=("ding_rna", "satpathy_atac"),
)

print(ratio.describe())
plt.figure(figsize=(4, 3))
plt.hist(ratio, bins=50)
plt.xlabel("dist(unimodal → nearest ref) / dist(unimodal → nearest unimodal)")
plt.ylabel("Count")
plt.title("Ref vs nearest-unimodal distance ratio")
plt.tight_layout()
plt.show()


# %% [code]
import numpy as np
from sklearn.neighbors import NearestNeighbors

def ref_fraction_null(Z, is_ref, k=30, n_perm=50, random_state=0):
    rng = np.random.default_rng(random_state)
    Z = np.asarray(Z, dtype=np.float32)
    is_ref = np.asarray(is_ref).astype(bool)

    nn = NearestNeighbors(n_neighbors=k+1).fit(Z)
    nbrs = nn.kneighbors(Z, return_distance=False)[:, 1:]

    frac_perm = []
    for _ in range(n_perm):
        perm = rng.permutation(len(is_ref))
        is_ref_perm = is_ref[perm]
        frac = (is_ref_perm[nbrs] & ~is_ref_perm[:, None]).mean(axis=1)
        frac_perm.append(frac[~is_ref_perm].mean())  # mean for unimodal cells
    return np.mean(frac_perm), np.std(frac_perm)

# Example usage (you’d fill these in):
# Z_all      = combo.obsm["X_univi"]
# is_ref_all = (combo.obs["dataset"] == "multiome").to_numpy()
# obs_mean   = combo.obs.loc[combo.obs["dataset"]!="multiome", "ref_neighbor_frac_k30"].mean()
# null_mean, null_std = ref_fraction_null(Z_all, is_ref_all, k=30, n_perm=100)
# print(obs_mean, null_mean, null_std)

uni_mask = combo.obs["dataset"] != "multiome"
df_uni = combo.obs.loc[uni_mask, ["dataset", "ref_neighbor_frac_k30"]]

print(df_uni.groupby("dataset")["ref_neighbor_frac_k30"].describe())

contact_rate = (df_uni["ref_neighbor_frac_k30"] > 0).mean()
print("Fraction of unimodal cells with ≥1 reference neighbor:", contact_rate)


# %% [code]
from sklearn.neighbors import NearestNeighbors
import numpy as np

Z = combo.obsm["X_univi"].astype(np.float32)

# Multiome bridge = reference
is_ref = combo.obs["dataset"].isin(["multiome_rna", "multiome_atac"]).to_numpy()

# Everything else = unimodal
is_uni = ~is_ref

Z_ref = Z[is_ref]
Z_uni = Z[is_uni]

print("Z_ref:", Z_ref.shape, "Z_uni:", Z_uni.shape)
print(combo.obs["dataset"].value_counts())


# %% [code]
print("Z_ref shape:", Z_ref.shape)
print("Z_uni shape:", Z_uni.shape)


# %% [code]
'''
if Z_ref.shape[0] == 0:
    # no reference cells here – define a convention and skip
    ratio = np.full(Z_uni.shape[0], np.nan)  # or np.inf
else:
    # nearest ref
    nn_ref = NearestNeighbors(n_neighbors=1).fit(Z_ref)
    d_ref, _ = nn_ref.kneighbors(Z_uni)
    d_ref = d_ref[:, 0]

    # nearest neighbor overall (excluding self, within unimodal set)
    nn_all = NearestNeighbors(n_neighbors=2).fit(Z_uni)
    d_all, idx_all = nn_all.kneighbors(Z_uni)
    d_nn = d_all[:, 1]

    ratio = d_ref / d_nn

print(pd.Series(ratio).describe())
'''

# %% [code]
# nearest ref
nn_ref = NearestNeighbors(n_neighbors=1).fit(Z_ref)
d_ref, _ = nn_ref.kneighbors(Z_uni)
d_ref = d_ref[:, 0]

# nearest neighbor overall (excluding self)
nn_all = NearestNeighbors(n_neighbors=2).fit(Z_uni)
d_all, idx_all = nn_all.kneighbors(Z_uni)
# first neighbor is self, second is true nearest other
d_nn = d_all[:, 1]

ratio = d_ref / d_nn
print(pd.Series(ratio).describe())


# %% [code]
# nearest neighbor overall (excluding self)
nn_all = NearestNeighbors(n_neighbors=2).fit(Z_uni)
d_all, idx_all = nn_all.kneighbors(Z_uni)
# first neighbor is self, second is true nearest other
d_nn = d_all[:, 1]

ratio = d_ref / d_nn
print(pd.Series(ratio).describe())


# %% [code]
ct_key = "celltype_harmonized"

# Quick composition table
pd.crosstab(combo.obs[ct_key], combo.obs["dataset"])

ref_datasets = ["multiome_rna", "multiome_atac"]
ref_labels = set(combo.obs.loc[combo.obs["dataset"].isin(ref_datasets), ct_key])
ding_labels = set(combo.obs.loc[combo.obs["dataset"] == "ding_rna", ct_key])

only_in_ding = sorted(ding_labels - ref_labels)
only_in_ref  = sorted(ref_labels - ding_labels)
shared       = sorted(ding_labels & ref_labels)

print("Labels only in Ding:", only_in_ding)
print("Labels only in Multiome:", only_in_ref)
print("Shared labels:", shared)


# %% [code]
import numpy as np
import pandas as pd
from sklearn.neighbors import KNeighborsClassifier
from sklearn.metrics import confusion_matrix, accuracy_score, f1_score


def knn_label_transfer_confusion_two_adata(
    adata_ref,
    adata_tgt,
    *,
    emb_key: str = "X_univi",
    ct_key: str = "celltype_harmonized",
    k: int = 15,
    drop_unknown: bool = True,
    unknown_labels=("Unknown", "Unassigned"),
    exclude_labels=None,  # e.g. ("Other",)
):
    """
    kNN label transfer from adata_ref -> adata_tgt in a shared embedding.

    Parameters
    ----------
    adata_ref : AnnData
        Reference cells (e.g. multiome_rna + multiome_atac).
    adata_tgt : AnnData
        Target cells (e.g. ding_rna + satpathy_atac).
    emb_key : str
        .obsm key for the shared embedding.
    ct_key : str
        .obs key with harmonized labels (fine or coarse).
    k : int
        Number of neighbors for kNN.
    drop_unknown : bool
        If True, drop `unknown_labels` from the *target* side.
    unknown_labels : tuple of str
        Labels in `ct_key` to consider as unknown/unassigned.
    exclude_labels : sequence of str or None
        Additional labels in `ct_key` to exclude from BOTH
        reference and target (e.g. ["Other"] for coarse types).

    Returns
    -------
    dict with keys:
        acc, macro_f1, classes, cm, cm_df, y_true, y_pred
    """
    # ------------- extract embedding + labels -------------
    Z_ref = np.asarray(adata_ref.obsm[emb_key], dtype=np.float32)
    Z_tgt = np.asarray(adata_tgt.obsm[emb_key], dtype=np.float32)

    y_ref = adata_ref.obs[ct_key].astype(str).to_numpy()
    y_tgt = adata_tgt.obs[ct_key].astype(str).to_numpy()

    print(f"Reference cells: {Z_ref.shape[0]} | Target cells: {Z_tgt.shape[0]}")

    if Z_ref.shape[0] == 0:
        raise ValueError("No reference cells found – check your subsetting / dataset names.")

    # ------------- optionally drop Unknown/Unassigned from TARGET -------------
    if drop_unknown and unknown_labels:
        mask_tgt = ~np.isin(y_tgt, unknown_labels)
        Z_tgt = Z_tgt[mask_tgt]
        y_tgt = y_tgt[mask_tgt]
        print(f"After dropping unknowns: target cells = {Z_tgt.shape[0]}")

    # ------------- optionally drop additional labels from BOTH sides -------------
    if exclude_labels is not None:
        exclude_labels = np.asarray(list(exclude_labels), dtype=str)

        mask_ref = ~np.isin(y_ref, exclude_labels)
        mask_tgt = ~np.isin(y_tgt, exclude_labels)

        Z_ref = Z_ref[mask_ref]
        y_ref = y_ref[mask_ref]

        Z_tgt = Z_tgt[mask_tgt]
        y_tgt = y_tgt[mask_tgt]

        print(
            f"After excluding labels {list(exclude_labels)}: "
            f"ref cells = {Z_ref.shape[0]}, tgt cells = {Z_tgt.shape[0]}"
        )

    if Z_ref.shape[0] == 0 or Z_tgt.shape[0] == 0:
        raise ValueError("No cells left after filtering – relax your filters or check labels.")

    # ------------- fit kNN on reference, predict on target -------------
    knn = KNeighborsClassifier(n_neighbors=k, weights="distance")
    knn.fit(Z_ref, y_ref)
    y_pred = knn.predict(Z_tgt)

    # ------------- metrics -------------
    classes = np.unique(np.concatenate([y_ref, y_tgt]))
    cm = confusion_matrix(y_tgt, y_pred, labels=classes)
    acc = accuracy_score(y_tgt, y_pred)
    macro_f1 = f1_score(y_tgt, y_pred, average="macro")

    cm_df = pd.DataFrame(cm, index=classes, columns=classes)

    return {
        "acc": acc,
        "macro_f1": macro_f1,
        "classes": classes,
        "cm": cm,
        "cm_df": cm_df,
        "y_true": y_tgt,
        "y_pred": y_pred,
    }


# %% [code]
# build separate ref / target AnnData from the *full* combo
#ref_mask = combo.obs["dataset"].isin(["multiome_rna", "multiome_atac"])
#tgt_mask = combo.obs["dataset"].isin(["ding_rna", "satpathy_atac"])
ref_mask = combo.obs["dataset"].isin(["ding_rna"])
tgt_mask = combo.obs["dataset"].isin(["satpathy_atac"])

adata_ref = combo[ref_mask].copy()
adata_tgt = combo[tgt_mask].copy()

'''
res = knn_label_transfer_confusion_two_adata(
    adata_ref,
    adata_tgt,
    emb_key="X_univi",
    ct_key="celltype_harmonized_coarse",
    k=15,
    drop_unknown=True,
    unknown_labels=("Unknown", "Unassigned"),
)
'''
res = knn_label_transfer_confusion_two_adata(
    adata_ref=adata_ref,
    adata_tgt=adata_tgt,
    emb_key="X_univi",
    ct_key="celltype_harmonized_coarse",
    k=15,
    exclude_labels=("Other", "Unknown", "Unassigned", "Megakaryocyte"),  # drop those coarse “other” cells
)


print("Acc:", res["acc"])
print("Macro F1:", res["macro_f1"])
display(res["cm_df"])


# %% [code]
import numpy as np
import matplotlib.pyplot as plt


def plot_knn_confusion_heatmap(
    res,
    *,
    normalize: str | None = "row",
    title: str | None = None,
    figsize=(6, 5),
    show_values: bool = True,
    value_fmt: str | None = None,
    rotation_x: int = 45,
    rotation_y: int = 0,
    text_color: str = "white",
):
    """
    Plot a confusion-matrix heatmap from the output of
    `knn_label_transfer_confusion_two_adata`.
    """
    cm = np.asarray(res["cm"], dtype=float)
    classes = np.asarray(res["classes"], dtype=str)

    # ------------- normalization -------------
    if normalize is not None:
        if normalize == "row":
            row_sums = cm.sum(axis=1, keepdims=True)
            row_sums[row_sums == 0] = 1.0
            cm_plot = cm / row_sums
        elif normalize == "col":
            col_sums = cm.sum(axis=0, keepdims=True)
            col_sums[col_sums == 0] = 1.0
            cm_plot = cm / col_sums
        else:
            raise ValueError("normalize must be one of {'row', 'col', None}")
    else:
        cm_plot = cm

    # default annotation format
    if value_fmt is None:
        value_fmt = ".2f" if normalize is not None else "d"

    nrows, ncols = cm_plot.shape

    # ------------- plotting (pcolormesh to avoid grid lines) -------------
    fig, ax = plt.subplots(figsize=figsize)

    mesh = ax.pcolormesh(
        cm_plot,
        edgecolors="face",   # no visible edges between cells
        linewidth=0.0,
        shading="auto",
    )
    mesh.set_rasterized(True)  # avoids hairlines in vector backends

    # ticks / labels (note: pcolormesh cell centers at i+0.5)
    ax.set_xticks(np.arange(ncols) + 0.5)
    ax.set_yticks(np.arange(nrows) + 0.5)
    ax.set_xticklabels(classes, rotation=rotation_x, ha="right")
    ax.set_yticklabels(classes, rotation=rotation_y)

    # keep [0, ncols] x [0, nrows] and put row 0 at top
    ax.set_xlim(0, ncols)
    ax.set_ylim(nrows, 0)

    ax.set_xlabel("Predicted label")
    ax.set_ylabel("True label")

    if title is not None:
        ax.set_title(title)

    # remove any axis grid
    ax.grid(False)

    # colorbar
    cbar = fig.colorbar(mesh, ax=ax)
    cbar.set_label("Fraction" if normalize is not None else "Count")

    # ------------- annotations -------------
    if show_values:
        annot_mat = cm_plot if normalize is not None else cm

        for i in range(nrows):
            for j in range(ncols):
                val = annot_mat[i, j]
                ax.text(
                    j + 0.5,
                    i + 0.5,
                    format(val, value_fmt),
                    ha="center",
                    va="center",
                    color=text_color,
                    fontsize=8,
                )

    fig.tight_layout()
    return fig, ax


# %% [code]
fig, ax = plot_knn_confusion_heatmap(
    res,
    normalize="row",  # show per-true-class fractions
    title="Ding → Satpathy label transfer (coarse, k=15)",
)
# fig.savefig("ding_to_satpathy_confusion_coarse.png", dpi=300)


# %% [code]
# build separate ref / target AnnData from the *full* combo
#ref_mask = combo.obs["dataset"].isin(["multiome_rna", "multiome_atac"])
#tgt_mask = combo.obs["dataset"].isin(["ding_rna", "satpathy_atac"])
ref_mask = combo.obs["dataset"].isin(["satpathy_atac"])
tgt_mask = combo.obs["dataset"].isin(["ding_rna"])

adata_ref = combo[ref_mask].copy()
adata_tgt = combo[tgt_mask].copy()

'''
res = knn_label_transfer_confusion_two_adata(
    adata_ref,
    adata_tgt,
    emb_key="X_univi",
    ct_key="celltype_harmonized_coarse",
    k=15,
    drop_unknown=True,
    unknown_labels=("Unknown", "Unassigned"),
)
'''
res_atac = knn_label_transfer_confusion_two_adata(
    adata_ref=adata_ref,
    adata_tgt=adata_tgt,
    emb_key="X_univi",
    ct_key="celltype_harmonized_coarse",
    k=15,
    exclude_labels=("Other", "Unknown", "Unassigned", "Megakaryocyte"),  # drop those coarse “other” cells
)


print("Acc:", res_atac["acc"])
print("Macro F1:", res_atac["macro_f1"])
display(res_atac["cm_df"])


# %% [code]
fig, ax = plot_knn_confusion_heatmap(
    res_atac,
    normalize="row",  # show per-true-class fractions
    title="Satpathy → Ding label transfer (coarse, k=15)",
)
# fig.savefig("ding_to_satpathy_confusion_coarse.png", dpi=300)


# %% [code]
# build separate ref / target AnnData from the *full* combo
#ref_mask = combo.obs["dataset"].isin(["multiome_rna", "multiome_atac"])
#tgt_mask = combo.obs["dataset"].isin(["ding_rna", "satpathy_atac"])
ref_mask = combo.obs["dataset"].isin(["satpathy_atac"])
tgt_mask = combo.obs["dataset"].isin(["ding_rna"])

adata_ref = combo[ref_mask].copy()
adata_tgt = combo[tgt_mask].copy()

res_fine_celltype_atac = knn_label_transfer_confusion_two_adata(
    adata_ref=adata_ref,
    adata_tgt=adata_tgt,
    emb_key="X_univi",
    ct_key="celltype_harmonized",
    k=15,
    #exclude_labels=("Other", "Unknown", "Unassigned", "Megakaryocyte"),  # drop those coarse “other” cells
)


# %% [code]
print("Acc:", res_fine_celltype_atac["acc"])
print("Macro F1:", res_fine_celltype_atac["macro_f1"])
display(res_fine_celltype_atac["cm_df"])
fig, ax = plot_knn_confusion_heatmap(
    res_fine_celltype_atac,
    normalize="row",  # show per-true-class fractions
    title="Satpathy → Ding label transfer (fine, k=15)",
)


# %% [code]
# build separate ref / target AnnData from the *full* combo
#ref_mask = combo.obs["dataset"].isin(["multiome_rna", "multiome_atac"])
#tgt_mask = combo.obs["dataset"].isin(["ding_rna", "satpathy_atac"])
ref_mask = combo.obs["dataset"].isin(["ding_rna"])
tgt_mask = combo.obs["dataset"].isin(["satpathy_atac"])

adata_ref = combo[ref_mask].copy()
adata_tgt = combo[tgt_mask].copy()

res_fine_celltype_rna = knn_label_transfer_confusion_two_adata(
    adata_ref=adata_ref,
    adata_tgt=adata_tgt,
    emb_key="X_univi",
    ct_key="celltype_harmonized",
    k=15,
    #exclude_labels=("Other", "Unknown", "Unassigned", "Megakaryocyte"),  # drop those coarse “other” cells
)


# %% [code]
print("Acc:", res_fine_celltype_rna["acc"])
print("Macro F1:", res_fine_celltype_rna["macro_f1"])
display(res_fine_celltype_rna["cm_df"])
fig, ax = plot_knn_confusion_heatmap(
    res_fine_celltype_rna,
    normalize="row",  # show per-true-class fractions
    title="Ding → Satpathy label transfer (fine, k=15)",
)
# fig.savefig("ding_to_satpathy_confusion_coarse.png", dpi=300)


# %% [code]
import numpy as np
import pandas as pd
from sklearn.neighbors import NearestNeighbors

Z = combo.obsm["X_univi"].astype(np.float32)

# Multiome bridge = reference
ref_datasets = ["multiome_rna", "multiome_atac"]
is_ref = combo.obs["dataset"].isin(ref_datasets).to_numpy(bool)
is_uni = ~is_ref

combo.obs["is_reference"] = is_ref
combo.obs["is_unimodal"] = is_uni

k = 30

# kNN on ALL cells in latent
nn = NearestNeighbors(n_neighbors=k + 1).fit(Z)
# drop the self-neighbor in column 0
nbrs = nn.kneighbors(Z, return_distance=False)[:, 1:]

# for each cell: which neighbors are reference, while the focal cell is unimodal?
nbr_is_ref = is_ref[nbrs] & is_uni[:, None]
ref_neighbor_frac = nbr_is_ref.mean(axis=1)

combo.obs["ref_neighbor_frac_k30"] = ref_neighbor_frac

def ref_fraction_null(Z, is_ref, k=30, n_perm=50, random_state=0):
    """
    Shuffle the is_ref labels and recompute the mean ref-neighbor fraction
    for unimodal cells, to get a null mean/std.
    """
    rng = np.random.default_rng(random_state)
    Z = np.asarray(Z, dtype=np.float32)
    is_ref = np.asarray(is_ref).astype(bool)

    nn = NearestNeighbors(n_neighbors=k + 1).fit(Z)
    nbrs = nn.kneighbors(Z, return_distance=False)[:, 1:]

    frac_perm = []
    for _ in range(n_perm):
        perm = rng.permutation(len(is_ref))
        is_ref_perm = is_ref[perm]

        # neighbors that are reference while focal cell is unimodal
        nbr_is_ref_perm = is_ref_perm[nbrs] & ~is_ref_perm[:, None]
        frac = nbr_is_ref_perm.mean(axis=1)

        # mean only over unimodal (non-reference) cells
        frac_perm.append(frac[~is_ref_perm].mean())

    return float(np.mean(frac_perm)), float(np.std(frac_perm))

obs_mean = combo.obs.loc[combo.obs["is_unimodal"], "ref_neighbor_frac_k30"].mean()
null_mean, null_std = ref_fraction_null(Z, is_ref, k=30, n_perm=100)
print("Observed unimodal mean:", obs_mean)
print("Null mean ± sd:", null_mean, null_std)

uni_mask = combo.obs["is_unimodal"]

df_uni = combo.obs.loc[
    uni_mask,
    ["dataset", "celltype_harmonized", "ref_neighbor_frac_k30", "is_reference"],
]

print(df_uni.groupby("dataset")["ref_neighbor_frac_k30"].describe())
print()
print(df_uni.groupby("celltype_harmonized")["ref_neighbor_frac_k30"].describe())

# global reference fraction (over ALL cells, not only unimodal)
frac_ref_global = combo.obs["is_reference"].mean()
print("Global reference fraction (all cells):", frac_ref_global)

# fraction of unimodal cells that have at least 1 reference neighbor
contact_rate = (df_uni["ref_neighbor_frac_k30"] > 0).mean()
print("Fraction of unimodal cells with ≥1 reference neighbor:", contact_rate)

from sklearn.neighbors import NearestNeighbors

Z_ref = Z[is_ref]
Z_uni = Z[is_uni]

print("Z_ref:", Z_ref.shape, "Z_uni:", Z_uni.shape)
print(combo.obs["dataset"].value_counts())

# nearest ref to each unimodal cell
nn_ref = NearestNeighbors(n_neighbors=1).fit(Z_ref)
d_ref, _ = nn_ref.kneighbors(Z_uni)
d_ref = d_ref[:, 0]

# nearest unimodal neighbor (excluding self)
nn_all = NearestNeighbors(n_neighbors=2).fit(Z_uni)
d_all, idx_all = nn_all.kneighbors(Z_uni)
d_nn = d_all[:, 1]

ratio = d_ref / d_nn
print(pd.Series(ratio).describe())


# %% [code]
print(uni_rna_pp)
print(set(uni_rna_pp.obs['tech']))


# %% [code]
# Subset combo to Ding cells
ding = combo[combo.obs["dataset"] == "ding_rna"].copy()

sc.pl.umap(
    ding,
    color="tech",
    size=20,
    alpha=0.65,
    legend_loc="right margin",  # or "on data"
    title="Ding RNA cells colored by sequencing technology",
)


# %% [code]
sc.pl.umap(
    ding,
    color="celltype_harmonized_coarse",
    size=20,
    alpha=0.65,
    legend_loc="right margin",  # or "on data"
    title="Ding RNA cells colored by sequencing technology",
)


# %% [code]
sc.pl.umap(
    ding,
    color="celltype_harmonized",
    size=20,
    alpha=0.65,
    legend_loc="right margin",  # or "on data"
    title="Ding RNA cells colored by sequencing technology",
)


# %% [code]
# Subset combo to Satpathy cells
satpathy = combo[combo.obs["dataset"] == "satpathy_atac"].copy()

sc.pl.umap(
    satpathy,
    color="celltype_harmonized_coarse",
    size=20,
    alpha=0.65,
    legend_loc="right margin",  # or "on data"
    title="Ding RNA cells colored by sequencing technology",
)


# %% [code]
sc.pl.umap(
    satpathy,
    color="celltype_harmonized",
    size=20,
    alpha=0.65,
    legend_loc="right margin",  # or "on data"
    title="Ding RNA cells colored by sequencing technology",
)


# %% [code]
# df_uni already has ref_neighbor_frac_k30 for unimodal cells
df_uni = combo.obs.loc[combo.obs["dataset"].isin(["ding_rna", "satpathy_atac"])].copy()

print(
    df_uni.groupby(["dataset", "tech"])["ref_neighbor_frac_k30"].describe()
)


# %% [code]
from sklearn.neighbors import NearestNeighbors
import numpy as np
import pandas as pd

ding = combo[combo.obs["dataset"] == "ding_rna"].copy()
Z = ding.obsm["X_univi"]
tech = ding.obs["tech"].astype(str).to_numpy()

nn = NearestNeighbors(n_neighbors=15).fit(Z)
idx = nn.kneighbors(return_distance=False)

# fraction of neighbors with same tech
same = (tech[idx] == tech[:, None]).mean(axis=1)
print("Median same-tech neighbor fraction:", np.median(same))

print(
    pd.Series(same, index=ding.obs_names)
      .groupby(ding.obs["tech"])
      .median()
)


# %% [code]


# %% [markdown]
### Use celltype classification head to fine-tune the bridge-trained model using the unimodal RNA data

# %% [code]
satpathy_atac_pp = uni_atac_lsi.copy()


# %% [code]
print(set(satpathy_atac_pp.obs['celltype_harmonized']))
print(set(uni_rna_pp.obs['celltype_harmonized']))


# %% [code]
# ============================================================
# UniVI: fine-tune bridge checkpoint with a shared celltype head
#   Supervision sources:
#     - labeled Ding RNA      (uni_rna_pp)
#     - labeled Satpathy ATAC (satpathy_atac_pp)
#
# Also includes:
#   - head predictions + confusion matrices (RNA+ATAC)
#   - encode new latent + concat + UMAP
#   - kNN label transfer eval in new latent
#   - per-dataset head eval (feature-aligned)
#   - paired multiome annotation (fused head probs)
# ============================================================

import copy
import numpy as np
import pandas as pd
import torch
from torch.utils.data import DataLoader
from sklearn.model_selection import StratifiedShuffleSplit
from sklearn.metrics import accuracy_score, f1_score, confusion_matrix

import anndata as ad
import scanpy as sc

from univi.config import UniVIConfig, ModalityConfig, ClassHeadConfig
from univi.models import UniVIMultiModalVAE
from univi.data import MultiModalDataset
from univi.evaluation import encode_adata


# -----------------------------
# USER CHOICES
# -----------------------------
#label_col = "celltype_harmonized_coarse"
label_col = "celltype_harmonized"
head_name = "celltype_higher_res"
exclude_labels_train = {"Unknown", "Unassigned", "General PBMC", "Megakaryocyte"}   # training-time exclusions

# supervised loss weights (balance RNA vs ATAC)
lambda_cls_rna  = 1.00
lambda_cls_atac = 1.00

'''
# Try:
lambda_gen = 2.0–10.0
lambda_cls_rna = lambda_cls_atac = 0.1–0.5
'''

# protect bridge generative objective
lambda_gen     = 2.00
lambda_uni_gen = 0.00    # optional: unimodal RNA gen term on RNA supervised steps (0..0.25-ish)
lambda_anchor  = 0.00    # optional: L2 anchor to ckpt (excluding head), ~1e-6..1e-4

# training regime
METHOD = "head_then_joint"   # "head_only" | "joint_mixed" | "head_then_joint"
n_epochs      = 1000
warmup_epochs = 300

# batch sizes
batch_size_bridge = 256
batch_size_lab_tr = 256
batch_size_lab_va = 512

# learning rates (param groups)
lr_backbone = 1e-5
lr_head     = 3e-4
weight_decay_backbone = 0.0
weight_decay_head     = 1e-4

# encoding / plotting
latent_choice = "moe_mean"   # "modality_mean" or "moe_mean"
out_key       = "X_univi_ft"
umap_n_neighbors = 30


# -----------------------------
# REQUIRED INPUTS
# -----------------------------
assert "uni_rna_pp" in globals(), "Need uni_rna_pp (Ding RNA preprocessed to model RNA input space)."
assert "satpathy_atac_pp" in globals(), "Need satpathy_atac_pp (Satpathy ATAC preprocessed to model ATAC input space)."
assert "rna_train_pp" in globals() and "atac_train_lsi" in globals(), "Need rna_train_pp + atac_train_lsi (bridge paired train)."
assert "output_dir" in globals() and "out_file" in globals(), "Need output_dir + out_file (bridge checkpoint path pieces)."
assert "device" in globals(), "Need device (e.g. 'cuda' or torch.device)."


# %% [code]
# ============================================================
# 1) Load bridge checkpoint + rebuild config
# ============================================================
ckpt = torch.load(output_dir + out_file, map_location=device)

cfg_dict = ckpt["univi_cfg"]
try:
    from omegaconf import DictConfig, OmegaConf
    if isinstance(cfg_dict, DictConfig):
        cfg_dict = OmegaConf.to_container(cfg_dict, resolve=True)
except ImportError:
    pass

modalities = [ModalityConfig(**m) for m in cfg_dict["modalities"]]
cfg_dict = {**cfg_dict, "modalities": modalities}
univi_cfg = UniVIConfig(**cfg_dict)


# %% [code]
# ============================================================
# 2) Build shared label vocab from UNION of labeled RNA + labeled ATAC
# ============================================================
def _labeled_mask_and_series(adata, label_col, exclude_labels):
    if label_col not in adata.obs:
        raise KeyError(f"adata.obs[{label_col!r}] not found")
    y = adata.obs[label_col].astype(str)
    m = (~y.str.startswith("unlabeled_")) & (~y.isin(list(exclude_labels)))
    return m.to_numpy(), y

mask_rna,  y_rna  = _labeled_mask_and_series(uni_rna_pp,       label_col, exclude_labels_train)
mask_atac, y_atac = _labeled_mask_and_series(satpathy_atac_pp, label_col, exclude_labels_train)

y_rna_lab  = y_rna[mask_rna]
y_atac_lab = y_atac[mask_atac]

classes = sorted(pd.unique(pd.concat([y_rna_lab, y_atac_lab], axis=0)))
label_to_id = {c: i for i, c in enumerate(classes)}
id_to_label = {i: c for c, i in label_to_id.items()}
n_classes = len(classes)
print(f"Supervised head classes (n={n_classes}):", classes)

def _make_codes(adata, y_raw, mask, label_to_id, head_name, ignore_index=-1):
    """Store integer codes in adata.obs[f'{head_name}_code']."""
    codes = np.full(adata.n_obs, ignore_index, dtype=np.int64)
    mask = np.asarray(mask, dtype=bool)
    if mask.sum() == 0:
        adata.obs[f"{head_name}_code"] = codes
        return codes

    idx = np.where(mask)[0]
    y = y_raw.iloc[idx].astype(str)
    mapped = y.map(label_to_id)          # float with NaN for unknown
    keep = mapped.notna().to_numpy()
    codes[idx[keep]] = mapped.to_numpy(np.int64)[keep]
    adata.obs[f"{head_name}_code"] = codes
    return codes

codes_rna  = _make_codes(uni_rna_pp,       y_rna,  mask_rna,  label_to_id, head_name)
codes_atac = _make_codes(satpathy_atac_pp, y_atac, mask_atac, label_to_id, head_name)


# %% [code]
# ============================================================
# 3) Attach NON-adversarial head + build model + load weights
# ============================================================
univi_cfg.class_heads = [
    ClassHeadConfig(
        name=head_name,
        n_classes=int(n_classes),
        loss_weight=1.0,
        ignore_index=-1,
        from_mu=True,
        warmup=0,
        adversarial=False,
    )
]
univi_cfg.validate()

model = UniVIMultiModalVAE(
    univi_cfg,
    loss_mode="v1", # or lite - but original model was trained with v1 so we'll stick with that
    v1_recon="avg",
    normalize_v1_terms=True,
).to(device)

missing, unexpected = model.load_state_dict(ckpt["state_dict"], strict=False)
print("Missing keys (expected new head params):", missing[:8], "...")
print("Unexpected keys:", unexpected[:8], "...")

# anchor target = post-load state dict (includes loaded backbone + fresh head)
base_state_dict = copy.deepcopy(model.state_dict())


# %% [code]
# ============================================================
# 4) Build loaders (RNA supervised, ATAC supervised, bridge paired)
# ============================================================
def _split_stratified(idx_all, y_codes, test_size=0.10, seed=42):
    sss = StratifiedShuffleSplit(n_splits=1, test_size=test_size, random_state=seed)
    tr_rel, va_rel = next(sss.split(idx_all, y_codes))
    return idx_all[tr_rel], idx_all[va_rel]

def _make_supervised_loaders(adata, codes, modality, batch_tr, batch_va):
    idx_lab = np.where(codes >= 0)[0]
    if len(idx_lab) == 0:
        raise ValueError(f"No labeled cells found for modality={modality!r}.")
    y_codes = codes[idx_lab]
    idx_tr, idx_va = _split_stratified(idx_lab, y_codes)

    ad_tr = adata[idx_tr].copy()
    ad_va = adata[idx_va].copy()
    y_tr = torch.from_numpy(codes[idx_tr].astype(np.int64))
    y_va = torch.from_numpy(codes[idx_va].astype(np.int64))

    ds_tr = MultiModalDataset(
        adata_dict={modality: ad_tr},
        X_key="X",
        labels={head_name: y_tr},
        paired=False,
        device=None,
    )
    ds_va = MultiModalDataset(
        adata_dict={modality: ad_va},
        X_key="X",
        labels={head_name: y_va},
        paired=False,
        device=None,
    )

    ld_tr = DataLoader(ds_tr, batch_size=batch_tr, shuffle=True,  num_workers=0)
    ld_va = DataLoader(ds_va, batch_size=batch_va, shuffle=False, num_workers=0)
    return ds_tr, ds_va, ld_tr, ld_va

train_ds_rna, val_ds_rna, cls_train_loader_rna, cls_val_loader_rna = _make_supervised_loaders(
    uni_rna_pp, codes_rna, "rna", batch_size_lab_tr, batch_size_lab_va
)
train_ds_atac, val_ds_atac, cls_train_loader_atac, cls_val_loader_atac = _make_supervised_loaders(
    satpathy_atac_pp, codes_atac, "atac", batch_size_lab_tr, batch_size_lab_va
)
print("Labeled RNA train/val:", len(train_ds_rna), len(val_ds_rna))
print("Labeled ATAC train/val:", len(train_ds_atac), len(val_ds_atac))

bridge_ds = MultiModalDataset(
    adata_dict={"rna": rna_train_pp, "atac": atac_train_lsi},
    X_key="X",
    paired=True,
    device=None,
)
bridge_loader = DataLoader(bridge_ds, batch_size=batch_size_bridge, shuffle=True, num_workers=0)


# %% [code]
# ============================================================
# 5) Training helpers (no missing functions / no global iterators)
# ============================================================
def _unwrap_x(batch):
    """MultiModalDataset may yield x or (x,y). Return x dict."""
    if isinstance(batch, (tuple, list)) and len(batch) == 2:
        return batch[0]
    return batch

def _cycle(loader):
    while True:
        for batch in loader:
            yield batch

def set_trainable(model, mode: str):
    if mode == "all":
        for _, p in model.named_parameters():
            p.requires_grad = True
    elif mode == "head_only":
        for n, p in model.named_parameters():
            p.requires_grad = n.startswith(f"class_heads.{head_name}.")
    else:
        raise ValueError(f"Unknown mode: {mode}")

def anchor_l2_penalty(model, base_state_dict, exclude_prefix=("class_heads.",)):
    loss = 0.0
    for k, v in model.state_dict().items():
        if any(k.startswith(pref) for pref in exclude_prefix):
            continue
        v0 = base_state_dict.get(k, None)
        if v0 is None:
            continue
        if not torch.is_floating_point(v) or not torch.is_floating_point(v0):
            continue
        loss = loss + torch.sum((v - v0.to(device)) ** 2)
    return loss

def build_optimizers(model):
    head_params, backbone_params = [], []
    for n, p in model.named_parameters():
        if not p.requires_grad:
            continue
        if n.startswith(f"class_heads.{head_name}."):
            head_params.append(p)
        else:
            backbone_params.append(p)

    opt_backbone = (
        torch.optim.AdamW(backbone_params, lr=lr_backbone, weight_decay=weight_decay_backbone)
        if backbone_params else None
    )
    opt_head = (
        torch.optim.AdamW(head_params, lr=lr_head, weight_decay=weight_decay_head)
        if head_params else None
    )
    return opt_backbone, opt_head

@torch.no_grad()
def eval_head_on_loader(model, loader):
    model.eval()
    y_true_all, y_pred_all = [], []
    for x_batch, y_batch in loader:
        x_batch = {k: v.to(device) for k, v in x_batch.items()}
        y_true = y_batch[head_name].to(device).view(-1)

        logits = model.predict_heads(
            x_batch,
            return_probs=False,
            use_mean=True,
            inject_label_expert=False,
        )[head_name]
        y_pred = logits.argmax(dim=-1)

        y_true_all.append(y_true.cpu().numpy())
        y_pred_all.append(y_pred.cpu().numpy())

    y_true_all = np.concatenate(y_true_all)
    y_pred_all = np.concatenate(y_pred_all)
    return float(accuracy_score(y_true_all, y_pred_all)), float(f1_score(y_true_all, y_pred_all, average="macro"))

def eval_both(model):
    rna_acc, rna_f1   = eval_head_on_loader(model, cls_val_loader_rna)
    atac_acc, atac_f1 = eval_head_on_loader(model, cls_val_loader_atac)
    mean_f1 = 0.5 * (rna_f1 + atac_f1)
    return rna_acc, rna_f1, atac_acc, atac_f1, mean_f1

def supervised_step_one_batch(model, opt_backbone, opt_head, epoch, which, it_rna, it_atac):
    """One supervised optimizer step on exactly ONE batch (rna or atac)."""
    model.train()

    if which == "rna":
        batch = next(it_rna)
        w = lambda_cls_rna
    elif which == "atac":
        batch = next(it_atac)
        w = lambda_cls_atac
    else:
        raise ValueError(which)

    x_batch, y_batch = batch
    x_batch = {k: v.to(device) for k, v in x_batch.items()}
    y_batch = {k: v.to(device) for k, v in y_batch.items()}

    out = model(x_batch, y=y_batch, epoch=epoch)
    loss = w * out["head_losses"][head_name]

    # optional: unimodal RNA gen term
    if lambda_uni_gen > 0 and which == "rna":
        out_gen = model(x_batch, epoch=epoch)
        loss = loss + (lambda_uni_gen * out_gen["loss"])

    if lambda_anchor > 0:
        loss = loss + (lambda_anchor * anchor_l2_penalty(model, base_state_dict))

    if opt_backbone is not None: opt_backbone.zero_grad(set_to_none=True)
    if opt_head is not None:     opt_head.zero_grad(set_to_none=True)
    loss.backward()
    if opt_backbone is not None: opt_backbone.step()
    if opt_head is not None:     opt_head.step()

    return float(loss.detach().cpu())

def supervised_epoch_alternating(model, opt_backbone, opt_head, epoch):
    """Run a supervised epoch by alternating rna/atac for 2*max(len(loader))."""
    it_rna  = _cycle(cls_train_loader_rna)
    it_atac = _cycle(cls_train_loader_atac)

    steps = max(len(cls_train_loader_rna), len(cls_train_loader_atac))
    losses = []
    for s in range(steps * 2):
        which = "rna" if (s % 2 == 0) else "atac"
        losses.append(supervised_step_one_batch(model, opt_backbone, opt_head, epoch, which, it_rna, it_atac))
    return float(np.mean(losses))

def train_head_only(model):
    set_trainable(model, "head_only")
    _, opt_head = build_optimizers(model)
    assert opt_head is not None, "No head params?"

    best = (-np.inf, None)
    for epoch in range(n_epochs):
        loss = supervised_epoch_alternating(model, opt_backbone=None, opt_head=opt_head, epoch=epoch)
        rna_acc, rna_f1, atac_acc, atac_f1, mean_f1 = eval_both(model)

        if mean_f1 > best[0]:
            best = (mean_f1, {k: v.detach().cpu().clone() for k, v in model.state_dict().items()})

        print(
            f"[epoch {epoch:03d}] head_only_loss={loss:.4f} | "
            f"RNA acc/F1={rna_acc:.3f}/{rna_f1:.3f} | "
            f"ATAC acc/F1={atac_acc:.3f}/{atac_f1:.3f} | meanF1={mean_f1:.3f}"
        )

    if best[1] is not None:
        model.load_state_dict(best[1])
        print("Restored best head-only with meanF1 =", best[0])

def train_joint_mixed(model):
    set_trainable(model, "all")
    opt_backbone, opt_head = build_optimizers(model)
    if opt_backbone is None and opt_head is None:
        raise RuntimeError("No trainable parameters found.")

    best = (-np.inf, None)

    it_bridge = _cycle(bridge_loader)
    it_rna    = _cycle(cls_train_loader_rna)
    it_atac   = _cycle(cls_train_loader_atac)

    steps = max(len(bridge_loader), len(cls_train_loader_rna), len(cls_train_loader_atac))

    for epoch in range(n_epochs):
        model.train()
        gen_losses, sup_losses = [], []

        for step in range(steps):
            # --- bridge gen: ONE batch ---
            batch = next(it_bridge)
            x_batch = _unwrap_x(batch)
            x_batch = {k: v.to(device) for k, v in x_batch.items()}

            out = model(x_batch, epoch=epoch)
            loss = lambda_gen * out["loss"]
            if lambda_anchor > 0:
                loss = loss + (lambda_anchor * anchor_l2_penalty(model, base_state_dict))

            if opt_backbone is not None: opt_backbone.zero_grad(set_to_none=True)
            if opt_head is not None:     opt_head.zero_grad(set_to_none=True)
            loss.backward()
            if opt_backbone is not None: opt_backbone.step()
            if opt_head is not None:     opt_head.step()

            gen_losses.append(float(loss.detach().cpu()))

            # --- supervised: ONE batch (alternate rna/atac) ---
            which = "rna" if (step % 2 == 0) else "atac"
            sup_losses.append(supervised_step_one_batch(model, opt_backbone, opt_head, epoch, which, it_rna, it_atac))

        rna_acc, rna_f1, atac_acc, atac_f1, mean_f1 = eval_both(model)
        if mean_f1 > best[0]:
            best = (mean_f1, {k: v.detach().cpu().clone() for k, v in model.state_dict().items()})

        print(
            f"[epoch {epoch:03d}] gen_loss={np.mean(gen_losses):.4f} | sup_loss={np.mean(sup_losses):.4f} | "
            f"RNA acc/F1={rna_acc:.3f}/{rna_f1:.3f} | ATAC acc/F1={atac_acc:.3f}/{atac_f1:.3f} | meanF1={mean_f1:.3f}"
        )

    if best[1] is not None:
        model.load_state_dict(best[1])
        print("Restored best joint-mixed with meanF1 =", best[0])

def train_head_then_joint(model):
    print("Warmup (head_only) ...")
    set_trainable(model, "head_only")
    _, opt_head = build_optimizers(model)
    assert opt_head is not None

    best = (-np.inf, None)
    for epoch in range(warmup_epochs):
        loss = supervised_epoch_alternating(model, opt_backbone=None, opt_head=opt_head, epoch=epoch)
        rna_acc, rna_f1, atac_acc, atac_f1, mean_f1 = eval_both(model)

        if mean_f1 > best[0]:
            best = (mean_f1, {k: v.detach().cpu().clone() for k, v in model.state_dict().items()})

        print(
            f"[warmup {epoch:03d}] head_loss={loss:.4f} | "
            f"RNA acc/F1={rna_acc:.3f}/{rna_f1:.3f} | ATAC acc/F1={atac_acc:.3f}/{atac_f1:.3f} | meanF1={mean_f1:.3f}"
        )

    if best[1] is not None:
        model.load_state_dict(best[1])
        print("Warmup restored best with meanF1 =", best[0])

    print("\nSwitching to joint_mixed (backbone unfrozen) ...\n")
    train_joint_mixed(model)


# %% [code]
# ============================================================
# 6) RUN TRAINING
# ============================================================
print("METHOD =", METHOD)
if METHOD == "head_only":
    train_head_only(model)
elif METHOD == "joint_mixed":
    train_joint_mixed(model)
elif METHOD == "head_then_joint":
    train_head_then_joint(model)
else:
    raise ValueError(f"Unknown METHOD: {METHOD}")

# keep a consistent name if you used model_cls elsewhere
model_cls = model


# %% [code]
# ============================================================
# 7) Predict head labels (RNA + ATAC) + confusion matrices
# ============================================================
@torch.no_grad()
def predict_head_codes_single_modality(model, adata, modality, *, head_name, device, batch_size=1024):
    model.eval()
    ds = MultiModalDataset(
        adata_dict={modality: adata},
        X_key="X",
        paired=False,
        device=None,
    )
    loader = DataLoader(ds, batch_size=batch_size, shuffle=False, num_workers=0)

    pred_codes = []
    for batch in loader:
        x_batch = _unwrap_x(batch)
        x_batch = {k: v.to(device) for k, v in x_batch.items()}
        logits = model.predict_heads(
            x_batch,
            return_probs=False,
            use_mean=True,
            inject_label_expert=False,
        )[head_name]
        pred_codes.append(logits.argmax(dim=-1).cpu().numpy())

    return np.concatenate(pred_codes).astype(np.int64)

# Ding RNA predictions
uni_rna_pp.obs[f"{head_name}_pred_code"] = predict_head_codes_single_modality(
    model_cls, uni_rna_pp, "rna", head_name=head_name, device=device, batch_size=1024
)
uni_rna_pp.obs[f"{head_name}_pred"] = pd.Categorical(
    [id_to_label[int(i)] for i in uni_rna_pp.obs[f"{head_name}_pred_code"].to_numpy()],
    categories=classes,
)

# Satpathy ATAC predictions
satpathy_atac_pp.obs[f"{head_name}_pred_code"] = predict_head_codes_single_modality(
    model_cls, satpathy_atac_pp, "atac", head_name=head_name, device=device, batch_size=1024
)
satpathy_atac_pp.obs[f"{head_name}_pred"] = pd.Categorical(
    [id_to_label[int(i)] for i in satpathy_atac_pp.obs[f"{head_name}_pred_code"].to_numpy()],
    categories=classes,
)

# Confusion on labeled cells only (RNA)
mask_lab_rna = (uni_rna_pp.obs[f"{head_name}_code"].to_numpy() >= 0)
y_true = uni_rna_pp.obs.loc[mask_lab_rna, f"{head_name}_code"].to_numpy().astype(int)
y_pred = uni_rna_pp.obs.loc[mask_lab_rna, f"{head_name}_pred_code"].to_numpy().astype(int)
acc = accuracy_score(y_true, y_pred)
f1  = f1_score(y_true, y_pred, average="macro")
cm  = confusion_matrix(y_true, y_pred, labels=np.arange(n_classes))
print("RNA head acc:", acc, "macroF1:", f1)
cm_df_rna = pd.DataFrame(cm, index=classes, columns=classes)
display(cm_df_rna)

# Confusion on labeled cells only (ATAC)
mask_lab_atac = (satpathy_atac_pp.obs[f"{head_name}_code"].to_numpy() >= 0)
y_true = satpathy_atac_pp.obs.loc[mask_lab_atac, f"{head_name}_code"].to_numpy().astype(int)
y_pred = satpathy_atac_pp.obs.loc[mask_lab_atac, f"{head_name}_pred_code"].to_numpy().astype(int)
acc = accuracy_score(y_true, y_pred)
f1  = f1_score(y_true, y_pred, average="macro")
cm  = confusion_matrix(y_true, y_pred, labels=np.arange(n_classes))
print("ATAC head acc:", acc, "macroF1:", f1)
cm_df_atac = pd.DataFrame(cm, index=classes, columns=classes)
display(cm_df_atac)


# %% [code]
print(latent_choice)


# %% [code]
# ============================================================
# 8) Encode new latent for (bridge + unimodal) and UMAP
#    (Uses encode_adata; stores in .obsm[out_key])
# ============================================================
def _ensure_obs_cols(a, dataset, modality, is_reference):
    a = a.copy()
    a.obs["dataset"] = str(dataset)
    a.obs["modality"] = str(modality)
    a.obs["is_reference"] = bool(is_reference)
    return a

# pick bridge datasets for visualization
bridge_rna  = globals().get("bridge_rna",  rna_train_pp)
bridge_atac = globals().get("bridge_atac", atac_train_lsi)

bridge_rna   = _ensure_obs_cols(bridge_rna,   dataset="multiome_rna",  modality="rna",  is_reference=True)
bridge_atac  = _ensure_obs_cols(bridge_atac,  dataset="multiome_atac", modality="atac", is_reference=True)
ding_rna     = _ensure_obs_cols(uni_rna_pp,   dataset="ding_rna",      modality="rna",  is_reference=False)
satpathy_atac= _ensure_obs_cols(satpathy_atac_pp, dataset="satpathy_atac", modality="atac", is_reference=False)

def _encode_into(adata, modality):
    Z = encode_adata(
        model_cls,
        adata,
        modality=modality,
        latent=latent_choice,
        device=device,
        batch_size=1024,
    )
    adata = adata.copy()
    adata.obsm[out_key] = np.asarray(Z, dtype=np.float32)
    return adata

bridge_rna    = _encode_into(bridge_rna, "rna")
bridge_atac   = _encode_into(bridge_atac, "atac")
ding_rna      = _encode_into(ding_rna, "rna")
satpathy_atac = _encode_into(satpathy_atac, "atac")

combo_ft = ad.concat(
    {"multiome_rna": bridge_rna, "multiome_atac": bridge_atac, "ding_rna": ding_rna, "satpathy_atac": satpathy_atac},
    axis=0,
    join="outer",
    label="dataset",
    index_unique=None,
)
assert out_key in combo_ft.obsm, f"Missing {out_key} in combo_ft.obsm"

sc.pp.neighbors(combo_ft, n_neighbors=umap_n_neighbors, use_rep=out_key, metric="euclidean")
sc.tl.umap(combo_ft, min_dist=0.3, spread=1.0)

sc.pl.umap(combo_ft, color=["dataset"],      title=[f"Dataset ({out_key})"], frameon=False, size=10, alpha=0.5)
sc.pl.umap(combo_ft, color=["modality"],     title=[f"Modality ({out_key})"], frameon=False, size=10, alpha=0.5)
sc.pl.umap(combo_ft, color=["is_reference"], title=[f"Reference vs unimodal ({out_key})"], frameon=False, size=10, alpha=0.5)

# labeled celltypes where they exist
has_labels = combo_ft.obs[label_col].notna() & ~combo_ft.obs[label_col].astype(str).str.startswith("unlabeled_")
if has_labels.sum() > 0:
    sc.pl.umap(combo_ft[has_labels], color=[label_col], title=[f"{label_col} ({out_key})"], frameon=False, size=10, alpha=0.5)


# %% [code]
sc.pl.umap(combo_ft, color=["tech"], title=[f"Tech ({out_key})"], frameon=False, size=10, alpha=0.5)


# %% [code]
print(combo_ft.uns['dataset_colors'])
print(combo_ft.uns['modality_colors'])
print(combo_ft)


# %% [code]
print(combo_ft.obs["dataset"].value_counts(dropna=False))
print(sorted(combo_ft.obs["dataset"].astype(str).unique()))


# %% [code]
import pandas as pd

# 0) clean labels (prevents invisible whitespace bugs)
combo_ft.obs["dataset"] = combo_ft.obs["dataset"].astype(str).str.strip()

# 1) the order you want in the legend (match your 2nd image)
dataset_order = ["multiome_rna", "multiome_atac", "ding_rna", "satpathy_atac"]

# 2) exact colors (match your 1st image)
palette = {
    #"multiome_rna":   "#1f77b4",  # blue
    "multiome_rna":   "#2ca02c",  # green
    #"multiome_atac":  "#ff7f0e",  # orange
    "multiome_atac":  "#d62728",  # red
    #"ding_rna":       "#2ca02c",  # green
    "ding_rna":       "#1f77b4",  # blue
    #"satpathy_atac":  "#d62728",  # red
    "satpathy_atac":  "#ff7f0e",  # orange
}

# enforce categorical order
combo_ft.obs["dataset"] = pd.Categorical(
    combo_ft.obs["dataset"],
    categories=dataset_order,
    ordered=True,
)

# force Scanpy's internal color list to match that order
combo_ft.uns["dataset_colors"] = [palette[k] for k in dataset_order]

sc.pl.umap(
    combo_ft,
    color="dataset",
    title=f"Dataset ({out_key})",
    frameon=False,
    size=10,
    alpha=0.5,
)


# %% [code]
# ============================================================
# 9) kNN label transfer in NEW latent (optional, requires your functions)
# ============================================================
# Requires: knn_label_transfer_confusion_two_adata, plot_knn_confusion_heatmap
if "knn_label_transfer_confusion_two_adata" in globals():
    exclude_labels_knn = ("Other", "Unknown", "Unassigned", "General PBMC", "Megakaryocyte")
    k = 15

    ding_mask = combo_ft.obs["dataset"].astype(str).eq("ding_rna").to_numpy()
    sat_mask  = combo_ft.obs["dataset"].astype(str).eq("satpathy_atac").to_numpy()
    adata_ding = combo_ft[ding_mask].copy()
    adata_sat  = combo_ft[sat_mask].copy()

    res_r2a = knn_label_transfer_confusion_two_adata(
        adata_ref=adata_ding,
        adata_tgt=adata_sat,
        emb_key=out_key,
        ct_key=label_col,
        k=k,
        exclude_labels=exclude_labels_knn,
    )
    print("Ding RNA → Satpathy ATAC", "Acc:", res_r2a["acc"], "MacroF1:", res_r2a["macro_f1"])
    display(res_r2a["cm_df"])
    if "plot_knn_confusion_heatmap" in globals():
        plot_knn_confusion_heatmap(res_r2a, normalize="row", title=f"Ding RNA → Satpathy ATAC (k={k}, {out_key})")

    res_a2r = knn_label_transfer_confusion_two_adata(
        adata_ref=adata_sat,
        adata_tgt=adata_ding,
        emb_key=out_key,
        ct_key=label_col,
        k=k,
        exclude_labels=exclude_labels_knn,
    )
    print("Satpathy ATAC → Ding RNA", "Acc:", res_a2r["acc"], "MacroF1:", res_a2r["macro_f1"])
    display(res_a2r["cm_df"])
    if "plot_knn_confusion_heatmap" in globals():
        plot_knn_confusion_heatmap(res_a2r, normalize="row", title=f"Satpathy ATAC → Ding RNA (k={k}, {out_key})")


# %% [code]
# ============================================================
# 10) Per-dataset head eval with feature alignment (robust)
# ============================================================
def _infer_modality_from_dataset_name(ds: str) -> str:
    s = str(ds).lower()
    if "atac" in s: return "atac"
    if "rna"  in s: return "rna"
    raise ValueError(f"Can't infer modality from dataset={ds!r}.")

def _align_to_expected_vars(adata, expected_vars, *, name=""):
    expected_vars = pd.Index(expected_vars)
    have = adata.var_names
    missing = expected_vars.difference(have)
    if len(missing) > 0:
        raise ValueError(
            f"{name}: missing {len(missing)} expected features. Example missing: {missing[:10].tolist()}"
        )
    return adata[:, expected_vars].copy()

@torch.no_grad()
def eval_head_on_subset(
    model,
    adata_sub,
    *,
    modality: str,
    head_name: str,
    label_col: str,
    label_to_id: dict,
    expected_vars_by_modality: dict,
    exclude_labels=(),
    batch_size: int = 1024,
    device="cuda",
):
    y_raw = adata_sub.obs[label_col].astype(str)
    mask = (
        (~y_raw.str.startswith("unlabeled_"))
        & (~y_raw.isin(list(exclude_labels)))
        & (y_raw.isin(label_to_id.keys()))
    )
    if mask.sum() == 0:
        return None

    ad0 = adata_sub[mask.to_numpy()].copy()
    ad0 = _align_to_expected_vars(ad0, expected_vars_by_modality[modality], name=f"{modality}/{head_name}")

    y = torch.from_numpy(y_raw[mask].map(label_to_id).to_numpy(np.int64))

    ds = MultiModalDataset(
        adata_dict={modality: ad0},
        X_key="X",
        labels={head_name: y},
        paired=False,
        device=None,
    )
    loader = DataLoader(ds, batch_size=batch_size, shuffle=False, num_workers=0)

    model.eval()
    y_true_all, y_pred_all = [], []
    for x_batch, y_batch in loader:
        x_batch = {k: v.to(device) for k, v in x_batch.items()}
        y_true = y_batch[head_name].to(device).view(-1)
        logits = model.predict_heads(x_batch, return_probs=False, use_mean=True, inject_label_expert=False)[head_name]
        y_pred = logits.argmax(dim=-1)
        y_true_all.append(y_true.cpu().numpy())
        y_pred_all.append(y_pred.cpu().numpy())

    y_true_all = np.concatenate(y_true_all)
    y_pred_all = np.concatenate(y_pred_all)
    acc = float(accuracy_score(y_true_all, y_pred_all))
    f1  = float(f1_score(y_true_all, y_pred_all, average="macro"))
    cm  = confusion_matrix(y_true_all, y_pred_all)
    return {"n": int(len(y_true_all)), "acc": acc, "macroF1": f1, "cm": cm}

expected_vars_by_modality = {
    "rna":  list(rna_train_pp.var_names),
    "atac": list(atac_train_lsi.var_names),
}

dataset_key = "dataset"
results = []
cms = {}

for ds_name in pd.unique(combo_ft.obs[dataset_key].astype(str)):
    ad_ds = combo_ft[combo_ft.obs[dataset_key].astype(str).eq(ds_name).to_numpy()].copy()
    modality = _infer_modality_from_dataset_name(ds_name)

    out = eval_head_on_subset(
        model_cls,
        ad_ds,
        modality=modality,
        head_name=head_name,
        label_col=label_col,
        label_to_id=label_to_id,
        expected_vars_by_modality=expected_vars_by_modality,
        exclude_labels=exclude_labels_train,
        batch_size=1024,
        device=device,
    )

    if out is None:
        results.append({"dataset": ds_name, "modality": modality, "n": 0, "acc": np.nan, "macroF1": np.nan})
    else:
        results.append({"dataset": ds_name, "modality": modality, "n": out["n"], "acc": out["acc"], "macroF1": out["macroF1"]})
        cms[ds_name] = out["cm"]

df_head_by_dataset = pd.DataFrame(results).sort_values(["modality", "acc"], ascending=[True, False])
display(df_head_by_dataset)


# %% [code]
print(bridge_rna)
print(bridge_atac)


# %% [code]
rna_multiome_pp = bridge_rna.copy()
atac_multiome_lsi = bridge_atac.copy()


# %% [code]
def _labeled_mask(adata, label_col, exclude_labels):
    if label_col not in adata.obs:
        raise KeyError(f"adata.obs[{label_col!r}] not found")
    y = adata.obs[label_col].astype(str)
    m = (~y.str.startswith("unlabeled_")) & (~y.isin(list(exclude_labels)))
    return m.to_numpy(), y

def _make_codes(adata, y_raw, mask, label_to_id, head_name, ignore_index=-1):
    codes = np.full(adata.n_obs, ignore_index, dtype=np.int64)
    mask = np.asarray(mask, dtype=bool)
    if mask.sum() == 0:
        adata.obs[f"{head_name}_code"] = codes
        return codes

    idx = np.where(mask)[0]
    y = y_raw.iloc[idx].astype(str)

    mapped = y.map(label_to_id)          # NaN for OOV
    keep = mapped.notna().to_numpy()

    codes[idx[keep]] = mapped.to_numpy(dtype=np.int64)[keep]
    adata.obs[f"{head_name}_code"] = codes
    return codes

def _split_stratified(idx_all, y_codes, test_size=0.10, seed=42):
    sss = StratifiedShuffleSplit(n_splits=1, test_size=test_size, random_state=seed)
    tr_rel, va_rel = next(sss.split(idx_all, y_codes))
    return idx_all[tr_rel], idx_all[va_rel]

def _make_supervised_loaders(adata, codes, modality, batch_tr, batch_va):
    idx_lab = np.where(codes >= 0)[0]
    if len(idx_lab) == 0:
        raise ValueError(f"No labeled cells for modality={modality!r}")
    y_codes = codes[idx_lab]
    idx_tr, idx_va = _split_stratified(idx_lab, y_codes)

    ad_tr = adata[idx_tr].copy()
    ad_va = adata[idx_va].copy()

    y_tr = torch.from_numpy(codes[idx_tr].astype(np.int64))
    y_va = torch.from_numpy(codes[idx_va].astype(np.int64))

    ds_tr = MultiModalDataset(
        adata_dict={modality: ad_tr},
        X_key="X",
        labels={head_name: y_tr},
        paired=False,
        device=None,
    )
    ds_va = MultiModalDataset(
        adata_dict={modality: ad_va},
        X_key="X",
        labels={head_name: y_va},
        paired=False,
        device=None,
    )

    ld_tr = DataLoader(ds_tr, batch_size=batch_tr, shuffle=True,  num_workers=0)
    ld_va = DataLoader(ds_va, batch_size=batch_va, shuffle=False, num_workers=0)
    return ds_tr, ds_va, ld_tr, ld_va

def _cycle(loader):
    while True:
        for batch in loader:
            yield batch

def _align_features_strict(adata, ref, modality_name):
    if ref is None:
        return adata
    if not np.array_equal(adata.var_names, ref.var_names):
        missing = ref.var_names.difference(adata.var_names)
        extra   = adata.var_names.difference(ref.var_names)
        if len(missing) or len(extra):
            raise ValueError(
                f"[{modality_name}] Feature mismatch vs reference.\n"
                f"  missing (need in adata): {len(missing)}\n"
                f"  extra   (in adata only): {len(extra)}\n"
                f"Fix preprocessing/projecting so var_names match."
            )
        adata = adata[:, ref.var_names].copy()
    return adata


# %% [code]
assert "rna_multiome_pp" in globals() and "atac_multiome_lsi" in globals(), "Provide multiome RNA+ATAC to annotate."

# Align cells
if not np.array_equal(rna_multiome_pp.obs_names, atac_multiome_lsi.obs_names):
    common = rna_multiome_pp.obs_names.intersection(atac_multiome_lsi.obs_names)
    rna_multiome_pp = rna_multiome_pp[common].copy()
    atac_multiome_lsi = atac_multiome_lsi[common].copy()
    atac_multiome_lsi = atac_multiome_lsi[rna_multiome_pp.obs_names].copy()

# Align features
rna_multiome_pp  = _align_features_strict(rna_multiome_pp,  rna_train_pp,  "rna")
atac_multiome_lsi = _align_features_strict(atac_multiome_lsi, atac_train_lsi, "atac")

@torch.no_grad()
def predict_fused_to_obs(model, rna_ad, atac_ad, *, head_name, id_to_label, batch_size=1024):
    ds = MultiModalDataset(
        adata_dict={"rna": rna_ad, "atac": atac_ad},
        X_key="X",
        paired=True,
        device=None,
    )
    loader = DataLoader(ds, batch_size=batch_size, shuffle=False, num_workers=0)

    model.eval()
    probs_all = []
    for batch in loader:
        x_batch = batch[0] if isinstance(batch, (tuple, list)) else batch
        x_batch = {k: v.to(device) for k, v in x_batch.items()}

        probs = model.predict_heads(
            x_batch,
            return_probs=True,
            use_mean=True,
            inject_label_expert=False,
        )[head_name]
        probs_all.append(probs.detach().cpu())

    probs_all = torch.cat(probs_all, dim=0).numpy()
    pred_id = probs_all.argmax(axis=1).astype(np.int64)
    conf = probs_all.max(axis=1).astype(np.float32)
    pred_label = np.array([id_to_label[int(i)] for i in pred_id], dtype=object)

    for ad_ in (rna_ad, atac_ad):
        ad_.obs[f"{head_name}_pred_code"] = pred_id
        ad_.obs[f"{head_name}_pred"] = pd.Categorical(pred_label, categories=list(id_to_label.values()))
        ad_.obs[f"{head_name}_conf"] = conf

    return pred_id, conf

pred_id_fused, conf_fused = predict_fused_to_obs(
    model, rna_multiome_pp, atac_multiome_lsi, head_name=head_name, id_to_label=id_to_label, batch_size=1024
)

# Optional confidence thresholding
CONF_THRESH = 0.25
low = conf_fused < CONF_THRESH
for ad_ in (rna_multiome_pp, atac_multiome_lsi):
    col = f"{head_name}_pred_thresh{CONF_THRESH:g}"
    tmp = ad_.obs[f"{head_name}_pred"].astype(str).to_numpy()
    tmp[low] = "Unassigned"
    ad_.obs[col] = pd.Categorical(tmp)

print("Annotated multiome cells:", rna_multiome_pp.n_obs, "| low-conf frac:", float(low.mean()))


# %% [code]
# Encode latents
#latent_choice = "modality_mean"
latent_choice = "moe_mean"
out_key = "X_univi_ft"

def _encode_into(adata, modality):
    Z = encode_adata(
        model,
        adata,
        modality=modality,
        latent=latent_choice,
        device=device,
        batch_size=1024,
    )
    adata.obsm[out_key] = np.asarray(Z, dtype=np.float32)
    return adata

# Make small wrappers for consistent metadata
def _ensure_obs_cols(a, dataset, modality, is_reference):
    a = a.copy()
    a.obs["dataset"] = str(dataset)
    a.obs["modality"] = str(modality)
    a.obs["is_reference"] = bool(is_reference)
    return a

bridge_rna   = _ensure_obs_cols(rna_train_pp,      "multiome_rna",  "rna",  True)
bridge_atac  = _ensure_obs_cols(atac_train_lsi,    "multiome_atac", "atac", True)
ding_rna     = _ensure_obs_cols(uni_rna_pp,        "ding_rna",      "rna",  False)
sat_atac     = _ensure_obs_cols(satpathy_atac_pp,  "satpathy_atac", "atac", False)

bridge_rna  = _encode_into(bridge_rna,  "rna")
bridge_atac = _encode_into(bridge_atac, "atac")
ding_rna    = _encode_into(ding_rna,    "rna")
sat_atac    = _encode_into(sat_atac,    "atac")

# Concatenate fresh (avoids accidentally reusing old neighbor/umap)
combo_ft = ad.concat(
    {"multiome_rna": bridge_rna, "multiome_atac": bridge_atac, "ding_rna": ding_rna, "satpathy_atac": sat_atac},
    axis=0,
    join="outer",
    label="dataset",
    index_unique=None,
)

assert out_key in combo_ft.obsm, f"Missing {out_key} in combo_ft.obsm"

# NEW UMAP stored under a new basis name
suffix = "univi_ft"
neighbors_key = f"neighbors_{suffix}"

sc.pp.neighbors(combo_ft, n_neighbors=30, use_rep=out_key, metric="euclidean", key_added=neighbors_key)
sc.tl.umap(combo_ft, neighbors_key=neighbors_key, min_dist=0.3, spread=1.0)

# stash it so you can plot without overwriting later
combo_ft.obsm[f"X_umap_{suffix}"] = combo_ft.obsm["X_umap"].copy()

# Plot the NEW one explicitly
sc.pl.embedding(combo_ft, basis=f"umap_{suffix}", color=["dataset"], frameon=False, size=2,
                title=[f"Dataset ({out_key}) [{suffix}]"])
sc.pl.embedding(combo_ft, basis=f"umap_{suffix}", color=["modality"], frameon=False, size=2,
                title=[f"Modality ({out_key}) [{suffix}]"])
sc.pl.embedding(combo_ft, basis=f"umap_{suffix}", color=["is_reference"], frameon=False, size=2,
                title=[f"Reference vs unimodal ({out_key}) [{suffix}]"])

if "tech" in combo_ft.obs.columns:
    sc.pl.embedding(combo_ft, basis=f"umap_{suffix}", color=["tech"], frameon=False, size=2,
                    title=[f"Tech ({out_key}) [{suffix}]"])

has_labels = combo_ft.obs[label_col].notna() & ~combo_ft.obs[label_col].astype(str).str.startswith("unlabeled_")
if has_labels.sum() > 0:
    sc.pl.embedding(combo_ft[has_labels], basis=f"umap_{suffix}", color=[label_col], frameon=False, size=2,
                    title=[f"{label_col} ({out_key}) [{suffix}]"])


# %% [code]
# assumes you already have knn_label_transfer_confusion_two_adata and plot_knn_confusion_heatmap
label_col_lt = "celltype_harmonized_coarse"
exclude_lt = ("Other", "Unknown", "Unassigned", "Megakaryocyte")
k = 15

ding_mask = combo_ft.obs["dataset"].astype(str).eq("ding_rna").to_numpy()
sat_mask  = combo_ft.obs["dataset"].astype(str).eq("satpathy_atac").to_numpy()

adata_ding = combo_ft[ding_mask].copy()
adata_sat  = combo_ft[sat_mask].copy()

res_r2a = knn_label_transfer_confusion_two_adata(
    adata_ref=adata_ding,
    adata_tgt=adata_sat,
    emb_key=out_key,
    ct_key=label_col_lt,
    k=k,
    exclude_labels=exclude_lt,
)
print("Ding RNA → Satpathy ATAC")
print("  Acc:", res_r2a["acc"])
print("  MacroF1:", res_r2a["macro_f1"])
display(res_r2a["cm_df"])
plot_knn_confusion_heatmap(res_r2a, normalize="row",
                           title=f"Ding RNA → Satpathy ATAC (k={k}, {out_key})")

res_a2r = knn_label_transfer_confusion_two_adata(
    adata_ref=adata_sat,
    adata_tgt=adata_ding,
    emb_key=out_key,
    ct_key=label_col_lt,
    k=k,
    exclude_labels=exclude_lt,
)
print("Satpathy ATAC → Ding RNA")
print("  Acc:", res_a2r["acc"])
print("  MacroF1:", res_a2r["macro_f1"])
display(res_a2r["cm_df"])
plot_knn_confusion_heatmap(res_a2r, normalize="row",
                           title=f"Satpathy ATAC → Ding RNA (k={k}, {out_key})")


# %% [code]
print(combo_ft)


# %% [code]
sc.pl.embedding(combo_ft, basis=f"X_umap_univi_ft", color=["celltype_harmonized"], frameon=False, size=3, alpha=0.65,
                title=[f"Reference vs unimodal by harmonized celltype"])


# %% [code]
sc.pl.embedding(combo_ft, basis=f"X_umap_univi_ft", color=["celltype_harmonized_coarse"], frameon=False, size=3, alpha=0.65,
                title=[f"Reference vs unimodal by harmonized coarse celltype"])


# %% [code]
combo_rna = combo_ft[combo_ft.obs['modality'] == 'rna']


# %% [code]
# (6a) UMAP overlays for a *small* set of markers (pick ~6–12 so it’s readable)
umap_genes = [g for g in ["MS4A1","CD79A","IL7R","CCR7","NKG7","GNLY","LST1","S100A8","LYZ","TRAC","FCER1A","CLEC10A"] if g in combo_rna.var_names]
#sc.pl.umap(combo_ft, color=umap_genes, frameon=False, size=20, alpha=0.65, ncols=2)


# %% [code]
#sc.pl.umap(combo_ft, color=umap_genes, frameon=False, size=20, alpha=0.65, ncols=2)


# %% [code]
#sc.pl.embedding(combo_ft, basis=f"X_umap_univi_ft", color=["celltype_pred"], frameon=False, size=2,
#                title=[f"Reference vs unimodal by harmonized coarse celltype"])


# %% [code]
sc.pl.embedding(combo_ft, basis=f"X_umap_univi_ft", color=["modality"], frameon=False, size=3, alpha=0.65,
                title=[f"Reference vs unimodal by harmonized coarse celltype"])


# %% [code]
sc.pl.embedding(combo_ft, basis=f"X_umap_univi_ft", color=["celltype_higher_res_pred"], frameon=False, size=3, alpha=0.65,
                title=[f"Reference vs unimodal by harmonized celltype"])


# %% [code]
sc.pl.embedding(combo_ft, basis=f"X_umap_univi_ft", color=["dataset"], frameon=False, size=3, alpha=0.65,
                title=[f"Reference vs unimodal by harmonized celltype"])


# %% [code]
sc.pl.embedding(combo_ft, basis=f"X_umap_univi_ft", color=["tech"], frameon=False, size=3, alpha=0.3,
                title=[f"Reference vs unimodal by harmonized celltype"])


# %% [code]
print(rna_multiome_pp)


# %% [code]
sc.pl.embedding(rna_multiome_pp, basis=f"X_umap", color=["celltype_higher_res_conf"], frameon=False, size=3, alpha=0.65,
                title=[f"Reference bridge data annotated by classification decoder head"])


# %% [code]
sc.pl.embedding(rna_multiome_pp, basis=f"X_umap", color=["celltype_higher_res_pred"], frameon=False, size=3, alpha=0.65,
                title=[f"Reference bridge data annotated by classification decoder head"])


# %% [code]
print(atac_multiome_lsi)


# %% [code]
sc.pl.embedding(atac_multiome_lsi, basis=f"X_univi_ft", color=["celltype_higher_res_pred"], frameon=False, size=3, alpha=0.65,
                title=[f"Reference bridge data annotated by classification decoder head"])


# %% [code]
import numpy as np
import pandas as pd
import scanpy as sc
import anndata as ad

from univi.evaluation import encode_adata

# -----------------------------
# Inputs you already have
# -----------------------------
# rna_multiome_pp : AnnData (RNA; preprocessed to match model input space)
# atac_multiome_lsi : AnnData (ATAC; preprocessed LSI to match model input space)
# model_cls : fine-tuned UniVI model (with head)
# device : "cuda" / "cpu"
#
# And you said both already have head outputs in .obs, e.g.:
#   obs[f"{head_name}_pred"], obs[f"{head_name}_conf"] etc.

# -----------------------------
# Settings
# -----------------------------
out_key = "X_univi_ft_multiome"   # where to store latent in .obsm
latent_choice = "moe_mean"        # "moe_mean" (shared) or "modality_mean"
head_name = "celltype_higher_res" # change if you used a different head name
anno_key = f"{head_name}_pred"    # your predicted annotation column in .obs

# -----------------------------
# (1) Align cells if needed (paired multiome)
# -----------------------------
if not np.array_equal(rna_multiome_pp.obs_names, atac_multiome_lsi.obs_names):
    common = rna_multiome_pp.obs_names.intersection(atac_multiome_lsi.obs_names)
    rna_multiome_pp = rna_multiome_pp[common].copy()
    atac_multiome_lsi = atac_multiome_lsi[common].copy()
    atac_multiome_lsi = atac_multiome_lsi[rna_multiome_pp.obs_names].copy()

# -----------------------------
# (2) Encode into latent with the fine-tuned model
# -----------------------------
model_cls.eval()

Z_rna = encode_adata(
    model_cls,
    rna_multiome_pp,
    modality="rna",
    latent=latent_choice,
    device=device,
    batch_size=1024,
)
rna_multiome_pp.obsm[out_key] = np.asarray(Z_rna, dtype=np.float32)

Z_atac = encode_adata(
    model_cls,
    atac_multiome_lsi,
    modality="atac",
    latent=latent_choice,
    device=device,
    batch_size=1024,
)
atac_multiome_lsi.obsm[out_key] = np.asarray(Z_atac, dtype=np.float32)

# -----------------------------
# (3) Make a combined AnnData for UMAP
# -----------------------------
rna_plot = rna_multiome_pp.copy()
atac_plot = atac_multiome_lsi.copy()

rna_plot.obs["modality"] = "rna"
atac_plot.obs["modality"] = "atac"

# ensure the annotation column exists (optional safeguard)
if anno_key not in rna_plot.obs.columns:
    raise KeyError(f"{anno_key!r} not found in rna_multiome_pp.obs")
if anno_key not in atac_plot.obs.columns:
    raise KeyError(f"{anno_key!r} not found in atac_multiome_lsi.obs")

combo = ad.concat(
    {"multiome_rna": rna_plot, "multiome_atac": atac_plot},
    axis=0,
    join="outer",
    label="dataset",
    index_unique=None,
)

# -----------------------------
# (4) Neighbors + UMAP on new latent
# -----------------------------
sc.pp.neighbors(combo, n_neighbors=30, use_rep=out_key, metric="euclidean")
sc.tl.umap(combo, min_dist=0.3, spread=1.0)

# -----------------------------
# (5) Plot UMAP colored by new annotations (+ modality)
# -----------------------------
sc.pl.umap(combo, color=["modality"], frameon=False, size=30, alpha=0.65, title=f"Multiome ({latent_choice}) by modality")
sc.pl.umap(combo, color=[anno_key], frameon=False, size=30, alpha=0.65, title=f"Multiome ({latent_choice}) by {anno_key}")

# optional: show confidence if you have it
conf_key = f"{head_name}_conf"
if conf_key in combo.obs.columns:
    sc.pl.umap(combo, color=[conf_key], frameon=False, size=30, alpha=0.65, title=f"Multiome ({latent_choice}) by {conf_key}")

# If you want the per-modality plots:
sc.pl.umap(combo[combo.obs["modality"] == "rna"],  color=[anno_key], frameon=False, size=30, alpha=0.65, title=f"RNA only ({latent_choice})")
sc.pl.umap(combo[combo.obs["modality"] == "atac"], color=[anno_key], frameon=False, size=30, alpha=0.65, title=f"ATAC only ({latent_choice})")


# %% [code]
# ------------------------------------------------
# 3.1 Set plotting defaults
# ------------------------------------------------

sc.set_figure_params(
    figsize=(10, 8),
    dpi=100,
    dpi_save=300,
    fontsize=12,
    frameon=False,
)
plt.rcParams.update({
    "figure.figsize": (10, 8),
    "figure.dpi": 100,
    "savefig.dpi": 300,
    "savefig.bbox": "tight",
    "savefig.pad_inches": 0.1,
})


# %% [code]
print(combo_ft.obs['dataset'])


# %% [code]
# -----------------------------
# (6) Quick marker genes (RNA only) for your predicted cell types
# -----------------------------
'''
marker_genes = {
    "B": ["MS4A1", "CD79A", "CD74"],
    "CD4 T": ["IL7R", "CCR7", "LTB"],
    "CD8/cytotoxic": ["NKG7", "GNLY", "GZMB", "PRF1"],
    "NK": ["NKG7", "FCGR3A", "TRAC"],   # TRAC sanity check that these are T/NK-like
    "Monocyte": ["LST1", "S100A8", "S100A9", "FCN1", "LYZ"],
    "Dendritic": ["CLEC10A"],  # LILRA4 more pDC-ish (if present)
}
'''
'''
marker_genes = {
    "B": ["MS4A1", "CD79A", "CD74"],
    "CD4 T": ["IL7R", "CCR7", "LTB", "TRAC"],      # T-lineage check
    "CD8/cytotoxic": ["NKG7", "GNLY", "GZMB", "PRF1", "TRAC"],  # helps distinguish cytotoxic T vs NK
    "NK": ["NKG7", "FCGR3A"],                      # NK markers only
    "TCR/CD3": ["TRAC", "CD3D", "CD3E"],           # optional: best for “NK should be negative”
    "Monocyte": ["LST1", "S100A8", "S100A9", "FCN1", "LYZ"],
    "Dendritic": ["CLEC10A"],
}
'''
'''
marker_genes = {
    # --- B cells (naive + memory + antigen presentation) ---
    "B": [
        "MS4A1", "CD79A", "CD74",
        "CD79B", "HLA-DRA", "HLA-DPA1", "HLA-DPB1",
        "CD22", "CD19",
        "BANK1", "CD83"
    ],

    # --- CD4 T (naive/central memory leaning) ---
    "CD4 T": [
        "IL7R", "CCR7", "LTB", "TRAC",
        "MAL", "TCF7", "LEF1", "LST1",  # (remove LST1 if you want zero myeloid bleed)
        "TRBC1", "ICOS", "IL32"
    ],

    # --- CD8 / cytotoxic program (overlaps NK on purpose) ---
    "CD8/cytotoxic": [
        "NKG7", "GNLY", "GZMB", "PRF1", "TRAC",
        "GZMK", "GZMH", "FGFBP2",
        "CTSW", "KLRD1", "KLRB1",
        "TRBC1", "CD8A", "CD8B"
    ],

    # --- NK (TRAC-/CD3-; split CD56bright vs CD16+) ---
    "NK": [
        "NKG7", "FCGR3A",
        "KLRD1", "TRDC",  # (drop TRDC if you don't want gamma-delta signal here)
        "TYROBP", "FCER1G", "XCL2", "KLRC1", "KLRC2",
        "IL7R"  # (optional; can highlight CD56bright-like NK if present)
    ],

    # --- TCR/CD3 lineage check (should be ~0 in true NK/mono) ---
    "TCR/CD3": [
        "TRAC", "TRBC1", "TRBC2",
        "CD247", "TRAT1"
    ],

    # --- Monocytes (classical + non-classical) ---
    "Monocyte": [
        "LST1", "S100A8", "S100A9", "FCN1", "LYZ",
        "CTSS", "LGALS3", "MNDA", "TYROBP",
        "MS4A7", "FCGR3A", "LILRB1",
        "VCAN", "IFITM3"
    ],

    # --- Dendritic (cDC2 + cDC1 + pDC coverage) ---
    "Dendritic": [
        "CST3", "ITGAX", "CLEC10A",
        "IL3RA", "GZMB",                  # pDC-ish
        "IRF7"                            # optional support depending on dataset
    ],
}
'''

marker_genes = {
    "B": ["MS4A1", "CD79A", "CD74", "CD37", "CD19"],

    "CD4 T": ["IL7R", "CCR7", "LTB", "TCF7", "LEF1"],

    "CD8/cytotoxic": ["CD8A", "NKG7", "GNLY", "GZMB", "PRF1", "FGFBP2"],

    "NK": ["FCGR3A", "KLRD1", "TYROBP", "FCER1G", "XCL1"],

    "TCR/CD3": ["TRAC", "TRBC1", "CD3D", "CD3E", "CD247"],

    "Monocyte": ["LST1", "S100A8", "S100A9", "FCN1", "LYZ", "MS4A7"],

    "Dendritic": ["FCER1A", "CLEC10A", "CST3", "ITGAX", "CLEC9A", "FCER1A"],

    # optional if you actually have pDC as a row:
    "pDC": ["IL3RA", "LILRA4", "GZMB", "IRF7"],
}

# subset to RNA rows for gene expression plots
combo_rna = combo[combo.obs["modality"] == "rna"].copy()
#combo_rna = combo_ft[combo_ft.obs['dataset'] == 'ding_rna'].copy()

# keep only genes that actually exist
flat = sorted({g for gs in marker_genes.values() for g in gs})
present = [g for g in flat if g in combo_rna.var_names]
missing = [g for g in flat if g not in combo_rna.var_names]
print(f"Marker genes present: {len(present)}/{len(flat)}")
if missing:
    print("Missing markers (ok):", missing)

# (6a) UMAP overlays for a *small* set of markers (pick ~6–12 so it’s readable)
#umap_genes = [g for g in ["MS4A1","CD79A","IL7R","TRAC","CD8A","NKG7","FCGR3A","LST1","S100A8","LYZ","FCER1A","CLEC10A"]
# if g in combo_rna.var_names]
#sc.pl.umap(combo_rna, color=umap_genes, frameon=False, size=20, alpha=0.65, ncols=2)

# (6b) Dotplot by predicted labels (clean “what’s activating in each class” view)
'''
sc.pl.dotplot(
    combo_rna,
    var_names=marker_genes,      # dict -> grouped gene panels
    groupby=anno_key,
    standard_scale="var",        # z-score genes for comparability
    dot_min=0.0,
    dot_max=0.7,
)
'''

# %% [code]
# (6a) UMAP overlays for a *small* set of markers (pick ~6–12 so it’s readable)
umap_genes = [g for g in ["MS4A1","CD79A","IL7R","TRAC","CD8A","NKG7","FCGR3A","LST1","S100A8","LYZ","FCER1A","CLEC10A"]
 if g in combo_rna.var_names]
sc.pl.umap(combo_rna, color=umap_genes, frameon=False, size=20, alpha=0.65, ncols=2)


# %% [code]
sc.pl.umap(combo_rna, color="celltype_harmonized", frameon=False, size=3, alpha=0.65, ncols=2)


# %% [code]
sc.pl.umap(combo_rna, color="tech", frameon=False, size=3, alpha=0.65, ncols=2)


# %% [code]
import pandas as pd

# pick an order that matches your marker panels left→right
desired_order = [
    "B",
    "CD4 T", "Naive CD4 T", "Memory CD4 T", "CD4 T helper", "Regulatory T",
    "Cytotoxic T", "Memory CD8 T", "Naive CD8 T",   # or your exact CD8 label
    "NK",
    "CD14+ monocyte", "CD16+ monocyte", "Monocyte",
    "Dendritic cell", "Plasmacytoid dendritic cell",
]

# keep only labels that exist, append leftovers at the end
present = combo_rna.obs[anno_key].astype(str).unique().tolist()
cats = [c for c in desired_order if c in present] + [c for c in present if c not in desired_order]

combo_rna.obs[anno_key] = pd.Categorical(combo_rna.obs[anno_key].astype(str), categories=cats, ordered=True)

# (important) filter marker dict so dotplot doesn't choke on missing genes
marker_genes_filt = {
    ct: [g for g in genes if g in combo_rna.var_names]
    for ct, genes in marker_genes.items()
}
marker_genes_filt = {ct: genes for ct, genes in marker_genes_filt.items() if len(genes) > 0}

sc.pl.dotplot(
    combo_rna,
    var_names=marker_genes_filt,
    groupby=anno_key,
    categories_order=cats,     # <- forces y-axis order
    standard_scale="var",
    dot_min=0.0,
    dot_max=1.0,
)


# %% [code]
# (6c) Optional: gene-set scores per lineage and visualize as violin (less noisy)
for ct, genes in marker_genes.items():
    g = [x for x in genes if x in combo_rna.var_names]
    if len(g) >= 2:
        sc.tl.score_genes(combo_rna, g, score_name=f"score_{ct}")

score_cols = [c for c in combo_rna.obs.columns if c.startswith("score_")]
sc.pl.violin(combo_rna, keys=score_cols, groupby=anno_key, stripplot=False, rotation=45)


# %% [code]


# %% [code]


# %% [code]


# %% [code]


# %% [code]


# %% [code]


# %% [code]


# %% [code]


# %% [code]


# %% [code]


# %% [code]
````
