## Supplemental Code for "Unifying multimodal single-cell data with a mixture-of-experts *β*-variational autoencoder framework": Fig06_TEAseq_trimodal.ipynb

````
# %% [markdown]
## UniVI manuscript - Figure 6 generation reproducible workflow
### TEA-seq RNA + ADT + ATAC latent embedding by cell type and modality; examples of predicted accessibility programs from RNA and ADT; predicted gene programs from ADT and ATAC; predicted protein expression from RNA and ATAC; alignment and reconstruction metrics

Andrew Ashford, Pathways + Omics Group, Oregon Health & Science University, Portland, OR - 12/14/2025

This Jupyter Notebook will house the end-to-end workflow to generate the panels in Figure 6 of our manuscript, "Unifying multimodal single-cell data with a mixture-of-experts β-variational autoencoder framework" which is currently being revised for Genome Research and is available currently on bioRxiv at the following link: https://www.biorxiv.org/content/10.1101/2025.02.28.640429v1.full

GitHub for the project - including a Quickstart guide - can be found at: https://github.com/Ashford-A/UniVI

Package is pip installable via the command: 
```bash
pip install univi
```


# %% [markdown]
#### Import modules

# %% [code]
# Import non-UniVI modules
import os
import sys
from pathlib import Path

import numpy as np
import pandas as pd
import scanpy as sc
import anndata as ad

import torch
from torch.utils.data import DataLoader, Subset

import scipy.sparse as sp

import snapatac2 as snap


# %% [code]
from sklearn.feature_extraction.text import TfidfTransformer
from sklearn.decomposition import TruncatedSVD
from sklearn.preprocessing import StandardScaler, normalize

from sklearn.neighbors import NearestNeighbors
from sklearn.metrics import confusion_matrix, accuracy_score, f1_score


# %% [code]
# Import required UniVI modules
from univi import (
    ModalityConfig,
    UniVIConfig,
    TrainingConfig,
    UniVIMultiModalVAE,
    matching,
    UniVITrainer,
    write_univi_latent,
    MultiModalDataset,
)

import univi as uv
import univi.evaluation as ue
import univi.plotting as up

# Double check UniVI module version
print("Installed version is univi v" + str(uv.__version__))


# %% [markdown]
### Specify device to use for model

Set "device" - preferably device should be "cuda" for speedier model implementation/training. Requires GPU and the correct packages/versions.


# %% [code]
print("torch:", torch.__version__)
print("torch.cuda.is_available():", torch.cuda.is_available())
device = "cuda" if torch.cuda.is_available() else "cpu"
print("Using device:", device)


# %% [markdown]
### Specify file paths

Where data lives.

# %% [code]
data_dir = Path("/home/groups/precepts/ashforda/UniVI_v2/UniVI_older-non_git/data/TEA-seq_data")

# Pattern for discovering RNA files → sample prefixes
RNA_SUFFIX = "_200M_cellranger-arc_filtered_feature_bc_matrix.h5"
ADT_SUFFIX = "_48M_adt_counts.csv.gz"
FRAG_SUFFIX = "_200M_atac_filtered_fragments.tsv.gz"
META_SUFFIX = "_200M_atac_filtered_metadata.csv.gz"

# Which sample to hold out (different patient)
# >>> CHANGE THIS TO THE ONE YOU WANT TO HOLD OUT <<<
HOLDOUT_PREFIX = "GSM5123953_X066-MP0C1W5_leukopak_perm-cells_tea"


# %% [markdown]
### Read in data

Read data into AnnData objects using the paths in the code chunk above.

# %% [code]
# ============================================================
# Helper functions
# ============================================================
def _to_dense(X):
    import numpy as np
    import scipy.sparse as sp
    if sp.issparse(X):
        return X.toarray()  # portable; don't use .A
    return np.asarray(X)

def strip_suffix(idx):
    """
    Drop trailing '-<number>' from 10x barcodes, e.g. 'AACT-1' -> 'AACT'.
    Works on any index-like object.
    """
    idx = pd.Index(idx.astype(str))
    return idx.str.replace(r"-\d+$", "", regex=True)

def find_sample_prefixes(data_dir: Path) -> list[str]:
    """Find all sample prefixes based on the RNA h5 naming convention."""
    rna_files = sorted(data_dir.glob(f"*{RNA_SUFFIX}"))
    prefixes = [f.name.replace(RNA_SUFFIX, "") for f in rna_files]
    return prefixes

def load_teaseq_sample(prefix: str, data_dir: Path):
    """
    Load RNA, ADT, ATAC (raw) for a single TEA-seq sample.

    Returns
    -------
    rna : AnnData
        Filtered RNA cells.
    adt : AnnData
        ADT counts restricted to the same cell barcodes as RNA.
    atac_raw : AnnData
        ATAC object after metadata join + basic QC (n_fragments >= 1500).
    """
    print(f"\n=== Loading sample: {prefix} ===")
    rna_h5         = data_dir / f"{prefix}{RNA_SUFFIX}"
    adt_counts_csv = data_dir / f"{prefix}{ADT_SUFFIX}"
    frag_tsv       = data_dir / f"{prefix}{FRAG_SUFFIX}"
    atac_meta_csv  = data_dir / f"{prefix}{META_SUFFIX}"

    # ---------- RNA ----------
    print("  [RNA] reading 10x h5 ...")
    m = sc.read_10x_h5(rna_h5)
    rna = m.copy()
    rna.var_names_make_unique()
    rna.obs_names = rna.obs_names.astype(str)
    print("  [RNA] shape:", rna.shape)

    # ---------- ADT ----------
    print("  [ADT] reading counts CSV ...")
    adt_df = pd.read_csv(adt_counts_csv, index_col=0)
    adt_df.index = adt_df.index.astype(str)
    print("  [ADT] raw shape:", adt_df.shape)

    # Align ADT to RNA barcodes
    rna_idx = pd.Index(rna.obs_names.astype(str))
    adt_idx = pd.Index(adt_df.index.astype(str))

    # 1) Try exact match first
    common = rna_idx.intersection(adt_idx)
    if len(common) > 0:
        common = common.sort_values()
        print(f"  [ADT] barcodes intersecting RNA (exact): {len(common)}")
        rna = rna[common].copy()
        adt_df = adt_df.loc[common].copy()
    else:
        # 2) Fall back to stripped barcodes (e.g. RNA has '-1', ADT doesn't)
        print("  [ADT] no exact barcode overlap; trying stripped barcodes ...")
        rna_stripped = strip_suffix(rna_idx)
        adt_stripped = strip_suffix(adt_idx)

        # Make sure mapping is unique (one row per stripped barcode)
        rna_map = pd.Series(rna_idx, index=rna_stripped)
        adt_map = pd.Series(adt_idx, index=adt_stripped)

        common_stripped = np.intersect1d(rna_stripped.unique(), adt_stripped.unique())
        print(f"  [ADT] barcodes intersecting RNA (stripped): {len(common_stripped)}")

        if len(common_stripped) == 0:
            raise ValueError(
                f"No overlapping barcodes between RNA and ADT for sample {prefix} "
                "(even after stripping suffixes)."
            )

        common_stripped = np.sort(common_stripped)

        rna_keep = rna_map.loc[common_stripped].values
        adt_keep = adt_map.loc[common_stripped].values

        rna = rna[rna_keep].copy()
        adt_df = adt_df.loc[adt_keep].copy()

        # store stripped barcode for debugging if you want
        rna.obs["barcode_stripped"] = common_stripped
        # for ADT we’ll carry the same order
        adt_df.index = common_stripped

    # Wrap ADT as AnnData
    adt = ad.AnnData(
        X=sp.csr_matrix(adt_df.values),
        obs=pd.DataFrame(index=adt_df.index.astype(str)),
        var=pd.DataFrame(index=adt_df.columns.astype(str)),
    )
    adt.var_names_make_unique()
    print("  [ADT] filtered shape (matching RNA):", adt.shape)

    # ---------- ATAC ----------
    print("  [ATAC] importing fragments via snapatac2 ...")
    atac_raw = snap.pp.import_data(
        fragment_file=str(frag_tsv),
        chrom_sizes=snap.genome.hg38,
        sorted_by_barcode=False,
    )
    print("  [ATAC] raw shape:", atac_raw.shape)

    meta = pd.read_csv(atac_meta_csv)
    meta = meta.set_index("barcodes")
    meta.index = meta.index.astype(str)

    common_ids = atac_raw.obs_names.intersection(meta.index)
    print("  [ATAC] cells with metadata:", len(common_ids), "of", atac_raw.n_obs)

    atac_raw = atac_raw[common_ids].copy()
    atac_raw.obs = atac_raw.obs.join(meta, how="left")

    # Simple QC
    if "n_fragments" in atac_raw.obs.columns:
        mask = atac_raw.obs["n_fragments"] >= 1500
        print("  [ATAC] keeping", mask.sum(), "cells with n_fragments >= 1500")
        atac_raw = atac_raw[mask].copy()

    # Tag sample id (after all filtering)
    rna.obs["sample_id"] = prefix
    adt.obs["sample_id"] = prefix
    atac_raw.obs["sample_id"] = prefix

    return rna, adt, atac_raw


# %% [code]
# ============================================================
# 1) Discover samples + split into train vs holdout
# ============================================================
all_prefixes = find_sample_prefixes(data_dir)
print("Found prefixes:", all_prefixes)

# Keep only TEA-seq samples for this pipeline
teaseq_prefixes = [p for p in all_prefixes if "_tea" in p]
print("TEA-seq prefixes:", teaseq_prefixes)

if HOLDOUT_PREFIX not in teaseq_prefixes:
    raise ValueError(
        f"HOLDOUT_PREFIX '{HOLDOUT_PREFIX}' not in TEA-seq prefixes: {teaseq_prefixes}"
    )

train_prefixes = [p for p in teaseq_prefixes if p != HOLDOUT_PREFIX]
print("Training prefixes:", train_prefixes)
print("Holdout prefix:", HOLDOUT_PREFIX)

rna_raw_dict  = {}
adt_raw_dict  = {}
atac_raw_dict = {}

for prefix in teaseq_prefixes:
    print(f"\n=== Loading sample: {prefix} ===")
    rna_s, adt_s, atac_s = load_teaseq_sample(prefix, data_dir)
    rna_raw_dict[prefix]  = rna_s
    adt_raw_dict[prefix]  = adt_s
    atac_raw_dict[prefix] = atac_s
    

# %% [code]
# ============================================================
# 2) Harmonize barcodes within each sample (RNA / ADT / ATAC)
#    and make them globally unique: <barcode>__<sample_id>
# ============================================================
def harmonize_barcodes_per_sample(rna, adt, atac_raw, sample_id: str):
    # strip 10x suffixes
    rna.obs_names = strip_suffix(rna.obs_names.to_series())
    adt.obs_names = strip_suffix(adt.obs_names.to_series())

    if "original_barcodes" not in atac_raw.obs.columns:
        raise KeyError("atac_raw.obs must contain 'original_barcodes' to map to 10x barcodes.")

    atac_raw.obs["barcode_10x"] = atac_raw.obs["original_barcodes"].astype(str)
    atac_raw.obs_names = strip_suffix(atac_raw.obs["barcode_10x"])

    # make globally unique obs_names
    rna.obs_names  = rna.obs_names + f"__{sample_id}"
    adt.obs_names  = adt.obs_names + f"__{sample_id}"
    atac_raw.obs_names = atac_raw.obs_names + f"__{sample_id}"

    return rna, adt, atac_raw


# %% [code]
for prefix in teaseq_prefixes:
    rna_raw_dict[prefix], adt_raw_dict[prefix], atac_raw_dict[prefix] = harmonize_barcodes_per_sample(
        rna_raw_dict[prefix],
        adt_raw_dict[prefix],
        atac_raw_dict[prefix],
        prefix,
    )


# %% [markdown]
### Preprocess each data type

# %% [code]
# ============================================================
# CONFIG
# ============================================================

# Preproc hyperparams
N_RNA_HVG       = 2000
N_ATAC_LSI      = 100
MIN_TILE_FRAC   = 0.005
MAX_TILE_FRAC   = 0.80
GAUSS_MAX_SCALE = 10.0
EPS             = 1e-6
RANDOM_STATE    = 42


# %% [code]
# ============================================================
# 3) RNA: learn HVGs from *training* samples only,
#    then apply to all (train + holdout)
# ============================================================
print("\n=== RNA preprocessing across samples ===")
rna_train_list = [rna_raw_dict[p].copy() for p in train_prefixes]

# Keep raw counts in .layers
for r in rna_train_list:
    if "counts" not in r.layers:
        r.layers["counts"] = r.X.copy()

# Concatenate training RNA for HVG selection
rna_train_concat = sc.concat(
    rna_train_list,
    join="outer",
    label="sample_id",
    keys=train_prefixes,
    index_unique=None,
)

print("  [RNA] concatenated training shape:", rna_train_concat.shape)

# Normalize + log1p
sc.pp.normalize_total(rna_train_concat, target_sum=1e4)
sc.pp.log1p(rna_train_concat)

# HVGs across training (Seurat v3 flavor, batched by sample)
sc.pp.highly_variable_genes(
    rna_train_concat,
    n_top_genes=N_RNA_HVG,
    flavor="seurat_v3",
    batch_key="sample_id",
)

rna_hvg_mask = rna_train_concat.var["highly_variable"].fillna(False)
rna_hvg_names = rna_train_concat.var_names[rna_hvg_mask].tolist()
print(f"  [RNA] selected {len(rna_hvg_names)} HVGs from training samples.")

# Scale on concatenated training HVGs (for Gaussian decoder)
rna_train_concat = rna_train_concat[:, rna_hvg_names].copy()
sc.pp.scale(rna_train_concat, max_value=GAUSS_MAX_SCALE)

# Get training means/stds for reuse (optional, nice but not strictly required)
X_rna_train = _to_dense(rna_train_concat.X).astype(np.float32)
rna_mean = X_rna_train.mean(axis=0, keepdims=True)
rna_std  = X_rna_train.std(axis=0, keepdims=True) + EPS


def preprocess_rna_with_hvgs(rna_raw: ad.AnnData, hvg_names, mean=None, std=None):
    rna = rna_raw.copy()
    if "counts" not in rna.layers:
        rna.layers["counts"] = rna.X.copy()

    sc.pp.normalize_total(rna, target_sum=1e4)
    sc.pp.log1p(rna)
    rna.layers["log1p"] = rna.X.copy()

    # Subset to HVGs (ignore genes missing in this sample)
    keep = [g for g in hvg_names if g in rna.var_names]
    rna = rna[:, keep].copy()

    # Scale; if global mean/std provided, reuse; else use per-sample
    if mean is not None and std is not None and rna.n_vars == mean.shape[1]:
        X = _to_dense(rna.X).astype(np.float32)
        Xz = (X - mean) / std
        rna.X = Xz.astype(np.float32)
    else:
        sc.pp.scale(rna, max_value=GAUSS_MAX_SCALE)

    return rna


rna_train_processed = []
for prefix in train_prefixes:
    print(f"  [RNA] preprocessing training sample {prefix}")
    rna_train_processed.append(preprocess_rna_with_hvgs(
        rna_raw_dict[prefix], rna_hvg_names, mean=rna_mean, std=rna_std
    ))

print(f"  [RNA] preprocessing holdout sample {HOLDOUT_PREFIX}")
rna_holdout = preprocess_rna_with_hvgs(
    rna_raw_dict[HOLDOUT_PREFIX], rna_hvg_names, mean=rna_mean, std=rna_std
)

# Concatenate processed training RNA
rna_train = ad.concat(
    rna_train_processed,
    join="outer",
    label="sample_id",
    keys=train_prefixes,
    index_unique=None,
)
print("Final rna_train shape:", rna_train.shape)
print("Holdout rna_holdout shape:", rna_holdout.shape)


# %% [code]
# ============================================================
# 4) ADT: CLR + z, shared panel based on training intersection
# ============================================================
print("\n=== ADT preprocessing across samples ===")

# First, figure out shared ADT panel from training samples
adt_train_list_raw = [adt_raw_dict[p] for p in train_prefixes]
adt_panels = [set(a.var_names) for a in adt_train_list_raw]
adt_panel = sorted(set.intersection(*adt_panels))
print(f"  [ADT] panel intersection across training samples: {len(adt_panel)} features")

def preprocess_adt_panel(adt_raw: ad.AnnData, panel: list[str]):
    adt = adt_raw.copy()
    # align to panel (drop anything else)
    keep = [f for f in panel if f in adt.var_names]
    adt = adt[:, keep].copy()
    if "counts" not in adt.layers:
        adt.layers["counts"] = adt.X.copy()

    X_counts = _to_dense(adt.layers["counts"]).astype(float)

    # CLR per cell
    X_log = np.log1p(X_counts + EPS)
    X_clr = X_log - X_log.mean(axis=1, keepdims=True)

    # Per-feature z across cells
    mean_adt = X_clr.mean(axis=0, keepdims=True)
    std_adt  = X_clr.std(axis=0, keepdims=True) + EPS
    X_clr_z  = (X_clr - mean_adt) / std_adt

    adt.layers["clr"]   = X_clr.astype(np.float32)
    adt.layers["clr_z"] = X_clr_z.astype(np.float32)
    adt.X = X_clr_z.astype(np.float32)
    return adt

adt_train_processed = []
for prefix in train_prefixes:
    print(f"  [ADT] preprocessing training sample {prefix}")
    adt_train_processed.append(preprocess_adt_panel(adt_raw_dict[prefix], adt_panel))

print(f"  [ADT] preprocessing holdout sample {HOLDOUT_PREFIX}")
adt_holdout = preprocess_adt_panel(adt_raw_dict[HOLDOUT_PREFIX], adt_panel)

adt_train = ad.concat(
    adt_train_processed,
    join="outer",
    label="sample_id",
    keys=train_prefixes,
    index_unique=None,
)
print("Final adt_train shape:", adt_train.shape)
print("Holdout adt_holdout shape:", adt_holdout.shape)


# %% [code]
# ============================================================
# 5) ATAC: global tiles → TF-IDF → LSI → z
#    SVD / IDF learned on *training* cells only, then applied to holdout
# ============================================================
print("\n=== ATAC preprocessing across samples (global TF-IDF + LSI from training) ===")

# 5.1 Add tile matrix for all samples (train + holdout) and check that
#     the number/order of tiles match across samples.
atac_all_prefixes = teaseq_prefixes

tile_mats = {}
n_cells_total_train = 0
global_tile_counts = None
tile_var_names_ref = None

for prefix in atac_all_prefixes:
    print(f"  [ATAC] add_tile_matrix for sample {prefix}")
    atac_raw = atac_raw_dict[prefix]
    snap.pp.add_tile_matrix(atac_raw)

    X = atac_raw.X
    if not sp.issparse(X):
        X = sp.csr_matrix(X)

    # store for later
    tile_mats[prefix] = (atac_raw, X)

    # sanity: ensure identical tile feature space across samples
    if tile_var_names_ref is None:
        tile_var_names_ref = atac_raw.var_names.copy()
    else:
        if not np.array_equal(tile_var_names_ref, atac_raw.var_names):
            raise ValueError(f"Tile names differ for sample {prefix}; "
                             "this code assumes a shared tile grid across samples.")

    # accumulate global tile counts / cells for *training* samples only
    if prefix in train_prefixes:
        n_cells = X.shape[0]
        n_cells_total_train += n_cells
        tile_on = (X > 0).astype(np.int64)
        counts = np.asarray(tile_on.sum(axis=0)).ravel()
        if global_tile_counts is None:
            global_tile_counts = counts
        else:
            global_tile_counts += counts

print("  [ATAC] total training cells for TF-IDF/LSI:", n_cells_total_train)

# 5.2 Global tile filtering based on training
tile_frac = global_tile_counts / float(n_cells_total_train)
keep_tiles = (tile_frac > MIN_TILE_FRAC) & (tile_frac < MAX_TILE_FRAC)
print(
    f"  [ATAC] Filtering tiles by freq (training only): "
    f"keeping {keep_tiles.sum()} / {len(tile_frac)} "
    f"({keep_tiles.sum() / len(tile_frac):.2%}) tiles"
)

tile_names_kept = tile_var_names_ref[keep_tiles]

# 5.3 Build big TF-IDF matrix for training cells only (kept tiles)
X_tfidf_train_list = []
for prefix in train_prefixes:
    atac_raw, X = tile_mats[prefix]
    X = X[:, keep_tiles]
    # TF: per-cell L1 norm
    tf = normalize(X, norm="l1", axis=1)
    X_tfidf_train_list.append(tf)

X_tfidf_train = sp.vstack(X_tfidf_train_list).tocsr()
n_cells_train, n_feats_kept = X_tfidf_train.shape
print(f"  [ATAC] TF-IDF training matrix shape: {n_cells_train} × {n_feats_kept}")

# Global IDF over training cells
df = np.asarray((X_tfidf_train > 0).sum(axis=0)).ravel()  # tf>0 is same as X>0 here
idf = np.log1p(n_cells_train / (1.0 + df))

# Apply IDF and per-cell L2 norm to training TF matrix
X_tfidf_train = X_tfidf_train.multiply(idf).astype(np.float32)
X_tfidf_train = normalize(X_tfidf_train, norm="l2", axis=1)

# 5.4 Fit global LSI (TruncatedSVD) on training TF-IDF
print(f"  [ATAC] Fitting TruncatedSVD with {N_ATAC_LSI} components on training TF-IDF ...")
svd = TruncatedSVD(n_components=N_ATAC_LSI, random_state=RANDOM_STATE)
lsi_train = svd.fit_transform(X_tfidf_train)  # (cells_train, N_ATAC_LSI)

# Center & compute global mean/std for LSI dims
lsi_train = lsi_train.astype(np.float32)
lsi_mean = lsi_train.mean(axis=0, keepdims=True)
lsi_std  = lsi_train.std(axis=0, keepdims=True) + EPS

# 5.5 Function to apply TF-IDF + global LSI + z to any sample
def preprocess_atac_with_global_lsi(atac_raw, X_tile, keep_tiles_mask, idf_vec, svd_model, lsi_mean, lsi_std):
    X = X_tile[:, keep_tiles_mask]
    if not sp.issparse(X):
        X = sp.csr_matrix(X)

    # TF
    tf = normalize(X, norm="l1", axis=1)
    # TF-IDF (global IDF from training)
    X_tfidf = tf.multiply(idf_vec).astype(np.float32)
    X_tfidf = normalize(X_tfidf, norm="l2", axis=1)

    # LSI transform using global SVD
    lsi = svd_model.transform(X_tfidf).astype(np.float32)

    # center & z-score using training stats
    lsi_centered = lsi - lsi_mean
    lsi_z = lsi_centered / lsi_std

    atac = ad.AnnData(
        X=lsi_z.astype(np.float32),
        obs=atac_raw.obs.copy(),
        var=pd.DataFrame(index=[f"LSI_{i+1}" for i in range(lsi.shape[1])]),
    )
    atac.obsm["X_lsi"] = lsi.astype(np.float32)
    return atac

# 5.6 Apply to each training sample
atac_train_processed = []
for prefix in train_prefixes:
    print(f"  [ATAC] preprocessing training sample {prefix}")
    atac_raw, X_tile = tile_mats[prefix]
    atac_proc = preprocess_atac_with_global_lsi(
        atac_raw, X_tile,
        keep_tiles_mask=keep_tiles,
        idf_vec=idf,
        svd_model=svd,
        lsi_mean=lsi_mean,
        lsi_std=lsi_std,
    )
    atac_train_processed.append(atac_proc)

print(f"  [ATAC] preprocessing holdout sample {HOLDOUT_PREFIX}")
atac_raw_holdout, X_tile_holdout = tile_mats[HOLDOUT_PREFIX]
atac_holdout = preprocess_atac_with_global_lsi(
    atac_raw_holdout, X_tile_holdout,
    keep_tiles_mask=keep_tiles,
    idf_vec=idf,
    svd_model=svd,
    lsi_mean=lsi_mean,
    lsi_std=lsi_std,
)

# 5.7 Concatenate processed training ATAC
atac_train = ad.concat(
    atac_train_processed,
    join="outer",
    label="sample_id",
    keys=train_prefixes,
    index_unique=None,
)

print("Final atac_train shape:", atac_train.shape)
print("Holdout atac_holdout shape:", atac_holdout.shape)


# %% [code]
# Subset all the overlapping cells in the same order for all modalities for train and holdout data
common_train = (rna_train.obs_names.intersection(adt_train.obs_names).intersection(atac_train.obs_names))
common_train = common_train.sort_values()

print(f"For the training data there are common {len(common_train)} tri-modal cells.")

rna_train  = rna_train[common_train].copy()
adt_train  = adt_train[common_train].copy()
atac_train = atac_train[common_train].copy()
    

# %% [code]
# Subset all the overlapping cells in the same order for all modalities for train and holdout data
common_holdout = (rna_holdout.obs_names.intersection(adt_holdout.obs_names).intersection(atac_holdout.obs_names))
common_holdout = common_holdout.sort_values()

print(f"For the holdout data there are common {len(common_holdout)} tri-modal cells.")

rna_holdout  = rna_holdout[common_holdout].copy()
adt_holdout  = adt_holdout[common_holdout].copy()
atac_holdout = atac_holdout[common_holdout].copy()


# %% [code]
# ============================================================
# 6) Final objects for UniVI training + held-out evaluation
# ============================================================
print("\n=== Final multimodal objects ===")
print("rna_train :", rna_train.shape)
print("adt_train :", adt_train.shape)
print("atac_train:", atac_train.shape)
print("rna_holdout :", rna_holdout.shape)
print("adt_holdout :", adt_holdout.shape)
print("atac_holdout:", atac_holdout.shape)

# For UniVI training:
adata_dict_train = {
    "rna":  rna_train,
    "adt":  adt_train,
    "atac": atac_train,
}

# For held-out evaluation (RNA→ADT / RNA→ATAC prediction):
adata_dict_holdout = {
    "rna":  rna_holdout,
    "adt":  adt_holdout,
    "atac": atac_holdout,
}


# %% [markdown]
#### Optionally, save the above processed AnnData objects

# %% [code]
'''
output_dir = './results/TEA-seq_integration_reproducibility/processed_data/'

processed_training_data_filename = 'combined_pp_teaseq_training_data.h5ad'
processed_heldout_data_filename = 'combined_pp_teaseq_heldout_data.h5ad'
'''

# %% [code]
# Add components to the AnnData objects that will make them easy to distinguish and split later
rna_train.var['feature_type'] = 'rna'
adt_train.var['feature_type'] = 'adt'
atac_train.var['feature_type'] = 'atac'

rna_holdout.var['feature_type'] = 'rna'
adt_holdout.var['feature_type'] = 'adt'
atac_holdout.var['feature_type'] = 'atac'


# %% [code]
# Combine objects and then save; make sure to merge along feature axis (columns), not cells
combo_train = ad.concat(
    {"rna": rna_train, "adt": adt_train, "atac": atac_train},
    axis=1,            # concatenate features
    join="outer",
    label="feature_type",  # this adds `var.feature_type`
    merge="same",
)

combo_holdout = ad.concat(
    {"rna": rna_holdout, "adt": adt_holdout, "atac": atac_holdout},
    axis=1,
    join="outer",
    label="feature_type",
    merge="same",
)

# verify modalities stored
print(combo_train)
print(combo_train.var["feature_type"].value_counts())


# %% [code]
#combo_train.write_h5ad(output_dir + processed_training_data_filename, compression='gzip')


# %% [code]
#combo_holdout.write_h5ad(output_dir + processed_heldout_data_filename, compression='gzip')


# %% [markdown]
### Instantiate model params etc..

# %% [code]
univi_cfg = UniVIConfig(
    latent_dim=30,
    #beta=1.30,
    beta=1.15,
    #gamma=1.30,
    gamma=1.45,
    encoder_dropout=0.10,
    decoder_dropout=0.00,
    encoder_batchnorm=True,
    decoder_batchnorm=False,
    kl_anneal_start=0,
    kl_anneal_end=50,         # ramp KL up over first 50 epochs
    align_anneal_start=25,    # let reconstructions stabilize a bit first
    align_anneal_end=75,
    modalities=[
        ModalityConfig(
            name="rna",
            input_dim=rna_train.n_vars,
            encoder_hidden=[512, 256, 128],
            decoder_hidden=[128, 256, 512],
            likelihood="gaussian",
            #recon_weight=1.00,
            recon_weight=1.00,
        ),
        ModalityConfig(
            name="adt",
            input_dim=adt_train.n_vars,
            encoder_hidden=[128, 64],
            decoder_hidden=[64, 128],
            likelihood="gaussian",
            recon_weight=2.00,
            #recon_weight=1.00,
        ),
        ModalityConfig(
            name="atac",
            input_dim=atac_train.n_vars,  # n_lsi
            encoder_hidden=[128, 64],
            decoder_hidden=[64, 128],
            likelihood="gaussian",
            recon_weight=1.45,
            #recon_weight=1.00,
        ),
    ],
)

train_cfg = TrainingConfig(
    n_epochs=3000,
    batch_size=256,
    lr=1e-4,
    weight_decay=1e-4,
    device=device,      # "cuda"
    log_every=20,
    grad_clip=5.0,
    num_workers=0,
    seed=42,
    early_stopping=True,
    patience=200,
    min_delta=0.0,
)


# %% [markdown]
### Set up Multiome RNA + ATAC bridge data train/val/test splits

# %% [code]
print(adata_dict_train)


# %% [code]
print(adata_dict_holdout)


# %% [code]
dataset = MultiModalDataset(
    adata_dict=adata_dict_train,
    X_key="X",
    device=train_cfg.device,
)

n_cells = dataset.n_cells
indices = np.arange(n_cells)
rng = np.random.default_rng(42)
rng.shuffle(indices)

frac_train = 0.8
frac_val   = 0.1
n_train = int(frac_train * n_cells)
n_val   = int(frac_val * n_cells)

train_idx = indices[:n_train]
val_idx   = indices[n_train:n_train + n_val]
test_idx  = indices[n_train + n_val:]

train_ds = Subset(dataset, train_idx)
val_ds   = Subset(dataset, val_idx)

train_loader = DataLoader(
    train_ds,
    batch_size=train_cfg.batch_size,
    shuffle=True,
    num_workers=train_cfg.num_workers,
)

val_loader = DataLoader(
    val_ds,
    batch_size=train_cfg.batch_size,
    shuffle=False,
    num_workers=train_cfg.num_workers,
)


# %% [code]
print(len(train_idx))
print(len(val_idx))
print(len(test_idx))


# %% [code]
#model = UniVIMultiModalVAE(univi_cfg).to(train_cfg.device)
model = UniVIMultiModalVAE(
    univi_cfg,
    loss_mode="v1",      # cross-recon + cross-posterior alignment
    #loss_mode="lite"
    #v1_recon="cross",   # full k→j cross-recon
    v1_recon="avg",
    #v1_recon_mix=0.5,
    normalize_v1_terms=True,
    #recon_normalize_by_dim=True,
    #recon_dim_power=0.33333,
).to(device)


# %% [code]
trainer = UniVITrainer(
    model=model,
    train_loader=train_loader,
    val_loader=val_loader,
    train_cfg=train_cfg,
    device=train_cfg.device,
)


# %% [markdown]
### Train model

# %% [code]
# ---------- train ----------
history = trainer.fit()


# %% [code]
import matplotlib.pyplot as plt

# Quick training curves
fig, ax = plt.subplots()
ax.plot(history["train_loss"], label="train")
ax.plot(history["val_loss"], label="val")
ax.set_xlabel("Epoch")
ax.set_ylabel("Loss")
ax.set_title("UniVI Multiome training curves")
ax.legend()
plt.tight_layout()
plt.show()

fig, ax = plt.subplots()
ax.plot(history["beta"], label="beta")
ax.plot(history["gamma"], label="gamma")
ax.set_xlabel("Epoch")
ax.set_ylabel("Weight")
ax.set_title("KL / alignment annealing")
ax.legend()
plt.tight_layout()
plt.show()


# %% [markdown]
### Save model/load to best epoch checkpoint

# %% [code]
beta_used = "1.15"
gamma_used = "1.45"
latent_dims_used = "30"


# %% [code]
output_dir = f'./results/univi_TEA-seq_Figure_6_reproducibility_beta-{beta_used}_gamma-{gamma_used}_latent_dims-{latent_dims_used}_gaussian_all/'
out_file = f"trained_model_beta-{beta_used}_gamma-{gamma_used}_latent_dims-{latent_dims_used}_gaussian_both.pt"


# %% [code]
from dataclasses import asdict

os.makedirs(output_dir, exist_ok=True)


# %% [code]
ckpt_path = output_dir + out_file


# %% [code]
rna = rna_holdout.copy()
adt = adt_holdout.copy()
atac = atac_holdout.copy()


# %% [code]
print(rna)


# %% [code]
n_hvg = len(rna.var_names)
target_sum = 1e4
n_lsi = len(atac.var_names)
rna_hvg_names = rna.var_names


# %% [code]
# after training
#history = trainer.fit()

import sys, platform, hashlib, numpy as np, torch

def _hash_list(x):
    h = hashlib.sha256()
    for s in x:
        h.update(str(s).encode())
        h.update(b"\n")
    return h.hexdigest()

ckpt = {
    # core
    "state_dict": trainer.model.state_dict(),
    "univi_cfg": asdict(univi_cfg),

    # best model info
    "best_epoch": trainer.best_epoch,
    "best_val_loss": float(trainer.best_val_loss),

    # splits (STORE THESE!)
    "splits": {
        "train": np.asarray(train_idx, dtype=np.int64),
        "val":   np.asarray(val_idx, dtype=np.int64),
        "test":  np.asarray(test_idx, dtype=np.int64),
    },

    # preprocessing + feature sets (examples; include what you actually use)
    "preproc": {
        "rna":  {"layer": "counts", "n_hvg": n_hvg, "target_sum": target_sum, "log1p": True, "zscore": True},
        "adt":  {"layer": "counts", "clr": True, "zscore": True},
        "atac": {"layer": "counts", "tfidf": True, "n_lsi": n_lsi, "lsi_drop_first": True, "zscore": True},
    },
    "features": {
        "rna_hvgs": list(rna_hvg_names),          # list[str]
        "adt_names": list(adt.var_names),         # list[str] (or your selected panel)
        "atac_peaks": list(atac.var_names),       # list[str] (or whatever LSI used)
    },

    # provenance checks
    "data_fingerprint": {
        "rna_obs_hash": _hash_list(rna.obs_names),
        "rna_var_hash": _hash_list(rna.var_names),
        "adt_obs_hash": _hash_list(adt.obs_names),
        "adt_var_hash": _hash_list(adt.var_names),
        "atac_obs_hash": _hash_list(atac.obs_names),
        "atac_var_hash": _hash_list(atac.var_names),
    },

    # environment versions
    "versions": {
        "python": sys.version,
        "platform": platform.platform(),
        "torch": torch.__version__,
        "numpy": np.__version__,
        "univi": uv.__version__,
        "anndata": ad.__version__,
        "scanpy": sc.__version__,
    },

    # history (optional)
    "history": history,
}

torch.save(ckpt, ckpt_path)
print("Saved best model to:", ckpt_path)


# %% [code]
# Later to reload model
import torch
from univi.config import UniVIConfig, ModalityConfig
from univi.models.univi import UniVIMultiModalVAE

print("torch:", torch.__version__)
print("torch.cuda.is_available():", torch.cuda.is_available())
device = "cuda" if torch.cuda.is_available() else "cpu"
print("Using device:", device)

#device = "cuda"  # or "cuda" if available


# %% [code]
ckpt = torch.load(
    #output_dir + out_file,
    ckpt_path,
    map_location=device,
    weights_only=False,
)


# %% [code]
# ---- Rebuild UniVIConfig, making sure modalities are ModalityConfig objects ----
cfg_dict = ckpt["univi_cfg"]

# If this is an OmegaConf object or similar, make sure it's a plain dict
try:
    from omegaconf import DictConfig, OmegaConf
    if isinstance(cfg_dict, DictConfig):
        cfg_dict = OmegaConf.to_container(cfg_dict, resolve=True)
except ImportError:
    pass


# Now rehydrate each modality
modalities = [ModalityConfig(**m) for m in cfg_dict["modalities"]]
cfg_dict = {**cfg_dict, "modalities": modalities}

univi_cfg_loaded = UniVIConfig(**cfg_dict)

# ---- Rebuild model + load weights ----
model_loaded = UniVIMultiModalVAE(univi_cfg_loaded).to(device)
model_loaded.load_state_dict(ckpt["state_dict"])

print("Best epoch was:", ckpt.get("best_epoch"), "val loss =", ckpt.get("best_val_loss"))


# %% [code]
print(sorted(ckpt.keys()))
#print(ckpt['splits'])


# %% [markdown]
#### Evaluate model

# %% [code]
# These should be the same cells across modalities
assert np.array_equal(rna.obs_names, adt.obs_names)
assert np.array_equal(rna.obs_names, atac.obs_names)


# %% [code]
rna_test_adata = rna.copy()
adt_test_adata = adt.copy()
atac_test_adata = atac.copy()


# %% [code]
print(rna_test_adata)
print(adt_test_adata)
print(atac_test_adata)


# %% [code]
import scanpy as sc

# Make plots a bit bigger / nicer
sc.settings.set_figure_params(dpi=100, figsize=(10, 8))
#sc.settings.set_figure_params()


# %% [code]
# ============================
# UniVI TEA-seq evaluation (RNA / ADT / ATAC) – label-free, tri-modal
# ============================

import matplotlib.pyplot as plt
import seaborn as sns

from univi import evaluation as univi_eval
from univi import plotting as univi_plot  # not strictly needed, but left for convenience

from sklearn.metrics import silhouette_score
from sklearn.neighbors import NearestNeighbors
from sklearn.metrics import pairwise_distances

# -----------------------------------------
# CONFIG
# -----------------------------------------
FIGDIR = output_dir + "/figures"
os.makedirs(FIGDIR, exist_ok=True)


# %% [code]
# -----------------------------------------
# 0. Sanity checks
# -----------------------------------------
assert (
    rna_test_adata.n_obs == adt_test_adata.n_obs == atac_test_adata.n_obs
), "RNA / ADT / ATAC TEST sets must have same #cells"
assert np.array_equal(rna_test_adata.obs_names, adt_test_adata.obs_names), (
    "RNA and ADT obs_names must match 1:1 for pairwise metrics."
)
assert np.array_equal(rna_test_adata.obs_names, atac_test_adata.obs_names), (
    "RNA and ATAC obs_names must match 1:1 for pairwise metrics."
)

print(f"Test cells: {rna_test_adata.n_obs}")


# %% [code]
'''
for m, x in x_dict.items():
    if x is None: 
        continue
    print(m, "x.shape[1] =", x.shape[1], "cfg.input_dim =", model_loaded.mod_cfg_by_name[m].input_dim)
'''

# %% [code]
# -----------------------------------------
# 1. Encode latent embeddings for test sets
# -----------------------------------------
print("\nEncoding test sets into UniVI latent space...")

z_rna  = univi_eval.encode_adata(model, rna_test_adata,  modality="rna",  device=device)
z_adt  = univi_eval.encode_adata(model, adt_test_adata,  modality="adt",  device=device)
z_atac = univi_eval.encode_adata(model, atac_test_adata, modality="atac", device=device)

rna_test_adata.obsm["X_univi"]  = z_rna
adt_test_adata.obsm["X_univi"]  = z_adt
atac_test_adata.obsm["X_univi"] = z_atac

print("Latent shapes (test):")
print("  RNA :", z_rna.shape)
print("  ADT :", z_adt.shape)
print("  ATAC:", z_atac.shape)


# %% [code]
# -----------------------------------------
# 2. FOSCTTM (pairwise alignment)
# -----------------------------------------
'''
print("\nComputing FOSCTTM for each modality pair (lower = better)...")
fos_rna_adt  = univi_eval.compute_foscttm(z_rna,  z_adt)
fos_rna_atac = univi_eval.compute_foscttm(z_rna,  z_atac)
fos_adt_atac = univi_eval.compute_foscttm(z_adt,  z_atac)

print(f"  RNA  vs ADT : {fos_rna_adt:.4f}")
print(f"  RNA  vs ATAC: {fos_rna_atac:.4f}")
print(f"  ADT  vs ATAC: {fos_adt_atac:.4f}")

plt.figure(figsize=(4, 4))
pairs = ["RNA–ADT", "RNA–ATAC", "ADT–ATAC"]
vals = [fos_rna_adt, fos_rna_atac, fos_adt_atac]
sns.barplot(x=pairs, y=vals)
plt.ylabel("FOSCTTM (mean)")
plt.title("Tri-modal FOSCTTM (lower = better)")
#plt.tight_layout()
plt.savefig(os.path.join(FIGDIR, "foscttm_barplot.png"))
plt.show()
plt.close()
'''

# Added error bars
def foscttm_per_cell(z_src, z_tgt, metric="euclidean"):
    """
    Compute per-cell FOSCTTM:
    fraction of target cells closer than the true match (1:1 pairing).
    Assumes z_src[i] ↔ z_tgt[i] is the true pair.
    """
    assert z_src.shape[0] == z_tgt.shape[0], "Need 1:1 pairing for FOSCTTM"
    dists = pairwise_distances(z_src, z_tgt, metric=metric)        # (n_src, n_tgt)
    n = dists.shape[0]
    true_d = dists[np.arange(n), np.arange(n)]                     # (n,)
    # For each i, fraction of j with d(i, j) < d(i, true_match)
    per_cell = (dists < true_d[:, None]).mean(axis=1)
    return per_cell


print("\nComputing FOSCTTM for each modality pair (lower = better)...")

fos_rna_adt_cells  = foscttm_per_cell(z_rna,  z_adt)
fos_rna_atac_cells = foscttm_per_cell(z_rna,  z_atac)
fos_adt_atac_cells = foscttm_per_cell(z_adt,  z_atac)

fos_rna_adt_mean  = fos_rna_adt_cells.mean()
fos_rna_atac_mean = fos_rna_atac_cells.mean()
fos_adt_atac_mean = fos_adt_atac_cells.mean()

# Standard error of the mean (SEM)
n = z_rna.shape[0]
fos_rna_adt_sem  = fos_rna_adt_cells.std(ddof=1)  / np.sqrt(n)
fos_rna_atac_sem = fos_rna_atac_cells.std(ddof=1) / np.sqrt(n)
fos_adt_atac_sem = fos_adt_atac_cells.std(ddof=1) / np.sqrt(n)

print(f"  RNA  vs ADT : {fos_rna_adt_mean:.4f} ± {fos_rna_adt_sem:.4f} (SEM)")
print(f"  RNA  vs ATAC: {fos_rna_atac_mean:.4f} ± {fos_rna_atac_sem:.4f} (SEM)")
print(f"  ADT  vs ATAC: {fos_adt_atac_mean:.4f} ± {fos_adt_atac_sem:.4f} (SEM)")

pairs = ["RNA–ADT", "RNA–ATAC", "ADT–ATAC"]
vals  = [fos_rna_adt_mean, fos_rna_atac_mean, fos_adt_atac_mean]
errs  = [fos_rna_adt_sem,  fos_rna_atac_sem,  fos_adt_atac_sem]

plt.figure(figsize=(4, 4))
plt.bar(pairs, vals, yerr=errs, capsize=5)
plt.ylabel("FOSCTTM (mean ± SEM)")
plt.title("Tri-modal FOSCTTM (lower = better)")
plt.savefig(os.path.join(FIGDIR, "foscttm_barplot.png"))
plt.show()
plt.close()


# %% [code]
x_dict = {
    "rna":  torch.tensor(rna_test_adata.X,  dtype=torch.float32, device=device),   # (B, n_genes)
    "adt":  torch.tensor(adt_test_adata.X,  dtype=torch.float32, device=device),   # (B, n_adts)
    "atac": torch.tensor(atac_test_adata.X, dtype=torch.float32, device=device),   # (B, n_peaks)
}

# %% [code]
out = model_loaded(x_dict, epoch=305)
print({k: float(v.detach().cpu()) for k,v in out["recon_per_modality"].items()})

'''
# ---- Rebuild UniVIConfig, making sure modalities are ModalityConfig objects ----
cfg_dict = ckpt["univi_cfg"]

# If this is an OmegaConf object or similar, make sure it's a plain dict
try:
    from omegaconf import DictConfig, OmegaConf
    if isinstance(cfg_dict, DictConfig):
        cfg_dict = OmegaConf.to_container(cfg_dict, resolve=True)
except ImportError:
    pass


# Now rehydrate each modality
modalities = [ModalityConfig(**m) for m in cfg_dict["modalities"]]
cfg_dict = {**cfg_dict, "modalities": modalities}

univi_cfg_loaded = UniVIConfig(**cfg_dict)

# ---- Rebuild model + load weights ----
model_loaded = UniVIMultiModalVAE(univi_cfg_loaded).to(device)
model_loaded.load_state_dict(ckpt["state_dict"])

print("Best epoch was:", ckpt.get("best_epoch"), "val loss =", ckpt.get("best_val_loss"))
'''

# %% [code]
# -----------------------------------------
# 3. Modality mixing (all three modalities)
# -----------------------------------------
Z_joint = np.concatenate([z_rna, z_adt, z_atac], axis=0)
modality_labels = np.array(
    ["rna"]  * z_rna.shape[0]
    + ["adt"]  * z_adt.shape[0]
    + ["atac"] * z_atac.shape[0]
)

mixing_score = univi_eval.compute_modality_mixing(
    Z_joint,
    modality_labels,
    k=50,
)
print(f"\nGlobal modality mixing score (RNA/ADT/ATAC, k=30): {mixing_score:.3f}")

# kNN modality composition heatmap
print("Computing kNN modality composition...")
k = 30
nn = NearestNeighbors(n_neighbors=k + 1)
nn.fit(Z_joint)
_, idx = nn.kneighbors(Z_joint)

idx_neighbors = idx[:, 1:]  # drop self
neighbor_mods = modality_labels[idx_neighbors]

modalities = np.array(["rna", "adt", "atac"])
comp_matrix = np.zeros((len(modalities), len(modalities)))  # row = center, col = neighbor

for i, m_center in enumerate(modalities):
    mask_center = modality_labels == m_center
    neigh_for_center = neighbor_mods[mask_center].reshape(-1)
    for j, m_neigh in enumerate(modalities):
        comp_matrix[i, j] = (neigh_for_center == m_neigh).mean()

same_mod_frac = (neighbor_mods == modality_labels[:, None]).mean()
print(f"  Fraction of neighbors with same modality (k={k}): {same_mod_frac:.3f}")


# %% [code]
plt.figure(figsize=(5, 4))
sns.heatmap(
    comp_matrix,
    annot=True,
    fmt=".2f",
    xticklabels=modalities,
    yticklabels=modalities,
    cmap="viridis",
)
plt.xlabel("Neighbor modality")
plt.ylabel("Center modality")
plt.title(f"kNN modality composition (k={k})")
#plt.tight_layout()
plt.savefig(os.path.join(FIGDIR, "knn_modality_composition.png"))
plt.show()
plt.close()


# %% [code]
# -----------------------------------------
# 4. UMAP on UniVI latent (tri-modal)
# -----------------------------------------
print("\nBuilding tri-modal UMAP on UniVI latent...")

# Tag each test set with modality
rna_tmp  = rna_test_adata.copy()
adt_tmp  = adt_test_adata.copy()
atac_tmp = atac_test_adata.copy()

rna_tmp.obs["univi_source"]  = "rna"
adt_tmp.obs["univi_source"]  = "adt"
atac_tmp.obs["univi_source"] = "atac"

combined = rna_tmp.concatenate(
    adt_tmp,
    atac_tmp,
    join="outer",
    batch_key="univi_batch",
    batch_categories=["rna", "adt", "atac"],
    index_unique=None,
)

# ensure latent is correctly stacked
combined.obsm["X_univi"] = np.vstack([
    rna_test_adata.obsm["X_univi"],
    adt_test_adata.obsm["X_univi"],
    atac_test_adata.obsm["X_univi"],
])

# neighbors / UMAP / Leiden
sc.pp.neighbors(combined, use_rep="X_univi", n_neighbors=30)
sc.tl.umap(combined)
sc.tl.leiden(combined, key_added="univi_leiden", resolution=0.45)


# %% [code]
# UMAP colored by modality
sc.pl.umap(
    combined,
    color="univi_source",
    size=3,
    alpha=0.8,
    show=False,
)
plt.savefig(os.path.join(FIGDIR, "umap_tri_modal_modality.png"), bbox_inches="tight")
plt.show()
plt.close()

# UMAP colored by Leiden clusters (pseudo-clusters)
sc.pl.umap(
    combined,
    color="univi_leiden",
    size=3,
    alpha=0.8,
    show=False,
)
plt.savefig(os.path.join(FIGDIR, "umap_tri_modal_leiden.png"), bbox_inches="tight")
plt.show()
plt.close()

# Per-modality UMAPs, colored by Leiden
for mod in ["rna", "adt", "atac"]:
    sub = combined[combined.obs["univi_source"] == mod].copy()
    sc.pl.umap(
        sub,
        color="univi_leiden",
        size=3,
        alpha=0.8,
        show=False,
    )
    plt.savefig(
        os.path.join(FIGDIR, f"umap_{mod}_only_leiden.png"),
        bbox_inches="tight",
    )
    plt.show()
    plt.close()
    

# %% [code]
# -----------------------------------------
# 5. Latent geometry diagnostics
# -----------------------------------------
print("\nLatent geometry diagnostics...")

def latent_norms(z, label):
    norms = np.linalg.norm(z, axis=1)
    return norms, np.repeat(label, len(norms))

norm_rna,  lab_rna  = latent_norms(z_rna,  "RNA")
norm_adt,  lab_adt  = latent_norms(z_adt,  "ADT")
norm_atac, lab_atac = latent_norms(z_atac, "ATAC")

norms_all = np.concatenate([norm_rna, norm_adt, norm_atac])
labs_all  = np.concatenate([lab_rna, lab_adt, lab_atac])

plt.figure(figsize=(5, 4))
sns.violinplot(x=labs_all, y=norms_all, inner="box")
plt.ylabel("‖z‖ (L2 norm)")
plt.xlabel("Modality")
plt.title("Latent L2-norm distribution by modality")
#plt.tight_layout()
plt.savefig(os.path.join(FIGDIR, "latent_norms_by_modality.png"))
plt.show()
plt.close()

# Latent correlation heatmap of RNA latent dims
corr_latent = np.corrcoef(z_rna, rowvar=False)
plt.figure(figsize=(6, 5))
sns.heatmap(corr_latent, cmap="vlag", center=0)
plt.title("RNA latent dimension correlation (test set)")
#plt.tight_layout()
plt.savefig(os.path.join(FIGDIR, "latent_corr_heatmap_rna.png"))
plt.show()
plt.close()

# Silhouette score on modality (lower = better mixing)
if len(np.unique(modality_labels)) > 1:
    sil_mod = silhouette_score(Z_joint, modality_labels)
else:
    sil_mod = np.nan
print(f"Silhouette (modality) on UniVI latent: {sil_mod:.3f}")


# %% [code]
# -----------------------------------------
# 6. Local modality entropy (kNN) + UMAP
# -----------------------------------------
print("\nComputing local modality entropy...")

n_neighbors_local = 30
nn_local = NearestNeighbors(n_neighbors=n_neighbors_local, metric="euclidean")
nn_local.fit(Z_joint)
_, idx_local = nn_local.kneighbors(Z_joint)

mods = modality_labels
local_modality_entropy = []

for i in range(Z_joint.shape[0]):
    neigh = idx_local[i, 1:]  # drop self
    neigh_mod = mods[neigh]

    # empirical distribution over modalities
    ent = 0.0
    for m in modalities:
        p = (neigh_mod == m).mean()
        if p > 0:
            ent -= p * np.log2(p)
    local_modality_entropy.append(ent)

local_modality_entropy = np.asarray(local_modality_entropy)
combined.obs["local_modality_entropy"] = local_modality_entropy

print(f"Mean local modality entropy (k={n_neighbors_local}): {local_modality_entropy.mean():.3f}")

plt.figure(figsize=(5, 4))
plt.hist(local_modality_entropy, bins=30)
plt.xlabel("Local modality entropy (bits)")
plt.ylabel("Cells")
plt.title("kNN modality entropy")
#plt.tight_layout()
plt.savefig(os.path.join(FIGDIR, "local_modality_entropy_hist.png"))
plt.show()
plt.close()

# UMAP colored by local modality entropy
sc.pl.umap(
    combined,
    color="local_modality_entropy",
    size=3,
    alpha=0.8,
    show=False,
)
plt.savefig(os.path.join(FIGDIR, "umap_local_modality_entropy.png"), bbox_inches="tight")
plt.show()
plt.close()


# %% [code]
# -----------------------------------------
# 7. Pairwise matching metrics (top-k) for all modality pairs
# -----------------------------------------
print("\nPairwise matching metrics (top-1 / top-5 / top-10 / top-25 / top-50 / top-75 / top-100)...")
'''
def topk_matching(z_src, z_tgt, pair_name: str, k_match: int = 100):
    nn = NearestNeighbors(n_neighbors=k_match, metric="euclidean")
    nn.fit(z_tgt)
    _, idx_knn = nn.kneighbors(z_src)

    true_idx = np.arange(z_src.shape[0])
    top1_hits  = (idx_knn[:, 0] == true_idx)
    top5_hits  = (idx_knn[:, :5] == true_idx[:, None]).any(axis=1)
    top10_hits = (idx_knn[:, :10] == true_idx[:, None]).any(axis=1)
    top25_hits = (idx_knn[:, :25] == true_idx[:, None]).any(axis=1)
    top50_hits = (idx_knn[:, :50] == true_idx[:, None]).any(axis=1)
    top75_hits = (idx_knn[:, :75] == true_idx[:, None]).any(axis=1)
    top100_hits = (idx_knn[:, :100] == true_idx[:, None]).any(axis=1)

    print(f"  {pair_name}:")
    print(f"    Top-1 accuracy:  {top1_hits.mean():.3f}")
    print(f"    Top-5 accuracy:  {top5_hits.mean():.3f}")
    print(f"    Top-10 accuracy: {top10_hits.mean():.3f}")
    print(f"    Top-25 accuracy: {top25_hits.mean():.3f}")    
    print(f"    Top-50 accuracy: {top50_hits.mean():.3f}")
    print(f"    Top-75 accuracy: {top75_hits.mean():.3f}")    
    print(f"    Top-100 accuracy: {top100_hits.mean():.3f}")

    plt.figure(figsize=(16, 14))
    plt.bar(
        ["Top-1", "Top-5", "Top-10", "Top-25", "Top-50", "Top-75", "Top-100"],
        [top1_hits.mean(), top5_hits.mean(), top10_hits.mean(), top25_hits.mean(), top50_hits.mean(), 
         top75_hits.mean(), top100_hits.mean()],
    )
    plt.ylabel("Fraction of correctly matched pairs")
    plt.title(f"Cross-modal matching accuracy ({pair_name})")
    #plt.tight_layout()
    fname = f"matching_{pair_name.replace(' ', '_').replace('→','to')}.png"
    plt.savefig(os.path.join(FIGDIR, fname))
    plt.show()
    plt.close()
'''

# Added error bars
def topk_matching(z_src, z_tgt, pair_name: str, k_match: int = 100):
    nn = NearestNeighbors(n_neighbors=k_match, metric="euclidean")
    nn.fit(z_tgt)
    _, idx_knn = nn.kneighbors(z_src)

    true_idx = np.arange(z_src.shape[0])
    n_cells  = z_src.shape[0]

    top1_hits   = (idx_knn[:, 0] == true_idx)
    top5_hits   = (idx_knn[:, :5]   == true_idx[:, None]).any(axis=1)
    top10_hits  = (idx_knn[:, :10]  == true_idx[:, None]).any(axis=1)
    top25_hits  = (idx_knn[:, :25]  == true_idx[:, None]).any(axis=1)
    top50_hits  = (idx_knn[:, :50]  == true_idx[:, None]).any(axis=1)
    top75_hits  = (idx_knn[:, :75]  == true_idx[:, None]).any(axis=1)
    top100_hits = (idx_knn[:, :100] == true_idx[:, None]).any(axis=1)

    # Means
    means = np.array([
        top1_hits.mean(),
        top5_hits.mean(),
        top10_hits.mean(),
        top25_hits.mean(),
        top50_hits.mean(),
        top75_hits.mean(),
        top100_hits.mean(),
    ])

    # Binomial standard error: sqrt(p * (1 - p) / n)
    ses = np.sqrt(means * (1.0 - means) / n_cells)

    print(f"  {pair_name}:")
    labels = ["Top-1", "Top-5", "Top-10", "Top-25", "Top-50", "Top-75", "Top-100"]
    for lab, m, se in zip(labels, means, ses):
        print(f"    {lab} accuracy: {m:.3f} ± {se:.3f} (SEM)")

    plt.figure(figsize=(8, 6))
    plt.bar(labels, means, yerr=ses, capsize=5)
    plt.ylabel("Fraction of correctly matched pairs")
    plt.title(f"Cross-modal matching accuracy ({pair_name})")
    fname = f"matching_{pair_name.replace(' ', '_').replace('→','to')}.png"
    plt.savefig(os.path.join(FIGDIR, fname))
    plt.show()
    plt.close()

topk_matching(z_rna,  z_adt,  "RNA→ADT")
topk_matching(z_adt,  z_rna,  "ADT→RNA")
topk_matching(z_rna,  z_atac, "RNA→ATAC")
topk_matching(z_atac, z_rna,  "ATAC→RNA")
topk_matching(z_adt,  z_atac, "ADT→ATAC")
topk_matching(z_atac, z_adt,  "ATAC→ADT")


# %% [code]
# -----------------------------------------
# 8. Cross-modal reconstruction metrics
# -----------------------------------------
def _to_dense(X):
    return X.toarray() if sp.issparse(X) else np.asarray(X)

def cross_modal_metrics(
    model,
    src_adata,
    tgt_adata,
    src_mod: str,
    tgt_mod: str,
    name_prefix: str,
    device: str,
):
    Xhat_tgt = univi_eval.cross_modal_predict(
        model,
        adata_src=src_adata,
        src_mod=src_mod,
        tgt_mod=tgt_mod,
        device=device,
        batch_size=512,
    )

    X_tgt = _to_dense(tgt_adata.X)

    mse_feat  = univi_eval.mse_per_feature(X_tgt, Xhat_tgt)
    corr_feat = univi_eval.pearson_corr_per_feature(X_tgt, Xhat_tgt)

    print(f"\nCross-modal reconstruction: {src_mod} → {tgt_mod}")
    print(f"  Mean feature MSE: {mse_feat.mean():.4f}")
    print(f"  Mean feature Pearson r: {corr_feat.mean():.3f}")

    # Histogram of per-feature correlation
    plt.figure(figsize=(5, 4))
    sns.histplot(corr_feat, bins=40, kde=False)
    plt.xlabel("Per-feature Pearson r")
    plt.ylabel("Count")
    plt.title(f"{src_mod} → {tgt_mod}: feature-wise correlation")
    plt.tight_layout()
    plt.savefig(os.path.join(FIGDIR, f"{name_prefix}_corr_hist.png"))
    plt.show()
    plt.close()

    # Histogram of per-feature MSE
    plt.figure(figsize=(5, 4))
    sns.histplot(mse_feat, bins=40, kde=False)
    plt.xlabel("Per-feature MSE")
    plt.ylabel("Count")
    plt.title(f"{src_mod} → {tgt_mod}: feature-wise MSE")
    plt.tight_layout()
    plt.savefig(os.path.join(FIGDIR, f"{name_prefix}_mse_hist.png"))
    plt.show()
    plt.close()

    return mse_feat, corr_feat

# Evaluate key directions on TEST set
_ = cross_modal_metrics(model, rna_test_adata, adt_test_adata,
                        src_mod="rna", tgt_mod="adt",
                        name_prefix="RNA_to_ADT_test", device=device)

_ = cross_modal_metrics(model, rna_test_adata, atac_test_adata,
                        src_mod="rna", tgt_mod="atac",
                        name_prefix="RNA_to_ATAC_test", device=device)

_ = cross_modal_metrics(model, adt_test_adata, rna_test_adata,
                        src_mod="adt", tgt_mod="rna",
                        name_prefix="ADT_to_RNA_test", device=device)

_ = cross_modal_metrics(model, atac_test_adata, rna_test_adata,
                        src_mod="atac", tgt_mod="rna",
                        name_prefix="ATAC_to_RNA_test", device=device)


# %% [code]
# ============================
# 9. Compute TEA-seq latent metrics (no return_details)
# ============================

print("Encoding TEA-seq test data into UniVI latent...")

# 1) Encode each modality into UniVI latent
Z_rna  = trainer.encode_modality(rna_test_adata,  modality="rna",  batch_size=1024)
Z_adt  = trainer.encode_modality(adt_test_adata,  modality="adt",  batch_size=1024)
Z_atac = trainer.encode_modality(atac_test_adata, modality="atac", batch_size=1024)

print("Latent shapes:", Z_rna.shape, Z_adt.shape, Z_atac.shape)

# 2) Pairwise FOSCTTM
fos_rna_adt  = univi_eval.compute_foscttm(Z_rna,  Z_adt)
fos_rna_atac = univi_eval.compute_foscttm(Z_rna,  Z_atac)
fos_adt_atac = univi_eval.compute_foscttm(Z_adt,  Z_atac)

print(f"FOSCTTM (RNA vs ADT):  {fos_rna_adt:.4f}")
print(f"FOSCTTM (RNA vs ATAC): {fos_rna_atac:.4f}")
print(f"FOSCTTM (ADT vs ATAC): {fos_adt_atac:.4f}")

# 3) Build joint latent + modality labels
Z_joint = np.vstack([Z_rna, Z_adt, Z_atac])
modality_labels = np.array(
    ["rna"]  * Z_rna.shape[0] +
    ["adt"]  * Z_adt.shape[0] +
    ["atac"] * Z_atac.shape[0]
)

# 4) Global modality mixing score from your current API
k = 30  # choose whatever k you want to use consistently
try:
    # if your version supports k as an argument
    mixing_score = univi_eval.compute_modality_mixing(Z_joint, modality_labels, k)
except TypeError:
    # older version: no k arg, just use default
    mixing_score = univi_eval.compute_modality_mixing(Z_joint, modality_labels)

print(f"Modality mixing score (k~{k}): {mixing_score:.4f}")

# 5) Manual kNN neighbors to get same_mod_frac, entropy, neighbor_mods
print("Computing neighbor-level modality stats...")

nn = NearestNeighbors(n_neighbors=k + 1, metric="euclidean")
nn.fit(Z_joint)
dists_all, idx_all = nn.kneighbors(Z_joint)

# drop self
dists_neighbors = dists_all[:, 1:]      # (n_cells, k)
idx_neighbors   = idx_all[:, 1:]        # (n_cells, k)

neighbor_mods = modality_labels[idx_neighbors]  # (n_cells, k)

# fraction of neighbors *with the same modality* (global)
same_mod_mask = (neighbor_mods == modality_labels[:, None])
same_mod_frac = float(same_mod_mask.mean())

# per-cell local modality entropy
local_modality_entropy = np.empty(Z_joint.shape[0], dtype=float)
n_cells = Z_joint.shape[0]
unique_mods = np.unique(modality_labels)

for i in range(n_cells):
    neigh = neighbor_mods[i]
    counts = np.array([(neigh == m).sum() for m in unique_mods], dtype=float)
    probs = counts / counts.sum()
    # avoid log(0)
    probs = probs[probs > 0]
    if probs.size == 0:
        local_modality_entropy[i] = 0.0
    else:
        # natural-log entropy; relative differences matter more than units
        local_modality_entropy[i] = -np.sum(probs * np.log(probs))

print(f"Same-modality neighbor fraction (k={k}): {same_mod_frac:.4f}")
print(
    f"Local modality entropy (k={k}): "
    f"mean={local_modality_entropy.mean():.4f}, "
    f"median={np.median(local_modality_entropy):.4f}"
)

# 6) Silhouette by modality
sil_mod = silhouette_score(Z_joint, modality_labels, metric="euclidean")
print(f"Silhouette score (modality labels): {sil_mod:.4f}")


# %% [code]
# ============================
# 10. Summarize TEA-seq metrics & save as JSON
# ============================
import json

teaseq_metrics = {
    # Pairwise FOSCTTM
    "foscttm_rna_adt": float(fos_rna_adt),
    "foscttm_rna_atac": float(fos_rna_atac),
    "foscttm_adt_atac": float(fos_adt_atac),

    # Global mixing
    "modality_mixing_k20": float(mixing_score),
    "same_modality_neighbor_frac_k20": float(same_mod_frac),

    # Latent geometry
    "silhouette_modality": float(sil_mod) if not np.isnan(sil_mod) else None,
    "mean_local_modality_entropy_k20": float(local_modality_entropy.mean()),
    "median_local_modality_entropy_k20": float(np.median(local_modality_entropy)),

    # Dataset sizes
    "n_cells_test": int(rna_test_adata.n_obs),
    "n_genes_rna": int(rna_test_adata.n_vars),
    "n_features_adt": int(adt_test_adata.n_vars),
    "n_features_atac": int(atac_test_adata.n_vars),
}

'''
metrics_path = os.path.join(FIGDIR, "teaseq_univi_metrics.json")
with open(metrics_path, "w") as f:
    json.dump(teaseq_metrics, f, indent=2)

print(f"\n[TEA-seq] Saved benchmark metrics to: {metrics_path}")
'''


# %% [code]
# ============================
# 11. kNN distance diagnostics: same vs cross-modality neighbors
# ============================
print("\nComputing kNN distance distributions (same vs cross-modality)...")

from sklearn.neighbors import NearestNeighbors

k_dist = 30
nn_dist = NearestNeighbors(n_neighbors=k_dist + 1, metric="euclidean")
nn_dist.fit(Z_joint)
dists_all, idx_all = nn_dist.kneighbors(Z_joint)

# drop self
dists_neighbors = dists_all[:, 1:]              # (n_cells, k_dist)
idx_neighbors_dist = idx_all[:, 1:]             # already had neighbor_mods from earlier
flat_dists = dists_neighbors.reshape(-1)

# Center & neighbor modalities (flattened to per-edge view)
center_mods_flat = np.repeat(modality_labels, k_dist)
neighbor_mods_flat = neighbor_mods.reshape(-1)

same_mod_edge = neighbor_mods_flat == center_mods_flat

plt.figure(figsize=(6, 4))
sns.kdeplot(flat_dists[same_mod_edge], label="same modality", fill=True, alpha=0.6)
sns.kdeplot(flat_dists[~same_mod_edge], label="different modality", fill=True, alpha=0.6)
plt.xlabel("Euclidean distance in UniVI latent")
plt.ylabel("Density")
plt.title(f"kNN distance distribution (k={k_dist})")
plt.legend()
plt.tight_layout()
plt.savefig(os.path.join(FIGDIR, "knn_distance_same_vs_cross_modality.png"), dpi=200)
plt.show()
plt.close()


# %% [code]
# ============================
# 12. Cross-modal neighbor fraction per cell (and UMAP)
# ============================
print("\nComputing per-cell cross-modal neighbor fraction...")

# fraction of neighbors NOT sharing the cell's modality
cross_mod_neighbor_frac = 1.0 - (neighbor_mods == modality_labels[:, None]).mean(axis=1)
combined.obs["cross_mod_neighbor_frac"] = cross_mod_neighbor_frac

plt.figure(figsize=(5, 4))
sns.histplot(cross_mod_neighbor_frac, bins=30)
plt.xlabel("Fraction of neighbors with different modality")
plt.ylabel("Cells")
plt.title(f"Cross-modal neighbor fraction (k={k})")
plt.tight_layout()
plt.savefig(os.path.join(FIGDIR, "cross_mod_neighbor_fraction_hist.png"), dpi=200)
plt.show()
plt.close()

# UMAP colored by cross-modal neighbor fraction
sc.pl.umap(
    combined,
    color="cross_mod_neighbor_frac",
    size=3,
    alpha=0.8,
    cmap="viridis",
    show=False,
)
plt.title("UMAP – cross-modal neighbor fraction")
plt.savefig(os.path.join(FIGDIR, "umap_cross_mod_neighbor_fraction.png"),
            bbox_inches="tight", dpi=200)
plt.show()
plt.close()


# %% [code]
# ============================
# 13. Per-cluster modality composition (Leiden clusters)
# ============================
print("\nComputing modality composition per Leiden cluster...")

if "univi_leiden" in combined.obs.columns:
    comp_ct = pd.crosstab(combined.obs["univi_leiden"], combined.obs["univi_source"])
    comp_prop = comp_ct.div(comp_ct.sum(axis=1), axis=0)  # row-normalize

    plt.figure(figsize=(7, 5))
    comp_prop.sort_index().plot(
        kind="bar",
        stacked=True,
        width=0.9,
        colormap="tab10",
    )
    plt.xlabel("UniVI Leiden cluster")
    plt.ylabel("Fraction of cells")
    plt.title("Modality composition per UniVI Leiden cluster")
    plt.legend(title="Modality", bbox_to_anchor=(1.05, 1), loc="upper left")
    plt.tight_layout()
    plt.savefig(os.path.join(FIGDIR, "cluster_modality_composition_stacked_bar.png"),
                dpi=200)
    plt.show()
    plt.close()
else:
    print("  WARNING: 'univi_leiden' not found in combined.obs; skipping cluster composition plot.")
    

# %% [code]
# ============================
# 14. Feature-level cross-modal scatter plots (RNA → ADT / ATAC)
# ============================
print("\nFeature-level scatter plots for cross-modal prediction (RNA→ADT / RNA→ATAC)...")

# Example: RNA → ADT for a small subset of cells & selected markers
# (adjust marker names to what you actually have in adt_test_adata.var_names)

n_scatter_cells = min(5000, adt_test_adata.n_obs)  # subsample if huge
cell_idx = np.random.default_rng(42).choice(adt_test_adata.n_obs, n_scatter_cells, replace=False)

# recompute predictions just for this subset to avoid reusing big arrays
Xhat_adt_sub = univi_eval.cross_modal_predict(
    model,
    adata_src=rna_test_adata[cell_idx],
    src_mod="rna",
    tgt_mod="adt",
    device=device,
    batch_size=512,
)
X_adt_sub = _to_dense(adt_test_adata[cell_idx].X)

adt_markers_to_plot = [
    # put your favorite ADT markers here
    # e.g. "CD3", "CD4", "CD8A", "CD56", ...
]

for marker in adt_markers_to_plot:
    if marker not in adt_test_adata.var_names:
        print(f"  [RNA→ADT] Marker '{marker}' not found in adt_test_adata.var_names; skipping.")
        continue

    j = np.where(adt_test_adata.var_names == marker)[0][0]
    y_true = X_adt_sub[:, j]
    y_pred = Xhat_adt_sub[:, j]

    plt.figure(figsize=(4.5, 4))
    plt.hexbin(y_true, y_pred, gridsize=50, mincnt=1)
    plt.xlabel(f"True ADT ({marker})")
    plt.ylabel(f"Predicted ADT ({marker})")
    plt.title(f"RNA→ADT prediction for {marker}")
    plt.tight_layout()
    fname = f"scatter_RNA_to_ADT_{marker}.png".replace(" ", "_")
    plt.savefig(os.path.join(FIGDIR, fname), dpi=200)
    plt.show()
    plt.close()


# Example: RNA → ATAC for a few peaks / gene-body features (if named)
# (This is more exploratory since ATAC features are often peaks; choose a few named ones if available.)

n_scatter_cells_atac = min(5000, atac_test_adata.n_obs)
cell_idx_atac = np.random.default_rng(123).choice(atac_test_adata.n_obs, n_scatter_cells_atac, replace=False)

Xhat_atac_sub = univi_eval.cross_modal_predict(
    model,
    adata_src=rna_test_adata[cell_idx_atac],
    src_mod="rna",
    tgt_mod="atac",
    device=device,
    batch_size=512,
)
X_atac_sub = _to_dense(atac_test_adata[cell_idx_atac].X)

# If you have named ATAC features (e.g. gene body aggregates), you can list them here.
atac_features_to_plot = [
    # e.g. "TNFRSF4_body", "IFNG_enh", ...
]

for feat in atac_features_to_plot:
    if feat not in atac_test_adata.var_names:
        print(f"  [RNA→ATAC] Feature '{feat}' not found in atac_test_adata.var_names; skipping.")
        continue

    j = np.where(atac_test_adata.var_names == feat)[0][0]
    y_true = X_atac_sub[:, j]
    y_pred = Xhat_atac_sub[:, j]

    plt.figure(figsize=(4.5, 4))
    plt.hexbin(y_true, y_pred, gridsize=50, mincnt=1)
    plt.xlabel(f"True ATAC ({feat})")
    plt.ylabel(f"Predicted ATAC ({feat})")
    plt.title(f"RNA→ATAC prediction for {feat}")
    plt.tight_layout()
    fname = f"scatter_RNA_to_ATAC_{feat}.png".replace(" ", "_")
    plt.savefig(os.path.join(FIGDIR, fname), dpi=200)
    plt.show()
    plt.close()


# %% [code]
# -----------------------------------------
# Helper: unimodal Leiden clustering (safer)
# -----------------------------------------
def compute_unimodal_leiden(
    adata,
    mod: str,
    key_added: str = None,
    resolution: float = 1.0,
    n_neighbors: int = 15,
    n_pcs: int = 30,
):
    """
    Compute unimodal Leiden clusters for a single AnnData.

    Parameters
    ----------
    adata : AnnData
        Modality-specific AnnData (e.g. RNA-only, ADT-only, ATAC-only).
    mod : {"rna", "adt", "atac"}
        Name of the modality (used to pick rep + default key).
    key_added : str, optional
        Name of the .obs column for clusters (default: f"{mod}_leiden").
    resolution : float
        Leiden resolution parameter.
    n_neighbors : int
        Number of neighbors for kNN graph.
    n_pcs : int
        Number of PCs to use when computing PCA (if needed).

    Returns
    -------
    adata : AnnData
        Same object with a new .obs[key_added] column.
    """
    if adata is None or adata.n_obs == 0:
        print(f"[{mod}] No cells; skipping Leiden.")
        return adata

    if key_added is None:
        key_added = f"{mod}_leiden"

    # Decide which representation to use
    use_rep = None

    # ---------- choose rep ----------
    if mod in ("rna", "adt"):
        if "X_pca" in adata.obsm_keys():
            use_rep = "X_pca"
            print(f"[{mod}] Using existing X_pca for neighbors/Leiden.")
        else:
            print(f"[{mod}] Computing PCA on existing .X for neighbors/Leiden (no extra scaling)...")
            tmp = adata.copy()

            # Clean NaNs / infs if present
            X = tmp.X.A if sp.issparse(tmp.X) else np.asarray(tmp.X)
            if not np.isfinite(X).all():
                n_bad = np.sum(~np.isfinite(X))
                print(f"[{mod}] Warning: found {n_bad} non-finite entries in .X; replacing with 0.")
                X = np.nan_to_num(X, nan=0.0, posinf=0.0, neginf=0.0)
                if sp.issparse(tmp.X):
                    tmp.X = sp.csr_matrix(X)
                else:
                    tmp.X = X

            # PCA directly on (already processed) X
            sc.tl.pca(tmp, n_comps=n_pcs)
            adata.obsm["X_pca"] = tmp.obsm["X_pca"]
            use_rep = "X_pca"

    elif mod == "atac":
        if "X_lsi" in adata.obsm_keys():
            use_rep = "X_lsi"
            print(f"[{mod}] Using existing X_lsi for neighbors/Leiden.")
        else:
            print(f"[{mod}] X_lsi not found; computing PCA on .X for neighbors/Leiden (no extra scaling)...")
            tmp = adata.copy()

            X = tmp.X.A if sp.issparse(tmp.X) else np.asarray(tmp.X)
            if not np.isfinite(X).all():
                n_bad = np.sum(~np.isfinite(X))
                print(f"[{mod}] Warning: found {n_bad} non-finite entries in .X; replacing with 0.")
                X = np.nan_to_num(X, nan=0.0, posinf=0.0, neginf=0.0)
                if sp.issparse(tmp.X):
                    tmp.X = sp.csr_matrix(X)
                else:
                    tmp.X = X

            sc.tl.pca(tmp, n_comps=n_pcs)
            adata.obsm["X_pca"] = tmp.obsm["X_pca"]
            use_rep = "X_pca"

    else:
        # Fallback for any other modality
        if "X_pca" in adata.obsm_keys():
            use_rep = "X_pca"
            print(f"[{mod}] Using existing X_pca for neighbors/Leiden.")
        else:
            print(f"[{mod}] Unknown modality; computing PCA on .X for neighbors/Leiden (no extra scaling)...")
            tmp = adata.copy()

            X = tmp.X.A if sp.issparse(tmp.X) else np.asarray(tmp.X)
            if not np.isfinite(X).all():
                n_bad = np.sum(~np.isfinite(X))
                print(f"[{mod}] Warning: found {n_bad} non-finite entries in .X; replacing with 0.")
                X = np.nan_to_num(X, nan=0.0, posinf=0.0, neginf=0.0)
                if sp.issparse(tmp.X):
                    tmp.X = sp.csr_matrix(X)
                else:
                    tmp.X = X

            sc.tl.pca(tmp, n_comps=n_pcs)
            adata.obsm["X_pca"] = tmp.obsm["X_pca"]
            use_rep = "X_pca"

    # ---------- neighbors + Leiden ----------
    print(f"[{mod}] Computing neighbors (n_neighbors={n_neighbors}, use_rep={use_rep})...")
    sc.pp.neighbors(adata, n_neighbors=n_neighbors, use_rep=use_rep)

    print(f"[{mod}] Running Leiden (resolution={resolution}) → obs['{key_added}']...")
    sc.tl.leiden(adata, key_added=key_added, resolution=resolution)

    n_clusters = adata.obs[key_added].nunique()
    print(f"[{mod}] Leiden done: {n_clusters} clusters in obs['{key_added}'].")

    return adata

# -----------------------------------------
# Run unimodal Leiden on test sets
# -----------------------------------------

# You can tune resolution per modality if you want:
resolutions = {
    #"rna":  3.0,
    #"adt":  3.8,
    #"atac": 2.45,
    "rna":  2.25,
    "adt":  1.75,
    "atac": 0.85,
}

specs = [
    ("rna",  locals().get("rna_test_adata",  None)),
    ("adt",  locals().get("adt_test_adata",  None)),
    ("atac", locals().get("atac_test_adata", None)),
]

for mod, adata in specs:
    if adata is None:
        print(f"[{mod}] No AnnData object found (e.g. rna_test is None); skipping.")
        continue

    compute_unimodal_leiden(
        adata,
        mod=mod,
        key_added=f"{mod}_leiden",
        resolution=resolutions.get(mod, 1.0),
        n_neighbors=30,
        n_pcs=40,
    )


# %% [code]
import os
import numpy as np
import pandas as pd
import scipy.sparse as sp
import seaborn as sns
import matplotlib.pyplot as plt


def _to_dense(X):
    return X.toarray() if sp.issparse(X) else np.asarray(X)


def sanitize_for_clustermap(df: pd.DataFrame) -> pd.DataFrame:
    """Drop inf/nan columns + zero-variance columns (prevents scipy linkage errors)."""
    df = df.replace([np.inf, -np.inf], np.nan)
    df = df.dropna(axis=1, how="any")
    if df.shape[1] > 0:
        v = df.var(axis=0, ddof=0)
        df = df.loc[:, v > 0]
    return df


def cluster_means_df(
    adata,
    *,
    layer: str | None,
    cluster_key: str,
    features: list[str],
) -> pd.DataFrame:
    if cluster_key not in adata.obs:
        raise KeyError(f"{cluster_key} not in adata.obs")
    if not features:
        raise ValueError("features must be a non-empty list (won't use all features).")

    X = adata.X if layer in (None, "X") else adata.layers[layer]
    X = _to_dense(X)

    present = [f for f in features if f in adata.var_names]
    if len(present) == 0:
        raise ValueError("None of the requested features are in adata.var_names")

    cols = adata.var_names.get_indexer(present)
    clusters = adata.obs[cluster_key].astype("category")
    cats = clusters.cat.categories

    rows = []
    for cl in cats:
        mask = (clusters == cl).values
        rows.append(X[mask][:, cols].mean(axis=0))

    df = pd.DataFrame(
        np.vstack(rows),
        index=cats.astype(str),
        columns=pd.Index(present, dtype=str),
    )
    df.index.name = cluster_key
    df.columns.name = "Features"
    return df


def plot_clustermap(
    df: pd.DataFrame,
    *,
    title: str,
    outpath: str | None = None,
    cmap="viridis",
    vmin=None,
    vmax=None,
    center=None,
    method="average",
    metric="correlation",
    figsize=None,
    show=True,
    row_cluster=True,
    col_cluster=False,

    # NEW: allow freezing cluster structure from a "template" plot
    row_linkage=None,
    col_linkage=None,

    xtick_rotation=90,
    ytick_rotation=0,
    tick_fontsize=8,
    title_fontsize=12,
    dendrogram_ratio=(0.12, 0.12),

    # ---- colorbar controls ----
    cbar_pos=(0.05, 0.80, 0.2, 0.15),
    cbar_label="z-scored mean",
    cbar_labelpad=6,
    cbar_ticklength=2,
    #cbar_aspect=2.0,

    margins=dict(left=0.12, right=0.98, bottom=0.22, top=0.92),
    dpi_save=150,
):
    df = sanitize_for_clustermap(df)
    if df.shape[0] == 0 or df.shape[1] == 0:
        raise ValueError("Nothing left to plot after sanitization (0 rows/cols).")

    if figsize is None:
        figsize = (0.28 * df.shape[1] + 4.0, 0.28 * df.shape[0] + 3.0)

    g = sns.clustermap(
        df,
        cmap=cmap,
        vmin=vmin,
        vmax=vmax,
        center=center,
        method=method,
        metric=metric,
        figsize=figsize,
        linewidths=0,
        linecolor=None,
        row_cluster=row_cluster,
        col_cluster=col_cluster,

        # NEW: reuse dendrograms if provided
        row_linkage=row_linkage,
        col_linkage=col_linkage,

        dendrogram_ratio=dendrogram_ratio,
        cbar_pos=cbar_pos,
        cbar_kws={"label": cbar_label},
    )

    g.fig.suptitle(title, y=0.98, fontsize=title_fontsize)
    g.fig.subplots_adjust(**margins)

    if getattr(g, "cax", None) is not None:
        g.cax.set_position(cbar_pos)

        left, bottom, width, height = cbar_pos
        horizontal = width > height
        if horizontal:
            g.cax.xaxis.set_label_position("top")
            g.cax.xaxis.tick_top()
            g.cax.set_ylabel("")
            g.cax.set_xlabel(cbar_label, labelpad=cbar_labelpad, fontsize=tick_fontsize)

        try:
            g.cax.set_aspect(cbar_aspect)
        except Exception:
            pass

        g.cax.tick_params(labelsize=tick_fontsize, length=cbar_ticklength)

    g.ax_heatmap.set_xlabel(df.columns.name or "Features")
    g.ax_heatmap.set_ylabel(df.index.name or "Clusters")
    plt.setp(g.ax_heatmap.get_xticklabels(), rotation=xtick_rotation, ha="right", fontsize=tick_fontsize)
    plt.setp(g.ax_heatmap.get_yticklabels(), rotation=ytick_rotation, fontsize=tick_fontsize)

    if outpath is not None:
        os.makedirs(os.path.dirname(outpath), exist_ok=True)
        g.savefig(outpath, dpi=dpi_save, bbox_inches="tight", pad_inches=0.15)

    if show:
        plt.show()
    plt.close(g.fig)
    return g


def _align_to_template(df: pd.DataFrame, template: pd.DataFrame) -> pd.DataFrame:
    """
    Force df to have exactly the same rows/cols as template (drops extras, keeps order).
    This is important when reusing linkages.
    """
    df = df.reindex(index=template.index, columns=template.columns)
    # if anything became NaN (e.g., missing feature/cluster), drop those columns/rows consistently
    df = df.dropna(axis=0, how="any")
    df = df.dropna(axis=1, how="any")
    return df


# %% [code]
RNA_MARKERS = {
    "B":        ["MS4A1","CD79A","CD74","HLA-DRA","CD22","BANK1","BLK","IGHM","IGKC","JCHAIN"],
    "T_core":   ["TRAC","TRBC1","CD3D","CD3E","LCK","ITK","TXK"],
    "CD4_T":    ["IL7R","CCR7","LTB","TCF7","LEF1","MAL"],
    "CD8_Cyto": ["NKG7","GNLY","PRF1","GZMB","GZMH","CTSW","CCL5","XCL2","RUNX3"],
    "NK":       ["NKG7","GNLY","PRF1","FCGR3A","TYROBP","KLRD1"],
    "Mono":     ["LYZ","S100A8","S100A9","FCN1","LST1","CTSS","LGALS3","SLC2A3","IL1B","VCAN"],
    "DC":       ["FCER1A","CST3","CLEC10A","ITGAX","CD1C","WDFY4"],
    "pDC":      ["GZMB","TCF4","IRF7","IL3RA"],
    "Megak":    ["PPBP","PF4","GP1BB"],
    "IFN":      ["ISG15","IFIT1","IFIT2","IFIT3","MX1","OAS1","STAT1","OASL"],
}


# %% [code]
ADT_MARKERS = {
    # T / NK core
    "T_core":      ["CD3", "TCR-a/b", "TCR-g/d"],
    "CD4_T":       ["CD4", "CD197", "CD127"],              # CCR7, IL7R-ish
    "CD8_T":       ["CD8a", "CD45RA", "CD45RO"],
    "MAIT/iNKT":   ["TCR-Va7.2", "TCR-Va24-Ja18"],         # MAIT / iNKT proxies
    "Activation":  ["CD25", "CD95", "CD39", "CD71"],
    "Exhaustion":  ["CD279"],                              # PD-1
    "NK":          ["CD56", "CD16", "KLRG1", "CD319"],      # CD319 = SLAMF7

    # B lineage
    "B_core":      ["CD19", "CD21", "CD24", "IgD", "IgM"],
    "Plasmablast": ["CD38", "CD27", "CD269"],              # CD269 = BCMA

    # Myeloid / DC
    "Monocyte":    ["CD14", "CD11b", "CD172a", "CD192"],   # CD192=CCR2
    "DC":          ["CD11c", "HLA-DR", "CD86", "CD80", "CD40"],
    "pDC_hint":    ["CD123", "CD304"],                     # IL3RA, NRp1/BDCA4
    "cDC1_hint":   ["CD141"],                              # BDCA3
    "FcERI":       ["FceRI"],

    # Neutrophil-ish / granulocyte
    "Granulocyte": ["CD66b"],

    # Isotypes / controls (don’t use for annotation, but useful to QC)
    "Isotypes":    ["IgG1-K-Isotype-Control", "total"],
}


# %% [code]
def flatten_marker_dict(d):
    out = []
    for _, genes in d.items():
        out.extend(genes)
    # preserve order, unique
    seen = set()
    uniq = []
    for g in out:
        if g not in seen:
            uniq.append(g); seen.add(g)
    return uniq


# %% [code]
# ---- panels ----
RNA_PANEL = flatten_marker_dict(RNA_MARKERS)

ADT_PANEL_ANNOT = sorted({x for k,v in ADT_MARKERS.items() for x in v if k not in ("Isotypes",)})
# If you want to include isotypes too, use: sorted({x for v in ADT_MARKERS.values() for x in v})

ATAC_PANEL = [f"LSI_{i}" for i in range(1, 51)]  # LSI_1..LSI_50


# %% [code]
print(rna_test_adata)


# %% [code]
print(Z_rna)
print(Z_adt)
print(Z_atac)


# %% [code]
X_rna = torch.as_tensor(np.asarray(rna_test_adata.X, dtype=np.float32), dtype=torch.float32, device=device)

print(X_rna)


# %% [code]
X_adt = torch.as_tensor(np.asarray(adt_test_adata.X, dtype=np.float32), dtype=torch.float32, device=device)

print(X_adt)


# %% [code]
X_atac = torch.as_tensor(np.asarray(atac_test_adata.X, dtype=np.float32), dtype=torch.float32, device=device)

print(X_atac)


# %% [code]
mu_rna_dict, logvar_rna_dict = model_loaded.encode_modalities({"rna": X_rna})
mu_adt_dict, logvar_adt_dict = model_loaded.encode_modalities({"adt": X_adt})
mu_atac_dict, logvar_atac_dict = model_loaded.encode_modalities({"atac": X_atac})
# Can encode all modalities into their own space
mu_all_separate_z_dict, logvar_all_separate_z_dict = model_loaded.encode_modalities({"rna": X_rna, "adt": X_adt, "atac": X_atac})
# Or can encode all into shared latent
mu_all_shared_z_dict, logvar_all_shared_z_dict, z = model_loaded.encode_fused({"rna": X_rna, "adt": X_adt, "atac": X_atac}, epoch=ckpt.get("best_epoch"), use_mean=True)


# %% [code]
#print(mu_rna_dict)
#print(mu_adt_dict)
#print(mu_atac_dict)
print(mu_all_separate_z_dict.keys())
print(mu_all_separate_z_dict)
print(mu_all_shared_z_dict)


# %% [code]
#rna_recons = model_loaded.decode_modalities(torch.as_tensor(mu_all_separate_z_dict['rna'], dtype=torch.float32, device=device))
#adt_recons = model_loaded.decode_modalities(torch.as_tensor(mu_all_separate_z_dict['adt'], dtype=torch.float32, device=device))
#atac_recons = model_loaded.decode_modalities(torch.as_tensor(mu_all_separate_z_dict['atac'], dtype=torch.float32, device=device))

rna_recons = model_loaded.decode_modalities(torch.as_tensor(mu_all_shared_z_dict, dtype=torch.float32, device=device))
adt_recons = model_loaded.decode_modalities(torch.as_tensor(mu_all_shared_z_dict, dtype=torch.float32, device=device))
atac_recons = model_loaded.decode_modalities(torch.as_tensor(mu_all_shared_z_dict, dtype=torch.float32, device=device))


# %% [code]
print(rna_recons)
print(adt_recons)
print(atac_recons)


# %% [code]
# same-modality “denoised” reconstructions
X_rna_hat  = rna_recons['rna'].detach().cpu().numpy() 
X_adt_hat  = adt_recons['adt'].detach().cpu().numpy()
X_atac_hat = atac_recons['atac'].detach().cpu().numpy()

# attach as layers for plotting
rna_test_adata.layers["univi_denoised"]  = X_rna_hat.astype(np.float32)
adt_test_adata.layers["univi_denoised"]  = X_adt_hat.astype(np.float32)
atac_test_adata.layers["univi_denoised"] = X_atac_hat.astype(np.float32)


# %% [code]
# ---- loop ----
FIGDIR = FIGDIR  # your existing output dir

denoise_specs = [
    dict(adata=rna_test_adata,  mod="rna",  tag="rna_test_adata",  cluster_key="rna_leiden",
         raw_layer=("scaled" if "scaled" in rna_test_adata.layers else None),
         den_layer="univi_denoised",
         #den_layer="X_univi",
         features=RNA_PANEL),

    dict(adata=adt_test_adata,  mod="adt",  tag="adt_test_adata",  cluster_key="adt_leiden",
         raw_layer=("scaled" if "scaled" in adt_test_adata.layers else None),
         den_layer="univi_denoised",
         #den_layer="X_univi",
         features=ADT_PANEL_ANNOT),

    dict(adata=atac_test_adata, mod="atac", tag="atac_test_adata", cluster_key="atac_leiden",
         raw_layer=("scaled" if "scaled" in atac_test_adata.layers else None),
         den_layer="univi_denoised",
         #den_layer="X_univi",
         features=ATAC_PANEL),
]


# %% [code]
#sc.set_figure_params(dpi=200)


# %% [code]
for spec in denoise_specs:
    adata = spec["adata"]
    mod = spec["mod"]
    tag = spec["tag"]
    cluster_key = spec["cluster_key"]
    raw_layer = spec["raw_layer"]
    den_layer = spec["den_layer"]
    features = spec["features"]

    if den_layer not in adata.layers:
        raise KeyError(f"[{mod}] missing layer '{den_layer}'")

    # --- build dfs ---
    df_raw = cluster_means_df(adata, layer=raw_layer, cluster_key=cluster_key, features=features)
    df_den = cluster_means_df(adata, layer=den_layer, cluster_key=cluster_key, features=features)

    # --- sanitize raw FIRST and use it as the template for everything ---
    df_raw_tpl = sanitize_for_clustermap(df_raw)

    # align denoised to the exact same rows/cols (and order) as raw template
    df_den_aligned = _align_to_template(df_den, df_raw_tpl)

    # also align raw to whatever survived alignment (keeps strict matching)
    df_raw_aligned = _align_to_template(df_raw_tpl, df_den_aligned)

    # delta on aligned frames
    df_delta = df_den_aligned - df_raw_aligned

    # shared scaling for raw/den; symmetric scaling for delta
    vmin = np.nanmin([df_raw_aligned.values.min(), df_den_aligned.values.min()])
    vmax = np.nanmax([df_raw_aligned.values.max(), df_den_aligned.values.max()])
    v_abs = float(np.nanmax(np.abs(df_delta.values)))

    raw_out = os.path.join(FIGDIR, f"clustermap_{tag}_{mod}_raw_by_{cluster_key}.png")
    den_out = os.path.join(FIGDIR, f"clustermap_{tag}_{mod}_denoised_by_{cluster_key}.png")
    del_out = os.path.join(FIGDIR, f"clustermap_{tag}_{mod}_delta_by_{cluster_key}.png")

    cbar_pos = (0.075, 0.80, 0.02, 0.15)
    
    # --- 1) RAW: compute clustering ONCE (this defines the dendrograms) ---
    g_raw = plot_clustermap(
        df_raw_aligned,
        cbar_pos=cbar_pos,
        title=f"{tag} ({mod}) – raw (cluster means)",
        outpath=raw_out,
        cmap="viridis",
        vmin=vmin,
        vmax=vmax,
        row_cluster=True,
        col_cluster=True,
        dpi_save=150,
        show=True,
    )

    # grab linkages (only valid if you clustered that axis)
    row_L = g_raw.dendrogram_row.linkage if getattr(g_raw, "dendrogram_row", None) is not None else None
    col_L = g_raw.dendrogram_col.linkage if getattr(g_raw, "dendrogram_col", None) is not None else None

    # --- 2) DENOISED: reuse the exact same clusters/order ---
    plot_clustermap(
        df_den_aligned.reindex(index=df_raw_aligned.index, columns=df_raw_aligned.columns),
        cbar_pos=cbar_pos,
        title=f"{tag} ({mod}) – denoised (cluster means)",
        outpath=den_out,
        cmap="viridis",
        vmin=vmin,
        vmax=vmax,
        row_cluster=True,
        col_cluster=True,
        row_linkage=row_L,
        col_linkage=col_L,
        dpi_save=150,
        show=True,
    )

    # --- 3) DELTA: reuse the exact same clusters/order ---
    plot_clustermap(
        df_delta.reindex(index=df_raw_aligned.index, columns=df_raw_aligned.columns),
        cbar_pos=cbar_pos,
        title=f"{tag} ({mod}) – Δ (den - raw)",
        outpath=del_out,
        cmap="vlag",
        center=0,
        vmin=-v_abs,
        vmax=v_abs,
        row_cluster=True,
        col_cluster=True,
        row_linkage=row_L,
        col_linkage=col_L,
        dpi_save=150,
        show=True,
    )
    

# %% [code]


# %% [code]
# Rebuild univi_source if missing
if "univi_source" not in combined.obs.columns:
    combined.obs["univi_source"] = np.concatenate([
        np.repeat("rna",  rna_test_adata.n_obs),
        np.repeat("adt",  adt_test_adata.n_obs),
        np.repeat("atac", atac_test_adata.n_obs),
    ])


# %% [code]
# -----------------------------------------
# After building `combined` and running UMAP / univi_leiden
# -----------------------------------------
# combined was made from rna_tmp, adt_tmp, atac_tmp and has:
#   combined.obs["univi_source"] in {"rna", "adt", "atac"}

# Initialize columns as NaN
for key in ["rna_leiden", "adt_leiden", "atac_leiden"]:
    combined.obs[key] = np.nan

# RNA clusters → rows in `combined` with univi_source == "rna"
if "rna_leiden" in rna_test_adata.obs:
    mask_rna = combined.obs["univi_source"] == "rna"
    n_rna_combined = mask_rna.sum()
    assert n_rna_combined == rna_test_adata.n_obs, (
        f"RNA counts mismatch: combined has {n_rna_combined}, "
        f"rna_test_adata has {rna_test_adata.n_obs}"
    )
    combined.obs.loc[mask_rna, "rna_leiden"] = (
        rna_test_adata.obs["rna_leiden"].astype(str).values
    )

# ADT clusters
if "adt_leiden" in adt_test_adata.obs:
    mask_adt = combined.obs["univi_source"] == "adt"
    n_adt_combined = mask_adt.sum()
    assert n_adt_combined == adt_test_adata.n_obs, (
        f"ADT counts mismatch: combined has {n_adt_combined}, "
        f"adt_test_adata has {adt_test_adata.n_obs}"
    )
    combined.obs.loc[mask_adt, "adt_leiden"] = (
        adt_test_adata.obs["adt_leiden"].astype(str).values
    )

# ATAC clusters
if "atac_leiden" in atac_test_adata.obs:
    mask_atac = combined.obs["univi_source"] == "atac"
    n_atac_combined = mask_atac.sum()
    assert n_atac_combined == atac_test_adata.n_obs, (
        f"ATAC counts mismatch: combined has {n_atac_combined}, "
        f"atac_test_adata has {atac_test_adata.n_obs}"
    )
    combined.obs.loc[mask_atac, "atac_leiden"] = (
        atac_test_adata.obs["atac_leiden"].astype(str).values
    )


# %% [code]
# -----------------------------------------
# 10. Overlap of unimodal clusters in UniVI latent space
# -----------------------------------------
print("\nVisualizing UniVI latent UMAP colored by unimodal clusters...")

# These should have been carried into `combined` via concatenate
for key in ["rna_leiden", "adt_leiden", "atac_leiden"]:
    if key in combined.obs.columns:
        print(f"  Plotting UMAP colored by {key}...")
        sc.pl.umap(
            combined,
            color=key,
            size=65,
            alpha=0.8,
            show=False,
        )
        plt.savefig(
            os.path.join(FIGDIR, f"umap_tri_modal_{key}.png"),
            bbox_inches="tight",
        )
        plt.show()
        plt.close()
    else:
        print(f"  [!] {key} not found in combined.obs – skipping this UMAP.")

# -----------------------------------------
# Cluster overlap tables: how do unimodal clusters align cell-by-cell?
# -----------------------------------------
import pandas as pd

def cluster_overlap_heatmap(
    adata_a,
    key_a: str,
    adata_b,
    key_b: str,
    pair_name: str,
    normalize: str = "index",
):
    """
    Make a normalized confusion matrix between two clusterings
    (rows = clusters in A, columns = clusters in B).
    """
    assert np.array_equal(adata_a.obs_names, adata_b.obs_names), (
        f"{pair_name}: obs_names do not match 1:1"
    )

    s_a = adata_a.obs[key_a].astype("category")
    s_b = adata_b.obs[key_b].astype("category")

    tab = pd.crosstab(s_a, s_b, normalize=normalize)

    plt.figure(figsize=(0.5 * tab.shape[1] + 4, 0.5 * tab.shape[0] + 4))
    sns.heatmap(
        tab,
        annot=False,
        cmap="viridis",
        cbar_kws={"label": f"Fraction (normalized by {normalize})"},
    )
    plt.xlabel(key_b)
    plt.ylabel(key_a)
    plt.title(f"Cluster overlap: {pair_name}")
    plt.tight_layout()
    fname = f"cluster_overlap_{pair_name.replace(' ', '_').replace('→','to')}.png"
    plt.savefig(os.path.join(FIGDIR, fname))
    plt.show()
    plt.close()

    return tab

# Only run if the keys exist in obs
if "rna_leiden" in rna_test_adata.obs and "adt_leiden" in adt_test_adata.obs:
    print("\nCluster overlap RNA vs ADT (unimodal Leiden)...")
    tab_rna_adt = cluster_overlap_heatmap(
        rna_test_adata, "rna_leiden",
        adt_test_adata, "adt_leiden",
        pair_name="RNA→ADT",
        normalize="index",
    )

if "rna_leiden" in rna_test_adata.obs and "atac_leiden" in atac_test_adata.obs:
    print("\nCluster overlap RNA vs ATAC (unimodal Leiden)...")
    tab_rna_atac = cluster_overlap_heatmap(
        rna_test_adata, "rna_leiden",
        atac_test_adata, "atac_leiden",
        pair_name="RNA→ATAC",
        normalize="index",
    )

if "adt_leiden" in adt_test_adata.obs and "atac_leiden" in atac_test_adata.obs:
    print("\nCluster overlap ADT vs ATAC (unimodal Leiden)...")
    tab_adt_atac = cluster_overlap_heatmap(
        adt_test_adata, "adt_leiden",
        atac_test_adata, "atac_leiden",
        pair_name="ADT→ATAC",
        normalize="index",
    )


# %% [code]
# -----------------------------------------
# 11. Marker exploration (input vs latent vs recon)
# -----------------------------------------
import os
import numpy as np
import scanpy as sc
import matplotlib.pyplot as plt
import seaborn as sns  # already imported above, but just in case


def ensure_umap_on_univi(
    adata,
    n_neighbors: int = 30,
    min_dist: float = 0.5,
):
    """
    Ensure adata has a UMAP in .obsm['X_umap'] using .obsm['X_univi'] as input.
    """
    if "X_univi" not in adata.obsm_keys():
        raise ValueError("adata.obsm['X_univi'] is missing; cannot build UMAP on UniVI latent.")
    if "X_umap" in adata.obsm_keys():
        # nothing to do
        return

    print(f"[umap] Computing neighbors/UMAP on UniVI latent (n_neighbors={n_neighbors}, min_dist={min_dist})...")
    sc.pp.neighbors(adata, use_rep="X_univi", n_neighbors=n_neighbors)
    sc.tl.umap(adata, min_dist=min_dist)


def plot_markers_on_umap(
    adata,
    marker_dict: dict,
    title_prefix: str,
    figdir: str = FIGDIR,
    n_neighbors: int = 30,
    min_dist: float = 0.5,
):
    """
    Plot UMAPs colored by marker sets for a given AnnData.

    - Uses adata.obsm['X_univi'] to build UMAP if needed.
    - marker_dict = { "group_name": [marker1, marker2, ...], ... }
    """
    os.makedirs(figdir, exist_ok=True)

    # Make sure we have a UMAP on UniVI latent
    ensure_umap_on_univi(adata, n_neighbors=n_neighbors, min_dist=min_dist)

    var_names = np.array(adata.var_names)

    for group, genes in marker_dict.items():
        # intersect with available features
        present = [g for g in genes if g in var_names]
        if len(present) == 0:
            print(f"[marker] In {title_prefix}, no markers from {group} present in var_names.")
            continue

        print(f"[marker] {title_prefix}: plotting {group} markers: {present}")
        sc.pl.umap(
            adata,
            color=present,
            size=50,
            alpha=0.8,
            show=False,
        )
        fname = f"umap_{title_prefix}_{group}_markers.png".replace(" ", "_")
        plt.savefig(os.path.join(figdir, fname), bbox_inches="tight")
        plt.show()
        plt.close()

def cluster_level_marker_deltas(
    adata,
    markers,
    cluster_key: str,
    mod: str,
    tag: str,
    figdir: str = FIGDIR,
):
    """
    For a set of markers, compare cluster-level raw vs denoised means.
    Assumes `adata.layers["univi_denoised"]` exists.
    """
    markers = [g for g in markers if g in adata.var_names]
    if not markers:
        print(f"[marker] No markers present for {tag} ({mod}) – skipping.")
        return

    X_raw = _to_dense(adata.X)
    X_den = _to_dense(adata.layers["univi_denoised"])

    idx = adata.var_names.get_indexer(markers)
    cluster_ids = sorted(adata.obs[cluster_key].unique())

    raw_means = []
    den_means = []

    for cl in cluster_ids:
        mask = adata.obs[cluster_key] == cl
        if mask.sum() == 0:
            raw_means.append(np.full(len(markers), np.nan))
            den_means.append(np.full(len(markers), np.nan))
        else:
            raw_means.append(X_raw[mask][:, idx].mean(axis=0))
            den_means.append(X_den[mask][:, idx].mean(axis=0))

    raw_means = np.vstack(raw_means)
    den_means = np.vstack(den_means)
    delta_means = den_means - raw_means

    # Cluster-level bar plots or heatmaps
    df_raw   = pd.DataFrame(raw_means,   index=cluster_ids, columns=markers)
    df_den   = pd.DataFrame(den_means,   index=cluster_ids, columns=markers)
    df_delta = pd.DataFrame(delta_means, index=cluster_ids, columns=markers)

    # Heatmap of raw marker expression
    plt.figure(figsize=(0.7 * len(markers) + 4, 0.4 * len(cluster_ids) + 3))
    sns.heatmap(df_raw, cmap="viridis", cbar_kws={"label": "Mean raw expression"})
    plt.xlabel("Markers")
    plt.ylabel(cluster_key)
    plt.title(f"{tag} ({mod}) – raw marker expression by cluster")
    plt.tight_layout()
    plt.savefig(os.path.join(figdir, f"markers_{tag}_{mod}_raw_by_{cluster_key}.png"))
    plt.show()
    plt.close()

    # Heatmap of delta (denoised - raw)
    v_abs = np.nanmax(np.abs(delta_means))
    plt.figure(figsize=(0.7 * len(markers) + 4, 0.4 * len(cluster_ids) + 3))
    sns.heatmap(
        df_delta,
        cmap="vlag",
        center=0,
        vmin=-v_abs,
        vmax=v_abs,
        cbar_kws={"label": "Δ denoised - raw"},
    )
    plt.xlabel("Markers")
    plt.ylabel(cluster_key)
    plt.title(f"{tag} ({mod}) – change in marker expression by cluster")
    plt.tight_layout()
    plt.savefig(os.path.join(figdir, f"markers_{tag}_{mod}_delta_by_{cluster_key}.png"))
    plt.show()
    plt.close()


# %% [code]
print(rna_test_adata.X.min())
print(rna_test_adata.X.max())


# %% [code]
import os
import numpy as np
import matplotlib.pyplot as plt
import seaborn as sns
import scanpy as sc

FIGDIR = "./figures/teaseq_univi_tri_modal_eval_reproducibility"
os.makedirs(FIGDIR, exist_ok=True)

UMAP_N_NEIGHBORS = 30
UMAP_RANDOM_STATE = 42

# ------------------------------------------------------
# 0) Build tri-modal combined object on shared UniVI latent
# ------------------------------------------------------
# Sanity: make sure UniVI latents exist
for adata, name in [
    (rna_test_adata,  "rna_test_adata"),
    (adt_test_adata,  "adt_test_adata"),
    (atac_test_adata, "atac_test_adata"),
]:
    if "X_univi" not in adata.obsm_keys():
        raise ValueError(f"{name} is missing 'X_univi'; run encoding cell first.")

# Temporary copies tagged by modality
rna_tmp  = rna_test_adata.copy()
adt_tmp  = adt_test_adata.copy()
atac_tmp = atac_test_adata.copy()

rna_tmp.obs["modality"]  = "rna"
adt_tmp.obs["modality"]  = "adt"
atac_tmp.obs["modality"] = "atac"

combined = rna_tmp.concatenate(
    adt_tmp,
    atac_tmp,
    join="outer",
    batch_key=None,
    index_unique=None,
)

# Stack UniVI latents in same order as concatenate
combined.obsm["X_univi"] = np.vstack([
    rna_test_adata.obsm["X_univi"],
    adt_test_adata.obsm["X_univi"],
    atac_test_adata.obsm["X_univi"],
])

# Neighbors / UMAP once on the **joint** latent
sc.pp.neighbors(combined, use_rep="X_univi", n_neighbors=UMAP_N_NEIGHBORS)
sc.tl.umap(combined, random_state=UMAP_RANDOM_STATE)

# Freeze this layout
combined.obsm["X_univi_umap"] = combined.obsm["X_umap"].copy()

# Split back to per-modality coordinates
mask_rna  = combined.obs["modality"] == "rna"
mask_adt  = combined.obs["modality"] == "adt"
mask_atac = combined.obs["modality"] == "atac"

rna_test_adata.obsm["X_univi_umap"]  = combined.obsm["X_univi_umap"][mask_rna, :].copy()
adt_test_adata.obsm["X_univi_umap"]  = combined.obsm["X_univi_umap"][mask_adt, :].copy()
atac_test_adata.obsm["X_univi_umap"] = combined.obsm["X_univi_umap"][mask_atac, :].copy()

# ------------------------------------------------------
# Helper so all downstream plots use the frozen layout
# ------------------------------------------------------
def ensure_umap_on_univi(adata):
    """
    Make sure adata.obsm['X_umap'] points to the *frozen* UniVI UMAP.

    We do NOT recompute neighbors/UMAP here, to keep the layout consistent.
    """
    if "X_univi_umap" not in adata.obsm_keys():
        raise ValueError("No 'X_univi_umap' found; run joint-UMAP cell first.")
    adata.obsm["X_umap"] = adata.obsm["X_univi_umap"]

# ------------------------------------------------------
# 0A) UMAPs colored by unimodal Leiden and modality
# ------------------------------------------------------

# RNA UniVI UMAP colored by unimodal RNA Leiden
ensure_umap_on_univi(rna_test_adata)
if "rna_leiden" in rna_test_adata.obs.columns:
    sc.pl.umap(
        rna_test_adata,
        color="rna_leiden",
        size=3,
        alpha=0.8,
        show=False,
    )
    plt.title("RNA UniVI UMAP – unimodal Leiden (rna_leiden)")
    plt.savefig(os.path.join(FIGDIR, "umap_rna_univi_rna_leiden.png"),
                bbox_inches="tight", dpi=200)
    plt.show()
    plt.close()
else:
    print("[WARN] 'rna_leiden' not found in rna_test_adata.obs; skipping RNA Leiden UMAP.")

# ADT UniVI UMAP colored by unimodal ADT Leiden
ensure_umap_on_univi(adt_test_adata)
if "adt_leiden" in adt_test_adata.obs.columns:
    sc.pl.umap(
        adt_test_adata,
        color="adt_leiden",
        size=3,
        alpha=0.8,
        show=False,
    )
    plt.title("ADT UniVI UMAP – unimodal Leiden (adt_leiden)")
    plt.savefig(os.path.join(FIGDIR, "umap_adt_univi_adt_leiden.png"),
                bbox_inches="tight", dpi=200)
    plt.show()
    plt.close()
else:
    print("[WARN] 'adt_leiden' not found in adt_test_adata.obs; skipping ADT Leiden UMAP.")

# ATAC UniVI UMAP colored by unimodal ATAC Leiden (if you have it)
ensure_umap_on_univi(atac_test_adata)
if "atac_leiden" in atac_test_adata.obs.columns:
    sc.pl.umap(
        atac_test_adata,
        color="atac_leiden",
        size=3,
        alpha=0.8,
        show=False,
    )
    plt.title("ATAC UniVI UMAP – unimodal Leiden (atac_leiden)")
    plt.savefig(os.path.join(FIGDIR, "umap_atac_univi_atac_leiden.png"),
                bbox_inches="tight", dpi=200)
    plt.show()
    plt.close()
else:
    print("[WARN] 'atac_leiden' not found in atac_test_adata.obs; skipping ATAC Leiden UMAP.")

# Joint tri-modal UMAP colored by modality
ensure_umap_on_univi(combined)  # points combined.obsm["X_umap"] to the frozen tri-modal layout
sc.pl.umap(
    combined,
    color="modality",
    size=3,
    alpha=0.8,
    show=False,
)
plt.title("UniVI UMAP – modality (RNA vs ADT vs ATAC)")
plt.savefig(os.path.join(FIGDIR, "umap_univi_rna_adt_atac_modality.png"),
            bbox_inches="tight", dpi=200)
plt.show()
plt.close()


# ------------------------------------------------------
# 1) Define richer marker panels (RNA + ADT)
# ------------------------------------------------------

rna_markers = {
    # CD4-ish / helper / memory T
    "CD4_like_T": [
        "CD4", "IL7R", "CCR7", "LEF1", "TCF7", "BCL11B", "IKZF2", "MEF2C", "RUNX3",
        "TNFRSF4", "TNFRSF18", "CXCR4", "CCR6"
    ],
    # Cytotoxic CD8 / NK-like
    "Cytotoxic_T_NK": [
        "CD8A", "CD8B", "PRF1", "GZMB", "GZMH", "GNLY", "NKG7", "IFNG", "KLRD1",
        "KLRK1", "CX3CR1", "CCL5"
    ],
    # B cells / plasmablasts
    "B_cell": [
        "MS4A1", "CD19", "CD22", "CD79A", "CD79B", "BANK1", "BLK", "PAX5",
        "CD74", "HLA-DRA", "HLA-DRB1", "HLA-DQB1",
        "IGKC", "IGLC1", "IGHD", "IGHM", "IGHA1", "IGHA2", "IGHE", "MZB1", "XBP1"
    ],
    # Mono / myeloid / DC
    "Mono_like": [
        "LYZ", "S100A8", "S100A9", "S100A12", "FCN1", "IL1B", "TNFAIP3", "NFKBIA",
        "LST1", "CSF2RA", "CSF3R", "ITGAX", "ITGAD", "CTSS", "LAPTM5", "SRGN",
        "CCR2", "CXCL8", "CD14"
    ],
    # General activation / exhaustion-ish markers
    "Activation": [
        "CD69", "IFNG", "TNFRSF9", "TNFRSF18", "TNFRSF4", "LAG3", "TIGIT", "PDCD1",
        "TOX", "BATF", "EGR1", "FOSB"
    ],
}

adt_markers = {
    "T_all": ["CD3", "TCR-a/b", "TCR-g/d"],
    "CD4_T": ["CD3", "CD4", "CD45RA", "CD45RO", "CD27", "CD127", "CD279"],
    "CD8_T": ["CD3", "CD8a", "CD45RA", "CD45RO", "KLRG1", "CD27", "CD279"],
    "NK": ["CD56", "CD16", "KLRG1"],
    "B_cell": ["CD19", "CD21", "CD24", "IgD", "IgM", "CD38"],
    "Mono_DC": ["CD14", "CD16", "CD11b", "CD11c", "HLA-DR", "CD141", "CD172a", "CD192", "CD304"],
    "Activation": ["CD25", "CD40", "CD80", "CD86", "CD71", "CD95", "CD278", "CD279"],
}

# ------------------------------------------------------
# 2) Helper: filter markers to those present in the AnnData
# ------------------------------------------------------

def filter_markers_to_var(marker_dict, adata, verbose=True, label=""):
    varset = set(adata.var_names)
    out = {}
    for grp, genes in marker_dict.items():
        present = [g for g in genes if g in varset]
        if verbose:
            missing = [g for g in genes if g not in varset]
            if present:
                print(f"[{label}] {grp}: using {len(present)} markers; missing: {missing}")
            else:
                print(f"[{label}] {grp}: no markers present; all missing: {missing}")
        if present:
            out[grp] = present
    return out

rna_markers_f = filter_markers_to_var(rna_markers, rna_test_adata, label="RNA")
adt_markers_f = filter_markers_to_var(adt_markers, adt_test_adata, label="ADT")

# ------------------------------------------------------
# 3) UMAP plotting of marker groups on UniVI latent UMAPs
# ------------------------------------------------------

def plot_markers_on_umap(adata, marker_dict, title_prefix, figdir=FIGDIR, size=200):
    ensure_umap_on_univi(adata)  # just points to frozen layout
    for group, genes in marker_dict.items():
        if not genes:
            continue
        print(f"[marker UMAP] {title_prefix}: {group} -> {genes}")
        sc.pl.umap(
            adata,
            color=genes,
            size=size,
            alpha=0.8,
            show=False,
        )
        fname = f"umap_{title_prefix}_{group}_markers.png".replace(" ", "_")
        plt.savefig(os.path.join(figdir, fname), bbox_inches="tight")
        plt.show()
        plt.close()

# RNA marker UMAPs (on UniVI latent)
plot_markers_on_umap(
    rna_test_adata,
    marker_dict=rna_markers_f,
    title_prefix="rna_univi_latent",
)

# ADT marker UMAPs (on UniVI latent)
plot_markers_on_umap(
    adt_test_adata,
    marker_dict=adt_markers_f,
    title_prefix="adt_univi_latent",
)

# ------------------------------------------------------
# 4) Cluster-level marker heatmaps (raw vs denoised) per modality
# ------------------------------------------------------

def cluster_marker_heatmaps(
    adata,
    marker_dict,
    cluster_key,
    mod: str,
    tag: str,
    layer_raw=None,             # None -> use .X
    layer_denoised="univi_denoised",
    figdir=FIGDIR,
):
    """
    For each marker group, compute cluster × marker mean (raw vs denoised)
    and plot heatmaps + delta heatmap.
    """
    if cluster_key not in adata.obs.columns:
        print(f"[{mod}] No cluster_key='{cluster_key}' in obs; skipping.")
        return

    # Get matrices
    if layer_raw is None:
        X_raw = _to_dense(adata.X)
        raw_label = "X"
    else:
        X_raw = _to_dense(adata.layers[layer_raw])
        raw_label = layer_raw

    if layer_denoised in adata.layers:
        X_den = _to_dense(adata.layers[layer_denoised])
        has_den = True
    else:
        X_den = None
        has_den = False
        print(f"[{mod}] layer '{layer_denoised}' not found; only raw heatmaps will be plotted.")

    var_index = pd.Index(adata.var_names)
    clusters = adata.obs[cluster_key].astype("category")
    cluster_categories = clusters.cat.categories

    for group, genes in marker_dict.items():
        if not genes:
            continue
        genes_present = [g for g in genes if g in var_index]
        if not genes_present:
            print(f"[{mod}] [{group}] no markers present; skipping.")
            continue

        cols = var_index.get_indexer(genes_present)

        # cluster × gene mean (raw)
        df_raw = []
        for cl in cluster_categories:
            mask = (clusters == cl).values
            if not mask.any():
                continue
            m = X_raw[mask][:, cols].mean(axis=0)
            df_raw.append(pd.Series(m, index=genes_present, name=str(cl)))
        df_raw = pd.DataFrame(df_raw)

        # Plot raw heatmap
        plt.figure(figsize=(0.5 * len(genes_present) + 4, 0.5 * len(df_raw) + 3))
        sns.heatmap(
            df_raw,
            cmap="viridis",
            cbar_kws={"label": f"Mean {raw_label}"},
        )
        plt.xlabel("Markers")
        plt.ylabel(cluster_key)
        plt.title(f"{tag} ({mod}) – {group} markers (raw)")
        plt.tight_layout()
        fname = f"heatmap_{tag}_{mod}_{group}_raw.png".replace(" ", "_")
        plt.savefig(os.path.join(figdir, fname))
        plt.show()
        plt.close()

        if has_den:
            # cluster × gene mean (denoised)
            df_den = []
            for cl in cluster_categories:
                mask = (clusters == cl).values
                if not mask.any():
                    continue
                m = X_den[mask][:, cols].mean(axis=0)
                df_den.append(pd.Series(m, index=genes_present, name=str(cl)))
            df_den = pd.DataFrame(df_den)

            # Denoised heatmap
            plt.figure(figsize=(0.5 * len(genes_present) + 4, 0.5 * len(df_den) + 3))
            sns.heatmap(
                df_den,
                cmap="viridis",
                cbar_kws={"label": f"Mean {layer_denoised}"},
            )
            plt.xlabel("Markers")
            plt.ylabel(cluster_key)
            plt.title(f"{tag} ({mod}) – {group} markers (denoised)")
            plt.tight_layout()
            fname = f"heatmap_{tag}_{mod}_{group}_denoised.png".replace(" ", "_")
            plt.savefig(os.path.join(figdir, fname))
            plt.show()
            plt.close()

            # Delta (denoised - raw)
            df_delta = df_den - df_raw
            plt.figure(figsize=(0.5 * len(genes_present) + 4, 0.5 * len(df_delta) + 3))
            sns.heatmap(
                df_delta,
                cmap="vlag",
                center=0,
                cbar_kws={"label": "Mean (denoised - raw)"},
            )
            plt.xlabel("Markers")
            plt.ylabel(cluster_key)
            plt.title(f"{tag} ({mod}) – {group} markers Δ (den - raw)")
            plt.tight_layout()
            fname = f"heatmap_{tag}_{mod}_{group}_delta.png".replace(" ", "_")
            plt.savefig(os.path.join(figdir, fname))
            plt.show()
            plt.close()

'''
# RNA cluster-level marker heatmaps
cluster_marker_heatmaps(
    rna_test_adata,
    marker_dict=rna_markers_f,
    cluster_key="rna_leiden",
    mod="rna",
    tag="rna_test_adata",
    layer_raw=None,              # uses .X (your working RNA space)
    layer_denoised="univi_denoised",
)

# ADT cluster-level marker heatmaps
cluster_marker_heatmaps(
    adt_test_adata,
    marker_dict=adt_markers_f,
    cluster_key="adt_leiden",
    mod="adt",
    tag="adt_test_adata",
    layer_raw="counts",          # your raw ADT (e.g. CLR or arcsinh before UniVI)
    layer_denoised="univi_denoised",
)
'''

# %% [code]
# ------------------------------------------------------
# 5) ATAC LSI ↔ RNA marker correlations
# ------------------------------------------------------

from scipy.stats import pearsonr

def correlate_lsi_with_rna_markers(atac_adata, rna_adata, genes, layer_rna=None, top_k=5):
    """
    For each gene, compute Pearson corr between gene expression (RNA)
    and each ATAC LSI dimension. Return full table + top_k per gene.
    """
    assert np.array_equal(atac_adata.obs_names, rna_adata.obs_names), \
        "ATAC and RNA obs_names must match 1:1."

    X_lsi = _to_dense(atac_adata.X)  # cells × n_lsi

    if layer_rna is None:
        X_rna = _to_dense(rna_adata.X)
    else:
        X_rna = _to_dense(rna_adata.layers[layer_rna])
    var_index = pd.Index(rna_adata.var_names)

    results = []
    for g in genes:
        if g not in var_index:
            print(f"[ATAC-RNA corr] {g} not in rna var_names; skipping.")
            continue
        g_idx = var_index.get_loc(g)
        g_vec = X_rna[:, g_idx].ravel()

        # skip all-constant genes
        if np.allclose(g_vec, g_vec[0]):
            print(f"[ATAC-RNA corr] {g} is constant across cells; skipping.")
            continue

        for k in range(X_lsi.shape[1]):
            z = X_lsi[:, k]
            r, _ = pearsonr(z, g_vec)
            results.append({"gene": g, "lsi_dim": k, "corr": r})

    df = pd.DataFrame(results)
    if df.empty:
        print("No correlations computed (no genes found).")
        return df, df

    df = df.assign(abs_corr=lambda d: d["corr"].abs())  # type: ignore
    df_top = (df
              .sort_values(["gene", "abs_corr"], ascending=[True, False])
              .groupby("gene")
              .head(top_k))
    return df, df_top

# Pick a marker subset to probe ATAC LSI
genes_for_atac = [
    # B cell / Ig
    "MS4A1", "CD74", "HLA-DRA", "IGKC", "IGHM",
    # Cytotoxic / NK
    "PRF1", "GZMH", "GNLY", "NKG7", "IFNG",
    # Mono / myeloid
    "S100A8", "S100A9", "FCN1", "IL1B", "LYZ",
]

df_corr, df_corr_top = correlate_lsi_with_rna_markers(
    atac_test_adata,
    rna_test_adata,
    genes_for_atac,
    layer_rna=None,  # or "log1p" etc if you prefer
    top_k=3,
)

print("\nTop LSI dims per marker gene:")
print(df_corr_top)

# Optionally: visualize some high-correlation LSI dims on an ATAC UMAP (UniVI latent)
ensure_umap_on_univi(atac_test_adata)

X_lsi = _to_dense(atac_test_adata.X)
lsi_dims_to_plot = sorted(df_corr_top["lsi_dim"].unique())[:6]

for d in lsi_dims_to_plot:
    col = f"LSI_dim_{d}"   # avoid conflict with var_names like "LSI_1"
    atac_test_adata.obs[col] = X_lsi[:, d]

    sc.pl.umap(
        atac_test_adata,
        color=col,
        size=3,
        alpha=0.8,
        show=False,
    )
    fname = f"umap_atac_univi_{col}.png"
    plt.savefig(os.path.join(FIGDIR, fname), bbox_inches="tight")
    plt.show()
    plt.close()


# %% [code]
print(atac_test_adata)


# %% [code]
# ------------------------------------------------------
# Plot top 3 ATAC LSI dims enriched in NK/cytotoxic clusters
# on the SAME UniVI UMAP you're already using for ATAC plots
# ------------------------------------------------------

import numpy as np
import pandas as pd
import scanpy as sc
import scipy.sparse as sp

def _to_dense(X):
    return X.toarray() if sp.issparse(X) else np.asarray(X)

def rank_lsi_dims_for_clusters(
    atac_adata: sc.AnnData,
    *,
    cluster_key: str,
    target_clusters,
    lsi_key: str = "X_lsi",
    top_n: int = 10,
):
    """
    Rank LSI dims by separation of target_clusters vs others using Cohen's d.
    Returns df sorted by abs(d) desc.
    """
    if cluster_key not in atac_adata.obs:
        raise KeyError(f"{cluster_key!r} not in atac_adata.obs")

    if lsi_key not in atac_adata.obsm:
        raise KeyError(f"{lsi_key!r} not in atac_adata.obsm (expected LSI embedding)")

    X_lsi = _to_dense(atac_adata.obsm[lsi_key])  # cells x n_lsi
    clust = atac_adata.obs[cluster_key].astype(str).to_numpy()

    target_set = set([str(x) for x in target_clusters])
    m = np.array([c in target_set for c in clust], dtype=bool)

    if m.sum() == 0:
        raise ValueError(f"No cells found in target_clusters={target_clusters} under {cluster_key!r}")

    X_in  = X_lsi[m]
    X_out = X_lsi[~m]

    mu_in  = X_in.mean(axis=0)
    mu_out = X_out.mean(axis=0)
    sd_in  = X_in.std(axis=0, ddof=1)
    sd_out = X_out.std(axis=0, ddof=1)
    pooled = np.sqrt((sd_in**2 + sd_out**2) / 2.0)

    d = (mu_in - mu_out) / np.maximum(pooled, 1e-8)

    df = pd.DataFrame({
        "lsi_dim": np.arange(X_lsi.shape[1]),
        "cohens_d": d,
        "abs_d": np.abs(d),
        "mu_in": mu_in,
        "mu_out": mu_out,
    }).sort_values("abs_d", ascending=False)

    return df

# ----------------------------
# Use THIS object (the one you already plot on)
# ----------------------------
atac = atac_test_adata

# Ensure this is the same UniVI UMAP you’ve been using
# (your helper should create/ensure .obsm["X_umap"])
ensure_umap_on_univi(atac)

# ----------------------------
# Pick the NK/cytotoxic clusters you suspect
# ----------------------------
cluster_key = "atac_leiden"
nk_clusters = ["1", "5", "6"]

# ----------------------------
# Rank LSI dims enriched in those clusters
# ----------------------------
df_rank = rank_lsi_dims_for_clusters(
    atac,
    cluster_key=cluster_key,
    target_clusters=nk_clusters,
    lsi_key="X_lsi",
    top_n=100,
)

print(df_rank.head(25))

#top3 = df_rank["lsi_dim"].iloc[:3].tolist()
#print("Top 3 LSI dims for clusters", nk_clusters, ":", top3)

# ----------------------------
# Attach those dims to atac.obs and plot on the SAME UMAP
# ----------------------------
X_lsi = _to_dense(atac.obsm["X_lsi"])

top3 = [3, 4, 9]

for d in top3:
    col = f"LSI_dim_{d}"
    atac.obs[col] = X_lsi[:, d]

sc.pl.umap(
    atac,
    color=[f"LSI_dim_{d}" for d in top3],
    ncols=3,
    size=3,
    alpha=0.8,
    cmap="viridis",
)


# %% [code]
# ------------------------------------------------------
# 6) Sample cells per unimodal cluster across modalities
# ------------------------------------------------------

def sample_per_cluster(adata, cluster_key, n_per_cluster=2000, random_state=0):
    """
    Sample up to n_per_cluster cells from each cluster in cluster_key.
    Returns a new AnnData subset.
    """
    if cluster_key not in adata.obs.columns:
        raise ValueError(f"cluster_key='{cluster_key}' not in adata.obs")

    rng = np.random.default_rng(random_state)
    clusters = adata.obs[cluster_key].astype("category")
    idx_keep = []

    for cl in clusters.cat.categories:
        mask = (clusters == cl).values
        cell_indices = np.where(mask)[0]
        if len(cell_indices) == 0:
            continue
        n_take = min(n_per_cluster, len(cell_indices))
        chosen = rng.choice(cell_indices, size=n_take, replace=False)
        idx_keep.extend(chosen)

    idx_keep = sorted(idx_keep)
    print(f"[sample_per_cluster] {cluster_key}: keeping {len(idx_keep)} cells "
          f"(<= {n_per_cluster} per cluster)")
    return adata[idx_keep].copy()

n_per_cluster = 2000  # upper bound; will just cap at cluster size if smaller

rna_sample  = sample_per_cluster(rna_test_adata,  "rna_leiden",  n_per_cluster, random_state=42)
adt_sample  = sample_per_cluster(adt_test_adata,  "adt_leiden",  n_per_cluster, random_state=42)
atac_sample = sample_per_cluster(atac_test_adata, "atac_leiden", n_per_cluster, random_state=42)

# Tag modality for sampled sets
rna_sample.obs["univi_source"]  = "rna"
adt_sample.obs["univi_source"]  = "adt"
atac_sample.obs["univi_source"] = "atac"

# ------------------------------------------------------
# 7) Build combined sampled object in UniVI latent
# ------------------------------------------------------

combined_sample = rna_sample.concatenate(
    adt_sample,
    atac_sample,
    join="outer",
    batch_key="univi_batch",
    batch_categories=["rna", "adt", "atac"],
    index_unique=None,
)

# Stack UniVI latents in the same order as concatenate
combined_sample.obsm["X_univi"] = np.vstack([
    rna_sample.obsm["X_univi"],
    adt_sample.obsm["X_univi"],
    atac_sample.obsm["X_univi"],
])

# Neighbors/UMAP on UniVI latent for the sampled cells
sc.pp.neighbors(combined_sample, use_rep="X_univi", n_neighbors=30)
sc.tl.umap(combined_sample)

# ------------------------------------------------------
# 8) UMAP visualizations: modality + unimodal clusters
# ------------------------------------------------------

# UMAP colored by modality
sc.pl.umap(
    combined_sample,
    color="univi_source",
    size=3,
    alpha=0.8,
    show=False,
)
plt.savefig(os.path.join(FIGDIR, "umap_sampled_tri_modal_by_modality.png"), bbox_inches="tight")
plt.show()
plt.close()

# UMAP colored by each unimodal Leiden labelling (pseudo-celltypes)
for key in ["rna_leiden", "adt_leiden", "atac_leiden"]:
    if key in combined_sample.obs.columns:
        print(f"[sampled UMAP] Coloring by {key}")
        sc.pl.umap(
            combined_sample,
            color=key,
            size=3,
            alpha=0.8,
            show=False,
        )
        fname = f"umap_sampled_tri_modal_{key}.png"
        plt.savefig(os.path.join(FIGDIR, fname), bbox_inches="tight")
        plt.show()
        plt.close()
    else:
        print(f"[sampled UMAP] {key} not in combined_sample.obs; skipping.")

# ------------------------------------------------------
# 9) (Optional) Use denoised reconstructions for sampled cells
# ------------------------------------------------------
# If you already ran univi_eval.denoise_adata on full test sets, the sampled
# objects will inherit the 'univi_denoised' layer. If not, you can run it here:

for adata_sample, mod, tag in [
    (rna_sample,  "rna",  "rna_sample"),
    (adt_sample,  "adt",  "adt_sample"),
    (atac_sample, "atac", "atac_sample"),
]:
    if "univi_denoised" not in adata_sample.layers:
        print(f"[denoise sampled] Running decoder for {tag} ({mod})...")
        univi_eval.denoise_adata(
            model,
            adata_sample,
            modality=mod,
            device=device,
            batch_size=512,
            out_layer="univi_denoised",
        )

# Now you can re-use cluster_marker_heatmaps on rna_sample/adt_sample
cluster_marker_heatmaps(
    rna_sample,
    marker_dict=rna_markers_f,
    cluster_key="rna_leiden",
    mod="rna",
    tag="rna_sample",
    layer_raw=None,
    layer_denoised="univi_denoised",
)

cluster_marker_heatmaps(
    adt_sample,
    marker_dict=adt_markers_f,
    cluster_key="adt_leiden",
    mod="adt",
    tag="adt_sample",
    layer_raw=None,
    #layer_raw='counts',
    layer_denoised="univi_denoised",
)


# %% [code]
import numpy as np
import pandas as pd
import matplotlib.pyplot as plt
import seaborn as sns

from univi import evaluation as univi_eval

# ------------------------------------------------------
# 1) Cross-modal predictions from RNA to ADT and ATAC
# ------------------------------------------------------

print("\n[Cross-modal] Predicting ADT and ATAC from RNA test data...")

# RNA → ADT
Xhat_adt_from_rna = univi_eval.cross_modal_predict(
    model,
    adata_src=rna_test_adata,
    src_mod="rna",
    tgt_mod="adt",
    device=device,
    batch_size=512,
)
# shape check
print("  RNA→ADT prediction shape:", Xhat_adt_from_rna.shape)

adt_test_adata.layers["univi_pred_from_rna"] = Xhat_adt_from_rna

# RNA → ATAC
Xhat_atac_from_rna = univi_eval.cross_modal_predict(
    model,
    adata_src=rna_test_adata,
    src_mod="rna",
    tgt_mod="atac",
    device=device,
    batch_size=512,
)
print("  RNA→ATAC prediction shape:", Xhat_atac_from_rna.shape)

atac_test_adata.layers["univi_pred_from_rna"] = Xhat_atac_from_rna

print("Stored predictions in:")
print("  adt_test_adata.layers['univi_pred_from_rna']")
print("  atac_test_adata.layers['univi_pred_from_rna']")


# %% [code]
# ------------------------------------------------------
# 2) ADT cluster-level marker heatmaps: true vs RNA-predicted
# ------------------------------------------------------
# Assumes:
#   - adt_markers_f (ADT marker dict filtered to var_names)
#   - cluster_marker_heatmaps(...) defined previously
#   - ADT "true" data live in .layers["counts"] (or whatever you used)

cluster_marker_heatmaps(
    adata=adt_test_adata,
    marker_dict=adt_markers_f,
    cluster_key="adt_leiden",
    mod="adt",
    tag="adt_test_from_rna",
    layer_raw=None,                        # true ADT signal
    layer_denoised="univi_pred_from_rna",  # RNA→ADT prediction
)


# %% [code]
# ------------------------------------------------------
# 3) ADT: nearest-centroid cluster prediction using RNA→ADT
# ------------------------------------------------------

def assign_clusters_by_centroid(
    adata,
    cluster_key: str,
    layer: str,
    out_key: str,
):
    """
    For each cell, assign the cluster whose centroid (in given layer) is
    closest in Euclidean distance.
    """
    if cluster_key not in adata.obs.columns:
        raise ValueError(f"{cluster_key} not in adata.obs")

    if layer not in adata.layers:
        raise ValueError(f"{layer} not in adata.layers")

    X = _to_dense(adata.layers[layer])  # cells × features
    clusters = adata.obs[cluster_key].astype("category")
    cats = clusters.cat.categories

    centroids = []
    for cl in cats:
        mask = (clusters == cl).values
        if not mask.any():
            centroids.append(np.nan)
            continue
        centroids.append(X[mask].mean(axis=0))
    centroids = np.vstack(centroids)  # n_clusters × features

    # distances: cells × clusters
    dists = np.linalg.norm(X[:, None, :] - centroids[None, :, :], axis=2)
    idx_min = np.argmin(dists, axis=1)
    assigned = cats[idx_min]

    adata.obs[out_key] = pd.Categorical(assigned, categories=cats)
    print(f"[assign_clusters_by_centroid] Wrote predicted clusters to obs['{out_key}'].")


def confusion_true_vs_pred(
    adata,
    true_key: str,
    pred_key: str,
    tag: str,
    normalize: str = "index",
    figdir=FIGDIR,
):
    """
    Confusion matrix (normalized crosstab) between true and predicted clusters.
    """
    s_true = adata.obs[true_key].astype("category")
    s_pred = adata.obs[pred_key].astype("category")

    tab = pd.crosstab(s_true, s_pred, normalize=normalize)

    plt.figure(figsize=(0.5 * tab.shape[1] + 4, 0.5 * tab.shape[0] + 4))
    sns.heatmap(
        tab,
        cmap="viridis",
        annot=False,
        cbar_kws={"label": f"Fraction (normalized by {normalize})"},
    )
    plt.xlabel(f"Predicted ({pred_key})")
    plt.ylabel(f"True ({true_key})")
    plt.title(f"Cluster confusion: {tag}")
    plt.tight_layout()
    fname = f"cluster_confusion_{tag}.png".replace(" ", "_")
    plt.savefig(os.path.join(figdir, fname))
    plt.show()
    plt.close()

    return tab

# Predict ADT clusters using RNA->ADT predicted profiles
assign_clusters_by_centroid(
    adata=adt_test_adata,
    cluster_key="adt_leiden",
    layer="univi_pred_from_rna",
    out_key="adt_leiden_pred_from_rna",
)

# Confusion matrix: how well RNA->ADT recovers ADT clusters?
tab_adt_conf = confusion_true_vs_pred(
    adata=adt_test_adata,
    true_key="adt_leiden",
    pred_key="adt_leiden_pred_from_rna",
    tag="ADT_true_vs_RNA_predicted",
    normalize="index",
)
print("\nADT cluster confusion (true vs RNA-predicted features):")
print(tab_adt_conf)


# %% [code]
print(adt_test_adata)
print(adt_test_adata.X.min())
print(adt_test_adata.X.max())

adt_test_adata.layers['scaled'] = adt_test_adata.X


# %% [code]
# ------------------------------------------------------
# 4) ADT: UMAPs of true vs RNA-predicted marker expression
# ------------------------------------------------------

def plot_true_vs_pred_markers_on_umap(
    adata,
    markers,
    layer_true="scaled",
    layer_pred="univi_pred_from_rna",
    title_prefix="adt_true_vs_rna_pred",
    figdir=FIGDIR,
    size=200,
):
    if layer_true not in adata.layers:
        raise ValueError(f"{layer_true} not in adata.layers")
    if layer_pred not in adata.layers:
        raise ValueError(f"{layer_pred} not in adata.layers")

    ensure_umap_on_univi(adata)  # UMAP on X_univi

    X_true = _to_dense(adata.layers[layer_true])
    X_pred = _to_dense(adata.layers[layer_pred])

    var_index = pd.Index(adata.var_names)

    for m in markers:
        if m not in var_index:
            print(f"[UMAP true vs pred] Marker {m} not in ADT var_names; skipping.")
            continue
        idx = var_index.get_loc(m)
        adata.obs[f"{m}_true"] = X_true[:, idx]
        adata.obs[f"{m}_pred"] = X_pred[:, idx]

        print(f"[UMAP true vs pred] Plotting {m} (true vs RNA-pred).")
        sc.pl.umap(
            adata,
            color=[f"{m}_true", f"{m}_pred"],
            size=size,
            alpha=0.8,
            show=False,
        )
        fname = f"umap_{title_prefix}_{m}.png".replace(" ", "_")
        plt.savefig(os.path.join(figdir, fname), bbox_inches="tight")
        plt.show()
        plt.close()

# example ADT markers to inspect
adt_markers_to_plot = ["CD3", "CD4", "CD8a", "CD19", "CD14", "CD56", "HLA-DR"]

plot_true_vs_pred_markers_on_umap(
    adt_test_adata,
    markers=adt_markers_to_plot,
    layer_true="scaled",
    layer_pred="univi_pred_from_rna",
    title_prefix="adt_true_vs_rna_pred",
)


# %% [code]
# ------------------------------------------------------
# 5) ATAC: cluster-level comparison of true vs RNA-predicted ATAC features
# ------------------------------------------------------

def atac_cluster_feature_heatmaps(
    atac_adata,
    cluster_key="atac_leiden",
    layer_true="X",                    # treat .X as "true"
    layer_pred="univi_pred_from_rna",
    tag="atac_test_from_rna",
    figdir=FIGDIR,
    max_features_for_plot=40,
):
    """
    Compare per-cluster feature means between true ATAC (.X) and RNA-predicted ATAC.
    Features = columns of atac_adata.X (e.g., LSI dims).
    """
    if cluster_key not in atac_adata.obs.columns:
        print(f"[ATAC] No cluster_key='{cluster_key}' in obs; skipping.")
        return
    if layer_pred not in atac_adata.layers:
        print(f"[ATAC] No predicted layer '{layer_pred}'; skipping.")
        return

    # True ATAC feature matrix
    if layer_true == "X":
        X_true = _to_dense(atac_adata.X)
        true_label = "X"
    else:
        X_true = _to_dense(atac_adata.layers[layer_true])
        true_label = layer_true

    X_pred = _to_dense(atac_adata.layers[layer_pred])

    clusters = atac_adata.obs[cluster_key].astype("category")
    cats = clusters.cat.categories

    # For plotting we may want to restrict to features with highest variance
    var_true = X_true.var(axis=0)
    idx_sorted = np.argsort(var_true)[::-1]
    idx_plot = idx_sorted[:max_features_for_plot]

    feature_names = [str(atac_adata.var_names[i]) for i in idx_plot]

    # cluster × feature means (true)
    df_true = []
    for cl in cats:
        mask = (clusters == cl).values
        if not mask.any():
            continue
        m = X_true[mask][:, idx_plot].mean(axis=0)
        df_true.append(pd.Series(m, index=feature_names, name=str(cl)))
    df_true = pd.DataFrame(df_true)

    # cluster × feature means (pred)
    df_pred = []
    for cl in cats:
        mask = (clusters == cl).values
        if not mask.any():
            continue
        m = X_pred[mask][:, idx_plot].mean(axis=0)
        df_pred.append(pd.Series(m, index=feature_names, name=str(cl)))
    df_pred = pd.DataFrame(df_pred)

    # True heatmap
    plt.figure(figsize=(0.4 * len(feature_names) + 4, 0.5 * len(df_true) + 3))
    sns.heatmap(
        df_true,
        cmap="viridis",
        cbar_kws={"label": f"Mean {true_label}"},
    )
    plt.xlabel("ATAC features (e.g., LSI dims)")
    plt.ylabel(cluster_key)
    plt.title(f"{tag} – ATAC (true)")
    plt.tight_layout()
    plt.savefig(os.path.join(figdir, f"heatmap_{tag}_true.png"))
    plt.show()
    plt.close()

    # Predicted heatmap
    plt.figure(figsize=(0.4 * len(feature_names) + 4, 0.5 * len(df_pred) + 3))
    sns.heatmap(
        df_pred,
        cmap="viridis",
        cbar_kws={"label": "Mean pred_from_rna"},
    )
    plt.xlabel("ATAC features (e.g., LSI dims)")
    plt.ylabel(cluster_key)
    plt.title(f"{tag} – ATAC (RNA-predicted)")
    plt.tight_layout()
    plt.savefig(os.path.join(figdir, f"heatmap_{tag}_pred.png"))
    plt.show()
    plt.close()

    # Delta heatmap
    df_delta = df_pred - df_true
    plt.figure(figsize=(0.4 * len(feature_names) + 4, 0.5 * len(df_delta) + 3))
    sns.heatmap(
        df_delta,
        cmap="vlag",
        center=0,
        cbar_kws={"label": "Mean (pred - true)"},
    )
    plt.xlabel("ATAC features (e.g., LSI dims)")
    plt.ylabel(cluster_key)
    plt.title(f"{tag} – ATAC Δ (pred - true)")
    plt.tight_layout()
    plt.savefig(os.path.join(figdir, f"heatmap_{tag}_delta.png"))
    plt.show()
    plt.close()

    return df_true, df_pred, df_delta

df_atac_true, df_atac_pred, df_atac_delta = atac_cluster_feature_heatmaps(
    atac_test_adata,
    cluster_key="atac_leiden",
    layer_true="X",                     # .X as true LSI/features
    layer_pred="univi_pred_from_rna",
    tag="atac_from_rna",
    max_features_for_plot=40,
)


# %% [code]
# ------------------------------------------------------
# 6) ATAC: UMAP of selected feature dims (true vs RNA-pred)
# ------------------------------------------------------

def plot_atac_dims_true_vs_pred_umap(
    atac_adata,
    dims,
    layer_pred="univi_pred_from_rna",
    title_prefix="atac_true_vs_rna_pred_dims",
    figdir=FIGDIR,
    size=200,
):
    """
    dims = list of integer feature indices (columns in atac_adata.X).
    """
    ensure_umap_on_univi(atac_adata)

    X_true = _to_dense(atac_adata.X)
    X_pred = _to_dense(atac_adata.layers[layer_pred])

    for d in dims:
        if d < 0 or d >= atac_adata.n_vars:
            print(f"[ATAC UMAP] dim {d} out of range; skipping.")
            continue
        name = str(atac_adata.var_names[d])
        atac_adata.obs[f"ATAC_dim_{d}_true"] = X_true[:, d]
        atac_adata.obs[f"ATAC_dim_{d}_pred"] = X_pred[:, d]

        sc.pl.umap(
            atac_adata,
            color=[f"ATAC_dim_{d}_true", f"ATAC_dim_{d}_pred"],
            size=size,
            alpha=0.8,
            show=False,
        )
        fname = f"umap_{title_prefix}_dim_{d}.png"
        plt.savefig(os.path.join(figdir, fname), bbox_inches="tight")
        plt.show()
        plt.close()

# Example: just plot the first few dims
dims_to_plot = [0, 1, 2, 3, 4, 5, 6, 7, 8, 9, 10, 49, 99]
plot_atac_dims_true_vs_pred_umap(
    atac_test_adata,
    dims=dims_to_plot,
)


# %% [code]


# %% [code]
# ------------------------------------------------------------------
# RNA-gene ↔ ADT-marker mapping, tailored to your RNA var_names
# (genes that I *can see* in your provided var set are listed first)
# ------------------------------------------------------------------

rna_gene_map_for_adt_markers = {
    # -------------------------
    # T cell core markers
    # -------------------------
    # CD3 complex – only CD247 (CD3ζ) is clearly in your HVGs
    "CD3": ["CD247", "CD3D", "CD3E", "CD3G"],

    # CD4 helper / naïve-like: no CD4 gene itself, so we use good proxies
    "CD4": ["GATA3", "TCF7", "LEF1", "BCL11B", "IL7R"],

    # CD8 / cytotoxic T: no CD8A in your set, but strong cytotoxic signature is present
    "CD8": ["NKG7", "GNLY", "PRF1", "GZMH", "STAT4", "CD8A", "CD8B"],

    # Early activation
    "CD69": ["CD69", "NR4A2", "TNFAIP3"],

    # Co-stimulatory-ish / surface T cell marker
    "CD6": ["CD6"],

    # -------------------------
    # NK / cytotoxic markers
    # -------------------------
    # CD56 = NCAM1; your RNA has NCAM1 + classic NK/cytotoxic genes
    "CD56": ["NCAM1", "NKG7", "GNLY", "PRF1", "GZMH"],

    # If you have an NKG2D ADT (sometimes in panels)
    "NKG2D": ["NKG7", "GNLY", "PRF1"],

    # Generic “NK / cytotoxic” channel if you have one
    "NK_signature": ["NKG7", "GNLY", "PRF1", "GZMH", "STAT4"],

    # -------------------------
    # B cell markers
    # -------------------------
    # Your RNA HVGs clearly include CD22, MS4A1 (CD20), BANK1, PAX5, FCER2, FCRLA, etc.
    "CD19": ["CD22", "MS4A1", "BANK1", "PAX5", "CD19"],
    "CD20": ["MS4A1", "BANK1", "PAX5"],
    "CD22": ["CD22", "BANK1", "PAX5"],
    "CD23": ["FCER2"],
    "CD74": ["CD74", "HLA-DRA", "HLA-DRB1"],

    # More B-cell-ish markers if you have them in ADT
    "FCRL1": ["FCRL1", "BANK1", "PAX5"],
    "IgM":   ["IGHM", "IGKC"],
    "IgA":   ["IGHA1", "IGHA2"],
    "IgD":   ["IGHD"],

    # -------------------------
    # Myeloid / mono / DC
    # -------------------------
    # Monocytes / neutrophils – CD14 gene itself is not in your list, so use myeloid proxies
    "CD14": ["LYZ", "S100A8", "S100A9", "FCN1"],

    # CD16 – FCGR3B is clearly in your var_names (neutrophils)
    "CD16": ["FCGR3B"],

    # CD11c – ITGAX is present as a gene
    "CD11c": ["ITGAX"],
    "ITGAX": ["ITGAX"],  # in case the ADT channel is actually named ITGAX

    # Chemokine receptors
    "CCR2": ["CCR2"],

    # Cross-presenting DCs (XCR1 ADT often maps well to XCL1/XCL2 expression)
    "XCR1": ["XCL1", "XCL2"],

    # -------------------------
    # MHC-II / antigen presentation
    # -------------------------
    # You have HLA-DRA, HLA-DRB1, HLA-DQB1, CD74
    "HLA-DR": ["HLA-DRA", "HLA-DRB1", "HLA-DQB1", "CD74"],
    "HLA-DQ": ["HLA-DQB1", "HLA-DQA1"],  # HLA-DQA1 may or may not be present
    "HLA-DP": ["HLA-DPA1", "HLA-DPB1"],  # optional; will be auto-dropped if absent

    # -------------------------
    # Activation / costim / checkpoints
    # (many of these genes may *not* be in your HVGs, but we include them
    # so your code can still use this dict with other datasets)
    # -------------------------
    "CD83": ["CD83"],
    "CD96": ["CD96"],

    # Classic T cell activation / exhaustion markers
    "OX40":  ["TNFRSF4"],
    "4-1BB": ["TNFRSF9"],
    "PD-1":  ["PDCD1"],
    "CTLA4": ["CTLA4"],
    "TIGIT": ["TIGIT"],

    # -------------------------
    # T-reg / Th skewing
    # -------------------------
    "FOXP3": ["FOXP3", "IL2RA"],
    "GATA3": ["GATA3"],
    "TCF7_hi": ["TCF7", "LEF1", "BACH2"],

    # -------------------------
    # Cytokines / effector molecules
    # -------------------------
    "IFNg": ["IFNG", "IFNG-AS1"],
    "IL1b": ["IL1B"],
    "TNFa": ["TNF"],  # may not be present in this HVG set, but common in other datasets
}


# %% [code]
import numpy as np
import pandas as pd
import scanpy as sc
import matplotlib.pyplot as plt

# ------------------------------------------------------
# 4) ADT: UMAPs of true vs RNA-predicted marker expression
#     + RNA expression (mapped from marker -> gene symbol[s])
# ------------------------------------------------------

def _guess_rna_genes_for_marker(marker: str) -> list[str]:
    """
    Very simple heuristic to propose RNA gene symbols from an ADT marker name.
    Used only if rna_gene_map does not specify anything.
    """
    cand = []

    # Raw marker
    cand.append(marker)

    # Uppercase
    up = marker.upper()
    if up not in cand:
        cand.append(up)

    # Remove hyphens/slashes
    stripped = up.replace("-", "").replace("/", "")
    if stripped not in cand:
        cand.append(stripped)

    # Handle common CD8a / CD8A / CD8α style
    if up.startswith("CD8") and up not in cand:
        cand.append("CD8A")

    # HLA-DR → HLA-DRA / HLA-DRB1 guesses
    if up.startswith("HLA-DR"):
        for g in ["HLA-DRA", "HLA-DRB1", "HLA-DRB5"]:
            if g not in cand:
                cand.append(g)

    return cand


def plot_true_vs_pred_markers_on_umap_with_rna(
    adt_adata,
    rna_adata,
    markers,
    rna_gene_map: dict[str, list[str]] | None = None,
    layer_true: str = "scaled",              # ADT "true" (e.g. CLR/scale)
    layer_pred: str = "univi_pred_from_rna", # ADT predicted from RNA
    rna_layer: str | None = "log1p",         # RNA layer for expression; if None, use .X
    title_prefix: str = "adt_true_vs_rna_pred_with_rna",
    figdir: str = FIGDIR,
    size: int = 200,
):
    """
    For each marker in `markers`, plot UMAPs (on ADT UniVI latent) colored by:
      - ADT true expression
      - ADT predicted from RNA (cross-modal decoder)
      - RNA expression of corresponding gene(s), if present in RNA var_names

    RNA mapping:
      * Primary: rna_gene_map[marker] -> list of gene symbols
      * Fallback: simple heuristics (_guess_rna_genes_for_marker)
    """
    if rna_gene_map is None:
        rna_gene_map = {}

    # Sanity: same cells in same order
    assert np.array_equal(
        adt_adata.obs_names, rna_adata.obs_names
    ), "ADT and RNA obs_names must match 1:1 for per-cell comparisons."

    # Check ADT layers
    if layer_true not in adt_adata.layers:
        raise ValueError(f"{layer_true} not in adt_adata.layers")
    if layer_pred not in adt_adata.layers:
        raise ValueError(f"{layer_pred} not in adt_adata.layers")

    # ADT matrices
    X_true_adt = _to_dense(adt_adata.layers[layer_true])
    X_pred_adt = _to_dense(adt_adata.layers[layer_pred])
    adt_var_index = pd.Index(adt_adata.var_names)

    # RNA matrix (either layer or X)
    if rna_layer is None:
        X_rna = _to_dense(rna_adata.X)
        rna_label = "X"
    else:
        if rna_layer not in rna_adata.layers:
            raise ValueError(f"{rna_layer} not in rna_adata.layers")
        X_rna = _to_dense(rna_adata.layers[rna_layer])
        rna_label = rna_layer
    rna_var_index = pd.Index(rna_adata.var_names)

    # Ensure we have a UMAP on the ADT UniVI latent
    ensure_umap_on_univi(adt_adata)  # UMAP on adt_adata.obsm["X_univi"]

    for m in markers:
        if m not in adt_var_index:
            print(f"[UMAP true vs pred + RNA] Marker {m} not in ADT var_names; skipping.")
            continue

        # ADT indices and per-cell values
        j_adt = adt_var_index.get_loc(m)
        adt_adata.obs[f"{m}_adt_true"] = X_true_adt[:, j_adt]
        adt_adata.obs[f"{m}_adt_pred_from_rna"] = X_pred_adt[:, j_adt]

        colors = [f"{m}_adt_true", f"{m}_adt_pred_from_rna"]

        # --- RNA mapping for this marker ---
        # 1) explicit map if provided
        candidates = list(rna_gene_map.get(m, []))

        # 2) heuristic guesses
        if not candidates:
            candidates = _guess_rna_genes_for_marker(m)

        # keep only genes actually present in rna_adata
        present_genes = [g for g in candidates if g in rna_var_index]

        if present_genes:
            print(
                f"[UMAP true vs pred + RNA] {m}: "
                f"using RNA genes {present_genes} (layer={rna_label})."
            )
            for g in present_genes:
                j_rna = rna_var_index.get_loc(g)
                colname = f"{m}_rna_{rna_label}_{g}"
                adt_adata.obs[colname] = X_rna[:, j_rna]
                colors.append(colname)
        else:
            print(
                f"[UMAP true vs pred + RNA] {m}: "
                "no matching RNA genes found in var_names; "
                "plotting only ADT true + ADT pred_from_rna."
            )

        # UMAP panels: ADT true, ADT predicted, (optional) RNA gene panels
        sc.pl.umap(
            adt_adata,
            color=colors,
            size=size,
            alpha=0.8,
            show=False,
        )
        fname = f"umap_{title_prefix}_{m}.png".replace(" ", "_")
        plt.savefig(os.path.join(figdir, fname), bbox_inches="tight")
        plt.show()
        plt.close()


# --------------------------------------------------------------------
# Example mapping from ADT markers -> RNA gene symbols
# (feel free to tweak based on what actually exists in rna_test_adata.var_names)
# --------------------------------------------------------------------

# Example ADT markers to inspect
#adt_markers_to_plot = ["CD3", "CD4", "CD8a", "CD19", "CD14", "CD56", "HLA-DR"]

'''
plot_true_vs_pred_markers_on_umap_with_rna(
    adt_adata=adt_test_adata,
    rna_adata=rna_test_adata,
    markers=adt_markers_to_plot,
    rna_gene_map=rna_gene_map_for_adt_markers,
    layer_true="scaled",                 # whatever you used for ADT "true"
    layer_pred="univi_pred_from_rna",    # RNA→ADT predictions (decoded marker space)
    rna_layer=None,                      # use RNA log1p layer if you have it
    title_prefix="adt_true_vs_rna_pred_with_rna",
)
'''


# %% [code]
plot_true_vs_pred_markers_on_umap_with_rna(
    adt_adata=adt_test_adata,
    rna_adata=rna_test_adata,
    markers=["CD3", "CD4", "CD8", "CD19", "CD20", "CD14", "CD16", "CD56", "HLA-DR",
             "CD22", "CD74", "CD69", "CD83", "CD96", "CD11c"],
    rna_gene_map=rna_gene_map_for_adt_markers,
    layer_true="scaled",
    layer_pred="univi_pred_from_rna",
    rna_layer=None,  # or None if you prefer raw .X
    title_prefix="adt_true_vs_rna_pred_with_rna",
)


# %% [code]


# %% [code]
import numpy as np
import pandas as pd
import scanpy as sc
import matplotlib.pyplot as plt
import os

# ------------------------------------------------------
# Helper: heuristic mapping ATAC feature name -> RNA genes
# ------------------------------------------------------

def _guess_rna_genes_for_atac_feature(feat: str) -> list[str]:
    """
    Heuristic to propose RNA gene symbols from an ATAC feature name.

    Handles common patterns like:
      - "TNFRSF4_body", "IFNG_prom", "CD4_enh"
      - "TNFRSF4_body_peak123"
      - Already-gene-like names (e.g. "IFNG", "CD4")
    """
    cand = []

    # raw feature name
    cand.append(feat)

    # uppercase
    up = feat.upper()
    if up not in cand:
        cand.append(up)

    # split on common delimiters and keep first token as gene candidate
    for sep in ["|", ";"]:
        if sep in up:
            token = up.split(sep)[0]
            if token not in cand:
                cand.append(token)

    # patterns like "GENE_body", "GENE_prom", "GENE_enh", "GENE_peak123"
    if "_" in up:
        first_token = up.split("_")[0]
        if first_token not in cand:
            cand.append(first_token)

    # if it already looks like a gene (letters/numbers only)
    stripped = "".join(ch for ch in up if ch.isalnum())
    if stripped and stripped not in cand:
        cand.append(stripped)

    return cand


# ------------------------------------------------------
# RNA → ATAC: UMAPs of true vs RNA-predicted ATAC feature signal
#             + RNA expression of mapped genes
# ------------------------------------------------------

def plot_true_vs_pred_atac_on_umap_with_rna(
    atac_adata,
    rna_adata,
    atac_features,
    rna_gene_map: dict[str, list[str]] | None = None,
    layer_true: str = "scaled",              # ATAC "true" (e.g. normalized gene-body / peaks)
    layer_pred: str = "univi_pred_from_rna", # ATAC predicted from RNA (cross-modal)
    rna_layer: str | None = "log1p",         # RNA layer for expression; if None, use .X
    title_prefix: str = "atac_true_vs_rna_pred_with_rna",
    figdir: str = FIGDIR,
    size: int = 200,
):
    """
    For each ATAC feature in `atac_features`, plot UMAPs (on ATAC UniVI latent) colored by:
      - ATAC true signal (from `layer_true`)
      - ATAC predicted from RNA (from `layer_pred`)
      - RNA expression of corresponding gene(s), if present in RNA var_names

    RNA mapping:
      * Primary: rna_gene_map[feat] -> list of gene symbols
      * Fallback: heuristic guesses via _guess_rna_genes_for_atac_feature
    """
    if rna_gene_map is None:
        rna_gene_map = {}

    # Sanity: same cells, same order
    assert np.array_equal(
        atac_adata.obs_names, rna_adata.obs_names
    ), "ATAC and RNA obs_names must match 1:1 for per-cell comparisons."

    # Check ATAC layers
    if layer_true not in atac_adata.layers:
        raise ValueError(f"{layer_true} not in atac_adata.layers")
    if layer_pred not in atac_adata.layers:
        raise ValueError(f"{layer_pred} not in atac_adata.layers")

    # ATAC matrices
    X_true_atac = _to_dense(atac_adata.layers[layer_true])
    X_pred_atac = _to_dense(atac_adata.layers[layer_pred])
    atac_var_index = pd.Index(atac_adata.var_names)

    # RNA matrix (either layer or X)
    if rna_layer is None:
        X_rna = _to_dense(rna_adata.X)
        rna_label = "X"
    else:
        if rna_layer not in rna_adata.layers:
            raise ValueError(f"{rna_layer} not in rna_adata.layers")
        X_rna = _to_dense(rna_adata.layers[rna_layer])
        rna_label = rna_layer
    rna_var_index = pd.Index(rna_adata.var_names)

    # Ensure we have a UMAP on the ATAC UniVI latent
    # (this should use your frozen layout if you wired ensure_umap_on_univi that way)
    ensure_umap_on_univi(atac_adata)

    for feat in atac_features:
        if feat not in atac_var_index:
            print(f"[UMAP ATAC true vs pred + RNA] Feature {feat} not in ATAC var_names; skipping.")
            continue

        # ATAC indices and per-cell values
        j_atac = atac_var_index.get_loc(feat)
        atac_adata.obs[f"{feat}_atac_true"] = X_true_atac[:, j_atac]
        atac_adata.obs[f"{feat}_atac_pred_from_rna"] = X_pred_atac[:, j_atac]

        colors = [f"{feat}_atac_true", f"{feat}_atac_pred_from_rna"]

        # --- RNA mapping for this ATAC feature ---
        # 1) explicit map if provided
        candidates = list(rna_gene_map.get(feat, []))

        # 2) heuristic guesses
        if not candidates:
            candidates = _guess_rna_genes_for_atac_feature(feat)

        # keep only genes actually present in rna_adata
        present_genes = [g for g in candidates if g in rna_var_index]

        if present_genes:
            print(
                f"[UMAP ATAC true vs pred + RNA] {feat}: "
                f"using RNA genes {present_genes} (layer={rna_label})."
            )
            for g in present_genes:
                j_rna = rna_var_index.get_loc(g)
                colname = f"{feat}_rna_{rna_label}_{g}"
                atac_adata.obs[colname] = X_rna[:, j_rna]
                colors.append(colname)
        else:
            print(
                f"[UMAP ATAC true vs pred + RNA] {feat}: "
                "no matching RNA genes found in var_names; "
                "plotting only ATAC true + ATAC pred_from_rna."
            )

        # UMAP panels: ATAC true, ATAC predicted, (optional) RNA gene panels
        sc.pl.umap(
            atac_adata,
            color=colors,
            size=size,
            alpha=0.8,
            show=False,
        )
        fname = f"umap_{title_prefix}_{feat}.png".replace(" ", "_")
        plt.savefig(os.path.join(figdir, fname), bbox_inches="tight")
        plt.show()
        plt.close()


# %% [code]
atac_test_adata.layers["scaled"] = atac_test_adata.X.copy()


# %% [code]
print(atac_test_adata.var_names)


# %% [code]
# Example: choose some gene-body ATAC features to visualize
atac_features_to_plot = [
    "IFNG_body",
    "TNFRSF4_body",
    "CD4_body",
    # whatever matches your atac_test_adata.var_names
]

# Optional explicit mapping from ATAC features → RNA genes
rna_gene_map_for_atac_features = {
    "IFNG_body": ["IFNG"],
    "TNFRSF4_body": ["TNFRSF4"],
    "CD4_body": ["CD4"],
}

plot_true_vs_pred_atac_on_umap_with_rna(
    atac_adata=atac_test_adata,
    rna_adata=rna_test_adata,
    atac_features=atac_features_to_plot,
    rna_gene_map=rna_gene_map_for_atac_features,
    layer_true="scaled",                 # or whatever you used for true ATAC
    layer_pred="univi_pred_from_rna",    # RNA→ATAC predictions
    rna_layer=None,                      # or None to use rna_test_adata.X
    title_prefix="atac_true_vs_rna_pred_with_rna",
)


# %% [code]


# %% [markdown]
### Predict ADT and ATAC using RNA from the heldout TEA-seq set, then validate with actual

# %% [code]
rna_test = rna_test_adata.copy()
adt_test = adt_test_adata.copy()
atac_test = atac_test_adata.copy()


# %% [code]
print(rna_test)
print(adt_test)
print(atac_test)


# %% [code]
import numpy as np
import torch
from univi.evaluation import encode_adata

device = torch.device("cuda" if torch.cuda.is_available() else "cpu")
model_loaded.to(device)
model_loaded.eval()

@torch.no_grad()
def predict_from_rna(
    model,
    rna_adata,
    *,
    out_modality: str,
    rna_layer: str | None,
    batch_size: int = 1024,
    latent_kind: str = "modality_mean",
) -> np.ndarray:
    # 1) encode RNA into latent (np array)
    Z_rna = encode_adata(
        model,
        rna_adata,
        modality="rna",
        device=device,
        layer=rna_layer,
        latent=latent_kind,
        batch_size=batch_size,
    )

    # 2) decode latent -> all modalities, then pick out_modality
    preds = []
    for i in range(0, Z_rna.shape[0], batch_size):
        z_batch = torch.from_numpy(Z_rna[i:i + batch_size]).to(device)

        xhat_dict = model.decode_modalities(z_batch)  # <-- UniVI API :contentReference[oaicite:3]{index=3}
        x_hat = xhat_dict[out_modality]

        # some decoders return tuples/dicts; handle common cases safely
        if isinstance(x_hat, (tuple, list)):
            x_hat = x_hat[0]
        elif isinstance(x_hat, dict) and "mean" in x_hat:
            x_hat = x_hat["mean"]

        preds.append(x_hat.detach().cpu().numpy())

    return np.vstack(preds)


# %% [code]
import numpy as np
import pandas as pd
import torch
from scipy.stats import pearsonr, spearmanr

from univi.evaluation import cross_modal_predict

device = torch.device("cuda" if torch.cuda.is_available() else "cpu")
model_loaded.to(device)
model_loaded.eval()

RNA_LAYER  = None   # None => use .X
ADT_LAYER  = None
ATAC_LAYER = None

def _to_dense(X):
    import scipy.sparse as sp
    return X.toarray() if sp.issparse(X) else np.asarray(X)

def get_X(adata, layer):
    return adata.X if layer is None else adata.layers[layer]

# --- predictions (dense np arrays) ---
X_adt_hat_from_rna = cross_modal_predict(
    model_loaded,
    adata_src=rna_test,
    src_mod="rna",
    tgt_mod="adt",
    device=device,
    layer=RNA_LAYER,
    batch_size=1024,
    use_moe=True,  # for src-only input this just reduces to the src posterior
)

X_atac_hat_from_rna = cross_modal_predict(
    model_loaded,
    adata_src=rna_test,
    src_mod="rna",
    tgt_mod="atac",
    device=device,
    layer=RNA_LAYER,
    batch_size=1024,
    use_moe=True,
)

# --- observed matrices in the evaluation layers ---
X_adt_obs  = _to_dense(get_X(adt_test,  ADT_LAYER))
X_atac_obs = _to_dense(get_X(atac_test, ATAC_LAYER))

print("Pred shapes:")
print("  ADT  obs / hat:", X_adt_obs.shape,  X_adt_hat_from_rna.shape)
print("  ATAC obs / hat:", X_atac_obs.shape, X_atac_hat_from_rna.shape)


# %% [code]
from scipy.stats import pearsonr, spearmanr

def feature_perf_df(X_obs: np.ndarray,
                    X_hat: np.ndarray,
                    feature_names,
                    target: str):
    rows = []
    for j, name in enumerate(feature_names):
        y_true = X_obs[:, j]
        y_pred = X_hat[:, j]

        # skip completely constant features
        if np.allclose(y_true, y_true[0]) or np.allclose(y_pred, y_pred[0]):
            continue

        r, _   = pearsonr(y_true, y_pred)
        rho, _ = spearmanr(y_true, y_pred)
        rows.append(
            {"feature": name, "pearson_r": r, "spearman_r": rho, "target": target}
        )
    return pd.DataFrame(rows)


adt_feature_names  = adt_test.var_names.to_list()
atac_feature_names = atac_test.var_names.to_list()

adt_perf  = feature_perf_df(X_adt_obs,  X_adt_hat_from_rna,  adt_feature_names,  target="adt")
atac_perf = feature_perf_df(X_atac_obs, X_atac_hat_from_rna, atac_feature_names, target="atac")

print("ADT prediction – mean Pearson r:",  adt_perf["pearson_r"].mean())
print("ATAC prediction – mean Pearson r:", atac_perf["pearson_r"].mean())


# %% [code]
def summarize_perf(df, top_n=10, label="adt"):
    d = df.sort_values("pearson_r", ascending=False)
    print(f"\nTop {top_n} {label} features by Pearson r:")
    print(d.head(top_n)[["feature", "pearson_r", "spearman_r"]])

summarize_perf(adt_perf,  top_n=50, label="adt")
summarize_perf(atac_perf, top_n=50, label="atac")


# %% [code]
import matplotlib.pyplot as plt

def plot_top_features(
    df,
    top_n: int = 15,
    metric: str = "pearson_r",
    label: str = "ADT",
    figsize=(7, 0.4 * 15 + 1.5),
):
    """
    Plot top-N features by a given metric (e.g. Pearson r).

    df must have columns: 'feature', metric, and optionally 'spearman_r'.
    """
    d = df.sort_values(metric, ascending=False).head(top_n).copy()

    # dynamic figure height based on number of features
    fig_height = 0.4 * len(d) + 1.5
    fig, ax = plt.subplots(figsize=(figsize[0], fig_height))

    ax.barh(d["feature"], d[metric])
    ax.invert_yaxis()  # highest on top
    ax.set_xlabel(metric)
    ax.set_title(f"Top {len(d)} {label} features by {metric}", pad=8)
    ax.grid(False)

    # If we have Spearman, print it as text on the right
    if "spearman_r" in d.columns:
        for i, (y, r_spear) in enumerate(zip(d["feature"], d["spearman_r"])):
            ax.text(
                d[metric].iloc[i],
                i,
                f"  ρ={r_spear:.2f}",
                va="center",
                ha="left",
            )

    plt.tight_layout()
    plt.show()


# Usage
plot_top_features(adt_perf,  top_n=50, metric="pearson_r", label="adt")
plot_top_features(atac_perf, top_n=50, metric="pearson_r", label="atac")


# %% [code]
plot_top_features(rna_perf, top_n=50, metric="pearson_r", label="rna")


# %% [code]


# %% [code]


# %% [code]


# %% [code]


# %% [code]


# %% [code]


# %% [code]


# %% [code]


# %% [code]


# %% [code]
````
