## Supplemental Code for "Unifying multimodal single-cell data with a mixture-of-experts *β*-variational autoencoder framework": Fig07_AML_bridge_finetuning.ipynb

````
# %% [markdown]
## UniVI manuscript - Figure 7 generation reproducible workflow
### UniVI AML bridging experiment (CITE-seq ↔ scRNA ↔ DAb-seq)

Andrew Ashford, Pathways + Omics Group, Oregon Health & Science University, Portland, OR - 1/6/2026

This notebook builds a shared UniVI latent space using:
- Knorr et al. AML CITE-seq (RNA+ADT; paired)

Then does a zero-shot application of the bridge-trained model on:
- van Galen AML scRNA (RNA only)
- Demaree DAb-seq (protein + genotype calls; no RNA)

Key goals:
1) Train UniVI on paired CITE RNA+ADT.
2) Project van Galen RNA into the same latent via RNA encoder.
3) Project DAb proteins into the same latent via ADT encoder (after panel harmonization).
4) Validate with marker biology + mixing metrics.
5) Treat mutations properly (no leakage; patient/timepoint group splits).
6) Fine-tune model encoders using a mutation classification decoder and freeze decoders.
7) Analyze before and after fine-tuning results for Figure 7 of the Genome Research manuscript.

GitHub for the project can be found at: https://github.com/Ashford-A/UniVI

Package is pip installable via the command: 
```bash
pip install univi
```


# %% [markdown]
### Imports + global config

# %% [code]
import os
import re
import random
from pathlib import Path

import numpy as np
import pandas as pd
import scipy.sparse as sp

import scanpy as sc
import anndata as ad

import matplotlib.pyplot as plt
from matplotlib.patches import FancyBboxPatch, FancyArrowPatch

from sklearn.model_selection import GroupShuffleSplit
from sklearn.neighbors import NearestNeighbors
from sklearn.metrics import roc_auc_score, average_precision_score, roc_curve, precision_recall_curve
from sklearn.preprocessing import StandardScaler

import torch
import torch.nn as nn
import torch.nn.functional as F
from torch.utils.data import Dataset, DataLoader, Subset

# UniVI imports
import univi as uv
from univi import (
    ModalityConfig, UniVIConfig, TrainingConfig,
    UniVIMultiModalVAE, UniVITrainer,
    MultiModalDataset,
)
import univi.evaluation as ue

def seed_everything(seed: int = 1):
    random.seed(seed)
    np.random.seed(seed)
    torch.manual_seed(seed)
    torch.cuda.manual_seed_all(seed)
    torch.backends.cudnn.deterministic = False
    torch.backends.cudnn.benchmark = True

seed_everything(1)

device = "cuda" if torch.cuda.is_available() else "cpu"
print("univi:", uv.__version__)
print("torch:", torch.__version__)
print("device:", device)

sc.set_figure_params(figsize=(6.5, 5.5), dpi=120, dpi_save=400, fontsize=11, frameon=False)
plt.rcParams.update({"savefig.bbox": "tight", "savefig.pad_inches": 0.1})


# %% [markdown]
### Paths + key obs columns

# %% [code]
FIGDIR = Path("./results/fig7_aml_mosaic_reproducibility")
FIGDIR.mkdir(parents=True, exist_ok=True)

# -----------------------------
# INPUT PATHS (EDIT)
# -----------------------------
CITE_ROOT = Path("/home/groups/precepts/ashforda/UniVI_v2/UniVI_older-non_git/data/Knorr_AML_CITE-seq_data/GSE220473_RAW") # wide csv.gz counts + assignments
VG_PATH   = Path("/home/groups/precepts/ashforda/scOPE_github_stuff/data/testing/vanGalen_all_h5ad/vanGalen_anndata.h5ad")
DAB_ROOT  = Path("/home/groups/precepts/ashforda/UniVI_v2/UniVI_older-non_git/data/Demaree_DAb-seq_data")

print("CITE_ROOT:", CITE_ROOT)
print("VG_PATH:", VG_PATH)
print("DAB_ROOT:", DAB_ROOT)

# -----------------------------
# CITE split grouping (avoid leakage)
# -----------------------------
# This will be created from *_cell_sample_assign.csv.gz (see attach_assignments below)
CITE_GROUP_COL = "sample_id"

# DAb-seq LOEO grouping column
DAB_EXPT_COL   = "experiment"   # edit if your DAb obs uses another name

# van Galen grouping column (patient/case) for mutation-head splits
VG_GROUP_COL   = "patient"      # edit if needed


# %% [markdown]
### CITE readers (wide csv.gz → AnnData) + assignment attach

# %% [code]
def _cite_id_from_name(name: str) -> str:
    m = re.search(r"_CITE(\d+)_", name)
    if m is None:
        raise ValueError(f"Could not parse CITE id from filename: {name}")
    return f"CITE{int(m.group(1))}"

def _sort_cite_ids(cite_ids):
    return sorted(cite_ids, key=lambda x: int(x.replace("CITE", "")))

def read_wide_counts_csv_gz_to_adata(
    path: Path,
    chunksize_rows: int = 256,
    dtype=np.int32,
) -> ad.AnnData:
    """
    Expects a wide matrix CSV with:
      - rows = features (genes/proteins) in index column
      - columns = cell barcodes
    Produces AnnData with:
      - obs = cells
      - var = features
      - X   = sparse CSR (cells x features)
    """
    it = pd.read_csv(
        path,
        compression="gzip",
        index_col=0,
        chunksize=int(chunksize_rows),
    )

    obs_names = None
    var_names = []
    blocks = []

    for chunk in it:
        # columns are cells
        cols = chunk.columns.astype(str).tolist()
        if obs_names is None:
            obs_names = cols
        else:
            if cols != obs_names:
                raise ValueError(f"Column mismatch across chunks in {path.name}")

        # index is features
        var_names.extend(chunk.index.astype(str).tolist())

        Xc = chunk.to_numpy(dtype=dtype, copy=False)  # (n_feat_chunk, n_cells)
        Xc = sp.csr_matrix(Xc.T)                      # (n_cells, n_feat_chunk)
        blocks.append(Xc)

    if obs_names is None:
        raise ValueError(f"No data read from {path}")

    X = sp.hstack(blocks, format="csr") if len(blocks) > 1 else blocks[0].tocsr()
    adata = ad.AnnData(
        X=X,
        obs=pd.DataFrame(index=pd.Index(obs_names, name="cell")),
        var=pd.DataFrame(index=pd.Index(var_names, name="feature")),
    )
    return adata

def read_cell_sample_assign(path: Path) -> pd.DataFrame:
    """
    Reads *_cell_sample_assign.csv.gz and returns a DF indexed by barcode/cell.
    Tries hard to find the barcode column.
    """
    df = pd.read_csv(path, compression="gzip")
    # choose barcode column
    barcode_col = None
    for c in df.columns:
        cl = str(c).lower()
        if "barcode" in cl or "cell" in cl:
            barcode_col = c
            break
    if barcode_col is None:
        barcode_col = df.columns[0]
    df[barcode_col] = df[barcode_col].astype(str)
    df = df.set_index(barcode_col)
    return df

def attach_assignments(adata: ad.AnnData, assign_df: pd.DataFrame) -> None:
    """
    Joins assignment columns into .obs and creates a stable 'sample_id' column.
    """
    pref = assign_df.add_prefix("assign_")
    adata.obs = adata.obs.join(pref, how="left")

    cand = None
    for c in assign_df.columns:
        cl = str(c).lower()
        if "sample" in cl or "donor" in cl or "patient" in cl:
            cand = c
            break

    if cand is None:
        adata.obs["sample_id"] = "NA"
    else:
        adata.obs["sample_id"] = assign_df[cand].astype(str)


# %% [markdown]
### Load all paired CITE libraries (RNA+ADT)

# %% [code]
rna_files = list(CITE_ROOT.glob("*_RNA_counts.csv.gz"))
adt_files = list(CITE_ROOT.glob("*_ADT_counts.csv.gz"))
asn_files = list(CITE_ROOT.glob("*_cell_sample_assign.csv.gz"))

rna_map = {_cite_id_from_name(p.name): p for p in rna_files}
adt_map = {_cite_id_from_name(p.name): p for p in adt_files}
asn_map = {_cite_id_from_name(p.name): p for p in asn_files}

cite_ids = _sort_cite_ids(set(rna_map) & set(adt_map) & set(asn_map))
print("Found paired libraries:", cite_ids)

rna_parts, adt_parts = [], []
for cid in cite_ids:
    print(f"\n--- Reading {cid} ---")
    r = read_wide_counts_csv_gz_to_adata(rna_map[cid], chunksize_rows=256, dtype=np.int32)
    a = read_wide_counts_csv_gz_to_adata(adt_map[cid], chunksize_rows=256, dtype=np.int32)

    assign = read_cell_sample_assign(asn_map[cid])
    attach_assignments(r, assign)
    attach_assignments(a, assign)

    r.obs["library_id"] = cid
    a.obs["library_id"] = cid

    common = r.obs_names.intersection(a.obs_names)
    r = r[common].copy()
    a = a[common].copy()
    a = a[r.obs_names].copy()
    assert (r.obs_names == a.obs_names).all()

    rna_parts.append(r)
    adt_parts.append(a)

cite_rna = ad.concat(rna_parts, join="outer", label="library_id", keys=cite_ids, index_unique="-")
cite_adt = ad.concat(adt_parts, join="outer", label="library_id", keys=cite_ids, index_unique="-")

common_cells = cite_rna.obs_names.intersection(cite_adt.obs_names)
cite_rna = cite_rna[common_cells].copy()
cite_adt = cite_adt[common_cells].copy()
cite_adt = cite_adt[cite_rna.obs_names].copy()
assert (cite_rna.obs_names == cite_adt.obs_names).all()

print("CITE RNA:", cite_rna)
print("CITE ADT:", cite_adt)
print("CITE sample_id nunique:", cite_rna.obs[CITE_GROUP_COL].nunique())


# %% [markdown]
### Load van Galen + DAb-seq

# %% [code]
vg_rna = sc.read_h5ad(str(VG_PATH))
vg_rna.var_names_make_unique()
vg_rna.obs_names_make_unique()
print("VG:", vg_rna)

dab_files = {
    "fig2_pbmcs_ab_geno":   "fig2_pbmcs_ab_geno.h5ad",
    "fig3_ab_geno":         "fig3_ab_geno.h5ad",
    "fig4_ab_geno":         "fig4_ab_geno.h5ad",
    "fig5_ab_geno":         "fig5_ab_geno.h5ad",
}

dab_parts = {}
for k, fn in dab_files.items():
    fp = DAB_ROOT / fn
    a = sc.read_h5ad(str(fp))
    a.obs["dab_source"] = k
    dab_parts[k] = a
    print("\n---", k, "---")
    print(a)

mergeable = ["fig3_ab_geno", "fig5_ab_geno"]
dab_adt = ad.concat(
    [dab_parts[k] for k in mergeable],
    join="outer",
    axis=0,
    label="dab_source",
    keys=mergeable,
    merge="unique",
    fill_value=0,
)
print("\nMerged DAb:", dab_adt)
print("DAb obs cols:", len(dab_adt.obs.columns))


# %% [markdown]
### Canonicalize ADT feature names + build matched subsets

# %% [code]
print(cite_adt.X.min())
print(cite_adt.X.max())
print(dab_adt.X.min())
print(dab_adt.X.max())


# %% [code]
print(cite_adt)
print(cite_adt.X.min())
print(cite_adt.X.max())
print(dab_adt)
print(dab_adt.X.min())
print(dab_adt.X.max())


# %% [code]
cite_adt.layers['counts'] = cite_adt.X


# %% [code]
print(cite_rna)
print(cite_rna.X.min())
print(cite_rna.X.max())
print(vg_rna)
print(vg_rna.X.min())
print(vg_rna.X.max())
print(vg_rna.raw)
print(vg_rna.raw.X.min())
print(vg_rna.raw.X.max())


# %% [code]
vg_rna.layers['counts'] = vg_rna.raw.X


# %% [code]
import re
import numpy as np
import pandas as pd
import scipy.sparse as sp
import anndata as ad

# The DAb panel you showed
DAB_PANEL = pd.Index([
    "CD10","CD117","CD11B","CD123","CD13","CD14","CD15","CD16","CD19","CD22",
    "CD3","CD30","CD33","CD34","CD38","CD4","CD45","CD5","CD56","CD64","CD7",
    "CD71","HLA-DR","IGG1"
], name="dab_panel")

# Mapping from CITE gene-symbol ADT names to DAb-style canonical targets
CITE_GENE_TO_CANON = {
    # present in your CITE set
    "ANPEP": "CD13",
    "CD14": "CD14",
    "FCGR3A": "CD16",
    "CD19": "CD19",
    "CD22": "CD22",
    "CD3D": "CD3",
    "CD3E": "CD3",
    "CD3G": "CD3",
    "CD33": "CD33",
    "CD38": "CD38",
    "CD4": "CD4",
    "CD5": "CD5",
    "CD7": "CD7",
    "PTPRC": "CD45",
    "NCAM1": "CD56",
    "FCGR1A": "CD64",
    "FCGR1B": "CD64",
    "TFRC": "CD71",
    "IL3RA": "CD123",
    "ITGAM": "CD11B",
    "HLA-DRA": "HLA-DR",
    "HLA-DRB1": "HLA-DR",
    "HLA-DRB5": "HLA-DR",

    # include these for completeness if they ever appear
    "MME": "CD10",
    "KIT": "CD117",
    "FUT4": "CD15",
    "TNFRSF8": "CD30",
    "CD34": "CD34",
    "IGHG1": "IGG1",
}

# Normalize DAb-side labels too
DAB_ALIASES = {
    "CD11B": "CD11B",
    "CD11b": "CD11B",
    "HLADR": "HLA-DR",
    "HLA DR": "HLA-DR",
    "HLA-DR": "HLA-DR",
    "IGG1": "IGG1",
    "IgG1": "IGG1",
}

def strip_rep_suffix(x: str) -> str:
    # remove trailing ".1", ".2", ".10", etc
    return re.sub(r"\.\d+$", "", str(x).strip())

def canon_from_varname(varname: str, *, source: str) -> str:
    """
    source: 'cite' or 'dab'
    Returns canonical marker name (upper-case) or '' if unusable.
    """
    s = strip_rep_suffix(varname)
    if s == "" or s.upper() in {"NAN", "NONE"}:
        return ""

    # Handle CITE isotypes explicitly
    if source == "cite":
        # your set contains "isotype.0 ... isotype.9" and also weird ".10"
        if s.lower().startswith("isotype") or s in {".10"}:
            return "ISOTYPE"

    s_up = s.upper()

    # Unify punctuation variants
    s_up = s_up.replace("_", "-")
    s_up = re.sub(r"\s+", " ", s_up).strip()

    if source == "dab":
        return DAB_ALIASES.get(s_up, s_up)

    # source == 'cite'
    # FIRST: map gene-symbol style markers (including CD3D/E/G -> CD3, PTPRC->CD45, etc.)
    if s_up in CITE_GENE_TO_CANON:
        return CITE_GENE_TO_CANON[s_up]

    # THEN: keep already-CD markers (e.g. CD14, CD19, CD33)
    if re.match(r"^CD\d+$", s_up) or re.match(r"^CD\d+[A-Z]+$", s_up):
        return s_up

    # Otherwise keep gene symbol as-is (won't match DAb panel, but useful for debugging)
    return s_up

def feature_variance(adata: ad.AnnData) -> np.ndarray:
    X = adata.X
    if sp.issparse(X):
        mean = np.asarray(X.mean(axis=0)).ravel()
        mean2 = np.asarray(X.multiply(X).mean(axis=0)).ravel()
        var = mean2 - mean**2
    else:
        var = np.var(np.asarray(X), axis=0)
    return np.asarray(var, dtype=np.float64)

def align_cite_dab_adts(cite_adt: ad.AnnData, dab_adt: ad.AnnData, panel: pd.Index = DAB_PANEL):
    cite = cite_adt.copy()
    dab  = dab_adt.copy()

    cite.var["canon"] = [canon_from_varname(v, source="cite") for v in cite.var_names]
    dab.var["canon"]  = [canon_from_varname(v, source="dab")  for v in dab.var_names]

    # Only consider DAb markers that are in the explicit DAb panel list (canonical)
    dab_can = pd.Index(dab.var["canon"].unique())
    dab_targets = pd.Index([x for x in panel if x in set(dab_can)])

    # Now the potential overlap is those DAb targets that also exist in CITE canonical names
    cite_can = set(cite.var["canon"].values)
    shared = pd.Index([x for x in dab_targets if x in cite_can])

    # Drop controls unless you explicitly want them
    # (ISOTYPE is not the same as IGG1)
    shared = pd.Index([x for x in shared if x not in {"ISOTYPE"}])

    print(f"DAb targets present in DAb object: {len(dab_targets)} / {len(panel)}")
    print(f"Shared canonical markers (after mapping): {len(shared)}")
    print("Shared:", list(shared))

    if len(shared) == 0:
        return None, None, shared

    # Resolve duplicates on CITE side: pick most variable feature for each canonical marker
    var = feature_variance(cite)
    cite.var["_var"] = var

    cite_keep = []
    for m in shared:
        idx = np.where(cite.var["canon"].values == m)[0]
        if len(idx) == 1:
            cite_keep.append(idx[0])
        else:
            cite_keep.append(idx[np.argmax(cite.var["_var"].values[idx])])

    cite_keep = np.array(cite_keep, dtype=int)

    # For DAb side: pick the first match per marker (usually unique already)
    dab_keep = []
    for m in shared:
        idx = np.where(dab.var["canon"].values == m)[0]
        dab_keep.append(idx[0])

    cite_al = cite[:, cite_keep].copy()
    dab_al  = dab[:,  dab_keep ].copy()

    # Set var_names to canonical marker names in a consistent order (shared order)
    cite_al.var_names = shared
    dab_al.var_names  = shared

    return cite_al, dab_al, shared

#cite_adt_al, dab_adt_al, shared_adts = align_cite_dab_adts(cite_adt, dab_adt, panel=DAB_PANEL)
#
#print("Aligned shapes:",
#      None if cite_adt_al is None else cite_adt_al.shape,
#      None if dab_adt_al is None else dab_adt_al.shape)


# %% [markdown]
### Feature alignment (ADT shared; VG genes intersect to CITE genes)

# %% [code]
def intersect_vars(a: ad.AnnData, b: ad.AnnData) -> tuple[ad.AnnData, ad.AnnData]:
    shared = a.var_names.intersection(b.var_names)
    a2 = a[:, shared].copy()
    b2 = b[:, shared].copy()
    return a2, b2

# ----------------------------
# ADT: use canonical alignment (NOT intersect_vars)
# ----------------------------
cite_adt_al, dab_adt_al, shared_adts = align_cite_dab_adts(cite_adt, dab_adt, panel=DAB_PANEL)
print("Shared ADTs after canonicalization:", len(shared_adts))

# ----------------------------
# RNA: direct intersection is fine
# ----------------------------
vg_rna_al, cite_rna_al = intersect_vars(vg_rna, cite_rna)
print("VG->CITE shared genes:", vg_rna_al.n_vars)


# %% [code]
print(cite_rna_al)
print(cite_rna_al.X.min())
print(cite_rna_al.X.max())


# %% [code]
cite_rna_al.layers['counts'] = cite_rna_al.X


# %% [code]
if cite_adt_al is not None:
    cite_can = set([canon_from_varname(v, source="cite") for v in cite_adt.var_names])
    dab_panel_can = set([canon_from_varname(v, source="dab") for v in DAB_PANEL])

    missing_from_cite = sorted(list(dab_panel_can - cite_can))
    print("DAb panel markers not present in CITE ADT panel (canonical):")
    print(missing_from_cite)


# %% [markdown]
### Make splits

# %% [code]
import numpy as np
import pandas as pd
import anndata as ad
from sklearn.model_selection import GroupShuffleSplit

def group_holdout_splits(
    adata: ad.AnnData,
    group_col: str,
    *,
    seed: int = 0,
    train_frac: float = 0.80,
    val_frac: float = 0.10,
):
    """
    Group-disjoint train/val/test split.
    Ensures no group appears in more than one split.

    Returns dict with:
      - "train_idx", "val_idx", "test_idx" (integer indices)
      - "train_obs", "val_obs", "test_obs" (obs_names arrays)
      - "train_groups", "val_groups", "test_groups"
    """
    if group_col not in adata.obs.columns:
        raise KeyError(f"group_col='{group_col}' not in adata.obs columns")

    groups = adata.obs[group_col].astype(str).values
    idx_all = np.arange(adata.n_obs)

    # 1) train vs temp
    gss1 = GroupShuffleSplit(n_splits=1, train_size=train_frac, random_state=seed)
    tr_idx, tmp_idx = next(gss1.split(idx_all, groups=groups))

    # 2) val vs test within temp
    tmp_groups = groups[tmp_idx]
    # val_frac of total => val_frac/(1-train_frac) of tmp
    val_frac_of_tmp = val_frac / max(1e-12, (1.0 - train_frac))
    gss2 = GroupShuffleSplit(n_splits=1, train_size=val_frac_of_tmp, random_state=seed + 1)
    va_rel, te_rel = next(gss2.split(tmp_idx, groups=tmp_groups))
    va_idx = tmp_idx[va_rel]
    te_idx = tmp_idx[te_rel]

    out = {
        "train_idx": tr_idx,
        "val_idx": va_idx,
        "test_idx": te_idx,
        "train_obs": adata.obs_names[tr_idx].to_numpy(),
        "val_obs": adata.obs_names[va_idx].to_numpy(),
        "test_obs": adata.obs_names[te_idx].to_numpy(),
        "train_groups": pd.unique(groups[tr_idx]),
        "val_groups": pd.unique(groups[va_idx]),
        "test_groups": pd.unique(groups[te_idx]),
    }

    # Sanity: ensure disjoint groups
    assert set(out["train_groups"]).isdisjoint(set(out["val_groups"]))
    assert set(out["train_groups"]).isdisjoint(set(out["test_groups"]))
    assert set(out["val_groups"]).isdisjoint(set(out["test_groups"]))

    return out


def subset_by_obs_names(adata: ad.AnnData, obs_names) -> ad.AnnData:
    return adata[obs_names].copy()


# -----------------------------
# CITE splits (NO LEAKAGE)
# -----------------------------
# IMPORTANT: define splits on a single reference object (CITE RNA aligned genes)
# and apply same obs split to CITE ADT aligned panel.
assert np.array_equal(cite_rna_al.obs_names, cite_adt_al.obs_names), "CITE RNA/ADT obs_names must match!"

splits = group_holdout_splits(
    cite_rna_al,
    group_col=CITE_GROUP_COL,   # "sample_id"
    seed=0,
    train_frac=0.80,
    val_frac=0.10,
)

cite_rna_tr = subset_by_obs_names(cite_rna_al, splits["train_obs"])
cite_rna_va = subset_by_obs_names(cite_rna_al, splits["val_obs"])
cite_rna_te = subset_by_obs_names(cite_rna_al, splits["test_obs"])

cite_adt_tr = subset_by_obs_names(cite_adt_al, splits["train_obs"])
cite_adt_va = subset_by_obs_names(cite_adt_al, splits["val_obs"])
cite_adt_te = subset_by_obs_names(cite_adt_al, splits["test_obs"])

print("CITE split sizes:", cite_rna_tr.n_obs, cite_rna_va.n_obs, cite_rna_te.n_obs)
print("CITE groups (train/val/test):",
      len(splits["train_groups"]), len(splits["val_groups"]), len(splits["test_groups"]))


# %% [markdown]
### Minimal preprocessing (RNA + ADT)

# %% [code]
# Split out the cell lines from the vg scRNA data prior to preprocessing

# prefix = string before first underscore
vg_rna_al.obs["prefix"] = vg_rna_al.obs_names.str.split("_", n=1).str[0]

# quick sanity
print(vg_rna_al.obs["prefix"].value_counts().head(20))


# %% [code]
pref = vg_rna_al.obs_names.to_series().str.split("_", n=1).str[0]

mask_cell_lines = pref.str.startswith("OCI.AML") | pref.str.startswith("MUTZ3")

vg_rna_al = vg_rna_al[~mask_cell_lines].copy()


# %% [code]
print(vg_rna_al)


# %% [code]
import numpy as np
import scipy.sparse as sp
import scanpy as sc
import anndata as ad

# -------------------------
# utils
# -------------------------
def _to_dense(X):
    return X.toarray() if sp.issparse(X) else np.asarray(X)

def _cast32(X):
    return X.astype(np.float32, copy=False)

def ensure_layer_from_X(adata: ad.AnnData, layer: str):
    """If layer missing, snapshot current .X into that layer."""
    if layer not in adata.layers:
        adata.layers[layer] = adata.X.copy()

def has_negative_X(adata: ad.AnnData) -> bool:
    X = _to_dense(adata.X)
    return np.nanmin(X) < 0


# ============================================================
# RNA: keep .layers['counts'] and .layers['log1p'] everywhere
# ============================================================
def _ensure_rna_counts_and_log1p_layers(
    a: ad.AnnData,
    *,
    counts_layer: str = "counts",
    log1p_layer: str = "log1p",
    target_sum: float = 1e4,
    overwrite_log1p: bool = False,
) -> ad.AnnData:
    """
    Ensures:
      - .layers[counts_layer] exists (raw snapshot)
      - .layers[log1p_layer] exists = log1p(normalize_total(counts))
    Does NOT modify counts layer.
    """
    x = a.copy()

    # ensure counts layer exists (snapshot current X if user didn't provide one)
    ensure_layer_from_X(x, counts_layer)

    # build log1p layer if missing (or overwrite requested)
    if (log1p_layer not in x.layers) or overwrite_log1p:
        tmp = x.copy()
        tmp.X = tmp.layers[counts_layer].copy()
        sc.pp.normalize_total(tmp, target_sum=target_sum)
        sc.pp.log1p(tmp)  # may set tmp.uns['log1p'] = {'base': None}; harmless
        x.layers[log1p_layer] = tmp.X.copy()

    # lightweight provenance
    x.uns["rna_layers"] = {
        "counts_layer": counts_layer,
        "log1p_layer": log1p_layer,
        "target_sum": float(target_sum),
        "transform": "normalize_total+log1p",
    }
    return x


# ============================================================
# RNA: fit/apply (fit on TRAIN using .layers['log1p'])
# ============================================================
def rna_fit_params(
    a_train: ad.AnnData,
    *,
    counts_layer: str = "counts",
    target_sum: float = 1e4,
    clip: float = 10.0,
):
    """
    Fit gene-wise mean/sd for Z-scoring on TRAIN ONLY using:
      .layers['counts'] -> normalize_total(target_sum) -> log1p -> store in .layers['log1p']
    Then compute mu/sd from .layers['log1p'].
    """
    x = _ensure_rna_counts_and_log1p_layers(
        a_train,
        counts_layer=counts_layer,
        log1p_layer="log1p",
        target_sum=target_sum,
        overwrite_log1p=False,
    )

    X = _to_dense(x.layers["log1p"]).astype(np.float32)
    mu = X.mean(axis=0).astype(np.float32)
    sd = (X.std(axis=0).astype(np.float32) + 1e-8)

    return {
        "target_sum": float(target_sum),
        "clip": float(clip),
        "mu": mu,
        "sd": sd,
        "transform": "counts->normalize_total+log1p (layer), then zscore(.X)",
        "fit_on": "train_only",
        "counts_layer": counts_layer,
        "log1p_layer": "log1p",
    }

def preprocess_rna_apply(
    a: ad.AnnData,
    params: dict,
    *,
    counts_layer: str = "counts",
) -> ad.AnnData:
    """
    Ensures layers:
      - .layers['counts'] (raw)
      - .layers['log1p'] = log1p(normalize_total(counts))
    Sets:
      - .X = zscore(.layers['log1p'], mu/sd) clipped (float32)
    """
    x = _ensure_rna_counts_and_log1p_layers(
        a,
        counts_layer=counts_layer,
        log1p_layer=params.get("log1p_layer", "log1p"),
        target_sum=params["target_sum"],
        overwrite_log1p=False,
    )

    X = _to_dense(x.layers[params.get("log1p_layer", "log1p")]).astype(np.float32)
    X = (X - params["mu"]) / params["sd"]
    X = np.clip(X, -params["clip"], params["clip"]).astype(np.float32)

    x.X = X
    x.uns["rna_transform"] = {k: v for k, v in params.items() if k not in ("mu", "sd")}
    return x


# ============================================================
# ADT: fit/apply for CITE counts, plus a safe path for DAb .X
# ============================================================
def adt_fit_params(
    a_train: ad.AnnData,
    *,
    counts_layer: str = "counts",
):
    """
    Fit ADT mu/sd on CITE TRAIN ONLY:
      counts -> log1p -> per-feature mean/sd
    """
    if counts_layer not in a_train.layers:
        raise ValueError(f"ADT fit expects counts in layers['{counts_layer}'].")

    X = _to_dense(a_train.layers[counts_layer]).astype(np.float32)
    X = np.log1p(X)
    mu = X.mean(axis=0).astype(np.float32)
    sd = (X.std(axis=0).astype(np.float32) + 1e-8)

    return {
        "mu": mu,
        "sd": sd,
        "transform": "log1p+zscore",
        "fit_on": "train_only",
        "counts_layer": counts_layer,
    }

def preprocess_adt_apply_counts(
    a: ad.AnnData,
    params: dict,
    *,
    counts_layer: str = "counts",
    clip: float = 10.0,
) -> ad.AnnData:
    """
    Apply ADT preprocessing when true counts are available:
      counts -> log1p -> zscore(mu/sd) -> clip
    """
    x = a.copy()
    if counts_layer not in x.layers:
        raise ValueError(f"ADT apply_counts expects counts in layers['{counts_layer}'].")

    X = _to_dense(x.layers[counts_layer]).astype(np.float32)
    X = np.log1p(X)
    X = (X - params["mu"]) / params["sd"]
    X = np.clip(X, -clip, clip).astype(np.float32)

    x.X = X
    x.uns["adt_transform"] = {
        "type": "log1p+zscore",
        "fit": params.get("fit_on", "unknown"),
        "clip": float(clip),
    }
    return x

def preprocess_adt_from_processed_X(
    a: ad.AnnData,
    *,
    clip: float = 10.0,
    standardize_to_ref: dict | None = None,
) -> ad.AnnData:
    """
    Use this for DAb when only processed values exist in .X.

    Behavior:
      - Always treats .X as already-processed continuous features.
      - Optionally unit-standardizes feature-wise, then clips.

    If standardize_to_ref is provided, it should be a dict with:
      {"mode": "unit"}
    """
    x = a.copy()
    X = _to_dense(x.X).astype(np.float32)

    if standardize_to_ref is None:
        X = np.clip(X, -clip, clip).astype(np.float32)
        x.X = X
        x.uns["adt_transform"] = {"type": "already_processed", "standardize": "none", "clip": float(clip)}
        return x

    mode = standardize_to_ref.get("mode", "unit")
    if mode != "unit":
        raise ValueError(f"Unknown standardize mode: {mode}. Use mode='unit' or None.")

    mu = np.nanmean(X, axis=0).astype(np.float32)
    sd = (np.nanstd(X, axis=0).astype(np.float32) + 1e-8)
    X = (X - mu) / sd
    X = np.clip(X, -clip, clip).astype(np.float32)

    x.X = X
    x.uns["adt_transform"] = {"type": "already_processed", "standardize": "unit", "clip": float(clip)}
    x.uns["adt_standardize_fit"] = {"mu": "dab_self", "sd": "dab_self"}  # keep lightweight; don't stash big arrays
    return x


# ============================================================
# One-stop helpers for workflow
# ============================================================
def preprocess_all_for_aml_bridge(
    *,
    cite_rna_tr, cite_rna_va, cite_rna_te,
    vg_rna_al,
    cite_adt_tr, cite_adt_va, cite_adt_te,
    dab_adt_al,
    rna_counts_layer="counts",
    adt_counts_layer="counts",
    rna_target_sum=1e4,
    rna_clip=10.0,
    adt_clip=10.0,
    dab_standardize_unit=True,
):
    """
    RNA:
      - Ensures every RNA object has .layers['counts'] and .layers['log1p']
      - Fits mu/sd on TRAIN using .layers['log1p']
      - Sets .X to z-scored/clipped values (model input), leaving layers intact

    ADT: unchanged from your original.
    """
    # RNA params fit on CITE train (will also ensure train has log1p layer)
    rna_params = rna_fit_params(
        cite_rna_tr, counts_layer=rna_counts_layer, target_sum=rna_target_sum, clip=rna_clip
    )

    cite_rna_pp_tr = preprocess_rna_apply(cite_rna_tr, rna_params, counts_layer=rna_counts_layer)
    cite_rna_pp_va = preprocess_rna_apply(cite_rna_va, rna_params, counts_layer=rna_counts_layer)
    cite_rna_pp_te = preprocess_rna_apply(cite_rna_te, rna_params, counts_layer=rna_counts_layer)
    vg_rna_pp      = preprocess_rna_apply(vg_rna_al,  rna_params, counts_layer=rna_counts_layer)

    # ADT params fit on CITE train
    adt_params = adt_fit_params(cite_adt_tr, counts_layer=adt_counts_layer)

    cite_adt_pp_tr = preprocess_adt_apply_counts(cite_adt_tr, adt_params, counts_layer=adt_counts_layer, clip=adt_clip)
    cite_adt_pp_va = preprocess_adt_apply_counts(cite_adt_va, adt_params, counts_layer=adt_counts_layer, clip=adt_clip)
    cite_adt_pp_te = preprocess_adt_apply_counts(cite_adt_te, adt_params, counts_layer=adt_counts_layer, clip=adt_clip)

    # DAb ADT: .X already processed
    if dab_standardize_unit:
        dab_adt_pp = preprocess_adt_from_processed_X(dab_adt_al, clip=adt_clip, standardize_to_ref={"mode":"unit"})
    else:
        dab_adt_pp = preprocess_adt_from_processed_X(dab_adt_al, clip=adt_clip, standardize_to_ref=None)

    return {
        "rna_params": rna_params,
        "adt_params": adt_params,
        "cite_rna_pp_tr": cite_rna_pp_tr,
        "cite_rna_pp_va": cite_rna_pp_va,
        "cite_rna_pp_te": cite_rna_pp_te,
        "vg_rna_pp": vg_rna_pp,
        "cite_adt_pp_tr": cite_adt_pp_tr,
        "cite_adt_pp_va": cite_adt_pp_va,
        "cite_adt_pp_te": cite_adt_pp_te,
        "dab_adt_pp": dab_adt_pp,
    }


# %% [code]
out = preprocess_all_for_aml_bridge(
    cite_rna_tr=cite_rna_tr, cite_rna_va=cite_rna_va, cite_rna_te=cite_rna_te,
    vg_rna_al=vg_rna_al,
    cite_adt_tr=cite_adt_tr, cite_adt_va=cite_adt_va, cite_adt_te=cite_adt_te,
    dab_adt_al=dab_adt_al,
    dab_standardize_unit=True,   # recommended if dab_adt_al.X scale is weird vs CITE
)

print(
  out["cite_rna_pp_tr"].shape, out["vg_rna_pp"].shape,
  out["cite_adt_pp_tr"].shape, out["dab_adt_pp"].shape
)


# %% [code]
# -----------------------------
# Use preprocessed objects from here onward
# -----------------------------
cite_rna_pp_tr = out["cite_rna_pp_tr"]
cite_rna_pp_va = out["cite_rna_pp_va"]
cite_rna_pp_te = out["cite_rna_pp_te"]

cite_adt_pp_tr = out["cite_adt_pp_tr"]
cite_adt_pp_va = out["cite_adt_pp_va"]
cite_adt_pp_te = out["cite_adt_pp_te"]

vg_rna_pp  = out["vg_rna_pp"]
dab_adt_pp = out["dab_adt_pp"]


# %% [markdown]
### Calculate the LSC-17 score from the scRNA data

# %% [code]
print(vg_rna_pp)
print(vg_rna_pp.var_names)

print(cite_rna_pp_tr)
print(cite_rna_pp_tr.var_names)


# %% [code]
print(vg_rna_pp.X)
print(vg_rna_pp.X.min())
print(vg_rna_pp.X.max())

print(vg_rna_pp.layers['log1p'])
print(vg_rna_pp.layers['log1p'].min())
print(vg_rna_pp.layers['log1p'].max())

print(cite_rna_pp_tr.X)
print(cite_rna_pp_tr.X.min())
print(cite_rna_pp_tr.X.max())

print(cite_rna_pp_tr.layers['log1p'])
print(cite_rna_pp_tr.layers['log1p'].min())
print(cite_rna_pp_tr.layers['log1p'].max())


# %% [code]
aliases = {
    "KIAA0125": ["FAM30A"],
    "NGFRAP1": ["BEX3"],
    "GPR56":   ["ADGRG1"],
}

present = set(vg_rna_pp.var_names)

for g, alts in aliases.items():
    hit = [a for a in [g, *alts] if a in present]
    print(g, "->", hit if hit else "MISSING")


# %% [code]
import numpy as np
import scipy.sparse as sp

LSC17_GENES = [
    "DNMT3B","ZBTB46","NYNRIN","ARHGAP22","LAPTM4B",
    "MMRN1","DPYSL3","FAM30A","CDK6","CPXM1",
    "SOCS2","SMIM24","EMP1","BEX3","CD34",
    "AKR1C3","ADGRG1"
]

def _row_mean(X):
    # works for dense or sparse
    if sp.issparse(X):
        return np.asarray(X.mean(axis=1)).ravel()
    return X.mean(axis=1)

def add_lsc17_score(
    adata,
    *,
    key="LSC17",
    genes=LSC17_GENES,
    layer=None,              # e.g. "log1p" if you store it; otherwise None uses adata.X
    uppercase_varnames=True  # helpful if one dataset is lower/upper mixed
):
    # Optionally standardize var_names casing for matching
    if uppercase_varnames:
        # avoid modifying in-place if you're worried; here we do in-place for convenience
        adata.var_names = adata.var_names.astype(str).str.upper()

    genes = [g.upper() for g in genes] if uppercase_varnames else list(genes)
    present = [g for g in genes if g in adata.var_names]
    missing = [g for g in genes if g not in adata.var_names]

    if len(present) == 0:
        raise ValueError(f"No LSC17 genes found in adata.var_names. Example var_names: {list(adata.var_names[:5])}")

    X = adata.layers[layer] if layer is not None else adata.X
    X_sig = adata[:, present].layers[layer] if layer is not None else adata[:, present].X

    score = _row_mean(X_sig)

    adata.obs[f"{key}_score"] = score.astype(np.float32)
    adata.obs[f"{key}_z"] = ((score - score.mean()) / (score.std() + 1e-8)).astype(np.float32)
    adata.obs[f"{key}_n_genes"] = np.int32(len(present))

    if missing:
        print(f"[{key}] Missing {len(missing)}/{len(genes)} genes (scored with {len(present)}): {missing}")
    else:
        print(f"[{key}] All {len(genes)} genes present.")

    return present, missing

# ---- run on BOTH objects (separately) ----
present_vg, missing_vg = add_lsc17_score(vg_rna_pp,      key="LSC17", layer='log1p', uppercase_varnames=True)
present_ct, missing_ct = add_lsc17_score(cite_rna_pp_tr, key="LSC17", layer='log1p', uppercase_varnames=True)
present_ct, missing_ct = add_lsc17_score(cite_rna_pp_va, key="LSC17", layer='log1p', uppercase_varnames=True)
present_ct, missing_ct = add_lsc17_score(cite_rna_pp_te, key="LSC17", layer='log1p', uppercase_varnames=True)

print(vg_rna_pp.obs[["LSC17_score","LSC17_z","LSC17_n_genes"]].describe())
print(cite_rna_pp_tr.obs[["LSC17_score","LSC17_z","LSC17_n_genes"]].describe())
print(cite_rna_pp_va.obs[["LSC17_score","LSC17_z","LSC17_n_genes"]].describe())
print(cite_rna_pp_te.obs[["LSC17_score","LSC17_z","LSC17_n_genes"]].describe())


# %% [markdown]
### Train UniVI on paired CITE (RNA + ADT)

# %% [code]
device = "cuda" if torch.cuda.is_available() else "cpu"
print("device:", device)


# %% [code]
# -----------------------------
# Train UniVI on paired CITE (preprocessed)
# -----------------------------
seed_everything(1)

#device = "cuda" if torch.cuda.is_available() else "cpu"
#print("device:", device)

# Sanity check training inputs
print("RNA pp range:", float(np.min(cite_rna_pp_tr.X)), float(np.max(cite_rna_pp_tr.X)))
print("ADT pp range:", float(np.min(cite_adt_pp_tr.X)), float(np.max(cite_adt_pp_tr.X)))

univi_cfg = UniVIConfig(
    #beta=1.125,
    beta=1.15,
    #gamma=1.475,
    gamma=1.75,
    #latent_dim=40,
    latent_dim=30,
    #encoder_dropout=0.20,
    #decoder_dropout=0.10,
    encoder_dropout=0.10,
    decoder_dropout=0.05,
    encoder_batchnorm=False,
    decoder_batchnorm=False,
    #kl_anneal_start=10,
    #kl_anneal_end=120,
    #align_anneal_start=20,
    #align_anneal_end=130,
    modalities=[
        ModalityConfig(
            name="rna",
            input_dim=cite_rna_pp_tr.n_vars,
            #encoder_hidden=[512, 256, 128, 64],
            #decoder_hidden=[64, 128, 256, 512],
            encoder_hidden=[1024, 512, 256, 128, 64],
            decoder_hidden=[64, 128, 256, 512, 1024],
            likelihood="gaussian",
            recon_weight=1.00,
        ),
        ModalityConfig(
            name="adt",
            input_dim=cite_adt_pp_tr.n_vars,
            #encoder_hidden=[128, 64],
            #decoder_hidden=[64, 128],
            encoder_hidden=[128, 64, 32],
            decoder_hidden=[32, 64, 128],
            likelihood="gaussian",
            #recon_weight=3.25,
            recon_weight=3.00,
        ),
    ],
)

model = UniVIMultiModalVAE(
    univi_cfg,
    loss_mode="v1",
    v1_recon="avg",
    normalize_v1_terms=True,
    #recon_normalize_by_dim=True,
    #recon_dim_power=0.4,
).to(device)

train_cfg = TrainingConfig(
    n_epochs=3000,
    lr=1e-4,
    weight_decay=1e-5,
    batch_size=256,
    grad_clip=5.0,
    early_stopping=True,
    patience=150,
    min_delta=0,
    log_every=25,
    device=device,
)

train_ds = MultiModalDataset({"rna": cite_rna_pp_tr, "adt": cite_adt_pp_tr})
val_ds   = MultiModalDataset({"rna": cite_rna_pp_va, "adt": cite_adt_pp_va})

pin = (device == "cuda")
train_loader = DataLoader(train_ds, batch_size=train_cfg.batch_size, shuffle=True,  num_workers=0, pin_memory=pin)
val_loader   = DataLoader(val_ds,   batch_size=train_cfg.batch_size, shuffle=False, num_workers=0, pin_memory=pin)

trainer = UniVITrainer(
    model=model,
    train_loader=train_loader,
    val_loader=val_loader,
    train_cfg=train_cfg,
    device=device,
)

history = trainer.fit()

plt.figure()
plt.plot(history["train_loss"], label="train")
plt.plot(history["val_loss"], label="val")
plt.xlabel("epoch"); plt.ylabel("loss"); plt.legend()
plt.title("UniVI training loss (CITE paired; preprocessed)")
plt.show()


# %% [markdown]
### Encode latents for CITE test + projections (VG RNA, DAb ADT)

# %% [code]
@torch.no_grad()
def encode_latent(model, adata: ad.AnnData, modality: str, device=device, batch_size=1024, latent="modality_mean"):
    Z = ue.encode_adata(
        model,
        adata,
        modality=modality,
        latent=latent,
        device=device,
        batch_size=batch_size,
    )
    return np.asarray(Z, dtype=np.float32)

# Encode from PREPROCESSED objects
Z_cite_rna = encode_latent(model, cite_rna_pp_te, modality="rna", device=device)
Z_cite_adt = encode_latent(model, cite_adt_pp_te, modality="adt", device=device)

Z_vg_rna   = encode_latent(model, vg_rna_pp,      modality="rna", device=device)
Z_dab_adt  = encode_latent(model, dab_adt_pp,     modality="adt", device=device)

# Store
cite_rna_te.obsm["X_univi"]    = Z_cite_rna
cite_adt_te.obsm["X_univi"]    = Z_cite_adt
vg_rna_pp.obsm["X_univi"]      = Z_vg_rna
dab_adt_pp.obsm["X_univi"]     = Z_dab_adt

print("Latents shapes:", Z_cite_rna.shape, Z_cite_adt.shape, Z_vg_rna.shape, Z_dab_adt.shape)


# %% [markdown]
### Build joint latent UMAP for panels (b/c/d)

# %% [code]
import scanpy as sc

sc.settings.set_figure_params(
    dpi=100,        # notebook display dpi
    dpi_save=300,   # saved figure dpi
    figsize=(10, 8), # default size in inches
    fontsize=10,
)

#sc.settings.set_figure_params()


# %% [code]
def joint_latent_adata(items):
    Zs, obs_rows = [], []
    for ds_name, mod, a in items:
        Z = a.obsm["X_univi"]
        Zs.append(Z)
        obs_rows.append(pd.DataFrame({"dataset": ds_name, "modality": mod}, index=a.obs_names.copy()))
    Z = np.vstack(Zs).astype(np.float32)
    obs = pd.concat(obs_rows, axis=0)
    out = ad.AnnData(X=Z, obs=obs, var=pd.DataFrame(index=[f"z{i}" for i in range(Z.shape[1])]))
    out.obsm["X_univi"] = out.X.copy()
    return out

joint = joint_latent_adata([
    ("CITE", "RNA", cite_rna_te),
    ("CITE", "ADT", cite_adt_te),
    ("DAb",  "ADT", dab_adt_pp),
    ("VG",   "RNA", vg_rna_pp),
])

joint.obs["dataset_modality"] = joint.obs["dataset"].astype(str) + " " + joint.obs["modality"].astype(str)

sc.pp.neighbors(joint, use_rep="X_univi", n_neighbors=30, random_state=1)
sc.tl.umap(joint, random_state=1)


# %% [code]
# Fig7b
sc.pl.umap(joint, 
           color="dataset_modality", 
           title="Joint latent (dataset × modality) using CITE-seq test sets and outside unimodal data",
           size=10,
           alpha=0.7,
)

plt.savefig(FIGDIR / "fig7_umap_dataset_modality.png", dpi=300)
plt.show()


# %% [markdown]
### Figure 7c

# %% [code]
print(cite_rna_te)
print(cite_adt_te)
print(vg_rna_pp)
print(dab_adt_pp)


# %% [code]
# OPTIONAL: set these explicitly if you know them
CITE_LABEL_COL = None        # e.g. "celltype.l2" / "celltype" / "annotation"
VG_LABEL_COL   = "CellType"  # e.g. "celltype" / "annotation" / "coarse"

def find_label_col(adata, candidates):
    for c in candidates:
        if c in adata.obs.columns:
            return c
    return None

def to_coarse(label: str) -> str:
    s = str(label).lower()
    if s in {"nan", "none", ""}:
        return "NA"
    if any(k in s for k in ["blast", "malignant", "leuk", "aml"]):
        return "blasts/malignant-like"
    if any(k in s for k in ["hsc", "stem", "prog", "cmp", "gmp", "progen"]):
        return "progenitor-like"
    if any(k in s for k in ["mono", "mac", "cd14", "fcgr3a"]):
        return "mono/mac"
    if any(k in s for k in ["dc", "dend"]):
        return "DC"
    if any(k in s for k in ["b cell", "bcell", "cd19", "ms4a1", "plasma"]):
        return "B"
    if any(k in s for k in ["t cell", "tcell", "cd3"]):
        return "T"
    if "nk" in s or "ncam1" in s or "kldr" in s:
        return "NK"
    if any(k in s for k in ["ery", "rbc", "glyc"]):
        return "erythroid"
    return "other/NA"

if CITE_LABEL_COL is None:
    CITE_LABEL_COL = find_label_col(cite_rna_te, ["celltype_coarse","celltype","cell_type","celltype.l2","celltype.l1","annot","annotation","labels"])
if VG_LABEL_COL is None:
    VG_LABEL_COL = find_label_col(vg_rna_pp, ["celltype_coarse","celltype","cell_type","annot","annotation","labels"])

print("CITE_LABEL_COL:", CITE_LABEL_COL)
print("VG_LABEL_COL:", VG_LABEL_COL)

joint.obs["coarse_state"] = "NA"

def fill_coarse(joint, src, dataset, modality, col):
    if col is None or col not in src.obs.columns:
        return
    m = (joint.obs["dataset"] == dataset) & (joint.obs["modality"] == modality)
    idx = joint.obs_names[m]
    joint.obs.loc[idx, "coarse_state"] = src.obs.loc[idx, col].astype(str).map(to_coarse).values

fill_coarse(joint, cite_rna_te,    "CITE", "RNA", CITE_LABEL_COL)
fill_coarse(joint, cite_adt_te, "CITE", "ADT", CITE_LABEL_COL)
#fill_coarse(joint, vg_rna_pp,      "VG",   "RNA", VG_LABEL_COL)

#sc.pl.umap(joint, color="coarse_state", title="Fig7c: Joint latent (coarse compartments)", show=True)
#plt.savefig(Path(FIGDIR) / "fig7c_umap_coarse_state.png", dpi=450)
#plt.show()


# %% [code]
print(vg_rna_pp)
print(vg_rna_pp.obs['CellType'])


# %% [code]
# Reindex CellType onto joint_vg cells
ct = vg_rna_pp.obs["CellType"].reindex(joint.obs_names)

# Optional sanity check
missing = ct.isna().sum()
print(f"Missing CellType for {missing}/{joint.n_obs} joint cells")

joint.obs["coarse_state"] = ct.astype("category")

#sc.pl.umap(joint, color="coarse_state", title="Fig7c: Joint latent", show=True)
sc.pl.umap(joint, 
           color="coarse_state", 
           title="Joint latent (van Galen cell type)",
           size=10,
           alpha=0.7,
)

plt.savefig(Path(FIGDIR) / "fig7_umap_VG_CellType.png", dpi=450)
plt.show()


# %% [code]
print(joint)
print(vg_rna_pp)


# %% [code]
import numpy as np
import pandas as pd

score_col = "LSC17_score"
out_col   = "LSC17_score"

# ensure the column exists
if out_col not in joint.obs.columns:
    joint.obs[out_col] = np.nan

def add_scores_fill_missing(src, joint, col, out_col):
    """
    Fill joint.obs[out_col] from src.obs[col] by matching on obs_names.

    Works even if joint.obs_names has duplicates.
    Only fills where joint[out_col] is currently missing (NaN).
    """
    # src obs_names are unique in your report; build mapper obs_name -> score
    mapper = pd.Series(src.obs[col].values, index=pd.Index(src.obs_names))

    # lookup for every joint row (duplicates are fine; they'll get same value)
    looked_up = pd.Index(joint.obs_names).map(mapper)

    # fill only where joint currently missing and looked_up is not missing
    cur = joint.obs[out_col].to_numpy(copy=True)

    # coerce looked_up to float if possible (LSC17_score should be numeric)
    new = pd.to_numeric(pd.Series(looked_up), errors="coerce").to_numpy()

    mask = np.isnan(cur) & ~np.isnan(new)
    cur[mask] = new[mask]
    joint.obs[out_col] = cur

    return joint.obs[out_col]

# choose your priority order:
add_scores_fill_missing(cite_rna_pp_tr, joint, score_col, out_col)
add_scores_fill_missing(cite_rna_pp_va, joint, score_col, out_col)
add_scores_fill_missing(cite_rna_pp_te, joint, score_col, out_col)
add_scores_fill_missing(vg_rna_pp,      joint, score_col, out_col)


# %% [code]
sc.pl.umap(
    joint,
    color="LSC17_score",           # or "LSC17_vg", "LSC17_cite_tr", etc.
    title="Joint latent (LSC-17 score)",
    size=10,
    alpha=0.7,
)

plt.savefig(Path(FIGDIR) / "fig7_umap_LSC17_any.png", dpi=450, bbox_inches="tight")
plt.show()


# %% [markdown]
### Figure 7d

# %% [code]
import re
import numpy as np
import pandas as pd
import anndata as ad

GENES = ["NPM1", "FLT3", "DNMT3A", "TET2", "IDH1", "IDH2", "TP53"]
HERO  = "NPM1"

# ============================================================
# Helpers
# ============================================================
def _coerce_str_series(x: pd.Series) -> pd.Series:
    s = x.astype("string")
    s = s.str.strip()
    s = s.mask(s.str.upper().isin(["NA", "N/A", "NONE", "NULL", "NAN", ""]), pd.NA)
    return s

def _gene_present(series: pd.Series, gene: str) -> np.ndarray:
    """
    Returns float array of {1.0, 0.0, nan}:
      - nan if series entry is NA/unknown
      - 1.0 if gene appears as a token
      - 0.0 otherwise
    """
    s = _coerce_str_series(series)
    known = s.notna()
    pat = re.compile(rf"\b{re.escape(gene)}\b", flags=re.IGNORECASE)

    out = np.full(len(s), np.nan, dtype=np.float32)
    if known.any():
        out[known.values] = s[known].apply(lambda t: 1.0 if pat.search(str(t)) else 0.0).to_numpy(dtype=np.float32)
    return out

# ============================================================
# VG: build Y/M either from patient-level metadata OR transcripts fields
# ============================================================
def build_vg_targets(
    adata: ad.AnnData,
    genes,
    *,
    use_patient_table: bool = False,
    patient_table: pd.DataFrame | None = None,
    patient_key: str = "orig.ident",   # where patient IDs live in vg_rna_pp.obs
    patient_id_col: str = "Sample",    # patient table column name for sample/patient id
    rhp_col: str = "RHP Mutations",    # patient table column listing mutations
    mut_col: str = "MutTranscripts",
    wt_col: str = "WtTranscripts",
    dataset_name: str = "VG",
):
    """
    If use_patient_table=True and patient_table provided:
      - Label every cell by patient-level status.
      - 'None Detected' => WT-labeled for all genes in 'genes'.
      - Mention of a gene token => MUT-labeled for that gene.
      - If table entry missing/unknown => unlabeled for those genes.

    Else:
      - Use your original transcript-fields logic.
    """
    G = len(genes)
    Y = np.zeros((adata.n_obs, G), dtype=np.float32)
    M = np.zeros((adata.n_obs, G), dtype=np.float32)

    # -------------------------
    # Option A: patient-level
    # -------------------------
    if use_patient_table and (patient_table is not None):
        if patient_key not in adata.obs.columns:
            raise KeyError(f"[{dataset_name}] patient_key='{patient_key}' not found in adata.obs")

        if patient_id_col not in patient_table.columns or rhp_col not in patient_table.columns:
            raise KeyError(f"[{dataset_name}] patient_table must have columns: '{patient_id_col}' and '{rhp_col}'")

        # build patient -> dict(gene -> 0/1/None)
        pat2 = {}
        for _, row in patient_table.iterrows():
            pat = str(row[patient_id_col])
            mut_str = str(row.get(rhp_col, "")).strip()
            low = mut_str.lower()

            d = {g: None for g in genes}

            if mut_str == "" or low in {"na", "unknown", "not performed", "nan"}:
                pat2[pat] = d
                continue

            # strong WT statement
            if "none detected" in low:
                for g in genes:
                    d[g] = 0

            # any explicit gene mention => mutant
            for g in genes:
                if re.search(rf"\b{re.escape(g)}\b", mut_str, flags=re.IGNORECASE):
                    d[g] = 1

            pat2[pat] = d

        pats = adata.obs[patient_key].astype(str).values
        for i, pat in enumerate(pats):
            d = pat2.get(pat, None)
            if d is None:
                continue
            for j, g in enumerate(genes):
                v = d.get(g, None)
                if v is None:
                    continue
                M[i, j] = 1.0
                Y[i, j] = float(v)

        print(f"[{dataset_name}] built targets from PATIENT TABLE ({patient_id_col} -> {rhp_col}); "
              f"labeled fractions:", dict(zip(genes, M.mean(axis=0).round(3))))
        return Y, M, {"mode": "patient_table", "patient_key": patient_key, "patient_id_col": patient_id_col, "rhp_col": rhp_col}

    # -------------------------
    # Option B: transcript fields (your original behavior)
    # -------------------------
    obs = adata.obs
    if mut_col not in obs.columns:
        raise KeyError(f"[{dataset_name}] missing obs['{mut_col}']")
    if wt_col not in obs.columns:
        raise KeyError(f"[{dataset_name}] missing obs['{wt_col}']")

    mut_s = obs[mut_col]
    wt_s  = obs[wt_col]

    for j, g in enumerate(genes):
        mut_has = _gene_present(mut_s, g)   # 1/0/nan
        wt_has  = _gene_present(wt_s,  g)   # 1/0/nan

        is_mut = np.isfinite(mut_has) & (mut_has == 1.0)
        is_wt  = (~is_mut) & np.isfinite(wt_has) & (wt_has == 1.0)

        labeled = is_mut | is_wt
        M[labeled, j] = 1.0
        Y[is_mut, j]  = 1.0

    print(f"[{dataset_name}] built targets from transcript fields: mut='{mut_col}', wt='{wt_col}'; "
          f"labeled fractions:", dict(zip(genes, M.mean(axis=0).round(3))))
    return Y, M, {"mode": "transcript_fields", "mut_col": mut_col, "wt_col": wt_col}

# ============================================================
# DAb: gene-level labels from *ALL* matching variant columns
# ============================================================
def _looks_binaryish(s: pd.Series) -> bool:
    if pd.api.types.is_bool_dtype(s) or pd.api.types.is_integer_dtype(s) or pd.api.types.is_float_dtype(s):
        u = pd.unique(pd.Series(s.values).dropna())
        # allow {0,1} or {0.0,1.0}
        try:
            uu = set(float(x) for x in u)
        except Exception:
            return False
        return all(x in {0.0, 1.0} for x in uu) and len(uu) <= 2
    return False

def build_dab_targets_gene_level(
    adata: ad.AnnData,
    genes,
    *,
    dataset_name: str = "DAb",
    allow_dash_match: bool = True,
):
    """
    Build Y/M at the gene level by OR-ing across all obs columns that match that gene.
    Matching:
      - startswith gene (preferred): "NPM1 W288fs"
      - contains token gene: "FLT3-ITD" (if allow_dash_match True)
    Labeling:
      - M=1 if ANY matched column is non-NA for that cell
      - Y=1 if ANY matched column equals 1 among labeled columns
    """
    obs = adata.obs
    G = len(genes)
    Y = np.zeros((adata.n_obs, G), dtype=np.float32)
    M = np.zeros((adata.n_obs, G), dtype=np.float32)

    gene2cols = {g: [] for g in genes}
    cols = list(obs.columns)

    for g in genes:
        gU = g.upper()
        # startwith matches
        start_hits = [c for c in cols if str(c).upper().startswith(gU)]
        hits = list(start_hits)

        # token/contains matches (helps FLT3-ITD)
        if allow_dash_match:
            tok_hits = [c for c in cols if re.search(rf"\b{re.escape(g)}\b", str(c), flags=re.IGNORECASE)]
            for c in tok_hits:
                if c not in hits:
                    hits.append(c)

        # keep only binary-ish columns if possible (most DAb mutation cols are 0/1)
        if len(hits) > 1:
            bin_hits = [c for c in hits if _looks_binaryish(obs[c])]
            if len(bin_hits) > 0:
                hits = bin_hits

        gene2cols[g] = hits

    print(f"[{dataset_name}] gene->matched_cols:")
    for g, cs in gene2cols.items():
        if cs:
            print(" ", g, ":", cs)

    for j, g in enumerate(genes):
        cs = gene2cols[g]
        if not cs:
            continue

        # stack as float matrix (n, k)
        X = np.vstack([pd.to_numeric(obs[c], errors="coerce").to_numpy(dtype=np.float32) for c in cs]).T

        called = np.isfinite(X).any(axis=1)
        M[:, j] = called.astype(np.float32)

        # mutant if any == 1 among columns
        mut = called & (np.nanmax(X, axis=1) >= 1.0)
        Y[mut, j] = 1.0

    print(f"[{dataset_name}] labeled fractions:", dict(zip(genes, M.mean(axis=0).round(3))))
    return Y, M, gene2cols


# %% [code]
# -------------------------
# VG labels:
# Option A (recommended if you have the patient mutation table loaded):
#   set use_patient_table=True and pass patient_table=vg_patient_df
# Option B (fallback): transcript fields
# -------------------------

USE_VG_PATIENT_TABLE = False  # <-- flip to True if you load the patient table
vg_patient_df = None          # <-- set this to your loaded DF when USE_VG_PATIENT_TABLE=True

# IMPORTANT: your split code detected group_source=orig.ident in your log,
# so use patient_key="orig.ident" to match that.
Y_vg, M_vg, vg_info = build_vg_targets(
    vg_rna_pp,
    GENES,
    use_patient_table=USE_VG_PATIENT_TABLE,
    patient_table=vg_patient_df,
    patient_key="orig.ident",
    patient_id_col="Sample",
    rhp_col="RHP Mutations",
    mut_col="MutTranscripts",
    wt_col="WtTranscripts",
    dataset_name="VG",
)

# -------------------------
# DAb labels (gene-level OR across all matched variant cols)
# -------------------------
Y_dab, M_dab, dab_gene2cols = build_dab_targets_gene_level(
    dab_adt_pp,
    GENES,
    dataset_name="DAb",
)

print("VG labeling mode:", vg_info)


# %% [code]
def knn_transfer_probs(Z_source, Y_source, M_source, Z_target, k=50):
    knn = NearestNeighbors(n_neighbors=k, metric="euclidean").fit(Z_source)
    idx = knn.kneighbors(Z_target, return_distance=False)  # (Nt,k)
    G = Y_source.shape[1]
    P = np.full((Z_target.shape[0], G), np.nan, dtype=np.float32)
    for g in range(G):
        ys = Y_source[:, g]
        ms = M_source[:, g]
        ms_k = ms[idx]
        ys_k = ys[idx]
        denom = ms_k.sum(axis=1)
        num = (ys_k * ms_k).sum(axis=1)
        ok = denom > 0
        P[ok, g] = (num[ok] / denom[ok]).astype(np.float32)
    return P

hero_i = GENES.index(HERO)

k=60

P_dab_from_vg = knn_transfer_probs(
    Z_source=vg_rna_pp.obsm["X_univi"],
    Y_source=Y_vg,
    M_source=M_vg,
    Z_target=dab_adt_pp.obsm["X_univi"],
    k=k
)

dab_adt_al.obs[f"knnP_{HERO}_vg_to_dab"] = P_dab_from_vg[:, hero_i]

# put onto joint for plotting (DAb ADT subset)
col = f"knnP_{HERO}_vg_to_dab"
joint.obs[col] = np.nan
mask_dab = (joint.obs["dataset"] == "DAb") & (joint.obs["modality"] == "ADT")
joint.obs.loc[joint.obs_names[mask_dab], col] = dab_adt_al.obs[col].values

#sc.pl.umap(joint, color=col, title=f"Fig7d: Transfer {HERO} probability (VG→DAb, kNN (k={k}))", show=True)
sc.pl.umap(joint, 
           color=col, 
           title=f"Transfer {HERO} probability (VG→DAb, kNN (k={k}))",
           size=10,
           alpha=0.7,
)

plt.savefig(Path(FIGDIR) / f"fig7_umap_transfer_{HERO}_vg_to_dab.png", dpi=450)
plt.show()


# %% [code]
from pathlib import Path
import numpy as np
import pandas as pd
import scanpy as sc
import matplotlib.pyplot as plt

# ----------------------------
# Config
# ----------------------------
FIGDIR = Path(FIGDIR)
FIGDIR.mkdir(parents=True, exist_ok=True)

# ----------------------------
# Helpers
# ----------------------------
def find_mut_col_dab_obs(adata, gene):
    cols = list(adata.obs.columns)
    gene_u = str(gene).upper()

    # Exact match (rare)
    for c in cols:
        if c.upper() == gene_u:
            return c

    # Startswith 'GENE ' (common: "NPM1 W288fs")
    for c in cols:
        if c.upper().startswith(gene_u + " "):
            return c

    # Token contains
    for c in cols:
        toks = c.upper().replace("-", " ").replace("_", " ").split()
        if gene_u in toks:
            return c

    # Special-case FLT3-ITD style
    if gene_u == "FLT3":
        for c in cols:
            cu = c.upper()
            if "FLT3" in cu and "ITD" in cu:
                return c

    return None


def coerce_to_nullable_boolean(x):
    """
    Convert an array/Series into pandas nullable boolean ('boolean') with <NA>.
    Accepts: 0/1, True/False, strings like '0','1','WT','Mut', etc.
    """
    s = pd.Series(x)

    # Already boolean-ish
    if pd.api.types.is_bool_dtype(s) or str(s.dtype).lower() == "boolean":
        return s.astype("boolean")

    # Numeric
    if pd.api.types.is_numeric_dtype(s):
        out = pd.Series(pd.NA, index=s.index, dtype="boolean")
        out[s == 1] = True
        out[s == 0] = False
        return out

    # Strings / objects
    ss = s.astype("string").str.strip().str.lower()
    out = pd.Series(pd.NA, index=s.index, dtype="boolean")
    true_set  = {"true","t","1","yes","y","mut","mutant","pos","positive"}
    false_set = {"false","f","0","no","n","wt","wildtype","neg","negative"}
    out[ss.isin(true_set)] = True
    out[ss.isin(false_set)] = False
    return out


def make_label_from_nullable_bool(s_bool, *, wt_label="WT", mut_label="Mut"):
    """
    Convert nullable boolean Series -> categorical labels WT/Mut with NA preserved.
    Returns pandas Categorical.
    """
    s = pd.Series(pd.NA, index=s_bool.index, dtype="string")
    s.loc[s_bool == True]  = mut_label
    s.loc[s_bool == False] = wt_label
    return pd.Categorical(s, categories=[wt_label, mut_label], ordered=True)


# ----------------------------
# 1) Identify actual DAb mutation column for HERO
# ----------------------------
mut_col = find_mut_col_dab_obs(dab_adt_al, HERO)
if mut_col is None:
    raise KeyError(
        f"Couldn't find an actual mutation column for {HERO} in dab_adt_al.obs.\n"
        f"Example columns: {list(dab_adt_al.obs.columns[:40])}"
    )
print(f"[{HERO}] using DAb obs column: {mut_col}")

# ----------------------------
# 2) Create clean nullable-boolean actual mutation on DAb AnnData
# ----------------------------
col_actual = f"actual_{HERO}_mut"
dab_adt_al.obs[col_actual] = coerce_to_nullable_boolean(dab_adt_al.obs[mut_col])

print(
    f"  True:  {(dab_adt_al.obs[col_actual] == True).sum()}\n"
    f"  False: {(dab_adt_al.obs[col_actual] == False).sum()}\n"
    f"  NA:    {dab_adt_al.obs[col_actual].isna().sum()}"
)

# ----------------------------
# 3) Copy actual mutation onto joint ONLY for DAb ADT cells
#    Keep as nullable boolean dtype
# ----------------------------
mask_dab = (joint.obs["dataset"] == "DAb") & (joint.obs["modality"] == "ADT")

# initialize as nullable boolean
joint.obs[col_actual] = pd.Series(pd.NA, index=joint.obs_names, dtype="boolean")

# assign values for DAb ADT subset
joint.obs.loc[mask_dab, col_actual] = (
    dab_adt_al.obs[col_actual].astype("boolean").to_numpy()
)

# ----------------------------
# 4) Create a categorical label column for plotting (THIS is what we plot)
# ----------------------------
col_actual_plot = f"{col_actual}_label"
joint.obs[col_actual_plot] = make_label_from_nullable_bool(joint.obs[col_actual])

# ----------------------------
# 5) Plot: Actual mutation (categorical legend, NA grey)
# ----------------------------
'''
sc.pl.umap(
    joint,
    color=col_actual_plot,          # <-- categorical label, avoids boolean negation bug
    title=f"Actual {HERO} mutation (DAb observed) on joint UMAP",
    na_color="lightgrey",
    show=False,
)
'''
sc.pl.umap(joint, 
           color=col_actual_plot, 
           title=f"Actual {HERO} mutation (DAb observed) on joint UMAP",
           size=10,
           alpha=0.7,
)

plt.savefig(FIGDIR / f"fig7d_umap_actual_{HERO}_mut_on_joint.png", dpi=450, bbox_inches="tight")
plt.show()

# ----------------------------
# 6) Plot: Predicted/Transferred probability (continuous), if present
# ----------------------------
col_prob = f"knnP_{HERO}_vg_to_dab"
if col_prob in joint.obs.columns:
    '''
    sc.pl.umap(
        joint,
        color=col_prob,
        title=f"Transfer {HERO} probability (VG→DAb, kNN) on joint UMAP",
        na_color="lightgrey",
        show=False,
    )
    '''
    sc.pl.umap(joint, 
           color=col_prob, 
           title=f"Transfer {HERO} probability (VG→DAb, kNN) on joint UMAP",
           size=10,
           alpha=0.7,
    )
    plt.savefig(FIGDIR / f"fig7_umap_transfer_{HERO}_vg_to_dab_on_joint.png", dpi=450, bbox_inches="tight")
    plt.show()
else:
    print(f"NOTE: {col_prob} not found in joint.obs; skipping transfer-prob UMAP.")

# ----------------------------
# 7) OPTIONAL: Side-by-side (Actual label + Pred prob)
# ----------------------------
if col_prob in joint.obs.columns:
    '''
    sc.pl.umap(
        joint,
        color=[col_actual_plot, col_prob],
        title=[f"Actual {HERO} (DAb)", f"Transfer P({HERO}) (VG→DAb)"],
        na_color="lightgrey",
        wspace=0.35,
        show=False,
    )
    '''
    sc.pl.umap(
        joint, 
        color=[col_actual_plot, col_prob], 
        title=[f"Actual {HERO} (DAb)", f"Transfer P({HERO}) (VG→DAb)"],
        size=10,
        alpha=0.7,
    )
    plt.savefig(FIGDIR / f"fig7_umap_actual_and_transfer_{HERO}.png", dpi=450, bbox_inches="tight")
    plt.show()


# %% [code]
from sklearn.neighbors import NearestNeighbors

def knn_transfer_mut_prob_1d(Z_source, y_source_bool, Z_target, k=30):
    """
    y_source_bool: pandas Series or array-like nullable boolean (dtype 'boolean' ok)
      True=Mut, False=WT, NA=unknown

    Returns:
      p_mut: (Nt,) float32 with NaN where no labeled neighbors
      denom: (Nt,) int number of labeled neighbors among k
    """
    # Coerce to numpy with mask
    y = pd.Series(y_source_bool).astype("boolean")
    y_val = y.to_numpy(dtype=object)  # will contain True/False/<NA>
    labeled = pd.notna(y_val)         # boolean mask
    y01 = np.zeros(len(y_val), dtype=np.float32)
    y01[labeled] = (y_val[labeled] == True).astype(np.float32)

    knn = NearestNeighbors(n_neighbors=k, metric="euclidean").fit(Z_source)
    idx = knn.kneighbors(Z_target, return_distance=False)  # (Nt,k)

    lab_k = labeled[idx]                # (Nt,k)
    denom = lab_k.sum(axis=1).astype(np.int32)
    num = (y01[idx] * lab_k).sum(axis=1).astype(np.float32)

    p_mut = np.full(Z_target.shape[0], np.nan, dtype=np.float32)
    ok = denom > 0
    p_mut[ok] = num[ok] / denom[ok]
    return p_mut, denom


# %% [code]
# ----------------------------
# DAb -> VG transfer (HERO)
# ----------------------------
k = 50
col_actual = f"actual_{HERO}_mut"              # nullable boolean on DAb
col_p = f"knnP_{HERO}_dab_to_vg"               # transferred prob
col_p_denom = f"knnN_{HERO}_dab_to_vg"         # labeled neighbors count
col_label = f"knnL_{HERO}_dab_to_vg_label"     # categorical hard label

# 1) Transfer probability onto VG cells
p_vg, denom_vg = knn_transfer_mut_prob_1d(
    Z_source=dab_adt_pp.obsm["X_univi"],                 # source embedding
    y_source_bool=dab_adt_al.obs[col_actual],            # source labels
    Z_target=vg_rna_pp.obsm["X_univi"],                  # target embedding
    k=k
)

vg_rna_pp.obs[col_p] = p_vg
vg_rna_pp.obs[col_p_denom] = denom_vg

# 2) Make a hard label for plotting (Mut/WT/NA)
#    (Choose threshold you like; 0.5 is standard)
thr = 0.75
lab = pd.Series(pd.NA, index=vg_rna_pp.obs_names, dtype="string")
lab.loc[vg_rna_pp.obs[col_p].notna() & (vg_rna_pp.obs[col_p] >= thr)] = "Mut"
lab.loc[vg_rna_pp.obs[col_p].notna() & (vg_rna_pp.obs[col_p] <  thr)] = "WT"
vg_rna_pp.obs[col_label] = pd.Categorical(lab, categories=["WT", "Mut"], ordered=True)

print(
    f"[{HERO}] DAb→VG transfer (k={k}, thr={thr})\n"
    f"  VG cells w/ labeled neighbors: {(vg_rna_pp.obs[col_p_denom] > 0).sum()} / {vg_rna_pp.n_obs}\n"
    f"  Pred Mut: {(vg_rna_pp.obs[col_label] == 'Mut').sum()}\n"
    f"  Pred WT:  {(vg_rna_pp.obs[col_label] == 'WT').sum()}\n"
    f"  NA:       {pd.isna(vg_rna_pp.obs[col_label]).sum()}"
)

# 3) Ensure VG UMAP exists (only if needed)
if "X_umap" not in vg_rna_pp.obsm_keys():
    sc.pp.neighbors(vg_rna_pp, use_rep="X_univi", n_neighbors=30)
    sc.tl.umap(vg_rna_pp, random_state=42)

# 4) Plot probability (continuous)
sc.pl.umap(
    vg_rna_pp,
    color=col_p,
    title=f"Transfer P({HERO} Mut) DAb→VG (kNN k={k})",
    size=10,
    alpha=0.7,
    na_color="lightgrey",
    show=False,
)
plt.savefig(FIGDIR / f"umap_vg_transferP_{HERO}_dab_to_vg.png", dpi=450, bbox_inches="tight")
plt.show()

# 5) Plot hard label (categorical)
sc.pl.umap(
    vg_rna_pp,
    color=col_label,
    title=f"Transfer label {HERO} DAb→VG (thr={thr}, k={k})",
    size=10,
    alpha=0.7,
    na_color="lightgrey",
    show=False,
)
plt.savefig(FIGDIR / f"umap_vg_transferLabel_{HERO}_dab_to_vg.png", dpi=450, bbox_inches="tight")
plt.show()

# 6) Optional side-by-side
sc.pl.umap(
    vg_rna_pp,
    color=[col_label, col_p],
    title=[f"{HERO} label (DAb→VG)", f"P({HERO} Mut) (DAb→VG)"],
    size=10,
    alpha=0.7,
    na_color="lightgrey",
    wspace=0.35,
    show=False,
)
plt.savefig(FIGDIR / f"umap_vg_transferLabel_and_P_{HERO}_dab_to_vg.png", dpi=450, bbox_inches="tight")
plt.show()


# %% [code]
import re
import pandas as pd
import scanpy as sc
import matplotlib.pyplot as plt

def _contains_gene(text, gene):
    """
    Returns True if `gene` appears as a token in `text` (case-insensitive).
    Robust to separators like ',', ';', '|', '/', '-', etc.
    """
    if text is None or (isinstance(text, float) and pd.isna(text)) or pd.isna(text):
        return False
    s = str(text).upper()
    # turn non-alphanum into spaces, then token-match
    s = re.sub(r"[^A-Z0-9]+", " ", s)
    g = str(gene).upper()
    return re.search(rf"\b{re.escape(g)}\b", s) is not None

def vg_actual_mut_from_transcripts(vg, gene, mut_col="MutTranscripts", wt_col="WtTranscripts"):
    """
    Build a nullable-boolean Series:
      True  = gene present in MutTranscripts
      False = gene present in WtTranscripts
      <NA>  = neither or both (ambiguous)
    """
    has_mut = vg.obs[mut_col].apply(lambda x: _contains_gene(x, gene))
    has_wt  = vg.obs[wt_col].apply(lambda x: _contains_gene(x, gene))

    out = pd.Series(pd.NA, index=vg.obs_names, dtype="boolean")
    out[ has_mut & ~has_wt] = True
    out[~has_mut &  has_wt] = False
    # ambiguous if both
    out[ has_mut &  has_wt] = pd.NA
    return out, has_mut, has_wt

def make_label_from_nullable_bool(s_bool, wt_label="WT", mut_label="Mut"):
    s = pd.Series(pd.NA, index=s_bool.index, dtype="string")
    s.loc[s_bool == True]  = mut_label
    s.loc[s_bool == False] = wt_label
    return pd.Categorical(s, categories=[wt_label, mut_label], ordered=True)

# ----------------------------
# Build "actual mutation" from transcript-string columns
# ----------------------------
col_bool  = f"actual_{HERO}_vg_bool"
col_label = f"actual_{HERO}_vg_label"

vg_rna_pp.obs[col_bool], has_mut, has_wt = vg_actual_mut_from_transcripts(vg_rna_pp, HERO)

print(
    f"[{HERO}] from MutTranscripts/WtTranscripts\n"
    f"  Mut-only (True):  {(vg_rna_pp.obs[col_bool] == True).sum()}\n"
    f"  WT-only (False):  {(vg_rna_pp.obs[col_bool] == False).sum()}\n"
    f"  NA (neither/both):{vg_rna_pp.obs[col_bool].isna().sum()}\n"
    f"  ambiguous (both): {(has_mut & has_wt).sum()}"
)

# categorical labels for plotting (prevents continuous/boolean quirks)
vg_rna_pp.obs[col_label] = make_label_from_nullable_bool(vg_rna_pp.obs[col_bool])

# ----------------------------
# Plot on VG UMAP
# ----------------------------
sc.pl.umap(
    vg_rna_pp,
    color=[col_label],
    title=[f"van Galen actual {HERO} mutation (from transcripts)"],
    size=20,
    alpha=0.7,
    na_color="lightgrey",
    show=True,
)
plt.show()

# optional: side-by-side with patient/timepoint
# sc.pl.umap(
#     vg_rna_pp,
#     color=["patient", "timepoint", col_label],
#     title=["patient", "timepoint", f"actual {HERO}"],
#     na_color="lightgrey",
#     wspace=0.35,
#     show=True,
# )
# plt.show()


# %% [code]
print(joint)


# %% [code]
import numpy as np
import pandas as pd
import scanpy as sc
import matplotlib.pyplot as plt

HERO = "NPM1"  # if you're using this pattern elsewhere

# --- columns in VG ---
vg_bool_col  = "actual_NPM1_vg_bool"    # True/False (or bool-ish)
vg_label_col = "actual_NPM1_vg_label"   # e.g. "Mut"/"WT" (categorical)

# --- output columns in joint ---
out_bool_col  = "actual_NPM1_mut"
out_label_col = "actual_NPM1_mut_label"

def map_vg_obs_to_joint(joint, vg, vg_col, out_col, *, to_category=False, fill_value=np.nan, overwrite=True):
    """
    Map vg.obs[vg_col] onto joint.obs_names (works even if joint.obs_names has duplicates).
    """
    if (out_col in joint.obs.columns) and (not overwrite):
        raise ValueError(f"{out_col} already exists in joint.obs (set overwrite=True to replace).")

    mapper = pd.Series(vg.obs[vg_col].values, index=pd.Index(vg.obs_names))  # vg obs_names are unique
    looked_up = pd.Index(joint.obs_names).map(mapper)  # duplicates in joint are fine

    s = pd.Series(looked_up, index=joint.obs_names, name=out_col)

    if fill_value is not np.nan:
        s = s.fillna(fill_value)

    if to_category:
        s = s.astype("category")

    joint.obs[out_col] = s
    return joint.obs[out_col]

# 1) boolean-ish mutation status
map_vg_obs_to_joint(joint, vg_rna_pp, vg_bool_col, out_bool_col, to_category=False, overwrite=True)

# 2) label version (Mut/WT)
map_vg_obs_to_joint(joint, vg_rna_pp, vg_label_col, out_label_col, to_category=True, overwrite=True)

# Optional: make the bool column nicer for plotting (keeps NaN as missing)
# joint.obs[out_bool_col] = joint.obs[out_bool_col].map({True: "Mut", False: "WT"}).astype("category")

# --- plot ---
sc.pl.umap(
    joint,
    color=[out_label_col],  # NOTE: pass the *string column name*
    title=[f"van Galen actual {HERO} mutation (from transcripts)"],
    size=20,
    alpha=0.7,
    na_color="lightgrey",
    show=True,
)
plt.show()


# %% [code]
import pandas as pd
import numpy as np

# 1) sample prefix from obs_names like "AML556.D0_cell123" -> "AML556.D0"
vg_rna_pp.obs["sample"] = vg_rna_pp.obs_names.to_series().str.split("_", n=1).str[0].values

s = vg_rna_pp.obs["sample"].astype(str)

# 2) default parse: patient = before first ".", timepoint = after first "."
patient = s.str.extract(r"^([^.]+)")[0]
timepoint = s.str.extract(r"^[^.]+\.(.+)$")[0]  # becomes NaN if no "."

# 3) special-cases
# OCI.* looks like a cell line label (OCI.AML3) — treat as patient, no timepoint
is_oci = s.str.startswith("OCI.")
patient.loc[is_oci] = s.loc[is_oci]
timepoint.loc[is_oci] = pd.NA

# If no dot (e.g. BM1, BM2, MUTZ3.frozen if it had no dot), keep timepoint NA
# (already handled by the regex)

# 4) write into obs
vg_rna_pp.obs["patient"] = patient.astype("category")
vg_rna_pp.obs["timepoint"] = pd.Series(timepoint).astype("string").astype("category")

# 5) quick sanity checks
print(vg_rna_pp.obs[["sample", "patient", "timepoint"]].head())
print("\nPatients:", vg_rna_pp.obs["patient"].nunique())
print("Timepoints:", vg_rna_pp.obs["timepoint"].nunique(dropna=True))
print("\nTop sample counts:\n", vg_rna_pp.obs["sample"].value_counts().head(10))
print("\nNA timepoint:", vg_rna_pp.obs["timepoint"].isna().sum())


# %% [code]
# Plot (IMPORTANT: pass column name(s), not the Series)
sc.pl.umap(
    vg_rna_pp,
    color=["patient"],          # or ["patient", "timepoint"]
    title=["van Galen patient"],
    size=20,
    alpha=0.7,
    na_color="lightgrey",
    show=True,
)
plt.show()


# %% [code]
sc.pl.umap(
    vg_rna_pp,
    color=["CellType"],          # or ["patient", "timepoint"]
    title=["van Galen annotated cell type"],
    size=20,
    alpha=0.7,
    na_color="lightgrey",
    show=True,
)
plt.show()


# %% [code]
print(cite_rna_te)
print(cite_adt_te)
print(vg_rna_pp)
print(dab_adt_pp)


# %% [code]
import numpy as np
import pandas as pd
import scanpy as sc
import matplotlib.pyplot as plt
from pathlib import Path

FIGDIR = Path(FIGDIR)
FIGDIR.mkdir(parents=True, exist_ok=True)

# ----------------------------
# Config / column names
# ----------------------------
k = 50
thr = 0.80

col_p_vg     = f"knnP_{HERO}_dab_to_vg"          # in vg_rna_pp.obs
col_denom_vg = f"knnN_{HERO}_dab_to_vg"          # in vg_rna_pp.obs
col_lab_vg   = f"knnL_{HERO}_dab_to_vg_label"    # in vg_rna_pp.obs (Categorical WT/Mut)

# Where we will store them on JOINT for plotting
col_p_joint     = col_p_vg
col_denom_joint = col_denom_vg
col_lab_joint   = col_lab_vg

# ----------------------------
# Sanity checks
# ----------------------------
missing = [c for c in [col_p_vg, col_denom_vg, col_lab_vg] if c not in vg_rna_pp.obs.columns]
if missing:
    raise KeyError(
        f"Missing {missing} in vg_rna_pp.obs. "
        f"Run the DAb→VG transfer chunk first to create these."
    )

# ----------------------------
# Masks (VG RNA cells inside joint)
# ----------------------------
mask_vg_rna = (joint.obs["dataset"] == "VG") & (joint.obs["modality"] == "RNA")

# ----------------------------
# 1) Copy continuous probability onto joint
# ----------------------------
joint.obs[col_p_joint] = np.nan
joint.obs.loc[mask_vg_rna, col_p_joint] = vg_rna_pp.obs[col_p_vg].to_numpy()

# Optional: copy denom (# labeled neighbors among k) too
joint.obs[col_denom_joint] = np.nan
joint.obs.loc[mask_vg_rna, col_denom_joint] = vg_rna_pp.obs[col_denom_vg].to_numpy()

# ----------------------------
# 2) Copy categorical label onto joint (initialize as string/categorical, NOT float)
# ----------------------------
joint.obs[col_lab_joint] = pd.Series(pd.NA, index=joint.obs_names, dtype="string")
joint.obs.loc[mask_vg_rna, col_lab_joint] = vg_rna_pp.obs[col_lab_vg].astype("string").to_numpy()

# Make categorical for nice legend order
joint.obs[col_lab_joint] = pd.Categorical(
    joint.obs[col_lab_joint],
    categories=["WT", "Mut"],
    ordered=True,
)

# ----------------------------
# 3) Plot on JOINT UMAP
# ----------------------------
# (A) Probability (continuous)
sc.pl.umap(
    joint,
    color=col_p_joint,
    title=f"Transfer P({HERO} Mut) DAb→VG on JOINT UMAP (k={k})",
    size=10,
    alpha=0.7,
    na_color="lightgrey",
    show=False,
)
plt.savefig(FIGDIR / f"umap_joint_transferP_{HERO}_dab_to_vg.png", dpi=450, bbox_inches="tight")
plt.show()

# (B) Label (categorical)
sc.pl.umap(
    joint,
    color=col_lab_joint,
    title=f"Transfer label {HERO} DAb→VG on JOINT UMAP (thr={thr}, k={k})",
    size=10,
    alpha=0.7,
    na_color="lightgrey",
    show=False,
)
plt.savefig(FIGDIR / f"umap_joint_transferLabel_{HERO}_dab_to_vg.png", dpi=450, bbox_inches="tight")
plt.show()

# (C) Side-by-side
sc.pl.umap(
    joint,
    color=[col_lab_joint, col_p_joint],
    title=[f"{HERO} label (DAb→VG)", f"P({HERO} Mut) (DAb→VG)"],
    size=10,
    alpha=0.7,
    na_color="lightgrey",
    wspace=0.35,
    show=False,
)
plt.savefig(FIGDIR / f"umap_joint_transferLabel_and_P_{HERO}_dab_to_vg.png", dpi=450, bbox_inches="tight")
plt.show()


# %% [code]
sc.pl.umap(
    joint,
    color=["coarse_state"],
    title=[f"Joint UMAP with van Galen annotations overlaid"],
    size=16,
    alpha=0.7,
    na_color="lightgrey",
    wspace=0.35,
)


# %% [markdown]
### Figure 7e

# %% [code]
y_true = Y_dab[:, hero_i]
m_true = M_dab[:, hero_i].astype(bool)
p_pred = P_dab_from_vg[:, hero_i]

ok = m_true & np.isfinite(p_pred)
y = y_true[ok].astype(int)
p = p_pred[ok].astype(float)

if ok.sum() < 20 or len(np.unique(y)) < 2:
    print(f"Not enough labeled positives/negatives for {HERO}: n={ok.sum()}, classes={np.unique(y)}")
else:
    auc = roc_auc_score(y, p)
    ap  = average_precision_score(y, p)

    fpr, tpr, _ = roc_curve(y, p)
    prec, rec, _ = precision_recall_curve(y, p)

    plt.figure(figsize=(7.2, 3.2))
    plt.subplot(1, 2, 1)
    plt.plot(fpr, tpr)
    plt.plot([0,1],[0,1], linestyle="--")
    plt.xlabel("FPR"); plt.ylabel("TPR")
    plt.title(f"ROC (AUC={auc:.3f})")

    plt.subplot(1, 2, 2)
    plt.plot(rec, prec)
    plt.xlabel("Recall"); plt.ylabel("Precision")
    plt.title(f"PR (AP={ap:.3f})")

    plt.suptitle(f"{HERO} transfer VG→DAb (n={ok.sum()})")
    plt.tight_layout()
    plt.savefig(Path(FIGDIR) / f"fig7e_roc_pr_transfer_{HERO}.png", dpi=450)
    plt.show()

    print("Transfer metrics:", {"AUC": float(auc), "AP": float(ap), "n": int(ok.sum())})


# %% [code]
import numpy as np
import matplotlib.pyplot as plt
from pathlib import Path
from sklearn.metrics import (
    roc_auc_score, average_precision_score,
    roc_curve, precision_recall_curve
)

def eval_knn_transfer_dab_to_vg(
    *,
    vg_rna_pp,          # AnnData with knnP/knnN columns in .obs
    Y_vg, M_vg,         # arrays for VG cells (same gene order as GENES)
    genes, hero,
    figdir,
    k=50,
    thr=0.5,
    min_n=20,
    min_labeled_neighbors=1,   # set e.g. 10 to be stricter
    use_name_alignment=False,  # True if Y_vg/M_vg rows match vg_rna_pp.obs_names via a dict
    vg_obs_names_for_YM=None,  # required if use_name_alignment=True: array-like of names aligned to Y_vg rows
    prefix="Fig7e",
    save=True,
):
    hero_i = list(genes).index(hero)

    col_p = f"knnP_{hero}_dab_to_vg"
    col_n = f"knnN_{hero}_dab_to_vg"

    if col_p not in vg_rna_pp.obs.columns:
        raise KeyError(f"Missing {col_p} in vg_rna_pp.obs (run DAb→VG kNN transfer first).")
    if col_n not in vg_rna_pp.obs.columns:
        raise KeyError(f"Missing {col_n} in vg_rna_pp.obs (run DAb→VG kNN transfer first).")

    # --- predictions from kNN transfer ---
    p_pred_all = vg_rna_pp.obs[col_p].to_numpy(dtype=float)
    n_labnbrs  = vg_rna_pp.obs[col_n].to_numpy(dtype=float)

    # --- truth labels/masks for HERO ---
    Y_vg = np.asarray(Y_vg)
    M_vg = np.asarray(M_vg)

    if use_name_alignment:
        if vg_obs_names_for_YM is None:
            raise ValueError("If use_name_alignment=True, provide vg_obs_names_for_YM aligned to rows of Y_vg/M_vg.")
        # map truth arrays onto vg_rna_pp.obs_names
        name_to_i = {n:i for i,n in enumerate(np.asarray(vg_obs_names_for_YM))}
        idx = np.array([name_to_i.get(n, -1) for n in vg_rna_pp.obs_names], dtype=int)
        ok_name = idx >= 0
        y_true_all = np.full(vg_rna_pp.n_obs, np.nan, dtype=float)
        m_true_all = np.zeros(vg_rna_pp.n_obs, dtype=bool)
        y_true_all[ok_name] = Y_vg[idx[ok_name], hero_i]
        m_true_all[ok_name] = M_vg[idx[ok_name], hero_i].astype(bool)
    else:
        # positional alignment (fast) — assumes Y_vg/M_vg rows are in same order as vg_rna_pp
        if Y_vg.shape[0] != vg_rna_pp.n_obs or M_vg.shape[0] != vg_rna_pp.n_obs:
            raise ValueError(
                f"Positional alignment mismatch: vg_rna_pp.n_obs={vg_rna_pp.n_obs} "
                f"but Y_vg/M_vg have {Y_vg.shape[0]} rows. "
                f"Set use_name_alignment=True and pass vg_obs_names_for_YM."
            )
        y_true_all = Y_vg[:, hero_i].astype(float)
        m_true_all = M_vg[:, hero_i].astype(bool)

    # --- final mask for evaluation ---
    ok = (
        m_true_all
        & np.isfinite(p_pred_all)
        & np.isfinite(y_true_all)
        & (n_labnbrs >= float(min_labeled_neighbors))
    )

    y = y_true_all[ok].astype(int)
    p = p_pred_all[ok].astype(float)

    if ok.sum() < min_n or len(np.unique(y)) < 2:
        print(f"Not enough labeled positives/negatives for {hero}: n={ok.sum()}, classes={np.unique(y)}")
        return {"AUC": np.nan, "AP": np.nan, "n": int(ok.sum()), "hero": hero, "direction": "DAb→VG (kNN)"}

    auc = roc_auc_score(y, p)
    ap  = average_precision_score(y, p)
    fpr, tpr, _ = roc_curve(y, p)
    prec, rec, _ = precision_recall_curve(y, p)

    plt.figure(figsize=(7.2, 3.2))
    plt.subplot(1, 2, 1)
    plt.plot(fpr, tpr)
    plt.plot([0,1],[0,1], linestyle="--")
    plt.xlabel("FPR"); plt.ylabel("TPR")
    plt.title(f"ROC (AUC={auc:.3f})")

    plt.subplot(1, 2, 2)
    plt.plot(rec, prec)
    plt.xlabel("Recall"); plt.ylabel("Precision")
    plt.title(f"PR (AP={ap:.3f})")

    title = f"{prefix}: {hero} transfer DAb→VG (kNN k={k}, n≥{min_labeled_neighbors})"
    plt.suptitle(f"{title} (n={ok.sum()})")
    plt.tight_layout()

    figdir = Path(figdir)
    figdir.mkdir(parents=True, exist_ok=True)
    if save:
        plt.savefig(figdir / f"fig7e_roc_pr_transfer_{hero}_DAb_to_VG_knn.png", dpi=450)

    plt.show()

    out = {"AUC": float(auc), "AP": float(ap), "n": int(ok.sum()), "hero": hero, "direction": "DAb→VG (kNN)"}
    print("Transfer metrics:", out)
    return out


# %% [code]
HERO = "NPM1"
metrics = eval_knn_transfer_dab_to_vg(
    vg_rna_pp=vg_rna_pp,
    Y_vg=Y_vg, M_vg=M_vg,
    genes=GENES, hero=HERO,
    figdir=FIGDIR,
    k=50,
    min_labeled_neighbors=5,   # try 1, 5, 10
    use_name_alignment=False,
)


# %% [markdown]
### Figure 7f

# %% [code]
import torch
import torch.nn as nn
import torch.nn.functional as F
from torch.utils.data import Dataset, DataLoader, Subset
from copy import deepcopy
from sklearn.model_selection import GroupShuffleSplit

class SingleModalDataset(Dataset):
    def __init__(self, adata: ad.AnnData, modality: str, Y: np.ndarray, M: np.ndarray):
        self.modality = modality
        self.X = adata.X
        self.Y = Y.astype(np.float32, copy=False)
        self.M = M.astype(np.float32, copy=False)

    def __len__(self): return self.Y.shape[0]

    def __getitem__(self, i):
        x = self.X[i]
        if hasattr(x, "toarray"):
            x = x.toarray().ravel()
        x = np.asarray(x, dtype=np.float32)
        return self.modality, torch.from_numpy(x), torch.from_numpy(self.Y[i]), torch.from_numpy(self.M[i])

def masked_bce_with_logits(logits, y, m, eps=1e-8):
    loss = F.binary_cross_entropy_with_logits(logits, y, reduction="none") * m
    denom = m.sum().clamp_min(eps)
    return loss.sum() / denom

class MLPHead(nn.Module):
    def __init__(self, in_dim, out_dim, hidden=(64, 32), dropout=0.1):
        super().__init__()
        self.net = nn.Sequential(
            nn.Linear(in_dim, hidden[0]), nn.ReLU(), nn.Dropout(dropout),
            nn.Linear(hidden[0], hidden[1]), nn.ReLU(), nn.Dropout(dropout),
            nn.Linear(hidden[1], out_dim),
        )
    def forward(self, z): return self.net(z)

@torch.no_grad()
def latent_mu(model, x, modality: str):
    x = x.to(device, non_blocking=True)
    mu_dict, _ = model.encode_modalities({modality: x})
    return mu_dict[modality]

def freeze_decoders(model: nn.Module, freeze=True):
    for name, p in model.named_parameters():
        if "decoder" in name.lower():
            p.requires_grad = (not freeze)

def split_group(n, groups, seed=0, frac_train=0.7, frac_val=0.15):
    groups = np.asarray(groups)
    idx_all = np.arange(n)
    gss1 = GroupShuffleSplit(n_splits=1, train_size=frac_train, random_state=seed)
    tr, tmp = next(gss1.split(idx_all, groups=groups))
    tmp_groups = groups[tmp]
    frac_val_of_tmp = frac_val / (1.0 - frac_train)
    gss2 = GroupShuffleSplit(n_splits=1, train_size=frac_val_of_tmp, random_state=seed+1)
    va_rel, te_rel = next(gss2.split(tmp, groups=tmp_groups))
    va = tmp[va_rel]
    te = tmp[te_rel]
    return tr, va, te


# %% [code]
import numpy as np
import torch
import torch.nn as nn
import torch.nn.functional as F
from copy import deepcopy

# ----------------------------
# Heads
# ----------------------------
class MLPHead(nn.Module):
    def __init__(self, in_dim, out_dim, hidden=(64, 64, 32), dropout=0.2):
        super().__init__()
        self.net = nn.Sequential(
            nn.Linear(in_dim, hidden[0]),
            nn.ReLU(),
            nn.Dropout(dropout),
            nn.Linear(hidden[0], hidden[1]),
            nn.ReLU(),
            nn.Dropout(dropout),
            nn.Linear(hidden[1], out_dim),
        )

    def forward(self, z):
        return self.net(z)


class PerGeneMLPHead(nn.Module):
    """Separate small MLP head per gene/mutation. Returns logits (B, G)."""
    def __init__(self, in_dim, out_dim, hidden=(64, 64, 32), dropout=0.2):
        super().__init__()
        self.out_dim = int(out_dim)
        self.heads = nn.ModuleList([
            MLPHead(in_dim, 1, hidden=hidden, dropout=dropout) for _ in range(self.out_dim)
        ])

    def forward(self, z):
        return torch.cat([h(z) for h in self.heads], dim=1)


# ----------------------------
# Helpers
# ----------------------------
def _as_modality_key(modality):
    """
    Dataloader may collate strings into list[str] of length B.
    Collapse to a single canonical modality key.
    """
    if isinstance(modality, torch.Tensor):
        modality = modality.detach().cpu().tolist() if modality.numel() > 1 else modality.detach().cpu().item()

    if isinstance(modality, (list, tuple)):
        if len(modality) == 0:
            raise ValueError("Empty modality list from dataloader.")
        return _as_modality_key(modality[0])

    if isinstance(modality, np.generic):
        modality = modality.item()

    if isinstance(modality, (int, np.integer)):
        return "rna" if int(modality) == 0 else ("adt" if int(modality) == 1 else str(modality))

    s = str(modality).strip().lower()
    s = s.replace("mod:", "").replace("modality:", "")

    if s in ("rna", "adt", "atac"):
        return s

    aliases = {
        "gene": "rna",
        "expression": "rna",
        "protein": "adt",
        "antibody": "adt",
        "proteins": "adt",
    }
    return aliases.get(s, s)


def _to_device(x, device):
    if isinstance(x, torch.Tensor):
        return x.to(device, non_blocking=True)
    if isinstance(x, np.ndarray):
        return torch.from_numpy(x).to(device, non_blocking=True)
    try:
        return torch.tensor(x, device=device)
    except Exception as e:
        raise TypeError(f"Unsupported batch type for x: {type(x)}") from e


def freeze_decoders(model: nn.Module, freeze=True):
    """Freeze any parameter whose name contains 'decoder'."""
    for name, p in model.named_parameters():
        if "decoder" in name.lower():
            p.requires_grad = (not freeze)


def latent_mu_student(model, x, modality: str):
    """
    Student path: MUST allow grads during finetuning.
    """
    mu_dict, _ = model.encode_modalities({modality: x})
    if modality not in mu_dict:
        raise KeyError(f"Student encode_modalities missing key '{modality}'. Got {list(mu_dict.keys())}")
    return mu_dict[modality]


@torch.no_grad()
def latent_mu_teacher(model, x, modality: str):
    """
    Teacher path: no grads.
    """
    mu_dict, _ = model.encode_modalities({modality: x})
    if modality not in mu_dict:
        raise KeyError(f"Teacher encode_modalities missing key '{modality}'. Got {list(mu_dict.keys())}")
    return mu_dict[modality]


def masked_bce_sum_and_denom(logits, y, m, pos_weight=None):
    """
    Returns (loss_sum, denom) for correct aggregation across batches/loaders.
    """
    y = y.float()
    m = m.float()

    # sanitize unlabeled entries
    y = torch.where(m > 0, y, torch.zeros_like(y)).clamp(0.0, 1.0)

    if pos_weight is not None:
        pw = pos_weight.view(1, -1).to(logits.device)
        loss = F.binary_cross_entropy_with_logits(logits, y, reduction="none", pos_weight=pw)
    else:
        loss = F.binary_cross_entropy_with_logits(logits, y, reduction="none")

    loss = loss * m
    denom = m.sum()
    return loss.sum(), denom


def masked_bce_with_logits_weighted(logits, y, m, pos_weight=None, eps=1e-8):
    s, d = masked_bce_sum_and_denom(logits, y, m, pos_weight=pos_weight)
    return s / (torch.clamp(d, min=1.0) + eps)


# ----------------------------
# Main finetune
# ----------------------------
def finetune_encoders_and_head(
    base_model,
    train_loaders,
    val_loaders,
    *,
    out_dim,
    genes,
    device,
    # optimization
    lr_head=2e-4,
    lr_encoder=None,
    weight_decay=1e-6,
    max_epochs=400,
    patience=40,
    grad_clip=5.0,
    # schedule
    warmup_epochs=10,
    lambda_preserve=1.0,
    use_per_gene_heads=True,
    # class imbalance
    pos_weight=None,   # torch tensor (G,)
    # logging
    verbose=True,
    log_every=1,
    # selection behavior
    start_best_after_unfreeze=False,
):
    if lr_encoder is None:
        lr_encoder = lr_head * 0.1

    # student + teacher
    model_ft = deepcopy(base_model).to(device)
    teacher  = deepcopy(base_model).to(device)
    teacher.eval()
    for p in teacher.parameters():
        p.requires_grad = False

    # freeze student decoders always
    for p in model_ft.parameters():
        p.requires_grad = True
    freeze_decoders(model_ft, freeze=True)

    if pos_weight is not None:
        pos_weight = pos_weight.to(device)
        if pos_weight.numel() != int(out_dim):
            raise ValueError(f"pos_weight has shape {tuple(pos_weight.shape)} but out_dim={out_dim}")

    # iterators + stash so we don't drop the first batch
    train_iters = [iter(L) for L in train_loaders]
    stash = [None for _ in train_loaders]
    steps = max(len(L) for L in train_loaders)

    def next_batch(i):
        nonlocal train_iters, stash
        if stash[i] is not None:
            b = stash[i]
            stash[i] = None
            return b
        try:
            return next(train_iters[i])
        except StopIteration:
            train_iters[i] = iter(train_loaders[i])
            return next(train_iters[i])

    # build head after we infer latent dim
    head = None
    opt = None

    def build_head(latent_dim):
        if use_per_gene_heads:
            return PerGeneMLPHead(latent_dim, out_dim, hidden=(64, 32), dropout=0.1).to(device)
        return MLPHead(latent_dim, out_dim, hidden=(64, 32), dropout=0.1).to(device)

    def set_train_mode(warmup: bool):
        # encoder off during warmup, on after; head always on
        if warmup:
            for p in model_ft.parameters():
                p.requires_grad = False
            freeze_decoders(model_ft, freeze=True)
        else:
            for p in model_ft.parameters():
                p.requires_grad = True
            freeze_decoders(model_ft, freeze=True)

        for p in head.parameters():
            p.requires_grad = True

    def make_optimizer(warmup: bool):
        if warmup:
            return torch.optim.AdamW([{"params": head.parameters(), "lr": lr_head}],
                                     weight_decay=weight_decay)
        enc_params = [p for p in model_ft.parameters() if p.requires_grad]
        return torch.optim.AdamW(
            [{"params": enc_params, "lr": lr_encoder},
             {"params": head.parameters(), "lr": lr_head}],
            weight_decay=weight_decay
        )

    # best tracking
    best = {"score": np.inf, "epoch": -1, "model_state": None, "head_state": None}
    bad = 0

    if verbose:
        print(
            f"[FT] start: out_dim={out_dim} heads={'per_gene' if use_per_gene_heads else 'shared'} "
            f"lr_head={lr_head:g} lr_enc={lr_encoder:g} wd={weight_decay:g} "
            f"lambda_preserve={lambda_preserve:g} warmup={warmup_epochs} "
            f"max_epochs={max_epochs} patience={patience}"
        )
        if pos_weight is not None:
            pw = pos_weight.detach().cpu().numpy()
            print("[FT] pos_weight:", {g: float(pw[i]) for i, g in enumerate(genes)})

    for epoch in range(max_epochs):
        warmup = (epoch < int(warmup_epochs))

        # init head + optimizer once
        if head is None:
            b0 = next_batch(0)
            stash[0] = b0
            modality0, x0, y0, m0 = b0
            mod0 = _as_modality_key(modality0)
            x0 = _to_device(x0, device)

            with torch.no_grad():
                z0 = latent_mu_teacher(model_ft, x0, mod0)
            head = build_head(int(z0.shape[1]))

            set_train_mode(warmup=warmup)
            opt = make_optimizer(warmup=warmup)
            if verbose:
                print(f"[FT] phase -> {'warmup(head-only)' if warmup else 'finetune(enc+head)'}")

        # boundary: rebuild optimizer after unfreeze
        if epoch == int(warmup_epochs):
            set_train_mode(warmup=False)
            opt = make_optimizer(warmup=False)
            if verbose:
                print("[FT] phase -> finetune(enc+head)")

        model_ft.train(); head.train()

        tr_losses, tr_cls, tr_pres = [], [], []

        for s in range(steps):
            for i in range(len(train_loaders)):
                modality, x, y, m = next_batch(i)
                mod = _as_modality_key(modality)

                x = _to_device(x, device)
                y = y.to(device, non_blocking=True)
                m = m.to(device, non_blocking=True)

                opt.zero_grad(set_to_none=True)

                z  = latent_mu_student(model_ft, x, mod)   # grads ON
                lg = head(z)

                loss_cls = masked_bce_with_logits_weighted(lg, y, m, pos_weight=pos_weight)

                if lambda_preserve and lambda_preserve > 0:
                    zt = latent_mu_teacher(teacher, x, mod)  # grads OFF
                    loss_pres = torch.mean((z - zt) ** 2)
                    loss = loss_cls + float(lambda_preserve) * loss_pres
                else:
                    loss_pres = None
                    loss = loss_cls

                loss.backward()

                if grad_clip and grad_clip > 0:
                    params_to_clip = []
                    for pg in opt.param_groups:
                        params_to_clip.extend([p for p in pg["params"] if p.grad is not None])
                    torch.nn.utils.clip_grad_norm_(params_to_clip, grad_clip)

                opt.step()

                tr_losses.append(float(loss.detach().cpu().item()))
                tr_cls.append(float(loss_cls.detach().cpu().item()))
                if loss_pres is not None:
                    tr_pres.append(float(loss_pres.detach().cpu().item()))

        tr  = float(np.mean(tr_losses)) if tr_losses else np.nan
        trc = float(np.mean(tr_cls)) if tr_cls else np.nan
        trp = float(np.mean(tr_pres)) if tr_pres else 0.0

        # ---- val (correct aggregation) ----
        model_ft.eval(); head.eval()
        loss_sum = 0.0
        denom_sum = 0.0

        with torch.no_grad():
            for L in val_loaders:
                for modality, x, y, m in L:
                    mod = _as_modality_key(modality)
                    x = _to_device(x, device)
                    y = y.to(device, non_blocking=True)
                    m = m.to(device, non_blocking=True)

                    z = latent_mu_teacher(model_ft, x, mod)  # no grads in val
                    lg = head(z)
                    s_loss, s_denom = masked_bce_sum_and_denom(lg, y, m, pos_weight=pos_weight)
                    loss_sum  += float(s_loss.detach().cpu().item())
                    denom_sum += float(s_denom.detach().cpu().item())

        va = loss_sum / max(denom_sum, 1.0)

        can_update_best = (not start_best_after_unfreeze) or (epoch >= int(warmup_epochs))
        improved = False
        if can_update_best and (va < best["score"] - 1e-6):
            improved = True
            best["score"] = va
            best["epoch"] = epoch
            best["model_state"] = {k: t.detach().cpu().clone() for k, t in model_ft.state_dict().items()}
            best["head_state"]  = {k: t.detach().cpu().clone() for k, t in head.state_dict().items()}
            bad = 0
        else:
            bad += 1

        if verbose and log_every and (epoch % log_every == 0 or improved or epoch == 0):
            star = "*" if improved else " "
            phase = "W" if warmup else "F"
            best_str = f"{best['score']:.4f}@{best['epoch']:03d}" if best["epoch"] >= 0 else "NA"
            if lambda_preserve and lambda_preserve > 0:
                print(f"[FT]{star} {phase} ep{epoch:03d} tr={tr:.4f} (cls={trc:.4f},pres={trp:.4f}) va={va:.4f} best={best_str} bad={bad}/{patience}")
            else:
                print(f"[FT]{star} {phase} ep{epoch:03d} tr={tr:.4f} va={va:.4f} best={best_str} bad={bad}/{patience}")

        if bad >= patience:
            if verbose:
                print(f"[FT] early stop at epoch {epoch} (no improvement for {patience} epochs).")
            break

    # restore best
    if best["model_state"] is not None:
        model_ft.load_state_dict(best["model_state"])
    if best["head_state"] is not None:
        head.load_state_dict(best["head_state"])

    best_summary = {
        "best_val": float(best["score"]) if best["epoch"] >= 0 else float(va),
        "best_epoch": int(best["epoch"]),
        "stopped_epoch": int(epoch),
        "lambda_preserve": float(lambda_preserve),
        "warmup_epochs": int(warmup_epochs),
        "lr_head": float(lr_head),
        "lr_encoder": float(lr_encoder),
        "heads": "per_gene" if use_per_gene_heads else "shared",
        "start_best_after_unfreeze": bool(start_best_after_unfreeze),
    }

    if verbose:
        if best_summary["best_epoch"] >= 0:
            print(f"[FT] done: restored best epoch {best_summary['best_epoch']} (val={best_summary['best_val']:.4f}).")
        else:
            print("[FT] done: no best snapshot recorded.")

    return model_ft, head, best_summary


# %% [code]
from sklearn.metrics import roc_auc_score, average_precision_score

import numpy as np
import torch
from sklearn.metrics import roc_auc_score, average_precision_score

def eval_head(model, head, loader, genes, *, min_labeled=10, extra_diag=True):
    """
    Evaluate mutation head on a loader.

    Returns dict:
      out[gene] = {
        "auc": float or nan,
        "ap": float or nan,
        "ap_baseline": float or nan,   # ~ prevalence
        "n_labeled": int,
        "n_pos": int,
        "n_neg": int,
        "prevalence": float or nan,
        "p_mean": float or nan,
        "p_std": float or nan,
        "frac_p_gt_0p9": float or nan,
        "frac_p_lt_0p1": float or nan,
        "status": str,
      }
    """
    model.eval(); head.eval()

    Ys, Ms, Ps = [], [], []
    with torch.no_grad():
        for modality, x, y, m in loader:
            mod = _as_modality_key(modality)
            x = _to_device(x, device)

            z = latent_mu(model, x, mod)
            logits = head(z)
            p = torch.sigmoid(logits)

            Ys.append(y.detach().cpu().numpy())
            Ms.append(m.detach().cpu().numpy())
            Ps.append(p.detach().cpu().numpy())

    if len(Ys) == 0:
        base = {"auc": np.nan, "ap": np.nan, "ap_baseline": np.nan,
                "n_labeled": 0, "n_pos": 0, "n_neg": 0,
                "prevalence": np.nan, "p_mean": np.nan, "p_std": np.nan,
                "frac_p_gt_0p9": np.nan, "frac_p_lt_0p1": np.nan,
                "status": "empty_loader"}
        return {g: dict(base) for g in genes}

    Y = np.vstack(Ys)
    M = np.vstack(Ms).astype(bool)
    P = np.vstack(Ps)

    out = {}
    for j, g in enumerate(genes):
        mask = M[:, j]
        n = int(mask.sum())

        if n == 0:
            out[g] = {
                "auc": np.nan, "ap": np.nan, "ap_baseline": np.nan,
                "n_labeled": 0, "n_pos": 0, "n_neg": 0,
                "prevalence": np.nan,
                "p_mean": np.nan, "p_std": np.nan,
                "frac_p_gt_0p9": np.nan, "frac_p_lt_0p1": np.nan,
                "status": "too_few_labeled",
            }
            continue

        yy = Y[mask, j].astype(int)
        pp = P[mask, j].astype(float)

        n_pos = int((yy == 1).sum())
        n_neg = int((yy == 0).sum())
        prev = float(n_pos / max(n, 1))

        # prediction diagnostics
        p_mean = float(np.mean(pp))
        p_std  = float(np.std(pp))
        frac_hi = float(np.mean(pp > 0.9))
        frac_lo = float(np.mean(pp < 0.1))

        if n < min_labeled:
            out[g] = {
                "auc": np.nan, "ap": np.nan, "ap_baseline": prev,
                "n_labeled": n, "n_pos": n_pos, "n_neg": n_neg,
                "prevalence": prev,
                "p_mean": p_mean, "p_std": p_std,
                "frac_p_gt_0p9": frac_hi, "frac_p_lt_0p1": frac_lo,
                "status": f"too_few_labeled(n<{min_labeled})",
            }
            continue

        if (n_pos == 0) or (n_neg == 0):
            out[g] = {
                "auc": np.nan, "ap": np.nan, "ap_baseline": prev,
                "n_labeled": n, "n_pos": n_pos, "n_neg": n_neg,
                "prevalence": prev,
                "p_mean": p_mean, "p_std": p_std,
                "frac_p_gt_0p9": frac_hi, "frac_p_lt_0p1": frac_lo,
                "status": f"one_class(pos={n_pos},neg={n_neg})",
            }
            continue

        out[g] = {
            "auc": float(roc_auc_score(yy, pp)),
            "ap":  float(average_precision_score(yy, pp)),
            "ap_baseline": prev,  # random AP ≈ prevalence
            "n_labeled": n,
            "n_pos": n_pos,
            "n_neg": n_neg,
            "prevalence": prev,
            "p_mean": p_mean,
            "p_std": p_std,
            "frac_p_gt_0p9": frac_hi,
            "frac_p_lt_0p1": frac_lo,
            "status": "ok",
        }

    return out


# %% [code]
import numpy as np
import pandas as pd
import re

def choose_vg_splits_auto(
    vg_adata,
    Y_vg,
    M_vg,
    GENES,
    *,
    preferred_group_col="patient",
    seed0=1,
    tries=500,
    min_test_labeled=10,
    frac_train=0.80,
    frac_val=0.10,
):
    """
    Returns:
      vg_tr, vg_va, vg_te : np.ndarray[int] indices into vg_adata (cell indices)
      vg_split_mode       : str describing which grouping strategy was used

    Labeled cell definition:
      labeled if M_vg[i, :].sum() > 0  (i.e., at least one gene label present)

    Strategy:
      1) If preferred_group_col exists -> group split by that.
      2) Else try to infer a patient/group column from common names.
      3) Else try parsing patient-ish token from sample_id / obs_names.
      4) Else fallback to random split.
    """
    n = vg_adata.n_obs
    if Y_vg is None or M_vg is None:
        raise ValueError("Y_vg and M_vg must be provided (even if used only for labeled-count checks).")
    if len(M_vg) != n:
        raise ValueError(f"M_vg has {len(M_vg)} rows but vg_adata has n_obs={n}.")

    # --- labeled cells: at least one label present in mask
    if hasattr(M_vg, "toarray"):  # sparse
        labeled = np.asarray(M_vg.sum(axis=1)).ravel() > 0
    else:
        labeled = np.asarray(M_vg).sum(axis=1) > 0
    n_labeled = int(labeled.sum())

    # --- helpers
    def _as_series(x):
        if isinstance(x, pd.Series):
            return x
        return pd.Series(np.asarray(x), index=vg_adata.obs_names)

    def _try_get_group_series():
        obs = vg_adata.obs

        # 1) preferred column
        if preferred_group_col in obs.columns:
            s = _as_series(obs[preferred_group_col])
            if s.notna().any():
                return s.astype(str), f"group:{preferred_group_col}"

        # 2) common alternatives
        for col in ["patient", "donor", "subject", "individual", "pt", "patient_id",
                    "donor_id", "subject_id", "orig.ident", "orig_ident", "sample",
                    "sample_id", "library_id", "batch", "study", "dataset"]:
            if col in obs.columns:
                s = _as_series(obs[col])
                if s.notna().any():
                    return s.astype(str), f"group:{col}"

        # 3) parse from sample_id if present
        for col in ["sample_id", "sample", "orig.ident", "orig_ident", "library_id"]:
            if col in obs.columns:
                raw = _as_series(obs[col]).astype(str)

                def parse_token(v):
                    # take leading token before common separators; keeps things stable
                    parts = re.split(r"[|,;/\s]+", v)
                    return parts[0] if parts and parts[0] != "" else v

                parsed = raw.map(parse_token)
                if parsed.nunique() > 1:
                    return parsed, f"parsed:{col}"

        # 4) parse from obs_names
        raw = pd.Series(vg_adata.obs_names.astype(str), index=vg_adata.obs_names)

        def parse_from_name(v):
            # common patterns: PATIENT_* , patient-*, PT123_* , AML556.D0 etc.
            # we take the first chunk before '_' or '-'
            parts = re.split(r"[_-]+", v)
            return parts[0] if parts and parts[0] != "" else v

        parsed = raw.map(parse_from_name)
        if parsed.nunique() > 1:
            return parsed, "parsed:obs_names"

        return None, "random"

    def _group_split_indices(group_s, rng, frac_train, frac_val):
        # group_s: pd.Series of group labels length n
        groups = group_s.values
        uniq = np.unique(groups)
        rng.shuffle(uniq)

        n_g = len(uniq)
        n_tr_g = max(1, int(round(frac_train * n_g)))
        n_va_g = max(1, int(round(frac_val * n_g)))
        # ensure non-empty test groups if possible
        if n_tr_g + n_va_g >= n_g and n_g >= 3:
            n_va_g = max(1, n_g - n_tr_g - 1)

        tr_groups = set(uniq[:n_tr_g])
        va_groups = set(uniq[n_tr_g:n_tr_g + n_va_g])
        te_groups = set(uniq[n_tr_g + n_va_g:])

        idx = np.arange(n)
        tr = idx[np.isin(groups, list(tr_groups))]
        va = idx[np.isin(groups, list(va_groups))]
        te = idx[np.isin(groups, list(te_groups))]

        # guard in case tiny group count collapses
        if len(te) == 0:
            # steal from val if needed
            if len(va) > 0:
                cut = max(1, int(0.5 * len(va)))
                te = va[:cut]
                va = va[cut:]
            else:
                # ultimate fallback: random 10% test
                rng.shuffle(idx)
                te = idx[: max(1, int(0.10 * n))]
                va = idx[max(1, int(0.10 * n)) : max(2, int(0.20 * n))]
                tr = idx[max(2, int(0.20 * n)) :]

        return tr, va, te

    def _random_split_indices(rng, frac_train, frac_val):
        idx = np.arange(n)
        rng.shuffle(idx)
        n_tr = int(round(frac_train * n))
        n_va = int(round(frac_val * n))
        tr = idx[:n_tr]
        va = idx[n_tr:n_tr + n_va]
        te = idx[n_tr + n_va:]
        return tr, va, te

    # --- main: pick a grouping strategy, then retry to meet min_test_labeled
    group_s, mode = _try_get_group_series()

    best = None
    best_score = -1

    for t in range(tries):
        rng = np.random.default_rng(seed0 + t)

        if group_s is not None and mode != "random":
            tr, va, te = _group_split_indices(group_s, rng, frac_train, frac_val)
        else:
            tr, va, te = _random_split_indices(rng, frac_train, frac_val)

        test_labeled = int(labeled[te].sum())
        # prefer higher labeled in test; but accept if meets threshold
        if test_labeled > best_score:
            best_score = test_labeled
            best = (tr, va, te)

        if test_labeled >= min_test_labeled:
            return tr, va, te, mode

    # If we get here, we failed to hit the min_test_labeled threshold.
    # Return the best attempt we saw, but be explicit in the mode.
    tr, va, te = best
    warn_mode = mode + f" (best_effort: test_labeled={best_score}/{n_labeled}, min_required={min_test_labeled})"
    return tr, va, te, warn_mode


# %% [code]
# ============================================================
# Balanced / constrained split search (group-aware) with fallbacks
#   What this does:
#     1) Builds Y,M labels from either:
#          - MutTranscripts / WtTranscripts string columns (VG-style), OR
#          - explicit obs columns (DAb-style), including fuzzy auto-mapping
#     2) Searches for group-aware splits satisfying HARD mins on TRAIN/VAL/TEST
#     3) If impossible, falls back in a controlled way:
#          A) best-effort group-aware split (maximizes labeled + balance)
#          B) if that fails, cell-level stratified split (optional)
# ============================================================

# ----------------------------
# Pretty printing
# ----------------------------
def _summarize_split(name, idx, Y, M, genes):
    idx = np.asarray(idx, dtype=int)
    print(f"[{name}] n={len(idx):,}")
    for j, g in enumerate(genes):
        lab = M[idx, j] > 0
        n_lab = int(lab.sum())
        if n_lab == 0:
            print(f"  {g:<8}: n_labeled=0")
            continue
        yy = Y[idx, j][lab]
        n_pos = int((yy > 0.5).sum())
        n_neg = int((yy <= 0.5).sum())
        prev = n_pos / max(n_lab, 1)
        print(f"  {g:<8}: n_labeled={n_lab:5d}  n_pos={n_pos:5d}  n_neg={n_neg:5d}  prev={prev:0.3f}")
    print()


# ============================================================
# 1) Label builders
# ============================================================

def _compile_gene_pat(g):
    # whole token match (robust to punctuation + underscores)
    g = re.escape(str(g).upper())
    return re.compile(rf"(?<![A-Z0-9]){g}(?![A-Z0-9])", flags=re.IGNORECASE)


def build_YM_from_mut_wt_strings(
    adata,
    genes,
    *,
    mut_col="MutTranscripts",
    wt_col="WtTranscripts",
    conflict_policy="mut_wins",  # "mut_wins" or "na"
):
    """
    VG-style.
    M=1 if either mut or wt mentions the gene; else 0.
    Y=1 if mut mentions gene; else 0 when labeled.
    """
    if mut_col not in adata.obs.columns:
        raise KeyError(f"mut_col={mut_col!r} not in adata.obs.columns")
    if wt_col not in adata.obs.columns:
        raise KeyError(f"wt_col={wt_col!r} not in adata.obs.columns")

    mut_s = adata.obs[mut_col].astype("string").fillna("").str.upper()
    wt_s  = adata.obs[wt_col].astype("string").fillna("").str.upper()

    N = adata.n_obs
    G = len(genes)
    Y = np.zeros((N, G), dtype=np.float32)
    M = np.zeros((N, G), dtype=np.float32)

    for j, g in enumerate(genes):
        pat = _compile_gene_pat(g)
        in_mut = mut_s.str.contains(pat, regex=True).to_numpy()
        in_wt  = wt_s.str.contains(pat, regex=True).to_numpy()

        conflict = in_mut & in_wt
        labeled  = in_mut | in_wt

        if conflict.any():
            if conflict_policy == "mut_wins":
                pass
            elif conflict_policy == "na":
                labeled = labeled & (~conflict)
                in_mut  = in_mut & (~conflict)
            else:
                raise ValueError("conflict_policy must be 'mut_wins' or 'na'")

        M[:, j] = labeled.astype(np.float32)
        Y[:, j] = in_mut.astype(np.float32)

    return Y, M


def _normalize_token(s: str) -> str:
    s = str(s).upper()
    # remove common separators
    s = re.sub(r"[\s\-_:/]+", "", s)
    return s


def infer_obs_col_map_by_gene_substring(adata, genes, *, prefer=None):
    """
    DAb-style helper: try to auto-map each gene to an obs column whose name contains it.
    Examples:
      gene='DNMT3A' matches 'DNMT3A R882H'
      gene='FLT3'   matches 'FLT3-ITD'
      gene='NPM1'   matches 'NPM1 W288fs'
    If multiple matches, pick:
      - any column that also contains one of `prefer` tokens (e.g. ['ITD','R882','W288'])
      - otherwise shortest column name
    Returns dict {gene: col} for genes where a match was found.
    """
    cols = list(adata.obs.columns)
    cols_norm = [_normalize_token(c) for c in cols]
    prefer = [p.upper() for p in (prefer or [])]

    out = {}
    for g in genes:
        g_norm = _normalize_token(g)
        hits = [cols[i] for i, cn in enumerate(cols_norm) if g_norm in cn]
        if len(hits) == 0:
            continue
        if len(hits) == 1:
            out[g] = hits[0]
            continue

        # prefer ones that contain any preferred token
        if prefer:
            hits2 = []
            for c in hits:
                cn = _normalize_token(c)
                if any(_normalize_token(p) in cn for p in prefer):
                    hits2.append(c)
            if len(hits2) == 1:
                out[g] = hits2[0]
                continue
            if len(hits2) > 1:
                hits = hits2

        # choose shortest (often the "main" mutation column)
        hits = sorted(hits, key=lambda x: len(str(x)))
        out[g] = hits[0]
    return out


def build_YM_from_obs_cols(
    adata,
    genes,
    *,
    col_map=None,          # dict {gene: obs_col}
    col_pattern=None,      # format string using {gene} or {gene} / {gene} replacement
    threshold=0.5,
    treat_nonzero_as_pos=True,
    allow_auto_map=True,   # if True and col_map/col_pattern missing, try substring map
    auto_map_prefer=None,  # optional list of tokens to break ties
):
    """
    DAb-style. Reads values from .obs columns.
      - labeled if value is not NA
      - positive if (value != 0) when treat_nonzero_as_pos else (value > threshold)
    """
    N = adata.n_obs
    G = len(genes)
    Y = np.zeros((N, G), dtype=np.float32)
    M = np.zeros((N, G), dtype=np.float32)

    if col_map is None and col_pattern is None and allow_auto_map:
        col_map = infer_obs_col_map_by_gene_substring(adata, genes, prefer=auto_map_prefer)

    if col_map is None and col_pattern is None:
        raise ValueError("Provide col_map or col_pattern, or set allow_auto_map=True.")

    for j, g in enumerate(genes):
        if col_map is not None:
            col = col_map.get(g, None)
        else:
            col = col_pattern.format(gene=g)

        if col is None or col not in adata.obs.columns:
            continue

        s = adata.obs[col]
        is_lab = ~pd.isna(s)
        vals = pd.to_numeric(s, errors="coerce").fillna(0).to_numpy(dtype=float)

        if treat_nonzero_as_pos:
            y = (vals != 0).astype(np.float32)
        else:
            y = (vals > float(threshold)).astype(np.float32)

        Y[:, j] = y
        M[:, j] = is_lab.astype(np.float32)

    return Y, M


# ============================================================
# 2) Scoring + constraints
# ============================================================

def _score_split(tr, va, te, Y, M, genes, prefer_balance=True, balance_weight=10.0):
    """
    Higher is better.
      - reward more labeled in val/test
      - optionally reward prevalence near 0.5 in val/test
    """
    score = 0.0
    G = len(genes)
    for j in range(G):
        score += float(M[va, j].sum()) + float(M[te, j].sum())

        if prefer_balance:
            for idx in (va, te):
                lab = M[idx, j] > 0
                n_lab = float(lab.sum())
                if n_lab <= 0:
                    continue
                p = float((Y[idx, j][lab] > 0.5).mean())
                score += (1.0 - abs(p - 0.5) / 0.5) * float(balance_weight)

    return float(score)


def _counts_for(idx, j, Y, M):
    lab = M[idx, j] > 0
    n_lab = int(lab.sum())
    if n_lab == 0:
        return 0, 0, 0
    yy = Y[idx, j][lab]
    n_pos = int((yy > 0.5).sum())
    n_neg = int((yy <= 0.5).sum())
    return n_lab, n_pos, n_neg


def _check_constraints(idx, j, Y, M, *, min_labeled, min_pos, min_neg):
    n_lab, n_pos, n_neg = _counts_for(idx, j, Y, M)
    return (n_lab >= min_labeled) and (n_pos >= min_pos) and (n_neg >= min_neg)


# ============================================================
# 3) Splitters
# ============================================================

def _group_split_indices(groups, g_tr, g_va, g_te):
    groups = np.asarray(groups).astype(str)
    tr = np.where(np.isin(groups, list(g_tr)))[0]
    va = np.where(np.isin(groups, list(g_va)))[0]
    te = np.where(np.isin(groups, list(g_te)))[0]
    return tr, va, te


def choose_splits_balanced_groupaware(
    n_cells,
    groups,
    Y,
    M,
    genes,
    *,
    seed0=0,
    tries=5000,
    frac=(0.70, 0.15, 0.15),

    # HARD constraints for TRAIN/VAL/TEST
    min_train_labeled=10,
    min_val_labeled=10,
    min_test_labeled=10,
    min_train_pos=1,
    min_train_neg=1,
    min_val_pos=1,
    min_val_neg=1,
    min_test_pos=1,
    min_test_neg=1,

    enforce_genes=None,
    prefer_balance=True,
    balance_weight=10.0,
    verbose=True,

    # fallback behavior inside group-aware
    soft_fallback=True,
):
    """
    Group-aware random search. Returns either:
      - HARD split satisfying all constraints for enforce_genes, OR
      - best-effort group-aware split (if soft_fallback=True)
    """
    Y = np.asarray(Y)
    M = (np.asarray(M) > 0).astype(np.int8)

    n_cells = int(n_cells)
    groups = np.asarray(groups).astype(str)
    uniq = np.unique(groups)

    gene_to_j = {g: j for j, g in enumerate(genes)}
    if enforce_genes is None:
        enforce_genes = list(genes)
    enforce_js = [gene_to_j[g] for g in enforce_genes if g in gene_to_j]

    ntr = max(1, int(round(frac[0] * len(uniq))))
    nva = max(1, int(round(frac[1] * len(uniq))))
    rng = np.random.RandomState(seed0)

    best_hard = None
    best_hard_score = -np.inf
    best_soft = None
    best_soft_score = -np.inf

    for _ in range(int(tries)):
        perm = uniq.copy()
        rng.shuffle(perm)
        g_tr = set(perm[:ntr])
        g_va = set(perm[ntr:ntr+nva])
        g_te = set(perm[ntr+nva:])

        tr, va, te = _group_split_indices(groups, g_tr, g_va, g_te)
        if len(tr) == 0 or len(va) == 0 or len(te) == 0:
            continue

        sc = _score_split(tr, va, te, Y, M, genes, prefer_balance=prefer_balance, balance_weight=balance_weight)
        if sc > best_soft_score:
            best_soft_score = sc
            best_soft = (tr, va, te)

        ok = True
        for j in enforce_js:
            if not _check_constraints(tr, j, Y, M, min_labeled=min_train_labeled, min_pos=min_train_pos, min_neg=min_train_neg):
                ok = False; break
            if not _check_constraints(va, j, Y, M, min_labeled=min_val_labeled, min_pos=min_val_pos, min_neg=min_val_neg):
                ok = False; break
            if not _check_constraints(te, j, Y, M, min_labeled=min_test_labeled, min_pos=min_test_pos, min_neg=min_test_neg):
                ok = False; break
        if not ok:
            continue

        if sc > best_hard_score:
            best_hard_score = sc
            best_hard = (tr, va, te)

    used_fallback = False
    if best_hard is not None:
        tr, va, te = best_hard
        final_score = best_hard_score
        msg = "group_level | HARD constraints satisfied"
    else:
        if not soft_fallback or best_soft is None:
            raise RuntimeError("No group-aware split found (even best-effort).")
        tr, va, te = best_soft
        final_score = best_soft_score
        used_fallback = True
        msg = "group_level | FALLBACK (no hard split found)"

    info = dict(
        split_mode="group_level",
        used_fallback=bool(used_fallback),
        score=float(final_score),
        tries=int(tries),
        frac=tuple(frac),
        enforce_genes=list(enforce_genes),
        constraints=dict(
            min_train_labeled=min_train_labeled, min_train_pos=min_train_pos, min_train_neg=min_train_neg,
            min_val_labeled=min_val_labeled,     min_val_pos=min_val_pos,     min_val_neg=min_val_neg,
            min_test_labeled=min_test_labeled,   min_test_pos=min_test_pos,   min_test_neg=min_test_neg,
        ),
    )

    if verbose:
        print(f"Split mode: {msg} | score={final_score:0.2f}")
        _summarize_split("train", tr, Y, M, genes)
        _summarize_split("val",   va, Y, M, genes)
        _summarize_split("test",  te, Y, M, genes)

    return tr, va, te, info


def choose_splits_cell_level_stratified(
    n_cells,
    Y,
    M,
    genes,
    *,
    seed0=0,
    frac=(0.70, 0.15, 0.15),
    strat_gene=None,      # gene name, e.g. "NPM1" (uses labeled cells only for strat)
    verbose=True,
):
    """
    Cell-level fallback. If strat_gene is provided, we stratify by (Y for that gene) among labeled cells.
    Unlabeled cells are assigned randomly preserving overall frac.
    """
    rng = np.random.RandomState(seed0)
    n_cells = int(n_cells)

    idx_all = np.arange(n_cells, dtype=int)
    rng.shuffle(idx_all)

    ntr = int(round(frac[0] * n_cells))
    nva = int(round(frac[1] * n_cells))
    tr = idx_all[:ntr]
    va = idx_all[ntr:ntr+nva]
    te = idx_all[ntr+nva:]

    info = dict(split_mode="cell_level", strat_gene=strat_gene, frac=tuple(frac), seed=int(seed0))

    if verbose:
        print("Split mode: cell_level | RANDOM" + (f" (strat_gene={strat_gene})" if strat_gene else ""))
        _summarize_split("train", tr, Y, M, genes)
        _summarize_split("val",   va, Y, M, genes)
        _summarize_split("test",  te, Y, M, genes)

    return tr, va, te, info


# ============================================================
# 4) One wrapper that tries group-aware first, then falls back
# ============================================================
def make_balanced_splits_for_adata(
    adata,
    genes,
    *,
    # grouping
    group_col=None,             # e.g. "patient" or "experiment"
    allow_cell_level_fallback=True,

    # label building
    label_mode="mut_wt_strings",     # "mut_wt_strings" | "obs_cols"
    mut_col="MutTranscripts",
    wt_col="WtTranscripts",
    conflict_policy="mut_wins",

    # obs-cols labeling (DAb)
    obs_col_map=None,
    obs_col_pattern=None,
    allow_auto_map=True,
    auto_map_prefer=None,
    treat_nonzero_as_pos=True,
    threshold=0.5,

    # search
    tries=5000,
    frac=(0.70, 0.15, 0.15),
    seed0=0,

    # constraints
    min_train_labeled=10,
    min_val_labeled=10,
    min_test_labeled=10,
    min_train_pos=1,
    min_train_neg=1,
    min_val_pos=1,
    min_val_neg=1,
    min_test_pos=1,
    min_test_neg=1,

    enforce_genes=None,
    prefer_balance=True,
    balance_weight=10.0,
    verbose=True,

    # fallbacks
    soft_fallback=True,         # within group-aware: return best-effort if hard impossible
    cell_level_strat_gene=None, # if we must do cell-level fallback, optional gene to stratify on
):
    # ----- build Y,M -----
    if label_mode == "mut_wt_strings":
        Y, M = build_YM_from_mut_wt_strings(
            adata, genes, mut_col=mut_col, wt_col=wt_col, conflict_policy=conflict_policy
        )
    elif label_mode == "obs_cols":
        Y, M = build_YM_from_obs_cols(
            adata,
            genes,
            col_map=obs_col_map,
            col_pattern=obs_col_pattern,
            threshold=threshold,
            treat_nonzero_as_pos=treat_nonzero_as_pos,
            allow_auto_map=allow_auto_map,
            auto_map_prefer=auto_map_prefer,
        )
    else:
        raise ValueError("label_mode must be 'mut_wt_strings' or 'obs_cols'")

    # ----- attempt group-aware -----
    if group_col is not None:
        if group_col not in adata.obs.columns:
            raise KeyError(f"group_col={group_col!r} not in adata.obs.columns")
        groups = adata.obs[group_col].astype(str).to_numpy()

        try:
            tr, va, te, info = choose_splits_balanced_groupaware(
                adata.n_obs,
                groups,
                Y, M, genes,
                seed0=seed0,
                tries=tries,
                frac=frac,

                min_train_labeled=min_train_labeled,
                min_val_labeled=min_val_labeled,
                min_test_labeled=min_test_labeled,
                min_train_pos=min_train_pos,
                min_train_neg=min_train_neg,
                min_val_pos=min_val_pos,
                min_val_neg=min_val_neg,
                min_test_pos=min_test_pos,
                min_test_neg=min_test_neg,

                enforce_genes=enforce_genes,
                prefer_balance=prefer_balance,
                balance_weight=balance_weight,
                verbose=verbose,
                soft_fallback=soft_fallback,
            )
            info["group_col"] = str(group_col)
            return tr, va, te, Y, M, info

        except RuntimeError as e:
            if not allow_cell_level_fallback:
                raise
            if verbose:
                print(f"[WARN] group-aware split failed: {e}")
                print("[WARN] falling back to cell-level split.")

    # ----- cell-level fallback -----
    tr, va, te, info = choose_splits_cell_level_stratified(
        adata.n_obs,
        Y, M, genes,
        seed0=seed0,
        frac=frac,
        strat_gene=cell_level_strat_gene,
        verbose=verbose,
    )
    return tr, va, te, Y, M, info


# ============================================================
# 5) Example usage
# ============================================================

# --- VG (patient-aware; transcript strings)
# vg_tr, vg_va, vg_te, Y_vg, M_vg, vg_info = make_balanced_splits_for_adata(
#     vg_rna_pp,
#     GENES,
#     group_col="patient",
#     label_mode="mut_wt_strings",
#     mut_col="MutTranscripts",
#     wt_col="WtTranscripts",
#     tries=8000,
#     frac=(0.70, 0.15, 0.15),
#     # with sparse labels, keep mins low or enforce fewer genes
#     min_train_labeled=10, min_train_pos=2, min_train_neg=2,
#     min_val_labeled=4,    min_val_pos=1,   min_val_neg=1,
#     min_test_labeled=4,   min_test_pos=1,  min_test_neg=1,
#     enforce_genes=["NPM1","DNMT3A","FLT3","TP53","NRAS"],  # pick feasible ones
#     soft_fallback=True,
#     allow_cell_level_fallback=True,
#     cell_level_strat_gene="NPM1",
#     seed0=42,
#     verbose=True,
# )

# --- DAb (experiment-aware; obs columns with auto-map)
# dab_tr, dab_va, dab_te, Y_dab, M_dab, dab_info = make_balanced_splits_for_adata(
#     dab_adt_pp,
#     ["NPM1","DNMT3A","FLT3"],
#     group_col="experiment",
#     label_mode="obs_cols",
#     allow_auto_map=True,          # will map to e.g. 'DNMT3A R882H', 'NPM1 W288fs', 'FLT3-ITD'
#     auto_map_prefer=["ITD","R882","W288"],  # optional tie-breakers
#     tries=8000,
#     frac=(0.70, 0.15, 0.15),
#     min_train_labeled=200, min_train_pos=20, min_train_neg=20,
#     min_val_labeled=50,    min_val_pos=5,    min_val_neg=5,
#     min_test_labeled=50,   min_test_pos=5,   min_test_neg=5,
#     enforce_genes=["NPM1","DNMT3A","FLT3"],
#     soft_fallback=True,
#     allow_cell_level_fallback=True,
#     cell_level_strat_gene="NPM1",
#     seed0=42,
#     verbose=True,
# )


# %% [code]
import re
import numpy as np
import pandas as pd

def _compile_gene_pattern(gene: str) -> re.Pattern:
    """
    Match gene as a token-ish substring, case-insensitive.
    Prevents IDH1 matching IDH2 etc. by requiring non-alnum/_ around it.
    """
    g = re.escape(gene.upper())
    # boundaries: not letter/number/underscore on either side
    return re.compile(rf"(?<![A-Z0-9_]){g}(?![A-Z0-9_])", flags=re.IGNORECASE)

def _to_text_series(x):
    """Convert to uppercase string Series, with NaN -> ''."""
    s = x.copy()
    s = s.astype("string")
    s = s.fillna("")
    return s.str.upper()

def mutation_counts_from_transcripts(
    adata,
    genes,
    *,
    mut_col="MutTranscripts",
    wt_col="WtTranscripts",
    prefix="tx_",
    conflict_policy="mut_wins",  # "mut_wins" | "na" | "error"
    write_per_cell_labels=False, # if True, write adata.obs[f"{prefix}{gene}_label"]
):
    """
    Returns:
      summary_df: per-gene counts of Mut / WT / NA and conflicts
      (optionally) writes per-cell labels into adata.obs.
    """
    if mut_col not in adata.obs.columns or wt_col not in adata.obs.columns:
        raise KeyError(f"Need obs columns '{mut_col}' and '{wt_col}'. Found: {list(adata.obs.columns)[:20]} ...")

    mut_s = _to_text_series(adata.obs[mut_col])
    wt_s  = _to_text_series(adata.obs[wt_col])

    n = adata.n_obs
    rows = []

    for gene in genes:
        pat = _compile_gene_pattern(gene)

        in_mut = mut_s.str.contains(pat, regex=True)
        in_wt  = wt_s.str.contains(pat, regex=True)

        conflict = in_mut & in_wt

        if conflict_policy == "mut_wins":
            mut = in_mut
            wt  = (~in_mut) & in_wt
            na  = (~in_mut) & (~in_wt)
        elif conflict_policy == "na":
            mut = in_mut & (~conflict)
            wt  = in_wt  & (~conflict)
            na  = (~mut) & (~wt)  # includes conflicts as NA
        elif conflict_policy == "error":
            if conflict.any():
                idx = adata.obs_names[conflict][0]
                raise ValueError(f"Conflict for {gene}: appears in BOTH {mut_col} and {wt_col} (e.g. cell {idx})")
            mut = in_mut
            wt  = (~in_mut) & in_wt
            na  = (~in_mut) & (~in_wt)
        else:
            raise ValueError("conflict_policy must be 'mut_wins', 'na', or 'error'")

        mut_n = int(mut.sum())
        wt_n  = int(wt.sum())
        na_n  = int(na.sum())
        conf_n = int(conflict.sum())

        rows.append({
            "gene": gene,
            "mut_n": mut_n,
            "wt_n": wt_n,
            "na_n": na_n,
            "conflict_n": conf_n,
            "mut_frac": mut_n / n,
            "wt_frac": wt_n / n,
            "na_frac": na_n / n,
        })

        if write_per_cell_labels:
            col = f"{prefix}{gene}_label"
            lab = np.full(n, "NA", dtype=object)
            lab[wt.to_numpy()] = "WT"
            lab[mut.to_numpy()] = "Mut"
            if conflict_policy == "na":
                lab[conflict.to_numpy()] = "NA_conflict"
            else:
                lab[conflict.to_numpy()] = "Mut_conflict"
            adata.obs[col] = pd.Categorical(lab, categories=["WT", "Mut", "NA", "NA_conflict", "Mut_conflict"])

    summary_df = pd.DataFrame(rows).sort_values("gene").reset_index(drop=True)
    return summary_df


# -------------------------
# Example usage
# -------------------------
GENES = ["NPM1","DNMT3A","FLT3","TP53","NRAS","TET2","IDH2"]

df = mutation_counts_from_transcripts(
    vg_rna_pp,          # or your AnnData
    GENES,
    mut_col="MutTranscripts",
    wt_col="WtTranscripts",
    conflict_policy="mut_wins",
    write_per_cell_labels=False
)

print(df.to_string(index=False, formatters={
    "mut_frac": "{:.3f}".format,
    "wt_frac": "{:.3f}".format,
    "na_frac": "{:.3f}".format,
}))


# %% [code]
# ============================================================
# Run the split functions defined above
# ============================================================

# --- VG (patient-aware; transcript strings)
vg_tr, vg_va, vg_te, Y_vg, M_vg, vg_info = make_balanced_splits_for_adata(
    vg_rna_pp,
    GENES,
    group_col="patient",
    label_mode="mut_wt_strings",
    mut_col="MutTranscripts",     
    wt_col="WtTranscripts",
    tries=8000,
    frac=(0.70, 0.15, 0.15),
    # with sparse labels, keep mins low or enforce fewer genes
    min_train_labeled=10, min_train_pos=2, min_train_neg=2,
    min_val_labeled=4,    min_val_pos=1,   min_val_neg=1,
    min_test_labeled=4,   min_test_pos=1,  min_test_neg=1,
    enforce_genes=["NPM1","DNMT3A","FLT3","TP53","NRAS","IDH2","TET2"],  # pick feasible ones
    soft_fallback=False,
    allow_cell_level_fallback=True,
    cell_level_strat_gene="NPM1",
    seed0=42,
    verbose=True,
)


# %% [code]
# --- DAb (experiment-aware; obs columns with auto-map)
dab_tr, dab_va, dab_te, Y_dab, M_dab, dab_info = make_balanced_splits_for_adata(
    dab_adt_pp,
    ["NPM1","DNMT3A","FLT3"],
    group_col="experiment",
    label_mode="obs_cols",
    allow_auto_map=True,          # will map to e.g. 'DNMT3A R882H', 'NPM1 W288fs', 'FLT3-ITD'
    auto_map_prefer=["ITD","R882","W288"],  # optional tie-breakers
    tries=8000,
    frac=(0.70, 0.15, 0.15),
    min_train_labeled=200, min_train_pos=20, min_train_neg=20,
    min_val_labeled=10,    min_val_pos=5,    min_val_neg=5,
    min_test_labeled=50,   min_test_pos=5,   min_test_neg=5,
    enforce_genes=["NPM1","DNMT3A","FLT3"],
    soft_fallback=False,
    allow_cell_level_fallback=True,
    cell_level_strat_gene="NPM1",
    seed0=42,
    verbose=True,
)


# %% [code]
# datasets/loaders (unchanged, but now using the new VG splits)
vg_ds  = SingleModalDataset(vg_rna_pp,  "rna", Y_vg,  M_vg)
dab_ds = SingleModalDataset(dab_adt_pp, "adt", Y_dab, M_dab)

device = 'cuda'

B = 128
pin = (device == "cuda")


# %% [code]
import numpy as np

def pad_YM_to_genes(*, Y, M, genes_in, genes_out):
    """
    Pad/reorder (Y, M) from genes_in -> genes_out.
    Missing genes in genes_in will be filled with Y=0 and M=0 (unlabeled).
    """
    genes_in = list(genes_in)
    genes_out = list(genes_out)

    Y = np.asarray(Y)
    M = np.asarray(M)

    if Y.ndim != 2 or M.ndim != 2:
        raise ValueError(f"Expected 2D arrays, got Y{Y.shape}, M{M.shape}")
    if Y.shape != M.shape:
        raise ValueError(f"Y and M must have same shape, got Y{Y.shape} vs M{M.shape}")
    if Y.shape[1] != len(genes_in):
        raise ValueError(f"Y/M cols ({Y.shape[1]}) must match len(genes_in) ({len(genes_in)})")

    n = Y.shape[0]
    Y_out = np.zeros((n, len(genes_out)), dtype=Y.dtype)
    M_out = np.zeros((n, len(genes_out)), dtype=M.dtype)

    idx_in = {g: j for j, g in enumerate(genes_in)}
    for j_out, g in enumerate(genes_out):
        j_in = idx_in.get(g, None)
        if j_in is None:
            # not present in this dataset -> keep Y=0, M=0 (unlabeled)
            continue
        Y_out[:, j_out] = Y[:, j_in]
        M_out[:, j_out] = M[:, j_in]

    return Y_out, M_out


def summarize_label_coverage(name, Y, M, genes, idx):
    idx = np.asarray(idx, dtype=int)
    Msub = M[idx]
    Ysub = Y[idx]

    print(f"\n[{name}] n={len(idx):,}")
    for j, g in enumerate(genes):
        n_lab = int(Msub[:, j].sum())
        if n_lab == 0:
            print(f"  {g:6s}: n_labeled=0")
        else:
            yy = Ysub[Msub[:, j] > 0, j]
            n_pos = int((yy > 0.5).sum())
            prev = n_pos / max(n_lab, 1)
            print(f"  {g:6s}: n_labeled={n_lab:5d}  n_pos={n_pos:5d}  prev={prev:.3f}")


# ----------------------------
# Example usage for your case:
# ----------------------------
# Global gene order you want everywhere:
GENES = ["NPM1","DNMT3A","FLT3","TP53","NRAS","TET2","IDH2"]

# The genes your DAb Y/M currently correspond to (likely 3):
GENES_DAB = ["NPM1","DNMT3A","FLT3"]

# Pad/reorder DAb to full GENES with M=0 for missing genes
Y_dab, M_dab = pad_YM_to_genes(Y=Y_dab, M=M_dab, genes_in=GENES_DAB, genes_out=GENES)

# Now summaries won't crash and will correctly show n_labeled=0 for missing genes
summarize_label_coverage("VG train",   Y_vg,  M_vg,  GENES, vg_tr)
summarize_label_coverage("VG val",     Y_vg,  M_vg,  GENES, vg_va)
summarize_label_coverage("VG test",    Y_vg,  M_vg,  GENES, vg_te)

summarize_label_coverage("DAb train",  Y_dab, M_dab, GENES, dab_tr)
summarize_label_coverage("DAb val",    Y_dab, M_dab, GENES, dab_va)
summarize_label_coverage("DAb test",   Y_dab, M_dab, GENES, dab_te)


# %% [code]
from torch.utils.data import DataLoader, Subset

# --- sanity: your DAb labels/masks MUST already be padded to len(GENES) ---
# (run your pad_YM_to_genes(...) before this)
assert Y_vg.shape[1]  == len(GENES) and M_vg.shape[1]  == len(GENES),  (Y_vg.shape,  M_vg.shape,  len(GENES))
assert Y_dab.shape[1] == len(GENES) and M_dab.shape[1] == len(GENES), (Y_dab.shape, M_dab.shape, len(GENES))

# --- patch the underlying dataset targets so Subset() yields (B, len(GENES)) ---
# This assumes vg_ds/dab_ds store targets on attributes named Y/M (common in our earlier code).
# If your dataset uses different attribute names, change them here once.
if hasattr(vg_ds, "Y"): vg_ds.Y = Y_vg
if hasattr(vg_ds, "M"): vg_ds.M = M_vg

if hasattr(dab_ds, "Y"): dab_ds.Y = Y_dab
if hasattr(dab_ds, "M"): dab_ds.M = M_dab

# (optional) if your dataset keeps torch tensors cached, you may also want:
# if hasattr(dab_ds, "Y_t"): dab_ds.Y_t = torch.from_numpy(Y_dab).float()
# if hasattr(dab_ds, "M_t"): dab_ds.M_t = torch.from_numpy(M_dab).float()

# --- build loaders ---
vg_train_loader  = DataLoader(Subset(vg_ds,  vg_tr),  batch_size=B, shuffle=True,  num_workers=0, pin_memory=pin)
vg_val_loader    = DataLoader(Subset(vg_ds,  vg_va),  batch_size=B, shuffle=False, num_workers=0, pin_memory=pin)
vg_test_loader   = DataLoader(Subset(vg_ds,  vg_te),  batch_size=B, shuffle=False, num_workers=0, pin_memory=pin)

dab_train_loader = DataLoader(Subset(dab_ds, dab_tr), batch_size=B, shuffle=True,  num_workers=0, pin_memory=pin)
dab_val_loader   = DataLoader(Subset(dab_ds, dab_va), batch_size=B, shuffle=False, num_workers=0, pin_memory=pin)
dab_test_loader  = DataLoader(Subset(dab_ds, dab_te), batch_size=B, shuffle=False, num_workers=0, pin_memory=pin)

print("VG split sizes:",  len(vg_tr),  len(vg_va),  len(vg_te))
print("DAb split sizes:", len(dab_tr), len(dab_va), len(dab_te), "group:", DAB_EXPT_COL)

# --- quick one-batch check (should print (B, len(GENES))) ---
b = next(iter(dab_train_loader))
# adjust keys if your batch is a tuple instead of dict
y_b = b["y"] if isinstance(b, dict) else b[1]
m_b = b["m"] if isinstance(b, dict) else b[2]
print("DAb batch y/m:", tuple(y_b.shape), tuple(m_b.shape), "expected (*,", len(GENES), ")")


# %% [code]
import numpy as np

def _as_modality_key(modality):
    """
    Normalize modality tokens coming from dataloaders into canonical keys.
    Handles batched modality lists produced by DataLoader collate.
    """
    # If batch-collated: modality is usually a list/tuple of identical strings
    if isinstance(modality, (list, tuple)) and len(modality) > 0:
        modality = modality[0]

    # If numpy array of strings (sometimes happens)
    if isinstance(modality, np.ndarray):
        if modality.ndim > 0 and modality.size > 0:
            modality = modality.flat[0]
        else:
            modality = modality.item()

    # torch scalar / numpy scalar -> python scalar
    try:
        import torch
        if isinstance(modality, torch.Tensor):
            # if it's a batch of modality ids, take first
            if modality.numel() > 1:
                modality = modality.flatten()[0].detach().cpu().item()
            else:
                modality = modality.detach().cpu().item()
    except Exception:
        pass

    if isinstance(modality, (np.generic,)):
        modality = modality.item()

    # If dataset returns a dict/batch with modality inside
    if isinstance(modality, dict):
        for k in ("modality", "mod", "mod_key"):
            if k in modality:
                modality = modality[k]
                break

    # ints sometimes used as modality ids
    if isinstance(modality, (int, np.integer)):
        if modality == 0:
            return "rna"
        if modality == 1:
            return "adt"
        return str(modality)

    # strings / objects
    s = str(modality).strip().lower()

    if s in ("rna", "adt", "atac"):
        return s

    aliases = {
        "vg": "rna",
        "vg_rna": "rna",
        "van_galen": "rna",
        "gene": "rna",
        "expression": "rna",

        "dab": "adt",
        "dab_adt": "adt",
        "protein": "adt",
        "proteins": "adt",
        "antibody": "adt",
        "adt_counts": "adt",
    }

    s2 = s.replace("mod:", "").replace("modality:", "")
    return aliases.get(s2, s2)

import torch

def _to_device(x, device):
    """
    Move a tensor (or numpy array) to device with sane defaults.
    - ensures float32
    - keeps non_blocking when CUDA + pinned memory
    """
    if isinstance(x, np.ndarray):
        x = torch.from_numpy(x)

    if not torch.is_tensor(x):
        # last resort: try to tensor it
        x = torch.tensor(x)

    # Make sure dtype is float32 for UniVI encoders
    if x.dtype != torch.float32:
        x = x.float()

    return x.to(device, non_blocking=(str(device).startswith("cuda")))


# %% [code]
import numpy as np
import torch

def compute_pos_weight_from_arrays(Y, M, eps=1e-6, clamp_max=100.0):
    """
    Y, M: numpy arrays shape (N, G)
      - Y: 0/1 labels (or floats where >0.5 treated as positive)
      - M: 0/1 mask for labeledness

    Returns:
      pos_weight: torch.FloatTensor (G,) where pos_weight[j] = n_neg / n_pos
      (clamped to [1.0, clamp_max] to avoid insane values when pos is tiny)
    """
    Y = np.asarray(Y)
    M = np.asarray(M).astype(bool)

    if Y.ndim != 2 or M.ndim != 2 or Y.shape != M.shape:
        raise ValueError(f"Y and M must be same shape (N,G). Got Y{Y.shape}, M{M.shape}")

    N, G = Y.shape
    pw = np.ones(G, dtype=np.float32)

    for j in range(G):
        mask = M[:, j]
        if not np.any(mask):
            pw[j] = 1.0
            continue

        yy = Y[mask, j]
        npos = float((yy > 0.5).sum())
        nneg = float((yy <= 0.5).sum())

        # if all one class, don't blow up training
        if npos < 1.0 or nneg < 1.0:
            pw[j] = 1.0
            continue

        val = (nneg + eps) / (npos + eps)
        # pos_weight < 1 is allowed, but often unstable; clamp to >=1
        val = max(1.0, val)
        if clamp_max is not None:
            val = min(float(clamp_max), val)
        pw[j] = val

    return torch.tensor(pw, dtype=torch.float32)


# %% [code]
# ============================================================
# TRAIN + EVAL
# ============================================================
'''
model_ft, head, best = finetune_encoders_and_head(
    model,
    train_loaders=[vg_train_loader, dab_train_loader],
    val_loaders=[vg_val_loader, dab_val_loader],
    out_dim=len(GENES),
    lr=2e-4,
    max_epochs=400,
    patience=40,
)

print("Fine-tune best:", best)
'''

# build pos_weight from TRAIN labeled cells only
Y_tr = np.vstack([Y_vg[vg_tr],  Y_dab[dab_tr]])
M_tr = np.vstack([M_vg[vg_tr],  M_dab[dab_tr]])

pos_weight = compute_pos_weight_from_arrays(Y_tr, M_tr).to(device)

model_ft, head, best = finetune_encoders_and_head(
    model,
    train_loaders=[vg_train_loader, dab_train_loader],
    val_loaders=[vg_val_loader, dab_val_loader],
    out_dim=len(GENES),
    genes=GENES,
    device=device,
    lr_head=1e-4,
    lr_encoder=1e-5,
    weight_decay=1e-6,
    lambda_preserve=5.0,
    warmup_epochs=10,
    max_epochs=1000,
    patience=50,
    grad_clip=5.0,
    pos_weight=pos_weight,
    use_per_gene_heads=True,
    start_best_after_unfreeze=False,
)


# %% [code]
dab_perf = eval_head(model_ft, head, dab_test_loader, GENES)
vg_perf  = eval_head(model_ft, head, vg_test_loader,  GENES)

print("DAb test:", dab_perf)
print("VG  test:", vg_perf)


# %% [code]
def summarize_ap_vs_prevalence(*, Y, M, te_idx, genes, eval_out, title=""):
    rows = []
    for j, g in enumerate(genes):
        te_mask = np.asarray(M[te_idx, j]).astype(bool)
        n = int(te_mask.sum())
        if n == 0:
            prev = np.nan
            n_pos = 0
        else:
            yy = np.asarray(Y[te_idx, j])[te_mask].astype(int)
            n_pos = int((yy == 1).sum())
            prev = n_pos / n

        ap = eval_out.get(g, {}).get("ap", np.nan)
        auc = eval_out.get(g, {}).get("auc", np.nan)
        status = eval_out.get(g, {}).get("status", "missing")

        fold = (ap / prev) if (np.isfinite(ap) and np.isfinite(prev) and prev > 0) else np.nan

        rows.append({
            "gene": g,
            "status": status,
            "n_labeled": n,
            "n_pos": n_pos,
            "prevalence(AP_random)": prev,
            "AP": ap,
            "AP_over_random": fold,
            "AUC": auc,
        })

    df = pd.DataFrame(rows).sort_values(
        by=["status", "AP_over_random", "AP"], ascending=[True, False, False]
    )

    if title:
        print("\n" + title)
    display(df) if "display" in globals() else print(df.to_string(index=False))
    return df

# Example usage (adapt names to your variables):
# dab_test_out = eval_head(model_ft, head, dab_test_loader, GENES)   # you already did this
# vg_test_out  = eval_head(model_ft, head, vg_test_loader,  GENES)

df_dab = summarize_ap_vs_prevalence(
    Y=Y_dab, M=M_dab, te_idx=dab_te, genes=GENES, eval_out=dab_perf, title="DAb TEST: AP vs prevalence"
)
df_vg = summarize_ap_vs_prevalence(
    Y=Y_vg, M=M_vg, te_idx=vg_te, genes=GENES, eval_out=vg_perf, title="VG TEST: AP vs prevalence"
)

# Quick readout for a single gene (e.g., NPM1)
print("\nVG NPM1 row:")
print(df_vg[df_vg["gene"] == "NPM1"].to_string(index=False))


# %% [code]
from pathlib import Path
import numpy as np
import pandas as pd
import scanpy as sc
import anndata as ad
import matplotlib.pyplot as plt

# ----------------------------
# Config
# ----------------------------
FIGDIR = Path(FIGDIR)
FIGDIR.mkdir(parents=True, exist_ok=True)

# ----------------------------
# 0) Encode FT latents on the SAME objects you put into joint
#    (important: don't encode cite_rna_pp_tr then store into cite_rna_tr)
# ----------------------------
# Choose which CITE split you want in joint; you used *_tr below.
# Here I assume these are the objects you want in joint:
cite_rna_joint = cite_rna_pp_tr      # or cite_rna_pp_tr if you prefer, but be consistent
cite_adt_joint = cite_adt_pp_tr      # or cite_adt_pp_tr
vg_rna_joint   = vg_rna_pp
dab_adt_joint  = dab_adt_pp

# Encode and store
cite_rna_joint.obsm["X_univi_ft"] = encode_latent(model_ft, cite_rna_joint, modality="rna", device=device)
cite_adt_joint.obsm["X_univi_ft"] = encode_latent(model_ft, cite_adt_joint, modality="adt", device=device)
vg_rna_joint.obsm["X_univi_ft"]   = encode_latent(model_ft, vg_rna_joint,   modality="rna", device=device)
dab_adt_joint.obsm["X_univi_ft"]  = encode_latent(model_ft, dab_adt_joint,  modality="adt", device=device)

print("Latents shapes:",
      cite_rna_joint.obsm["X_univi_ft"].shape,
      cite_adt_joint.obsm["X_univi_ft"].shape,
      vg_rna_joint.obsm["X_univi_ft"].shape,
      dab_adt_joint.obsm["X_univi_ft"].shape)


# %% [code]
# ----------------------------
# 1) Build joint AnnData in FT space
# ----------------------------
def joint_latent_adata(items, rep_key="X_univi_ft"):
    Zs, obs_rows = [], []
    for ds_name, mod, a in items:
        Z = a.obsm[rep_key]
        Zs.append(Z)
        obs_rows.append(
            pd.DataFrame({"dataset": ds_name, "modality": mod}, index=a.obs_names.copy())
        )
    Z = np.vstack(Zs).astype(np.float32)
    obs = pd.concat(obs_rows, axis=0)
    out = ad.AnnData(
        X=Z,
        obs=obs,
        var=pd.DataFrame(index=[f"z{i}" for i in range(Z.shape[1])]),
    )
    out.obsm[rep_key] = out.X.copy()
    return out

joint = joint_latent_adata([
    ("CITE", "rna", cite_rna_joint),
    ("CITE", "adt", cite_adt_joint),
    ("DAb",  "adt", dab_adt_joint),
    ("VG",   "rna", vg_rna_joint),
], rep_key="X_univi_ft")

joint.obs["dataset_modality"] = joint.obs["dataset"].astype(str) + " " + joint.obs["modality"].astype(str)


# %% [code]
# ----------------------------------
# 2) Compute joint UMAP in FT space
# ----------------------------------
sc.pp.neighbors(joint, use_rep="X_univi_ft", n_neighbors=30)  # random_state not used in neighbors
sc.tl.umap(joint, random_state=1)


# %% [code]
# ----------------------------------------
# 3) Copy useful metadata to joint object
# ----------------------------------------
def _assign_block(joint, *, dataset, modality, values, colname):
    mask = (joint.obs["dataset"] == dataset) & (joint.obs["modality"] == modality)
    n = int(mask.sum())
    v = np.asarray(values)
    if v.shape[0] != n:
        raise ValueError(f"{colname}: trying to assign {v.shape[0]} values into {n} rows for {dataset} {modality}")
    joint.obs.loc[mask, colname] = v

# ----------------------------
# (A) CITE: sample_id, library_id -> joint (mask-based)
# ----------------------------
if "sample_id" in cite_rna_joint.obs.columns:
    _assign_block(
        joint,
        dataset="CITE", modality="rna",
        values=cite_rna_joint.obs["sample_id"].astype(str).to_numpy(),
        colname="cite_sample_id",
    )
if "library_id" in cite_rna_joint.obs.columns:
    _assign_block(
        joint,
        dataset="CITE", modality="rna",
        values=cite_rna_joint.obs["library_id"].astype(str).to_numpy(),
        colname="cite_library_id",
    )

if "sample_id" in cite_adt_joint.obs.columns:
    _assign_block(
        joint,
        dataset="CITE", modality="adt",
        values=cite_adt_joint.obs["sample_id"].astype(str).to_numpy(),
        colname="cite_sample_id",
    )
if "library_id" in cite_adt_joint.obs.columns:
    _assign_block(
        joint,
        dataset="CITE", modality="adt",
        values=cite_adt_joint.obs["library_id"].astype(str).to_numpy(),
        colname="cite_library_id",
    )

# AML vs Control label (works for both CITE RNA/ADT once cite_sample_id is set)
sid = joint.obs.get("cite_sample_id", pd.Series(pd.NA, index=joint.obs_names)).astype("string")
ac = pd.Series(pd.NA, index=joint.obs_names, dtype="string")
ac.loc[sid.str.contains("aml", case=False, na=False)] = "AML"
ac.loc[sid.str.contains("control", case=False, na=False)] = "Control"
joint.obs["cite_aml_vs_control"] = pd.Categorical(ac, categories=["Control", "AML"], ordered=True)

# ----------------------------
# (B) DAb actual mutation label -> joint (mask-based)
# ----------------------------
HERO = str(HERO)
col_actual_bool = f"actual_{HERO}_mut"
col_actual_lab  = f"actual_{HERO}_dab_label"

if col_actual_bool in dab_adt_al.obs.columns:
    s = dab_adt_al.obs[col_actual_bool].astype("boolean")
    lab = pd.Series(pd.NA, index=dab_adt_al.obs_names, dtype="string")
    lab.loc[s == True]  = "Mut"
    lab.loc[s == False] = "WT"
    lab = pd.Categorical(lab, categories=["WT", "Mut"], ordered=True)

    # align to dab_adt_joint order (important)
    lab_aligned = pd.Series(lab, index=dab_adt_al.obs_names).reindex(dab_adt_joint.obs_names).astype("string").to_numpy()

    _assign_block(
        joint,
        dataset="DAb", modality="adt",
        values=lab_aligned,
        colname=col_actual_lab,
    )
    joint.obs[col_actual_lab] = pd.Categorical(joint.obs[col_actual_lab], categories=["WT", "Mut"], ordered=True)
else:
    print(f"NOTE: {col_actual_bool} not found in dab_adt_al.obs; skipping DAb actual mutation copy.")

# ----------------------------
# (C) VG truth label (if you already made it on vg_rna_joint)
# ----------------------------
col_vg_truth = f"actual_{HERO}_vg_label"
if col_vg_truth in vg_rna_joint.obs.columns:
    _assign_block(
        joint,
        dataset="VG", modality="rna",
        values=vg_rna_joint.obs[col_vg_truth].astype("string").to_numpy(),
        colname=col_vg_truth,
    )
    joint.obs[col_vg_truth] = pd.Categorical(joint.obs[col_vg_truth], categories=["WT", "Mut"], ordered=True)

# ----------------------------
# (D) Optional: copy a probability column from a block into joint
# ----------------------------
def copy_obs_col_block(src_adata, *, dataset, modality, col):
    if col not in src_adata.obs.columns:
        print(f"NOTE: {col} not in src; skipping")
        return
    _assign_block(
        joint,
        dataset=dataset, modality=modality,
        values=src_adata.obs[col].to_numpy(),
        colname=col,
    )

# examples (only if these exist)
copy_obs_col_block(vg_rna_joint,  dataset="VG",   modality="rna", col=f"knnP_{HERO}_dab_to_vg_ft")
copy_obs_col_block(dab_adt_joint, dataset="DAb",  modality="adt", col=f"knnP_{HERO}_vg_to_dab")
copy_obs_col_block(cite_rna_joint, dataset="CITE", modality="rna", col=f"headP_{HERO}_cite_rna")
copy_obs_col_block(cite_adt_joint, dataset="CITE", modality="adt", col=f"headP_{HERO}_cite_adt")


# %% [code]
# ----------------------------
# 4) Plot joint FT UMAP panels
# ----------------------------
# Fig7b: dataset × modality
sc.pl.umap(
    joint,
    color="dataset_modality",
    title="Joint UMAP in FT space (dataset × modality)",
    size=15,
    alpha=0.7,
)
plt.savefig(FIGDIR / "fig7b_umap_dataset_modality_ft.png", dpi=450, bbox_inches="tight")
plt.show()

# CITE: AML vs control
if "cite_aml_vs_control" in joint.obs.columns:
    sc.pl.umap(
        joint,
        color=["cite_sample_id", "cite_aml_vs_control"],
        title=["CITE sample_id", "CITE AML vs Control"],
        na_color="lightgrey",
        wspace=0.35,
        size=15,
        alpha=0.7,
    )
    plt.savefig(FIGDIR / "joint_umap_ft_cite_sample_and_aml_control.png", dpi=450, bbox_inches="tight")
    plt.show()

# DAb: actual mutation label on joint (only DAb points have labels)
if col_actual_lab in joint.obs.columns:
    sc.pl.umap(
        joint,
        color=col_actual_lab,
        title=f"DAb observed {HERO} (WT/Mut) on joint FT UMAP",
        na_color="lightgrey",
        size=15,
        alpha=0.7,
    )
    plt.savefig(FIGDIR / f"joint_umap_ft_dab_actual_{HERO}.png", dpi=450, bbox_inches="tight")
    plt.show()

# Optional: VG truth + VG transferred prob (if present)
cols = []
if col_vg_truth in joint.obs.columns: cols.append(col_vg_truth)
pcol = f"knnP_{HERO}_dab_to_vg_ft"
if pcol in joint.obs.columns: cols.append(pcol)

if len(cols) > 0:
    sc.pl.umap(
        joint,
        color=cols,
        title=[f"VG truth {HERO}" if c==col_vg_truth else f"Transfer P({HERO}) DAb→VG (FT)" for c in cols],
        na_color="lightgrey",
        size=15,
        alpha=0.7,
        wspace=0.35,
    )
    plt.savefig(FIGDIR / f"joint_umap_ft_vg_truth_and_transfer_{HERO}.png", dpi=450, bbox_inches="tight")
    plt.show()


# %% [code]
# pick the VG RNA rows inside joint
if "dataset_modality" in joint.obs.columns:
    vg_mask = (joint.obs["dataset_modality"] == "VG rna")
else:
    vg_mask = (joint.obs["dataset"].astype(str).str.contains("VG")) & (joint.obs["modality"] == "rna")


# %% [code]
cols = ["patient", "CellType", "timepoint"]  # edit names if needed
src = vg_rna_pp.obs[cols].copy()

# align by cell id (= obs_names)
aligned = src.reindex(joint.obs_names)

# write into joint only for VG RNA rows
for c in cols:
    joint.obs.loc[vg_mask, c] = aligned.loc[joint.obs_names[vg_mask], c].values


# %% [code]
sc.pl.umap(
    joint,
    color=['patient'],
    title=[f"Joint fine-tuned latent colored by VG patient"],
    na_color="lightgrey",
    size=15,
    alpha=0.7,
    wspace=0.35,
)

# %% [code]
sc.pl.umap(
    joint,
    color=['CellType'],
    title=[f"Joint fine-tuned latent colored by VG cell type"],
    na_color="lightgrey",
    size=15,
    alpha=0.7,
    wspace=0.35,
)


# %% [code]
print(joint)
print(joint.obs['CellType'])


# %% [code]
import numpy as np
import pandas as pd
from sklearn.neighbors import NearestNeighbors

# =========================
# CONFIG: adjust these keys
# =========================
LATENT_KEY = "X_univi_ft"          # or "X_univi_ft" if you stored FT latents separately
DATASET_COL = "dataset_modality" # or ("dataset","modality") if you have them split
VG_TAG = "VG rna"
DAB_TAG = "DAb adt"

VG_CELLTYPE_COL = "CellType"    # the column you copied from vg_rna_pp.obs into joint.obs

K = 30                          # neighborhood size

# =========================
# Pull arrays + masks
# =========================
Z = np.asarray(joint.obsm[LATENT_KEY])

vg_mask  = (joint.obs[DATASET_COL].astype(str) == VG_TAG).values
dab_mask = (joint.obs[DATASET_COL].astype(str) == DAB_TAG).values

Z_vg  = Z[vg_mask]
Z_dab = Z[dab_mask]

vg_ct = joint.obs.loc[vg_mask, VG_CELLTYPE_COL].astype(str).values

print("VG cells:", Z_vg.shape[0], " DAb cells:", Z_dab.shape[0])
print("VG celltype NA fraction:", np.mean((vg_ct == "NA") | (vg_ct == "nan") | (vg_ct == "None")))

# =========================
# kNN from DAb -> VG
# =========================
knn = NearestNeighbors(n_neighbors=K, metric="euclidean")
knn.fit(Z_vg)
idx = knn.kneighbors(Z_dab, return_distance=False)   # (n_dab, K) indices into VG

# count VG celltypes in each DAb neighborhood
ct_levels = pd.Index(sorted(pd.unique(vg_ct)))
ct_to_i = {c:i for i,c in enumerate(ct_levels)}

counts = np.zeros((idx.shape[0], len(ct_levels)), dtype=np.int32)
for r in range(idx.shape[0]):
    for j in idx[r]:
        counts[r, ct_to_i[vg_ct[j]]] += 1

frac = counts / float(K)

# overall mapping: average fraction of VG celltypes around DAb cells
avg = frac.mean(axis=0)
df = pd.DataFrame({"VG_celltype": ct_levels, "avg_frac_in_DAb_kNN": avg})
df = df.sort_values("avg_frac_in_DAb_kNN", ascending=False)

print("\nTop VG celltypes surrounding DAb (avg over DAb cells):")
print(df.head(15).to_string(index=False))

# optional: how "PB-like" vs "marrow-progenitor-like" is DAb in this latent?
pb_like = set(["T","CTL","NK","B","Mono","Mono-like","cDC","cDC-like","pDC","Plasma","ProB"])
progen_like = set(["HSC","HSC-like","GMP","GMP-like","Prog","Prog-like","earlyEry","lateEry","ProMono","ProMono-like"])

pb_score = df[df["VG_celltype"].isin(pb_like)]["avg_frac_in_DAb_kNN"].sum()
prog_score = df[df["VG_celltype"].isin(progen_like)]["avg_frac_in_DAb_kNN"].sum()

print(f"\nDAb neighbor composition summary (from VG labels):")
print(f"  PB-like fraction ≈ {pb_score:.3f}")
print(f"  Progen/Ery-like fraction ≈ {prog_score:.3f}")


# %% [code]
import numpy as np
import pandas as pd
from sklearn.neighbors import NearestNeighbors

# --- assume you still have: joint, vg_mask, dab_mask, Z_vg, Z_dab, vg_ct, ct_levels, K ---
# If not, just re-run your previous block up to `frac`.

pb_like = set(["T","CTL","NK","B","Mono","Mono-like","cDC","cDC-like","pDC","Plasma","ProB"])
progen_like = set(["HSC","HSC-like","GMP","GMP-like","Prog","Prog-like","earlyEry","lateEry","ProMono","ProMono-like"])

# indices in ct_levels
pb_idx = np.array([i for i,c in enumerate(ct_levels) if c in pb_like], dtype=int)
pr_idx = np.array([i for i,c in enumerate(ct_levels) if c in progen_like], dtype=int)

pb_per_cell = frac[:, pb_idx].sum(axis=1)
pr_per_cell = frac[:, pr_idx].sum(axis=1)

print("DAb PB-like per-cell:  median", float(np.median(pb_per_cell)), "  p10/p90",
      float(np.quantile(pb_per_cell,0.1)), float(np.quantile(pb_per_cell,0.9)))
print("DAb Progen/Ery per-cell: median", float(np.median(pr_per_cell)), "  p10/p90",
      float(np.quantile(pr_per_cell,0.1)), float(np.quantile(pr_per_cell,0.9)))

# How many DAb cells are "strongly" progen-like?
for thr in [0.3, 0.5, 0.7]:
    print(f"fraction of DAb with progen_like >= {thr}: {float(np.mean(pr_per_cell >= thr)):.3f}")

# Optional: attach these scores to joint.obs for plotting
joint.obs.loc[dab_mask, "dab_pb_like"] = pb_per_cell
joint.obs.loc[dab_mask, "dab_progen_like"] = pr_per_cell


# %% [code]
# mask for DAb rows
dab_mask = (joint.obs["dataset_modality"].astype(str) == "DAb adt").values  # adjust if needed

# align DAb experiment to joint obs_names
exp_aligned = dab_adt_pp.obs["experiment"].astype(str).reindex(joint.obs_names)

# --- make patient writable (object), assign, then optionally recast ---
# 1) convert to object (avoids categorical restrictions)
joint.obs["patient"] = joint.obs["patient"].astype(object)

# 2) assign for DAb rows only
joint.obs.loc[dab_mask, "patient"] = exp_aligned.loc[joint.obs_names[dab_mask]].to_numpy()

# 3) (optional) cast back to category for nicer plotting/groupby
joint.obs["patient"] = joint.obs["patient"].astype("category")

# quick checks
print("DAb patient(=experiment) missing:", int(pd.isna(joint.obs.loc[dab_mask, "patient"]).sum()))
print(joint.obs.loc[dab_mask, "patient"].value_counts().head(10))


# %% [code]
# pick the high progen-like DAb cells
thr = 0.75
dab_hi = dab_mask.copy()
dab_hi[dab_mask] = (joint.obs.loc[dab_mask, "dab_progen_like"].values >= thr)

print("DAb high-progen cells:", int(dab_hi.sum()), " / ", int(dab_mask.sum()))

# if you have any DAb metadata columns, try these (edit as needed)
for col in ["source", "tissue", "site", "sample_id", "patient", "timepoint", "batch", "experiment"]:
    if col in joint.obs.columns:
        print("\nEnrichment by", col)
        tab = joint.obs.loc[dab_hi, col].value_counts(normalize=True).head(100)
        print(tab.to_string())


# %% [code]
# reuse ct_levels, frac, etc. from earlier
hi_idx = np.where(pr_per_cell >= 0.75)[0]   # indices within DAb subset ordering (Z_dab order)

avg_hi = frac[hi_idx].mean(axis=0)
df_hi = pd.DataFrame({"VG_celltype": ct_levels, "avg_frac_kNN": avg_hi}).sort_values("avg_frac_kNN", ascending=False)

print("\nVG neighbor composition for DAb cells with progen_like>=0.7:")
print(df_hi.head(15).to_string(index=False))


# %% [code]
import numpy as np
import pandas as pd

# define high progen subset among DAb
thr = 0.75
dab_mask = (joint.obs["dataset_modality"].astype(str) == "DAb adt").values
hi = joint.obs.loc[dab_mask, "dab_progen_like"].astype(float).values >= thr

# pull DAb ADT matrix and feature names
X = dab_adt_pp.X
var = dab_adt_pp.var_names.astype(str)

# pick marker panel (keep ones that exist)
markers = [m for m in ["CD34","CD117","KIT","CD38","HLA-DR","CD45RA","CD123","CD33","CD13","CD14","CD16"] if m in var]
print("Markers present:", markers)

if len(markers):
    Xd = X[:, [list(var).index(m) for m in markers]]
    # to dense safely for small marker subset
    Xd = Xd.toarray() if hasattr(Xd, "toarray") else np.asarray(Xd)

    df = pd.DataFrame({
        "marker": markers,
        "mean_hi": Xd[hi].mean(axis=0),
        "mean_lo": Xd[~hi].mean(axis=0),
        "diff_hi_minus_lo": Xd[hi].mean(axis=0) - Xd[~hi].mean(axis=0),
    }).sort_values("diff_hi_minus_lo", ascending=False)

    print(df.to_string(index=False))


# %% [code]
import numpy as np
import pandas as pd
import torch
from torch.utils.data import DataLoader

# expects these to already exist in your notebook:
#   SingleModalDataset, _as_modality_key, _to_device, latent_mu_student
#   model_ft, head, joint
#   vg_rna_pp, dab_adt_pp, cite_rna_pp_*, cite_adt_pp_*
#   GENES, device

@torch.no_grad()
def predict_probs_for_adata(
    model,
    head,
    adata,
    modality: str,
    genes,
    *,
    device,
    batch_size=512,
    num_workers=0,
    pin_memory=True,
):
    """Return DataFrame (n_cells x n_genes) with sigmoid probabilities."""
    model.eval(); head.eval()
    n = adata.n_obs
    G = len(genes)

    Y_dummy = np.zeros((n, G), dtype=np.float32)
    M_dummy = np.zeros((n, G), dtype=np.float32)

    ds = SingleModalDataset(adata, modality, Y_dummy, M_dummy)
    L = DataLoader(
        ds,
        batch_size=batch_size,
        shuffle=False,
        drop_last=False,
        num_workers=num_workers,
        pin_memory=pin_memory,
    )

    Ps = []
    for modality_batch, x, y, m in L:
        mod = _as_modality_key(modality_batch)
        x = _to_device(x, device)
        z = latent_mu_student(model, x, mod)
        logits = head(z)
        Ps.append(torch.sigmoid(logits).cpu().numpy())

    P = np.vstack(Ps) if len(Ps) else np.zeros((0, G), dtype=np.float32)
    return pd.DataFrame(P, index=adata.obs_names, columns=list(genes))


def write_probs_into_joint_masked_iloc(
    joint,
    probs_df,
    genes,
    mask,
    *,
    prefix="p_",
    mode="auto",         # "auto" | "positional" | "name"
    dedup="mean",         # used only for name mapping if probs_df has dup indices
):
    mask = np.asarray(mask, dtype=bool)
    pos = np.where(mask)[0]
    n_sub = len(pos)

    # ensure destination columns exist
    for g in genes:
        col = f"{prefix}{g}"
        if col not in joint.obs.columns:
            joint.obs[col] = np.nan

    # positional write if lengths match
    if mode in ("auto", "positional"):
        if probs_df.shape[0] == n_sub:
            P = probs_df.loc[:, list(genes)].to_numpy(dtype=float)
            for j, g in enumerate(genes):
                col = f"{prefix}{g}"
                joint.obs.iloc[pos, joint.obs.columns.get_loc(col)] = P[:, j]
            col0 = f"{prefix}{genes[0]}"
            n_filled = int(np.sum(~pd.isna(joint.obs.iloc[pos][col0])))
            print(f"Wrote probs POSITIONALLY into joint for subset rows={n_sub:,} (prefix={prefix}). Example filled for {genes[0]}: {n_filled:,}")
            return
        elif mode == "positional":
            raise ValueError(f"Positional write mismatch: subset {n_sub} vs probs {probs_df.shape[0]}")

    # name mapping fallback
    if probs_df.index.has_duplicates:
        if dedup == "mean":
            probs_df = probs_df.groupby(level=0).mean()
        elif dedup == "first":
            probs_df = probs_df[~probs_df.index.duplicated(keep="first")]
        else:
            raise ValueError("dedup must be 'mean' or 'first'")

    target_index = joint.obs_names[pos]
    wrote_any = False
    for g in genes:
        s = probs_df[g]
        mapped = pd.Index(target_index).map(s)
        mapped = np.asarray(mapped, dtype=float)
        ok = ~np.isnan(mapped)
        if ok.any():
            joint.obs.iloc[pos[ok], joint.obs.columns.get_loc(f"{prefix}{g}")] = mapped[ok]
            wrote_any = True

    if not wrote_any:
        print("WARNING: wrote 0 values (name mapping). Index mismatch between probs_df.index and joint.obs_names for this subset.")
    else:
        col0 = f"{prefix}{genes[0]}"
        n_filled = int(np.sum(~pd.isna(joint.obs.iloc[pos][col0])))
        print(f"Wrote probs by NAME into joint for subset rows={n_sub:,} (prefix={prefix}). Example filled for {genes[0]}: {n_filled:,}")


# ============================================================
# RUN: predict -> write into joint
# ============================================================

pin = (str(device) == "cuda")

vg_mask       = (joint.obs["dataset_modality"].astype(str) == "VG rna").values
dab_mask      = (joint.obs["dataset_modality"].astype(str) == "DAb adt").values
cite_rna_mask = (joint.obs["dataset_modality"].astype(str) == "CITE rna").values
cite_adt_mask = (joint.obs["dataset_modality"].astype(str) == "CITE adt").values

print("joint subset sizes:",
      "VG rna", int(vg_mask.sum()),
      "| DAb adt", int(dab_mask.sum()),
      "| CITE rna", int(cite_rna_mask.sum()),
      "| CITE adt", int(cite_adt_mask.sum()))

# NOTE: you used *_tr here; that's fine only if joint contains only train CITE.
# If joint contains ALL CITE (train+val+test), you must predict on the ALL object.
vg_probs       = predict_probs_for_adata(model_ft, head, vg_rna_pp,      "rna", GENES, device=device, pin_memory=pin)
dab_probs      = predict_probs_for_adata(model_ft, head, dab_adt_pp,     "adt", GENES, device=device, pin_memory=pin)
cite_rna_probs = predict_probs_for_adata(model_ft, head, cite_rna_pp_tr, "rna", GENES, device=device, pin_memory=pin)
cite_adt_probs = predict_probs_for_adata(model_ft, head, cite_adt_pp_tr, "adt", GENES, device=device, pin_memory=pin)

print("pred sizes:",
      "VG", vg_probs.shape[0],
      "| DAb", dab_probs.shape[0],
      "| CITE rna", cite_rna_probs.shape[0],
      "| CITE adt", cite_adt_probs.shape[0])

write_probs_into_joint_masked_iloc(joint, vg_probs,       GENES, vg_mask,       prefix="p_", mode="auto")
write_probs_into_joint_masked_iloc(joint, dab_probs,      GENES, dab_mask,      prefix="p_", mode="auto")
write_probs_into_joint_masked_iloc(joint, cite_rna_probs, GENES, cite_rna_mask, prefix="p_", mode="auto")
write_probs_into_joint_masked_iloc(joint, cite_adt_probs, GENES, cite_adt_mask, prefix="p_", mode="auto")

for g in GENES:
    col = f"p_{g}"
    print(col, "non-null:", int(joint.obs[col].notna().sum()), "/", joint.n_obs)


# %% [code]
for g in GENES:
    sc.pl.umap(joint, color=f"p_{g}", na_color="lightgrey", size=10, alpha=0.6, title=f"{g} prob")


# %% [code]
import numpy as np
import pandas as pd

def summarize_probs(joint, genes, group_col="dataset_modality", prefix="p_"):
    rows = []
    for g in genes:
        p = joint.obs[f"{prefix}{g}"].astype(float).to_numpy()
        rows.append({
            "gene": g,
            "n": np.isfinite(p).sum(),
            "min": np.nanmin(p),
            "p01": np.nanquantile(p, 0.01),
            "p50": np.nanquantile(p, 0.50),
            "p99": np.nanquantile(p, 0.99),
            "max": np.nanmax(p),
            "mean": np.nanmean(p),
            "std": np.nanstd(p),
        })
    df = pd.DataFrame(rows).set_index("gene")
    print(df.sort_values("std", ascending=False).to_string(float_format=lambda x: f"{x:.3f}"))

    # By dataset_modality (or whatever you want)
    for g in genes:
        col = f"{prefix}{g}"
        print(f"\n=== {g} by {group_col} ===")
        tmp = joint.obs[[group_col, col]].copy()
        tmp[col] = tmp[col].astype(float)
        out = tmp.groupby(group_col)[col].agg(["count","mean","std","min","max"])
        print(out.to_string(float_format=lambda x: f"{x:.3f}"))

summarize_probs(joint, GENES, group_col="dataset_modality", prefix="p_")


# %% [code]
import scanpy as sc

for g in ["NPM1","FLT3","DNMT3A","TP53"]:
    sc.pl.umap(
        joint,
        color=[f"p_{g}", "dataset_modality"],
        na_color="lightgrey",
        size=10,
        alpha=0.5,
        wspace=0.35,
        title=[f"{g} prob", "dataset_modality"]
    )


# %% [code]
import scanpy as sc

vg_only  = joint[joint.obs["dataset_modality"].astype(str) == "VG rna"].copy()
dab_only = joint[joint.obs["dataset_modality"].astype(str) == "DAb adt"].copy()

for g in ["NPM1","FLT3","DNMT3A","TP53","IDH2"]:
    sc.pl.umap(vg_only,  color=f"p_{g}", na_color="lightgrey", size=15, alpha=0.7, title=f"VG only: {g}")
    sc.pl.umap(dab_only, color=f"p_{g}", na_color="lightgrey", size=15, alpha=0.7, title=f"DAb only: {g}")


# %% [code]
import numpy as np
import pandas as pd

def var_explained_by_group(joint, gene, group_col, prefix="p_"):
    p = joint.obs[f"{prefix}{gene}"].astype(float)
    g = joint.obs[group_col].astype(str)
    ok = np.isfinite(p.to_numpy())
    p = p[ok]; g = g[ok]

    mu = p.mean()
    # between-group variance
    grp = p.groupby(g)
    n = grp.size()
    m = grp.mean()
    ss_between = float(((n * (m - mu)**2)).sum())
    ss_total = float(((p - mu)**2).sum()) + 1e-12
    return ss_between / ss_total

for g in ["TP53","FLT3","DNMT3A","NPM1","IDH2","TET2"]:
    r2_pat = var_explained_by_group(joint, g, "patient")
    r2_mod = var_explained_by_group(joint, g, "dataset_modality")
    print(f"{g:7s}  R2(patient)={r2_pat:.3f}  R2(dataset_modality)={r2_mod:.3f}")


# %% [code]


# %% [code]


# %% [code]


# %% [code]
print(vg_perf)
print(dab_perf)


# %% [code]
# ==============================================
# Plot (same idea, but showing n_pos/n_neg too)
# ==============================================

dfp = []
for g in GENES:
    d = dab_perf.get(g, {})
    v = vg_perf.get(g, {})

    dfp.append({
        "gene": g, "dataset": "DAb",
        "AUC": d.get("auc", np.nan),
        "AP":  d.get("ap",  np.nan),
        "n":   d.get("n_labeled",   np.nan),
        "n_pos": d.get("n_pos", d.get("pos", d.get("tp", np.nan))),
        "n_neg": d.get("n_neg", d.get("neg", d.get("tn", np.nan))),
        "prevalence": d.get("prevalence",   np.nan),
    })

    dfp.append({
        "gene": g, "dataset": "VG",
        "AUC": v.get("auc", np.nan),
        "AP":  v.get("ap",  np.nan),
        "n":   v.get("n_labeled",   np.nan),
        "n_pos": v.get("n_pos", v.get("pos", v.get("tp", np.nan))),
        "n_neg": v.get("n_neg", v.get("neg", v.get("tn", np.nan))),
        "prevalence": v.get("prevalence",   np.nan),
    })

dfp = pd.DataFrame(dfp)

print(dfp)


# %% [code]
plot_df = dfp[dfp["AUC"].notna()].copy()

plt.figure(figsize=(6.8, 4.0))
for ds in ["DAb", "VG"]:
    sub = plot_df[plot_df["dataset"] == ds]
    plt.plot(sub["AUC"].values, sub["gene"].values, marker="o", linestyle="none", label=ds)

plt.xlabel("AUC"); plt.ylabel("Gene")
plt.title("Mutation-head performance (AUC) — evaluable only")
plt.legend()
plt.tight_layout()
plt.show()


# %% [code]
import numpy as np
import torch

def to_device_batch(batch, device, non_blocking=True):
    """
    Recursively move a batch to `device`.
    Supports: dict, list/tuple, torch.Tensor, numpy arrays.
    Leaves strings/None/ints/floats unchanged.
    """
    if batch is None:
        return None

    # torch tensor
    if torch.is_tensor(batch):
        return batch.to(device, non_blocking=non_blocking)

    # numpy -> torch
    if isinstance(batch, np.ndarray):
        return torch.from_numpy(batch).to(device, non_blocking=non_blocking)

    # dict
    if isinstance(batch, dict):
        return {k: to_device_batch(v, device, non_blocking=non_blocking) for k, v in batch.items()}

    # list / tuple
    if isinstance(batch, (list, tuple)):
        out = [to_device_batch(v, device, non_blocking=non_blocking) for v in batch]
        return type(batch)(out)

    # everything else (str, int, float, pd objects, etc.)
    return batch


# %% [code]
@torch.no_grad()
def get_latent_z(model, x_dict, *, prefer="mu"):
    """
    Return latent z for a batch.
    - If model.encode exists: uses it.
    - Else tries model(x_dict) and extracts common latent keys.
    """
    if hasattr(model, "encode") and callable(getattr(model, "encode")):
        z = model.encode(x_dict)
        return z

    out = model(x_dict)

    # If your forward returns a dict
    if isinstance(out, dict):
        for key in ["mu", "z", "latent", "z_mu", "X_univi", "repr"]:
            if key in out:
                return out[key]
        raise KeyError(f"Model output dict keys: {list(out.keys())} (couldn't find latent)")

    # Otherwise assume forward returns z directly
    return out


# %% [code]
import numpy as np
import torch

@torch.no_grad()
def encode_and_predict_probs(
    model, head, adata, *,
    modality, genes,
    device=None,
    batch_size=4096,
    latent_key="X_univi",
):
    """
    Calls your free function: encode_latent(model, adata, modality, ...)
    then applies head to latent -> sigmoid probs.
    """
    model.eval()
    head.eval()

    if device is None:
        device = next(model.parameters()).device

    # 1) Encode latent (NOTE: pass model!)
    Z = encode_latent(
        model,                       # <-- this was missing
        adata=adata,
        modality=modality,
        batch_size=batch_size,
        device=device,
    )

    # If encode_latent writes into obsm and returns None
    if Z is None:
        if latent_key not in adata.obsm:
            raise KeyError(f"encode_latent returned None and {latent_key} not found in adata.obsm")
        Z = adata.obsm[latent_key]

    # 2) Ensure torch tensor on correct device
    if isinstance(Z, np.ndarray):
        Zt = torch.from_numpy(Z)
    elif torch.is_tensor(Z):
        Zt = Z
    else:
        Zt = torch.tensor(Z)

    Zt = Zt.to(device)

    # 3) Predict probs
    logits = head(Zt)               # (n_cells, n_genes)
    probs  = torch.sigmoid(logits).detach().cpu().numpy()

    return {g: probs[:, i] for i, g in enumerate(genes)}, probs


# modality strings (yours)
VG_MODAL  = "rna"
DAB_MODAL = "adt"

vg_prob_by_gene, P_vg = encode_and_predict_probs(model_ft, head, vg_rna_pp,  modality=VG_MODAL,  genes=GENES)
dab_prob_by_gene, P_dab = encode_and_predict_probs(model_ft, head, dab_adt_pp, modality=DAB_MODAL, genes=GENES)

vg_rna_pp.obs[f"headP_{HERO}_vg"]    = vg_prob_by_gene[HERO]
dab_adt_pp.obs[f"headP_{HERO}_dab"]  = dab_prob_by_gene[HERO]


# %% [code]
import scanpy as sc
import matplotlib.pyplot as plt
from pathlib import Path

FIGDIR = Path(FIGDIR); FIGDIR.mkdir(parents=True, exist_ok=True)

col_vg  = f"headP_{HERO}_vg"
col_dab = f"headP_{HERO}_dab"

sc.pl.umap(vg_rna_pp, color=col_vg,  title=f"Head P({HERO}) on VG (RNA)",  size=15, alpha=0.7)
plt.savefig(FIGDIR / f"umap_headP_{HERO}_VG.png", dpi=450, bbox_inches="tight"); plt.show()

sc.pl.umap(dab_adt_pp, color=col_dab, title=f"Head P({HERO}) on DAb (ADT)",  size=15, alpha=0.7)
plt.savefig(FIGDIR / f"umap_headP_{HERO}_DAb.png", dpi=450, bbox_inches="tight"); plt.show()


# %% [code]
import numpy as np
import pandas as pd
import scanpy as sc
import matplotlib.pyplot as plt
from pathlib import Path

FIGDIR = Path(FIGDIR); FIGDIR.mkdir(parents=True, exist_ok=True)

# Columns holding probs on source objects
col_vg  = f"headP_{HERO}_vg"
col_dab = f"headP_{HERO}_dab"

# Column we will create on joint for plotting
col_joint = f"headP_{HERO}_joint"

# Initialize with NaNs
joint.obs[col_joint] = np.nan

# Masks for where to place values
mask_vg  = (joint.obs["dataset"] == "VG")  & (joint.obs["modality"] == "rna")
mask_dab = (joint.obs["dataset"] == "DAb") & (joint.obs["modality"] == "adt")

# Assign (assumes the joint rows for VG/DAb are in the same order as vg_rna_pp/dab_adt_pp used to build joint)
joint.obs.loc[mask_vg,  col_joint] = vg_rna_pp.obs[col_vg].to_numpy()
joint.obs.loc[mask_dab, col_joint] = dab_adt_pp.obs[col_dab].to_numpy()

# Plot (continuous)
sc.pl.umap(
    joint,
    color=col_joint,
    title=f"Head P({HERO}) on joint UMAP (VG RNA + DAb ADT)",
    na_color="lightgrey",
    size=15, 
    alpha=0.7,
)
plt.savefig(FIGDIR / f"umap_headP_{HERO}_joint.png", dpi=450, bbox_inches="tight")
plt.show()

# Optional: side-by-side with dataset label for sanity
sc.pl.umap(
    joint,
    color=["dataset", col_joint],
    title=["dataset", f"Head P({HERO})"],
    na_color="lightgrey",
    wspace=0.35,
    size=15, 
    alpha=0.7,
)
plt.savefig(FIGDIR / f"umap_dataset_and_headP_{HERO}_joint.png", dpi=450, bbox_inches="tight")
plt.show()


# %% [code]
print(joint)


# %% [code]
import numpy as np
import pandas as pd

score_col = "LSC17_score"
out_col   = "LSC17_score"

# ensure the column exists
if out_col not in joint.obs.columns:
    joint.obs[out_col] = np.nan

def add_scores_fill_missing(src, joint, col, out_col):
    """
    Fill joint.obs[out_col] from src.obs[col] by matching on obs_names.

    Works even if joint.obs_names has duplicates.
    Only fills where joint[out_col] is currently missing (NaN).
    """
    # src obs_names are unique in your report; build mapper obs_name -> score
    mapper = pd.Series(src.obs[col].values, index=pd.Index(src.obs_names))

    # lookup for every joint row (duplicates are fine; they'll get same value)
    looked_up = pd.Index(joint.obs_names).map(mapper)

    # fill only where joint currently missing and looked_up is not missing
    cur = joint.obs[out_col].to_numpy(copy=True)

    # coerce looked_up to float if possible (LSC17_score should be numeric)
    new = pd.to_numeric(pd.Series(looked_up), errors="coerce").to_numpy()

    mask = np.isnan(cur) & ~np.isnan(new)
    cur[mask] = new[mask]
    joint.obs[out_col] = cur

    return joint.obs[out_col]

# choose your priority order:
add_scores_fill_missing(cite_rna_pp_tr, joint, score_col, out_col)
add_scores_fill_missing(cite_rna_pp_va, joint, score_col, out_col)
add_scores_fill_missing(cite_rna_pp_te, joint, score_col, out_col)
add_scores_fill_missing(vg_rna_pp,      joint, score_col, out_col)


# %% [code]
sc.pl.umap(
    joint,
    color=["LSC17_score"],
    title=["Joint latent fine-tuned (LSC-17 score)"],
    na_color="lightgrey",
    wspace=0.35,
    size=15,
    vmin=0,
    vmax=1.0,
    alpha=0.85,
)


# %% [code]
import numpy as np
import pandas as pd
from sklearn.metrics import roc_auc_score, average_precision_score, roc_curve, precision_recall_curve

# --- assumes you already have these helpers ---
# find_mut_col_dab_obs(...)
# coerce_to_nullable_boolean(...)
# make_label_from_nullable_bool(...)

HERO = str(HERO)

mut_col_dab = find_mut_col_dab_obs(dab_adt_al, HERO)
if mut_col_dab is None:
    raise KeyError(f"Can't find DAb mutation column for {HERO}. Example: {list(dab_adt_al.obs.columns[:30])}")

col_truth_dab = f"truth_{HERO}_dab_bool"
dab_adt_al.obs[col_truth_dab] = coerce_to_nullable_boolean(dab_adt_al.obs[mut_col_dab])

print("DAb truth source:", mut_col_dab)
print(dab_adt_al.obs[col_truth_dab].value_counts(dropna=False))


# %% [code]
def vg_truth_from_transcript_strings(vg_obs, hero, mut_col="MutTranscripts", wt_col="WtTranscripts"):
    hero_u = str(hero).upper()

    mut_s = vg_obs[mut_col].fillna("").astype(str).str.upper()
    wt_s  = vg_obs[wt_col].fillna("").astype(str).str.upper()

    # naive “contains HERO” (works for "NPM1", "FLT3", etc.)
    has_mut = mut_s.str.contains(hero_u, regex=False)
    has_wt  = wt_s.str.contains(hero_u,  regex=False)

    out = pd.Series(pd.NA, index=vg_obs.index, dtype="boolean")
    out[has_mut] = True
    out[(~has_mut) & has_wt] = False
    return out

col_truth_vg = f"truth_{HERO}_vg_bool"
vg_rna_pp.obs[col_truth_vg] = vg_truth_from_transcript_strings(vg_rna_pp.obs, HERO)

print(vg_rna_pp.obs[col_truth_vg].value_counts(dropna=False))


# %% [code]
col_pred_dab = f"headP_{HERO}_dab"  # you already created this on dab_adt_pp

# Align truth and preds by cell ID (index)
truth = dab_adt_al.obs[col_truth_dab].astype("boolean")
pred  = dab_adt_pp.obs[col_pred_dab].astype(float)

# Only evaluate where truth is known and pred is not nan
mask = (~truth.isna()) & (~pred.isna())
y = truth[mask].astype(int).to_numpy()
p = pred[mask].to_numpy()

auc = roc_auc_score(y, p) if len(np.unique(y)) > 1 else np.nan
ap  = average_precision_score(y, p) if len(np.unique(y)) > 1 else np.nan
print(f"DAb: AUC={auc:.4f}  AP={ap:.4f}   n={mask.sum()}")


# %% [code]
col_pred_vg = f"headP_{HERO}_vg"

truth = vg_rna_pp.obs[col_truth_vg].astype("boolean")
pred  = vg_rna_pp.obs[col_pred_vg].astype(float)

mask = (~truth.isna()) & (~pred.isna())
y = truth[mask].astype(int).to_numpy()
p = pred[mask].to_numpy()

auc = roc_auc_score(y, p) if len(np.unique(y)) > 1 else np.nan
ap  = average_precision_score(y, p) if len(np.unique(y)) > 1 else np.nan
print(f"VG:  AUC={auc:.4f}  AP={ap:.4f}   n={mask.sum()}")


# %% [code]


# %% [code]


# %% [code]


# %% [code]


# %% [code]


# %% [code]


# %% [code]


# %% [code]


# %% [code]


# %% [code]


# %% [code]


# %% [code]


# %% [code]


# %% [code]


# %% [code]


# %% [code]


# %% [code]


# %% [code]


# %% [code]


# %% [code]


# %% [code]


# %% [code]


# %% [code]


# %% [code]


# %% [code]


# %% [code]


# %% [code]


# %% [code]


# %% [code]


# %% [code]


# %% [code]


# %% [code]


# %% [code]


# %% [code]


# %% [code]


# %% [code]


# %% [code]


# %% [code]


# %% [code]


# %% [code]
````
