## Supplemental Code for "Unifying multimodal single-cell data with a mixture-of-experts *β*-variational autoencoder framework": Fig08_benchmark_merge_plot.ipynb

```
# %% [markdown]
## UniVI manuscript – Commonly-used data integration tool benchmarking test for Genome Research manuscript revisions
### UniVI benchmark (Multiome PBMC): Merging and plotting PyTorch and R tool CV sweep metrics

Andrew Ashford, Pathways + Omics Group, Oregon Health & Science University, Portland, OR – 1/25/2026


This notebook:
- loads summary tables output by the PyTorch and R integration method CV sweep notebooks
- generates plots of each of the metrics of all combines tools used for Multiome method comparison evaluations
  
Manuscript: https://www.biorxiv.org/content/10.1101/2025.02.28.640429v1.full
Code: https://github.com/Ashford-A/UniVI


# %% [markdown]
#### Import modules and specify file paths

# %% [code]
import numpy as np
import pandas as pd
from pathlib import Path

# -----------------------------
# Paths
# -----------------------------
#P_CV_LONG = Path("/home/groups/precepts/ashforda/runs_multiome_py_1-24-2026/runs/cv_sweep_py_1-24-2026/cv_sweep_long.csv")
P_CV_LONG = Path("/home/groups/precepts/ashforda/runs_multiome_py_1-31-2026/runs/cv_sweep_py_1-31-2026/cv_sweep_long.csv")
#P_CV_RUNS = Path("/home/groups/precepts/ashforda/univi_bench/runs_multiome_R_1-30-2026_WORKING/runs/cv_sweep_R_1-30-2026_WORKING/cv_runs.tsv")
P_CV_RUNS = Path("/home/groups/precepts/ashforda/univi_bench/runs_multiome_R_1-31-2026/runs/cv_sweep_R_1-31-2026/cv_runs.tsv")

OUT_DIR = Path("./Figure_9-method_comparison/violin_plots")
OUT_DIR.mkdir(parents=True, exist_ok=True)


# %% [markdown]
#### Load tables

# %% [code]
print("Reading:", P_CV_LONG)
print("Reading:", P_CV_RUNS)

df_long = pd.read_csv(P_CV_LONG)
df_runs = pd.read_csv(P_CV_RUNS, sep="\t")

print("df_long:", df_long.shape, "cols=", len(df_long.columns))
print("df_runs:", df_runs.shape, "cols=", len(df_runs.columns))


# %% [markdown]
#### Harmonize method tables

# %% [code]
# -----------------------------
# Standardization maps
# -----------------------------
# If df_runs ever contains verbose names, this maps them to short keys.
# If df_runs already contains short keys (like seurat_wnn), it will pass through unchanged.
RUNS_METHOD_MAP = {
    "Seurat/Signac WNN (Seurat 5.4.0; Signac 1.16.0)": "seurat_wnn",
    "Seurat CCA RNA vs GeneActivity (Seurat 5.4.0; Signac 1.16.0)": "seurat_cca",
    "Seurat v5 Bridge (Seurat 5.4.0; Signac 1.16.0; SeuratObject 5.3.0)": "seurat_bridge",
    "Harmony-modality (harmony 1.2.4)": "harmony",
    "LIGER gene-activity iNMF (rliger 2.2.1; SeuratObject 5.3.0)": "liger",
    "MOFA2 on PCA+LSI views (MOFA2 1.16.0)": "mofa2",
}

# short keys -> display names
DISPLAY_NAME = {
    "univi": "UniVI",
    "scglue": "scGLUE",
    "scjoint": "scJoint",
    "cobolt": "Cobolt",
    "multivi": "MultiVI",
    "peakvi": "PeakVI",
    "scmomat": "scMoMaT",
    "deepcca": "DeepCCA",
    "multimap": "MultiMap",

    "harmony": "Harmony",

    # robust: accept both keys if they appear
    "seurat_wnn": "SeuratWNN",
    "wnn": "SeuratWNN",

    "seurat_bridge": "SeuratBridge",
    "seurat_cca": "SeuratCCA",
    "liger": "LIGER",
    "mofa2": "MOFA2",
}

# raw column names -> standardized names (apply to BOTH tables)
METRIC_RENAME = {
    "PAIR(test)/FOSCTTM_mean": "FOSCTTM_mean_test",
    "PAIR(test)/FOSCTTM_MRR": "FOSCTTM_mrr_mean_test",
    "PAIR(test)/FOSCTTM_Recall@1": "FOSCTTM_recall@1_mean_test",
    "PAIR(test)/FOSCTTM_Recall@10": "FOSCTTM_recall@10_mean_test",
    "PAIR(test)/FOSCTTM_Recall@25": "FOSCTTM_recall@25_mean_test",
    "PAIR(test)/FOSCTTM_Recall@50": "FOSCTTM_recall@50_mean_test",
    "PAIR(test)/FOSCTTM_Recall@100": "FOSCTTM_recall@100_mean_test",
    "PAIR(test)/Label transfer acc mean": "label_transfer_acc_mean_test",
    "PAIR(test)/Label transfer macroF1 mean": "label_transfer_macroF1_mean_test",

    "MIX(test)/Mixing score": "mixing_score_test",

    "Fused kNN acc": "fused_knn_acc_test",
    "Fused kNN macroF1": "fused_knn_macroF1_test",
    "Fused kNN balanced acc": "fused_knn_balanced_acc_test",

    "Fused(fusedZ,test) silhouette": "fused_silhouette_test",
    "Fused(fusedZ,test) label purity": "fused_knn_label_purity_test",
    "Fused(fusedZ,test) k-means ARI": "fused_kmeans_ari_test",
    "Fused(fusedZ,test) k-means NMI": "fused_kmeans_nmi_test",
}

METHOD_ORDER = [
    "UniVI","scGLUE","Cobolt","MultiVI","scJoint","PeakVI","scMoMaT","MultiMap","DeepCCA",
    "Harmony","MOFA2","LIGER","SeuratWNN","SeuratBridge","SeuratCCA",
]


# %% [code]
# -----------------------------
# Helpers
# -----------------------------
def _clean_str(x):
    if pd.isna(x):
        return np.nan
    return str(x).strip()

def _clean_lower(x):
    if pd.isna(x):
        return np.nan
    return str(x).strip().lower()

def standardize_method_series(s: pd.Series) -> pd.Series:
    """
    Works for:
      - already-short keys like seurat_wnn
      - verbose strings that appear in RUNS_METHOD_MAP
    """
    m = s.map(_clean_str)

    # Map verbose -> short if applicable
    m2 = m.map(RUNS_METHOD_MAP)
    m = m2.fillna(m)

    # Lowercase keys
    m = m.map(_clean_lower)

    # Pretty display names if known
    m = m.map(DISPLAY_NAME).fillna(m)
    return m

def ensure_status(df: pd.DataFrame, failed_col="failed") -> pd.DataFrame:
    df = df.copy()
    if "status" in df.columns:
        df["status"] = df["status"].astype(str)
        return df
    if failed_col in df.columns:
        df["status"] = np.where(df[failed_col].astype(bool), "failed", "ok")
    else:
        df["status"] = "ok"
    return df

def to_long(df: pd.DataFrame, metric_cols: list[str], source_label: str) -> pd.DataFrame:
    keep = ["method"] + [m for m in metric_cols if m in df.columns]
    out = df[keep].copy()
    for m in keep[1:]:
        out[m] = pd.to_numeric(out[m], errors="coerce")
    out = out.melt(id_vars=["method"], var_name="metric", value_name="value")
    out["source"] = source_label
    out = out.dropna(subset=["method", "value"])
    return out


# %% [code]
# -----------------------------
# Standardize BOTH tables
# -----------------------------
df_long_std = df_long.copy()
df_runs_std = df_runs.copy()

# rename metrics on both so they can match 1:1
df_long_std = df_long_std.rename(columns=METRIC_RENAME)
df_runs_std = df_runs_std.rename(columns=METRIC_RENAME)

# standardize method names on both
if "method" in df_long_std.columns:
    df_long_std["method"] = standardize_method_series(df_long_std["method"])
else:
    df_long_std["method"] = "unknown"

if "method" in df_runs_std.columns:
    df_runs_std["method"] = standardize_method_series(df_runs_std["method"])
else:
    df_runs_std["method"] = "unknown"

# status on both
df_long_std = ensure_status(df_long_std)
df_runs_std = ensure_status(df_runs_std)

# keep only successful
df_long_ok = df_long_std[df_long_std["status"].str.lower().eq("ok")].copy()
df_runs_ok = df_runs_std[df_runs_std["status"].str.lower().eq("ok")].copy()

# keep only methods that exist
METHOD_ORDER_USE = [m for m in METHOD_ORDER if m in set(pd.concat([df_long_ok["method"], df_runs_ok["method"]]).unique())]
print("Final METHOD_ORDER:", METHOD_ORDER_USE)


# %% [code]
# -----------------------------
# Pick metrics: union of numeric-like cols across both
# -----------------------------
exclude_cols = {
    "method","seed","fold","status","error","failed","transductive","uses_labels",
    "n_train","n_val","n_test","latent_dim","reason","n_features","k_view","lr","dropout",
    "weight_decay","batch_size","max_epochs","patience","reg","best_val","nbatches",
    "n_latent","n_latent_requested","joint_dim_returned","atac_tfidf",
    "patched_scipy_sparse_vstack_scoped","latent_cap","lr_used","source","skipped",
    "__eval_error__","PAIR(test)/FOSCTTM_n_eval",
}

def numeric_metric_cols(df: pd.DataFrame) -> list[str]:
    cols = []
    for c in df.columns:
        if c in exclude_cols:
            continue
        if pd.api.types.is_numeric_dtype(df[c]):
            cols.append(c)
        else:
            v = pd.to_numeric(df[c], errors="coerce")
            if v.notna().any():
                cols.append(c)
    return cols

metric_cols = sorted(set(numeric_metric_cols(df_long_ok)).union(numeric_metric_cols(df_runs_ok)))

headline = [
    "FOSCTTM_mean_test",
    "label_transfer_acc_mean_test",
    "label_transfer_macroF1_mean_test",
    "mixing_score_test",
    "fused_knn_acc_test",
    "fused_kmeans_nmi_test",
    "fused_kmeans_ari_test",
    "fused_silhouette_test",
]
metric_cols = [m for m in headline if m in metric_cols] + [m for m in metric_cols if m not in headline]

print("Detected metrics:", len(metric_cols))
print(metric_cols[:40])


# %% [code]
# -----------------------------
# Build one long plotting table
# -----------------------------
long_py = to_long(df_long_ok, metric_cols, source_label="py")
long_r  = to_long(df_runs_ok, metric_cols, source_label="r")
df_plot = pd.concat([long_py, long_r], ignore_index=True)


# %% [code]
print(df_plot.groupby(["source","method"]).size().sort_values(ascending=False).head(30))
print("Metrics shared by both sources:", len(set(long_py.metric) & set(long_r.metric)))


# %% [code]
print("py runs per method:")
print(df_long_ok.groupby("method").size().sort_values(ascending=False))

print("\nr runs per method:")
print(df_runs_ok.groupby("method").size().sort_values(ascending=False))


# %% [code]
m = "FOSCTTM_mean_test"  # change as needed
tmp = df_plot[df_plot["metric"] == m]
print(tmp.groupby(["source","method"]).size().sort_values(ascending=False))


# %% [code]
import matplotlib as mpl
import matplotlib.pyplot as plt

# Bigger default figure size + readable fonts
mpl.rcParams["figure.figsize"] = (16, 14)    # default size for all new figures
mpl.rcParams["figure.dpi"] = 300             # notebook display dpi
mpl.rcParams["savefig.dpi"] = 300            # saved image dpi (paper-ready)
mpl.rcParams["font.size"] = 12
mpl.rcParams["axes.titlesize"] = 14
mpl.rcParams["axes.labelsize"] = 13
mpl.rcParams["xtick.labelsize"] = 11
mpl.rcParams["ytick.labelsize"] = 11
mpl.rcParams["legend.fontsize"] = 11
mpl.rcParams["lines.linewidth"] = 1.5


# %% [code]
# -----------------------------
# Plot: matched methods per metric on the same axis/order
# -----------------------------
def violin_metric_matched(
    df_plot_long: pd.DataFrame,
    metric: str,
    *,
    method_order: list[str],
    title: str | None = None,
    savepath: str | Path | None = None,
    figsize_base=(6, 4.5),
    min_n: int = 1,
):
    dd = df_plot_long[df_plot_long["metric"] == metric].copy()
    if dd.empty:
        print(f"[plot] no data for metric={metric}")
        return

    counts = dd.groupby("method")["value"].size()
    keep_methods = counts[counts >= min_n].index.tolist()
    dd = dd[dd["method"].isin(keep_methods)]
    if dd.empty:
        print(f"[plot] no methods with >= {min_n} points for metric={metric}")
        return

    present = list(pd.unique(dd["method"]))
    order_use = [m for m in method_order if m in present] + [m for m in present if m not in method_order]

    data = [dd.loc[dd["method"] == m, "value"].to_numpy(dtype=float) for m in order_use]
    means = np.array([np.mean(x) for x in data], dtype=float)
    sems  = np.array([np.std(x, ddof=1) / np.sqrt(len(x)) if len(x) > 1 else np.nan for x in data], dtype=float)
    medians = np.array([np.median(x) for x in data], dtype=float)

    fig_w = max(figsize_base[0], 0.85 * len(order_use))
    fig, ax = plt.subplots(figsize=(fig_w, figsize_base[1]))

    ax.violinplot(
        data,
        positions=np.arange(1, len(order_use) + 1),
        showmeans=False,
        showextrema=False,
        showmedians=False,
    )

    xs = np.arange(1, len(order_use) + 1)
    ax.errorbar(xs, means, yerr=sems, fmt="o", capsize=4, linewidth=1.5)
    ax.scatter(xs, medians, marker="_", s=300)

    ax.set_xticks(xs)
    ax.set_xticklabels(order_use, rotation=30, ha="right")
    ax.set_ylabel(metric)
    ax.set_title(title or f"{metric} (matched methods on same plot)")
    ax.grid(True, axis="y", alpha=0.3)
    fig.tight_layout()

    if savepath is not None:
        savepath = Path(savepath)
        savepath.parent.mkdir(parents=True, exist_ok=True)
        fig.savefig(savepath, dpi=200, bbox_inches="tight")

    plt.show()

for metric in metric_cols:
    violin_metric_matched(
        df_plot,
        metric,
        method_order=METHOD_ORDER_USE,
        title=f"{metric} across seed×fold (Py + R together; matched methods)",
        savepath=OUT_DIR / f"violin_matched_{metric}.png",
        min_n=1,
    )


# %% [code]


# %% [code]
import numpy as np
import pandas as pd
import matplotlib.pyplot as plt
from pathlib import Path

# -----------------------------
# Utility: decide direction automatically (overrideable)
# -----------------------------
def _infer_higher_is_better(metric_name: str) -> bool:
    """
    Heuristic:
      - FOSCTTM_mean: lower is better
      - loss/error/dist/etc: usually lower is better
      - otherwise assume higher is better
    """
    m = str(metric_name).lower()
    if "foscttm_mean" in m:
        return False
    if any(k in m for k in ["loss", "error", "distance", "dist", "rmse", "mse"]):
        return False
    return True

# -----------------------------
# Plot: best -> worst (per metric) across both sources
# -----------------------------
def violin_metric_best_to_worst(
    df_plot_long: pd.DataFrame,
    metric: str,
    *,
    title: str | None = None,
    savepath: str | Path | None = None,
    figsize_base=(6, 4.5),
    agg: str = "mean",               # "mean" or "median" for sorting
    higher_is_better: bool | None = None,
    min_n: int = 1,                  # drop methods with < min_n obs for this metric
    tie_breaker: str = "median",     # used when agg="mean" and means tie closely
    tie_eps: float = 1e-12,
    show_n_in_labels: bool = True,
):
    dd = df_plot_long[df_plot_long["metric"] == metric].copy()
    if dd.empty:
        print(f"[plot] no data for metric={metric}")
        return None

    # robust numeric conversion
    dd["value"] = pd.to_numeric(dd["value"], errors="coerce")
    dd = dd.dropna(subset=["value", "method"])
    if dd.empty:
        print(f"[plot] no numeric data for metric={metric}")
        return None

    # counts + filter
    gb = dd.groupby("method")["value"]
    counts = gb.size()
    keep = counts[counts >= min_n].index
    dd = dd[dd["method"].isin(keep)]
    if dd.empty:
        print(f"[plot] no methods with >= {min_n} points for metric={metric}")
        return None

    gb = dd.groupby("method")["value"]
    means = gb.mean()
    medians = gb.median()

    # pick scoring function for ordering
    if agg == "mean":
        score = means.copy()
        # optional tie-breaker
        if tie_breaker == "median":
            # stable-ish: sort by mean, then median
            score_tbl = pd.DataFrame({"mean": means, "median": medians, "n": gb.size()})
        else:
            score_tbl = pd.DataFrame({"mean": means, "median": medians, "n": gb.size()})
    elif agg == "median":
        score = medians.copy()
        score_tbl = pd.DataFrame({"median": medians, "mean": means, "n": gb.size()})
    else:
        raise ValueError("agg must be 'mean' or 'median'")

    if higher_is_better is None:
        higher_is_better = _infer_higher_is_better(metric)

    # sort best -> worst
    ascending = not higher_is_better

    # sort worst -> best
    #ascending = higher_is_better

    if agg == "mean":
        # sort by mean, then median as a deterministic tie-breaker
        score_tbl = score_tbl.sort_values(
            by=["mean", "median"] if higher_is_better else ["mean", "median"],
            ascending=[ascending, ascending],
        )
        # If you want “real” tie detection:
        # (kept simple; the sort above already gives deterministic ordering)
        order_use = score_tbl.index.tolist()
    else:
        # sort by median, then mean as tie-breaker
        score_tbl = score_tbl.sort_values(
            by=["median", "mean"],
            ascending=[ascending, ascending],
        )
        order_use = score_tbl.index.tolist()

    # gather per-method arrays in that order
    data = [dd.loc[dd["method"] == m, "value"].to_numpy(dtype=float) for m in order_use]

    # compute summary stats for errorbars/markers
    mean_vals = np.array([np.mean(x) for x in data], dtype=float)
    sem_vals  = np.array([np.std(x, ddof=1) / np.sqrt(len(x)) if len(x) > 1 else np.nan for x in data], dtype=float)
    med_vals  = np.array([np.median(x) for x in data], dtype=float)

    # x tick labels (optionally annotate n)
    if show_n_in_labels:
        ns = score_tbl.loc[order_use, "n"].to_numpy()
        xticklabels = [f"{m}\n(n={int(n)})" for m, n in zip(order_use, ns)]
    else:
        xticklabels = order_use

    # figure
    fig_w = max(figsize_base[0], 0.85 * len(order_use))
    fig, ax = plt.subplots(figsize=(fig_w, figsize_base[1]))

    ax.violinplot(
        data,
        positions=np.arange(1, len(order_use) + 1),
        showmeans=False,
        showextrema=False,
        showmedians=False,
    )

    xs = np.arange(1, len(order_use) + 1)
    ax.errorbar(xs, mean_vals, yerr=sem_vals, fmt="o", capsize=4, linewidth=1.5)
    ax.scatter(xs, med_vals, marker="_", s=300)

    direction = "higher=better" if higher_is_better else "lower=better"
    ax.set_xticks(xs)
    ax.set_xticklabels(xticklabels, rotation=30, ha="right")
    ax.set_ylabel(metric)
    ax.set_title(title or f"{metric} (best→worst by {agg}; {direction})")
    ax.grid(True, axis="y", alpha=0.3)
    fig.tight_layout()

    if savepath is not None:
        savepath = Path(savepath)
        savepath.parent.mkdir(parents=True, exist_ok=True)
        fig.savefig(savepath, dpi=200, bbox_inches="tight")

    plt.show()

    # return ranking table (for reporting)
    out = score_tbl.copy()
    out.insert(0, "method", out.index)
    out = out.reset_index(drop=True)
    out.insert(1, "metric", metric)
    out.insert(2, "higher_is_better", higher_is_better)
    return out


# -----------------------------
# Run for all metrics (save plots + collect rankings)
# -----------------------------
rank_tables = []
for metric in metric_cols:
    rank = violin_metric_best_to_worst(
        df_plot,
        metric,
        agg="mean",                       # or "median"
        higher_is_better=None,            # auto; override if you want
        min_n=1,
        title=f"{metric} across seed×fold (Py + R together; best→worst)",
        savepath=OUT_DIR / f"violin_best-to-worst_{metric}.png",
        show_n_in_labels=True,
    )
    if rank is not None:
        rank_tables.append(rank)

df_rank_all = pd.concat(rank_tables, ignore_index=True) if rank_tables else pd.DataFrame()
print("df_rank_all:", df_rank_all.shape)
display(df_rank_all.head(20))


# %% [code]


# %% [code]


# %% [code]


# %% [code]


# %% [code]


# %% [code]


# %% [code]


# %% [code]


# %% [code]


# %% [code]


# %% [code]


# %% [code]


# %% [code]
```
