## Supplemental Code for "Unifying multimodal single-cell data with a mixture-of-experts *β*-variational autoencoder framework": Fig08_benchmark_PyTorch.ipynb

````
# %% [markdown]
## UniVI manuscript - Figure 8 generation reproducible workflow
### Commonly-used data integration tool benchmarking test for Genome Research manuscript revisions - UniVI benchmark (Multiome PBMC): UniVI vs Python baselines that use PyTorch

Andrew Ashford, Pathways + Omics Group, Oregon Health & Science University, Portland, OR – 12/17/2025

This notebook:
- loads paired Multiome PBMC (RNA+ATAC)
- runs integration baselines available in a PyTorch/Python environment:
  - UniVI (our method)
  - scvi-tools MultiVI (paired RNA+ATAC)
  - MultiMAP (Teichlab)
  - scGLUE (optional; requires a guidance graph)
- evaluates with:
  - FOSCTTM (paired alignment)
  - modality mixing (kNN modality entropy-ish)
  - kNN label transfer (RNA→ATAC, ATAC→RNA)
- saves embeddings + a summary table

You can see our manuscript, "Unifying multimodal single-cell data with a mixture-of-experts β-variational autoencoder framework" which is currently being revised for Genome Research and is available on bioRxiv at the following link: https://www.biorxiv.org/content/10.1101/2025.02.28.640429v1.full

GitHub for the project - including a Quickstart guide - can be found at: https://github.com/Ashford-A/UniVI

Package is pip installable via the command: 
```bash
pip install univi
```


# %% [markdown]
### Setting cache directories

# %% [code]
import os
from pathlib import Path

# Put these on your larger-quota filesystem (RDS). Adjust if needed.
RDS = Path("/home/groups/precepts/ashforda")
CACHE = RDS / "cache"
WORK = RDS / "runs_multiome_py_1-31-2026"       # outputs go here
# Should have used WORK directory:
#WORK = RDS / "univi_bench/runs_multiome_py_1-24-2026"
DATA = RDS / "data"                             # where your .h5mu / 10x files live

for p in [CACHE, WORK, DATA]:
    p.mkdir(parents=True, exist_ok=True)

os.environ["XDG_CACHE_HOME"] = str(CACHE / "xdg")
os.environ["MPLCONFIGDIR"] = str(CACHE / "mpl")
os.environ["NUMBA_CACHE_DIR"] = str(CACHE / "numba")
os.environ["HF_HOME"] = str(CACHE / "hf")
os.environ["TORCH_HOME"] = str(CACHE / "torch")
os.environ["JUPYTER_PLATFORM_DIRS"] = "1"

print("WORK:", WORK)


# %% [markdown]
### Import modules

# %% [code]
import sys
import time
import json
import numpy as np
import pandas as pd

import anndata as ad
import scanpy as sc

from sklearn.feature_extraction.text import TfidfTransformer
from sklearn.decomposition import TruncatedSVD, PCA
from sklearn.preprocessing import StandardScaler, normalize
from sklearn.neighbors import NearestNeighbors
from sklearn.metrics import confusion_matrix, accuracy_score, f1_score, pairwise_distances, adjusted_rand_score, normalized_mutual_info_score, silhouette_score, balanced_accuracy_score
from sklearn.model_selection import train_test_split
from sklearn.cluster import KMeans

import scipy.sparse as sp

import torch
from torch.utils.data import DataLoader, Subset

import matplotlib.pyplot as plt
import seaborn as sns


# %% [code]
from scipy import special

x = np.linspace(0, np.pi, 5)
print(special.sph_legendre_p(2, 1, x))


# %% [code]
import importlib.metadata as im
for pkg in ["jax", "jaxlib"]:
    try:
        dist = im.distribution(pkg)
        print(pkg, dist.version, "installer:", dist.read_text("INSTALLER"))
    except Exception as e:
        print(pkg, "not found:", e)


# %% [code]
import shutil

env_bin = str(Path(sys.executable).resolve().parent)  # .../univi-bench-py/bin
os.environ["PATH"] = env_bin + ":" + os.environ.get("PATH", "")

print("env_bin:", env_bin)
print("bedtools which:", shutil.which("bedtools"))
!bedtools --version


# %% [markdown]
### Universal helper functions - timing, subsampling, saving, etc..

# %% [code]
def now():
    return time.perf_counter()

def set_seed(seed=0):
    import random
    random.seed(seed)
    np.random.seed(seed)
    torch.manual_seed(seed)
    torch.cuda.manual_seed_all(seed)

def subsample_idx(n, k, seed=0):
    if k is None or k >= n:
        return np.arange(n)
    rng = np.random.default_rng(seed)
    return rng.choice(n, size=int(k), replace=False)

def save_npz(path, **arrays):
    path = str(path)
    np.savez_compressed(path, **arrays)
    return path


# %% [markdown]
### Load Multiome data

# %% [code]
DATA_ROOT = Path("/home/groups/precepts/ashforda/UniVI_v2/UniVI_older-non_git/data/PBMC_10x_Multiome_data/10x_Genomics_Multiome_data")

RNA_PATH = DATA_ROOT / "10x-Multiome-Pbmc10k-RNA.h5ad"
ATAC_PATH = DATA_ROOT / "10x-Multiome-Pbmc10k-ATAC.h5ad"

print("RNA file:", RNA_PATH)
print("ATAC file:", ATAC_PATH)


# %% [code]
# Where to save plots/metrics (Path-safe, uses WORK)
BENCH_DIR = WORK / "benchmark_eval"
FIGDIR = BENCH_DIR / "figures"
RUNDIR = WORK / "runs_multiome"   # keep consistent with your earlier variable name

BENCH_DIR.mkdir(parents=True, exist_ok=True)
FIGDIR.mkdir(parents=True, exist_ok=True)
RUNDIR.mkdir(parents=True, exist_ok=True)

# neighbor params
K_MIX = 30
K_LT = 15
#K_LT = 3

# FOSCTTM
FOSCTTM_SUBSAMPLE_N = 3000  # None = all
#RNG_SEED = 0
RNG_SEED = 67

sc.settings.set_figure_params(dpi=200, figsize=(8, 6))

print("BENCH_DIR:", BENCH_DIR)
print("FIGDIR:", FIGDIR)
print("RUNDIR:", RUNDIR)


# %% [code]
rna = sc.read_h5ad(RNA_PATH)
atac = sc.read_h5ad(ATAC_PATH)

print(rna)
print(atac)

print("RNA obs names head:", rna.obs_names[:5].tolist())
print("ATAC obs names head:", atac.obs_names[:5].tolist())

print("paired cells:", rna.n_obs, "shared:", rna.n_obs == atac.n_obs)


# %% [code]
import numpy as np
import scipy.sparse as sp

def _is_integerish(X, *, tol=1e-6, max_check=200_000, seed=0):
    """
    True if values look like integers (within tol). Works for dense/sparse.
    For big matrices, randomly samples up to max_check entries.
    """
    rng = np.random.default_rng(seed)

    if sp.issparse(X):
        data = X.data
        if data.size == 0:
            return True  # all zeros
        # sample nonzeros
        if data.size > max_check:
            idx = rng.choice(data.size, size=max_check, replace=False)
            data = data[idx]
        return np.all(np.abs(data - np.round(data)) <= tol)

    arr = np.asarray(X)
    if arr.size == 0:
        return True
    flat = arr.ravel()
    if flat.size > max_check:
        idx = rng.choice(flat.size, size=max_check, replace=False)
        flat = flat[idx]
    # ignore NaNs if any
    flat = flat[np.isfinite(flat)]
    if flat.size == 0:
        return True
    return np.all(np.abs(flat - np.round(flat)) <= tol)

def _quick_matrix_info(X):
    if sp.issparse(X):
        return {
            "type": type(X).__name__,
            "shape": X.shape,
            "dtype": str(X.dtype),
            "nnz": int(X.nnz),
            "min_nonzero": float(X.data.min()) if X.nnz else 0.0,
            "max_nonzero": float(X.data.max()) if X.nnz else 0.0,
        }
    arr = np.asarray(X)
    return {
        "type": type(arr).__name__,
        "shape": arr.shape,
        "dtype": str(arr.dtype),
        "min": float(np.nanmin(arr)),
        "max": float(np.nanmax(arr)),
    }

def inspect_modality(ad, name):
    print(f"\n=== {name} ===")
    print("layers:", list(ad.layers.keys()))
    print("raw:", ad.raw is not None)
    print("X info:", _quick_matrix_info(ad.X))
    print("X integer-ish?", _is_integerish(ad.X))
    for k in ad.layers.keys():
        print(f"layer {k} info:", _quick_matrix_info(ad.layers[k]))
        print(f"layer {k} integer-ish?", _is_integerish(ad.layers[k]))

inspect_modality(rna, "RNA")
inspect_modality(atac, "ATAC")


# %% [code]
print("RNA layers:", list(rna.layers.keys()))
print("ATAC layers:", list(atac.layers.keys()))
print("RNA raw:", rna.raw is not None)
print("RNA X integer-ish?", _is_integerish(rna.X))
for k in rna.layers.keys():
    print("RNA layer", k, "integer-ish?", _is_integerish(rna.layers[k]))
    

# %% [code]
print(rna)
print(atac)


# %% [code]
LABEL_KEY = "cell_type"  # <-- change to your obs column
assert LABEL_KEY in rna.obs.columns, f"{LABEL_KEY} not in rna.obs"
assert (rna.obs[LABEL_KEY].values == atac.obs[LABEL_KEY].values).all(), "Labels differ between paired modalities!"


# %% [markdown]
### Data preprocessing/splits

# %% [code]
from sklearn.model_selection import train_test_split

def make_shared_splits(n, labels, *, seed=0, train_frac=0.8, val_frac=0.1):
    idx_all = np.arange(n)
    y = np.asarray(labels).astype(str)

    idx_train, idx_tmp, y_train, y_tmp = train_test_split(
        idx_all, y,
        test_size=(1.0 - train_frac),
        random_state=seed,
        stratify=y,
    )

    test_frac = 1.0 - train_frac - val_frac
    val_frac_of_tmp = val_frac / (val_frac + test_frac)

    idx_val, idx_test = train_test_split(
        idx_tmp,
        test_size=(1.0 - val_frac_of_tmp),
        random_state=seed,
        stratify=y_tmp,
    )

    return {"train": np.sort(idx_train), "val": np.sort(idx_val), "test": np.sort(idx_test)}

labels_all = rna.obs[LABEL_KEY].astype(str).to_numpy()
splits = make_shared_splits(rna.n_obs, labels_all, seed=RNG_SEED, train_frac=0.8, val_frac=0.1)

print({k: len(v) for k, v in splits.items()})


# %% [code]
rna[splits["train"]].obs[LABEL_KEY].value_counts().to_frame("n")


# %% [code]
rna[splits["val"]].obs[LABEL_KEY].value_counts().to_frame("n")


# %% [code]
rna[splits["test"]].obs[LABEL_KEY].value_counts().to_frame("n")


# %% [code]
import numpy as np
import pandas as pd
import anndata as ad
import scanpy as sc
import scipy.sparse as sp

from sklearn.feature_extraction.text import TfidfTransformer
from sklearn.decomposition import TruncatedSVD
from sklearn.preprocessing import StandardScaler, normalize


# -------------------------
# small utilities
# -------------------------
def _is_integerish(X) -> bool:
    """Heuristic: treat sparse/dense as integer-ish if all stored values are near integers."""
    if sp.issparse(X):
        d = X.data
        if d.size == 0:
            return True
        return np.all(np.isfinite(d)) and np.all(np.abs(d - np.rint(d)) < 1e-6)
    X = np.asarray(X)
    return np.all(np.isfinite(X)) and np.all(np.abs(X - np.rint(X)) < 1e-6)


def ensure_counts_layer(adata, layer="counts"):
    adata = adata.copy()
    if layer in adata.layers and _is_integerish(adata.layers[layer]):
        return adata
    if adata.raw is not None and _is_integerish(adata.raw.X):
        adata.layers[layer] = adata.raw.X.copy()
        return adata
    adata.layers[layer] = adata.X.copy()
    return adata


def _to_csr_float32(X):
    if sp.issparse(X):
        X = X.tocsr(copy=False)
        return X.astype(np.float32, copy=False) if X.dtype != np.float32 else X
    return sp.csr_matrix(np.asarray(X, dtype=np.float32))


def binarize_csr(X):
    X = X.tocsr(copy=True)
    if X.data.size:
        X.data[:] = 1.0
    return X


# -------------------------
# RNA: HVGs on TRAIN, then (counts subset) + (log1p subset)
# -------------------------
def fit_hvgs_on_train(rna, train_idx, *, counts_layer="counts", n_hvg=2000, seed=0):
    rna = ensure_counts_layer(rna, layer=counts_layer)
    rna_tr = rna[train_idx].copy()
    rna_tr.X = rna_tr.layers[counts_layer]
    try:
        sc.pp.highly_variable_genes(rna_tr, n_top_genes=int(n_hvg), flavor="seurat_v3", layer=None)
    except Exception:
        sc.pp.highly_variable_genes(rna_tr, n_top_genes=int(n_hvg), flavor="seurat", layer=None)

    hvg = rna_tr.var_names[rna_tr.var["highly_variable"].to_numpy()].astype(str).tolist()
    if len(hvg) == 0:
        raise RuntimeError("HVG selection returned 0 genes.")
    return hvg


def transform_rna_log_hvg(rna, hvg, *, counts_layer="counts", target_sum=1e4):
    rna = ensure_counts_layer(rna, layer=counts_layer)
    a = rna[:, hvg].copy()
    a.layers["log1p"] = a.layers[counts_layer].copy()
    sc.pp.normalize_total(a, target_sum=float(target_sum), layer="log1p")
    sc.pp.log1p(a, layer="log1p")
    a.X = a.layers["log1p"]
    return a


# -------------------------
# ATAC: TF-IDF + SVD fit on TRAIN, apply to ALL (LSI)
# -------------------------
def fit_atac_lsi_on_train(
    atac, train_idx, *,
    counts_layer="counts",
    n_lsi=50,
    seed=0,
    tfidf_kwargs=None,
    svd_kwargs=None,
    do_l2_norm=False,
    do_scale=True,
):
    atac = ensure_counts_layer(atac, layer=counts_layer)
    X = _to_csr_float32(atac.layers[counts_layer])

    tfidf_kwargs = tfidf_kwargs or dict(norm=None, use_idf=True, smooth_idf=True, sublinear_tf=False)
    tfidf = TfidfTransformer(**tfidf_kwargs)

    Xtr_t = tfidf.fit_transform(X[train_idx])

    svd_kwargs = svd_kwargs or dict(algorithm="randomized", n_iter=7, random_state=int(seed))
    svd = TruncatedSVD(n_components=int(n_lsi), **svd_kwargs)
    Ztr = svd.fit_transform(Xtr_t)

    if do_l2_norm:
        Ztr = normalize(Ztr, norm="l2", axis=1)

    scaler = None
    if do_scale:
        scaler = StandardScaler(with_mean=True, with_std=True)
        Ztr = scaler.fit_transform(Ztr)

    return tfidf, svd, scaler


def transform_atac_lsi(atac, tfidf, svd, scaler, *, counts_layer="counts", n_lsi=None, do_l2_norm=False):
    atac = ensure_counts_layer(atac, layer=counts_layer)
    X = _to_csr_float32(atac.layers[counts_layer])

    Xt = tfidf.transform(X)
    Z = svd.transform(Xt)

    if do_l2_norm:
        Z = normalize(Z, norm="l2", axis=1)

    if scaler is not None:
        Z = scaler.transform(Z)

    Z = Z.astype(np.float32, copy=False)
    if n_lsi is not None:
        Z = Z[:, : int(n_lsi)].copy()

    atac_lsi = ad.AnnData(
        X=Z,
        obs=atac.obs.copy(),
        var=pd.DataFrame(index=[f"LSI_{i}" for i in range(Z.shape[1])]),
    )
    atac_lsi.obsm["X_lsi"] = Z
    return atac_lsi


# -------------------------
# ATAC peaks for MultiVI/PeakVI/scMoMaT/etc:
# FIXED: select peaks using TRAIN-only detection-rate window + variability
# -------------------------
def fit_peaks_by_detection_window_on_train(
    atac_counts, train_idx, *,
    counts_layer="counts",
    dr_min=0.01,
    dr_max=0.30,
    n_peaks=4000,
    prefer_var="bernoulli",  # "bernoulli" (p*(1-p)) or "sample_var"
):
    """
    Select peaks using TRAIN only:
      1) binarize on the fly
      2) filter by detection rate window (dr_min..dr_max)
      3) rank by variability (default Bernoulli var p(1-p))
    Returns: list of peak names (var_names)
    """
    train_idx = np.asarray(train_idx, dtype=int)
    atac_counts = ensure_counts_layer(atac_counts, layer=counts_layer)

    X = _to_csr_float32(atac_counts.layers[counts_layer])
    Xtr = X[train_idx]
    Xtr_bin = (Xtr > 0).astype(np.float32)

    dr = np.asarray(Xtr_bin.mean(axis=0)).ravel()  # fraction of train cells with peak open
    keep = (dr >= float(dr_min)) & (dr <= float(dr_max))
    keep_idx = np.where(keep)[0]
    if keep_idx.size == 0:
        qs = np.quantile(dr, [0, 0.01, 0.05, 0.1, 0.5, 0.9, 0.95, 0.99, 1.0])
        raise RuntimeError(
            "No peaks pass detection-rate window on TRAIN.\n"
            f"Try loosening dr_min/dr_max. Current dr_min={dr_min}, dr_max={dr_max}\n"
            f"TRAIN detection-rate quantiles: {qs}"
        )

    if str(prefer_var).lower() == "sample_var":
        # variance of binary across train (equivalent-ish to p(1-p), but computed directly)
        # NOTE: toarray is heavy; but for 4k selection, we only do it on kept peaks
        Xk = Xtr_bin[:, keep_idx].toarray()
        score = Xk.var(axis=0)
    else:
        p = dr[keep_idx]
        score = p * (1.0 - p)  # Bernoulli variance

    # take top n_peaks by score
    k = int(min(n_peaks, keep_idx.size))
    order = np.argsort(-score)[:k]
    top = keep_idx[order]
    top = np.sort(top)

    return atac_counts.var_names[top].astype(str).tolist()


def subset_to_chr_features(adata, *, chrom_col_candidates=("chrom", "interval")):
    for col in chrom_col_candidates:
        if col in adata.var.columns:
            s = adata.var[col].astype(str)
            mask = s.str.startswith("chr")
            return adata[:, mask].copy()
    return adata


def summarize_peak_detection(atac_bin, train_idx, *, label="ATAC", n_q=9):
    train_idx = np.asarray(train_idx, dtype=int)
    X = atac_bin.X
    X = X.tocsr(copy=False) if sp.issparse(X) else sp.csr_matrix(np.asarray(X))
    Xtr = X[train_idx]
    dr = np.asarray((Xtr > 0).mean(axis=0)).ravel()
    qs = np.quantile(dr, [0, .01, .05, .10, .50, .90, .95, .99, 1.0])
    print(f"{label} TRAIN detection-rate quantiles (min/1/5/10/50/90/95/99/max): {qs}")
    return dr


# -------------------------
# build everything
# -------------------------
def build_shared_inputs(
    rna, atac, splits, *,
    rna_counts_layer="counts",
    atac_counts_layer="counts",
    n_hvg=2000,
    target_sum=1e4,
    n_lsi=100,
    # peak selection knobs (IMPORTANT)
    n_peaks_multivi=4002,
    dr_min=0.01,
    dr_max=0.30,
    seed=0,
):
    tr = np.asarray(splits["train"], dtype=int)

    # ---- RNA ----
    hvg = fit_hvgs_on_train(rna, tr, counts_layer=rna_counts_layer, n_hvg=n_hvg, seed=seed)

    rna = ensure_counts_layer(rna, layer=rna_counts_layer)
    rna_counts_hvg = rna[:, hvg].copy()
    rna_counts_hvg.X = rna_counts_hvg.layers[rna_counts_layer].copy()

    rna_log_hvg = transform_rna_log_hvg(rna, hvg, counts_layer=rna_counts_layer, target_sum=target_sum)

    # ---- ATAC (counts + LSI) ----
    atac = ensure_counts_layer(atac, layer=atac_counts_layer)
    tfidf, svd, scaler = fit_atac_lsi_on_train(
        atac, tr, counts_layer=atac_counts_layer, n_lsi=n_lsi, seed=seed,
        do_l2_norm=False, do_scale=True,
    )
    atac_lsi = transform_atac_lsi(atac, tfidf, svd, scaler, counts_layer=atac_counts_layer, n_lsi=n_lsi)

    # ---- ATAC (binary counts for peak-based models) ----
    atac_counts_bin = atac.copy()
    X = _to_csr_float32(atac_counts_bin.layers[atac_counts_layer])
    atac_counts_bin.X = binarize_csr(X)

    # ---- FIXED peak selection: TRAIN-only DR window + variability ----
    peaks = fit_peaks_by_detection_window_on_train(
        atac, tr,
        counts_layer=atac_counts_layer,
        dr_min=dr_min,
        dr_max=dr_max,
        n_peaks=n_peaks_multivi,
        prefer_var="bernoulli",
    )
    atac_counts_bin_hv = atac_counts_bin[:, peaks].copy()
    atac_counts_bin_hv = subset_to_chr_features(atac_counts_bin_hv)

    # quick sanity print (so we never again accidentally pick ubiquitous peaks)
    _ = summarize_peak_detection(atac_counts_bin_hv, tr, label="ATAC selected-peaks")

    return dict(
        hvg=hvg,
        peaks=peaks,
        tfidf=tfidf,
        svd=svd,
        scaler=scaler,
        rna_counts_hvg=rna_counts_hvg,
        rna_log_hvg=rna_log_hvg,
        atac_counts_bin=atac_counts_bin,
        atac_counts_bin_hv=atac_counts_bin_hv,
        atac_lsi=atac_lsi,
    )


# ---- build everything ONCE ----
shared = build_shared_inputs(
    rna, atac, splits,
    n_hvg=2000,
    n_peaks_multivi=4002,
    dr_min=0.01,   # try 0.005 if too strict
    dr_max=0.30,   # try 0.20-0.40 depending on PBMC resolution
    n_lsi=101,
    seed=RNG_SEED,
)

rna_counts_hvg      = shared["rna_counts_hvg"]
rna_log_hvg         = shared["rna_log_hvg"]
atac_counts_bin     = shared["atac_counts_bin"]
atac_counts_bin_hv  = shared["atac_counts_bin_hv"]
atac_lsi            = shared["atac_lsi"]

print("ATAC full peaks (bin):", atac_counts_bin.n_vars)
print("ATAC selected peaks (bin):", atac_counts_bin_hv.n_vars)

# Canonical per-method inputs (as you were doing)
rna_univi     = rna_log_hvg
atac_univi    = atac_lsi
atac_multivi  = atac_counts_bin_hv
rna_multimap  = rna_log_hvg
atac_multimap = atac_lsi
atac_peakvi   = atac_counts_bin_hv

print("rna_counts_hvg:", rna_counts_hvg.shape, "X dtype:", rna_counts_hvg.X.dtype)
print("rna_log_hvg:", rna_log_hvg.shape, "X dtype:", rna_log_hvg.X.dtype)
print("atac_counts_bin:", atac_counts_bin.shape, "X type:", type(atac_counts_bin.X))
print("atac_counts_bin_hv:", atac_counts_bin_hv.shape, "X type:", type(atac_counts_bin_hv.X))
print("atac_lsi:", atac_lsi.shape, "X dtype:", atac_lsi.X.dtype, "obsm['X_lsi']:", atac_lsi.obsm["X_lsi"].shape)


# %% [code]
import scipy.sparse as sp
X = atac_counts_bin.X
assert sp.issparse(X)
print("min/max data:", (X.data.min() if X.data.size else 0), (X.data.max() if X.data.size else 0))
print("unique values in data (up to 10):", np.unique(X.data)[:10])
# Expect max==1.0 and unique subset of {1.0}


# %% [markdown]
## Unified "runner" interface for each method

# %% [code]
# Helper for some of the outputs of all the functions
def _standard_extra_json(*, transductive: bool, uses_labels: bool, **kw):
    d = dict(transductive=bool(transductive), uses_labels=bool(uses_labels))
    d.update(kw)
    return d


# %% [code]
def standard_flags(*, transductive: bool, uses_labels: bool, **extra):
    """
    Standard metadata for fairness/comparability.
    transductive=True  -> model fit used non-train (e.g., all cells incl test/val)
    uses_labels=True   -> ground-truth labels used during training (semi-supervised)
    """
    d = {"transductive": bool(transductive), "uses_labels": bool(uses_labels)}
    d.update(extra)
    return d


def ensure_flags(out: dict, *, default_transductive=True, default_uses_labels=False):
    """
    Guarantees flags exist, even if a runner forgot.
    """
    out = dict(out) if out is not None else {}
    ej = dict(out.get("extra_json", {}) or {})
    ej.setdefault("transductive", bool(default_transductive))
    ej.setdefault("uses_labels", bool(default_uses_labels))
    out["extra_json"] = ej
    return out


# %% [markdown]
### 1) UniVI

# %% [code]
def run_univi(rna_adata, atac_adata, *, out_dir, splits, seed=0, X_key="X", loss_mode="v1"):
    set_seed(seed)
    out_dir = Path(out_dir); out_dir.mkdir(parents=True, exist_ok=True)

    import univi
    from univi import UniVIConfig, ModalityConfig, TrainingConfig, UniVIMultiModalVAE
    from univi.data import MultiModalDataset, align_paired_obs_names
    from univi.trainer import UniVITrainer

    adata_dict = {"rna": rna_adata, "atac": atac_adata}
    adata_dict = align_paired_obs_names(adata_dict)
    rna_adata = adata_dict["rna"]
    atac_adata = adata_dict["atac"]

    cfg = UniVIConfig(
        latent_dim=30,
        #beta=1.0, gamma=1.25,
        #beta=1.25, gamma=4.25,
        # Best so far:
        beta=1.35, gamma=3.75,
        #beta=1.45, gamma=3.25,
        # Best so far:
        #encoder_dropout=0.05, decoder_dropout=0.00,
        encoder_dropout=0.10, decoder_dropout=0.05,
        encoder_batchnorm=True, decoder_batchnorm=False,
        #encoder_batchnorm=False, decoder_batchnorm=False,
        #kl_anneal_start=0, kl_anneal_end=15,
        #align_anneal_start=10, align_anneal_end=25,
        # Best so far:
        kl_anneal_start=25, kl_anneal_end=75,
        align_anneal_start=45, align_anneal_end=95,
        #kl_anneal_start=0, kl_anneal_end=30,
        #align_anneal_start=15, align_anneal_end=45,
        modalities=[
            ModalityConfig("rna",  rna_adata.n_vars,  [512, 256, 128], [128, 256, 512], likelihood="gaussian"),
            ModalityConfig("atac", atac_adata.n_vars, [256, 128, 64],  [64, 128, 256],  likelihood="gaussian"),
        ],
    )

    device = "cuda" if torch.cuda.is_available() else "cpu"

    train_cfg = TrainingConfig(
        n_epochs=200, batch_size=256, lr=1e-3, weight_decay=1e-4,
        device=device, early_stopping=True, patience=50, log_every=100
    )

    dataset = MultiModalDataset(adata_dict=adata_dict, X_key=X_key, device=None)

    train_loader = DataLoader(Subset(dataset, splits["train"]), batch_size=train_cfg.batch_size, shuffle=True,  num_workers=0)
    val_loader   = DataLoader(Subset(dataset, splits["val"]),   batch_size=train_cfg.batch_size, shuffle=False, num_workers=0)

    model = UniVIMultiModalVAE(cfg, loss_mode=loss_mode, v1_recon="avg", normalize_v1_terms=True).to(device)
    #model = UniVIMultiModalVAE(cfg, loss_mode=loss_mode, v1_recon="avg", recon_normalize_by_dim=True, recon_dim_power=0.40, normalize_v1_terms=True).to(device)
    
    trainer = UniVITrainer(model=model, train_loader=train_loader, val_loader=val_loader, train_cfg=train_cfg, device=device)

    t0 = now()
    _  = trainer.fit()
    t1 = now()

    # Encode ALL cells (evaluation will slice test)
    model.eval()
    full_loader = DataLoader(dataset, batch_size=train_cfg.batch_size, shuffle=False, num_workers=0)

    Z_rna, Z_atac, Z_fused = [], [], []
    with torch.no_grad():
        for batch in full_loader:
            x_dict = {k: v.to(device) for k, v in batch.items()}
            mu_dict, _ = model.encode_modalities(x_dict)
            Z_rna.append(mu_dict["rna"].detach().cpu().numpy())
            Z_atac.append(mu_dict["atac"].detach().cpu().numpy())
            mu_z, _, _ = model.encode_fused(x_dict, use_mean=True)
            Z_fused.append(mu_z.detach().cpu().numpy())
            
    fit_seconds = float(t1 - t0)
    
    return ensure_flags({
        "Z_rna": np.concatenate(Z_rna, axis=0),
        "Z_atac": np.concatenate(Z_atac, axis=0),
        "Z_fused": np.concatenate(Z_fused, axis=0),
        "fit_seconds": fit_seconds,
        "extra_json": standard_flags(transductive=False, uses_labels=False),
    })


# %% [markdown]
### 2) MultiVI (scvi-tools)

# %% [code]
import os, sys, glob, importlib.util

print("CWD:", os.getcwd())
print("sys.path[0]:", sys.path[0])
print("Any local jax* files?:", glob.glob("jax*") + glob.glob("**/jax*", recursive=False))

spec = importlib.util.find_spec("jax")
print("jax spec:", spec)
print("jax origin:", None if spec is None else spec.origin)

spec2 = importlib.util.find_spec("jaxlib")
print("jaxlib spec:", spec2)
print("jaxlib origin:", None if spec2 is None else spec2.origin)


# %% [code]
def scvi_train_with_patience(
    model,
    *,
    max_epochs=200,
    patience=50,
    monitor="elbo_validation",
    mode="min",
    **train_kwargs,
):
    import inspect
    from lightning.pytorch.callbacks import EarlyStopping

    sig = inspect.signature(model.train)

    # 1) Force scvi internal early stopping OFF (prevents TrainRunner double-passing)
    if "early_stopping" in sig.parameters:
        train_kwargs["early_stopping"] = False

    # 2) Remove early-stopping args from the top-level kwargs
    for k in ("early_stopping_patience", "early_stopping_monitor", "early_stopping_min_delta"):
        train_kwargs.pop(k, None)

    # 3) ALSO remove them from scvi plan kwargs containers (where duplicates often come from)
    for plan_key in ("plan_kwargs", "training_plan_kwargs"):
        if plan_key in sig.parameters and plan_key in train_kwargs and train_kwargs[plan_key] is not None:
            d = dict(train_kwargs[plan_key])
            for k in ("early_stopping_patience", "early_stopping_monitor", "early_stopping_min_delta"):
                d.pop(k, None)
            train_kwargs[plan_key] = d

    # 4) Add Lightning EarlyStopping callback via trainer_kwargs (or callbacks if supported)
    cb = EarlyStopping(monitor=monitor, patience=int(patience), mode=mode)

    if "trainer_kwargs" in sig.parameters:
        tk = dict(train_kwargs.get("trainer_kwargs") or {})
        cbs = list(tk.get("callbacks") or [])
        cbs.append(cb)
        tk["callbacks"] = cbs
        train_kwargs["trainer_kwargs"] = tk
    elif "callbacks" in sig.parameters:
        cbs = list(train_kwargs.get("callbacks") or [])
        cbs.append(cb)
        train_kwargs["callbacks"] = cbs
    else:
        # last resort: just train without early stopping rather than crashing
        # (you can still set max_epochs lower)
        pass

    return model.train(max_epochs=int(max_epochs), **train_kwargs)

def multivi_latents(model, mdata):
    try:
        Z_joint = np.asarray(model.get_latent_representation(mdata, modality="joint"))
    except Exception:
        Z_joint = np.asarray(model.get_latent_representation(mdata))
    Z_expr = np.asarray(model.get_latent_representation(mdata, modality="rna"))
    Z_acc  = np.asarray(model.get_latent_representation(mdata, modality="atac"))
    return Z_expr, Z_acc, Z_joint


# %% [code]
def setup_multivi_mudata(MULTIVI, mdata):
    import inspect
    sig = inspect.signature(MULTIVI.setup_mudata)

    # try the most common patterns
    if "rna_layer" in sig.parameters or "atac_layer" in sig.parameters:
        return MULTIVI.setup_mudata(mdata, rna_layer=None, atac_layer=None)

    if "modalities" in sig.parameters:
        # often expects mapping from modality type -> mudata key
        return MULTIVI.setup_mudata(mdata, modalities={"expression": "rna", "accessibility": "atac"})

    # last resort
    return MULTIVI.setup_mudata(mdata)


# %% [code]
from __future__ import annotations

from pathlib import Path
import time
import numpy as np

def _align_by_obs_names(a, b):
    """
    Return (a2, b2) restricted to overlapping obs_names, **preserving `a`'s order**.
    This avoids index drift (e.g., from np.intersect1d sorting) and keeps splits valid
    as long as they were made on `a`'s current order.
    """
    a_names = np.asarray(a.obs_names, dtype=str)
    b_names = np.asarray(b.obs_names, dtype=str)

    b_set = set(b_names.tolist())

    # Preserve the order of `a` (critical!)
    common_in_a_order = [x for x in a_names if x in b_set]
    if len(common_in_a_order) == 0:
        raise ValueError("RNA/ATAC have zero overlapping obs_names. Cannot run paired methods.")

    a2 = a[common_in_a_order].copy()
    b2 = b[common_in_a_order].copy()

    # Final sanity: exact same names + order
    if not np.array_equal(np.asarray(a2.obs_names, dtype=str), np.asarray(b2.obs_names, dtype=str)):
        raise RuntimeError("Post-align mismatch in obs_names order between RNA and ATAC.")

    return a2, b2
    

def run_multivi(
    adata_rna,
    adata_atac,
    *,
    out_dir,
    splits,
    seed=0,
    n_latent=30,
    max_epochs=200,
    patience=50,
    batch_key=None,
    rna_layer="counts",
    atac_layer="counts",
    encode_batch_size=256,
):
    """
    scvi-tools MULTIVI runner (leakage-aware):
      - trains ONLY on (train+val)
      - reloads model onto FULL data to compute embeddings for ALL cells
      - returns Z_fused and (if supported) Z_rna / Z_atac
    """
    import inspect
    import shutil
    import muon as mu
    import scvi
    from scvi.model import MULTIVI

    t0 = time.perf_counter()
    out_dir = Path(out_dir)
    out_dir.mkdir(parents=True, exist_ok=True)

    scvi.settings.seed = int(seed)

    # -------------------------
    # align cells by obs_names
    # -------------------------
    rna0, atac0 = _align_by_obs_names(adata_rna, adata_atac)

    # -------------------------
    # build FULL MuData
    # -------------------------
    rna = rna0.copy()
    atac = atac0.copy()

    # Only create layers if they don't already exist
    if rna_layer is not None and rna_layer not in rna.layers:
        rna.layers[rna_layer] = rna.X.copy()
    if atac_layer is not None and atac_layer not in atac.layers:
        atac.layers[atac_layer] = atac.X.copy()

    m_full = mu.MuData({"rna": rna, "atac": atac})
    m_full.update()

    # -------------------------
    # define FIT idx = train + val (your split)
    # -------------------------
    tr = np.asarray(splits["train"], dtype=int)
    va = np.asarray(splits.get("val", []), dtype=int)
    fit_idx = np.concatenate([tr, va]) if va.size else tr
    fit_idx = np.asarray(fit_idx, dtype=int)

    # Make FIT mudata (no test cells seen by training)
    m_fit = m_full[fit_idx].copy()
    m_fit.update()

    # -------------------------
    # setup_mudata (API drift tolerant)
    # -------------------------
    setup_err = None
    try:
        MULTIVI.setup_mudata(
            m_fit,
            rna_layer=rna_layer,
            atac_layer=atac_layer,
            batch_key=batch_key,
        )
    except Exception as e1:
        setup_err = e1
        # Some versions require modalities mapping
        try:
            modalities = {"rna_layer": "rna", "atac_layer": "atac"}
            MULTIVI.setup_mudata(
                m_fit,
                rna_layer=rna_layer,
                atac_layer=atac_layer,
                batch_key=batch_key,
                modalities=modalities,
            )
        except Exception as e2:
            raise RuntimeError(
                "MultiVI setup_mudata failed.\n"
                f"m_fit.mod keys={list(m_fit.mod.keys())}\n"
                f"Original error: {repr(setup_err)}\n"
                f"Fallback error: {repr(e2)}"
            ) from e2

    # -------------------------
    # train on FIT only
    # -------------------------
    model = MULTIVI(m_fit, n_latent=int(n_latent))

    # Prefer your helper if it exists; else fall back to model.train
    if "scvi_train_with_patience" in globals():
        scvi_train_with_patience(
            model,
            max_epochs=max_epochs,
            patience=patience,
            monitor="elbo_validation",
            mode="min",
            check_val_every_n_epoch=1,
        )
    else:
        # pure scvi fallback
        train_sig = inspect.signature(model.train)
        train_kwargs = dict(
            max_epochs=int(max_epochs),
            early_stopping=True,
            check_val_every_n_epoch=1,
        )
        # only pass patience if supported
        for k in ("early_stopping_patience", "patience"):
            if k in train_sig.parameters:
                train_kwargs[k] = int(patience)
                break
        model.train(**train_kwargs)

    # -------------------------
    # save + reload onto FULL data to encode ALL cells
    # (this is the key to leakage-aware training but full-data embedding)
    # -------------------------
    save_dir = out_dir / "multivi_model"
    if save_dir.exists():
        shutil.rmtree(save_dir)
    model.save(str(save_dir), overwrite=True)

    # Setup FULL mudata with same config *before* loading (important)
    # If you don't do this, some versions won't attach the right AnnDataManager.
    setup_err2 = None
    try:
        MULTIVI.setup_mudata(
            m_full,
            rna_layer=rna_layer,
            atac_layer=atac_layer,
            batch_key=batch_key,
        )
    except Exception as e1:
        setup_err2 = e1
        try:
            modalities = {"rna_layer": "rna", "atac_layer": "atac"}
            MULTIVI.setup_mudata(
                m_full,
                rna_layer=rna_layer,
                atac_layer=atac_layer,
                batch_key=batch_key,
                modalities=modalities,
            )
        except Exception as e2:
            raise RuntimeError(
                "MultiVI setup_mudata on FULL data failed (needed for reload).\n"
                f"Original error: {repr(setup_err2)}\n"
                f"Fallback error: {repr(e2)}"
            ) from e2

    # load signature differs across scvi versions
    load_fn = getattr(MULTIVI, "load", None)
    if load_fn is None or not callable(load_fn):
        raise RuntimeError("Your scvi-tools MULTIVI has no .load(...). Cannot reload onto full data.")

    load_sig = inspect.signature(load_fn)
    load_kwargs = {}
    # common: MULTIVI.load(path, adata=...) OR MULTIVI.load(path, mudata=...)
    if "adata" in load_sig.parameters:
        load_kwargs["adata"] = m_full
    elif "mudata" in load_sig.parameters:
        load_kwargs["mudata"] = m_full
    else:
        # worst case: load(path, ...) but no adata param exposed
        # try calling without and hope it was saved with data
        load_kwargs = {}

    model_full = MULTIVI.load(str(save_dir), **load_kwargs)

    # -------------------------
    # latent extraction (API drift tolerant)
    # -------------------------
    def _get_latent(modality=None):
        fn = model_full.get_latent_representation
        sig = inspect.signature(fn)
        kwargs = {}
        if "adata" in sig.parameters:
            kwargs["adata"] = m_full
        if "batch_size" in sig.parameters:
            kwargs["batch_size"] = int(encode_batch_size)
        if "modality" in sig.parameters and modality is not None:
            kwargs["modality"] = modality
        return np.asarray(fn(**kwargs), dtype=np.float32)

    #Z_fused = _get_latent(modality=None)
    # prefer true joint
    try:
        Z_fused = _get_latent(modality="joint")
    except Exception:
        Z_fused = _get_latent(modality=None)
    
    Z_rna = None
    Z_atac = None
    try:
        #Z_rna = _get_latent(modality="rna")
        Z_rna = _get_latent(modality="expression")
    except Exception:
        pass
    try:
        #Z_atac = _get_latent(modality="atac")
        Z_atac = _get_latent(modality="accessibility")
    except Exception:
        pass

    fit_seconds = float(time.perf_counter() - t0)

    return ensure_flags({
        "Z_rna": Z_rna,
        "Z_atac": Z_atac,
        "Z_fused": Z_fused,
        "fit_seconds": fit_seconds,
        "extra_json": standard_flags(
            transductive=False,  # <-- IMPORTANT: trained on train+val only
            uses_labels=False,
            note="Trained on train+val only; model reloaded onto full data to encode all cells."
        ),
    }, default_transductive=False, default_uses_labels=False)


# %% [markdown]
### 3) scGLUE (RNA-ATAC integration)

# %% [markdown]
#### a) Calculate guidance-graph

# %% [code]
import re
import gzip

def _open_textmaybe_gz(path):
    path = str(path)
    if path.endswith(".gz"):
        return gzip.open(path, "rt")
    return open(path, "rt")

def parse_peaks_from_var(atac, *, chrom_col=None, start_col=None, end_col=None):
    """
    Returns DataFrame with columns: peak, chrom, start, end, mid
    Supports:
      - atac.var has chrom/start/end columns
      - atac.var_names formatted like 'chr1:123-456' or 'chr1_123_456'
    """
    var = atac.var.copy()
    if chrom_col and start_col and end_col and all(c in var.columns for c in [chrom_col, start_col, end_col]):
        df = pd.DataFrame({
            "peak": atac.var_names.astype(str),
            "chrom": var[chrom_col].astype(str).values,
            "start": var[start_col].astype(int).values,
            "end":   var[end_col].astype(int).values,
        })
    elif all(c in var.columns for c in ["chrom", "start", "end"]):
        df = pd.DataFrame({
            "peak": atac.var_names.astype(str),
            "chrom": var["chrom"].astype(str).values,
            "start": var["start"].astype(int).values,
            "end":   var["end"].astype(int).values,
        })
    else:
        peaks = atac.var_names.astype(str)
        chrom = []
        start = []
        end = []
        for p in peaks:
            m = re.match(r"^(chr[^:_]+)[:_](\d+)[\-_](\d+)$", p)
            if m is None:
                raise ValueError(f"Can't parse peak name: {p!r}. Provide chrom/start/end columns in atac.var.")
            chrom.append(m.group(1))
            start.append(int(m.group(2)))
            end.append(int(m.group(3)))
        df = pd.DataFrame({"peak": peaks, "chrom": chrom, "start": start, "end": end})

    df["mid"] = ((df["start"].values + df["end"].values) // 2).astype(int)
    return df

def parse_gtf_genes(gtf_path, *, gene_id_key="gene_id", gene_name_key="gene_name"):
    """
    Minimal GTF parser: returns DataFrame for 'gene' features with:
    gene_id, gene_name, chrom, start, end, strand, tss
    """
    rows = []
    with _open_textmaybe_gz(gtf_path) as f:
        for line in f:
            if not line or line.startswith("#"):
                continue
            parts = line.rstrip("\n").split("\t")
            if len(parts) != 9:
                continue
            chrom, source, feature, start, end, score, strand, frame, attrs = parts
            if feature != "gene":
                continue

            # attribute parsing: key "value";
            def get_attr(key):
                m = re.search(rf'{re.escape(key)}\s+"([^"]+)"', attrs)
                return m.group(1) if m else None

            gid = get_attr(gene_id_key)
            gname = get_attr(gene_name_key)

            if gid is None and gname is None:
                continue

            start_i = int(start)
            end_i = int(end)
            tss = start_i if strand == "+" else end_i

            rows.append({
                "gene_id": gid,
                "gene_name": gname,
                "chrom": chrom,
                "start": start_i,
                "end": end_i,
                "strand": strand,
                "tss": int(tss),
            })

    genes = pd.DataFrame(rows)
    if genes.empty:
        raise ValueError(f"No gene records parsed from: {gtf_path}")
    return genes

import numpy as np
import scipy.sparse as sp

def restrict_features_to_fit_cells(rna_counts, atac_counts, fit_idx):
    """
    Leakage-aware feature filtering:
    keep only genes/peaks that are observed (nnz>0) in FIT cells,
    then apply that feature subset to ALL cells.
    """
    fit_idx = np.asarray(fit_idx)

    rna_fit = rna_counts[fit_idx]
    atac_fit = atac_counts[fit_idx]

    Xr = rna_fit.X
    Xa = atac_fit.X

    if sp.issparse(Xr):
        gene_nnz = np.asarray((Xr > 0).sum(axis=0)).ravel()
    else:
        gene_nnz = (np.asarray(Xr) > 0).sum(axis=0)

    if sp.issparse(Xa):
        peak_nnz = np.asarray((Xa > 0).sum(axis=0)).ravel()
    else:
        peak_nnz = (np.asarray(Xa) > 0).sum(axis=0)

    keep_genes = gene_nnz > 0
    keep_peaks = peak_nnz > 0

    # subset ALL cells to the features discovered on FIT cells
    rna2 = rna_counts[:, keep_genes].copy()
    atac2 = atac_counts[:, keep_peaks].copy()

    return rna2, atac2


# %% [markdown]
You need the guidance graph node names to match rna.var_names and atac.var_names.
- If rna.var_names are Ensembl IDs → use gene_id.
- If rna.var_names are gene symbols → use gene_name.
- If you have both in rna.var, you can map.

# %% [code]
def pick_gene_key(rna):
    # Heuristic: Ensembl IDs often start with ENS
    sample = rna.var_names.astype(str)[:100]
    ens_like = np.mean([s.startswith("ENS") for s in sample])
    return "gene_id" if ens_like > 0.5 else "gene_name"

GENE_KEY = pick_gene_key(rna)
print("Using GENE_KEY:", GENE_KEY, "(must match rna.var_names)")


# %% [code]
print(rna.var_names)
print(GENE_KEY)


# %% [code]
# 1) Are var_names unique?
print("rna var_names unique:", rna.var_names.is_unique)

# 2) After annotation, how many are missing?
cols = ["chrom", "chromStart", "chromEnd"]
print("missing coords:", rna.var[cols].isna().any(axis=1).sum(), "of", rna.n_vars)


# %% [markdown]
Build a bounded distance-based peak↔gene graph

# %% [code]
def parse_peaks_from_var(atac, chrom_col=None, start_col=None, end_col=None):
    # If you *already* have structured columns, prefer them
    if chrom_col in (atac.var.columns if hasattr(atac, "var") else []) and \
       start_col in atac.var.columns and end_col in atac.var.columns:
        df = atac.var[[chrom_col, start_col, end_col]].copy()
        df.columns = ["chrom", "start", "end"]
        df["start"] = df["start"].astype(int)
        df["end"] = df["end"].astype(int)
        df["peak"] = atac.var_names.astype(str)
    else:
        peaks = atac.var_names.astype(str)

        chrom, start, end = [], [], []
        # accepts: chr1:1-2, 1:1-2, KI270727.1:52331-52752, chr1_1-2, etc.
        pat = re.compile(r"^([^:_]+)[:_](\d+)[\-_](\d+)$")

        for p in peaks:
            m = pat.match(p)
            if m is None:
                raise ValueError(
                    f"Can't parse peak name: {p!r}. "
                    f"Expected something like 'chr1:100-200' or 'KI270727.1:100-200'."
                )
            chrom.append(m.group(1))
            start.append(int(m.group(2)))
            end.append(int(m.group(3)))

        df = pd.DataFrame({"chrom": chrom, "start": start, "end": end, "peak": peaks})

    df["mid"] = ((df["start"].values + df["end"].values) // 2).astype(int)
    return df


# %% [code]
def harmonize_chr_prefix(genes, peaks):
    genes_has_chr = (genes["chrom"].astype(str).str.startswith("chr")).mean() > 0.5
    peaks_has_chr = (peaks["chrom"].astype(str).str.startswith("chr")).mean() > 0.5

    if genes_has_chr and not peaks_has_chr:
        # add chr to canonical chroms only
        canon = re.compile(r"^(?:\d+|X|Y|M|MT)$")
        peaks["chrom"] = peaks["chrom"].astype(str).map(lambda c: ("chr" + c) if canon.match(c) else c)

    elif peaks_has_chr and not genes_has_chr:
        genes["chrom"] = genes["chrom"].astype(str).str.replace(r"^chr", "", regex=True)

    return genes, peaks


# %% [code]
import networkx as nx

def build_guidance_graph_distance(
    rna, atac, gtf_path,
    *,
    window=200_000,
    k_nearest=10,
    promoter_width=3_000,
    gene_id_key="gene_id",
    gene_name_key="gene_name",
    gene_key_for_nodes=None,
):
    """
    Build a guidance graph (networkx.Graph) where nodes are:
      - genes (named exactly like rna.var_names)
      - peaks (named exactly like atac.var_names)

    Edges: peak -- gene with attributes:
      weight: float
      sign: +1
      distance: int
      in_promoter: bool
    """
    gene_key_for_nodes = gene_key_for_nodes or pick_gene_key(rna)

    # Parse and restrict genes to those present in RNA
    genes = parse_gtf_genes(gtf_path, gene_id_key=gene_id_key, gene_name_key=gene_name_key)

    # Which column will be the "node name" to match rna.var_names?
    node_col = "gene_id" if gene_key_for_nodes == "gene_id" else "gene_name"
    genes = genes.dropna(subset=[node_col]).copy()
    genes["node"] = genes[node_col].astype(str)

    present_genes = pd.Index(rna.var_names.astype(str))
    genes = genes[genes["node"].isin(present_genes)].copy()
    if genes.empty:
        raise ValueError("No genes matched between GTF and rna.var_names. Check GENE_KEY / annotation.")

    # Parse peaks
    peaks = parse_peaks_from_var(atac)
    peaks["node"] = peaks["peak"].astype(str)

    # >>> ADD THIS BLOCK RIGHT HERE <<<
    genes, peaks = harmonize_chr_prefix(genes, peaks)

    # Optional but helpful: keep only peaks on chroms present in the GTF genes
    valid_chroms = set(genes["chrom"].astype(str).unique())
    peaks = peaks[peaks["chrom"].astype(str).isin(valid_chroms)].copy()

    # Build edges efficiently by chromosome with sorting + two-pointer
    G = nx.Graph()
    # add nodes
    for g in genes["node"].values:
        G.add_node(g, kind="gene")
    for p in peaks["node"].values:
        G.add_node(p, kind="peak")

    genes_by_chr = {c: df.sort_values("tss") for c, df in genes.groupby("chrom", sort=False)}
    peaks_by_chr = {c: df.sort_values("mid") for c, df in peaks.groupby("chrom", sort=False)}

    def _edge_weight(dist):
        # smooth decay; tweak if you like
        return float(np.exp(-dist / 50_000))

    total_edges = 0
    for chrom, pdf in peaks_by_chr.items():
        gdf = genes_by_chr.get(chrom, None)
        if gdf is None or gdf.empty or pdf.empty:
            continue

        g_tss = gdf["tss"].values
        g_nodes = gdf["node"].values

        # for each peak, find genes within window using binary search range
        for peak_node, mid in zip(pdf["node"].values, pdf["mid"].values):
            lo = np.searchsorted(g_tss, mid - window, side="left")
            hi = np.searchsorted(g_tss, mid + window, side="right")
            if hi <= lo:
                continue

            # distances for candidates
            cand_tss = g_tss[lo:hi]
            cand_nodes = g_nodes[lo:hi]
            dists = np.abs(cand_tss - mid)

            # choose up to k nearest
            if len(dists) > k_nearest:
                idx = np.argpartition(dists, k_nearest)[:k_nearest]
                dists = dists[idx]
                cand_nodes = cand_nodes[idx]

            # add edges
            for gene_node, dist in zip(cand_nodes, dists):
                in_prom = bool(dist <= promoter_width)
                w = _edge_weight(int(dist))
                if in_prom:
                    w = float(min(1.0, w * 2.0))  # bump promoter edges

                G.add_edge(
                    peak_node, gene_node,
                    weight=w,
                    sign=1,
                    distance=int(dist),
                    in_promoter=in_prom,
                )
                total_edges += 1

    if total_edges == 0:
        raise ValueError("Built graph has 0 edges. Likely chr naming mismatch (e.g. '1' vs 'chr1').")

    return G

# TODO: set your GTF path
GTF = Path("/home/groups/precepts/ashforda/scOPE_github_stuff/data/reference/Homo_sapiens_GRCh38.p13.gencode.annotation.gtf")

guidance_graph = build_guidance_graph_distance(
    rna_counts_hvg, atac_counts_bin, GTF,
    window=200_000,
    k_nearest=15,
    promoter_width=3_000,
    gene_key_for_nodes=pick_gene_key(rna_counts_hvg),
)

'''
# build graph on HV peaks instead of full peaks
guidance_graph = build_guidance_graph_distance(
    rna_counts_hvg, atac_counts_bin_hv, GTF,
    window=150_000, k_nearest=5, promoter_width=2_000,
    gene_key_for_nodes=pick_gene_key(rna_counts_hvg),
)
'''

print("Graph nodes prior to adding self-loops:", guidance_graph.number_of_nodes())
print("Graph edges prior to adding self-loops:", guidance_graph.number_of_edges())

def add_self_loops(G):
    G = G.copy()
    for n in G.nodes():
        if not G.has_edge(n, n):
            G.add_edge(n, n, weight=1.0, sign=1)
    return G

guidance_graph = add_self_loops(guidance_graph)

print("Graph nodes:", guidance_graph.number_of_nodes())
print("Graph edges:", guidance_graph.number_of_edges())


# %% [markdown]
(optional) save/reload the guidance graph

# %% [code]
import pickle

GG_PATH = WORK / "scglue_guidance_graph.pkl"

with open(GG_PATH, "wb") as f:
    pickle.dump(guidance_graph, f)

print("Saved:", GG_PATH)


# %% [markdown]
#### b) Run scGLUE

# %% [code]
def subset_to_graph(rna_adata, atac_adata, G):
    gene_nodes = [n for n, a in G.nodes(data=True) if a.get("kind") == "gene"]
    peak_nodes = [n for n, a in G.nodes(data=True) if a.get("kind") == "peak"]

    rna2  = rna_adata[:, rna_adata.var_names.isin(gene_nodes)].copy()
    atac2 = atac_adata[:, atac_adata.var_names.isin(peak_nodes)].copy()

    if rna2.n_vars == 0 or atac2.n_vars == 0:
        raise ValueError(f"After subsetting to graph nodes: rna vars={rna2.n_vars}, atac vars={atac2.n_vars}")

    return rna2, atac2


# %% [code]
import numpy as np
import scipy.sparse as sp

def compute_hvg_on_fit_cells(rna, fit_idx, *, n_top_genes=2000, flavor="seurat_v3", batch_key=None):
    """
    Compute HVGs using FIT cells only, then write rna.var['highly_variable'] for ALL cells.
    """
    import scanpy as sc
    fit_idx = np.asarray(fit_idx)

    rna_fit = rna[fit_idx].copy()
    sc.pp.highly_variable_genes(
        rna_fit,
        n_top_genes=int(min(n_top_genes, rna_fit.n_vars)),
        flavor=flavor,
        batch_key=batch_key,
        subset=False,
        inplace=True,
    )
    hv = rna_fit.var["highly_variable"].reindex(rna.var_names).fillna(False).astype(bool).values
    rna.var["highly_variable"] = hv
    return hv


def fit_pca_train_apply_all(rna, fit_idx, *, n_pca=100, target_sum=1e4, seed=0, layer="counts"):
    """
    Fit scaler+PCA on FIT cells only, then apply to ALL cells, storing rna.obsm['X_pca'].
    Uses rna.layers[layer] if present, else rna.X.
    """
    import scanpy as sc
    from sklearn.decomposition import PCA
    from sklearn.preprocessing import StandardScaler

    fit_idx = np.asarray(fit_idx)

    tmp = rna.copy()
    if layer is not None and layer in tmp.layers:
        tmp.X = tmp.layers[layer].copy()
    else:
        tmp.X = tmp.X.copy()

    sc.pp.normalize_total(tmp, target_sum=target_sum)
    sc.pp.log1p(tmp)

    X = tmp.X.toarray() if sp.issparse(tmp.X) else np.asarray(tmp.X)
    X = np.asarray(X, dtype=np.float32)

    scaler = StandardScaler(with_mean=True, with_std=True)
    X_fit = scaler.fit_transform(X[fit_idx])

    ncomp = int(min(n_pca, max(2, X_fit.shape[1] - 1)))
    pca = PCA(n_components=ncomp, random_state=int(seed))
    pca.fit(X_fit)

    rna.obsm["X_pca"] = pca.transform(scaler.transform(X)).astype(np.float32, copy=False)
    return scaler, pca


def _tfidf_fit_transform(X_fit_csr: sp.csr_matrix):
    from sklearn.preprocessing import normalize
    tf = normalize(X_fit_csr, norm="l1", axis=1)
    df = np.asarray((X_fit_csr > 0).sum(axis=0)).ravel().astype(np.float64)
    N = X_fit_csr.shape[0]
    idf = np.log1p(N / (1.0 + df))
    return tf.multiply(idf).tocsr(), idf


def _tfidf_transform(X_all_csr: sp.csr_matrix, idf: np.ndarray):
    from sklearn.preprocessing import normalize
    tf = normalize(X_all_csr, norm="l1", axis=1)
    return tf.multiply(idf).tocsr()


def compute_hv_peaks_on_fit_cells(atac, fit_idx, *, n_top_peaks=20000):
    """
    Compute HV peaks using FIT cells only (variance in TF-IDF space),
    then write atac.var['highly_variable'] for ALL cells.
    """
    fit_idx = np.asarray(fit_idx)

    X_all = atac.X
    if not sp.issparse(X_all):
        X_all = sp.csr_matrix(np.asarray(X_all))
    else:
        X_all = X_all.tocsr(copy=False)
    X_all = X_all.astype(np.float32, copy=False)

    X_fit = X_all[fit_idx]
    X_tfidf_fit, _idf = _tfidf_fit_transform(X_fit)

    mean = np.asarray(X_tfidf_fit.mean(axis=0)).ravel()
    mean2 = np.asarray(X_tfidf_fit.power(2).mean(axis=0)).ravel()
    var = np.maximum(mean2 - mean**2, 0.0)

    n_top = int(min(n_top_peaks, atac.n_vars))
    top_idx = np.argpartition(var, -n_top)[-n_top:]
    hv = np.zeros(atac.n_vars, dtype=bool)
    hv[top_idx] = True

    atac.var["highly_variable"] = hv
    return hv


def fit_lsi_train_apply_all(atac, fit_idx, *, n_lsi=100, seed=0, l2norm=True, drop_first=True):
    """
    Fit TF-IDF + TruncatedSVD on FIT cells only, apply to ALL cells, store atac.obsm['X_lsi'].

    If drop_first=True, we fit n_lsi+1 components and drop component 0 (LSI1),
    returning exactly n_lsi dims in atac.obsm['X_lsi'].
    """
    from sklearn.decomposition import TruncatedSVD
    from sklearn.preprocessing import normalize
    import numpy as np
    import scipy.sparse as sp

    fit_idx = np.asarray(fit_idx, dtype=int)

    X_all = atac.X
    if not sp.issparse(X_all):
        X_all = sp.csr_matrix(np.asarray(X_all))
    else:
        X_all = X_all.tocsr(copy=False)
    X_all = X_all.astype(np.float32, copy=False)

    X_fit = X_all[fit_idx]
    X_tfidf_fit, idf = _tfidf_fit_transform(X_fit)

    n_components = int(n_lsi) + (1 if drop_first else 0)
    svd = TruncatedSVD(n_components=n_components, random_state=int(seed))
    svd.fit(X_tfidf_fit)

    Z_all = svd.transform(_tfidf_transform(X_all, idf))

    if drop_first:
        Z_all = Z_all[:, 1:]  # drop LSI1

    if l2norm:
        Z_all = normalize(Z_all, norm="l2", axis=1)

    atac.obsm["X_lsi"] = Z_all.astype(np.float32, copy=False)
    return idf, svd


# %% [code]
import scglue, ignite, torch
print("scglue:", getattr(scglue, "__version__", "unknown"))
print("ignite:", getattr(ignite, "__version__", "unknown"))
print("torch :", torch.__version__)
print("python:", __import__("sys").version)

# %% [code]
# ============================
# SCGLUE SIMPLE (robust + debug)
# - fixes indentation + undefined vars
# - passes init_kws/compile_kws/fit_kws/balance_kws safely
# - directory is always a real path
# ============================

from __future__ import annotations

from pathlib import Path
import time
import numpy as np
import scipy.sparse as sp


def _annotate_peaks_from_varnames(atac):
    """
    Robust peak parser:
      - supports 'chr1:123-456'
      - supports 'chr1_123_456'
    Drops peaks that cannot be parsed (prevents IntCastingNaNError).
    """
    import pandas as pd

    peaks = pd.Index(atac.var_names.astype(str))

    # pattern A: chr1:123-456
    m = peaks.to_series().str.extract(
        r"^(?P<chrom>[^:]+):(?P<start>\d+)-(?P<end>\d+)$"
    )

    # pattern B: chr1_123_456
    bad = m["start"].isna()
    if bad.any():
        m2 = peaks[bad].to_series().str.extract(
            r"^(?P<chrom>[^_]+)_(?P<start>\d+)_(?P<end>\d+)$"
        )
        m.loc[bad, ["chrom", "start", "end"]] = m2[["chrom", "start", "end"]].values

    ok = m["start"].notna() & m["end"].notna() & m["chrom"].notna()
    n_bad = int((~ok).sum())
    if n_bad:
        print(f"[scglue] peak parsing: dropping {n_bad}/{len(peaks)} peaks with unparseable var_names")
        atac = atac[:, ok.values].copy()
        m = m.loc[ok].copy()

    atac.var["chrom"] = m["chrom"].astype(str).values
    atac.var["chromStart"] = m["start"].astype(np.int64).values
    atac.var["chromEnd"] = m["end"].astype(np.int64).values
    return atac


def _fit_scglue_robust(scglue, adatas, graph, *, model_class=None,
                      skip_balance=False, init_kws=None, compile_kws=None,
                      fit_kws=None, balance_kws=None):
    """
    Call scglue.models.fit_SCGLUE across signature drift safely.
    Your install: (adatas, graph, model=..., skip_balance=..., init_kws=..., compile_kws=..., fit_kws=..., balance_kws=...)
    """
    import inspect

    init_kws = {} if init_kws is None else init_kws
    compile_kws = {} if compile_kws is None else compile_kws
    fit_kws = {} if fit_kws is None else fit_kws
    balance_kws = {} if balance_kws is None else balance_kws

    if not isinstance(init_kws, dict):
        raise TypeError(f"init_kws must be dict, got {type(init_kws)}: {init_kws!r}")
    if not isinstance(compile_kws, dict):
        raise TypeError(f"compile_kws must be dict, got {type(compile_kws)}: {compile_kws!r}")
    if not isinstance(fit_kws, dict):
        raise TypeError(f"fit_kws must be dict, got {type(fit_kws)}: {fit_kws!r}")
    if not isinstance(balance_kws, dict):
        raise TypeError(f"balance_kws must be dict, got {type(balance_kws)}: {balance_kws!r}")

    sig = inspect.signature(scglue.models.fit_SCGLUE)
    params = sig.parameters

    kwargs = {}
    # model / skip_balance are top-level in your signature
    if "model" in params and model_class is not None:
        kwargs["model"] = model_class
    if "skip_balance" in params:
        kwargs["skip_balance"] = bool(skip_balance)

    # nested kw dicts in your signature
    if "init_kws" in params:
        kwargs["init_kws"] = init_kws
    if "compile_kws" in params:
        kwargs["compile_kws"] = compile_kws
    if "fit_kws" in params:
        kwargs["fit_kws"] = fit_kws
    if "balance_kws" in params:
        kwargs["balance_kws"] = balance_kws

    # fallback for older forks that *don't* nest
    if "init_kws" not in params:
        kwargs.update(init_kws)
    if "compile_kws" not in params:
        kwargs.update(compile_kws)
    if "fit_kws" not in params:
        kwargs.update(fit_kws)
    if "balance_kws" not in params:
        kwargs.update(balance_kws)

    return scglue.models.fit_SCGLUE(adatas, graph, **kwargs)


def run_scglue_simple(
    rna_raw,
    atac_raw,
    *,
    gtf_path,
    splits,
    out_dir,
    seed=0,
    latent_dim=30,
    max_epochs=200,
    val_split=0.1,
    n_hvg=2000,
    n_pca=100,
    n_lsi=101,
    fuse="mean",                 # "mean" | "rna" | "atac" | "concat" | None
    model_class=None,            # e.g. scglue.models.PairedSCGLUEModel (optional)
    skip_balance=False,          # set True to skip balancing weights
    patience=15,                 # set 0 to avoid restore logic entirely
    reduce_lr_patience=8,
    debug_print_kws=True,
):
    import random
    import scanpy as sc
    import scglue
    import torch

    # -------------------------
    # seeds
    # -------------------------
    random.seed(seed)
    np.random.seed(seed)
    torch.manual_seed(seed)
    if torch.cuda.is_available():
        torch.cuda.manual_seed_all(seed)

    # -------------------------
    # FIT cells (train+val only; test held out)
    # -------------------------
    tr = np.asarray(splits["train"], dtype=int)
    va = np.asarray(splits.get("val", []), dtype=int)
    fit_idx = np.concatenate([tr, va]) if va.size else tr

    # -------------------------
    # Restrict features to those observed in FIT cells
    # -------------------------
    def _nnz_mask(X):
        if sp.issparse(X):
            return np.asarray((X > 0).sum(axis=0)).ravel() > 0
        X = np.asarray(X)
        return (X > 0).sum(axis=0) > 0

    keep_genes = _nnz_mask(rna_raw[fit_idx].X)
    keep_peaks = _nnz_mask(atac_raw[fit_idx].X)

    rna = rna_raw[:, keep_genes].copy()
    atac = atac_raw[:, keep_peaks].copy()

    # -------------------------
    # Genomic coords
    # -------------------------
    gtf_path = str(Path(gtf_path))

    v = rna.var_names.astype(str)
    frac_ens = np.mean([s.startswith("ENS") for s in v[: min(200, len(v))]])
    gtf_by = "gene_id" if frac_ens > 0.5 else "gene_name"

    scglue.data.get_gene_annotation(rna, gtf=gtf_path, gtf_by=gtf_by)

    gene_ok = rna.var[["chrom", "chromStart", "chromEnd"]].notna().all(axis=1).to_numpy()
    if gene_ok.sum() < rna.n_vars:
        print(f"[scglue] gene annotation: dropping {rna.n_vars - gene_ok.sum()}/{rna.n_vars} genes without coords")
        rna = rna[:, gene_ok].copy()

    atac = _annotate_peaks_from_varnames(atac)

    # -------------------------
    # RNA preprocessing (HVGs fit on FIT only)
    # -------------------------
    rna.layers["counts"] = rna.X.copy()

    rna_fit2 = rna[fit_idx].copy()
    sc.pp.highly_variable_genes(
        rna_fit2,
        n_top_genes=int(min(n_hvg, rna_fit2.n_vars)),
        flavor="seurat_v3",
    )
    hv = rna_fit2.var["highly_variable"].reindex(rna.var_names).fillna(False).to_numpy(dtype=bool)
    rna.var["highly_variable"] = hv

    sc.pp.normalize_total(rna, target_sum=1e4)
    sc.pp.log1p(rna)
    sc.pp.scale(rna, max_value=10, zero_center=False)

    try:
        sc.tl.pca(rna, n_comps=int(n_pca), svd_solver="auto", mask_var="highly_variable")
    except TypeError:
        sc.tl.pca(rna, n_comps=int(n_pca), svd_solver="auto", use_highly_variable=True)

    # -------------------------
    # ATAC preprocessing
    # -------------------------
    scglue.data.lsi(atac, n_components=int(n_lsi), n_iter=15)

    # -------------------------
    # Guidance graph + check
    # -------------------------
    guidance = scglue.genomics.rna_anchored_guidance_graph(rna, atac)
    scglue.graph.check_graph(guidance, [rna, atac])

    # -------------------------
    # Configure dataset for GLUE
    # -------------------------
    scglue.models.configure_dataset(
        rna, "NB",
        use_highly_variable=True,
        use_layer="counts",
        use_rep="X_pca",
        use_obs_names=True,
    )
    scglue.models.configure_dataset(
        atac, "NB",
        use_highly_variable=True,
        use_rep="X_lsi",
        use_obs_names=True,
    )

    # -------------------------
    # run directory (must exist)
    # -------------------------
    out_dir = Path(out_dir)
    out_dir.mkdir(parents=True, exist_ok=True)
    run_dir = out_dir / "scglue_train" / time.strftime("%Y%m%d_%H%M%S")
    run_dir.mkdir(parents=True, exist_ok=False)

    fit_kws = dict(
        max_epochs=int(max_epochs),
        val_split=float(val_split),
        directory=str(run_dir),
        patience=int(patience),
        reduce_lr_patience=int(reduce_lr_patience),
    )
    init_kws = dict(latent_dim=int(latent_dim), random_seed=int(seed))
    compile_kws = {}   # keep explicit; some forks accept optimizer/lr here
    balance_kws = {}   # keep explicit; some forks accept params here

    def _ck(name, obj):
        if isinstance(obj, str):
            s = obj if len(obj) <= 120 else (obj[:120] + "…")
            print(f"[scglue][debug] {name}: {type(obj)} {s!r}")
        else:
            print(f"[scglue][debug] {name}: {type(obj)} {obj}")

    if debug_print_kws:
        _ck("init_kws", init_kws)
        _ck("compile_kws", compile_kws)
        _ck("fit_kws", fit_kws)
        _ck("balance_kws", balance_kws)
        _ck("model_class", model_class)

    # -------------------------
    # Fit (only on FIT cells)
    # -------------------------
    glue = _fit_scglue_robust(
        scglue,
        {"rna": rna[fit_idx].copy(), "atac": atac[fit_idx].copy()},
        guidance,
        model_class=model_class,
        skip_balance=skip_balance,
        init_kws=init_kws,
        compile_kws=compile_kws,
        fit_kws=fit_kws,
        balance_kws=balance_kws,
    )

    # -------------------------
    # Encode ALL cells (evaluation will slice test)
    # -------------------------
    Z_rna = np.asarray(glue.encode_data("rna", rna), dtype=np.float32)
    Z_atac = np.asarray(glue.encode_data("atac", atac), dtype=np.float32)

    if fuse == "mean":
        Z_fused = 0.5 * (Z_rna + Z_atac)
    elif fuse == "rna":
        Z_fused = Z_rna
    elif fuse == "atac":
        Z_fused = Z_atac
    elif fuse == "concat":
        Z_fused = np.concatenate([Z_rna, Z_atac], axis=1)
    elif fuse is None:
        Z_fused = None
    else:
        raise ValueError(f"Unknown fuse={fuse!r}")

    return dict(
        Z_fused=Z_fused,
        Z_rna=Z_rna,
        Z_atac=Z_atac,
        fit_idx=fit_idx,
        gtf_by=gtf_by,
        n_genes=rna.n_vars,
        n_peaks=atac.n_vars,
        run_dir=str(run_dir),
        fit_kws=fit_kws,
        init_kws=init_kws,
        extra_json={"transductive": False, "uses_labels": False},
    )


# %% [code]
from __future__ import annotations

from pathlib import Path
import time
import numpy as np
import scipy.sparse as sp


def _annotate_peaks_from_varnames(atac):
    import pandas as pd

    peaks = pd.Index(atac.var_names.astype(str))

    m = peaks.to_series().str.extract(r"^(?P<chrom>[^:]+):(?P<start>\d+)-(?P<end>\d+)$")
    bad = m["start"].isna()
    if bad.any():
        m2 = peaks[bad].to_series().str.extract(r"^(?P<chrom>[^_]+)_(?P<start>\d+)_(?P<end>\d+)$")
        m.loc[bad, ["chrom", "start", "end"]] = m2[["chrom", "start", "end"]].values

    ok = m["start"].notna() & m["end"].notna() & m["chrom"].notna()
    n_bad = int((~ok).sum())
    if n_bad:
        print(f"[scglue] peak parsing: dropping {n_bad}/{len(peaks)} peaks with unparseable var_names")
        atac = atac[:, ok.values].copy()
        m = m.loc[ok].copy()

    atac.var["chrom"] = m["chrom"].astype(str).values
    atac.var["chromStart"] = m["start"].astype(np.int64).values
    atac.var["chromEnd"] = m["end"].astype(np.int64).values
    return atac


def _nnz_mask_cellslice(X):
    """Return boolean mask over features: feature has at least 1 nonzero in this cell slice."""
    if sp.issparse(X):
        return np.asarray((X > 0).sum(axis=0)).ravel() > 0
    X = np.asarray(X)
    return (X > 0).sum(axis=0) > 0


def _topk_by_counts(X, k: int):
    """Pick top-k features by total counts (sparse-safe)."""
    if sp.issparse(X):
        s = np.asarray(X.sum(axis=0)).ravel()
    else:
        s = np.asarray(X).sum(axis=0)
    k = int(min(k, s.size))
    if k <= 0:
        return np.zeros(s.size, dtype=bool)
    idx = np.argpartition(-s, kth=k - 1)[:k]
    m = np.zeros(s.size, dtype=bool)
    m[idx] = True
    return m


def _fit_scglue_robust(scglue, adatas, graph, *, model_class=None,
                      skip_balance=False, init_kws=None, compile_kws=None,
                      fit_kws=None, balance_kws=None):
    import inspect

    init_kws = {} if init_kws is None else init_kws
    compile_kws = {} if compile_kws is None else compile_kws
    fit_kws = {} if fit_kws is None else fit_kws
    balance_kws = {} if balance_kws is None else balance_kws

    for nm, obj in [("init_kws", init_kws), ("compile_kws", compile_kws), ("fit_kws", fit_kws), ("balance_kws", balance_kws)]:
        if not isinstance(obj, dict):
            raise TypeError(f"{nm} must be dict, got {type(obj)}: {obj!r}")

    sig = inspect.signature(scglue.models.fit_SCGLUE)
    params = sig.parameters

    kwargs = {}
    if "model" in params and model_class is not None:
        kwargs["model"] = model_class
    if "skip_balance" in params:
        kwargs["skip_balance"] = bool(skip_balance)

    if "init_kws" in params:
        kwargs["init_kws"] = init_kws
    else:
        kwargs.update(init_kws)

    if "compile_kws" in params:
        kwargs["compile_kws"] = compile_kws
    else:
        kwargs.update(compile_kws)

    if "fit_kws" in params:
        kwargs["fit_kws"] = fit_kws
    else:
        kwargs.update(fit_kws)

    if "balance_kws" in params:
        kwargs["balance_kws"] = balance_kws
    else:
        kwargs.update(balance_kws)

    return scglue.models.fit_SCGLUE(adatas, graph, **kwargs)


def run_scglue_simple_v2(
    rna_raw,
    atac_raw,
    *,
    gtf_path,
    splits,
    out_dir,
    seed=0,
    latent_dim=30,
    max_epochs=200,

    # HV / PCA / LSI
    n_hvg=2000,
    n_pca=100,
    n_lsi=101,
    n_hvpeaks=50000,          # IMPORTANT: define HV peaks for ATAC if using use_highly_variable=True

    # training/early stopping behavior
    val_split=0.1,            # internal val fraction (must be >0 for many scglue versions)
    patience=30,
    reduce_lr_patience=15,

    # fusion
    fuse="mean",              # "mean" | "rna" | "atac" | "concat" | None

    # model options
    model_class=None,
    skip_balance=False,

    # debug
    debug=True,
):
    import random
    import scanpy as sc
    import scglue
    import torch

    # -------------------------
    # seeds
    # -------------------------
    random.seed(int(seed))
    np.random.seed(int(seed))
    torch.manual_seed(int(seed))
    if torch.cuda.is_available():
        torch.cuda.manual_seed_all(int(seed))

    # -------------------------
    # FIT cells: train (+ optional val) but test held out
    # -------------------------
    tr = np.asarray(splits["train"], dtype=int)
    va = np.asarray(splits.get("val", []), dtype=int)
    fit_idx = np.concatenate([tr, va]) if va.size else tr

    if fit_idx.size < 100:
        raise ValueError(f"[scglue] Too few fit cells ({fit_idx.size}). Something is off in splits.")

    # -------------------------
    # sanity: ensure X is counts-like
    # -------------------------
    def _assert_counts_like(adata, name):
        X = adata.X
        if not sp.issparse(X):
            X = np.asarray(X)
        # allow float, but must be nonnegative
        mn = X.min() if not sp.issparse(X) else X.min()
        if mn < 0:
            raise ValueError(f"[scglue] {name}.X has negative values (min={mn}). "
                             f"Feed raw counts in .X; put TFIDF/LSI in obsm, not .X.")
    _assert_counts_like(rna_raw, "rna_raw")
    _assert_counts_like(atac_raw, "atac_raw")

    # -------------------------
    # Restrict to features observed in FIT cells
    # -------------------------
    keep_genes = _nnz_mask_cellslice(rna_raw[fit_idx].X)
    keep_peaks = _nnz_mask_cellslice(atac_raw[fit_idx].X)

    rna = rna_raw[:, keep_genes].copy()
    atac = atac_raw[:, keep_peaks].copy()

    if rna.n_vars < 500:
        raise ValueError(f"[scglue] After FIT filtering, only {rna.n_vars} genes remain.")
    if atac.n_vars < 1000:
        raise ValueError(f"[scglue] After FIT filtering, only {atac.n_vars} peaks remain.")

    # -------------------------
    # Gene + peak genomic coords
    # -------------------------
    gtf_path = str(Path(gtf_path))

    v = rna.var_names.astype(str)
    frac_ens = np.mean([s.startswith("ENS") for s in v[: min(200, len(v))]])
    gtf_by = "gene_id" if frac_ens > 0.5 else "gene_name"

    scglue.data.get_gene_annotation(rna, gtf=gtf_path, gtf_by=gtf_by)

    gene_ok = rna.var[["chrom", "chromStart", "chromEnd"]].notna().all(axis=1).to_numpy()
    if gene_ok.sum() < rna.n_vars:
        drop = int(rna.n_vars - gene_ok.sum())
        print(f"[scglue] gene annotation: dropping {drop}/{rna.n_vars} genes without coords")
        rna = rna[:, gene_ok].copy()

    atac = _annotate_peaks_from_varnames(atac)

    # -------------------------
    # RNA preprocessing (HVGs fit on FIT only)
    # -------------------------
    rna.layers["counts"] = rna.X.copy()

    rna_fit = rna[fit_idx].copy()
    sc.pp.highly_variable_genes(
        rna_fit,
        n_top_genes=int(min(n_hvg, rna_fit.n_vars)),
        flavor="seurat_v3",
    )
    hv = rna_fit.var["highly_variable"].reindex(rna.var_names).fillna(False).to_numpy(dtype=bool)
    rna.var["highly_variable"] = hv

    sc.pp.normalize_total(rna, target_sum=1e4)
    sc.pp.log1p(rna)
    sc.pp.scale(rna, max_value=10, zero_center=False)

    try:
        sc.tl.pca(rna, n_comps=int(n_pca), svd_solver="auto", mask_var="highly_variable")
    except TypeError:
        sc.tl.pca(rna, n_comps=int(n_pca), svd_solver="auto", use_highly_variable=True)

    # -------------------------
    # ATAC HV peaks + LSI
    # -------------------------
    atac.layers["counts"] = atac.X.copy()

    # define HV peaks ON FIT cells (counts-based; robust + fast)
    hvp = _topk_by_counts(atac[fit_idx].X, k=int(min(n_hvpeaks, atac.n_vars)))
    atac.var["highly_variable"] = hvp

    # LSI uses counts in .X (or counts layer depending on scglue version)
    scglue.data.lsi(atac, n_components=int(n_lsi), n_iter=15)

    # -------------------------
    # Guidance graph + check
    # -------------------------
    guidance = scglue.genomics.rna_anchored_guidance_graph(rna, atac)
    scglue.graph.check_graph(guidance, [rna, atac])

    # -------------------------
    # Configure datasets
    # -------------------------
    scglue.models.configure_dataset(
        rna, "NB",
        use_highly_variable=True,
        use_layer="counts",
        use_rep="X_pca",
        use_obs_names=True,
    )
    scglue.models.configure_dataset(
        atac, "NB",
        use_highly_variable=True,      # NOW SAFE because we created atac.var["highly_variable"]
        use_layer="counts",
        use_rep="X_lsi",
        use_obs_names=True,
    )

    # -------------------------
    # run directory
    # -------------------------
    out_dir = Path(out_dir)
    out_dir.mkdir(parents=True, exist_ok=True)
    run_dir = out_dir / "scglue_train" / time.strftime("%Y%m%d_%H%M%S")
    run_dir.mkdir(parents=True, exist_ok=False)

    # enforce strictly-positive val_split (some scglue versions require this)
    val_split_eff = float(val_split)
    if not (0.0 < val_split_eff < 1.0):
        # choose something sane + safe
        val_split_eff = min(0.1, max(0.01, 1.0 / float(fit_idx.size)))
        if debug:
            print(f"[scglue] val_split was {val_split!r}; using safe val_split_eff={val_split_eff:.4g}")

    fit_kws = dict(
        max_epochs=int(max_epochs),
        val_split=float(val_split_eff),
        directory=str(run_dir),
        patience=int(patience),
        reduce_lr_patience=int(reduce_lr_patience),
    )
    init_kws = dict(latent_dim=int(latent_dim), random_seed=int(seed))
    compile_kws = {}
    balance_kws = {}

    if debug:
        print(f"[scglue][dbg] fit cells={fit_idx.size} genes={rna.n_vars} peaks={atac.n_vars}")
        print(f"[scglue][dbg] RNA HVGs={int(rna.var['highly_variable'].sum())} "
              f"ATAC HVpeaks={int(atac.var['highly_variable'].sum())}")
        print(f"[scglue][dbg] run_dir={run_dir}")
        print(f"[scglue][dbg] init_kws={init_kws}")
        print(f"[scglue][dbg] fit_kws={fit_kws}")

    # -------------------------
    # Fit (on FIT cells)
    # -------------------------
    glue = _fit_scglue_robust(
        scglue,
        {"rna": rna[fit_idx].copy(), "atac": atac[fit_idx].copy()},
        guidance,
        model_class=model_class,
        skip_balance=skip_balance,
        init_kws=init_kws,
        compile_kws=compile_kws,
        fit_kws=fit_kws,
        balance_kws=balance_kws,
    )

    # -------------------------
    # Encode ALL cells
    # -------------------------
    Z_rna = np.asarray(glue.encode_data("rna", rna), dtype=np.float32)
    Z_atac = np.asarray(glue.encode_data("atac", atac), dtype=np.float32)

    if fuse == "mean":
        Z_fused = 0.5 * (Z_rna + Z_atac)
    elif fuse == "rna":
        Z_fused = Z_rna
    elif fuse == "atac":
        Z_fused = Z_atac
    elif fuse == "concat":
        Z_fused = np.concatenate([Z_rna, Z_atac], axis=1)
    elif fuse is None:
        Z_fused = None
    else:
        raise ValueError(f"Unknown fuse={fuse!r}")

    return dict(
        Z_fused=Z_fused,
        Z_rna=Z_rna,
        Z_atac=Z_atac,
        fit_idx=fit_idx,
        gtf_by=gtf_by,
        n_genes=int(rna.n_vars),
        n_peaks=int(atac.n_vars),
        run_dir=str(run_dir),
        fit_kws=fit_kws,
        init_kws=init_kws,
        extra_json={"transductive": False, "uses_labels": False},
    )


# %% [code]
# =========================
# NEW: scGLUE silencing utils
# =========================
import os
import re
import logging
import warnings
from contextlib import contextmanager

@contextmanager
def silence_scglue(
    *,
    verbose: bool = False,
    keep_user_prints: bool = True,
):
    """
    Quiet scGLUE/scanpy/lightning + common torch pin_memory deprecation spam.
    - verbose=True => do nothing (full logs)
    - keep_user_prints kept for clarity (we don't redirect stdout here)
    """
    if verbose:
        yield
        return

    # Progress bars: many libs respect these
    os.environ.setdefault("TQDM_DISABLE", "1")
    os.environ.setdefault("TF_CPP_MIN_LOG_LEVEL", "3")

    # Clamp noisy loggers
    noisy = [
        "scglue",
        "scanpy",
        "anndata",
        "muon",
        "pytorch_lightning",
        "lightning",
        "torch",
    ]
    old_levels = {}
    for name in noisy:
        lg = logging.getLogger(name)
        old_levels[name] = lg.level
        lg.setLevel(logging.ERROR)

    # Warnings: keep it surgical
    with warnings.catch_warnings():
        warnings.filterwarnings("ignore", category=FutureWarning)
        warnings.filterwarnings("ignore", category=UserWarning)
        warnings.filterwarnings("ignore", category=ResourceWarning)

        # This is the ultra-spammy one in your log:
        warnings.filterwarnings(
            "ignore",
            category=DeprecationWarning,
            module=r"torch\.utils\.data\._utils\.pin_memory",
        )

        # Often seen with lightning/torch compile churn
        warnings.filterwarnings("ignore", category=DeprecationWarning)

        # Scanpy verbosity if available
        try:
            import scanpy as sc
            sc.settings.verbosity = 0
        except Exception:
            pass

        try:
            yield
        finally:
            for name, lvl in old_levels.items():
                logging.getLogger(name).setLevel(lvl)


def _filter_kwargs_to_callable(fn, kwargs: dict) -> dict:
    """Return subset of kwargs accepted by fn's signature (API-drift tolerant)."""
    import inspect
    try:
        sig = inspect.signature(fn)
    except Exception:
        return dict(kwargs)
    return {k: v for k, v in kwargs.items() if k in sig.parameters}


def _set_scglue_trainer_quiet(fit_kws: dict, *, verbose: bool) -> dict:
    """
    scGLUE fit_SCGLUE doesn't necessarily accept Lightning Trainer args.
    So we DO NOT add enable_progress_bar/log_every_n_steps here.
    Quieting is handled via `silence_scglue(...)` (warnings/loggers/tqdm env).
    """
    # If you want to be extra safe, strip keys that might sneak in from elsewhere:
    banned = {"enable_progress_bar", "log_every_n_steps", "enable_model_summary", "trainer_kwargs", "callbacks"}
    out = {k: v for k, v in fit_kws.items() if k not in banned}
    return out


def _ensure_counts_layers_for_scglue(rna, atac, *, rna_layer="counts", atac_layer="counts"):
    """
    Ensure layers exist for scglue.configure_dataset(use_layer=...).
    Tries to avoid guessing counts, but will fall back to copying .X.
    """
    if rna_layer not in rna.layers:
        if getattr(rna, "raw", None) is not None and getattr(rna.raw, "X", None) is not None:
            rna.layers[rna_layer] = rna.raw.X.copy()
        else:
            rna.layers[rna_layer] = rna.X.copy()

    if atac_layer not in atac.layers:
        if getattr(atac, "raw", None) is not None and getattr(atac.raw, "X", None) is not None:
            atac.layers[atac_layer] = atac.raw.X.copy()
        else:
            atac.layers[atac_layer] = atac.X.copy()

    return rna, atac


# %% [code]
# =========================
# REWRITE: scGLUE runner (quiet-by-default)
# =========================
from pathlib import Path
import time
import random
import numpy as np
import scipy.sparse as sp

def run_scglue_fair(
    rna_raw,
    atac_raw,
    *,
    gtf_path,
    splits,
    out_dir,
    seed=0,
    latent_dim=30,
    max_epochs=200,
    # feature dims
    n_hvg=2000,
    n_pca=100,
    n_lsi=101,
    n_hvpeaks=50000,
    # scglue training behavior
    val_split=0.1,
    patience=30,
    reduce_lr_patience=15,
    fuse="mean",     # mean|rna|atac|concat|None
    model_class=None,
    skip_balance=False,
    # layers (explicit)
    rna_counts_layer="counts",
    atac_counts_layer="counts",
    # verbosity
    verbose=False,
    debug=False,
):
    """
    Leakage-aware scGLUE runner (quiet by default):
      - Restricts features (genes/peaks) using FIT cells only (train+val)
      - Fits HVG+PCA on FIT only -> applies to ALL
      - Fits HV-peaks+LSI on FIT only -> applies to ALL
      - Trains GLUE on FIT only -> encodes ALL (test inference-only)

    Requires your existing helpers:
      - _annotate_peaks_from_varnames(atac)
      - compute_hvg_on_fit_cells(...)
      - fit_pca_train_apply_all(...)
      - compute_hv_peaks_on_fit_cells(...)
      - fit_lsi_train_apply_all(...)
      - _fit_scglue_robust(...)
      - ensure_flags(...)
      - standard_flags(...)
    """
    import scanpy as sc
    import scglue
    import torch

    # -------------------------
    # seeds
    # -------------------------
    random.seed(int(seed))
    np.random.seed(int(seed))
    torch.manual_seed(int(seed))
    if torch.cuda.is_available():
        torch.cuda.manual_seed_all(int(seed))

    out_dir = Path(out_dir)
    out_dir.mkdir(parents=True, exist_ok=True)

    tr = np.asarray(splits["train"], dtype=int)
    va = np.asarray(splits.get("val", []), dtype=int)
    fit_idx = np.concatenate([tr, va]) if va.size else tr
    fit_idx = np.asarray(fit_idx, dtype=int)

    # keep val strictly inside FIT; no test leakage
    val_split_eff = float(val_split)
    if not (0.0 < val_split_eff < 1.0):
        val_split_eff = min(0.1, max(0.01, 1.0 / float(max(1, fit_idx.size))))

    # -------------------------
    # small utilities
    # -------------------------
    def _as_csr(X):
        if sp.issparse(X):
            return X.tocsr()
        return sp.csr_matrix(np.asarray(X))

    def _nnz_mask(X):
        X = _as_csr(X)
        return np.asarray((X > 0).sum(axis=0)).ravel() > 0

    t0 = time.perf_counter()

    with silence_scglue(verbose=bool(verbose)):
        # -------------------------
        # feature restriction based on FIT cells only
        # -------------------------
        keep_genes = _nnz_mask(rna_raw[fit_idx].X)
        keep_peaks = _nnz_mask(atac_raw[fit_idx].X)

        rna = rna_raw[:, keep_genes].copy()
        atac = atac_raw[:, keep_peaks].copy()

        # -------------------------
        # annotate genes + peaks
        # -------------------------
        gtf_path = str(Path(gtf_path))
        frac_ens = np.mean([str(s).startswith("ENS") for s in rna.var_names[: min(200, rna.n_vars)]])
        gtf_by = "gene_id" if frac_ens > 0.5 else "gene_name"

        scglue.data.get_gene_annotation(rna, gtf=gtf_path, gtf_by=gtf_by)
        gene_ok = rna.var[["chrom", "chromStart", "chromEnd"]].notna().all(axis=1).to_numpy()
        if gene_ok.sum() < rna.n_vars:
            if debug:
                print(f"[scglue] dropping {rna.n_vars - gene_ok.sum()}/{rna.n_vars} genes without coords")
            rna = rna[:, gene_ok].copy()

        atac = _annotate_peaks_from_varnames(atac)  # your robust parser

        # -------------------------
        # ensure counts layers
        # -------------------------
        rna, atac = _ensure_counts_layers_for_scglue(
            rna, atac,
            rna_layer=rna_counts_layer,
            atac_layer=atac_counts_layer,
        )

        # -------------------------
        # HVGs / PCA fit on FIT only -> apply to ALL
        # -------------------------
        compute_hvg_on_fit_cells(
            rna, fit_idx,
            n_top_genes=int(n_hvg),
            flavor="seurat_v3",
            batch_key=None,
        )
        fit_pca_train_apply_all(
            rna, fit_idx,
            n_pca=int(n_pca),
            target_sum=1e4,
            seed=int(seed),
            layer=rna_counts_layer,
        )

        # -------------------------
        # HV-peaks / LSI fit on FIT only -> apply to ALL
        # -------------------------
        compute_hv_peaks_on_fit_cells(atac, fit_idx, n_top_peaks=int(n_hvpeaks))
        fit_lsi_train_apply_all(atac, fit_idx, n_lsi=int(n_lsi), seed=int(seed), l2norm=True)

        # -------------------------
        # build guidance graph + check
        # -------------------------
        guidance = scglue.genomics.rna_anchored_guidance_graph(rna, atac)
        scglue.graph.check_graph(guidance, [rna, atac])

        # -------------------------
        # configure datasets (use reps we computed leakage-safe)
        # -------------------------
        scglue.models.configure_dataset(
            rna, "NB",
            use_highly_variable=True,
            use_layer=rna_counts_layer,
            use_rep="X_pca",
            use_obs_names=True,
        )
        scglue.models.configure_dataset(
            atac, "NB",
            use_highly_variable=True,
            use_layer=atac_counts_layer,
            use_rep="X_lsi",
            use_obs_names=True,
        )

        # -------------------------
        # train on FIT only
        # -------------------------
        run_dir = out_dir / "scglue_train" / time.strftime("%Y%m%d_%H%M%S")
        run_dir.mkdir(parents=True, exist_ok=False)

        init_kws = dict(latent_dim=int(latent_dim), random_seed=int(seed))
        fit_kws = dict(
            max_epochs=int(max_epochs),
            val_split=float(val_split_eff),
            directory=str(run_dir),
            patience=int(patience),
            reduce_lr_patience=int(reduce_lr_patience),
        )
        fit_kws = _set_scglue_trainer_quiet(fit_kws, verbose=bool(verbose))

        glue = _fit_scglue_robust(
            scglue,
            {"rna": rna[fit_idx].copy(), "atac": atac[fit_idx].copy()},
            guidance,
            model_class=model_class,
            skip_balance=skip_balance,
            init_kws=init_kws,
            compile_kws={},
            fit_kws=fit_kws,
            balance_kws={},
        )

        # -------------------------
        # encode ALL cells (test is inference-only)
        # -------------------------
        Z_rna = np.asarray(glue.encode_data("rna", rna), dtype=np.float32)
        Z_atac = np.asarray(glue.encode_data("atac", atac), dtype=np.float32)

        if fuse == "mean":
            Z_fused = 0.5 * (Z_rna + Z_atac)
        elif fuse == "rna":
            Z_fused = Z_rna
        elif fuse == "atac":
            Z_fused = Z_atac
        elif fuse == "concat":
            Z_fused = np.concatenate([Z_rna, Z_atac], axis=1)
        elif fuse is None:
            Z_fused = None
        else:
            raise ValueError(f"Unknown fuse={fuse!r}")

    fit_seconds = float(time.perf_counter() - t0)

    if debug and not verbose:
        # minimal one-liner summary
        print(f"[scglue] done. fit_seconds={fit_seconds:.2f}  Z_rna={Z_rna.shape}  Z_atac={Z_atac.shape}")

    return ensure_flags({
        "Z_fused": Z_fused,
        "Z_rna": Z_rna,
        "Z_atac": Z_atac,
        "fit_seconds": fit_seconds,
        "extra_json": standard_flags(
            transductive=False,
            uses_labels=False,
            note="HVG/HVpeaks + PCA/LSI fit on train+val only; GLUE trained on train+val only; test is inference-only.",
        ),
    }, default_transductive=False, default_uses_labels=False)


# %% [code]
#import inspect, scglue, scglue.models
#src = inspect.getsource(scglue.models.fit_SCGLUE)
#print(src)


# %% [markdown]
### 4) MultiMAP (robust wrapper + API autodetect)

# %% [code]
def run_multimap(rna_log_hvg, atac_lsi, *, out_dir, splits, seed=0, latent_dim=30):
    import numpy as np
    import scipy.sparse as sp
    from sklearn.decomposition import PCA
    from sklearn.preprocessing import StandardScaler

    if not hasattr(np, "infty"):
        np.infty = np.inf  # NumPy 2 compat shim

    from MultiMAP.matrix import MultiMAP as multimap_matrix

    # ------------------------------------------------------------
    # 0) Align modalities to the SAME cells in the SAME order
    # ------------------------------------------------------------
    rna0 = rna_log_hvg
    atac0 = atac_lsi

    shared = rna0.obs_names.intersection(atac0.obs_names)
    if shared.size == 0:
        raise ValueError("MultiMAP: RNA/ATAC have no shared cells (obs_names intersection empty).")

    # canonical order = RNA order
    rna = rna0[shared].copy()
    atac = atac0[shared].copy()
    if not rna.obs_names.equals(atac.obs_names):
        raise ValueError("MultiMAP: failed to align RNA/ATAC obs_names after subsetting.")

    n_obs = int(rna.n_obs)

    # ------------------------------------------------------------
    # 1) Remap splits (original RNA index space -> aligned shared space)
    # ------------------------------------------------------------
    tr0 = np.asarray(splits["train"], dtype=int)

    # map original RNA row index -> aligned row index (or -1 if dropped)
    orig_pos = rna0.obs_names.get_indexer(shared)  # positions of shared cells in original RNA
    inv = np.full(int(rna0.n_obs), -1, dtype=int)
    inv[orig_pos] = np.arange(shared.size, dtype=int)

    tr = inv[tr0]
    tr = tr[tr >= 0]
    if tr.size == 0:
        raise ValueError(
            "MultiMAP: no TRAIN cells remain after aligning modalities. "
            "Train split likely contains cells missing from one modality."
        )

    # ------------------------------------------------------------
    # 2) Build matrices (dense), PCA-on-train for RNA, then scale
    # ------------------------------------------------------------
    Xr = rna.X
    if sp.issparse(Xr):
        Xr = Xr.toarray()
    Xr = np.asarray(Xr, dtype=np.float32)

    Xa = atac.X
    if sp.issparse(Xa):
        Xa = Xa.toarray()
    X2 = np.asarray(Xa, dtype=np.float32)

    # PCA fit on aligned TRAIN only; transform all aligned cells
    n_pcs = int(X2.shape[1])
    pca = PCA(n_components=n_pcs, random_state=int(seed))
    pca.fit(Xr[tr])
    X1 = pca.transform(Xr).astype(np.float32, copy=False)

    # train-fit scaling per modality
    sc1 = StandardScaler(with_mean=True, with_std=True)
    sc2 = StandardScaler(with_mean=True, with_std=True)
    sc1.fit(X1[tr])
    sc2.fit(X2[tr])
    X1s = sc1.transform(X1).astype(np.float32, copy=False)
    X2s = sc2.transform(X2).astype(np.float32, copy=False)

    if not (np.isfinite(X1s).all() and np.isfinite(X2s).all()):
        raise ValueError("MultiMAP: non-finite values detected after scaling.")

    # ------------------------------------------------------------
    # 3) Run MultiMAP (effectively transductive)
    # ------------------------------------------------------------
    t0 = now()
    params, graph, Z = multimap_matrix(
        [X1s, X2s],
        n_components=int(latent_dim),
        random_state=int(seed),
    )
    t1 = now()

    Z = np.asarray(Z, dtype=np.float32)

    if Z.ndim != 2:
        raise ValueError(f"MultiMAP: unexpected Z ndim={Z.ndim} (shape={Z.shape})")
    if Z.shape[1] != int(latent_dim):
        raise ValueError(f"MultiMAP: Z dim ({Z.shape[1]}) != latent_dim ({int(latent_dim)})")

    # ------------------------------------------------------------
    # 4) Handle MultiMAP output shape
    #    - Some versions return (n_obs, d) fused
    #    - The matrix API often returns (sum_i n_i, d) concatenated
    #      which here is (2*n_obs, d)
    # ------------------------------------------------------------
    if Z.shape[0] == n_obs:
        Zr = Z
        Za = None
        Zf = Z
        mode = "fused"
    elif Z.shape[0] == 2 * n_obs:
        Zr = Z[:n_obs, :]
        Za = Z[n_obs:2 * n_obs, :]
        Zf = 0.5 * (Zr + Za)
        mode = "concat_by_modality"
    else:
        raise ValueError(
            f"MultiMAP: unexpected Z rows ({Z.shape[0]}). "
            f"Expected {n_obs} (fused) or {2*n_obs} (concat for 2 modalities)."
        )

    return ensure_flags({
        "Z_fused": Zf,
        "Z_rna": Zr,
        "Z_atac": Za,
        "fit_seconds": float(t1 - t0),
        "extra_json": standard_flags(
            transductive=True,
            uses_labels=False,
            n_shared=int(n_obs),
            n_train_shared=int(tr.size),
            output_mode=mode,
            note="Aligned RNA/ATAC by shared obs_names; PCA+scaling fit on aligned TRAIN subset; MultiMAP matrix output handled."
        ),
    })


# %% [markdown]
### 5) scMoMaT

# %% [code]
# ============================================================
# scMoMaT runner (fork-tolerant, auto-orientation, robust parsing)
# - Builds per-batch inputs with shared cell order across modalities
# - Auto-detects whether your fork expects (cells x features) or (features x cells)
# - Parses extract_cell_factors() across common fork return formats
# - Always returns: Z_fused, and Z_rna/Z_atac/Z_adt when available
# ============================================================

from __future__ import annotations

import time
import json
import inspect
from pathlib import Path
from typing import Dict, List, Tuple, Optional

import numpy as np
import scipy.sparse as sp
import torch

# --- SciPy >=1.11 compatibility: some scMoMaT forks use len(sparse) internally ---
try:
    def _len_shape0(self):
        return self.shape[0]
    sp.spmatrix.__len__ = _len_shape0  # type: ignore[attr-defined]
except Exception:
    pass


def run_scmomat_fair(
    *,
    rna=None,
    atac=None,
    adt=None,
    out_dir=None,
    splits=None,  # unused (transductive); kept for signature compatibility
    batch_key=None,                  # if None => single batch
    K=30,
    layers_by_mod=None,              # e.g. {"rna":"counts","atac":"counts"}
    seed=0,
    T=2000,
    lr=1e-4,
    lamb=1e-3,
    interval=200,
    batch_size=0.1,
    device="cuda",
    verbose=False,

    # Orientation handling:
    # - "auto": detect which axis is "cells" by matching across modalities, then adapt for constructor
    # - False: pass (cells x features) as-is
    # - True:  pass (features x cells) (i.e., transpose)
    transpose_to_features_by_cells: bool | str = "auto",

    ensure_nonnegative: bool = False,
    **_ignored,
):
    """
    Notes:
      - scMoMaT is transductive: fits on all cells it embeds.
      - For multi-batch, this runner intersects cells within each batch across modalities.
      - For fork quirks, we:
          * force dense float32 contiguous arrays
          * auto-orient inputs ("auto") so the model sees consistent cell axis
          * parse extract_cell_factors() in many formats (batch-major, modality-major, single-array)
    """
    # -------------------------
    # imports + helpers
    # -------------------------
    def _safe_import_scmomat_model():
        import scmomat
        if hasattr(scmomat, "scmomat_model"):
            return scmomat.scmomat_model
        from scmomat.model import scmomat_model
        return scmomat_model

    def _call_with_signature(fn, /, **kwargs):
        sig = inspect.signature(fn)
        allowed = {k: v for k, v in kwargs.items() if k in sig.parameters}
        return fn(**allowed)

    def _batch_levels(*adatas, batch_key):
        if batch_key is None:
            return ["__single_batch__"]
        batches = set()
        for ad in adatas:
            if ad is None:
                continue
            if batch_key not in ad.obs:
                raise KeyError(f"batch_key={batch_key!r} not found in adata.obs")
            batches |= set(ad.obs[batch_key].astype(str).unique().tolist())
        return sorted(batches)

    def _subset_batch(ad, batch_key, b):
        if ad is None:
            return None
        if batch_key is None:
            return ad
        mask = (ad.obs[batch_key].astype(str).values == str(b))
        if not np.any(mask):
            return None
        return ad[mask]

    def _intersect_obs_names(adatas: List):
        present = [ad for ad in adatas if ad is not None]
        if not present:
            return []
        common = set(present[0].obs_names.tolist())
        for ad in present[1:]:
            common &= set(ad.obs_names.tolist())
        # IMPORTANT: preserve a deterministic order. We'll use the reference modality order later anyway.
        return sorted(common)

    def _get_mat(ad, *, mod, layers_by_mod):
        if ad is None:
            return None
        layer = (layers_by_mod or {}).get(mod, None)
        if layer is None:
            return ad.X
        if layer not in ad.layers:
            raise KeyError(f"layers_by_mod[{mod!r}]={layer!r} not found in adata.layers")
        return ad.layers[layer]

    def _to_dense_f32(X):
        if sp.issparse(X):
            X = X.toarray()
        X = np.asarray(X, dtype=np.float32)
        if ensure_nonnegative:
            X = np.maximum(X, 0.0)
        return np.ascontiguousarray(X)

    def _shape_tree(x, depth=0, max_items=4):
        pad = "  " * depth
        if isinstance(x, dict):
            keys = list(x.keys())
            s = f"{pad}dict keys={keys[:max_items]}{'...' if len(keys)>max_items else ''}\n"
            for k in keys[:max_items]:
                s += f"{pad}  [{k}] -> " + _shape_tree(x[k], depth+2, max_items)
            return s
        if isinstance(x, (list, tuple)):
            s = f"{pad}{type(x).__name__}[len={len(x)}]\n"
            for i, xi in enumerate(x[:max_items]):
                s += _shape_tree(xi, depth+1, max_items)
            return s
        try:
            a = np.asarray(x)
            return f"{pad}array shape={getattr(a,'shape',None)} dtype={getattr(a,'dtype',None)}\n"
        except Exception:
            return f"{pad}{type(x).__name__}\n"

    def _canon_cell_factors(cf):
        # common forks return dicts
        if isinstance(cf, dict):
            for key in ("cell_factors", "Z", "H", "factors"):
                if key in cf:
                    return cf[key]
            return cf
        # some return (something, factors)
        if isinstance(cf, (tuple, list)) and len(cf) == 2:
            a, b = cf
            if isinstance(a, (list, tuple, dict)):
                return a
            if isinstance(b, (list, tuple, dict)):
                return b
        return cf

    def _unwrap_singletons(x, max_depth=8):
        y = x
        for _ in range(max_depth):
            if isinstance(y, (list, tuple)) and len(y) == 1:
                y = y[0]
            else:
                break
        return y

    def _try_parse_factors(cf_raw, mod_names: List[str], n_batches: int, K: int):
        """
        Returns:
          Z_by_mod: dict name -> list[batches] -> (cells,K)
          had_modality_specific: bool
        Accepts:
          - modality-major: cf[mod_i][batch_i] = (cells,K)
          - batch-major:    cf[batch_i] = (cells,K)
          - SINGLE output:  cf is (cells,K) OR [ (cells,K) ]  (common when nbatches=1)
          - dict formats (per-mod keys or fused)
        """
        cf = _canon_cell_factors(cf_raw)
        cf = _unwrap_singletons(cf)

        # ----- Case 0: direct (cells,K) array -----
        try:
            arr = np.asarray(cf)
            if arr.ndim == 2 and arr.shape[1] == K:
                # if nbatches>1, some forks concatenate batches; we can't safely split without sizes
                if n_batches != 1:
                    raise RuntimeError(
                        f"extract_cell_factors returned a single array (n={arr.shape[0]},K={K}) "
                        f"but nbatches={n_batches}. This fork likely concatenates batches; "
                        f"need batch sizes to split."
                    )
                return {"fused": [arr.astype(np.float32, copy=False)]}, False
        except Exception:
            pass

        # ----- Case A: modality-major list -----
        if isinstance(cf, (list, tuple)) and len(cf) == len(mod_names):
            ok = True
            for mi in range(len(mod_names)):
                vi = _unwrap_singletons(cf[mi])
                if not (isinstance(vi, (list, tuple)) and len(vi) == n_batches):
                    ok = False
                    break
                z00 = np.asarray(_unwrap_singletons(vi[0]))
                if z00.ndim != 2 or z00.shape[1] != K:
                    ok = False
                    break
            if ok:
                Z_by_mod = {}
                for mi, mname in enumerate(mod_names):
                    vi = _unwrap_singletons(cf[mi])
                    Z_by_mod[mname] = [
                        np.asarray(_unwrap_singletons(vi[bi]), dtype=np.float32)
                        for bi in range(n_batches)
                    ]
                return Z_by_mod, True

        # ----- Case B: batch-major fused list -----
        if isinstance(cf, (list, tuple)) and len(cf) == n_batches:
            z0 = np.asarray(_unwrap_singletons(cf[0]))
            if z0.ndim == 2 and z0.shape[1] == K:
                return {
                    "fused": [np.asarray(_unwrap_singletons(cf[bi]), dtype=np.float32) for bi in range(n_batches)]
                }, False

        # ----- Case C: dict with per-mod keys or fused -----
        if isinstance(cf, dict):
            # per-mod keys
            hits = [m for m in mod_names if m in cf]
            if len(hits) == len(mod_names):
                Z_by_mod = {}
                ok = True
                for m in mod_names:
                    v = _unwrap_singletons(_canon_cell_factors(cf[m]))
                    if not (isinstance(v, (list, tuple)) and len(v) == n_batches):
                        ok = False
                        break
                    z0 = np.asarray(_unwrap_singletons(v[0]))
                    if z0.ndim != 2 or z0.shape[1] != K:
                        ok = False
                        break
                    Z_by_mod[m] = [np.asarray(_unwrap_singletons(v[bi]), dtype=np.float32) for bi in range(n_batches)]
                if ok:
                    return Z_by_mod, True

            # fused key
            if "fused" in cf:
                v = _unwrap_singletons(_canon_cell_factors(cf["fused"]))
                if isinstance(v, (list, tuple)) and len(v) == n_batches:
                    z0 = np.asarray(_unwrap_singletons(v[0]))
                    if z0.ndim == 2 and z0.shape[1] == K:
                        return {"fused": [np.asarray(_unwrap_singletons(v[bi]), dtype=np.float32) for bi in range(n_batches)]}, False

        raise RuntimeError("Could not parse extract_cell_factors() into recognized structure.")

    def _auto_orient_counts(counts: Dict[str, List[np.ndarray]], mod_names: List[str], *, verbose=False):
        """
        Make orientation consistent across modalities:
          - Decide if "cells" is axis0 or axis1 by checking which axis matches across modalities.
          - If transpose_to_features_by_cells is True/False, honor it.
          - If "auto", choose an orientation where "cells" matches across modalities.
        Returns: (counts, mode_string)
        """
        # Inspect first batch (assume consistent across batches)
        b0_shapes = {m: counts[m][0].shape for m in mod_names}

        # axis0 cells hypothesis: all counts[m][0].shape[0] equal
        axis0_ok = len({b0_shapes[m][0] for m in mod_names}) == 1
        # axis1 cells hypothesis: all counts[m][0].shape[1] equal
        axis1_ok = len({b0_shapes[m][1] for m in mod_names}) == 1

        if verbose:
            print("[scmomat][orient] shapes(batch0):", b0_shapes)
            print("[scmomat][orient] axis0_ok=", axis0_ok, "axis1_ok=", axis1_ok)

        # If user forced a choice
        if transpose_to_features_by_cells is True:
            # force features x cells (transpose everything)
            for m in mod_names:
                counts[m] = [np.ascontiguousarray(x.T) for x in counts[m]]
            return counts, "forced_features_by_cells"
        if transpose_to_features_by_cells is False:
            # force cells x features (as-is)
            return counts, "forced_cells_by_features"

        # AUTO: prefer an arrangement with cells on axis0 (common), else axis1, else bail
        if axis0_ok and not axis1_ok:
            return counts, "auto_kept_cells_by_features"
        if axis1_ok and not axis0_ok:
            # transpose so cells become axis0
            for m in mod_names:
                counts[m] = [np.ascontiguousarray(x.T) for x in counts[m]]
            return counts, "auto_transposed_to_cells_by_features"
        if axis0_ok and axis1_ok:
            # ambiguous (same #cells == #features for all mods is rare, but handle)
            return counts, "auto_ambiguous_kept_cells_by_features"

        raise ValueError(
            "scMoMaT input orientation mismatch across modalities: neither axis0 nor axis1 matches as 'cells'. "
            f"batch0 shapes={b0_shapes}. This usually means you transposed only one modality or changed cell sets."
        )

    # -------------------------
    # checks + setup
    # -------------------------
    if rna is None and atac is None and adt is None:
        raise ValueError("Provide at least one modality AnnData: rna, atac, or adt.")

    if out_dir is not None:
        out_dir = Path(out_dir)
        out_dir.mkdir(parents=True, exist_ok=True)

    ref_mod = "rna" if rna is not None else ("atac" if atac is not None else "adt")
    ref_adata = {"rna": rna, "atac": atac, "adt": adt}[ref_mod]
    n_ref = int(ref_adata.n_obs)

    # seeds
    np.random.seed(int(seed))
    torch.manual_seed(int(seed))
    if torch.cuda.is_available():
        torch.cuda.manual_seed_all(int(seed))

    mods_present = [(m, ad) for (m, ad) in (("rna", rna), ("atac", atac), ("adt", adt)) if ad is not None]
    mod_names = [m for m, _ in mods_present]
    batches = _batch_levels(rna, atac, adt, batch_key=batch_key)

    mats: Dict[str, List[np.ndarray]] = {m: [] for m in mod_names}
    ref_batch_meta: List[Tuple[str, np.ndarray]] = []

    # -------------------------
    # build per-batch matrices with shared cells across modalities
    # -------------------------
    for b in batches:
        sub = {m: _subset_batch(ad, batch_key, b) for m, ad in mods_present}
        if any(sub[m] is None for m in sub):
            continue

        common = _intersect_obs_names([sub[m] for m in sub])
        if len(common) == 0:
            continue

        # Use the *reference modality order* for stable mapping + correct filling later
        # (sorted(common) can reorder; so reorder to ref subset order)
        ref_sub = sub[ref_mod]
        assert ref_sub is not None
        common_set = set(common)
        common_in_ref_order = [c for c in ref_sub.obs_names.tolist() if c in common_set]
        if len(common_in_ref_order) == 0:
            continue

        sub2 = {m: sub[m][common_in_ref_order] for m in sub}
        ref_idx_global = ref_adata.obs_names.get_indexer(common_in_ref_order).astype(int)
        if np.any(ref_idx_global < 0):
            continue

        for m, _ad in mods_present:
            Xb = _get_mat(sub2[m], mod=m, layers_by_mod=layers_by_mod)
            Xb = _to_dense_f32(Xb)  # dense float32 contiguous
            mats[m].append(Xb)

        ref_batch_meta.append((str(b), ref_idx_global))

    if len(ref_batch_meta) == 0:
        raise RuntimeError("No usable batches after subsetting+intersection across modalities.")

    counts = {m: mats[m] for m in mod_names}
    counts["nbatches"] = int(len(ref_batch_meta))

    # -------------------------
    # auto/forced orientation handling (this prevents the "sizes aren't the same" fork pain)
    # -------------------------
    counts, orient_mode = _auto_orient_counts(counts, mod_names, verbose=bool(verbose))
    if verbose:
        for m in mod_names:
            x0 = counts[m][0]
            print(f"[scmomat] mod={m} batch0 shape={x0.shape} dtype={x0.dtype} contiguous={x0.flags['C_CONTIGUOUS']}")
        print(f"[scmomat] nbatches={counts['nbatches']} orient_mode={orient_mode}")

    # -------------------------
    # build model
    # -------------------------
    dev_obj = device
    if isinstance(device, str):
        try:
            dev_obj = torch.device(device)
        except Exception:
            dev_obj = device

    Model = _safe_import_scmomat_model()
    try:
        model = Model(
            counts,
            K=int(K),
            batch_size=float(batch_size),
            interval=int(interval),
            lr=float(lr),
            lamb=float(lamb),
            seed=int(seed),
            device=dev_obj,
        )
    except Exception as e:
        raise TypeError(
            "scMoMaT construction failed.\n"
            f"counts keys={sorted(list(counts.keys()))}\n"
            f"nbatches={counts.get('nbatches')}\n"
            f"orient_mode={orient_mode}\n"
            f"Error: {type(e).__name__}({e})"
        ) from e

    # -------------------------
    # train
    # -------------------------
    t0 = time.time()
    if hasattr(model, "train_func") and callable(getattr(model, "train_func")):
        _call_with_signature(
            model.train_func,
            T=int(T),
            n_epochs=int(T),
            lr=float(lr),
            learning_rate=float(lr),
            lamb=float(lamb),
            interval=int(interval),
            verbose=bool(verbose),
            seed=int(seed),
            random_seed=int(seed),
        )
    runtime_sec = float(time.time() - t0)

    # -------------------------
    # extract factors
    # -------------------------
    if not hasattr(model, "extract_cell_factors"):
        raise AttributeError("scMoMaT model has no extract_cell_factors() in this fork.")

    cf_raw = model.extract_cell_factors()
    if verbose:
        print("[scmomat] extract_cell_factors() structure:")
        print(_shape_tree(cf_raw))

    n_batches = len(ref_batch_meta)
    Z_by_mod, had_modality_specific = _try_parse_factors(cf_raw, mod_names=mod_names, n_batches=n_batches, K=int(K))

    # -------------------------
    # fill full matrices in ref cell order
    # -------------------------
    def _fill_full(Z_full: np.ndarray, Z_batches: List[np.ndarray]):
        for (bname, ref_idx_global), Zb in zip(ref_batch_meta, Z_batches):
            Zb = np.asarray(Zb, dtype=np.float32)
            if Zb.ndim != 2 or Zb.shape[1] != int(K):
                raise RuntimeError(f"Batch {bname}: expected (cells,K)=(*,{K}) got {Zb.shape}")
            if Zb.shape[0] != len(ref_idx_global):
                raise RuntimeError(f"Batch {bname}: Z rows {Zb.shape[0]} != n_cells {len(ref_idx_global)}")
            Z_full[ref_idx_global] = Zb

    Z_rna_full = Z_atac_full = Z_adt_full = None

    if had_modality_specific:
        if "rna" in mod_names:
            Z_rna_full = np.full((n_ref, int(K)), np.nan, dtype=np.float32)
            _fill_full(Z_rna_full, Z_by_mod["rna"])
        if "atac" in mod_names:
            Z_atac_full = np.full((n_ref, int(K)), np.nan, dtype=np.float32)
            _fill_full(Z_atac_full, Z_by_mod["atac"])
        if "adt" in mod_names:
            Z_adt_full = np.full((n_ref, int(K)), np.nan, dtype=np.float32)
            _fill_full(Z_adt_full, Z_by_mod["adt"])

        # benchmark "fused" rule: average RNA/ATAC if both exist
        if (Z_rna_full is not None) and (Z_atac_full is not None):
            Z_fused = 0.5 * (Z_rna_full + Z_atac_full)
        else:
            if ref_mod == "rna" and Z_rna_full is not None:
                Z_fused = Z_rna_full
            elif ref_mod == "atac" and Z_atac_full is not None:
                Z_fused = Z_atac_full
            elif ref_mod == "adt" and Z_adt_full is not None:
                Z_fused = Z_adt_full
            else:
                first = mod_names[0]
                Z_fused = np.full((n_ref, int(K)), np.nan, dtype=np.float32)
                _fill_full(Z_fused, Z_by_mod[first])
    else:
        Z_fused = np.full((n_ref, int(K)), np.nan, dtype=np.float32)
        if "fused" in Z_by_mod:
            _fill_full(Z_fused, Z_by_mod["fused"])
        else:
            first = mod_names[0]
            _fill_full(Z_fused, Z_by_mod[first])

    # -------------------------
    # output
    # -------------------------
    out = {
        "Z": Z_fused,
        "Z_fused": Z_fused,
        "Z_rna": Z_rna_full,
        "Z_atac": Z_atac_full,
        "Z_adt": Z_adt_full,
        "runtime_sec": float(runtime_sec),
        "fit_seconds": float(runtime_sec),
        "transductive": True,
        "uses_labels": False,
        "notes": "Transductive: scMoMaT fits on all cells it embeds.",
        "mods_used": mod_names,
        "batches_used": [b for (b, _) in ref_batch_meta],
        "out_dir": str(out_dir) if out_dir is not None else None,
        "extra_json": {
            "transductive": True,
            "uses_labels": False,
            "notes": "Transductive: scMoMaT fits on all cells it embeds.",
            "had_modality_specific": bool(had_modality_specific),
            "orient_mode": orient_mode,
        },
    }

    if out_dir is not None:
        out_dir = Path(out_dir)
        np.save(out_dir / "Z_fused.npy", Z_fused)
        if Z_rna_full is not None:
            np.save(out_dir / "Z_rna.npy", Z_rna_full)
        if Z_atac_full is not None:
            np.save(out_dir / "Z_atac.npy", Z_atac_full)
        if Z_adt_full is not None:
            np.save(out_dir / "Z_adt.npy", Z_adt_full)
        with open(out_dir / "run_info.json", "w") as f:
            json.dump({k: v for k, v in out.items() if not k.startswith("Z")}, f, indent=2)

    return out
    


# %% [code]
from __future__ import annotations

import time
from pathlib import Path
from typing import Optional, Dict

import numpy as np
import scipy.sparse as sp


# --- SciPy >=1.11 compatibility: scMoMaT uses len(sparse) internally ---
try:
    def _len_shape0(self):
        return self.shape[0]
    sp.spmatrix.__len__ = _len_shape0
except Exception:
    pass


def run_scmomat_docstyle(
    *,
    rna=None,
    atac=None,
    adt=None,
    layers_by_mod: Optional[Dict[str, str]] = None,
    batch_key: Optional[str] = None,
    K: int = 30,
    T: int = 2000,
    lr: float = 1e-4,
    lamb: float = 1e-3,
    interval: int = 200,
    batch_size: float = 0.1,
    device: str = "cuda",
    seed: int = 0,
    out_dir: Optional[str] = None,
    verbose: bool = True,
) -> dict:
    """
    scMoMaT runner that matches the orientation your fork is enforcing:
      - matrices are (cells, features)
      - batches are lists (one array per batch)
    """

    if rna is None and atac is None and adt is None:
        raise ValueError("Provide at least one modality AnnData: rna, atac, or adt.")

    # pick a reference modality for output row order
    ref_mod = "rna" if rna is not None else ("atac" if atac is not None else "adt")
    ref = {"rna": rna, "atac": atac, "adt": adt}[ref_mod]
    n_ref = int(ref.n_obs)

    np.random.seed(int(seed))

    def _get_X(ad, mod: str):
        if ad is None:
            return None
        layer = (layers_by_mod or {}).get(mod, None)
        if layer is None:
            X = ad.X
        else:
            if layer not in ad.layers:
                raise KeyError(f"layers_by_mod[{mod!r}]={layer!r} not in ad.layers")
            X = ad.layers[layer]
        if sp.issparse(X):
            X = X.toarray()
        X = np.asarray(X, dtype=np.float32)
        return np.ascontiguousarray(X)  # (cells, features)

    def _batches(*ads):
        if batch_key is None:
            return ["__single_batch__"]
        vals = set()
        for ad in ads:
            if ad is None:
                continue
            if batch_key not in ad.obs:
                raise KeyError(f"batch_key={batch_key!r} not in ad.obs")
            vals |= set(ad.obs[batch_key].astype(str).unique().tolist())
        return sorted(vals)

    def _subset(ad, b):
        if ad is None or batch_key is None:
            return ad
        m = (ad.obs[batch_key].astype(str).values == str(b))
        return ad[m] if np.any(m) else None

    def _common_cells(ad_list):
        present = [a for a in ad_list if a is not None]
        common = set(present[0].obs_names.tolist())
        for a in present[1:]:
            common &= set(a.obs_names.tolist())
        return sorted(common)

    # build per-batch lists
    mods = [(m, x) for m, x in [("rna", rna), ("atac", atac), ("adt", adt)] if x is not None]
    mod_names = [m for m, _ in mods]
    batch_ids = _batches(rna, atac, adt)

    mats = {m: [] for m in mod_names}
    ref_batch_meta = []  # (batch_name, ref_global_indices)

    for b in batch_ids:
        sub = {m: _subset(ad, b) for m, ad in mods}
        if any(sub[m] is None for m in sub):
            continue

        common = _common_cells([sub[m] for m in sub])
        if len(common) == 0:
            continue

        sub = {m: sub[m][common] for m in sub}
        ref_idx = ref.obs_names.get_indexer(common).astype(int)
        if np.any(ref_idx < 0):
            continue

        for m in mod_names:
            mats[m].append(_get_X(sub[m], m))  # (cells, features)
        ref_batch_meta.append((str(b), ref_idx))

    if len(ref_batch_meta) == 0:
        raise RuntimeError("No usable batches after intersection.")

    counts = {m: mats[m] for m in mod_names}
    counts["nbatches"] = int(len(ref_batch_meta))

    if verbose:
        shapes0 = {m: mats[m][0].shape for m in mod_names}
        print(f"[scmomat] shapes(batch0)={shapes0} nbatches={counts['nbatches']} (cells x features)")

    # import + train
    import scmomat
    Model = scmomat.scmomat_model if hasattr(scmomat, "scmomat_model") else __import__("scmomat.model", fromlist=["scmomat_model"]).scmomat_model

    model = Model(
        counts,
        K=int(K),
        batch_size=float(batch_size),
        interval=int(interval),
        lr=float(lr),
        lamb=float(lamb),
        seed=int(seed),
        device=device,
    )

    t0 = time.time()
    model.train_func(T=int(T))
    runtime_sec = float(time.time() - t0)

    cf = model.extract_cell_factors()

    # Your fork returns: list[len=nbatches] of (cells, K)
    if isinstance(cf, (list, tuple)) and len(cf) == len(ref_batch_meta):
        Z_batches = [np.asarray(z, dtype=np.float32) for z in cf]
    else:
        # also handle singleton cases: [array] or array
        if isinstance(cf, (list, tuple)) and len(cf) == 1:
            Z_batches = [np.asarray(cf[0], dtype=np.float32)]
        else:
            Z_batches = [np.asarray(cf, dtype=np.float32)]

    Z_full = np.full((n_ref, int(K)), np.nan, dtype=np.float32)
    for (bname, ref_idx), Zb in zip(ref_batch_meta, Z_batches):
        if Zb.shape[0] != len(ref_idx):
            raise RuntimeError(f"Batch {bname}: Z rows {Zb.shape[0]} != n_cells {len(ref_idx)}")
        Z_full[ref_idx] = Zb

    if out_dir is not None:
        outp = Path(out_dir)
        outp.mkdir(parents=True, exist_ok=True)
        np.save(outp / "Z_fused.npy", Z_full)

    return {
        "Z_fused": Z_full,
        "Z": Z_full,
        "fit_seconds": runtime_sec,
        "transductive": True,
        "uses_labels": False,
        "extra_json": {"transductive": True, "mods_used": mod_names, "nbatches": counts["nbatches"]},
    }


# %% [markdown]
### 6) scJoint

# %% [code]
import numpy as np
import scipy.sparse as sp

def build_gene_activity_from_peaks(
    atac,
    rna,
    *,
    gene_upstream=2000,
    gene_downstream=2000,
    layer_out="gene_activity",
    dtype=np.float32,
):
    """
    Build ATAC gene activity by summing peak counts overlapping gene body windows.
    Uses rna.var['chrom','chromStart','chromEnd'] and atac.var['chrom','chromStart','chromEnd'].
    Writes atac.layers[layer_out] as CSR (cells x genes in RNA var order).
    """
    required_gene_cols = {"chrom", "chromStart", "chromEnd"}
    required_peak_cols = {"chrom", "chromStart", "chromEnd"}
    if not required_gene_cols.issubset(set(rna.var.columns)):
        raise KeyError(f"RNA var missing columns: {required_gene_cols - set(rna.var.columns)}")
    if not required_peak_cols.issubset(set(atac.var.columns)):
        raise KeyError(f"ATAC var missing columns: {required_peak_cols - set(atac.var.columns)}")

    # peak coords
    p_chr = np.asarray(atac.var["chrom"]).astype(str)
    p_start = np.asarray(atac.var["chromStart"]).astype(np.int64)
    p_end   = np.asarray(atac.var["chromEnd"]).astype(np.int64)

    # gene coords (RNA order)
    g_chr = np.asarray(rna.var["chrom"]).astype(str)
    g_start = np.asarray(rna.var["chromStart"]).astype(np.int64) - int(gene_upstream)
    g_end   = np.asarray(rna.var["chromEnd"]).astype(np.int64)   + int(gene_downstream)
    g_start = np.maximum(g_start, 0)

    # build COO edges: peak_idx -> gene_idx if intervals overlap
    rows = []
    cols = []

    # per-chrom sweep for overlap edges
    for chrom in np.unique(np.intersect1d(np.unique(p_chr), np.unique(g_chr))):
        p_idx = np.where(p_chr == chrom)[0]
        g_idx = np.where(g_chr == chrom)[0]
        if p_idx.size == 0 or g_idx.size == 0:
            continue

        # sort peaks by start
        p_ord = p_idx[np.argsort(p_start[p_idx], kind="mergesort")]
        ps = p_start[p_ord]
        pe = p_end[p_ord]

        # sort genes by start
        g_ord = g_idx[np.argsort(g_start[g_idx], kind="mergesort")]
        gs = g_start[g_ord]
        ge = g_end[g_ord]

        j0 = 0
        for gi, (s, e) in zip(g_ord, zip(gs, ge)):
            # advance j0 until peak end >= gene start (can't overlap before this)
            while j0 < p_ord.size and pe[j0] < s:
                j0 += 1
            j = j0
            # walk forward while peak start <= gene end
            while j < p_ord.size and ps[j] <= e:
                # overlap condition
                if pe[j] >= s:
                    rows.append(p_ord[j])  # peak index
                    cols.append(gi)        # gene index (RNA var index)
                j += 1

    if len(rows) == 0:
        raise RuntimeError("No peak↔gene overlaps found; cannot build gene activity.")

    rows = np.asarray(rows, dtype=np.int64)
    cols = np.asarray(cols, dtype=np.int64)
    data = np.ones(rows.shape[0], dtype=np.int8)

    # peak->gene mapping (peaks x genes)
    P2G = sp.coo_matrix((data, (rows, cols)), shape=(atac.n_vars, rna.n_vars)).tocsr()

    Xp = atac.X
    if not sp.issparse(Xp):
        Xp = sp.csr_matrix(np.asarray(Xp))
    Xp = Xp.tocsr()

    # gene activity (cells x genes)
    Xg = (Xp @ P2G).astype(dtype).tocsr()

    # put it in obsm (allowed to be cells x genes)
    atac.obsm[layer_out] = Xg

    # optionally store gene names used for columns
    atac.uns[f"{layer_out}__gene_names"] = rna.var_names.to_numpy()

    return atac


# %% [code]
atac_for_scjoint = atac.copy()
atac_for_scjoint = build_gene_activity_from_peaks(atac_for_scjoint, rna, layer_out="gene_activity", gene_upstream=50000, gene_downstream=50000)

print("gene_activity layer:", atac_for_scjoint.obsm["gene_activity"].shape)


# %% [code]
import numpy as np
import pandas as pd
import scipy.sparse as sp

def restrict_gene_activity_to_rna_features(
    atac,
    rna_feat_adata,
    *,
    ga_key="gene_activity",
    out_key="gene_activity_hvg",
):
    """
    Takes atac.obsm[ga_key] (cells x genes_in_full_rna_space) and slices columns
    to match rna_feat_adata.var_names exactly (order preserved).

    Requires atac.uns[f"{ga_key}__gene_names"] that you saved in build_gene_activity_from_peaks().
    Writes atac.obsm[out_key] and atac.uns[f"{out_key}__gene_names"].
    Returns out_key.
    """
    if ga_key not in atac.obsm:
        raise KeyError(f"{ga_key!r} not in atac.obsm. Available: {list(atac.obsm.keys())}")

    all_genes = atac.uns.get(f"{ga_key}__gene_names", None)
    if all_genes is None:
        raise KeyError(f"Missing atac.uns['{ga_key}__gene_names']; can't map gene-activity columns.")

    X = atac.obsm[ga_key]
    if not sp.issparse(X):
        X = sp.csr_matrix(np.asarray(X))
    X = X.tocsr()

    target_genes = pd.Index(rna_feat_adata.var_names.astype(str))
    all_genes = pd.Index(np.asarray(all_genes).astype(str))

    col_idx = all_genes.get_indexer(target_genes)
    missing = np.where(col_idx < 0)[0]
    if missing.size:
        ex = target_genes[missing[:10]].tolist()
        raise ValueError(
            f"{missing.size} / {len(target_genes)} RNA features not found in gene-activity gene list. "
            f"Examples: {ex}"
        )

    atac.obsm[out_key] = X[:, col_idx].astype(np.float32).tocsr()
    atac.uns[f"{out_key}__gene_names"] = target_genes.to_numpy()

    return out_key


# %% [code]
# Slice gene activity to match the RNA HVG space used for scJoint
ga_key_for_scjoint = restrict_gene_activity_to_rna_features(
    atac_for_scjoint, rna_counts_hvg, ga_key="gene_activity", out_key="gene_activity_hvg"
)

print("ATAC gene activity (HVG) shape:", atac_for_scjoint.obsm[ga_key_for_scjoint].shape)
print("RNA HVG shape:", rna_counts_hvg.n_obs, rna_counts_hvg.n_vars)


# %% [code]
# --- scJoint runner (robust + notebook-friendly; patched for all known issues) ---
from __future__ import annotations

import os, sys, time, shutil, subprocess
from pathlib import Path

import numpy as np
import scipy.sparse as sp


def _as_csr_float32(X):
    if sp.issparse(X):
        X = X.tocsr()
        if X.dtype != np.float32:
            X = X.astype(np.float32)
        return X
    return sp.csr_matrix(np.asarray(X, dtype=np.float32))


def run_scjoint(
    adata_rna,
    adata_atac,
    *,
    labels_key,                       # obs col name OR array-like labels
    atac_gene_activity_layer,          # REQUIRED: gene activity in atac.obsm or atac.layers
    scjoint_repo,                     # path to your cloned SydneyBioX/scJoint
    out_dir,
    seed=0,
    embedding_size=30,
    batch_size=256,
    gpu=0,
    epochs_stage1=50,
    epochs_stage3=50,
    lr_stage1=1e-3,
    lr_stage3=1e-3,
    momentum=0.9,
    center_weight=1,
    p=0.8,
    with_crossentropy=True,
    threads=12,
):
    """
    Runs scJoint via subprocess in a way that works on clusters / notebooks.

    Key requirements:
      - ATAC input must be GENE-ACTIVITY with same #features as RNA input
      - We patch: np.float removal, non-tty stty progress bar, torch dynamo/compile
    """
    t0 = time.perf_counter()
    out_dir = Path(out_dir)
    out_dir.mkdir(parents=True, exist_ok=True)

    repo = Path(scjoint_repo)
    if not repo.exists():
        raise FileNotFoundError(f"scJoint repo not found at: {repo}")

    main_src = repo / "main.py"
    if not main_src.exists():
        raise FileNotFoundError(f"Couldn't find main.py in scJoint repo: {main_src}")

    # ---- labels ----
    if isinstance(labels_key, str):
        y = adata_rna.obs[labels_key].astype(str).to_numpy()
    else:
        y = np.asarray(labels_key).astype(str)

    classes, y_int = np.unique(y, return_inverse=True)
    n_classes = int(len(classes))

    # ---- matrices ----
    X_rna = _as_csr_float32(adata_rna.X)

    if atac_gene_activity_layer is None:
        raise ValueError("atac_gene_activity_layer is required for scJoint (must be gene-space matrix).")

    if atac_gene_activity_layer in adata_atac.obsm:
        X_atac = _as_csr_float32(adata_atac.obsm[atac_gene_activity_layer])
    elif atac_gene_activity_layer in adata_atac.layers:
        X_atac = _as_csr_float32(adata_atac.layers[atac_gene_activity_layer])
    else:
        raise KeyError(
            f"Couldn't find atac_gene_activity_layer='{atac_gene_activity_layer}' in atac.obsm or atac.layers. "
            f"Available obsm={list(adata_atac.obsm.keys())[:10]}..., layers={list(adata_atac.layers.keys())}"
        )

    if X_atac.shape[0] != X_rna.shape[0]:
        raise ValueError(f"RNA/ATAC cell count mismatch: RNA {X_rna.shape}, ATAC {X_atac.shape}")

    if X_atac.shape[1] != X_rna.shape[1]:
        raise ValueError(
            "scJoint requires ATAC gene activity to have SAME number of features as RNA.\n"
            f"Got RNA features={X_rna.shape[1]}, ATAC gene-activity features={X_atac.shape[1]}.\n"
            "Fix by constructing ATAC gene activity in the same gene space/order as your RNA features (e.g. HVGs)."
        )

    input_size = int(X_rna.shape[1])

    # ---- write inputs ----
    data_dir = out_dir / "data"
    data_dir.mkdir(exist_ok=True)

    rna_npz = data_dir / "rna.npz"
    atac_npz = data_dir / "atac_gene_activity.npz"
    labels_txt = data_dir / "labels.txt"

    sp.save_npz(str(rna_npz), X_rna)
    sp.save_npz(str(atac_npz), X_atac)
    labels_txt.write_text("\n".join(map(str, y_int.tolist())) + "\n")

    # ---- write config.py expected by scJoint main.py ----
    # Must define class Config with required attributes (threads, rna_protein_paths, atac_protein_paths, etc.)
    cfg_py = out_dir / "config.py"
    cfg_py.write_text(
        f"""
import torch

class Config(object):
    def __init__(self):
        self.use_cuda = True
        self.threads = {int(threads)}

        if not self.use_cuda:
            self.device = torch.device('cpu')
        else:
            self.device = torch.device('cuda:0')

        # dataset info
        self.number_of_class = {int(n_classes)}
        self.input_size = {int(input_size)}

        self.rna_paths = ["{rna_npz}"]
        self.rna_labels = ["{labels_txt}"]

        self.atac_paths = ["{atac_npz}"]
        self.atac_labels = []  # optional

        # MUST exist even if unused
        self.rna_protein_paths = []
        self.atac_protein_paths = []

        # training
        self.batch_size = {int(batch_size)}
        self.lr_stage1 = {float(lr_stage1)}
        self.lr_stage3 = {float(lr_stage3)}
        self.lr_decay_epoch = 20
        self.epochs_stage1 = {int(epochs_stage1)}
        self.epochs_stage3 = {int(epochs_stage3)}
        self.p = {float(p)}
        self.embedding_size = {int(embedding_size)}
        self.momentum = {float(momentum)}
        self.center_weight = {float(center_weight)}
        self.with_crossentorpy = {bool(with_crossentropy)}
        self.seed = {int(seed)}
        self.checkpoint = ""

        # some forks read this
        self.save_dir = "{(out_dir / "output")}"
"""
    )

    # ---- copy main.py into out_dir so it imports our config.py ----
    main_dst = out_dir / "main.py"
    shutil.copy2(str(main_src), str(main_dst))

    # ---- sitecustomize.py patches for numpy + stty progress bar ----
    # - numpy 2.x removed np.float / np.int / np.bool
    # - scJoint progress bar tries: os.popen("stty size") and crashes in non-tty
    sitecustomize = out_dir / "sitecustomize.py"
    sitecustomize.write_text(
        "import os, io\n"
        "import numpy as np\n"
        "\n"
        "# ---- NumPy 2.x compatibility (scJoint uses np.float etc.) ----\n"
        "if not hasattr(np, 'float'): np.float = float\n"
        "if not hasattr(np, 'int'):   np.int = int\n"
        "if not hasattr(np, 'bool'):  np.bool = bool\n"
        "\n"
        "# ---- scJoint progress bar fix for non-TTY ----\n"
        "_orig_popen = os.popen\n"
        "def _patched_popen(cmd, mode='r', *args, **kwargs):\n"
        "    try:\n"
        "        s = cmd.strip() if isinstance(cmd, str) else ''\n"
        "        if s.startswith('stty size'):\n"
        "            return io.StringIO('24 120\\n')\n"
        "    except Exception:\n"
        "        pass\n"
        "    return _orig_popen(cmd, mode, *args, **kwargs)\n"
        "os.popen = _patched_popen\n"
    )

    # ---- env for subprocess ----
    env = os.environ.copy()
    env["PYTHONHASHSEED"] = str(int(seed))

    # Disable torch dynamo/compile (avoids sympy / torch._dynamo / fx symbolic shapes issues)
    env["TORCH_DISABLE_DYNAMO"] = "1"
    env["TORCHDYNAMO_DISABLE"] = "1"
    env["TORCH_COMPILE_DISABLE"] = "1"

    # Make sure sitecustomize is found FIRST (out_dir must precede repo on PYTHONPATH)
    env["PYTHONPATH"] = str(out_dir) + os.pathsep + str(repo) + os.pathsep + env.get("PYTHONPATH", "")

    # GPU selection
    if gpu is None:
        env["CUDA_VISIBLE_DEVICES"] = ""
    else:
        env["CUDA_VISIBLE_DEVICES"] = str(int(gpu))

    # ---- run ----
    proc = subprocess.run(
        [sys.executable, str(main_dst)],
        cwd=str(out_dir),
        env=env,
        stdout=subprocess.PIPE,
        stderr=subprocess.STDOUT,
        text=True,
    )

    log_path = out_dir / "scjoint_run.log"
    log_path.write_text(proc.stdout)

    if proc.returncode != 0:
        tail = "\n".join(proc.stdout.splitlines()[-80:])
        raise RuntimeError(
            "scJoint failed. See scjoint_run.log for details.\n"
            f"Last 80 lines:\n{tail}"
        )

    # ---- find embeddings ----
    # Different forks dump in different places; search widely.
    cand_dirs = [out_dir / "output", out_dir, out_dir / "results", out_dir / "Result", out_dir / "log"]
    emb_files = []
    for d in cand_dirs:
        if d.exists():
            emb_files.extend(list(d.rglob("*embeddings*.txt")))
            emb_files.extend(list(d.rglob("*embedding*.txt")))
            emb_files.extend(list(d.rglob("*latent*.txt")))
    emb_files = sorted({p.resolve() for p in emb_files})

    if len(emb_files) < 2:
        # dump some directory listing hints
        existing = []
        for d in cand_dirs:
            if d.exists():
                existing.extend([str(p.relative_to(d)) for p in d.rglob("*")][:50])
        raise RuntimeError(
            "scJoint finished but I couldn't find 2 embedding files.\n"
            f"Searched in: {[str(d) for d in cand_dirs]}\n"
            f"Found: {[str(p) for p in emb_files]}\n"
            f"Sample files under output dirs: {existing}\n"
            f"Log: {log_path}"
        )

    # Heuristic: take the two largest embedding-ish files (usually RNA/ATAC)
    emb_files = sorted(emb_files, key=lambda p: p.stat().st_size, reverse=True)[:2]
    Z0 = np.loadtxt(str(emb_files[0])).astype(np.float32)
    Z1 = np.loadtxt(str(emb_files[1])).astype(np.float32)

    if Z0.shape != Z1.shape:
        raise RuntimeError(f"Embedding shape mismatch: {emb_files[0]} {Z0.shape} vs {emb_files[1]} {Z1.shape}")

    Z_rna, Z_atac = Z0, Z1
    Z_fused = 0.5 * (Z_rna + Z_atac)

    fit_seconds = float(time.perf_counter() - t0)

    return ensure_flags({
        "Z_rna": Z_rna,
        "Z_atac": Z_atac,
        "Z_fused": Z_fused,
        "fit_seconds": fit_seconds,
        "extra_json": standard_flags(
            transductive=True,
            uses_labels=True,
            note="scJoint trained with RNA labels; fit included all cells."
        ),
    }, default_transductive=True, default_uses_labels=True)


# %% [code]
from __future__ import annotations

import os, sys, time, shutil, subprocess
from pathlib import Path

import numpy as np
import scipy.sparse as sp


def _as_csr_float32(X):
    if sp.issparse(X):
        X = X.tocsr(copy=False)
        return X.astype(np.float32, copy=False) if X.dtype != np.float32 else X
    return sp.csr_matrix(np.asarray(X, dtype=np.float32))


def _scrub_finite(Z: np.ndarray, *, name: str, verbose: bool = True) -> np.ndarray:
    """Replace NaN/Inf with 0.0."""
    Z = np.asarray(Z, dtype=np.float32)
    bad = ~np.isfinite(Z)
    if bad.any():
        if verbose:
            n_nan = int(np.isnan(Z).sum())
            n_inf = int(np.isinf(Z).sum())
            print(f"[scJoint] WARNING: {name} had non-finite: nan={n_nan} inf={n_inf} -> set to 0")
        Z = Z.copy()
        Z[bad] = 0.0
    return Z


def run_scjoint_split_aware(
    adata_rna,
    adata_atac,
    *,
    splits,
    labels_key,
    atac_gene_activity_layer,
    scjoint_repo,
    out_dir,
    seed=0,
    latent_dim=30,
    batch_size=256,
    gpu=0,
    epochs_stage1=50,
    epochs_stage3=50,
    lr_stage1=1e-3,
    lr_stage3=1e-3,
    momentum=0.9,
    center_weight=1,
    p=0.8,
    with_crossentropy=True,
    threads=12,
    verbose=True,
    # -------------------------
    # NEW knobs
    # -------------------------
    allow_transductive_fallback: bool = False,
    fill_missing: str = "nan",  # "nan" (recommended) or "zeros"
):
    """
    scJoint runner with explicit behavior:

    - Inductive (default): train on FIT=train(+val) only.
        * FIT rows get embeddings
        * non-FIT rows are missing (NaN by default, or zeros if fill_missing="zeros")
        * Metrics that evaluate on TEST will be NaN if TEST is entirely missing (expected)

    - Transductive fallback (optional): if allow_transductive_fallback=True and a test split exists,
      we train on ALL cells to produce embeddings for TEST (sets transductive=True in flags).
    """
    t0 = time.perf_counter()
    out_dir = Path(out_dir)
    out_dir.mkdir(parents=True, exist_ok=True)

    fill_missing = str(fill_missing).lower()
    if fill_missing not in ("nan", "zeros"):
        raise ValueError("fill_missing must be 'nan' or 'zeros'")

    # splits
    n_full = int(adata_rna.n_obs)
    tr = np.asarray(splits["train"], dtype=int)
    va = np.asarray(splits.get("val", []), dtype=int)
    te = np.asarray(splits.get("test", []), dtype=int)

    fit_idx = np.concatenate([tr, va]) if va.size else tr
    fit_idx = np.asarray(fit_idx, dtype=int)

    transductive = False
    if allow_transductive_fallback and te.size > 0:
        # train on ALL so scJoint produces embeddings for everyone
        fit_idx = np.arange(n_full, dtype=int)
        transductive = True

    # subset for training
    rna_fit = adata_rna[fit_idx].copy()
    atac_fit = adata_atac[fit_idx].copy()

    repo = Path(scjoint_repo)
    if not repo.exists():
        raise FileNotFoundError(f"scJoint repo not found at: {repo}")
    main_src = repo / "main.py"
    if not main_src.exists():
        raise FileNotFoundError(f"Couldn't find main.py in scJoint repo: {main_src}")

    # ---- labels on FIT ----
    if isinstance(labels_key, str):
        if labels_key not in rna_fit.obs:
            raise KeyError(f"labels_key={labels_key!r} not in rna_fit.obs")
        y_fit = rna_fit.obs[labels_key].astype(str).to_numpy()
    else:
        y_all = np.asarray(labels_key).astype(str)
        if y_all.shape[0] != n_full:
            raise ValueError(f"labels array len={y_all.shape[0]} != n_full={n_full}")
        y_fit = y_all[fit_idx]

    classes, y_int = np.unique(y_fit, return_inverse=True)
    n_classes = int(len(classes))

    # ---- matrices ----
    X_rna = _as_csr_float32(rna_fit.X)

    if atac_gene_activity_layer is None:
        raise ValueError("atac_gene_activity_layer is required for scJoint (gene-space matrix).")

    if atac_gene_activity_layer in atac_fit.obsm:
        X_atac = _as_csr_float32(atac_fit.obsm[atac_gene_activity_layer])
    elif atac_gene_activity_layer in atac_fit.layers:
        X_atac = _as_csr_float32(atac_fit.layers[atac_gene_activity_layer])
    else:
        raise KeyError(
            f"Couldn't find atac_gene_activity_layer='{atac_gene_activity_layer}' in atac.obsm or atac.layers. "
            f"obsm={list(atac_fit.obsm.keys())[:10]}..., layers={list(atac_fit.layers.keys())}"
        )

    if X_atac.shape[0] != X_rna.shape[0]:
        raise ValueError(f"RNA/ATAC cell count mismatch: RNA {X_rna.shape}, ATAC {X_atac.shape}")
    if X_atac.shape[1] != X_rna.shape[1]:
        raise ValueError(
            "scJoint requires ATAC gene activity to have SAME features as RNA.\n"
            f"RNA p={X_rna.shape[1]}, ATAC gene-activity p={X_atac.shape[1]}"
        )

    input_size = int(X_rna.shape[1])

    # ---- write inputs ----
    data_dir = out_dir / "data"
    data_dir.mkdir(exist_ok=True)

    rna_npz = data_dir / "rna.npz"
    atac_npz = data_dir / "atac_gene_activity.npz"
    labels_txt = data_dir / "labels.txt"

    sp.save_npz(str(rna_npz), X_rna)
    sp.save_npz(str(atac_npz), X_atac)
    labels_txt.write_text("\n".join(map(str, y_int.tolist())) + "\n")

    # ---- config.py ----
    cfg_py = out_dir / "config.py"
    cfg_py.write_text(
        f"""
import torch
class Config(object):
    def __init__(self):
        self.use_cuda = True
        self.threads = {int(threads)}
        self.device = torch.device('cuda:0') if self.use_cuda else torch.device('cpu')

        self.number_of_class = {int(n_classes)}
        self.input_size = {int(input_size)}

        self.rna_paths = ["{rna_npz}"]
        self.rna_labels = ["{labels_txt}"]
        self.atac_paths = ["{atac_npz}"]
        self.atac_labels = []

        self.rna_protein_paths = []
        self.atac_protein_paths = []

        self.batch_size = {int(batch_size)}
        self.lr_stage1 = {float(lr_stage1)}
        self.lr_stage3 = {float(lr_stage3)}
        self.lr_decay_epoch = 20
        self.epochs_stage1 = {int(epochs_stage1)}
        self.epochs_stage3 = {int(epochs_stage3)}
        self.p = {float(p)}
        self.embedding_size = {int(latent_dim)}
        self.momentum = {float(momentum)}
        self.center_weight = {float(center_weight)}
        self.with_crossentorpy = {bool(with_crossentropy)}
        self.seed = {int(seed)}
        self.checkpoint = ""
        self.save_dir = "{(out_dir / "output")}"
"""
    )

    # ---- copy main.py ----
    main_dst = out_dir / "main.py"
    shutil.copy2(str(main_src), str(main_dst))

    # ---- sitecustomize.py patches ----
    sitecustomize = out_dir / "sitecustomize.py"
    sitecustomize.write_text(
        "import os, io\n"
        "import numpy as np\n"
        "if not hasattr(np, 'float'): np.float = float\n"
        "if not hasattr(np, 'int'):   np.int = int\n"
        "if not hasattr(np, 'bool'):  np.bool = bool\n"
        "_orig_popen = os.popen\n"
        "def _patched_popen(cmd, mode='r', *args, **kwargs):\n"
        "    try:\n"
        "        s = cmd.strip() if isinstance(cmd, str) else ''\n"
        "        if s.startswith('stty size'):\n"
        "            return io.StringIO('24 120\\n')\n"
        "    except Exception:\n"
        "        pass\n"
        "    return _orig_popen(cmd, mode, *args, **kwargs)\n"
        "os.popen = _patched_popen\n"
    )

    # ---- env ----
    env = os.environ.copy()
    env["PYTHONHASHSEED"] = str(int(seed))
    env["TORCH_DISABLE_DYNAMO"] = "1"
    env["TORCHDYNAMO_DISABLE"] = "1"
    env["TORCH_COMPILE_DISABLE"] = "1"
    env["PYTHONPATH"] = str(out_dir) + os.pathsep + str(repo) + os.pathsep + env.get("PYTHONPATH", "")
    env["CUDA_VISIBLE_DEVICES"] = "" if gpu is None else str(int(gpu))

    # ---- run ----
    proc = subprocess.run(
        [sys.executable, str(main_dst)],
        cwd=str(out_dir),
        env=env,
        stdout=subprocess.PIPE,
        stderr=subprocess.STDOUT,
        text=True,
        check=False,
    )

    log_path = out_dir / "scjoint_run.log"
    log_path.write_text(proc.stdout)

    if proc.returncode != 0:
        tail = "\n".join(proc.stdout.splitlines()[-120:])
        raise RuntimeError(
            "scJoint failed. See scjoint_run.log for details.\n"
            f"Last 120 lines:\n{tail}"
        )

    # ---- find embeddings ----
    cand_dirs = [out_dir / "output", out_dir, out_dir / "results", out_dir / "Result", out_dir / "log"]
    emb_files = []
    for d in cand_dirs:
        if d.exists():
            emb_files.extend(list(d.rglob("*embeddings*.txt")))
            emb_files.extend(list(d.rglob("*embedding*.txt")))
            emb_files.extend(list(d.rglob("*latent*.txt")))
    emb_files = sorted({p.resolve() for p in emb_files})

    if len(emb_files) < 2:
        raise RuntimeError(f"scJoint finished but couldn't find embeddings txts under {cand_dirs}. Log: {log_path}")

    def _score(p: Path):
        s = p.name.lower()
        score = 0
        if "rna" in s: score += 10
        if "atac" in s or "acc" in s: score += 10
        if "joint" in s or "fused" in s: score += 2
        return score

    emb_sorted = sorted(emb_files, key=lambda p: (_score(p), p.stat().st_size), reverse=True)
    p0, p1 = emb_sorted[0], emb_sorted[1]

    Z0 = np.loadtxt(str(p0)).astype(np.float32)
    Z1 = np.loadtxt(str(p1)).astype(np.float32)

    fit_n = len(fit_idx)
    if Z0.ndim != 2 or Z1.ndim != 2 or Z0.shape[0] != fit_n or Z1.shape[0] != fit_n:
        # fallback: pick any two matching fit_n
        cands = []
        for p in emb_files:
            try:
                z = np.loadtxt(str(p))
                if z.ndim == 2 and z.shape[0] == fit_n:
                    cands.append((p, z.astype(np.float32)))
            except Exception:
                pass
        if len(cands) < 2:
            raise RuntimeError(
                f"Found embedding txts but none matched expected n_cells={fit_n}. "
                f"Example {p0.name}={Z0.shape}"
            )
        cands = sorted(cands, key=lambda t: t[0].stat().st_size, reverse=True)[:2]
        p0, p1 = cands[0][0], cands[1][0]
        Z0, Z1 = cands[0][1], cands[1][1]

    if Z0.shape != Z1.shape:
        raise RuntimeError(f"Embedding shape mismatch: {p0} {Z0.shape} vs {p1} {Z1.shape}")

    Z0 = _scrub_finite(Z0, name=f"Z0({p0.name})", verbose=verbose)
    Z1 = _scrub_finite(Z1, name=f"Z1({p1.name})", verbose=verbose)

    emb_dim = int(Z0.shape[1])

    # outputs
    embed_mask = np.zeros(n_full, dtype=bool)
    embed_mask[fit_idx] = True

    if fill_missing == "zeros":
        Z_full  = np.zeros((n_full, emb_dim), dtype=np.float32)
        Zr_full = np.zeros_like(Z_full)
        Za_full = np.zeros_like(Z_full)
    else:
        Z_full  = np.full((n_full, emb_dim), np.nan, dtype=np.float32)
        Zr_full = np.full_like(Z_full, np.nan)
        Za_full = np.full_like(Z_full, np.nan)

    Zr_full[fit_idx] = Z0
    Za_full[fit_idx] = Z1
    Z_full[fit_idx]  = 0.5 * (Z0 + Z1)

    # sanity: embedded rows must be finite
    if not np.isfinite(Z_full[fit_idx]).all():
        bad = int((~np.isfinite(Z_full[fit_idx])).sum())
        raise RuntimeError(f"[scJoint] BUG: embedded rows have non-finite after scrub: {bad}")

    fit_seconds = float(time.perf_counter() - t0)

    if verbose:
        n_embedded = int(embed_mask.sum())
        n_missing = int(n_full - n_embedded)
        miss_desc = "zeros" if fill_missing == "zeros" else "NaN"
        tr_desc = "ALL (transductive)" if transductive else "train(+val) only"
        print(
            f"[scJoint] emb_dim={emb_dim} | trained_on={tr_desc}: embedded {n_embedded}/{n_full} "
            f"(missing={n_missing}; missing_as={miss_desc}) | files=({p0.name}, {p1.name})"
        )

    # NOTE: keep your ensure_flags/standard_flags calls
    return ensure_flags({
        "Z_rna": Zr_full,
        "Z_atac": Za_full,
        "Z_fused": Z_full,
        "embed_mask": embed_mask,
        "fit_seconds": fit_seconds,
        "extra_json": standard_flags(
            transductive=transductive,
            uses_labels=True,
            note=(
                "Trained on ALL cells (transductive) to embed test."
                if transductive else
                f"Trained on train(+val) only; non-FIT cells are {miss_desc}."
            ) + f" emb_dim={emb_dim}."
        ),
    }, default_transductive=transductive, default_uses_labels=True)


# %% [markdown]
### 7) DeepCCA

# %% [code]
class _MLP(torch.nn.Module):
    def __init__(self, in_dim, hidden, out_dim, dropout=0.1):
        super().__init__()
        layers = []
        d = in_dim
        for h in hidden:
            layers += [torch.nn.Linear(d, h), torch.nn.ReLU(), torch.nn.Dropout(dropout)]
            d = h
        layers += [torch.nn.Linear(d, out_dim)]
        self.net = torch.nn.Sequential(*layers)

    def forward(self, x):
        return self.net(x)


def _inv_sqrtm_psd(A, eps=1e-6):
    # A: (d,d) symmetric PSD
    # returns A^{-1/2} using eigendecomp
    w, V = torch.linalg.eigh(A)
    w = torch.clamp(w, min=eps)
    return (V * (w.rsqrt())) @ V.T

def deepcca_loss_cholesky(H1, H2, reg=1e-2, eps=1e-6, max_tries=6):
    """
    Stable DeepCCA loss: uses Cholesky whitening instead of eigendecomp.
    Returns negative sum of canonical correlations (minimize).
    H1,H2: (b,d)
    """
    # Use float64 for stability; cast back not needed for loss scalar
    H1 = H1.double()
    H2 = H2.double()

    H1 = H1 - H1.mean(dim=0, keepdim=True)
    H2 = H2 - H2.mean(dim=0, keepdim=True)

    b = H1.shape[0]
    I1 = torch.eye(H1.shape[1], device=H1.device, dtype=H1.dtype)
    I2 = torch.eye(H2.shape[1], device=H2.device, dtype=H2.dtype)

    C11 = (H1.T @ H1) / (b - 1)
    C22 = (H2.T @ H2) / (b - 1)
    C12 = (H1.T @ H2) / (b - 1)

    # Robust Cholesky with increasing jitter
    L1 = L2 = None
    for t in range(max_tries):
        jitter = float(reg) + (10.0**t) * float(eps)
        try:
            L1 = torch.linalg.cholesky(C11 + jitter * I1)
            L2 = torch.linalg.cholesky(C22 + jitter * I2)
            break
        except RuntimeError:
            continue
    if L1 is None or L2 is None:
        # last-resort fallback: add big jitter and try once more
        jitter = float(reg) + (10.0**max_tries) * float(eps)
        L1 = torch.linalg.cholesky(C11 + jitter * I1)
        L2 = torch.linalg.cholesky(C22 + jitter * I2)

    # T = L1^{-1} C12 L2^{-T}
    # Solve L1 * X = C12
    X = torch.linalg.solve_triangular(L1, C12, upper=False)
    # Solve L2 * Y = X^T  -> Y = L2^{-1} X^T, then transpose back gives X L2^{-T}
    Y = torch.linalg.solve_triangular(L2, X.T, upper=False).T
    T = Y

    s = torch.linalg.svdvals(T)
    corr = torch.sum(s)
    return -corr

def deepcca_loss(H1, H2, reg=1e-3, eps=1e-6):
    """
    Negative sum of canonical correlations (to minimize).
    H1,H2: (b,d)
    """
    H1 = H1 - H1.mean(dim=0, keepdim=True)
    H2 = H2 - H2.mean(dim=0, keepdim=True)

    b = H1.shape[0]
    C11 = (H1.T @ H1) / (b - 1) + reg * torch.eye(H1.shape[1], device=H1.device)
    C22 = (H2.T @ H2) / (b - 1) + reg * torch.eye(H2.shape[1], device=H2.device)
    C12 = (H1.T @ H2) / (b - 1)

    C11_inv_sqrt = _inv_sqrtm_psd(C11, eps=eps)
    C22_inv_sqrt = _inv_sqrtm_psd(C22, eps=eps)

    T = C11_inv_sqrt @ C12 @ C22_inv_sqrt
    # sum singular values = total correlation
    s = torch.linalg.svdvals(T)
    corr = torch.sum(s)
    return -corr


def run_deepcca(
    rna_log_hvg,
    atac_lsi,
    *,
    out_dir,
    splits,
    seed=0,
    latent_dim=30,
    hidden=(512, 256, 128),
    dropout=0.05,
    lr=1e-3,
    weight_decay=1e-4,
    batch_size=256,
    max_epochs=200,
    patience=50,
    reg=1e-3,
    clip_grad=5.0,
    #align_weight=0.0,   # set to ~0.05–0.2 if you want stronger geometric mixing
    align_weight=0.05,
):
    """
    DeepCCA baseline.
    - Aligns RNA/ATAC by shared obs_names (critical).
    - Remaps splits from original RNA index space -> aligned shared index space.
    - Standardizes each modality using TRAIN stats only.
    - Trains on TRAIN, early-stops on VAL using the same Cholesky CCA loss.
    - Returns Z_rna, Z_atac, Z_fused=mean(Zr,Za).
    """
    from pathlib import Path
    import numpy as np
    import scipy.sparse as sp
    import torch
    from sklearn.preprocessing import StandardScaler

    set_seed(seed)
    out_dir = Path(out_dir)
    out_dir.mkdir(parents=True, exist_ok=True)
    device = "cuda" if torch.cuda.is_available() else "cpu"

    # ------------------------------------------------------------
    # 0) Align modalities to SAME cells in SAME order
    # ------------------------------------------------------------
    rna0 = rna_log_hvg
    atac0 = atac_lsi

    shared = rna0.obs_names.intersection(atac0.obs_names)
    if shared.size == 0:
        raise ValueError("DeepCCA: RNA/ATAC have no shared cells (obs_names intersection empty).")

    # canonical order = RNA order
    rna = rna0[shared].copy()
    atac = atac0[shared].copy()
    if not rna.obs_names.equals(atac.obs_names):
        raise ValueError("DeepCCA: failed to align RNA/ATAC obs_names after subsetting.")

    n_obs = int(rna.n_obs)

    # ------------------------------------------------------------
    # 1) Remap splits (original RNA index space -> aligned shared space)
    # ------------------------------------------------------------
    tr0 = np.asarray(splits["train"], dtype=int)
    va0 = np.asarray(splits["val"], dtype=int)

    orig_pos = rna0.obs_names.get_indexer(shared)  # positions of shared cells in original RNA
    if np.any(orig_pos < 0):
        raise ValueError("DeepCCA: internal error computing RNA indexer for shared cells.")

    inv = np.full(int(rna0.n_obs), -1, dtype=int)
    inv[orig_pos] = np.arange(shared.size, dtype=int)

    tr = inv[tr0]
    tr = tr[tr >= 0]
    va = inv[va0]
    va = va[va >= 0]

    if tr.size == 0:
        raise ValueError("DeepCCA: no TRAIN cells remain after aligning modalities.")
    if va.size == 0:
        raise ValueError("DeepCCA: no VAL cells remain after aligning modalities.")

    if int(batch_size) < int(latent_dim) + 8:
        raise ValueError(f"DeepCCA: batch_size ({batch_size}) too small for latent_dim ({latent_dim}).")

    # ------------------------------------------------------------
    # 2) Build dense matrices
    # ------------------------------------------------------------
    Xr = rna.X
    Xa = atac.X
    if sp.issparse(Xr):
        Xr = Xr.toarray()
    if sp.issparse(Xa):
        Xa = Xa.toarray()
    Xr = np.asarray(Xr, dtype=np.float32)
    Xa = np.asarray(Xa, dtype=np.float32)

    if Xr.shape[0] != n_obs or Xa.shape[0] != n_obs:
        raise ValueError(f"DeepCCA: n_obs mismatch after align: RNA {Xr.shape[0]} vs ATAC {Xa.shape[0]}")

    # ------------------------------------------------------------
    # 3) Train-fit scaling per modality
    # ------------------------------------------------------------
    sc_r = StandardScaler(with_mean=True, with_std=True)
    sc_a = StandardScaler(with_mean=True, with_std=True)
    sc_r.fit(Xr[tr])
    sc_a.fit(Xa[tr])
    Xr_s = sc_r.transform(Xr).astype(np.float32, copy=False)
    Xa_s = sc_a.transform(Xa).astype(np.float32, copy=False)

    if not (np.isfinite(Xr_s).all() and np.isfinite(Xa_s).all()):
        raise ValueError("DeepCCA: non-finite values detected after scaling.")

    # ------------------------------------------------------------
    # 4) Models + optimizer
    # ------------------------------------------------------------
    enc_r = _MLP(Xr_s.shape[1], list(hidden), int(latent_dim), dropout=dropout).to(device)
    enc_a = _MLP(Xa_s.shape[1], list(hidden), int(latent_dim), dropout=dropout).to(device)

    opt = torch.optim.AdamW(
        list(enc_r.parameters()) + list(enc_a.parameters()),
        lr=float(lr),
        weight_decay=float(weight_decay),
    )

    def _iter_batches(idxs):
        idxs = np.asarray(idxs, dtype=int).copy()
        rng = np.random.default_rng(int(seed))
        rng.shuffle(idxs)
        for i in range(0, len(idxs), int(batch_size)):
            yield idxs[i:i + int(batch_size)]

    # consistent loss (Cholesky CCA)
    def _cca_loss(hr, ha):
        return deepcca_loss_cholesky(hr, ha, reg=float(reg), eps=1e-6)

    best_val = np.inf
    best_state = None
    bad = 0

    # ------------------------------------------------------------
    # 5) Train loop
    # ------------------------------------------------------------
    t0 = now()
    for epoch in range(int(max_epochs)):
        enc_r.train()
        enc_a.train()

        for b in _iter_batches(tr):
            xr = torch.from_numpy(Xr_s[b]).to(device)
            xa = torch.from_numpy(Xa_s[b]).to(device)

            hr = enc_r(xr)
            ha = enc_a(xa)

            loss = _cca_loss(hr, ha)
            if float(align_weight) > 0:
                loss = loss + float(align_weight) * torch.mean((hr - ha) ** 2)

            opt.zero_grad(set_to_none=True)
            loss.backward()

            if clip_grad is not None and float(clip_grad) > 0:
                torch.nn.utils.clip_grad_norm_(
                    list(enc_r.parameters()) + list(enc_a.parameters()),
                    max_norm=float(clip_grad),
                )

            opt.step()

        # --------------------------------------------------------
        # Validation (same loss as train)
        # --------------------------------------------------------
        enc_r.eval()
        enc_a.eval()
        with torch.no_grad():
            vals = []
            for b in _iter_batches(va):
                xr = torch.from_numpy(Xr_s[b]).to(device)
                xa = torch.from_numpy(Xa_s[b]).to(device)
                hr = enc_r(xr)
                ha = enc_a(xa)
                v = _cca_loss(hr, ha)
                if float(align_weight) > 0:
                    v = v + float(align_weight) * torch.mean((hr - ha) ** 2)
                vals.append(float(v.cpu()))
            val_loss = float(np.mean(vals)) if len(vals) else np.inf

        if val_loss < best_val - 1e-6:
            best_val = val_loss
            best_state = (enc_r.state_dict(), enc_a.state_dict())
            bad = 0
        else:
            bad += 1
            if bad >= int(patience):
                break

    if best_state is not None:
        enc_r.load_state_dict(best_state[0])
        enc_a.load_state_dict(best_state[1])

    # ------------------------------------------------------------
    # 6) Encode all aligned cells
    # ------------------------------------------------------------
    enc_r.eval()
    enc_a.eval()
    with torch.no_grad():
        Zr = enc_r(torch.from_numpy(Xr_s).to(device)).cpu().numpy().astype(np.float32)
        Za = enc_a(torch.from_numpy(Xa_s).to(device)).cpu().numpy().astype(np.float32)

    t1 = now()

    if Zr.shape != Za.shape or Zr.shape[0] != n_obs:
        raise ValueError(f"DeepCCA: unexpected embedding shapes: Zr {Zr.shape}, Za {Za.shape}, n_obs={n_obs}")

    Zf = 0.5 * (Zr + Za)

    return ensure_flags({
        "Z_rna": Zr,
        "Z_atac": Za,
        "Z_fused": Zf,
        "fit_seconds": float(t1 - t0),
        "extra_json": standard_flags(
            transductive=False,
            uses_labels=False,
            model="DeepCCA",
            latent_dim=int(latent_dim),
            hidden=list(hidden),
            dropout=float(dropout),
            lr=float(lr),
            weight_decay=float(weight_decay),
            batch_size=int(batch_size),
            max_epochs=int(max_epochs),
            patience=int(patience),
            reg=float(reg),
            clip_grad=float(clip_grad) if clip_grad is not None else None,
            align_weight=float(align_weight),
            n_shared=int(n_obs),
            n_train_shared=int(tr.size),
            n_val_shared=int(va.size),
            best_val=float(best_val),
            note="Aligned RNA/ATAC by shared obs_names; splits remapped; standardized on TRAIN; Cholesky CCA loss used for train+val."
        ),
    }, default_transductive=False, default_uses_labels=False)


# %% [markdown]
### 8) PeakVI

# %% [code]
def run_peakvi(
    atac_counts_bin_hv,
    *,
    out_dir,
    splits,
    seed=0,
    n_latent=30,
    max_epochs=200,
    patience=50,
    batch_key=None,
    layer=None,
):
    """
    PEAKVI baseline (ATAC-only).

    Trains on TRAIN+VAL only (so: transductive=False).
    Returns:
      Z_atac  : latent embedding for ATAC (all cells; inference for test)
      Z_fused : same as Z_atac (unimodal)
      Z_rna   : None
    """
    import numpy as np
    from pathlib import Path
    import scvi
    from scvi.model import PEAKVI

    set_seed(seed)
    out_dir = Path(out_dir); out_dir.mkdir(parents=True, exist_ok=True)

    tr = np.asarray(splits["train"])
    va = np.asarray(splits["val"])
    order_tv = np.concatenate([tr, va])

    # -----------------------
    # Train on train+val only
    # -----------------------
    atac_tv = atac_counts_bin_hv[order_tv].copy()

    PEAKVI.setup_anndata(
        atac_tv,
        layer=layer,
        batch_key=batch_key,
    )

    t0 = now()
    model = PEAKVI(atac_tv, n_latent=int(n_latent))

    scvi_train_with_patience(
        model,
        max_epochs=int(max_epochs),
        patience=int(patience),
        monitor="elbo_validation",
        train_size=len(tr) / len(order_tv),
        validation_size=len(va) / len(order_tv),
        shuffle_set_split=False,
    )
    t1 = now()

    # -----------------------
    # Embed ALL cells (no fit)
    # -----------------------
    atac_all = atac_counts_bin_hv.copy()

    # IMPORTANT: do NOT call PEAKVI.setup_anndata(atac_all, ...) again.
    # Just pass it to the model for inference.
    Z = np.asarray(model.get_latent_representation(adata=atac_all), dtype=np.float32)

    return ensure_flags({
        "Z_rna": None,
        "Z_atac": Z,
        "Z_fused": Z,
        "fit_seconds": float(t1 - t0),
        "extra_json": standard_flags(
            transductive=False,
            uses_labels=False,
            scvi_version=scvi.__version__,
            model="PEAKVI",
            n_latent=int(n_latent),
            trained_on="train+val",
            layer=layer,
            batch_key=batch_key,
            note="ATAC-only; Z_fused == Z_atac. Test cells are inference only."
        ),
    }, default_transductive=False, default_uses_labels=False)


# %% [code]
def run_peakvi_fair(
    atac_counts_bin_hv,
    *,
    out_dir,
    splits,
    seed=0,
    n_latent=30,
    max_epochs=200,
    patience=50,
    batch_key=None,
    layer=None,
    encode_batch_size=256,
):
    import numpy as np
    import shutil
    from pathlib import Path
    import scvi
    from scvi.model import PEAKVI

    set_seed(seed)
    out_dir = Path(out_dir); out_dir.mkdir(parents=True, exist_ok=True)

    tr = np.asarray(splits["train"], dtype=int)
    va = np.asarray(splits["val"], dtype=int)
    order_tv = np.concatenate([tr, va])

    # ---- train on train+val only ----
    atac_tv = atac_counts_bin_hv[order_tv].copy()
    PEAKVI.setup_anndata(atac_tv, layer=layer, batch_key=batch_key)

    t0 = now()
    model = PEAKVI(atac_tv, n_latent=int(n_latent))
    scvi_train_with_patience(
        model,
        max_epochs=int(max_epochs),
        patience=int(patience),
        monitor="elbo_validation",
        train_size=len(tr) / len(order_tv),
        validation_size=len(va) / len(order_tv),
        shuffle_set_split=False,
        check_val_every_n_epoch=1,
    )
    t1 = now()

    # ---- save + reload onto FULL data for stable inference ----
    save_dir = out_dir / "peakvi_model"
    if save_dir.exists():
        shutil.rmtree(save_dir)
    model.save(str(save_dir), overwrite=True)

    atac_full = atac_counts_bin_hv.copy()
    PEAKVI.setup_anndata(atac_full, layer=layer, batch_key=batch_key)

    model_full = PEAKVI.load(str(save_dir), adata=atac_full)

    Z = np.asarray(
        model_full.get_latent_representation(adata=atac_full, batch_size=int(encode_batch_size)),
        dtype=np.float32,
    )

    return ensure_flags({
        "Z_rna": None,
        "Z_atac": Z,
        "Z_fused": Z,
        "fit_seconds": float(t1 - t0),
        "extra_json": standard_flags(
            transductive=False,
            uses_labels=False,
            scvi_version=scvi.__version__,
            model="PEAKVI",
            n_latent=int(n_latent),
            trained_on="train+val",
            note="Saved+reloaded onto full AnnData for inference; training used train+val only."
        ),
    }, default_transductive=False, default_uses_labels=False)


# %% [markdown]
### 9) CoBOLT

# %% [code]
import importlib, pkgutil, sys
print("cobolt in pkgutil?", any(m.name == "cobolt" for m in pkgutil.iter_modules()))
import cobolt
print("cobolt module file:", cobolt.__file__)


# %% [code]
from __future__ import annotations

import time
import inspect
import traceback
from pathlib import Path
from typing import Optional, Dict, Any, List, Sequence

import numpy as np
import scipy.sparse as sp
import torch


def run_cobolt_working(
    rna_adata,
    atac_adata,
    *,
    splits: Optional[dict] = None,
    rna_key: Optional[str] = "counts",
    atac_key: Optional[str] = None,
    prefer_layer: bool = True,
    n_latent: int = 30,
    max_epochs: int = 200,
    batch_size: int = 256,
    device: str = "cuda",
    seed: int = 0,
    require_paired: bool = True,

    # ATAC TF-IDF
    atac_tfidf: bool = False,
    atac_tfidf_l2_norm: bool = False,
    tfidf_smooth_idf: bool = False,

    # NEW: RNA normalization (highly recommended for CoBOLT stability)
    rna_cp10k_log1p: bool = False,
    rna_target_sum: float = 1e4,

    out_dir: Optional[str] = None,
    verbose: bool = True,
    latent_cap: int = 60,

    # LR control + retry
    lr: Optional[float] = None,
    lr_try: Optional[Sequence[float]] = (5e-3, 1e-3, 3e-4, 1e-4, 3e-5, 1e-5, 3e-6, 1e-6),
    max_lr_retries: int = 15,
) -> dict:
    """
    CoBOLT runner (your version) with:
      - scoped scipy.sparse.vstack patch
      - optional ATAC TF-IDF
      - NEW: RNA cp10k + log1p (sparse-safe) to prevent divergence
      - LR retry ladder
      - latent extraction via bool masks len=2:
           [True,True]=joint, [True,False]=rna, [False,True]=atac
    """

    def log(msg: str):
        if verbose:
            print(msg, flush=True)

    # -------------------------
    # Safe manual CSR vstack
    # -------------------------
    def _csr_vstack(mats):
        mats = list(mats)
        if len(mats) == 0:
            return sp.csr_matrix((0, 0), dtype=np.float32)

        mats = [m.tocsr(copy=False) for m in mats]
        n_cols = int(mats[0].shape[1])
        total_rows = int(sum(m.shape[0] for m in mats))
        if total_rows == 0:
            return sp.csr_matrix((0, n_cols), dtype=np.float32)

        data = np.concatenate([m.data for m in mats]) if mats else np.array([], dtype=np.float32)
        indices = np.concatenate([m.indices for m in mats]) if mats else np.array([], dtype=np.int32)

        indptr = np.empty(total_rows + 1, dtype=np.int64)
        indptr[0] = 0
        row_pos = 0
        nnz_off = 0
        for m in mats:
            r = int(m.shape[0])
            indptr[row_pos + 1: row_pos + r + 1] = nnz_off + m.indptr[1:]
            row_pos += r
            nnz_off += int(m.indptr[-1])

        return sp.csr_matrix((data, indices, indptr), shape=(total_rows, n_cols))

    # -------------------------
    # Scoped patch: scipy.sparse.vstack
    # -------------------------
    class _ScopedPatchScipyVstack:
        def __init__(self, verbose: bool = True):
            self.verbose = verbose
            self._saved = []

        def __enter__(self):
            import scipy.sparse as _sp
            try:
                import scipy.sparse._construct as _c
            except Exception:
                _c = None

            def vstack_safe(blocks, *args, **kwargs):
                flat = []
                stack = [blocks]
                while stack:
                    x = stack.pop()
                    if x is None:
                        continue
                    if sp.issparse(x):
                        flat.append(x)
                        continue
                    if isinstance(x, np.ndarray):
                        if x.dtype == object or x.ndim != 2:
                            try:
                                elems = x.reshape(-1).tolist()
                            except Exception:
                                elems = [x]
                            stack.extend(reversed(elems))
                        else:
                            flat.append(sp.csr_matrix(x))
                        continue
                    if isinstance(x, (list, tuple)):
                        stack.extend(reversed(x))
                        continue
                    arr = np.asarray(x)
                    if arr.ndim == 2 and arr.dtype != object:
                        flat.append(sp.csr_matrix(arr))
                    else:
                        raise TypeError(f"vstack_safe: unsupported block type {type(x)} shape={getattr(x,'shape',None)}")
                return _csr_vstack(flat)

            self._saved.append((_sp, "vstack", _sp.vstack))
            _sp.vstack = vstack_safe

            if _c is not None and hasattr(_c, "vstack"):
                self._saved.append((_c, "vstack", _c.vstack))
                _c.vstack = vstack_safe

            if self.verbose:
                print("[cobolt][patch] scoped scipy.sparse.vstack -> safe manual csr_vstack", flush=True)
            return self

        def __exit__(self, exc_type, exc, tb):
            for obj, name, old in reversed(self._saved):
                try:
                    setattr(obj, name, old)
                except Exception:
                    pass
            self._saved.clear()
            if self.verbose:
                print("[cobolt][patch] restored scipy.sparse.vstack", flush=True)
            return False

    # -------------------------
    # helpers
    # -------------------------
    def _resolve_matrix(adata, key: Optional[str]):
        if key is None:
            return adata.X, "X"
        layers = getattr(adata, "layers", {})
        obsm = getattr(adata, "obsm", {})
        if prefer_layer and key in layers:
            return layers[key], f"layers[{key}]"
        if key in obsm:
            return obsm[key], f"obsm[{key}]"
        if key in layers:
            return layers[key], f"layers[{key}]"
        raise KeyError(f"{key!r} not found in .layers or .obsm")

    def _as_csr_f32_2d(X, src: str):
        if sp.issparse(X):
            X = X.tocsr(copy=False)
            if X.ndim != 2:
                raise ValueError(f"{src} sparse not 2D: shape={X.shape}")
            return X.astype(np.float32, copy=False)
        X = np.asarray(X)
        if X.ndim != 2:
            raise ValueError(f"{src} not 2D: shape={X.shape}")
        if X.dtype == object:
            raise ValueError(f"{src} dtype=object (ragged?) shape={X.shape}")
        return sp.csr_matrix(X.astype(np.float32, copy=False))

    def _rna_cp10k_log1p(X: sp.csr_matrix, target_sum: float = 1e4, eps: float = 1e-12) -> sp.csr_matrix:
        """Sparse-safe: scale rows to target_sum then log1p(data) (in-place-ish on CSR data)."""
        X = X.tocsr(copy=True).astype(np.float32, copy=False)
        rs = np.asarray(X.sum(axis=1)).ravel().astype(np.float32)
        scale = (float(target_sum) / np.maximum(rs, eps)).astype(np.float32)

        # apply row scaling without densifying: multiply each row's data by scale[row]
        indptr = X.indptr
        for i in range(X.shape[0]):
            start, end = indptr[i], indptr[i + 1]
            if start != end:
                X.data[start:end] *= scale[i]

        # log1p on stored nonzeros
        np.log1p(X.data, out=X.data)
        return X

    def _tfidf_fit_idf(X_train_csr: sp.csr_matrix, *, smooth_idf: bool = True) -> np.ndarray:
        X = X_train_csr.tocsr(copy=False)
        N = int(X.shape[0])
        df = np.asarray((X > 0).sum(axis=0)).ravel().astype(np.float64)
        if smooth_idf:
            idf = np.log1p(N / (1.0 + df)) + 1.0
        else:
            df = np.maximum(df, 1.0)
            idf = np.log(N / df) + 1.0
        return idf.astype(np.float32)

    def _tfidf_apply(X_csr: sp.csr_matrix, *, idf: np.ndarray, l2_norm: bool = True, eps: float = 1e-12) -> sp.csr_matrix:
        X = X_csr.tocsr(copy=False).astype(np.float32, copy=False)
        rs = np.asarray(X.sum(axis=1)).ravel().astype(np.float32)
        inv_rs = (1.0 / np.maximum(rs, eps)).astype(np.float32)
        tf = X.multiply(inv_rs[:, None])
        tfidf = tf.multiply(np.asarray(idf, dtype=np.float32)[None, :])
        if not l2_norm:
            return tfidf.tocsr(copy=False)
        row_sq = np.asarray(tfidf.multiply(tfidf).sum(axis=1)).ravel().astype(np.float32)
        inv_norm = (1.0 / np.sqrt(np.maximum(row_sq, eps))).astype(np.float32)
        return tfidf.multiply(inv_norm[:, None]).tocsr(copy=False)

    def _cap(Z: np.ndarray, name: str) -> np.ndarray:
        if latent_cap is not None and Z.shape[1] > int(latent_cap):
            log(f"[cobolt][safe] {name} dim {Z.shape[1]} > {latent_cap}; truncating to {latent_cap}")
            return Z[:, : int(latent_cap)].copy()
        return Z

    def _to_full(Z_fit: np.ndarray, n_full: int, fit_idx: np.ndarray) -> np.ndarray:
        Z_full = np.full((n_full, Z_fit.shape[1]), np.nan, dtype=np.float32)
        Z_full[fit_idx] = Z_fit
        return Z_full

    def _looks_like_divergence(e: Exception) -> bool:
        s = (str(e) or "").lower()
        return ("diverged" in s) or ("diverged." in s) or ("try a smaller learning rate" in s) or ("nan" in s and "loss" in s)

    def _train_with_lr(model, epochs: int, lr_value: Optional[float]) -> None:
        train_fn = model.train
        sig = inspect.signature(train_fn)
        params = sig.parameters

        kwargs = {}
        if "num_epochs" in params:
            kwargs["num_epochs"] = int(epochs)
        elif "epochs" in params:
            kwargs["epochs"] = int(epochs)

        if lr_value is not None:
            if "lr" in params:
                kwargs["lr"] = float(lr_value)
            elif "learning_rate" in params:
                kwargs["learning_rate"] = float(lr_value)

        if kwargs:
            train_fn(**kwargs)
        else:
            train_fn(int(epochs))

    def _check_finite_sparse(X: sp.csr_matrix, name: str):
        if X.nnz == 0:
            return
        if not np.isfinite(X.data).all():
            bad = np.where(~np.isfinite(X.data))[0][:10]
            raise ValueError(f"{name} contains non-finite values in .data (first bad idx: {bad}).")

    try:
        # seed
        import random
        random.seed(seed)
        np.random.seed(seed)
        torch.manual_seed(seed)
        if torch.cuda.is_available():
            torch.cuda.manual_seed_all(seed)

        # pairing checks
        if require_paired:
            if rna_adata.n_obs != atac_adata.n_obs:
                raise ValueError(f"RNA/ATAC n_obs differ: {rna_adata.n_obs} vs {atac_adata.n_obs}")
            if not np.all(rna_adata.obs_names.astype(str) == atac_adata.obs_names.astype(str)):
                raise ValueError("RNA and ATAC obs_names differ. Reorder/subset to match for paired runs.")

        # subset to FIT
        n_full = int(rna_adata.n_obs)
        if splits is None:
            fit_idx = np.arange(n_full, dtype=int)
            train_note = "all"
        else:
            tr = np.asarray(splits["train"], dtype=int)
            va = np.asarray(splits.get("val", []), dtype=int)
            fit_idx = np.concatenate([tr, va]) if va.size else tr
            fit_idx = np.asarray(fit_idx, dtype=int)
            train_note = "train+val"

        rna_fit = rna_adata[fit_idx].copy()
        atac_fit = atac_adata[fit_idx].copy()

        # matrices
        Xr_raw, rna_src = _resolve_matrix(rna_fit, rna_key)
        Xa_raw, atac_src = _resolve_matrix(atac_fit, atac_key)
        Xr = _as_csr_f32_2d(Xr_raw, rna_src)
        Xa = _as_csr_f32_2d(Xa_raw, atac_src)

        # NEW: normalize RNA
        if rna_cp10k_log1p:
            Xr = _rna_cp10k_log1p(Xr, target_sum=float(rna_target_sum))
            rna_src = rna_src + " -> cp10k+log1p"

        # ATAC TF-IDF
        if atac_tfidf:
            if splits is not None:
                tr = np.asarray(splits["train"], dtype=int)
                tr_set = set(tr.tolist())
                tr_in_fit = np.array([i in tr_set for i in fit_idx.tolist()], dtype=bool)
                if tr_in_fit.sum() < 2:
                    tr_in_fit[:] = True
            else:
                tr_in_fit = np.ones(Xa.shape[0], dtype=bool)

            idf = _tfidf_fit_idf(Xa[tr_in_fit], smooth_idf=bool(tfidf_smooth_idf))
            Xa = _tfidf_apply(Xa, idf=idf, l2_norm=bool(atac_tfidf_l2_norm))
            atac_src = atac_src + " -> TFIDF"

        _check_finite_sparse(Xr, "RNA matrix")
        _check_finite_sparse(Xa, "ATAC matrix")

        log(f"[cobolt] training_on={train_note} n_fit={rna_fit.n_obs}")
        log(f"[cobolt] RNA source={rna_src} shape={Xr.shape} nnz={Xr.nnz}")
        log(f"[cobolt] ATAC source={atac_src} shape={Xa.shape} nnz={Xa.nnz}")

        # import CoBOLT components
        from cobolt.utils.data import SingleData, MultiData
        from cobolt.utils.dataset import MultiomicDataset
        from cobolt.model import Cobolt

        def _make_singledata(*, X, barcodes, features, dataset_name: str, feature_name: str):
            barcodes = np.asarray(barcodes, dtype=str)
            features = np.asarray(features, dtype=str)

            sig = inspect.signature(SingleData.__init__)
            params = set(sig.parameters.keys()) - {"self"}

            base = {
                "count": X,
                "feature": features,
                "barcode": barcodes,
                "dataset_name": str(dataset_name),
                "feature_name": str(feature_name),
                "dataset": np.zeros(len(barcodes), dtype=np.int64),
            }
            aliases = {"counts": "count", "barcodes": "barcode", "genes": "feature"}

            kw: Dict[str, Any] = {}
            for k, v in base.items():
                if k in params:
                    kw[k] = v
            for k, src in aliases.items():
                if k in params and src in base and k not in kw:
                    kw[k] = base[src]
            return SingleData(**kw)

        bc = np.asarray(rna_fit.obs_names, dtype=str)
        rna_feat = np.asarray(rna_fit.var_names, dtype=str)
        atac_feat = np.asarray(atac_fit.var_names, dtype=str)

        a = _make_singledata(X=Xr, barcodes=bc, features=rna_feat, dataset_name="joint", feature_name="rna")
        b = _make_singledata(X=Xa, barcodes=bc, features=atac_feat, dataset_name="joint", feature_name="atac")

        with _ScopedPatchScipyVstack(verbose=verbose):
            md = MultiData(a, b)
            ds = MultiomicDataset(md)

            model = Cobolt(dataset=ds, n_latent=int(n_latent), device=device, batch_size=int(batch_size))

            # training with LR retries
            lr_candidates: List[Optional[float]] = []
            if lr is not None:
                lr_candidates.append(float(lr))
            if lr_try is not None:
                for x in lr_try:
                    xx = float(x)
                    if not any((c is not None and abs(c - xx) < 1e-20) for c in lr_candidates):
                        lr_candidates.append(xx)
            if not lr_candidates:
                lr_candidates = [None]

            last_train_err = None
            used_lr = None

            t0 = time.perf_counter()
            for attempt, lr_value in enumerate(lr_candidates[: max_lr_retries]):
                try:
                    used_lr = lr_value
                    if lr_value is None:
                        log("[cobolt] training with cobolt default lr")
                    else:
                        log(f"[cobolt] training with lr={lr_value:g}")
                    _train_with_lr(model, int(max_epochs), lr_value)
                    last_train_err = None
                    break
                except Exception as e:
                    last_train_err = e
                    if _looks_like_divergence(e):
                        log(f"[cobolt][warn] training diverged (attempt {attempt+1}); will retry smaller lr")
                        continue
                    raise

            fit_seconds = float(time.perf_counter() - t0)
            if last_train_err is not None:
                raise RuntimeError(f"CoBOLT training failed after LR retries. Last error: {last_train_err!r}") from last_train_err

            # latent extraction (bool list len 2)
            sig_lat = inspect.signature(model.get_latent)
            log(f"[cobolt] get_latent signature: {sig_lat}")
            has_data_kw = ("data" in sig_lat.parameters)
            data_candidates = ["all", "full", "train"] if has_data_kw else [None]

            def _get_latent_or_raise(comb: List[bool], kind: str):
                last_err = None
                for dname in data_candidates:
                    for rb in (False, True):
                        try:
                            kwargs = {"return_barcode": rb}
                            if has_data_kw and dname is not None:
                                kwargs["data"] = dname
                            out = model.get_latent(comb, **kwargs)
                            out0 = out[0] if (isinstance(out, tuple) and len(out) >= 1) else out
                            if torch.is_tensor(out0):
                                out0 = out0.detach().cpu().numpy()
                            Z = np.asarray(out0, dtype=np.float32)
                            if Z.ndim != 2:
                                raise RuntimeError(f"get_latent({kind}) returned non-2D: shape={Z.shape}")
                            if Z.shape[0] != rna_fit.n_obs:
                                raise RuntimeError(
                                    f"get_latent({kind}) rows {Z.shape[0]} != n_fit {rna_fit.n_obs} "
                                    f"(comb={comb}, data={dname!r}, rb={rb})"
                                )
                            log(f"[cobolt] got_latent {kind} comb={comb} data={dname!r} return_barcode={rb} -> {Z.shape}")
                            return Z, (dname if dname is not None else "default")
                        except Exception as e:
                            last_err = e
                raise RuntimeError(f"get_latent({kind}) failed for comb={comb}. Last error: {last_err!r}") from last_err

            Z_joint_fit, data_used_joint = _get_latent_or_raise([True, True],  "joint")
            Z_rna_fit,   data_used_rna   = _get_latent_or_raise([True, False], "rna")
            Z_atac_fit,  data_used_atac  = _get_latent_or_raise([False, True], "atac")

        Z_joint_fit = _cap(Z_joint_fit, "Z_joint")
        Z_rna_fit   = _cap(Z_rna_fit,   "Z_rna")
        Z_atac_fit  = _cap(Z_atac_fit,  "Z_atac")

        Z_joint = _to_full(Z_joint_fit, n_full, fit_idx)
        Z_rna   = _to_full(Z_rna_fit,   n_full, fit_idx)
        Z_atac  = _to_full(Z_atac_fit,  n_full, fit_idx)

        if out_dir is not None:
            outp = Path(out_dir)
            outp.mkdir(parents=True, exist_ok=True)
            np.save(outp / "Z_fused.npy", Z_joint)
            np.save(outp / "Z_rna.npy", Z_rna)
            np.save(outp / "Z_atac.npy", Z_atac)

        return {
            "Z_fused": Z_joint,
            "Z": Z_joint,
            "Z_rna": Z_rna,
            "Z_atac": Z_atac,
            "fit_seconds": float(fit_seconds),
            "transductive": True,
            "uses_labels": False,
            "extra_json": {
                "trained_on": train_note,
                "n_latent_requested": int(n_latent),
                "joint_dim_returned": int(Z_joint_fit.shape[1]),
                "rna_preproc": f"cp{rna_target_sum:g}+log1p" if rna_cp10k_log1p else "none",
                "atac_tfidf": bool(atac_tfidf),
                "patched_scipy_sparse_vstack_scoped": True,
                "data_used": {"joint": data_used_joint, "rna": data_used_rna, "atac": data_used_atac},
                "latent_cap": int(latent_cap) if latent_cap is not None else None,
                "lr_used": used_lr,
                "lr_try": list(lr_try) if lr_try is not None else None,
            },
        }

    except Exception:
        print("========== [cobolt] FULL TRACEBACK ==========")
        print(traceback.format_exc())
        print("========== [cobolt] END TRACEBACK ==========")
        raise


# %% [markdown]
### Orchestrate + evaluate everything

# %% [code]
torch.set_float32_matmul_precision("high")

(Path(os.environ["XDG_CACHE_HOME"]) / "torch" / "kernels").mkdir(parents=True, exist_ok=True)

import warnings
warnings.filterwarnings(
    "ignore",
    message="The argument 'device' of Tensor\\.pin_memory\\(\\) is deprecated.*",
    category=DeprecationWarning,
)
warnings.filterwarnings(
    "ignore",
    message="The argument 'device' of Tensor\\.is_pinned\\(\\) is deprecated.*",
    category=DeprecationWarning,
)


# %% [markdown]
### 1) Run + evaluate

# %% [markdown]
### Helper/evaluation/run all function definitions

# %% [code]
import numpy as np
from sklearn.neighbors import NearestNeighbors
from sklearn.metrics import (
    accuracy_score, f1_score, balanced_accuracy_score,
    silhouette_score, adjusted_rand_score, normalized_mutual_info_score
)
from sklearn.cluster import KMeans


# -------------------------
# NaN helpers
# -------------------------
def _finite_row_mask(Z):
    Z = np.asarray(Z)
    if Z.ndim != 2:
        raise ValueError(f"Expected 2D embedding, got shape={Z.shape}")
    return np.isfinite(Z).all(axis=1)

def _subset_finite_rows(Z, idx):
    """Return (Z_sub, idx_sub) keeping only finite rows within idx."""
    Z = np.asarray(Z)
    idx = np.asarray(idx, dtype=int)
    m = _finite_row_mask(Z[idx])
    idx2 = idx[m]
    return Z[idx2], idx2

def _paired_finite_on_idx(Za, Zb, idx):
    """Return (Za_sub, Zb_sub, idx_sub) where BOTH are finite."""
    Za = np.asarray(Za); Zb = np.asarray(Zb)
    idx = np.asarray(idx, dtype=int)
    ma = _finite_row_mask(Za[idx])
    mb = _finite_row_mask(Zb[idx])
    m = ma & mb
    idx2 = idx[m]
    return Za[idx2], Zb[idx2], idx2

def _seed_int(seed, default=0):
    """Return a real int seed, even if seed is None / nan / weird."""
    if seed is None:
        return int(default)
    try:
        # handle numpy scalars, floats, etc.
        if isinstance(seed, (float, np.floating)) and not np.isfinite(seed):
            return int(default)
        return int(seed)
    except Exception:
        return int(default)


# -------------------------
# FOSCTTM (now None/NaN-safe)
# -------------------------
def foscttm_values(Za, Zb, *, subsample=3000, seed=0, topk=(1, 5, 10, 50, 100)):
    Za = np.asarray(Za); Zb = np.asarray(Zb)
    assert Za.shape[0] == Zb.shape[0]

    n = int(Za.shape[0])
    if n < 2:
        return {
            "idx": np.arange(n, dtype=int),
            "fos_a2b": np.array([], dtype=np.float32),
            "fos_b2a": np.array([], dtype=np.float32),
            "fos_mean_bidir": np.array([], dtype=np.float32),
            "mean": np.nan, "sem": np.nan,
            "mrr_a2b": np.nan, "mrr_b2a": np.nan, "mrr_mean": np.nan,
            **{f"recall@{k}_a2b": np.nan for k in topk},
            **{f"recall@{k}_b2a": np.nan for k in topk},
            **{f"recall@{k}_mean": np.nan for k in topk},
        }

    # subsample handling:
    # - None => use all
    # - <=0  => return NaNs (nothing to compute)
    if subsample is None:
        m = n
    else:
        subsample = int(subsample)
        if subsample <= 0:
            return {
                "idx": np.array([], dtype=int),
                "fos_a2b": np.array([], dtype=np.float32),
                "fos_b2a": np.array([], dtype=np.float32),
                "fos_mean_bidir": np.array([], dtype=np.float32),
                "mean": np.nan, "sem": np.nan,
                "mrr_a2b": np.nan, "mrr_b2a": np.nan, "mrr_mean": np.nan,
                **{f"recall@{k}_a2b": np.nan for k in topk},
                **{f"recall@{k}_b2a": np.nan for k in topk},
                **{f"recall@{k}_mean": np.nan for k in topk},
            }
        m = min(subsample, n)

    rng = np.random.default_rng(int(seed) if seed is not None else 0)
    idx = rng.choice(n, size=m, replace=False)

    A = Za[idx]; B = Zb[idx]
    m = int(A.shape[0])
    if m < 2:
        # can happen if n==1, but we already guarded; keep safe anyway
        return {
            "idx": idx.astype(int),
            "fos_a2b": np.array([], dtype=np.float32),
            "fos_b2a": np.array([], dtype=np.float32),
            "fos_mean_bidir": np.array([], dtype=np.float32),
            "mean": np.nan, "sem": np.nan,
            "mrr_a2b": np.nan, "mrr_b2a": np.nan, "mrr_mean": np.nan,
            **{f"recall@{k}_a2b": np.nan for k in topk},
            **{f"recall@{k}_b2a": np.nan for k in topk},
            **{f"recall@{k}_mean": np.nan for k in topk},
        }

    A2 = np.sum(A*A, axis=1, keepdims=True)
    B2 = np.sum(B*B, axis=1, keepdims=True).T
    D = np.maximum(A2 + B2 - 2.0*(A @ B.T), 0.0)

    diag = np.diag(D)
    rank_a2b = (D < diag[:, None]).sum(axis=1) + 1
    rank_b2a = (D.T < diag[:, None]).sum(axis=1) + 1

    fos_a2b = (rank_a2b - 1) / (m - 1)
    fos_b2a = (rank_b2a - 1) / (m - 1)
    fos_mean = 0.5 * (fos_a2b + fos_b2a)

    out = {
        "idx": idx.astype(int),
        "fos_a2b": fos_a2b,
        "fos_b2a": fos_b2a,
        "fos_mean_bidir": fos_mean,
        "mean": float(np.mean(fos_mean)),
        "sem": float(np.std(fos_mean, ddof=1) / np.sqrt(m)),
        "mrr_a2b": float(np.mean(1.0 / rank_a2b)),
        "mrr_b2a": float(np.mean(1.0 / rank_b2a)),
        "mrr_mean": float(0.5*(np.mean(1.0/rank_a2b) + np.mean(1.0/rank_b2a))),
    }
    for k in topk:
        k = int(k)
        out[f"recall@{k}_a2b"] = float(np.mean(rank_a2b <= k))
        out[f"recall@{k}_b2a"] = float(np.mean(rank_b2a <= k))
        out[f"recall@{k}_mean"] = float(0.5*(out[f"recall@{k}_a2b"] + out[f"recall@{k}_b2a"]))
    return out


# -------------------------
# Label transfer (NaN-safe + shape-safe)
# -------------------------
def knn_label_transfer(Z_ref, y_ref, Z_query, idx_query=None, *, k=15, metric="euclidean"):
    """
    Returns:
      y_pred_used: predicted labels for USED query rows (finite)
      idx_used: indices into the ORIGINAL query (global indices if idx_query provided)
    """
    Z_ref = np.asarray(Z_ref)
    Z_query = np.asarray(Z_query)
    y_ref = np.asarray(y_ref).astype(str)

    if Z_ref.ndim != 2 or Z_query.ndim != 2:
        raise ValueError(f"Z_ref/Z_query must be 2D. Got {Z_ref.shape}, {Z_query.shape}")

    if idx_query is None:
        idx_query = np.arange(Z_query.shape[0], dtype=int)
    idx_query = np.asarray(idx_query, dtype=int)

    # filter finite rows
    ref_mask = _finite_row_mask(Z_ref)
    qry_mask = _finite_row_mask(Z_query[idx_query])

    if ref_mask.sum() == 0 or qry_mask.sum() == 0:
        return np.array([], dtype=str), np.array([], dtype=int)

    Zr = Z_ref[ref_mask]
    yr = y_ref[ref_mask]

    idx_used = idx_query[qry_mask]
    Zq = Z_query[idx_used]

    k_eff = min(int(k), Zr.shape[0])
    nbrs = NearestNeighbors(n_neighbors=k_eff, metric=metric).fit(Zr)
    nn = nbrs.kneighbors(Zq, return_distance=False)

    y_pred = []
    for neigh in nn:
        vals, counts = np.unique(yr[neigh], return_counts=True)
        y_pred.append(vals[np.argmax(counts)])
    return np.asarray(y_pred, dtype=str), idx_used


def label_transfer_metrics_split(Z_rna, Z_atac, labels, *, train_idx, test_idx, k=15):
    """
    Scores ONLY on the test cells that were actually predicted (finite query rows).
    """
    y = np.asarray(labels).astype(str)
    tr = np.asarray(train_idx, dtype=int)
    te = np.asarray(test_idx, dtype=int)

    # Train sets must be finite in the reference space too (we pass Z_*[tr] into knn_label_transfer,
    # which internally filters finite rows, so OK)

    # RNA -> ATAC
    yhat_r2a, te_used_r2a = knn_label_transfer(Z_rna[tr], y[tr], Z_atac, idx_query=te, k=k)
    if len(te_used_r2a):
        acc_r2a = accuracy_score(y[te_used_r2a], yhat_r2a)
        f1_r2a  = f1_score(y[te_used_r2a], yhat_r2a, average="macro")
    else:
        acc_r2a = np.nan
        f1_r2a = np.nan

    # ATAC -> RNA
    yhat_a2r, te_used_a2r = knn_label_transfer(Z_atac[tr], y[tr], Z_rna, idx_query=te, k=k)
    if len(te_used_a2r):
        acc_a2r = accuracy_score(y[te_used_a2r], yhat_a2r)
        f1_a2r  = f1_score(y[te_used_a2r], yhat_a2r, average="macro")
    else:
        acc_a2r = np.nan
        f1_a2r = np.nan

    acc_mean = np.nanmean([acc_r2a, acc_a2r])
    f1_mean  = np.nanmean([f1_r2a, f1_a2r])

    return {
        "acc_rna_to_atac": float(acc_r2a) if np.isfinite(acc_r2a) else np.nan,
        "macroF1_rna_to_atac": float(f1_r2a) if np.isfinite(f1_r2a) else np.nan,
        "acc_atac_to_rna": float(acc_a2r) if np.isfinite(acc_a2r) else np.nan,
        "macroF1_atac_to_rna": float(f1_a2r) if np.isfinite(f1_a2r) else np.nan,
        "acc_mean": float(acc_mean) if np.isfinite(acc_mean) else np.nan,
        "macroF1_mean": float(f1_mean) if np.isfinite(f1_mean) else np.nan,
    }


# -------------------------
# Mixing score (NaN-safe)
# -------------------------
def modality_mixing_score(Z_rna, Z_atac, k=20):
    Z_rna = np.asarray(Z_rna)
    Z_atac = np.asarray(Z_atac)
    assert Z_rna.shape[0] == Z_atac.shape[0], "paired n required"

    idx = np.arange(Z_rna.shape[0], dtype=int)
    Zr, Za, _ = _paired_finite_on_idx(Z_rna, Z_atac, idx)
    if Zr.shape[0] < 2:
        return np.nan

    Z = np.vstack([Zr, Za])
    modality = np.array(["rna"] * Zr.shape[0] + ["atac"] * Za.shape[0])

    k_eff = min(int(k) + 1, Z.shape[0])
    nbrs = NearestNeighbors(n_neighbors=k_eff).fit(Z)
    nn = nbrs.kneighbors(Z, return_distance=False)[:, 1:]

    frac_other = np.mean([np.mean(modality[nn[i]] != modality[i]) for i in range(Z.shape[0])])
    return float(frac_other)


# -------------------------
# Fused-only metrics (NaN-safe on TRAIN and TEST)
# -------------------------
def knn_predict_labels(Z_train, y_train, Z_query, *, k=15, metric="euclidean"):
    Z_train = np.asarray(Z_train)
    Z_query = np.asarray(Z_query)
    y_train = np.asarray(y_train).astype(str)

    if Z_train.shape[0] == 0 or Z_query.shape[0] == 0:
        return np.asarray([], dtype=str)

    k_eff = min(int(k), Z_train.shape[0])
    nbrs = NearestNeighbors(n_neighbors=k_eff, metric=metric).fit(Z_train)
    nn = nbrs.kneighbors(Z_query, return_distance=False)

    y_pred = []
    for neigh in nn:
        vals, counts = np.unique(y_train[neigh], return_counts=True)
        y_pred.append(vals[np.argmax(counts)])
    return np.asarray(y_pred, dtype=str)


def fused_knn_metrics(Z_fused, labels, *, splits, k=15):
    Zf = np.asarray(Z_fused)
    y  = np.asarray(labels).astype(str)
    tr = np.asarray(splits["train"], dtype=int)
    te = np.asarray(splits["test"], dtype=int)

    # finite-only TRAIN and TEST
    Ztr, tr_used = _subset_finite_rows(Zf, tr)
    Zte, te_used = _subset_finite_rows(Zf, te)

    if len(tr_used) == 0 or len(te_used) == 0:
        return {
            "fused_knn_acc_test": np.nan,
            "fused_knn_macroF1_test": np.nan,
            "fused_knn_balanced_acc_test": np.nan,
        }

    yhat = knn_predict_labels(Ztr, y[tr_used], Zte, k=k)
    if yhat.shape[0] != len(te_used):
        # hard safety: never allow inconsistent sample lengths
        return {
            "fused_knn_acc_test": np.nan,
            "fused_knn_macroF1_test": np.nan,
            "fused_knn_balanced_acc_test": np.nan,
        }

    y_true = y[te_used]
    return {
        "fused_knn_acc_test": float(accuracy_score(y_true, yhat)),
        "fused_knn_macroF1_test": float(f1_score(y_true, yhat, average="macro")),
        "fused_knn_balanced_acc_test": float(balanced_accuracy_score(y_true, yhat)),
    }


def fused_silhouette_by_label(Z_fused, labels, *, splits):
    Zf = np.asarray(Z_fused)
    y  = np.asarray(labels).astype(str)
    te = np.asarray(splits["test"], dtype=int)

    Zte, te_used = _subset_finite_rows(Zf, te)
    if len(te_used) < 3:
        return {"fused_silhouette_test": np.nan}

    y_te = y[te_used]
    n_unique = len(np.unique(y_te))
    if n_unique < 2 or n_unique >= len(y_te):
        return {"fused_silhouette_test": np.nan}

    try:
        s = silhouette_score(Zte, y_te, metric="euclidean")
    except Exception:
        s = np.nan
    return {"fused_silhouette_test": float(s) if np.isfinite(s) else np.nan}


def knn_label_purity(Z, labels, *, splits, k=30):
    Z = np.asarray(Z)
    y = np.asarray(labels).astype(str)
    te = np.asarray(splits["test"], dtype=int)

    Zte, te_used = _subset_finite_rows(Z, te)
    if len(te_used) < 3:
        return {"fused_knn_label_purity_test": np.nan}

    k_eff = min(int(k) + 1, Zte.shape[0])
    nbrs = NearestNeighbors(n_neighbors=k_eff).fit(Zte)
    nn = nbrs.kneighbors(Zte, return_distance=False)[:, 1:]
    yt = y[te_used]
    purity = np.mean([(yt[nn[i]] == yt[i]).mean() for i in range(len(yt))])
    return {"fused_knn_label_purity_test": float(purity)}


def fused_ari_nmi_kmeans(Z_fused, labels, *, splits, seed=0):
    Z = np.asarray(Z_fused)
    y = np.asarray(labels).astype(str)
    te = np.asarray(splits["test"], dtype=int)

    Zte, te_used = _subset_finite_rows(Z, te)
    if len(te_used) < 3:
        return {"fused_kmeans_ari_test": np.nan, "fused_kmeans_nmi_test": np.nan}

    yt = y[te_used]
    K = len(np.unique(yt))
    if K < 2:
        return {"fused_kmeans_ari_test": np.nan, "fused_kmeans_nmi_test": np.nan}

    #cl = KMeans(n_clusters=K, n_init="auto", random_state=int(seed)).fit_predict(Zte)
    cl = KMeans(n_clusters=K, n_init="auto", random_state=_seed_int(seed)).fit_predict(Zte)
    
    return {
        "fused_kmeans_ari_test": float(adjusted_rand_score(yt, cl)),
        "fused_kmeans_nmi_test": float(normalized_mutual_info_score(yt, cl)),
    }


# -------------------------
# Main evaluation (NaN-safe everywhere)
# -------------------------
def _has_pair(Zr, Za):
    return (
        Zr is not None and Za is not None
        and np.asarray(Zr).ndim == 2 and np.asarray(Za).ndim == 2
        and np.asarray(Zr).shape[0] == np.asarray(Za).shape[0]
    )

def evaluate_embeddings_split(
    out,
    labels,
    *,
    splits,
    foscttm_sub=3000,
    k_mix=20,
    k_lt=15,
    k_fused=15,
    seed=0,
):

    seed = _seed_int(seed)

    tr = np.asarray(splits["train"], dtype=int)
    te = np.asarray(splits["test"], dtype=int)
    y  = np.asarray(labels).astype(str)

    Zr = out.get("Z_rna", None)
    Za = out.get("Z_atac", None)
    Zf = out.get("Z_fused", None)

    row = {}

    # paired metrics on TEST, only where BOTH are finite
    if _has_pair(Zr, Za):
        Zr_te, Za_te, _ = _paired_finite_on_idx(Zr, Za, te)
        if Zr_te.shape[0] >= 2:
            fos = foscttm_values(Zr_te, Za_te, subsample=foscttm_sub, seed=seed, topk=(1, 10))
            row["FOSCTTM_mean_test"] = fos["mean"]
            row["FOSCTTM_sem_test"]  = fos["sem"]
            row["FOSCTTM_mrr_mean_test"] = fos["mrr_mean"]
            row["FOSCTTM_recall@1_mean_test"]  = fos.get("recall@1_mean", np.nan)
            row["FOSCTTM_recall@10_mean_test"] = fos.get("recall@10_mean", np.nan)

            lt = label_transfer_metrics_split(Zr, Za, y, train_idx=tr, test_idx=te, k=k_lt)
            row["label_transfer_acc_rna_to_atac_test"]     = lt["acc_rna_to_atac"]
            row["label_transfer_macroF1_rna_to_atac_test"] = lt["macroF1_rna_to_atac"]
            row["label_transfer_acc_atac_to_rna_test"]     = lt["acc_atac_to_rna"]
            row["label_transfer_macroF1_atac_to_rna_test"] = lt["macroF1_atac_to_rna"]
            row["label_transfer_acc_mean_test"]            = lt["acc_mean"]
            row["label_transfer_macroF1_mean_test"]        = lt["macroF1_mean"]

            row["mixing_score_test"] = modality_mixing_score(Zr_te, Za_te, k=k_mix)
        else:
            row.update({
                "FOSCTTM_mean_test": np.nan, "FOSCTTM_sem_test": np.nan, "FOSCTTM_mrr_mean_test": np.nan,
                "FOSCTTM_recall@1_mean_test": np.nan, "FOSCTTM_recall@10_mean_test": np.nan,
                "label_transfer_acc_rna_to_atac_test": np.nan, "label_transfer_macroF1_rna_to_atac_test": np.nan,
                "label_transfer_acc_atac_to_rna_test": np.nan, "label_transfer_macroF1_atac_to_rna_test": np.nan,
                "label_transfer_acc_mean_test": np.nan, "label_transfer_macroF1_mean_test": np.nan,
                "mixing_score_test": np.nan,
            })
    else:
        row.update({
            "FOSCTTM_mean_test": np.nan, "FOSCTTM_sem_test": np.nan, "FOSCTTM_mrr_mean_test": np.nan,
            "FOSCTTM_recall@1_mean_test": np.nan, "FOSCTTM_recall@10_mean_test": np.nan,
            "label_transfer_acc_rna_to_atac_test": np.nan, "label_transfer_macroF1_rna_to_atac_test": np.nan,
            "label_transfer_acc_atac_to_rna_test": np.nan, "label_transfer_macroF1_atac_to_rna_test": np.nan,
            "label_transfer_acc_mean_test": np.nan, "label_transfer_macroF1_mean_test": np.nan,
            "mixing_score_test": np.nan,
        })

    # fused-only metrics (NaN-safe on TRAIN and TEST)
    if Zf is not None and np.asarray(Zf).ndim == 2 and np.asarray(Zf).size > 0:
        row.update(fused_knn_metrics(Zf, y, splits=splits, k=k_fused))
        row.update(fused_silhouette_by_label(Zf, y, splits=splits))
        row.update(knn_label_purity(Zf, y, splits=splits, k=30))
        row.update(fused_ari_nmi_kmeans(Zf, y, splits=splits, seed=seed))
    else:
        row.update({
            "fused_knn_acc_test": np.nan,
            "fused_knn_macroF1_test": np.nan,
            "fused_knn_balanced_acc_test": np.nan,
            "fused_silhouette_test": np.nan,
            "fused_knn_label_purity_test": np.nan,
            "fused_kmeans_ari_test": np.nan,
            "fused_kmeans_nmi_test": np.nan,
        })

    return row


# %% [markdown]
### Specify the methods/their inputs/where to save their results

# %% [code]
import numpy as np
import scipy.sparse as sp

def _minmax(X):
    """Return (min, max, nnz, shape, dtype, frac_nonint, frac_neg, frac_naninf)."""
    if sp.issparse(X):
        data = X.data
        nnz = int(data.size)
        # sparse min/max need care for implicit zeros
        dmin = float(data.min()) if nnz else 0.0
        dmax = float(data.max()) if nnz else 0.0
        xmin = min(0.0, dmin)   # because implicit zeros exist
        xmax = max(0.0, dmax)
        arr = data
    else:
        arr = np.asarray(X).ravel()
        nnz = int(np.count_nonzero(arr))
        xmin = float(np.nanmin(arr)) if arr.size else np.nan
        xmax = float(np.nanmax(arr)) if arr.size else np.nan

    arr = np.asarray(arr)
    finite = np.isfinite(arr)
    frac_naninf = float(1.0 - (finite.mean() if arr.size else 1.0))
    arr_f = arr[finite] if arr.size else arr

    if arr_f.size:
        frac_nonint = float(np.mean(np.abs(arr_f - np.round(arr_f)) > 1e-6))
        frac_neg = float(np.mean(arr_f < 0))
    else:
        frac_nonint = 0.0
        frac_neg = 0.0

    return xmin, xmax, nnz, tuple(getattr(X, "shape", (len(arr),))), str(getattr(X, "dtype", arr.dtype)), frac_nonint, frac_neg, frac_naninf


def _layer_or_X(adata, layer=None):
    if layer is None or layer == "X":
        return adata.X, "X"
    if hasattr(adata, "layers") and layer in adata.layers:
        return adata.layers[layer], f"layers['{layer}']"
    if hasattr(adata, "obsm") and layer in adata.obsm:
        return adata.obsm[layer], f"obsm['{layer}']"
    raise KeyError(f"{adata!r} has no layer/obsm key '{layer}'")


def describe_adata(adata, name, *, layers=("X", "counts")):
    print(f"\n=== {name} ===")
    print(adata)
    # basic obs/var info
    try:
        print(f"  n_obs={adata.n_obs} n_vars={adata.n_vars}")
    except Exception:
        pass

    for layer in layers:
        try:
            X, src = _layer_or_X(adata, None if layer == "X" else layer)
        except KeyError:
            continue
        xmin, xmax, nnz, shape, dtype, frac_nonint, frac_neg, frac_naninf = _minmax(X)
        print(f"  {src}: shape={shape} dtype={dtype} nnz={nnz}")
        print(f"    min={xmin:.4g}  max={xmax:.4g}  frac_nonint={frac_nonint:.4f}  frac_neg={frac_neg:.4f}  frac_naninf={frac_naninf:.4f}")


# %% [code]
# ---------------------------------------------------------------------------
# Run the data checks to make sure intended counts going to each method etc.
# ---------------------------------------------------------------------------

describe_adata(rna_univi, "rna_univi", layers=("X", "counts"))
describe_adata(atac_univi, "atac_univi", layers=("X", "counts"))

describe_adata(rna_counts_hvg, "rna_counts_hvg_multivi", layers=("X", "counts"))
describe_adata(atac_multivi, "atac_multivi", layers=("X", "counts"))

describe_adata(rna_log_hvg, "rna_log_hvg_multimap", layers=("X", "counts"))
describe_adata(atac_lsi, "atac_lsi_multimap", layers=("X", "counts"))

describe_adata(rna, "rna (raw input)", layers=("X", "counts"))
describe_adata(atac, "atac (raw input)", layers=("X", "counts"))

describe_adata(rna_counts_hvg, "rna_counts_hvg_deepcca", layers=("X", "counts"))
describe_adata(atac_counts_bin_hv, "atac_counts_bin_hv_deepcca", layers=("X", "counts"))

describe_adata(rna_counts_hvg, "rna_counts_hvg_scmomat", layers=("X", "counts"))
describe_adata(atac_counts_bin_hv, "rna_counts_hvg_scmomat", layers=("X", "counts"))

describe_adata(rna_counts_hvg, "rna_counts_hvg_scjoint", layers=("X", "counts"))
describe_adata(atac_for_scjoint, "atac_for_scjoint", layers=("X", "counts", "gene_activity_hvg"))

describe_adata(atac_peakvi, "atac_peakvi", layers=("X", "counts"))

describe_adata(rna_counts_hvg, "rna_counts_hvg_cobolt", layers=("X", "counts"))
describe_adata(atac_counts_bin_hv, "atac_counts_bin_hv_cobolt", layers=("X", "counts"))


# %% [code]
'''
METHODS = {}

METHODS["univi"]    = lambda: run_univi(rna_univi, atac_univi, out_dir=WORK/"runs/univi", splits=splits, seed=RNG_SEED)
METHODS["multivi"]  = lambda: run_multivi(rna_counts_hvg, atac_multivi, out_dir=WORK/"runs/multivi", splits=splits, seed=RNG_SEED, n_latent=30, max_epochs=200, 
                                          patience=50, rna_layer="counts", atac_layer=None) #atac_layer="counts")
METHODS["multimap"] = lambda: run_multimap(rna_log_hvg, atac_lsi, out_dir=WORK/"runs/multimap", splits=splits, seed=RNG_SEED)
METHODS["scglue"]   = lambda: run_scglue_fair(rna_raw=rna, atac_raw=atac, gtf_path=GTF, out_dir=WORK/"runs/scglue", splits=splits, seed=RNG_SEED, 
                                              latent_dim=30, max_epochs=200, val_split=0.1, fuse="mean", verbose=False, debug=True)
METHODS["deepcca"]  = lambda: run_deepcca(rna_log_hvg, atac_lsi, out_dir=WORK/"runs/deepcca", splits=splits, seed=RNG_SEED, latent_dim=30, reg=1e-3)
METHODS["scmomat"]  = lambda: run_scmomat_docstyle(rna=rna_counts_hvg, atac=atac_counts_bin_hv, out_dir=WORK/"runs/scmomat", batch_key=None, 
                                                   layers_by_mod={"rna": "counts", "atac": "counts"}, K=30, T=4000, lr=1e-2, lamb=0.001, seed=RNG_SEED, 
                                                   interval=1000, verbose=True)
METHODS["scjoint"]  = lambda: run_scjoint_split_aware(rna_counts_hvg, atac_for_scjoint, splits=splits, labels_key=LABEL_KEY, 
                                                      atac_gene_activity_layer="gene_activity_hvg", 
                                                      scjoint_repo="/home/groups/precepts/ashforda/external_github_packages/scJoint", 
                                                      out_dir=WORK / "runs" / "scjoint", seed=RNG_SEED, latent_dim=30, batch_size=256, gpu=0, 
                                                      allow_transductive_fallback=True, fill_missing="nan")
METHODS["peakvi"]   = lambda: run_peakvi_fair(atac_peakvi, out_dir=WORK/"runs/peakvi", splits=splits, seed=RNG_SEED, n_latent=30, max_epochs=200, patience=50, 
                                              layer=None)
METHODS["cobolt"]   = lambda: run_cobolt_working(rna_counts_hvg, atac_counts_bin_hv, splits=None, rna_key="counts", atac_key=None, n_latent=30, max_epochs=200,
                                                 batch_size=256, device="cuda", seed=RNG_SEED, rna_cp10k_log1p=False, lr=5e-3, verbose=True)
'''

# %% [code]
# ============================================================
# Method registry (callables that return ensure_flags(...) dicts)
# ============================================================
METHODS = {}

# Core
METHODS["univi"] = lambda: run_univi(
    rna_univi,
    atac_univi,
    out_dir=WORK / "runs" / "univi",
    splits=splits,
    seed=RNG_SEED,
)

# MultiVI (split-aware; layers explicit)
METHODS["multivi"] = lambda: run_multivi(
    rna_counts_hvg,
    atac_multivi,
    out_dir=WORK / "runs" / "multivi",
    splits=splits,
    seed=RNG_SEED,
    n_latent=30,
    max_epochs=200,
    patience=50,
    rna_layer="counts",
    atac_layer=None,  # set to "counts" ONLY if atac_multivi.layers["counts"] exists
)

# MultiMAP (transductive; make sure run_multimap densifies sparse + train-fit scales)
METHODS["multimap"] = lambda: run_multimap(
    rna_log_hvg,
    atac_lsi,
    out_dir=WORK / "runs" / "multimap",
    splits=splits,
    seed=RNG_SEED,
    latent_dim=30,
)

# scGLUE (split-aware)
METHODS["scglue"] = lambda: run_scglue_fair(
    rna_raw=rna,
    atac_raw=atac,
    gtf_path=GTF,
    out_dir=WORK / "runs" / "scglue",
    splits=splits,
    seed=RNG_SEED,
    latent_dim=30,
    max_epochs=200,
    val_split=0.1,
    fuse="mean",
    verbose=False,
    debug=False,  # flip to True only when debugging; can change behavior in some implementations
)

# DeepCCA (split-aware)
METHODS["deepcca"] = lambda: run_deepcca(
    rna_log_hvg,
    atac_lsi,
    out_dir=WORK / "runs" / "deepcca",
    splits=splits,
    seed=RNG_SEED,
    latent_dim=30,
)

# scMoMaT (doc-style runner; uses layers_by_mod)
METHODS["scmomat"] = lambda: run_scmomat_docstyle(
    rna=rna_counts_hvg,
    atac=atac_counts_bin_hv,
    out_dir=WORK / "runs" / "scmomat",
    batch_key=None,
    layers_by_mod={"rna": "counts", "atac": "counts"},
    K=30,
    T=4000,
    lr=1e-2,
    lamb=0.001,
    seed=RNG_SEED,
    interval=1000,
    verbose=True,
)

# scJoint (split-aware; beware fill_missing="nan" if anything downstream can't handle NaNs)
METHODS["scjoint"] = lambda: run_scjoint_split_aware(
    rna_counts_hvg,
    atac_for_scjoint,
    splits=splits,
    labels_key=LABEL_KEY,
    atac_gene_activity_layer="gene_activity_hvg",
    scjoint_repo="/home/groups/precepts/ashforda/external_github_packages/scJoint",
    out_dir=WORK / "runs" / "scjoint",
    seed=RNG_SEED,
    latent_dim=30,
    batch_size=256,
    gpu=0,
    allow_transductive_fallback=True,
    fill_missing="nan",
)

# PeakVI (ATAC-only baseline)
METHODS["peakvi"] = lambda: run_peakvi_fair(
    atac_peakvi,
    out_dir=WORK / "runs" / "peakvi",
    splits=splits,
    seed=RNG_SEED,
    n_latent=30,
    max_epochs=200,
    patience=50,
    layer=None,
)

# CoBOLT (Can't pass splits.. ; CoBOLT runner is effectively transductive, mark it inside ensure_flags)
METHODS["cobolt"] = lambda: run_cobolt_working(
    rna_counts_hvg,
    atac_counts_bin_hv,
    splits=None,
    rna_key="counts",
    atac_key=None,
    n_latent=30,
    max_epochs=200,
    batch_size=256,
    device="cuda",
    seed=RNG_SEED,
    rna_cp10k_log1p=False,
    lr=5e-3,
    verbose=True,
)


# %% [markdown]
### Run all methods/save results/calculate evaluations

# %% [code]
# -----------------------------
# Make it shush (warnings + logs + tqdm)
# Put this at the very top of the notebook.
# -----------------------------
import os, warnings, logging

# 1) Warnings (target the annoying ones)
warnings.filterwarnings("ignore", message="The pynvml package is deprecated.*", category=FutureWarning)
warnings.filterwarnings("ignore", message="The argument 'device' of Tensor\\.pin_memory\\(\\) is deprecated.*", category=DeprecationWarning)
warnings.filterwarnings("ignore", message="The argument 'device' of Tensor\\.is_pinned\\(\\) is deprecated.*", category=DeprecationWarning)
warnings.filterwarnings("ignore", category=ResourceWarning)   # subprocess still running
# If you're truly done with warnings:
warnings.filterwarnings("ignore")

# 2) Logging (scglue / ignite / lightning / scvi etc.)
for name in [
    "scglue", "ignite", "lightning", "pytorch_lightning", "scvi",
    "muon", "anndata", "scanpy", "matplotlib"
]:
    logging.getLogger(name).setLevel(logging.ERROR)

# also quiet the root logger
logging.getLogger().setLevel(logging.ERROR)

# 3) tqdm progress bars (best-effort global disable)
os.environ["TQDM_DISABLE"] = "1"
os.environ["TOKENIZERS_PARALLELISM"] = "false"  # also reduces random HF noise if it appears


# %% [code]
import json
import numpy as np
import pandas as pd

labels_all = rna.obs[LABEL_KEY].astype(str).to_numpy()

def extract_flags(out):
    ej = (out or {}).get("extra_json", {}) or {}
    return {
        "transductive": bool(ej.get("transductive", True)),   # conservative default
        "uses_labels": bool(ej.get("uses_labels", False)),    # conservative default
    }

rows = []
all_embeddings = {}

for method, fn in METHODS.items():
    print(f"\n=== Running {method} ===")
    t0 = now()

    try:
        out = fn()
        out = ensure_flags(out)  # must set out["extra_json"]["transductive"/"uses_labels"]
        fit_seconds = float(out.get("fit_seconds", now() - t0))

        metrics = evaluate_embeddings_split(
            out,
            labels_all,
            splits=splits,
            foscttm_sub=FOSCTTM_SUBSAMPLE_N,
            k_mix=K_MIX,
            k_lt=K_LT,
            k_fused=K_LT,
            seed=RNG_SEED,
        )

        flags = extract_flags(out)

        row = {
            "method": method,
            "failed": False,
            "fit_seconds": fit_seconds,
            **flags,
            **metrics,
            # always store as JSON string for consistency
            "extra_json": json.dumps(out.get("extra_json", {}), default=str),
        }

        rows.append(row)
        all_embeddings[method] = out
        print(pd.Series(row))

    except Exception as e:
        fit_seconds = float(now() - t0)
        row = {
            "method": method,
            "failed": True,
            "error": repr(e),
            "fit_seconds": fit_seconds,
            # keep schema stable
            "transductive": True,
            "uses_labels": False,
            "extra_json": json.dumps({"transductive": True, "uses_labels": False}, default=str),
        }
        rows.append(row)
        print(f"[{method}] FAILED:", repr(e))

results = pd.DataFrame(rows)

results


# %% [markdown]
### 2) Save results + embeddings

# %% [code]
def _safe_np(x):
    if x is None:
        return np.zeros((0, 0), dtype=np.float32)
    x = np.asarray(x)
    return x

results_path = WORK / "results_multiome.csv"
results.to_csv(results_path, index=False)
print("Saved:", results_path)

emb_dir = WORK / "embeddings_multiome"
emb_dir.mkdir(parents=True, exist_ok=True)

for name, emb in all_embeddings.items():
    path = emb_dir / f"{name}.npz"
    save_npz(
        path,
        Z_rna=_safe_np(emb.get("Z_rna")),
        Z_atac=_safe_np(emb.get("Z_atac")),
        Z_fused=_safe_np(emb.get("Z_fused")),
    )
    print("Saved:", path)


# %% [markdown]
### All method comparison plots

# %% [code]
FIGDIR = WORK / "figures_multiome"
FIGDIR.mkdir(parents=True, exist_ok=True)

print("Saving figures to:", FIGDIR)


# %% [code]
import numpy as np
import anndata as ad
import scanpy as sc
import matplotlib.pyplot as plt
from pathlib import Path

def compute_umap_coords(Z, *, n_neighbors=15, min_dist=0.5, seed=0, metric="euclidean"):
    """
    Compute UMAP coordinates from an embedding Z (n,d). Returns (n,2).
    """
    Z = np.asarray(Z, dtype=np.float32)
    tmp = ad.AnnData(X=np.zeros((Z.shape[0], 1), dtype=np.float32))
    tmp.obsm["X_emb"] = Z

    sc.pp.neighbors(tmp, use_rep="X_emb", n_neighbors=int(n_neighbors), metric=metric)
    sc.tl.umap(tmp, min_dist=float(min_dist), random_state=int(seed))
    return np.asarray(tmp.obsm["X_umap"], dtype=np.float32)


# %% [code]
def plot_modality_overlay_umap(
    Z_rna, Z_atac,
    *,
    title="Modality overlay UMAP",
    n_neighbors=15,
    min_dist=0.5,
    seed=0,
    s=3,
    alpha=0.5,
    savepath=None,
):
    """
    UMAP on stacked [Z_rna; Z_atac], then scatter plot colored by modality.
    Uses matplotlib defaults (no explicit colors).
    """
    Z_rna = np.asarray(Z_rna)
    Z_atac = np.asarray(Z_atac)
    assert Z_rna.shape[0] == Z_atac.shape[0], "RNA/ATAC must have same n (paired)."

    Z = np.vstack([Z_rna, Z_atac])
    um = compute_umap_coords(Z, n_neighbors=n_neighbors, min_dist=min_dist, seed=seed)

    n = Z_rna.shape[0]
    um_rna = um[:n]
    um_atac = um[n:]

    plt.figure(figsize=(6.5, 5.5))
    plt.scatter(um_rna[:, 0], um_rna[:, 1], s=s, alpha=alpha, label="RNA")
    plt.scatter(um_atac[:, 0], um_atac[:, 1], s=s, alpha=alpha, label="ATAC")
    plt.title(title)
    plt.xlabel("UMAP1")
    plt.ylabel("UMAP2")
    plt.legend(markerscale=3)
    plt.tight_layout()

    if savepath is not None:
        savepath = Path(savepath)
        savepath.parent.mkdir(parents=True, exist_ok=True)
        plt.savefig(savepath, dpi=250)
    plt.show()

    return {"umap_stacked": um}


# %% [code]
def foscttm_values_from_pairwise(Z_rna, Z_atac, *, subsample=3000, seed=0, metric="euclidean_sq"):
    """
    Returns per-cell FOSCTTM values for RNA->ATAC and ATAC->RNA on a subsample.
    FOSCTTM(i) = fraction of "wrong" matches closer than the true match.
    """
    Z_rna = np.asarray(Z_rna)
    Z_atac = np.asarray(Z_atac)
    n = Z_rna.shape[0]
    assert n == Z_atac.shape[0], "RNA/ATAC must have same n (paired)."

    rng = np.random.default_rng(seed)
    m = min(int(subsample), n)
    idx = rng.choice(n, size=m, replace=False)

    A = Z_rna[idx]
    B = Z_atac[idx]

    # Pairwise distances (m x m). Use squared Euclidean by default (fast + stable).
    # D[i,j] = dist(A[i], B[j])
    if metric == "euclidean_sq":
        # (a-b)^2 = a^2 + b^2 - 2ab
        A2 = np.sum(A * A, axis=1, keepdims=True)          # (m,1)
        B2 = np.sum(B * B, axis=1, keepdims=True).T        # (1,m)
        D = A2 + B2 - 2.0 * (A @ B.T)
        D = np.maximum(D, 0.0)
    elif metric == "euclidean":
        A2 = np.sum(A * A, axis=1, keepdims=True)
        B2 = np.sum(B * B, axis=1, keepdims=True).T
        D = np.sqrt(np.maximum(A2 + B2 - 2.0 * (A @ B.T), 0.0))
    else:
        raise ValueError(f"Unknown metric: {metric}")

    # True match is diagonal: i matches i
    diag = np.diag(D).copy()

    # RNA -> ATAC: for each i, fraction of j where D[i,j] < D[i,i]
    # subtract 1 if you count itself (but strict < means diag not counted anyway)
    fos_rna_to_atac = (D < diag[:, None]).sum(axis=1) / (m - 1)

    # ATAC -> RNA is the same using transpose: D[j,i] comparisons
    diag2 = diag  # same diagonal
    fos_atac_to_rna = (D.T < diag2[:, None]).sum(axis=1) / (m - 1)

    return {
        "idx": idx,
        "fos_rna_to_atac": np.asarray(fos_rna_to_atac, dtype=float),
        "fos_atac_to_rna": np.asarray(fos_atac_to_rna, dtype=float),
        "fos_mean_bidir": 0.5 * (fos_rna_to_atac + fos_atac_to_rna),
    }


# %% [code]
def plot_foscttm_distribution(fos_dict, *, title="FOSCTTM distribution", saveprefix=None):
    """
    fos_dict: output of foscttm_values_from_pairwise
    """
    v1 = fos_dict["fos_rna_to_atac"]
    v2 = fos_dict["fos_atac_to_rna"]
    vb = fos_dict["fos_mean_bidir"]

    # Histogram
    plt.figure(figsize=(7, 4.5))
    plt.hist(v1, bins=40, alpha=0.6, label="RNA→ATAC")
    plt.hist(v2, bins=40, alpha=0.6, label="ATAC→RNA")
    plt.hist(vb, bins=40, alpha=0.6, label="Mean (bidir)")
    plt.title(title)
    plt.xlabel("FOSCTTM (lower = better)")
    plt.ylabel("Count")
    plt.legend()
    plt.tight_layout()
    if saveprefix:
        plt.savefig(f"{saveprefix}_foscttm_hist.png", dpi=250)
    plt.show()

    # Violin + box overlay (simple)
    data = [v1, v2, vb]
    labels = ["RNA→ATAC", "ATAC→RNA", "Mean"]
    plt.figure(figsize=(7, 4.5))
    parts = plt.violinplot(data, showmeans=False, showmedians=True, showextrema=False)
    plt.boxplot(data, widths=0.2, showfliers=False)
    plt.xticks([1, 2, 3], labels)
    plt.title(title)
    plt.ylabel("FOSCTTM (lower = better)")
    plt.tight_layout()
    if saveprefix:
        plt.savefig(f"{saveprefix}_foscttm_violin.png", dpi=250)
    plt.show()


# %% [code]
import warnings
import numpy as np
import pandas as pd
import matplotlib.pyplot as plt


def plot_umap_from_embedding(
    Z,
    obs_df,
    color_key=None,
    *,
    title="",
    n_neighbors=15,
    min_dist=0.5,
    seed=0,
    s=4,
    alpha=0.8,
    legend=True,
    ax=None,
):
    """
    Compute UMAP from an embedding matrix Z (n_cells x d) and scatter-plot it.
    - obs_df: pandas DataFrame (e.g., adata.obs or adata.obs.iloc[idx])
    - color_key: column in obs_df to color by (categorical or continuous)
    Returns: dict with 'U' (umap coords), 'fig', 'ax'
    """
    import numpy as np
    import pandas as pd
    import matplotlib.pyplot as plt

    Z = np.asarray(Z)
    if Z.ndim != 2:
        raise ValueError(f"Z must be 2D (n_cells x d); got shape={Z.shape}")

    # UMAP
    try:
        import umap
        reducer = umap.UMAP(
            n_neighbors=int(n_neighbors),
            min_dist=float(min_dist),
            metric="euclidean",
            random_state=int(seed),
        )
        U = reducer.fit_transform(Z)
    except Exception as e:
        raise RuntimeError(
            "UMAP failed. Do you have `umap-learn` installed? "
            "Try: pip install umap-learn"
        ) from e

    if ax is None:
        fig, ax = plt.subplots(figsize=(6, 5))
    else:
        fig = ax.figure

    # Plot
    if color_key is None:
        ax.scatter(U[:, 0], U[:, 1], s=s, alpha=alpha)
    else:
        if color_key not in obs_df.columns:
            raise KeyError(f"color_key={color_key!r} not in obs_df.columns")

        y = obs_df[color_key].to_numpy()

        # Continuous vs categorical
        if np.issubdtype(pd.Series(y).dtype, np.number):
            sc = ax.scatter(U[:, 0], U[:, 1], s=s, alpha=alpha, c=y)
            fig.colorbar(sc, ax=ax, fraction=0.046, pad=0.04)
        else:
            y = pd.Series(y).astype(str).fillna("NA").to_numpy()
            cats = pd.unique(y)
            for c in cats:
                m = (y == c)
                ax.scatter(U[m, 0], U[m, 1], s=s, alpha=alpha, label=c)
            if legend:
                ax.legend(
                    loc="upper left",
                    bbox_to_anchor=(1.02, 1),
                    frameon=False,
                    fontsize=7,
                    markerscale=2.5,
                )

    ax.set_title(title)
    ax.set_xlabel("UMAP1")
    ax.set_ylabel("UMAP2")
    ax.set_xticks([])
    ax.set_yticks([])

    return {"U": U, "fig": fig, "ax": ax}


# %% [code]
def dashboard_save_all(
    all_embeddings,
    results_df=None,
    *,
    figdir=FIGDIR,
    foscttm_subsample=3000,
    seed=0,
    n_neighbors=15,
    min_dist=0.5,
    splits=None,
):
    figdir = Path(figdir)
    figdir.mkdir(parents=True, exist_ok=True)

    te = splits["test"] if splits is not None else None

    def _row_subset(Z, idx):
        if Z is None or idx is None:
            return Z
        # pandas DataFrame / Series
        if hasattr(Z, "iloc"):
            return Z.iloc[idx]
        # torch Tensor
        try:
            import torch
            if isinstance(Z, torch.Tensor):
                return Z[idx]
        except Exception:
            pass
        # numpy / list-like
        return np.asarray(Z)[idx]

    fos_by_method = {}

    for method, out in all_embeddings.items():
        Zr = out.get("Z_rna", None)
        Za = out.get("Z_atac", None)
        Zf = out.get("Z_fused", None)

        # Prefer modality overlay only if Zr/Za exist
        if Zr is not None and Za is not None:
            #Zr_plot = Zr[te] if te is not None else Zr
            #Za_plot = Za[te] if te is not None else Za
            Zr_plot = _row_subset(Zr, te)
            Za_plot = _row_subset(Za, te)
            
            overlay_path = figdir / f"{method}__modality_overlay_umap.png"
            plot_modality_overlay_umap(
                Zr_plot, Za_plot,
                title=f"{method}: modality overlay (RNA vs ATAC)" + (" [TEST]" if te is not None else ""),
                n_neighbors=n_neighbors,
                min_dist=min_dist,
                seed=seed,
                savepath=overlay_path,
            )

            fos = foscttm_values_from_pairwise(
                Zr_plot, Za_plot,
                subsample=foscttm_subsample,
                seed=seed,
            )
            fos_by_method[method] = fos["fos_mean_bidir"]

            fos_prefix = figdir / f"{method}"
            plot_foscttm_distribution(
                fos,
                title=f"{method}: FOSCTTM distribution" + (" [TEST]" if te is not None else ""),
                saveprefix=str(fos_prefix),
            )

        # If only fused exists, save a fused UMAP (no FOSCTTM, no overlay)
        elif Zf is not None:
            #Zf_plot = Zf[te] if te is not None else Zf
            Zf_plot = _row_subset(Zf, te)
            '''
            tmp = plot_umap_from_embedding(
                Zf_plot,
                rna.obs.iloc[te] if te is not None else rna.obs,
                LABEL_KEY,
                title=f"{method}: fused UMAP" + (" [TEST]" if te is not None else ""),
            )
            '''
            
            obs_plot = rna.obs.iloc[te] if te is not None else rna.obs
            tmp = plot_umap_from_embedding(
                Zf_plot,
                obs_plot,
                LABEL_KEY,
                title=f"{method}: fused UMAP" + (" [TEST]" if te is not None else ""),
            )
            
            plt.savefig(figdir / f"{method}__fused_umap.png", dpi=250)
            plt.show()

        else:
            print(f"[{method}] no embeddings found, skipping plots")

    # Combined violin across methods (only those with true paired foscttm)
    if len(fos_by_method) > 0:
        methods = list(fos_by_method.keys())
        data = [fos_by_method[m] for m in methods]

        plt.figure(figsize=(max(8, 0.6 * len(methods)), 5))
        plt.violinplot(data, showmeans=False, showmedians=True, showextrema=False)
        plt.boxplot(data, widths=0.2, showfliers=False)
        plt.xticks(range(1, len(methods) + 1), methods, rotation=45, ha="right")
        plt.title("FOSCTTM (bidir mean) distribution across methods")
        plt.ylabel("FOSCTTM (lower = better)")
        plt.tight_layout()
        outpath = figdir / "ALL__foscttm_violin.png"
        plt.savefig(outpath, dpi=250)
        plt.show()

    if results_df is not None:
        results_df.to_csv(figdir / "results_metrics.csv", index=False)
        with open(figdir / "results_metrics.json", "w") as f:
            json.dump(results_df.to_dict(orient="records"), f, indent=2)

    print("Saved figures to:", figdir)
    return {"figdir": str(figdir), "fos_by_method": fos_by_method}


# %% [code]
dash = dashboard_save_all(
    all_embeddings,
    results_df=results,
    figdir=FIGDIR,
    #foscttm_subsample=FOSCTTM_SUBSAMPLE_N,
    foscttm_subsample=3000,
    seed=RNG_SEED,
    splits=splits,
)
dash


# %% [code]
'''
# With random seed = 67
    method	failed	fit_seconds	transductive	uses_labels	FOSCTTM_mean_test	FOSCTTM_sem_test	FOSCTTM_mrr_mean_test	FOSCTTM_recall@1_mean_test	FOSCTTM_recall@10_mean_test	...	label_transfer_macroF1_mean_test	mixing_score_test	fused_knn_acc_test	fused_knn_macroF1_test	fused_knn_balanced_acc_test	fused_silhouette_test	fused_knn_label_purity_test	fused_kmeans_ari_test	fused_kmeans_nmi_test	extra_json
0	univi	False	172.071677	False	False	0.018010	0.000823	0.267465	0.135892	0.548755	...	0.874115	0.473278	0.954357	0.940906	0.934369	0.351260	0.837725	0.508302	0.760567	{"transductive": false, "uses_labels": false}
1	multivi	False	223.625776	False	False	0.052636	0.001936	0.098820	0.033195	0.227178	...	0.689088	0.490519	0.866183	0.732153	0.746310	0.392727	0.775622	0.486352	0.734174	{"transductive": false, "uses_labels": false, ...
2	multimap	False	46.769239	True	False	NaN	NaN	NaN	NaN	NaN	...	NaN	NaN	0.876556	0.782693	0.790407	0.215923	0.745574	0.400929	0.664981	{"transductive": true, "uses_labels": false}
3	scglue	False	1038.876083	False	False	0.036658	0.001968	0.194465	0.093361	0.415456	...	0.762499	0.491836	0.942946	0.920576	0.910559	0.246982	0.819398	0.534926	0.773158	{"transductive": false, "uses_labels": false}
4	deepcca	False	32.100153	False	False	0.487022	0.008665	0.009648	0.000519	0.015041	...	0.045115	0.174471	0.931535	0.840092	0.841512	0.199901	0.792911	0.518514	0.673216	{"transductive": false, "uses_labels": false, ...
5	scmomat	False	68.720901	True	False	NaN	NaN	NaN	NaN	NaN	...	NaN	NaN	0.725104	0.566274	0.555586	-0.015417	0.543396	0.261847	0.532138	{"transductive": true, "uses_labels": false, "...
6	scjoint	False	107.798547	True	True	0.071130	0.002187	0.097969	0.036307	0.217324	...	0.688012	0.452396	0.946058	0.875449	0.874119	0.352216	0.850657	0.822849	0.844262	{"transductive": true, "uses_labels": true, "n...
7	peakvi	False	113.293390	False	False	NaN	NaN	NaN	NaN	NaN	...	NaN	NaN	0.876556	0.818412	0.819604	0.221018	0.759647	0.453958	0.718532	{"transductive": false, "uses_labels": false, ...
8	cobolt	False	808.664924	True	False	NaN	NaN	NaN	NaN	NaN	...	NaN	NaN	0.918050	0.831186	0.818098	0.089372	0.710961	0.458295	0.704632	{"transductive": true, "uses_labels": false}
9 rows × 25 columns
'''


# %% [code]
from __future__ import annotations

from pathlib import Path
import numpy as np
import pandas as pd
import matplotlib.pyplot as plt


# ----------------------------
# Metric metadata (edit freely)
# ----------------------------

DEFAULT_METRICS = [
    # paired alignment
    ("FOSCTTM_mean_test", "FOSCTTM mean (↓)", "lower"),
    ("FOSCTTM_mrr_mean_test", "FOSCTTM MRR (↑)", "higher"),
    ("FOSCTTM_recall@1_mean_test", "FOSCTTM Recall@1 (↑)", "higher"),
    ("FOSCTTM_recall@10_mean_test", "FOSCTTM Recall@10 (↑)", "higher"),
    # label transfer / mixing
    ("label_transfer_acc_mean_test", "Label transfer acc mean (↑)", "higher"),
    ("label_transfer_macroF1_mean_test", "Label transfer macroF1 mean (↑)", "higher"),
    ("mixing_score_test", "Mixing score (↑)", "higher"),
    # fused quality
    ("fused_knn_acc_test", "Fused kNN acc (↑)", "higher"),
    ("fused_knn_macroF1_test", "Fused kNN macroF1 (↑)", "higher"),
    ("fused_knn_balanced_acc_test", "Fused kNN balanced acc (↑)", "higher"),
    ("fused_silhouette_test", "Fused silhouette (↑)", "higher"),
    ("fused_knn_label_purity_test", "Fused label purity (↑)", "higher"),
    ("fused_kmeans_ari_test", "Fused k-means ARI (↑)", "higher"),
    ("fused_kmeans_nmi_test", "Fused k-means NMI (↑)", "higher"),
    # runtime
    ("fit_seconds", "Fit seconds (↓)", "lower"),
]


# ----------------------------
# Helpers
# ----------------------------

def _ensure_df(results_df) -> pd.DataFrame:
    if results_df is None:
        raise ValueError("results_df is required")
    df = results_df.copy()
    if "method" not in df.columns:
        raise ValueError("results_df must contain a 'method' column.")
    # coerce metric columns to numeric where possible
    for c in df.columns:
        if c == "method":
            continue
        if df[c].dtype == object:
            # try numeric coercion; ignore failures
            df[c] = pd.to_numeric(df[c], errors="ignore")
    return df


def _ok_methods(df: pd.DataFrame, *, include_failed: bool = False) -> pd.DataFrame:
    if "failed" not in df.columns or include_failed:
        return df
    return df.loc[~df["failed"].astype(bool)].copy()


def _score_for_sort(values: np.ndarray, direction: str) -> np.ndarray:
    # used only for ordering methods
    if direction == "higher":
        return values
    if direction == "lower":
        return -values
    return values


def _method_order_for_metric(df: pd.DataFrame, metric: str, direction: str) -> list[str]:
    d = df[["method", metric]].dropna()
    if d.empty:
        return df["method"].tolist()
    v = d[metric].to_numpy()
    s = _score_for_sort(v, direction)
    order = d.iloc[np.argsort(-s)]["method"].tolist()
    # include methods that were NaN at the end
    rest = [m for m in df["method"].tolist() if m not in order]
    return order + rest


# ----------------------------
# 1) Scalar metric leaderboard plots
# ----------------------------

def plot_metric_leaderboard(
    results_df,
    metric: str,
    *,
    title: str | None = None,
    direction: str = "higher",   # "higher" or "lower"
    figdir: str | Path | None = None,
    filename: str | None = None,
    include_failed: bool = False,
    annotate: bool = True,
    logy: bool = False,
):
    """
    One number per method -> dot plot / lollipop.

    direction:
      - "higher": best at top
      - "lower": best at top
    """
    df = _ensure_df(results_df)
    df = _ok_methods(df, include_failed=include_failed)

    if metric not in df.columns:
        print(f"[plot_metric_leaderboard] missing metric: {metric}")
        return None

    d = df[["method", metric]].copy()
    d[metric] = pd.to_numeric(d[metric], errors="coerce")

    order = _method_order_for_metric(d, metric, direction)
    d["method"] = pd.Categorical(d["method"], categories=order, ordered=True)
    d = d.sort_values("method")

    y = d["method"].astype(str).tolist()
    x = d[metric].to_numpy()

    plt.figure(figsize=(8, max(4, 0.45 * len(y))))
    # lollipop: line from 0 to x, then dot
    for i, val in enumerate(x):
        if np.isfinite(val):
            plt.plot([0, val], [i, i], linewidth=1)
            plt.scatter([val], [i], s=40)

    plt.yticks(range(len(y)), y)
    plt.xlabel(metric)
    plt.title(title or f"{metric} across methods")
    if logy:
        # log y doesn't make sense; log x axis does
        plt.xscale("log")
        plt.xlabel(metric + " (log scale)")
    plt.grid(axis="x", alpha=0.2)
    plt.tight_layout()

    if annotate:
        for i, val in enumerate(x):
            if np.isfinite(val):
                plt.text(val, i, f" {val:.4g}", va="center", fontsize=9)

    outpath = None
    if figdir is not None:
        figdir = Path(figdir)
        figdir.mkdir(parents=True, exist_ok=True)
        outname = filename or f"ALL__leaderboard__{metric}.png"
        outpath = figdir / outname
        plt.savefig(outpath, dpi=250)
    plt.show()
    return outpath


def plot_all_scalar_leaderboards(
    results_df,
    *,
    metrics=DEFAULT_METRICS,
    figdir: str | Path,
    include_failed: bool = False,
):
    """
    Make one leaderboard plot per scalar metric.
    """
    for metric, pretty, direction in metrics:
        logx = (metric == "fit_seconds")
        plot_metric_leaderboard(
            results_df,
            metric,
            title=pretty,
            direction=direction,
            figdir=figdir,
            filename=f"ALL__leaderboard__{metric}.png",
            include_failed=include_failed,
            annotate=True,
            logy=logx,
        )


# ----------------------------
# 2) Distribution violins (per-method arrays)
# ----------------------------

def plot_distribution_violin(
    dist_by_method: dict[str, np.ndarray],
    *,
    title: str,
    ylabel: str,
    figdir: str | Path | None = None,
    filename: str | None = None,
    order: list[str] | None = None,
    show_box: bool = True,
    clip_quantiles: tuple[float, float] | None = None,  # e.g. (0.01, 0.99)
):
    """
    dist_by_method: {method: 1D array}
    """
    # filter empties / nan-only
    clean = {}
    for m, arr in dist_by_method.items():
        if arr is None:
            continue
        a = np.asarray(arr, dtype=np.float32).ravel()
        a = a[np.isfinite(a)]
        if a.size == 0:
            continue
        if clip_quantiles is not None:
            lo, hi = np.quantile(a, clip_quantiles)
            a = a[(a >= lo) & (a <= hi)]
        if a.size:
            clean[m] = a

    if not clean:
        print("[plot_distribution_violin] nothing to plot (all empty).")
        return None

    methods = list(clean.keys())
    if order is not None:
        methods = [m for m in order if m in clean] + [m for m in methods if m not in (order or [])]

    data = [clean[m] for m in methods]

    plt.figure(figsize=(max(8, 0.6 * len(methods)), 5))
    plt.violinplot(data, showmeans=False, showmedians=True, showextrema=False)
    if show_box:
        plt.boxplot(data, widths=0.2, showfliers=False)
    plt.xticks(range(1, len(methods) + 1), methods, rotation=45, ha="right")
    plt.title(title)
    plt.ylabel(ylabel)
    plt.tight_layout()

    outpath = None
    if figdir is not None:
        figdir = Path(figdir)
        figdir.mkdir(parents=True, exist_ok=True)
        outname = filename or (title.replace(" ", "_") + ".png")
        outpath = figdir / outname
        plt.savefig(outpath, dpi=250)
    plt.show()
    return outpath


# ----------------------------
# 3) Metric matrix view (heatmap-like, no seaborn)
# ----------------------------

def plot_metric_matrix(
    results_df,
    *,
    metrics: list[tuple[str, str, str]] = DEFAULT_METRICS,
    figdir: str | Path | None = None,
    filename: str = "ALL__metric_matrix.png",
    include_failed: bool = False,
    normalize: bool = True,
):
    """
    Heatmap-ish matrix:
      rows = methods
      cols = metrics
    normalize=True: per-metric min-max (inverts for "lower is better") so higher = better (0..1)
    """
    df = _ensure_df(results_df)
    df = _ok_methods(df, include_failed=include_failed)

    methods = df["method"].astype(str).tolist()
    cols = []
    pretty = []
    direction = []
    for m, p, d in metrics:
        if m in df.columns:
            cols.append(m)
            pretty.append(p)
            direction.append(d)

    if not cols:
        print("[plot_metric_matrix] none of the requested metrics exist in results_df.")
        return None

    M = df[cols].apply(pd.to_numeric, errors="coerce").to_numpy(dtype=np.float32)

    # normalize per column
    if normalize:
        M2 = M.copy()
        for j in range(M.shape[1]):
            col = M[:, j]
            ok = np.isfinite(col)
            if ok.sum() < 2:
                continue
            v = col[ok]
            mn, mx = float(v.min()), float(v.max())
            if mx <= mn + 1e-12:
                continue
            scaled = (col - mn) / (mx - mn)
            # invert if lower is better, so "higher score is better"
            if direction[j] == "lower":
                scaled = 1.0 - scaled
            M2[:, j] = scaled
        M = M2

    # plot with imshow
    plt.figure(figsize=(max(10, 0.75 * len(cols)), max(6, 0.45 * len(methods))))
    im = plt.imshow(M, aspect="auto", interpolation="nearest")
    plt.colorbar(im, fraction=0.03, pad=0.02, label=("normalized score (higher=better)" if normalize else "raw value"))
    plt.xticks(range(len(cols)), pretty, rotation=45, ha="right")
    plt.yticks(range(len(methods)), methods)
    plt.title("Metric matrix across methods")
    plt.tight_layout()

    outpath = None
    if figdir is not None:
        figdir = Path(figdir)
        figdir.mkdir(parents=True, exist_ok=True)
        outpath = figdir / filename
        plt.savefig(outpath, dpi=250)
    plt.show()
    return outpath


# ----------------------------
# 4) Wrapper: make all comparison plots
# ----------------------------

def save_all_method_comparison_plots(
    results_df,
    *,
    figdir: str | Path,
    fos_by_method: dict[str, np.ndarray] | None = None,
    include_failed: bool = False,
):
    """
    Produces:
      - leaderboards for each scalar metric
      - metric matrix heatmap-ish view
      - violin for FOSCTTM distribution (if provided)
    """
    figdir = Path(figdir)
    figdir.mkdir(parents=True, exist_ok=True)

    # scalar leaderboards
    plot_all_scalar_leaderboards(results_df, metrics=DEFAULT_METRICS, figdir=figdir, include_failed=include_failed)

    # matrix view
    plot_metric_matrix(results_df, metrics=DEFAULT_METRICS, figdir=figdir, include_failed=include_failed, normalize=True)

    # distribution violin(s)
    if fos_by_method is not None and len(fos_by_method) > 0:
        plot_distribution_violin(
            fos_by_method,
            title="FOSCTTM distribution across methods",
            ylabel="FOSCTTM (lower = better)",
            figdir=figdir,
            filename="ALL__foscttm_violin.png",
            show_box=True,
        )

    print("[comparison plots] saved to:", figdir)
    return {"figdir": str(figdir)}


# %% [code]
dash = dashboard_save_all(
    all_embeddings,
    results_df=results,
    figdir=FIGDIR,
    foscttm_subsample=3000,
    seed=RNG_SEED,
    splits=splits,
)

save_all_method_comparison_plots(
    results,
    figdir=FIGDIR,
    fos_by_method=dash["fos_by_method"],  # violin uses distributions
)


# %% [markdown]
### Run across several random seeds for cross-validation

# %% [code]
# ============================================================
# Seed-sweep robustness (Multiome-ready)
# - varies *splits* across seeds (shared across methods)
# - REBUILDS any preprocessing that is "fit on TRAIN" per seed:
#     * RNA HVGs
#     * ATAC TFIDF+SVD/LSI
#     * ATAC HV-peaks (for MultiVI / PeakVI style)
# - does NOT change any existing run_* functions:
#     it just updates globals (RNG_SEED, splits, shared inputs) that your METHODS thunks use
# ============================================================

import time, json
from pathlib import Path
import numpy as np
import pandas as pd

# -----------------------------
# 0) configure the sweep
# -----------------------------
SEEDS = list(range(5))
SWEEP_TAG = "cv_sweep_py_1-31-2026"
OUT_DIR = (Path(WORK) / "runs" / SWEEP_TAG) if "WORK" in globals() else (Path("./runs") / SWEEP_TAG)
OUT_DIR.mkdir(parents=True, exist_ok=True)

# split hyperparams (match your make_shared_splits signature)
SPLIT_KWS = dict(
    train_frac=0.8,
    val_frac=0.1,
)

# preprocessing hyperparams (match your build_shared_inputs)
PREP_KWS = dict(
    rna_counts_layer="counts",
    atac_counts_layer="counts",
    n_hvg=2000,
    target_sum=1e4,
    n_lsi=101,
    n_hvpeaks_multivi=4002,
)

# -----------------------------
# 1) build splits for a given seed
# -----------------------------
def make_splits_for_seed(seed: int):
    # uses your existing function
    return make_shared_splits(
        rna.n_obs,
        labels_all,
        seed=int(seed),
        **SPLIT_KWS,
    )

# -----------------------------
# 2) (optional but recommended) set RNG seeds too
# -----------------------------
def set_all_seeds(seed: int):
    np.random.seed(int(seed))
    try:
        import random
        random.seed(int(seed))
    except Exception:
        pass
    try:
        import torch
        torch.manual_seed(int(seed))
        torch.cuda.manual_seed_all(int(seed))
        torch.backends.cudnn.deterministic = True
        torch.backends.cudnn.benchmark = False
    except Exception:
        pass

# -----------------------------
# 3) metric extraction extended
# -----------------------------
import time, json
from pathlib import Path

import numpy as np
import pandas as pd

from sklearn.model_selection import StratifiedKFold, StratifiedShuffleSplit
from sklearn.cluster import KMeans
from sklearn.metrics import adjusted_rand_score, normalized_mutual_info_score

def _extract_metrics(out):
    if out is None:
        return {}

    candidates = []
    if isinstance(out, dict):
        candidates.append(out)
        for k in ["summary", "metrics", "scores", "eval", "results", "extra_json"]:
            if k in out and isinstance(out[k], dict):
                candidates.append(out[k])

    merged = {}
    for c in candidates:
        merged.update(c)

    flat = {}
    for k, v in merged.items():
        if isinstance(v, (int, float, np.integer, np.floating)) and np.isfinite(v):
            flat[k] = float(v)
        elif isinstance(v, (list, tuple, np.ndarray)) and len(v) == 1:
            try:
                flat[k] = float(v[0])
            except Exception:
                pass
    return flat

def _maybe_add_ari_nmi(row_metrics, out, labels_test, *, k=None, seed=0):
    """
    If the method didn't already report ARI/NMI, compute from Z_fused on test.
    Expects out["Z_fused"] as (cells x d) with rownames aligned to full data,
    OR already subset to test in the runner.
    """
    if ("fused_kmeans_ari_test" in row_metrics) and ("fused_kmeans_nmi_test" in row_metrics):
        return row_metrics

    if not (isinstance(out, dict) and ("Z_fused" in out) and (out["Z_fused"] is not None)):
        return row_metrics

    Z = out["Z_fused"]
    # handle pandas / numpy
    if hasattr(Z, "values"):
        Z = Z.values
    Z = np.asarray(Z)

    if Z.ndim != 2 or Z.shape[0] != len(labels_test):
        # If runner returned full Z for all cells, you can adapt here,
        # but we can't guess mapping safely without names.
        return row_metrics

    if k is None:
        k = len(np.unique(labels_test))
    k = int(max(2, k))

    km = KMeans(n_clusters=k, n_init=10, random_state=int(seed))
    pred = km.fit_predict(Z)

    row_metrics["fused_kmeans_ari_test"] = float(adjusted_rand_score(labels_test, pred))
    row_metrics["fused_kmeans_nmi_test"] = float(normalized_mutual_info_score(labels_test, pred))
    return row_metrics

# -----------------------------
# fold builder: train/val/test indices
# -----------------------------
def make_folds(labels_all, n_splits=5, seed=0):
    labels_all = np.asarray(labels_all)
    skf = StratifiedKFold(n_splits=int(n_splits), shuffle=True, random_state=int(seed))

    for fold, (fit_idx, test_idx) in enumerate(skf.split(np.zeros_like(labels_all), labels_all)):
        # carve val out of fit_idx
        y_fit = labels_all[fit_idx]
        sss = StratifiedShuffleSplit(n_splits=1, test_size=0.1, random_state=int(seed + 10_000 + fold))
        tr_rel, va_rel = next(sss.split(np.zeros_like(y_fit), y_fit))
        train_idx = fit_idx[tr_rel]
        val_idx   = fit_idx[va_rel]

        yield fold, {
            "train": train_idx.tolist(),
            "val":   val_idx.tolist(),
            "test":  test_idx.tolist(),
            "unused": []
        }

# -----------------------------
# your "train-fit preprocessing" wrapper
# -----------------------------
def drop_first_lsi(atac_lsi: ad.AnnData) -> ad.AnnData:
    """
    Given atac_lsi with shape (cells, 101), return a new AnnData with shape (cells, 100)
    containing cols 1..100 (i.e. LSI 2-101 in 1-based terms).
    """
    X = np.asarray(atac_lsi.X)
    if X.ndim != 2:
        raise ValueError(f"Expected atac_lsi.X to be 2D, got {X.shape}")
    if X.shape[1] < 2:
        raise ValueError(f"Need at least 2 LSI dims to drop first, got {X.shape[1]}")

    X2 = X[:, 1:].astype(np.float32, copy=False)

    # keep obs, new var names
    out = ad.AnnData(
        X=X2,
        obs=atac_lsi.obs.copy(),
        var=pd.DataFrame(index=[f"LSI_{i}" for i in range(1, X2.shape[1] + 1)]),
    )
    return out
    
def rebuild_shared_inputs_for_split(seed: int, splits: dict):
    shared = build_shared_inputs(
        rna, atac, splits,
        seed=int(seed),
        **PREP_KWS,
    )

    globals()["shared"] = shared

    # ---- keep everything you already export ----
    globals()["rna_counts_hvg"]      = shared["rna_counts_hvg"]
    globals()["rna_log_hvg"]         = shared["rna_log_hvg"]
    globals()["atac_counts_bin"]     = shared["atac_counts_bin"]
    globals()["atac_counts_bin_hv"]  = shared["atac_counts_bin_hv"]

    # ---- ATAC LSI: compute 101, then make drop-first variant ----
    atac_lsi_101 = shared["atac_lsi"]            # shape: (cells, 101)
    atac_lsi_drop1 = drop_first_lsi(atac_lsi_101)  # shape: (cells, 100)

    # store both for debugging / reuse
    globals()["atac_lsi_101"] = atac_lsi_101
    globals()["atac_lsi_drop1"] = atac_lsi_drop1

    # IMPORTANT: choose which one each method family should see
    # - UniVI / MultiMAP: use drop-first (LSI 2-101)
    # - If any method truly wants the raw 101, point it to atac_lsi_101 instead
    globals()["atac_lsi"] = atac_lsi_drop1  # <-- default global LSI used by many thunks

    # method-specific "shared inputs" wiring (same as your original intent)
    globals()["rna_univi"]     = globals()["rna_log_hvg"]
    globals()["atac_univi"]    = globals()["atac_lsi_drop1"]     # <-- drop first

    globals()["rna_multimap"]  = globals()["rna_log_hvg"]
    globals()["atac_multimap"] = globals()["atac_lsi_drop1"]     # <-- drop first

    # methods that use peaks/binary matrices are unchanged
    globals()["atac_multivi"]  = globals()["atac_counts_bin_hv"]
    globals()["atac_peakvi"]   = globals()["atac_counts_bin_hv"]

    return shared
    

# -----------------------------
# main CV sweep
# -----------------------------
def run_cv_sweep(
    *,
    seeds,
    n_folds,
    out_dir,
    methods,
    labels_all,
    add_ari_nmi_if_missing=True,
):
    out_dir = Path(out_dir)
    out_dir.mkdir(parents=True, exist_ok=True)

    rows = []
    raw_out_path = out_dir / "raw_outputs.jsonl"

    with raw_out_path.open("w") as f_jsonl:
        for seed in seeds:
            # (optional) set seeds globally if you want determinism
            np.random.seed(int(seed))
            try:
                import random
                random.seed(int(seed))
            except Exception:
                pass
            try:
                import torch
                torch.manual_seed(int(seed))
                torch.cuda.manual_seed_all(int(seed))
            except Exception:
                pass

            for fold, splits in make_folds(labels_all, n_splits=n_folds, seed=seed):
                globals()["RNG_SEED"] = int(seed)
                globals()["splits"] = splits

                # IMPORTANT: rebuild train-fit preprocessing for THIS fold
                rebuild_shared_inputs_for_split(seed, splits)

                print(f"\n=== seed={seed} fold={fold} sizes ===", {k: len(v) for k, v in splits.items()})

                for method_name, thunk in methods.items():
                    t0 = time.time()
                    status, err, out = "ok", None, None

                    try:
                        out = thunk()
                    except Exception as e:
                        status = "fail"
                        err = repr(e)

                    dt = time.time() - t0
                    metrics = _extract_metrics(out)

                    # optional: compute ARI/NMI if not present AND runner returned Z_fused for test
                    if status == "ok" and add_ari_nmi_if_missing:
                        labels_test = np.asarray(labels_all)[splits["test"]]
                        metrics = _maybe_add_ari_nmi(metrics, out, labels_test, seed=seed)

                    row = {
                        "seed": int(seed),
                        "fold": int(fold),
                        "method": str(method_name),
                        "status": status,
                        "seconds": float(dt),
                        "error": err,
                        "n_train": int(len(splits["train"])),
                        "n_val": int(len(splits["val"])),
                        "n_test": int(len(splits["test"])),
                        **metrics,
                    }
                    rows.append(row)

                    f_jsonl.write(json.dumps({
                        "seed": int(seed),
                        "fold": int(fold),
                        "method": str(method_name),
                        "status": status,
                        "seconds": float(dt),
                        "error": err,
                        "metrics": metrics,
                        "out_keys": sorted(list(out.keys())) if isinstance(out, dict) else None,
                    }) + "\n")

    df = pd.DataFrame(rows)
    df.to_csv(out_dir / "cv_sweep_long.csv", index=False)

    ok = df[df["status"] == "ok"].copy()
    house = {"seed","fold","method","status","seconds","error","n_train","n_val","n_test"}
    metric_cols = [c for c in ok.columns if c not in house and pd.api.types.is_numeric_dtype(ok[c])]

    if metric_cols:
        # summarize over (seed,fold) replicates
        summ = (ok.groupby("method")[metric_cols]
                  .agg(["mean","std","median","count"])
                  .sort_values((metric_cols[0], "mean"), ascending=False))
        summ.to_csv(out_dir / "cv_sweep_summary.csv")
    else:
        summ = ok.groupby("method").size().to_frame("n_ok")
        summ.to_csv(out_dir / "cv_sweep_summary.csv")

    print("\nSaved:")
    print(" -", out_dir / "cv_sweep_long.csv")
    print(" -", out_dir / "cv_sweep_summary.csv")
    print(" -", raw_out_path)

    return df, summ


# %% [code]
def _get_Zmod(out):
    Z_mod = out.get("Z_atac", None)
    if Z_mod is None:
        Z_mod = out.get("Z_adt", None)
    return Z_mod

def _pair_test_metrics(out, labels_all, splits, *, metric="euclidean", recall_ks=(1,10,25,50,100),
                       max_pair_n=3000, seed=0, block=512):
    """
    Test-only paired retrieval metrics + permutation sanity control.
    Returns keys prefixed with "TEST/" and "PERM/".
    """
    M = {}
    if not isinstance(out, dict):
        return M

    Z_rna = out.get("Z_rna", None)
    Z_mod = _get_Zmod(out)
    if Z_rna is None or Z_mod is None:
        return M

    Z_rna = np.asarray(Z_rna, np.float32)
    Z_mod = np.asarray(Z_mod, np.float32)
    n = len(labels_all)

    if Z_rna.shape[0] != n or Z_mod.shape[0] != n:
        # can't trust pairing by index
        M["TEST/pair_shape_mismatch"] = 1.0
        M["TEST/n"] = float(min(Z_rna.shape[0], Z_mod.shape[0]))
        return M

    te = np.asarray(splits["test"], dtype=int)
    Z1 = Z_rna[te]
    Z2 = Z_mod[te]
    m = np.isfinite(Z1).all(1) & np.isfinite(Z2).all(1)
    Z1 = Z1[m]; Z2 = Z2[m]
    n_te = Z1.shape[0]
    M["TEST/n"] = float(n_te)

    if n_te < 10:
        return M

    # true test pairing metrics
    trueM = pair_ranking_metrics_exact(
        Z1, Z2, metric=metric, ks=recall_ks, max_n=max_pair_n, seed=seed, block=block
    )
    for k, v in trueM.items():
        M[f"TEST/{k}"] = v

    # permutation control (should ~chance)
    rng = np.random.default_rng(int(seed) + 999)
    perm = rng.permutation(n_te)
    shufM = pair_ranking_metrics_exact(
        Z1, Z2[perm], metric=metric, ks=recall_ks, max_n=max_pair_n, seed=seed, block=block
    )
    for k, v in shufM.items():
        M[f"PERM/{k}"] = v

    # quick “is permutation near chance?” flag for Recall@1
    # chance ~ 1/n_eval
    n_eval = shufM.get("FOSCTTM_n_eval", n_te)
    if n_eval and n_eval > 0:
        chance_r1 = 1.0 / float(n_eval)
        perm_r1 = shufM.get("FOSCTTM_Recall@1", np.nan)
        M["PERM/chance_Recall@1"] = float(chance_r1)
        M["PERM/Recall@1_over_chance"] = float(perm_r1 / chance_r1) if np.isfinite(perm_r1) else np.nan

    return M

def _pair_id_alignment_check(out):
    """
    If a method returns per-modality ids, report match rate.
    Looks for common key names; you can expand this list.
    """
    if not isinstance(out, dict):
        return {}
    keysets = [
        ("rna_ids", "atac_ids"),
        ("rna_ids", "adt_ids"),
        ("obs_names_rna", "obs_names_atac"),
        ("obs_names_rna", "obs_names_adt"),
    ]
    for kr, km in keysets:
        if (kr in out) and (km in out):
            r = np.asarray(out[kr]).astype(str)
            m = np.asarray(out[km]).astype(str)
            if r.shape == m.shape and r.size > 0:
                return {"PAIR/id_match_rate": float(np.mean(r == m)), "PAIR/id_n": int(r.size)}
            return {"PAIR/id_shape_mismatch": 1.0}
    return {}  # silently skip if ids absent


# %% [code]
def fit_preproc_on_train(
    rna, atac, train_idx, *,
    rna_counts_layer="counts",
    atac_counts_layer="counts",
    n_hvg=2000,
    target_sum=1e4,
    n_lsi=101,
    n_peaks_multivi=4002,
    dr_min=0.01,
    dr_max=0.30,
    seed=0,
):
    # RNA HVGs (TRAIN only)
    hvg = fit_hvgs_on_train(rna, train_idx, counts_layer=rna_counts_layer, n_hvg=n_hvg, seed=seed)

    # ATAC TFIDF+SVD(+scaler) (TRAIN only)
    tfidf, svd, scaler = fit_atac_lsi_on_train(
        atac, train_idx, counts_layer=atac_counts_layer, n_lsi=n_lsi, seed=seed,
        do_l2_norm=False, do_scale=True,
    )

    # ATAC peak selection (TRAIN only)
    peaks = fit_peaks_by_detection_window_on_train(
        atac, train_idx,
        counts_layer=atac_counts_layer,
        dr_min=dr_min,
        dr_max=dr_max,
        n_peaks=n_peaks_multivi,
        prefer_var="bernoulli",
    )

    return dict(hvg=hvg, tfidf=tfidf, svd=svd, scaler=scaler, peaks=peaks)


def transform_with_preproc(
    rna, atac, artifacts, *,
    rna_counts_layer="counts",
    atac_counts_layer="counts",
    target_sum=1e4,
    n_lsi=101,
    subset_chr=True,
):
    hvg   = artifacts["hvg"]
    tfidf = artifacts["tfidf"]
    svd   = artifacts["svd"]
    scaler= artifacts["scaler"]
    peaks = artifacts["peaks"]

    # RNA counts subset (HVG)
    rna = ensure_counts_layer(rna, layer=rna_counts_layer)
    rna_counts_hvg = rna[:, hvg].copy()
    rna_counts_hvg.X = rna_counts_hvg.layers[rna_counts_layer].copy()

    # RNA log subset (HVG)
    rna_log_hvg = transform_rna_log_hvg(rna, hvg, counts_layer=rna_counts_layer, target_sum=target_sum)

    # ATAC LSI (ALL using train-fit tfidf/svd/scaler)
    atac_lsi = transform_atac_lsi(atac, tfidf, svd, scaler, counts_layer=atac_counts_layer, n_lsi=n_lsi)

    # ATAC binarized (ALL) + selected peaks (from train)
    atac = ensure_counts_layer(atac, layer=atac_counts_layer)
    atac_counts_bin = atac.copy()
    X = _to_csr_float32(atac_counts_bin.layers[atac_counts_layer])
    atac_counts_bin.X = binarize_csr(X)

    atac_counts_bin_hv = atac_counts_bin[:, peaks].copy()
    if subset_chr:
        atac_counts_bin_hv = subset_to_chr_features(atac_counts_bin_hv)

    return dict(
        rna_counts_hvg=rna_counts_hvg,
        rna_log_hvg=rna_log_hvg,
        atac_lsi=atac_lsi,
        atac_counts_bin=atac_counts_bin,
        atac_counts_bin_hv=atac_counts_bin_hv,
        # also return artifacts for debugging
        hvg=hvg, peaks=peaks, tfidf=tfidf, svd=svd, scaler=scaler,
    )


# %% [code]
# ============================================================
# Seed-sweep robustness + full metric suite (Multiome-ready)
# - varies splits across seeds (shared across methods)
# - rebuilds any preprocessing fit-on-train per fold
# - does NOT change run_* methods: uses globals RNG_SEED/splits/shared inputs
# - computes: FOSCTTM mean, MRR, Recall@1/10, label transfer acc/macroF1,
#             mixing score, fused kNN acc/macroF1/balanced acc,
#             fused silhouette, label purity, kmeans ARI/NMI, fit seconds
# ============================================================

import time, json
from pathlib import Path
import numpy as np
import pandas as pd

from sklearn.model_selection import StratifiedKFold, StratifiedShuffleSplit
from sklearn.neighbors import NearestNeighbors
from sklearn.cluster import KMeans
from sklearn.metrics import (
    accuracy_score, f1_score, balanced_accuracy_score,
    adjusted_rand_score, normalized_mutual_info_score, silhouette_score
)

# -----------------------------
# 0) configure the sweep
# -----------------------------
SEEDS = list(range(5))
SWEEP_TAG = "seed_sweep_multiome_fullmetrics"

OUT_DIR = (Path(WORK) / "runs" / SWEEP_TAG) if "WORK" in globals() else (Path("./runs") / SWEEP_TAG)
OUT_DIR.mkdir(parents=True, exist_ok=True)

SPLIT_KWS = dict(
    train_frac=0.8,
    val_frac=0.1,
)

PREP_KWS = dict(
    rna_counts_layer="counts",
    atac_counts_layer="counts",
    n_hvg=2000,
    target_sum=1e4,
    n_lsi=101,
    n_hvpeaks_multivi=4002,
)

# evaluation hyperparams
EVAL_KWS = dict(
    metric="euclidean",                # "euclidean" or "cosine"
    k_label=15,                        # kNN for label transfer + fused kNN
    k_mixing=30,                       # kNN for mixing fraction
    recall_ks=(1, 10, 25, 50, 100),    # Recall@1/10/25/50/100
    max_pair_n=3000,                   # subsample size for exact pair retrieval metrics
    max_sil_n=3000,                    # subsample size for silhouette
    seed=0,
    block=512,                         # block size for pairwise ranking
)

# -----------------------------
# 1) optional: seed everything
# -----------------------------
def set_all_seeds(seed: int):
    np.random.seed(int(seed))
    try:
        import random
        random.seed(int(seed))
    except Exception:
        pass
    try:
        import torch
        torch.manual_seed(int(seed))
        torch.cuda.manual_seed_all(int(seed))
        torch.backends.cudnn.deterministic = True
        torch.backends.cudnn.benchmark = False
    except Exception:
        pass
        

# -----------------------------
# 2) fold builder (train/val/test)
# -----------------------------
from sklearn.model_selection import StratifiedShuffleSplit
import numpy as np

def make_folds(labels_all, n_splits=5, seed=0, train_frac=0.8, val_frac=0.1, test_frac=0.1):
    """
    Repeated stratified holdout splits with fractions of TOTAL data.
    n_splits = number of repeats (what you currently call folds).
    """
    labels_all = np.asarray(labels_all)
    n = len(labels_all)

    train_frac = float(train_frac)
    val_frac   = float(val_frac)
    test_frac  = float(test_frac)
    if abs((train_frac + val_frac + test_frac) - 1.0) > 1e-8:
        raise ValueError("train_frac + val_frac + test_frac must equal 1.")

    # 1) split off train vs (val+test)
    sss1 = StratifiedShuffleSplit(
        n_splits=int(n_splits),
        test_size=(val_frac + test_frac),
        random_state=int(seed),
    )

    for fold, (train_idx, temp_idx) in enumerate(sss1.split(np.zeros(n), labels_all)):
        y_temp = labels_all[temp_idx]

        # 2) split temp into val vs test (stratified)
        # val is val_frac / (val_frac + test_frac) of temp
        val_prop_of_temp = val_frac / (val_frac + test_frac)

        sss2 = StratifiedShuffleSplit(
            n_splits=1,
            test_size=(1.0 - val_prop_of_temp),   # portion of temp that becomes test
            random_state=int(seed + 10_000 + fold),
        )
        val_rel, test_rel = next(sss2.split(np.zeros(len(temp_idx)), y_temp))

        val_idx  = temp_idx[val_rel]
        test_idx = temp_idx[test_rel]

        yield fold, {
            "train": train_idx.tolist(),
            "val":   val_idx.tolist(),
            "test":  test_idx.tolist(),
            "unused": []
        }


# -----------------------------
# 3) rebuild preprocessing per fold (your wrapper)
# -----------------------------
def _call_fit_preproc_on_train_safe(fit_preproc_on_train, rna, atac, tr, *, seed, prep_kws, **extra):
    """
    Calls fit_preproc_on_train but removes keys it doesn't accept.
    Also maps common alias names.
    """
    import inspect

    kws = dict(prep_kws) if prep_kws is not None else {}

    # ---- alias mapping (if you store alt names in PREP_KWS) ----
    # If PREP_KWS uses n_hvpeaks_multivi, map it to the argument your function expects.
    # If your fit_preproc_on_train expects n_peaks_multivi, this will work.
    if "n_hvpeaks_multivi" in kws and "n_peaks_multivi" not in kws:
        kws["n_peaks_multivi"] = kws.pop("n_hvpeaks_multivi")

    # ---- remove any keys not in signature ----
    sig = inspect.signature(fit_preproc_on_train)
    allowed = set(sig.parameters.keys())

    # keep only kwargs that are accepted (or function has **kwargs)
    has_var_kw = any(p.kind == inspect.Parameter.VAR_KEYWORD for p in sig.parameters.values())
    if not has_var_kw:
        kws = {k: v for k, v in kws.items() if k in allowed}

    # extra explicit overrides win
    kws.update(extra)

    return fit_preproc_on_train(rna, atac, tr, seed=int(seed), **kws)


def rebuild_shared_inputs_for_split(seed: int, splits: dict):
    tr = np.asarray(splits["train"], dtype=int)

    artifacts = _call_fit_preproc_on_train_safe(
        fit_preproc_on_train,
        rna, atac, tr,
        seed=int(seed),
        prep_kws=PREP_KWS,
        # your extra knobs here (only if fit_preproc_on_train supports them)
        dr_min=0.01,
        dr_max=0.30,
        n_peaks_multivi=PREP_KWS.get("n_hvpeaks_multivi", 4002),  # still fine
    )

    shared = transform_with_preproc(
        rna, atac, artifacts,
        target_sum=PREP_KWS.get("target_sum", 1e4),
        subset_chr=True,
    )

    globals()["shared"] = shared


# ============================================================
# 4) Metric helpers (FULL suite)
# ============================================================
def _is_labeled(y):
    y = np.asarray(y).astype(str)
    bad = (y == "nan") | (y == "None") | (y == "NA") | (y == "")
    return ~bad


def _subsample_idx(n, max_n, seed=0):
    if (max_n is None) or (n <= max_n):
        return np.arange(n, dtype=int)
    rng = np.random.default_rng(int(seed))
    return rng.choice(n, size=int(max_n), replace=False)


def _dist_block(A, B, metric="euclidean"):
    A = np.asarray(A, np.float32)
    B = np.asarray(B, np.float32)
    if metric == "euclidean":
        # squared euclidean; ranking invariant vs euclidean
        A2 = (A * A).sum(1, keepdims=True)              # (a,1)
        B2 = (B * B).sum(1, keepdims=True).T            # (1,b)
        D2 = A2 + B2 - 2.0 * (A @ B.T)
        return np.maximum(D2, 0.0)
    elif metric == "cosine":
        An = A / (np.linalg.norm(A, axis=1, keepdims=True) + 1e-8)
        Bn = B / (np.linalg.norm(B, axis=1, keepdims=True) + 1e-8)
        # cosine distance = 1 - cos sim
        return 1.0 - (An @ Bn.T)
    else:
        raise ValueError("metric must be 'euclidean' or 'cosine'")


def pair_ranking_metrics_exact(Z1, Z2, *, metric="euclidean", ks=(1,10), max_n=3000, seed=0, block=512):
    """
    Exact (on subsample) FOSCTTM mean, MRR, Recall@k based on rank of true pair.
    Z1[i] paired with Z2[i].
    Uses blockwise distance comparisons; does NOT materialize full n×n matrix.
    """
    Z1 = np.asarray(Z1, np.float32)
    Z2 = np.asarray(Z2, np.float32)
    assert Z1.shape == Z2.shape
    n0 = Z1.shape[0]

    idx = _subsample_idx(n0, max_n, seed=seed)
    Z1 = Z1[idx]; Z2 = Z2[idx]
    n = Z1.shape[0]

    # true distances (diag)
    true_d = np.empty(n, dtype=np.float32)
    for i in range(0, n, block):
        sl = slice(i, min(n, i+block))
        D = _dist_block(Z1[sl], Z2[sl], metric=metric)
        true_d[sl] = np.diag(D)

    # count how many targets are closer than the true pair for each query
    closer = np.zeros(n, dtype=np.int32)

    for j in range(0, n, block):
        Bj = Z2[j:min(n, j+block)]
        D = _dist_block(Z1, Bj, metric=metric)  # (n, b)
        closer += (D < true_d[:, None]).sum(axis=1).astype(np.int32)

    rank = closer + 1  # 1-based rank
    foscttm = (rank - 1) / (n - 1 if n > 1 else 1)
    mrr = (1.0 / rank).mean()

    out = {
        "FOSCTTM_mean": float(foscttm.mean()),   # ↓ better
        "FOSCTTM_MRR": float(mrr),               # ↑ better
        "FOSCTTM_n_eval": int(n),
    }
    for k in ks:
        out[f"FOSCTTM_Recall@{int(k)}"] = float((rank <= int(k)).mean())
    return out


def label_transfer_knn(Z_src, y_src, Z_tgt, y_tgt, *, k=15, metric="euclidean"):
    y_src = np.asarray(y_src).astype(str)
    y_tgt = np.asarray(y_tgt).astype(str)

    msrc = _is_labeled(y_src) & np.isfinite(Z_src).all(1)
    mtgt = _is_labeled(y_tgt) & np.isfinite(Z_tgt).all(1)
    if msrc.sum() < 10 or mtgt.sum() < 10:
        return np.nan, np.nan

    Zs = np.asarray(Z_src[msrc], np.float32)
    Zt = np.asarray(Z_tgt[mtgt], np.float32)
    ys = y_src[msrc]
    yt = y_tgt[mtgt]

    nn = NearestNeighbors(n_neighbors=int(k), metric=metric)
    nn.fit(Zs)
    ind = nn.kneighbors(Zt, return_distance=False)

    preds = []
    for nbrs in ind:
        labs = ys[nbrs]
        vals, counts = np.unique(labs, return_counts=True)
        preds.append(vals[np.argmax(counts)])
    preds = np.asarray(preds, dtype=str)

    acc = float(accuracy_score(yt, preds))
    f1  = float(f1_score(yt, preds, average="macro"))
    return acc, f1


def mixing_score(Z, domain, *, k=30, metric="euclidean"):
    Z = np.asarray(Z, np.float32)
    domain = np.asarray(domain).astype(str)

    if Z.shape[0] < (k + 5):
        return np.nan, {}

    nn = NearestNeighbors(n_neighbors=int(k)+1, metric=metric)
    nn.fit(Z)
    ind = nn.kneighbors(Z, return_distance=False)[:, 1:]
    dom_nbr = domain[ind]
    frac_other = (dom_nbr != domain[:, None]).mean(axis=1)

    per_dom = {d: float(frac_other[domain == d].mean()) for d in np.unique(domain)}
    return float(frac_other.mean()), per_dom


def fused_knn_metrics(Z, y, train_idx, test_idx, *, k=15, metric="euclidean"):
    Z = np.asarray(Z, np.float32)
    y = np.asarray(y).astype(str)
    tr = np.asarray(train_idx, dtype=int)
    te = np.asarray(test_idx, dtype=int)

    mtr = _is_labeled(y[tr]) & np.isfinite(Z[tr]).all(1)
    mte = _is_labeled(y[te]) & np.isfinite(Z[te]).all(1)
    if mtr.sum() < 20 or mte.sum() < 20:
        return {"Fused kNN acc": np.nan, "Fused kNN macroF1": np.nan, "Fused kNN balanced acc": np.nan}

    Ztr, ytr = Z[tr][mtr], y[tr][mtr]
    Zte, yte = Z[te][mte], y[te][mte]

    nn = NearestNeighbors(n_neighbors=int(k), metric=metric)
    nn.fit(Ztr)
    ind = nn.kneighbors(Zte, return_distance=False)

    preds = []
    for nbrs in ind:
        labs = ytr[nbrs]
        vals, counts = np.unique(labs, return_counts=True)
        preds.append(vals[np.argmax(counts)])
    preds = np.asarray(preds, dtype=str)

    return {
        "Fused kNN acc": float(accuracy_score(yte, preds)),
        "Fused kNN macroF1": float(f1_score(yte, preds, average="macro")),
        "Fused kNN balanced acc": float(balanced_accuracy_score(yte, preds)),
    }
    

def clustering_metrics(Z, y, *, seed=0, max_sil_n=3000):
    Z = np.asarray(Z, np.float32)
    y = np.asarray(y).astype(str)
    m = _is_labeled(y) & np.isfinite(Z).all(1)
    Z, y = Z[m], y[m]
    if Z.shape[0] < 50 or len(np.unique(y)) < 2:
        return {
            "Fused silhouette": np.nan,
            "Fused label purity": np.nan,
            "Fused k-means ARI": np.nan,
            "Fused k-means NMI": np.nan,
        }

    k = max(2, len(np.unique(y)))
    km = KMeans(n_clusters=int(k), n_init=10, random_state=int(seed))
    cl = km.fit_predict(Z)

    # purity
    purity = 0.0
    for c in np.unique(cl):
        labs = y[cl == c]
        vals, counts = np.unique(labs, return_counts=True)
        purity += counts.max()
    purity = float(purity / len(y))

    ari = float(adjusted_rand_score(y, cl))
    nmi = float(normalized_mutual_info_score(y, cl))

    # silhouette on subsample
    idx = _subsample_idx(len(y), max_sil_n, seed=seed)
    sil = float(silhouette_score(Z[idx], y[idx], metric="euclidean"))

    return {
        "Fused silhouette": sil,
        "Fused label purity": purity,
        "Fused k-means ARI": ari,
        "Fused k-means NMI": nmi,
    }


def _extract_metrics(out):
    """Keep your old behavior: pull numeric scalars from out or out['metrics']/etc."""
    if out is None:
        return {}
    candidates = []
    if isinstance(out, dict):
        candidates.append(out)
        for k in ["summary", "metrics", "scores", "eval", "results", "extra_json"]:
            if k in out and isinstance(out[k], dict):
                candidates.append(out[k])
    merged = {}
    for c in candidates:
        merged.update(c)
    flat = {}
    for k, v in merged.items():
        if isinstance(v, (int, float, np.integer, np.floating)) and np.isfinite(v):
            flat[k] = float(v)
        elif isinstance(v, (list, tuple, np.ndarray)) and len(v) == 1:
            try:
                flat[k] = float(v[0])
            except Exception:
                pass
    return flat


def evaluate_out_full(
    out, labels_all, splits, *,
    metric="euclidean", k_label=15, k_mixing=30,
    recall_ks=(1,10), max_pair_n=3000, max_sil_n=3000,
    seed=0, block=512,
    pair_eval_split="all",          # NEW: "all" or "test"
    mixing_eval_split="all",        # NEW: "all" or "test" (optional)
):
    """
    Computes full metric suite using:
      - Z_rna + Z_atac (or Z_adt) if present (paired, aligned)
      - Z_fused if present (aligned)

    pair_eval_split:
      - "all": compute paired retrieval/label-transfer/mixing on all cells (old behavior)
      - "test": compute those only on splits["test"] (strict CV)

    mixing_eval_split:
      - defaults to "all" to preserve old behavior
      - set to "test" if you want mixing/clustering on test only as well
    """
    labels_all = np.asarray(labels_all).astype(str)
    n = len(labels_all)
    M = {}

    if not isinstance(out, dict):
        return M

    # unify naming
    Z_rna = out.get("Z_rna", None)
    Z_mod = out.get("Z_atac", None)
    if Z_mod is None:
        Z_mod = out.get("Z_adt", None)
    Z_fused = out.get("Z_fused", None)

    # ---------- choose index subset for paired metrics ----------
    if pair_eval_split not in ("all", "test"):
        raise ValueError("pair_eval_split must be 'all' or 'test'")
    if mixing_eval_split not in ("all", "test"):
        raise ValueError("mixing_eval_split must be 'all' or 'test'")

    idx_pair = np.arange(n, dtype=int) if pair_eval_split == "all" else np.asarray(splits["test"], dtype=int)
    idx_mix  = np.arange(n, dtype=int) if mixing_eval_split == "all" else np.asarray(splits["test"], dtype=int)

    # ---------- paired metrics require both modality embeddings ----------
    if (Z_rna is not None) and (Z_mod is not None):
        Z_rna = np.asarray(Z_rna, np.float32)
        Z_mod = np.asarray(Z_mod, np.float32)

        if Z_rna.shape[0] == n and Z_mod.shape[0] == n:
            # subset first, then finite-mask
            Z1 = Z_rna[idx_pair]
            Z2 = Z_mod[idx_pair]
            y  = labels_all[idx_pair]

            m = np.isfinite(Z1).all(1) & np.isfinite(Z2).all(1)
            Z1, Z2, y = Z1[m], Z2[m], y[m]

            if Z1.shape[0] >= 10:
                pairM = pair_ranking_metrics_exact(
                    Z1, Z2,
                    metric=metric, ks=recall_ks, max_n=max_pair_n,
                    seed=seed, block=block
                )
                # prefix so it's obvious what split you used
                pref = "PAIR(all)/" if pair_eval_split == "all" else "PAIR(test)/"
                for k, v in pairM.items():
                    M[pref + k] = v

                # label transfer both directions (mean)
                acc12, f112 = label_transfer_knn(Z1, y, Z2, y, k=k_label, metric=metric)
                acc21, f121 = label_transfer_knn(Z2, y, Z1, y, k=k_label, metric=metric)
                M[pref + "Label transfer acc mean"] = float(np.nanmean([acc12, acc21]))
                M[pref + "Label transfer macroF1 mean"] = float(np.nanmean([f112, f121]))

            # mixing/clustering on stacked embedding (optionally subset)
            Z1m = Z_rna[idx_mix]
            Z2m = Z_mod[idx_mix]
            ym  = labels_all[idx_mix]
            mm  = np.isfinite(Z1m).all(1) & np.isfinite(Z2m).all(1)
            Z1m, Z2m, ym = Z1m[mm], Z2m[mm], ym[mm]

            if Z1m.shape[0] >= (k_mixing + 5) and len(np.unique(ym)) >= 2:
                Zstack = np.vstack([Z1m, Z2m])
                ystack = np.concatenate([ym, ym])
                domain = np.array(["RNA"] * len(ym) + ["MOD"] * len(ym), dtype=str)

                mix, per_dom = mixing_score(Zstack, domain, k=k_mixing, metric=metric)
                mpref = "MIX(all)/" if mixing_eval_split == "all" else "MIX(test)/"
                M[mpref + "Mixing score"] = mix
                M[mpref + "Mixing score (RNA)"] = per_dom.get("RNA", np.nan)
                M[mpref + "Mixing score (MOD)"] = per_dom.get("MOD", np.nan)

                # clustering-style metrics on stacked embedding
                cm = clustering_metrics(Zstack, ystack, seed=seed, max_sil_n=max_sil_n)
                for k, v in cm.items():
                    M[mpref + k] = v

        # else: shapes not aligned -> skip pair metrics

    # ---------- fused metrics ----------
    if Z_fused is not None:
        Zf = np.asarray(Z_fused, np.float32)
        if Zf.shape[0] == n:
            # NOTE: this is already strict train->test by construction
            M.update(fused_knn_metrics(Zf, labels_all, splits["train"], splits["test"], k=k_label, metric=metric))

            # clustering metrics on fused embedding (ALL labeled by default)
            #M.update({k.replace("Fused ", "Fused(fusedZ) "): v
            #          for k, v in clustering_metrics(Zf, labels_all, seed=seed, max_sil_n=max_sil_n).items()})
                       
            # clustering metrics on fused embedding (TEST only)
            te = np.asarray(splits["test"], dtype=int)
            Zf_te = Zf[te]
            y_te  = labels_all[te]
            M.update({k.replace("Fused ", "Fused(fusedZ,test) "): v
                       for k, v in clustering_metrics(Zf_te, y_te, seed=seed, max_sil_n=max_sil_n).items()})

    return M
    

# ============================================================
# 5) main CV sweep (rewritten)
# ============================================================
def run_cv_sweep_fullmetrics(
    *,
    seeds,
    n_folds,
    out_dir,
    methods,
    labels_all,
    eval_kws=None,
):
    eval_kws = dict(EVAL_KWS if eval_kws is None else eval_kws)

    out_dir = Path(out_dir)
    out_dir.mkdir(parents=True, exist_ok=True)

    rows = []
    raw_out_path = out_dir / "raw_outputs.jsonl"

    with raw_out_path.open("w") as f_jsonl:
        for seed in seeds:
            set_all_seeds(int(seed))

            for fold, splits in make_folds(labels_all, n_splits=n_folds, seed=seed):
                globals()["RNG_SEED"] = int(seed)
                globals()["splits"] = splits

                rebuild_shared_inputs_for_split(seed, splits)

                print(f"\n=== seed={seed} fold={fold} sizes ===", {k: len(v) for k, v in splits.items()})

                for method_name, thunk in methods.items():
                    t0 = time.time()
                    status, err, out = "ok", None, None

                    try:
                        out = thunk()
                    except Exception as e:
                        status = "fail"
                        err = repr(e)

                    dt = time.time() - t0

                    # start with whatever the runner already reported
                    metrics = _extract_metrics(out)

                    # add full metric suite + sanity checks
                    if status == "ok":
                        try:
                            eval_kws2 = dict(eval_kws)
                            eval_kws2.pop("seed", None)

                            # ---- your existing suite (note: this currently uses ALL cells for pair metrics) ----
                            #addM = evaluate_out_full(out, labels_all, splits, seed=seed, **eval_kws2)
                            #addM = evaluate_out_full(out, labels_all, splits, seed=seed, pair_eval_split="test", **eval_kws2)
                            # can add: mixing_eval_split="test", # <- optional; leave off if you want mixing on all
                            addM = evaluate_out_full(
                                out, labels_all, splits,
                                seed=seed,
                                pair_eval_split="test",
                                mixing_eval_split="test",
                                **eval_kws2
                            )
                            
                            for k, v in addM.items():
                                if k not in metrics:
                                    metrics[k] = v

                            # ---- sanity: if ids are available, check 1:1 pairing by name ----
                            for k, v in _pair_id_alignment_check(out).items():
                                if k not in metrics:
                                    metrics[k] = v

                            # ---- sanity: TEST-only pair metrics + permutation control ----
                            # (this is the big one for “is FOSCTTM real?”)
                            test_pairM = _pair_test_metrics(out, labels_all, splits, seed=seed, **eval_kws2)
                            for k, v in test_pairM.items():
                                if k not in metrics:
                                    metrics[k] = v

                        except Exception as e:
                            metrics["__eval_error__"] = repr(e)


                    row = {
                        "seed": int(seed),
                        "fold": int(fold),
                        "method": str(method_name),
                        "status": status,
                        "fit_seconds": float(dt),
                        "error": err,
                        "n_train": int(len(splits["train"])),
                        "n_val": int(len(splits["val"])),
                        "n_test": int(len(splits["test"])),
                        **metrics,
                    }
                    rows.append(row)

                    f_jsonl.write(json.dumps({
                        "seed": int(seed),
                        "fold": int(fold),
                        "method": str(method_name),
                        "status": status,
                        "fit_seconds": float(dt),
                        "error": err,
                        "metrics": metrics,
                        "out_keys": sorted(list(out.keys())) if isinstance(out, dict) else None,
                    }) + "\n")

    df = pd.DataFrame(rows)
    df.to_csv(out_dir / "cv_sweep_long.csv", index=False)

    ok = df[df["status"] == "ok"].copy()
    house = {"seed","fold","method","status","fit_seconds","error","n_train","n_val","n_test"}
    metric_cols = [c for c in ok.columns if c not in house and pd.api.types.is_numeric_dtype(ok[c])]

    if metric_cols:
        summ = (ok.groupby("method")[metric_cols]
                  .agg(["mean","std","median","count"]))
        # pick a stable sort if present
        sort_key = ("FOSCTTM_mean", "mean") if ("FOSCTTM_mean" in ok.columns) else (metric_cols[0], "mean")
        summ = summ.sort_values(sort_key, ascending=True if sort_key[0].startswith("FOSCTTM_mean") else False)
        summ.to_csv(out_dir / "cv_sweep_summary.csv")
    else:
        summ = ok.groupby("method").size().to_frame("n_ok")
        summ.to_csv(out_dir / "cv_sweep_summary.csv")

    print("\nSaved:")
    print(" -", out_dir / "cv_sweep_long.csv")
    print(" -", out_dir / "cv_sweep_summary.csv")
    print(" -", raw_out_path)

    return df, summ

# ------------------------------------------------------------
# Run it:
# df_long, df_summary = run_cv_sweep_fullmetrics(
#     seeds=SEEDS,
#     n_folds=5,
#     out_dir=OUT_DIR,
#     methods=methods,
#     labels_all=labels_all,
# )
# ------------------------------------------------------------


# %% [code]
#SEEDS = list(range(10))
SEEDS = [67, 1985, 789, 3, 99]
N_FOLDS = 3
OUT_DIR = Path(WORK) / "runs" / "cv_sweep_py_1-31-2026"
OUT_DIR.mkdir(parents=True, exist_ok=True)

df, summ = run_cv_sweep_fullmetrics(
    seeds=SEEDS,
    n_folds=N_FOLDS,
    out_dir=OUT_DIR,
    methods=METHODS,
    labels_all=labels_all,
)

ok = df[df["status"] == "ok"].copy()

for m in ["FOSCTTM_mean_test", "fused_kmeans_ari_test", "fused_kmeans_nmi_test"]:
    if m in ok.columns:
        violin_metric(ok, m, savepath=OUT_DIR / f"violin_{m}.png")


# %% [code]
import numpy as np
import pandas as pd
import matplotlib.pyplot as plt

# -----------------------------
# plotting
# -----------------------------
import matplotlib.pyplot as plt

def violin_metric(df_ok: pd.DataFrame, metric: str, *, title=None, savepath=None):
    dd = df_ok[["method", metric]].dropna().copy()
    if dd.empty:
        print(f"[plot] no data for metric={metric}")
        return

    order = dd.groupby("method")[metric].mean().sort_values(ascending=False).index.tolist()
    data = [dd.loc[dd["method"] == m, metric].to_numpy() for m in order]

    means = np.array([np.mean(x) for x in data], dtype=float)
    sems  = np.array([np.std(x, ddof=1) / np.sqrt(len(x)) if len(x) > 1 else np.nan for x in data], dtype=float)
    medians = np.array([np.median(x) for x in data], dtype=float)

    fig, ax = plt.subplots(figsize=(max(6, 0.85 * len(order)), 4.5))
    ax.violinplot(data, positions=np.arange(1, len(order) + 1),
                  showmeans=False, showextrema=False, showmedians=False)
    xs = np.arange(1, len(order) + 1)
    ax.errorbar(xs, means, yerr=sems, fmt="o", capsize=4, linewidth=1.5)
    ax.scatter(xs, medians, marker="_", s=300)

    ax.set_xticks(xs)
    ax.set_xticklabels(order, rotation=30, ha="right")
    ax.set_ylabel(metric)
    ax.set_title(title or f"{metric} across CV folds (mean±SEM, median tick)")
    ax.grid(True, axis="y", alpha=0.3)
    fig.tight_layout()
    if savepath is not None:
        sp = Path(savepath)
        sp.parent.mkdir(parents=True, exist_ok=True)  # make dirs if needed
        fig.savefig(sp, dpi=200)                      # overwrites if exists
        
    plt.show()


# %% [code]
house = {"seed",'fold',"method","status","error","n_train","n_val","n_test","n_unused",'transductive','uses_labels','n_genes',
        'n_peaks','latent_dim','dropout','lr','weight_decay','batch_size','max_epochs','patience','reg','best_val','T','lamb',
        'nbatches','n_latent'}


# %% [code]
cols = [c for c in df.columns if "FOSCTTM" in c or "Mixing" in c or "Label transfer" in c or "Fused" in c]
df[["seed","fold","method","status"] + cols].head()


# %% [code]
print(ok.columns)


# %% [code]
print(summ.columns)


# %% [code]
metric_cols = [c for c in ok.columns if c not in house and pd.api.types.is_numeric_dtype(ok[c])]


# %% [code]
for m in ["FOSCTTM_mean_test", "fused_kmeans_ari_test", "fused_kmeans_nmi_test"]:
    if m in ok.columns:
        violin_metric(ok, m, savepath=OUT_DIR / f"violin_{m}.png")
        

# %% [code]
# pick metrics you care about (or use your auto-detected metric_cols)
for metric in metric_cols:
    violin_metric(ok, metric, savepath=OUT_DIR / f"violin_{metric}.png")
    

# %% [code]


# %% [code]


# %% [code]


# %% [code]


# %% [code]


# %% [code]


# %% [code]


# %% [code]


# %% [code]


# %% [code]
````
