## Supplemental Code for "Unifying multimodal single-cell data with a mixture-of-experts *β*-variational autoencoder framework": Fig08_benchmark_R.ipynb

```
# %% [markdown]
## UniVI manuscript - Figure 6 generation reproducible workflow
### Commonly-used data integration tool benchmarking test for Genome Research manuscript revisions - UniVI benchmark (Multiome PBMC): comparing UniVI against widely-used R baselines


Andrew Ashford, Pathways + Omics Group, Oregon Health & Science University, Portland, OR – 1/25/2026


This notebook:
- loads paired Multiome PBMC (RNA+ATAC)
- runs integration baselines in an R environment:
  - Seurat/Signac WNN (canonical Multiome baseline)
  - Harmony (batch correction baseline; requires ≥2 batches to be meaningful)
  - LIGER (iNMF; gene-activity-based if annotation is available)
  - MOFA2 (factor model on PCA+LSI views; low pain + well-cited)
- evaluates with:
  - FOSCTTM (paired alignment; only for methods returning both RNA+ATAC embeddings)
  - modality mixing (kNN entropy proxy; needs modality-specific embeddings)
  - kNN label transfer (RNA→ATAC, ATAC→RNA proxies via fused embedding)
- saves embeddings + a summary table
  
Manuscript: https://www.biorxiv.org/content/10.1101/2025.02.28.640429v1.full
Code: https://github.com/Ashford-A/UniVI


# %% [markdown]
### Setting cache directories

# %% [code]
# ---- paths ----
RDS  <- "/home/groups/precepts/ashforda"
WORK <- file.path(RDS, "univi_bench", "runs_multiome_R_1-31-2026")
dir.create(WORK, recursive = TRUE, showWarnings = FALSE)

DATA_ROOT <- "/home/groups/precepts/ashforda/UniVI_v2/UniVI_older-non_git/data/PBMC_10x_Multiome_data/10x_Genomics_Multiome_data/"
RNA_PATH  <- file.path(DATA_ROOT, "10x-Multiome-Pbmc10k-RNA.h5ad")
ATAC_PATH <- file.path(DATA_ROOT, "10x-Multiome-Pbmc10k-ATAC.h5ad")

# Optional:
FRAG_PATH <- file.path(DATA_ROOT, "pbmc10k_multiome/pbmc_granulocyte_sorted_10k_atac_fragments.tsv.gz")

cat("WORK:", WORK, "\n")
cat("RNA :", RNA_PATH, "\n")
cat("ATAC:", ATAC_PATH, "\n")
cat("FRAG_PATH (optional):", FRAG_PATH, "\n")


# %% [markdown]
### Import modules

# %% [code]
requireNamespace("aricode", quietly = TRUE)
packageVersion("aricode")


# %% [code]
n <- as.integer(Sys.getenv("SLURM_CPUS_PER_TASK", "1"))
n <- ifelse(is.na(n) || n < 1L, 1L, n)

Sys.setenv(
  OMP_NUM_THREADS = n,
  OPENBLAS_NUM_THREADS = n,
  MKL_NUM_THREADS = n,
  VECLIB_MAXIMUM_THREADS = n,
  NUMEXPR_NUM_THREADS = n
)

if (requireNamespace("RhpcBLASctl", quietly = TRUE)) {
  RhpcBLASctl::blas_set_num_threads(n)
  RhpcBLASctl::omp_set_num_threads(n)
}

message("[threads] n=", n,
        " OMP=", Sys.getenv("OMP_NUM_THREADS"),
        " OPENBLAS=", Sys.getenv("OPENBLAS_NUM_THREADS"),
        " MKL=", Sys.getenv("MKL_NUM_THREADS"))


# %% [code]
# ============================================================
# Cluster-safe R deps bootstrap (micromamba-first)
# - Avoid building heavy packages from source on compute nodes
# - Prefer micromamba/conda-forge/bioconda for big deps
# - Allow optional methods (MOFA2, ArchR) to be skipped cleanly
# ============================================================

.msg <- function(...) message(...)
ENV_NAME <- {
  pref <- Sys.getenv("CONDA_PREFIX", "")
  if (nzchar(pref)) basename(pref) else Sys.getenv("CONDA_DEFAULT_ENV", "univi-bench-r")
}

mm_hint <- function(pkgs, env_name = ENV_NAME, channels = c("conda-forge", "bioconda")) {
  ch <- paste(paste0("-c ", channels), collapse = " ")
  paste0("micromamba install -n ", env_name, " ", ch, " ", paste(pkgs, collapse = " "))
}

require_or_stop <- function(pkgs, install_hint = NULL) {
  missing <- pkgs[!vapply(pkgs, requireNamespace, logical(1), quietly = TRUE)]
  if (length(missing)) {
    msg <- paste0("[deps] Missing packages: ", paste(missing, collapse = ", "))
    if (!is.null(install_hint)) msg <- paste0(msg, "\n", install_hint)
    stop(msg, call. = FALSE)
  }
  invisible(TRUE)
}

# ---- core required deps ----
core_pkgs <- c(
  "Matrix", "data.table", "ggplot2",
  "Seurat", "SeuratObject",
  "Signac",
  "SingleCellExperiment", "zellkonverter",
  "FNN", "RcppAnnoy"
)

core_pkgs <- unique(c(core_pkgs, "GenomicRanges", "IRanges"))

# Note: SummarizedExperiment is used implicitly by zellkonverter outputs
core_pkgs <- unique(c(core_pkgs, "SummarizedExperiment"))

# Recommend installing these via micromamba (heavy)
hint_core <- mm_hint(c(
  "r-matrix", "r-data.table", "r-ggplot2",
  "r-seurat", "r-seuratobject", "r-signac",
  "r-fnn", "r-rcppannoy",
  "bioconductor-singlecellexperiment", "bioconductor-zellkonverter",
  "bioconductor-summarizedexperiment", "bioconductor-genomicranges", "bioconductor-iranges"
))

require_or_stop(core_pkgs, install_hint = paste0("Suggested micromamba install:\n  ", hint_core))

suppressPackageStartupMessages({
  library(Matrix)
  library(data.table)
  library(ggplot2)
  library(Seurat)
  library(SeuratObject)
  library(Signac)
  library(SingleCellExperiment)
  library(zellkonverter)
  library(SummarizedExperiment)
  library(GenomicRanges)
  library(IRanges)
})

# ---- optional deps ----
has_harmony <- requireNamespace("harmony", quietly = TRUE)
has_liger   <- requireNamespace("rliger", quietly = TRUE)
has_mofa2   <- requireNamespace("MOFA2", quietly = TRUE)

if (!has_harmony) .msg(paste0("[optional] harmony missing. Install:\n  ", mm_hint("r-harmony")))
if (!has_liger)   .msg(paste0("[optional] rliger missing. Install:\n  ", mm_hint("r-rliger")))
if (!has_mofa2)   .msg(paste0("[optional] MOFA2 missing. Install:\n  ", mm_hint("bioconductor-mofa2")))


# %% [code]
parallel::detectCores()
future::availableCores()
system("nproc", intern = TRUE)
system("grep Cpus_allowed_list /proc/self/status", intern = TRUE)


# %% [code]
# -----------------------------
# core-limit (R CMD check / envs)
# -----------------------------
# This env var forces future/parallel to pretend you only have 2 cores.
# It does NOT reflect your actual cgroup CPU mask.
Sys.unsetenv("_R_CHECK_LIMIT_CORES_")
Sys.unsetenv("R_CHECK_LIMIT_CORES")

# If future already cached the old value, reset later when future is loaded:
.reset_future_cores <- function() {
  if (requireNamespace("future", quietly = TRUE)) {
    try(future::availableCores(reset = TRUE), silent = TRUE)
  }
  invisible(NULL)
}

# -----------------------------
# threads (BLAS/OpenMP)
# -----------------------------
# Default: keep BLAS conservative to avoid oversubscription,
# then explicitly open it up only when you mean to.
set_blas_threads <- function(n = 1L) {
  n <- as.integer(n)
  if (!is.finite(n) || n < 1L) n <- 1L
  Sys.setenv(
    OMP_NUM_THREADS = as.character(n),
    OPENBLAS_NUM_THREADS = as.character(n),
    MKL_NUM_THREADS = as.character(n),
    VECLIB_MAXIMUM_THREADS = as.character(n),
    NUMEXPR_NUM_THREADS = as.character(n)
  )
  invisible(n)
}

# Start safe (1 thread)
set_blas_threads(1L)

# -----------------------------
# future controls
# -----------------------------
if (requireNamespace("future", quietly = TRUE)) {
  .reset_future_cores()

  # Global default: sequential (avoids Seurat NormalizeData() globals explosion)
  future::plan(future::sequential)

  # Raise in case some part still uses futures internally
  options(future.globals.maxSize = 200 * 1024^3)  # 200 GiB (adjust if needed)
}

# Helper: workers from Slurm (fallback default)
slurm_workers <- function(default = 1L) {
  x <- suppressWarnings(as.integer(Sys.getenv("SLURM_CPUS_PER_TASK", unset = as.character(default))))
  if (!is.finite(x) || x < 1L) x <- default
  x
}

# Helper: run a function under a temporary future plan, then restore
with_future_plan <- function(plan, workers = NULL, fun, ...) {
  if (!requireNamespace("future", quietly = TRUE)) return(fun(...))
  old <- future::plan()
  on.exit(future::plan(old), add = TRUE)

  .reset_future_cores()
  if (!is.null(workers)) {
    future::plan(plan, workers = workers)
  } else {
    future::plan(plan)
  }
  fun(...)
}


# %% [code]
parallel::detectCores()
future::availableCores()
system("nproc", intern = TRUE)
system("grep Cpus_allowed_list /proc/self/status", intern = TRUE)


# %% [markdown]
### Helper methods - timing, subsampling, saving, etc..

# %% [code]
`%||%` <- function(a, b) {
  if (is.null(a)) return(b)
  if (length(a) == 0) return(b)
  if (is.character(a) && length(a) == 1L && !nzchar(a)) return(b)
  a
}

# ------------------------------------------------------------
# Seurat v4/v5 compatibility: GetAssayData(slot=) -> layer=
# ------------------------------------------------------------
get_layer <- function(seurat_obj, layer = "counts", assay = NULL) {
  if (!is.null(assay)) Seurat::DefaultAssay(seurat_obj) <- assay
  ass <- Seurat::DefaultAssay(seurat_obj)

  # ---- Seurat v5 path: LayerData (preferred) ----
  if (exists("LayerData", where = asNamespace("SeuratObject"), inherits = FALSE)) {
    # If the requested layer doesn't exist, be explicit (helps debugging)
    lay_names <- tryCatch(
      SeuratObject::Layers(seurat_obj[[ass]]),
      error = function(e) character(0)
    )
    if (length(lay_names) > 0 && !(layer %in% lay_names)) {
      stop(sprintf(
        "[get_layer] assay='%s' has layers: %s ; requested layer='%s' not found",
        ass, paste(lay_names, collapse = ", "), layer
      ), call. = FALSE)
    }

    return(SeuratObject::LayerData(seurat_obj[[ass]], layer = layer))
  }

  # ---- Older Seurat path: GetAssayData(slot=...) ----
  return(SeuratObject::GetAssayData(seurat_obj, assay = ass, slot = layer))
}

# ------------------------------------------------------------
# Global Seurat v4/v5 compatibility: assay matrix getter
# - Works for SeuratObject v5 layers and older slot-based assays
# ------------------------------------------------------------
get_assay_mat <- function(obj, assay, layer = "counts") {
  a <- obj[[assay]]

  # Try SeuratObject v5 "layer=" first
  out <- tryCatch(
    SeuratObject::GetAssayData(a, layer = layer),
    error = function(e) e
  )
  if (!inherits(out, "error")) return(out)

  msg <- conditionMessage(out)

  # If 'layer' arg isn't supported (older SeuratObject), fall back to slot=
  if (grepl("unused argument.*layer|formal argument.*layer|\\blayer\\b.*not used", msg, ignore.case = TRUE)) {
    slot <- layer
    if (layer %in% c("scale", "scale.data")) slot <- "scale.data"

    out2 <- tryCatch(
      SeuratObject::GetAssayData(a, slot = slot),
      error = function(e) e
    )
    if (inherits(out2, "error")) {
      stop(
        "GetAssayData failed for assay=", assay,
        " layer/slot=", layer,
        " (layer error: ", msg,
        "; slot error: ", conditionMessage(out2), ")",
        call. = FALSE
      )
    }
    return(out2)
  }

  # Otherwise it's a real error (e.g., missing layer)
  stop(
    "GetAssayData failed for assay=", assay,
    " layer=", layer, " (", msg, ")",
    call. = FALSE
  )
}


# %% [code]
l2norm_rows <- function(X) {
  X <- as.matrix(X)
  d <- sqrt(rowSums(X * X)) + 1e-12
  X / d
}

knn_idx_annoy <- function(X, k=50, n_trees=50) {
  if (!requireNamespace("RcppAnnoy", quietly = TRUE)) stop("Need RcppAnnoy installed.")
  X <- as.matrix(X)
  a <- RcppAnnoy::AnnoyEuclidean$new(ncol(X))
  for (i in seq_len(nrow(X))) a$addItem(i - 1L, X[i, ])
  a$build(n_trees)

  idx <- vector("list", nrow(X))
  for (i in seq_len(nrow(X))) {
    nn <- a$getNNsByItem(i - 1L, k + 1L, includeDistances = FALSE)
    nn <- nn[nn != (i - 1L)][1:k]
    idx[[i]] <- nn + 1L
  }
  idx
}

modality_mixing_entropy <- function(Z, modality, k=50, idx_eval=NULL) {
  if (!is.null(idx_eval)) {
    Z <- Z[idx_eval, , drop=FALSE]
    modality <- modality[idx_eval]
  }
  nn <- knn_idx_annoy(Z, k=k)
  ent <- numeric(length(nn))
  for (i in seq_along(nn)) {
    m <- modality[unlist(nn[[i]])]
    p <- table(m) / length(m)
    ent[i] <- -sum(p * log(p + 1e-12))
  }
  mean(ent)
}

knn_predict_labels <- function(Z_train, y_train, Z_query, k=15) {
  if (!requireNamespace("FNN", quietly = TRUE)) stop("Need FNN installed.")
  nn <- FNN::get.knnx(data=as.matrix(Z_train), query=as.matrix(Z_query), k=k)$nn.index
  apply(nn, 1, function(ii) {
    yy <- y_train[ii]
    names(sort(table(yy), decreasing=TRUE))[1]
  })
}

label_transfer_metrics <- function(Z_fused, labels, splits, k=15) {
  tr <- splits$train
  te <- splits$test
  yhat <- knn_predict_labels(Z_fused[tr, , drop=FALSE], labels[tr], Z_fused[te, , drop=FALSE], k=k)
  list(acc_test = mean(yhat == labels[te]))
}

foscttm_values <- function(Za, Zb, idx_eval, subsample_n=3000, seed=0) {
  set.seed(seed)
  idx <- idx_eval
  if (!is.null(subsample_n) && length(idx) > subsample_n) idx <- sample(idx, subsample_n)

  A <- as.matrix(Za[idx, , drop=FALSE])
  B <- as.matrix(Zb[idx, , drop=FALSE])

  A2 <- rowSums(A * A)
  B2 <- rowSums(B * B)
  D  <- outer(A2, B2, "+") - 2 * (A %*% t(B))
  D[D < 0] <- 0

  ranks <- numeric(nrow(D))
  for (i in seq_len(nrow(D))) {
    ranks[i] <- rank(D[i, ], ties.method="average")[i]
  }
  (ranks - 1) / (nrow(D) - 1)
}

mixing_proxy <- function(Zr, Za, idx_eval, k=50) {
  # Build joint point cloud by duplicating cells (RNA points + ATAC points)
  Z_joint <- rbind(Zr, Za)
  mod <- c(rep("RNA", nrow(Zr)), rep("ATAC", nrow(Za)))
  idx2 <- c(idx_eval, idx_eval + nrow(Zr))
  modality_mixing_entropy(Z_joint, mod, k=k, idx_eval=idx2)
}

mixing_proxy_rann <- function(Zr, Za, idx_eval, k=50, normalized=TRUE) {
  if (!requireNamespace("RANN", quietly=TRUE)) {
    stop("Need RANN installed for mixing_proxy_rann(). Install.packages('RANN')")
  }
  if (is.null(idx_eval) || length(idx_eval) < 2) return(NA_real_)

  # subset to eval cells FIRST so kNN is well-defined and fast
  Zr <- Zr[idx_eval, , drop=FALSE]
  Za <- Za[idx_eval, , drop=FALSE]

  # joint matrix: (2*n_eval) x d
  Z_joint <- rbind(Zr, Za)
  mod <- c(rep("rna", nrow(Zr)), rep("atac", nrow(Za)))

  # sanity: no NA/Inf
  if (!all(is.finite(Z_joint))) {
    stop("Z_joint contains NA/Inf; cannot compute mixing.")
  }

  n <- nrow(Z_joint)
  # kNN needs k+1 <= n; we remove self neighbor later
  k_eff <- min(as.integer(k), n - 2L)
  if (k_eff < 2L) return(NA_real_)

  nn <- RANN::nn2(data=Z_joint, query=Z_joint, k=k_eff + 1L)
  idx <- nn$nn.idx

  # drop self neighbor: typically first column is self (distance 0)
  idx <- idx[, -1, drop=FALSE]

  # compute modality entropy per point
  ent <- numeric(n)
  for (i in seq_len(n)) {
    neigh_mod <- mod[idx[i, ]]
    p <- table(neigh_mod) / length(neigh_mod)
    ent[i] <- -sum(p * log(p))
  }

  if (isTRUE(normalized)) ent <- ent / log(2)  # for 2 modalities -> [0,1]
  mean(ent)
}


# %% [code]
# -------------------------
# Helpers: canonicalize + align paired embeddings
# -------------------------
.canon_cell_ids <- function(x) {
  x <- as.character(x)
  x <- gsub("\\s+", "", x)
  x
}

.align_paired_latents <- function(Zr, Za, min_common = 50L, enforce_order = TRUE) {
  if (is.null(Zr) || is.null(Za)) return(list(Zr = NULL, Za = NULL, cells = NULL, reason = "missing"))
  Zr <- as.matrix(Zr); Za <- as.matrix(Za)
  if (is.null(rownames(Zr)) || is.null(rownames(Za))) {
    return(list(Zr = NULL, Za = NULL, cells = NULL, reason = "missing_rownames"))
  }

  cr <- .canon_cell_ids(rownames(Zr))
  ca <- .canon_cell_ids(rownames(Za))
  rownames(Zr) <- cr
  rownames(Za) <- ca

  common <- intersect(cr, ca)
  if (length(common) < min_common) {
    return(list(Zr = NULL, Za = NULL, cells = NULL, reason = paste0("too_few_common=", length(common))))
  }

  # Preserve RNA order by default (important for reproducibility)
  if (enforce_order) {
    common <- cr[cr %in% common]
  }

  Zr2 <- Zr[common, , drop = FALSE]
  Za2 <- Za[common, , drop = FALSE]

  # Ensure identical rownames after subsetting
  if (!identical(rownames(Zr2), rownames(Za2))) {
    # fall back to explicit ordering using match()
    Za2 <- Za2[match(rownames(Zr2), rownames(Za2)), , drop = FALSE]
    if (!identical(rownames(Zr2), rownames(Za2))) {
      return(list(Zr = NULL, Za = NULL, cells = NULL, reason = "cannot_align_order"))
    }
  }

  list(Zr = Zr2, Za = Za2, cells = rownames(Zr2), reason = NA_character_)
}

.l2norm_rows <- function(Z, eps = 1e-12) {
  Z <- as.matrix(Z)
  nrm <- sqrt(rowSums(Z * Z))
  nrm <- pmax(nrm, eps)
  Z / nrm
}


# %% [code]
.assert_fold_cells_ok <- function(fold_cells, ctx = "") {
  if (!is.list(fold_cells)) {
    stop(sprintf("%s fold_cells must be a list(train,val,test).", ctx))
  }
  need <- c("train", "val", "test")
  if (!all(need %in% names(fold_cells))) {
    stop(sprintf("%s fold_cells must have names: %s", ctx, paste(need, collapse = ",")))
  }

  # coerce to character vectors, drop empties, unique
  for (nm in need) {
    x <- fold_cells[[nm]]
    x <- unique(as.character(x))
    x <- x[nzchar(x)]
    fold_cells[[nm]] <- x
  }

  # sanity: no overlaps (this is what usually matters for CV)
  ov_tv <- intersect(fold_cells$train, fold_cells$val)
  ov_tt <- intersect(fold_cells$train, fold_cells$test)
  ov_vt <- intersect(fold_cells$val, fold_cells$test)
  if (length(ov_tv) || length(ov_tt) || length(ov_vt)) {
    msg <- c()
    if (length(ov_tv)) msg <- c(msg, sprintf("train∩val=%d", length(ov_tv)))
    if (length(ov_tt)) msg <- c(msg, sprintf("train∩test=%d", length(ov_tt)))
    if (length(ov_vt)) msg <- c(msg, sprintf("val∩test=%d", length(ov_vt)))
    stop(sprintf("%s fold_cells splits overlap: %s", ctx, paste(msg, collapse = " | ")))
  }

  # sanity: non-empty
  if (length(fold_cells$train) < 1 || length(fold_cells$val) < 1 || length(fold_cells$test) < 1) {
    stop(sprintf(
      "%s fold_cells has empty split(s): train=%d val=%d test=%d",
      ctx, length(fold_cells$train), length(fold_cells$val), length(fold_cells$test)
    ))
  }

  invisible(TRUE)
}


# %% [code]
.subset_seurat_safe <- function(obj, cells, ctx = "") {
  if (!requireNamespace("SeuratObject", quietly = TRUE)) stop("Need SeuratObject installed.")
  if (!requireNamespace("Seurat", quietly = TRUE)) stop("Need Seurat installed.")

  all_cells <- tryCatch(SeuratObject::Cells(obj), error = function(e) colnames(obj))
  cells <- unique(as.character(cells))
  cells <- cells[nzchar(cells)]
  keep <- intersect(cells, all_cells)

  if (!length(keep)) {
    stop(sprintf("%s subset: 0/%d requested cells found in object.", ctx, length(cells)))
  }

  # Seurat::subset is usually safest across versions; use cells= for Seurat objects
  out <- tryCatch(
    Seurat::subset(obj, cells = keep),
    error = function(e1) {
      # fallback: bracket subset if subset() fails for some reason
      tryCatch(obj[, keep, drop = FALSE], error = function(e2) e2)
    }
  )

  if (inherits(out, "error")) {
    stop(sprintf("%s subset failed: %s", ctx, conditionMessage(out)))
  }
  out
}


# %% [code]
# -----------------------------
# Latent alignment helpers
# -----------------------------
.as_mat <- function(Z) {
  if (is.null(Z)) return(NULL)
  if (inherits(Z, "dgCMatrix") || inherits(Z, "Matrix")) return(as.matrix(Z))
  as.matrix(Z)
}

.ensure_rownames <- function(Z, cell_ids, name = "Z") {
  Z <- .as_mat(Z)
  if (is.null(Z)) return(NULL)

  # If no rownames, assume it's in the same order as cell_ids
  if (is.null(rownames(Z))) {
    if (!is.null(cell_ids) && length(cell_ids) == nrow(Z)) {
      rownames(Z) <- cell_ids
    } else {
      stop(sprintf("[%s] Missing rownames and cannot infer (nrow=%d, len(cell_ids)=%s).",
                   name, nrow(Z), ifelse(is.null(cell_ids), "NULL", length(cell_ids))))
    }
  }
  Z
}

.reindex_rows <- function(Z, cell_ids, name = "Z", allow_drop = TRUE) {
  Z <- .as_mat(Z)
  if (is.null(Z)) return(NULL)
  if (is.null(rownames(Z))) stop(sprintf("[%s] rownames required for reindexing.", name))

  keep <- intersect(cell_ids, rownames(Z))
  if (length(keep) == 0) {
    stop(sprintf("[%s] No overlap with requested cell_ids (len=%d).", name, length(cell_ids)))
  }
  if (!allow_drop && length(keep) != length(cell_ids)) {
    miss <- setdiff(cell_ids, rownames(Z))
    stop(sprintf("[%s] Missing %d required rows (e.g. %s).",
                 name, length(miss), paste(head(miss, 3), collapse = ", ")))
  }

  Z2 <- Z[cell_ids[cell_ids %in% rownames(Z)], , drop = FALSE]
  Z2
}

.check_latents <- function(Z_rna, Z_atac, Z_fused, ss = NULL, tag = "") {
  tr <- character(0)
  te <- character(0)
  fold <- character(0)

  if (is.list(ss)) {
    tr   <- as.character(ss$train_cells %||% character(0))
    te   <- as.character(ss$test_cells  %||% character(0))
    fold <- as.character(ss$fold_cells  %||% character(0))
  }

  .one <- function(Z, nm) {
    if (is.null(Z)) {
      cat(tag, nm, ": NULL\n")
      return()
    }
    rn <- rownames(Z)
    cat(
      tag, nm,
      " dim=", paste(dim(Z), collapse = "x"),
      " rn=", !is.null(rn),
      " fold_overlap=", if (!is.null(rn) && length(fold)) length(intersect(fold, rn)) else NA_integer_,
      " test_overlap=", if (!is.null(rn) && length(te))   length(intersect(te, rn))   else NA_integer_,
      " train_overlap=",if (!is.null(rn) && length(tr))   length(intersect(tr, rn))   else NA_integer_,
      "\n"
    )
  }

  .one(Z_rna,  "Z_rna")
  .one(Z_atac, "Z_atac")
  .one(Z_fused,"Z_fused")
}

# Inductive RNA PCA: fit on TRAIN only, project ALL fold cells
.compute_rna_pca_inductive <- function(
  obj,
  fold_cells,
  train_cells,
  npcs = 30L,
  nfeatures = 2000L,
  verbose = TRUE
) {
  if (!requireNamespace("Matrix", quietly = TRUE)) stop("Need Matrix installed.")
  if (!requireNamespace("Seurat", quietly = TRUE)) stop("Need Seurat installed.")

  .msg <- function(...) if (isTRUE(verbose)) cat(..., "\n")

  fold_cells  <- intersect(as.character(fold_cells), colnames(obj))
  train_cells <- intersect(as.character(train_cells), fold_cells)
  if (length(fold_cells) < 50) stop("[RNA-PCA] too few fold cells")
  if (length(train_cells) < 50) stop("[RNA-PCA] too few train cells")

  obj_fold <- subset(obj, cells = fold_cells)
  Seurat::DefaultAssay(obj_fold) <- "RNA"

  # ensure normalized data exists
  obj_fold <- Seurat::NormalizeData(obj_fold, verbose = FALSE)

  # HVGs on TRAIN only
  obj_tr <- subset(obj_fold, cells = train_cells)
  obj_tr <- Seurat::FindVariableFeatures(obj_tr, nfeatures = as.integer(nfeatures), verbose = FALSE)
  feats <- Seurat::VariableFeatures(obj_tr)
  if (length(feats) < 50) stop("[RNA-PCA] too few HVGs on TRAIN")

  # log-normalized data matrix (features x cells)
  X_all <- .get_assay_data_compat(obj_fold, assay = "RNA", layer = "data")
  X_tr  <- X_all[feats, train_cells, drop = FALSE]
  X_all <- X_all[feats, fold_cells,  drop = FALSE]

  # prcomp wants cell x feature
  Xt_tr <- t(as.matrix(X_tr))
  mu  <- colMeans(Xt_tr)
  sdv <- apply(Xt_tr, 2, stats::sd)
  sdv[!is.finite(sdv) | sdv <= 1e-8] <- 1.0

  Xs_tr <- sweep(Xt_tr, 2, mu, FUN = "-")
  Xs_tr <- sweep(Xs_tr, 2, sdv, FUN = "/")

  k <- min(as.integer(npcs), ncol(Xs_tr))
  if (k < 2) stop("[RNA-PCA] npcs too small after processing")

  .msg(sprintf("[RNA-PCA] fit prcomp on TRAIN: n=%d feats=%d npcs=%d", nrow(Xs_tr), ncol(Xs_tr), k))
  pc <- stats::prcomp(Xs_tr, center = FALSE, scale. = FALSE, rank. = k)

  Xt_all <- t(as.matrix(X_all))
  Xs_all <- sweep(Xt_all, 2, mu, FUN = "-")
  Xs_all <- sweep(Xs_all, 2, sdv, FUN = "/")
  Z_all  <- Xs_all %*% pc$rotation[, seq_len(k), drop = FALSE]

  rownames(Z_all) <- fold_cells
  colnames(Z_all) <- paste0("PC", seq_len(k))
  Z_all
}

.safeguard_bridge_future <- function(max_gb = 20, sequential = TRUE, verbose = TRUE) {
  .msg <- function(...) if (isTRUE(verbose)) cat(..., "\n")

  if (!requireNamespace("future", quietly = TRUE)) {
    .msg("[Bridge] NOTE: package 'future' not installed; cannot set future.globals.maxSize / plan.")
    return(invisible(NULL))
  }

  # Increase allowed export size (some Seurat/Signac internals can trip this)
  max_bytes <- as.numeric(max_gb) * 1024^3
  old_opt <- getOption("future.globals.maxSize")
  options(future.globals.maxSize = max_bytes)

  if (isTRUE(sequential)) {
    # Most reliable to avoid gigantic globals being exported to workers
    old_plan <- tryCatch(future::plan(), error = function(e) NULL)
    suppressWarnings(try(future::plan("sequential"), silent = TRUE))
    .msg(sprintf("[Bridge] future safeguard: globals.maxSize=%.1f GiB; plan=sequential", max_gb))
    return(invisible(list(old_opt = old_opt, old_plan = old_plan)))
  } else {
    .msg(sprintf("[Bridge] future safeguard: globals.maxSize=%.1f GiB (plan unchanged)", max_gb))
    return(invisible(list(old_opt = old_opt, old_plan = NULL)))
  }
}

.get_assay_data_compat <- function(obj, assay = "RNA", layer = "data") {
  if (!requireNamespace("SeuratObject", quietly = TRUE)) stop("Need SeuratObject installed.")
  a <- obj[[assay]]
  # SeuratObject v5+: layer=
  out <- tryCatch(SeuratObject::GetAssayData(a, layer = layer), error = function(e) e)
  if (!inherits(out, "error")) return(out)

  # Older SeuratObject: slot=
  out2 <- tryCatch(SeuratObject::GetAssayData(a, slot = layer), error = function(e) e)
  if (inherits(out2, "error")) {
    stop(sprintf("GetAssayData failed for assay=%s layer=%s: %s",
                 assay, layer, conditionMessage(out)))
  }
  out2
}

.ensure_counts_layer <- function(obj, assay) {
  a <- obj[[assay]]
  lys <- tryCatch(SeuratObject::Layers(a), error = function(e) character(0))

  if (!("counts" %in% lys)) {
    # try to pull counts from existing storage
    x <- tryCatch(SeuratObject::GetAssayData(a, layer = "counts"), error = function(e) NULL)
    if (is.null(x)) {
      x <- tryCatch(SeuratObject::GetAssayData(a, slot = "counts"), error = function(e) NULL)
    }
    if (is.null(x)) stop("[Bridge] cannot materialize counts for assay=", assay)

    # write it as a layer
    SeuratObject::LayerData(a, layer = "counts") <- x
    obj[[assay]] <- a
  }
  obj
}


# %% [code]
# -----------------------------
# .finalize_latents() (R syntax)
# -----------------------------
.finalize_latents <- function(
  Z_rna,
  Z_atac,
  Z_fused,
  fold_cells,
  labels = NULL,
  splits = NULL,
  ss = NULL,
  method_str = "method",
  extra_json = list(),
  verbose = FALSE
) {
  .msg <- function(...) if (isTRUE(verbose)) { cat(..., "\n"); flush.console() }

  # Prefer ss labels/splits if provided (your old behavior)
  if (!is.null(ss)) {
    if (is.null(labels) && !is.null(ss$labels)) labels <- ss$labels
    if (is.null(splits) && !is.null(ss$splits)) splits <- ss$splits
  }

  # -------------------------
  # Normalize fold_cells into list(train/val/test)
  # Accept BOTH:
  #   - list(train,val,test)  [preferred]
  #   - character vector of cell IDs (common inside runners)
  # -------------------------
  needed <- c("train", "val", "test")

  if (is.null(fold_cells)) {
    stop("[finalize] fold_cells is NULL")
  }

  if (is.list(fold_cells)) {
    if (is.null(names(fold_cells))) stop("[finalize] fold_cells list must be named")
    if (!all(needed %in% names(fold_cells))) {
      stop("[finalize] fold_cells list must contain names: train, val, test")
    }
    # ok
  } else {
    # treat as a character vector of fold universe cells
    if (!is.character(fold_cells)) {
      stop(sprintf("[finalize] fold_cells must be list(train/val/test) or character; got: %s",
                   paste(class(fold_cells), collapse = ",")))
    }

    fold_univ <- unique(as.character(fold_cells))
    if (!length(fold_univ)) stop("[finalize] fold_cells character vector is empty")

    # splits may be indices OR cell IDs; normalize to cell IDs if possible
    tr_cells <- character(0)
    te_cells <- character(0)

    if (is.list(splits)) {
      if (!is.null(splits$train)) {
        if (is.numeric(splits$train)) {
          idx <- as.integer(splits$train)
          idx <- idx[!is.na(idx) & idx >= 1 & idx <= length(fold_univ)]
          tr_cells <- fold_univ[idx]
        } else {
          tr_cells <- intersect(as.character(splits$train), fold_univ)
        }
      }
      if (!is.null(splits$test)) {
        if (is.numeric(splits$test)) {
          idx <- as.integer(splits$test)
          idx <- idx[!is.na(idx) & idx >= 1 & idx <= length(fold_univ)]
          te_cells <- fold_univ[idx]
        } else {
          te_cells <- intersect(as.character(splits$test), fold_univ)
        }
      }
    }

    # If splits weren't usable, default: train=all, test=empty
    if (!length(tr_cells) && !length(te_cells)) {
      tr_cells <- fold_univ
      te_cells <- character(0)
    }

    # val = whatever is left
    val_cells <- setdiff(fold_univ, union(tr_cells, te_cells))

    fold_cells <- list(train = tr_cells, val = val_cells, test = te_cells)

    .msg(sprintf(
      "[finalize] coerced fold_cells from character -> list: train=%d val=%d test=%d",
      length(fold_cells$train), length(fold_cells$val), length(fold_cells$test)
    ))
  }

  # -------------------------
  # Validate latents + rownames
  # -------------------------
  .check_latent <- function(Z, nm) {
    if (is.null(Z)) return(list(ok = FALSE, why = "NULL"))
    Zm <- tryCatch(as.matrix(Z), error = function(e) NULL)
    if (is.null(Zm) || !is.matrix(Zm)) return(list(ok = FALSE, why = "not matrix-coercible"))
    if (nrow(Zm) < 2 || ncol(Zm) < 2) return(list(ok = FALSE, why = sprintf("bad shape %dx%d", nrow(Zm), ncol(Zm))))
    rn <- rownames(Zm)
    if (is.null(rn) || any(!nzchar(rn))) return(list(ok = FALSE, why = "missing/empty rownames"))
    list(ok = TRUE, Z = Zm)
  }

  ckf <- .check_latent(Z_fused, "Z_fused")
  if (!ckf$ok) {
    .msg(sprintf("[finalize] %s invalid: %s", method_str, ckf$why))
    return(list(
      Z_rna = NULL, Z_atac = NULL, Z_fused = NULL,
      extra_json = c(list(method = method_str, skipped = TRUE, reason = paste0("[finalize] Z_fused invalid: ", ckf$why)), extra_json)
    ))
  }

  Z_fused <- ckf$Z

  # Keep only fold universe that exists in Z_fused
  fold_univ <- unique(c(fold_cells$train, fold_cells$val, fold_cells$test))
  fold_univ <- intersect(fold_univ, rownames(Z_fused))

  if (length(fold_univ) < 50) {
    .msg(sprintf("[finalize] %s: too few fold_univ cells in Z_fused: %d", method_str, length(fold_univ)))
    .msg(sprintf("[finalize] Z_fused rows=%d ; train=%d val=%d test=%d",
                 nrow(Z_fused), length(fold_cells$train), length(fold_cells$val), length(fold_cells$test)))
    return(list(
      Z_rna = NULL, Z_atac = NULL, Z_fused = NULL,
      extra_json = c(list(method = method_str, skipped = TRUE,
                          reason = paste0("[finalize] too few fold cells present in Z_fused: ", length(fold_univ))),
                     extra_json)
    ))
  }

  # Reorder fused to fold_univ for consistency
  Z_fused <- Z_fused[fold_univ, , drop = FALSE]

  # OPTIONAL: keep paired latents if they are compatible
  Z_rna_ok  <- .check_latent(Z_rna,  "Z_rna")
  Z_atac_ok <- .check_latent(Z_atac, "Z_atac")

  if (Z_rna_ok$ok) {
    Z_rna <- Z_rna_ok$Z
    common <- intersect(rownames(Z_rna), fold_univ)
    Z_rna <- Z_rna[common, , drop = FALSE]
  } else {
    Z_rna <- NULL
  }

  if (Z_atac_ok$ok) {
    Z_atac <- Z_atac_ok$Z
    common <- intersect(rownames(Z_atac), fold_univ)
    Z_atac <- Z_atac[common, , drop = FALSE]
  } else {
    Z_atac <- NULL
  }

  # Final fold_cells cleanup to match kept universe
  fold_cells <- lapply(fold_cells, function(x) intersect(as.character(x), fold_univ))

  .msg(sprintf(
    "[finalize] %s: kept cells=%d | train=%d val=%d test=%d | Zf=%dx%d Zr=%s Za=%s",
    method_str,
    length(fold_univ),
    length(fold_cells$train), length(fold_cells$val), length(fold_cells$test),
    nrow(Z_fused), ncol(Z_fused),
    if (is.null(Z_rna)) "NULL" else paste(dim(Z_rna), collapse = "x"),
    if (is.null(Z_atac)) "NULL" else paste(dim(Z_atac), collapse = "x")
  ))

  list(
    Z_rna = Z_rna,
    Z_atac = Z_atac,
    Z_fused = Z_fused,
    fold_cells = fold_cells,
    labels = labels,
    splits = splits,
    extra_json = c(list(method = method_str, skipped = FALSE, reason = NA_character_), extra_json)
  )
}


# %% [markdown]
### Specify LSI and PCA dims and latent dims globally to use for methods that use these

# %% [code]
.clamp_dims <- function(dims, n_avail) {
  dims <- as.integer(dims)
  dims[dims >= 1L & dims <= as.integer(n_avail)]
}

.safe_embed <- function(obj, red, dims) {
  Z <- Seurat::Embeddings(obj, red)
  d <- .clamp_dims(dims, ncol(Z))
  if (length(d) < 2) {
    stop(sprintf("[%s] Not enough dims after clamp. requested=[%s], avail=%d",
                 red, paste(dims, collapse=","), ncol(Z)), call.=FALSE)
  }
  Z[, d, drop=FALSE]
}


# %% [code]
DIMS_LSI <- 2:101   # ATAC
DIMS_PCA <- 1:100   # RNA

clamp_dims <- function(dims, n_avail) intersect(as.integer(dims), seq_len(n_avail))

LATENT_K  <- 30


# %% [markdown]
### Load Multiome data

# %% [code]
# Read as SingleCellExperiment
sce_rna  <- zellkonverter::readH5AD(RNA_PATH)
sce_atac <- zellkonverter::readH5AD(ATAC_PATH)

cat("sce_rna :", dim(sce_rna), "\n")
cat("sce_atac:", dim(sce_atac), "\n")

# ---------------------------
# RNA -> Seurat
# ---------------------------
rna_counts <- SummarizedExperiment::assay(sce_rna, "X")
if (!inherits(rna_counts, "dgCMatrix")) rna_counts <- as(rna_counts, "dgCMatrix")

obj_rna <- CreateSeuratObject(
  counts  = rna_counts,
  assay   = "RNA",
  project = "pbmc_multiome"
)

# ---------------------------
# ATAC -> Seurat (ChromatinAssay from var columns)
# ---------------------------
atac_counts <- SummarizedExperiment::assay(sce_atac, "X")
if (!inherits(atac_counts, "dgCMatrix")) atac_counts <- as(atac_counts, "dgCMatrix")

rd_atac <- as.data.frame(SummarizedExperiment::rowData(sce_atac))

need_cols <- c("chrom", "chromStart", "chromEnd")
missing_cols <- setdiff(need_cols, colnames(rd_atac))
if (length(missing_cols)) {
  stop("ATAC var is missing required columns: ", paste(missing_cols, collapse=", "))
}

# Signac expects 1-based start in GRanges; many exports store 0-based starts.
# We'll do a conservative fix: if any chromStart==0, shift by +1.
start1 <- as.integer(rd_atac$chromStart)
end1   <- as.integer(rd_atac$chromEnd)
if (any(start1 == 0L, na.rm=TRUE)) start1 <- start1 + 1L

gr_peaks <- GenomicRanges::GRanges(
  seqnames = rd_atac$chrom,
  ranges   = IRanges::IRanges(start = start1, end = end1)
)

# Give peaks stable names if not present
peak_names <- rownames(atac_counts)
if (is.null(peak_names) || any(!nzchar(peak_names))) {
  peak_names <- paste0(rd_atac$chrom, ":", start1, "-", end1)
  rownames(atac_counts) <- peak_names
}
names(gr_peaks) <- peak_names

chrom_assay <- CreateChromatinAssay(
  counts = atac_counts,
  ranges = gr_peaks,
  genome = NULL   # <- IMPORTANT: avoid seqinfo error
)

# after CreateChromatinAssay(...)
Signac::Fragments(chrom_assay) <- Signac::CreateFragmentObject(
  path = FRAG_PATH,
  cells = colnames(atac_counts)   # cells in the ATAC matrix
)

obj_atac <- CreateSeuratObject(
  counts  = chrom_assay,
  assay   = "ATAC",
  project = "pbmc_multiome"
)

# ---------------------------
# Attach shared metadata (obs)
# ---------------------------
md_rna  <- as.data.frame(SummarizedExperiment::colData(sce_rna))
md_atac <- as.data.frame(SummarizedExperiment::colData(sce_atac))
common_md_cols <- intersect(colnames(md_rna), colnames(md_atac))
if (length(common_md_cols) > 0) {
  obj_rna  <- AddMetaData(obj_rna,  metadata = md_rna[, common_md_cols, drop=FALSE])
  obj_atac <- AddMetaData(obj_atac, metadata = md_atac[, common_md_cols, drop=FALSE])
}

# ---------------------------
# Align paired cells
# ---------------------------
common_cells <- intersect(colnames(obj_rna), colnames(obj_atac))
stopifnot(length(common_cells) > 0)

obj_rna  <- obj_rna[, common_cells]
obj_atac <- obj_atac[, common_cells]
obj_atac <- obj_atac[, colnames(obj_rna)]

cat("paired n_cells:", ncol(obj_rna),
    " (RNA==ATAC order:", identical(colnames(obj_rna), colnames(obj_atac)), ")\n")

# ---------------------------
# Combine into one multi-assay object
# ---------------------------
obj <- obj_rna
obj[["ATAC"]] <- obj_atac[["ATAC"]]


# %% [code]
pick_node_scratch <- function() {
  cands <- c(Sys.getenv("SLURM_TMPDIR"), Sys.getenv("TMPDIR"), file.path("/tmp", Sys.getenv("USER")))
  cands <- cands[nzchar(cands)]
  for (d in cands) {
    dir.create(d, recursive = TRUE, showWarnings = FALSE)
    ok <- tryCatch({
      tf <- tempfile(tmpdir = d)
      writeLines("ok", tf)
      unlink(tf)
      TRUE
    }, error = function(e) FALSE)
    if (isTRUE(ok)) return(d)
  }
  stop("No writable node-local scratch found (tried SLURM_TMPDIR, TMPDIR, /tmp/$USER).")
}

stage_fragments_to_node <- function(frag_src, scratch_dir = NULL, verbose = TRUE) {
  .msg <- function(...) if (isTRUE(verbose)) cat(..., "\n")

  if (!file.exists(frag_src)) stop("frag_src does not exist: ", frag_src)

  scratch_dir <- scratch_dir %||% pick_node_scratch()
  frags_dir   <- file.path(scratch_dir, "frags")
  dir.create(frags_dir, recursive = TRUE, showWarnings = FALSE)

  frag_dst <- file.path(frags_dir, basename(frag_src))
  if (!file.exists(frag_dst)) {
    .msg("[stage] copying fragments to node scratch:")
    .msg("  src: ", frag_src)
    .msg("  dst: ", frag_dst)
    ok <- file.copy(frag_src, frag_dst, overwrite = FALSE)
    if (!isTRUE(ok)) stop("Failed to copy fragments to node scratch.")
  } else {
    .msg("[stage] fragments already present -> reusing: ", frag_dst)
  }

  # stage index if present
  tbi_src1 <- paste0(frag_src, ".tbi")
  tbi_src2 <- sub("\\.gz$", ".tbi", frag_src)

  tbi_dst1 <- paste0(frag_dst, ".tbi")
  tbi_dst2 <- sub("\\.gz$", ".tbi", frag_dst)

  if (file.exists(tbi_src1) && !file.exists(tbi_dst1)) file.copy(tbi_src1, tbi_dst1, overwrite = FALSE)
  if (file.exists(tbi_src2) && !file.exists(tbi_dst2)) file.copy(tbi_src2, tbi_dst2, overwrite = FALSE)

  has_tbi <- file.exists(tbi_dst1) || file.exists(tbi_dst2)
  if (!has_tbi) {
    stop(
      "Missing .tbi index next to staged fragment.\n",
      "Create it on the node shell: tabix -p bed ", shQuote(frag_dst),
      call. = FALSE
    )
  }

  frag_dst
}

assert_fragments_ok <- function(frag_path) {
  if (!file.exists(frag_path)) stop("Fragment file not found: ", frag_path)
  if (file.info(frag_path)$size < 1e6) {
    warning("Fragment file looks very small (<1MB). Path might be wrong: ", frag_path)
  }
  tbi1 <- paste0(frag_path, ".tbi")
  tbi2 <- sub("\\.gz$", ".tbi", frag_path)
  if (!(file.exists(tbi1) || file.exists(tbi2))) {
    stop("Fragment index (.tbi) not found next to: ", frag_path)
  }
  invisible(TRUE)
}

compute_gene_activity_cached <- function(
  obj,
  frag_src,
  assay_atac = "ATAC",
  assay_out  = "ACTIVITY",
  features   = NULL,
  do_normalize = FALSE,
  attach_cells = FALSE,   # set TRUE only if barcodes mismatch / filtering needed
  scratch_dir = NULL,
  verbose = TRUE
) {
  .msg <- function(...) if (isTRUE(verbose)) cat(..., "\n")

  if (!requireNamespace("Signac", quietly = TRUE)) stop("Need Signac installed.")
  if (!requireNamespace("Seurat", quietly = TRUE)) stop("Need Seurat installed.")

  if (assay_out %in% names(obj@assays)) {
    .msg("[GeneActivity] ", assay_out, " already present -> reuse.")
    return(list(obj = obj, frag_local = NA_character_))
  }

  frag_local <- stage_fragments_to_node(frag_src, scratch_dir = scratch_dir, verbose = verbose)
  assert_fragments_ok(frag_local)

  DefaultAssay(obj) <- assay_atac

  frags <- tryCatch(Signac::Fragments(obj), error = function(e) NULL)
  if (is.null(frags) || length(frags) == 0) {
    .msg("[GeneActivity] Attaching fragments: ", frag_local)
    Signac::Fragments(obj) <- Signac::CreateFragmentObject(
      path  = frag_local,
      cells = if (isTRUE(attach_cells)) colnames(obj) else NULL
    )
  }

  if (!is.null(features)) .msg("[GeneActivity] features n=", length(features))
  .msg("[GeneActivity] Computing gene activity…")
  ga_counts <- Signac::GeneActivity(obj, features = features, verbose = FALSE)

  obj[[assay_out]] <- Seurat::CreateAssayObject(counts = ga_counts)

  if (isTRUE(do_normalize)) {
    obj[[assay_out]] <- Seurat::NormalizeData(
      obj[[assay_out]],
      normalization.method = "LogNormalize",
      scale.factor = 1e4,
      verbose = FALSE
    )
  }

  .msg("[GeneActivity] Done. ", assay_out, " dim=", paste(dim(obj[[assay_out]]), collapse="x"))
  list(obj = obj, frag_local = frag_local)
}


ensure_node_local_fragments <- function(
  obj,
  frag_local,
  assay_atac = "ATAC",
  attach_cells = FALSE,
  verbose = TRUE
) {
  .msg <- function(...) if (isTRUE(verbose)) cat(..., "\n")
  .stop <- function(...) stop(paste0(...), call. = FALSE)

  if (!requireNamespace("Signac", quietly = TRUE)) .stop("Need Signac.")
  if (!inherits(obj, "Seurat")) .stop("obj must be Seurat.")
  if (!(assay_atac %in% names(obj@assays))) .stop("missing assay: ", assay_atac)
  if (!file.exists(frag_local)) .stop("frag_local not found: ", frag_local)

  DefaultAssay(obj) <- assay_atac

  frags <- tryCatch(Signac::Fragments(obj), error = function(e) NULL)
  if (!is.null(frags) && length(frags) > 0) {
    cur <- tryCatch(frags[[1]]@path, error = function(e) NA_character_)
    if (is.character(cur) && nzchar(cur) &&
        identical(normalizePath(cur), normalizePath(frag_local))) {
      .msg("[frags] already attached + node-local -> reuse")
      return(obj)
    }
    .msg("[frags] attached but path differs; switching to node-local: ", frag_local)
  } else {
    .msg("[frags] none attached; attaching node-local: ", frag_local)
  }

  frag_obj <- Signac::CreateFragmentObject(
    path = frag_local,
    cells = if (isTRUE(attach_cells)) Seurat::Cells(obj) else NULL,
    validate.fragments = FALSE
  )
  obj[[assay_atac]]@fragments <- list(frag_obj)
  obj
}


pick_ga_features <- function(obj, n = 4000L, prefer_hvg = TRUE) {
  # 1) VariableFeatures(RNA)
  hv <- tryCatch(Seurat::VariableFeatures(obj[["RNA"]]), error = function(e) character(0))
  hv <- unique(hv[!is.na(hv) & nzchar(hv)])
  if (prefer_hvg && length(hv) >= 10) return(head(hv, n))

  # 2) rownames(RNA)
  rn <- tryCatch(rownames(obj[["RNA"]]), error = function(e) character(0))
  rn <- unique(rn[!is.na(rn) & nzchar(rn)])
  if (length(rn) >= 10) return(head(rn, n))

  # 3) Annotation(ATAC)$gene_name (fallback)
  ann <- tryCatch(Signac::Annotation(obj[["ATAC"]]), error = function(e) NULL)
  if (!is.null(ann) && requireNamespace("S4Vectors", quietly = TRUE)) {
    if ("gene_name" %in% names(S4Vectors::mcols(ann))) {
      g <- unique(S4Vectors::mcols(ann)$gene_name)
      g <- g[!is.na(g) & nzchar(g)]
      if (length(g) >= 10) return(head(g, n))
    }
  }

  character(0)
}
                  

# %% [code]
compute_gene_activity_on_obj <- function(
  obj,
  assay_atac = "ATAC",
  assay_out = "ACTIVITY",
  features,
  extend_upstream = 2000,
  extend_downstream = 0,
  process_n = 8000,
  max_width = 5e5,
  biotypes = "protein_coding",
  do_normalize = FALSE,
  verbose = TRUE
) {
  .msg <- function(...) if (isTRUE(verbose)) cat(..., "\n")
  .stop <- function(...) stop(paste0(...), call. = FALSE)

  if (!requireNamespace("Signac", quietly = TRUE)) .stop("Need Signac.")
  if (!requireNamespace("Seurat", quietly = TRUE)) .stop("Need Seurat.")
  if (!inherits(obj, "Seurat")) .stop("obj must be Seurat.")
  if (!(assay_atac %in% names(obj@assays))) .stop("missing assay_atac=", assay_atac)

  if (assay_out %in% names(obj@assays)) {
    .msg("[GA] ", assay_out, " already present -> reuse")
    return(obj)
  }

  features <- unique(features[!is.na(features) & nzchar(features)])
  if (length(features) < 10) .stop("[GA] too few features: ", length(features))

  DefaultAssay(obj) <- assay_atac
  .msg("[GA] Computing gene activity… (cells=", ncol(obj), ", feats=", length(features), ")")
  ga_counts <- Signac::GeneActivity(
    object = obj,
    assay = assay_atac,
    features = features,
    extend.upstream = extend_upstream,
    extend.downstream = extend_downstream,
    biotypes = biotypes,
    max.width = max_width,
    process_n = process_n,
    verbose = FALSE
  )

  obj[[assay_out]] <- Seurat::CreateAssayObject(counts = ga_counts)

  if (isTRUE(do_normalize)) {
    obj[[assay_out]] <- Seurat::NormalizeData(
      obj[[assay_out]],
      normalization.method = "LogNormalize",
      scale.factor = 1e4,
      verbose = FALSE
    )
  }

  .msg("[GA] Done. ", assay_out, " dim=", paste(dim(obj[[assay_out]]), collapse = "x"))
  obj
}


# %% [code]
stage_fragments_tmp_user <- function(frag_src, verbose = TRUE) {
  .msg <- function(...) if (isTRUE(verbose)) cat(..., "\n")
  if (!file.exists(frag_src)) stop("frag_src not found: ", frag_src, call. = FALSE)

  user <- Sys.getenv("USER")
  tmp_root <- file.path("/tmp", user, "frags")
  dir.create(tmp_root, recursive = TRUE, showWarnings = FALSE)

  frag_dst <- file.path(tmp_root, basename(frag_src))
  if (!file.exists(frag_dst)) {
    .msg("[stage] copying fragments to node scratch:")
    .msg("  src:  ", frag_src)
    .msg("  dst:  ", frag_dst)
    ok <- file.copy(frag_src, frag_dst, overwrite = FALSE)
    if (!isTRUE(ok)) stop("Failed to copy fragments to ", frag_dst, call. = FALSE)
  } else {
    .msg("[stage] reusing node-local fragments: ", frag_dst)
  }

  # copy index if present
  tbi_src1 <- paste0(frag_src, ".tbi")
  tbi_src2 <- sub("\\.gz$", ".tbi", frag_src)
  tbi_dst1 <- paste0(frag_dst, ".tbi")
  tbi_dst2 <- sub("\\.gz$", ".tbi", frag_dst)

  if (file.exists(tbi_src1) && !file.exists(tbi_dst1)) file.copy(tbi_src1, tbi_dst1, overwrite = FALSE)
  if (file.exists(tbi_src2) && !file.exists(tbi_dst2)) file.copy(tbi_src2, tbi_dst2, overwrite = FALSE)

  if (!(file.exists(tbi_dst1) || file.exists(tbi_dst2))) {
    stop("Missing .tbi index next to staged fragment: ", frag_dst,
         "\nCreate on node: tabix -p bed ", shQuote(frag_dst),
         call. = FALSE)
  }

  frag_dst
}

ensure_activity_per_seed <- function(
  obj,
  seed,
  frag_src,
  assay_atac = "ATAC",
  assay_out  = "ACTIVITY",
  features   = NULL,
  n_features_default = 4000,
  extend_upstream = 2000,
  extend_downstream = 0,
  process_n = 8000,
  max_width = 5e5,
  biotypes = "protein_coding",
  do_normalize = FALSE,
  attach_cells = FALSE,   # set TRUE only if barcode mismatch
  verbose = TRUE
) {
  .msg <- function(...) if (isTRUE(verbose)) cat(..., "\n")

  if (!requireNamespace("Seurat", quietly = TRUE)) stop("Need Seurat.")
  if (!requireNamespace("Signac", quietly = TRUE)) stop("Need Signac.")
  if (!inherits(obj, "Seurat")) stop("obj must be Seurat.", call. = FALSE)
  if (!(assay_atac %in% names(obj@assays))) stop("missing assay_atac=", assay_atac, call. = FALSE)

  set.seed(seed)

  if (assay_out %in% names(obj@assays)) {
    .msg("[seed ", seed, "] ", assay_out, " already present -> reuse")
    return(obj)
  }

  # choose features (keep bounded)
  if (is.null(features)) {
    hv <- tryCatch(Seurat::VariableFeatures(obj, assay = "RNA"), error = function(e) character(0))
    hv <- hv[!is.na(hv) & nzchar(hv)]
    if (!length(hv)) stop("[seed ", seed, "] no RNA VariableFeatures found; pass features explicitly.", call. = FALSE)
    features <- head(unique(hv), n_features_default)
    .msg("[seed ", seed, "] using RNA HVGs n=", length(features))
  } else {
    features <- unique(features[!is.na(features) & nzchar(features)])
    if (length(features) > n_features_default) features <- head(features, n_features_default)
    .msg("[seed ", seed, "] using provided features n=", length(features))
  }

  # stage fragments to /tmp/$USER/frags
  frag_local <- stage_fragments_tmp_user(frag_src, verbose = verbose)

  # attach fragments to ATAC assay (pointing at local path)
  DefaultAssay(obj) <- assay_atac
  frags <- tryCatch(Signac::Fragments(obj[[assay_atac]]), error = function(e) NULL)
  if (is.null(frags) || length(frags) == 0) {
    .msg("[seed ", seed, "] attaching fragments: ", frag_local)
    frag_obj <- Signac::CreateFragmentObject(
      path = frag_local,
      cells = if (isTRUE(attach_cells)) Seurat::Cells(obj) else NULL,
      validate.fragments = FALSE
    )
    obj[[assay_atac]]@fragments <- list(frag_obj)
  } else {
    # overwrite path to the staged copy to avoid hitting network FS
    .msg("[seed ", seed, "] replacing fragment path with node-local copy: ", frag_local)
    frag_obj <- Signac::CreateFragmentObject(
      path = frag_local,
      cells = if (isTRUE(attach_cells)) Seurat::Cells(obj) else NULL,
      validate.fragments = FALSE
    )
    obj[[assay_atac]]@fragments <- list(frag_obj)
  }

  .msg("[seed ", seed, "] Computing gene activity…")
  ga <- Signac::GeneActivity(
    object = obj,
    assay = assay_atac,
    features = features,
    extend.upstream = extend_upstream,
    extend.downstream = extend_downstream,
    biotypes = biotypes,
    max.width = max_width,
    process_n = process_n,
    verbose = FALSE
  )

  obj[[assay_out]] <- Seurat::CreateAssayObject(counts = ga)

  if (isTRUE(do_normalize)) {
    obj[[assay_out]] <- Seurat::NormalizeData(
      obj[[assay_out]],
      normalization.method = "LogNormalize",
      scale.factor = 1e4,
      verbose = FALSE
    )
  }

  .msg("[seed ", seed, "] Done. ", assay_out, " dim=", paste(dim(obj[[assay_out]]), collapse="x"))
  obj
}

                    # ---- helpers this chunk depends on (match your "fast staged fragments" style) ----
pick_node_scratch <- function() {
  cands <- c(Sys.getenv("SLURM_TMPDIR"), Sys.getenv("TMPDIR"), file.path("/tmp", Sys.getenv("USER")))
  cands <- cands[nzchar(cands)]
  for (d in cands) {
    dir.create(d, recursive = TRUE, showWarnings = FALSE)
    ok <- tryCatch({
      tf <- tempfile(tmpdir = d)
      writeLines("ok", tf)
      unlink(tf)
      TRUE
    }, error = function(e) FALSE)
    if (isTRUE(ok)) return(d)
  }
  stop("No writable node-local scratch found (tried SLURM_TMPDIR, TMPDIR, /tmp/$USER).", call. = FALSE)
}

stage_fragments_to_node <- function(frag_src, scratch_dir = NULL, verbose = TRUE) {
  .msg <- function(...) if (isTRUE(verbose)) cat(..., "\n")
  if (!nzchar(frag_src) || !file.exists(frag_src)) stop("frag_src does not exist: ", frag_src, call. = FALSE)

  scratch_dir <- scratch_dir %||% pick_node_scratch()
  frags_dir   <- file.path(scratch_dir, "frags")
  dir.create(frags_dir, recursive = TRUE, showWarnings = FALSE)

  frag_dst <- file.path(frags_dir, basename(frag_src))
  if (!file.exists(frag_dst)) {
    .msg("[stage] copying fragments to node scratch:")
    .msg("  src: ", frag_src)
    .msg("  dst: ", frag_dst)
    ok <- file.copy(frag_src, frag_dst, overwrite = FALSE)
    if (!isTRUE(ok)) stop("Failed to copy fragments to node scratch.", call. = FALSE)
  } else {
    .msg("[stage] fragments already present -> reusing: ", frag_dst)
  }

  # stage index if present
  tbi_src1 <- paste0(frag_src, ".tbi")
  tbi_src2 <- sub("\\.gz$", ".tbi", frag_src)
  tbi_dst1 <- paste0(frag_dst, ".tbi")
  tbi_dst2 <- sub("\\.gz$", ".tbi", frag_dst)

  if (file.exists(tbi_src1) && !file.exists(tbi_dst1)) file.copy(tbi_src1, tbi_dst1, overwrite = FALSE)
  if (file.exists(tbi_src2) && !file.exists(tbi_dst2)) file.copy(tbi_src2, tbi_dst2, overwrite = FALSE)

  if (!(file.exists(tbi_dst1) || file.exists(tbi_dst2))) {
    stop(
      "Missing .tbi index next to staged fragment.\n",
      "Create it on the node shell: tabix -p bed ", shQuote(frag_dst),
      call. = FALSE
    )
  }
  frag_dst
}

assert_fragments_ok <- function(frag_path) {
  if (!file.exists(frag_path)) stop("Fragment file not found: ", frag_path, call. = FALSE)
  if (!is.finite(file.info(frag_path)$size) || file.info(frag_path)$size < 1e6) {
    warning("Fragment file looks very small (<1MB). Path might be wrong: ", frag_path, call. = FALSE)
  }
  tbi1 <- paste0(frag_path, ".tbi")
  tbi2 <- sub("\\.gz$", ".tbi", frag_path)
  if (!(file.exists(tbi1) || file.exists(tbi2))) stop("Fragment index (.tbi) not found next to: ", frag_path, call. = FALSE)
  invisible(TRUE)
}

# Attach fragments pointing to *node-local* path.
# - If fragments already attached but point elsewhere, replace.
# - Use attach_cells=TRUE for subset objects (folds) to avoid barcode mismatch.
ensure_node_local_fragments <- function(
  obj,
  frag_local,
  assay_atac = "ATAC",
  attach_cells = FALSE,
  verbose = TRUE
) {
  .msg <- function(...) if (isTRUE(verbose)) cat(..., "\n")

  if (!requireNamespace("Signac", quietly = TRUE)) stop("Need Signac.")
  if (!requireNamespace("Seurat", quietly = TRUE)) stop("Need Seurat.")
  if (!inherits(obj, "Seurat")) stop("obj must be Seurat.")
  if (!(assay_atac %in% names(obj@assays))) stop("Missing assay_atac=", assay_atac)

  if (!file.exists(frag_local)) stop("frag_local not found: ", frag_local)

  DefaultAssay(obj) <- assay_atac

  # Try to detect existing fragment object path
  frags <- tryCatch(Signac::Fragments(obj[[assay_atac]]), error = function(e) NULL)

  # Helper: get first fragment path if possible
  get_path1 <- function(frags) {
    if (is.null(frags) || length(frags) == 0) return(NA_character_)
    # Fragment objects typically store @path; but not always exposed
    p <- tryCatch(frags[[1]]@path, error = function(e) NA_character_)
    as.character(p)
  }

  cur_path <- get_path1(frags)

  if (!is.na(cur_path) && identical(normalizePath(cur_path), normalizePath(frag_local))) {
    .msg("[frags] already attached + node-local -> reuse")
    return(obj)
  }

  if (!is.na(cur_path)) {
    .msg("[frags] attached but path differs; switching to node-local: ", frag_local)
  } else {
    .msg("[frags] none attached; attaching node-local: ", frag_local)
  }

  # Recreate fragment object pointing at node-local
  frag_obj <- Signac::CreateFragmentObject(
    path = frag_local,
    cells = if (isTRUE(attach_cells)) Seurat::Cells(obj) else NULL,
    validate.fragments = FALSE
  )
  obj[[assay_atac]]@fragments <- list(frag_obj)
  obj
}

# Robust feature picker (seed or fold fallback)
pick_ga_features <- function(obj, n = 4000L, prefer_hvg = TRUE) {
  if (!requireNamespace("Seurat", quietly = TRUE)) stop("Need Seurat.", call. = FALSE)
  if (!requireNamespace("Signac", quietly = TRUE)) stop("Need Signac.", call. = FALSE)

  # 1) VariableFeatures(RNA)
  if (prefer_hvg && ("RNA" %in% names(obj@assays))) {
    hv <- tryCatch(Seurat::VariableFeatures(obj[["RNA"]]), error = function(e) character(0))
    hv <- unique(hv[!is.na(hv) & nzchar(hv)])
    if (length(hv) >= 10) return(head(hv, n))
  }

  # 2) rownames(RNA)
  if ("RNA" %in% names(obj@assays)) {
    rn <- tryCatch(rownames(obj[["RNA"]]), error = function(e) character(0))
    rn <- unique(rn[!is.na(rn) & nzchar(rn)])
    if (length(rn) >= 10) return(head(rn, n))
  }

  # 3) Annotation(ATAC)$gene_name
  if ("ATAC" %in% names(obj@assays)) {
    ann <- tryCatch(Signac::Annotation(obj[["ATAC"]]), error = function(e) NULL)
    if (!is.null(ann) && requireNamespace("S4Vectors", quietly = TRUE)) {
      if ("gene_name" %in% names(S4Vectors::mcols(ann))) {
        g <- unique(S4Vectors::mcols(ann)$gene_name)
        g <- g[!is.na(g) & nzchar(g)]
        if (length(g) >= 10) return(head(g, n))
      }
    }
  }

  character(0)
}

# Compute GA on a Seurat object that already has node-local fragments attached
compute_gene_activity_on_obj <- function(
  obj,
  assay_atac = "ATAC",
  assay_out  = "ACTIVITY",
  features,
  extend_upstream = 2000,
  extend_downstream = 0,
  process_n = 8000,
  max_width = 5e5,
  biotypes = "protein_coding",
  do_normalize = FALSE,
  verbose = TRUE,
  drop_existing = TRUE
) {
  .msg <- function(...) if (isTRUE(verbose)) cat(..., "\n")
  if (!requireNamespace("Signac", quietly = TRUE)) stop("Need Signac.")
  if (!requireNamespace("Seurat", quietly = TRUE)) stop("Need Seurat.")
  if (!inherits(obj, "Seurat")) stop("obj must be Seurat.")
  if (!(assay_atac %in% names(obj@assays))) stop("Missing assay_atac=", assay_atac)

  features <- unique(features[!is.na(features) & nzchar(features)])
  if (length(features) < 10) stop("Too few GA features: ", length(features))

  if (isTRUE(drop_existing) && assay_out %in% names(obj@assays)) {
    obj[[assay_out]] <- NULL
  }

  DefaultAssay(obj) <- assay_atac

  .msg(sprintf("[GeneActivity] cells=%d features=%d (example=%s)",
               ncol(obj), length(features), features[[1]]))

  ga_counts <- Signac::GeneActivity(
    object = obj,
    assay = assay_atac,
    features = features,
    extend.upstream = extend_upstream,
    extend.downstream = extend_downstream,
    biotypes = biotypes,
    max.width = max_width,
    process_n = process_n,
    verbose = FALSE
  )

  obj[[assay_out]] <- Seurat::CreateAssayObject(counts = ga_counts)

  if (isTRUE(do_normalize)) {
    obj[[assay_out]] <- Seurat::NormalizeData(
      obj[[assay_out]],
      normalization.method = "LogNormalize",
      scale.factor = 1e4,
      verbose = FALSE
    )
  }

  .msg(sprintf("[GeneActivity] done. %s dim=%dx%d",
               assay_out, nrow(obj[[assay_out]]), ncol(obj[[assay_out]])))
  obj
}


# ------------------------------------------------------------
# GA feature picker (fixed, vector-safe)
# ------------------------------------------------------------
pick_ga_features_fixed_n <- function(
  obj_in,
  hvgs_fold = NULL,
  n = 4000L,
  source = c("hvg_only","hvg_then_rna","rna","hvg_then_atac","atac_annot"),
  filter_to_atac_annotation = TRUE
) {
  source <- match.arg(source)
  n <- as.integer(n)
  if (!is.finite(n) || n < 1L) stop("[GA] n must be >= 1", call. = FALSE)

  # RNA genes
  rna_all <- tryCatch(rownames(obj_in[["RNA"]]), error = function(e) character(0))
  rna_all <- unique(as.character(rna_all))
  rna_all <- rna_all[!is.na(rna_all) & nzchar(rna_all)]

  # ATAC annotation genes (if available)
  ann_genes <- character(0)
  ann <- tryCatch(Signac::Annotation(obj_in[["ATAC"]]), error = function(e) NULL)
  if (!is.null(ann) && requireNamespace("S4Vectors", quietly = TRUE)) {
    mc <- tryCatch(S4Vectors::mcols(ann), error = function(e) NULL)
    if (!is.null(mc) && "gene_name" %in% names(mc)) {
      ann_genes <- unique(as.character(mc$gene_name))
      ann_genes <- ann_genes[!is.na(ann_genes) & nzchar(ann_genes)]
    }
  }

  # HVGs (fold) — IMPORTANT: never use %||% here; handle NULL explicitly
  hv <- if (is.null(hvgs_fold)) character(0) else unique(as.character(hvgs_fold))
  hv <- hv[!is.na(hv) & nzchar(hv)]

  restrict <- function(x) {
    x <- unique(as.character(x))
    x <- x[!is.na(x) & nzchar(x)]
    if (isTRUE(filter_to_atac_annotation) && length(ann_genes)) {
      x <- intersect(x, ann_genes)
    }
    unique(x)
  }

  rna_u <- restrict(rna_all)
  ann_u <- restrict(ann_genes)
  hv_u  <- restrict(hv)

  take_n <- function(x) head(x, min(length(x), n))

  feats <- switch(
    source,
    "rna"          = take_n(rna_u),
    "atac_annot"   = take_n(ann_u),
    "hvg_only"     = take_n(hv_u),
    "hvg_then_rna" = take_n(c(hv_u, setdiff(rna_u, hv_u))),
    "hvg_then_atac"= take_n(c(hv_u, setdiff(ann_u, hv_u)))
  )

  feats <- unique(feats)
  feats <- feats[!is.na(feats) & nzchar(feats)]
  feats
}


# %% [code]
# ============================================================
# Build + attach Signac-compatible gene annotation to ATAC assay
# - Requires: chrom, chromStart, chromEnd from RNA rowData
# - Adds required mcols: gene_name, gene_id, gene_biotype, type
# - Also adds tx_id/transcript_id for compatibility with older checks
# ============================================================
rd_rna <- as.data.frame(SummarizedExperiment::rowData(sce_rna))

# If mixed feature types exist, keep only gene expression rows (optional but safer)
if ("feature_types" %in% colnames(rd_rna)) {
  rd_rna <- rd_rna[rd_rna$feature_types %in% c("Gene Expression", "gene", "Gene"), , drop = FALSE]
}

need <- c("chrom", "chromStart", "chromEnd")
miss <- setdiff(need, colnames(rd_rna))
if (length(miss)) {
  stop("[annotation] RNA rowData missing: ", paste(miss, collapse = ", "))
}

# coords
g_start <- as.integer(rd_rna$chromStart)
g_end   <- as.integer(rd_rna$chromEnd)
if (any(g_start == 0L, na.rm = TRUE)) g_start <- g_start + 1L  # 0->1 based if needed

# required fields
gene_name <- if ("gene_name" %in% colnames(rd_rna)) as.character(rd_rna$gene_name) else rownames(rd_rna)

gene_id <- NULL
if ("gene_ids" %in% colnames(rd_rna)) gene_id <- as.character(rd_rna$gene_ids)
if (is.null(gene_id) && "gene_id" %in% colnames(rd_rna)) gene_id <- as.character(rd_rna$gene_id)
if (is.null(gene_id)) gene_id <- as.character(rownames(rd_rna))

gene_biotype <- NULL
if ("gene_type" %in% colnames(rd_rna)) gene_biotype <- as.character(rd_rna$gene_type)
if (is.null(gene_biotype) && "gene_biotype" %in% colnames(rd_rna)) gene_biotype <- as.character(rd_rna$gene_biotype)
if (is.null(gene_biotype)) gene_biotype <- rep("unknown", length(gene_id))

# Signac wants a 'type' column (e.g., "gene")
type <- rep("gene", length(gene_id))

# build GRanges
gr_genes <- GenomicRanges::GRanges(
  seqnames = rd_rna$chrom,
  ranges   = IRanges::IRanges(start = g_start, end = g_end)
)

# attach required metadata columns
S4Vectors::mcols(gr_genes)$gene_name    <- gene_name
S4Vectors::mcols(gr_genes)$gene_id      <- gene_id
S4Vectors::mcols(gr_genes)$gene_biotype <- gene_biotype
S4Vectors::mcols(gr_genes)$type         <- type

# compatibility cols (some Signac code paths check these)
S4Vectors::mcols(gr_genes)$tx_id         <- gene_id
S4Vectors::mcols(gr_genes)$transcript_id <- gene_id

# clean up invalid rows
ok <- !is.na(GenomicRanges::seqnames(gr_genes)) &
  !is.na(IRanges::start(gr_genes)) &
  !is.na(IRanges::end(gr_genes)) &
  IRanges::start(gr_genes) <= IRanges::end(gr_genes)
gr_genes <- gr_genes[ok]

# name the ranges (nice-to-have)
names(gr_genes) <- make.unique(as.character(S4Vectors::mcols(gr_genes)$gene_name))

# attach
Signac::Annotation(obj[["ATAC"]]) <- gr_genes
cat("[annotation] Attached to ATAC assay. n_genes =", length(gr_genes), "\n")


# %% [code]
FRAG_SRC <- "/home/groups/precepts/ashforda/UniVI_v2/UniVI_older-non_git/data/PBMC_10x_Multiome_data/10x_Genomics_Multiome_data/pbmc10k_multiome/pbmc_granulocyte_sorted_10k_atac_fragments.tsv.gz"

features_fast <- head(Seurat::VariableFeatures(obj[["RNA"]]), 5000)


# %% [code]
# ---------------------------
# Compute GeneActivity once (cache as ACTIVITY assay)
# ---------------------------
if (!requireNamespace("Signac", quietly = TRUE)) stop("Need Signac installed.")
if (!("ATAC" %in% Seurat::Assays(obj))) stop("obj has no ATAC assay")

# sanity: fragments exist on ATAC assay
frs <- Signac::Fragments(obj[["ATAC"]])
if (length(frs) == 0) stop("ATAC assay has no fragments attached (needed for GeneActivity).")

# choose features for gene activity (use RNA HVGs; compute if missing)
if (length(Seurat::VariableFeatures(obj[["RNA"]])) == 0) {
  obj <- Seurat::FindVariableFeatures(obj, assay = "RNA", nfeatures = 10000, verbose = FALSE)
}
ga_feats <- Seurat::VariableFeatures(obj[["RNA"]])

obj <- compute_gene_activity_on_obj(
  obj,
  assay_atac = "ATAC",
  assay_out  = "ACTIVITY",
  features   = ga_feats,
  process_n  = 25000,
  max_width  = 5e5,
  do_normalize = FALSE,
  verbose = TRUE
)

stopifnot("ACTIVITY" %in% Seurat::Assays(obj))
cat("[OK] assays now:", paste(Seurat::Assays(obj), collapse = ","), "\n")


# %% [markdown]
### Data preprocessing

# %% [code]
ensure_data_from_counts <- function(obj, assay = "RNA", target_sum = 1e4, clip = Inf) {
  if (!requireNamespace("SeuratObject", quietly = TRUE)) stop("Need SeuratObject.")
  if (!requireNamespace("Matrix", quietly = TRUE)) stop("Need Matrix.")

  A <- obj[[assay]]

  # pull counts (v5 layer first, fallback to slot)
  counts <- tryCatch(SeuratObject::GetAssayData(A, layer = "counts"), error = function(e) NULL)
  if (is.null(counts)) {
    counts <- tryCatch(SeuratObject::GetAssayData(A, slot = "counts"), error = function(e) NULL)
  }
  if (is.null(counts)) stop("No counts found for assay=", assay)

  if (!inherits(counts, "dgCMatrix")) counts <- as(counts, "dgCMatrix")

  cs <- Matrix::colSums(counts)
  sf <- target_sum / pmax(cs, 1e-12)

  X <- counts
  X@x <- X@x * rep.int(sf, diff(X@p))
  X@x <- log1p(X@x)
  if (is.finite(clip)) X@x <- pmin(X@x, clip)

  # write to data layer
  SeuratObject::LayerData(A, layer = "data") <- X
  obj[[assay]] <- A
  obj
}


# %% [code]
# ============================================================
# Python-matching preprocessing (train-fit / all-apply) — NO LEAKAGE
# - RNA: HVGs fit on train (after LogNormalize+log1p); apply lognorm+log1p to all HVGs;
#        fit gene-wise mu/sd on TRAIN; z-score ALL; fit SVD on TRAIN; project ALL -> PCA
# - ATAC: binarize; TF-IDF with sklearn-like smooth_idf (IDF fit on TRAIN); apply ALL;
#         fit SVD on TRAIN; project ALL -> LSI; StandardScaler fit on TRAIN; scale ALL
# Stores results as obj[["pca"]] and obj[["lsi"]]
#
# Rewrite goals:
#   - keep exact math the same
#   - avoid accidental densification / huge memory retention
#   - be robust to drop=TRUE surprises and Seurat v5 layer quirks
# ============================================================

require_or_stop(c("Matrix"))
if (!requireNamespace("irlba", quietly = TRUE)) {
  stop("[deps] Missing irlba. Install via micromamba: ", mm_hint("r-irlba"), call. = FALSE)
}
if (!requireNamespace("Seurat", quietly = TRUE)) {
  stop("[deps] Missing Seurat. Install via micromamba/CRAN.", call. = FALSE)
}

# -------------------------
# helpers
# -------------------------
.as_dgc <- function(X) {
  if (inherits(X, "dgCMatrix")) return(X)
  as(X, "dgCMatrix")
}

# scale columns of dgCMatrix (genes/peaks x cells) by sf (length ncol)
.scale_cols_sparse <- function(X_dgc, sf) {
  X_dgc <- .as_dgc(X_dgc)
  stopifnot(length(sf) == ncol(X_dgc))
  X_dgc@x <- X_dgc@x * rep.int(sf, diff(X_dgc@p))
  X_dgc
}

# LogNormalize + log1p on sparse counts (genes x cells)
.lognorm_log1p <- function(counts_dgc, target_sum = 1e4, clip = Inf) {
  counts_dgc <- .as_dgc(counts_dgc)
  cs <- Matrix::colSums(counts_dgc)
  sf <- target_sum / pmax(cs, 1e-12)
  X <- .scale_cols_sparse(counts_dgc, sf)
  X@x <- log1p(X@x)
  if (is.finite(clip)) X@x <- pmin(X@x, clip)
  X
}

# Fit HVGs on TRAIN ONLY after Seurat LogNormalize (train-only subset)
.fit_hvgs_on_train_seurat <- function(obj, train_cells, n_hvg = 2000, target_sum = 1e4) {
  Seurat::DefaultAssay(obj) <- "RNA"
  # Seurat's `[` does NOT use drop=, so do NOT pass it
  obj_tr <- obj[, train_cells]
  obj_tr <- Seurat::NormalizeData(
    obj_tr,
    normalization.method = "LogNormalize",
    scale.factor = target_sum,
    verbose = FALSE
  )
  obj_tr <- Seurat::FindVariableFeatures(obj_tr, nfeatures = n_hvg, verbose = FALSE)
  Seurat::VariableFeatures(obj_tr)
}


# gene-wise mu/sd across TRAIN cells; input is genes x train_cells
.fit_mu_sd_gene <- function(X_genes_x_cells) {
  X <- as.matrix(X_genes_x_cells)
  mu  <- rowMeans(X)
  ex2 <- rowMeans(X * X)
  sd  <- sqrt(pmax(ex2 - mu * mu, 0)) + 1e-8
  list(mu = mu, sd = sd)
}

# apply z-score gene-wise to genes x all_cells
.apply_zscore_gene <- function(X_genes_x_cells, mu, sd) {
  X <- as.matrix(X_genes_x_cells)
  X <- sweep(X, 1, mu, "-")
  X <- sweep(X, 1, sd, "/")
  X
}

# ---- TF-IDF sklearn-like (smooth_idf=TRUE, sublinear_tf=FALSE, norm=None)
# Fit on TRAIN ONLY
.tfidf_fit_sklearn_smooth <- function(bin_peaks_x_cells_train) {
  Xtr <- .as_dgc(bin_peaks_x_cells_train)
  n_tr <- ncol(Xtr)
  df_tr <- Matrix::rowSums(Xtr > 0)
  idf <- log((1 + n_tr) / (1 + df_tr)) + 1
  list(idf = idf)
}

# Apply to ALL using TRAIN-fit idf
.tfidf_apply_sklearn_smooth <- function(bin_peaks_x_cells_all, idf) {
  X <- .as_dgc(bin_peaks_x_cells_all)
  cs <- Matrix::colSums(X)
  inv_cs <- 1 / pmax(cs, 1e-12)
  Tf <- .scale_cols_sparse(X, inv_cs)
  TfIdf <- Tf
  TfIdf@x <- TfIdf@x * idf[TfIdf@i + 1L]
  TfIdf
}

# ---- SVD: fit on TRAIN ONLY, project ALL
.svd_fit_irlba <- function(cells_x_features_train, n_comp = 30, seed = 0) {
  set.seed(seed)
  s <- irlba::irlba(cells_x_features_train, nv = n_comp, nu = n_comp)
  list(V = s$v, d = s$d)
}

# IMPORTANT: do NOT multiply by d again
.svd_project <- function(cells_x_features_all, V, prefix = "dim_") {
  Z <- as.matrix(cells_x_features_all %*% V)  # already U %*% diag(d)
  colnames(Z) <- paste0(prefix, seq_len(ncol(Z)))
  Z
}

.standardize_by_train <- function(Z_all, train_idx) {
  Ztr <- Z_all[train_idx, , drop = FALSE]
  mu <- colMeans(Ztr)
  sd <- apply(Ztr, 2, stats::sd) + 1e-8
  Zs <- sweep(sweep(Z_all, 2, mu, "-"), 2, sd, "/")
  list(Z = Zs, mu = mu, sd = sd)
}

# -------------------------
# main: call this PER FOLD
# -------------------------
preprocess_py_style <- function(
  obj,
  train_idx,
  seed = 0,
  n_hvg = 2000,
  target_sum = 1e4,
  clip_rna = Inf,
  n_pca = 100,
  n_lsi = 101
) {
  stopifnot(length(train_idx) > 10)

  cells <- Seurat::Cells(obj)
  if (max(train_idx) > length(cells) || min(train_idx) < 1) {
    stop("[py-pre] train_idx out of range for fold object (n_cells=", length(cells), ").", call. = FALSE)
  }
  train_cells <- cells[train_idx]

  # ---------- RNA ----------
  Seurat::DefaultAssay(obj) <- "RNA"
  rna_counts <- .as_dgc(get_layer(obj, layer = "counts", assay = "RNA"))
  if (is.null(rna_counts) || nrow(rna_counts) == 0 || ncol(rna_counts) == 0) {
    stop("[py-pre] could not load RNA counts layer (counts).", call. = FALSE)
  }

  hvgs <- .fit_hvgs_on_train_seurat(obj, train_cells, n_hvg = n_hvg, target_sum = target_sum)
  hvgs <- intersect(hvgs, rownames(rna_counts))
  if (length(hvgs) < 200) {
    stop("[py-pre] too few HVGs after intersect: ", length(hvgs), call. = FALSE)
  }

  # lognorm+log1p applied to ALL cells for HVGs (stays sparse)
  X_rna_log <- .lognorm_log1p(rna_counts[hvgs, , drop = FALSE], target_sum = target_sum, clip = clip_rna)

  # fit mu/sd on TRAIN ONLY; apply to ALL (dense matrix of size hvgs x cells)
  ms_rna <- .fit_mu_sd_gene(X_rna_log[, train_idx, drop = FALSE])
  X_rna_z <- .apply_zscore_gene(X_rna_log, ms_rna$mu, ms_rna$sd)

  # fit SVD on TRAIN ONLY; project ALL
  fit_rna <- .svd_fit_irlba(t(X_rna_z)[train_idx, , drop = FALSE], n_comp = n_pca, seed = seed)
  Z_rna <- .svd_project(t(X_rna_z), V = fit_rna$V, prefix = "PC_")
  rownames(Z_rna) <- cells

  # aggressively free big intermediates before ATAC
  rm(X_rna_z); rm(X_rna_log); gc()

  # ---------- ATAC ----------
  Seurat::DefaultAssay(obj) <- "ATAC"
  atac_counts <- .as_dgc(get_layer(obj, layer = "counts", assay = "ATAC"))
  if (is.null(atac_counts) || nrow(atac_counts) == 0 || ncol(atac_counts) == 0) {
    stop("[py-pre] could not load ATAC counts layer (counts).", call. = FALSE)
  }

  # binarize (sparse) WITHOUT deprecated lgCMatrix casts
  atac_bin <- 1 * (atac_counts > 0)   # sparse numeric
  # keep as dgCMatrix (no need to cast to dMatrix; avoid surprises)
  atac_bin <- .as_dgc(atac_bin)

  # TF-IDF: fit IDF on TRAIN ONLY; apply to ALL
  tfidf_fit <- .tfidf_fit_sklearn_smooth(atac_bin[, train_idx, drop = FALSE])
  atac_tfidf_all <- .tfidf_apply_sklearn_smooth(atac_bin, idf = tfidf_fit$idf)

  # free binarized peaks if memory is tight
  rm(atac_bin); gc()

  # SVD/LSI: fit on TRAIN ONLY; project ALL
  fit_atac <- .svd_fit_irlba(t(atac_tfidf_all)[train_idx, , drop = FALSE], n_comp = n_lsi, seed = seed)
  Z_atac_raw <- .svd_project(t(atac_tfidf_all), V = fit_atac$V, prefix = "LSI_")
  rownames(Z_atac_raw) <- cells

  rm(atac_tfidf_all); gc()

  # StandardScaler: fit on TRAIN ONLY; apply to ALL
  scaled_atac <- .standardize_by_train(Z_atac_raw, train_idx)
  Z_atac <- scaled_atac$Z

  # ---------- attach reductions ----------
  obj[["pca"]] <- Seurat::CreateDimReducObject(
    embeddings = Z_rna,
    key = "PC_",
    assay = "RNA"
  )
  obj[["lsi"]] <- Seurat::CreateDimReducObject(
    embeddings = Z_atac,
    key = "LSI_",
    assay = "ATAC"
  )

  # Return: keep only what you actually need downstream.
  # (You can always add back rna_V/atac_V if you truly use them.)
  list(
    obj = obj,
    hvgs = hvgs,
    rna_mu = ms_rna$mu,
    rna_sd = ms_rna$sd,
    atac_idf = tfidf_fit$idf,
    atac_mu = scaled_atac$mu,
    atac_sd = scaled_atac$sd
  )
}

cat(
  "[py-pre] Loaded preprocess_py_style() (train-fit/all-apply, no leakage).\n",
  "Per fold: pp <- preprocess_py_style(obj_fold, train_idx=splits$train, seed=...)\n",
  sep = ""
)


# %% [markdown]
## Unified "runner" interface for each method

# %% [markdown]
### 1) Seurat/Signac WNN

# %% [code]
run_seurat_wnn <- function(
  obj,
  latent_dim = 30,
  rna_red = "pca",
  atac_red = "lsi",
  rna_dims = 1:100,
  atac_dims = 2:101,
  splits = NULL,
  fold_cells = NULL,   # MUST be list(train,val,test)
  verbose = TRUE,
  ...
) {
  if (!requireNamespace("Seurat", quietly = TRUE)) stop("Need Seurat installed.")

  .msg <- function(...) if (isTRUE(verbose)) { cat(..., "\n"); flush.console() }
  method_str <- "Seurat WNN"

  .skip <- function(reason, extra = list()) {
    .msg("[WNN] SKIP:", reason)
    list(Z_rna=NULL, Z_atac=NULL, Z_fused=NULL,
         extra_json=c(list(method=method_str, skipped=TRUE, reason=reason), extra))
  }

  .clamp_dims <- function(dims, n_avail) {
    dims <- as.integer(dims)
    dims <- dims[is.finite(dims)]
    dims <- dims[dims >= 1 & dims <= as.integer(n_avail)]
    dims <- unique(dims)
    if (!length(dims)) stop("[WNN] no valid dims after clamping")
    dims
  }

  .zscore_cols <- function(M) {
    # z-score each column; keep zeros if sd=0
    M <- as.matrix(M)
    mu <- colMeans(M, na.rm = TRUE)
    sdv <- apply(M, 2, stats::sd, na.rm = TRUE)
    sdv[!is.finite(sdv) | sdv == 0] <- 1
    sweep(sweep(M, 2, mu, "-"), 2, sdv, "/")
  }

  if (!is.list(fold_cells) || !all(c("train","val","test") %in% names(fold_cells))) {
    stop("[WNN] fold_cells must be a list(train,val,test)")
  }

  fold_univ <- unique(c(fold_cells$train, fold_cells$val, fold_cells$test))
  fold_univ <- intersect(fold_univ, Seurat::Cells(obj))
  if (length(fold_univ) < 50) return(.skip("too few fold cells"))

  # work on fold-universe only (safe even if obj already subset)
  if (length(fold_univ) != length(Seurat::Cells(obj))) {
    obj <- subset(obj, cells = fold_univ)
  }

  n <- length(Seurat::Cells(obj))
  .msg(sprintf("[WNN] TRANSDUCTIVE: WNN+fusion on ALL cells (n=%d)", n))
  .msg("[WNN] FindMultiModalNeighbors...")

  # WNN graph
  obj <- Seurat::FindMultiModalNeighbors(
    obj,
    reduction.list = list(rna_red, atac_red),
    dims.list = list(rna_dims, atac_dims),
    verbose = FALSE
  )

  # Pull embeddings used by WNN (and clamp dims to what's available)
  Er_all <- tryCatch(Seurat::Embeddings(obj, reduction = rna_red), error=function(e) NULL)
  Ea_all <- tryCatch(Seurat::Embeddings(obj, reduction = atac_red), error=function(e) NULL)
  if (is.null(Er_all) || is.null(Ea_all)) return(.skip("missing embeddings for rna_red/atac_red"))

  dr <- .clamp_dims(rna_dims,  ncol(Er_all))
  da <- .clamp_dims(atac_dims, ncol(Ea_all))

  Er <- Er_all[, dr, drop = FALSE]
  Ea <- Ea_all[, da, drop = FALSE]

  # align rows (cell IDs)
  common <- intersect(rownames(Er), rownames(Ea))
  if (length(common) < 50) return(.skip("too few paired cells between RNA/ATAC embeddings"))
  common <- common[order(match(common, rownames(Er)))]
  Er <- Er[common, , drop=FALSE]
  Ea <- Ea[common, , drop=FALSE]

  # ---- Mechanistic improvement: standardize each view before concatenation ----
  Ers <- .zscore_cols(Er)
  Eas <- .zscore_cols(Ea)

  X <- cbind(Ers, Eas)
  X[!is.finite(X)] <- 0

  # PCA for fused latent (deterministic; no sampling)
  pc <- stats::prcomp(X, center = TRUE, scale. = FALSE)
  K <- min(as.integer(latent_dim), ncol(pc$x))
  Zf <- pc$x[, seq_len(K), drop=FALSE]
  rownames(Zf) <- rownames(X)

  # For paired metrics, return Z_rna/Z_atac as the modality embeddings used by WNN
  Zr <- Er
  Za <- Ea

  .finalize_latents(
    Z_rna   = Zr,
    Z_atac  = Za,
    Z_fused = Zf,
    fold_cells = fold_cells,
    labels = NULL,
    splits = splits,
    method_str = method_str,
    extra_json = list(
      transductive = TRUE, uses_labels = FALSE, paired_latents = TRUE,
      rna_dims_used = dr, atac_dims_used = da,
      fused_note = "Fused latent = PCA on z-scored (PCA||LSI) embeddings"
    ),
    verbose = verbose
  )
}


# %% [markdown]
### 2) Seurat CCA

# %% [code]
run_seurat_cca_geneactivity <- function(
  obj,
  latent_dim = 30,
  nfeatures  = 2000,
  verbose    = TRUE,
  splits     = NULL,
  fold_cells = NULL,     # list(train,val,test) from CV driver
  min_split_n = 50L,
  target_sum = 1e4,
  # --- NEW (safe defaults) ---
  min_detect_frac = 0.01,      # require detection in >=1% of fit cells in BOTH modalities
  center_scale = TRUE,         # z-score features using fit-cell stats (recommended)
  eps_sd = 1e-6                # avoid div-by-zero / tiny sd
) {
  if (!requireNamespace("Seurat", quietly = TRUE)) stop("Need Seurat installed.")
  if (!requireNamespace("SeuratObject", quietly = TRUE)) stop("Need SeuratObject installed.")
  if (!requireNamespace("Matrix", quietly = TRUE)) stop("Need Matrix installed.")

  v_seurat <- as.character(packageVersion("Seurat"))
  .msg <- function(...) if (isTRUE(verbose)) cat(..., "\n")
  method_str <- paste0("Seurat CCA RNA vs GeneActivity (Seurat ", v_seurat, ")")

  .skip <- function(reason, extra = list()) {
    if (isTRUE(verbose)) cat("[CCA] SKIP:", reason, "\n")
    list(
      Z_rna = NULL, Z_atac = NULL, Z_fused = NULL,
      extra_json = c(list(method = method_str, skipped = TRUE, reason = reason), extra)
    )
  }

  if (!("RNA" %in% names(obj@assays)))      return(.skip("no RNA assay"))
  if (!("ACTIVITY" %in% names(obj@assays))) return(.skip("no ACTIVITY assay (compute/cache GeneActivity upstream)"))

  # ---- decide fold universe ----
  if (is.list(fold_cells) && all(c("train","val","test") %in% names(fold_cells))) {
    fold_cells_list <- fold_cells
    fold_cells_vec  <- unique(c(fold_cells$train, fold_cells$val, fold_cells$test))
    fold_cells_vec  <- intersect(fold_cells_vec, colnames(obj))
    if (length(fold_cells_vec) < min_split_n) return(.skip("too few fold cells"))
    ss <- list(
      fold_cells = fold_cells_vec,
      splits = splits,
      labels = NULL,
      train_cells = intersect(fold_cells$train, colnames(obj)),
      test_cells  = intersect(fold_cells$test, colnames(obj))
    )
  } else {
    ss <- .get_split_cells(obj, splits, labels = NULL, verbose = FALSE)
    fold_cells_vec <- intersect(ss$fold_cells, colnames(obj))
    fold_cells_list <- list(train = fold_cells_vec, val = character(0), test = character(0))
  }

  # ---- layer getters ----
  .get_layer <- function(assay_obj, layer) {
    out <- tryCatch(SeuratObject::GetAssayData(assay_obj, layer = layer), error = function(e) NULL)
    if (is.null(out)) return(NULL)
    if (nrow(out) == 0 || ncol(out) == 0) return(NULL)
    out
  }
  .is_nonempty <- function(M) {
    if (is.null(M)) return(FALSE)
    if (inherits(M, "dgCMatrix")) return(Matrix::nnzero(M) > 0)
    any(M != 0)
  }
  .log1p_cp <- function(counts, target_sum = 1e4) {
    if (!inherits(counts, "dgCMatrix")) counts <- Matrix::Matrix(counts, sparse = TRUE)
    cs <- Matrix::colSums(counts); cs[cs == 0] <- 1
    sf <- target_sum / cs
    out <- counts
    out@x <- out@x * rep.int(sf, diff(out@p))
    out@x <- log1p(out@x)
    out
  }
  .get_expr <- function(obj, assay_name, cells_use) {
    A <- obj[[assay_name]]
    X_data <- .get_layer(A, "data")
    if (.is_nonempty(X_data)) return(list(mat = X_data[, cells_use, drop = FALSE], source = "data"))
    X_counts <- .get_layer(A, "counts")
    if (.is_nonempty(X_counts)) {
      Xc <- X_counts[, cells_use, drop = FALSE]
      return(list(mat = .log1p_cp(Xc, target_sum = target_sum), source = "log1p_cp10k_from_counts"))
    }
    X_scale <- .get_layer(A, "scale.data")
    if (.is_nonempty(X_scale)) return(list(mat = X_scale[, cells_use, drop = FALSE], source = "scale.data"))
    list(mat = NULL, source = NA_character_)
  }

  xr0 <- .get_expr(obj, "RNA",      fold_cells_vec)
  xa0 <- .get_expr(obj, "ACTIVITY", fold_cells_vec)
  if (is.null(xr0$mat)) return(.skip("could not access usable RNA layer"))
  if (is.null(xa0$mat)) return(.skip("could not access usable ACTIVITY layer"))

  Xr <- xr0$mat
  Xa <- xa0$mat
  common_cells <- intersect(colnames(Xr), colnames(Xa))
  common_cells <- intersect(fold_cells_vec, common_cells)
  if (length(common_cells) < min_split_n) return(.skip(paste0("too few paired cells: ", length(common_cells))))

  Xr <- Xr[, common_cells, drop=FALSE]
  Xa <- Xa[, common_cells, drop=FALSE]

  # Fit on TRAIN if meaningful, else fold
  train_cells <- intersect(ss$train_cells %||% common_cells, common_cells)
  test_cells  <- intersect(ss$test_cells  %||% character(0), common_cells)
  use_inductive <- length(train_cells) >= min_split_n && length(test_cells) >= min_split_n
  fit_cells <- if (use_inductive) train_cells else common_cells

  # ---- helpers for sparse feature stats ----
  .row_var_sparse <- function(M) {
    # Var across columns for each row, using E[x^2] - E[x]^2
    mu  <- Matrix::rowMeans(M)
    mu2 <- Matrix::rowMeans(M * M)
    vv  <- pmax(mu2 - mu * mu, 0)
    as.numeric(vv)
  }
  .row_detect_frac_sparse <- function(M) {
    # fraction of columns where value != 0 (works for sparse + dense)
    if (inherits(M, "dgCMatrix")) {
      # rowSums(M != 0) is ok; keep it reasonably efficient
      as.numeric(Matrix::rowSums(M != 0)) / ncol(M)
    } else {
      rowMeans(M != 0)
    }
  }

  # features
  hvgs_rna <- tryCatch(Seurat::VariableFeatures(obj[["RNA"]]),      error = function(e) character(0))
  hvgs_act <- tryCatch(Seurat::VariableFeatures(obj[["ACTIVITY"]]), error = function(e) character(0))
  feats <- intersect(hvgs_rna, hvgs_act)
  feats <- intersect(feats, intersect(rownames(Xr), rownames(Xa)))

  # If HVG overlap is weak, pick top-variable common genes using BOTH modalities + detection filter
  if (length(feats) < 200) {
    common_genes <- intersect(rownames(Xr), rownames(Xa))
    if (length(common_genes) < 200) return(.skip(paste0("too few common genes: ", length(common_genes))))

    Xr_fit0 <- Xr[common_genes, fit_cells, drop=FALSE]
    Xa_fit0 <- Xa[common_genes, fit_cells, drop=FALSE]

    # detection filter in BOTH
    dr <- .row_detect_frac_sparse(Xr_fit0)
    da <- .row_detect_frac_sparse(Xa_fit0)
    keep <- which(dr >= min_detect_frac & da >= min_detect_frac)
    if (length(keep) < 200) {
      # don’t hard-fail if dataset is extremely sparse; relax gently by keeping what we have
      keep <- which((dr >= min_detect_frac) & (da > 0))  # slight relaxation
    }
    if (length(keep) < 50) return(.skip(paste0("too few detectable common genes after filter: ", length(keep))))

    common2 <- common_genes[keep]
    Xr_fit <- Xr[common2, fit_cells, drop=FALSE]
    Xa_fit <- Xa[common2, fit_cells, drop=FALSE]

    vr <- .row_var_sparse(Xr_fit)
    va <- .row_var_sparse(Xa_fit)
    v  <- (vr + va) / 2

    ord <- order(v, decreasing = TRUE)
    feats <- common2[ord][seq_len(min(as.integer(nfeatures), length(ord)))]
    .msg("[CCA] HVG intersection small; using top-variable common genes (bimodal): ", length(feats),
         " | min_detect_frac=", min_detect_frac)
  }

  feats <- intersect(feats, intersect(rownames(Xr), rownames(Xa)))
  if (length(feats) < 50) return(.skip(paste0("too few usable features: ", length(feats))))

  # ---- build FIT matrices ----
  .scrub <- function(M) { M[!is.finite(M)] <- 0; M }

  # Start sparse -> dense only at the last moment (cancor needs dense)
  Xr_fit_mat <- t(as.matrix(Xr[feats, fit_cells, drop=FALSE]))
  Xa_fit_mat <- t(as.matrix(Xa[feats, fit_cells, drop=FALSE]))
  Xr_fit_mat <- .scrub(Xr_fit_mat)
  Xa_fit_mat <- .scrub(Xa_fit_mat)

  if (nrow(Xr_fit_mat) < min_split_n) return(.skip("too few FIT cells"))

  # Drop near-zero variance features (either modality) to improve conditioning
  vr_fit <- apply(Xr_fit_mat, 2, stats::var)
  va_fit <- apply(Xa_fit_mat, 2, stats::var)
  keepj <- which(is.finite(vr_fit) & is.finite(va_fit) & (vr_fit > 0) & (va_fit > 0))
  if (length(keepj) < 50) return(.skip(paste0("too few nonzero-variance features after filter: ", length(keepj))))
  if (length(keepj) < ncol(Xr_fit_mat)) {
    Xr_fit_mat <- Xr_fit_mat[, keepj, drop=FALSE]
    Xa_fit_mat <- Xa_fit_mat[, keepj, drop=FALSE]
    feats_used <- feats[keepj]
  } else {
    feats_used <- feats
  }

  # Optional: z-score each feature using FIT stats, applied consistently to FIT + PROJ
  if (isTRUE(center_scale)) {
    mr <- colMeans(Xr_fit_mat); sr <- apply(Xr_fit_mat, 2, stats::sd); sr[sr < eps_sd | !is.finite(sr)] <- 1
    ma <- colMeans(Xa_fit_mat); sa <- apply(Xa_fit_mat, 2, stats::sd); sa[sa < eps_sd | !is.finite(sa)] <- 1

    Xr_fit_mat <- sweep(Xr_fit_mat, 2, mr, "-"); Xr_fit_mat <- sweep(Xr_fit_mat, 2, sr, "/")
    Xa_fit_mat <- sweep(Xa_fit_mat, 2, ma, "-"); Xa_fit_mat <- sweep(Xa_fit_mat, 2, sa, "/")
  } else {
    mr <- sr <- ma <- sa <- NULL
  }

  .msg("[CCA] cancor() fit on n_cells=", nrow(Xr_fit_mat), " feats=", ncol(Xr_fit_mat),
       if (use_inductive) " [INDUCTIVE]" else " [TRANSDUCTIVE]")

  # Capture warnings (so you can record them)
  cca_warn <- character(0)
  cc <- withCallingHandlers(
    tryCatch(stats::cancor(Xr_fit_mat, Xa_fit_mat), error = function(e) e),
    warning = function(w) {
      cca_warn <<- c(cca_warn, conditionMessage(w))
      invokeRestart("muffleWarning")
    }
  )
  if (inherits(cc, "error")) return(.skip(paste0("cancor failed: ", conditionMessage(cc))))

  K <- min(as.integer(latent_dim), ncol(cc$xcoef), ncol(cc$ycoef))
  if (!is.finite(K) || K < 2) return(.skip("cancor returned <2 usable dims"))

  # ---- projection on ALL common_cells using same feats_used and same scaling ----
  Xr_proj <- t(as.matrix(Xr[feats_used, common_cells, drop=FALSE])); Xr_proj <- .scrub(Xr_proj)
  Xa_proj <- t(as.matrix(Xa[feats_used, common_cells, drop=FALSE])); Xa_proj <- .scrub(Xa_proj)

  if (isTRUE(center_scale)) {
    Xr_proj <- sweep(Xr_proj, 2, mr[match(feats_used, feats_used)], "-")
    Xr_proj <- sweep(Xr_proj, 2, sr[match(feats_used, feats_used)], "/")
    Xa_proj <- sweep(Xa_proj, 2, ma[match(feats_used, feats_used)], "-")
    Xa_proj <- sweep(Xa_proj, 2, sa[match(feats_used, feats_used)], "/")
  }

  Zr0 <- Xr_proj %*% cc$xcoef[, seq_len(K), drop=FALSE]
  Za0 <- Xa_proj %*% cc$ycoef[, seq_len(K), drop=FALSE]
  rownames(Zr0) <- common_cells; rownames(Za0) <- common_cells
  colnames(Zr0) <- paste0("CC", seq_len(K)); colnames(Za0) <- paste0("CC", seq_len(K))

  al <- .align_paired_latents(Zr0, Za0, min_common = min_split_n, enforce_order = TRUE)
  if (is.null(al$Zr)) return(.skip(paste0("paired latent alignment failed: ", al$reason)))

  Zr_lat <- al$Zr
  Za_lat <- al$Za
  Zf <- .l2norm_rows((.l2norm_rows(Zr_lat) + .l2norm_rows(Za_lat)) / 2)

  .finalize_latents(
    Z_rna = Zr_lat,
    Z_atac = Za_lat,
    Z_fused = Zf,
    fold_cells = fold_cells_list,
    labels = NULL,
    splits = splits,
    method_str = method_str,
    extra_json = list(
      transductive = !use_inductive,
      uses_labels = FALSE,
      paired_latents = TRUE,
      sources = list(rna = xr0$source, activity = xa0$source),
      n_features_used = length(feats_used),
      fit_n_cells = nrow(Xr_fit_mat),
      proj_n_cells = nrow(Zf),
      center_scale = isTRUE(center_scale),
      min_detect_frac = min_detect_frac,
      cca_warn_n = length(cca_warn),
      cca_warn = if (length(cca_warn)) unique(cca_warn) else NULL
    ),
    verbose = verbose
  )
}


# %% [markdown]
### 3) Seurat v5 bridge integration

# %% [code]
.manual_prepare_bridge_reference <- function(
  reference,
  bridge,
  reference.reduction = "pca",
  reference.dims      = 1:50,
  normalization.method = "LogNormalize",
  reference.assay = "RNA",
  bridge.ref.assay = "RNA",
  bridge.query.assay = NULL,                 # unused here; kept for compatibility
  supervised.reduction = NULL,               # unused here; kept for compatibility
  bridge.query.reduction = NULL,             # unused here; kept for compatibility
  bridge.query.features = NULL,              # unused here; kept for compatibility
  laplacian.reduction.dims = 1:50,           # unused here; kept for compatibility
  k_anchor_try = c(50L, 30L, 20L, 10L, 5L),
  k_score_try  = c(50L, 30L, 20L),
  k_filter_try = list(NA, 200L, 100L, 50L),
  k_weight_try = c(20L, 10L, 5L, 2L, 1L),
  lap_dims_try = list(1:50),
  verbose = TRUE
) {
  if (!requireNamespace("Seurat", quietly = TRUE)) stop("Need Seurat installed.")
  if (!requireNamespace("SeuratObject", quietly = TRUE)) stop("Need SeuratObject installed.")

  .msg(sprintf("[BridgePrep] starting manual prep: ref=%d cells, bridge=%d cells",
               ncol(reference), ncol(bridge)), verbose = verbose)

  # ---- Ensure RNA is sane on both ----
  Seurat::DefaultAssay(reference) <- reference.assay
  Seurat::DefaultAssay(bridge)    <- bridge.ref.assay

  feats_ref <- rownames(.get_assay_mat(reference, reference.assay, "counts", verbose = verbose))
  feats_br  <- rownames(.get_assay_mat(bridge,    bridge.ref.assay, "counts", verbose = verbose))
  feats     <- intersect(feats_ref, feats_br)
  .assert(length(feats) >= 50,
          "[BridgePrep] too few shared RNA genes between reference and bridge (n=%d)",
          length(feats), verbose = verbose)

  # Prefer reference VariableFeatures if present
  vf <- tryCatch(SeuratObject::VariableFeatures(reference), error = function(e) character(0))
  vf <- intersect(vf, feats)
  if (length(vf) < 50) vf <- feats

  reference <- .ensure_rna_ready(reference, rna_assay = reference.assay, feats = vf, verbose = verbose)
  bridge    <- .ensure_rna_ready(bridge,    rna_assay = bridge.ref.assay, feats = vf, verbose = verbose)

  reference <- .harden_assay_layers(reference, reference.assay, verbose = verbose, create_data_features = vf)
  bridge    <- .harden_assay_layers(bridge,    bridge.ref.assay, verbose = verbose, create_data_features = vf)

  reference <- .set_default_layer(reference, reference.assay, "counts", verbose = verbose)
  bridge    <- .set_default_layer(bridge,    bridge.ref.assay, "counts", verbose = verbose)

  # ---- Make sure reference reduction exists ----
  reds_ref <- tryCatch(SeuratObject::Reductions(reference), error = function(e) character(0))
  .assert(reference.reduction %in% reds_ref,
          "[BridgePrep] reference reduction '%s' not found (have: %s)",
          reference.reduction, paste(reds_ref, collapse=", "), verbose = verbose)

  Eref <- Seurat::Embeddings(reference, reference.reduction)
  dims_use <- reference.dims[reference.dims >= 1 & reference.dims <= ncol(Eref)]
  if (length(dims_use) < 2) dims_use <- seq_len(min(50L, ncol(Eref)))
  .assert(length(dims_use) >= 2, "[BridgePrep] no usable dims for anchors", verbose = verbose)

  # ---- Find anchors (reference -> bridge) with retries ----
  anchors <- NULL
  last_err <- NULL

  for (k_anchor in as.integer(k_anchor_try)) {
    for (k_score in as.integer(k_score_try)) {
      for (k_filter in k_filter_try) {
        anchors_try <- tryCatch(
          Seurat::FindTransferAnchors(
            reference = reference,
            query = bridge,
            reference.assay = reference.assay,
            query.assay = bridge.ref.assay,
            reference.reduction = reference.reduction,
            normalization.method = normalization.method,
            dims = dims_use,
            features = vf,
            k.anchor = k_anchor,
            k.score  = k_score,
            k.filter = k_filter,
            recompute.residuals = TRUE,
            verbose = isTRUE(verbose)
          ),
          error = function(e) e
        )
        if (!inherits(anchors_try, "error")) {
          anchors <- anchors_try
          break
        } else {
          last_err <- anchors_try
        }
      }
      if (!is.null(anchors)) break
    }
    if (!is.null(anchors)) break
  }

  if (is.null(anchors)) {
    stop(sprintf("[BridgePrep] FindTransferAnchors(reference->bridge) failed: %s",
                 conditionMessage(last_err)), call. = FALSE)
  }

  # ---- Map bridge onto reference (debug only) ----
  mapped_bridge <- .map_query_robust(
    anchorset = anchors,
    query = bridge,
    reference = reference,
    k_weight_try = k_weight_try,
    store.weights = FALSE,
    verbose = verbose
  )
  if (inherits(mapped_bridge, "error")) {
    stop(sprintf("[BridgePrep] MapQuery(reference->bridge) failed: %s",
                 conditionMessage(mapped_bridge)), call. = FALSE)
  }

  # ---- Return "prep" ----
  prep <- reference
  prep@misc$bridge_mapped <- mapped_bridge

  Seurat::DefaultAssay(prep) <- reference.assay
  prep <- .harden_assay_layers(prep, reference.assay, verbose = verbose, create_data_features = vf)
  prep <- .set_default_layer(prep, reference.assay, "counts", verbose = verbose)

  # IMPORTANT FOR STABILITY: ensure the reference reduction is still present on prep
  # (Some Seurat operations can drop reductions unexpectedly.)
  reds_prep <- tryCatch(SeuratObject::Reductions(prep), error = function(e) character(0))
  .assert(reference.reduction %in% reds_prep,
          "[BridgePrep] prep lost reduction '%s' unexpectedly", reference.reduction,
          verbose = verbose)

  .msg("[BridgePrep] OK: returning prep (reference) with bridge_mapped stored in prep@misc$bridge_mapped",
       verbose = verbose)
  prep
}


# %% [code]
# ============================================================
# Seurat v5 Bridge — FULL CHUNK (REWRITE)
#
# Fixes the NEW failure you hit:
#   Cannot ensure canonical layer 'data' for assay=RNA ... layers=counts
#
# Why it happens:
#   In Seurat v5/Assay5, CreateSeuratObject() often starts with ONLY a "counts"
#   layer. Depending on Seurat/SeuratObject versions + defaults, NormalizeData()
#   may not create a literal layer named exactly "data" (or may stash it in a
#   non-layer path). Your previous .ensure_layer() required the "data" layer to
#   already exist, so it hard-failed.
#
# What this rewrite changes:
#   .ensure_layer() can now *create* a missing "data" layer (for RNA) by
#    computing log1p-normalized data from counts (layer-safe).
#   .ensure_rna_ready() no longer assumes NormalizeData will create "data";
#    it falls back to manual lognorm reliably.
#   Removes the accidental in-function redefinition of .make_anchor_safe_rna_only
#    (you already defined it globally).
#
# Everything else (TFIDF/LSI, MapQuery robustness, anchor-safe DietSeurat) stays
# aligned with your design.
# ============================================================

`%||%` <- function(x, y) if (!is.null(x)) x else y

# ---------------------------
# logging / assertions
# ---------------------------
.msg <- function(..., verbose = TRUE) {
  if (isTRUE(verbose)) { cat(..., "\n"); flush.console() }
}
.stopf <- function(fmt, ..., obj = NULL, verbose = TRUE) {
  msg <- sprintf(fmt, ...)
  .msg("[BridgeDBG] ERROR:", msg, verbose = verbose)
  if (!is.null(obj)) { .msg("[BridgeDBG] str(obj):", verbose = verbose); try(str(obj), silent = TRUE) }
  stop(msg, call. = FALSE)
}
.assert <- function(cond, fmt, ..., obj = NULL, verbose = TRUE) {
  if (!isTRUE(cond)) .stopf(fmt, ..., obj = obj, verbose = verbose)
  invisible(TRUE)
}
.dbg_mat <- function(x, name = "x", verbose = TRUE, nnz = TRUE) {
  if (!isTRUE(verbose)) return(invisible(TRUE))
  d <- tryCatch(dim(x), error = function(e) NULL)
  cls <- paste(class(x), collapse="|")
  dn_r <- tryCatch(!is.null(rownames(x)), error=function(e) FALSE)
  dn_c <- tryCatch(!is.null(colnames(x)), error=function(e) FALSE)
  extra <- ""
  if (isTRUE(nnz) && inherits(x, "dgCMatrix")) extra <- sprintf(" nnz=%d", length(x@x))
  .msg(sprintf("[BridgeDBG] %s: class=%s dim=%s rownames=%s colnames=%s%s",
               name, cls,
               if (is.null(d)) "NULL" else paste(d, collapse="x"),
               dn_r, dn_c, extra),
       verbose = verbose)
  invisible(TRUE)
}
.is_2d_matlike <- function(x) {
  if (is.null(x)) return(FALSE)
  d <- tryCatch(dim(x), error=function(e) NULL)
  if (is.null(d) || length(d) != 2L || any(d <= 0)) return(FALSE)
  is.matrix(x) || inherits(x, "Matrix")
}
.set_dimnames_safe <- function(x, rn = NULL, cn = NULL, name = "x", verbose = TRUE) {
  d <- tryCatch(dim(x), error = function(e) NULL)
  .assert(!is.null(d) && length(d) == 2L, "%s not 2D before dimnames", name, obj = d, verbose = verbose)
  if (!is.null(rn)) {
    .assert(length(rn) == nrow(x), "%s rownames length mismatch (%d vs %d)", name, length(rn), nrow(x),
            obj = list(dim=dim(x)), verbose = verbose)
    rownames(x) <- rn
  }
  if (!is.null(cn)) {
    .assert(length(cn) == ncol(x), "%s colnames length mismatch (%d vs %d)", name, length(cn), ncol(x),
            obj = list(dim=dim(x)), verbose = verbose)
    colnames(x) <- cn
  }
  x
}
.assert_finite_matrix <- function(X, name = "X", verbose = TRUE, max_show = 5) {
  .assert(!is.null(X), "%s is NULL", name, verbose = verbose)
  .assert(is.matrix(X) || inherits(X, "Matrix"),
          "%s not matrix-like (class=%s)", name, paste(class(X), collapse="|"),
          obj = class(X), verbose = verbose)
  X2 <- if (inherits(X, "Matrix")) as.matrix(X) else X
  .assert(length(dim(X2)) == 2L, "%s not 2D (dim=%s)", name, paste(dim(X2), collapse="x"),
          obj = dim(X2), verbose = verbose)
  bad <- !is.finite(X2)
  if (any(bad)) {
    idx <- which(bad, arr.ind = TRUE)
    idx <- idx[seq_len(min(nrow(idx), max_show)), , drop = FALSE]
    .stopf("%s contains NA/Inf (showing up to %d bad indices).",
           name, max_show,
           obj = list(dim = dim(X2), n_bad = sum(bad), first_bad_rc = idx),
           verbose = verbose)
  }
  invisible(TRUE)
}

# ============================================================
# SeuratObject v5 layer-safe access
# ============================================================
.SO_version <- function() {
  if (!requireNamespace("SeuratObject", quietly = TRUE)) return(package_version("0.0.0"))
  tryCatch(packageVersion("SeuratObject"), error = function(e) package_version("0.0.0"))
}
.SO_GE_5 <- function() .SO_version() >= package_version("5.0.0")

.layers_safe <- function(a) {
  if (!requireNamespace("SeuratObject", quietly = TRUE)) return(character(0))
  out <- tryCatch(SeuratObject::Layers(a), error = function(e) NULL)
  if (is.null(out)) out <- tryCatch(names(a@layers), error = function(e) character(0))
  unique(as.character(out %||% character(0)))
}

.pick_good_layer <- function(a, want) {
  want <- as.character(want)
  L <- .layers_safe(a)
  if (!length(L)) return(NULL)

  cand <- unique(c(
    want,
    L[startsWith(L, paste0(want, "."))],
    L[startsWith(L, want)],
    L[grepl(want, L, fixed = TRUE)]
  ))
  cand <- cand[cand %in% L]
  if (!length(cand)) return(NULL)

  for (lyr in cand) {
    x <- tryCatch(SeuratObject::LayerData(a, layer = lyr), error = function(e) NULL)
    if (.is_2d_matlike(x)) return(lyr)
  }
  NULL
}

.set_default_layer <- function(obj, assay, layer, verbose = TRUE) {
  a <- obj[[assay]]
  if (is.null(a)) return(obj)
  if (!.SO_GE_5()) return(obj)
  if (!exists("DefaultLayer", where = asNamespace("SeuratObject"), inherits = FALSE)) return(obj)
  ok <- tryCatch({
    SeuratObject::DefaultLayer(a) <- layer
    TRUE
  }, error = function(e) FALSE)
  if (isTRUE(ok)) {
    obj[[assay]] <- a
    .msg(sprintf("[BridgeDBG] set DefaultLayer(%s)='%s'", assay, layer), verbose = verbose)
  }
  obj
}

.get_assay_mat <- function(obj, assay, what = c("counts","data","scale.data"), verbose = TRUE) {
  what <- match.arg(what)
  if (!requireNamespace("Seurat", quietly = TRUE)) stop("Need Seurat installed.")
  if (!requireNamespace("SeuratObject", quietly = TRUE)) stop("Need SeuratObject installed.")

  a <- obj[[assay]]
  .assert(!is.null(a), "Assay '%s' not found on object", assay, verbose = verbose)

  if (.SO_GE_5()) {
    lyr <- .pick_good_layer(a, what)
    if (!is.null(lyr)) {
      out <- tryCatch(Seurat::GetAssayData(obj, assay = assay, layer = lyr), error = function(e) e)
      if (!inherits(out, "error") && .is_2d_matlike(out)) return(out)
    }
    # fallback attempt: sometimes data exists in slot-like path even if Layers() doesn't show it
    out2 <- tryCatch(Seurat::GetAssayData(obj, assay = assay, slot = what), error = function(e) NULL)
    if (.is_2d_matlike(out2)) return(out2)

    .stopf("Assay=%s has no VALID 2D matrix for '%s' (layers=%s)",
           assay, what, paste(.layers_safe(a), collapse=", "),
           obj = list(assay=assay, want=what, layers=.layers_safe(a)),
           verbose = verbose)
  }

  # pre-v5
  out3 <- tryCatch(Seurat::GetAssayData(obj, assay = assay, slot = what), error = function(e) e)
  if (inherits(out3, "error")) .stopf("GetAssayData(slot=) failed for assay=%s slot=%s: %s",
                                      assay, what, conditionMessage(out3), verbose = verbose)
  .assert(.is_2d_matlike(out3),
          "GetAssayData(slot=) returned non-2D for assay=%s slot=%s (class=%s)",
          assay, what, paste(class(out3), collapse="|"),
          obj = out3, verbose = verbose)
  out3
}

.set_assay_mat <- function(obj, assay, what = c("counts","data","scale.data"), mat, verbose = TRUE) {
  what <- match.arg(what)
  if (!requireNamespace("SeuratObject", quietly = TRUE)) stop("Need SeuratObject installed.")
  if (!requireNamespace("Seurat", quietly = TRUE)) stop("Need Seurat installed.")
  .assert(.is_2d_matlike(mat),
          "Refusing to write non-2D object into assay=%s layer=%s (class=%s)",
          assay, what, paste(class(mat), collapse="|"), obj = mat, verbose = verbose)

  a <- obj[[assay]]
  .assert(!is.null(a), "Assay '%s' not found for write", assay, verbose = verbose)

  if (.SO_GE_5()) {
    ok <- FALSE
    tryCatch({
      SeuratObject::LayerData(a, layer = what) <- mat
      obj[[assay]] <- a
      ok <- TRUE
    }, error = function(e) {
      tryCatch({
        a@layers[[what]] <- mat
        obj[[assay]] <- a
        ok <- TRUE
      }, error = function(e2) ok <<- FALSE)
    })
    .assert(ok, "Failed to write layer '%s' for assay=%s", what, assay, verbose = verbose)
    return(obj)
  }

  out <- tryCatch(Seurat::SetAssayData(obj, assay = assay, slot = what, new.data = mat), error = function(e) e)
  if (inherits(out, "error")) .stopf("SetAssayData(slot=) failed for assay=%s slot=%s: %s",
                                     assay, what, conditionMessage(out), verbose = verbose)
  out
}

# ============================================================
# NEW: reliable manual LogNormalize -> "data" layer creator
# ============================================================
.normalize_log1p_to_data <- function(obj, assay, features = NULL, scale_factor = 1e4, verbose = TRUE) {
  if (!requireNamespace("Matrix", quietly = TRUE)) stop("Need Matrix installed.")

  cnt <- .get_assay_mat(obj, assay, "counts", verbose = verbose)
  if (!inherits(cnt, "dgCMatrix")) cnt <- Matrix::Matrix(cnt, sparse = TRUE)

  feats_all <- rownames(cnt)
  if (is.null(features)) features <- feats_all
  features <- unique(as.character(features)); features <- features[nzchar(features)]
  features <- intersect(features, feats_all)
  .assert(length(features) >= 2L, "normalize: <2 features after clamp", obj = length(features), verbose = verbose)

  cnt2 <- cnt[features, , drop = FALSE]
  cs <- Matrix::colSums(cnt2); cs[cs <= 0] <- 1
  norm <- cnt2 %*% Matrix::Diagonal(x = as.numeric(scale_factor / cs))
  norm@x <- log1p(norm@x)
  norm <- .set_dimnames_safe(norm, rn = features, cn = colnames(cnt2), name = sprintf("data[%s]", assay), verbose = verbose)

  obj <- .set_assay_mat(obj, assay, "data", norm, verbose = verbose)
  obj
}

# ============================================================
# UPDATED: ensure_layer() can CREATE missing "data" for RNA
# ============================================================
.ensure_layer <- function(obj, assay, what = c("counts","data","scale.data"),
                          fallback = what, verbose = TRUE,
                          force_sparse = NULL,
                          allow_create_data = TRUE,
                          create_data_features = NULL) {
  what <- match.arg(what)
  if (!requireNamespace("SeuratObject", quietly = TRUE)) stop("Need SeuratObject installed.")
  if (!requireNamespace("Seurat", quietly = TRUE)) stop("Need Seurat installed.")
  if (!requireNamespace("Matrix", quietly = TRUE)) stop("Need Matrix installed.")

  a <- obj[[assay]]
  .assert(!is.null(a), "Assay '%s' missing", assay, verbose = verbose)

  if (is.null(force_sparse)) force_sparse <- (what %in% c("counts","data"))

  if (.SO_GE_5()) {
    L <- .layers_safe(a)

    # already present + valid?
    if (what %in% L) {
      x0 <- tryCatch(SeuratObject::LayerData(a, layer = what), error = function(e) NULL)
      if (.is_2d_matlike(x0)) {
        if (isTRUE(force_sparse) && !inherits(x0, "dgCMatrix") && what %in% c("counts","data")) {
          x0s <- Matrix::Matrix(x0, sparse = TRUE)
          x0s <- .set_dimnames_safe(x0s, rn = rownames(x0), cn = colnames(x0), name=paste0(what,"[",assay,"]"), verbose = verbose)
          obj <- .set_assay_mat(obj, assay, what, x0s, verbose = verbose)
        }
        return(obj)
      }
    }

    # special: if "data" missing, create it deterministically from counts
    if (identical(what, "data") && isTRUE(allow_create_data)) {
      .msg(sprintf("[BridgeDBG] layer '%s' missing for assay=%s; creating via manual lognorm", what, assay),
           verbose = verbose)
      # create data from counts
      obj <- .normalize_log1p_to_data(
        obj, assay = assay,
        features = create_data_features %||% rownames(.get_assay_mat(obj, assay, "counts", verbose = verbose)),
        verbose = verbose
      )
      # optional coercion
      x_new <- .get_assay_mat(obj, assay, "data", verbose = verbose)
      if (isTRUE(force_sparse) && !inherits(x_new, "dgCMatrix")) {
        x_new <- Matrix::Matrix(x_new, sparse = TRUE)
        x_new <- .set_dimnames_safe(x_new, rn = rownames(x_new), cn = colnames(x_new), name=paste0("data[",assay,"]"), verbose = verbose)
        obj <- .set_assay_mat(obj, assay, "data", x_new, verbose = verbose)
      }
      return(obj)
    }

    # otherwise: copy from some existing "like" layer if possible
    src <- .pick_good_layer(a, what)
    if (is.null(src)) src <- .pick_good_layer(a, fallback)

    .assert(!is.null(src),
            "Cannot ensure canonical layer '%s' for assay=%s (fallback=%s). layers=%s",
            what, assay, fallback, paste(L, collapse=", "),
            obj = list(assay=assay, want=what, layers=L),
            verbose = verbose)

    m <- tryCatch(SeuratObject::LayerData(a, layer = src), error = function(e) e)
    if (inherits(m, "error") || !.is_2d_matlike(m)) {
      .stopf("Source layer '%s' for assay=%s is not a valid 2D matrix-like (class=%s)",
             src, assay, paste(class(m), collapse="|"), obj = m, verbose = verbose)
    }
    if (isTRUE(force_sparse) && !inherits(m, "dgCMatrix") && what %in% c("counts","data")) {
      m <- Matrix::Matrix(m, sparse = TRUE)
      m <- .set_dimnames_safe(m, rn = rownames(m), cn = colnames(m), name=paste0(src,"[",assay,"]"), verbose = verbose)
    }
    obj <- .set_assay_mat(obj, assay, what, m, verbose = verbose)
    .msg(sprintf("[BridgeDBG] ensured canonical layer '%s' for assay=%s (src=%s)", what, assay, src), verbose = verbose)
    return(obj)
  }

  # pre-v5: no-op
  obj
}

.harden_assay_layers <- function(obj, assay, verbose = TRUE, create_data_features = NULL) {
  if (!requireNamespace("SeuratObject", quietly = TRUE)) stop("Need SeuratObject installed.")
  if (!requireNamespace("Seurat", quietly = TRUE)) stop("Need Seurat installed.")
  if (!requireNamespace("Matrix", quietly = TRUE)) stop("Need Matrix installed.")
  if (!(assay %in% SeuratObject::Assays(obj))) return(obj)

  obj <- .ensure_layer(obj, assay, "counts", verbose = verbose, force_sparse = TRUE)
  obj <- .ensure_layer(obj, assay, "data",   verbose = verbose, force_sparse = TRUE,
                       allow_create_data = TRUE, create_data_features = create_data_features)

  obj <- .set_default_layer(obj, assay, "counts", verbose = verbose)

  cts <- .get_assay_mat(obj, assay, "counts", verbose = verbose)
  dat <- .get_assay_mat(obj, assay, "data",   verbose = verbose)
  .assert(.is_2d_matlike(cts), "counts[%s] not 2D after harden", assay, obj = cts, verbose = verbose)
  .assert(.is_2d_matlike(dat), "data[%s] not 2D after harden",   assay, obj = dat, verbose = verbose)

  obj
}

# ============================================================
# RNA prep helpers (UPDATED to never assume NormalizeData creates a "data" layer)
# ============================================================
.ensure_rna_ready <- function(obj, rna_assay, feats = NULL, verbose = TRUE) {
  if (!requireNamespace("Seurat", quietly = TRUE)) stop("Need Seurat installed.")
  if (!requireNamespace("SeuratObject", quietly = TRUE)) stop("Need SeuratObject installed.")

  .assert(rna_assay %in% SeuratObject::Assays(obj), "RNA assay '%s' not present", rna_assay, verbose = verbose)

  Seurat::DefaultAssay(obj) <- rna_assay

  # clamp feats now so harden can use them if it must create data
  cnt0 <- .get_assay_mat(obj, rna_assay, "counts", verbose = verbose)
  feats_all <- rownames(cnt0)
  if (is.null(feats)) feats <- feats_all
  feats <- unique(as.character(feats)); feats <- feats[nzchar(feats)]
  feats <- intersect(feats, feats_all)
  .assert(length(feats) >= 2L, "ensure_rna_ready: <2 feats after clamp", obj = length(feats), verbose = verbose)

  # harden counts + ensure data exists (creating data if needed)
  obj <- .harden_assay_layers(obj, rna_assay, verbose = verbose, create_data_features = feats)

  # If user wants NormalizeData semantics, we *try* it, but we don't trust it to make "data"
  # (Some v5 paths keep things in a non-layer place.)
  if (.SO_GE_5()) {
    L <- .layers_safe(obj[[rna_assay]])
    if (!("data" %in% L)) {
      .msg("[BridgeDBG] RNA 'data' layer still not visible; trying NormalizeData then re-harden", verbose = verbose)
      obj2 <- tryCatch(Seurat::NormalizeData(obj, assay = rna_assay, verbose = FALSE), error = function(e) e)
      if (!inherits(obj2, "error")) obj <- obj2
      obj <- .harden_assay_layers(obj, rna_assay, verbose = verbose, create_data_features = feats)
    }
  }

  SeuratObject::VariableFeatures(obj) <- feats
  obj <- .harden_assay_layers(obj, rna_assay, verbose = verbose, create_data_features = feats)
  obj
}

.get_counts_sparse <- function(obj, assay, cells = NULL, verbose = TRUE) {
  if (!requireNamespace("Matrix", quietly = TRUE)) stop("Need Matrix installed.")
  obj <- .harden_assay_layers(obj, assay, verbose = verbose)

  m <- .get_assay_mat(obj, assay, "counts", verbose = verbose)
  if (!inherits(m, "dgCMatrix")) m <- Matrix::Matrix(m, sparse = TRUE)
  if (!is.null(cells)) {
    cells <- intersect(unique(as.character(cells)), colnames(m))
    .assert(length(cells) > 0, "no requested cells present in counts for assay=%s", assay, verbose = verbose)
    m <- m[, cells, drop = FALSE]
  }
  .dbg_mat(m, sprintf("counts[%s]", assay), verbose = verbose)
  m
}

.manual_scale_to_scale_data <- function(obj, assay, features, verbose = TRUE) {
  if (!requireNamespace("matrixStats", quietly = TRUE)) stop("Need matrixStats installed.")
  obj <- .harden_assay_layers(obj, assay, verbose = verbose, create_data_features = features)

  X <- .get_assay_mat(obj, assay, "data", verbose = verbose)
  features <- intersect(unique(as.character(features)), rownames(X))
  .assert(length(features) >= 2L, "manual-scale: <2 features", obj = length(features), verbose = verbose)

  X <- X[features, , drop = FALSE]
  cn_cells <- colnames(X)

  .msg(sprintf("[Bridge] manual-scale: data %dx%d class=%s -> densify",
               nrow(X), ncol(X), paste(class(X), collapse="|")),
       verbose = verbose)

  Xd <- as.matrix(X)
  rm(X); invisible(gc())

  mu <- matrixStats::rowMeans2(Xd)
  sd <- matrixStats::rowSds(Xd); sd[sd < 1e-8] <- 1.0
  Xd <- (Xd - mu) / sd
  Xd <- .set_dimnames_safe(Xd, rn = features, cn = cn_cells, name = "scale.data", verbose = verbose)

  obj <- .set_assay_mat(obj, assay, "scale.data", Xd, verbose = verbose)
  obj
}

# ============================================================
# TFIDF/LSI + MapQuery robust (unchanged)
# ============================================================
.tfidf_from_counts_sparse <- function(counts, verbose = TRUE) {
  if (!requireNamespace("Matrix", quietly = TRUE)) stop("Need Matrix installed.")
  if (!inherits(counts, "dgCMatrix")) counts <- Matrix::Matrix(counts, sparse = TRUE)

  cs <- Matrix::colSums(counts); cs[cs <= 0] <- 1
  tf <- counts %*% Matrix::Diagonal(x = 1 / cs)

  df <- Matrix::rowSums(counts > 0); df[df <= 0] <- 1
  N <- ncol(counts)
  idf <- log1p(N / df)

  tfidf <- Matrix::Diagonal(x = as.numeric(idf)) %*% tf
  rownames(tfidf) <- rownames(counts)
  colnames(tfidf) <- colnames(counts)
  tfidf
}

.add_lsi_reduction_from_tfidf <- function(obj, tfidf, assay, n_components = 101L,
                                         reduction_name = "lsi", key = "LSI_", verbose = TRUE) {
  if (!requireNamespace("irlba", quietly = TRUE)) stop("Need irlba installed.")
  if (!requireNamespace("Seurat", quietly = TRUE)) stop("Need Seurat installed.")

  d <- dim(tfidf)
  min_dim <- min(d[1], d[2])
  k_req <- as.integer(n_components)
  k <- min(k_req, max(2L, min_dim - 1L))

  .msg(sprintf("[Bridge] LSI via irlba: TFIDF %dx%d | requested k=%d -> using k=%d",
               d[1], d[2], k_req, k), verbose = verbose)

  sv <- irlba::irlba(tfidf, nv = k, nu = k)
  U <- sv$u; V <- sv$v
  colnames(U) <- paste0(key, seq_len(k))
  colnames(V) <- paste0(key, seq_len(k))
  rownames(U) <- rownames(tfidf)
  rownames(V) <- colnames(tfidf)

  dr <- Seurat::CreateDimReducObject(embeddings = V, loadings = U, assay = assay, key = key)
  obj[[reduction_name]] <- dr
  list(obj = obj, loadings_u = U)
}

.project_lsi_query_from_u <- function(obj_q, tfidf_q, loadings_u, assay,
                                      reduction_name = "lsi", key = "LSI_", verbose = TRUE) {
  if (!requireNamespace("Matrix", quietly = TRUE)) stop("Need Matrix installed.")
  if (!requireNamespace("Seurat", quietly = TRUE)) stop("Need Seurat installed.")

  feats <- intersect(rownames(loadings_u), rownames(tfidf_q))
  tfidf_q2 <- tfidf_q[feats, , drop = FALSE]
  U2 <- loadings_u[feats, , drop = FALSE]
  prod <- Matrix::t(tfidf_q2) %*% as.matrix(U2)

  emb_q <- as.matrix(prod)
  colnames(emb_q) <- colnames(U2)
  rownames(emb_q) <- colnames(tfidf_q2)

  dr <- Seurat::CreateDimReducObject(embeddings = emb_q, loadings = U2, assay = assay, key = key)
  obj_q[[reduction_name]] <- dr
  obj_q
}

.map_query_robust <- function(anchorset, query, reference,
                             k_weight_try = c(20L, 10L, 5L, 2L, 1L),
                             store.weights = TRUE, verbose = TRUE) {
  if (!requireNamespace("Seurat", quietly = TRUE)) stop("Need Seurat installed.")
  out0 <- tryCatch(
    Seurat::MapQuery(anchorset = anchorset, query = query, reference = reference,
                     store.weights = store.weights, verbose = isTRUE(verbose)),
    error = function(e) e
  )
  if (!inherits(out0, "error")) return(out0)

  msg0 <- conditionMessage(out0)
  if (!grepl("k\\.weight.*smaller than the number of anchors", msg0)) return(out0)

  for (kw in as.integer(k_weight_try)) {
    out <- tryCatch(
      Seurat::MapQuery(anchorset = anchorset, query = query, reference = reference,
                       k.weight = kw, store.weights = store.weights, verbose = isTRUE(verbose)),
      error = function(e) e
    )
    if (!inherits(out, "error")) return(out)
  }
  out0
}

# ============================================================
# Anchor-safe wrapper (unchanged; keep ONE definition)
# ============================================================
.make_anchor_safe_rna_only <- function(obj, assay, keep_reductions = character(0), verbose = TRUE) {
  if (!requireNamespace("SeuratObject", quietly = TRUE)) stop("Need SeuratObject installed.")
  if (!requireNamespace("Seurat", quietly = TRUE)) stop("Need Seurat installed.")

  Seurat::DefaultAssay(obj) <- assay
  obj <- .ensure_rna_ready(obj, rna_assay = assay, feats = rownames(.get_assay_mat(obj, assay, "counts", verbose = verbose)), verbose = verbose)
  obj <- .harden_assay_layers(obj, assay, verbose = verbose)
  obj <- .set_default_layer(obj, assay, "counts", verbose = verbose)

  out <- tryCatch(
    SeuratObject::DietSeurat(
      obj,
      assays = assay,
      dimreducs = keep_reductions,
      graphs = NULL,
      misc = FALSE
    ),
    error = function(e) e
  )
  if (inherits(out, "error")) {
    .msg("[BridgeDBG] DietSeurat failed; proceeding without DietSeurat (still hardened)", verbose = verbose)
    out <- obj
  }

  Seurat::DefaultAssay(out) <- assay
  out <- .harden_assay_layers(out, assay, verbose = verbose)
  out <- .set_default_layer(out, assay, "counts", verbose = verbose)
  out
}


# %% [code]
run_seurat_bridge_v5_safe <- function(
  obj_bridge,
  fold_cells,
  hvgs = NULL,
  seed = 0,
  rna_assay = "RNA",
  atac_assay = NULL,
  ref_n = 4000,
  bridge_n = 4000,
  query_n = NULL,
  latent_dim = 30,
  reference_reduction = "pca",
  reference_dims = 1:100,
  laplacian_reduction_dims = 1:50,
  atac_top_features_n = 20000L,
  verbose = TRUE,
  pca_approx = TRUE,
  force_sequential = TRUE,
  npcs_min = 50L,
  n_hvg = 2000L,
  k_weight_try = c(20L, 10L, 5L, 2L, 1L),
  k_anchor_try = c(50L, 30L, 20L, 10L, 5L),
  k_score_try  = c(50L, 30L, 20L),
  k_filter_try = list(NA, 200L, 100L, 50L),
  lap_dims_try = list(2:101, 1:100, 2:51, 1:50, 2:31, 1:30, 2:21, 1:20),
  ...
) {
  if (!requireNamespace("Seurat", quietly = TRUE)) stop("Need Seurat installed.")
  if (!requireNamespace("SeuratObject", quietly = TRUE)) stop("Need SeuratObject installed.")
  if (!requireNamespace("Signac", quietly = TRUE)) stop("Need Signac installed.")
  if (!requireNamespace("Matrix", quietly = TRUE)) stop("Need Matrix installed.")
  if (!requireNamespace("future", quietly = TRUE)) stop("Need future installed.")

  .skip_local <- function(reason, extra = list()) {
    .msg("[Bridge] SKIP:", reason, verbose = verbose)
    list(
      Z_fused = NULL, Z_rna = NULL, Z_atac = NULL,
      extra_json = c(list(method="seurat_bridge", skipped=TRUE, reason=as.character(reason)), extra)
    )
  }

  # --- NEW: helpers for fused embedding (mechanics only) ---
  .zscore_cols <- function(M) {
    M <- as.matrix(M)
    mu <- colMeans(M, na.rm = TRUE)
    sdv <- apply(M, 2, stats::sd, na.rm = TRUE)
    sdv[!is.finite(sdv) | sdv == 0] <- 1
    sweep(sweep(M, 2, mu, "-"), 2, sdv, "/")
  }

  .fused_pca_concat <- function(Zr, Za, k) {
    # Zr/Za: same rows (cells), any #cols
    Zr2 <- .zscore_cols(Zr)
    Za2 <- .zscore_cols(Za)
    X <- cbind(Zr2, Za2)
    X[!is.finite(X)] <- 0
    pc <- stats::prcomp(X, center = TRUE, scale. = FALSE)
    K <- min(as.integer(k), ncol(pc$x))
    Zf <- pc$x[, seq_len(K), drop = FALSE]
    rownames(Zf) <- rownames(X)
    Zf
  }

  # validate fold_cells
  if (!is.list(fold_cells) || !all(c("train","val","test") %in% names(fold_cells))) {
    return(.skip_local("fold_cells must be list(train,val,test)"))
  }
  for (nm in c("train","val","test")) {
    fold_cells[[nm]] <- unique(as.character(fold_cells[[nm]]))
    fold_cells[[nm]] <- fold_cells[[nm]][nzchar(fold_cells[[nm]])]
  }

  aa <- SeuratObject::Assays(obj_bridge)
  if (!(rna_assay %in% aa)) return(.skip_local("no RNA assay"))

  # infer ATAC assay if not provided
  if (is.null(atac_assay)) {
    chrom_like <- aa[sapply(aa, function(a) inherits(obj_bridge[[a]], "ChromatinAssay"))]
    if (length(chrom_like) > 0) atac_assay <- chrom_like[[1]]
    if (is.null(atac_assay) && "ATAC" %in% aa) atac_assay <- "ATAC"
  }
  if (is.null(atac_assay) || !(atac_assay %in% aa)) return(.skip_local("no ATAC/Chromatin assay"))

  all_cells <- SeuratObject::Cells(obj_bridge)
  fold_univ <- unique(c(fold_cells$train, fold_cells$val, fold_cells$test))
  fold_univ <- intersect(fold_univ, all_cells)
  if (length(fold_univ) < 50) return(.skip_local("too few fold cells"))

  train_cells <- intersect(fold_cells$train, fold_univ)
  if (length(train_cells) < 50) train_cells <- fold_univ

  set.seed(as.integer(seed))
  ref_cells    <- if (length(train_cells) > ref_n)    sample(train_cells, ref_n)    else train_cells
  bridge_cells <- if (length(train_cells) > bridge_n) sample(train_cells, bridge_n) else train_cells
  query_cells  <- fold_univ
  if (!is.null(query_n) && length(query_cells) > query_n) query_cells <- sample(query_cells, query_n)

  .msg(sprintf("[Bridge] TRAIN=%d -> ref=%d bridge=%d | query=%d",
               length(train_cells), length(ref_cells), length(bridge_cells), length(query_cells)),
       verbose = verbose)

  # force sequential
  old_plan <- future::plan()
  on.exit({ try(future::plan(old_plan), silent = TRUE) }, add = TRUE)
  if (isTRUE(force_sequential)) future::plan("sequential")

  # extract RNA counts (layer-safe)
  .msg("[Bridge] extracting RNA counts (sparse) ...", verbose = verbose)
  rna_ref_full <- .get_counts_sparse(obj_bridge, rna_assay, ref_cells,    verbose = verbose)
  rna_br_full  <- .get_counts_sparse(obj_bridge, rna_assay, bridge_cells, verbose = verbose)
  rna_q_full   <- .get_counts_sparse(obj_bridge, rna_assay, query_cells,  verbose = verbose)

  feats <- NULL
  if (!is.null(hvgs) && length(hvgs)) feats <- unique(as.character(hvgs))
  feats <- feats %||% rownames(rna_ref_full)
  feats <- unique(as.character(feats)); feats <- feats[nzchar(feats)]
  feats <- intersect(feats, rownames(rna_ref_full))
  feats <- intersect(feats, rownames(rna_br_full))
  if (length(feats) > as.integer(n_hvg)) feats <- feats[seq_len(as.integer(n_hvg))]
  if (length(feats) < 50) return(.skip_local(paste0("no usable shared RNA features (n=", length(feats), ")")))

  rna_ref <- rna_ref_full[feats, , drop = FALSE]
  rna_br  <- rna_br_full[feats, , drop = FALSE]
  rna_q   <- rna_q_full[feats, , drop = FALSE]
  rm(rna_ref_full, rna_br_full, rna_q_full); invisible(gc())
  .msg(sprintf("[Bridge] RNA restricted: genes=%d | ref nnz=%d", length(feats), length(rna_ref@x)), verbose = verbose)

  # extract ATAC counts (layer-safe)
  .msg("[Bridge] extracting ATAC counts (sparse) ...", verbose = verbose)
  atac_br_full <- .get_counts_sparse(obj_bridge, atac_assay, bridge_cells, verbose = verbose)
  atac_q_full  <- .get_counts_sparse(obj_bridge, atac_assay, query_cells,  verbose = verbose)

  .msg(sprintf("[Bridge] ATAC bridge: peaks=%d cells=%d nnz=%d",
               nrow(atac_br_full), ncol(atac_br_full), length(atac_br_full@x)), verbose = verbose)

  .msg(sprintf("[Bridge] selecting top ATAC features (n=%d) ...", as.integer(atac_top_features_n)), verbose = verbose)
  peak_sums <- Matrix::rowSums(atac_br_full)
  ord <- order(as.numeric(peak_sums), decreasing = TRUE)
  keep_n <- min(as.integer(atac_top_features_n), length(ord))
  keep_peaks <- rownames(atac_br_full)[ord[seq_len(keep_n)]]
  keep_peaks <- intersect(keep_peaks, rownames(atac_q_full))
  if (length(keep_peaks) < 1000) return(.skip_local(paste0("too few shared ATAC peaks (n=", length(keep_peaks), ")")))

  atac_br <- atac_br_full[keep_peaks, , drop = FALSE]
  atac_q  <- atac_q_full[keep_peaks, , drop = FALSE]
  rm(atac_br_full, atac_q_full, peak_sums, ord); invisible(gc())

  # build minimal objects
  .msg("[Bridge] building minimal Seurat objects ...", verbose = verbose)
  obj_ref <- Seurat::CreateSeuratObject(counts = rna_ref, assay = rna_assay)
  obj_br  <- Seurat::CreateSeuratObject(counts = rna_br,  assay = rna_assay)
  obj_q   <- Seurat::CreateSeuratObject(counts = rna_q,   assay = rna_assay)

  obj_br[[atac_assay]] <- Signac::CreateChromatinAssay(counts = atac_br, fragments = NULL)
  obj_q[[atac_assay]]  <- Signac::CreateChromatinAssay(counts = atac_q,  fragments = NULL)

  SeuratObject::VariableFeatures(obj_ref) <- feats
  SeuratObject::VariableFeatures(obj_br)  <- feats
  SeuratObject::VariableFeatures(obj_q)   <- feats

  rm(rna_ref, rna_br, rna_q, atac_br, atac_q); invisible(gc())

  # Ensure RNA layers
  obj_ref <- .ensure_rna_ready(obj_ref, rna_assay = rna_assay, feats = feats, verbose = verbose)
  obj_br  <- .ensure_rna_ready(obj_br,  rna_assay = rna_assay, feats = feats, verbose = verbose)
  obj_q   <- .ensure_rna_ready(obj_q,   rna_assay = rna_assay, feats = feats, verbose = verbose)

  # reference PCA
  need_pcs <- max(as.integer(npcs_min), as.integer(latent_dim), max(reference_dims, na.rm = TRUE))
  .msg(sprintf("[Bridge] ref: MANUAL_SCALE + RunPCA | genes=%d cells=%d npcs=%d approx=%s",
               length(feats), ncol(obj_ref), need_pcs, if (isTRUE(pca_approx)) "TRUE" else "FALSE"),
       verbose = verbose)

  obj_ref <- .manual_scale_to_scale_data(obj_ref, assay = rna_assay, features = feats, verbose = verbose)
  obj_ref <- Seurat::RunPCA(
    obj_ref,
    assay = rna_assay,
    features = feats,
    npcs = need_pcs,
    approx = isTRUE(pca_approx),
    verbose = FALSE,
    reduction.name = reference_reduction
  )

  Eref <- tryCatch(Seurat::Embeddings(obj_ref, reduction = reference_reduction), error = function(e) NULL)
  if (is.null(Eref) || !is.matrix(Eref) || ncol(Eref) < 2) return(.skip_local("reference PCA embeddings invalid"))

  reference_dims2 <- reference_dims[reference_dims >= 1 & reference_dims <= ncol(Eref)]
  if (length(reference_dims2) < 2) return(.skip_local("reference.dims has <2 after clamp"))

  # TFIDF+LSI on bridge ATAC + project query
  .msg(sprintf("[Bridge] bridge: manual TFIDF+LSI on ATAC (peaks=%d cells=%d)", length(keep_peaks), ncol(obj_br)),
       verbose = verbose)

  obj_br <- .harden_assay_layers(obj_br, atac_assay, verbose = verbose)
  obj_q  <- .harden_assay_layers(obj_q,  atac_assay, verbose = verbose)

  atac_counts_br <- .get_counts_sparse(obj_br, atac_assay, SeuratObject::Cells(obj_br), verbose = verbose)
  tfidf_br <- .tfidf_from_counts_sparse(atac_counts_br, verbose = verbose)
  rm(atac_counts_br); invisible(gc())

  n_lsi_req <- max(as.integer(npcs_min), as.integer(latent_dim), max(unlist(lap_dims_try), na.rm = TRUE))
  lsi_out <- .add_lsi_reduction_from_tfidf(
    obj = obj_br, tfidf = tfidf_br, assay = atac_assay,
    n_components = n_lsi_req, reduction_name = "lsi", key = "LSI_", verbose = verbose
  )
  obj_br <- lsi_out$obj
  U <- lsi_out$loadings_u
  rm(lsi_out, tfidf_br); invisible(gc())

  atac_counts_q <- .get_counts_sparse(obj_q, atac_assay, SeuratObject::Cells(obj_q), verbose = verbose)
  tfidf_q <- .tfidf_from_counts_sparse(atac_counts_q, verbose = FALSE)
  rm(atac_counts_q); invisible(gc())

  obj_q <- .project_lsi_query_from_u(
    obj_q = obj_q, tfidf_q = tfidf_q, loadings_u = U,
    assay = atac_assay, reduction_name = "lsi", key = "LSI_", verbose = verbose
  )
  rm(tfidf_q, U); invisible(gc())

  # manual PrepareBridgeReference + MapQuery(reference->bridge)
  .msg("[Bridge] manual PrepareBridgeReference + MapQuery ...", verbose = verbose)
  prep <- .manual_prepare_bridge_reference(
    reference = obj_ref,
    bridge    = obj_br,
    reference.reduction = reference_reduction,
    reference.dims      = reference_dims2,
    normalization.method = "LogNormalize",
    reference.assay = rna_assay,
    bridge.ref.assay = rna_assay,
    bridge.query.assay = atac_assay,
    supervised.reduction = NULL,
    bridge.query.reduction = "lsi",
    bridge.query.features = keep_peaks,
    laplacian.reduction.dims = laplacian_reduction_dims,
    k_anchor_try = k_anchor_try,
    k_score_try  = k_score_try,
    k_filter_try = k_filter_try,
    k_weight_try = k_weight_try,
    lap_dims_try = lap_dims_try,
    verbose = verbose
  )
  if (inherits(prep, "error")) return(.skip_local(paste0("manual_prepare_bridge_reference failed: ", conditionMessage(prep))))

  # Harden prep RNA
  Seurat::DefaultAssay(prep) <- rna_assay
  prep <- .ensure_rna_ready(prep, rna_assay = rna_assay, feats = feats, verbose = verbose)
  prep <- .harden_assay_layers(prep, rna_assay, verbose = verbose, create_data_features = feats)

  # query -> prep anchors + MapQuery
  query_to_prep_map <- function(prep, obj_q, rna_assay, feats,
                                reference_reduction = "pca",
                                reference_dims2 = 1:50,
                                k_weight_try = c(20L, 10L, 5L, 2L, 1L),
                                verbose = TRUE) {

    Seurat::DefaultAssay(prep)  <- rna_assay
    Seurat::DefaultAssay(obj_q) <- rna_assay

    prep  <- .ensure_rna_ready(prep,  rna_assay = rna_assay, feats = feats, verbose = verbose)
    obj_q <- .ensure_rna_ready(obj_q, rna_assay = rna_assay, feats = feats, verbose = verbose)

    prep  <- .harden_assay_layers(prep,  rna_assay, verbose = verbose, create_data_features = feats)
    obj_q <- .harden_assay_layers(obj_q, rna_assay, verbose = verbose, create_data_features = feats)

    prep_reds <- tryCatch(SeuratObject::Reductions(prep), error = function(e) character(0))
    pref_reds <- c(paste0("ref.", reference_reduction), reference_reduction, "pca")
    red_use <- pref_reds[pref_reds %in% prep_reds]
    if (!length(red_use) && length(prep_reds)) red_use <- prep_reds
    if (!length(red_use)) stop("[Bridge] prepared object has no reductions for anchor projection")
    red_use <- red_use[[1]]

    prep_safe  <- .make_anchor_safe_rna_only(prep,  rna_assay, keep_reductions = red_use, verbose = verbose)
    query_safe <- .make_anchor_safe_rna_only(obj_q, rna_assay, keep_reductions = character(0), verbose = verbose)

    shared_q_feats <- intersect(
      rownames(.get_assay_mat(prep_safe,  rna_assay, "counts", verbose = verbose)),
      rownames(.get_assay_mat(query_safe, rna_assay, "counts", verbose = verbose))
    )
    shared_q_feats <- intersect(shared_q_feats, feats)
    if (length(shared_q_feats) < 50) stop(paste0("[Bridge] too few shared RNA feats for query->prep (n=", length(shared_q_feats), ")"))

    Ep <- Seurat::Embeddings(prep_safe, red_use)
    dims2 <- reference_dims2[reference_dims2 >= 1 & reference_dims2 <= ncol(Ep)]
    if (length(dims2) < 2) dims2 <- seq_len(min(50L, ncol(Ep)))
    if (length(dims2) < 2) stop("[Bridge] no usable dims for query->prep anchors")

    anchors_q <- Seurat::FindTransferAnchors(
      reference = prep_safe,
      query = query_safe,
      reference.assay = rna_assay,
      query.assay = rna_assay,
      reference.reduction = red_use,
      normalization.method = "LogNormalize",
      dims = dims2,
      recompute.residuals = TRUE,
      features = shared_q_feats,
      k.filter = NA,
      verbose = isTRUE(verbose)
    )

    .map_query_robust(
      anchorset = anchors_q,
      query = query_safe,
      reference = prep_safe,
      k_weight_try = k_weight_try,
      store.weights = FALSE,
      verbose = verbose
    )
  }

  mapped <- tryCatch(
    query_to_prep_map(
      prep = prep,
      obj_q = obj_q,
      rna_assay = rna_assay,
      feats = feats,
      reference_reduction = reference_reduction,
      reference_dims2 = reference_dims2,
      k_weight_try = k_weight_try,
      verbose = verbose
    ),
    error = function(e) e
  )
  if (inherits(mapped, "error")) return(.skip_local(paste0("FindTransferAnchors(query->prep) failed: ", conditionMessage(mapped))))

  # ============================================================
  # OUTPUTS (unchanged for Z_rna / Z_atac)
  #   Z_rna  = mapped query embedding in ref space (ref.pca preferred)
  #   Z_atac = query lsi
  #   Z_fused = NOW JOINT: PCA([z(Z_rna)||z(Z_atac)])
  # ============================================================
  red_avail <- tryCatch(SeuratObject::Reductions(mapped), error = function(e) character(0))
  pref_rna <- c("ref.pca", paste0("ref.", reference_reduction), reference_reduction, "pca")
  red_rna_use <- pref_rna[pref_rna %in% red_avail]
  if (!length(red_rna_use)) red_rna_use <- red_avail
  if (!length(red_rna_use)) return(.skip_local("MapQuery succeeded but no reductions present"))

  Zr <- NULL
  red_rna_chosen <- NA_character_
  for (r in red_rna_use) {
    Zcand <- tryCatch(Seurat::Embeddings(mapped, reduction = r), error = function(e) NULL)
    if (!is.null(Zcand) && is.matrix(Zcand) && nrow(Zcand) >= 10 && ncol(Zcand) >= 2) {
      Zr <- Zcand
      red_rna_chosen <- r
      break
    }
  }
  if (is.null(Zr)) return(.skip_local("no usable RNA-aligned embeddings after MapQuery"))

  Za <- tryCatch(Seurat::Embeddings(obj_q, reduction = "lsi"), error = function(e) NULL)
  if (is.null(Za) || !is.matrix(Za) || nrow(Za) < 10 || ncol(Za) < 2) {
    return(.skip_local("ATAC LSI embeddings missing/invalid on obj_q"))
  }

  qc <- as.character(query_cells)

  if (is.null(rownames(Zr)) || any(!nzchar(rownames(Zr)))) rownames(Zr) <- SeuratObject::Cells(mapped)
  if (is.null(rownames(Za)) || any(!nzchar(rownames(Za)))) rownames(Za) <- SeuratObject::Cells(obj_q)

  miss_r <- setdiff(qc, rownames(Zr))
  miss_a <- setdiff(qc, rownames(Za))
  if (length(miss_r)) stop(sprintf("[Bridge] Z_rna missing %d/%d query cells; example: %s",
                                  length(miss_r), length(qc), paste(head(miss_r, 5), collapse=", ")))
  if (length(miss_a)) stop(sprintf("[Bridge] Z_atac missing %d/%d query cells; example: %s",
                                  length(miss_a), length(qc), paste(head(miss_a, 5), collapse=", ")))

  Zr <- Zr[qc, , drop = FALSE]
  Za <- Za[qc, , drop = FALSE]

  d_use <- as.integer(latent_dim)
  d_use_final <- min(d_use, ncol(Zr), ncol(Za))
  if (!is.finite(d_use_final) || d_use_final < 2L) return(.skip_local("latent_dim clamp left <2 dims"))

  Zr <- Zr[, seq_len(d_use_final), drop = FALSE]
  Za <- Za[, seq_len(d_use_final), drop = FALSE]

  # ---- FIX A: fused = PCA on concatenated z-scored [Z_rna || Z_atac] ----
  Zf <- .fused_pca_concat(Zr, Za, k = d_use_final)
  Zf <- Zf[qc, , drop = FALSE]  # ensure exact order

  .msg(sprintf("[Bridge] OK: Z_rna=%dx%d (%s) | Z_atac=%dx%d (lsi) | Z_fused=%dx%d | query=%d",
               nrow(Zr), ncol(Zr), red_rna_chosen,
               nrow(Za), ncol(Za),
               nrow(Zf), ncol(Zf),
               length(qc)),
       verbose = verbose)

  list(
    Z_fused = Zf,
    Z_rna   = Zr,
    Z_atac  = Za,
    extra_json = list(
      method = "seurat_bridge",
      skipped = FALSE,
      reason = NA_character_,
      rna_reduction = red_rna_chosen,
      atac_reduction = "lsi",
      latent_dim = d_use_final,
      n_features = length(feats),
      n_peaks = length(keep_peaks),
      n_query = length(qc),
      seed = as.integer(seed),
      pca_approx = isTRUE(pca_approx),
      force_sequential = isTRUE(force_sequential),
      manual_scaled = TRUE,
      manual_lsi = TRUE,
      n_lsi = as.integer(n_lsi_req),
      fused_construction = "pca(concat(z(Z_rna), z(Z_atac)))",
      lsi_dim1_policy = "lap_dims_try prefers drop dim1 (2:...) first"
    )
  )
}


# %% [markdown]
### 4) Harmony

# %% [code]
run_harmony_dual <- function(
  obj,
  latent_dim = 30,
  rna_red = "pca",
  atac_red = "lsi",
  rna_dims = 1:100,
  atac_dims = 2:101,
  splits = NULL,
  fold_cells = NULL,   # MUST be list(train,val,test)
  seed = 0,
  verbose = TRUE,
  ...
) {
  if (!requireNamespace("Seurat", quietly = TRUE)) stop("Need Seurat installed.")
  if (!requireNamespace("harmony", quietly = TRUE)) stop("Need harmony installed.")

  .msg <- function(...) if (isTRUE(verbose)) { cat(..., "\n"); flush.console() }
  method_str <- "Harmony (dual stacked)"

  .skip <- function(reason, extra = list()) {
    .msg("[harmony] SKIP:", reason)
    list(
      Z_rna=NULL, Z_atac=NULL, Z_fused=NULL,
      extra_json=c(list(method=method_str, skipped=TRUE, reason=reason), extra)
    )
  }

  .zscore_cols <- function(M) {
    M <- as.matrix(M)
    mu <- colMeans(M, na.rm = TRUE)
    sdv <- apply(M, 2, stats::sd, na.rm = TRUE)
    sdv[!is.finite(sdv) | sdv == 0] <- 1
    sweep(sweep(M, 2, mu, "-"), 2, sdv, "/")
  }

  .assert_fold_cells_ok(fold_cells, ctx = "[harmony]")

  # Restrict to fold universe
  all_cells <- colnames(obj)  # canonical
  fold_univ <- unique(c(fold_cells$train, fold_cells$val, fold_cells$test))
  fold_univ <- intersect(fold_univ, all_cells)
  if (length(fold_univ) < 50) return(.skip("too few fold cells"))

  if (length(fold_univ) != length(all_cells)) {
    obj <- .subset_seurat_safe(obj, cells = fold_univ)
  }

  set.seed(as.integer(seed))
  n <- ncol(obj)
  .msg(sprintf("[harmony] TRANSDUCTIVE: fit Harmony on ALL cells (n=%d)", n))
  .msg("[harmony] HarmonyMatrix batch='modality' on stacked embeddings (unique rownames)...")

  Er <- tryCatch(Seurat::Embeddings(obj, reduction = rna_red), error=function(e) NULL)
  Ea <- tryCatch(Seurat::Embeddings(obj, reduction = atac_red), error=function(e) NULL)
  if (is.null(Er) || is.null(Ea)) return(.skip("missing embeddings for rna_red/atac_red"))

  # bounds-check dims
  rna_dims  <- rna_dims[rna_dims >= 1 & rna_dims <= ncol(Er)]
  atac_dims <- atac_dims[atac_dims >= 1 & atac_dims <= ncol(Ea)]
  if (length(rna_dims) == 0)  return(.skip("rna_dims empty after bounds check"))
  if (length(atac_dims) == 0) return(.skip("atac_dims empty after bounds check"))

  Er <- Er[, rna_dims, drop=FALSE]
  Ea <- Ea[, atac_dims, drop=FALSE]

  common <- intersect(rownames(Er), rownames(Ea))
  if (length(common) < 50) return(.skip("too few paired cells between RNA/ATAC embeddings"))
  common <- common[order(match(common, rownames(Er)))]
  Er <- Er[common, , drop=FALSE]
  Ea <- Ea[common, , drop=FALSE]

  # ---- FIX A: z-score each modality embedding before stacking ----
  Ers <- .zscore_cols(Er)
  Eas <- .zscore_cols(Ea)

  # ---- FIX: make stacked rownames unique (avoid duplicate row.names crash) ----
  rn_rna  <- paste0(common, "::RNA")
  rn_atac <- paste0(common, "::ATAC")

  Z_all <- rbind(Ers, Eas)
  rownames(Z_all) <- c(rn_rna, rn_atac)

  meta2 <- data.frame(
    modality = rep(c("RNA","ATAC"), each = nrow(Ers)),
    row.names = c(rn_rna, rn_atac)
  )

  Z_h <- tryCatch(
    harmony::HarmonyMatrix(
      data_mat = Z_all,
      meta_data = meta2,
      vars_use = "modality",
      do_pca = FALSE,
      verbose = FALSE
    ),
    error = function(e) e
  )
  if (inherits(Z_h, "error")) return(.skip(paste0("HarmonyMatrix failed: ", conditionMessage(Z_h))))

  # split back using the unique rownames
  Zr_h <- Z_h[rn_rna, , drop=FALSE]
  Za_h <- Z_h[rn_atac, , drop=FALSE]
  rownames(Zr_h) <- common
  rownames(Za_h) <- common

  K <- min(as.integer(latent_dim), ncol(Z_h))
  if (K < 2) return(.skip("latent_dim clamp left <2 dims"))

  # ---- FIX B: stable fused construction ----
  # L2-normalize each modality then average, then L2-normalize again
  Zr_k <- Zr_h[, seq_len(K), drop=FALSE]
  Za_k <- Za_h[, seq_len(K), drop=FALSE]
  Zf <- .l2norm_rows((.l2norm_rows(Zr_k) + .l2norm_rows(Za_k)) / 2)
  rownames(Zf) <- common

  .finalize_latents(
    Z_rna   = Zr_h,
    Z_atac  = Za_h,
    Z_fused = Zf,
    fold_cells = fold_cells,
    labels = NULL,
    splits = splits,
    method_str = method_str,
    extra_json = list(
      transductive = TRUE,
      uses_labels = FALSE,
      paired_latents = TRUE,
      zscore_before_stack = TRUE,
      fused_construction = "l2norm(mean(l2norm(Zr_h[1:K]), l2norm(Za_h[1:K])))"
    ),
    verbose = verbose
  )
}


# %% [markdown]
### 5) LIGER

# %% [code]
run_liger_gene_activity <- function(
  obj,
  latent_dim = 30,
  verbose = TRUE,
  splits = NULL,
  assay_rna = "RNA",
  assay_activity = "ACTIVITY",
  min_common = 200L
) {
  if (!requireNamespace("rliger", quietly = TRUE)) stop("Need rliger installed.")
  if (!requireNamespace("Seurat", quietly = TRUE)) stop("Need Seurat installed.")
  if (!requireNamespace("SeuratObject", quietly = TRUE)) stop("Need SeuratObject installed.")
  if (!requireNamespace("Matrix", quietly = TRUE)) stop("Need Matrix installed.")

  v_liger <- as.character(packageVersion("rliger"))
  v_so    <- as.character(packageVersion("SeuratObject"))
  method_str <- paste0("LIGER gene-activity iNMF (rliger ", v_liger, "; SeuratObject ", v_so, ")")
  .msg <- function(...) if (isTRUE(verbose)) cat(..., "\n")

  .skip <- function(reason) {
    list(
      Z_rna = NULL, Z_atac = NULL, Z_fused = NULL,
      extra_json = list(method = method_str, skipped = TRUE, reason = reason)
    )
  }

  ss <- .get_split_cells(obj, splits)
  within_fold <- !is.null(splits)

  fold_cells <- Seurat::Cells(obj)
  if (within_fold) {
    fold_cells <- intersect(ss$fold_cells, fold_cells)
    if (length(fold_cells) < min_common) return(.skip(paste0("too few fold cells: ", length(fold_cells))))
    obj <- subset(obj, cells = fold_cells)
  }

  if (!(assay_rna %in% names(obj@assays))) return(.skip(paste0("no ", assay_rna, " assay")))
  if (!(assay_activity %in% names(obj@assays))) return(.skip("ACTIVITY assay missing"))

  get_counts <- function(assay) {
    a <- obj[[assay]]
    x <- tryCatch(SeuratObject::GetAssayData(a, layer = "counts"), error = function(e) NULL)
    if (is.null(x)) x <- SeuratObject::GetAssayData(a, slot = "counts")
    as(x, "dgCMatrix")
  }

  rna <- get_counts(assay_rna)
  ga  <- get_counts(assay_activity)

  common_cells <- intersect(colnames(rna), colnames(ga))
  if (length(common_cells) < min_common) return(.skip(paste0("too few common cells RNA vs ACTIVITY: ", length(common_cells))))
  rna <- rna[, common_cells, drop = FALSE]
  ga  <- ga [, common_cells, drop = FALSE]

  common_genes <- intersect(rownames(rna), rownames(ga))
  if (length(common_genes) < min_common) return(.skip(paste0("too few common genes: ", length(common_genes))))
  rna <- rna[common_genes, , drop = FALSE]
  ga  <- ga [common_genes, , drop = FALSE]

  K <- as.integer(latent_dim)
  if (is.na(K) || K < 2) return(.skip("latent_dim must be >= 2"))

  .msg("[LIGER] iNMF on n_cells=", ncol(rna), " n_genes=", nrow(rna), " k=", K)

  lig <- tryCatch({
    x <- rliger::createLiger(list(rna = rna, atacGA = ga))
    x <- rliger::normalize(x)
    x <- rliger::selectGenes(x)
    x <- rliger::scaleNotCenter(x)
    x <- rliger::runINMF(x, k = K)
    if ("quantileNorm" %in% getNamespaceExports("rliger")) x <- rliger::quantileNorm(x) else x <- rliger::quantile_norm(x)
    x
  }, error = function(e) e)
  if (inherits(lig, "error")) return(.skip(paste0("LIGER failed: ", conditionMessage(lig))))

  H <- lig@H.norm %||% lig@H
  if (is.null(H)) return(.skip("LIGER returned no H/H.norm"))
  H <- as.matrix(H)

  # Parse modality + barcode from rownames(H)
  rn <- rownames(H)
  mod <- ifelse(grepl("^rna[\\|:_\\-\\.]", rn), "RNA",
         ifelse(grepl("^atacGA[\\|:_\\-\\.]", rn), "ATAC", NA_character_))
  bc  <- rn
  bc  <- sub("^rna[\\|:_\\-\\.]+",    "", bc)
  bc  <- sub("^atacGA[\\|:_\\-\\.]+", "", bc)

  ok <- !is.na(mod) & !is.na(bc) & nzchar(bc)
  H <- H[ok, , drop = FALSE]
  mod <- mod[ok]
  bc  <- bc[ok]

  # Build paired matrices where possible
  # (If duplicates exist within a modality/barcode, average them)
  build_lat <- function(target_mod) {
    rows <- which(mod == target_mod)
    if (!length(rows)) return(NULL)
    Hm <- H[rows, , drop = FALSE]
    b  <- bc[rows]
    ix <- split(seq_len(nrow(Hm)), b)
    out <- matrix(NA_real_, nrow = length(ix), ncol = ncol(Hm),
                  dimnames = list(names(ix), colnames(Hm)))
    for (i in seq_along(ix)) {
      rr <- ix[[i]]
      out[i, ] <- if (length(rr) == 1) Hm[rr, ] else colMeans(Hm[rr, , drop = FALSE])
    }
    out
  }

  Zr <- build_lat("RNA")
  Za <- build_lat("ATAC")

  # Keep only cells in the Seurat object’s order
  cn <- Seurat::Cells(obj)

  # Paired intersection for Zr/Za
  common_pair <- intersect(rownames(Zr %||% matrix(nrow=0,ncol=0)), rownames(Za %||% matrix(nrow=0,ncol=0)))
  common_pair <- intersect(common_pair, cn)

  # If we have paired: return paired + fused (average)
  if (length(common_pair) >= min_common) {
    common_pair <- cn[cn %in% common_pair]
    Zr <- Zr[common_pair, , drop = FALSE]
    Za <- Za[common_pair, , drop = FALSE]
    d <- min(K, ncol(Zr), ncol(Za))
    Zr <- Zr[, seq_len(d), drop = FALSE]
    Za <- Za[, seq_len(d), drop = FALSE]
    Zf <- .l2norm_rows((.l2norm_rows(Zr) + .l2norm_rows(Za)) / 2)

    return(.finalize_latents(
      Z_rna = Zr,
      Z_atac = Za,
      Z_fused = Zf,
      fold_cells = cn,
      labels = ss$labels,
      splits = ss$splits,
      method_str = method_str,
      extra_json = list(
        transductive = TRUE,               # LIGER fit uses all fold cells; no true inductive projection here
        uses_labels = FALSE,
        paired_latents = TRUE,
        fit_cells_mode = if (within_fold) "all_fold_cells" else "all_cells",
        note = "LIGER: returned paired (RNA vs ACTIVITY) factors when both modalities exist per barcode; fused = average."
      ),
      verbose = verbose
    ))
  }

  # Otherwise: fall back to fused-only (still useful)
  # Use whichever modality has more rows
  Zf <- Zr %||% Za
  if (is.null(Zf) || nrow(Zf) < min_common) return(.skip("could not build sufficient paired or fused factors"))
  keep <- intersect(cn, rownames(Zf))
  keep <- cn[cn %in% keep]
  Zf <- Zf[keep, , drop = FALSE]
  Zf <- Zf[, seq_len(min(K, ncol(Zf))), drop = FALSE]

  list(
    Z_rna = NULL,
    Z_atac = NULL,
    Z_fused = Zf,
    extra_json = list(
      method = method_str,
      transductive = TRUE,
      uses_labels = FALSE,
      paired_latents = FALSE,
      fit_cells_mode = if (within_fold) "all_fold_cells" else "all_cells",
      note = "LIGER: paired factors not available for enough cells; returning fused-only from the dominant modality."
    )
  )
}


# %% [markdown]
### 6) MOFA2

# %% [code]
run_mofa2_pca_lsi <- function(
  obj,
  latent_dim = 30,
  rna_dims = 1:100,
  atac_dims = 2:101,
  verbose = TRUE,
  splits = NULL,
  min_common = 50L,
  seed = 1L,                 # NEW: reproducibility (mechanical)
  maxiter = 1000L,           # NEW: explicit, still defaulting to your 1000
  convergence_mode = "fast"  # NEW: explicit
) {
  if (!requireNamespace("MOFA2", quietly = TRUE)) stop("Need MOFA2 installed.")
  if (!requireNamespace("Seurat", quietly = TRUE)) stop("Need Seurat installed.")

  v_mofa <- as.character(packageVersion("MOFA2"))
  .msg <- function(...) if (isTRUE(verbose)) { cat(..., "\n"); flush.console() }

  ss <- .get_split_cells(obj, splits)
  within_fold <- !is.null(splits)

  if (within_fold) {
    fold_cells <- intersect(ss$fold_cells, colnames(obj))
    if (length(fold_cells) < 200) stop("[MOFA2] too few fold cells.")
    obj <- subset(obj, cells = fold_cells)
    .msg(sprintf("[MOFA2] within-fold TRANSDUCTIVE: fit on all fold cells (n=%d)", ncol(obj)))
  } else {
    fold_cells <- colnames(obj)
    .msg(sprintf("[MOFA2] TRANSDUCTIVE: fit on all cells (n=%d)", ncol(obj)))
  }

  if (!("pca" %in% names(obj@reductions))) stop("Need PCA on RNA (obj[['pca']]).")
  if (!("lsi" %in% names(obj@reductions))) stop("Need LSI on ATAC (obj[['lsi']]).")

  .clamp_dims <- function(dims, n_avail) {
    dims <- as.integer(dims)
    dims <- dims[is.finite(dims)]
    dims <- dims[dims >= 1 & dims <= as.integer(n_avail)]
    dims <- unique(dims)
    if (!length(dims)) stop("no valid dims after clamping")
    dims
  }

  Zr_all <- Seurat::Embeddings(obj, "pca")
  Za_all <- Seurat::Embeddings(obj, "lsi")
  dr <- .clamp_dims(rna_dims,  ncol(Zr_all))
  da <- .clamp_dims(atac_dims, ncol(Za_all))

  Zr0 <- Zr_all[, dr, drop = FALSE]
  Za0 <- Za_all[, da, drop = FALSE]

  al <- .align_paired_latents(Zr0, Za0, min_common = min_common, enforce_order = TRUE)
  if (is.null(al$Zr)) stop("[MOFA2] cannot align PCA/LSI inputs: ", al$reason)

  Zr <- al$Zr
  Za <- al$Za
  common <- rownames(Zr)

  # ---- Mechanistic tweak: don't unnecessarily truncate both to k_view unless you really want symmetry ----
  # We'll keep each view as-is; MOFA can handle different numbers of features per view.
  # If you *want* symmetry, set k_view manually outside and pass dims that match.

  # MOFA expects features x samples
  # scale(): center+scale each feature (dimension) across cells; keep safe for sd=0
  .scale_safe <- function(M) {
    M <- as.matrix(M)
    mu <- rowMeans(M, na.rm = TRUE)
    sdv <- apply(M, 1, stats::sd, na.rm = TRUE)
    sdv[!is.finite(sdv) | sdv == 0] <- 1
    sweep(sweep(M, 1, mu, "-"), 1, sdv, "/")
  }

  view_rna  <- .scale_safe(t(Zr))  # dims x cells
  view_atac <- .scale_safe(t(Za))  # dims x cells
  data_list <- list(RNA_PCA = view_rna, ATAC_LSI = view_atac)

  .msg("[MOFA2] create/prepare...")
  mofa <- MOFA2::create_mofa(data_list)

  model_opts <- MOFA2::get_default_model_options(mofa)
  model_opts$num_factors <- as.integer(latent_dim)

  train_opts <- MOFA2::get_default_training_options(mofa)
  train_opts$maxiter <- as.integer(maxiter)
  train_opts$convergence_mode <- as.character(convergence_mode)

  # ---- Mechanistic tweak: reproducibility ----
  # This makes factors much less “jittery” across folds/runs.
  set.seed(as.integer(seed))

  mofa <- MOFA2::prepare_mofa(mofa, model_options = model_opts, training_options = train_opts)

  outfile <- file.path(tempdir(), paste0("MOFA2__K", latent_dim, "__", format(Sys.time(), "%Y%m%d_%H%M%S"), ".hdf5"))
  .msg("[MOFA2] training -> ", outfile)
  mofa <- MOFA2::run_mofa(mofa, outfile = outfile, use_basilisk = TRUE)

  factors_list <- MOFA2::get_factors(mofa, factors = "all")
  group_name <- names(factors_list)[1]
  Zf <- as.matrix(factors_list[[group_name]])  # samples x K
  if (is.null(rownames(Zf))) rownames(Zf) <- common

  # align everything to the fold cell universe/order
  keep <- intersect(fold_cells, Reduce(intersect, list(rownames(Zr), rownames(Za), rownames(Zf))))
  if (length(keep) < min_common) stop("[MOFA2] too few common cells after factor extraction")

  keep <- fold_cells[fold_cells %in% keep]
  Zr <- Zr[keep, , drop = FALSE]
  Za <- Za[keep, , drop = FALSE]
  Zf <- Zf[keep, , drop = FALSE]

  list(
    mofa = mofa,
    Z_rna = Zr,
    Z_atac = Za,
    Z_fused = Zf,
    extra_json = list(
      method = paste0("MOFA2 on PCA+LSI views (MOFA2 ", v_mofa, ")"),
      transductive = TRUE,
      uses_labels = FALSE,
      paired_latents = TRUE,
      fit_cells_mode = if (within_fold) "all_fold_cells" else "all_cells",
      note = "Returns paired view latents as PCA/LSI inputs + joint MOFA2 factors (fused). View latents are baseline PCA/LSI, not MOFA-specific view latents.",
      rna_dims_used = dr,
      atac_dims_used = da,
      latent_dim = as.integer(latent_dim),
      n_cells = length(keep),
      seed = as.integer(seed),
      maxiter = as.integer(maxiter),
      convergence_mode = as.character(convergence_mode)
    )
  )
}


# %% [markdown]
### Split + labels code

# %% [code]
# ---------------------------
# Seed + labels + train/test split
# ---------------------------
seed <- 0L
set.seed(seed)

# canonical cell IDs / order for everything downstream
cell_ids <- colnames(obj)
stopifnot(length(cell_ids) == ncol(obj))

md <- obj@meta.data
stopifnot(all(cell_ids %in% rownames(md)))

# ---- choose a label column if available ----
label_candidates <- c(
  "celltype", "cell_type", "celltype.l2", "celltype.l1",
  "seurat_annotations", "predicted.celltype", "celltype_final"
)

label_key <- label_candidates[label_candidates %in% colnames(md)][1]
if (is.na(label_key) || length(label_key) == 0) label_key <- NA_character_

if (!is.na(label_key)) {
  # pull labels from meta.data and reorder to match obj column order
  labels_named <- setNames(as.character(md[[label_key]]), rownames(md))
  labels <- labels_named[cell_ids]

  if (anyNA(labels)) {
    bad <- sum(is.na(labels))
    stop(sprintf("[labels] %d labels are NA after aligning '%s' to obj cells", bad, label_key))
  }

  cat("[labels] using meta.data column:", label_key, "\n")
} else {
  # fallback: RNA clustering labels
  cat("[labels] no label column found; using RNA clusters as fallback labels\n")

  Seurat::DefaultAssay(obj) <- "RNA"

  # ensure PCA exists
  if (!("pca" %in% names(obj@reductions))) {
    obj <- Seurat::NormalizeData(obj, verbose = FALSE)
    obj <- Seurat::FindVariableFeatures(obj, nfeatures = 2000, verbose = FALSE)
    obj <- Seurat::ScaleData(obj, verbose = FALSE)
    obj <- Seurat::RunPCA(obj, npcs = 30, verbose = FALSE)
  }

  obj <- Seurat::FindNeighbors(obj, reduction = "pca", dims = 1:30, verbose = FALSE)
  obj <- Seurat::FindClusters(obj, resolution = 0.5, verbose = FALSE)

  labels <- as.character(obj$seurat_clusters)
  label_key <- "seurat_clusters"
}

# ---- ALWAYS name labels with canonical cell IDs ----
stopifnot(length(labels) == length(cell_ids))
names(labels) <- cell_ids
stopifnot(identical(names(labels), cell_ids))

cat("[labels] n=", length(labels),
    " unique=", length(unique(labels)),
    " key=", label_key, "\n")

# ---- train/test split in index space (paired; one index space) ----
n <- length(cell_ids)
idx_all <- sample.int(n, size = n, replace = FALSE)

n_train <- floor(0.8 * n)
splits <- list(
  train = idx_all[1:n_train],
  test  = idx_all[(n_train + 1):n]
)

cat("[splits] n_train=", length(splits$train),
    " n_test=", length(splits$test),
    " (seed=", seed, ")\n")


# %% [markdown]
### Evaluation loop function(s)

# %% [code]
library(data.table)

# -----------------------------
# small helpers
# -----------------------------
`%||%` <- if (exists("%||%", mode = "function")) get("%||%") else function(a, b) {
  if (is.null(a) || length(a) == 0) b else a
}

.clamp_k <- function(k, n) {
  if (is.null(n) || n < 3) return(NA_integer_)
  as.integer(min(as.integer(k), as.integer(n) - 2L))
}

.is_finite_rows <- function(Z) {
  Z <- as.matrix(Z)
  if (nrow(Z) == 0) return(logical(0))
  rowSums(!is.finite(Z)) == 0L
}

.mode_chr <- function(x) {
  x <- as.character(x)
  x <- x[!is.na(x)]
  if (!length(x)) return(NA_character_)
  tab <- table(x)
  names(tab)[which.max(tab)]
}

.macro_f1 <- function(y_true, y_pred) {
  y_true <- as.character(y_true)
  y_pred <- as.character(y_pred)
  labs <- sort(unique(y_true[!is.na(y_true)]))
  if (!length(labs)) return(NA_real_)

  f1s <- vapply(labs, function(lv) {
    tp <- sum(y_true == lv & y_pred == lv, na.rm = TRUE)
    fp <- sum(y_true != lv & y_pred == lv, na.rm = TRUE)
    fn <- sum(y_true == lv & y_pred != lv, na.rm = TRUE)
    prec <- if ((tp + fp) > 0) tp / (tp + fp) else 0
    rec  <- if ((tp + fn) > 0) tp / (tp + fn) else 0
    if ((prec + rec) > 0) 2 * prec * rec / (prec + rec) else 0
  }, numeric(1))

  mean(f1s, na.rm = TRUE)
}

.balanced_acc <- function(y_true, y_pred) {
  y_true <- as.character(y_true)
  y_pred <- as.character(y_pred)
  labs <- sort(unique(y_true[!is.na(y_true)]))
  if (!length(labs)) return(NA_real_)

  recs <- vapply(labs, function(lv) {
    tp <- sum(y_true == lv & y_pred == lv, na.rm = TRUE)
    fn <- sum(y_true == lv & y_pred != lv, na.rm = TRUE)
    if ((tp + fn) > 0) tp / (tp + fn) else NA_real_
  }, numeric(1))

  mean(recs, na.rm = TRUE)
}

.safe <- function(expr, on_error = NULL) {
  tryCatch(expr, error = function(e) on_error)
}

# -----------------------------
# Helper: call a function but silently drop unsupported named args
# - if fn has ..., pass everything
# - otherwise, only pass named args it supports
# (positional args are always passed as-is)
# -----------------------------
.call_supported <- function(fn, ...) {
  dots <- list(...)
  nms  <- names(dots); if (is.null(nms)) nms <- rep("", length(dots))

  fmls <- names(formals(fn)); if (is.null(fmls)) fmls <- character(0)
  has_dots <- "..." %in% fmls

  # split positional vs named
  is_named <- nzchar(nms)
  pos <- dots[!is_named]
  nam <- dots[is_named]

  if (has_dots) return(do.call(fn, c(pos, nam)))

  keep <- intersect(names(nam), fmls)
  do.call(fn, c(pos, nam[keep]))
}

# -----------------------------
# PAIRED FOSCTTM wrapper
# Uses idx_eval, subsample_n (or fos_sub), seed if supported by foscttm_values
# Returns mean, sem, plus optional extras if foscttm_values returns them
# -----------------------------
.foscttm_metrics <- function(Z_rna, Z_atac, idx_eval, fos_sub = 3000L, seed = 0L) {
  if (!exists("foscttm_values", mode = "function")) stop("foscttm_values() not found")
  if (is.null(Z_rna) || is.null(Z_atac)) stop("Z_rna/Z_atac required")
  if (is.null(idx_eval) || length(idx_eval) < 2) stop("idx_eval missing/too small")

  Z_rna  <- as.matrix(Z_rna)
  Z_atac <- as.matrix(Z_atac)
  if (nrow(Z_rna) != nrow(Z_atac)) stop("Z_rna/Z_atac nrow mismatch")

  idx_eval <- as.integer(idx_eval)
  idx_eval <- idx_eval[idx_eval >= 1 & idx_eval <= nrow(Z_rna)]
  if (length(idx_eval) < 2) stop("idx_eval too small after clamping")

  fos_sub_eff <- as.integer(min(as.integer(fos_sub), length(idx_eval)))

  f <- get("foscttm_values", mode = "function")

  # try a few calling conventions, but ALWAYS include idx_eval and subsample if supported
  out <- .safe(.call_supported(f, Z_rna, Z_atac,
                              idx_eval = idx_eval,
                              subsample_n = fos_sub_eff,
                              fos_sub = fos_sub_eff,
                              seed = as.integer(seed)),
              on_error = NULL)
  if (is.null(out)) out <- .safe(.call_supported(f, Z1 = Z_rna, Z2 = Z_atac,
                                                idx_eval = idx_eval,
                                                subsample_n = fos_sub_eff,
                                                fos_sub = fos_sub_eff,
                                                seed = as.integer(seed)),
                                on_error = NULL)
  if (is.null(out)) out <- .safe(.call_supported(f, A = Z_rna, B = Z_atac,
                                                idx_eval = idx_eval,
                                                subsample_n = fos_sub_eff,
                                                fos_sub = fos_sub_eff,
                                                seed = as.integer(seed)),
                                on_error = NULL)
  if (is.null(out)) stop("foscttm_values call failed (signature mismatch)")

  vec <- NULL
  mrr <- NA_real_
  rec1 <- rec10 <- rec25 <- rec50 <- rec100 <- NA_real_

  if (is.numeric(out)) {
    vec <- as.numeric(out)
  } else if (is.list(out)) {
    for (cand in c("foscttm", "foscttm_values", "values")) {
      if (!is.null(out[[cand]]) && is.numeric(out[[cand]])) {
        vec <- as.numeric(out[[cand]])
        break
      }
    }
    if (!is.null(out$mrr) && is.numeric(out$mrr)) mrr <- mean(out$mrr, na.rm = TRUE)
    if (!is.null(out$MRR) && is.numeric(out$MRR)) mrr <- mean(out$MRR, na.rm = TRUE)

    if (!is.null(out$recall) && is.list(out$recall)) {
      rec <- out$recall
      getr <- function(k) rec[[as.character(k)]] %||% rec[[paste0("k", k)]] %||% rec[[paste0("@", k)]]
      rec1   <- as.numeric(getr(1)   %||% NA_real_)
      rec10  <- as.numeric(getr(10)  %||% NA_real_)
      rec25  <- as.numeric(getr(25)  %||% NA_real_)
      rec50  <- as.numeric(getr(50)  %||% NA_real_)
      rec100 <- as.numeric(getr(100) %||% NA_real_)
    }
  }

  if (is.null(vec) || !length(vec)) stop("foscttm_values returned no numeric vector")
  vec <- vec[is.finite(vec)]
  if (!length(vec)) stop("foscttm vector all non-finite")

  mu  <- mean(vec)
  sem <- stats::sd(vec) / sqrt(max(1L, length(vec)))

  list(
    mean = mu,
    sem  = sem,
    mrr_mean = mrr,
    recall1  = rec1,
    recall10 = rec10,
    recall25 = rec25,
    recall50 = rec50,
    recall100= rec100,
    n_eval   = length(vec)
  )
}

# -----------------------------
# PAIRED mixing score wrapper
# Prefers mixing_proxy_rann(Zr, Za, idx_eval=..., k=...) if present
# -----------------------------
.mixing_score_paired <- function(Z_rna, Z_atac, idx_eval, k = 50L) {
  if (is.null(Z_rna) || is.null(Z_atac)) return(NA_real_)
  if (is.null(idx_eval) || length(idx_eval) < 2) return(NA_real_)

  Z_rna  <- as.matrix(Z_rna)
  Z_atac <- as.matrix(Z_atac)
  if (nrow(Z_rna) != nrow(Z_atac) || nrow(Z_rna) < 2) return(NA_real_)

  idx_eval <- as.integer(idx_eval)
  idx_eval <- idx_eval[idx_eval >= 1 & idx_eval <= nrow(Z_rna)]
  if (length(idx_eval) < 2) return(NA_real_)

  # true paired mixing
  if (exists("mixing_proxy_rann", mode = "function")) {
    f <- get("mixing_proxy_rann", mode = "function")
    return(.safe(as.numeric(.call_supported(f, Z_rna, Z_atac, idx_eval = idx_eval, k = as.integer(k))),
                 on_error = NA_real_))
  }

  # sometimes mixing_proxy is also paired
  if (exists("mixing_proxy", mode = "function")) {
    f <- get("mixing_proxy", mode = "function")
    out <- .safe(as.numeric(.call_supported(f, Z_rna, Z_atac, idx_eval = idx_eval, k = as.integer(k))),
                 on_error = NA_real_)
    return(out)
  }

  NA_real_
}

# -----------------------------
# kNN label transfer
# -----------------------------
.knn_transfer <- function(Z_train, y_train, Z_test, y_test, k = 15L) {
  if (!requireNamespace("RANN", quietly = TRUE)) {
    return(list(acc = NA_real_, macroF1 = NA_real_, y_pred = NULL,
                n_train = NA_integer_, n_test = NA_integer_))
  }

  Z_train <- as.matrix(Z_train)
  Z_test  <- as.matrix(Z_test)

  ok_tr <- .is_finite_rows(Z_train)
  ok_te <- .is_finite_rows(Z_test)

  Z_train <- Z_train[ok_tr, , drop = FALSE]
  y_train <- as.character(y_train[ok_tr])

  Z_test  <- Z_test[ok_te, , drop = FALSE]
  y_test2 <- as.character(y_test[ok_te])

  if (nrow(Z_train) < 10 || nrow(Z_test) < 5) {
    return(list(acc = NA_real_, macroF1 = NA_real_, y_pred = NULL,
                n_train = nrow(Z_train), n_test = nrow(Z_test)))
  }

  k_eff <- .clamp_k(as.integer(k), nrow(Z_train))
  if (is.na(k_eff) || k_eff < 1) {
    return(list(acc = NA_real_, macroF1 = NA_real_, y_pred = NULL,
                n_train = nrow(Z_train), n_test = nrow(Z_test)))
  }

  nn <- RANN::nn2(data = Z_train, query = Z_test, k = k_eff)$nn.idx
  y_pred <- apply(nn, 1, function(ii) .mode_chr(y_train[ii]))

  acc <- mean(y_pred == y_test2, na.rm = TRUE)
  list(
    acc = acc,
    macroF1 = .macro_f1(y_test2, y_pred),
    y_pred = y_pred,
    y_true = y_test2,
    n_train = nrow(Z_train),
    n_test  = nrow(Z_test)
  )
}

# -----------------------------
# fused embedding metrics (robust)
# -----------------------------
.fused_metrics <- function(Zf, labels_named, tr_cells, te_cells,
                           k = 15L, silhouette_max_n = 3000L, seed = 0L) {

  out <- list(
    fused_knn_acc_test = NA_real_,
    fused_knn_macroF1_test = NA_real_,
    fused_knn_balanced_acc_test = NA_real_,
    fused_silhouette_test = NA_real_,
    fused_knn_label_purity_test = NA_real_,
    fused_kmeans_ari_test = NA_real_,
    fused_kmeans_nmi_test = NA_real_
  )

  if (is.null(Zf) || is.null(labels_named) || is.null(names(labels_named))) return(out)
  if (is.null(tr_cells) || is.null(te_cells)) return(out)

  Zf <- as.matrix(Zf)
  if (is.null(rownames(Zf))) return(out)

  common <- intersect(rownames(Zf), names(labels_named))
  if (length(common) < 50) return(out)

  Zf <- Zf[common, , drop = FALSE]
  y  <- as.character(labels_named[common]); names(y) <- common

  ok <- .is_finite_rows(Zf)
  if (!all(ok)) {
    Zf <- Zf[ok, , drop = FALSE]
    y  <- y[rownames(Zf)]
  }

  tr <- intersect(tr_cells, rownames(Zf))
  te <- intersect(te_cells, rownames(Zf))
  if (length(tr) < 20 || length(te) < 20) return(out)

  Z_train <- Zf[tr, , drop = FALSE]; y_train <- y[tr]
  Z_test  <- Zf[te, , drop = FALSE]; y_test  <- y[te]

  kt <- .knn_transfer(Z_train, y_train, Z_test, y_test, k = k)
  out$fused_knn_acc_test <- kt$acc
  out$fused_knn_macroF1_test <- kt$macroF1
  if (!is.null(kt$y_pred) && !is.null(kt$y_true)) {
    out$fused_knn_balanced_acc_test <- .balanced_acc(kt$y_true, kt$y_pred)
  }

  # label purity (TEST)
  if (requireNamespace("RANN", quietly = TRUE) && nrow(Z_test) >= 10) {
    k_eff <- .clamp_k(as.integer(k) + 1L, nrow(Z_test))
    if (!is.na(k_eff) && k_eff >= 3) {
      nn <- RANN::nn2(data = Z_test, query = Z_test, k = k_eff)$nn.idx
      nn <- nn[, -1, drop = FALSE]
      purity <- vapply(seq_len(nrow(nn)), function(i) {
        mean(y_test[nn[i, ]] == y_test[i], na.rm = TRUE)
      }, numeric(1))
      out$fused_knn_label_purity_test <- mean(purity, na.rm = TRUE)
    }
  }

  # silhouette (TEST)
  if (requireNamespace("cluster", quietly = TRUE)) {
    set.seed(seed)
    te_use <- te
    if (length(te_use) > silhouette_max_n) te_use <- sample(te_use, size = silhouette_max_n, replace = FALSE)

    y_s <- y[te_use]
    Z_s <- Zf[te_use, , drop = FALSE]
    y_s <- y_s[!is.na(y_s)]
    Z_s <- Z_s[names(y_s), , drop = FALSE]

    if (length(unique(y_s)) >= 2 && nrow(Z_s) >= 10) {
      D <- stats::dist(Z_s)
      sil <- cluster::silhouette(as.integer(as.factor(y_s)), D)
      out$fused_silhouette_test <- mean(sil[, "sil_width"], na.rm = TRUE)
    }
  }

  # kmeans ARI + NMI (TEST), k = nclasses in TRAIN
  k_clust <- length(unique(y_train[!is.na(y_train)]))
  if (k_clust >= 2 && nrow(Z_test) >= k_clust) {
    set.seed(seed)
    yt <- y_test
    oky <- !is.na(yt)
    Zk <- Z_test[oky, , drop = FALSE]
    yt <- yt[oky]

    if (nrow(Zk) >= k_clust && length(unique(yt)) >= 2) {
      km <- .safe(stats::kmeans(Zk, centers = k_clust, nstart = 10), on_error = NULL)
      if (!is.null(km)) {
        if (requireNamespace("mclust", quietly = TRUE)) {
          out$fused_kmeans_ari_test <- mclust::adjustedRandIndex(
            as.integer(as.factor(yt)),
            as.integer(km$cluster)
          )
        }
        if (requireNamespace("aricode", quietly = TRUE)) {
          out$fused_kmeans_nmi_test <- aricode::NMI(
            as.integer(as.factor(yt)),
            as.integer(km$cluster)
          )
        }
      }
    }
  }

  out
}

# -----------------------------
# evaluate_one: COMPUTES + RETURNS BOTH legacy and canonical columns
# -----------------------------
evaluate_one <- function(name, out, labels, splits,
                         k_mix = 30L, k_lt = 15L, fos_sub = 3000L,
                         seed = 0L, verbose = FALSE) {

  .msg <- function(...) if (isTRUE(verbose)) message(...)

  # pull embeddings
  Zf <- out$Z_fused %||% out$Zf %||% out$latents$fused %||% NULL
  Zr <- out$Z_rna   %||% out$Zr %||% out$latents$rna   %||% NULL
  Za <- out$Z_atac  %||% out$Za %||% out$latents$atac  %||% NULL

  extra <- out$extra_json %||% out$extra %||% list()
  method_str  <- extra$method %||% extra$method_str %||% name
  fit_seconds <- out$fit_seconds %||% extra$fit_seconds %||% NA_real_

  status_in <- out$status %||% extra$status %||% NA_character_
  skipped_in <- isTRUE(extra$skipped)
  reason_in  <- extra$reason %||% out$reason %||% NA_character_

  error_flag <- isTRUE(out$error) || isTRUE(out$errored) || isTRUE(out$failed) ||
    identical(status_in, "failed") || identical(status_in, "error")

  failed  <- isTRUE(skipped_in) || isTRUE(error_flag)
  skipped <- failed

  trained_on   <- extra$trained_on %||% NA_character_
  transductive <- extra$transductive %||% (!is.na(trained_on) && trained_on %in% c("all", "full"))
  uses_labels  <- extra$uses_labels %||% FALSE

  .record_err <- function(key, e) {
    extra[[key]] <- paste0(class(e)[1], ": ", conditionMessage(e))
    .msg(paste0("[", name, "] ", key, " -> ", extra[[key]]))
    invisible(NULL)
  }

  # legacy columns (what your dt_runs currently expects)
  FOSCTTM_mean_test <- NA_real_
  FOSCTTM_sem_test  <- NA_real_
  FOSCTTM_mrr_mean_test <- NA_real_
  `FOSCTTM_recall@1_mean_test`  <- NA_real_
  `FOSCTTM_recall@10_mean_test` <- NA_real_
  mixing_score_test <- NA_real_
  label_transfer_acc_mean_test <- NA_real_

  # canonical columns (optional / for your future rename)
  `PAIR(test)/FOSCTTM_mean` <- NA_real_
  `PAIR(test)/FOSCTTM_MRR`  <- NA_real_
  `PAIR(test)/FOSCTTM_n_eval` <- NA_real_
  `PAIR(test)/FOSCTTM_Recall@1`   <- NA_real_
  `PAIR(test)/FOSCTTM_Recall@10`  <- NA_real_
  `PAIR(test)/FOSCTTM_Recall@25`  <- NA_real_
  `PAIR(test)/FOSCTTM_Recall@50`  <- NA_real_
  `PAIR(test)/FOSCTTM_Recall@100` <- NA_real_

  `PAIR(test)/Label transfer acc mean` <- NA_real_
  `PAIR(test)/Label transfer macroF1 mean` <- NA_real_
  `MIX(test)/Mixing score` <- NA_real_
  `Fused kNN acc` <- NA_real_
  `Fused kNN macroF1` <- NA_real_
  `Fused kNN balanced acc` <- NA_real_
  `MIX(test)/Fused silhouette` <- NA_real_
  `MIX(test)/Fused label purity` <- NA_real_
  `MIX(test)/Fused k-means ARI` <- NA_real_
  `MIX(test)/Fused k-means NMI` <- NA_real_
  `__eval_error__` <- NA_character_

  # name labels by cell IDs if needed
  if (!is.null(labels) && is.null(names(labels))) {
    cn <- NULL
    if (!is.null(Zf) && !is.null(rownames(Zf))) cn <- rownames(Zf)
    else if (!is.null(Zr) && !is.null(rownames(Zr))) cn <- rownames(Zr)
    else if (!is.null(Za) && !is.null(rownames(Za))) cn <- rownames(Za)
    if (!is.null(cn) && length(labels) == length(cn)) names(labels) <- cn
  }

  # build split cell IDs
  sp_cells <- tryCatch(
    build_split_cells(splits, labels_named = labels),
    error = function(e) { .record_err("split_build_error", e); NULL }
  )
  tr_cells <- sp_cells$train_cells %||% character(0)
  te_cells <- sp_cells$test_cells  %||% character(0)

  # return early if failed
  if (isTRUE(skipped)) {
    return(data.table(
      method = name,
      failed = TRUE,
      fit_seconds = fit_seconds,
      transductive = as.numeric(isTRUE(transductive)),
      uses_labels  = as.numeric(isTRUE(uses_labels)),

      # legacy
      FOSCTTM_mean_test = FOSCTTM_mean_test,
      FOSCTTM_sem_test  = FOSCTTM_sem_test,
      FOSCTTM_mrr_mean_test = FOSCTTM_mrr_mean_test,
      `FOSCTTM_recall@1_mean_test` = `FOSCTTM_recall@1_mean_test`,
      `FOSCTTM_recall@10_mean_test` = `FOSCTTM_recall@10_mean_test`,
      mixing_score_test = mixing_score_test,
      label_transfer_acc_mean_test = label_transfer_acc_mean_test,

      # canonical
      `PAIR(test)/FOSCTTM_mean` = `PAIR(test)/FOSCTTM_mean`,
      `PAIR(test)/FOSCTTM_MRR`  = `PAIR(test)/FOSCTTM_MRR`,
      `PAIR(test)/FOSCTTM_n_eval` = `PAIR(test)/FOSCTTM_n_eval`,
      `PAIR(test)/FOSCTTM_Recall@1` = `PAIR(test)/FOSCTTM_Recall@1`,
      `PAIR(test)/FOSCTTM_Recall@10` = `PAIR(test)/FOSCTTM_Recall@10`,
      `PAIR(test)/FOSCTTM_Recall@25` = `PAIR(test)/FOSCTTM_Recall@25`,
      `PAIR(test)/FOSCTTM_Recall@50` = `PAIR(test)/FOSCTTM_Recall@50`,
      `PAIR(test)/FOSCTTM_Recall@100` = `PAIR(test)/FOSCTTM_Recall@100`,
      `PAIR(test)/Label transfer acc mean` = `PAIR(test)/Label transfer acc mean`,
      `PAIR(test)/Label transfer macroF1 mean` = `PAIR(test)/Label transfer macroF1 mean`,
      `MIX(test)/Mixing score` = `MIX(test)/Mixing score`,
      `Fused kNN acc` = `Fused kNN acc`,
      `Fused kNN macroF1` = `Fused kNN macroF1`,
      `Fused kNN balanced acc` = `Fused kNN balanced acc`,
      `MIX(test)/Fused silhouette` = `MIX(test)/Fused silhouette`,
      `MIX(test)/Fused label purity` = `MIX(test)/Fused label purity`,
      `MIX(test)/Fused k-means ARI` = `MIX(test)/Fused k-means ARI`,
      `MIX(test)/Fused k-means NMI` = `MIX(test)/Fused k-means NMI`,
      `__eval_error__` = "skipped_or_fit_failed",

      extra_json = jsonlite::toJSON(extra, auto_unbox = TRUE),
      skipped = TRUE,
      reason = reason_in,
      method_str = method_str
    ))
  }

  # -----------------------------
  # 1) paired metrics on TEST
  # -----------------------------
  if (!is.null(Zr) && !is.null(Za) &&
      !is.null(rownames(Zr)) && !is.null(rownames(Za))) {

    common_cells <- intersect(rownames(Zr), rownames(Za))
    if (length(common_cells) >= 50) {

      ok_r <- .is_finite_rows(Zr[common_cells, , drop = FALSE])
      ok_a <- .is_finite_rows(Za[common_cells, , drop = FALSE])
      finite_cells <- common_cells[ok_r & ok_a]

      te_cells_eff <- intersect(te_cells, finite_cells)

      if (length(finite_cells) >= 50 && length(te_cells_eff) >= 50) {
        Zr2 <- as.matrix(Zr[finite_cells, , drop = FALSE])
        Za2 <- as.matrix(Za[finite_cells, , drop = FALSE])

        te_idx <- match(te_cells_eff, finite_cells)
        te_idx <- te_idx[!is.na(te_idx)]

        # paired mixing (THIS is what your pipeline used before)
        mix_pair <- tryCatch(
          .mixing_score_paired(Zr2, Za2, idx_eval = te_idx, k = as.integer(k_mix)),
          error = function(e) { .record_err("mixing_pair_error", e); NA_real_ }
        )
        mixing_score_test <- mix_pair
        `MIX(test)/Mixing score` <- mix_pair

        # paired FOSCTTM
        fm <- tryCatch(
          .foscttm_metrics(Zr2, Za2, idx_eval = te_idx, fos_sub = as.integer(fos_sub), seed = seed),
          error = function(e) { .record_err("foscttm_error", e); NULL }
        )
        if (!is.null(fm)) {
          # legacy
          FOSCTTM_mean_test <- fm$mean %||% NA_real_
          FOSCTTM_sem_test  <- fm$sem  %||% NA_real_
          FOSCTTM_mrr_mean_test <- fm$mrr_mean %||% NA_real_
          `FOSCTTM_recall@1_mean_test`  <- fm$recall1  %||% NA_real_
          `FOSCTTM_recall@10_mean_test` <- fm$recall10 %||% NA_real_

          # canonical
          `PAIR(test)/FOSCTTM_mean` <- fm$mean %||% NA_real_
          `PAIR(test)/FOSCTTM_MRR`  <- fm$mrr_mean %||% NA_real_
          `PAIR(test)/FOSCTTM_n_eval` <- as.numeric(fm$n_eval %||% NA_real_)
          `PAIR(test)/FOSCTTM_Recall@1`   <- fm$recall1  %||% NA_real_
          `PAIR(test)/FOSCTTM_Recall@10`  <- fm$recall10 %||% NA_real_
          `PAIR(test)/FOSCTTM_Recall@25`  <- fm$recall25 %||% NA_real_
          `PAIR(test)/FOSCTTM_Recall@50`  <- fm$recall50 %||% NA_real_
          `PAIR(test)/FOSCTTM_Recall@100` <- fm$recall100%||% NA_real_
        }

        # label transfer mean accuracy (paired)
        if (!is.null(names(labels))) {
          y_finite <- as.character(labels[finite_cells]); names(y_finite) <- finite_cells
          tr_cells_eff <- intersect(tr_cells, finite_cells)

          tr_cells_lab <- tr_cells_eff[!is.na(y_finite[tr_cells_eff])]
          te_cells_lab <- te_cells_eff[!is.na(y_finite[te_cells_eff])]

          if (length(tr_cells_lab) >= 50 && length(te_cells_lab) >= 20) {
            tr_idx <- match(tr_cells_lab, finite_cells); tr_idx <- tr_idx[!is.na(tr_idx)]
            te_idx2 <- match(te_cells_lab, finite_cells); te_idx2 <- te_idx2[!is.na(te_idx2)]

            k_eff <- .clamp_k(as.integer(k_lt), length(tr_idx))
            if (!is.na(k_eff) && k_eff >= 1) {

              lt_ra <- tryCatch(
                .knn_transfer(
                  Z_train = Zr2[tr_idx, , drop = FALSE],
                  y_train = y_finite[finite_cells[tr_idx]],
                  Z_test  = Za2[te_idx2, , drop = FALSE],
                  y_test  = y_finite[finite_cells[te_idx2]],
                  k = k_eff
                ),
                error = function(e) { .record_err("lt_rna_to_atac_error", e); NULL }
              )

              lt_ar <- tryCatch(
                .knn_transfer(
                  Z_train = Za2[tr_idx, , drop = FALSE],
                  y_train = y_finite[finite_cells[tr_idx]],
                  Z_test  = Zr2[te_idx2, , drop = FALSE],
                  y_test  = y_finite[finite_cells[te_idx2]],
                  k = k_eff
                ),
                error = function(e) { .record_err("lt_atac_to_rna_error", e); NULL }
              )

              acc_mean <- mean(c(lt_ra$acc %||% NA_real_, lt_ar$acc %||% NA_real_), na.rm = TRUE)
              f1_mean  <- mean(c(lt_ra$macroF1 %||% NA_real_, lt_ar$macroF1 %||% NA_real_), na.rm = TRUE)

              label_transfer_acc_mean_test <- acc_mean
              `PAIR(test)/Label transfer acc mean` <- acc_mean
              `PAIR(test)/Label transfer macroF1 mean` <- f1_mean
            }
          }
        }

      } else {
        extra$paired_skip_reason <- paste0(
          "Not enough paired finite TEST cells: finite=", length(finite_cells),
          " test_eff=", length(te_cells_eff)
        )
      }
    } else {
      extra$paired_skip_reason <- paste0("Not enough paired cells: common=", length(common_cells))
    }
  } else {
    extra$paired_skip_reason <- "Z_rna/Z_atac missing or missing rownames"
  }

  # -----------------------------
  # 2) fused embedding metrics on TEST
  # -----------------------------
  if (!is.null(Zf) && !is.null(rownames(Zf)) && !is.null(names(labels))) {
    zf_cells <- rownames(Zf)
    tr_eff <- intersect(tr_cells, zf_cells)
    te_eff <- intersect(te_cells, zf_cells)

    if (length(tr_eff) >= 50 && length(te_eff) >= 20) {
      fm2 <- tryCatch(
        .fused_metrics(Zf = Zf, labels_named = labels, tr_cells = tr_eff, te_cells = te_eff,
                      k = k_lt, silhouette_max_n = 3000L, seed = seed),
        error = function(e) { .record_err("fused_metrics_error", e); NULL }
      )
      if (!is.null(fm2)) {
        `Fused kNN acc` <- fm2$fused_knn_acc_test
        `Fused kNN macroF1` <- fm2$fused_knn_macroF1_test
        `Fused kNN balanced acc` <- fm2$fused_knn_balanced_acc_test
        `MIX(test)/Fused silhouette` <- fm2$fused_silhouette_test
        `MIX(test)/Fused label purity` <- fm2$fused_knn_label_purity_test
        `MIX(test)/Fused k-means ARI` <- fm2$fused_kmeans_ari_test
        `MIX(test)/Fused k-means NMI` <- fm2$fused_kmeans_nmi_test
      }
    }
  }

  if (!is.null(extra$foscttm_error) || !is.null(extra$mixing_pair_error) || !is.null(extra$fused_metrics_error)) {
    `__eval_error__` <- extra$foscttm_error %||% extra$mixing_pair_error %||% extra$fused_metrics_error
  }

  # IMPORTANT: return BOTH sets of columns
  data.table(
    method = name,
    failed = FALSE,
    fit_seconds = fit_seconds,
    transductive = as.numeric(isTRUE(transductive)),
    uses_labels  = as.numeric(isTRUE(uses_labels)),

    # legacy cols (your TSV currently expects these)
    FOSCTTM_mean_test = FOSCTTM_mean_test,
    FOSCTTM_sem_test  = FOSCTTM_sem_test,
    FOSCTTM_mrr_mean_test = FOSCTTM_mrr_mean_test,
    `FOSCTTM_recall@1_mean_test` = `FOSCTTM_recall@1_mean_test`,
    `FOSCTTM_recall@10_mean_test` = `FOSCTTM_recall@10_mean_test`,
    mixing_score_test = mixing_score_test,
    label_transfer_acc_mean_test = label_transfer_acc_mean_test,

    # canonical cols (optional)
    `PAIR(test)/FOSCTTM_mean` = `PAIR(test)/FOSCTTM_mean`,
    `PAIR(test)/FOSCTTM_MRR`  = `PAIR(test)/FOSCTTM_MRR`,
    `PAIR(test)/FOSCTTM_n_eval` = `PAIR(test)/FOSCTTM_n_eval`,
    `PAIR(test)/FOSCTTM_Recall@1` = `PAIR(test)/FOSCTTM_Recall@1`,
    `PAIR(test)/FOSCTTM_Recall@10` = `PAIR(test)/FOSCTTM_Recall@10`,
    `PAIR(test)/FOSCTTM_Recall@25` = `PAIR(test)/FOSCTTM_Recall@25`,
    `PAIR(test)/FOSCTTM_Recall@50` = `PAIR(test)/FOSCTTM_Recall@50`,
    `PAIR(test)/FOSCTTM_Recall@100` = `PAIR(test)/FOSCTTM_Recall@100`,
    `PAIR(test)/Label transfer acc mean` = `PAIR(test)/Label transfer acc mean`,
    `PAIR(test)/Label transfer macroF1 mean` = `PAIR(test)/Label transfer macroF1 mean`,
    `MIX(test)/Mixing score` = `MIX(test)/Mixing score`,
    `Fused kNN acc` = `Fused kNN acc`,
    `Fused kNN macroF1` = `Fused kNN macroF1`,
    `Fused kNN balanced acc` = `Fused kNN balanced acc`,
    `MIX(test)/Fused silhouette` = `MIX(test)/Fused silhouette`,
    `MIX(test)/Fused label purity` = `MIX(test)/Fused label purity`,
    `MIX(test)/Fused k-means ARI` = `MIX(test)/Fused k-means ARI`,
    `MIX(test)/Fused k-means NMI` = `MIX(test)/Fused k-means NMI`,
    `__eval_error__` = `__eval_error__`,

    extra_json = jsonlite::toJSON(extra, auto_unbox = TRUE),
    skipped = FALSE,
    reason = reason_in,
    method_str = method_str
  )
}


# %% [code]
build_split_cells <- function(splits, labels_named) {
  if (is.null(names(labels_named))) stop("labels_named must have names(cell IDs)")
  cn <- names(labels_named)

  to_cells <- function(x) {
    if (is.null(x)) return(character(0))
    if (is.numeric(x)) {
      x <- as.integer(x)
      x <- x[!is.na(x) & x >= 1L & x <= length(cn)]
      return(cn[x])
    }
    intersect(as.character(x), cn)
  }

  tr <- to_cells(splits$train)
  va <- to_cells(splits$val)
  te <- to_cells(splits$test)

  list(train_cells = tr, val_cells = va, test_cells = te)
}

.get_split_cells <- function(split_obj, which = c("train", "val", "test")) {
  which <- match.arg(which)
  if (is.null(split_obj)) stop("[split] split_obj is NULL")

  # Accept either a list with $cells or a list with $train/$val/$test directly
  if (!is.null(split_obj$cells) && !is.null(split_obj$cells[[which]])) {
    return(split_obj$cells[[which]])
  }
  if (!is.null(split_obj[[which]])) {
    return(split_obj[[which]])
  }

  stop(sprintf("[split] couldn't find cells for '%s' in split_obj", which))
}


# %% [code]


# %% [code]
library(data.table)

`%||%` <- if (exists("%||%", mode = "function")) get("%||%") else function(a, b) {
  if (is.null(a) || length(a) == 0) b else a
}

# ============================================================
# 0) Column schema (base + adds per-latent metrics)
# ============================================================
PY_EVAL_COLS_BASE <- c(
  "seed","fold","method","status","fit_seconds","error",
  "n_train","n_val","n_test","transductive","uses_labels",
  "PAIR(test)/FOSCTTM_mean","PAIR(test)/FOSCTTM_MRR","PAIR(test)/FOSCTTM_n_eval",
  "PAIR(test)/FOSCTTM_Recall@1","PAIR(test)/FOSCTTM_Recall@10","PAIR(test)/FOSCTTM_Recall@25",
  "PAIR(test)/FOSCTTM_Recall@50","PAIR(test)/FOSCTTM_Recall@100",
  "PAIR(test)/Label transfer acc mean","PAIR(test)/Label transfer macroF1 mean",
  "MIX(test)/Mixing score","MIX(test)/Mixing score (RNA)","MIX(test)/Mixing score (ATAC)",
  "MIX(test)/Fused silhouette","MIX(test)/Fused label purity","MIX(test)/Fused k-means ARI","MIX(test)/Fused k-means NMI",
  "Fused kNN acc","Fused kNN macroF1","Fused kNN balanced acc",
  "Fused(fusedZ,test) silhouette","Fused(fusedZ,test) label purity","Fused(fusedZ,test) k-means ARI","Fused(fusedZ,test) k-means NMI",
  "__eval_error__",
  "latent_dim","dropout","lr","weight_decay","batch_size","max_epochs","patience","reg","best_val","nbatches",
  "n_latent","n_latent_requested","joint_dim_returned",
  "atac_tfidf","patched_scipy_sparse_vstack_scoped","latent_cap","lr_used"
)

# NEW: explicit single-latent columns (so you can summarize/plot them)
PY_EVAL_COLS_SINGLE <- c(
  "RNA(test)/kNN acc","RNA(test)/kNN macroF1","RNA(test)/kNN balanced acc",
  "RNA(test)/silhouette","RNA(test)/label purity","RNA(test)/kmeans ARI","RNA(test)/kmeans NMI",
  "ATAC(test)/kNN acc","ATAC(test)/kNN macroF1","ATAC(test)/kNN balanced acc",
  "ATAC(test)/silhouette","ATAC(test)/label purity","ATAC(test)/kmeans ARI","ATAC(test)/kmeans NMI"
)

PY_EVAL_COLS <- c(PY_EVAL_COLS_BASE, PY_EVAL_COLS_SINGLE)

# ============================================================
# 1) basic utils
# ============================================================
.as_matrix <- function(x) {
  if (is.null(x)) return(NULL)
  if (inherits(x, "dgCMatrix")) x <- as.matrix(x)
  if (is.data.frame(x)) x <- as.matrix(x)
  if (!is.matrix(x)) x <- tryCatch(as.matrix(x), error = function(e) NULL)
  x
}

.ensure_rownames <- function(Z, cells_ref = NULL) {
  Z <- .as_matrix(Z)
  if (is.null(Z)) return(NULL)
  rn <- rownames(Z)
  if (!is.null(rn) && length(rn) == nrow(Z) && all(nzchar(rn))) return(Z)
  if (!is.null(cells_ref) && length(cells_ref) == nrow(Z)) {
    rownames(Z) <- as.character(cells_ref)
    return(Z)
  }
  rownames(Z) <- paste0("cell_", seq_len(nrow(Z)))
  Z
}

# Enforce consistent rownames for Zr/Za/Zf, using obj_cells as the source of truth
.harden_latents <- function(out, obj_cells) {
  out <- out %||% list()

  Zf <- out$Z_fused %||% out$Zf %||% out$latents$fused %||% NULL
  Zr <- out$Z_rna   %||% out$Zr %||% out$latents$rna   %||% NULL
  Za <- out$Z_atac  %||% out$Za %||% out$latents$atac  %||% NULL

  Zf <- .ensure_rownames(Zf, obj_cells)
  Zr <- .ensure_rownames(Zr, obj_cells)
  Za <- .ensure_rownames(Za, obj_cells)

  # If same nrow but essentially no overlap, assume same order and force names to match
  if (!is.null(Zr) && !is.null(Za) && nrow(Zr) == nrow(Za)) {
    if (length(intersect(rownames(Zr), rownames(Za))) < 10) rownames(Za) <- rownames(Zr)
  }
  if (!is.null(Zf) && !is.null(Zr) && nrow(Zf) == nrow(Zr)) {
    if (length(intersect(rownames(Zf), rownames(Zr))) < 10) rownames(Zf) <- rownames(Zr)
  }

  out$Z_fused <- Zf
  out$Z_rna   <- Zr
  out$Z_atac  <- Za
  out
}

.clamp_k <- function(k, n) {
  if (is.null(n) || n < 3) return(NA_integer_)
  as.integer(min(as.integer(k), as.integer(n) - 2L))
}

.is_finite_rows <- function(Z) {
  Z <- as.matrix(Z)
  if (nrow(Z) == 0) return(logical(0))
  rowSums(!is.finite(Z)) == 0L
}

.mode_chr <- function(x) {
  x <- as.character(x)
  x <- x[!is.na(x)]
  if (!length(x)) return(NA_character_)
  tab <- table(x)
  names(tab)[which.max(tab)]
}

.macro_f1 <- function(y_true, y_pred) {
  y_true <- as.character(y_true)
  y_pred <- as.character(y_pred)
  labs <- sort(unique(y_true[!is.na(y_true)]))
  if (!length(labs)) return(NA_real_)

  f1s <- vapply(labs, function(lv) {
    tp <- sum(y_true == lv & y_pred == lv, na.rm = TRUE)
    fp <- sum(y_true != lv & y_pred == lv, na.rm = TRUE)
    fn <- sum(y_true == lv & y_pred != lv, na.rm = TRUE)
    prec <- if ((tp + fp) > 0) tp / (tp + fp) else 0
    rec  <- if ((tp + fn) > 0) tp / (tp + fn) else 0
    if ((prec + rec) > 0) 2 * prec * rec / (prec + rec) else 0
  }, numeric(1))

  mean(f1s, na.rm = TRUE)
}

.balanced_acc <- function(y_true, y_pred) {
  y_true <- as.character(y_true)
  y_pred <- as.character(y_pred)
  labs <- sort(unique(y_true[!is.na(y_true)]))
  if (!length(labs)) return(NA_real_)

  recs <- vapply(labs, function(lv) {
    tp <- sum(y_true == lv & y_pred == lv, na.rm = TRUE)
    fn <- sum(y_true == lv & y_pred != lv, na.rm = TRUE)
    if ((tp + fn) > 0) tp / (tp + fn) else NA_real_
  }, numeric(1))

  mean(recs, na.rm = TRUE)
}

.safe <- function(expr, on_error = NULL) tryCatch(expr, error = function(e) on_error)

# Call fn but only pass supported named args (positional always passed)
.call_supported <- function(fn, ...) {
  dots <- list(...)
  nms  <- names(dots); if (is.null(nms)) nms <- rep("", length(dots))
  fmls <- tryCatch(names(formals(fn)), error = function(e) character(0))
  if (!length(fmls) || "..." %in% fmls) return(do.call(fn, dots))
  is_named <- nzchar(nms)
  pos <- dots[!is_named]
  nam <- dots[is_named]
  keep <- intersect(names(nam), fmls)
  do.call(fn, c(pos, nam[keep]))
}

# ------------------------------------------------------------
# NEW: kmeans helper (prevents "did not converge in 10 iterations")
# - increases iter.max and nstart
# - suppresses warnings (still returns km; you can inspect km$iter)
# - lightly standardizes columns to improve stability across embeddings
# ------------------------------------------------------------
.kmeans_safe <- function(X, centers, seed = 0L, nstart = 25L, iter.max = 100L) {
  X <- as.matrix(X)
  if (!is.finite(centers) || is.na(centers) || centers < 2L) return(NULL)
  if (nrow(X) < centers) return(NULL)

  ok <- rowSums(!is.finite(X)) == 0L
  if (!all(ok)) X <- X[ok, , drop = FALSE]
  if (nrow(X) < centers) return(NULL)

  sds <- apply(X, 2, stats::sd)
  sds[!is.finite(sds) | sds == 0] <- 1
  X <- scale(X, center = TRUE, scale = sds)

  set.seed(as.integer(seed))
  suppressWarnings(
    tryCatch(
      stats::kmeans(
        X,
        centers   = as.integer(centers),
        nstart    = as.integer(nstart),
        iter.max  = as.integer(iter.max),
        algorithm = "Hartigan-Wong"
      ),
      error = function(e) NULL
    )
  )
}

# ============================================================
# 2) Split mapping (FIXED: uses cell_ids as ground truth)
# ============================================================
build_split_cells <- function(splits, cell_ids, labels_named = NULL) {
  if (is.null(cell_ids) || !length(cell_ids)) stop("cell_ids required")
  cn <- as.character(cell_ids)

  to_cells <- function(x) {
    if (is.null(x)) return(character(0))
    if (is.numeric(x)) {
      x <- as.integer(x)
      x <- x[!is.na(x) & x >= 1L & x <= length(cn)]
      return(cn[x])
    }
    intersect(as.character(x), cn)
  }

  tr <- to_cells(splits$train)
  va <- to_cells(splits$val)
  te <- to_cells(splits$test)

  list(train_cells = tr, val_cells = va, test_cells = te)
}

# ============================================================
# 3) Metrics primitives
# ============================================================

# -----------------------------
# PAIRED FOSCTTM wrapper (robust)
# - supports foscttm_values(Za, Zb, idx_eval, subsample_n, seed)
# - ALWAYS computes MRR + Recall@K from ranks (even if foscttm_values doesn't return them)
# -----------------------------
.foscttm_metrics <- function(Z_rna, Z_atac, idx_eval, fos_sub = 3000L, seed = 0L) {
  if (is.null(Z_rna) || is.null(Z_atac)) stop("Z_rna/Z_atac required")
  Za <- as.matrix(Z_rna)
  Zb <- as.matrix(Z_atac)
  if (nrow(Za) != nrow(Zb)) stop("Z_rna/Z_atac nrow mismatch")
  n <- nrow(Za)
  if (n < 2) stop("Too few rows for FOSCTTM")

  idx_eval <- as.integer(idx_eval)
  idx_eval <- idx_eval[is.finite(idx_eval) & !is.na(idx_eval)]
  idx_eval <- idx_eval[idx_eval >= 1L & idx_eval <= n]
  idx_eval <- unique(idx_eval)
  if (length(idx_eval) < 2) stop("idx_eval too small after clamping")

  subsample_n <- as.integer(min(as.integer(fos_sub), length(idx_eval)))
  if (!is.finite(subsample_n) || is.na(subsample_n) || subsample_n < 2L) {
    stop("subsample_n too small after clamping")
  }

  # --- subsample eval indices for speed ---
  set.seed(as.integer(seed))
  if (subsample_n < length(idx_eval)) {
    idx_eval_sub <- sort(sample(idx_eval, size = subsample_n, replace = FALSE))
  } else {
    idx_eval_sub <- idx_eval
  }

  # ------------------------------------------------------------
  # 1) Try calling foscttm_values() for the raw FOSCTTM vector (optional)
  # ------------------------------------------------------------
  vec <- NULL
  if (exists("foscttm_values", mode = "function")) {
    f <- get("foscttm_values", mode = "function")
    out <- tryCatch(
      f(Za = Za, Zb = Zb, idx_eval = idx_eval_sub,
        subsample_n = as.integer(length(idx_eval_sub)),
        seed = as.integer(seed)),
      error = function(e) e
    )

    if (!inherits(out, "error")) {
      if (is.numeric(out)) {
        vec <- as.numeric(out)
      } else if (is.list(out)) {
        for (cand in c("foscttm", "foscttm_values", "values")) {
          if (!is.null(out[[cand]]) && is.numeric(out[[cand]])) {
            vec <- as.numeric(out[[cand]])
            break
          }
        }
      }
    }
  }

  # ------------------------------------------------------------
  # 2) Compute ranks directly (this gives MRR + Recall@K reliably)
  # ------------------------------------------------------------
  Za_e <- Za[idx_eval_sub, , drop = FALSE]
  Zb_all <- Zb

  a2 <- rowSums(Za_e^2)
  b2 <- rowSums(Zb_all^2)

  G <- Za_e %*% t(Zb_all)
  D2 <- outer(a2, b2, "+") - 2 * G
  D2[!is.finite(D2)] <- Inf

  ranks <- integer(nrow(D2))
  for (ii in seq_len(nrow(D2))) {
    true_j <- idx_eval_sub[ii]
    d_true <- D2[ii, true_j]
    ranks[ii] <- 1L + sum(D2[ii, ] < d_true, na.rm = TRUE)
  }

  # ------------------------------------------------------------
  # 3) If foscttm_values didn't give vec, derive it from rank
  # ------------------------------------------------------------
  if (is.null(vec) || !length(vec)) {
    vec <- (ranks - 1) / (n - 1)
  } else {
    vec <- vec[is.finite(vec)]
    if (length(vec) != length(ranks)) {
      vec <- (ranks - 1) / (n - 1)
    }
  }

  mu  <- mean(vec, na.rm = TRUE)
  sem <- stats::sd(vec, na.rm = TRUE) / sqrt(max(1L, sum(is.finite(vec))))

  mrr <- mean(1 / ranks, na.rm = TRUE)
  recall_k <- function(k) mean(ranks <= k, na.rm = TRUE)

  list(
    mean = mu,
    sem  = sem,
    mrr_mean = mrr,
    recall1   = recall_k(1L),
    recall10  = recall_k(10L),
    recall25  = recall_k(25L),
    recall50  = recall_k(50L),
    recall100 = recall_k(100L),
    n_eval    = length(ranks)
  )
}

# --- PAIRED mixing (self-contained)
.mixing_score_paired <- function(Z_rna, Z_atac, idx_eval, k = 50L) {
  if (is.null(Z_rna) || is.null(Z_atac)) return(NA_real_)
  if (!requireNamespace("RANN", quietly = TRUE)) return(NA_real_)

  Zr <- as.matrix(Z_rna)
  Za <- as.matrix(Z_atac)
  if (nrow(Zr) != nrow(Za) || nrow(Zr) < 10) return(NA_real_)

  idx_eval <- as.integer(idx_eval)
  idx_eval <- idx_eval[is.finite(idx_eval) & !is.na(idx_eval)]
  idx_eval <- idx_eval[idx_eval >= 1L & idx_eval <= nrow(Zr)]
  idx_eval <- unique(idx_eval)
  if (length(idx_eval) < 5) return(NA_real_)

  n <- nrow(Zr)
  k_eff <- .clamp_k(as.integer(k), 2L * n)
  if (is.na(k_eff) || k_eff < 5) return(NA_real_)

  Zpool <- rbind(Zr, Za)
  q_rna  <- idx_eval
  q_atac <- idx_eval + n

  nn_r <- RANN::nn2(data = Zpool, query = Zpool[q_rna, , drop = FALSE], k = k_eff)$nn.idx
  nn_a <- RANN::nn2(data = Zpool, query = Zpool[q_atac, , drop = FALSE], k = k_eff)$nn.idx

  is_atac <- function(j) j > n
  is_rna  <- function(j) j <= n

  frac_r <- vapply(seq_len(nrow(nn_r)), function(ii) mean(is_atac(nn_r[ii, ])), numeric(1))
  frac_a <- vapply(seq_len(nrow(nn_a)), function(ii) mean(is_rna(nn_a[ii, ])), numeric(1))

  mean(c(frac_r, frac_a), na.rm = TRUE)
}

# --- kNN label transfer
.knn_transfer <- function(Z_train, y_train, Z_test, y_test, k = 15L) {
  if (!requireNamespace("RANN", quietly = TRUE)) {
    return(list(acc = NA_real_, macroF1 = NA_real_, y_pred = NULL, y_true = NULL))
  }
  Z_train <- as.matrix(Z_train); Z_test <- as.matrix(Z_test)

  ok_tr <- .is_finite_rows(Z_train)
  ok_te <- .is_finite_rows(Z_test)
  Z_train <- Z_train[ok_tr, , drop = FALSE]
  y_train <- as.character(y_train[ok_tr])
  Z_test  <- Z_test[ok_te, , drop = FALSE]
  y_test2 <- as.character(y_test[ok_te])

  if (nrow(Z_train) < 10 || nrow(Z_test) < 5) return(list(acc = NA_real_, macroF1 = NA_real_, y_pred = NULL, y_true = NULL))
  k_eff <- .clamp_k(as.integer(k), nrow(Z_train))
  if (is.na(k_eff) || k_eff < 1) return(list(acc = NA_real_, macroF1 = NA_real_, y_pred = NULL, y_true = NULL))

  nn <- RANN::nn2(data = Z_train, query = Z_test, k = k_eff)$nn.idx
  y_pred <- apply(nn, 1, function(ii) .mode_chr(y_train[ii]))

  list(
    acc = mean(y_pred == y_test2, na.rm = TRUE),
    macroF1 = .macro_f1(y_test2, y_pred),
    y_pred = y_pred,
    y_true = y_test2
  )
}

# --- Single-latent metrics on TEST (kNN, purity, silhouette, kmeans ARI/NMI)
.single_latent_metrics <- function(Z, labels_named, tr_cells, te_cells, k = 15L, silhouette_max_n = 3000L, seed = 0L) {
  out <- list(
    knn_acc = NA_real_, knn_macroF1 = NA_real_, knn_balanced_acc = NA_real_,
    silhouette = NA_real_, purity = NA_real_, kmeans_ari = NA_real_, kmeans_nmi = NA_real_
  )
  if (is.null(Z) || is.null(rownames(Z)) || is.null(labels_named) || is.null(names(labels_named))) return(out)

  Z <- as.matrix(Z)
  common <- intersect(rownames(Z), names(labels_named))
  if (length(common) < 50) return(out)

  Z <- Z[common, , drop = FALSE]
  y <- as.character(labels_named[common]); names(y) <- common

  ok <- .is_finite_rows(Z)
  if (!all(ok)) { Z <- Z[ok, , drop = FALSE]; y <- y[rownames(Z)] }

  tr <- intersect(tr_cells, rownames(Z))
  te <- intersect(te_cells, rownames(Z))
  if (length(tr) < 50 || length(te) < 20) return(out)

  y_tr <- y[tr]; y_te <- y[te]
  ok_tr <- !is.na(y_tr); ok_te <- !is.na(y_te)
  tr <- tr[ok_tr]; te <- te[ok_te]
  if (length(tr) < 50 || length(te) < 20) return(out)

  Ztr <- Z[tr, , drop = FALSE]; Zte <- Z[te, , drop = FALSE]
  ytr <- y[tr]; yte <- y[te]

  kt <- .knn_transfer(Ztr, ytr, Zte, yte, k = k)
  out$knn_acc <- kt$acc
  out$knn_macroF1 <- kt$macroF1
  if (!is.null(kt$y_pred) && !is.null(kt$y_true)) out$knn_balanced_acc <- .balanced_acc(kt$y_true, kt$y_pred)

  # purity on TEST (self-kNN)
  if (requireNamespace("RANN", quietly = TRUE) && nrow(Zte) >= 10) {
    k_eff <- .clamp_k(as.integer(k) + 1L, nrow(Zte))
    if (!is.na(k_eff) && k_eff >= 3) {
      nn <- RANN::nn2(data = Zte, query = Zte, k = k_eff)$nn.idx
      nn <- nn[, -1, drop = FALSE]
      pur <- vapply(seq_len(nrow(nn)), function(i) mean(yte[nn[i, ]] == yte[i], na.rm = TRUE), numeric(1))
      out$purity <- mean(pur, na.rm = TRUE)
    }
  }

  # silhouette on TEST
  if (requireNamespace("cluster", quietly = TRUE)) {
    set.seed(as.integer(seed))
    te_use <- te
    if (length(te_use) > silhouette_max_n) te_use <- sample(te_use, size = silhouette_max_n, replace = FALSE)
    ys <- y[te_use]; Zs <- Z[te_use, , drop = FALSE]
    oky <- !is.na(ys); ys <- ys[oky]; Zs <- Zs[oky, , drop = FALSE]
    if (length(unique(ys)) >= 2 && nrow(Zs) >= 10) {
      D <- stats::dist(Zs)
      sil <- cluster::silhouette(as.integer(as.factor(ys)), D)
      out$silhouette <- mean(sil[, "sil_width"], na.rm = TRUE)
    }
  }

  # kmeans ARI/NMI on TEST (FIXED: safer k + stable kmeans)
  k_tr <- length(unique(ytr[!is.na(ytr)]))
  k_te <- length(unique(yte[!is.na(yte)]))
  k_clust <- min(k_tr, k_te, nrow(Zte) - 1L)

  if (is.finite(k_clust) && k_clust >= 2 && length(unique(yte)) >= 2) {
    km <- .kmeans_safe(Zte, centers = k_clust, seed = seed, nstart = 25L, iter.max = 100L)
    if (!is.null(km)) {
      if (requireNamespace("mclust", quietly = TRUE)) {
        out$kmeans_ari <- mclust::adjustedRandIndex(as.integer(as.factor(yte)), as.integer(km$cluster))
      }
      if (requireNamespace("aricode", quietly = TRUE)) {
        out$kmeans_nmi <- aricode::NMI(as.integer(as.factor(yte)), as.integer(km$cluster))
      }
    }
  }

  out
}

# ============================================================
# 4) Fused metrics (kept same; only kmeans made robust)
# ============================================================
.fused_metrics <- function(Zf, labels_named, tr_cells, te_cells,
                           k = 15L, silhouette_max_n = 3000L, seed = 0L) {
  out <- list(
    fused_knn_acc_test = NA_real_,
    fused_knn_macroF1_test = NA_real_,
    fused_knn_balanced_acc_test = NA_real_,
    fused_silhouette_test = NA_real_,
    fused_knn_label_purity_test = NA_real_,
    fused_kmeans_ari_test = NA_real_,
    fused_kmeans_nmi_test = NA_real_
  )
  if (is.null(Zf) || is.null(labels_named) || is.null(names(labels_named))) return(out)
  Zf <- as.matrix(Zf)
  if (is.null(rownames(Zf))) return(out)

  common <- intersect(rownames(Zf), names(labels_named))
  if (length(common) < 50) return(out)
  Zf <- Zf[common, , drop = FALSE]
  y  <- as.character(labels_named[common]); names(y) <- common

  ok <- .is_finite_rows(Zf)
  if (!all(ok)) { Zf <- Zf[ok, , drop = FALSE]; y <- y[rownames(Zf)] }

  tr <- intersect(tr_cells, rownames(Zf))
  te <- intersect(te_cells, rownames(Zf))
  if (length(tr) < 20 || length(te) < 20) return(out)

  Z_train <- Zf[tr, , drop = FALSE]; y_train <- y[tr]
  Z_test  <- Zf[te, , drop = FALSE]; y_test  <- y[te]

  kt <- .knn_transfer(Z_train, y_train, Z_test, y_test, k = k)
  out$fused_knn_acc_test <- kt$acc
  out$fused_knn_macroF1_test <- kt$macroF1
  if (!is.null(kt$y_pred) && !is.null(kt$y_true)) out$fused_knn_balanced_acc_test <- .balanced_acc(kt$y_true, kt$y_pred)

  # purity (TEST)
  if (requireNamespace("RANN", quietly = TRUE) && nrow(Z_test) >= 10) {
    k_eff <- .clamp_k(as.integer(k) + 1L, nrow(Z_test))
    if (!is.na(k_eff) && k_eff >= 3) {
      nn <- RANN::nn2(data = Z_test, query = Z_test, k = k_eff)$nn.idx
      nn <- nn[, -1, drop = FALSE]
      purity <- vapply(seq_len(nrow(nn)), function(i) mean(y_test[nn[i, ]] == y_test[i], na.rm = TRUE), numeric(1))
      out$fused_knn_label_purity_test <- mean(purity, na.rm = TRUE)
    }
  }

  # silhouette (TEST)
  if (requireNamespace("cluster", quietly = TRUE)) {
    set.seed(seed)
    te_use <- te
    if (length(te_use) > silhouette_max_n) te_use <- sample(te_use, size = silhouette_max_n, replace = FALSE)
    y_s <- y[te_use]; Z_s <- Zf[te_use, , drop = FALSE]
    oky <- !is.na(y_s); y_s <- y_s[oky]; Z_s <- Z_s[oky, , drop = FALSE]
    if (length(unique(y_s)) >= 2 && nrow(Z_s) >= 10) {
      D <- stats::dist(Z_s)
      sil <- cluster::silhouette(as.integer(as.factor(y_s)), D)
      out$fused_silhouette_test <- mean(sil[, "sil_width"], na.rm = TRUE)
    }
  }

  # kmeans ARI/NMI (TEST) (FIXED: safer k + stable kmeans)
  yt <- y_test
  oky <- !is.na(yt)
  Zk <- Z_test[oky, , drop = FALSE]
  yt <- yt[oky]

  k_tr <- length(unique(y_train[!is.na(y_train)]))
  k_te <- length(unique(yt[!is.na(yt)]))
  k_clust <- min(k_tr, k_te, nrow(Zk) - 1L)

  if (is.finite(k_clust) && k_clust >= 2 && nrow(Zk) >= k_clust && length(unique(yt)) >= 2) {
    km <- .kmeans_safe(Zk, centers = k_clust, seed = seed, nstart = 25L, iter.max = 100L)
    if (!is.null(km)) {
      if (requireNamespace("mclust", quietly = TRUE)) {
        out$fused_kmeans_ari_test <- mclust::adjustedRandIndex(as.integer(as.factor(yt)), as.integer(km$cluster))
      }
      if (requireNamespace("aricode", quietly = TRUE)) {
        out$fused_kmeans_nmi_test <- aricode::NMI(as.integer(as.factor(yt)), as.integer(km$cluster))
      }
    }
  }

  out
}

# ============================================================
# 5) Evaluate one method -> returns PYTHON-SCHEMA row (unchanged)
# ============================================================
evaluate_one_python_schema <- function(
  method_name,
  out,
  labels,
  splits,
  cell_ids,
  seed,
  fold,
  status = "ok",
  error_msg = NA_character_,
  fit_seconds = NA_real_,
  k_mix = 30L,
  k_lt  = 15L,
  fos_sub = 3000L,
  verbose = FALSE
) {
  .msg <- function(...) if (isTRUE(verbose)) message(...)

  # Ensure labels are named by cell_ids
  if (!is.null(labels)) {
    if (is.null(names(labels))) {
      if (length(labels) == length(cell_ids)) names(labels) <- cell_ids
    }
  }

  sp <- tryCatch(build_split_cells(splits, cell_ids = cell_ids, labels_named = labels),
                 error = function(e) {
                   list(train_cells=character(0), val_cells=character(0), test_cells=character(0),
                        .err=paste0("split_build_error: ", conditionMessage(e)))
                 })
  tr_cells <- sp$train_cells %||% character(0)
  va_cells <- sp$val_cells   %||% character(0)
  te_cells <- sp$test_cells  %||% character(0)

  row <- as.list(rep(NA, length(PY_EVAL_COLS)))
  names(row) <- PY_EVAL_COLS

  row$seed <- as.integer(seed)
  row$fold <- as.integer(fold)
  row$method <- as.character(method_name)
  row$status <- as.character(status)
  row$fit_seconds <- as.numeric(fit_seconds)
  row$error <- as.character(error_msg)

  row$n_train <- as.integer(length(tr_cells))
  row$n_val   <- as.integer(length(va_cells))
  row$n_test  <- as.integer(length(te_cells))

  extra <- out$extra_json %||% out$extra %||% list()
  row$transductive <- as.numeric(isTRUE(extra$transductive %||% out$transductive %||% FALSE))
  row$uses_labels  <- as.numeric(isTRUE(extra$uses_labels  %||% out$uses_labels  %||% FALSE))

  out2 <- .harden_latents(out, obj_cells = cell_ids)
  Zf <- out2$Z_fused
  Zr <- out2$Z_rna
  Za <- out2$Z_atac

  eval_errs <- character(0)
  .record <- function(tag, e) eval_errs <<- c(eval_errs, paste0(tag, ": ", conditionMessage(e)))

  # ----------------------------------------------------------
  # A) PAIRED metrics (on TEST overlap)
  # ----------------------------------------------------------
  if (!is.null(Zr) && !is.null(Za) && !is.null(rownames(Zr)) && !is.null(rownames(Za))) {
    common <- intersect(rownames(Zr), rownames(Za))
    te_eff <- intersect(te_cells, common)

    if (length(te_eff) >= 50) {
      Zr_te <- Zr[te_eff, , drop = FALSE]
      Za_te <- Za[te_eff, , drop = FALSE]

      row[["MIX(test)/Mixing score"]] <- tryCatch(
        .mixing_score_paired(Zr_te, Za_te, idx_eval = seq_len(nrow(Zr_te)), k = as.integer(k_mix)),
        error = function(e) { .record("mixing_pair_error", e); NA_real_ }
      )

      fm <- tryCatch(
        .foscttm_metrics(Zr_te, Za_te, idx_eval = seq_len(nrow(Zr_te)),
                         fos_sub = as.integer(min(fos_sub, nrow(Zr_te))), seed = as.integer(seed)),
        error = function(e) { .record("foscttm_error", e); NULL }
      )
      if (!is.null(fm)) {
        row[["PAIR(test)/FOSCTTM_mean"]]       <- as.numeric(fm$mean %||% NA_real_)
        row[["PAIR(test)/FOSCTTM_MRR"]]        <- as.numeric(fm$mrr_mean %||% NA_real_)
        row[["PAIR(test)/FOSCTTM_n_eval"]]     <- as.numeric(fm$n_eval %||% NA_real_)
        row[["PAIR(test)/FOSCTTM_Recall@1"]]   <- as.numeric(fm$recall1 %||% NA_real_)
        row[["PAIR(test)/FOSCTTM_Recall@10"]]  <- as.numeric(fm$recall10 %||% NA_real_)
        row[["PAIR(test)/FOSCTTM_Recall@25"]]  <- as.numeric(fm$recall25 %||% NA_real_)
        row[["PAIR(test)/FOSCTTM_Recall@50"]]  <- as.numeric(fm$recall50 %||% NA_real_)
        row[["PAIR(test)/FOSCTTM_Recall@100"]] <- as.numeric(fm$recall100 %||% NA_real_)
      }

      # CROSS-MODALITY label transfer
      if (!is.null(labels) && !is.null(names(labels))) {
        common_lab <- common[!is.na(labels[common])]
        tr_eff <- intersect(tr_cells, common_lab)
        te_eff2 <- intersect(te_cells, common_lab)

        if (length(tr_eff) >= 50 && length(te_eff2) >= 20) {
          k_eff <- .clamp_k(as.integer(k_lt), length(tr_eff))

          lt_r2a <- tryCatch(
            .knn_transfer(
              Z_train = Zr[tr_eff, , drop = FALSE],
              y_train = labels[tr_eff],
              Z_test  = Za[te_eff2, , drop = FALSE],
              y_test  = labels[te_eff2],
              k = k_eff
            ),
            error = function(e) { .record("lt_rna_to_mod_error", e); NULL }
          )

          lt_a2r <- tryCatch(
            .knn_transfer(
              Z_train = Za[tr_eff, , drop = FALSE],
              y_train = labels[tr_eff],
              Z_test  = Zr[te_eff2, , drop = FALSE],
              y_test  = labels[te_eff2],
              k = k_eff
            ),
            error = function(e) { .record("lt_mod_to_rna_error", e); NULL }
          )

          acc_mean <- mean(c(lt_r2a$acc %||% NA_real_, lt_a2r$acc %||% NA_real_), na.rm = TRUE)
          f1_mean  <- mean(c(lt_r2a$macroF1 %||% NA_real_, lt_a2r$macroF1 %||% NA_real_), na.rm = TRUE)

          row[["PAIR(test)/Label transfer acc mean"]]     <- as.numeric(acc_mean)
          row[["PAIR(test)/Label transfer macroF1 mean"]] <- as.numeric(f1_mean)
        }
      }
    } else {
      .msg("Paired eval skipped: too few paired TEST cells (", length(te_eff), ")")
    }
  } else {
    .msg("Paired eval skipped: missing Z_rna/Z_atac or missing rownames()")
  }

  # ----------------------------------------------------------
  # B) FUSED metrics
  # ----------------------------------------------------------
  if (!is.null(Zf) && !is.null(rownames(Zf)) && !is.null(labels) && !is.null(names(labels))) {
    tr_eff <- intersect(tr_cells, rownames(Zf))
    te_eff <- intersect(te_cells, rownames(Zf))
    if (length(tr_eff) >= 50 && length(te_eff) >= 20) {
      fm2 <- tryCatch(
        .fused_metrics(
          Zf = Zf, labels_named = labels, tr_cells = tr_eff, te_cells = te_eff,
          k = as.integer(k_lt), silhouette_max_n = 3000L, seed = as.integer(seed)
        ),
        error = function(e) { .record("fused_metrics_error", e); NULL }
      )
      if (!is.null(fm2)) {
        row[["Fused kNN acc"]]                 <- as.numeric(fm2$fused_knn_acc_test %||% NA_real_)
        row[["Fused kNN macroF1"]]             <- as.numeric(fm2$fused_knn_macroF1_test %||% NA_real_)
        row[["Fused kNN balanced acc"]]        <- as.numeric(fm2$fused_knn_balanced_acc_test %||% NA_real_)
        row[["MIX(test)/Fused silhouette"]]    <- as.numeric(fm2$fused_silhouette_test %||% NA_real_)
        row[["MIX(test)/Fused label purity"]]  <- as.numeric(fm2$fused_knn_label_purity_test %||% NA_real_)
        row[["MIX(test)/Fused k-means ARI"]]   <- as.numeric(fm2$fused_kmeans_ari_test %||% NA_real_)
        row[["MIX(test)/Fused k-means NMI"]]   <- as.numeric(fm2$fused_kmeans_nmi_test %||% NA_real_)

        row[["Fused(fusedZ,test) silhouette"]]    <- row[["MIX(test)/Fused silhouette"]]
        row[["Fused(fusedZ,test) label purity"]]  <- row[["MIX(test)/Fused label purity"]]
        row[["Fused(fusedZ,test) k-means ARI"]]   <- row[["MIX(test)/Fused k-means ARI"]]
        row[["Fused(fusedZ,test) k-means NMI"]]   <- row[["MIX(test)/Fused k-means NMI"]]
      }
    }
  } else {
    .msg("Fused eval skipped: missing Z_fused or missing labels names()")
  }

  # ----------------------------------------------------------
  # C) SINGLE-LATENT metrics (LABEL-BASED)
  # ----------------------------------------------------------
  # NOTE:
  # - .single_latent_metrics() returns label-based quality metrics (kNN transfer, silhouette by label, purity, kmeans ARI/NMI)
  # - It does NOT (and cannot) compute "RNA-only" or "ATAC-only" *modality mixing* because each embedding is single-modality.
  # - Cross-modality mixing is computed separately as MIX(test)/Mixing score using BOTH Zr and Za.
  #
  # Therefore: DO NOT write purity into MIX(test)/Mixing score (RNA/ATAC).
  row[["MIX(test)/Mixing score (RNA)"]]  <- NA_real_
  row[["MIX(test)/Mixing score (ATAC)"]] <- NA_real_

  if (!is.null(labels) && !is.null(names(labels))) {

    if (!is.null(Zr) && !is.null(rownames(Zr))) {
      m_r <- tryCatch(
        .single_latent_metrics(
          Z = Zr,
          labels_named = labels,
          tr_cells = tr_cells,
          te_cells = te_cells,
          k = as.integer(k_lt),
          seed = as.integer(seed)
        ),
        error = function(e) { .record("single_rna_error", e); NULL }
      )

      if (!is.null(m_r)) {
        row[["RNA(test)/kNN acc"]]           <- as.numeric(m_r$knn_acc)
        row[["RNA(test)/kNN macroF1"]]       <- as.numeric(m_r$knn_macroF1)
        row[["RNA(test)/kNN balanced acc"]]  <- as.numeric(m_r$knn_balanced_acc)
        row[["RNA(test)/silhouette"]]        <- as.numeric(m_r$silhouette)
        row[["RNA(test)/label purity"]]      <- as.numeric(m_r$purity)
        row[["RNA(test)/kmeans ARI"]]        <- as.numeric(m_r$kmeans_ari)
        row[["RNA(test)/kmeans NMI"]]        <- as.numeric(m_r$kmeans_nmi)
      }
    }

    if (!is.null(Za) && !is.null(rownames(Za))) {
      m_a <- tryCatch(
        .single_latent_metrics(
          Z = Za,
          labels_named = labels,
          tr_cells = tr_cells,
          te_cells = te_cells,
          k = as.integer(k_lt),
          seed = as.integer(seed)
        ),
        error = function(e) { .record("single_atac_error", e); NULL }
      )

      if (!is.null(m_a)) {
        row[["ATAC(test)/kNN acc"]]          <- as.numeric(m_a$knn_acc)
        row[["ATAC(test)/kNN macroF1"]]      <- as.numeric(m_a$knn_macroF1)
        row[["ATAC(test)/kNN balanced acc"]] <- as.numeric(m_a$knn_balanced_acc)
        row[["ATAC(test)/silhouette"]]       <- as.numeric(m_a$silhouette)
        row[["ATAC(test)/label purity"]]     <- as.numeric(m_a$purity)
        row[["ATAC(test)/kmeans ARI"]]       <- as.numeric(m_a$kmeans_ari)
        row[["ATAC(test)/kmeans NMI"]]       <- as.numeric(m_a$kmeans_nmi)
      }
    }
  }

  # ----------------------------------------------------------
  # D) Hyperparam fields if present
  # ----------------------------------------------------------
  hp <- extra %||% list()
  for (k in c("latent_dim","dropout","lr","weight_decay","batch_size","max_epochs","patience","reg",
             "best_val","nbatches","n_latent","n_latent_requested","joint_dim_returned",
             "atac_tfidf","patched_scipy_sparse_vstack_scoped","latent_cap","lr_used")) {
    if (!is.null(hp[[k]])) row[[k]] <- hp[[k]]
  }

  row[["__eval_error__"]] <- if (length(eval_errs)) paste(eval_errs, collapse=" | ") else NA_character_
  if (!is.null(sp$.err)) row[["__eval_error__"]] <- paste(row[["__eval_error__"]] %||% "", sp$.err, sep=" | ")

  data.table::as.data.table(row)[, ..PY_EVAL_COLS]
}

# ============================================================
# 6) Wrapper you call per run
# ============================================================
run_and_eval_one <- function(
  method_name,
  res_out,
  obj_cells,
  labels,
  splits_idx,
  seed,
  fold,
  status = "ok",
  error_msg = NA_character_,
  fit_seconds = NA_real_,
  k_mix = 30L,
  k_lt  = 15L,
  fos_sub = 3000L,
  verbose = FALSE
) {
  evaluate_one_python_schema(
    method_name = method_name,
    out = res_out,
    labels = labels,
    splits = splits_idx,
    cell_ids = obj_cells,
    seed = seed,
    fold = fold,
    status = status,
    error_msg = error_msg,
    fit_seconds = fit_seconds,
    k_mix = k_mix,
    k_lt  = k_lt,
    fos_sub = fos_sub,
    verbose = verbose
  )
}


# %% [code]


# %% [markdown]
### Run CV/seed sweeps using the same seeds/folds as the PyTorch tool Multiome evaluation

# %% [code]
# -----------------------------
# 1) make stratified folds
# -----------------------------
make_stratified_folds <- function(labels, k = 3, seed = 0) {
  set.seed(seed)
  y <- as.character(labels)
  n <- length(y)

  # if only one class (or all NA), fall back to random K-fold
  uy <- unique(y[!is.na(y)])
  if (length(uy) < 2) {
    idx <- sample(seq_len(n))
    fold_id <- rep(seq_len(k), length.out = n)
    out <- split(idx, fold_id)  # idx already permuted
    return(lapply(out, function(v) sort(unique(as.integer(v)))))
  }

  # stratify within each class
  by_class <- split(seq_len(n), y)
  folds <- vector("list", k)
  for (kk in seq_len(k)) folds[[kk]] <- integer(0)

  for (cls in names(by_class)) {
    ii <- sample(by_class[[cls]])
    parts <- split(ii, rep(seq_len(k), length.out = length(ii)))
    for (kk in seq_len(k)) {
      add <- parts[[as.character(kk)]]
      if (!is.null(add)) folds[[kk]] <- c(folds[[kk]], add)
    }
  }

  lapply(folds, function(v) sort(unique(as.integer(v))))
}

fold_splits_from_folds <- function(n, folds, fold_idx, seed=0, val_frac_of_non_test=0.10) {
  te <- folds[[fold_idx]]
  non_test <- setdiff(seq_len(n), te)

  set.seed(seed + 1000L * as.integer(fold_idx))

  n_val <- max(1L, floor(val_frac_of_non_test * length(non_test)))
  va <- sample(non_test, size = n_val, replace = FALSE)
  tr <- setdiff(non_test, va)

  list(train = tr, val = va, test = te)
}


# %% [code]
.get_split_cells <- function(obj, splits = NULL, labels = NULL, verbose = FALSE) {
  .msg <- function(...) if (isTRUE(verbose)) cat("[.get_split_cells]", ..., "\n")

  cn <- Seurat::Cells(obj)
  n  <- length(cn)

  # --- normalize labels to a named vector over cn (if provided) ---
  labels_named <- NULL
  if (!is.null(labels)) {
    if (is.null(names(labels))) {
      if (length(labels) != n) stop("[.get_split_cells] labels has no names and length != n_cells")
      labels_named <- labels
      names(labels_named) <- cn
    } else {
      # reorder to cn where possible
      labels_named <- labels[intersect(cn, names(labels))]
      # if labels contains all cn, reorder to cn
      if (all(cn %in% names(labels))) labels_named <- labels[cn]
    }
  }

  # If no splits, fold is all cells. Also return a "splits" object = NULL.
  if (is.null(splits)) {
    .msg("no splits; fold = all cells (n=", n, ")")
    return(list(
      train_cells = NULL, val_cells = NULL, test_cells = NULL,
      fold_cells  = cn,
      labels      = labels_named,
      splits      = NULL
    ))
  }

  # Convert split spec (indices or cell IDs) -> cell IDs
  to_cells <- function(x) {
    if (is.null(x)) return(character(0))
    if (is.logical(x)) {
      x <- which(isTRUE(x))
      x <- as.integer(x)
    }
    if (is.numeric(x)) {
      x <- as.integer(x)
      x <- x[!is.na(x)]
      x <- x[x >= 1 & x <= n]
      return(cn[x])
    }
    # assume character
    intersect(as.character(x), cn)
  }

  tr <- to_cells(splits$train)
  va <- to_cells(splits$val)
  te <- to_cells(splits$test)

  fold <- unique(c(tr, va, te))
  if (length(fold) == 0) {
    fold <- cn
    .msg("splits parsed to empty; using fold=all cells")
  }

  # Canonical "splits indices into labels order":
  # We define the label/split universe order to be fold_cells.
  # (This also matches how you want latents reindexed.)
  fold_cells <- fold

  splits_out <- list(
    train = match(tr, fold_cells),
    val   = match(va, fold_cells),
    test  = match(te, fold_cells)
  )
  splits_out$train <- splits_out$train[!is.na(splits_out$train)]
  splits_out$val   <- splits_out$val[!is.na(splits_out$val)]
  splits_out$test  <- splits_out$test[!is.na(splits_out$test)]

  .msg(sprintf("parsed: train=%d val=%d test=%d fold=%d (n_cells=%d)",
               length(tr), length(va), length(te), length(fold_cells), n))

  # If labels were provided, subset/reorder to fold_cells
  if (!is.null(labels_named)) {
    if (!all(fold_cells %in% names(labels_named))) {
      # allow missing labels, but keep names aligned where present
      labels_fold <- labels_named[intersect(fold_cells, names(labels_named))]
      # make a full vector aligned to fold_cells with NAs where missing
      tmp <- rep(NA, length(fold_cells)); names(tmp) <- fold_cells
      tmp[names(labels_fold)] <- labels_fold
      labels_named <- tmp
    } else {
      labels_named <- labels_named[fold_cells]
    }
  }

  list(
    train_cells = tr, val_cells = va, test_cells = te,
    fold_cells  = fold_cells,
    labels      = labels_named,
    splits      = splits_out
  )
}


# %% [code]
# ------------------------------------------------------------
# Helper: call fn but only pass named args it supports
# (positional args are not used here; we pass named only)
# ------------------------------------------------------------
.call_supported_named <- function(fn, ...) {
  dots <- list(...)
  fml  <- tryCatch(names(formals(fn)), error = function(e) character(0))
  if (!length(fml) || "..." %in% fml) return(do.call(fn, dots))
  keep <- intersect(names(dots), fml)
  do.call(fn, dots[keep])
}

# ------------------------------------------------------------
# Standard CV wrapper signature:
# function(obj, splits, fold_cells, labels, hvgs, ga_features, seed, verbose, ...)
# Every method MUST accept these, even if it ignores most of them.
# ------------------------------------------------------------
METHODS <- list(

  seurat_wnn = function(
    obj, splits = NULL, fold_cells = NULL, labels = NULL, hvgs = NULL, ga_features = NULL,
    seed = 0, verbose = FALSE, ...
  ) {
    set.seed(as.integer(seed))
    # DO NOT forward ... unless run_seurat_wnn supports it
    run_seurat_wnn(
      obj = obj,
      latent_dim = 30,
      rna_red = "pca",
      atac_red = "lsi",
      rna_dims = 1:100,
      atac_dims = 2:101,
      splits = splits,
      fold_cells = fold_cells,
      verbose = verbose
    )
  },

  seurat_cca = function(
    obj, splits = NULL, fold_cells = NULL, labels = NULL, hvgs = NULL, ga_features = NULL,
    seed = 0, verbose = FALSE, ...
  ) {
    set.seed(as.integer(seed))
    # Key: accept labels/future_workers/blas_threads/pca_approx via ... but ignore them here.
    # Also: only pass what run_seurat_cca_geneactivity actually supports.
    .call_supported_named(
      run_seurat_cca_geneactivity,
      obj = obj,
      latent_dim = 30,
      splits = splits,
      fold_cells = fold_cells,
      hvgs = hvgs,
      ga_features = ga_features,
      verbose = verbose
    )
  },

  seurat_bridge = function(
    obj, splits = NULL, fold_cells = NULL, labels = NULL, hvgs = NULL, ga_features = NULL,
    seed = 0, verbose = FALSE,
    # consume these if they come in via ...
    future_workers = NULL, blas_threads = NULL, pca_approx = NULL,
    ...
  ) {
    set.seed(as.integer(seed))

    # Map your global knobs onto what your bridge runner expects
    # (your run_seurat_bridge_v5_safe uses bridge_workers/bridge_blas_threads/pca_approx-ish flags)
    bridge_workers      <- if (!is.null(future_workers)) as.integer(future_workers) else NULL
    bridge_blas_threads <- if (!is.null(blas_threads))   as.integer(blas_threads)   else NULL
    pca_approx_flag     <- if (!is.null(pca_approx))     isTRUE(pca_approx)         else TRUE

    .call_supported_named(
      run_seurat_bridge_v5_safe,
      obj_bridge = obj,
      splits = splits,
      fold_cells = fold_cells,
      hvgs = hvgs,
      seed = seed,
      verbose = verbose,

      # common flags you already use
      pca_approx = pca_approx_flag,
      force_sequential = TRUE,
      manual_scale = TRUE,

      # only passed if the underlying function supports them
      bridge_workers = bridge_workers,
      bridge_blas_threads = bridge_blas_threads
    )
  },

  harmony = function(
    obj, splits = NULL, fold_cells = NULL, labels = NULL, hvgs = NULL, ga_features = NULL,
    seed = 0, verbose = FALSE, ...
  ) {
    set.seed(as.integer(seed))
    # DO NOT forward ... unless run_harmony_dual supports it
    run_harmony_dual(
      obj = obj,
      latent_dim = 30,
      rna_red = "pca",
      atac_red = "lsi",
      rna_dims = 1:100,
      atac_dims = 2:101,
      splits = splits,
      fold_cells = fold_cells,
      labels = labels,
      seed = seed,
      verbose = verbose
    )
  },

  liger = function(
    obj, splits = NULL, fold_cells = NULL, labels = NULL, hvgs = NULL, ga_features = NULL,
    seed = 0, verbose = FALSE, ...
  ) {
    set.seed(as.integer(seed))
    # LIGER runners differ a LOT by version; pass only the safest args
    .call_supported_named(
      run_liger_gene_activity,
      obj = obj,
      latent_dim = 30,
      fold_cells = fold_cells,
      assay_rna = "RNA",
      assay_activity = "ACTIVITY",
      verbose = verbose
      # do NOT pass splits/labels/... unless your function supports it
    )
  },

  mofa2 = function(
    obj, splits = NULL, fold_cells = NULL, labels = NULL, hvgs = NULL, ga_features = NULL,
    seed = 0, verbose = FALSE, ...
  ) {
    set.seed(as.integer(seed))
    .call_supported_named(
      run_mofa2_pca_lsi,
      obj = obj,
      latent_dim = 30,
      rna_dims = 1:100,
      atac_dims = 2:101,
      splits = splits,
      verbose = verbose
      # do NOT pass fold_cells/labels/hvgs/...
    )
  }
)


# %% [code]
cat("labels named? ", !is.null(names(labels)), "\n")
cat("labels length / ncol(obj): ", length(labels), " / ", ncol(obj), "\n")
cat("splits train/test: ", length(splits$train), " / ", length(splits$test), "\n")

# translate splits to cell IDs via the labels mapping
tr_cells_dbg <- names(labels)[splits$train]
te_cells_dbg <- names(labels)[splits$test]
cat("dbg tr/te cells: ", length(tr_cells_dbg), " / ", length(te_cells_dbg), "\n")

# check overlaps with embeddings *if you already computed some reduction*
if ("pca" %in% names(obj@reductions)) {
  zr <- Seurat::Embeddings(obj, "pca")
  cat("dbg test overlap with pca rownames: ", length(intersect(te_cells_dbg, rownames(zr))), "\n")
}


# %% [code]
# -----------------------------
# 3) run one (seed, fold, method)
# -----------------------------
run_one <- function(method_name, method_fun, obj, labels, splits, seed, fold,
                    k_mix=30, k_lt=15, fos_sub=3000, verbose=FALSE) {

  `%||%` <- function(a, b) {
    if (is.null(a)) return(b)
    if (length(a) == 0) return(b)
    if (is.atomic(a) && length(a) == 1 && is.na(a)) return(b)
    a
  }

  # helper: call method_fun with only args it supports
  # IMPORTANT: if fun has "..." in formals, pass everything through
  .call_method <- function(fun, ...) {
    args <- list(...)
    fml  <- names(formals(fun))
    if (is.null(fml)) fml <- character(0)

    if ("..." %in% fml) {
      return(do.call(fun, args))
    } else {
      args <- args[names(args) %in% fml]
      return(do.call(fun, args))
    }
  }

  # ---- FIT timing ----
  t0 <- proc.time()[["elapsed"]]
  status <- "ok"
  err <- NA_character_

  out <- tryCatch(
    .call_method(
      method_fun,
      obj = obj,
      splits = splits,
      seed = seed,
      verbose = verbose
    ),
    error = function(e) {
      status <<- "error"
      err <<- conditionMessage(e)
      list(
        Z_rna = NULL, Z_atac = NULL, Z_fused = NULL,
        extra_json = list(
          method = method_name,
          skipped = TRUE,
          reason = err
        )
      )
    }
  )
  t1 <- proc.time()[["elapsed"]]
  fit_seconds <- as.numeric(t1 - t0)

  # normalize/ensure extra_json exists
  if (is.null(out$extra_json)) out$extra_json <- list(method = method_name)

  # derive skipped/reason from extra_json (so runners can intentionally skip)
  skipped <- isTRUE(out$extra_json$skipped)
  reason  <- out$extra_json$reason %||% NA_character_

  # If error happened, force skipped=TRUE so downstream code doesn't assume embeddings exist
  if (identical(status, "error")) {
    skipped <- TRUE
    reason  <- reason %||% err
    out$extra_json$skipped <- TRUE
    out$extra_json$reason  <- reason
  }

  # ---- EVAL timing ----
  t2 <- proc.time()[["elapsed"]]

  ev <- tryCatch(
    evaluate_one(
      name = method_name,
      out = out,
      labels = labels,
      splits = splits,
      k_mix = k_mix,
      k_lt = k_lt,
      fos_sub = fos_sub,
      seed = seed,
      verbose = verbose
    ),
    error = function(e) {
      status <<- "error"
      err2 <- conditionMessage(e)
      data.table::data.table(
        method = out$extra_json$method %||% method_name,
        failed = TRUE,
        skipped = TRUE,
        reason = paste0("evaluate_one failed: ", err2)
      )
    }
  )

  t3 <- proc.time()[["elapsed"]]
  eval_seconds <- as.numeric(t3 - t2)

  # ---- attach bookkeeping + timings ----
  if (!data.table::is.data.table(ev)) ev <- data.table::as.data.table(ev)

  if (!("method"   %in% names(ev))) ev[, method := (out$extra_json$method %||% method_name)]
  if (!("failed"   %in% names(ev))) ev[, failed := identical(status, "error")]
  if (!("skipped"  %in% names(ev))) ev[, skipped := skipped]
  if (!("reason"   %in% names(ev))) ev[, reason := reason]

  if (identical(status, "error")) {
    ev[, `:=`(failed = TRUE, skipped = TRUE)]
    if (is.na(ev$reason[1])) ev[, reason := (reason %||% err)]
  }

  ev[, `:=`(
    seed = seed,
    fold = fold,
    status = status,
    error = err,
    fit_seconds = fit_seconds,
    eval_seconds = eval_seconds,
    total_seconds = fit_seconds + eval_seconds
  )]

  ev
}


# %% [code]
# ============================================================
# FIXED HELPERS (separate)
# - stratified_split_total: more stable rounding; guarantees disjoint+exhaustive
# - .flatten_extra_json: safe even if jsonlite missing (falls back to as.character)
# - summarize_metrics_all: robust SEM and numeric columns
# ============================================================
stratified_split_total <- function(labels, seed,
                                   train_frac = 0.8,
                                   val_frac   = 0.1,
                                   test_frac  = 0.1,
                                   na_label   = "__NA__") {
  stopifnot(abs((train_frac + val_frac + test_frac) - 1) < 1e-8)
  set.seed(seed)

  y <- as.character(labels)
  y[is.na(y)] <- na_label
  labs <- factor(y)

  n_all <- length(labs)
  idx_all <- seq_len(n_all)

  train_idx <- integer(0)
  val_idx   <- integer(0)
  test_idx  <- integer(0)

  for (lv in levels(labs)) {
    idx <- idx_all[labs == lv]
    n <- length(idx)
    if (n == 0) next
    idx <- sample(idx, size = n, replace = FALSE)

    # Allocate test/val with rounding but keep feasible
    n_test <- as.integer(round(test_frac * n))
    n_val  <- as.integer(round(val_frac  * n))

    # Enforce feasibility: keep at least 1 train when possible
    if (n >= 3) {
      n_test <- max(1L, min(n_test, n - 2L))
      n_val  <- max(1L, min(n_val,  n - n_test - 1L))
    } else if (n == 2) {
      n_test <- min(1L, n_test)
      n_val  <- 0L
    } else {
      n_test <- 0L
      n_val  <- 0L
    }

    # If rounding overflowed, trim val first then test
    if (n_test + n_val > n) {
      overflow <- (n_test + n_val) - n
      drop_val <- min(overflow, n_val)
      n_val <- n_val - drop_val
      overflow <- overflow - drop_val
      if (overflow > 0) n_test <- max(0L, n_test - overflow)
    }

    n_train <- n - n_test - n_val

    if (n_test  > 0) test_idx  <- c(test_idx,  idx[seq_len(n_test)])
    if (n_val   > 0) val_idx   <- c(val_idx,   idx[seq.int(n_test + 1L, n_test + n_val)])
    if (n_train > 0) train_idx <- c(train_idx, idx[seq.int(n_test + n_val + 1L, n)])
  }

  # Shuffle within splits for randomness
  train_idx <- sample(train_idx)
  val_idx   <- sample(val_idx)
  test_idx  <- sample(test_idx)

  # Hard invariants
  stopifnot(length(intersect(train_idx, val_idx))  == 0L)
  stopifnot(length(intersect(train_idx, test_idx)) == 0L)
  stopifnot(length(intersect(val_idx, test_idx))   == 0L)
  stopifnot(length(union(union(train_idx, val_idx), test_idx)) == n_all)

  list(train = train_idx, val = val_idx, test = test_idx)
}


.flatten_extra_json <- function(extra) {
  if (is.null(extra)) return(list())
  if (!is.list(extra)) return(list(extra_json = as.character(extra)))

  out <- list()
  has_json <- requireNamespace("jsonlite", quietly = TRUE)

  for (nm in names(extra)) {
    v <- extra[[nm]]
    if (is.null(v)) next

    if (is.atomic(v) && length(v) == 1L) {
      out[[nm]] <- v
    } else {
      out[[nm]] <- if (has_json) jsonlite::toJSON(v, auto_unbox = TRUE) else as.character(v)
    }
  }
  out
}


summarize_metrics_all <- function(dt, by = "method") {
  stopifnot(by %in% names(dt))
  if (!requireNamespace("data.table", quietly = TRUE)) stop("Need data.table installed.")

  if (!("skipped" %in% names(dt))) dt[, skipped := FALSE]
  if (!("error"   %in% names(dt))) dt[, error := FALSE]

  ok <- dt[status == "ok" & !skipped]
  if (!nrow(ok)) return(data.table::data.table())

  id_cols <- c("seed","fold","method","status","error","skipped","reason","method_str","extra_json")
  id_cols <- intersect(id_cols, names(ok))

  num_cols <- names(ok)[vapply(ok, is.numeric, logical(1))]
  num_cols <- setdiff(num_cols, id_cols)
  if (!length(num_cols)) return(data.table::data.table())

  sem <- function(x) {
    x <- x[is.finite(x)]
    if (length(x) <= 1) return(NA_real_)
    stats::sd(x) / sqrt(length(x))
  }

  ok[, c(
    list(n = .N),
    setNames(lapply(num_cols, function(m) mean(get(m), na.rm = TRUE)), paste0(num_cols, "_mean")),
    setNames(lapply(num_cols, function(m) stats::sd(get(m), na.rm = TRUE)), paste0(num_cols, "_sd")),
    setNames(lapply(num_cols, function(m) sem(get(m))), paste0(num_cols, "_sem"))
  ), by = by]
}


# %% [code]
# Helper: run a function under a temporary future plan, then restore.
# (This avoids the "obj_fold not found" quoting/eval mess and avoids giant FUN globals.)
with_future_plan_call <- function(plan, workers = NULL, fun, ...) {
  if (!requireNamespace("future", quietly = TRUE)) {
    return(do.call(fun, list(...)))
  }
  old <- future::plan()
  on.exit(future::plan(old), add = TRUE)

  if (!is.null(workers)) future::plan(plan, workers = workers) else future::plan(plan)
  do.call(fun, list(...))
}

# Helper: choose workers from Slurm (fallback default)
slurm_workers <- function(default = 1L) {
  x <- suppressWarnings(as.integer(Sys.getenv("SLURM_CPUS_PER_TASK", unset = as.character(default))))
  if (!is.finite(x) || x < 1L) x <- default
  x
}

# Optional: control BLAS threads (if RhpcBLASctl is available)
set_blas_threads <- function(n = 1L) {
  n <- as.integer(n)
  if (!is.finite(n) || n < 1L) n <- 1L
  if (requireNamespace("RhpcBLASctl", quietly = TRUE)) {
    RhpcBLASctl::blas_set_num_threads(n)
    RhpcBLASctl::omp_set_num_threads(n)
  } else {
    Sys.setenv(
      OMP_NUM_THREADS = as.character(n),
      OPENBLAS_NUM_THREADS = as.character(n),
      MKL_NUM_THREADS = as.character(n),
      VECLIB_MAXIMUM_THREADS = as.character(n),
      NUMEXPR_NUM_THREADS = as.character(n)
    )
  }
  invisible(n)
}


# %% [code]
library(future)
#plan(multisession, workers = 24)
#print(future::plan())
#print(future::nbrOfWorkers())


# %% [code]
requireNamespace("future.apply", quietly = TRUE)


# %% [code]
# ============================================================
# Fold-local cache: keep a fold-level "obj with ACTIVITY" so we
# compute GeneActivity at most once per (fold, seed)
# ============================================================
.make_fold_cache <- function() {
  new.env(parent = emptyenv())
}

.get_fold_cache_key <- function(fold, seed, feat_hash) {
  paste0("fold=", fold, "|seed=", seed, "|feat=", feat_hash)
}


# %% [code]
stamp <- function() format(Sys.time(), "%H:%M:%S")

mem_gb <- function() {
  # Linux-friendly; returns NA on systems without /proc
  rss_kb <- tryCatch({
    as.numeric(readLines("/proc/self/status")[grep("^VmRSS:", readLines("/proc/self/status"))] |>
                 sub("VmRSS:\\s+", "", x = _) |>
                 sub("\\s+kB", "", x = _))
  }, error = function(e) NA_real_)
  if (is.na(rss_kb)) return(NA_real_)
  rss_kb / (1024^2)
}

timed <- function(label, verbose = TRUE, expr) {
  t0_wall <- Sys.time()
  t0_cpu  <- proc.time()
  out <- force(expr)
  dt_wall <- as.numeric(difftime(Sys.time(), t0_wall, units = "secs"))
  dt_cpu  <- (proc.time() - t0_cpu)[["elapsed"]]
  if (isTRUE(verbose)) {
    cat(sprintf("[%s] %s | wall=%.1fs cpu=%.1fs rss=%.2fGB\n",
                stamp(), label, dt_wall, dt_cpu, mem_gb()))
  }
  out
}


# %% [code]
# ============================================================
# Seed + CV sweep for the R benchmarking notebook
# - Stratified K-fold by labels (fallback: random K-fold if only 1 label)
# - Re-fits each method per (seed, fold) and evaluates on that fold’s test split
# - Saves: per-run TSV + (mean, sd, sem) summary TSV
# ============================================================
library(data.table)

# -----------------------------
# configure the sweep
# -----------------------------
SEEDS        <- c(67, 1985, 789, 3, 99)
#SEEDS        <- c(67)
#SEEDS        <- c(67, 1985)
N_FOLDS      <- 3
#N_FOLDS      <- 2
#N_FOLDS      <- 1
SWEEP_TAG    <- "cv_sweep_R_1-31-2026"
OUT_DIR      <- file.path(WORK %||% ".", "runs", SWEEP_TAG)
dir.create(OUT_DIR, recursive = TRUE, showWarnings = FALSE)

cat("[cv] OUT_DIR=", OUT_DIR, "\n")


# %% [code]
methods_need_activity <- function(method_names) {
  # anything that touches GeneActivity / ACTIVITY assay
  mn <- as.character(method_names)
  grepl("seurat_cca|liger", mn, ignore.case = TRUE)
}


# %% [code]
Sys.setenv(OMP_NUM_THREADS="24",
           OPENBLAS_NUM_THREADS="24",
           MKL_NUM_THREADS="24",
           VECLIB_MAXIMUM_THREADS="24",
           NUMEXPR_NUM_THREADS="24")
if (requireNamespace("RhpcBLASctl", quietly=TRUE)) {
  RhpcBLASctl::blas_set_num_threads(24)
  RhpcBLASctl::omp_set_num_threads(24)
}


# %% [code]
if (requireNamespace("RhpcBLASctl", quietly = TRUE)) {
  RhpcBLASctl::blas_set_num_threads(24)
  RhpcBLASctl::omp_set_num_threads(24)
  cat("after set: blas=", RhpcBLASctl::blas_get_num_procs(), "\n")
  cat("after set: omp =", RhpcBLASctl::omp_get_num_procs(), "\n")
}


# %% [code]
# -----------------------------
# Inject variables into a function's environment temporarily
# (FIXED: lock bindings so runners can't overwrite fold_cells)
# -----------------------------
.with_injected_env <- function(fn, inject, expr_call) {
  env0 <- environment(fn)
  old_vals <- list()
  had <- logical()

  nms <- names(inject)
  had <- vapply(nms, function(k) exists(k, envir = env0, inherits = FALSE), logical(1))
  old_vals <- lapply(nms, function(k) if (exists(k, envir = env0, inherits = FALSE)) get(k, envir = env0, inherits = FALSE) else NULL)
  names(old_vals) <- nms

  # inject + LOCK so runner can't clobber them
  for (k in nms) {
    assign(k, inject[[k]], envir = env0)
    if (!bindingIsLocked(k, env0)) lockBinding(k, env0)
  }

  on.exit({
    # unlock + restore
    for (k in nms) {
      if (bindingIsLocked(k, env0)) unlockBinding(k, env0)
    }
    for (i in seq_along(nms)) {
      k <- nms[[i]]
      if (had[[i]]) {
        assign(k, old_vals[[k]], envir = env0)
      } else {
        if (exists(k, envir = env0, inherits = FALSE)) rm(list = k, envir = env0)
      }
    }
  }, add = TRUE)

  eval(expr_call, envir = parent.frame())
}

# -----------------------------
# call function but drop unsupported args
# -----------------------------
.call_supported <- function(fn, ...) {
  dots <- list(...)
  if (is.null(fn)) stop("fn is NULL")
  fml <- tryCatch(names(formals(fn)), error = function(e) character(0))
  if (!length(fml) || "..." %in% fml) return(do.call(fn, dots))
  keep <- intersect(names(dots), fml)
  do.call(fn, dots[keep])
}

# -----------------------------
# SAFE wrappers that ensure fold_cells is visible to runners
# (FIXED: WNN wrapper does NOT pass labels, because run_seurat_wnn doesn't take it)
# -----------------------------
run_seurat_wnn_safe <- function(
  obj, splits = NULL, fold_cells = NULL,
  latent_dim = 30, rna_dims = 1:100, atac_dims = 2:101,
  verbose = FALSE, ...
) {
  if (!requireNamespace("Seurat", quietly = TRUE)) stop("Need Seurat installed.")
  .msg <- function(...) if (isTRUE(verbose)) { cat(..., "\n"); flush.console() }

  .assert_fold_cells_ok(fold_cells, ctx = "[WNN_SAFE]")
  .msg(sprintf(
    "[WNN_SAFE] obj cells=%d | train=%d val=%d test=%d | assays=%s",
    ncol(obj), length(fold_cells$train), length(fold_cells$val), length(fold_cells$test),
    paste(Seurat::Assays(obj), collapse = ",")
  ))

  out <- .call_supported(
    run_seurat_wnn,
    obj = obj,
    latent_dim = latent_dim,
    rna_dims = rna_dims,
    atac_dims = atac_dims,
    splits = splits,
    fold_cells = fold_cells,   # <-- THIS was missing
    verbose = verbose,
    ...
  )
  out
}

run_harmony_dual_safe <- function(
  obj, splits = NULL, fold_cells = NULL, labels = NULL,
  latent_dim = 30, rna_dims = 1:100, atac_dims = 2:101,
  seed = 0, verbose = FALSE, ...
) {
  if (!requireNamespace("Seurat", quietly = TRUE)) stop("Need Seurat installed.")
  .msg <- function(...) if (isTRUE(verbose)) { cat(..., "\n"); flush.console() }

  .assert_fold_cells_ok(fold_cells, ctx = "[HARMONY_SAFE]")
  .msg(sprintf(
    "[HARMONY_SAFE] obj cells=%d | train=%d val=%d test=%d | labels=%s",
    ncol(obj), length(fold_cells$train), length(fold_cells$val), length(fold_cells$test),
    if (is.null(labels)) "NULL" else sprintf("len=%d", length(labels))
  ))

  out <- .call_supported(
    run_harmony_dual,
    obj = obj,
    latent_dim = latent_dim,
    rna_dims = rna_dims,
    atac_dims = atac_dims,
    splits = splits,
    fold_cells = fold_cells,   # <-- THIS was missing
    labels = labels,
    seed = seed,
    verbose = verbose,
    ...
  )
  out
}

# ============================================================
# CV SWEEP FIX PACK (paste-in)
# (kept as-is except dispatcher + sweep fixes)
# ============================================================

.available_cores_safe <- function() {
  n <- NA_integer_
  if (requireNamespace("future", quietly = TRUE)) {
    n <- suppressWarnings(tryCatch(future::availableCores(), error = function(e) NA_integer_))
  }
  if (!is.finite(n) || is.na(n)) {
    x <- tryCatch(system("grep Cpus_allowed_list /proc/self/status", intern = TRUE), error = function(e) character(0))
    if (length(x)) {
      s <- sub("^.*:\\s*", "", x[1])
      parts <- unlist(strsplit(s, ","))
      cnt <- 0L
      for (p in parts) {
        if (grepl("-", p)) {
          ab <- as.integer(strsplit(p, "-", fixed = TRUE)[[1]])
          if (length(ab) == 2 && all(is.finite(ab))) cnt <- cnt + (ab[2] - ab[1] + 1L)
        } else {
          if (nzchar(p)) cnt <- cnt + 1L
        }
      }
      n <- cnt
    }
  }
  if (!is.finite(n) || is.na(n) || n < 1L) n <- 1L
  as.integer(n)
}

.make_fold_cells <- function(cell_ids, splits_idx, val_frac = 0.1, seed = 0) {
  set.seed(as.integer(seed))
  n <- length(cell_ids)

  tr <- as.integer(splits_idx$train %||% integer(0))
  te <- as.integer(splits_idx$test  %||% integer(0))
  tr <- tr[tr >= 1 & tr <= n]
  te <- te[te >= 1 & te <= n]

  n_val <- floor(as.numeric(val_frac) * length(tr))
  if (is.finite(n_val) && n_val >= 1L && length(tr) >= (n_val + 2L)) {
    val_idx <- sample(tr, size = n_val, replace = FALSE)
    tr_idx  <- setdiff(tr, val_idx)
  } else {
    val_idx <- integer(0)
    tr_idx  <- tr
  }

  list(
    fold_cells = list(
      train = cell_ids[tr_idx],
      val   = cell_ids[val_idx],
      test  = cell_ids[te]
    ),
    splits_idx = list(
      train = tr_idx,
      val   = val_idx,
      test  = te
    )
  )
}

.align_labels_to_cells <- function(labels, cell_ids) {
  if (is.null(labels)) return(NULL)
  if (!is.null(names(labels))) {
    out <- labels[cell_ids]
    names(out) <- cell_ids
    return(out)
  }
  if (length(labels) != length(cell_ids)) {
    stop("[labels] labels length does not match n_cells and labels are not named.")
  }
  names(labels) <- cell_ids
  labels
}

# -----------------------------
# METHOD DISPATCHER (FIXED: no more unused-arg explosions)
# - Always passes fold_cells list to safe wrappers for WNN/Harmony
# - Uses .call_supported for other methods
# -----------------------------
.run_method_safe <- function(
  method_spec,
  method_name,
  obj_fold,
  fold_cells,
  splits_idx,
  labels,
  hvgs = NULL,
  ga_features = NULL,
  latent_dim = 30,
  rna_dims = 1:100,
  atac_dims = 2:101,
  verbose = TRUE,
  seed = 0,
  ...
) {
  if (!requireNamespace("Seurat", quietly = TRUE)) stop("Need Seurat installed.")
  .msg <- function(...) if (isTRUE(verbose)) { cat(..., "\n"); flush.console() }

  .assert_fold_cells_ok(fold_cells, ctx = "[dispatch]")

  splits_cells <- list(train = fold_cells$train, test = fold_cells$test)

  .msg(sprintf(
    "[dispatch] %s | obj_fold n=%d | train=%d val=%d test=%d | assays=%s",
    method_name, ncol(obj_fold),
    length(fold_cells$train), length(fold_cells$val), length(fold_cells$test),
    paste(Seurat::Assays(obj_fold), collapse = ",")
  ))

  .standardize <- function(x) {
    list(
      Z_rna   = x$Z_rna   %||% NULL,
      Z_atac  = x$Z_atac  %||% NULL,
      Z_fused = x$Z_fused %||% NULL,
      extra_json = x$extra_json %||% list()
    )
  }

  method <- as.character(method_spec)[1]

  if (identical(method, "seurat_wnn")) {
    x <- run_seurat_wnn_safe(
      obj = obj_fold,
      latent_dim = latent_dim,
      rna_dims = rna_dims,
      atac_dims = atac_dims,
      splits = splits_cells,
      fold_cells = fold_cells,
      verbose = verbose,
      ...
    )
    return(.standardize(x))
  }

  if (identical(method, "harmony")) {
    x <- run_harmony_dual_safe(
      obj = obj_fold,
      latent_dim = latent_dim,
      rna_dims = rna_dims,
      atac_dims = atac_dims,
      splits = splits_cells,
      fold_cells = fold_cells,
      labels = labels,
      seed = seed,
      verbose = verbose,
      ...
    )
    return(.standardize(x))
  }

  if (identical(method, "seurat_cca")) {
    .msg(sprintf("[dispatch] ACTIVITY assay present? %s", "ACTIVITY" %in% Seurat::Assays(obj_fold)))
    x <- .call_supported(
      run_seurat_cca_geneactivity,
      obj = obj_fold,
      latent_dim = latent_dim,
      splits = splits_cells,
      verbose = verbose,
      fold_cells = fold_cells,
      hvgs = hvgs,
      ga_features = ga_features,
      ...
    )
    return(.standardize(x))
  }

  if (identical(method, "seurat_bridge")) {
    x <- .call_supported(
      run_seurat_bridge_v5_safe,
      obj_bridge = obj_fold,
      splits = splits_cells,
      fold_cells = fold_cells,
      hvgs = hvgs,
      seed = seed,
      latent_dim = latent_dim,
      verbose = verbose,
      ...
    )
    return(.standardize(x))
  }

  if (identical(method, "liger")) {
    .msg(sprintf("[dispatch] ACTIVITY assay present? %s", "ACTIVITY" %in% Seurat::Assays(obj_fold)))
    x <- .call_supported(
      run_liger_gene_activity,
      obj = obj_fold,
      latent_dim = latent_dim,
      splits = splits_cells,
      verbose = verbose,
      fold_cells = fold_cells,
      hvgs = hvgs,
      ga_features = ga_features,
      ...
    )
    return(.standardize(x))
  }

  if (identical(method, "mofa2")) {
    x <- .call_supported(
      run_mofa2_pca_lsi,
      obj = obj_fold,
      latent_dim = latent_dim,
      rna_dims = rna_dims,
      atac_dims = atac_dims,
      splits = splits_cells,
      verbose = verbose,
      fold_cells = fold_cells,
      ...
    )
    return(.standardize(x))
  }

  stop("Unknown method: ", method_name %||% method)
}


# %% [code]
# ============================================================
# Make labels safe for evaluation
# - ensures named character/factor vector aligned to obj_cells
# - returns NULL if unusable
# ============================================================
.harden_labels_for_eval <- function(labels, obj_cells, min_classes = 2L) {
  if (is.null(labels) || is.null(obj_cells) || length(obj_cells) == 0L) return(NULL)

  # If labels is unnamed but same length as obj_cells, name it.
  if (is.null(names(labels)) && length(labels) == length(obj_cells)) {
    names(labels) <- obj_cells
  }

  # If still unnamed, can't align reliably.
  if (is.null(names(labels))) return(NULL)

  # Align to obj_cells
  y <- labels[obj_cells]
  if (is.null(y) || length(y) != length(obj_cells)) return(NULL)

  # Sanitize values
  y <- as.character(y)
  y[!is.finite(match(y, y))] <- y  # no-op; keeps non-NA strings stable
  y[is.na(y) | y %in% c("NA","NaN","nan","None","")] <- NA_character_

  # Require at least 2 classes among non-NA labels
  y2 <- y[!is.na(y)]
  if (length(unique(y2)) < as.integer(min_classes)) return(NULL)

  names(y) <- obj_cells
  y
}


# %% [code]
library(data.table)

# ------------------------------------------------------------
# helpers (drop-in)
# ------------------------------------------------------------
`%||%` <- if (exists("%||%", mode = "function")) get("%||%") else function(a, b) {
  if (is.null(a) || length(a) == 0) b else a
}

.as_one_row_list <- function(x) {
  # x can be: data.table/data.frame/list
  if (is.null(x)) return(list())
  if (data.table::is.data.table(x) || is.data.frame(x)) {
    if (nrow(x) < 1) return(list())
    return(as.list(x[1]))
  }
  if (is.list(x)) return(x)
  list()
}

# ---- FOSCTTM compatibility wrapper ----
# Handles "signature mismatch" when your call site and foscttm_values() definition diverged.
.foscttm_values_compat <- function(...) {
  dots <- list(...)
  out <- try(do.call(foscttm_values, dots), silent = TRUE)
  if (!inherits(out, "try-error")) return(out)

  fn  <- foscttm_values
  fml <- names(formals(fn))
  dots2 <- dots[names(dots) %in% fml]
  out2 <- try(do.call(fn, dots2), silent = TRUE)
  if (!inherits(out2, "try-error")) return(out2)

  # rethrow the original error for debugging
  stop(out)
}

# ---- Latent normalizer ----
# Makes per-modality latents discoverable across methods (WNN/Harmony/Bridge/LIGER/MOFA2/etc)
.normalize_latents <- function(res) {
  if (is.null(res) || !is.list(res)) return(res)

  # fused
  if (is.null(res$Z_fused) && !is.null(res$Zf)) res$Z_fused <- res$Zf

  # RNA
  if (is.null(res$Z_rna)) {
    res$Z_rna <- res$Zr %||% res$Z_RNA %||% res$Zrna %||% NULL
  }

  # second modality: call it "mod" (often ATAC or GA-derived)
  if (is.null(res$Z_mod)) {
    res$Z_mod <- res$Z_atac %||% res$Za %||% res$Z_ATAC %||% res$Zatac %||% NULL
  }

  res
}

# ---- Safe subsetting to split cells that exist in a given latent ----
.pick_split_cells <- function(cell_ids, splits_idx, which = c("train", "val", "test")) {
  which <- match.arg(which)
  idx <- splits_idx[[which]] %||% integer(0)
  if (is.integer(idx) || is.numeric(idx)) {
    idx <- as.integer(idx)
    idx <- idx[idx >= 1L & idx <= length(cell_ids)]
    cell_ids[idx]
  } else {
    # already cell names
    as.character(idx)
  }
}

# ---- Map from "pretty" eval columns -> legacy columns used by CV table ----
.normalize_eval_columns <- function(metrics_any) {
  m <- .as_one_row_list(metrics_any)

  legacy_keys <- c(
    "FOSCTTM_mean_test", "FOSCTTM_sem_test", "FOSCTTM_mrr_mean_test",
    "FOSCTTM_recall@1_mean_test", "FOSCTTM_recall@10_mean_test",
    "FOSCTTM_recall@25_mean_test", "FOSCTTM_recall@50_mean_test",
    "FOSCTTM_recall@100_mean_test",
    "mixing_score_test",
    "label_transfer_acc_mean_test",
    "label_transfer_macroF1_mean_test"
  )

  has_any_legacy <- any(vapply(legacy_keys, function(k) !is.null(m[[k]]), logical(1)))

  if (!has_any_legacy) {
    # Pair / foscttm
    m[["FOSCTTM_mean_test"]]     <- m[["PAIR(test)/FOSCTTM_mean"]] %||% NA_real_
    m[["FOSCTTM_mrr_mean_test"]] <- m[["PAIR(test)/FOSCTTM_MRR"]]  %||% NA_real_

    # SEM (not present in pretty output unless you add it)
    m[["FOSCTTM_sem_test"]] <- m[["PAIR(test)/FOSCTTM_SEM"]] %||% NA_real_

    # Recalls
    m[["FOSCTTM_recall@1_mean_test"]]   <- m[["PAIR(test)/FOSCTTM_Recall@1"]]   %||% NA_real_
    m[["FOSCTTM_recall@10_mean_test"]]  <- m[["PAIR(test)/FOSCTTM_Recall@10"]]  %||% NA_real_
    m[["FOSCTTM_recall@25_mean_test"]]  <- m[["PAIR(test)/FOSCTTM_Recall@25"]]  %||% NA_real_
    m[["FOSCTTM_recall@50_mean_test"]]  <- m[["PAIR(test)/FOSCTTM_Recall@50"]]  %||% NA_real_
    m[["FOSCTTM_recall@100_mean_test"]] <- m[["PAIR(test)/FOSCTTM_Recall@100"]] %||% NA_real_

    # Mixing
    m[["mixing_score_test"]] <- m[["MIX(test)/Mixing score"]] %||% NA_real_

    # Label transfer
    m[["label_transfer_acc_mean_test"]] <-
      m[["PAIR(test)/Label transfer acc mean"]] %||% NA_real_
    m[["label_transfer_macroF1_mean_test"]] <-
      m[["PAIR(test)/Label transfer macroF1 mean"]] %||% NA_real_
  }

  for (k in legacy_keys) {
    if (is.null(m[[k]])) m[[k]] <- NA_real_
    if (!is.na(m[[k]])) {
      if (length(m[[k]]) > 1) m[[k]] <- m[[k]][1]
    }
  }

  data.table::data.table(
    FOSCTTM_mean_test = as.numeric(m[["FOSCTTM_mean_test"]]),
    FOSCTTM_sem_test  = as.numeric(m[["FOSCTTM_sem_test"]]),
    FOSCTTM_mrr_mean_test = as.numeric(m[["FOSCTTM_mrr_mean_test"]]),
    `FOSCTTM_recall@1_mean_test`  = as.numeric(m[["FOSCTTM_recall@1_mean_test"]]),
    `FOSCTTM_recall@10_mean_test` = as.numeric(m[["FOSCTTM_recall@10_mean_test"]]),
    `FOSCTTM_recall@25_mean_test` = as.numeric(m[["FOSCTTM_recall@25_mean_test"]]),
    `FOSCTTM_recall@50_mean_test` = as.numeric(m[["FOSCTTM_recall@50_mean_test"]]),
    `FOSCTTM_recall@100_mean_test`= as.numeric(m[["FOSCTTM_recall@100_mean_test"]]),
    mixing_score_test = as.numeric(m[["mixing_score_test"]]),
    label_transfer_acc_mean_test = as.numeric(m[["label_transfer_acc_mean_test"]]),
    label_transfer_macroF1_mean_test = as.numeric(m[["label_transfer_macroF1_mean_test"]])
  )
}

# ------------------------------------------------------------
# Back-compat adapter:
# - normalizes out latents (Z_fused/Z_rna/Z_mod)
# - calls your evaluate_one()
# - normalizes into the legacy metric columns your CV table expects
# - also patches FOSCTTM call by injecting compat wrapper if evaluate_one uses it
# ------------------------------------------------------------
evaluate_one_safe <- function(
  res, labels, splits_idx,
  k_mix = 30L, k_lt = 15L, fos_sub = 3000L,
  seed = 0L, verbose = FALSE
) {
  if (!requireNamespace("jsonlite", quietly = TRUE)) {
    stop("Need jsonlite (used only to parse res$extra_json if it's a JSON string).")
  }

  # Normalize latent keys so evaluate_one can find individual latents consistently
  res <- .normalize_latents(res)

  # Determine method name robustly
  ex <- res$extra_json %||% res$extra %||% NULL
  if (is.character(ex) && length(ex) == 1L && nzchar(ex)) {
    ex <- tryCatch(jsonlite::fromJSON(ex), error = function(e) NULL)
  }
  name <- (ex$method %||% ex$method_str %||% res$method %||% "method")

  # If your evaluate_one() calls foscttm_values() internally, this makes it robust:
  # temporarily mask foscttm_values in the local env with a compat wrapper.
  # (No-op if evaluate_one doesn't use foscttm_values.)
  old_fos <- NULL
  have_fos <- exists("foscttm_values", mode = "function")
  if (have_fos) old_fos <- get("foscttm_values", mode = "function")
  on.exit({
    if (have_fos && !is.null(old_fos)) {
      assign("foscttm_values", old_fos, envir = .GlobalEnv)
    }
  }, add = TRUE)
  if (have_fos) {
    assign("foscttm_values", function(...) .foscttm_values_compat(...), envir = .GlobalEnv)
  }

  raw_metrics <- evaluate_one(
    name    = name,
    out     = res,
    labels  = labels,
    splits  = splits_idx,
    k_mix   = k_mix,
    k_lt    = k_lt,
    fos_sub = fos_sub,
    seed    = seed,
    verbose = verbose
  )

  .normalize_eval_columns(raw_metrics)
}


# %% [code]
# ============================================================
# Helpers
# ============================================================
`%||%` <- if (exists("%||%", mode="function")) get("%||%") else function(a,b) {
  if (is.null(a) || length(a) == 0) b else a
}

.msg <- function(..., verbose=TRUE) {
  if (isTRUE(verbose)) { cat(..., "\n"); flush.console() }
}

.assert_named_cells <- function(x, all_cells, ctx="") {
  if (is.null(x)) return(invisible(TRUE))
  if (!is.character(x)) {
    stop(ctx, " expected character cell names, got ", class(x)[1], " (looks like indices leaking in).")
  }
  bad <- setdiff(x, all_cells)
  if (length(bad) > 0) stop(ctx, " contains cells not in object: e.g. ", bad[[1]])
  invisible(TRUE)
}

.assert_fold_cells_ok_names <- function(obj, fold_cells, ctx="") {
  if (!is.list(fold_cells) || !all(c("train","val","test") %in% names(fold_cells))) {
    stop(ctx, " fold_cells must be list(train,val,test)")
  }
  all_cells <- Seurat::Cells(obj)
  .assert_named_cells(fold_cells$train %||% character(0), all_cells, paste0(ctx, " fold_cells$train"))
  .assert_named_cells(fold_cells$val   %||% character(0), all_cells, paste0(ctx, " fold_cells$val"))
  .assert_named_cells(fold_cells$test  %||% character(0), all_cells, paste0(ctx, " fold_cells$test"))
  TRUE
}

.fix_labels_names <- function(labels, obj_cells) {
  if (is.null(labels)) return(NULL)
  if (!is.null(names(labels))) return(labels)
  if (length(labels) == length(obj_cells)) {
    names(labels) <- obj_cells
    return(labels)
  }
  labels
}

.fix_latent_rownames <- function(Z, obj_cells) {
  if (is.null(Z)) return(NULL)
  Z <- as.matrix(Z)
  if (is.null(rownames(Z)) || any(!nzchar(rownames(Z)))) {
    if (nrow(Z) == length(obj_cells)) rownames(Z) <- obj_cells
  }
  Z
}

# ============================================================
# Repeated holdout splits (GLOBAL indices 1..n)
# - stratified if labels usable
# - transductive only when n_folds==1 && test_frac<=0
# ============================================================
.make_repeated_holdout_splits <- function(
  n,
  labels_all = NULL,
  n_folds = 5L,
  seed = 0L,
  train_frac = 0.8,
  val_frac   = 0.1,
  test_frac  = 0.1
) {
  n <- as.integer(n)
  if (!is.finite(n) || is.na(n) || n < 2L) stop("[cv] need >=2 cells")

  n_folds <- as.integer(n_folds)
  if (!is.finite(n_folds) || is.na(n_folds) || n_folds < 1L) n_folds <- 1L

  # transductive: no holdout
  if (n_folds == 1L && (!is.finite(test_frac) || is.na(test_frac) || test_frac <= 0)) {
    return(list(list(train=integer(0), val=integer(0), test=integer(0), transductive=TRUE)))
  }

  train_frac <- as.numeric(train_frac); val_frac <- as.numeric(val_frac); test_frac <- as.numeric(test_frac)
  if (!is.finite(train_frac + val_frac + test_frac) ||
      abs((train_frac + val_frac + test_frac) - 1.0) > 1e-8) {
    stop("[cv] train_frac + val_frac + test_frac must equal 1 (unless transductive)")
  }

  # labels eligibility
  .as_y <- function(y) {
    if (is.null(y)) return(NULL)
    y <- as.character(y)
    if (length(y) != n) return(NULL)
    y[is.na(y) | y %in% c("NA","NaN","nan","None","")] <- NA_character_
    y
  }
  y <- .as_y(labels_all)

  .can_stratify <- function(y) {
    if (is.null(y)) return(FALSE)
    y2 <- y[!is.na(y)]
    if (length(y2) < 50) return(FALSE)
    tab <- table(y2)
    length(tab) >= 2 && all(tab >= 2)
  }
  do_strat <- .can_stratify(y)

  .sample_unstrat <- function(idx, size, seed_i) {
    set.seed(as.integer(seed_i))
    idx <- as.integer(idx)
    if (size >= length(idx)) return(sort(idx))
    sort(sample(idx, size=as.integer(size), replace=FALSE))
  }

  .sample_strat <- function(idx, y, size, seed_i) {
    set.seed(as.integer(seed_i))
    idx <- as.integer(idx)
    y_idx <- y[idx]
    ok <- !is.na(y_idx)
    if (!any(ok)) return(.sample_unstrat(idx, size, seed_i))

    idx_ok <- idx[ok]
    y_ok <- y_idx[ok]

    tab <- table(y_ok)
    props <- as.numeric(tab / sum(tab))
    want <- pmax(0L, floor(props * size))

    while (sum(want) < size) {
      j <- which.max(props - (want / pmax(1, size)))
      want[j] <- want[j] + 1L
    }
    while (sum(want) > size) {
      j <- which.max(want)
      want[j] <- want[j] - 1L
    }

    out <- integer(0)
    for (j in seq_along(tab)) {
      lv <- names(tab)[j]
      pool <- idx_ok[y_ok == lv]
      if (!length(pool)) next
      take <- min(as.integer(want[j]), length(pool))
      if (take > 0) out <- c(out, sample(pool, size=take, replace=FALSE))
    }
    if (length(out) < size) {
      left <- setdiff(idx, out)
      need <- size - length(out)
      out <- c(out, sample(left, size=need, replace=FALSE))
    }
    sort(unique(out))
  }

  n_train <- max(1L, floor(train_frac * n))
  n_temp <- n - n_train
  if (n_temp < 2L) stop("[cv] too few left for val+test; increase n or train_frac")

  val_prop_temp <- val_frac / (val_frac + test_frac)
  n_val  <- max(1L, floor(val_prop_temp * n_temp))
  n_test <- n_temp - n_val
  if (n_test < 1L) { n_test <- 1L; n_val <- n_temp - 1L }

  all_idx <- seq_len(n)
  folds <- vector("list", n_folds)

  for (fold in seq_len(n_folds)) {
    train_idx <- if (do_strat) .sample_strat(all_idx, y, n_train, as.integer(seed) + fold)
                 else          .sample_unstrat(all_idx, n_train, as.integer(seed) + fold)

    temp_idx <- setdiff(all_idx, train_idx)

    val_idx <- if (do_strat) .sample_strat(temp_idx, y, n_val, as.integer(seed) + 10000L + fold)
               else          .sample_unstrat(temp_idx, n_val, as.integer(seed) + 10000L + fold)

    test_idx <- setdiff(temp_idx, val_idx)

    folds[[fold]] <- list(train=train_idx, val=val_idx, test=test_idx, transductive=FALSE)
  }

  folds
}

# ============================================================
# Map GLOBAL split indices -> FOLD-LOCAL indices
# - split indices are LOCAL to obj_cells (cell_ids_fold)
# ============================================================
.global_to_fold_local_splits <- function(splits_global, cell_ids_global, cell_ids_fold) {
  if (isTRUE(splits_global$transductive %||% FALSE)) {
    return(list(train=integer(0), val=integer(0), test=integer(0), transductive=TRUE))
  }

  to_cells <- function(ii) cell_ids_global[as.integer(ii)]
  tr_cells <- to_cells(splits_global$train)
  va_cells <- to_cells(splits_global$val)
  te_cells <- to_cells(splits_global$test)

  tr_cells <- intersect(tr_cells, cell_ids_fold)
  va_cells <- intersect(va_cells, cell_ids_fold)
  te_cells <- intersect(te_cells, cell_ids_fold)

  list(
    train = match(tr_cells, cell_ids_fold),
    val   = match(va_cells, cell_ids_fold),
    test  = match(te_cells, cell_ids_fold),
    transductive = FALSE
  )
}


# %% [code]
# ============================================================
# run_cv_sweep_R_fixed
# - repeated holdout (GLOBAL)
# - subset obj to fold_univ (within-fold transductive)
# - evaluation uses FOLD-LOCAL indices (splits_fold)
# - methods receive fold_cells as NAMES (not indices)
# ============================================================
run_cv_sweep_R_fixed <- function(
  seeds,
  n_folds,
  obj,
  labels = NULL,
  methods,
  out_dir,
  k_mix = 30,
  k_lt  = 15,
  fos_sub = 3000,
  verbose = TRUE,
  train_frac = 0.8,
  val_frac   = 0.1,
  test_frac  = 0.1,
  latent_dim = 30,
  rna_dims   = 1:100,
  atac_dims  = 2:101,

  bridge_future_plan = "multisession",
  bridge_workers = 8,
  bridge_blas_threads = 8,
  bridge_future_globals_max_gb = NULL,
  future_globals_max_gb = 50,

  ...
) {
  if (!requireNamespace("data.table", quietly = TRUE)) stop("Need data.table")
  if (!requireNamespace("future", quietly = TRUE)) stop("Need future")
  if (!requireNamespace("Seurat", quietly = TRUE)) stop("Need Seurat")
  if (!dir.exists(out_dir)) dir.create(out_dir, recursive = TRUE, showWarnings = FALSE)

  # normalize methods to a named list of functions
  if (is.character(methods)) {
    stop("[methods] pass a named list of wrappers (recommended) so dispatch is explicit.")
  }
  if (!is.list(methods) || is.null(names(methods)) || any(!nzchar(names(methods)))) {
    stop("[methods] must be a named list of functions")
  }
  if (!all(vapply(methods, is.function, logical(1)))) stop("[methods] all entries must be functions")
  method_names <- names(methods)

  cell_ids_global <- Seurat::Cells(obj)
  n_global <- length(cell_ids_global)
  if (n_global < 2L) stop("[cv] need >=2 cells")

  # labels aligned to GLOBAL cells (named)
  labels_global <- NULL
  if (!is.null(labels)) {
    if (!is.null(names(labels))) {
      labels_global <- tryCatch(.align_labels_to_cells(labels, cell_ids_global), error=function(e) NULL)
    } else if (length(labels) == n_global) {
      labels_global <- labels
      names(labels_global) <- cell_ids_global
    }
  }
  labels_global <- .fix_labels_names(labels_global, cell_ids_global)

  # future globals size
  options(future.globals.maxSize = as.numeric(future_globals_max_gb) * 1024^3)

  .with_future_plan <- function(plan_name, workers, globals_gb, expr) {
    old_plan <- future::plan()
    old_opt  <- getOption("future.globals.maxSize")
    on.exit({
      future::plan(old_plan)
      options(future.globals.maxSize = old_opt)
    }, add = TRUE)

    options(future.globals.maxSize = as.numeric(globals_gb) * 1024^3)
    future::plan(plan_name, workers = as.integer(workers))
    force(expr)
  }

  runs <- list(); rr <- 1L

  for (seed in seeds) {
    .msg("\n=== seed = ", seed, " ===", verbose=verbose)

    fold_splits_global <- .make_repeated_holdout_splits(
      n = n_global,
      labels_all = if (!is.null(labels_global)) unname(labels_global) else NULL,
      n_folds = n_folds,
      seed = seed,
      train_frac = train_frac,
      val_frac = val_frac,
      test_frac = test_frac
    )

    for (fold in seq_along(fold_splits_global)) {
      .msg("\n--- fold = ", fold, " ---", verbose=verbose)

      splits_global <- fold_splits_global[[fold]]
      transductive_mode <- isTRUE(splits_global$transductive %||% FALSE)

      # fold cell names (for methods)
      if (transductive_mode) {
        fold_cells <- list(train=character(0), val=character(0), test=character(0))
        fold_univ <- cell_ids_global
      } else {
        fold_cells <- list(
          train = cell_ids_global[splits_global$train],
          val   = cell_ids_global[splits_global$val],
          test  = cell_ids_global[splits_global$test]
        )
        .assert_fold_cells_ok_names(obj, fold_cells, ctx="[cv]")
        fold_univ <- unique(c(fold_cells$train, fold_cells$val, fold_cells$test))
      }

      # subset to fold universe
      obj_fold <- .subset_seurat_safe(obj, cells = fold_univ)
      cell_ids_fold <- Seurat::Cells(obj_fold)

      # fold-local splits for evaluation (indices into cell_ids_fold)
      splits_fold <- .global_to_fold_local_splits(splits_global, cell_ids_global, cell_ids_fold)

      # fold-local labels (named by cell id)
      labels_fold <- NULL
      if (!is.null(labels_global)) {
        labels_fold <- labels_global[cell_ids_fold]
        labels_fold <- .fix_labels_names(labels_fold, cell_ids_fold)
      }

      .msg(sprintf(
        "[cv] obj_fold cells=%d | train=%d val=%d test=%d | transductive=%s",
        length(cell_ids_fold),
        length(splits_fold$train), length(splits_fold$val), length(splits_fold$test),
        if (transductive_mode) "TRUE" else "FALSE"
      ), verbose=verbose)

      .msg(sprintf("[cv] assays in obj_fold: %s", paste(Seurat::Assays(obj_fold), collapse=",")), verbose=verbose)

      # preprocess: use TRAIN only if inductive, else all
      train_idx_fold <- if (!transductive_mode && length(splits_fold$train) > 0L) splits_fold$train else seq_along(cell_ids_fold)
      if (length(train_idx_fold) <= 10L) train_idx_fold <- seq_along(cell_ids_fold)

      hvgs <- NULL
      if (exists("preprocess_py_style", mode="function")) {
        pp <- preprocess_py_style(obj_fold, train_idx = train_idx_fold, seed = seed)
        obj_fold <- pp$obj
        hvgs <- pp$hvgs %||% NULL
        .msg(sprintf("[cv] preprocess_py_style hvgs=%s", if (is.null(hvgs)) "NULL" else length(hvgs)), verbose=verbose)
      }

      ga_features <- NULL

      for (method_name in method_names) {
        .msg(sprintf("[cv] ENTER method=%s seed=%d fold=%d", method_name, seed, fold), verbose=verbose)

        t0 <- proc.time()[[3]]
        status <- "ok"; err <- NA_character_

        out <- tryCatch({
          # IMPORTANT: pass splits_fold (fold-local indices) AND fold_cells (names)
          methods[[method_name]](
            obj = obj_fold,
            splits = splits_fold,
            fold_cells = fold_cells,
            labels = labels_fold,
            hvgs = hvgs,
            ga_features = ga_features,
            seed = seed,
            verbose = verbose,
            ...
          )
        }, error=function(e) {
          status <<- "failed"
          err <<- conditionMessage(e)
          NULL
        })

        fit_seconds <- proc.time()[[3]] - t0

        # harden output matrices if present
        if (!is.null(out)) {
          out$Z_rna   <- .fix_latent_rownames(out$Z_rna,   cell_ids_fold)
          out$Z_atac  <- .fix_latent_rownames(out$Z_atac,  cell_ids_fold)
          out$Z_fused <- .fix_latent_rownames(out$Z_fused, cell_ids_fold)
        }

        if (identical(status, "ok")) {
          skipped_flag <- isTRUE((out$extra_json$skipped %||% out$extra$skipped) %||% FALSE)
          no_latent <- is.null(out) || is.null(out$Z_fused)
          if (skipped_flag || no_latent) {
            status <- "skipped"
            if (is.na(err)) err <- (out$extra_json$reason %||% out$extra$reason) %||% "no fused latent returned"
          }
        }

        ev <- tryCatch(
          run_and_eval_one(
            method_name = method_name,
            res_out     = out,
            obj_cells   = cell_ids_fold,
            labels      = labels_fold,
            splits_idx  = splits_fold,
            seed        = seed,
            fold        = fold,
            status      = status,
            error_msg   = err,
            fit_seconds = fit_seconds,
            k_mix       = k_mix,
            k_lt        = k_lt,
            fos_sub     = fos_sub,
            verbose     = verbose
          ),
          error=function(e) {
            run_and_eval_one(
              method_name = method_name,
              res_out     = NULL,
              obj_cells   = cell_ids_fold,
              labels      = labels_fold,
              splits_idx  = splits_fold,
              seed        = seed,
              fold        = fold,
              status      = "failed",
              error_msg   = paste0("run_and_eval_one failed: ", conditionMessage(e)),
              fit_seconds = fit_seconds,
              k_mix       = k_mix,
              k_lt        = k_lt,
              fos_sub     = fos_sub,
              verbose     = FALSE
            )
          }
        )

        runs[[rr]] <- data.table::as.data.table(ev); rr <- rr + 1L

        if (identical(status, "ok")) {
          .msg(sprintf("[cv] EXIT  method=%s seed=%d fold=%d OK (%.1fs)", method_name, seed, fold, fit_seconds), verbose=verbose)
        } else {
          .msg(sprintf("[cv] EXIT  method=%s seed=%d fold=%d %s: %s", method_name, seed, fold, toupper(status), err %||% ""), verbose=verbose)
        }
      }
    }
  }

  data.table::rbindlist(runs, fill=TRUE)
}


# %% [code]
options(future.stdout = TRUE)
options(future.conditions = "message")
Sys.setenv(R_FUTURE_STDOUT = "true")


# %% [code]
cv_out <- run_cv_sweep_R_fixed(
  seeds = SEEDS,
  n_folds = N_FOLDS,
  obj = obj,
  labels = labels,
  methods = METHODS,
  out_dir = OUT_DIR,
  future_globals_max_gb = 50,
  verbose = TRUE,
  
  # these will flow into ... and reach bridge
  future_workers = 24,
  blas_threads   = 24,
  pca_approx     = TRUE
)


# %% [code]
dt_runs <- cv_out
names(dt_runs)


# %% [code]
summ <- summarize_metrics_all(dt_runs, by = "method")


# %% [code]
f_runs <- file.path(OUT_DIR, "cv_runs.tsv")
f_summ <- file.path(OUT_DIR, "cv_summary.tsv")

fwrite(dt_runs, f_runs, sep="\t")
fwrite(summ, f_summ, sep="\t")

cat("Saved per-run:", f_runs, "\n")
cat("Saved summary:", f_summ, "\n")


# %% [code]
print(head(dt_runs))
#print(summ[order(-fit_seconds_mean)])


# %% [code]


# %% [code]


# %% [code]


# %% [code]


# %% [code]


# %% [code]


# %% [code]


# %% [code]


# %% [code]


# %% [code]


# %% [code]


# %% [code]


# %% [code]


# %% [code]


# %% [code]


# %% [code]


# %% [code]


# %% [code]


# %% [code]


# %% [code]


# %% [code]


# %% [code]
```
