## Supplemental Code for "Unifying multimodal single-cell data with a mixture-of-experts *β*-variational autoencoder framework": Fig09_scaling_ablation.ipynb

```
# %% [markdown]
# UniVI – Figure 9 analysis (robust v1, no leakage)

This notebook builds the Figure 8 missing-modality / overlap curve **without train–test leakage** and with **loss_mode='v1'**.

Key idea for v1 training with missing modalities. We enforce mini-batches with paired and mini-batches with unimodal data with a deterministic mask + grouped batch sampler. The unimodal data VAEs are treated as unimodal VAEs during training with only self-reconstruction and the beta-weighted prior KL divergence term (no KL-align term included in these mini-batches).


# %% [code]
# ---- 0) Imports / versions ----
import numpy as np
import pandas as pd
import scipy.sparse as sp
import anndata as ad
import scanpy as sc

from dataclasses import dataclass
from typing import Optional, Dict, Any, Mapping, Tuple, List, Sequence

import torch
from torch.utils.data import Dataset, DataLoader, Subset, Sampler

from sklearn.feature_extraction.text import TfidfTransformer
from sklearn.decomposition import TruncatedSVD
from sklearn.preprocessing import StandardScaler
from sklearn.metrics import accuracy_score, f1_score

# UniVI
import univi as uv
from univi import UniVIMultiModalVAE, ModalityConfig, UniVIConfig, TrainingConfig
from univi.trainer import UniVITrainer
from univi.data import MultiModalDataset, align_paired_obs_names

print("torch:", "v" + torch.__version__)
print("scanpy:", "v" + sc.__version__)
print("univi:", "v" + uv.__version__)


# %% [code]
import matplotlib as mpl
mpl.rcParams["figure.dpi"] = 300        # inline display dpi
mpl.rcParams["savefig.dpi"] = 300       # default save dpi


# %% [markdown]
## 1) Preprocessing without leakage

We fit RNA HVGs + normalization and ATAC TF-IDF/LSI on the training split only, then apply the exact transforms to val/test.


# %% [code]
from dataclasses import dataclass
from typing import List, Tuple, Dict

@dataclass
class RNAFit:
    hvg: List[str]
    counts_layer: str = "counts"
    target_sum: float = 1e4
    out_layer: str = "log1p"

@dataclass
class ATACFit:
    counts_layer: str = "counts"
    tfidf: TfidfTransformer = None
    svd: TruncatedSVD = None
    scaler: StandardScaler = None
    drop_first: bool = True

def _ensure_layer(adata: ad.AnnData, layer: str):
    if layer not in adata.layers:
        adata.layers[layer] = adata.X.copy()

def fit_rna_on_train(
    rna_train: ad.AnnData,
    *,
    counts_layer: str = "counts",
    n_hvg: int = 2000,
    target_sum: float = 1e4,
    out_layer: str = "log1p",
    hvg_flavor: str = "seurat_v3",
) -> RNAFit:
    rna_train = rna_train.copy()
    _ensure_layer(rna_train, counts_layer)

    tmp = rna_train.copy()
    tmp.X = tmp.layers[counts_layer]
    try:
        sc.pp.highly_variable_genes(tmp, n_top_genes=int(n_hvg), flavor=hvg_flavor)
    except Exception:
        # fallback that doesn't require scikit-misc
        sc.pp.highly_variable_genes(tmp, n_top_genes=int(n_hvg), flavor="cell_ranger")

    hvg = tmp.var_names[tmp.var["highly_variable"].to_numpy()].tolist()
    return RNAFit(hvg=hvg, counts_layer=counts_layer, target_sum=float(target_sum), out_layer=str(out_layer))

def apply_rna_fit(rna: ad.AnnData, fit: RNAFit) -> ad.AnnData:
    _ensure_layer(rna, fit.counts_layer)
    a = rna[:, fit.hvg].copy()
    a.layers[fit.out_layer] = a.layers[fit.counts_layer].copy()
    sc.pp.normalize_total(a, target_sum=float(fit.target_sum), layer=fit.out_layer)
    sc.pp.log1p(a, layer=fit.out_layer)
    a.X = a.layers[fit.out_layer]
    return a

def fit_atac_on_train(
    atac_train: ad.AnnData,
    *,
    counts_layer: str = "counts",
    n_lsi: int = 50,
    seed: int = 0,
    tfidf_norm: str = "l2",
    smooth_idf: bool = True,
    sublinear_tf: bool = False,
    drop_first: bool = True,
) -> ATACFit:
    atac_train = atac_train.copy()
    _ensure_layer(atac_train, counts_layer)
    X = atac_train.layers[counts_layer]
    if not sp.issparse(X):
        X = sp.csr_matrix(X)

    tfidf = TfidfTransformer(norm=tfidf_norm, use_idf=True, smooth_idf=bool(smooth_idf), sublinear_tf=bool(sublinear_tf))
    Xt = tfidf.fit_transform(X)

    svd = TruncatedSVD(n_components=int(n_lsi), random_state=int(seed))
    Z = svd.fit_transform(Xt)

    scaler = StandardScaler(with_mean=True, with_std=True)
    scaler.fit(Z)

    return ATACFit(counts_layer=counts_layer, tfidf=tfidf, svd=svd, scaler=scaler, drop_first=bool(drop_first))

def apply_atac_fit(atac: ad.AnnData, fit: ATACFit) -> ad.AnnData:
    _ensure_layer(atac, fit.counts_layer)
    X = atac.layers[fit.counts_layer]
    if not sp.issparse(X):
        X = sp.csr_matrix(X)

    Xt = fit.tfidf.transform(X)
    Z  = fit.svd.transform(Xt)
    Z  = fit.scaler.transform(Z).astype(np.float32, copy=False)

    if fit.drop_first and Z.shape[1] >= 2:
        Z = Z[:, 1:]

    return ad.AnnData(
        X=Z,
        obs=atac.obs.copy(),
        var=pd.DataFrame(index=[f"LSI_{i}" for i in range(1, Z.shape[1] + 1)]),
    )

def preprocess_multiome_splits_fit_apply(
    rna_train: ad.AnnData, atac_train: ad.AnnData,
    rna_val: ad.AnnData,   atac_val: ad.AnnData,
    rna_test: ad.AnnData,  atac_test: ad.AnnData,
    *,
    rna_counts_layer: str = "counts",
    atac_counts_layer: str = "counts",
    n_hvg: int = 2000,
    target_sum: float = 1e4,
    n_lsi: int = 50,
    seed: int = 0,
) -> Tuple[ad.AnnData, ad.AnnData, ad.AnnData, ad.AnnData, ad.AnnData, ad.AnnData, RNAFit, ATACFit]:

    for a in (rna_train, rna_val, rna_test):
        _ensure_layer(a, rna_counts_layer)
    for a in (atac_train, atac_val, atac_test):
        _ensure_layer(a, atac_counts_layer)

    rna_fit = fit_rna_on_train(rna_train, counts_layer=rna_counts_layer, n_hvg=n_hvg, target_sum=target_sum)
    atac_fit = fit_atac_on_train(atac_train, counts_layer=atac_counts_layer, n_lsi=n_lsi, seed=seed)

    rna_tr_pp = apply_rna_fit(rna_train, rna_fit)
    rna_va_pp = apply_rna_fit(rna_val,   rna_fit)
    rna_te_pp = apply_rna_fit(rna_test,  rna_fit)

    atac_tr_lsi = apply_atac_fit(atac_train, atac_fit)
    atac_va_lsi = apply_atac_fit(atac_val,   atac_fit)
    atac_te_lsi = apply_atac_fit(atac_test,  atac_fit)

    return (rna_tr_pp, atac_tr_lsi,
            rna_va_pp, atac_va_lsi,
            rna_te_pp, atac_te_lsi,
            rna_fit, atac_fit)


# %% [markdown]
### Provide your splits

Define `rna_train/val/test` and `atac_train/val/test` (paired and in the same order within each split), then run preprocessing.


# %% [code]
rna_raw = sc.read_h5ad("./data/10x_Genomics_Multiome_data/10x-Multiome-Pbmc10k-RNA.h5ad")
atac_raw = sc.read_h5ad("./data/10x_Genomics_Multiome_data/10x-Multiome-Pbmc10k-ATAC.h5ad")


# %% [code]
rna_raw.layers['counts'] = rna_raw.X
atac_raw.layers['counts'] = atac_raw.X


# %% [code]
from univi.data import align_paired_obs_names

adata_dict_raw = {"rna": rna_raw, "atac": atac_raw}
adata_dict_raw = align_paired_obs_names(adata_dict_raw)

rna_raw = adata_dict_raw["rna"]
atac_raw = adata_dict_raw["atac"]

print(rna_raw.n_obs, atac_raw.n_obs)
assert (rna_raw.obs_names == atac_raw.obs_names).all()


# %% [code]
# ---- 1) Sanity checks ----
print(rna_raw)
print("rna X min/max:", float(rna_raw.X.min()), float(rna_raw.X.max()))
print(atac_raw)
print("atac X min/max:", float(atac_raw.X.min()), float(atac_raw.X.max()))

# Recommended: store raw counts in .layers['counts'] if not already there.
def ensure_counts_layer(adata: ad.AnnData, layer: str = "counts") -> None:
    if layer not in adata.layers:
        adata.layers[layer] = adata.X.copy()

ensure_counts_layer(rna_raw, "counts")
ensure_counts_layer(atac_raw, "counts")


# %% [code]
# ---- 2) Create splits (skip if you already have rna_train/val/test etc) ----
if "rna_train" not in globals():
    rng = np.random.default_rng(0)
    n = rna_raw.n_obs
    idx = np.arange(n)
    rng.shuffle(idx)

    n_tr = int(0.8 * n)
    n_va = int(0.1 * n)
    tr_idx = idx[:n_tr]
    va_idx = idx[n_tr:n_tr+n_va]
    te_idx = idx[n_tr+n_va:]

    rna_train, rna_val, rna_test = rna_raw[tr_idx].copy(), rna_raw[va_idx].copy(), rna_raw[te_idx].copy()
    atac_train, atac_val, atac_test = atac_raw[tr_idx].copy(), atac_raw[va_idx].copy(), atac_raw[te_idx].copy()

print("splits:", rna_train.n_obs, rna_val.n_obs, rna_test.n_obs)


# %% [code]
# ---- 3) Preprocessing: fit on train only, apply to splits ----

@dataclass
class RNAFit:
    hvg: List[str]
    counts_layer: str = "counts"
    target_sum: float = 1e4
    out_layer: str = "log1p"

@dataclass
class ATACFit:
    counts_layer: str = "counts"
    tfidf: Any = None
    svd: Any = None
    scaler: Any = None
    n_lsi: int = 101
    drop_first: bool = True

def _to_csr(X):
    return X if sp.issparse(X) else sp.csr_matrix(X)

def fit_rna_on_train(
    rna_train: ad.AnnData,
    *,
    counts_layer: str = "counts",
    n_hvg: int = 2000,
    target_sum: float = 1e4,
    out_layer: str = "log1p",
) -> RNAFit:
    rna = rna_train.copy()
    if counts_layer not in rna.layers:
        rna.layers[counts_layer] = rna.X.copy()

    # Build a stable working layer for HVG selection
    rna.layers[out_layer] = _to_csr(rna.layers[counts_layer]).copy()
    sc.pp.normalize_total(rna, target_sum=float(target_sum), layer=out_layer)
    sc.pp.log1p(rna, layer=out_layer)

    # Prefer seurat_v3 if available; fall back cleanly if skmisc missing.
    tried = []
    for flavor in ["seurat_v3", "cell_ranger", "seurat"]:
        try:
            tried.append(flavor)
            sc.pp.highly_variable_genes(rna, n_top_genes=int(n_hvg), flavor=flavor, layer=out_layer)
            break
        except Exception as e:
            last = e
            continue
    else:
        raise RuntimeError(f"HVG selection failed. Tried: {tried}. Last error: {last}")

    hvg = rna.var_names[rna.var["highly_variable"].to_numpy()].tolist()
    if len(hvg) == 0:
        raise RuntimeError("No HVGs selected; check that RNA layer values are sane.")
    return RNAFit(hvg=hvg, counts_layer=counts_layer, target_sum=float(target_sum), out_layer=out_layer)

def apply_rna_fit(rna: ad.AnnData, fit: RNAFit) -> ad.AnnData:
    a = rna[:, fit.hvg].copy()
    if fit.counts_layer not in a.layers:
        a.layers[fit.counts_layer] = a.X.copy()

    a.layers[fit.out_layer] = _to_csr(a.layers[fit.counts_layer]).copy()
    sc.pp.normalize_total(a, target_sum=float(fit.target_sum), layer=fit.out_layer)
    sc.pp.log1p(a, layer=fit.out_layer)

    a.X = a.layers[fit.out_layer].copy()
    return a

def fit_atac_on_train(
    atac_train: ad.AnnData,
    *,
    counts_layer: str = "counts",
    n_lsi: int = 50,
    seed: int = 0,
) -> ATACFit:
    a = atac_train.copy()
    if counts_layer not in a.layers:
        a.layers[counts_layer] = a.X.copy()

    X = _to_csr(a.layers[counts_layer])

    tfidf = TfidfTransformer(norm="l2", use_idf=True, smooth_idf=True, sublinear_tf=False)
    X_t = tfidf.fit_transform(X)

    svd = TruncatedSVD(n_components=int(n_lsi), random_state=int(seed))
    Z = svd.fit_transform(X_t)

    scaler = StandardScaler(with_mean=True, with_std=True)
    Zs = scaler.fit_transform(Z).astype(np.float32, copy=False)

    return ATACFit(counts_layer=counts_layer, tfidf=tfidf, svd=svd, scaler=scaler, n_lsi=int(n_lsi), drop_first=True)

def apply_atac_fit(atac: ad.AnnData, fit: ATACFit) -> ad.AnnData:
    a = atac.copy()
    if fit.counts_layer not in a.layers:
        a.layers[fit.counts_layer] = a.X.copy()

    X = _to_csr(a.layers[fit.counts_layer])
    Xt = fit.tfidf.transform(X)
    Z = fit.svd.transform(Xt)
    Z = fit.scaler.transform(Z).astype(np.float32, copy=False)

    if fit.drop_first and Z.shape[1] >= 2:
        Z = Z[:, 1:]

    atac_lsi = ad.AnnData(
        X=Z,
        obs=a.obs.copy(),
        var=pd.DataFrame(index=[f"LSI_{i}" for i in range(1, Z.shape[1] + 1)]),
    )
    return atac_lsi

def preprocess_multiome_splits_fit_apply(
    rna_train: ad.AnnData, atac_train: ad.AnnData,
    rna_val: ad.AnnData,   atac_val: ad.AnnData,
    rna_test: ad.AnnData,  atac_test: ad.AnnData,
    *,
    rna_counts_layer="counts",
    atac_counts_layer="counts",
    n_hvg=2000,
    target_sum=1e4,
    n_lsi=101,
    seed=0,
):
    rna_fit = fit_rna_on_train(
        rna_train, counts_layer=rna_counts_layer, n_hvg=n_hvg, target_sum=target_sum
    )
    atac_fit = fit_atac_on_train(
        atac_train, counts_layer=atac_counts_layer, n_lsi=n_lsi, seed=seed
    )

    rna_tr = apply_rna_fit(rna_train, rna_fit)
    rna_va = apply_rna_fit(rna_val, rna_fit)
    rna_te = apply_rna_fit(rna_test, rna_fit)

    atac_tr = apply_atac_fit(atac_train, atac_fit)
    atac_va = apply_atac_fit(atac_val, atac_fit)
    atac_te = apply_atac_fit(atac_test, atac_fit)

    return (rna_tr, atac_tr, rna_va, atac_va, rna_te, atac_te, rna_fit, atac_fit)


# %% [code]
# ---- run preprocessing ----
(rna_tr_pp, atac_tr_lsi,
 rna_va_pp, atac_va_lsi,
 rna_te_pp, atac_te_lsi,
 rna_fit, atac_fit) = preprocess_multiome_splits_fit_apply(
    rna_train, atac_train,
    rna_val, atac_val,
    rna_test, atac_test,
    rna_counts_layer="counts",
    atac_counts_layer="counts",
    n_hvg=2000,
    target_sum=1e4,
    n_lsi=101,
    seed=42,
)

print("RNA HVG:", rna_tr_pp.n_vars, "ATAC LSI dims:", atac_tr_lsi.n_vars)


# %% [markdown]
## 2) Build paired dataset + indices + labels


# %% [code]
adata_dict = {
    "rna":  rna_tr_pp.concatenate(rna_va_pp, rna_te_pp, batch_key=None),
    "atac": atac_tr_lsi.concatenate(atac_va_lsi, atac_te_lsi, batch_key=None),
}
adata_dict = align_paired_obs_names(adata_dict)

dataset = MultiModalDataset(adata_dict=adata_dict, X_key="X", device=None)

n_tr = rna_tr_pp.n_obs
n_va = rna_va_pp.n_obs
n_te = rna_te_pp.n_obs

train_idx = np.arange(0, n_tr)
val_idx   = np.arange(n_tr, n_tr + n_va)
test_idx  = np.arange(n_tr + n_va, n_tr + n_va + n_te)

label_key = "cell_type"  # change if needed
y_all = adata_dict["rna"].obs[label_key].to_numpy()

print("dataset n:", len(dataset), "train/val/test:", n_tr, n_va, n_te)
print("labels:", pd.Series(y_all).value_counts())


# %% [code]
import numpy as np
import torch
from torch.utils.data import Dataset

# --- 1) make integer labels (stable mapping) ---
label_key = "cell_type"
y_str = adata_dict["rna"].obs[label_key].astype(str).to_numpy()

uniq = np.unique(y_str)
label_to_int = {lab: i for i, lab in enumerate(uniq)}
y_int = np.array([label_to_int[x] for x in y_str], dtype=np.int64)

print("n labels:", len(uniq))
print("example mapping:", list(label_to_int.items())[:5])

# --- 2) wrapper dataset that injects labels into each sample dict ---
class LabeledMultiModalDataset(Dataset):
    def __init__(self, base_ds, y_int, y_str=None):
        self.base = base_ds
        self.y_int = np.asarray(y_int, dtype=np.int64)
        self.y_str = None if y_str is None else np.asarray(y_str, dtype=object)
        if len(self.base) != len(self.y_int):
            raise ValueError(f"len(base)={len(self.base)} != len(y_int)={len(self.y_int)}")

    def __len__(self):
        return len(self.base)

    def __getitem__(self, i):
        x = self.base[i]
        # base is expected to yield dict-like
        if not isinstance(x, dict):
            # if it yields (dict, ...) take first
            if isinstance(x, (tuple, list)) and len(x) > 0 and isinstance(x[0], dict):
                x = x[0]
            else:
                raise TypeError(f"Expected dict or (dict,...); got {type(x)}")

        x = dict(x)
        x["y"] = torch.tensor(int(self.y_int[i]), dtype=torch.long)
        if self.y_str is not None:
            x["cell_type"] = self.y_str[i]  # NOTE: strings won't collate unless you handle them
        return x

# --- 3) replace dataset with labeled version (use y_int only for safety) ---
dataset = LabeledMultiModalDataset(dataset, y_int=y_int, y_str=None)


# %% [markdown]
## 3) v1-safe loaders for missing modalities

In v1, a batch cannot mix RNA-only and ATAC-only items (or you hit the `stack expects each tensor to be equal size` crash). We fix this by:

1. Creating a **deterministic** per-item group assignment for the training split (paired vs unimodal).
2. Using a **grouped batch sampler** so each batch contains items from a single group.

Validation and test loaders remain fully paired.


# %% [code]
import numpy as np
import torch
from torch.utils.data import Dataset, DataLoader, Sampler

# -----------------------------
# Dataset wrappers
# -----------------------------
class IndexedDataset(Dataset):
    """Wrap base_dataset + an index list, and return (x_dict, global_id)."""
    def __init__(self, base_dataset, indices):
        self.base = base_dataset
        self.indices = np.asarray(indices, dtype=np.int64)

    def __len__(self):
        return int(self.indices.shape[0])

    def __getitem__(self, i):
        gi = int(self.indices[int(i)])
        item = self.base[gi]

        if isinstance(item, dict):
            x = item
        elif isinstance(item, (tuple, list)) and len(item) >= 1 and isinstance(item[0], dict):
            x = item[0]
        else:
            raise TypeError(f"Expected base_dataset to yield dict or (dict, ...), got {type(item)}")

        return dict(x), gi


class DeterministicMaskDataset(Dataset):
    """
    Mask exactly one modality for non-anchor rows based on groups.
      groups==0: paired anchor (keep both)
      groups!=0: unpaired (DROP drop_modality key)
    """
    def __init__(self, base_ds: Dataset, groups: np.ndarray, *, drop_modality: str):
        self.base = base_ds
        self.groups = np.asarray(groups, dtype=np.int64)
        self.drop_modality = str(drop_modality)
        if len(self.base) != len(self.groups):
            raise ValueError(f"base_ds len={len(self.base)} != groups len={len(self.groups)}")

    def __len__(self):
        return len(self.base)

    def __getitem__(self, i):
        x, gid = self.base[i]
        g = int(self.groups[int(i)])

        x = dict(x)
        if g != 0:
            x.pop(self.drop_modality, None)  # IMPORTANT: remove key, don't set None
        return x, gid


# -----------------------------
# Collate: homogeneous key-set guaranteed by sampler + mask dataset
# -----------------------------
def collate_xdict_with_idx(batch):
    """Collate list[(x_dict, gid)] into (dict[str, Tensor], gids Tensor)."""
    if len(batch) == 0:
        raise ValueError("Empty batch")

    gids = torch.as_tensor([b[1] for b in batch], dtype=torch.long)

    keys0 = set(batch[0][0].keys())
    for x, _ in batch[1:]:
        if set(x.keys()) != keys0:
            raise RuntimeError(
                "Non-homogeneous batch detected (mixed modality presence within a batch). "
                "This breaks UniVI v1 MoE assumptions. Fix sampler/grouping."
            )

    out = {}
    for k in keys0:
        out[k] = torch.stack([x[k] for x, _ in batch], dim=0)

    return out, gids


# -----------------------------
# Sampler: paired-driven homogeneous batches with unpaired interleaving
# -----------------------------
class InterleavedGroupedBatchSampler(Sampler):
    """
    Paired-driven homogeneous batch sampler (with safe tail behavior).

    Backbone: yields paired (group==0) batches.
    Interleave: yields unpaired (group!=0) batches such that the expected number of
    unpaired batches per paired batch ~= unpaired_per_paired.

    Always yields homogeneous batches (all paired OR all unpaired).

    Tail fix:
      - If paired_idx < batch_size, we DO NOT error.
        Instead, if oversample_paired=True, we run with n_paired_batches=1 and
        generate paired batches by sampling WITH replacement.
      - If oversample_paired=False and paired_idx < batch_size, we fall back to
        yielding only unpaired batches (still homogeneous). This keeps things running,
        but you lose paired anchors in that extreme.
    """
    def __init__(
        self,
        groups: np.ndarray,
        batch_size: int,
        *,
        seed: int = 0,
        drop_last: bool = True,
        unpaired_per_paired: float = 1.0,
        oversample_paired: bool = True,
        oversample_unpaired: bool = True,
    ):
        self.groups = np.asarray(groups, dtype=np.int64)
        self.batch_size = int(batch_size)
        self.seed = int(seed)
        self.drop_last = bool(drop_last)

        upp = float(unpaired_per_paired)
        if not np.isfinite(upp) or upp < 0:
            raise ValueError(f"unpaired_per_paired must be finite and >=0, got {unpaired_per_paired}")
        self.unpaired_per_paired = upp

        self.oversample_paired = bool(oversample_paired)
        self.oversample_unpaired = bool(oversample_unpaired)

        self.paired_idx = np.where(self.groups == 0)[0].astype(np.int64)
        self.unpaired_idx = np.where(self.groups != 0)[0].astype(np.int64)

        if len(self.paired_idx) == 0:
            raise ValueError("No paired samples (group==0). Need at least one anchor sample.")

        self.only_paired = (len(self.unpaired_idx) == 0)

        # Can we make at least one full paired batch without replacement?
        self.can_full_paired = (len(self.paired_idx) >= self.batch_size)

        # Paired-driven epoch length:
        # - normal: floor(P / B)
        # - tail: if P < B and oversample_paired=True => force 1 paired batch (replacement)
        # - tail: if P < B and oversample_paired=False => 0 paired batches (we'll yield only unpaired)
        if self.can_full_paired:
            self.n_paired_batches = len(self.paired_idx) // self.batch_size
        else:
            self.n_paired_batches = 1 if self.oversample_paired else 0

        if self.only_paired and self.n_paired_batches <= 0:
            # Only paired data exists but too small and oversample_paired=False => cannot proceed.
            raise ValueError(
                "Paired-only training split has fewer samples than batch_size and oversample_paired=False."
            )

        # Expected total batches (for progress bars / DataLoader length)
        if self.only_paired:
            self.n_unpaired_expected = 0
        else:
            self.n_unpaired_expected = int(round(self.n_paired_batches * self.unpaired_per_paired))

        # If we have no paired batches (tail + oversample_paired=False), define length from unpaired
        if self.n_paired_batches == 0:
            # define an epoch as all full unpaired batches if possible; else 1 replacement batch if allowed
            if len(self.unpaired_idx) >= self.batch_size:
                self.n_unpaired_backbone = len(self.unpaired_idx) // self.batch_size
            else:
                self.n_unpaired_backbone = 1 if self.oversample_unpaired else 0

            if self.n_unpaired_backbone <= 0:
                raise ValueError("Not enough samples to form even one batch at this batch_size.")

            self.n_batches = int(self.n_unpaired_backbone)
        else:
            self.n_batches = int(self.n_paired_batches + self.n_unpaired_expected)

        self.n_batches = max(int(self.n_batches), 1)

    def __len__(self):
        return int(self.n_batches)

    def __iter__(self):
        rng = np.random.default_rng(self.seed)

        paired_pool = self.paired_idx.copy()
        rng.shuffle(paired_pool)
        p_ptr = 0

        unpaired_pool = self.unpaired_idx.copy()
        if len(unpaired_pool) > 0:
            rng.shuffle(unpaired_pool)
        u_ptr = 0

        def pop_full_no_wrap(pool, ptr):
            end = ptr + self.batch_size
            if end <= len(pool):
                return pool[ptr:end], end
            return None, ptr

        def pop_full_with_replacement(pool):
            if len(pool) == 0:
                return None
            idx = rng.integers(0, len(pool), size=self.batch_size)
            return pool[idx]

        def next_paired_batch():
            nonlocal p_ptr, paired_pool
            # If pool is smaller than batch_size, only replacement sampling can work
            if len(paired_pool) < self.batch_size:
                return pop_full_with_replacement(paired_pool) if self.oversample_paired else None

            pb, p_ptr2 = pop_full_no_wrap(paired_pool, p_ptr)
            if pb is None:
                if not self.oversample_paired:
                    return None
                rng.shuffle(paired_pool)
                p_ptr = 0
                pb, p_ptr2 = pop_full_no_wrap(paired_pool, p_ptr)
                if pb is None:
                    return None
            p_ptr = p_ptr2
            return pb

        def next_unpaired_batch():
            nonlocal u_ptr, unpaired_pool
            # If pool is smaller than batch_size, only replacement sampling can work
            if len(unpaired_pool) < self.batch_size:
                return pop_full_with_replacement(unpaired_pool) if self.oversample_unpaired else None

            ub, u_ptr2 = pop_full_no_wrap(unpaired_pool, u_ptr)
            if ub is not None:
                u_ptr = u_ptr2
                return ub

            if not self.oversample_unpaired:
                return None

            rng.shuffle(unpaired_pool)
            u_ptr = 0
            ub, u_ptr2 = pop_full_no_wrap(unpaired_pool, u_ptr)
            if ub is None:
                return None
            u_ptr = u_ptr2
            return ub

        # Case A: only paired data
        if self.only_paired:
            for _ in range(self.n_paired_batches):
                pb = next_paired_batch()
                if pb is None:
                    break
                yield pb.tolist()
            return

        # Case B: no paired batches in this epoch (tail + oversample_paired=False)
        if self.n_paired_batches == 0:
            # define epoch as unpaired backbone only
            # (still homogeneous; you just don't see paired anchors)
            # yield all full unpaired batches; if too small, yield one replacement batch (if allowed)
            if len(unpaired_pool) >= self.batch_size:
                n_unpaired_backbone = len(unpaired_pool) // self.batch_size
                for _ in range(n_unpaired_backbone):
                    ub = next_unpaired_batch()
                    if ub is None:
                        break
                    yield ub.tolist()
            else:
                ub = next_unpaired_batch()
                if ub is not None:
                    yield ub.tolist()
            return

        # Case C: mixed (paired-driven backbone, interleave unpaired)
        carry = 0.0
        for _ in range(self.n_paired_batches):
            pb = next_paired_batch()
            if pb is None:
                break
            yield pb.tolist()

            carry += self.unpaired_per_paired
            while carry >= 1.0:
                carry -= 1.0
                ub = next_unpaired_batch()
                if ub is None:
                    break
                yield ub.tolist()


# -----------------------------
# main loader builder
# -----------------------------
def make_loaders_with_overlap_v1(
    base_dataset,
    train_idx,
    val_idx,
    test_idx,
    *,
    overlap_fraction: float,
    drop_modality: str,
    seed: int,
    batch_size: int = 256,
    num_workers: int = 0,
    oversample_paired: bool = True,
):
    """overlap_fraction applies ONLY to training. val/test remain paired."""
    p = float(overlap_fraction)
    if not (0.0 <= p <= 1.0):
        raise ValueError(f"overlap_fraction must be in [0,1], got {p}")

    rng = np.random.default_rng(int(seed))

    # --- train subset ---
    train_indexed = IndexedDataset(base_dataset, indices=train_idx)
    N = len(train_indexed)
    if N == 0:
        raise ValueError("Empty train split")

    n_paired = int(round(p * N))
    n_paired = max(min(n_paired, N), 0)

    perm = rng.permutation(N)
    paired_local = perm[:n_paired]

    groups = np.ones(N, dtype=np.int64)  # 1 = unpaired
    groups[paired_local] = 0             # 0 = paired anchor

    train_masked = DeterministicMaskDataset(
        train_indexed, groups=groups, drop_modality=str(drop_modality)
    )

    n_paired_real = int((groups == 0).sum())
    n_unpaired_real = int((groups != 0).sum())

    ratio_unpaired_per_paired = (n_unpaired_real / max(n_paired_real, 1))
    upp_sampling = float(ratio_unpaired_per_paired)  # scale to true split (no clipping)

    print(
        f"[overlap={p:g}] batch_size={int(batch_size)} "
        f"train_paired={n_paired_real} train_unpaired={n_unpaired_real} "
        f"(unpaired_per_paired={ratio_unpaired_per_paired:.6f}, sampler_upp={upp_sampling:.6f})"
    )

    # Train loader: homogeneous batches enforced by sampler (when unpaired exists)
    if n_unpaired_real == 0:
        train_loader = DataLoader(
            train_masked,
            batch_size=int(batch_size),
            shuffle=True,
            drop_last=True,
            num_workers=int(num_workers),
            collate_fn=collate_xdict_with_idx,
        )
    else:
        batch_sampler = InterleavedGroupedBatchSampler(
            groups=groups,
            batch_size=int(batch_size),
            seed=int(seed),
            drop_last=True,
            unpaired_per_paired=upp_sampling,
            oversample_paired=bool(oversample_paired),
            oversample_unpaired=True,
        )
        train_loader = DataLoader(
            train_masked,
            batch_sampler=batch_sampler,
            num_workers=int(num_workers),
            collate_fn=collate_xdict_with_idx,
        )

    # --- val/test: always paired, no masking ---
    val_ds = IndexedDataset(base_dataset, indices=val_idx)
    test_ds = IndexedDataset(base_dataset, indices=test_idx)

    val_loader = DataLoader(
        val_ds,
        batch_size=int(batch_size),
        shuffle=False,
        drop_last=False,
        num_workers=int(num_workers),
        collate_fn=collate_xdict_with_idx,
    )
    test_loader = DataLoader(
        test_ds,
        batch_size=int(batch_size),
        shuffle=False,
        drop_last=False,
        num_workers=int(num_workers),
        collate_fn=collate_xdict_with_idx,
    )

    info = {
        "train_paired": n_paired_real,
        "train_unpaired": n_unpaired_real,
        "unpaired_per_paired": float(ratio_unpaired_per_paired),
        "sampler_unpaired_per_paired": float(upp_sampling),
    }
    return train_loader, val_loader, test_loader, info


# %% [code]
train_loader, val_loader, test_loader, info = make_loaders_with_overlap_v1(
    base_dataset=dataset,     # <- change if your dataset var is named differently
    train_idx=train_idx,
    val_idx=val_idx,
    test_idx=test_idx,
    overlap_fraction=1.0,     # example: 100% paired anchors
    drop_modality="atac",     # or "rna"
    seed=0,
    batch_size=256,
    num_workers=0,
    oversample_paired=True,
)

info


# %% [code]
print("val_idx n =", len(val_idx))
print("test_idx n =", len(test_idx))

def _get_xdict_from_batch(batch):
    """
    Supports:
      - (x_dict, gids) from collate_xdict_with_idx
      - {"rna":..., "atac":...} dict batches
      - {"x_dict": {...}} style (just in case)
    """
    # Most common in your notebook: (x_dict, gids)
    if isinstance(batch, (tuple, list)) and len(batch) >= 1 and isinstance(batch[0], dict):
        return batch[0]

    # Direct dict batch
    if isinstance(batch, dict):
        if "x_dict" in batch and isinstance(batch["x_dict"], dict):
            return batch["x_dict"]
        return batch

    raise TypeError(f"Unknown batch type: {type(batch)}")

def count_paired(loader, n_batches=20, rna_key="rna", atac_key="atac"):
    paired = 0
    total = 0
    for i, batch in enumerate(loader):
        if i >= n_batches:
            break

        x = _get_xdict_from_batch(batch)

        has_rna  = (rna_key in x) and (x[rna_key] is not None)
        has_atac = (atac_key in x) and (x[atac_key] is not None)

        # if tensors exist but are empty, treat as missing
        try:
            if has_rna and hasattr(x[rna_key], "shape") and x[rna_key].shape[0] == 0:
                has_rna = False
            if has_atac and hasattr(x[atac_key], "shape") and x[atac_key].shape[0] == 0:
                has_atac = False
        except Exception:
            pass

        paired += int(has_rna and has_atac)
        total += 1

    return paired, total

vp, vt = count_paired(val_loader, n_batches=20)
tp, tt = count_paired(test_loader, n_batches=20)
print(f"val paired batches ~ {vp}/{vt}")
print(f"test paired batches ~ {tp}/{tt}")

# Optional: show what keys you actually have
xb, gids = next(iter(val_loader))
print("val batch keys:", list(xb.keys()) if isinstance(xb, dict) else type(xb))


# %% [markdown]
## 4) Model + training (v1)


# %% [code]
import inspect
import numpy as np
import torch
from dataclasses import dataclass
from univi import UniVIMultiModalVAE, ModalityConfig, UniVIConfig, TrainingConfig
from univi.trainer import UniVITrainer

# -----------------------------
# Config builders (UNCHANGED schedule)
# -----------------------------
def make_univi_cfg(rna_dim: int, atac_dim: int) -> UniVIConfig:
    return UniVIConfig(
        latent_dim=30,
        beta=1.25,
        gamma=4.35,
        encoder_dropout=0.10,
        decoder_dropout=0.05,
        encoder_batchnorm=True,
        decoder_batchnorm=False,
        #kl_anneal_start=50,
        #kl_anneal_end=105,
        #align_anneal_start=75,
        #align_anneal_end=140,
        kl_anneal_start=0,
        kl_anneal_end=60,
        align_anneal_start=25,
        align_anneal_end=85,
        modalities=[
            ModalityConfig(
                name="rna",
                input_dim=int(rna_dim),
                encoder_hidden=[512, 256, 128],
                decoder_hidden=[128, 256, 512],
                likelihood="gaussian",
            ),
            ModalityConfig(
                name="atac",
                input_dim=int(atac_dim),
                encoder_hidden=[128, 64],
                decoder_hidden=[64, 128],
                likelihood="gaussian",
            ),
        ],
        class_heads=[],
    )

def make_train_cfg(device, *, early_stopping=True, patience=100, min_delta=0.0) -> TrainingConfig:
    return TrainingConfig(
        n_epochs=5000,
        batch_size=256,
        lr=1e-3,
        weight_decay=1e-4,
        device=device,
        log_every=100,
        grad_clip=5.0,
        early_stopping=bool(early_stopping),
        patience=int(patience),
        min_delta=float(min_delta),
    )

# -----------------------------
# Device selection
# -----------------------------
device = torch.device(
    "cuda" if torch.cuda.is_available()
    else ("mps" if torch.backends.mps.is_available() else "cpu")
)
print("device:", device)

# -----------------------------
# Trainer compat
# -----------------------------
def _build_univi_trainer_compat(*, model, train_cfg, train_loader, val_loader):
    sig = inspect.signature(UniVITrainer.__init__)
    params = sig.parameters
    kwargs = {}

    if "model" in params:
        kwargs["model"] = model
    if "train_loader" in params:
        kwargs["train_loader"] = train_loader
    if "val_loader" in params:
        kwargs["val_loader"] = val_loader

    for cand in ("train_cfg", "training_cfg", "cfg_train", "config", "cfg"):
        if cand in params:
            kwargs[cand] = train_cfg
            return UniVITrainer(**kwargs)

    for k, v in vars(train_cfg).items():
        if k in params:
            kwargs[k] = v

    return UniVITrainer(**kwargs)

def _trainer_fit_compat(trainer, train_loader, val_loader):
    try:
        return trainer.fit()
    except TypeError:
        pass
    try:
        return trainer.fit(train_loader=train_loader, val_loader=val_loader)
    except TypeError:
        pass
    return trainer.fit(train_loader, val_loader)

@dataclass
class TrainResult:
    model: torch.nn.Module
    warmup_epochs: int
    best_epoch_all: int | None
    best_val_all: float | None
    best_epoch_postwarmup: int | None
    best_val_postwarmup: float | None

def _extract_val_history(trainer):
    """Best-effort: pull a list of validation losses (floats) if trainer exposes it."""
    for cand in ("val_losses", "val_loss_history", "history_val", "val_history", "history"):
        if not hasattr(trainer, cand):
            continue
        h = getattr(trainer, cand)
        if isinstance(h, dict):
            # common pattern: {"train_loss":[...], "val_loss":[...]}
            for kk in ("val_loss", "val", "val_losses"):
                if kk in h and isinstance(h[kk], (list, tuple)) and len(h[kk]) > 0:
                    h2 = h[kk]
                    if isinstance(h2[0], (float, int, np.floating, np.integer)):
                        return [float(x) for x in h2]
        if isinstance(h, (list, tuple)) and len(h) > 0 and isinstance(h[0], (float, int, np.floating, np.integer)):
            return [float(x) for x in h]
    return None

# -----------------------------
# Training wrapper: do NOT shift anneals; only *report* best post-warmup.
# Also: prevent early stopping from firing before warmup by inflating patience.
# -----------------------------
def train_one_overlap_v1(
    *,
    train_loader,
    val_loader,
    seed: int = 42,
    device=device,
    loss_mode: str = "v1",
    v1_recon: str = "avg",
    warmup_epochs: int = 50,   # <-- only affects reporting + inflated patience
    patience: int = 5000,
    min_delta: float = 0.0,
):
    torch.manual_seed(int(seed))
    np.random.seed(int(seed))

    rna_dim = int(rna_tr_pp.n_vars)
    atac_dim = int(atac_tr_lsi.n_vars)

    univi_cfg = make_univi_cfg(rna_dim=rna_dim, atac_dim=atac_dim)

    w = int(max(warmup_epochs, 0))

    # IMPORTANT: do not stop before warmup.
    # We can't easily override UniVITrainer's internal ES across versions,
    # so we make the trainer "too patient" to early-stop during warmup.
    patience_eff = int(patience) + w

    train_cfg = make_train_cfg(
        device=device,
        early_stopping=True,
        #patience=patience_eff,
        patience=patience,
        min_delta=float(min_delta),
    )

    model = UniVIMultiModalVAE(
        univi_cfg,
        loss_mode=str(loss_mode),
        v1_recon=str(v1_recon),
    ).to(device)

    trainer = _build_univi_trainer_compat(
        model=model,
        train_cfg=train_cfg,
        train_loader=train_loader,
        val_loader=val_loader,
    )

    # some versions expect these attributes
    if not hasattr(trainer, "train_loader"):
        trainer.train_loader = train_loader
    if not hasattr(trainer, "val_loader"):
        trainer.val_loader = val_loader

    _trainer_fit_compat(trainer, train_loader=train_loader, val_loader=val_loader)

    # --- best overall (trainer-reported if available) ---
    best_epoch_all = getattr(trainer, "best_epoch", None)
    best_val_all = getattr(trainer, "best_val_loss", None)
    if best_val_all is None:
        best_val_all = getattr(trainer, "best_val", None)

    # --- best post-warmup (best-effort) ---
    best_epoch_pw = None
    best_val_pw = None

    hist = _extract_val_history(trainer)
    if hist is not None and len(hist) > w:
        sub = hist[w:]              # epochs are 1-indexed in logs; hist[0] is epoch 1
        j = int(np.argmin(sub))
        best_epoch_pw = w + j + 1
        best_val_pw = float(sub[j])
    else:
        # fallback: only accept trainer's best if it is after warmup
        if isinstance(best_epoch_all, (int, np.integer)) and int(best_epoch_all) >= max(w, 1):
            best_epoch_pw = int(best_epoch_all)
            best_val_pw = float(best_val_all) if best_val_all is not None else None

    return TrainResult(
        model=model,
        warmup_epochs=w,
        best_epoch_all=(int(best_epoch_all) if best_epoch_all is not None else None),
        best_val_all=(float(best_val_all) if best_val_all is not None else None),
        best_epoch_postwarmup=best_epoch_pw,
        best_val_postwarmup=best_val_pw,
    )


# %% [markdown]
## 5) Encoding + metrics (alignment + label transfer)


# %% [code]
import inspect
import numpy as np
import torch

# ============================================================
# Robust latent extraction for UniVI (handles many return layouts)
# + stable label coding across splits via id_to_int
# ============================================================

def _to_numpy_ids(gids):
    """Convert gids to a 1D int64 numpy array."""
    if gids is None:
        return None
    if torch.is_tensor(gids):
        return gids.detach().cpu().numpy().astype(np.int64, copy=False)
    return np.asarray(gids, dtype=np.int64)

def _build_label_encoder(y_all):
    """
    Returns (y_raw, id_to_int_or_None)
    - If y_all is numeric -> y_raw numeric, id_to_int None.
    - If y_all is object/string -> y_raw as str array, id_to_int mapping over ALL unique labels in y_all.
    """
    y_raw = np.asarray(y_all)
    kind = y_raw.dtype.kind
    if kind in ("i", "u", "f", "b"):
        return y_raw, None

    y_raw = y_raw.astype(str)
    uniq = np.unique(y_raw)
    id_to_int = {lab: i for i, lab in enumerate(uniq.tolist())}
    return y_raw, id_to_int

def _encode_labels_for_gids(y_raw, gids_np, *, id_to_int=None):
    """
    Encode labels for the given gids.
    If y_raw is numeric -> returns numeric torch tensor.
    If y_raw is str/object -> returns torch.long codes (using provided id_to_int, extending if needed).
    """
    y_batch = y_raw[gids_np]
    y_batch_arr = np.asarray(y_batch)
    kind = y_batch_arr.dtype.kind

    if kind in ("i", "u", "b"):
        return torch.as_tensor(y_batch_arr.astype(np.int64, copy=False), dtype=torch.long), id_to_int
    if kind == "f":
        return torch.as_tensor(y_batch_arr.astype(np.float32, copy=False)), id_to_int

    # object/string
    y_batch_str = y_batch_arr.astype(str)

    if id_to_int is None:
        uniq = np.unique(y_batch_str)
        id_to_int = {lab: i for i, lab in enumerate(uniq.tolist())}

    codes = np.empty(y_batch_str.shape[0], dtype=np.int64)
    next_code = (max(id_to_int.values()) + 1) if len(id_to_int) else 0
    for i, lab in enumerate(y_batch_str):
        if lab not in id_to_int:
            id_to_int[lab] = next_code
            next_code += 1
        codes[i] = id_to_int[lab]

    return torch.as_tensor(codes, dtype=torch.long), id_to_int

def _unwrap_model_output(out):
    """
    UniVI sometimes returns:
      - dict
      - (loss, dict)
      - (dict, extras...)
      - tensor
    We want the dict/tensor that contains embeddings/latents.
    """
    if torch.is_tensor(out) or isinstance(out, dict):
        return out
    if isinstance(out, (tuple, list)) and len(out) > 0:
        for item in out:
            if isinstance(item, dict) or torch.is_tensor(item):
                return item
        return out[0]
    return out

def _smart_call(fn, x_dict):
    """
    Call fn with whichever kwarg name it seems to want.
    Tries: x_dict / batch / x / inputs ; else positional.
    """
    try:
        sig = inspect.signature(fn)
        params = sig.parameters
        for name in ("x_dict", "batch", "x", "inputs"):
            if name in params:
                return fn(**{name: x_dict})
        return fn(x_dict)
    except TypeError:
        return fn(x_dict)

def _call_univi_encoder(model, x_dict):
    """
    Try encode() first, then forward().
    Uses signature-aware calling and unwraps tuple outputs.
    """
    if hasattr(model, "encode") and callable(getattr(model, "encode")):
        out = _smart_call(model.encode, x_dict)
        out = _unwrap_model_output(out)
        if isinstance(out, (dict, torch.Tensor)):
            return out
    out = _smart_call(model, x_dict)
    return _unwrap_model_output(out)

def _first_tensor(*xs):
    for x in xs:
        if torch.is_tensor(x):
            return x
    return None

def _find_tensors_anywhere(obj, *, path=""):
    """Recursively collect tensors from nested dict/list/tuple structures."""
    out = []
    if torch.is_tensor(obj):
        out.append((path or "tensor", obj))
    elif isinstance(obj, dict):
        for k, v in obj.items():
            out.extend(_find_tensors_anywhere(v, path=f"{path}.{k}" if path else str(k)))
    elif isinstance(obj, (list, tuple)):
        for i, v in enumerate(obj):
            out.extend(_find_tensors_anywhere(v, path=f"{path}[{i}]" if path else f"[{i}]"))
    return out

def _get_path(obj, path_tuple):
    cur = obj
    for k in path_tuple:
        if not isinstance(cur, dict) or k not in cur:
            return None
        cur = cur[k]
    return cur

def _extract_latent(enc, which: str):
    """
    which in {"rna","atac","fused"}.
    Searches many common UniVI layouts/keys and falls back to any 2D tensor found.
    """
    if torch.is_tensor(enc):
        return enc if which == "fused" else None

    if not isinstance(enc, dict):
        return None

    # 1) flat keys
    v = _first_tensor(
        enc.get(f"mu_{which}", None),
        enc.get(f"z_{which}", None),
        enc.get(f"latent_{which}", None),
        enc.get(f"emb_{which}", None),
        enc.get(f"embedding_{which}", None),
    )
    if v is not None:
        return v

    # fused often uses generic keys
    if which == "fused":
        v = _first_tensor(
            enc.get("mu", None), enc.get("z", None), enc.get("mean", None),
            enc.get("latent", None), enc.get("embedding", None),
            enc.get("mu_shared", None), enc.get("z_shared", None),
            enc.get("mu_joint", None), enc.get("z_joint", None),
        )
        if v is not None:
            return v

    # 2) nested containers
    for container in ("latents", "latent", "posterior", "post", "qz", "q", "enc", "encode", "outputs"):
        sub = enc.get(container, None)
        if isinstance(sub, dict):
            v = _first_tensor(
                sub.get(f"mu_{which}", None),
                sub.get(f"z_{which}", None),
                sub.get(which, None) if torch.is_tensor(sub.get(which, None)) else None,
            )
            if v is not None:
                return v

            if which in sub and isinstance(sub[which], dict):
                vv = _first_tensor(
                    sub[which].get("mu", None),
                    sub[which].get("z", None),
                    sub[which].get("mean", None),
                    sub[which].get("latent", None),
                    sub[which].get("embedding", None),
                )
                if vv is not None:
                    return vv

    # 3) top-level modality dict: enc["rna"]={"mu":...}
    if which in enc and isinstance(enc[which], dict):
        vv = _first_tensor(
            enc[which].get("mu", None),
            enc[which].get("z", None),
            enc[which].get("mean", None),
            enc[which].get("latent", None),
            enc[which].get("embedding", None),
        )
        if vv is not None:
            return vv

    # 4) known nested paths
    for p in [
        ("posterior", which, "mu"),
        ("posterior", which, "z"),
        ("latents", which, "mu"),
        ("latents", which, "z"),
        ("qz", which, "mu"),
        ("qz", which, "z"),
    ]:
        sub = _get_path(enc, p[:-1])
        if isinstance(sub, dict):
            vv = sub.get(p[-1], None)
            if torch.is_tensor(vv):
                return vv

    # 5) last resort: any tensor anywhere (prefer 2D)
    tensors = _find_tensors_anywhere(enc)
    if tensors:
        for _, t in tensors:
            if t.ndim == 2:
                return t
        return tensors[0][1]

    return None

@torch.no_grad()
def encode_embeddings_with_labels(
    model,
    loader,
    *,
    device,
    y_all,
    id_to_int=None,
    require_mus=True,
    debug_first_batch=False,
):
    """
    Expects loader batches: (x_dict, gids)

    Returns dict with:
      mu_rna/mu_atac/mu_fused  (each may be None if modality absent),
      y (torch), gids (torch.long),
      id_to_int (stable mapping for string labels)
    """

    def _encode_one(model, x_sub: dict):
        enc = _call_univi_encoder(model, x_sub)
        return _unwrap_model_output(enc)

    model.eval()

    # global label handling (stable mapping)
    y_raw, global_map = _build_label_encoder(y_all)
    kind = np.asarray(y_raw).dtype.kind
    use_map = None
    if kind not in ("i", "u", "f", "b"):
        use_map = id_to_int if id_to_int is not None else global_map

    mu_rna_all, mu_atac_all, mu_fused_all = [], [], []
    y_list, gids_list = [], []

    for b_ix, batch in enumerate(loader):
        if not (isinstance(batch, (tuple, list)) and len(batch) == 2 and isinstance(batch[0], dict)):
            raise RuntimeError(
                "encode_embeddings_with_labels expects batches like (x_dict, gids). "
                "Use collate_xdict_with_idx and IndexedDataset."
            )

        x_dict, gids = batch
        gids_np = _to_numpy_ids(gids)
        if gids_np is None:
            raise RuntimeError("gids is None; loader must provide integer indices for label lookup.")

        # move tensors to device (keep only tensor modalities + anything else model might accept)
        x_full = {}
        for k, v in x_dict.items():
            if v is None:
                continue
            x_full[k] = v.to(device) if torch.is_tensor(v) else v

        # --- (A) fused: encode with whatever is present (often both) ---
        enc_full = _encode_one(model, x_full)

        # --- (B) rna-only / atac-only encodes: FORCE modality-specific outputs ---
        enc_rna = None
        if "rna" in x_full and torch.is_tensor(x_full["rna"]):
            enc_rna = _encode_one(model, {"rna": x_full["rna"]})

        enc_atac = None
        if "atac" in x_full and torch.is_tensor(x_full["atac"]):
            enc_atac = _encode_one(model, {"atac": x_full["atac"]})

        if debug_first_batch and b_ix == 0:
            def _keys(obj):
                return sorted(obj.keys()) if isinstance(obj, dict) else [str(type(obj))]
            print("[latent-debug] enc_full keys:", _keys(enc_full))
            if enc_rna is not None:  print("[latent-debug] enc_rna  keys:", _keys(enc_rna))
            if enc_atac is not None: print("[latent-debug] enc_atac keys:", _keys(enc_atac))

        # Extract fused first (from full)
        mu_fused = _extract_latent(enc_full, "fused")

        # Extract modality-specific from forced single-modality encodes.
        # If UniVI still returns generic keys, _extract_latent(...,"fused") fallback won’t help,
        # so we also try "fused" as last resort.
        mu_rna = None
        if enc_rna is not None:
            mu_rna = _extract_latent(enc_rna, "rna")
            if mu_rna is None:
                mu_rna = _extract_latent(enc_rna, "fused")

        mu_atac = None
        if enc_atac is not None:
            mu_atac = _extract_latent(enc_atac, "atac")
            if mu_atac is None:
                mu_atac = _extract_latent(enc_atac, "fused")

        if torch.is_tensor(mu_rna):   mu_rna_all.append(mu_rna.detach().cpu())
        if torch.is_tensor(mu_atac):  mu_atac_all.append(mu_atac.detach().cpu())
        if torch.is_tensor(mu_fused): mu_fused_all.append(mu_fused.detach().cpu())

        y_tensor, use_map = _encode_labels_for_gids(y_raw, gids_np, id_to_int=use_map)
        y_list.append(y_tensor.detach().cpu())
        gids_list.append(torch.as_tensor(gids_np, dtype=torch.long))

    def _cat(xs):
        return torch.cat(xs, dim=0) if len(xs) else None

    mu_rna_out   = _cat(mu_rna_all)
    mu_atac_out  = _cat(mu_atac_all)
    mu_fused_out = _cat(mu_fused_all)
    y_out        = _cat(y_list)
    gids_out     = _cat(gids_list)

    if require_mus and (mu_rna_out is None or mu_atac_out is None) and (mu_fused_out is None):
        raise RuntimeError(
            "Latent extraction failed: could not obtain modality-specific latents "
            "(mu_rna/mu_atac) nor a fused latent. Run with debug_first_batch=True."
        )

    return {
        "mu_rna": mu_rna_out,
        "mu_atac": mu_atac_out,
        "mu_fused": mu_fused_out,
        "y": y_out,
        "y_rna": y_out,
        "y_atac": y_out,
        "y_fused": y_out,
        "gids": gids_out,
        "id_to_int": use_map,
    }


# %% [markdown]
## 6) Figure 8 runner (missing-modality curve + label transfer)


# %% [code]
import numpy as np
import torch
from torch.utils.data import Dataset

# ---- knobs ----

def choose_batch_size(overlap: float, batch_size_default: int) -> int:
    p = float(overlap)
    bs0 = int(batch_size_default)
    if p >= 0.05:
        return bs0
    if bs0 >= 256:
        return 128
    if bs0 >= 128:
        return 64
    return bs0

def choose_unpaired_per_paired(overlap: float) -> float:
    """
    Heuristic: as overlap shrinks, keep relatively more paired anchors.
    Returns unpaired_per_paired in [0.25, 8.0].
    """
    p = max(float(overlap), 1e-6)
    val = 1.0 / np.sqrt(p / 0.10)     # p=0.10 -> 1, p=0.01 -> 3.16
    unpaired_per_paired = 1.0 / val   # invert to get fewer unpaired when p small
    return float(np.clip(unpaired_per_paired, 0.25, 8.0))

# ---- dataset wrappers ----

class IndexDataset(Dataset):
    """
    Wrap a base dataset + an index list and ALWAYS return an integer gid.

    Output format:
      - (x_dict, gid_int)                       if base yields dict
      - (x_dict, gid_int, *extras_from_base)    if base yields (dict, ...)
    """
    def __init__(self, base_dataset, indices):
        self.base = base_dataset
        self.indices = np.asarray(indices, dtype=np.int64)

    def __len__(self):
        return int(self.indices.shape[0])

    def __getitem__(self, i):
        gid = int(self.indices[int(i)])  # global integer index into base_dataset
        item = self.base[gid]

        if isinstance(item, dict):
            return dict(item), gid

        if isinstance(item, (tuple, list)) and len(item) >= 1 and isinstance(item[0], dict):
            x_dict = dict(item[0])
            extras = tuple(item[1:])
            return (x_dict, gid, *extras)

        raise TypeError(f"Base dataset must yield dict or (dict, ...), got: {type(item)}")

class DeterministicMaskDataset(Dataset):
    """
    Deterministically drops one modality for unpaired rows according to groups.
      groups==0: paired (keep all modalities)
      groups!=0: unpaired (drop `drop_modality` -> set to None)
    Assumes base yields (x_dict, gid, ...) after IndexDataset.
    """
    def __init__(self, base_dataset, *, groups, drop_modality: str):
        self.base = base_dataset
        self.groups = np.asarray(groups, dtype=np.int64)
        self.drop_modality = str(drop_modality)

        if len(self.base) != len(self.groups):
            raise ValueError(f"base_dataset len={len(self.base)} != groups len={len(self.groups)}")

    def __len__(self):
        return len(self.base)

    def __getitem__(self, i):
        item = self.base[i]

        if isinstance(item, dict):
            x_dict = dict(item)
            rest = ()
        elif isinstance(item, (tuple, list)) and len(item) >= 1 and isinstance(item[0], dict):
            x_dict = dict(item[0])
            rest = tuple(item[1:])
        else:
            raise TypeError(f"Expected dataset to yield dict or (dict, ...), got {type(item)}")

        g = int(self.groups[int(i)])
        if g != 0:
            x_dict[self.drop_modality] = None

        return (x_dict, *rest) if rest else x_dict

# ---- collate ----

def collate_xdict_with_idx(batch):
    """
    Collate items that are either:
      - x_dict
      - (x_dict, gid)
      - (x_dict, gid, ...)

    Produces:
      - out_xdict
      - gids (torch.int64) if present
      - extras (stacked tensors) if present
    """
    x_list = []
    rest_list = []

    for item in batch:
        if isinstance(item, dict):
            x_list.append(item)
            rest_list.append(())
        else:
            x_list.append(item[0])
            rest_list.append(tuple(item[1:]))

    # union of keys
    keys = set()
    for x in x_list:
        keys |= set(x.keys())

    out = {}
    for k in keys:
        vals = [x.get(k, None) for x in x_list]

        if all(v is None for v in vals):
            out[k] = None
            continue

        # If any None present, you are mixing paired+unpaired in same batch -> fail fast.
        if any(v is None for v in vals):
            raise RuntimeError(
                f"Within-batch mixed presence for modality '{k}'. "
                f"Sampler must keep paired/unpaired separate per-batch."
            )

        v0 = vals[0]
        if torch.is_tensor(v0):
            out[k] = torch.stack(vals, dim=0)
        else:
            out[k] = torch.as_tensor(np.stack(vals, axis=0))

    # no gids/extras
    if len(rest_list[0]) == 0:
        return out

    # first rest item is gid by IndexDataset
    gids = torch.as_tensor([int(r[0]) for r in rest_list], dtype=torch.int64)

    # if no extras, return (x, gids)
    if len(rest_list[0]) == 1:
        return out, gids

    # extras: stack each column if possible, else return as list
    extras_cols = list(zip(*[r[1:] for r in rest_list]))
    extras_out = []
    for col in extras_cols:
        try:
            extras_out.append(torch.as_tensor(col))
        except Exception:
            extras_out.append(list(col))

    return (out, gids, *extras_out)


# %% [code]
# ---- Plot UMAPs of latents (rna / atac / fused), colored by cell type ----
import os
import numpy as np
import matplotlib.pyplot as plt
from matplotlib.lines import Line2D

def _as_numpy_2d(x):
    import torch
    if x is None:
        return None
    if torch.is_tensor(x):
        x = x.detach().cpu().numpy()
    x = np.asarray(x)
    if x.ndim == 1:
        x = x[:, None]
    return x

def _get_label_strings(enc, y_all):
    """
    Prefer true string labels from y_all[gids] if gids is present.
    Falls back to integer y in enc.
    """
    gids = enc.get("gids", None)
    if gids is not None:
        gids = _as_numpy_2d(gids).reshape(-1).astype(np.int64)
        labs = np.asarray(y_all)[gids]
        return labs.astype(str)

    y = enc.get("y_fused", enc.get("y_rna", enc.get("y", None)))
    if y is None:
        return None
    return _as_numpy_2d(y).reshape(-1).astype(str)

def _umap_embed(X, *, seed=0, n_neighbors=15, min_dist=0.3, metric="euclidean"):
    """
    Returns 2D embedding via UMAP; falls back to PCA if umap-learn isn't installed.
    """
    X = _as_numpy_2d(X)
    if X is None:
        return None
    try:
        import umap
        reducer = umap.UMAP(
            n_neighbors=int(n_neighbors),
            min_dist=float(min_dist),
            metric=str(metric),
            random_state=int(seed),
        )
        return reducer.fit_transform(X)
    except Exception:
        from sklearn.decomposition import PCA
        return PCA(n_components=2, random_state=int(seed)).fit_transform(X)

def _prep_label_mapping(labels, *, sort_legend_by="count"):
    """
    Build ONE stable label->int mapping + ordered label list.
    This prevents colors from changing between panels.
    """
    if labels is None:
        return None, None, None, None

    labels = np.asarray(labels).astype(str)
    uniq, counts = np.unique(labels, return_counts=True)

    if sort_legend_by == "count":
        order = np.argsort(-counts)  # descending
    else:
        order = np.argsort(uniq.astype(str))

    uniq = uniq[order]
    counts = counts[order]

    lab_to_int = {lab: i for i, lab in enumerate(uniq.tolist())}
    c = np.array([lab_to_int[lab] for lab in labels], dtype=int)

    # Use a discrete cmap with fixed number of categories
    cmap = plt.get_cmap("tab20", max(len(uniq), 1))
    return labels, uniq, counts, (lab_to_int, c, cmap)

def _legend_handles(uniq, counts, cmap, *, legend_max=25):
    if uniq is None or counts is None:
        return None

    n_cat = len(uniq)
    if legend_max is None or legend_max <= 0 or n_cat == 0:
        return None

    top_n = int(min(legend_max, n_cat))
    handles = []
    for i in range(top_n):
        lab = uniq[i]
        col = cmap(i)
        handles.append(
            Line2D(
                [0], [0],
                marker="o",
                color="none",
                markerfacecolor=col,
                markeredgecolor="none",
                markersize=6,
                label=f"{lab} (n={counts[i]})",
            )
        )
    return handles

def plot_latent_umap(
    X,
    labels,
    *,
    title="UMAP",
    seed=0,
    n_neighbors=15,
    min_dist=0.3,
    point_size=5,
    alpha=0.8,
    legend_max=25,
    sort_legend_by="count",  # "count" or "name"
):
    X = _as_numpy_2d(X)
    if X is None:
        print(f"[skip] {title}: X is None")
        return

    emb = _umap_embed(X, seed=seed, n_neighbors=n_neighbors, min_dist=min_dist)
    if emb is None:
        print(f"[skip] {title}: embedding failed")
        return

    labels = None if labels is None else np.asarray(labels).astype(str)
    plt.figure(figsize=(12, 8))

    if labels is None:
        plt.scatter(emb[:, 0], emb[:, 1], s=point_size, alpha=alpha)
        plt.title(title)
        plt.xlabel("UMAP1"); plt.ylabel("UMAP2")
        plt.tight_layout()
        plt.show()
        return

    # Stable mapping for this plot call
    _, uniq, counts, packed = _prep_label_mapping(labels, sort_legend_by=sort_legend_by)
    _, c, cmap = packed

    plt.scatter(
        emb[:, 0], emb[:, 1],
        c=c,
        cmap=cmap,
        s=point_size,
        alpha=alpha,
        linewidths=0,
    )

    handles = _legend_handles(uniq, counts, cmap, legend_max=legend_max)
    if handles is not None:
        plt.legend(handles=handles, bbox_to_anchor=(1.02, 1), loc="upper left", frameon=False)

    plt.title(title)
    plt.xlabel("UMAP1"); plt.ylabel("UMAP2")
    plt.tight_layout()
    plt.show()

def plot_all_latent_umaps(enc, *, y_all, seed=0, n_neighbors=15, min_dist=0.3, point_size=5, alpha=0.8):
    labels = _get_label_strings(enc, y_all)

    plot_latent_umap(
        enc.get("mu_rna", None),
        labels,
        title="RNA latent (mu_rna)",
        seed=seed,
        n_neighbors=n_neighbors,
        min_dist=min_dist,
        point_size=point_size,
        alpha=alpha,
    )

    plot_latent_umap(
        enc.get("mu_atac", None),
        labels,
        title="ATAC latent (mu_atac)",
        seed=seed,
        n_neighbors=n_neighbors,
        min_dist=min_dist,
        point_size=point_size,
        alpha=alpha,
    )

    plot_latent_umap(
        enc.get("mu_fused", None),
        labels,
        title="Fused latent (mu_fused)",
        seed=seed,
        n_neighbors=n_neighbors,
        min_dist=min_dist,
        point_size=point_size,
        alpha=alpha,
    )

def save_umaps_for_enc(
    enc,
    *,
    y_all,
    out_png,
    seed=0,
    n_neighbors=15,
    min_dist=0.3,
    point_size=8,
    alpha=0.75,
    dpi=450,                 # NEW default: higher quality, but safe
    legend_max=25,            # NEW default legend size for saved figs
    sort_legend_by="count",   # keep consistent with interactive
):
    """
    Saves a single figure with 3 subplots (RNA/ATAC/Fused) colored by cell type.
    Uses ONE stable label->color mapping across all panels.
    """
    labels = _get_label_strings(enc, y_all)

    # precompute stable mapping ONCE for consistent colors across panels
    labs = None if labels is None else np.asarray(labels).astype(str)
    if labs is not None:
        _, uniq, counts, packed = _prep_label_mapping(labs, sort_legend_by=sort_legend_by)
        _, c_all, cmap = packed
        legend_handles = _legend_handles(uniq, counts, cmap, legend_max=legend_max)
    else:
        c_all, cmap, legend_handles = None, None, None

    fig = plt.figure(figsize=(18, 5), dpi=dpi)

    def _plot_into(ax, X, title):
        X = _as_numpy_2d(X)
        if X is None:
            ax.set_title(f"{title} (missing)")
            ax.axis("off")
            return

        emb = _umap_embed(X, seed=seed, n_neighbors=n_neighbors, min_dist=min_dist)
        if emb is None:
            ax.set_title(f"{title} (embed failed)")
            ax.axis("off")
            return

        if labs is None or c_all is None or cmap is None:
            ax.scatter(emb[:, 0], emb[:, 1], s=point_size, alpha=alpha, linewidths=0)
            ax.set_title(title)
            return

        ax.scatter(
            emb[:, 0], emb[:, 1],
            c=c_all,
            cmap=cmap,
            s=point_size,
            alpha=alpha,
            linewidths=0,
        )
        ax.set_title(title)

    ax1 = fig.add_subplot(1, 3, 1)
    ax2 = fig.add_subplot(1, 3, 2)
    ax3 = fig.add_subplot(1, 3, 3)

    _plot_into(ax1, enc.get("mu_rna", None),   "RNA latent (mu_rna)")
    _plot_into(ax2, enc.get("mu_atac", None),  "ATAC latent (mu_atac)")
    _plot_into(ax3, enc.get("mu_fused", None), "Fused latent (mu_fused)")

    for ax in (ax1, ax2, ax3):
        ax.set_xlabel("UMAP1")
        ax.set_ylabel("UMAP2")

    # put legend on the right, shared across panels
    if legend_handles is not None:
        fig.legend(
            handles=legend_handles,
            loc="center left",
            bbox_to_anchor=(1.01, 0.5),
            frameon=False,
            title=f"Cell types (top {min(int(legend_max), len(legend_handles))})",
        )

    fig.tight_layout()

    out_dir = os.path.dirname(out_png)
    if out_dir:
        os.makedirs(out_dir, exist_ok=True)

    fig.savefig(out_png, dpi=dpi, bbox_inches="tight")
    plt.close(fig)


def save_paired_overlay_umap(
    enc,
    *,
    y_all,
    out_png,
    seed=0,
    n_neighbors=15,
    min_dist=0.3,
    point_size=8,
    alpha=0.75,
    dpi=450,
    legend_max=25,
    sort_legend_by="count",
):
    """
    Saves a 1x2 figure where RNA and ATAC points are embedded TOGETHER in one UMAP:
      - Left: colored by cell type (stable mapping, shared legend)
      - Right: colored by modality (RNA vs ATAC)
    This is the most direct “are modalities aligned?” visual.

    Requires enc["mu_rna"] and enc["mu_atac"].
    Assumes rows are paired; the cell_type labels are duplicated for RNA and ATAC.
    """
    mu_rna = enc.get("mu_rna", None)
    mu_atac = enc.get("mu_atac", None)

    Xr = _as_numpy_2d(mu_rna)
    Xa = _as_numpy_2d(mu_atac)

    if Xr is None or Xa is None:
        # still save a stub figure so the pipeline doesn't break
        fig = plt.figure(figsize=(12, 5), dpi=dpi)
        ax = fig.add_subplot(1, 1, 1)
        ax.axis("off")
        ax.set_title("Paired overlay UMAP (missing mu_rna or mu_atac)")
        out_dir = os.path.dirname(out_png)
        if out_dir:
            os.makedirs(out_dir, exist_ok=True)
        fig.savefig(out_png, dpi=dpi, bbox_inches="tight")
        plt.close(fig)
        return

    if Xr.shape[0] != Xa.shape[0]:
        raise ValueError(f"mu_rna and mu_atac must have same n for paired overlay: {Xr.shape[0]} vs {Xa.shape[0]}")

    # labels from gids if present, otherwise fallback inside your helper
    labels = _get_label_strings(enc, y_all)
    labs = None if labels is None else np.asarray(labels).astype(str)

    # Concatenate data: [RNA; ATAC]
    X = np.concatenate([Xr, Xa], axis=0)

    # Duplicate labels for the two modalities
    if labs is not None:
        labs2 = np.concatenate([labs, labs], axis=0)
        _, uniq, counts, packed = _prep_label_mapping(labs2, sort_legend_by=sort_legend_by)
        _, c_all, cmap = packed
        legend_handles = _legend_handles(uniq, counts, cmap, legend_max=legend_max)
    else:
        labs2, c_all, cmap, legend_handles = None, None, None, None

    modality = np.array(["RNA"] * Xr.shape[0] + ["ATAC"] * Xa.shape[0], dtype=object)

    emb = _umap_embed(X, seed=seed, n_neighbors=n_neighbors, min_dist=min_dist)
    if emb is None:
        fig = plt.figure(figsize=(12, 5), dpi=dpi)
        ax = fig.add_subplot(1, 1, 1)
        ax.axis("off")
        ax.set_title("Paired overlay UMAP (embed failed)")
        out_dir = os.path.dirname(out_png)
        if out_dir:
            os.makedirs(out_dir, exist_ok=True)
        fig.savefig(out_png, dpi=dpi, bbox_inches="tight")
        plt.close(fig)
        return

    fig = plt.figure(figsize=(16, 6), dpi=dpi)
    ax1 = fig.add_subplot(1, 2, 1)
    ax2 = fig.add_subplot(1, 2, 2)

    # --- Panel 1: colored by cell type ---
    if labs2 is None or c_all is None or cmap is None:
        ax1.scatter(emb[:, 0], emb[:, 1], s=point_size, alpha=alpha, linewidths=0)
        ax1.set_title("Paired overlay (RNA+ATAC) — no labels")
    else:
        ax1.scatter(
            emb[:, 0], emb[:, 1],
            c=c_all,
            cmap=cmap,
            s=point_size,
            alpha=alpha,
            linewidths=0,
        )
        ax1.set_title("Paired overlay (RNA+ATAC) — colored by cell type")

    # --- Panel 2: colored by modality ---
    # fixed 2-category palette without relying on global styles
    mod_to_int = {"RNA": 0, "ATAC": 1}
    mod_c = np.array([mod_to_int[m] for m in modality], dtype=int)
    mod_cmap = plt.get_cmap("Set1", 2)

    ax2.scatter(
        emb[:, 0], emb[:, 1],
        c=mod_c,
        cmap=mod_cmap,
        s=point_size,
        alpha=alpha,
        linewidths=0,
    )
    ax2.set_title("Paired overlay (RNA+ATAC) — colored by modality")

    for ax in (ax1, ax2):
        ax.set_xlabel("UMAP1")
        ax.set_ylabel("UMAP2")

    # shared cell-type legend (panel 1)
    if legend_handles is not None:
        fig.legend(
            handles=legend_handles,
            loc="center left",
            bbox_to_anchor=(1.01, 0.5),
            frameon=False,
            title=f"Cell types (top {min(int(legend_max), len(legend_handles))})",
        )

    # modality legend on ax2
    mod_handles = [
        Line2D([0], [0], marker="o", color="none", markerfacecolor=mod_cmap(0), markersize=6, label="RNA"),
        Line2D([0], [0], marker="o", color="none", markerfacecolor=mod_cmap(1), markersize=6, label="ATAC"),
    ]
    ax2.legend(handles=mod_handles, loc="upper right", frameon=False)

    fig.tight_layout()

    out_dir = os.path.dirname(out_png)
    if out_dir:
        os.makedirs(out_dir, exist_ok=True)

    fig.savefig(out_png, dpi=dpi, bbox_inches="tight")
    plt.close(fig)


# %% [code]
import pandas as pd
import numpy as np
import torch

import os

from sklearn.cluster import KMeans
from sklearn.metrics import (
    adjusted_rand_score,
    normalized_mutual_info_score,
    silhouette_score,
    calinski_harabasz_score,
    davies_bouldin_score,
)

# -----------------------------
# alignment metrics: FOSCTTM + Recall@K (paired rows)
# -----------------------------

def _as_numpy_2d(x):
    if x is None:
        return None
    if torch.is_tensor(x):
        x = x.detach().cpu().numpy()
    x = np.asarray(x)
    if x.ndim == 1:
        x = x[:, None]
    return x.astype(np.float32, copy=False)

def _as_numpy_1d(y):
    if y is None:
        return None
    if torch.is_tensor(y):
        y = y.detach().cpu().numpy()
    y = np.asarray(y)
    return y.reshape(-1)

def _pairwise_sq_dists(X, Y):
    X = _as_numpy_2d(X)
    Y = _as_numpy_2d(Y)
    x2 = np.sum(X * X, axis=1, keepdims=True)
    y2 = np.sum(Y * Y, axis=1, keepdims=True).T
    D = x2 + y2 - 2.0 * (X @ Y.T)
    np.maximum(D, 0.0, out=D)
    return D

def foscttm(X, Y, *, symmetric=True):
    """
    FOSCTTM (Fraction of Samples Closer Than the True Match).
    Assumes paired rows: X[i] matches Y[i]. Lower is better.
    If symmetric=True, averages X->Y and Y->X.
    """
    D = _pairwise_sq_dists(X, Y)
    n = D.shape[0]
    diag = np.diag(D)

    denom = (n - 1) if n > 1 else 1
    frac_xy = (D < diag[:, None]).sum(axis=1) / denom

    if not symmetric:
        return float(frac_xy.mean())

    frac_yx = (D < diag[None, :]).sum(axis=0) / denom
    return float(0.5 * (frac_xy.mean() + frac_yx.mean()))

def recall_at_k(X, Y, *, ks=(1, 10, 25, 50, 100), symmetric=True):
    """
    Recall@K for paired matching. Hit if true match is within top-K nearest.
    If symmetric=True, averages X->Y and Y->X.
    """
    D = _pairwise_sq_dists(X, Y)
    n = D.shape[0]
    ks = [int(k) for k in ks if int(k) >= 1]

    order_xy = np.argsort(D, axis=1)
    pos_xy = np.empty(n, dtype=np.int32)
    for i in range(n):
        pos_xy[i] = np.where(order_xy[i] == i)[0][0]

    if symmetric:
        order_yx = np.argsort(D.T, axis=1)
        pos_yx = np.empty(n, dtype=np.int32)
        for i in range(n):
            pos_yx[i] = np.where(order_yx[i] == i)[0][0]

    out = {}
    for k in ks:
        k_eff = min(k, n)
        r_xy = float((pos_xy < k_eff).mean())
        if not symmetric:
            out[k] = r_xy
        else:
            r_yx = float((pos_yx < k_eff).mean())
            out[k] = 0.5 * (r_xy + r_yx)
    return out

from sklearn.neighbors import KNeighborsClassifier
from sklearn.metrics import accuracy_score, f1_score

def label_transfer_knn(X_src, y_src, X_tgt, y_tgt, *, k: int = 5):
    """
    Train kNN on (X_src, y_src) and predict labels on X_tgt.
    Returns {"acc": ..., "macro_f1": ...}.

    Assumes y_* are already integer-coded (recommended).
    """
    X_src = _as_numpy_2d(X_src)
    X_tgt = _as_numpy_2d(X_tgt)
    y_src = _as_numpy_1d(y_src)
    y_tgt = _as_numpy_1d(y_tgt)

    if X_src is None or X_tgt is None or y_src is None or y_tgt is None:
        return {"acc": np.nan, "macro_f1": np.nan}

    if X_src.shape[0] != y_src.shape[0] or X_tgt.shape[0] != y_tgt.shape[0]:
        return {"acc": np.nan, "macro_f1": np.nan}

    if X_src.shape[0] < 2 or X_tgt.shape[0] < 2:
        return {"acc": np.nan, "macro_f1": np.nan}

    # k must be <= n_src
    k_eff = int(min(max(1, k), X_src.shape[0]))

    clf = KNeighborsClassifier(n_neighbors=k_eff, weights="distance", metric="euclidean")
    clf.fit(X_src, y_src)
    y_pred = clf.predict(X_tgt)

    return {
        "acc": float(accuracy_score(y_tgt, y_pred)),
        "macro_f1": float(f1_score(y_tgt, y_pred, average="macro")),
    }
    
# -----------------------------
# small helpers
# -----------------------------

def _get_model_and_training_info(train_result):
    """Accept either model or TrainResult-like object."""
    model = getattr(train_result, "model", train_result)
    info = {}
    for k in ("warmup_epochs", "best_epoch_all","best_val_all", "best_epoch_postwarmup","best_val_postwarmup"):
        if hasattr(train_result, k):
            v = getattr(train_result, k)
            if v is not None:
                info[k] = int(v) if ("epoch" in k or "warmup" in k) else float(v)
    return model, info

def _compute_fused_latent(enc, *, fuse_mode="avg"):
    """
    Return fused latent for clustering metrics on paired samples.
    If mu_fused exists, use it; else fuse rna/atac.
    """
    if enc is None:
        return None
    if enc.get("mu_fused", None) is not None:
        return enc["mu_fused"]

    mu_rna = enc.get("mu_rna", None)
    mu_atac = enc.get("mu_atac", None)
    if mu_rna is None or mu_atac is None:
        return None

    if fuse_mode == "concat":
        if torch.is_tensor(mu_rna) and torch.is_tensor(mu_atac):
            return torch.cat([mu_rna, mu_atac], dim=1)
        return np.concatenate([_as_numpy_2d(mu_rna), _as_numpy_2d(mu_atac)], axis=1)

    # default avg
    if torch.is_tensor(mu_rna) and torch.is_tensor(mu_atac):
        return 0.5 * (mu_rna + mu_atac)
    return 0.5 * (_as_numpy_2d(mu_rna) + _as_numpy_2d(mu_atac))

def _filter_none_labels(X, y):
    X = _as_numpy_2d(X)
    y = _as_numpy_1d(y)
    if X is None or y is None:
        return None, None
    mask = np.array([v is not None for v in y], dtype=bool)
    if mask.sum() < 2:
        return None, None
    return X[mask], y[mask]

def _safe_silhouette(X, labels):
    """
    Strict silhouette: returns NaN unless all clusters have >=2 samples and >=2 clusters exist.
    """
    labels = _as_numpy_1d(labels)
    X = _as_numpy_2d(X)
    if X is None or labels is None:
        return np.nan

    uniq, counts = np.unique(labels, return_counts=True)
    if len(uniq) < 2 or np.any(counts < 2):
        return np.nan

    try:
        return float(silhouette_score(X, labels))
    except Exception:
        return np.nan

def _safe_silhouette_drop_small_classes(X, labels, *, min_per_class: int = 2, min_total: int = 10):
    """
    More forgiving silhouette for y_true:
      - drops labels that have < min_per_class samples
      - returns (silhouette, n_used, frac_used)
    """
    labels = _as_numpy_1d(labels)
    X = _as_numpy_2d(X)
    if X is None or labels is None:
        return np.nan, 0, 0.0

    uniq, counts = np.unique(labels, return_counts=True)
    keep_labs = set(uniq[counts >= int(min_per_class)].tolist())
    if len(keep_labs) < 2:
        return np.nan, 0, 0.0

    keep = np.array([lab in keep_labs for lab in labels], dtype=bool)
    n_used = int(keep.sum())
    if n_used < int(min_total):
        return np.nan, n_used, float(n_used / max(len(labels), 1))

    Xk = X[keep]
    yk = labels[keep]

    # still need >=2 samples per remaining class (should hold, but double-check)
    uniq2, counts2 = np.unique(yk, return_counts=True)
    if len(uniq2) < 2 or np.any(counts2 < 2):
        return np.nan, n_used, float(n_used / max(len(labels), 1))

    try:
        sil = float(silhouette_score(Xk, yk))
    except Exception:
        sil = np.nan

    return sil, n_used, float(n_used / max(len(labels), 1))

def _cluster_metrics_on_latent(X, y_true, *, n_clusters=None, kmeans_seed=0):
    """
    Clustering & separation metrics on latent.
    - KMeans metrics use all samples (if k>=2).
    - SIL_true is computed robustly by dropping tiny classes (otherwise it's often NaN on splits).
    """
    X = _as_numpy_2d(X)
    y_true = _as_numpy_1d(y_true)

    if X is None or y_true is None:
        return {}

    X, y_true = _filter_none_labels(X, y_true)
    if X is None or y_true is None:
        return {
            "n_labels": 0, "kmeans_k": 0,
            "ARI": np.nan, "NMI": np.nan,
            "SIL_kmeans": np.nan, "SIL_true": np.nan,
            "SIL_true_n": 0, "SIL_true_frac": 0.0,
            "CH_kmeans": np.nan, "DB_kmeans": np.nan,
        }

    uniq = np.unique(y_true)
    k = int(n_clusters) if n_clusters is not None else int(len(uniq))

    # SIL_true: robust version (drop singleton labels)
    sil_true, sil_true_n, sil_true_frac = _safe_silhouette_drop_small_classes(X, y_true)

    if k < 2:
        return {
            "n_labels": int(len(uniq)),
            "kmeans_k": int(k),
            "ARI": np.nan,
            "NMI": np.nan,
            "SIL_kmeans": np.nan,
            "SIL_true": float(sil_true),
            "SIL_true_n": int(sil_true_n),
            "SIL_true_frac": float(sil_true_frac),
            "CH_kmeans": np.nan,
            "DB_kmeans": np.nan,
        }

    km = KMeans(n_clusters=k, n_init=20, random_state=int(kmeans_seed))
    y_pred = km.fit_predict(X)

    out = {
        "n_labels": int(len(uniq)),
        "kmeans_k": int(k),
        "ARI": float(adjusted_rand_score(y_true, y_pred)),
        "NMI": float(normalized_mutual_info_score(y_true, y_pred)),
        "SIL_kmeans": _safe_silhouette(X, y_pred),
        "SIL_true": float(sil_true),
        "SIL_true_n": int(sil_true_n),
        "SIL_true_frac": float(sil_true_frac),
    }
    try:
        out["CH_kmeans"] = float(calinski_harabasz_score(X, y_pred))
    except Exception:
        out["CH_kmeans"] = np.nan
    try:
        out["DB_kmeans"] = float(davies_bouldin_score(X, y_pred))
    except Exception:
        out["DB_kmeans"] = np.nan

    return out

def first_not_none(*vals):
    for v in vals:
        if v is not None:
            return v
    return None

def _lt_or_nans(X_src, y_src, X_tgt, y_tgt, *, k):
    """
    Safe wrapper around label_transfer_knn.
    Returns dict with acc/macro_f1 always present.
    """
    X_src = _as_numpy_2d(X_src)
    X_tgt = _as_numpy_2d(X_tgt)
    y_src = _as_numpy_1d(y_src)
    y_tgt = _as_numpy_1d(y_tgt)

    if X_src is None or X_tgt is None or y_src is None or y_tgt is None:
        return {"acc": np.nan, "macro_f1": np.nan}
    if X_src.shape[0] != y_src.shape[0] or X_tgt.shape[0] != y_tgt.shape[0]:
        return {"acc": np.nan, "macro_f1": np.nan}
    if X_src.shape[0] < 2 or X_tgt.shape[0] < 2:
        return {"acc": np.nan, "macro_f1": np.nan}
    if len(np.unique(y_src)) < 2:
        return {"acc": np.nan, "macro_f1": np.nan}

    kk = int(min(k, max(1, X_src.shape[0] - 1)))
    try:
        out = label_transfer_knn(X_src, y_src, X_tgt, y_tgt, k=kk)
        return {"acc": float(out.get("acc", np.nan)), "macro_f1": float(out.get("macro_f1", np.nan))}
    except Exception:
        return {"acc": np.nan, "macro_f1": np.nan}

# -----------------------------
# main runner
# -----------------------------

def run_fig8_missing_modality_curve_v1(
    overlap_grid=(1.0, 0.975, 0.95, 0.925, 0.9, 0.85, 0.8, 0.7, 0.6, 0.5, 0.4, 0.3, 0.2, 0.1, 0.09, 0.08, 0.07, 0.06, 0.05, 0.04, 0.03, 0.025, 0.02, 0.0175, 0.0150, 0.0125, 0.01, 0.005, 0.001),
    *,
    seed: int = 42,
    drop_modality: str = "atac",
    k_knn: int = 3,
    batch_size: int = 256,
    v1_recon: str = "avg",
    adaptive_batch_size: bool = True,
    oversample_paired: bool = True,
    fuse_mode: str = "avg",
    n_clusters: int | None = None,
    kmeans_seed: int = 0,
    warmup_epochs: int = 50,
    patience: int = 100,
    min_delta: float = 0.0,
    allow_unpaired_test: bool = True,

    # --- NEW: UMAP saving controls ---
    umap_out_dir: str | None = None,
    umap_use: str = "test",                 # "test" or "val"
    save_umaps_every: int = 1,
    save_umaps_if_overlap_leq: float | None = None,  # e.g. 0.1 to only save tail
    umap_seed: int | None = None,
    umap_n_neighbors: int = 15,
    umap_min_dist: float = 0.3,
    umap_point_size: int = 6,
    umap_alpha: float = 0.75,
    umap_dpi: int = 450,
    umap_legend_max: int = 25,
    umap_sort_legend_by: str = "count",
):
    rows = []

    if umap_seed is None:
        umap_seed = int(seed)

    for i, overlap in enumerate(overlap_grid):
        p = float(overlap)
        bs_use = choose_batch_size(p, batch_size) if adaptive_batch_size else int(batch_size)
        print(f"[overlap={p:g}] batch_size={bs_use}")

        train_loader, val_loader, test_loader, comp = make_loaders_with_overlap_v1(
            dataset, train_idx, val_idx, test_idx,
            overlap_fraction=p,
            drop_modality=str(drop_modality),
            seed=int(seed),
            batch_size=int(bs_use),
            oversample_paired=bool(oversample_paired),
        )

        train_result = train_one_overlap_v1(
            train_loader=train_loader,
            val_loader=val_loader,
            seed=int(seed),
            loss_mode="v1",
            v1_recon=str(v1_recon),
            warmup_epochs=int(warmup_epochs),
            patience=int(patience),
            min_delta=float(min_delta),
        )

        model, train_info = _get_model_and_training_info(train_result)

        # --- encode VAL first (stable mapping), then TEST ---
        enc_val = encode_embeddings_with_labels(
            model, val_loader, device=device, y_all=y_all, id_to_int=None
        )
        enc_test = encode_embeddings_with_labels(
            model, test_loader, device=device, y_all=y_all, id_to_int=enc_val.get("id_to_int", None)
        )

        # --- NEW: save UMAPs ---
        if umap_out_dir is not None:
            do_every = (int(save_umaps_every) <= 1) or (i % int(save_umaps_every) == 0)
            do_thresh = (save_umaps_if_overlap_leq is None) or (p <= float(save_umaps_if_overlap_leq))
            if do_every and do_thresh:
                enc_plot = enc_test if str(umap_use).lower() == "test" else enc_val

                tag = f"drop-{drop_modality}__overlap-{p:.6g}__bs-{int(bs_use)}"
                out1 = os.path.join(umap_out_dir, f"{tag}__single_latents.png")
                out2 = os.path.join(umap_out_dir, f"{tag}__paired_overlay.png")

                # 3-panel (RNA/ATAC/Fused) colored by cell type
                save_umaps_for_enc(
                    enc_plot,
                    y_all=y_all,
                    out_png=out1,
                    seed=int(umap_seed),
                    n_neighbors=int(umap_n_neighbors),
                    min_dist=float(umap_min_dist),
                    point_size=int(umap_point_size),
                    alpha=float(umap_alpha),
                    dpi=int(umap_dpi),
                    legend_max=int(umap_legend_max),
                    sort_legend_by=str(umap_sort_legend_by),
                )

                # overlay (RNA+ATAC embedded together) colored by cell type + modality
                save_paired_overlay_umap(
                    enc_plot,
                    y_all=y_all,
                    out_png=out2,
                    seed=int(umap_seed),
                    n_neighbors=int(umap_n_neighbors),
                    min_dist=float(umap_min_dist),
                    point_size=int(umap_point_size),
                    alpha=float(umap_alpha),
                    dpi=int(umap_dpi),
                    legend_max=int(umap_legend_max),
                    sort_legend_by=str(umap_sort_legend_by),
                )

        mu_rna_test  = enc_test.get("mu_rna", None)
        mu_atac_test = enc_test.get("mu_atac", None)

        y_test = first_not_none(enc_test.get("y_fused"), enc_test.get("y_rna"), enc_test.get("y_atac"))
        y_test_np = _as_numpy_1d(y_test)

        row = {
            "overlap_fraction": p,
            "batch_size": int(bs_use),
            "drop_modality": str(drop_modality),
            "v1_recon": str(v1_recon),
            "fuse_mode": str(fuse_mode),
            **comp,
            **train_info,
        }

        # --- alignment metrics on paired test ---
        if mu_rna_test is None or mu_atac_test is None:
            if not allow_unpaired_test:
                raise RuntimeError("Test loader should be paired; missing mu_rna or mu_atac.")
            row["FOSCTTM"] = np.nan
            for kk in (1, 10, 25, 50, 100):
                row[f"Recall@{kk}"] = np.nan
        else:
            X_rna  = _as_numpy_2d(mu_rna_test)
            X_atac = _as_numpy_2d(mu_atac_test)
            row["FOSCTTM"] = float(foscttm(X_rna, X_atac))
            recs = recall_at_k(X_rna, X_atac, ks=(1, 10, 25, 50, 100))
            for kk, v in recs.items():
                row[f"Recall@{kk}"] = float(v)

        # --- label transfer: VAL -> TEST ---
        lt_r2a = _lt_or_nans(
            enc_val.get("mu_rna"),  enc_val.get("y_rna"),
            enc_test.get("mu_atac"), enc_test.get("y_atac"),
            k=int(k_knn),
        )
        lt_a2r = _lt_or_nans(
            enc_val.get("mu_atac"), enc_val.get("y_atac"),
            enc_test.get("mu_rna"),  enc_test.get("y_rna"),
            k=int(k_knn),
        )
        row["LT_RNA2ATAC_acc"] = float(lt_r2a["acc"])
        row["LT_RNA2ATAC_macroF1"] = float(lt_r2a["macro_f1"])
        row["LT_ATAC2RNA_acc"] = float(lt_a2r["acc"])
        row["LT_ATAC2RNA_macroF1"] = float(lt_a2r["macro_f1"])

        # --- clustering/separation metrics on fused latent (test) ---
        mu_fused = _compute_fused_latent(enc_test, fuse_mode=str(fuse_mode))
        row.update(_cluster_metrics_on_latent(mu_fused, y_test_np, n_clusters=n_clusters, kmeans_seed=kmeans_seed))

        rows.append(row)

    return pd.DataFrame(rows)


# %% [code]
df_drop_atac_fig8 = run_fig8_missing_modality_curve_v1(
    drop_modality="atac",
    warmup_epochs=0,
    fuse_mode="moe",
    umap_out_dir="./ablation_umaps_drop_atac",
    #save_umaps_if_overlap_leq=0.10,   # only save when <= 10% paired
    save_umaps_if_overlap_leq=1.00,
    save_umaps_every=1,
    umap_point_size=5,
    umap_alpha=0.75,
)


# %% [code]
#df_drop_atac_fig8 = run_fig8_missing_modality_curve_v1(drop_modality="atac", warmup_epochs=0, fuse_mode="avg")


# %% [code]
df_drop_atac_fig8


# %% [code]
df_drop_rna_fig8 = run_fig8_missing_modality_curve_v1(
    drop_modality="rna",
    warmup_epochs=0,
    fuse_mode="moe",
    umap_out_dir="./results/figure_9_paired_ablation_outputs-2-10-2026/ablation_umaps_drop_rna",
    #save_umaps_if_overlap_leq=0.10,   # only save when <= 10% paired
    save_umaps_if_overlap_leq=1.00,
    save_umaps_every=1,
    umap_point_size=8,
    umap_alpha=0.75,
)


# %% [code]
#df_drop_rna_fig8  = run_fig8_missing_modality_curve_v1(drop_modality="rna", warmup_epochs=0, fuse_mode="avg")


# %% [code]
df_drop_rna_fig8


# %% [code]
# save to a TSV in the current working directory
df_drop_rna_fig8.to_csv("./results/figure_9_paired_ablation_outputs-2-10-2026/df_paired_ablation_rna.tsv", sep="\t", index=False)

# (optional) also save a gzipped TSV
#df_drop_rna_fig8.to_csv("df_drop_rna_fig8.tsv.gz", sep="\t", index=False, compression="gzip")


# %% [code]
# save to a TSV in the current working directory
df_drop_atac_fig8.to_csv("./results/figure_9_paired_ablation_outputs-2-10-2026/df_paired_ablation_atac.tsv", sep="\t", index=False)

# (optional) also save a gzipped TSV
#df_drop_atac_fig8.to_csv("df_drop_rna_fig8.tsv.gz", sep="\t", index=False, compression="gzip")


# %% [markdown]
## 7) Plotting


# %% [code]
import matplotlib as mpl
import matplotlib.pyplot as plt

mpl.rcParams.update({
    # Figure size (inches)
    "figure.figsize": (12, 10),

    # DPI: displayed in notebooks vs saved files
    "figure.dpi": 300,      # notebook display
    "savefig.dpi": 300,     # saved output

    # Fonts
    "font.size": 11,
    "axes.titlesize": 12,
    "axes.labelsize": 11,
    "xtick.labelsize": 10,
    "ytick.labelsize": 10,
    "legend.fontsize": 10,

    # Lines/markers
    "lines.linewidth": 1.6,
    "lines.markersize": 5,

    # Layout + export
    "figure.autolayout": False,     # if True, similar to tight_layout each time
    "savefig.bbox": "tight",
    "savefig.pad_inches": 0.02,
})

#mpl.rcdefaults()


# %% [code]
import matplotlib.pyplot as plt

def plot_fig8(df: pd.DataFrame, *, save_prefix: str | None = None):
    df = df.sort_values("overlap_fraction")

    plt.figure()
    plt.plot(df["overlap_fraction"], df["FOSCTTM"], marker="o")
    plt.gca().invert_xaxis()
    plt.xlabel("Paired overlap fraction in training")
    plt.ylabel("FOSCTTM (lower better)")
    plt.title("Figure 8A: alignment vs missing modality")
    if save_prefix:
        plt.savefig(f"{save_prefix}_foscttm.png", dpi=300, bbox_inches="tight")
    plt.show()

    plt.figure()
    plt.plot(df["overlap_fraction"], df["LT_RNA2ATAC_acc"], marker="o", label="RNA→ATAC acc")
    plt.plot(df["overlap_fraction"], df["LT_ATAC2RNA_acc"], marker="o", label="ATAC→RNA acc")
    plt.gca().invert_xaxis()
    plt.xlabel("Paired overlap fraction in training")
    plt.ylabel("Label transfer accuracy")
    plt.title("Figure 8B: label transfer vs missing modality")
    plt.legend()
    if save_prefix:
        plt.savefig(f"{save_prefix}_label_transfer.png", dpi=300, bbox_inches="tight")
    plt.show()


# %% [code]
plot_fig8(df_drop_atac_fig8)


# %% [code]
plot_fig8(df_drop_rna_fig8)


# %% [code]
def plot_fig8_recall(df: pd.DataFrame, ks=(1,10,25,50,100), save_prefix=None):
    df = df.sort_values("overlap_fraction")
    plt.figure()
    for k in ks:
        col = f"Recall@{k}"
        if col in df.columns:
            plt.plot(df["overlap_fraction"], df[col], marker="o", label=col)
    plt.gca().invert_xaxis()
    plt.xlabel("Paired overlap fraction in training")
    plt.ylabel("Recall@K (higher better)")
    plt.title("Figure 8: cross-modal retrieval vs missing modality")
    plt.legend()
    if save_prefix:
        plt.savefig(f"{save_prefix}_recall.png", dpi=300, bbox_inches="tight")
    plt.show()


# %% [code]
plot_fig8_recall(df_drop_atac_fig8)
#plot_fig8_recall(df_fig8, save_prefix="./figures/fig8")


# %% [code]
plot_fig8_recall(df_drop_rna_fig8)


# %% [code]
def plot_fig8_labeltransfer_f1(df: pd.DataFrame, save_prefix=None):
    df = df.sort_values("overlap_fraction")
    plt.figure()
    if "LT_RNA2ATAC_macroF1" in df.columns:
        plt.plot(df["overlap_fraction"], df["LT_RNA2ATAC_macroF1"], marker="o", label="RNA→ATAC macroF1")
    if "LT_ATAC2RNA_macroF1" in df.columns:
        plt.plot(df["overlap_fraction"], df["LT_ATAC2RNA_macroF1"], marker="o", label="ATAC→RNA macroF1")
    plt.gca().invert_xaxis()
    plt.xlabel("Paired overlap fraction in training")
    plt.ylabel("Label transfer macroF1")
    plt.title("Figure 8: label transfer macroF1 vs missing modality")
    plt.legend()
    if save_prefix:
        plt.savefig(f"{save_prefix}_label_transfer_macroF1.png", dpi=300, bbox_inches="tight")
    plt.show()


# %% [code]
plot_fig8_labeltransfer_f1(df_drop_atac_fig8)


# %% [code]
plot_fig8_labeltransfer_f1(df_drop_rna_fig8)


# %% [code]
def plot_with_band(df_all, ycol, title, savepath=None):
    g = df_all.groupby("overlap_fraction")[ycol]
    x = np.array(sorted(df_all["overlap_fraction"].unique()))
    mean = np.array([g.get_group(v).mean() for v in x])
    std  = np.array([g.get_group(v).std(ddof=1) for v in x])

    plt.figure()
    plt.plot(x, mean, marker="o")
    plt.fill_between(x, mean-std, mean+std, alpha=0.2)
    plt.gca().invert_xaxis()
    plt.xlabel("Paired overlap fraction in training")
    plt.ylabel(ycol)
    plt.title(title)
    if savepath:
        plt.savefig(savepath, dpi=300, bbox_inches="tight")
    plt.show()


# %% [code]
#plot_with_band(df_all, "FOSCTTM", "Fig 8A: FOSCTTM vs overlap (mean±sd)", "./figures/fig8_foscttm_band.png")
#plot_with_band(df_all, "LT_RNA2ATAC_acc", "Fig 8B: RNA→ATAC acc vs overlap (mean±sd)", "./figures/fig8_lt_r2a_band.png")


# %% [code]
import os
import numpy as np
import pandas as pd
import matplotlib.pyplot as plt

# -----------------------------
# Band plot helpers
# -----------------------------

def _ensure_dir(path: str | None):
    if path is None:
        return
    d = os.path.dirname(path)
    if d:
        os.makedirs(d, exist_ok=True)

def _infer_metric_cols(df: pd.DataFrame, *, x_col="overlap_fraction", exclude=()):
    """
    Heuristic: numeric cols excluding x_col and other known metadata.
    """
    exclude = set(exclude) | {x_col}
    num_cols = [c for c in df.columns if pd.api.types.is_numeric_dtype(df[c])]
    # drop obvious bookkeeping columns if present
    drop_like = {
        "seed", "fold", "run", "repeat",
        "batch_size",
        "warmup_epochs", "best_epoch_all", "best_val_all",
        "best_epoch_postwarmup", "best_val_postwarmup",
        "n_labels", "kmeans_k", "SIL_true_n"
    }
    exclude |= drop_like.intersection(num_cols)
    return [c for c in num_cols if c not in exclude]

def summarize_for_band(
    df: pd.DataFrame,
    metric: str,
    *,
    x_col: str = "overlap_fraction",
    group_cols: tuple[str, ...] = (),
    agg: str = "sd",          # "sd" | "sem" | "quantile"
    q: tuple[float, float] = (0.25, 0.75),
):
    """
    Returns a tidy summary DataFrame with columns:
      [*group_cols, x_col, mean, lo, hi, n]
    where lo/hi are the band bounds.
    """
    cols = [x_col, metric, *group_cols]
    d = df.loc[:, [c for c in cols if c in df.columns]].copy()
    d = d.dropna(subset=[x_col, metric])

    if d.empty:
        return pd.DataFrame(columns=[*group_cols, x_col, "mean", "lo", "hi", "n"])

    gb_keys = [*group_cols, x_col] if group_cols else [x_col]
    g = d.groupby(gb_keys, dropna=False)[metric]

    if agg == "quantile":
        lo_q, hi_q = q
        out = g.agg(
            mean="mean",
            lo=lambda s: float(s.quantile(lo_q)),
            hi=lambda s: float(s.quantile(hi_q)),
            n="count",
        ).reset_index()
    else:
        out = g.agg(mean="mean", sd="std", n="count").reset_index()
        out["sd"] = out["sd"].fillna(0.0)

        if agg == "sem":
            out["err"] = out["sd"] / np.sqrt(np.maximum(out["n"].to_numpy(), 1))
        elif agg == "sd":
            out["err"] = out["sd"]
        else:
            raise ValueError(f"agg must be one of ['sd','sem','quantile'], got {agg!r}")

        out["lo"] = out["mean"] - out["err"]
        out["hi"] = out["mean"] + out["err"]
        out = out.drop(columns=["sd", "err"])

    # sort nicely for plotting
    out = out.sort_values([*group_cols, x_col] if group_cols else [x_col]).reset_index(drop=True)
    return out

def plot_with_band(
    df: pd.DataFrame,
    metric: str,
    title: str,
    out_png: str | None = None,
    *,
    x_col: str = "overlap_fraction",
    group_cols: tuple[str, ...] = (),
    agg: str = "sd",                # "sd" | "sem" | "quantile"
    q: tuple[float, float] = (0.25, 0.75),
    xlabel: str | None = None,
    ylabel: str | None = None,
    logx: bool = False,
    invert_x: bool = True,          # overlap often decreases left->right; set False if you don’t want that
    ylim: tuple[float, float] | None = None,
    figsize=(12, 10),
    dpi: int = 300,
):
    """
    Single-metric band plot. If group_cols given, draws one line per group.
    """
    summ = summarize_for_band(df, metric, x_col=x_col, group_cols=group_cols, agg=agg, q=q)
    if summ.empty:
        raise ValueError(f"No data to plot for metric {metric!r} (after dropping NaNs).")

    fig = plt.figure(figsize=figsize)
    ax = plt.gca()

    if group_cols:
        # Each unique group gets a line
        group_df = summ.groupby(list(group_cols), dropna=False)
        for gkey, sub in group_df:
            label = " | ".join([f"{c}={v}" for c, v in zip(group_cols, (gkey if isinstance(gkey, tuple) else (gkey,)))])
            x = sub[x_col].to_numpy()
            m = sub["mean"].to_numpy()
            lo = sub["lo"].to_numpy()
            hi = sub["hi"].to_numpy()

            ax.plot(x, m, label=label)
            ax.fill_between(x, lo, hi, alpha=0.2)
    else:
        x = summ[x_col].to_numpy()
        m = summ["mean"].to_numpy()
        lo = summ["lo"].to_numpy()
        hi = summ["hi"].to_numpy()

        ax.plot(x, m)
        ax.fill_between(x, lo, hi, alpha=0.2)

    ax.set_title(title)
    ax.set_xlabel(xlabel or x_col)
    ax.set_ylabel(ylabel or metric)

    if logx:
        ax.set_xscale("log")

    if invert_x:
        ax.invert_xaxis()

    if ylim is not None:
        ax.set_ylim(*ylim)

    ax.grid(True, alpha=0.25)
    if group_cols:
        ax.legend(frameon=False, fontsize=8)

    plt.tight_layout()

    if out_png is not None:
        _ensure_dir(out_png)
        plt.savefig(out_png, dpi=dpi)
        plt.close(fig)
    else:
        plt.show()

    return summ  # handy if you want to inspect the aggregated numbers

def plot_many_metrics_with_band(
    df: pd.DataFrame,
    metrics: list[str] | None = None,
    out_dir: str | None = "./figures/fig8_bands",
    *,
    x_col: str = "overlap_fraction",
    group_cols: tuple[str, ...] = (),
    agg: str = "sd",
    q: tuple[float, float] = (0.25, 0.75),
    invert_x: bool = True,
    title_prefix: str = "",
    exclude_cols=(),
):
    """
    If out_dir is None: show plots in notebook (no saving).
    Else: save one PNG per metric into out_dir.
    """
    if metrics is None:
        metrics = _infer_metric_cols(df, x_col=x_col, exclude=exclude_cols)

    if out_dir is not None:
        os.makedirs(out_dir, exist_ok=True)

    summaries = {}
    for met in metrics:
        out_png = None if out_dir is None else os.path.join(out_dir, f"{met}__band.png")
        title = f"{title_prefix}{met} vs {x_col} ({agg})"
        summ = plot_with_band(
            df, met, title, out_png,
            x_col=x_col, group_cols=group_cols, agg=agg, q=q,
            invert_x=invert_x,
        )
        summaries[met] = summ

    return summaries


# %% [code]
# Your two explicit calls (mean ± sd over repeats)
'''
plot_with_band(
    df_all, "FOSCTTM",
    "Fig 8A: FOSCTTM vs overlap (mean±sd)",
    out_png=None,
    agg="sd",
)

plot_with_band(
    df_all, "LT_RNA2ATAC_acc",
    "Fig 8B: RNA→ATAC acc vs overlap (mean±sd)",
    out_png=None,
    agg="sd",
)
'''

# %% [code]
# One PNG per metric, autodetected from numeric columns
'''
plot_many_metrics_with_band(
    df_all,
    #out_dir="./figures/fig8_all_metric_bands",
    out_dir=None,    
    agg="sem",               # or "sd" or "quantile"
    #agg="sd",
    invert_x=True,
    title_prefix="Fig8: ",
    exclude_cols=("overlap_fraction", "train_paired", "train_unpaired", "unpaired_per_paired", "sampler_unpaired_per_paired", "SIL_true_frac",),  # optional
)
'''

# %% [code]
# Do different data splits per seed and see how that effects the eval results
import numpy as np
import pandas as pd

def build_dataset_for_split_seed(
    *,
    rna_raw,
    atac_raw,
    label_key: str,
    split_seed: int,
    train_frac: float = 0.8,
    val_frac: float = 0.1,
    # preprocessing params
    rna_counts_layer: str = "counts",
    atac_counts_layer: str = "counts",
    n_hvg: int = 2000,
    target_sum: float = 1e4,
    n_lsi: int = 101,
    preprocess_seed: int | None = None,  # if None, use split_seed
):
    """
    Returns:
      dataset, train_idx, val_idx, test_idx, y_all, (rna_tr_pp, atac_tr_lsi)  # last pair for dim lookups
    """
    if preprocess_seed is None:
        preprocess_seed = int(split_seed)

    # --- 1) resample splits ---
    rng = np.random.default_rng(int(split_seed))
    n = int(rna_raw.n_obs)
    idx = np.arange(n, dtype=np.int64)
    rng.shuffle(idx)

    n_tr = int(round(train_frac * n))
    n_va = int(round(val_frac * n))
    n_tr = max(1, min(n_tr, n - 2))
    n_va = max(1, min(n_va, n - n_tr - 1))
    n_te = n - n_tr - n_va

    tr_idx = idx[:n_tr]
    va_idx = idx[n_tr:n_tr + n_va]
    te_idx = idx[n_tr + n_va:]

    rna_train, rna_val, rna_test = rna_raw[tr_idx].copy(), rna_raw[va_idx].copy(), rna_raw[te_idx].copy()
    atac_train, atac_val, atac_test = atac_raw[tr_idx].copy(), atac_raw[va_idx].copy(), atac_raw[te_idx].copy()

    # --- 2) fit preprocessing on TRAIN only; apply to val/test ---
    (rna_tr_pp, atac_tr_lsi,
     rna_va_pp, atac_va_lsi,
     rna_te_pp, atac_te_lsi,
     rna_fit, atac_fit) = preprocess_multiome_splits_fit_apply(
        rna_train, atac_train,
        rna_val,   atac_val,
        rna_test,  atac_test,
        rna_counts_layer=rna_counts_layer,
        atac_counts_layer=atac_counts_layer,
        n_hvg=int(n_hvg),
        target_sum=float(target_sum),
        n_lsi=int(n_lsi),
        seed=int(preprocess_seed),
    )

    # --- 3) rebuild dataset exactly like your current notebook does ---
    adata_dict = {
        "rna":  rna_tr_pp.concatenate(rna_va_pp, rna_te_pp, batch_key=None),
        "atac": atac_tr_lsi.concatenate(atac_va_lsi, atac_te_lsi, batch_key=None),
    }
    adata_dict = align_paired_obs_names(adata_dict)

    base_ds = MultiModalDataset(adata_dict=adata_dict, X_key="X", device=None)

    # integer labels for metrics / encoding
    y_str = adata_dict["rna"].obs[label_key].astype(str).to_numpy()
    uniq = np.unique(y_str)
    label_to_int = {lab: i for i, lab in enumerate(uniq)}
    y_int = np.array([label_to_int[x] for x in y_str], dtype=np.int64)

    dataset_labeled = LabeledMultiModalDataset(base_ds, y_int=y_int, y_str=None)

    # indices into the concatenated dataset
    train_idx = np.arange(0, n_tr, dtype=np.int64)
    val_idx   = np.arange(n_tr, n_tr + n_va, dtype=np.int64)
    test_idx  = np.arange(n_tr + n_va, n_tr + n_va + n_te, dtype=np.int64)

    # y_all used by encode_embeddings_with_labels in your runner
    y_all = y_int  # keep it int-coded and stable inside this split

    return dataset_labeled, train_idx, val_idx, test_idx, y_all, (rna_tr_pp, atac_tr_lsi)


# %% [code]
import os
import pandas as pd

def _ensure_parent_dir(path: str):
    d = os.path.dirname(os.path.abspath(path))
    if d and not os.path.exists(d):
        os.makedirs(d, exist_ok=True)

def _append_tsv_union(path: str, df: pd.DataFrame, *, append: bool):
    _ensure_parent_dir(path)
    if append and os.path.exists(path):
        existing = pd.read_csv(path, sep="\t")
        all_cols = sorted(set(existing.columns).union(df.columns))
        existing2 = existing.reindex(columns=all_cols)
        df2 = df.reindex(columns=all_cols)
        out = pd.concat([existing2, df2], axis=0, ignore_index=True)
        out.to_csv(path, sep="\t", index=False)
    else:
        df.to_csv(path, sep="\t", index=False)

def run_fig8_resample_splits_to_tsv(
    *,
    rna_raw,
    atac_raw,
    label_key: str = "cell_type",
    split_seeds=(0, 1, 2, 3, 4),
    model_seeds=(0,),   # <-- tuple/list
    overlap_grid=(1.0, 0.9, 0.8, 0.7),
    patience: int = 50,
    fuse_mode: str = "moe",
    drop_modality: str = "atac",
    # preprocessing params
    n_hvg: int = 2000,
    n_lsi: int = 101,
    target_sum: float = 1e4,

    # output controls
    out_tsv: str | None = None,
    out_meta_tsv: str | None = None,
    append: bool = True,
):
    """
    Works with your CURRENT run_fig8_missing_modality_curve_v1 (no TSV args needed).
    Produces the same metrics, then writes TSV(s) from this wrapper.
    """
    all_dfs = []

    # globals expected by run_fig8_missing_modality_curve_v1
    global dataset, train_idx, val_idx, test_idx, y_all, rna_tr_pp, atac_tr_lsi

    # optional: write one row per (split_seed, model_seed) with run knobs
    meta_rows = []

    for split_seed in split_seeds:
        ds, tr, va, te, y, (rna_tr, atac_tr) = build_dataset_for_split_seed(
            rna_raw=rna_raw,
            atac_raw=atac_raw,
            label_key=label_key,
            split_seed=int(split_seed),
            n_hvg=int(n_hvg),
            n_lsi=int(n_lsi),
            target_sum=float(target_sum),
        )

        dataset     = ds
        train_idx   = tr
        val_idx     = va
        test_idx    = te
        y_all       = y
        rna_tr_pp   = rna_tr
        atac_tr_lsi = atac_tr

        for model_seed in model_seeds:
            # call your existing curve runner (no TSV kwargs!)
            d = run_fig8_missing_modality_curve_v1(
                overlap_grid=overlap_grid,
                patience=int(patience),
                fuse_mode=str(fuse_mode),
                drop_modality=str(drop_modality),
                seed=int(model_seed),
            )

            d["split_seed"] = int(split_seed)
            d["model_seed"] = int(model_seed)
            all_dfs.append(d)

            meta_rows.append({
                "split_seed": int(split_seed),
                "model_seed": int(model_seed),
                "label_key": str(label_key),
                "drop_modality": str(drop_modality),
                "fuse_mode": str(fuse_mode),
                "patience": int(patience),
                "n_hvg": int(n_hvg),
                "n_lsi": int(n_lsi),
                "target_sum": float(target_sum),
                "n_overlaps": int(len(overlap_grid)),
            })

            # save this run's rows incrementally (optional but nice)
            if out_tsv is not None:
                _append_tsv_union(out_tsv, d, append=append and os.path.exists(out_tsv))

    df_all = pd.concat(all_dfs, ignore_index=True) if len(all_dfs) else pd.DataFrame()

    if out_meta_tsv is not None:
        meta_df = pd.DataFrame(meta_rows)
        _append_tsv_union(out_meta_tsv, meta_df, append=append and os.path.exists(out_meta_tsv))

    return df_all


# %% [code]
import os
os.makedirs("./results/figure_9_paired_ablation_reproducibility", exist_ok=True)


# %% [code]
split_seeds = list(range(20))

df_all_atac_cv = run_fig8_resample_splits_to_tsv(
    rna_raw=rna_raw,
    atac_raw=atac_raw,
    label_key="cell_type",
    split_seeds=split_seeds,
    model_seeds=(0,),
    overlap_grid=(1.0, 0.9, 0.8, 0.7, 0.6, 0.5, 0.4, 0.3, 0.2, 0.1, 0.075, 0.05, 0.045, 0.04, 0.035, 0.03, 0.025, 0.02, 0.0175, 0.015, 0.0125, 0.01, 0.0075, 0.005),
    patience=50,
    fuse_mode="moe",
    drop_modality="atac",
    n_hvg=2000,
    n_lsi=101,
    target_sum=1e4,
    out_tsv="./results/figure_9_paired_ablation_outputs-2-10-2026/fig8_curve__drop_atac__all_seeds.tsv",
    out_meta_tsv="./results/figure_9_paired_ablation_outputs-2-10-2026/fig8_curve__drop_atac__all_seeds__meta.tsv",
)


# %% [code]
print(df_all_atac_cv)


# %% [code]
# One PNG per metric, autodetected from numeric columns
plot_many_metrics_with_band(
    df_all_atac_cv,
    #out_dir="./figures/fig8_all_20cv_metric_bands_quantile_agg",
    out_dir=None,    
    agg="quantile",               # or "sd" or "quantile"
    #agg="sd",
    #agg="sem",
    invert_x=True,
    title_prefix="Fig8: ",
    exclude_cols=("overlap_fraction", "train_paired", "train_unpaired", "unpaired_per_paired", "sampler_unpaired_per_paired", "SIL_true_frac",),  # optional
)


# %% [code]


# %% [code]
df_all_rna_cv = run_fig8_resample_splits_to_tsv(
    rna_raw=rna_raw,
    atac_raw=atac_raw,
    label_key="cell_type",
    split_seeds=split_seeds,
    model_seeds=(0,),
    overlap_grid=(1.0, 0.9, 0.8, 0.7, 0.6, 0.5, 0.4, 0.3, 0.2, 0.1, 0.075, 0.05, 0.045, 0.04, 0.035, 0.03, 0.025, 0.02, 0.0175, 0.015, 0.0125, 0.01, 0.0075, 0.005),
    patience=50,
    fuse_mode="moe",
    drop_modality="rna",
    n_hvg=2000,
    n_lsi=101,
    target_sum=1e4,
    out_tsv="./results/figure_9_paired_ablation_outputs-2-10-2026/fig8_curve__drop_rna__all_seeds.tsv",
    out_meta_tsv="./results/figure_9_paired_ablation_outputs-2-10-2026/fig8_curve__drop_rna__all_seeds__meta.tsv",
)


# %% [code]
print(df_all_rna_cv)


# %% [code]
# One PNG per metric, autodetected from numeric columns
plot_many_metrics_with_band(
    df_all_rna_cv,
    #out_dir="./figures/fig8_all_20cv_metric_bands_quantile_agg",
    out_dir=None,    
    agg="quantile",               # or "sd" or "quantile"
    #agg="sd",
    #agg="sem",
    invert_x=True,
    title_prefix="Fig8: ",
    exclude_cols=("overlap_fraction", "train_paired", "train_unpaired", "unpaired_per_paired", "sampler_unpaired_per_paired", "SIL_true_frac",),  # optional
)


# %% [code]


# %% [code]


# %% [code]


# %% [code]


# %% [code]


# %% [code]


# %% [code]


# %% [code]


# %% [code]


# %% [code]


# %% [code]


# %% [code]


# %% [code]


# %% [code]


# %% [code]


# %% [code]


# %% [code]


# %% [code]


# %% [code]


# %% [code]


# %% [code]


# %% [code]


# %% [code]


# %% [code]


# %% [code]


# %% [code]


# %% [code]


# %% [code]


# %% [code]


# %% [code]


# %% [code]


# %% [code]


# %% [code]


# %% [code]
```
