## Supplemental Code for "Unifying multimodal single-cell data with a mixture-of-experts *β*-variational autoencoder framework": Fig09_scaling_compile.ipynb

```
# %% [code]
import os
import pandas as pd

# ---- set this ----
path = "./ablation_analysis_results_fixed_MoE_UMAP_modality_colors/celltype_ablation_grid_metrics_COARSE.csv"

# ---- loader ----
ext = os.path.splitext(path)[1].lower()
if ext in [".tsv", ".txt"]:
    df = pd.read_csv(path, sep="\t")
elif ext == ".csv":
    df = pd.read_csv(path)
elif ext == ".parquet":
    df = pd.read_parquet(path)
else:
    raise ValueError(f"Unsupported file type: {ext}")

print(df.shape)
df.head()


# %% [code]
import numpy as np
import pandas as pd
import matplotlib.pyplot as plt

cond_col  = "drop_modality"        # 'rna' or 'atac'
label_col = "ablated_cell_type"    # coarse label


# %% [code]
def summarize_metric(df, metric_col, *, cond_col="drop_modality", label_col="ablated_cell_type",
                     mode_col=None, mode_value=None, agg="mean"):
    d = df.copy()

    # optional filter by fuse_mode (or other column)
    if mode_col is not None and mode_value is not None and mode_col in d.columns:
        d = d[d[mode_col].astype(str) == str(mode_value)].copy()

    # coerce numeric metric
    d[metric_col] = pd.to_numeric(d[metric_col], errors="coerce")
    d = d.dropna(subset=[metric_col, cond_col, label_col]).copy()
    if len(d) == 0:
        raise ValueError(f"No valid rows for metric_col={metric_col} after filtering.")

    # aggregate (handles repeats)
    if agg == "mean":
        g = d.groupby([cond_col, label_col], as_index=False)[metric_col].mean()
    elif agg == "median":
        g = d.groupby([cond_col, label_col], as_index=False)[metric_col].median()
    else:
        raise ValueError("agg must be 'mean' or 'median'")

    # pivot to: rows=drop_modality, cols=cell type
    P = g.pivot(index=cond_col, columns=label_col, values=metric_col)

    # enforce row order if present
    row_order = [r for r in ["rna", "atac"] if r in P.index.astype(str).tolist()]
    if row_order:
        P = P.reindex(row_order)

    return P  # DataFrame shape: (2, n_celltypes)


# %% [code]
def order_celltypes(P, how="mean", ref="rna"):
    """
    P: rows=drop_modality, cols=cell types
    how:
      - 'mean' : sort by mean across modalities (descending)
      - 'ref'  : sort by chosen ref row (descending)
      - 'diff' : sort by |rna - atac| (descending)
    """
    cols = P.columns.tolist()
    if how == "mean":
        score = P.mean(axis=0)
        return score.sort_values(ascending=False).index.tolist()
    if how == "ref":
        if ref not in P.index:
            return cols
        score = P.loc[ref]
        return score.sort_values(ascending=False).index.tolist()
    if how == "diff":
        if all(r in P.index for r in ["rna", "atac"]):
            score = (P.loc["rna"] - P.loc["atac"]).abs()
            return score.sort_values(ascending=False).index.tolist()
        return cols
    return cols


# %% [code]
def plot_two_modality_lines(P, *, title, y_label, ylim=None,
                            sort_how="mean", ref="rna",
                            figsize=(14, 12), dpi=300,
                            show_points=True):
    """
    P: DataFrame rows=drop_modality, cols=cell types
    """
    order = order_celltypes(P, how=sort_how, ref=ref)
    P = P[order]

    x = np.arange(P.shape[1])

    plt.figure(figsize=figsize, dpi=dpi)

    # plot each modality as a line
    for row in P.index:
        y = P.loc[row].to_numpy(dtype=float)
        plt.plot(x, y, marker="o" if show_points else None, linewidth=2, label=str(row))

    plt.xticks(x, [str(c) for c in P.columns], rotation=90)
    plt.xlabel("Ablated population (coarse label)")
    plt.ylabel(y_label)
    plt.title(title)
    plt.grid(True, axis="y", alpha=0.25)

    if ylim is not None:
        plt.ylim(*ylim)

    plt.legend(title="drop_modality")
    plt.tight_layout()
    plt.show()


# %% [code]
metric_col = "Recall@10"
P = summarize_metric(df, metric_col, cond_col=cond_col, label_col=label_col, agg="mean")

plot_two_modality_lines(
    P,
    title="Fig 10(c): correspondence degradation under targeted modality dropout (Recall@10)",
    y_label="Recall@10 (higher is better)",
    ylim=(0, 1),          # Recall is in [0,1]
    sort_how="mean",      # try: "mean", "diff", or "ref"
    ref="rna",
    figsize=(14, 12),
    dpi=300,
    show_points=True
)


# %% [code]
metric_col = "Recall@10"
P = summarize_metric(df, metric_col, cond_col=cond_col, label_col=label_col, agg="mean")

plot_two_modality_lines(
    P,
    title="Fig 10(c): correspondence degradation under targeted modality dropout (Recall@10)",
    y_label="Recall@10 (higher is better)",
    ylim=(0, 1),          # Recall is in [0,1]
    sort_how="mean",      # try: "mean", "diff", or "ref"
    ref="rna",
    figsize=(14, 12),
    dpi=300,
    show_points=True
)


# %% [code]
for how in ["mean", "diff", "ref"]:
    plot_two_modality_lines(
        P,
        title=f"Fig 10(c): Recall@10 (sorted by {how})",
        y_label="Recall@10",
        ylim=(0.8, 1),
        sort_how=how,
        ref="rna",
        figsize=(8, 6),
        dpi=300
    )


# %% [code]
metric_col = "FOSCTTM"
P = summarize_metric(df, metric_col, cond_col=cond_col, label_col=label_col, agg="mean")

plot_two_modality_lines(
    P,
    title="Fig 10(c): correspondence degradation under targeted modality dropout (FOSCTTM)",
    y_label="FOSCTTM (lower is better)",
    #ylim=None,            # let it autoscale
    ylim=(0, .175),
    sort_how="mean",
    figsize=(8, 6),
    dpi=300
)

# optional: invert y-axis so "better" is visually higher
plt.gca().invert_yaxis()
plt.show()


# %% [code]
metric_col = "ACC_fused_macroF1"
P = summarize_metric(df, metric_col, cond_col=cond_col, label_col=label_col, agg="mean")

plot_two_modality_lines(
    P,
    title="Fig 10(d): semantic stability under targeted modality dropout (macro-F1)",
    y_label="Macro-F1 (higher is better)",
    ylim=(0.9, 1),
    sort_how="mean",
    figsize=(8, 6),
    dpi=300
)


# %% [code]


# %% [code]


# %% [code]


# %% [code]


# %% [code]


# %% [code]


# %% [code]


# %% [code]


# %% [code]


# %% [code]


# %% [code]


# %% [code]


# %% [code]


# %% [code]


# %% [code]


# %% [code]


# %% [code]


# %% [code]


# %% [code]


# %% [code]


# %% [code]


# %% [code]


# %% [code]


# %% [code]


# %% [code]


# %% [code]


# %% [code]


# %% [code]


# %% [code]
```
