## Supplemental Code for "Unifying multimodal single-cell data with a mixture-of-experts *β*-variational autoencoder framework": Fig10_population_ablation_MoE.ipynb

```
# %% [markdown]
## UniVI – Figure 10 analysis (robust v1, no leakage)

This notebook performs the Figure 10 coarse + fine celltype ablation analysis using PBMC Multiome data.

 - organized, single source of truth (no duplicates)
 - robust latent extraction
 - MoE fused latent computed ONCE in encoder (MoE-first)
 - STRICT: if MoE latent not available, FAIL (no avg/joint fallback)
 - grouped batching for ablated TRAIN to avoid mixed missingness in a batch


# %% [code]
# =============================================================================
# ---- 0) Imports / versions ----
# =============================================================================
import os
import gc
import copy
import inspect
from dataclasses import dataclass
from typing import Any, Dict, List, Optional, Sequence, Tuple

import numpy as np
import pandas as pd
import scipy.sparse as sp
import anndata as ad
import scanpy as sc

import torch
from torch.utils.data import Dataset, DataLoader, Subset, Sampler

from sklearn.feature_extraction.text import TfidfTransformer
from sklearn.decomposition import TruncatedSVD
from sklearn.preprocessing import StandardScaler

from sklearn.metrics import (
    adjusted_rand_score,
    normalized_mutual_info_score,
    silhouette_score,
    accuracy_score,
    f1_score,
)
from sklearn.cluster import KMeans
from sklearn.neighbors import KNeighborsClassifier

import matplotlib.pyplot as plt
from matplotlib.lines import Line2D

# UniVI
from univi.data import MultiModalDataset, align_paired_obs_names
from univi import UniVIMultiModalVAE, ModalityConfig, UniVIConfig, TrainingConfig
from univi.trainer import UniVITrainer

print("scanpy:", sc.__version__)
print("torch:", torch.__version__)


# %% [code]
# =============================================================================
# ---- 1) User config ----
# =============================================================================
RNA_PATH  = "./data/10x_Genomics_Multiome_data/10x-Multiome-Pbmc10k-RNA.h5ad"
ATAC_PATH = "./data/10x_Genomics_Multiome_data/10x-Multiome-Pbmc10k-ATAC.h5ad"

CELLTYPE_KEY = "cell_type"
COUNTS_LAYER = "counts"

SEED = 42
DEVICE = "mps"  # "cuda" / "cpu" also ok

# preprocessing
N_HVG = 2000
RNA_TARGET_SUM = 1e4
N_LSI = 101

# training
BATCH_SIZE = 128
RECALL_KS = (1, 5, 10, 25, 50, 100)

# STRICT behavior
STRICT_MOE = True  # if True: require mu_moe, else crash
# If you want to allow model-fused fallback (NOT avg), set STRICT_MOE=False and use fuse_mode="fused"


# %% [code]
# =============================================================================
# ---- 2) Small utilities (single source of truth) ----
# =============================================================================
def pick_device():
    return torch.device(
        "cuda" if torch.cuda.is_available()
        else ("mps" if torch.backends.mps.is_available() else "cpu")
    )

def ensure_counts_layer(adata: ad.AnnData, layer: str = "counts") -> None:
    if layer not in adata.layers:
        adata.layers[layer] = adata.X.copy()

def _to_csr(X):
    return X if sp.issparse(X) else sp.csr_matrix(X)

def _as_numpy_2d(x):
    if x is None:
        return None
    if torch.is_tensor(x):
        x = x.detach().cpu().numpy()
    x = np.asarray(x)
    if x.ndim == 1:
        x = x[:, None]
    return x.astype(np.float32, copy=False)

def _as_numpy_1d(y):
    if y is None:
        return None
    if torch.is_tensor(y):
        y = y.detach().cpu().numpy()
    y = np.asarray(y)
    return y.reshape(-1)

def _to_numpy_ids(gids):
    if gids is None:
        return None
    if torch.is_tensor(gids):
        return gids.detach().cpu().numpy().astype(np.int64, copy=False)
    return np.asarray(gids, dtype=np.int64)

def _unwrap_model_output(out):
    if torch.is_tensor(out) or isinstance(out, dict):
        return out
    if isinstance(out, (tuple, list)) and len(out) > 0:
        for item in out:
            if isinstance(item, dict) or torch.is_tensor(item):
                return item
        return out[0]
    return out

def _smart_call(fn, x_dict):
    try:
        sig = inspect.signature(fn)
        params = sig.parameters
        for name in ("x_dict", "batch", "x", "inputs"):
            if name in params:
                return fn(**{name: x_dict})
        return fn(x_dict)
    except TypeError:
        return fn(x_dict)

def _call_univi_encoder(model, x_dict):
    if hasattr(model, "encode") and callable(getattr(model, "encode")):
        out = _smart_call(model.encode, x_dict)
        return _unwrap_model_output(out)
    out = _smart_call(model, x_dict)
    return _unwrap_model_output(out)
    

# %% [code]
# =============================================================================
# ---- 3) Robust latent extraction + MoE fusion (THE FIX) ----
# =============================================================================
def _first_tensor(*xs):
    for x in xs:
        if torch.is_tensor(x):
            return x
    return None

def _find_tensors_anywhere(obj):
    out = []
    if torch.is_tensor(obj):
        out.append(obj)
    elif isinstance(obj, dict):
        for v in obj.values():
            out.extend(_find_tensors_anywhere(v))
    elif isinstance(obj, (list, tuple)):
        for v in obj:
            out.extend(_find_tensors_anywhere(v))
    return out

def _extract_latent(enc, which: str):
    """
    which in {"rna","atac","fused"}.
    Tries common UniVI layouts/keys; last resort: any 2D tensor.
    """
    if enc is None:
        return None

    if torch.is_tensor(enc):
        # if model directly returns a tensor, treat it as "fused"
        return enc if which == "fused" else None

    if not isinstance(enc, dict):
        return None

    # ---- direct keys like mu_rna / z_atac etc ----
    v = _first_tensor(
        enc.get(f"mu_{which}", None),
        enc.get(f"z_{which}", None),
        enc.get(f"latent_{which}", None),
        enc.get(f"embedding_{which}", None),
        enc.get(f"emb_{which}", None),
    )
    if v is not None:
        return v

    # ---- dict-of-modality patterns (mu_dict / z_dict) ----
    for dkey in ("mu_dict", "z_dict", "latent_dict", "emb_dict"):
        sub = enc.get(dkey, None)
        if isinstance(sub, dict) and which in sub and torch.is_tensor(sub[which]):
            return sub[which]

    # ---- fused special keys (UniVI commonly returns mu_z / z as fused posterior) ----
    if which == "fused":
        v = _first_tensor(
            enc.get("mu_moe", None),
            enc.get("mu_joint", None),
            enc.get("mu_fused", None),
            enc.get("mu_shared", None),

            # common UniVI fused outputs
            enc.get("mu_z", None),   # fused mean
            enc.get("z", None),      # fused sample

            enc.get("mu", None),
            enc.get("mean", None),
            enc.get("latent", None),
            enc.get("embedding", None),
        )
        if v is not None:
            return v

    # ---- nested dict possibilities ----
    for container in (
        "latents", "latent", "posterior", "post", "qz", "q",
        "enc", "encode", "outputs", "moe", "mix", "router", "extras"
    ):
        sub = enc.get(container, None)
        if isinstance(sub, dict):
            v = _first_tensor(
                sub.get(f"mu_{which}", None),
                sub.get(f"z_{which}", None),
                sub.get(f"latent_{which}", None),
                sub.get(f"embedding_{which}", None),
            )
            if v is not None:
                return v

            if which == "fused":
                v = _first_tensor(
                    sub.get("mu_moe", None),
                    sub.get("mu_joint", None),
                    sub.get("mu_fused", None),
                    sub.get("mu_z", None),
                    sub.get("z", None),
                    sub.get("mu", None),
                )
                if v is not None:
                    return v

    # last resort: any 2D tensor
    ts = _find_tensors_anywhere(enc)
    for t in ts:
        if torch.is_tensor(t) and t.ndim == 2:
            return t
    return ts[0] if ts else None

def _extract_gating_weights_from_enc_full(enc_full):
    """
    Precision-derived implicit MoE weights from per-modality log-variances.

    Returns:
      (W, mods)
        W: torch.Tensor (B, M) with per-cell modality weights (mean over latent dims),
           where M = number of modalities with a tensor logvar.
        mods: list[str] modality names in the same column order as W

    If not available, returns (None, None).
    """
    if not isinstance(enc_full, dict):
        return None, None

    lv = enc_full.get("logvar_dict", None)
    if not isinstance(lv, dict) or len(lv) == 0:
        return None, None

    # keep only modalities with tensor logvars
    mods = [k for k, v in lv.items() if (v is not None and torch.is_tensor(v))]
    if len(mods) < 2:
        return None, None

    # stack precisions: (M, B, D)
    prec = torch.stack([torch.exp(-lv[m]) for m in mods], dim=0)

    # normalize across modalities: (M, B, D)
    denom = prec.sum(dim=0).clamp_min(1e-8)
    w = prec / denom

    # reduce over latent dims -> (M, B)
    w_mean = w.mean(dim=-1)

    # transpose -> (B, M)
    W = w_mean.transpose(0, 1).contiguous()
    return W, mods

def _pick_explicit_moe_tensor(enc_full):
    """
    If UniVI already returns a fused latent explicitly, prefer it.

    IMPORTANT:
    - In many UniVI versions, the fused posterior mean is returned as mu_z (NOT mu_moe).
    - If gating is not exposed, mu_z is still the correct fused latent (not an average).
    """
    if enc_full is None:
        return None
    if torch.is_tensor(enc_full):
        return None
    if not isinstance(enc_full, dict):
        return None

    v = _first_tensor(
        enc_full.get("mu_moe", None),
        enc_full.get("mu_joint", None),
        enc_full.get("mu_z", None),
    )
    if v is not None:
        return v

    for container in ("moe", "mix", "mixture", "router", "extras", "outputs", "enc", "latent", "latents", "posterior", "post"):
        sub = enc_full.get(container, None)
        if isinstance(sub, dict):
            v = _first_tensor(
                sub.get("mu_moe", None),
                sub.get("mu_joint", None),
                sub.get("mu_z", None),
            )
            if v is not None:
                return v

    return None


# %% [code]
# =============================================================================
# ---- 4) Label coding + paired alignment metrics ----
# =============================================================================
def _build_label_encoder(y_all):
    y_raw = np.asarray(y_all)
    kind = y_raw.dtype.kind
    if kind in ("i", "u", "f", "b"):
        return y_raw, None
    y_raw = y_raw.astype(str)
    uniq = np.unique(y_raw)
    id_to_int = {lab: i for i, lab in enumerate(uniq.tolist())}
    return y_raw, id_to_int

def _encode_labels_for_gids(y_raw, gids_np, *, id_to_int=None):
    y_batch = y_raw[gids_np]
    y_batch_arr = np.asarray(y_batch)
    kind = y_batch_arr.dtype.kind

    if kind in ("i", "u", "b"):
        return torch.as_tensor(y_batch_arr.astype(np.int64, copy=False), dtype=torch.long), id_to_int
    if kind == "f":
        return torch.as_tensor(y_batch_arr.astype(np.float32, copy=False)), id_to_int

    y_batch_str = y_batch_arr.astype(str)
    if id_to_int is None:
        uniq = np.unique(y_batch_str)
        id_to_int = {lab: i for i, lab in enumerate(uniq.tolist())}

    codes = np.empty(y_batch_str.shape[0], dtype=np.int64)
    next_code = (max(id_to_int.values()) + 1) if len(id_to_int) else 0
    for i, lab in enumerate(y_batch_str):
        if lab not in id_to_int:
            id_to_int[lab] = next_code
            next_code += 1
        codes[i] = id_to_int[lab]

    return torch.as_tensor(codes, dtype=torch.long), id_to_int

def _pairwise_sq_dists(X, Y):
    X = _as_numpy_2d(X)
    Y = _as_numpy_2d(Y)
    x2 = np.sum(X * X, axis=1, keepdims=True)
    y2 = np.sum(Y * Y, axis=1, keepdims=True).T
    D = x2 + y2 - 2.0 * (X @ Y.T)
    np.maximum(D, 0.0, out=D)
    return D

def foscttm(X, Y, *, symmetric=True):
    D = _pairwise_sq_dists(X, Y)
    n = D.shape[0]
    if n <= 1:
        return float(np.nan)
    diag = np.diag(D)
    denom = (n - 1)
    frac_xy = (D < diag[:, None]).sum(axis=1) / denom
    if not symmetric:
        return float(frac_xy.mean())
    frac_yx = (D < diag[None, :]).sum(axis=0) / denom
    return float(0.5 * (frac_xy.mean() + frac_yx.mean()))

def recall_at_k(X, Y, *, ks=(1, 10, 25, 50, 100), symmetric=True):
    D = _pairwise_sq_dists(X, Y)
    n = D.shape[0]
    if n == 0:
        return {int(k): float(np.nan) for k in ks}
    ks = [int(k) for k in ks if int(k) >= 1]
    order_xy = np.argsort(D, axis=1)

    pos_xy = np.empty(n, dtype=np.int32)
    for i in range(n):
        pos_xy[i] = np.where(order_xy[i] == i)[0][0]

    if symmetric:
        order_yx = np.argsort(D.T, axis=1)
        pos_yx = np.empty(n, dtype=np.int32)
        for i in range(n):
            pos_yx[i] = np.where(order_yx[i] == i)[0][0]

    out = {}
    for k in ks:
        k_eff = min(k, n)
        r_xy = float((pos_xy < k_eff).mean())
        if not symmetric:
            out[k] = r_xy
        else:
            r_yx = float((pos_yx < k_eff).mean())
            out[k] = 0.5 * (r_xy + r_yx)
    return out


# %% [code]
# =============================================================================
# ---- 5) Preprocess RNA/ATAC (fit on TRAIN only, apply to val/test) ----
# =============================================================================
def _is_integer_like_sparse(X, max_check=200000) -> bool:
    if sp.issparse(X):
        data = X.data
    else:
        data = np.asarray(X).ravel()
    if data.size == 0:
        return True
    if data.size > max_check:
        data = data[:max_check]
    return np.all(np.isfinite(data)) and np.all(np.abs(data - np.round(data)) < 1e-6)

@dataclass
class RNAFit:
    hvg: List[str]
    counts_layer: str = "counts"
    target_sum: float = 1e4
    out_layer: str = "log1p"

@dataclass
class ATACFit:
    counts_layer: str = "counts"
    tfidf: Any = None
    svd: Any = None
    scaler: Any = None
    n_lsi: int = 101
    drop_first: bool = True

def fit_rna_on_train(
    rna_train: ad.AnnData,
    *,
    counts_layer: str = "counts",
    n_hvg: int = 2000,
    target_sum: float = 1e4,
    out_layer: str = "log1p",
) -> RNAFit:
    rna = rna_train.copy()
    if counts_layer not in rna.layers:
        rna.layers[counts_layer] = rna.X.copy()

    rna.layers[out_layer] = _to_csr(rna.layers[counts_layer]).copy()
    sc.pp.normalize_total(rna, target_sum=float(target_sum), layer=out_layer)
    sc.pp.log1p(rna, layer=out_layer)

    counts_ok = _is_integer_like_sparse(rna.layers[counts_layer])
    flavors = (["seurat_v3"] if counts_ok else []) + ["cell_ranger", "seurat"]

    last = None
    for flavor in flavors:
        try:
            sc.pp.highly_variable_genes(
                rna,
                layer=out_layer,
                n_top_genes=int(n_hvg),
                flavor=flavor,
                subset=False,
            )
            if "highly_variable" in rna.var.columns and int(rna.var["highly_variable"].sum()) > 0:
                break
        except Exception as e:
            last = e
            continue
    else:
        raise RuntimeError(f"HVG selection failed. Tried {flavors}. Last error: {last}")

    hvg = rna.var_names[rna.var["highly_variable"].to_numpy()].tolist()
    if len(hvg) == 0:
        raise RuntimeError("No HVGs selected; check RNA values.")
    return RNAFit(hvg=hvg, counts_layer=counts_layer, target_sum=float(target_sum), out_layer=out_layer)

def apply_rna_fit(rna: ad.AnnData, fit: RNAFit) -> ad.AnnData:
    a = rna[:, fit.hvg].copy()
    if fit.counts_layer not in a.layers:
        a.layers[fit.counts_layer] = a.X.copy()

    a.layers[fit.out_layer] = _to_csr(a.layers[fit.counts_layer]).copy()
    sc.pp.normalize_total(a, target_sum=float(fit.target_sum), layer=fit.out_layer)
    sc.pp.log1p(a, layer=fit.out_layer)
    a.X = a.layers[fit.out_layer]
    return a

def fit_atac_on_train(
    atac_train: ad.AnnData,
    *,
    counts_layer: str = "counts",
    n_lsi: int = 101,
    seed: int = 0,
) -> ATACFit:
    a = atac_train.copy()
    if counts_layer not in a.layers:
        a.layers[counts_layer] = a.X.copy()
    X = _to_csr(a.layers[counts_layer])

    tfidf = TfidfTransformer()
    X_tfidf = tfidf.fit_transform(X)

    svd = TruncatedSVD(n_components=int(n_lsi), random_state=int(seed))
    X_lsi = svd.fit_transform(X_tfidf)

    scaler = StandardScaler(with_mean=True, with_std=True)
    X_lsi = scaler.fit_transform(X_lsi)

    return ATACFit(counts_layer=counts_layer, tfidf=tfidf, svd=svd, scaler=scaler, n_lsi=int(n_lsi), drop_first=True)

def apply_atac_fit(atac: ad.AnnData, fit: ATACFit) -> ad.AnnData:
    a = atac.copy()
    if fit.counts_layer not in a.layers:
        a.layers[fit.counts_layer] = a.X.copy()

    X = _to_csr(a.layers[fit.counts_layer])
    X_tfidf = fit.tfidf.transform(X)
    X_lsi = fit.svd.transform(X_tfidf)
    X_lsi = fit.scaler.transform(X_lsi)

    if fit.drop_first and X_lsi.shape[1] > 1:
        X_lsi = X_lsi[:, 1:]

    n_feats = X_lsi.shape[1]
    var = pd.DataFrame(index=[f"LSI_{i+1}" for i in range(n_feats)])
    atac_lsi = ad.AnnData(
        X=X_lsi.astype(np.float32, copy=False),
        obs=a.obs.copy(),
        var=var,
    )
    atac_lsi.uns["lsi"] = {"n_components": int(n_feats), "drop_first": bool(fit.drop_first)}
    return atac_lsi

def preprocess_splits(
    rna: ad.AnnData,
    atac: ad.AnnData,
    train_idx: np.ndarray,
    val_idx: np.ndarray,
    test_idx: np.ndarray,
    *,
    counts_layer: str = "counts",
    n_hvg: int = 2000,
    target_sum: float = 1e4,
    n_lsi: int = 101,
    seed: int = 0,
):
    rna_train, rna_val, rna_test = rna[train_idx].copy(), rna[val_idx].copy(), rna[test_idx].copy()
    atac_train, atac_val, atac_test = atac[train_idx].copy(), atac[val_idx].copy(), atac[test_idx].copy()

    rna_fit  = fit_rna_on_train(rna_train, counts_layer=counts_layer, n_hvg=n_hvg, target_sum=target_sum)
    atac_fit = fit_atac_on_train(atac_train, counts_layer=counts_layer, n_lsi=n_lsi, seed=seed)

    rna_tr = apply_rna_fit(rna_train, rna_fit)
    rna_va = apply_rna_fit(rna_val,   rna_fit)
    rna_te = apply_rna_fit(rna_test,  rna_fit)

    atac_tr = apply_atac_fit(atac_train, atac_fit)
    atac_va = apply_atac_fit(atac_val,   atac_fit)
    atac_te = apply_atac_fit(atac_test,  atac_fit)

    return rna_tr, atac_tr, rna_va, atac_va, rna_te, atac_te, rna_fit, atac_fit
    

# %% [code]
# =============================================================================
# ---- 6) Load paired data + split + preprocess + build dataset ----
# =============================================================================
rna_raw  = sc.read_h5ad(RNA_PATH)
atac_raw = sc.read_h5ad(ATAC_PATH)
print(rna_raw)
print(atac_raw)

adata_dict_raw = {"rna": rna_raw, "atac": atac_raw}
adata_dict_raw = align_paired_obs_names(adata_dict_raw)
rna_raw  = adata_dict_raw["rna"]
atac_raw = adata_dict_raw["atac"]

assert (rna_raw.obs_names == atac_raw.obs_names).all(), "RNA/ATAC obs_names not aligned"
ensure_counts_layer(rna_raw, COUNTS_LAYER)
ensure_counts_layer(atac_raw, COUNTS_LAYER)

assert CELLTYPE_KEY in rna_raw.obs.columns, f"{CELLTYPE_KEY} missing in rna_raw.obs"
assert CELLTYPE_KEY in atac_raw.obs.columns, f"{CELLTYPE_KEY} missing in atac_raw.obs"

rng = np.random.default_rng(SEED)
y = rna_raw.obs[CELLTYPE_KEY].to_numpy()
idx_all = np.arange(rna_raw.n_obs)

train_frac, val_frac = 0.75, 0.10
train_idx, val_idx, test_idx = [], [], []
for ct in np.unique(y):
    idx_ct = idx_all[y == ct]
    rng.shuffle(idx_ct)
    n_ct = len(idx_ct)
    n_tr = int(np.floor(train_frac * n_ct))
    n_va = int(np.floor(val_frac * n_ct))
    tr = idx_ct[:n_tr]
    va = idx_ct[n_tr:n_tr+n_va]
    te = idx_ct[n_tr+n_va:]
    train_idx.append(tr); val_idx.append(va); test_idx.append(te)

train_idx = np.concatenate(train_idx)
val_idx   = np.concatenate(val_idx)
test_idx  = np.concatenate(test_idx)

rng.shuffle(train_idx); rng.shuffle(val_idx); rng.shuffle(test_idx)

print("train/val/test:", len(train_idx), len(val_idx), len(test_idx))
print("train label counts:\n", pd.Series(y[train_idx]).value_counts())
print("val label counts:\n", pd.Series(y[val_idx]).value_counts())
print("test label counts:\n", pd.Series(y[test_idx]).value_counts())

rna_tr, atac_tr, rna_va, atac_va, rna_te, atac_te, rna_fit, atac_fit = preprocess_splits(
    rna_raw, atac_raw,
    train_idx, val_idx, test_idx,
    counts_layer=COUNTS_LAYER,
    n_hvg=N_HVG,
    target_sum=RNA_TARGET_SUM,
    n_lsi=N_LSI,
    seed=SEED,
)

adata_dict = {
    "rna":  rna_tr.concatenate(rna_va, rna_te, batch_key=None),
    "atac": atac_tr.concatenate(atac_va, atac_te, batch_key=None),
}
adata_dict = align_paired_obs_names(adata_dict)
dataset = MultiModalDataset(adata_dict=adata_dict, X_key="X", device=None)

n_tr = rna_tr.n_obs
n_va = rna_va.n_obs
n_te = rna_te.n_obs

TRAIN_IDX = np.arange(0, n_tr)
VAL_IDX   = np.arange(n_tr, n_tr + n_va)
TEST_IDX  = np.arange(n_tr + n_va, n_tr + n_va + n_te)

y_all = adata_dict["rna"].obs[CELLTYPE_KEY].to_numpy()
print("dataset n:", len(dataset), "train/val/test:", n_tr, n_va, n_te)
print("n unique cell types:", len(np.unique(y_all)))


# %% [code]
# =============================================================================
# ---- 7) Ablation masking + grouped batching (TRAIN only) ----
# =============================================================================
class IndexedDataset(Dataset):
    def __init__(self, base_ds, *, global_indices):
        self.base = base_ds
        self.global_indices = np.asarray(global_indices)

    def __len__(self):
        return len(self.base)

    def __getitem__(self, i):
        x = self.base[i]
        gid = int(self.global_indices[i])
        return x, gid

class DeterministicMaskDataset(Dataset):
    # group=0: paired; group=1: RNA-only; group=2: ATAC-only
    def __init__(self, base_ds: Dataset, groups: np.ndarray):
        self.base = base_ds
        self.groups = np.asarray(groups).astype(int)
        assert len(self.base) == len(self.groups)

    def __len__(self):
        return len(self.base)

    def __getitem__(self, i):
        x, gid = self.base[i]
        g = int(self.groups[i])
        if g == 0:
            return x, gid
        x2 = dict(x)
        if g == 1:
            x2.pop("atac", None)
        elif g == 2:
            x2.pop("rna", None)
        else:
            raise ValueError(f"Unknown group {g}")
        return x2, gid

class GroupedBatchSampler(Sampler[List[int]]):
    """
    Samples batches from a single group at a time (no mixed missingness inside a batch).
    """
    def __init__(self, groups: np.ndarray, batch_size: int, seed: int = 0, drop_last: bool = True):
        self.groups = np.asarray(groups).astype(int)
        self.batch_size = int(batch_size)
        self.seed = int(seed)
        self.drop_last = bool(drop_last)

        self.group_to_indices = {}
        for g in np.unique(self.groups):
            self.group_to_indices[int(g)] = np.where(self.groups == g)[0].tolist()

        self.nonempty_groups = [g for g, idxs in self.group_to_indices.items() if len(idxs) > 0]
        if len(self.nonempty_groups) == 0:
            raise ValueError("No samples in any group.")

    def __iter__(self):
        rng = np.random.default_rng(self.seed)
        pools = {}
        for g in self.nonempty_groups:
            idxs = self.group_to_indices[g].copy()
            rng.shuffle(idxs)
            pools[g] = idxs

        while True:
            available = [g for g in self.nonempty_groups if len(pools[g]) >= self.batch_size]
            if len(available) == 0:
                break
            chosen = int(rng.choice(available))
            batch = [pools[chosen].pop() for _ in range(self.batch_size)]
            yield batch

        if not self.drop_last:
            for g in self.nonempty_groups:
                while len(pools[g]) > 0:
                    b = pools[g][: self.batch_size]
                    pools[g] = pools[g][len(b):]
                    yield b

    def __len__(self):
        return sum(len(self.group_to_indices[g]) // self.batch_size for g in self.nonempty_groups)

def _stack_collate(batch):
    x0 = batch[0][0]
    keys = list(x0.keys())
    x_dict = {k: torch.stack([b[0][k] for b in batch], 0) for k in keys}
    gids = torch.tensor([b[1] for b in batch], dtype=torch.long)
    return x_dict, gids

def make_loaders_for_celltype_ablation(
    base_dataset,
    train_idx,
    val_idx,
    test_idx,
    *,
    y_all: np.ndarray,
    ablate_cell_type: str,
    drop_modality: str,   # "rna" or "atac"
    seed: int = 0,
    batch_size: int = 128,
    num_workers: int = 0,
    disable_ablation: bool = False,
):
    train_base = Subset(base_dataset, train_idx)
    val_base   = Subset(base_dataset, val_idx)
    test_base  = Subset(base_dataset, test_idx)

    train_indexed = IndexedDataset(train_base, global_indices=train_idx)
    val_indexed   = IndexedDataset(val_base,   global_indices=val_idx)
    test_indexed  = IndexedDataset(test_base,  global_indices=test_idx)

    y_train = y_all[train_idx]
    groups = np.zeros(len(train_indexed), dtype=int)

    if not disable_ablation:
        is_ct = (y_train == ablate_cell_type)
        if str(drop_modality) == "atac":
            groups[is_ct] = 1  # RNA-only
        elif str(drop_modality) == "rna":
            groups[is_ct] = 2  # ATAC-only
        else:
            raise ValueError("drop_modality must be 'atac' or 'rna'")

    train_masked = DeterministicMaskDataset(train_indexed, groups=groups)
    batch_sampler = GroupedBatchSampler(groups=groups, batch_size=int(batch_size), seed=int(seed), drop_last=True)

    train_loader = DataLoader(
        train_masked,
        batch_sampler=batch_sampler,
        num_workers=int(num_workers),
        collate_fn=_stack_collate,
    )

    val_loader = DataLoader(
        val_indexed,
        batch_size=int(batch_size),
        shuffle=False,
        num_workers=int(num_workers),
        collate_fn=_stack_collate,
    )
    test_loader = DataLoader(
        test_indexed,
        batch_size=int(batch_size),
        shuffle=False,
        num_workers=int(num_workers),
        collate_fn=_stack_collate,
    )
    return train_loader, val_loader, test_loader


# %% [code]
# =============================================================================
# ---- 8) UniVI training helpers (Fig8-matched + warmup + compat) ----
# =============================================================================
def make_univi_cfg(rna_dim: int, atac_dim: int) -> UniVIConfig:
    return UniVIConfig(
        latent_dim=30,
        beta=1.25,
        gamma=4.35,
        encoder_dropout=0.10,
        decoder_dropout=0.05,
        encoder_batchnorm=True,
        decoder_batchnorm=False,
        kl_anneal_start=50,
        kl_anneal_end=100,
        align_anneal_start=75,
        align_anneal_end=125,
        modalities=[
            ModalityConfig(
                name="rna",
                input_dim=int(rna_dim),
                encoder_hidden=[512, 256, 128],
                decoder_hidden=[128, 256, 512],
                likelihood="gaussian",
            ),
            ModalityConfig(
                name="atac",
                input_dim=int(atac_dim),
                encoder_hidden=[128, 64],
                decoder_hidden=[64, 128],
                likelihood="gaussian",
            ),
        ],
        class_heads=[],
    )

def make_train_cfg(device) -> TrainingConfig:
    return TrainingConfig(
        n_epochs=5000,
        batch_size=int(BATCH_SIZE),
        lr=1e-3,
        weight_decay=1e-4,
        device=device,
        log_every=25,
        grad_clip=5.0,
        early_stopping=True,
        patience=50,
    )

def _patch_trainer_best_epoch_warmup(trainer, warmup_epochs: int):
    warmup_epochs = int(warmup_epochs or 0)
    if warmup_epochs <= 0:
        return trainer

    candidate_method_names = [
        "_update_best", "_maybe_update_best", "_maybe_save_best", "_save_best",
        "_early_stopping_step", "_check_early_stopping", "_maybe_early_stop",
        "update_best", "maybe_save_best", "early_stopping_step", "check_early_stopping",
    ]

    def _infer_epoch(args, kwargs):
        if "epoch" in kwargs and kwargs["epoch"] is not None:
            return int(kwargs["epoch"])
        for attr in ("epoch", "current_epoch", "global_epoch"):
            v = getattr(trainer, attr, None)
            if isinstance(v, (int, np.integer)):
                return int(v)
        for a in args:
            if isinstance(a, (int, np.integer)):
                return int(a)
        return None

    patched_any = False
    for name in candidate_method_names:
        fn = getattr(trainer, name, None)
        if fn is None or not callable(fn):
            continue
        orig = fn

        def wrapped(*args, __orig=orig, **kwargs):
            ep = _infer_epoch(args, kwargs)
            if ep is not None and ep < warmup_epochs:
                return None
            return __orig(*args, **kwargs)

        setattr(trainer, name, wrapped)
        patched_any = True

    trainer._best_epoch_warmup_epochs = warmup_epochs
    trainer._best_epoch_warmup_patched = bool(patched_any)
    return trainer

def _build_univi_trainer_compat(*, model, train_cfg, train_loader, val_loader):
    sig = inspect.signature(UniVITrainer.__init__)
    params = sig.parameters
    kwargs = {}
    if "model" in params:
        kwargs["model"] = model
    if "train_loader" in params:
        kwargs["train_loader"] = train_loader
    if "val_loader" in params:
        kwargs["val_loader"] = val_loader

    cfg_key = None
    for cand in ("train_cfg", "training_cfg", "cfg_train", "config", "cfg", "training_config"):
        if cand in params:
            cfg_key = cand
            break
    if cfg_key is not None:
        kwargs[cfg_key] = train_cfg
        return UniVITrainer(**kwargs)

    for k, v in vars(train_cfg).items():
        if k in params:
            kwargs[k] = v
    return UniVITrainer(**kwargs)

def _trainer_fit_compat(trainer, train_loader, val_loader):
    try:
        return trainer.fit()
    except TypeError:
        pass
    try:
        return trainer.fit(train_loader=train_loader, val_loader=val_loader)
    except TypeError:
        pass
    return trainer.fit(train_loader, val_loader)

def train_one_model(
    *,
    train_loader,
    val_loader,
    rna_dim: int,
    atac_dim: int,
    seed: int = 0,
    device=None,
    loss_mode: str = "v1",
    v1_recon: str = "moe",
    best_epoch_warmup: int = 0,
):
    if device is None:
        device = pick_device()
    torch.manual_seed(int(seed))
    np.random.seed(int(seed))

    univi_cfg = make_univi_cfg(rna_dim=int(rna_dim), atac_dim=int(atac_dim))
    train_cfg = make_train_cfg(device=device)

    model = UniVIMultiModalVAE(
        univi_cfg,
        loss_mode=str(loss_mode),
        v1_recon=str(v1_recon),
    ).to(device)

    trainer = _build_univi_trainer_compat(
        model=model,
        train_cfg=train_cfg,
        train_loader=train_loader,
        val_loader=val_loader,
    )
    if not hasattr(trainer, "train_loader"):
        trainer.train_loader = train_loader
    if not hasattr(trainer, "val_loader"):
        trainer.val_loader = val_loader

    _patch_trainer_best_epoch_warmup(trainer, warmup_epochs=int(best_epoch_warmup))
    _trainer_fit_compat(trainer, train_loader=train_loader, val_loader=val_loader)
    return model


# %% [code]
# =============================================================================
# ---- 9) Encoding (MoE-first, robust) + STRICT MoE accessor ----
# =============================================================================
@torch.no_grad()
def encode_embeddings_with_labels(
    model,
    loader,
    *,
    device,
    y_all,
    id_to_int=None,
    debug_first_batch=False,
):
    """
    Returns dict with:
      mu_rna, mu_atac,
      mu_moe: STRICT fused posterior mean if available (prefers mu_z / mu_moe / mu_joint),
      mu_fused: model-provided fused latent (via _extract_latent(enc_full,"fused")),
      moe_gating: (n_cells, n_modalities) precision-derived weights if logvar_dict is present,
      moe_gating_mods: list[str] naming the columns of moe_gating,
      y / gids / id_to_int,
      moe_status
    """
    model.eval()

    y_raw, global_map = _build_label_encoder(y_all)
    kind = np.asarray(y_raw).dtype.kind
    use_map = id_to_int if id_to_int is not None else (global_map if kind not in ("i", "u", "f", "b") else None)

    mu_rna_all, mu_atac_all, mu_moe_all, mu_fused_all = [], [], [], []
    moe_gating_all = []
    y_list, gids_list = [], []

    saw_gating = False
    saw_explicit_moe = False
    gating_shapes = set()

    # store a single consistent column order for gating
    gating_mods = None

    for b_ix, (x_dict, gids) in enumerate(loader):
        gids_np = _to_numpy_ids(gids)
        if gids_np is None:
            raise RuntimeError("Loader must provide gids for label lookup.")

        # move tensors to device
        x_full = {}
        for k, v in x_dict.items():
            if v is None:
                continue
            x_full[k] = v.to(device) if torch.is_tensor(v) else v

        # full encode (with whatever modalities are present)
        enc_full = _unwrap_model_output(_call_univi_encoder(model, x_full))

        # unimodal encodes (for alignment / plots)
        enc_rna = None
        if "rna" in x_full and torch.is_tensor(x_full["rna"]):
            enc_rna = _unwrap_model_output(_call_univi_encoder(model, {"rna": x_full["rna"]}))
        enc_atac = None
        if "atac" in x_full and torch.is_tensor(x_full["atac"]):
            enc_atac = _unwrap_model_output(_call_univi_encoder(model, {"atac": x_full["atac"]}))

        # ---- gating weights (precision-derived from logvar_dict) ----
        W, mods = _extract_gating_weights_from_enc_full(enc_full)
        if torch.is_tensor(W):
            saw_gating = True
            gating_shapes.add(tuple(W.shape))
            moe_gating_all.append(W.detach().cpu())

            if gating_mods is None:
                gating_mods = list(mods)
            else:
                # if it changes mid-run, something is wrong; fail loud
                if list(mods) != list(gating_mods):
                    raise RuntimeError(
                        f"moe_gating_mods changed across batches.\n"
                        f"first={gating_mods}\nnow={list(mods)}"
                    )

        if debug_first_batch and b_ix == 0:
            def _keys(obj):
                return sorted(obj.keys()) if isinstance(obj, dict) else [str(type(obj))]
            print("[latent-debug] enc_full keys:", _keys(enc_full))
            print("[latent-debug] gating found:", torch.is_tensor(W),
                  ("shape=" + str(tuple(W.shape)) if torch.is_tensor(W) else ""),
                  ("mods=" + str(mods) if mods is not None else ""))

        # ---- extract unimodal means ----
        mu_rna = _extract_latent(enc_rna, "rna")
        if mu_rna is None:
            mu_rna = _extract_latent(enc_rna, "fused")

        mu_atac = _extract_latent(enc_atac, "atac")
        if mu_atac is None:
            mu_atac = _extract_latent(enc_atac, "fused")

        # model-provided fused latent (not avg fallback)
        mu_fused_from_model = _extract_latent(enc_full, "fused")

        # ---- STRICT fused posterior mean ("MoE latent") ----
        mu_moe = _pick_explicit_moe_tensor(enc_full)
        if torch.is_tensor(mu_moe):
            saw_explicit_moe = True
        else:
            mu_moe = None

        # ---- accumulate latents ----
        if torch.is_tensor(mu_rna):
            mu_rna_all.append(mu_rna.detach().cpu())
        if torch.is_tensor(mu_atac):
            mu_atac_all.append(mu_atac.detach().cpu())
        if torch.is_tensor(mu_moe):
            mu_moe_all.append(mu_moe.detach().cpu())
        if torch.is_tensor(mu_fused_from_model):
            mu_fused_all.append(mu_fused_from_model.detach().cpu())

        # ---- labels ----
        y_tensor, use_map = _encode_labels_for_gids(y_raw, gids_np, id_to_int=use_map)
        y_list.append(y_tensor.detach().cpu())
        gids_list.append(torch.as_tensor(gids_np, dtype=torch.long))

    def _cat(xs):
        return torch.cat(xs, dim=0) if len(xs) else None

    mu_moe_cat = _cat(mu_moe_all)
    mu_fused_cat = _cat(mu_fused_all)
    moe_gating_cat = _cat(moe_gating_all)

    enc_out = {
        "mu_rna": _cat(mu_rna_all),
        "mu_atac": _cat(mu_atac_all),

        "mu_moe": mu_moe_cat,
        "mu_fused": mu_fused_cat,

        # NEW: gating matrix + column order
        "moe_gating": moe_gating_cat,
        "moe_gating_mods": (list(gating_mods) if gating_mods is not None else None),

        "y": _cat(y_list),
        "y_fused": _cat(y_list),
        "gids": _cat(gids_list),
        "id_to_int": use_map,
        "moe_status": {
            "saw_gating": bool(saw_gating),
            "gating_shapes": sorted(list(gating_shapes)),
            "moe_gating_mods": (list(gating_mods) if gating_mods is not None else None),
            "saw_explicit_moe": bool(saw_explicit_moe),
            "has_mu_moe": (mu_moe_cat is not None),
            "has_mu_fused": (mu_fused_cat is not None),
            "has_moe_gating": (moe_gating_cat is not None),
        },
    }
    return enc_out

def report_moe_status(enc, *, prefix="[moe_status]"):
    st = (enc or {}).get("moe_status", {})
    print(
        prefix,
        "saw_gating:", st.get("saw_gating", False),
        "| gating_shapes:", st.get("gating_shapes", []),
        "| moe_gating_mods:", st.get("moe_gating_mods", None),
        "| saw_explicit_moe:", st.get("saw_explicit_moe", False),
        "| has_mu_moe:", st.get("has_mu_moe", False),
        "| has_mu_fused:", st.get("has_mu_fused", False),
        "| has_moe_gating:", st.get("has_moe_gating", False),
    )

def compute_moe_latent_strict(enc: dict) -> np.ndarray:
    """
    STRICT fused latent accessor.

    Accepts either:
      - the batch encoder output dict containing mu_z / mu_fused / z, OR
      - the enc_out dict returned by encode_embeddings_with_labels (contains mu_moe / mu_fused).

    Preference order:
      1) mu_moe
      2) mu_z / mu_fused
      3) z
    """
    if enc is None or not isinstance(enc, dict):
        raise RuntimeError("enc must be a dict")

    for k in ("mu_moe", "mu_z", "mu_fused"):
        if enc.get(k, None) is not None:
            X = _as_numpy_2d(enc[k])
            if X is not None:
                return X

    if enc.get("z", None) is not None:
        X = _as_numpy_2d(enc["z"])
        if X is not None:
            return X

    st = (enc or {}).get("moe_status", {})
    raise RuntimeError(
        "Requested STRICT fused latent but none was found.\n"
        "Looked for keys: mu_moe, mu_z, mu_fused, z.\n"
        f"moe_status={st}\n"
    )

def compute_fused_latent_model(enc: dict) -> np.ndarray:
    """
    Returns model-provided fused latent (mu_fused). No avg fallback.
    """
    X = _as_numpy_2d(enc.get("mu_fused", None)) if isinstance(enc, dict) else None
    if X is None:
        raise RuntimeError("Requested model fused latent but enc['mu_fused'] is missing.")
    return X


# %% [code]
# =============================================================================
# ---- 10) kNN ACC + silhouette helper ----
# =============================================================================
def knn_acc_from_reference(X_ref, y_ref, X_query, y_query, *, k: int = 3):
    X_ref = _as_numpy_2d(X_ref)
    X_query = _as_numpy_2d(X_query)
    y_ref = _as_numpy_1d(y_ref)
    y_query = _as_numpy_1d(y_query)
    if X_ref is None or X_query is None or y_ref is None or y_query is None:
        return {"acc": np.nan, "macro_f1": np.nan}
    if X_ref.shape[0] < 2 or X_query.shape[0] < 2:
        return {"acc": np.nan, "macro_f1": np.nan}
    k_eff = int(min(max(1, k), X_ref.shape[0]))
    clf = KNeighborsClassifier(n_neighbors=k_eff, weights="distance", metric="euclidean")
    clf.fit(X_ref, y_ref)
    y_hat = clf.predict(X_query)
    return {"acc": float(accuracy_score(y_query, y_hat)), "macro_f1": float(f1_score(y_query, y_hat, average="macro"))}

def safe_silhouette_by_true_labels(X, y_true, *, min_per_class=2, min_total=10):
    X = _as_numpy_2d(X)
    y = _as_numpy_1d(y_true)
    if X is None or y is None:
        return np.nan, 0, 0.0
    uniq, counts = np.unique(y, return_counts=True)
    keep = set(uniq[counts >= int(min_per_class)].tolist())
    if len(keep) < 2:
        return np.nan, 0, 0.0
    mask = np.array([lab in keep for lab in y], dtype=bool)
    n_used = int(mask.sum())
    if n_used < int(min_total):
        return np.nan, n_used, float(n_used / max(len(y), 1))
    Xk = X[mask]
    yk = y[mask]
    uniq2, counts2 = np.unique(yk, return_counts=True)
    if len(uniq2) < 2 or np.any(counts2 < 2):
        return np.nan, n_used, float(n_used / max(len(y), 1))
    try:
        return float(silhouette_score(Xk, yk)), n_used, float(n_used / max(len(y), 1))
    except Exception:
        return np.nan, n_used, float(n_used / max(len(y), 1))
        

# %% [code]
# =============================================================================
# ---- 11) UMAP plotting (SELF-CONTAINED, TWO COMPOSITE FIGURES ONLY) ----
#   Outputs ONLY:
#     1) <base>__spaces_celltype.png   (3 cols: RNA / ATAC / MoE, colored by cell type)
#     2) <base>__summary_4col.png      (4 cols: overlap mod / overlap ct / MoE gradient / MoE ct)
# =============================================================================
import os
import numpy as np
import matplotlib.pyplot as plt
from matplotlib.lines import Line2D
from matplotlib.colors import TwoSlopeNorm

# -----------------------------------------------------------------------------
# Minimal numeric helpers (self-contained)
# -----------------------------------------------------------------------------
def _as_numpy_1d(x):
    if x is None:
        return None
    try:
        import torch
        if torch.is_tensor(x):
            x = x.detach().cpu().numpy()
    except Exception:
        pass
    x = np.asarray(x)
    return x.reshape(-1)

def _as_numpy_2d(x):
    if x is None:
        return None
    try:
        import torch
        if torch.is_tensor(x):
            x = x.detach().cpu().numpy()
    except Exception:
        pass
    x = np.asarray(x)
    if x.ndim == 1:
        x = x[:, None]
    return x.astype(np.float32, copy=False)

def _get_celltype_labels_from_enc(enc, y_all):
    """
    Returns per-row cell-type string labels for the *paired* cells.
    Priority:
      1) enc['gids'] indexes into y_all
      2) enc['y_fused'] or enc['y'] already stores labels
    """
    if enc is None or not isinstance(enc, dict):
        return None

    gids = enc.get("gids", None)
    if gids is not None:
        gids = _as_numpy_1d(gids)
        if gids is None:
            return None
        gids = gids.astype(np.int64, copy=False)
        labs = np.asarray(y_all)[gids]
        return np.asarray(labs).astype(str)

    y = enc.get("y_fused", enc.get("y", None))
    if y is None:
        return None
    y = _as_numpy_1d(y)
    return None if y is None else np.asarray(y).astype(str)

# -----------------------------------------------------------------------------
# UMAP embedding (with PCA fallback)
# -----------------------------------------------------------------------------
def _umap_embed(X, *, seed=0, n_neighbors=15, min_dist=0.3, metric="euclidean"):
    X = _as_numpy_2d(X)
    if X is None:
        return None
    try:
        import umap
        reducer = umap.UMAP(
            n_neighbors=int(n_neighbors),
            min_dist=float(min_dist),
            metric=str(metric),
            random_state=int(seed),
        )
        return reducer.fit_transform(X)
    except Exception:
        from sklearn.decomposition import PCA
        return PCA(n_components=2, random_state=int(seed)).fit_transform(X)

# -----------------------------------------------------------------------------
# Color/legend helpers
# -----------------------------------------------------------------------------
def _make_color_codes(labels, *, sort_legend_by="count", cmap_name="tab20"):
    if labels is None:
        return None, None, None, None, None
    labels = np.asarray(labels).astype(str)

    uniq, counts = np.unique(labels, return_counts=True)
    if str(sort_legend_by).lower() == "count":
        order = np.argsort(-counts)
    else:
        order = np.argsort(uniq.astype(str))
    uniq = uniq[order]
    counts = counts[order]

    lab_to_int = {lab: i for i, lab in enumerate(uniq.tolist())}
    c = np.array([lab_to_int[lab] for lab in labels], dtype=int)
    cmap = plt.get_cmap(str(cmap_name), max(len(uniq), 1))
    return c, uniq, counts, cmap, lab_to_int

def _legend_handles_from_uniq(uniq, counts, cmap, *, legend_max=25):
    if uniq is None or counts is None or cmap is None:
        return None
    n_cat = len(uniq)
    if legend_max is None or int(legend_max) <= 0 or n_cat == 0:
        return None
    top_n = int(min(int(legend_max), n_cat))
    handles = []
    for i in range(top_n):
        lab = uniq[i]
        col = cmap(i)
        handles.append(
            Line2D([0], [0], marker="o", color="none",
                   markerfacecolor=col, markeredgecolor="none",
                   markersize=6, label=f"{lab} (n={counts[i]})")
        )
    return handles

def _plot_umap_scatter(
    ax,
    emb,
    *,
    color_codes=None,
    cmap=None,
    title="UMAP",
    point_size=8,
    alpha=0.75,
):
    if emb is None:
        ax.set_title(f"{title} (embed failed)")
        ax.axis("off")
        return None
    if color_codes is None or cmap is None:
        sc = ax.scatter(emb[:, 0], emb[:, 1], s=point_size, alpha=alpha, linewidths=0)
    else:
        sc = ax.scatter(emb[:, 0], emb[:, 1], c=color_codes, cmap=cmap,
                        s=point_size, alpha=alpha, linewidths=0)
    ax.set_title(title)
    ax.set_xlabel("UMAP1")
    ax.set_ylabel("UMAP2")
    return sc

# -----------------------------------------------------------------------------
# Overlap helper: stack RNA+ATAC
# -----------------------------------------------------------------------------
def _stack_for_overlap(enc):
    """
    Returns:
      X_stack: (2n, d) stacked [rna; atac]
      modality_labels: (2n,) ["rna"]*n + ["atac"]*n
    """
    if enc is None or not isinstance(enc, dict):
        return None, None
    Xr = _as_numpy_2d(enc.get("mu_rna", None))
    Xa = _as_numpy_2d(enc.get("mu_atac", None))
    if Xr is None or Xa is None:
        return None, None
    if Xr.shape[1] != Xa.shape[1]:
        raise ValueError(f"Overlap requires same dim: mu_rna {Xr.shape} vs mu_atac {Xa.shape}")
    n = int(min(Xr.shape[0], Xa.shape[0]))
    X = np.vstack([Xr[:n], Xa[:n]])
    mod = np.array(["rna"] * n + ["atac"] * n, dtype=object)
    return X, mod

# -----------------------------------------------------------------------------
# MoE gating gradient utilities (robust to missing mods)
# -----------------------------------------------------------------------------
def _get_moe_gating(enc):
    """
    Returns:
      G: (N, M) float32
      mods: list[str] length M (best-effort)
    or (None, None) if missing.
    """
    if not isinstance(enc, dict):
        return None, None

    G = _as_numpy_2d(enc.get("moe_gating", None))
    if G is None or G.ndim != 2 or G.shape[1] < 2:
        return None, None

    mods = enc.get("moe_gating_mods", None)
    if mods is None:
        # Best-effort default for common 2-modality case
        if G.shape[1] == 2:
            mods = ["rna", "atac"]
        else:
            mods = [f"mod{i}" for i in range(G.shape[1])]
    else:
        mods = [str(x) for x in list(mods)]
        if len(mods) < G.shape[1]:
            mods = mods + [f"mod{i}" for i in range(len(mods), G.shape[1])]

    return G.astype(np.float32, copy=False), mods

def _rna_atac_delta_from_gating(enc):
    """
    delta = w_rna - w_atac in [-1,1]
    Returns (delta, w_rna, w_atac) or (None,None,None) if unavailable.
    """
    G, mods = _get_moe_gating(enc)
    if G is None:
        return None, None, None

    if "rna" in mods and "atac" in mods:
        i_rna = mods.index("rna")
        i_atac = mods.index("atac")
    else:
        # fallback (only if you ever have unnamed mods)
        i_rna, i_atac = 0, 1

    w_rna = G[:, i_rna].astype(np.float32, copy=False)
    w_atac = G[:, i_atac].astype(np.float32, copy=False)

    s = (w_rna + w_atac).clip(1e-8)
    w_rna = w_rna / s
    w_atac = w_atac / s

    delta = (w_rna - w_atac).astype(np.float32, copy=False)
    delta = np.clip(delta, -1.0, 1.0)
    return delta, w_rna, w_atac

# -----------------------------------------------------------------------------
# Main saver: ONLY TWO FIGURES
# -----------------------------------------------------------------------------
def save_umaps_for_enc(
    enc,
    *,
    y_all,
    out_png,  # base path; we'll append suffixes
    seed=0,
    n_neighbors=15,
    min_dist=0.3,
    point_size=8,
    alpha=0.75,
    dpi=450,
    legend_max=25,
    sort_legend_by="count",
    overlap_metric: str = "euclidean",

    # --- NEW: make gradient visible ---
    gradient_cmap: str = "bwr",
    gradient_alpha: float = 0.95,     # higher opacity for gradient panel
    gradient_point_size: int = None,  # default: 1.2x point_size
    gradient_gain: float = 6.0,       # boost (w_rna - w_atac) before clipping
    gradient_vlim: float = 0.5,       # clip range; smaller => more saturated
):
    """
    Outputs ONLY:
      1) <base>__spaces_celltype.png   (3 cols: RNA / ATAC / MoE, colored by cell type)
      2) <base>__summary_4col.png      (4 cols: overlap mod / overlap ct / MoE gradient / MoE ct)
    """
    if enc is None or not isinstance(enc, dict):
        return

    out_dir = os.path.dirname(out_png)
    if out_dir:
        os.makedirs(out_dir, exist_ok=True)
    base = out_png[:-4] if out_png.lower().endswith(".png") else out_png

    # ---- latents ----
    X_rna  = _as_numpy_2d(enc.get("mu_rna", None))
    X_atac = _as_numpy_2d(enc.get("mu_atac", None))

    X_moe = enc.get("mu_moe", None)
    if X_moe is None:
        X_moe = enc.get("mu_fused", None)
    X_moe = _as_numpy_2d(X_moe)

    # ---- cell type labels ----
    labs_ct = _get_celltype_labels_from_enc(enc, y_all)
    if labs_ct is not None:
        c_ct, uniq_ct, counts_ct, cmap_ct, _ = _make_color_codes(
            labs_ct, sort_legend_by=sort_legend_by, cmap_name="tab20"
        )
        legend_ct = _legend_handles_from_uniq(uniq_ct, counts_ct, cmap_ct, legend_max=legend_max)
    else:
        c_ct, cmap_ct, legend_ct = None, None, None

    # =============================================================================
    # FIGURE 1: spaces (RNA / ATAC / MoE), colored by cell type
    # =============================================================================
    fig = plt.figure(figsize=(18, 5), dpi=int(dpi))
    axes = [fig.add_subplot(1, 3, i + 1) for i in range(3)]

    for ax, X, title in [
        (axes[0], X_rna,  "RNA latent (mu_rna)"),
        (axes[1], X_atac, "ATAC latent (mu_atac)"),
        (axes[2], X_moe,  "MoE fused latent (mu_moe)"),
    ]:
        if X is None:
            ax.set_title(f"{title} (missing)")
            ax.axis("off")
            continue
        emb = _umap_embed(X, seed=seed, n_neighbors=n_neighbors, min_dist=min_dist, metric="euclidean")
        _plot_umap_scatter(
            ax, emb,
            color_codes=c_ct, cmap=cmap_ct,
            title=title,
            point_size=point_size,
            alpha=alpha
        )

    if legend_ct is not None:
        fig.legend(
            handles=legend_ct, loc="center left", bbox_to_anchor=(1.01, 0.5),
            frameon=False, title=f"Cell types (top {min(int(legend_max), len(legend_ct))})"
        )

    fig.tight_layout()
    fig.savefig(base + "__spaces_celltype.png", dpi=int(dpi), bbox_inches="tight")
    plt.close(fig)

    # =============================================================================
    # FIGURE 2: summary 4 columns
    #   col1: overlap UMAP colored by modality
    #   col2: overlap UMAP colored by cell type
    #   col3: MoE UMAP colored by RNA<->ATAC gating gradient (delta=w_rna-w_atac)
    #   col4: MoE UMAP colored by cell type
    # =============================================================================
    X_overlap, overlap_mod_labels = _stack_for_overlap(enc)
    emb_overlap = None
    if X_overlap is not None:
        emb_overlap = _umap_embed(
            X_overlap,
            seed=seed,
            n_neighbors=n_neighbors,
            min_dist=min_dist,
            metric=str(overlap_metric),
        )

    c_ov_mod, cmap_ov_mod, legend_ov_mod = None, None, None
    if overlap_mod_labels is not None:
        c_ov_mod, uniq_ov_mod, counts_ov_mod, cmap_ov_mod, _ = _make_color_codes(
            overlap_mod_labels, sort_legend_by="alpha", cmap_name="Set1"
        )
        legend_ov_mod = _legend_handles_from_uniq(uniq_ov_mod, counts_ov_mod, cmap_ov_mod, legend_max=10)

    c_ov_ct, cmap_ov_ct = None, None
    if emb_overlap is not None and labs_ct is not None:
        labs_overlap_ct = np.concatenate([labs_ct, labs_ct], axis=0)
        c_ov_ct, _, _, cmap_ov_ct, _ = _make_color_codes(
            labs_overlap_ct, sort_legend_by=sort_legend_by, cmap_name="tab20"
        )

    emb_moe = None
    if X_moe is not None:
        emb_moe = _umap_embed(
            X_moe,
            seed=seed,
            n_neighbors=n_neighbors,
            min_dist=min_dist,
            metric="euclidean",
        )

    # gradient values for MoE panel
    delta, w_rna, w_atac = _rna_atac_delta_from_gating(enc)
    if emb_moe is not None and delta is not None:
        m = int(min(len(delta), emb_moe.shape[0]))
        delta = delta[:m]
        emb_moe_use = emb_moe[:m]
    else:
        emb_moe_use = emb_moe

    # --- visibility controls for gradient panel ---
    g_alpha = float(gradient_alpha)
    g_size = int(max(1, (int(point_size) if gradient_point_size is None else int(gradient_point_size))))
    if gradient_point_size is None:
        g_size = int(np.ceil(1.25 * g_size))

    # boost + clip to make colors visible even if delta is near 0
    vlim = float(gradient_vlim)
    gain = float(gradient_gain)

    fig = plt.figure(figsize=(28, 5), dpi=int(dpi))
    ax1 = fig.add_subplot(1, 4, 1)
    ax2 = fig.add_subplot(1, 4, 2)
    ax3 = fig.add_subplot(1, 4, 3)
    ax4 = fig.add_subplot(1, 4, 4)

    # col1: overlap by modality
    if emb_overlap is None:
        ax1.set_title("Overlap (RNA+ATAC stacked) — colored by modality (missing)")
        ax1.axis("off")
    else:
        _plot_umap_scatter(
            ax1, emb_overlap,
            color_codes=c_ov_mod, cmap=cmap_ov_mod,
            title="Overlap (RNA+ATAC stacked) — colored by modality",
            point_size=point_size, alpha=alpha
        )
        if legend_ov_mod is not None:
            ax1.legend(handles=legend_ov_mod, bbox_to_anchor=(1.02, 1), loc="upper left", frameon=False)

    # col2: overlap by cell type
    if emb_overlap is None:
        ax2.set_title("Overlap (RNA+ATAC stacked) — colored by cell type (missing)")
        ax2.axis("off")
    else:
        title2 = "Overlap (RNA+ATAC stacked) — colored by cell type"
        if labs_ct is None:
            title2 += " (labels missing)"
        _plot_umap_scatter(
            ax2, emb_overlap,
            color_codes=c_ov_ct if labs_ct is not None else None,
            cmap=cmap_ov_ct if labs_ct is not None else None,
            title=title2,
            point_size=point_size, alpha=alpha
        )

    # col3: MoE gating gradient (VISIBLE)
    if emb_moe_use is None:
        ax3.set_title("MoE fused — RNA↔ATAC gating gradient (missing)")
        ax3.axis("off")
    else:
        if delta is None:
            ax3.set_title("MoE fused — RNA↔ATAC gating gradient (gating missing)")
            ax3.scatter(emb_moe_use[:, 0], emb_moe_use[:, 1], s=g_size, alpha=max(g_alpha, 0.85), linewidths=0)
            ax3.set_xlabel("UMAP1"); ax3.set_ylabel("UMAP2")
        else:
            # boost + clip, then two-slope normalize around 0
            score = (delta.astype(np.float32, copy=False) * gain)
            score = np.clip(score, -vlim, vlim)
            norm = TwoSlopeNorm(vmin=-vlim, vcenter=0.0, vmax=vlim)

            sc3 = ax3.scatter(
                emb_moe_use[:, 0], emb_moe_use[:, 1],
                c=score,
                cmap=str(gradient_cmap),
                norm=norm,
                s=g_size,
                alpha=g_alpha,
                linewidths=0,
            )
            ax3.set_title("MoE fused — gating gradient (w_RNA - w_ATAC)")
            ax3.set_xlabel("UMAP1"); ax3.set_ylabel("UMAP2")
            cb = fig.colorbar(sc3, ax=ax3, fraction=0.046, pad=0.04)
            cb.set_label("w_RNA - w_ATAC  (red=RNA, blue=ATAC)")

    # col4: MoE by cell type
    if emb_moe_use is None:
        ax4.set_title("MoE fused — colored by cell type (missing)")
        ax4.axis("off")
    else:
        _plot_umap_scatter(
            ax4, emb_moe_use,
            color_codes=c_ct, cmap=cmap_ct,
            title="MoE fused — colored by cell type",
            point_size=point_size, alpha=alpha
        )

    if legend_ct is not None:
        fig.legend(
            handles=legend_ct, loc="center left", bbox_to_anchor=(1.01, 0.5),
            frameon=False, title=f"Cell types (top {min(int(legend_max), len(legend_ct))})"
        )

    fig.tight_layout()
    fig.savefig(base + "__summary_4col.png", dpi=int(dpi), bbox_inches="tight")
    plt.close(fig)


# %% [code]
# =============================================================================
# ---- 12) One ablation run (STRICT MoE only; no avg fallback) ----
# =============================================================================
def run_one_ablation(
    ablate_cell_type=None,
    drop_modality: str = "atac",
    *,
    seed: int = 0,
    fuse_mode: str = "moe",     # kept for API; "moe" is what we enforce
    return_enc: bool = False,
    debug_first_batch: bool = False,
    acc_knn_k: int = 3,

    save_umaps: bool = True,
    umap_dir: str = "results/fig9_ablation_analysis_results_MoE/figures",
    umap_seed: Optional[int] = None,
    umap_n_neighbors: int = 15,
    umap_min_dist: float = 0.3,
    umap_point_size: int = 6,
    umap_alpha: float = 0.75,
    umap_dpi: int = 450,
    umap_legend_max: int = 25,
    sort_legend_by: str = "count",

    print_moe_status: bool = True,
):
    baseline = (ablate_cell_type is None) or (str(ablate_cell_type).lower() in {"none", "__none__", "__baseline__", "baseline"})
    dummy = "__NO_SUCH_CELLTYPE__"

    train_loader, val_loader, test_loader = make_loaders_for_celltype_ablation(
        dataset, TRAIN_IDX, VAL_IDX, TEST_IDX,
        y_all=y_all,
        ablate_cell_type=(dummy if baseline else str(ablate_cell_type)),
        drop_modality=str(drop_modality),
        seed=int(seed),
        batch_size=int(BATCH_SIZE),
        disable_ablation=bool(baseline),
    )

    rna_dim  = int(adata_dict["rna"].X.shape[1])
    atac_dim = int(adata_dict["atac"].X.shape[1])
    dev = torch.device(str(DEVICE)) if isinstance(DEVICE, str) else DEVICE

    model = train_one_model(
        train_loader=train_loader,
        val_loader=val_loader,
        rna_dim=rna_dim,
        atac_dim=atac_dim,
        seed=int(seed),
        device=dev,
        loss_mode="v1",
        v1_recon="moe",          # IMPORTANT: train with moe recon
        best_epoch_warmup=50,
    )

    # ---- ENCODE (this is where you encode) ----
    enc_val = encode_embeddings_with_labels(
        model, val_loader, device=dev, y_all=y_all, id_to_int=None, debug_first_batch=bool(debug_first_batch),
    )
    enc_test = encode_embeddings_with_labels(
        model, test_loader, device=dev, y_all=y_all, id_to_int=enc_val.get("id_to_int", None), debug_first_batch=False,
    )

    if print_moe_status:
        ab_name = "NONE" if baseline else str(ablate_cell_type)
        print(f"[ablation] drop={drop_modality} ablate={ab_name} seed={seed}")
        report_moe_status(enc_test)

    # ---- STRICT: require MoE latent or crash ----
    if bool(STRICT_MOE):
        Z_val  = compute_moe_latent_strict(enc_val)
        Z_test = compute_moe_latent_strict(enc_test)
    else:
        # explicit non-avg fallback (model fused)
        Z_val  = compute_fused_latent_model(enc_val)
        Z_test = compute_fused_latent_model(enc_test)

    # ---- Save UMAPs (MoE only) ----
    if bool(save_umaps):
        u_seed = int(seed if umap_seed is None else umap_seed)
        ab_name = "NONE" if baseline else str(ablate_cell_type)
        ab_name_safe = ab_name.replace("/", "_").replace(" ", "_")
        drop_safe = str(drop_modality).replace("/", "_").replace(" ", "_")
        os.makedirs(umap_dir, exist_ok=True)

        out_png = os.path.join(
            umap_dir,
            f"drop-{drop_safe}__ablate-{ab_name_safe}__seed-{int(seed)}__MOE_latents.png"
        )
        # replace mu_fused plot with mu_moe plot
        enc_for_plot = dict(enc_test)
        # ensure mu_moe exists if strict
        enc_for_plot["mu_moe"] = enc_test.get("mu_moe", None)

        save_umaps_for_enc(
            enc_for_plot,
            y_all=y_all,
            out_png=out_png,
            seed=u_seed,
            n_neighbors=int(umap_n_neighbors),
            min_dist=float(umap_min_dist),
            point_size=int(umap_point_size),
            alpha=float(umap_alpha),
            dpi=int(umap_dpi),
            legend_max=int(umap_legend_max),
            sort_legend_by=str(sort_legend_by),
        )

    # ---- Metrics ----
    mu_rna_test  = enc_test.get("mu_rna", None)
    mu_atac_test = enc_test.get("mu_atac", None)

    y_val  = enc_val.get("y_fused", enc_val.get("y", None))
    y_test = enc_test.get("y_fused", enc_test.get("y", None))

    row = {
        "drop_modality": str(drop_modality),
        "ablated_cell_type": ("NONE" if baseline else str(ablate_cell_type)),
        "fuse_mode": "moe_strict" if STRICT_MOE else "model_fused",
        "ACC_fused_mode": f"knn_val_to_test_k{int(acc_knn_k)}",
    }

    # Alignment metrics on paired TEST (still based on mu_rna/mu_atac)
    if mu_rna_test is None or mu_atac_test is None:
        row["FOSCTTM"] = np.nan
        for k in RECALL_KS:
            row[f"Recall@{k}"] = np.nan
    else:
        row["FOSCTTM"] = float(foscttm(mu_rna_test, mu_atac_test))
        for k, v in recall_at_k(mu_rna_test, mu_atac_test, ks=RECALL_KS).items():
            row[f"Recall@{k}"] = float(v)

    # Fused (MoE) metrics
    if Z_test is None or y_test is None:
        row.update({
            "ARI_fused": np.nan,
            "NMI_fused": np.nan,
            "ACC_fused": np.nan,
            "ACC_fused_macroF1": np.nan,
            "SIL_fused_celltype": np.nan,
            "SIL_fused_n": 0,
            "SIL_fused_frac": 0.0,
        })
    else:
        X_test = _as_numpy_2d(Z_test)
        y_test_np = _as_numpy_1d(y_test)

        # KMeans ARI/NMI
        try:
            n_clusters = int(len(np.unique(y_test_np)))
            if n_clusters >= 2 and X_test.shape[0] >= n_clusters:
                km = KMeans(n_clusters=n_clusters, random_state=int(seed), n_init=20)
                y_pred = km.fit_predict(X_test)
                row["ARI_fused"] = float(adjusted_rand_score(y_test_np, y_pred))
                row["NMI_fused"] = float(normalized_mutual_info_score(y_test_np, y_pred))
            else:
                row["ARI_fused"] = np.nan
                row["NMI_fused"] = np.nan
        except Exception:
            row["ARI_fused"] = np.nan
            row["NMI_fused"] = np.nan

        # kNN ACC (val -> test)
        knn_out = {"acc": np.nan, "macro_f1": np.nan}
        if Z_val is not None and y_val is not None:
            try:
                knn_out = knn_acc_from_reference(Z_val, y_val, Z_test, y_test, k=int(acc_knn_k))
            except Exception:
                knn_out = {"acc": np.nan, "macro_f1": np.nan}

        row["ACC_fused"] = float(knn_out.get("acc", np.nan))
        row["ACC_fused_macroF1"] = float(knn_out.get("macro_f1", np.nan))

        sil, sil_n, sil_frac = safe_silhouette_by_true_labels(X_test, y_test_np, min_per_class=2, min_total=10)
        row["SIL_fused_celltype"] = float(sil)
        row["SIL_fused_n"] = int(sil_n)
        row["SIL_fused_frac"] = float(sil_frac)

    if return_enc:
        return row, {"enc_val": enc_val, "enc_test": enc_test, "model": model}
    return row


# %% [code]
# =============================================================================
# ---- 13) Ablation grid runner ----
# =============================================================================
def run_ablation_grid(
    *,
    drop_modalities=("rna", "atac"),
    seed: int = 0,
    save_umaps: bool = True,
    umap_dir: str = "results/fig9_ablation_analysis_results_MoE/figures",
):
    cell_types = sorted(pd.unique(y_all))
    rows = []
    for dm in drop_modalities:
        for ct in cell_types:
            print(f"[grid] drop_modality={dm:>4s}  ablate_cell_type={ct}")
            row, _out = run_one_ablation(
                ct, dm,
                seed=int(seed),
                return_enc=bool(save_umaps),
                save_umaps=bool(save_umaps),
                umap_dir=str(umap_dir),
            )
            rows.append(row)
    return pd.DataFrame(rows)


# %% [code]
# =============================================================================
# ---- 14) Example sanity checks ----
# =============================================================================
# Baseline (should PASS only if enc['mu_moe'] exists; otherwise it will crash loudly)
row_base, out_base = run_one_ablation(None, "atac", seed=SEED, return_enc=True, debug_first_batch=True, save_umaps=True)
print(row_base)


# %% [code]
# Smoke test: ablate the first cell type
cell_types = sorted(pd.unique(y_all))
ct = cell_types[0]
print(f"Performing a smoke-test by ablating {ct} from atac.")

row_smoke, out_smoke = run_one_ablation(ct, "atac", seed=SEED, return_enc=True, save_umaps=False)
print(row_smoke)


# %% [code]
# =============================================================================
# ---- 15) Run grid + save ----
# =============================================================================
df_grid = run_ablation_grid(
    drop_modalities=("rna", "atac"),
    seed=SEED,
    save_umaps=True,
    umap_dir="results/fig9_ablation_analysis_results_MoE/figures",
)
print(df_grid.head())

OUTDIR = "./results/fig9_ablation_analysis_results_MoE"
os.makedirs(OUTDIR, exist_ok=True)
csv_path = os.path.join(OUTDIR, "celltype_ablation_grid_metrics.csv")
df_grid.to_csv(csv_path, index=False)
print("Wrote:", csv_path)


# %% [code]
# =============================================================================
# ---- 16) REST OF ANALYSIS: heatmaps + coarse labels + saving ----
# =============================================================================
def plot_metric_heatmap(
    df: pd.DataFrame,
    metric: str,
    drop_modality: str,
    *,
    title_prefix: str = "",
    sort_by: str = "ablated_cell_type",
    dpi: int = 250,
):
    sub = df[df["drop_modality"] == drop_modality].copy()
    if sub.empty:
        print(f"[skip] no rows for drop_modality={drop_modality}")
        return
    if metric not in sub.columns:
        print(f"[skip] metric {metric} not in df")
        return

    if sort_by in sub.columns:
        sub[sort_by] = sub[sort_by].astype(str)
        sub = sub.sort_values(sort_by)

    x = sub["ablated_cell_type"].astype(str).to_list()
    vals = pd.to_numeric(sub[metric], errors="coerce").to_numpy(dtype=float)
    M = vals[None, :]

    plt.figure(figsize=(max(10, 0.45 * len(x)), 2.4), dpi=int(dpi))
    im = plt.imshow(M, aspect="auto")
    plt.yticks([0], [metric])
    plt.xticks(np.arange(len(x)), x, rotation=90)
    plt.colorbar(im, fraction=0.03, pad=0.02)

    prefix = (title_prefix + " — ") if title_prefix else ""
    plt.title(f"{prefix}{metric} — ablate {drop_modality} in TRAIN (by cell type)")
    plt.tight_layout()
    plt.show()

def save_metric_heatmap(
    df: pd.DataFrame,
    metric: str,
    drop_modality: str,
    *,
    title_prefix: str = "",
    outdir: str = OUTDIR,
    dpi: int = 250,
):
    sub = df[df["drop_modality"] == drop_modality].copy()
    if sub.empty or metric not in sub.columns:
        return None

    sub["ablated_cell_type"] = sub["ablated_cell_type"].astype(str)
    sub = sub.sort_values("ablated_cell_type")

    x = sub["ablated_cell_type"].to_list()
    vals = pd.to_numeric(sub[metric], errors="coerce").to_numpy(dtype=float)
    M = vals[None, :]

    plt.figure(figsize=(max(10, 0.45 * len(x)), 2.4), dpi=int(dpi))
    im = plt.imshow(M, aspect="auto")
    plt.yticks([0], [metric])
    plt.xticks(np.arange(len(x)), x, rotation=90)
    plt.colorbar(im, fraction=0.03, pad=0.02)

    prefix = (title_prefix + " — ") if title_prefix else ""
    plt.title(f"{prefix}{metric} — ablate {drop_modality} in TRAIN (by cell type)")
    plt.tight_layout()

    fname = f"heatmap_{title_prefix}_{drop_modality}_{metric}".replace(" ", "_").replace("@", "at").replace("/", "_")
    path = os.path.join(outdir, fname + ".png")
    plt.savefig(path, bbox_inches="tight")
    plt.close()
    return path

def plot_big_heatmap_shared_scale(
    df: pd.DataFrame,
    metrics,
    drop_modality: str,
    *,
    title: str = "",
    sort_by: str = "ablated_cell_type",
    vmin: float = -0.1,
    vmax: float = 0.9,
    cmap: str = "viridis",
    show_nan_as: Optional[float] = None,
    dpi: int = 200,
):
    sub = df[df["drop_modality"] == drop_modality].copy()
    if sub.empty:
        print(f"[skip] no rows for drop_modality={drop_modality}")
        return

    if sort_by in sub.columns:
        sub[sort_by] = sub[sort_by].astype(str)
        sub = sub.sort_values(sort_by)

    xlabels = sub["ablated_cell_type"].astype(str).to_list()
    metrics_use = [m for m in metrics if m in sub.columns]
    if not metrics_use:
        print("[skip] none of the requested metrics exist in df columns")
        return

    M = np.full((len(metrics_use), len(xlabels)), np.nan, dtype=float)
    for i, m in enumerate(metrics_use):
        M[i, :] = pd.to_numeric(sub[m], errors="coerce").to_numpy(dtype=float)

    if show_nan_as is not None:
        M = np.where(np.isfinite(M), M, float(show_nan_as))

    fig_w = max(10, 0.30 * len(xlabels))
    fig_h = max(4, 0.50 * len(metrics_use))
    plt.figure(figsize=(fig_w, fig_h), dpi=int(dpi))

    im = plt.imshow(M, aspect="auto", vmin=vmin, vmax=vmax, cmap=cmap)
    plt.colorbar(im, label=f"value (shared scale [{vmin}, {vmax}])")

    plt.yticks(np.arange(len(metrics_use)), metrics_use)
    plt.xticks(np.arange(len(xlabels)), xlabels, rotation=90)

    plt.title(title or f"BIG heatmap — drop {drop_modality} — shared scale [{vmin}, {vmax}]")
    plt.tight_layout()
    plt.show()

METRICS_TO_PLOT = [
    "FOSCTTM",
    "Recall@1", "Recall@10", "Recall@25", "Recall@50", "Recall@100",
    "ARI_fused", "NMI_fused", "ACC_fused", "SIL_fused_celltype",
]
'''
for dm in ("rna", "atac"):
    for m in METRICS_TO_PLOT:
        if m in df_grid.columns:
            plot_metric_heatmap(df_grid, m, dm, title_prefix="FINE")

for dm in ("rna", "atac"):
    for m in METRICS_TO_PLOT:
        p = save_metric_heatmap(df_grid, m, dm, title_prefix="FINE")
        if p is not None:
            print("Saved:", p)
'''
for dm in ("rna", "atac"):
    plot_big_heatmap_shared_scale(
        df_grid, METRICS_TO_PLOT, drop_modality=dm,
        title=f"FINE — drop {dm} — shared scale [-0.05, 1.0]",
        vmin=-0.05, vmax=1.0
    )


# %% [code]
# --- coarse labels ---
COARSE_MAP = {
    "CD14 Mono": "Monocyte",
    "CD16 Mono": "Monocyte",
    "CD4 Naive": "CD4 T",
    "CD4 TCM":   "CD4 T",
    "CD4 TEM":   "CD4 T",
    "CD8 Naive": "CD8 T",
    "CD8 TEM_1": "CD8 T",
    "CD8 TEM_2": "CD8 T",
    "Treg":      "Other T",
    "MAIT":      "Other T",
    "gdT":       "Other T",
    "NK": "NK",
    "Naive B":         "B",
    "Memory B":        "B",
    "Intermediate B":  "B",
    "cDC": "DC",
    "pDC": "DC",
    "HSPC":  "HSPC",
    "Plasma":"Plasma",
}

def collapse_celltypes(y_labels, mapping=COARSE_MAP, unknown="Other"):
    y = np.asarray(y_labels).astype(str)
    return np.array([mapping.get(lbl, unknown) for lbl in y], dtype=object)

y_all_coarse = collapse_celltypes(y_all)
print("fine unique:", len(pd.unique(y_all)))
print("coarse unique:", len(pd.unique(y_all_coarse)))
print(pd.Series(y_all_coarse).value_counts())

def run_ablation_grid_with_labels(
    y_labels,
    *,
    drop_modalities=("rna", "atac"),
    seed: int = 0,
):
    global y_all
    y_all_orig = y_all
    try:
        y_all = np.asarray(y_labels)
        cell_types = sorted(pd.unique(y_all))
        rows = []
        for dm in drop_modalities:
            for ct in cell_types:
                print(f"[grid] labels=COARSE  drop_modality={dm:>4s}  ablate_cell_type={ct}")
                rows.append(run_one_ablation(ct, dm, seed=seed, acc_knn_k=3))
        return pd.DataFrame(rows)
    finally:
        y_all = y_all_orig

df_grid_coarse = run_ablation_grid_with_labels(y_all_coarse, drop_modalities=("rna", "atac"), seed=SEED)
coarse_csv = os.path.join(OUTDIR, "celltype_ablation_grid_metrics_COARSE.csv")
df_grid_coarse.to_csv(coarse_csv, index=False)
print("Wrote:", coarse_csv)
'''
for dm in ("rna", "atac"):
    for m in METRICS_TO_PLOT:
        if m in df_grid_coarse.columns:
            plot_metric_heatmap(df_grid_coarse, m, dm, title_prefix="COARSE")

for dm in ("rna", "atac"):
    for m in METRICS_TO_PLOT:
        p = save_metric_heatmap(df_grid_coarse, m, dm, title_prefix="COARSE")
        if p is not None:
            print("Saved:", p)
'''
for dm in ("rna", "atac"):
    plot_big_heatmap_shared_scale(
        df_grid_coarse, METRICS_TO_PLOT, drop_modality=dm,
        title=f"COARSE — drop {dm} — shared scale [-0.5, 1.0]",
        vmin=-0.05, vmax=1.0
    )

print("Done ✅")


# %% [code]


# %% [code]


# %% [code]


# %% [code]


# %% [code]


# %% [code]


# %% [code]


# %% [code]


# %% [code]


# %% [code]


# %% [code]


# %% [code]


# %% [code]


# %% [code]


# %% [code]


# %% [code]


# %% [code]


# %% [code]


# %% [code]


# %% [code]


# %% [code]


# %% [code]


# %% [code]


# %% [code]


# %% [code]


# %% [code]


# %% [code]


# %% [code]


# %% [code]


# %% [code]


# %% [code]


# %% [code]


# %% [code]


# %% [code]


# %% [code]


# %% [code]


# %% [code]


# %% [code]


# %% [code]


# %% [code]


# %% [code]


# %% [code]


# %% [code]


# %% [code]


# %% [code]


# %% [code]


# %% [code]


# %% [code]
```
