## Supplemental Code for "Unifying multimodal single-cell data with a mixture-of-experts *β*-variational autoencoder framework": Fig10_population_compile.ipynb

```
# %% [code]
import numpy as np
import pandas as pd
import matplotlib.pyplot as plt

PATH_DROP_ATAC = "./results/figure_9_paired_ablation_outputs-2-10-2026/fig8_curve__drop_atac__all_seeds.tsv"
PATH_DROP_RNA  = "./results/figure_9_paired_ablation_outputs-2-10-2026/fig8_curve__drop_rna__all_seeds.tsv"

PATH_META_ATAC = "./results/figure_9_paired_ablation_outputs-2-10-2026/fig8_curve__drop_atac__all_seeds__meta.tsv"
PATH_META_RNA  = "./results/figure_9_paired_ablation_outputs-2-10-2026/fig8_curve__drop_rna__all_seeds__meta.tsv"


# %% [code]
drop_atac = pd.read_csv(PATH_DROP_ATAC, sep="\t")
drop_rna  = pd.read_csv(PATH_DROP_RNA,  sep="\t")

meta_atac = pd.read_csv(PATH_META_ATAC, sep="\t")
meta_rna  = pd.read_csv(PATH_META_RNA,  sep="\t")

print(drop_atac.shape, drop_rna.shape)
print("overlap_fraction unique (drop_atac):", drop_atac["overlap_fraction"].nunique())
print("overlap_fraction unique (drop_rna): ", drop_rna["overlap_fraction"].nunique())
print("splits (drop_atac):", drop_atac["split_seed"].nunique(), "model_seeds:", drop_atac["model_seed"].nunique())
print("splits (drop_rna): ", drop_rna["split_seed"].nunique(),  "model_seeds:", drop_rna["model_seed"].nunique())


# %% [code]
def add_derived(df: pd.DataFrame) -> pd.DataFrame:
    df = df.copy()
    df["LT_bidir_macroF1"] = 0.5 * (df["LT_ATAC2RNA_macroF1"] + df["LT_RNA2ATAC_macroF1"])
    return df

drop_atac = add_derived(drop_atac)
drop_rna  = add_derived(drop_rna)


# %% [code]
def summarize_overlaps(df: pd.DataFrame, metric: str) -> pd.DataFrame:
    g = df.groupby("overlap_fraction")[metric]
    out = g.agg(["mean", "std", "count"]).reset_index()
    out["sem"] = out["std"] / np.sqrt(out["count"])
    out["ci95"] = 1.96 * out["sem"]
    return out.sort_values("overlap_fraction")

def prep_summary_tables(drop_atac: pd.DataFrame, drop_rna: pd.DataFrame, metric: str):
    s_atac = summarize_overlaps(drop_atac, metric)
    s_rna  = summarize_overlaps(drop_rna,  metric)
    return s_atac, s_rna


# %% [code]
def plot_metric_vs_overlap(
    ax,
    df_atac, df_rna,
    metric: str,
    y_label: str,
    title: str,
    logx: bool = False,
    show_raw: bool = True,
    alpha_raw: float = 0.08,
):
    # raw traces (optional): one line per (split_seed, model_seed)
    if show_raw:
        for (ss, ms), d in df_atac.groupby(["split_seed", "model_seed"]):
            d = d.sort_values("overlap_fraction")
            ax.plot(d["overlap_fraction"], d[metric], linewidth=1, alpha=alpha_raw)
        for (ss, ms), d in df_rna.groupby(["split_seed", "model_seed"]):
            d = d.sort_values("overlap_fraction")
            ax.plot(d["overlap_fraction"], d[metric], linewidth=1, alpha=alpha_raw)

    s_atac, s_rna = prep_summary_tables(df_atac, df_rna, metric)

    # mean + CI (two bold curves)
    ax.plot(s_atac["overlap_fraction"], s_atac["mean"], linewidth=2.5, label="Drop ATAC")
    ax.fill_between(
        s_atac["overlap_fraction"],
        s_atac["mean"] - s_atac["ci95"],
        s_atac["mean"] + s_atac["ci95"],
        alpha=0.2,
        linewidth=0,
    )

    ax.plot(s_rna["overlap_fraction"], s_rna["mean"], linewidth=2.5, label="Drop RNA")
    ax.fill_between(
        s_rna["overlap_fraction"],
        s_rna["mean"] - s_rna["ci95"],
        s_rna["mean"] + s_rna["ci95"],
        alpha=0.2,
        linewidth=0,
    )

    ax.set_title(title)
    ax.set_xlabel("Overlap fraction")
    ax.set_ylabel(y_label)
    ax.grid(True, alpha=0.3)

    if logx:
        ax.set_xscale("log")
        # avoid weird limits if overlap includes 0 (it shouldn't)
        xmin = min(df_atac["overlap_fraction"].min(), df_rna["overlap_fraction"].min())
        ax.set_xlim(left=max(xmin, 1e-4))

    ax.legend(frameon=False)


# %% [code]
import matplotlib as mpl

mpl.rcParams['figure.dpi'] =  400     # display DPI in notebook
mpl.rcParams['savefig.dpi'] = 400     # when you do plt.savefig(...)


# %% [code]
fig, ax = plt.subplots(figsize=(10, 8))
plot_metric_vs_overlap(
    ax,
    drop_atac, drop_rna,
    metric="FOSCTTM",
    y_label="FOSCTTM (lower is better)",
    title="(a) Paired correspondence vs overlap",
    #logx=True,          # overlap spans tiny values; log-x usually looks nicer
    logx=False,
    show_raw=True,
)
plt.tight_layout()
plt.show()


# %% [code]
fig, ax = plt.subplots(figsize=(10, 8))
plot_metric_vs_overlap(
    ax,
    drop_atac, drop_rna,
    metric="Recall@1",
    y_label="Recall@1 (higher is better)",
    title="(b1) Paired retrieval (Recall@1) vs overlap",
    #logx=True,
    logx=False,
    show_raw=True,
)
plt.tight_layout()
plt.show()

fig, ax = plt.subplots(figsize=(10, 8))
plot_metric_vs_overlap(
    ax,
    drop_atac, drop_rna,
    metric="Recall@10",
    y_label="Recall@10 (higher is better)",
    title="(b2) Paired retrieval (Recall@10) vs overlap",
    #logx=True,
    logx=False,
    show_raw=True,
)
plt.tight_layout()
plt.show()


# %% [code]
fig, ax = plt.subplots(figsize=(10, 8))
plot_metric_vs_overlap(
    ax,
    drop_atac, drop_rna,
    metric="LT_bidir_macroF1",
    y_label="Bidirectional kNN label transfer (macro-F1)",
    title="(c) Cross-modality semantic consistency vs overlap",
    #logx=True,
    logx=False,
    show_raw=True,
)
plt.tight_layout()
plt.show()


# %% [code]
fig, ax = plt.subplots(figsize=(10, 8))
plot_metric_vs_overlap(
    ax,
    drop_atac, drop_rna,
    metric="NMI",
    y_label="k-means NMI (higher is better)",
    title="(d) Fused-space biological structure vs overlap",
    logx=False,
    show_raw=True,
)
plt.tight_layout()
plt.show()


# %% [code]
fig, ax = plt.subplots(figsize=(16, 3))

# Choose regime boundaries (edit these if you want them to match your narrative exactly)
low_anchor_max = 0.03
modest_max = 0.10

ax.axvspan(1e-4, low_anchor_max, alpha=0.15)
ax.axvspan(low_anchor_max, modest_max, alpha=0.10)

ax.set_xscale("log")
ax.set_xlim(1e-3, 1.0)
ax.set_ylim(0, 1)

ax.set_yticks([])
ax.set_xlabel("Overlap fraction (log scale)")
ax.set_title("(e) Regimes: correspondence collapse vs structure stabilization")

ax.text(0.002, 0.6, "Low-anchor regime:\nstrict correspondence\ncollapses", fontsize=11)
ax.text(0.04, 0.6, "Modest overlap:\ndownstream structure\nstabilizes", fontsize=11)
ax.text(0.25, 0.6, "High overlap:\nall metrics\nsaturate", fontsize=11)

ax.grid(True, alpha=0.25)
plt.tight_layout()
plt.show()


# %% [code]
import matplotlib.pyplot as plt
from mpl_toolkits.axes_grid1.inset_locator import inset_axes, mark_inset

fig, ax = plt.subplots(figsize=(13, 3))

low_anchor_max = 0.03
modest_max = 0.10

# Main axis
ax.axvspan(0.0, low_anchor_max, alpha=0.15)
ax.axvspan(low_anchor_max, modest_max, alpha=0.10)
ax.set_xlim(0.0, 1.0)
ax.set_ylim(0, 1)
ax.set_yticks([])
ax.set_xlabel("Overlap fraction")
ax.set_title("(e) Regimes: correspondence collapse vs structure stabilization")
ax.grid(True, alpha=0.25)

# Put the "high overlap" label on the main axis normally
ax.text(0.55, 0.70, "High overlap:\nall metrics\nsaturate",
        ha="center", va="center", fontsize=11)

# Inset: zoom into 0–0.12
axins = inset_axes(ax, width="75%", height="65%", loc="upper left",
                   bbox_to_anchor=(0.02, 0.05, 0.5, 0.9),
                   bbox_transform=ax.transAxes, borderpad=0.6)

axins.axvspan(0.0, low_anchor_max, alpha=0.15)
axins.axvspan(low_anchor_max, modest_max, alpha=0.10)
axins.set_xlim(0.0, 0.12)
axins.set_ylim(0, 1)
axins.set_xticks([0.00, 0.03, 0.06, 0.09, 0.12])
axins.set_yticks([])
axins.grid(True, alpha=0.2)

# Labels inside inset (now they fit!)
axins.text(low_anchor_max/2, 0.72,
           "Low-anchor:\ncorrespondence\ncollapses",
           ha="center", va="center", fontsize=10)

axins.text((low_anchor_max+modest_max)/2, 0.35,
           "Modest overlap:\nstructure\nstabilizes",
           ha="center", va="center", fontsize=10)

# Draw connectors showing inset corresponds to left region
mark_inset(ax, axins, loc1=2, loc2=4, fc="none", ec="0.4", lw=1)

plt.tight_layout()
plt.show()


# %% [code]


# %% [code]


# %% [code]


# %% [code]


# %% [code]


# %% [code]


# %% [code]


# %% [code]


# %% [code]


# %% [code]


# %% [code]


# %% [code]


# %% [code]


# %% [code]


# %% [code]


# %% [code]


# %% [code]


# %% [code]


# %% [code]


# %% [code]


# %% [code]


# %% [code]


# %% [code]


# %% [code]


# %% [code]


# %% [code]


# %% [code]


# %% [code]


# %% [code]


# %% [code]


# %% [code]


# %% [code]


# %% [code]


# %% [code]


# %% [code]


# %% [code]


# %% [code]


# %% [code]


# %% [code]
```
