## Supplemental Code for "Unifying multimodal single-cell data with a mixture-of-experts *β*-variational autoencoder framework": Supp_grid_sweep.ipynb

```
# %% [markdown]
# UniVI – Model parameter grid sweep supplemental figure analysis (robust v1, no leakage)

This notebook builds the UniVI hyperameter grid sweep supplemental figures  **without train–test leakage** and with **loss_mode='v1'**.


# %% [code]
# =============================================================================
# ---- 0) Imports / versions / globals ----
# =============================================================================
import os, gc, time, math, copy, inspect
from dataclasses import dataclass
from typing import Any, Dict, Iterable, Optional, Tuple

import numpy as np
import pandas as pd
import scipy.sparse as sp
import anndata as ad
import scanpy as sc

import torch
from torch.utils.data import Dataset, DataLoader, Sampler

from sklearn.feature_extraction.text import TfidfTransformer
from sklearn.decomposition import TruncatedSVD
from sklearn.preprocessing import normalize
from sklearn.neighbors import NearestNeighbors
from sklearn.cluster import KMeans
from sklearn.metrics import (
    adjusted_rand_score,
    normalized_mutual_info_score,
    silhouette_score,
    calinski_harabasz_score,
    davies_bouldin_score,
    f1_score,
    accuracy_score,
)

# UniVI
import univi as uv
from univi import UniVIMultiModalVAE, ModalityConfig, UniVIConfig, TrainingConfig
from univi.trainer import UniVITrainer
from univi.data import MultiModalDataset, align_paired_obs_names

print("torch:", torch.__version__)
print("scanpy:", sc.__version__)
print("univi:", uv.__version__)

GLOBAL_SEED = 0

def seed_all(seed: int):
    seed = int(seed)
    np.random.seed(seed)
    torch.manual_seed(seed)
    if torch.cuda.is_available():
        torch.cuda.manual_seed_all(seed)

seed_all(GLOBAL_SEED)

device = torch.device(
    "cuda" if torch.cuda.is_available()
    else ("mps" if torch.backends.mps.is_available() else "cpu")
)
print("device:", device)

DTYPE = np.float32


# %% [code]
# =============================================================================
# ---- 1) Load data (paths match your Figure 8 notebook) ----
# =============================================================================
RNA_PATH  = "./data/10x_Genomics_Multiome_data/10x-Multiome-Pbmc10k-RNA.h5ad"
ATAC_PATH = "./data/10x_Genomics_Multiome_data/10x-Multiome-Pbmc10k-ATAC.h5ad"

rna_raw  = sc.read_h5ad(RNA_PATH)
atac_raw = sc.read_h5ad(ATAC_PATH)

def ensure_counts_layer(a: ad.AnnData, layer: str = "counts"):
    if layer not in a.layers:
        a.layers[layer] = a.X.copy()
    return a

rna_raw  = ensure_counts_layer(rna_raw, "counts")
atac_raw = ensure_counts_layer(atac_raw, "counts")

print("RNA:", rna_raw)
print("ATAC:", atac_raw)


# %% [code]
# =============================================================================
# ---- 2) Align paired obs_names ----
# =============================================================================
adata_dict_raw = {"rna": rna_raw, "atac": atac_raw}
adata_dict_raw = align_paired_obs_names(adata_dict_raw)

rna_raw  = adata_dict_raw["rna"]
atac_raw = adata_dict_raw["atac"]

if not (rna_raw.obs_names == atac_raw.obs_names).all():
    raise RuntimeError("obs_names mismatch after align_paired_obs_names()")
print("Aligned n_obs:", rna_raw.n_obs)


# %% [code]
# =============================================================================
# ---- 3) Train/val/test split ----
# =============================================================================
N = int(rna_raw.n_obs)
rng = np.random.default_rng(GLOBAL_SEED)
perm = rng.permutation(N)

frac_train = 0.75
frac_val   = 0.10

n_tr = int(round(frac_train * N))
n_va = int(round(frac_val   * N))
n_te = int(N - n_tr - n_va)

idx_tr = perm[:n_tr]
idx_va = perm[n_tr:n_tr + n_va]
idx_te = perm[n_tr + n_va:]

rna_tr  = rna_raw[idx_tr].copy()
rna_va  = rna_raw[idx_va].copy()
rna_te  = rna_raw[idx_te].copy()

atac_tr = atac_raw[idx_tr].copy()
atac_va = atac_raw[idx_va].copy()
atac_te = atac_raw[idx_te].copy()

print("Splits:", rna_tr.n_obs, rna_va.n_obs, rna_te.n_obs)


# %% [code]
# =============================================================================
# ---- 4) RNA preprocessing (fit on train, apply to val/test) ----
# =============================================================================
def _get_layer_X(a: ad.AnnData, layer: str) -> Any:
    if layer not in a.layers:
        raise KeyError(f"Layer '{layer}' not found. Available layers: {list(a.layers.keys())}")
    X = a.layers[layer]
    return X.tocsr() if sp.issparse(X) else X

def _normalize_log1p_dense_inplace(a: ad.AnnData, *, target_sum: float, out_dtype=DTYPE):
    sc.pp.normalize_total(a, target_sum=float(target_sum))
    sc.pp.log1p(a)
    if sp.issparse(a.X):
        a.X = a.X.toarray()
    a.X = np.asarray(a.X, dtype=out_dtype)

def _apply_zscore_clip_inplace(a: ad.AnnData, mean: np.ndarray, std: np.ndarray, *, clip: float = 10.0):
    X = np.asarray(a.X, dtype=DTYPE)
    X = (X - mean) / std
    if clip is not None:
        X = np.clip(X, -float(clip), float(clip))
    a.X = X

def preprocess_rna_train(
    adata: ad.AnnData,
    *,
    counts_layer: str = "counts",
    n_hvg: int = 2000,
    target_sum: float = 1e4,
    flavor: str = "seurat_v3",
    out_dtype=DTYPE,
) -> Tuple[ad.AnnData, list, np.ndarray, np.ndarray]:
    a = adata.copy()
    a.X = _get_layer_X(a, counts_layer)

    sc.pp.highly_variable_genes(a, n_top_genes=int(n_hvg), flavor=str(flavor))
    hvg = a.var_names[a.var["highly_variable"]].tolist()
    if len(hvg) == 0:
        raise RuntimeError("No HVGs selected. Check your RNA AnnData .X/.layers['counts'].")

    a = a[:, hvg].copy()
    _normalize_log1p_dense_inplace(a, target_sum=target_sum, out_dtype=out_dtype)

    mean = a.X.mean(axis=0).astype(out_dtype, copy=False)
    std  = a.X.std(axis=0, ddof=0).astype(out_dtype, copy=False)
    std  = np.where(std == 0, 1.0, std).astype(out_dtype, copy=False)

    _apply_zscore_clip_inplace(a, mean, std, clip=10.0)
    return a, hvg, mean, std

def preprocess_rna_apply(
    adata: ad.AnnData,
    hvg: list,
    mean: np.ndarray,
    std: np.ndarray,
    *,
    counts_layer: str = "counts",
    target_sum: float = 1e4,
    out_dtype=DTYPE,
) -> ad.AnnData:
    a = adata.copy()
    a.X = _get_layer_X(a, counts_layer)

    # align genes to training HVG order; fill missing with zeros
    hvg = list(hvg)
    present = [g for g in hvg if g in a.var_names]
    a_sub = a[:, present].copy()

    _normalize_log1p_dense_inplace(a_sub, target_sum=target_sum, out_dtype=out_dtype)

    X_full = np.zeros((a_sub.n_obs, len(hvg)), dtype=out_dtype)
    col_map = {g: j for j, g in enumerate(a_sub.var_names.tolist())}
    for j, g in enumerate(hvg):
        jj = col_map.get(g, None)
        if jj is not None:
            X_full[:, j] = a_sub.X[:, jj]

    out = ad.AnnData(X=X_full, obs=a_sub.obs.copy())
    out.obs_names = a_sub.obs_names.copy()
    out.var_names = np.array(hvg, dtype=str)

    _apply_zscore_clip_inplace(out, mean, std, clip=10.0)
    out.X = np.asarray(out.X, dtype=out_dtype)
    return out

# ---- run ----
rna_tr_pp, hvg, rna_mean, rna_std = preprocess_rna_train(rna_tr, n_hvg=2000, target_sum=1e4)
rna_va_pp = preprocess_rna_apply(rna_va, hvg, rna_mean, rna_std, target_sum=1e4)
rna_te_pp = preprocess_rna_apply(rna_te, hvg, rna_mean, rna_std, target_sum=1e4)

print("RNA dims:", rna_tr_pp.shape, "|", rna_va_pp.shape, "|", rna_te_pp.shape)
assert list(rna_va_pp.var_names) == list(hvg)
assert list(rna_te_pp.var_names) == list(hvg)


# %% [code]
# =============================================================================
# ---- 5) ATAC preprocessing (TF-IDF + LSI) ----
# =============================================================================
def _get_counts_csr(a: ad.AnnData, counts_layer: str = "counts") -> sp.csr_matrix:
    X = _get_layer_X(a, counts_layer)
    return X.tocsr() if sp.issparse(X) else sp.csr_matrix(X)

def preprocess_atac_lsi_train(
    adata: ad.AnnData,
    *,
    counts_layer: str = "counts",
    n_lsi: int = 101,
    drop_first: bool = True,
    binarize: bool = False,
    seed: int = 0,
    out_dtype=DTYPE,
):
    a = adata.copy()
    X = _get_counts_csr(a, counts_layer)

    if binarize:
        X = X.copy()
        X.data[:] = 1.0

    tfidf = TfidfTransformer(norm="l2", use_idf=True, smooth_idf=True, sublinear_tf=True)
    X_tfidf = tfidf.fit_transform(X)

    svd = TruncatedSVD(n_components=int(n_lsi), random_state=int(seed))
    X_lsi = svd.fit_transform(X_tfidf)

    if drop_first:
        if X_lsi.shape[1] < 2:
            raise RuntimeError(f"Cannot drop_first with n_lsi={n_lsi} (need >=2).")
        X_lsi = X_lsi[:, 1:]

    X_lsi = normalize(X_lsi, norm="l2")
    X_lsi = np.asarray(X_lsi, dtype=out_dtype)

    out = ad.AnnData(X=X_lsi, obs=a.obs.copy())
    out.obs_names = a.obs_names.copy()
    return out, tfidf, svd

def preprocess_atac_lsi_apply(
    adata: ad.AnnData,
    tfidf: TfidfTransformer,
    svd: TruncatedSVD,
    *,
    counts_layer: str = "counts",
    drop_first: bool = True,
    binarize: bool = False,
    out_dtype=DTYPE,
) -> ad.AnnData:
    a = adata.copy()
    X = _get_counts_csr(a, counts_layer)

    if binarize:
        X = X.copy()
        X.data[:] = 1.0

    X_tfidf = tfidf.transform(X)
    X_lsi = svd.transform(X_tfidf)

    if drop_first:
        X_lsi = X_lsi[:, 1:]

    X_lsi = normalize(X_lsi, norm="l2")
    X_lsi = np.asarray(X_lsi, dtype=out_dtype)

    out = ad.AnnData(X=X_lsi, obs=a.obs.copy())
    out.obs_names = a.obs_names.copy()
    return out

# ---- run ----
atac_tr_lsi, tfidf_atac, svd_atac = preprocess_atac_lsi_train(
    atac_tr, n_lsi=101, drop_first=True, seed=GLOBAL_SEED
)
atac_va_lsi = preprocess_atac_lsi_apply(atac_va, tfidf_atac, svd_atac, drop_first=True)
atac_te_lsi = preprocess_atac_lsi_apply(atac_te, tfidf_atac, svd_atac, drop_first=True)

print("ATAC dims:", atac_tr_lsi.shape, "|", atac_va_lsi.shape, "|", atac_te_lsi.shape)


# %% [code]
# =============================================================================
# ---- 6) Build MultiModalDataset ----
# =============================================================================
def _concat_three(a1: ad.AnnData, a2: ad.AnnData, a3: ad.AnnData) -> ad.AnnData:
    # Use scanpy concat to avoid deprecated AnnData.concatenate behavior differences
    return sc.concat([a1, a2, a3], axis=0, join="outer", merge="same", label=None, index_unique=None)

adata_dict = {
    "rna":  _concat_three(rna_tr_pp, rna_va_pp, rna_te_pp),
    "atac": _concat_three(atac_tr_lsi, atac_va_lsi, atac_te_lsi),
}
adata_dict = align_paired_obs_names(adata_dict)

dataset = MultiModalDataset(adata_dict=adata_dict, X_key="X", device=None)

n_tr = int(rna_tr_pp.n_obs)
n_va = int(rna_va_pp.n_obs)
n_te = int(rna_te_pp.n_obs)

train_idx = np.arange(0, n_tr, dtype=np.int64)
val_idx   = np.arange(n_tr, n_tr + n_va, dtype=np.int64)
test_idx  = np.arange(n_tr + n_va, n_tr + n_va + n_te, dtype=np.int64)

label_key = "cell_type"  # adjust if needed
if label_key not in adata_dict["rna"].obs.columns:
    print("Available RNA obs columns:", list(adata_dict["rna"].obs.columns)[:50])
    raise KeyError(f"label_key='{label_key}' not found in RNA obs")

y_all = adata_dict["rna"].obs[label_key].astype(str).to_numpy()

print("dataset n:", len(dataset), "train/val/test:", n_tr, n_va, n_te)
print(pd.Series(y_all).value_counts().head(20))


# %% [code]
# =============================================================================
# ---- 7) Overlap loaders with homogeneous batches ----
# =============================================================================
class IndexedDataset(Dataset):
    """Wrap base_dataset + a list of global indices, returning (x_dict, global_id)."""
    def __init__(self, base_dataset: Dataset, indices: np.ndarray):
        self.base = base_dataset
        self.indices = np.asarray(indices, dtype=np.int64)
        if self.indices.ndim != 1:
            raise ValueError("indices must be 1D")

    def __len__(self) -> int:
        return int(self.indices.shape[0])

    def __getitem__(self, i: int):
        gi = int(self.indices[int(i)])
        item = self.base[gi]
        if isinstance(item, dict):
            x = item
        elif isinstance(item, (tuple, list)) and len(item) >= 1 and isinstance(item[0], dict):
            x = item[0]
        else:
            raise TypeError(f"Expected base_dataset to yield dict or (dict,...); got {type(item)}")
        return dict(x), gi

class DeterministicMaskDataset(Dataset):
    """
    groups==0: paired (keep both)
    groups!=0: unpaired (DROP drop_modality key)
    """
    def __init__(self, base_ds: Dataset, groups: np.ndarray, *, drop_modality: str):
        self.base = base_ds
        self.groups = np.asarray(groups, dtype=np.int64)
        self.drop_modality = str(drop_modality)
        if len(self.base) != len(self.groups):
            raise ValueError(f"len(base)={len(self.base)} != len(groups)={len(self.groups)}")

    def __len__(self) -> int:
        return len(self.base)

    def __getitem__(self, i: int):
        x, gid = self.base[i]
        g = int(self.groups[int(i)])
        x = dict(x)
        if g != 0:
            x.pop(self.drop_modality, None)
        return x, gid

def collate_xdict_with_idx(batch):
    if len(batch) == 0:
        raise ValueError("Empty batch")
    gids = torch.as_tensor([b[1] for b in batch], dtype=torch.long)

    keys0 = set(batch[0][0].keys())
    for x, _ in batch[1:]:
        if set(x.keys()) != keys0:
            raise RuntimeError(
                "Non-homogeneous batch (mixed modality keys inside batch). "
                "Fix grouping/sampler."
            )

    out = {}
    for k in keys0:
        out[k] = torch.stack([x[k] for x, _ in batch], dim=0)
    return out, gids

class InterleavedGroupedBatchSampler(Sampler):
    """Yield homogeneous batches: all paired (group==0) or all unpaired (group!=0)."""
    def __init__(
        self,
        groups: np.ndarray,
        batch_size: int,
        *,
        seed: int = 0,
        unpaired_per_paired: float = 1.0,
        oversample_paired: bool = True,
        oversample_unpaired: bool = True,
    ):
        self.groups = np.asarray(groups, dtype=np.int64)
        self.batch_size = int(batch_size)
        self.seed = int(seed)

        upp = float(unpaired_per_paired)
        if not np.isfinite(upp) or upp < 0:
            raise ValueError(f"unpaired_per_paired must be finite and >=0, got {upp}")
        self.unpaired_per_paired = upp

        self.oversample_paired = bool(oversample_paired)
        self.oversample_unpaired = bool(oversample_unpaired)

        self.paired_idx = np.where(self.groups == 0)[0].astype(np.int64)
        self.unpaired_idx = np.where(self.groups != 0)[0].astype(np.int64)
        if len(self.paired_idx) == 0:
            raise ValueError("No paired samples (group==0). Need at least one anchor.")

        self.only_paired = (len(self.unpaired_idx) == 0)

        # Paired-driven epoch length (in batches)
        if len(self.paired_idx) >= self.batch_size:
            self.n_paired_batches = len(self.paired_idx) // self.batch_size
        else:
            self.n_paired_batches = 1 if self.oversample_paired else 0

        self.n_unpaired_expected = 0 if self.only_paired else int(round(self.n_paired_batches * self.unpaired_per_paired))
        self.n_batches = max(int(self.n_paired_batches + self.n_unpaired_expected), 1)

    def __len__(self) -> int:
        return int(self.n_batches)

    def __iter__(self):
        rng = np.random.default_rng(self.seed)

        paired_pool = self.paired_idx.copy()
        rng.shuffle(paired_pool)
        p_ptr = 0

        unpaired_pool = self.unpaired_idx.copy()
        if len(unpaired_pool) > 0:
            rng.shuffle(unpaired_pool)
        u_ptr = 0

        def sample_with_replacement(pool):
            sel = rng.integers(0, len(pool), size=self.batch_size)
            return pool[sel]

        def next_batch(pool, ptr, oversample):
            end = ptr + self.batch_size
            if end <= len(pool):
                return pool[ptr:end], end
            if not oversample:
                return None, ptr
            rng.shuffle(pool)
            ptr = 0
            end = ptr + self.batch_size
            if end <= len(pool):
                return pool[ptr:end], end
            return sample_with_replacement(pool), ptr

        carry = 0.0
        for _ in range(self.n_paired_batches):
            if len(paired_pool) < self.batch_size:
                if not self.oversample_paired:
                    break
                pb = sample_with_replacement(paired_pool)
            else:
                pb, p_ptr = next_batch(paired_pool, p_ptr, self.oversample_paired)
                if pb is None:
                    break
            yield pb.tolist()

            if self.only_paired:
                continue

            carry += self.unpaired_per_paired
            while carry >= 1.0 and len(unpaired_pool) > 0:
                carry -= 1.0
                if len(unpaired_pool) < self.batch_size:
                    if not self.oversample_unpaired:
                        break
                    ub = sample_with_replacement(unpaired_pool)
                else:
                    ub, u_ptr = next_batch(unpaired_pool, u_ptr, self.oversample_unpaired)
                    if ub is None:
                        break
                yield ub.tolist()

def make_loaders_with_overlap_v1(
    base_dataset,
    train_idx,
    val_idx,
    test_idx,
    *,
    overlap_fraction: float,
    drop_modality: str,
    seed: int,
    batch_size: int = 256,
    num_workers: int = 0,
    oversample_paired: bool = True,
):
    """overlap_fraction applies ONLY to training. val/test remain paired."""
    p = float(overlap_fraction)
    if not (0.0 <= p <= 1.0):
        raise ValueError(f"overlap_fraction must be in [0,1], got {p}")

    rng = np.random.default_rng(int(seed))

    train_indexed = IndexedDataset(base_dataset, indices=train_idx)
    N = len(train_indexed)
    if N == 0:
        raise ValueError("Empty train split")

    n_paired = int(round(p * N))
    n_paired = max(min(n_paired, N), 0)

    perm = rng.permutation(N)
    paired_local = perm[:n_paired]

    groups = np.ones(N, dtype=np.int64)
    groups[paired_local] = 0

    train_masked = DeterministicMaskDataset(train_indexed, groups=groups, drop_modality=str(drop_modality))

    n_paired_real = int((groups == 0).sum())
    n_unpaired_real = int((groups != 0).sum())
    upp_sampling = float(n_unpaired_real / max(n_paired_real, 1))

    print(
        f"[overlap={p:g}] batch_size={int(batch_size)} "
        f"train_paired={n_paired_real} train_unpaired={n_unpaired_real} "
        f"(sampler_upp={upp_sampling:.6f}) drop_modality={drop_modality}"
    )

    if n_unpaired_real == 0:
        train_loader = DataLoader(
            train_masked,
            batch_size=int(batch_size),
            shuffle=True,
            drop_last=True,
            num_workers=int(num_workers),
            collate_fn=collate_xdict_with_idx,
        )
    else:
        batch_sampler = InterleavedGroupedBatchSampler(
            groups=groups,
            batch_size=int(batch_size),
            seed=int(seed),
            unpaired_per_paired=upp_sampling,
            oversample_paired=bool(oversample_paired),
            oversample_unpaired=True,
        )
        train_loader = DataLoader(
            train_masked,
            batch_sampler=batch_sampler,
            num_workers=int(num_workers),
            collate_fn=collate_xdict_with_idx,
        )

    val_loader = DataLoader(
        IndexedDataset(base_dataset, indices=val_idx),
        batch_size=int(batch_size),
        shuffle=False,
        drop_last=False,
        num_workers=int(num_workers),
        collate_fn=collate_xdict_with_idx,
    )
    test_loader = DataLoader(
        IndexedDataset(base_dataset, indices=test_idx),
        batch_size=int(batch_size),
        shuffle=False,
        drop_last=False,
        num_workers=int(num_workers),
        collate_fn=collate_xdict_with_idx,
    )

    info = {
        "train_paired": n_paired_real,
        "train_unpaired": n_unpaired_real,
        "sampler_unpaired_per_paired": float(upp_sampling),
        "overlap_fraction": float(p),
        "drop_modality": str(drop_modality),
    }
    return train_loader, val_loader, test_loader, info


# %% [code]
# =============================================================================
# ---- 8) UniVI config + trainer compat + training wrapper (ES-gated, seed-fixed) ----
# =============================================================================
def make_univi_cfg(
    rna_dim: int,
    atac_dim: int,
    *,
    latent_dim: int = 30,
    beta: float = 1.25,
    gamma: float = 4.35,
    kl_anneal_start: int = 0,
    kl_anneal_end: int = 60,
    align_anneal_start: int = 25,
    align_anneal_end: int = 85,
    encoder_dropout: float = 0.10,
    decoder_dropout: float = 0.05,
) -> UniVIConfig:
    return UniVIConfig(
        latent_dim=int(latent_dim),
        beta=float(beta),
        gamma=float(gamma),
        encoder_dropout=float(encoder_dropout),
        decoder_dropout=float(decoder_dropout),
        encoder_batchnorm=True,
        decoder_batchnorm=False,
        kl_anneal_start=int(kl_anneal_start),
        kl_anneal_end=int(kl_anneal_end),
        align_anneal_start=int(align_anneal_start),
        align_anneal_end=int(align_anneal_end),
        modalities=[
            ModalityConfig(
                name="rna",
                input_dim=int(rna_dim),
                encoder_hidden=[512, 256, 128],
                decoder_hidden=[128, 256, 512],
                likelihood="gaussian",
            ),
            ModalityConfig(
                name="atac",
                input_dim=int(atac_dim),
                encoder_hidden=[128, 64],
                decoder_hidden=[64, 128],
                likelihood="gaussian",
            ),
        ],
        class_heads=[],
    )

def make_train_cfg(
    device,
    *,
    seed: int,
    n_epochs: int = 5000,
    batch_size: int = 256,
    lr: float = 1e-3,
    weight_decay: float = 1e-4,
    log_every: int = 25,
    grad_clip: float = 5.0,
    early_stopping: bool = True,
    patience: int = 100,
    min_delta: float = 0.0,
    num_workers: int = 0,
) -> TrainingConfig:
    return TrainingConfig(
        n_epochs=int(n_epochs),
        batch_size=int(batch_size),
        lr=float(lr),
        weight_decay=float(weight_decay),
        device=device,
        log_every=int(log_every),
        grad_clip=float(grad_clip),
        num_workers=int(num_workers),
        seed=int(seed),
        early_stopping=bool(early_stopping),
        patience=int(patience),
        min_delta=float(min_delta),
    )

def _build_univi_trainer_compat(*, model, train_cfg, train_loader, val_loader):
    sig = inspect.signature(UniVITrainer.__init__)
    params = sig.parameters
    kwargs = {}

    if "model" in params: kwargs["model"] = model
    if "train_loader" in params: kwargs["train_loader"] = train_loader
    if "val_loader" in params: kwargs["val_loader"] = val_loader

    for cand in ("train_cfg", "training_cfg", "cfg_train", "config", "cfg"):
        if cand in params:
            kwargs[cand] = train_cfg
            return UniVITrainer(**kwargs)

    for k, v in vars(train_cfg).items():
        if k in params:
            kwargs[k] = v

    return UniVITrainer(**kwargs)

def _trainer_fit_compat(trainer, train_loader, val_loader):
    try:
        return trainer.fit()
    except TypeError:
        pass
    try:
        return trainer.fit(train_loader=train_loader, val_loader=val_loader)
    except TypeError:
        pass
    return trainer.fit(train_loader, val_loader)

@dataclass
class TrainResult:
    model: torch.nn.Module
    best_epoch: Optional[int]
    best_val: Optional[float]
    wall_seconds: float
    peak_mem_mb: Optional[float]

def _filter_kwargs_for_init(cls_or_fn, kwargs: dict, *, tag: str = "") -> dict:
    if not kwargs:
        return {}
    sig = inspect.signature(cls_or_fn)
    params = sig.parameters
    if any(p.kind == inspect.Parameter.VAR_KEYWORD for p in params.values()):
        return dict(kwargs)
    keep, dropped = {}, []
    for k, v in kwargs.items():
        if k in params:
            keep[k] = v
        else:
            dropped.append(k)
    if dropped:
        prefix = f"[{tag}] " if tag else ""
        print(f"{prefix}dropping unsupported kwargs: {dropped}")
    return keep

def train_one(
    *,
    train_loader,
    val_loader,
    univi_cfg: UniVIConfig,
    seed: int,
    device,
    loss_mode: str = "v1",
    v1_recon: str = "moe",
    batch_size: int = 256,
    lr: float = 1e-3,
    weight_decay: float = 1e-4,
    n_epochs: int = 5000,
    early_stopping: bool = True,
    patience: int = 100,
    min_delta: float = 0.0,
    min_epochs: int = 0,
    grad_clip: float = 5.0,
    log_every: int = 25,
    num_workers: int = 0,
    extra_model_kwargs: Optional[dict] = None,
):
    seed_all(int(seed))

    model_kwargs = _filter_kwargs_for_init(
        UniVIMultiModalVAE.__init__,
        dict(extra_model_kwargs or {}),
        tag="model",
    )

    model = UniVIMultiModalVAE(
        univi_cfg,
        loss_mode=str(loss_mode),
        v1_recon=str(v1_recon),
        **model_kwargs,
    ).to(device)

    # ---- FIX: real early-stopping gate
    # If trainer uses patience as "epochs since best", increase patience by min_epochs
    # so it cannot trip before min_epochs has elapsed.
    min_epochs = int(max(min_epochs, 0))
    patience = int(patience)
    patience_eff = patience + min_epochs if bool(early_stopping) else patience

    train_cfg = make_train_cfg(
        device=device,
        seed=int(seed),
        n_epochs=int(n_epochs),
        batch_size=int(batch_size),
        lr=float(lr),
        weight_decay=float(weight_decay),
        log_every=int(log_every),
        grad_clip=float(grad_clip),
        early_stopping=bool(early_stopping),
        patience=int(patience_eff),
        min_delta=float(min_delta),
        num_workers=int(num_workers),
    )

    trainer = _build_univi_trainer_compat(
        model=model,
        train_cfg=train_cfg,
        train_loader=train_loader,
        val_loader=val_loader,
    )

    if not hasattr(trainer, "train_loader"):
        trainer.train_loader = train_loader
    if not hasattr(trainer, "val_loader"):
        trainer.val_loader = val_loader

    t0 = time.perf_counter()
    _trainer_fit_compat(trainer, train_loader=train_loader, val_loader=val_loader)
    t1 = time.perf_counter()

    peak_mem_mb = getattr(trainer, "peak_mem_mb", None)
    best_epoch = getattr(trainer, "best_epoch", None)
    best_val = getattr(trainer, "best_val_loss", None)
    if best_val is None:
        best_val = getattr(trainer, "best_val", None)

    if hasattr(trainer, "restore_best"):
        try:
            trainer.restore_best()
        except Exception:
            pass

    return TrainResult(
        model=model,
        best_epoch=(int(best_epoch) if best_epoch is not None else None),
        best_val=(float(best_val) if best_val is not None else None),
        wall_seconds=float(t1 - t0),
        peak_mem_mb=(float(peak_mem_mb) if peak_mem_mb is not None else None),
    )


# %% [code]
# =============================================================================
# ---- 9) Robust per-modality latent extraction (FIXED for mu_dict outputs) ----
# =============================================================================
def _to_numpy(x):
    if isinstance(x, np.ndarray):
        return x
    if torch.is_tensor(x):
        return x.detach().cpu().numpy()
    return np.asarray(x)

def _pick_tensor(obj):
    if torch.is_tensor(obj):
        return obj
    if isinstance(obj, (tuple, list)):
        for x in obj[::-1]:
            if torch.is_tensor(x):
                return x
        return None
    if isinstance(obj, dict):
        for k in ("z", "latent", "mu", "z_shared", "emb", "repr"):
            if k in obj and torch.is_tensor(obj[k]):
                return obj[k]
        for v in obj.values():
            if torch.is_tensor(v):
                return v
        return None
    return None

def _extract_modality_latent(out: Any, mod: str) -> Optional[torch.Tensor]:
    """
    Handles UniVI outputs like the ones you showed:
      keys include: 'mu_dict', 'logvar_dict', 'z', ...
    Prefer mu_dict[mod] as a stable modality-specific representation.
    """
    if not isinstance(out, dict):
        return None

    # 1) Your current UniVI version: mu_dict / logvar_dict
    if "mu_dict" in out and isinstance(out["mu_dict"], dict) and mod in out["mu_dict"]:
        t = out["mu_dict"][mod]
        return t if torch.is_tensor(t) else None

    # 2) Common alternatives in other versions
    for key in (f"mu_{mod}", f"z_{mod}", mod):
        if key in out:
            t = _pick_tensor(out[key])
            if t is not None:
                return t

    if "latents" in out and isinstance(out["latents"], dict):
        t = _pick_tensor(out["latents"].get(mod, None))
        if t is not None:
            return t

    if "z_dict" in out and isinstance(out["z_dict"], dict) and mod in out["z_dict"]:
        t = out["z_dict"][mod]
        return t if torch.is_tensor(t) else None

    return None

def _extract_fused_latent(out: Any) -> Optional[torch.Tensor]:
    if not isinstance(out, dict):
        return _pick_tensor(out)
    for k in ("z", "z_shared", "latent", "mu", "mu_z"):
        if k in out:
            t = _pick_tensor(out[k])
            if t is not None:
                return t
    return _pick_tensor(out)

@torch.no_grad()
def _model_call_any(model, xb: dict):
    # Prefer inference/encode if present; otherwise forward
    if hasattr(model, "inference"):
        try:
            return model.inference(xb)
        except Exception:
            pass
    if hasattr(model, "encode"):
        try:
            return model.encode(xb)
        except Exception:
            pass
    return model(xb)

@torch.no_grad()
def encode_modality_mu(model, xb: dict, *, device, mod: str) -> torch.Tensor:
    if mod not in xb:
        raise KeyError(f"Batch missing modality '{mod}'. Keys: {list(xb.keys())}")
    xb_mod = {mod: xb[mod].to(device)}
    out = _model_call_any(model, xb_mod)
    z = _extract_modality_latent(out, mod)
    if z is None:
        keys = list(out.keys()) if isinstance(out, dict) else None
        raise RuntimeError(
            f"Could not extract modality latent for mod='{mod}'. "
            f"out_type={type(out)} keys={keys}"
        )
    return z.detach().cpu()

@torch.no_grad()
def encode_fused_z(model, xb: dict, *, device) -> torch.Tensor:
    xb2 = {k: v.to(device) for k, v in xb.items()}
    out = _model_call_any(model, xb2)
    z = _extract_fused_latent(out)
    if z is None:
        keys = list(out.keys()) if isinstance(out, dict) else None
        raise RuntimeError(f"Could not extract fused latent. out_type={type(out)} keys={keys}")
    return z.detach().cpu()

@torch.no_grad()
def collect_test_latents(model, loader, *, device):
    """
    For paired loader (xb has 'rna' and 'atac'):
      returns Z_fused, Z_rna(mu), Z_atac(mu), gids
    """
    model.eval()
    Zf, Zr, Za, G = [], [], [], []

    for xb, gids in loader:
        if "rna" not in xb or "atac" not in xb:
            raise RuntimeError(f"Expected paired xb keys ['rna','atac']; got {list(xb.keys())}")

        zf = encode_fused_z(model, xb, device=device)
        zr = encode_modality_mu(model, xb, device=device, mod="rna")
        za = encode_modality_mu(model, xb, device=device, mod="atac")

        Zf.append(zf); Zr.append(zr); Za.append(za)
        G.append(gids.detach().cpu())

    return torch.cat(Zf, 0), torch.cat(Zr, 0), torch.cat(Za, 0), torch.cat(G, 0)


# %% [code]
# =============================================================================
# ---- 10) Eval helpers (kNN, clustering, mixing) ----
# =============================================================================
import numpy as np
from sklearn.cluster import KMeans
from sklearn.metrics import (
    adjusted_rand_score,
    normalized_mutual_info_score,
    silhouette_score,
    calinski_harabasz_score,
    davies_bouldin_score,
    accuracy_score,
    f1_score,
    balanced_accuracy_score,
)
from sklearn.neighbors import NearestNeighbors


def _to_numpy(x):
    """Torch tensor -> numpy, otherwise np.asarray."""
    try:
        import torch
        if isinstance(x, torch.Tensor):
            return x.detach().cpu().numpy()
    except Exception:
        pass
    return np.asarray(x)


def l2_normalize_rows(X, eps=1e-12):
    X = _to_numpy(X).astype(np.float32, copy=False)
    n = np.linalg.norm(X, axis=1, keepdims=True)
    return X / np.maximum(n, eps)


def knn_majority_vote_predict(y_neighbors):
    """
    y_neighbors: (n, k) int labels
    Returns: (n,) predicted labels by majority vote; ties -> smallest label id.
    """
    y_neighbors = np.asarray(y_neighbors)
    pred = np.empty(y_neighbors.shape[0], dtype=y_neighbors.dtype)
    for i in range(y_neighbors.shape[0]):
        vals, cnts = np.unique(y_neighbors[i], return_counts=True)
        pred[i] = vals[np.argmax(cnts)]
    return pred


def knn_label_transfer(Z_src, y_src, Z_tgt, *, k=5, metric="cosine", normalize=True):
    """
    Predict labels for target points using kNN in source space (source->target).
    This is true "transfer": y_src are labels for the Z_src points.

    Returns: pred_tgt labels (len = n_tgt)
    """
    Z_src = _to_numpy(Z_src).astype(np.float32, copy=False)
    Z_tgt = _to_numpy(Z_tgt).astype(np.float32, copy=False)
    y_src = np.asarray(y_src, dtype=np.int64)

    if normalize:
        Z_src = l2_normalize_rows(Z_src)
        Z_tgt = l2_normalize_rows(Z_tgt)

    nn = NearestNeighbors(n_neighbors=int(k), metric=str(metric))
    nn.fit(Z_src)
    idx = nn.kneighbors(Z_tgt, return_distance=False)
    votes = y_src[idx]
    return knn_majority_vote_predict(votes)


def knn_loo_accuracy(Z, y, *, k=5, metric="cosine", normalize=True):
    """
    Leave-one-out kNN accuracy within a single embedding space.
    Uses k+1 neighbors and drops self.
    """
    Z = _to_numpy(Z).astype(np.float32, copy=False)
    y = np.asarray(y, dtype=np.int64)

    if normalize:
        Z = l2_normalize_rows(Z)

    nn = NearestNeighbors(n_neighbors=int(k) + 1, metric=str(metric))
    nn.fit(Z)
    idx = nn.kneighbors(Z, return_distance=False)[:, 1:]  # drop self
    pred = knn_majority_vote_predict(y[idx])
    return float(accuracy_score(y, pred)), float(f1_score(y, pred, average="macro")), float(balanced_accuracy_score(y, pred))


def clustering_metrics(Z, y_true, *, kmeans_k, seed=0):
    """
    KMeans clusters on Z, compare to y_true.
    Returns: ARI, NMI, SIL_kmeans, CH_kmeans, DB_kmeans, SIL_true (silhouette by true labels).
    """
    Z = _to_numpy(Z).astype(np.float32, copy=False)
    y_true = np.asarray(y_true, dtype=np.int64)

    out = {"kmeans_k": int(kmeans_k)}

    if int(kmeans_k) < 2:
        out.update({"ARI": np.nan, "NMI": np.nan, "SIL_kmeans": np.nan, "CH_kmeans": np.nan, "DB_kmeans": np.nan})
        out["SIL_true"] = np.nan
        return out

    km = KMeans(n_clusters=int(kmeans_k), random_state=int(seed), n_init=10)
    y_km = km.fit_predict(Z)

    out["ARI"] = float(adjusted_rand_score(y_true, y_km))
    out["NMI"] = float(normalized_mutual_info_score(y_true, y_km))

    if len(np.unique(y_km)) > 1 and Z.shape[0] > int(kmeans_k):
        out["SIL_kmeans"] = float(silhouette_score(Z, y_km, metric="euclidean"))
        out["CH_kmeans"]  = float(calinski_harabasz_score(Z, y_km))
        out["DB_kmeans"]  = float(davies_bouldin_score(Z, y_km))
    else:
        out["SIL_kmeans"] = np.nan
        out["CH_kmeans"]  = np.nan
        out["DB_kmeans"]  = np.nan

    if len(np.unique(y_true)) > 1 and Z.shape[0] > len(np.unique(y_true)):
        out["SIL_true"] = float(silhouette_score(Z, y_true, metric="euclidean"))
    else:
        out["SIL_true"] = np.nan

    return out


def modality_mixing_score(Z, modality_labels, *, k=15, metric="cosine", normalize=True):
    """
    1 - mean(frac of same-modality neighbors). Higher is better mixing.
    """
    Z = _to_numpy(Z).astype(np.float32, copy=False)
    m = np.asarray(modality_labels, dtype=np.int64)

    if normalize:
        Z = l2_normalize_rows(Z)

    nn = NearestNeighbors(n_neighbors=int(k) + 1, metric=str(metric))
    nn.fit(Z)
    idx = nn.kneighbors(Z, return_distance=False)[:, 1:]
    same = (m[idx] == m[:, None]).mean(axis=1)
    return float(1.0 - same.mean())


# %% [code]
# =============================================================================
# ---- 11) Paired eval metrics (FOSCTTM + recall@k + evaluate_on_test) ----
# =============================================================================
def foscttm_rank_fraction(Za, Zb):
    """
    Classic FOSCTTM (lower is better): for each i, fraction of non-match pairs
    that are closer than the true match, averaged across i.
    Uses cosine similarity via L2-normalized dot product.
    """
    A = l2_normalize_rows(Za)
    B = l2_normalize_rows(Zb)
    S = A @ B.T  # cosine sim
    D = 1.0 - S  # cosine distance proxy

    d_true = np.diag(D)  # true pair distance
    n = D.shape[0]
    mask = ~np.eye(n, dtype=bool)

    frac = (D < d_true[:, None]) & mask
    return float(frac.sum(axis=1).mean() / max(1, (n - 1)))


def recall_at_k(Za, Zb, *, ks=(1, 5, 10, 25, 50, 100), symmetric=True):
    """
    Recall@k retrieval in cosine space after L2 normalization.
    recall_A2B@k: true match in top-k when querying A against B
    recall_B2A@k: reverse
    recall_sym@k: average of the two
    """
    A = l2_normalize_rows(Za)
    B = l2_normalize_rows(Zb)
    S = A @ B.T

    n = S.shape[0]
    true = np.arange(n)[:, None]
    out = {}

    order = np.argsort(-S, axis=1)
    for k in ks:
        out[f"recall_A2B@{int(k)}"] = float((order[:, :int(k)] == true).any(axis=1).mean())

    if symmetric:
        order2 = np.argsort(-S.T, axis=1)
        for k in ks:
            out[f"recall_B2A@{int(k)}"] = float((order2[:, :int(k)] == true).any(axis=1).mean())
        for k in ks:
            out[f"recall_sym@{int(k)}"] = float(0.5 * (out[f"recall_A2B@{int(k)}"] + out[f"recall_B2A@{int(k)}"]))
    return out


def foscttm_at_k(Za, Zb, *, ks=(1, 5, 10, 25, 50, 100)):
    """
    "FOSCTTM@k" as an error-rate derived from symmetric recall@k:
        FOSCTTM@k = 1 - recall_sym@k
    Lower is better, consistent with classic FOSCTTM.
    """
    rec = recall_at_k(Za, Zb, ks=tuple(int(x) for x in ks), symmetric=True)
    out = {}
    for k in ks:
        out[f"FOSCTTM@{int(k)}"] = float(1.0 - rec[f"recall_sym@{int(k)}"])
    # back-compat: FOSCTTM means classic rank fraction (keep both!)
    return out, rec


def make_label_codes(y_all):
    """
    Robust label encoding. Keeps deterministic mapping via sorted unique strings.
    """
    y = np.asarray(y_all, dtype=str)
    uniq = np.unique(y)
    mapping = {lab: i for i, lab in enumerate(uniq.tolist())}
    y_int = np.array([mapping[v] for v in y], dtype=np.int64)
    return y_int, mapping


def evaluate_on_test(
    model,
    test_loader,
    *,
    device,
    y_all,
    k_knn=3,
    seed=0,
    fos_ks=(1, 5, 10, 25, 50, 100),
    mix_k=15,
):
    """
    Paired-only evaluation:
      - Requires collect_test_latents() returning (Zf, Zr, Za, gids).
      - y_all indexed by gids gives per-cell labels (strings or ints).

    Outputs include:
      - classic FOSCTTM (rank fraction)
      - recall@k (A2B, B2A, sym)
      - FOSCTTM@k = 1 - recall_sym@k
      - within-embedding LOO kNN acc/macroF1/bal_acc for Zr, Za, Zf
      - KMeans ARI/NMI + SIL/CH/DB + SIL_true on Zf
      - modality mixing score on stacked [Zr; Za]
    """
    Zf_t, Zr_t, Za_t, gids_t = collect_test_latents(model, test_loader, device=device)

    Zf = _to_numpy(Zf_t)
    Zr = _to_numpy(Zr_t)
    Za = _to_numpy(Za_t)
    gids = _to_numpy(gids_t).astype(np.int64)

    y_int_all, _ = make_label_codes(y_all)
    y_true = y_int_all[gids]

    out = {}
    out["n_test"] = int(len(gids))
    out["latent_dim"] = int(Zf.shape[1]) if Zf is not None else -1
    out["k_knn"] = int(k_knn)

    # ---- Paired retrieval metrics (RNA vs ATAC latents) ----
    out["FOSCTTM_rankfrac"] = foscttm_rank_fraction(Zr, Za)

    fosk, rec = foscttm_at_k(Zr, Za, ks=tuple(int(x) for x in fos_ks))
    out.update(fosk)      # FOSCTTM@{...}
    out.update(rec)       # recall_*@k

    # keep legacy key if you want it:
    out["FOSCTTM"] = out["FOSCTTM_rankfrac"]

    # ---- kNN accuracy within each space (LOO) ----
    # (This is the clean "kNN ACC" people usually mean.)
    acc_r, f1_r, bal_r = knn_loo_accuracy(Zr, y_true, k=int(k_knn), metric="cosine", normalize=True)
    acc_a, f1_a, bal_a = knn_loo_accuracy(Za, y_true, k=int(k_knn), metric="cosine", normalize=True)
    acc_f, f1_f, bal_f = knn_loo_accuracy(Zf, y_true, k=int(k_knn), metric="cosine", normalize=True)

    out["kNN_LOO_acc_rna"] = float(acc_r)
    out["kNN_LOO_macroF1_rna"] = float(f1_r)
    out["kNN_LOO_balacc_rna"] = float(bal_r)

    out["kNN_LOO_acc_atac"] = float(acc_a)
    out["kNN_LOO_macroF1_atac"] = float(f1_a)
    out["kNN_LOO_balacc_atac"] = float(bal_a)

    out["kNN_LOO_acc_fused"] = float(acc_f)
    out["kNN_LOO_macroF1_fused"] = float(f1_f)
    out["kNN_LOO_balacc_fused"] = float(bal_f)

    # ---- Optional: cross-space "transfer" (source labels must come from SOURCE set) ----
    # In paired-only, the "source" and "target" are the same cells, so this becomes
    # more of a neighborhood-consistency score than true transfer.
    # We keep it anyway since you had it.
    pred_r2a = knn_label_transfer(Z_src=Zr, y_src=y_true, Z_tgt=Za, k=int(k_knn), metric="cosine", normalize=True)
    pred_a2r = knn_label_transfer(Z_src=Za, y_src=y_true, Z_tgt=Zr, k=int(k_knn), metric="cosine", normalize=True)

    out["LT_RNA2ATAC_acc"] = float(accuracy_score(y_true, pred_r2a))
    out["LT_ATAC2RNA_acc"] = float(accuracy_score(y_true, pred_a2r))
    out["LT_RNA2ATAC_macroF1"] = float(f1_score(y_true, pred_r2a, average="macro"))
    out["LT_ATAC2RNA_macroF1"] = float(f1_score(y_true, pred_a2r, average="macro"))

    # ---- Clustering + silhouette on fused ----
    kmeans_k = int(len(np.unique(y_true)))
    out.update(clustering_metrics(Zf, y_true, kmeans_k=kmeans_k, seed=int(seed)))

    # ---- Mixing score: stack rna + atac and measure neighbor modality purity ----
    Z_mix = np.vstack([Zr, Za])
    m_lab = np.array([0] * Zr.shape[0] + [1] * Za.shape[0], dtype=np.int64)
    out[f"mixing_1minus_samefrac_k{int(mix_k)}"] = modality_mixing_score(
        Z_mix, m_lab, k=int(mix_k), metric="cosine", normalize=True
    )

    # handy counts
    out["n_labels"] = int(len(np.unique(y_true)))
    out["kmeans_k"] = int(kmeans_k)

    return out


# %% [code]
# =============================================================================
# ---- 12) Param sweep spec ----
# =============================================================================
from __future__ import annotations
from dataclasses import dataclass
from typing import Optional

@dataclass(frozen=True)
class SweepPoint:
    name: str
    cfg_patch: dict
    model_kwargs: dict
    loss_mode: str = "v1"
    v1_recon: str = "moe"

def _fmt(x: float) -> str:
    # 0.25 -> "0p25", -1.0 -> "m1p0"
    #return str(float(x)).replace(".", "p").replace("-", "m")
    return str(float(x)).replace("p", ".").replace("m", "-")

def _sp(
    name: str,
    *,
    cfg_patch: Optional[dict] = None,
    model_kwargs: Optional[dict] = None,
    loss_mode: str = "v1",
    v1_recon: str = "moe",
) -> SweepPoint:
    return SweepPoint(
        name=name,
        cfg_patch={} if cfg_patch is None else dict(cfg_patch),
        model_kwargs={} if model_kwargs is None else dict(model_kwargs),
        loss_mode=str(loss_mode),
        v1_recon=str(v1_recon),
    )

def _filter_cfg_patch(cfg_patch: dict, *, allowed_keys: Optional[set[str]] = None) -> dict:
    """
    Optional safety: if you pass allowed_keys (e.g. from UniVIConfig.__dataclass_fields__),
    we drop unknown keys so sweeps don't crash on older config versions.
    """
    if not cfg_patch:
        return {}
    if allowed_keys is None:
        return dict(cfg_patch)
    out = {}
    for k, v in cfg_patch.items():
        if k in allowed_keys:
            out[k] = v
    return out


# -----------------------------------------------------------------------------
# Value grids (easy to edit)
# -----------------------------------------------------------------------------
BETA_GRID   = [0.0, 0.25, 0.5, 0.75, 1.0, 1.25, 1.5, 2.0, 3.0, 4.0, 4.35, 6.0, 8.0, 12.0, 18.0, 24.0, 48.0, 64.0, 128.0]
GAMMA_GRID  = [0.0, 0.25, 0.5, 0.75, 1.0, 1.25, 1.5, 2.0, 3.0, 4.0, 4.35, 6.0, 8.0, 12.0, 18.0, 24.0, 48.0, 64.0, 128.0]
LATENT_DIMS = [2, 4, 6, 8, 10, 12, 14, 16, 18, 20, 24, 30, 36, 48, 64, 128]

# Dropout grids
ENC_DROPOUT_GRID = [0.0, 0.025, 0.05, 0.075, 0.10, 0.125, 0.15, 0.175, 0.20, 0.25, 0.35, 0.50]
DEC_DROPOUT_GRID = [0.0, 0.025, 0.05, 0.075, 0.10, 0.125, 0.15, 0.175, 0.20, 0.25, 0.35, 0.50]
ENCDEC_COUPLED   = [0.0, 0.025, 0.05, 0.075, 0.10, 0.125, 0.15, 0.175, 0.20, 0.25, 0.35, 0.50]


def make_sweep_points(*, cfg_allowed_keys: Optional[set[str]] = None) -> list[SweepPoint]:
    """
    If you pass cfg_allowed_keys, cfg_patch keys will be filtered (nice for backward compatibility).
    Example:
        cfg_allowed_keys = set(UniVIConfig.__dataclass_fields__.keys())
        SWEEP_POINTS = make_sweep_points(cfg_allowed_keys=cfg_allowed_keys)
    """
    pts: list[SweepPoint] = []

    # --- baselines ---
    pts += [
        _sp("baseline", v1_recon="moe"),
        _sp("no_moe_agg", v1_recon="avg"),
    ]

    # --- alignment off ---
    # In this UniVI version there is no symmetric_align model kwarg.
    # So "no alignment" is gamma=0.
    pts.append(_sp("no_align", cfg_patch=_filter_cfg_patch({"gamma": 0.0}, allowed_keys=cfg_allowed_keys), v1_recon="moe"))
    '''
    # --- beta sweep ---
    for b in BETA_GRID:
        pts.append(_sp(
            f"beta_{_fmt(b)}",
            cfg_patch=_filter_cfg_patch({"beta": float(b)}, allowed_keys=cfg_allowed_keys),
            v1_recon="moe",
        ))

    # --- gamma sweep ---
    for g in GAMMA_GRID:
        pts.append(_sp(
            f"gamma_{_fmt(g)}",
            cfg_patch=_filter_cfg_patch({"gamma": float(g)}, allowed_keys=cfg_allowed_keys),
            v1_recon="moe",
        ))
    '''
    
    # --- beta x gamma grid (ALL combos) ---
    # Optionally skip the (baseline beta, baseline gamma) combo to avoid duplicating "baseline"
    BASE_BETA = float(getattr(UniVIConfig, "__dummy__", 0.0) or 1.25)   # optional: just for naming logic
    BASE_GAMMA = float(getattr(UniVIConfig, "__dummy__", 0.0) or 4.35) # optional: just for naming logic

    SKIP_BASELINE_COMBO = False  # set True if you want to not include baseline beta/gamma combos

    for b in BETA_GRID:
        for g in GAMMA_GRID:
            if SKIP_BASELINE_COMBO and (float(b) == float(BASE_BETA)) and (float(g) == float(BASE_GAMMA)):
                continue

            pts.append(_sp(
                f"bg_beta_{_fmt(b)}__gamma_{_fmt(g)}",
                cfg_patch=_filter_cfg_patch({"beta": float(b), "gamma": float(g)}, allowed_keys=cfg_allowed_keys),
                v1_recon="moe",
            ))


    # --- annealing variants ---
    pts += [
        _sp("no_prior_anneal",
            cfg_patch=_filter_cfg_patch({"kl_anneal_start": 0, "kl_anneal_end": 0}, allowed_keys=cfg_allowed_keys),
            v1_recon="moe"),
        _sp("no_align_anneal",
            cfg_patch=_filter_cfg_patch({"align_anneal_start": 0, "align_anneal_end": 0}, allowed_keys=cfg_allowed_keys),
            v1_recon="moe"),
        _sp("very_early_anneal",
            cfg_patch=_filter_cfg_patch({
                "kl_anneal_start": 0, "kl_anneal_end": 25,
                "align_anneal_start": 10, "align_anneal_end": 35,
            }, allowed_keys=cfg_allowed_keys),
            v1_recon="moe"),
        _sp("early_anneal",
            cfg_patch=_filter_cfg_patch({
                "kl_anneal_start": 0, "kl_anneal_end": 55,
                "align_anneal_start": 25, "align_anneal_end": 90,
            }, allowed_keys=cfg_allowed_keys),
            v1_recon="moe"),
        _sp("late_anneal",
            cfg_patch=_filter_cfg_patch({
                "kl_anneal_start": 50, "kl_anneal_end": 105,
                "align_anneal_start": 75, "align_anneal_end": 140,
            }, allowed_keys=cfg_allowed_keys),
            v1_recon="moe"),
        _sp("very_late_anneal",
            cfg_patch=_filter_cfg_patch({
                "kl_anneal_start": 100, "kl_anneal_end": 155,
                "align_anneal_start": 125, "align_anneal_end": 190,
            }, allowed_keys=cfg_allowed_keys),
            v1_recon="moe"),
    ]

    # --- latent dim sweep ---
    for d in LATENT_DIMS:
        pts.append(_sp(
            f"latent_{d}",
            cfg_patch=_filter_cfg_patch({"latent_dim": int(d)}, allowed_keys=cfg_allowed_keys),
            v1_recon="moe",
        ))

    # --- dropout sweeps ---
    # NOTE: key names must match UniVIConfig. Common possibilities:
    #   encoder_dropout / decoder_dropout
    #   enc_dropout / dec_dropout
    # Adjust the keys below to match your config.
    ENC_KEY = "encoder_dropout"
    DEC_KEY = "decoder_dropout"

    for p in ENC_DROPOUT_GRID:
        pts.append(_sp(
            f"drop_enc_{_fmt(p)}",
            cfg_patch=_filter_cfg_patch({ENC_KEY: float(p)}, allowed_keys=cfg_allowed_keys),
            v1_recon="moe",
        ))

    for p in DEC_DROPOUT_GRID:
        pts.append(_sp(
            f"drop_dec_{_fmt(p)}",
            cfg_patch=_filter_cfg_patch({DEC_KEY: float(p)}, allowed_keys=cfg_allowed_keys),
            v1_recon="moe",
        ))

    for p in ENCDEC_COUPLED:
        pts.append(_sp(
            f"drop_encdec_{_fmt(p)}",
            cfg_patch=_filter_cfg_patch({ENC_KEY: float(p), DEC_KEY: float(p)}, allowed_keys=cfg_allowed_keys),
            v1_recon="moe",
        ))

    # --- sanity check: unique names ---
    names = [p.name for p in pts]
    if len(names) != len(set(names)):
        dupes = sorted({n for n in names if names.count(n) > 1})
        raise ValueError(f"Duplicate sweep point names: {dupes}")

    return pts


# Usage (recommended):
# cfg_allowed = set(UniVIConfig.__dataclass_fields__.keys())
# SWEEP_POINTS = make_sweep_points(cfg_allowed_keys=cfg_allowed)
SWEEP_POINTS = make_sweep_points(cfg_allowed_keys=None)

print("n sweep points:", len(SWEEP_POINTS))
print("example:", SWEEP_POINTS[:5])


# %% [code]
# =============================================================================
# ---- 13) Grid runner (paired-only) + outputs (uses min_epochs ES gating) ----
# =============================================================================
OUTDIR = "./results/fig10_ablation_scaling_all_combos"
os.makedirs(OUTDIR, exist_ok=True)

def apply_cfg_patch(cfg: UniVIConfig, patch: dict) -> UniVIConfig:
    cfg2 = copy.deepcopy(cfg)
    for k, v in (patch or {}).items():
        if hasattr(cfg2, k):
            setattr(cfg2, k, v)
        else:
            print(f"[warn] UniVIConfig has no attribute '{k}' (skipping patch)")
    return cfg2

def save_tsv(df: pd.DataFrame, path: str):
    df.to_csv(path, sep="\t", index=False)
    print("[wrote]", path)

def make_paired_loaders(*, base_dataset, train_idx, val_idx, test_idx, batch_size: int = 256, num_workers: int = 0):
    train_ds = IndexedDataset(base_dataset, indices=train_idx)
    val_ds   = IndexedDataset(base_dataset, indices=val_idx)
    test_ds  = IndexedDataset(base_dataset, indices=test_idx)

    train_loader = DataLoader(train_ds, batch_size=int(batch_size), shuffle=True,  drop_last=True,
                              num_workers=int(num_workers), collate_fn=collate_xdict_with_idx)
    val_loader   = DataLoader(val_ds,   batch_size=int(batch_size), shuffle=False, drop_last=False,
                              num_workers=int(num_workers), collate_fn=collate_xdict_with_idx)
    test_loader  = DataLoader(test_ds,  batch_size=int(batch_size), shuffle=False, drop_last=False,
                              num_workers=int(num_workers), collate_fn=collate_xdict_with_idx)

    info = {"train_paired": int(len(train_ds)), "train_unpaired": 0, "overlap_fraction": 1.0, "drop_modality": "none"}
    return train_loader, val_loader, test_loader, info

def _infer_min_epochs_from_cfg(cfg: UniVIConfig) -> int:
    kl_end = getattr(cfg, "kl_anneal_end", 0)
    al_end = getattr(cfg, "align_anneal_end", 0)
    try:
        return int(max(int(kl_end), int(al_end), 0))
    except Exception:
        return 0

def run_grid_paired_only(
    *,
    sweep_points,
    seed=0,
    batch_size=256,
    patience=100,
    min_delta=0.0,
    k_knn=5,
    fuse_mode="avg",
    num_workers=0,
    gate_es_by_anneal_end: bool = True,
    extra_min_epochs: int = 0,
):
    rna_dim  = int(rna_tr_pp.n_vars)
    atac_dim = int(atac_tr_lsi.n_vars)

    train_loader, val_loader, test_loader, info = make_paired_loaders(
        base_dataset=dataset,
        train_idx=train_idx,
        val_idx=val_idx,
        test_idx=test_idx,
        batch_size=int(batch_size),
        num_workers=int(num_workers),
    )

    xb0, _ = next(iter(val_loader))
    if not ("rna" in xb0 and "atac" in xb0):
        raise RuntimeError(f"Expected paired batches with keys ['rna','atac']; got {list(xb0.keys())}")

    rows = []
    for spoint in sweep_points:
        cfg0 = make_univi_cfg(rna_dim=rna_dim, atac_dim=atac_dim)
        cfg  = apply_cfg_patch(cfg0, spoint.cfg_patch)

        min_epochs = _infer_min_epochs_from_cfg(cfg) if gate_es_by_anneal_end else 0
        min_epochs = int(max(min_epochs + int(extra_min_epochs), 0))

        tr = train_one(
            train_loader=train_loader,
            val_loader=val_loader,
            univi_cfg=cfg,
            seed=int(seed),
            device=device,
            loss_mode=str(spoint.loss_mode),
            v1_recon=str(spoint.v1_recon),
            batch_size=int(batch_size),
            patience=int(patience),
            min_delta=float(min_delta),
            min_epochs=int(min_epochs),
            extra_model_kwargs=dict(spoint.model_kwargs),
        )

        met = evaluate_on_test(
            tr.model,
            test_loader,
            device=device,
            y_all=y_all,
            k_knn=int(k_knn),
            seed=int(seed),
        )

        row = {
            "sweep": spoint.name,
            "loss_mode": spoint.loss_mode,
            "v1_recon": spoint.v1_recon,
            "seed": int(seed),
            "batch_size": int(batch_size),
            "patience": int(patience),
            "min_delta": float(min_delta),
            "min_epochs_gate": int(min_epochs),
            "k_knn": int(k_knn),
            "fuse_mode": str(fuse_mode),
            "best_epoch": tr.best_epoch,
            "best_val": tr.best_val,
            "wall_seconds": tr.wall_seconds,
            "gpu_peak_mem_mb": tr.peak_mem_mb,
            "latent_dim": int(getattr(cfg, "latent_dim", -1)),
            "beta": float(getattr(cfg, "beta", np.nan)),
            "gamma": float(getattr(cfg, "gamma", np.nan)),
            "kl_anneal_start": int(getattr(cfg, "kl_anneal_start", -1)),
            "kl_anneal_end": int(getattr(cfg, "kl_anneal_end", -1)),
            "align_anneal_start": int(getattr(cfg, "align_anneal_start", -1)),
            "align_anneal_end": int(getattr(cfg, "align_anneal_end", -1)),
        }
        row.update(info)
        row.update(met)
        rows.append(row)

        fos = row.get("FOSCTTM", np.nan)
        fos_str = "nan" if (fos is None or (isinstance(fos, float) and np.isnan(fos))) else f"{fos:.4f}"
        print(f"[done] sweep={spoint.name:>20}  min_epochs={min_epochs:>4}  FOSCTTM={fos_str}  wall={tr.wall_seconds:.1f}s")

        del tr
        gc.collect()
        if torch.cuda.is_available():
            torch.cuda.empty_cache()

    return pd.DataFrame(rows)

df_grid = run_grid_paired_only(
    sweep_points=SWEEP_POINTS,
    seed=0,
    batch_size=256,
    patience=50,
    min_delta=0.0,
    k_knn=3,
    fuse_mode="moe",
    num_workers=0,
    gate_es_by_anneal_end=True,
    extra_min_epochs=0,
)

save_tsv(df_grid, os.path.join(OUTDIR, "param_grid_paired_only.tsv"))
df_grid.to_csv(os.path.join(OUTDIR, "param_grid_paired_only.csv"), index=False)
df_grid.head()


# %% [code]
print(SWEEP_POINTS[0])


# %% [code]
# =============================================================================
# ---- 14) Scaling curve (paired-only; uses min_epochs ES gating)
#      Self-contained utilities + optional oversampling beyond available train cells.
#      - MPS: torch.mps current_allocated_memory + driver_allocated_memory (if available)
#      - CUDA: sampled allocated/reserved + true peak stats
#      - CPU: RSS + CPU% via psutil (optional)
# =============================================================================
import os, time, gc, threading
import numpy as np
import pandas as pd
import torch

# --- psutil (optional, but recommended) ---
try:
    import psutil
    _proc = psutil.Process(os.getpid())
except Exception:
    psutil = None
    _proc = None

def get_cpu_rss_mb():
    if _proc is None:
        return None
    try:
        return float(_proc.memory_info().rss / (1024**2))
    except Exception:
        return None

def _prime_cpu_percent():
    if psutil is None or _proc is None:
        return
    try:
        _proc.cpu_percent(interval=None)
    except Exception:
        pass

def resolve_device(device=None):
    """
    Supports: None, 'mps', 'cpu', 'cuda', 'cuda:0', torch.device(...)
    Default preference: MPS (if available) -> CUDA -> CPU
    """
    if device is None:
        if hasattr(torch.backends, "mps") and torch.backends.mps.is_available():
            device = torch.device("mps")
        elif torch.cuda.is_available():
            device = torch.device("cuda:0")
        else:
            device = torch.device("cpu")
    elif isinstance(device, str):
        device = torch.device(device)

    if device.type == "cuda":
        idx = 0 if device.index is None else int(device.index)
        try:
            torch.cuda.set_device(idx)
        except Exception:
            pass
        device = torch.device(f"cuda:{idx}")

    return device

def _mps_mem_mb():
    """
    Returns (allocated_mb, driver_allocated_mb) for MPS if available else (None, None)
    """
    if not (hasattr(torch, "mps") and hasattr(torch.mps, "current_allocated_memory")):
        return (None, None)
    try:
        a = float(torch.mps.current_allocated_memory() / (1024**2))
        d = float(torch.mps.driver_allocated_memory() / (1024**2)) if hasattr(torch.mps, "driver_allocated_memory") else None
        return (a, d)
    except Exception:
        return (None, None)

def _cuda_reset_peaks(device):
    if device.type != "cuda" or not torch.cuda.is_available():
        return
    try:
        torch.cuda.reset_peak_memory_stats(device)
        torch.cuda.empty_cache()
    except Exception:
        pass

def _cuda_peaks_mb(device):
    """
    Returns (peak_alloc_mb, peak_reserved_mb) for CUDA else (None, None)
    """
    if device.type != "cuda" or not torch.cuda.is_available():
        return (None, None)
    try:
        alloc = float(torch.cuda.max_memory_allocated(device) / (1024**2))
        reserv = float(torch.cuda.max_memory_reserved(device) / (1024**2))
        return (alloc, reserv)
    except Exception:
        return (None, None)

class ResourceMonitor:
    """
    Samples CPU% + RSS, and (if possible) GPU memory.
    - MPS: samples torch.mps current_allocated_memory + driver_allocated_memory
    - CUDA: samples torch.cuda memory_allocated + memory_reserved
    """
    def __init__(self, device, sample_every_s: float = 0.2):
        self.device = device
        self.sample_every_s = float(sample_every_s)

        self._stop = threading.Event()
        self._thr = None

        # CPU samples
        self.cpu_percent = []
        self.rss_mb = []

        # GPU samples (backend-dependent)
        self.gpu_alloc_mb = []     # MPS allocated / CUDA allocated
        self.gpu_driver_mb = []    # MPS driver_allocated (None on CUDA)
        self.gpu_reserved_mb = []  # CUDA reserved (None on MPS)

        _prime_cpu_percent()

    def start(self):
        self._thr = threading.Thread(target=self._run, daemon=True)
        self._thr.start()
        return self

    def stop(self):
        self._stop.set()
        if self._thr is not None:
            self._thr.join(timeout=2.0)

    def _run(self):
        while not self._stop.is_set():
            # CPU
            if psutil is not None and _proc is not None:
                try:
                    self.cpu_percent.append(float(_proc.cpu_percent(interval=None)))
                    self.rss_mb.append(get_cpu_rss_mb())
                except Exception:
                    pass

            # GPU
            if self.device.type == "mps":
                a, d = _mps_mem_mb()
                self.gpu_alloc_mb.append(a)
                self.gpu_driver_mb.append(d)
            elif self.device.type == "cuda" and torch.cuda.is_available():
                try:
                    self.gpu_alloc_mb.append(float(torch.cuda.memory_allocated(self.device) / (1024**2)))
                    self.gpu_reserved_mb.append(float(torch.cuda.memory_reserved(self.device) / (1024**2)))
                except Exception:
                    pass

            time.sleep(self.sample_every_s)

    @staticmethod
    def _clean(xs):
        return [x for x in xs if x is not None and not (isinstance(x, float) and np.isnan(x))]

    def summary(self):
        def _mean(xs):
            xs = self._clean(xs)
            return float(np.mean(xs)) if xs else None

        def _max(xs):
            xs = self._clean(xs)
            return float(np.max(xs)) if xs else None

        return {
            "cpu_percent_mean": _mean(self.cpu_percent),
            "cpu_percent_max": _max(self.cpu_percent),
            "cpu_rss_peak_mb": _max(self.rss_mb),

            "gpu_alloc_mean_mb": _mean(self.gpu_alloc_mb),
            "gpu_alloc_peak_mb": _max(self.gpu_alloc_mb),

            # MPS-only
            "mps_driver_mean_mb": _mean(self.gpu_driver_mb),
            "mps_driver_peak_mb": _max(self.gpu_driver_mb),

            # CUDA-only (sampled)
            "cuda_reserved_mean_mb_sampled": _mean(self.gpu_reserved_mb),
            "cuda_reserved_peak_mb_sampled": _max(self.gpu_reserved_mb),
        }

def sample_train_indices(idx, n_target, seed=0, allow_oversample=True):
    """
    Returns indices of length n_target.
    - If n_target <= len(idx): sample without replacement.
    - If n_target >  len(idx): if allow_oversample, sample WITH replacement; else return idx (cap).
    """
    rng = np.random.default_rng(int(seed))
    idx = np.asarray(idx, dtype=np.int64)
    n_target = int(n_target)

    if n_target <= 0:
        return idx[:0]

    if n_target <= len(idx):
        out = rng.choice(idx, size=n_target, replace=False)
        return out.astype(np.int64)

    # n_target > len(idx)
    if not allow_oversample:
        return idx.astype(np.int64)

    out = rng.choice(idx, size=n_target, replace=True)
    return out.astype(np.int64)

def _empty_backend_cache(device):
    if device.type == "cuda" and torch.cuda.is_available():
        torch.cuda.empty_cache()
    elif device.type == "mps" and hasattr(torch, "mps") and hasattr(torch.mps, "empty_cache"):
        try:
            torch.mps.empty_cache()
        except Exception:
            pass

def run_scaling_paired_only(
    *,
    n_cells_grid=(50, 100, 250, 500, 1000, 2000, 4000, 6000, 8000, 10000),
    seed=0,
    batch_size=256,
    spoint=None,
    patience=100,
    min_delta=0.0,
    num_workers=0,
    gate_es_by_anneal_end: bool = True,
    extra_min_epochs: int = 0,
    device=None,
    sample_every_s: float = 0.2,
    allow_oversample: bool = True,
):
    if spoint is None:
        spoint = SweepPoint(name="baseline", cfg_patch={}, model_kwargs={}, loss_mode="v1", v1_recon="moe")

    device = resolve_device(device)

    rna_dim  = int(rna_tr_pp.n_vars)
    atac_dim = int(atac_tr_lsi.n_vars)

    rows = []
    for n_cells in n_cells_grid:
        # --- choose training indices (possibly oversampled) ---
        tr_idx2 = sample_train_indices(
            train_idx,
            n_target=int(n_cells),
            seed=int(seed),
            allow_oversample=bool(allow_oversample),
        )
        n_eff = int(len(tr_idx2))
        n_unique = int(np.unique(tr_idx2).size)
        oversample_factor = float(n_eff / max(n_unique, 1))

        train_loader, val_loader, test_loader, _info = make_paired_loaders(
            base_dataset=dataset,
            train_idx=tr_idx2,
            val_idx=val_idx,
            test_idx=test_idx,
            batch_size=int(batch_size),
            num_workers=int(num_workers),
        )

        cfg0 = make_univi_cfg(rna_dim=rna_dim, atac_dim=atac_dim)
        cfg  = apply_cfg_patch(cfg0, spoint.cfg_patch)

        min_epochs = _infer_min_epochs_from_cfg(cfg) if gate_es_by_anneal_end else 0
        min_epochs = int(max(min_epochs + int(extra_min_epochs), 0))

        cpu_before = get_cpu_rss_mb()

        # Reset CUDA peaks if on CUDA (no-op on MPS/CPU)
        _cuda_reset_peaks(device)

        # Start monitor thread
        mon = ResourceMonitor(device=device, sample_every_s=sample_every_s).start()

        t0 = time.perf_counter()
        tr = train_one(
            train_loader=train_loader,
            val_loader=val_loader,
            univi_cfg=cfg,
            seed=int(seed),
            device=device,
            loss_mode=str(spoint.loss_mode),
            v1_recon=str(spoint.v1_recon),
            batch_size=int(batch_size),
            patience=int(patience),
            min_delta=float(min_delta),
            min_epochs=int(min_epochs),
            extra_model_kwargs=dict(spoint.model_kwargs),
        )
        t1 = time.perf_counter()

        mon.stop()
        mon_s = mon.summary()

        cpu_after = get_cpu_rss_mb()

        # CUDA-only peaks (None on MPS/CPU)
        cuda_peak_alloc_mb, cuda_peak_reserved_mb = _cuda_peaks_mb(device)

        # MPS "snapshot" after run (helpful if sampling missed a spike)
        mps_alloc_after_mb, mps_driver_after_mb = _mps_mem_mb() if device.type == "mps" else (None, None)

        tr_peak = getattr(tr, "peak_mem_mb", None)  # optional field from your trainer

        rows.append({
            # --- scaling axes ---
            "n_train_cells_eff": n_eff,          # may include duplicates
            "n_train_cells_unique": n_unique,    # unique cells
            "oversample_factor": oversample_factor,
            "allow_oversample": bool(allow_oversample),

            # --- sweep/config ---
            "sweep": spoint.name,
            "loss_mode": spoint.loss_mode,
            "v1_recon": spoint.v1_recon,
            "seed": int(seed),
            "batch_size": int(batch_size),
            "patience": int(patience),
            "min_delta": float(min_delta),
            "min_epochs_gate": int(min_epochs),

            # --- timing ---
            "wall_seconds": float(t1 - t0),

            # --- device/backend ---
            "device": str(device),
            "backend": device.type,

            # --- GPU mem (backend-aware; sampled) ---
            "gpu_alloc_peak_mb_sampled": mon_s["gpu_alloc_peak_mb"],  # CUDA allocated / MPS allocated
            "gpu_alloc_mean_mb_sampled": mon_s["gpu_alloc_mean_mb"],
            "mps_driver_peak_mb_sampled": mon_s["mps_driver_peak_mb"],  # MPS only
            "mps_driver_mean_mb_sampled": mon_s["mps_driver_mean_mb"],
            "mps_alloc_after_mb": mps_alloc_after_mb,   # MPS only
            "mps_driver_after_mb": mps_driver_after_mb, # MPS only

            # --- CUDA-only peaks/stats ---
            "cuda_peak_alloc_mb": cuda_peak_alloc_mb,
            "cuda_peak_reserved_mb": cuda_peak_reserved_mb,
            "cuda_reserved_peak_mb_sampled": mon_s["cuda_reserved_peak_mb_sampled"],
            "cuda_reserved_mean_mb_sampled": mon_s["cuda_reserved_mean_mb_sampled"],

            # --- optional trainer-reported value ---
            "gpu_peak_mem_mb_trainone": tr_peak,

            # --- CPU metrics ---
            "cpu_rss_mb_before": cpu_before,
            "cpu_rss_mb_after": cpu_after,
            "cpu_rss_mb_delta": (None if (cpu_before is None or cpu_after is None) else float(cpu_after - cpu_before)),
            "cpu_rss_peak_mb_sampled": mon_s["cpu_rss_peak_mb"],
            "cpu_percent_mean": mon_s["cpu_percent_mean"],
            "cpu_percent_max": mon_s["cpu_percent_max"],

            # --- training outputs ---
            "best_epoch": getattr(tr, "best_epoch", None),
            "best_val": getattr(tr, "best_val", None),

            # --- bookkeeping ---
            "train_paired": int(n_eff),
            "train_unpaired": 0,
            "overlap_fraction": 1.0,
            "drop_modality": "none",
            "latent_dim": int(getattr(cfg, "latent_dim", -1)),
            "beta": float(getattr(cfg, "beta", np.nan)),
            "gamma": float(getattr(cfg, "gamma", np.nan)),
            "kl_anneal_start": int(getattr(cfg, "kl_anneal_start", -1)),
            "kl_anneal_end": int(getattr(cfg, "kl_anneal_end", -1)),
            "align_anneal_start": int(getattr(cfg, "align_anneal_start", -1)),
            "align_anneal_end": int(getattr(cfg, "align_anneal_end", -1)),
        })

        print(
            f"[scale] n_eff={n_eff:>6} n_unique={n_unique:>6} x{oversample_factor:>4.1f} "
            f"min_epochs={min_epochs:>4} wall={t1-t0:>7.1f}s backend={device.type} "
            f"gpu_alloc_peak={mon_s['gpu_alloc_peak_mb']}MB mps_driver_peak={mon_s['mps_driver_peak_mb']}MB "
            f"cpu_rss_peak={mon_s['cpu_rss_peak_mb']}MB"
        )

        # cleanup
        del tr
        gc.collect()
        _empty_backend_cache(device)

    return pd.DataFrame(rows)

# --- example call ---
df_scale = run_scaling_paired_only(
    n_cells_grid=(10, 25, 50, 100, 200, 300, 400, 500, 600, 700, 800, 900, 1000, 1500, 2000, 3000, 4000, 5000, 6000, 7000, 8000, 9000, 10000, 15000, 20000, 30000, 40000, 50000, 100000, 500000),
    #n_cells_grid=(50000, 100000),
    seed=0,
    batch_size=256,
    spoint=SWEEP_POINTS[0],
    patience=50,
    min_delta=0.0,
    num_workers=0,
    gate_es_by_anneal_end=True,
    extra_min_epochs=0,
    device="mps",
    sample_every_s=0.2,
    allow_oversample=True,   # <- set False to keep the original "cap at max unique" behavior
)

save_tsv(df_scale, os.path.join(OUTDIR, "scaling_curve.tsv"))
df_scale.to_csv(os.path.join(OUTDIR, "scaling_curve.csv"), index=False)
df_scale


# %% [code]
# =============================================================================
# ---- 15) Plotting (paired-only) ----
# =============================================================================
import matplotlib.pyplot as plt
import matplotlib as mpl

# ---- global defaults ----
mpl.rcParams["figure.figsize"] = (14.0, 12.0)     # default size for plt.figure()
mpl.rcParams["figure.dpi"] = 300                  # DPI for on-screen display
mpl.rcParams["savefig.dpi"] = 300                 # default DPI for saved files (if you don't override)
mpl.rcParams["savefig.bbox"] = "tight"            # nice default cropping

def _savefig(savepath):
    if savepath:
        plt.savefig(savepath, dpi=300)
        print("[wrote]", savepath)

def barplot_best(df, metric, *, top_n=30, ascending=None, savepath=None, title=None):
    if metric not in df.columns:
        print(f"[skip] missing metric: {metric}")
        return
    d = df.copy()
    d[metric] = pd.to_numeric(d[metric], errors="coerce")
    d = d.dropna(subset=[metric])

    if ascending is None:
        m = metric.lower()
        ascending = any(s in m for s in ["loss", "error", "dist", "foscttm", "rmse", "mae"])

    d = d.sort_values(metric, ascending=bool(ascending)).head(int(top_n))

    # ---- plot ----
    fig, ax = plt.subplots(figsize=(10, 0.32 * len(d) + 2))

    y = d["sweep"].astype(str).to_numpy()
    x = d[metric].astype(float).to_numpy()

    ax.barh(y, x, height=0.9)  # slightly thicker bars (optional but looks nicer)

    ax.set_xlabel(metric)
    ax.set_ylabel("sweep")
    ax.set_title(title if title is not None else f"Top {top_n}: {metric}")

    # ---- remove the big top/bottom gap ----
    ax.margins(y=0)                 # kill categorical padding
    ax.set_ylim(-0.5, len(d) - 0.5) # clamp to first/last bar

    # If you want labels in the same order as your sorted rows (best at top):
    ax.invert_yaxis()

    fig.tight_layout()
    _savefig(savepath)
    plt.show()

def scatter_tradeoff(df, x, y, *, annotate_top=0, savepath=None, title=None):
    if x not in df.columns or y not in df.columns:
        print(f"[skip] missing: {x} or {y}")
        return

    d = df[["sweep", x, y]].copy()
    d[x] = pd.to_numeric(d[x], errors="coerce")
    d[y] = pd.to_numeric(d[y], errors="coerce")
    d = d.dropna()

    plt.figure(figsize=(6.5, 5))
    plt.scatter(d[x].astype(float), d[y].astype(float))
    plt.xlabel(x)
    plt.ylabel(y)
    plt.title(title if title is not None else f"{y} vs {x}")
    plt.tight_layout()

    if int(annotate_top) > 0:
        y_asc = any(s in y.lower() for s in ["loss", "error", "dist", "foscttm", "rmse", "mae"])
        dd = d.sort_values(y, ascending=bool(y_asc)).head(int(annotate_top))
        for _, r in dd.iterrows():
            plt.annotate(str(r["sweep"]), (float(r[x]), float(r[y])), fontsize=8)

    _savefig(savepath)
    plt.show()

def heatmap_hparams_vs_metric(df, metric, hp_x="beta", hp_y="gamma", *, agg="median", savepath=None, title=None):
    needed = [metric, hp_x, hp_y]
    for c in needed:
        if c not in df.columns:
            print(f"[skip] missing col: {c}")
            return

    d = df[[hp_x, hp_y, metric]].copy()
    d[hp_x] = pd.to_numeric(d[hp_x], errors="coerce")
    d[hp_y] = pd.to_numeric(d[hp_y], errors="coerce")
    d[metric] = pd.to_numeric(d[metric], errors="coerce")
    d = d.dropna()

    if agg == "mean":
        pv = d.groupby([hp_y, hp_x])[metric].mean().unstack(hp_x)
    elif agg == "min":
        pv = d.groupby([hp_y, hp_x])[metric].min().unstack(hp_x)
    elif agg == "max":
        pv = d.groupby([hp_y, hp_x])[metric].max().unstack(hp_x)
    else:
        pv = d.groupby([hp_y, hp_x])[metric].median().unstack(hp_x)

    plt.figure(figsize=(1.2*max(6, pv.shape[1]), 1.0*max(4, pv.shape[0])))
    plt.imshow(pv.values, aspect="auto")
    plt.xticks(range(pv.shape[1]), [str(x) for x in pv.columns], rotation=45, ha="right")
    plt.yticks(range(pv.shape[0]), [str(y) for y in pv.index])
    plt.xlabel(hp_x)
    plt.ylabel(hp_y)
    plt.title(title if title is not None else f"{metric} heatmap ({agg})")
    plt.colorbar(label=metric)
    plt.tight_layout()
    _savefig(savepath)
    plt.show()

def plot_scaling_multi(df, *, saveprefix=None):
    if "n_train_cells" in df.columns and "wall_seconds" in df.columns:
        d = df.sort_values("n_train_cells")
        plt.figure(figsize=(6,4))
        plt.plot(d["n_train_cells"], d["wall_seconds"], marker="o")
        plt.xlabel("# train cells")
        plt.ylabel("wall seconds (train)")
        plt.title("Scaling: runtime vs #cells")
        plt.tight_layout()
        _savefig(None if saveprefix is None else f"{saveprefix}_runtime.png")
        plt.show()

        plt.figure(figsize=(6,4))
        thr = d["n_train_cells"].astype(float) / np.maximum(d["wall_seconds"].astype(float), 1e-9)
        plt.plot(d["n_train_cells"], thr, marker="o")
        plt.xlabel("# train cells")
        plt.ylabel("train throughput (cells/sec)")
        plt.title("Scaling: throughput vs #cells")
        plt.tight_layout()
        _savefig(None if saveprefix is None else f"{saveprefix}_throughput.png")
        plt.show()

    if "n_train_cells" in df.columns and "gpu_peak_mem_mb" in df.columns:
        d = df.sort_values("n_train_cells")
        plt.figure(figsize=(6,4))
        plt.plot(d["n_train_cells"], d["gpu_peak_mem_mb"], marker="o")
        plt.xlabel("# train cells")
        plt.ylabel("GPU peak mem (MB)")
        plt.title("Scaling: GPU peak memory vs #cells")
        plt.tight_layout()
        _savefig(None if saveprefix is None else f"{saveprefix}_gpu_mem.png")
        plt.show()

    if "n_train_cells" in df.columns and "cpu_rss_mb_delta" in df.columns:
        d = df.sort_values("n_train_cells")
        plt.figure(figsize=(6,4))
        plt.plot(d["n_train_cells"], d["cpu_rss_mb_delta"], marker="o")
        plt.xlabel("# train cells")
        plt.ylabel("CPU RSS delta (MB)")
        plt.title("Scaling: CPU memory delta vs #cells")
        plt.tight_layout()
        _savefig(None if saveprefix is None else f"{saveprefix}_cpu_mem_delta.png")
        plt.show()

# ---- leaderboards ----
barplot_best(
    df_grid, "FOSCTTM", top_n=50,
    savepath=os.path.join(OUTDIR, "figures", "grid_top_FOSCTTM.png"),
    title="Grid leaderboard: FOSCTTM (lower is better)"
)

for m in ["LT_RNA2ATAC_macroF1", "LT_ATAC2RNA_macroF1", "mixing_1minus_samefrac_k15"]:
    if m in df_grid.columns:
        barplot_best(
            df_grid, m, top_n=50, ascending=False,
            savepath=os.path.join(OUTDIR, "figures", f"grid_top_{m}.png")
        )

# ---- tradeoffs ----
if "FOSCTTM" in df_grid.columns and "LT_RNA2ATAC_macroF1" in df_grid.columns:
    scatter_tradeoff(
        df_grid, "FOSCTTM", "LT_RNA2ATAC_macroF1",
        annotate_top=8,
        savepath=os.path.join(OUTDIR, "figures", "tradeoff_LTmacroF1_vs_FOSCTTM.png"),
        title="Tradeoff: label transfer vs FOSCTTM"
    )

if "FOSCTTM" in df_grid.columns and "mixing_1minus_samefrac_k15" in df_grid.columns:
    scatter_tradeoff(
        df_grid, "FOSCTTM", "mixing_1minus_samefrac_k15",
        annotate_top=8,
        savepath=os.path.join(OUTDIR, "figures", "tradeoff_mixing_vs_FOSCTTM.png"),
        title="Tradeoff: mixing vs FOSCTTM"
    )

# ---- hyperparameter maps ----
if "beta" in df_grid.columns and "gamma" in df_grid.columns:
    if "FOSCTTM" in df_grid.columns:
        heatmap_hparams_vs_metric(
            df_grid, "FOSCTTM", hp_x="beta", hp_y="gamma", agg="median",
            savepath=os.path.join(OUTDIR, "figures", "heatmap_FOSCTTM_beta_gamma.png"),
            title="FOSCTTM vs (beta, gamma)"
        )
    if "LT_RNA2ATAC_macroF1" in df_grid.columns:
        heatmap_hparams_vs_metric(
            df_grid, "LT_RNA2ATAC_macroF1", hp_x="beta", hp_y="gamma", agg="median",
            savepath=os.path.join(OUTDIR, "figures", "heatmap_LT_RNA2ATAC_macroF1_beta_gamma.png"),
            title="LT_RNA2ATAC_macroF1 vs (beta, gamma)"
        )

# ---- scaling plots ----
plot_scaling_multi(df_scale, saveprefix=os.path.join(OUTDIR, "scaling"))


# %% [code]
# =============================================================================
# ---- Tradeoff visualization suite (clean, scalable, paper-friendly) ----
# Produces 4 plots per (x, y):
#   A) density (hexbin) + optional highlight
#   B) scatter + Pareto front + few labels
#   C) scatter colored by beta
#   D) scatter colored by gamma
#
# Key ideas:
#   - Use hexbin for the "many points" overview (no clutter).
#   - Use Pareto front to identify true tradeoffs (not just top-by-y).
#   - Label only a few points (non-overlapping, near points) for readability.
#   - Only one continuous colormap per panel (beta OR gamma), never both.
# =============================================================================
import os
import numpy as np
import pandas as pd
import matplotlib.pyplot as plt
import matplotlib as mpl

mpl.rcParams["figure.dpi"] = 300
mpl.rcParams["savefig.dpi"] = 300
mpl.rcParams["savefig.bbox"] = "tight"

# ---------- helpers ----------
def _ensure_dir(p):
    if p:
        os.makedirs(os.path.dirname(p), exist_ok=True)

def _tight_limits(ax, x, y, pad_frac=0.03, top_pad_frac=0.05):
    x = np.asarray(x, dtype=float)
    y = np.asarray(y, dtype=float)
    xr = float(np.nanmax(x) - np.nanmin(x)) if len(x) else 1.0
    yr = float(np.nanmax(y) - np.nanmin(y)) if len(y) else 1.0
    xr = xr if xr > 0 else 1.0
    yr = yr if yr > 0 else 1.0
    ax.set_xlim(float(np.nanmin(x) - pad_frac * xr), float(np.nanmax(x) + pad_frac * xr))
    ax.set_ylim(float(np.nanmin(y) - pad_frac * yr), float(np.nanmax(y) + top_pad_frac * yr))

def _objective_direction(metric_name: str) -> str:
    """
    Return 'min' if lower is better, else 'max'. You can extend this list.
    """
    m = metric_name.lower()
    if any(s in m for s in ["loss", "error", "dist", "foscttm", "rmse", "mae", "rankfrac"]):
        return "min"
    return "max"

def _pareto_front(df, x, y, x_dir="min", y_dir="max"):
    """
    Compute Pareto-efficient points for two objectives.
      - x_dir: 'min' or 'max'
      - y_dir: 'min' or 'max'
    Returns df subset (Pareto front), sorted for nicer plotting.
    """
    d = df[[ "sweep", x, y ]].copy()

    # Convert to a "minimize both" problem:
    xx = d[x].to_numpy(dtype=float)
    yy = d[y].to_numpy(dtype=float)
    if x_dir == "max":
        xx = -xx
    if y_dir == "max":
        yy = -yy

    # Sort by xx then keep those with best (lowest) yy so far
    order = np.argsort(xx)
    xx_s = xx[order]
    yy_s = yy[order]
    keep = np.zeros(len(d), dtype=bool)

    best_y = np.inf
    for k, (xs, ys) in enumerate(zip(xx_s, yy_s)):
        if ys < best_y:
            best_y = ys
            keep[order[k]] = True

    front = d.loc[keep].copy()

    # Sort front for line plotting (by x in original direction)
    front = front.sort_values(x, ascending=(x_dir == "min"))
    return front

def _pick_nonoverlapping_points(ax, xs, ys, k, min_dist_px=28):
    """
    Greedy select up to k points separated by at least min_dist_px in DISPLAY coords.
    Preserves order of xs/ys (so provide "best-first" ordering).
    """
    pts = ax.transData.transform(np.c_[xs, ys])  # pixels
    chosen = []
    for i, p in enumerate(pts):
        if len(chosen) >= k:
            break
        ok = True
        for j in chosen:
            if np.linalg.norm(p - pts[j]) < float(min_dist_px):
                ok = False
                break
        if ok:
            chosen.append(i)
    return chosen

def _prep_xy(df, x, y, extra_cols=()):
    cols = ["sweep", x, y] + list(extra_cols)
    for c in ["sweep", x, y]:
        if c not in df.columns:
            raise KeyError(f"missing column: {c}")

    d = df[cols].copy()
    for c in [x, y] + list(extra_cols):
        if c in d.columns:
            d[c] = pd.to_numeric(d[c], errors="coerce")
    d = d.dropna(subset=[x, y])
    return d

# ---------- plotters ----------
def plot_tradeoff_density(
    df, x, y, *,
    savepath=None, title=None,
    gridsize=45,
    mincnt=1,
    figsize=(6.8, 5.0),
):
    """
    Big-picture plot: density via hexbin. Great when there are many points.
    """
    d = _prep_xy(df, x, y)

    fig, ax = plt.subplots(figsize=figsize)
    hb = ax.hexbin(
        d[x].to_numpy(dtype=float),
        d[y].to_numpy(dtype=float),
        gridsize=int(gridsize),
        mincnt=int(mincnt),
        linewidths=0.0,
    )

    ax.set_xlabel(x)
    ax.set_ylabel(y)
    ax.set_title(title if title is not None else f"{y} vs {x} (density)", pad=10)
    _tight_limits(ax, d[x].to_numpy(), d[y].to_numpy())

    cbar = fig.colorbar(hb, ax=ax, pad=0.02)
    cbar.set_label("count")

    fig.tight_layout()
    if savepath:
        _ensure_dir(savepath)
        plt.savefig(savepath, dpi=300, bbox_inches="tight")
        print("[wrote]", savepath)
    plt.show()

def plot_tradeoff_pareto(
    df, x, y, *,
    savepath=None, title=None,
    figsize=(6.8, 5.0),
    s=26, alpha=0.45,
    edge_alpha=0.12, edge_lw=0.25,
    label_k=10,
    label_candidates=40,
    label_min_dist_px=30,
):
    """
    Scatter + Pareto front + a few labels from the Pareto front.
    This is the most decision-useful view.
    """
    d = _prep_xy(df, x, y)

    x_dir = _objective_direction(x)   # usually x=FOSCTTM => 'min'
    y_dir = _objective_direction(y)   # usually y=macroF1 => 'max'
    front = _pareto_front(d, x, y, x_dir=x_dir, y_dir=y_dir)

    fig, ax = plt.subplots(figsize=figsize)

    # Background cloud
    ax.scatter(
        d[x].to_numpy(dtype=float),
        d[y].to_numpy(dtype=float),
        s=s, alpha=alpha,
        edgecolors=(0,0,0,edge_alpha),
        linewidths=edge_lw,
        rasterized=True,  # makes saved PDFs smaller if you export to pdf later
    )

    # Pareto front overlay (line + points)
    ax.plot(front[x].to_numpy(dtype=float), front[y].to_numpy(dtype=float), linewidth=1.2)
    ax.scatter(front[x].to_numpy(dtype=float), front[y].to_numpy(dtype=float), s=max(38, s+8), alpha=0.95)

    ax.set_xlabel(x)
    ax.set_ylabel(y)
    ax.set_title(title if title is not None else f"{y} vs {x} (Pareto front)", pad=10)
    _tight_limits(ax, d[x].to_numpy(), d[y].to_numpy())

    # Label a subset of Pareto points (best distributed)
    if int(label_k) > 0 and len(front) > 0:
        # choose candidate Pareto points (e.g., first N along front)
        cand = front.copy()
        if len(cand) > int(label_candidates):
            cand = cand.head(int(label_candidates))

        fig.canvas.draw()  # needed for coordinate transforms
        xs = cand[x].to_numpy(dtype=float)
        ys = cand[y].to_numpy(dtype=float)

        keep = _pick_nonoverlapping_points(ax, xs, ys, k=int(label_k), min_dist_px=float(label_min_dist_px))
        cand = cand.iloc[keep].copy()

        ymin, ymax = ax.get_ylim()
        y_thr = ymax - 0.10 * (ymax - ymin)  # flip label below point if near top

        for _, r in cand.iterrows():
            px, py = float(r[x]), float(r[y])
            dx, dy, va = 6, 6, "bottom"
            if py >= y_thr:
                dx, dy, va = 6, -10, "top"
            ax.annotate(
                str(r["sweep"]),
                xy=(px, py),
                xytext=(dx, dy),
                textcoords="offset points",
                fontsize=8,
                ha="left", va=va,
                bbox=dict(boxstyle="round,pad=0.15", fc="white", ec="none", alpha=0.75),
                clip_on=True,
            )

    fig.tight_layout()
    if savepath:
        _ensure_dir(savepath)
        plt.savefig(savepath, dpi=300, bbox_inches="tight")
        print("[wrote]", savepath)
    plt.show()

def plot_tradeoff_colored(
    df, x, y, *, color_by,
    savepath=None, title=None,
    figsize=(6.8, 5.0),
    s=26, alpha=0.65,
    edge_alpha=0.15, edge_lw=0.25,
    cmap="viridis",
    vmin=None, vmax=None,
):
    """
    One continuous variable per panel (beta OR gamma OR gpu mem). Clean + readable.
    """
    d = _prep_xy(df, x, y, extra_cols=(color_by,))
    if color_by not in d.columns:
        print(f"[skip] missing: {color_by}")
        return
    d = d.dropna(subset=[color_by])
    if len(d) == 0:
        print("[skip] no finite rows after cleaning.")
        return

    cvals = d[color_by].to_numpy(dtype=float)
    if vmin is None: vmin = float(np.nanmin(cvals))
    if vmax is None: vmax = float(np.nanmax(cvals))

    fig, ax = plt.subplots(figsize=figsize)
    sc = ax.scatter(
        d[x].to_numpy(dtype=float),
        d[y].to_numpy(dtype=float),
        c=cvals,
        s=s, alpha=alpha,
        cmap=cmap,
        vmin=vmin, vmax=vmax,
        edgecolors=(0,0,0,edge_alpha),
        linewidths=edge_lw,
        rasterized=True,
    )

    ax.set_xlabel(x)
    ax.set_ylabel(y)
    ax.set_title(title if title is not None else f"{y} vs {x} (colored by {color_by})", pad=10)
    _tight_limits(ax, d[x].to_numpy(), d[y].to_numpy())

    cbar = fig.colorbar(sc, ax=ax, pad=0.02)
    cbar.set_label(color_by)

    fig.tight_layout()
    if savepath:
        _ensure_dir(savepath)
        plt.savefig(savepath, dpi=300, bbox_inches="tight")
        print("[wrote]", savepath)
    plt.show()

# ---------- one-call suite ----------
def tradeoff_viz_suite(
    df, x, y, *,
    outdir,
    stem,
    figsize=(6.8, 5.0),
    do_density=True,
    do_pareto=True,
    do_beta=True,
    do_gamma=True,
):
    os.makedirs(outdir, exist_ok=True)

    if do_density:
        plot_tradeoff_density(
            df, x, y,
            figsize=figsize,
            savepath=os.path.join(outdir, f"{stem}__density.png"),
            title=f"{y} vs {x} (density)",
        )

    if do_pareto:
        plot_tradeoff_pareto(
            df, x, y,
            figsize=figsize,
            savepath=os.path.join(outdir, f"{stem}__pareto.png"),
            title=f"{y} vs {x} (Pareto front)",
            label_k=10,
            label_candidates=60,
            label_min_dist_px=32,
        )

    # Fixed scales for comparability across multiple plots with same df
    if do_beta and ("beta" in df.columns):
        bmin, bmax = float(pd.to_numeric(df["beta"], errors="coerce").min()), float(pd.to_numeric(df["beta"], errors="coerce").max())
        plot_tradeoff_colored(
            df, x, y, color_by="beta",
            figsize=figsize,
            vmin=bmin, vmax=bmax,
            savepath=os.path.join(outdir, f"{stem}__beta.png"),
            title=f"{y} vs {x} (colored by beta)",
        )

    if do_gamma and ("gamma" in df.columns):
        gmin, gmax = float(pd.to_numeric(df["gamma"], errors="coerce").min()), float(pd.to_numeric(df["gamma"], errors="coerce").max())
        plot_tradeoff_colored(
            df, x, y, color_by="gamma",
            figsize=figsize,
            vmin=gmin, vmax=gmax,
            savepath=os.path.join(outdir, f"{stem}__gamma.png"),
            title=f"{y} vs {x} (colored by gamma)",
        )

# =============================================================================
# ---- Example usage (your main tradeoff) ----
# =============================================================================
tradeoff_viz_suite(
    df_grid,
    x="FOSCTTM",
    y="LT_RNA2ATAC_macroF1",
    outdir=os.path.join(OUTDIR, "figures"),
    stem="tradeoff_LTmacroF1_vs_FOSCTTM",
    figsize=(14, 12),
    do_density=True,
    do_pareto=True,
    do_beta=True,
    do_gamma=True,
)


# %% [code]
# =============================================================================
# ---- 15) Plotting (paired-only) ----
# =============================================================================
import numpy as np
import pandas as pd
import matplotlib.pyplot as plt


# ---- metric metadata ----
# Higher-is-better by default unless listed.
LOWER_BETTER_SUBSTR = ["loss", "error", "dist", "foscttm", "rmse", "mae", "db_"]
DEFAULT_KS = (1, 10, 25, 50, 100)

def _is_lower_better(metric: str) -> bool:
    m = metric.lower()
    return any(s in m for s in LOWER_BETTER_SUBSTR)

def _coerce_numeric(df, cols):
    d = df.copy()
    for c in cols:
        if c in d.columns:
            d[c] = pd.to_numeric(d[c], errors="coerce")
    return d

def _savefig(savepath):
    if savepath:
        plt.savefig(savepath, dpi=300)
        print("[wrote]", savepath)


# -----------------------------------------------------------------------------
# 1) Leaderboard barh
# -----------------------------------------------------------------------------
def barplot_best(df, metric, *, top_n=30, ascending=None, savepath=None, title=None):
    if metric not in df.columns:
        print(f"[skip] missing metric: {metric}")
        return
    d = _coerce_numeric(df, [metric]).dropna(subset=[metric]).copy()

    if ascending is None:
        ascending = _is_lower_better(metric)

    d = d.sort_values(metric, ascending=bool(ascending)).head(int(top_n))

    plt.figure(figsize=(10, 0.32 * len(d) + 2))
    plt.barh(d["sweep"].astype(str), d[metric].astype(float))
    plt.xlabel(metric)
    plt.ylabel("sweep")
    plt.title(title if title is not None else f"Top {top_n}: {metric} ({'lower' if ascending else 'higher'} is better)")
    plt.tight_layout()
    _savefig(savepath)
    plt.show()


# -----------------------------------------------------------------------------
# 2) Tradeoff scatter + optional Pareto highlight
# -----------------------------------------------------------------------------
def pareto_mask(x, y, *, x_lower_better=False, y_lower_better=False):
    """
    Returns boolean mask of Pareto-optimal points for 2D.
    """
    x = np.asarray(x, float)
    y = np.asarray(y, float)

    # Convert to "higher is better" by flipping if needed
    xx = -x if x_lower_better else x
    yy = -y if y_lower_better else y

    n = len(xx)
    keep = np.ones(n, dtype=bool)
    for i in range(n):
        if not keep[i]:
            continue
        dominated = (xx >= xx[i]) & (yy >= yy[i]) & ((xx > xx[i]) | (yy > yy[i]))
        dominated[i] = False
        keep[dominated] = False
    return keep

def scatter_tradeoff(df, x, y, *, annotate_top=0, savepath=None, title=None,
                     max_labels=12, label_fontsize=8):
    if x not in df.columns or y not in df.columns:
        print(f"[skip] missing: {x} or {y}")
        return

    d = df[["sweep", x, y]].copy()
    d[x] = pd.to_numeric(d[x], errors="coerce")
    d[y] = pd.to_numeric(d[y], errors="coerce")
    d = d.dropna()

    fig, ax = plt.subplots(figsize=(6.5, 5))

    # points: smaller + slightly transparent helps a lot
    ax.scatter(d[x].astype(float), d[y].astype(float), s=30, alpha=0.85)

    ax.set_xlabel(x)
    ax.set_ylabel(y)
    ax.set_title(title if title is not None else f"{y} vs {x}")

    # ---- tighten margins (like the barplots) ----
    ax.margins(x=0.02, y=0.02)  # small breathing room but not the big default padding

    # If you want hard clamps based on data (optional but nice):
    xvals = d[x].astype(float).to_numpy()
    yvals = d[y].astype(float).to_numpy()
    xr = float(xvals.max() - xvals.min()) if len(xvals) else 1.0
    yr = float(yvals.max() - yvals.min()) if len(yvals) else 1.0
    ax.set_xlim(float(xvals.min() - 0.03 * xr), float(xvals.max() + 0.03 * xr))
    ax.set_ylim(float(yvals.min() - 0.03 * yr), float(yvals.max() + 0.03 * yr))

    # ---- annotations: pick points, then repel text a bit ----
    if int(annotate_top) > 0:
        # if "lower is better" for y, sort ascending; else descending
        y_lower_better = any(s in y.lower() for s in ["loss", "error", "dist", "foscttm", "rmse", "mae", "rankfrac"])
        dd = d.sort_values(y, ascending=bool(y_lower_better)).head(int(annotate_top))
        if len(dd) > int(max_labels):
            dd = dd.head(int(max_labels))

        # initial annotation placement
        texts = []
        for _, r in dd.iterrows():
            txt = ax.text(float(r[x]), float(r[y]), str(r["sweep"]),
                          fontsize=label_fontsize, ha="left", va="bottom")
            texts.append(txt)

        # simple label repulsion: iteratively nudge labels apart in display coords
        fig.canvas.draw()  # need renderer for accurate text extents
        renderer = fig.canvas.get_renderer()

        def _overlap(b1, b2, pad=2.0):
            return not (b1.x1 + pad < b2.x0 or b1.x0 - pad > b2.x1 or
                        b1.y1 + pad < b2.y0 or b1.y0 - pad > b2.y1)

        # nudge loop
        for _ in range(80):
            moved = False
            bbs = [t.get_window_extent(renderer=renderer) for t in texts]
            for i in range(len(texts)):
                for j in range(i + 1, len(texts)):
                    if _overlap(bbs[i], bbs[j], pad=2.0):
                        # move j a bit up/right in data units proportional to axis ranges
                        xi, yi = texts[j].get_position()
                        texts[j].set_position((xi + 0.008 * xr, yi + 0.012 * yr))
                        moved = True
            if not moved:
                break
            fig.canvas.draw()
            renderer = fig.canvas.get_renderer()

    fig.tight_layout()
    _savefig(savepath)
    plt.show()


# -----------------------------------------------------------------------------
# 3) Heatmap of hp grid vs metric
# -----------------------------------------------------------------------------
def heatmap_hparams_vs_metric(df, metric, hp_x="beta", hp_y="gamma", *, agg="median", savepath=None, title=None):
    needed = [metric, hp_x, hp_y]
    for c in needed:
        if c not in df.columns:
            print(f"[skip] missing col: {c}")
            return

    d = _coerce_numeric(df[[hp_x, hp_y, metric]], [hp_x, hp_y, metric]).dropna().copy()

    if agg == "mean":
        pv = d.groupby([hp_y, hp_x])[metric].mean().unstack(hp_x)
    elif agg == "min":
        pv = d.groupby([hp_y, hp_x])[metric].min().unstack(hp_x)
    elif agg == "max":
        pv = d.groupby([hp_y, hp_x])[metric].max().unstack(hp_x)
    else:
        pv = d.groupby([hp_y, hp_x])[metric].median().unstack(hp_x)

    # sort axes numerically when possible
    try:
        pv = pv.sort_index(axis=0).sort_index(axis=1)
    except Exception:
        pass

    plt.figure(figsize=(1.1 * max(6, pv.shape[1]), 1.0 * max(4, pv.shape[0])))
    plt.imshow(pv.values, aspect="auto")
    plt.xticks(range(pv.shape[1]), [str(x) for x in pv.columns], rotation=45, ha="right")
    plt.yticks(range(pv.shape[0]), [str(y) for y in pv.index])
    plt.xlabel(hp_x)
    plt.ylabel(hp_y)
    plt.title(title if title is not None else f"{metric} heatmap ({agg})")
    plt.colorbar(label=metric)
    plt.tight_layout()
    _savefig(savepath)
    plt.show()


# -----------------------------------------------------------------------------
# 4) Retrieval curve: recall_sym@k (or FOSCTTM@k) across k
# -----------------------------------------------------------------------------
def plot_retrieval_curve(df, *, which="recall_sym", ks=DEFAULT_KS, top_n=10, by="best", savepath=None, title=None):
    """
    Plots recall_sym@k (higher better) OR FOSCTTM@k (lower better) across k.
    Picks top_n runs by recall_sym@1 or by FOSCTTM_rankfrac depending on 'which'.
    """
    ks = [int(k) for k in ks]
    cols = [f"{which}@{k}" for k in ks]
    for c in cols:
        if c not in df.columns:
            print(f"[skip] missing col: {c}")
            return

    d = _coerce_numeric(df[["sweep"] + cols + (["FOSCTTM_rankfrac"] if "FOSCTTM_rankfrac" in df.columns else [])], cols).dropna().copy()

    # choose ranking metric
    if which.lower().startswith("recall"):
        rank_metric = f"{which}@{ks[0]}"
        ascending = False
    else:
        # FOSCTTM@k lower better
        rank_metric = f"{which}@{ks[0]}"
        ascending = True

    d = d.sort_values(rank_metric, ascending=bool(ascending)).head(int(top_n))

    plt.figure(figsize=(7.2, 4.8))
    for _, r in d.iterrows():
        y = [float(r[f"{which}@{k}"]) for k in ks]
        plt.plot(ks, y, marker="o", alpha=0.85, label=str(r["sweep"]))

    plt.xlabel("k")
    plt.ylabel(which)
    plt.title(title if title is not None else f"Retrieval curve: {which}@k (top {top_n} by {rank_metric})")
    plt.xticks(ks)
    plt.tight_layout()

    # Legend can get huge; only show if small
    if len(d) <= 12:
        plt.legend(fontsize=8, loc="best")

    _savefig(savepath)
    plt.show()


# -----------------------------------------------------------------------------
# 5) Correlation heatmap (redundancy check)
# -----------------------------------------------------------------------------
def corr_heatmap(df, metrics, *, savepath=None, title="Metric correlation (Spearman)"):
    keep = [m for m in metrics if m in df.columns]
    if len(keep) < 2:
        print("[skip] need >=2 metrics present")
        return

    d = _coerce_numeric(df[keep], keep).dropna()
    if len(d) < 3:
        print("[skip] too few rows after dropna")
        return

    C = d.corr(method="spearman").values

    plt.figure(figsize=(0.55 * len(keep) + 3, 0.55 * len(keep) + 3))
    plt.imshow(C, aspect="auto", vmin=-1, vmax=1)
    plt.xticks(range(len(keep)), keep, rotation=45, ha="right")
    plt.yticks(range(len(keep)), keep)
    plt.title(title)
    plt.colorbar(label="Spearman rho")
    plt.tight_layout()
    _savefig(savepath)
    plt.show()


# -----------------------------------------------------------------------------
# 6) Scaling plots (kept, minor robustness)
# -----------------------------------------------------------------------------
def plot_scaling_multi(df, *, x="n_train_cells", saveprefix=None):
    if df is None or len(df) == 0:
        print("[skip] empty df_scale")
        return

    if x not in df.columns:
        print(f"[skip] missing x: {x}")
        return

    d = df.copy()
    d = _coerce_numeric(d, [x, "wall_seconds", "gpu_peak_mem_mb", "cpu_rss_mb_delta"])
    d = d.sort_values(x)

    if "wall_seconds" in d.columns:
        plt.figure(figsize=(6,4))
        plt.plot(d[x], d["wall_seconds"], marker="o")
        plt.xlabel(x)
        plt.ylabel("wall seconds (train)")
        plt.title("Scaling: runtime vs size")
        plt.tight_layout()
        _savefig(None if saveprefix is None else f"{saveprefix}_runtime.png")
        plt.show()

        plt.figure(figsize=(6,4))
        thr = d[x].astype(float) / np.maximum(d["wall_seconds"].astype(float), 1e-9)
        plt.plot(d[x], thr, marker="o")
        plt.xlabel(x)
        plt.ylabel("train throughput (items/sec)")
        plt.title("Scaling: throughput vs size")
        plt.tight_layout()
        _savefig(None if saveprefix is None else f"{saveprefix}_throughput.png")
        plt.show()

    if "gpu_peak_mem_mb" in d.columns:
        plt.figure(figsize=(6,4))
        plt.plot(d[x], d["gpu_peak_mem_mb"], marker="o")
        plt.xlabel(x)
        plt.ylabel("GPU peak mem (MB)")
        plt.title("Scaling: GPU peak memory vs size")
        plt.tight_layout()
        _savefig(None if saveprefix is None else f"{saveprefix}_gpu_mem.png")
        plt.show()

    if "cpu_rss_mb_delta" in d.columns:
        plt.figure(figsize=(6,4))
        plt.plot(d[x], d["cpu_rss_mb_delta"], marker="o")
        plt.xlabel(x)
        plt.ylabel("CPU RSS delta (MB)")
        plt.title("Scaling: CPU memory delta vs size")
        plt.tight_layout()
        _savefig(None if saveprefix is None else f"{saveprefix}_cpu_mem_delta.png")
        plt.show()


# %% [code]
barplot_best(df_grid, "recall_sym@1", top_n=50, ascending=False,
            savepath=os.path.join(OUTDIR, "figures", "grid_top_recall_sym_at1.png"),
            title="Grid leaderboard: recall_sym@1 (higher is better)")

barplot_best(df_grid, "FOSCTTM_rankfrac", top_n=50,
            savepath=os.path.join(OUTDIR, "figures", "grid_top_FOSCTTM_rankfrac.png"),
            title="Grid leaderboard: FOSCTTM_rankfrac (lower is better)")

for m in ["ARI", "NMI", "kNN_LOO_acc_fused", "kNN_LOO_macroF1_fused", "mixing_1minus_samefrac_k15"]:
    if m in df_grid.columns:
        barplot_best(df_grid, m, top_n=50, ascending=False,
                    savepath=os.path.join(OUTDIR, "figures", f"grid_top_{m}.png"))


# %% [code]
scatter_tradeoff(df_grid, "recall_sym@1", "ARI", annotate_top=25,
                 savepath=os.path.join(OUTDIR, "figures", "tradeoff_ARI_vs_recall1.png"),
                 title="Tradeoff: fused biology vs pairing quality")

scatter_tradeoff(df_grid, "recall_sym@1", "mixing_1minus_samefrac_k15", annotate_top=25,
                 savepath=os.path.join(OUTDIR, "figures", "tradeoff_mixing_vs_recall1.png"),
                 title="Tradeoff: mixing vs pairing quality")


# %% [code]
plot_retrieval_curve(df_grid, which="recall_sym", ks=(1,10,25,50,100), top_n=25,
                     savepath=os.path.join(OUTDIR, "figures", "retrieval_curve_recall_sym.png"),
                     title="Paired retrieval: recall_sym@k")


# %% [code]
plot_retrieval_curve(df_grid, which="FOSCTTM", ks=(1,10,25,50,100), top_n=25,
                     savepath=os.path.join(OUTDIR, "figures", "retrieval_curve_FOSCTTM_at_k.png"),
                     title="Paired retrieval: FOSCTTM@k (lower is better)")


# %% [code]
heatmap_hparams_vs_metric(df_grid, "recall_sym@1", hp_x="beta", hp_y="gamma", agg="median",
                          savepath=os.path.join(OUTDIR, "figures", "heatmap_recall1_beta_gamma.png"),
                          title="recall_sym@1 vs (beta, gamma)")

heatmap_hparams_vs_metric(df_grid, "ARI", hp_x="beta", hp_y="gamma", agg="median",
                          savepath=os.path.join(OUTDIR, "figures", "heatmap_ARI_beta_gamma.png"),
                          title="ARI vs (beta, gamma)")


# %% [code]
corr_heatmap(
    df_grid,
    metrics=[
        "recall_sym@1", "recall_sym@10", "FOSCTTM_rankfrac",
        "ARI", "NMI", "SIL_true",
        "kNN_LOO_acc_fused", "kNN_LOO_macroF1_fused",
        "mixing_1minus_samefrac_k15"
    ],
    savepath=os.path.join(OUTDIR, "figures", "corr_metrics.png")
)


# %% [code]


# %% [code]


# %% [code]


# %% [code]


# %% [code]


# %% [code]


# %% [code]


# %% [code]


# %% [code]


# %% [code]


# %% [code]


# %% [code]


# %% [code]


# %% [code]


# %% [code]


# %% [code]


# %% [code]


# %% [code]


# %% [code]


# %% [code]


# %% [code]


# %% [code]


# %% [code]


# %% [code]


# %% [code]


# %% [code]


# %% [code]


# %% [code]


# %% [code]


# %% [code]


# %% [code]


# %% [code]


# %% [code]


# %% [code]


# %% [code]


# %% [code]


# %% [code]


# %% [code]


# %% [code]


# %% [code]


# %% [code]


# %% [code]


# %% [code]


# %% [code]


# %% [code]


# %% [code]


# %% [code]


# %% [code]


# %% [code]


# %% [code]


# %% [code]


# %% [code]


# %% [code]


# %% [code]


# %% [code]


# %% [code]


# %% [code]


# %% [code]


# %% [code]


# %% [code]


# %% [code]


# %% [code]


# %% [code]


# %% [code]


# %% [code]


# %% [code]


# %% [code]


# %% [code]


# %% [code]


# %% [code]


# %% [code]


# %% [code]


# %% [code]


# %% [code]
```
