## Supplemental Code for "Unifying multimodal single-cell data with a mixture-of-experts *β*-variational autoencoder framework": Supp_grid_sweep_compile.ipynb

```
# %% [code]
# --- Read TSV + make 4-panel hyperparam heatmaps (beta x gamma) ---
# Panels:
# (a) FOSCTTM (lower better)
# (b) Recall@1 (higher better)
# (c) Label transfer macro-F1 (worse direction if both available; higher better)
# (d) ARI (higher better)
#
# Output: shows in-notebook at 8x6 inches, and saves a 300 dpi PNG.

import numpy as np
import pandas as pd
import matplotlib.pyplot as plt

# -----------------------------
# 1) Load
# -----------------------------
RESULTS_DIR = "./results/fig10_ablation_scaling_all_combos/"
TSV_PATH = RESULTS_DIR + "param_grid_paired_only.tsv"
df = pd.read_csv(TSV_PATH, sep="\t")
print(df.shape)
display(df.head())


# %% [code]
# -----------------------------
# 2) Pick / derive the requested metrics robustly
# -----------------------------
def pick_first_existing(cols):
    for c in cols:
        if c in df.columns:
            return c
    return None

fos_col = pick_first_existing(["FOSCTTM", "FOSCTTM_rankfrac"])
recall_col = pick_first_existing(["recall_sym@10", "recall_A2B@1", "recall_B2A@1"])

# Label transfer macro-F1: use worse direction if both exist (or just the one you have)
lt_r2a = "LT_RNA2ATAC_macroF1" if "LT_RNA2ATAC_macroF1" in df.columns else None
lt_a2r = "LT_ATAC2RNA_macroF1" if "LT_ATAC2RNA_macroF1" in df.columns else None

if lt_r2a and lt_a2r:
    df["LT_macroF1_worse_dir"] = np.minimum(
        pd.to_numeric(df[lt_r2a], errors="coerce"),
        pd.to_numeric(df[lt_a2r], errors="coerce"),
    )
    lt_col = "LT_macroF1_worse_dir"
else:
    lt_col = pick_first_existing([lt_r2a, lt_a2r, "LT_RNA2ATAC_acc", "LT_ATAC2RNA_acc"])

ari_col = pick_first_existing(["NMI", "ARI"])  # prefer NMI; fallback ARI

needed = {"beta", "gamma", fos_col, recall_col, lt_col, ari_col}
missing = [c for c in needed if c is None or c not in df.columns]
if missing:
    raise ValueError(f"Missing expected columns: {missing}")

# Ensure numeric
for c in [fos_col, recall_col, lt_col, ari_col, "beta", "gamma"]:
    df[c] = pd.to_numeric(df[c], errors="coerce")


# %% [code]
# -----------------------------
# 3) Aggregate across seeds (median), then take "best" across other hyperparams per (beta, gamma)
# -----------------------------
# Group over "config columns" excluding seed-like / run-logging columns.
# This gives you a robust per-config score across random seeds,
# then collapses to a beta/gamma summary.
exclude_cols = {
    "seed", "wall_seconds", "gpu_peak_mem_mb", "best_epoch", "best_val", "sweep",
    "n_test",
}
# keep only columns that exist
exclude_cols = {c for c in exclude_cols if c in df.columns}

# Choose grouping columns as: all non-metric, non-excluded columns except seed
metric_cols = {fos_col, recall_col, lt_col, ari_col}
group_cols = [c for c in df.columns if c not in metric_cols and c not in exclude_cols]

# Make sure beta/gamma are included
for c in ["beta", "gamma"]:
    if c not in group_cols:
        group_cols.append(c)

df_cfg = (
    df.dropna(subset=["beta", "gamma", fos_col, recall_col, lt_col, ari_col])
      .groupby(group_cols, dropna=False, as_index=False)[[fos_col, recall_col, lt_col, ari_col]]
      .median(numeric_only=True)
)

# Collapse to beta/gamma by selecting best across configs
# (min for FOSCTTM; max for the others)
def collapse_best(df_cfg, value_col, higher_is_better: bool):
    agg = df_cfg.groupby(["beta", "gamma"], as_index=False)[value_col]
    out = agg.max() if higher_is_better else agg.min()
    return out

fos_bg = collapse_best(df_cfg, fos_col, higher_is_better=False)
rec_bg = collapse_best(df_cfg, recall_col, higher_is_better=True)
lt_bg  = collapse_best(df_cfg, lt_col,  higher_is_better=True)
ari_bg = collapse_best(df_cfg, ari_col, higher_is_better=True)


# %% [code]
# -----------------------------
# 4) Helper to pivot into a matrix and plot heatmap
# -----------------------------
def to_matrix(bg_df, value_col):
    betas = np.array(sorted(bg_df["beta"].unique()))
    gammas = np.array(sorted(bg_df["gamma"].unique()))
    mat = np.full((len(gammas), len(betas)), np.nan, dtype=float)
    b_index = {b:i for i,b in enumerate(betas)}
    g_index = {g:i for i,g in enumerate(gammas)}

    for _, r in bg_df.iterrows():
        i = g_index[r["gamma"]]
        j = b_index[r["beta"]]
        mat[i, j] = r[value_col]
    return betas, gammas, mat

def plot_heat(ax, betas, gammas, mat, title, lower_is_better=False):
    im = ax.imshow(mat, origin="lower", aspect="auto", interpolation="nearest")
    ax.set_title(title)
    ax.set_xlabel("beta")
    ax.set_ylabel("gamma")

    ax.set_xticks(np.arange(len(betas)))
    ax.set_xticklabels([str(b) for b in betas], rotation=45, ha="right")
    ax.set_yticks(np.arange(len(gammas)))
    ax.set_yticklabels([str(g) for g in gammas])

    cbar = plt.colorbar(im, ax=ax, fraction=0.046, pad=0.04)
    cbar.set_label("value")

    # Optional: mark the best cell
    if np.isfinite(mat).any():
        if lower_is_better:
            best = np.nanargmin(mat)
        else:
            best = np.nanargmax(mat)
        iy, ix = np.unravel_index(best, mat.shape)
        ax.scatter([ix], [iy], marker="o", s=60, facecolors="none", edgecolors="black", linewidths=2)


def plot_heat(ax, betas, gammas, mat, title, lower_is_better=False, forbid_beta_zero=True):
    im = ax.imshow(mat, origin="lower", aspect="auto", interpolation="nearest")
    ax.set_title(title)
    ax.set_xlabel("beta")
    ax.set_ylabel("gamma")

    ax.set_xticks(np.arange(len(betas)))
    ax.set_xticklabels([str(b) for b in betas], rotation=45, ha="right")
    ax.set_yticks(np.arange(len(gammas)))
    ax.set_yticklabels([str(g) for g in gammas])

    cbar = plt.colorbar(im, ax=ax, fraction=0.046, pad=0.04)
    cbar.set_label("value")

    # ---- mark best cell, optionally excluding beta==0 ----
    if np.isfinite(mat).any():
        mat_search = mat.copy()

        if forbid_beta_zero:
            betas_f = np.asarray(betas, dtype=float)
            zero_cols = np.where(np.isclose(betas_f, 0.0, atol=1e-12, rtol=0.0))[0]
            if zero_cols.size > 0:
                mat_search[:, zero_cols] = np.nan  # exclude beta=0 from "best" search

        # If everything is NaN after masking, just skip marking best
        if np.isfinite(mat_search).any():
            best = np.nanargmin(mat_search) if lower_is_better else np.nanargmax(mat_search)
            iy, ix = np.unravel_index(best, mat_search.shape)
            ax.scatter([ix], [iy], marker="o", s=60,
                       facecolors="none", edgecolors="black", linewidths=2, zorder=5)
        else:
            ax.text(0.01, 0.01, "best excluded (β=0 only)",
                    transform=ax.transAxes, fontsize=8, va="bottom", ha="left")


# %% [code]
# -----------------------------
# 5) Build the 4-panel figure (8x6 inches) + save at 300 dpi
# -----------------------------
fig, axes = plt.subplots(2, 2, figsize=(18, 16), constrained_layout=True)

b, g, m = to_matrix(fos_bg, fos_col)
plot_heat(axes[0, 0], b, g, m, f"(a) {fos_col} (lower is better)", lower_is_better=True)

b, g, m = to_matrix(rec_bg, recall_col)
plot_heat(axes[0, 1], b, g, m, f"(b) {recall_col} (higher is better)", lower_is_better=False)

b, g, m = to_matrix(lt_bg, lt_col)
plot_heat(axes[1, 0], b, g, m, f"(c) {lt_col} (higher is better)", lower_is_better=False)

b, g, m = to_matrix(ari_bg, ari_col)
plot_heat(axes[1, 1], b, g, m, f"(d) {ari_col} (higher is better)", lower_is_better=False)

out_png = "paired_only__beta_gamma__4panel.png"
fig.savefig(out_png, dpi=400, bbox_inches="tight")
plt.show()

print(f"Saved: {out_png} (400 dpi)")


# %% [code]
def plot_heat(
    ax, betas, gammas, mat, title,
    lower_is_better=False,
    forbid_beta_zero=True,
    baseline_beta=1.25,
    baseline_gamma=4.35,
    baseline_marker="*",
):
    im = ax.imshow(mat, origin="lower", aspect="auto", interpolation="nearest")
    ax.set_title(title)
    ax.set_xlabel("beta")
    ax.set_ylabel("gamma")

    ax.set_xticks(np.arange(len(betas)))
    ax.set_xticklabels([str(b) for b in betas], rotation=45, ha="right")
    ax.set_yticks(np.arange(len(gammas)))
    ax.set_yticklabels([str(g) for g in gammas])

    # IMPORTANT: attach colorbar to the figure that owns this Axes
    fig = ax.figure
    cbar = fig.colorbar(im, ax=ax, fraction=0.046, pad=0.04)
    cbar.set_label("value")

    # ---- mark best cell, optionally excluding beta==0 ----
    if np.isfinite(mat).any():
        mat_search = mat.copy()

        if forbid_beta_zero:
            betas_f = np.asarray(betas, dtype=float)
            zero_cols = np.where(np.isclose(betas_f, 0.0, atol=1e-12, rtol=0.0))[0]
            if zero_cols.size > 0:
                mat_search[:, zero_cols] = np.nan

        if np.isfinite(mat_search).any():
            best = np.nanargmin(mat_search) if lower_is_better else np.nanargmax(mat_search)
            iy, ix = np.unravel_index(best, mat_search.shape)
            ax.scatter(
                [ix], [iy],
                marker="o", s=200,
                facecolors="none", edgecolors="black",
                linewidths=2, zorder=5
            )
        else:
            ax.text(
                0.01, 0.01, "best excluded (β=0 only)",
                transform=ax.transAxes, fontsize=8,
                va="bottom", ha="left"
            )

    # ---- star the baseline (beta=1.25, gamma=4.35) if present ----
    if baseline_beta is not None and baseline_gamma is not None:
        betas_f = np.asarray(betas, dtype=float)
        gammas_f = np.asarray(gammas, dtype=float)

        bx = np.where(np.isclose(betas_f, float(baseline_beta), atol=1e-12, rtol=0.0))[0]
        gy = np.where(np.isclose(gammas_f, float(baseline_gamma), atol=1e-12, rtol=0.0))[0]

        if bx.size > 0 and gy.size > 0:
            ix = int(bx[0])
            iy = int(gy[0])
            ax.scatter(
                [ix], [iy],
                marker=baseline_marker, s=150,
                color="black", linewidths=1.0,
                zorder=6
            )
        else:
            ax.text(
                0.01, 0.97, f"baseline (β={baseline_beta}, γ={baseline_gamma}) not in grid",
                transform=ax.transAxes, fontsize=7,
                va="top", ha="left"
            )


# %% [code]
plt.rcParams["figure.dpi"] = 500


# %% [code]
plt.close("all")

fig, axes = plt.subplots(2, 2, figsize=(18, 16), constrained_layout=True)

b, g, m = to_matrix(fos_bg, fos_col)
plot_heat(axes[0, 0], b, g, m, f"(a) {fos_col} (lower is better)",
          lower_is_better=True, baseline_beta=1.25, baseline_gamma=4.35)

b, g, m = to_matrix(rec_bg, recall_col)
plot_heat(axes[0, 1], b, g, m, f"(b) {recall_col} (higher is better)",
          lower_is_better=False, baseline_beta=1.25, baseline_gamma=4.35)

b, g, m = to_matrix(lt_bg, lt_col)
plot_heat(axes[1, 0], b, g, m, f"(c) {lt_col} (higher is better)",
          lower_is_better=False, baseline_beta=1.25, baseline_gamma=4.35)

b, g, m = to_matrix(ari_bg, ari_col)
plot_heat(axes[1, 1], b, g, m, f"(d) {ari_col} (higher is better)",
          lower_is_better=False, baseline_beta=1.25, baseline_gamma=4.35)


# %% [code]
# =============================================================================
# Supplemental plots (USES YOUR REAL COLUMN NAMES)
#   1) latent dim sweep (from param_grid_paired_only.tsv)
#   2) dropout sweeps (encoder/decoder) parsed from sweep labels like drop_enc_0.1
#   3) scaling curve (from scaling_curve.tsv)
#
# FIXES:
#   - Panel (b): CPU forced blue (C0), GPU/MPS forced orange (C1)
#   - Panel (b): legend placed INSIDE panel (b) (no constrained_layout blow-up)
#   - Panel titles get a little padding to avoid collisions
# =============================================================================

import re
import numpy as np
import pandas as pd
import matplotlib.pyplot as plt

# -----------------------------
# Paths
# -----------------------------
SWEEP_TSV   = RESULTS_DIR + "param_grid_paired_only.tsv"
SCALING_TSV = RESULTS_DIR + "scaling_curve.tsv"

# -----------------------------
# Settings
# -----------------------------
BASELINE_BETA  = 1.25
BASELINE_GAMMA = 4.35

FOS_COL    = "FOSCTTM_rankfrac"   # lower is better
RECALL_COL = "recall_sym@10"      # higher is better
#ARI_COL    = "ARI"                # higher is better
ARI_COL    = "NMI"                # higher is better

DEFAULT_LATENT    = 30
DEFAULT_DROP_ENC  = 0.10
DEFAULT_DROP_DEC  = 0.05
DEFAULT_DROP_TIED = None  # unused in this 2-row dropout plot

SAVE_DPI = 500

# -----------------------------
# Helpers
# -----------------------------
def to_num(df, cols):
    for c in cols:
        if c in df.columns:
            df[c] = pd.to_numeric(df[c], errors="coerce")
    return df

def filter_baseline_bg(df):
    out = df.copy()
    if "beta" in out.columns:
        out = out[np.isclose(out["beta"].astype(float), float(BASELINE_BETA), atol=1e-12, rtol=0.0)]
    if "gamma" in out.columns:
        out = out[np.isclose(out["gamma"].astype(float), float(BASELINE_GAMMA), atol=1e-12, rtol=0.0)]
    return out

def add_lt_worse_dir(df):
    r2a = "LT_RNA2ATAC_macroF1"
    a2r = "LT_ATAC2RNA_macroF1"
    if (r2a in df.columns) and (a2r in df.columns):
        df = to_num(df, [r2a, a2r])
        df["LT_macroF1_worse_dir"] = np.minimum(df[r2a].to_numpy(), df[a2r].to_numpy())
        return df, "LT_macroF1_worse_dir"
    raise ValueError("Expected LT_RNA2ATAC_macroF1 and LT_ATAC2RNA_macroF1 columns.")

def plot_curve(ax, x, y, title, xlab, ylab, star_x=None, star_y=None):
    ax.plot(x, y, marker="o", linewidth=1.6, markersize=4)
    ax.set_title(title)
    ax.set_xlabel(xlab)
    ax.set_ylabel(ylab)
    ax.grid(True, linewidth=0.3, alpha=0.5)
    if (star_x is not None) and (star_y is not None) and np.isfinite(star_y):
        ax.scatter([star_x], [star_y], marker="*", s=160, edgecolors="black", linewidths=1.0, zorder=5)

def star_from_exact(x, y, x0):
    if x0 is None:
        return None, None
    x = np.asarray(x, dtype=float)
    y = np.asarray(y, dtype=float)
    m = np.isclose(x, float(x0), atol=1e-12, rtol=0.0)
    if m.any():
        return float(x0), float(y[m][0])
    return None, None

def agg_1d(df, xcol, ycols, agg="median"):
    d = df.dropna(subset=[xcol]).copy()
    for c in ycols:
        d = d.dropna(subset=[c])
    g = d.groupby(xcol, as_index=False)[ycols]
    out = g.median(numeric_only=True) if agg == "median" else g.mean(numeric_only=True)
    return out.sort_values(xcol)

# --- parse dropout from sweep labels ---
_drop_re = re.compile(r"^drop_(encdec|enc|dec)_(\d*\.?\d+)$")
def with_parsed_dropout(df):
    kinds, ps = [], []
    for s in df["sweep"].astype(str).tolist():
        m = _drop_re.match(s)
        if m:
            kinds.append(m.group(1))
            ps.append(float(m.group(2)))
        else:
            kinds.append(np.nan)
            ps.append(np.nan)
    out = df.copy()
    out["drop_kind"] = kinds
    out["drop_p"] = ps
    return out

# =============================================================================
# Load sweep TSV
# =============================================================================
df = pd.read_csv(SWEEP_TSV, sep="\t")
print("Sweep TSV:", df.shape)
display(df.head(3))

df = to_num(df, ["latent_dim", "beta", "gamma", "seed", FOS_COL, RECALL_COL, ARI_COL, "NMI"])
df, LT_COL = add_lt_worse_dir(df)

for c in ["sweep", "latent_dim", FOS_COL, RECALL_COL, LT_COL, ARI_COL]:
    if c not in df.columns:
        raise ValueError(f"Missing required column: {c}")

# =============================================================================
# 1) Latent dimension sweep (2x2)
# =============================================================================
d_lat = df[df["sweep"].astype(str).str.contains("latent", case=False, regex=False)].copy()
if d_lat.empty:
    d_lat = df[df["latent_dim"].notna()].copy()

d_lat = filter_baseline_bg(d_lat)
lat = agg_1d(d_lat, "latent_dim", [FOS_COL, RECALL_COL, LT_COL, ARI_COL], agg="median")
x = lat["latent_dim"].astype(float).to_numpy()

#fig, axes = plt.subplots(2, 2, figsize=(8, 6), constrained_layout=True)
fig, axes = plt.subplots(2, 2, figsize=(14, 12), constrained_layout=True)
axes = axes.ravel()

y = lat[FOS_COL].astype(float).to_numpy()
sx, sy = star_from_exact(x, y, DEFAULT_LATENT)
plot_curve(axes[0], x, y, "(a) FOSCTTM_rankfrac (↓)", "latent_dim", FOS_COL, sx, sy)

y = lat[RECALL_COL].astype(float).to_numpy()
sx, sy = star_from_exact(x, y, DEFAULT_LATENT)
plot_curve(axes[1], x, y, f"(b) {RECALL_COL} (↑)", "latent_dim", RECALL_COL, sx, sy)

y = lat[LT_COL].astype(float).to_numpy()
sx, sy = star_from_exact(x, y, DEFAULT_LATENT)
plot_curve(axes[2], x, y, "(c) LT macroF1 (worse-dir) (↑)", "latent_dim", LT_COL, sx, sy)

y = lat[ARI_COL].astype(float).to_numpy()
sx, sy = star_from_exact(x, y, DEFAULT_LATENT)
plot_curve(axes[3], x, y, "(d) NMI (↑)", "latent_dim", ARI_COL, sx, sy)

out_lat = "supp_latent_dim_sweep.png"
fig.savefig(out_lat, dpi=SAVE_DPI, bbox_inches="tight")
plt.show()
print(f"Saved: {out_lat} ({SAVE_DPI} dpi)")

# =============================================================================
# 2) Dropout sweeps (enc / dec) parsed from sweep labels
# =============================================================================
d_do = with_parsed_dropout(df)
d_do = d_do.dropna(subset=["drop_kind", "drop_p"]).copy()
d_do = filter_baseline_bg(d_do)

if d_do.empty:
    print("[skip] No dropout sweeps found matching sweep labels like drop_enc_0.1 / drop_dec_0.05")
else:
    METRICS = [
        (FOS_COL, "FOSCTTM_rankfrac (↓)", False),
        (LT_COL,  "LT macroF1 worse-dir (↑)", True),
    ]
    kinds = [
        ("enc", DEFAULT_DROP_ENC, "Encoder dropout"),
        ("dec", DEFAULT_DROP_DEC, "Decoder dropout"),
    ]

    nrows, ncols = 2, len(METRICS)
    #fig, axes = plt.subplots(nrows, ncols, figsize=(8, 3*nrows), constrained_layout=True)
    fig, axes = plt.subplots(nrows, ncols, figsize=(14, 3*nrows), constrained_layout=True)    
    axes = np.asarray(axes)
    if axes.ndim == 1:
        axes = axes[:, None]

    for r, (kind, x0, rowname) in enumerate(kinds):
        dk = d_do[d_do["drop_kind"] == kind].copy()
        if dk.empty:
            for c in range(ncols):
                axes[r, c].axis("off")
            axes[r, 0].set_title(f"{rowname}\n(no rows)")
            continue

        ycols = [m for (m, _, _) in METRICS]
        dd = agg_1d(dk, "drop_p", ycols, agg="median")
        x = dd["drop_p"].astype(float).to_numpy()

        for c, (mcol, mlabel, _) in enumerate(METRICS):
            y = dd[mcol].astype(float).to_numpy()
            sx, sy = star_from_exact(x, y, x0)
            plot_curve(axes[r, c], x, y, f"{rowname}: {mlabel}", "dropout p", mcol, sx, sy)

    out_do = "supp_dropout_sweeps.png"
    fig.savefig(out_do, dpi=SAVE_DPI, bbox_inches="tight")
    plt.show()
    print(f"Saved: {out_do} ({SAVE_DPI} dpi)")

# =============================================================================
# 3) Scaling curve (scaling_curve.tsv)  [CLEAN: spacing + legend + colors]
# =============================================================================
sc = pd.read_csv(SCALING_TSV, sep="\t")
print("Scaling TSV:", sc.shape)
display(sc.head(3))

X = "n_train_cells_eff"
if X not in sc.columns:
    raise ValueError(f"Missing expected scaling x-axis column: {X}")

TIME     = "wall_seconds"            if "wall_seconds"            in sc.columns else None
CPU_PEAK = "cpu_rss_peak_mb_sampled" if "cpu_rss_peak_mb_sampled" in sc.columns else None
VAL      = "best_val"                if "best_val"                in sc.columns else None
CPU_MEAN = "cpu_percent_mean"        if "cpu_percent_mean"        in sc.columns else None

sc = to_num(sc, [X, TIME, CPU_PEAK, VAL, CPU_MEAN])
scp = sc.dropna(subset=[X]).copy().sort_values(X)
x = scp[X].astype(float).to_numpy()

# --- Make the figure wider + reserve right margin for legend
#fig, axes = plt.subplots(2, 2, figsize=(11.5, 6.5))
fig, axes = plt.subplots(2, 2, figsize=(16, 8))
axes = axes.ravel()

# Manual spacing (THIS is what fixes title collisions)
#fig.subplots_adjust(left=0.08, right=0.76, top=0.92, bottom=0.10, wspace=0.55, hspace=0.45)
fig.subplots_adjust(left=0.08, right=0.70, top=0.92, bottom=0.10, wspace=0.55, hspace=0.45)

TITLE_KW = dict(fontsize=13, pad=10)

# (a) time
if TIME is not None:
    y = pd.to_numeric(scp[TIME], errors="coerce").to_numpy()
    axes[0].plot(x, y, marker="o", linewidth=1.6, markersize=4)
    axes[0].set_title("(a) Wall-clock time vs effective training size", **TITLE_KW)
    axes[0].set_xlabel(X)
    axes[0].set_ylabel(TIME)
    axes[0].grid(True, linewidth=0.3, alpha=0.5)
else:
    axes[0].axis("off")
    axes[0].set_title("(a) time\n(missing)", **TITLE_KW)

# (b) memory overlay
ax = axes[1]
ax.set_title("(b) Peak memory vs effective training size", **TITLE_KW)
ax.set_xlabel(X)
ax.grid(True, linewidth=0.3, alpha=0.5)

# CPU series (blue, left axis)
cpu_series = None
if CPU_PEAK is not None and CPU_PEAK in scp.columns:
    cpu_series = pd.to_numeric(scp[CPU_PEAK], errors="coerce").to_numpy()
have_cpu = cpu_series is not None and np.isfinite(cpu_series).any()

# GPU/MPS (orange, right axis) — pick best available
backend = str(scp["backend"].iloc[0]) if "backend" in scp.columns and len(scp) else ""
device  = str(scp["device"].iloc[0])  if "device"  in scp.columns and len(scp) else ""

mps_candidates     = ["mps_driver_peak_mb_sampled", "gpu_alloc_peak_mb_sampled", "mps_alloc_after_mb"]
cuda_candidates    = ["cuda_peak_alloc_mb", "cuda_peak_reserved_mb", "cuda_reserved_peak_mb_sampled"]
generic_candidates = ["gpu_peak_mem_mb_trainone", "gpu_alloc_peak_mb_sampled"]

if backend == "mps" or device == "mps":
    candidates = mps_candidates + generic_candidates
elif backend == "cuda" or device == "cuda":
    candidates = cuda_candidates + generic_candidates
else:
    candidates = generic_candidates

gpu_series, gpu_label = None, None
for c in candidates:
    if c in scp.columns:
        arr = pd.to_numeric(scp[c], errors="coerce").to_numpy()
        if np.isfinite(arr).any():
            gpu_series, gpu_label = arr, c
            break
have_gpu = gpu_series is not None and np.isfinite(gpu_series).any()

legend_handles = []
legend_labels  = []

if (not have_cpu) and (not have_gpu):
    ax.axis("off")
    ax.set_title("(b) peak memory\n(missing)", **TITLE_KW)
else:
    if have_cpu:
        ln_cpu = ax.plot(
            x, cpu_series,
            marker="o", linewidth=1.6, markersize=4,
            color="C0", zorder=3
        )[0]
        ax.set_ylabel("CPU MB", color="C0")
        ax.tick_params(axis="y", labelcolor="C0")
        legend_handles.append(ln_cpu)
        legend_labels.append("CPU peak (MB)")

    ax2 = None
    if have_gpu:
        ax2 = ax.twinx()
        ln_gpu = ax2.plot(
            x, gpu_series,
            marker="o", linewidth=1.6, markersize=4,
            color="C1", zorder=4
        )[0]
        ax2.set_ylabel("GPU/MPS MB", color="C1")
        ax2.tick_params(axis="y", labelcolor="C1")
        legend_handles.append(ln_gpu)
        legend_labels.append(f"GPU/MPS peak (MB): {gpu_label}")

# (c) best val
if VAL is not None:
    y = pd.to_numeric(scp[VAL], errors="coerce").to_numpy()
    axes[2].plot(x, y, marker="o", linewidth=1.6, markersize=4)
    axes[2].set_title("(c) Best validation loss vs size", **TITLE_KW)
    axes[2].set_xlabel(X)
    axes[2].set_ylabel(VAL)
    axes[2].grid(True, linewidth=0.3, alpha=0.5)
else:
    axes[2].axis("off")
    axes[2].set_title("(c) best_val\n(missing)", **TITLE_KW)

# (d) cpu%
if CPU_MEAN is not None:
    y = pd.to_numeric(scp[CPU_MEAN], errors="coerce").to_numpy()
    axes[3].plot(x, y, marker="o", linewidth=1.6, markersize=4)
    axes[3].set_title("(d) Mean CPU% vs size", **TITLE_KW)
    axes[3].set_xlabel(X)
    axes[3].set_ylabel(CPU_MEAN)
    axes[3].grid(True, linewidth=0.3, alpha=0.5)
else:
    axes[3].axis("off")
    axes[3].set_title("(d) CPU%\n(missing)", **TITLE_KW)

# Figure-level legend (never overlaps any panel)
if legend_handles:
    fig.legend(
        legend_handles, legend_labels,
        loc="upper left",
        bbox_to_anchor=(0.78, 0.92),
        fontsize=9,
        frameon=True,
        framealpha=0.95
    )

out_sc = "supp_scaling_curve.png"
fig.savefig(out_sc, dpi=SAVE_DPI, bbox_inches="tight")
plt.show()
print(f"Saved: {out_sc} ({SAVE_DPI} dpi)")


# %% [code]


# %% [code]


# %% [code]


# %% [code]


# %% [code]


# %% [code]


# %% [code]


# %% [code]


# %% [code]


# %% [code]


# %% [code]


# %% [code]


# %% [code]


# %% [code]


# %% [code]


# %% [code]


# %% [code]


# %% [code]


# %% [code]


# %% [code]


# %% [code]


# %% [code]


# %% [code]


# %% [code]
```
