## Supplemental Code for "Unifying multimodal single-cell data with a mixture-of-experts *β*-variational autoencoder framework": Supp_mouse_skin_SHAREseq.ipynb

````
# %% [markdown]
## UniVI cross-tissue generalization — SHARE-seq mouse skin (Ma et al. 2020)
### Train a bimodal UniVI model (RNA + ATAC) on paired SHARE-seq mouse skin hair follicle data to assess cross-tissue robustness of UniVI outside the hematopoietic context.

Andrew Ashford, Pathways + Omics Group, Oregon Health & Science University, Portland, OR

This notebook applies the UniVI RNA–ATAC pipeline (same architecture and hyperparameter regime as the 10x Multiome PBMC analysis) to the Ma et al. 2020 SHARE-seq mouse skin dataset (GEO: GSE140203). The goal is to demonstrate that UniVI's paired posterior alignment generalizes to a non-hematopoietic tissue with distinct cell-type architecture (cycling epithelium, outer root sheath, inner root sheath, medulla, interfollicular epidermis, dermal fibroblasts). Metrics reported here feed directly into the supplemental cross-tissue robustness figure.

**Data source:** Ma S et al., *Cell* 2020. GSE140203 — extract the mouse skin samples from `GSE140203_RAW.tar`.

GitHub for the project — including a Quickstart guide — can be found at: https://github.com/Ashford-A/UniVI

Package is pip installable via:
```bash
pip install univi
```

# %% [markdown]
#### Import modules

# %% [code]
# Import non-UniVI modules
import os
import re
import sys
import json
import gzip
from pathlib import Path

import numpy as np
import pandas as pd
import scipy.sparse as sp
import anndata as ad
import scanpy as sc

import torch
from torch.utils.data import DataLoader, Subset


# %% [code]
from sklearn.decomposition import TruncatedSVD
from sklearn.preprocessing import normalize
from sklearn.neighbors import NearestNeighbors
from sklearn.metrics import silhouette_score, accuracy_score, f1_score
from sklearn.metrics import pairwise_distances


# %% [code]
import matplotlib.pyplot as plt
import seaborn as sns


# %% [code]
# Import required UniVI modules
from univi import (
    ModalityConfig,
    UniVIConfig,
    TrainingConfig,
    UniVIMultiModalVAE,
    matching,
    UniVITrainer,
    write_univi_latent,
    MultiModalDataset,
)

import univi as uv
import univi.evaluation as univi_eval
import univi.plotting as up
from univi.data import MultiModalDataset, align_paired_obs_names

print("Installed version is univi v" + str(uv.__version__))


# %% [markdown]
### Specify device

# %% [code]
print("torch:", torch.__version__)
print("torch.cuda.is_available():", torch.cuda.is_available())
device = "cuda" if torch.cuda.is_available() else ("mps" if torch.backends.mps.is_available() else "cpu")
print("Using device:", device)


# %% [markdown]
### Specify file paths

**GEO download instructions:**
1. Download `GSE140203_RAW.tar` from https://www.ncbi.nlm.nih.gov/geo/query/acc.cgi?acc=GSE140203
2. Extract: `tar -xf GSE140203_RAW.tar -C ./data/SHARE-seq_skin/`
3. Keep only skin samples — look for files with `skin` in the name. Run the discovery cell below to confirm filenames.
4. (Optional) Download cell metadata/cluster labels from the paper's supplement or the authors' GitHub if available.

Expected files (exact names may vary by GEO submission version):
- RNA: `*skin*rna*counts*.txt.gz` or `*skin*RNA*.txt.gz`
- ATAC peaks: `*skin*atac*counts*.txt.gz` or `*skin*ATAC*.txt.gz`
- Cell metadata (optional): `*skin*metadata*.txt.gz` or `*skin*barcodes*.txt.gz`

# %% [markdown]
### Extract GSE140203_RAW.tar and identify skin files

Run the cell below once to extract the archive and list the skin-relevant files.
The full tar is ~7.4 GB so extraction takes a few minutes.
After extraction, the cell prints every filename containing `skin` so you can
confirm exact names before the `find_file` cell runs.

# %% [code]
DATA_DIR = "./data/SHARE-seq_skin"


# %% [code]
from pathlib import Path
import tarfile

# Coerce to Path regardless of how DATA_DIR was defined upstream
DATA_DIR = Path(DATA_DIR)

TAR_PATH     = DATA_DIR / "GSE140203_RAW.tar"
EXTRACT_ROOT = DATA_DIR / "GSE140203_RAW"   # extracted files land here
EXTRACT_ROOT.mkdir(exist_ok=True)

# ── 1. Check what's already extracted ──────────────────────────────────────
already = list(EXTRACT_ROOT.rglob("*"))
if already:
    print(f"Extraction directory already contains {len(already)} item(s) — skipping re-extract.")
    print("Delete or empty EXTRACT_ROOT to force re-extraction.")
else:
    print(f"Extracting {TAR_PATH.name} → {EXTRACT_ROOT} ...")
    print("(~7.4 GB, expect 2–5 minutes)")
    with tarfile.open(TAR_PATH, "r") as tar:
        tar.extractall(path=EXTRACT_ROOT)
    print("Extraction complete.")

# ── 2. Walk the extracted tree and report all files ────────────────────────
all_files = sorted(EXTRACT_ROOT.rglob("*"))
print(f"\nTotal files/dirs in {EXTRACT_ROOT.name}: {len(all_files)}")

# Show every file that mentions 'skin' (case-insensitive)
skin_files = [p for p in all_files if p.is_file() and "skin" in p.name.lower()]
print(f"\nFiles containing 'skin' ({len(skin_files)}):")
for p in skin_files:
    print(" ", p.relative_to(DATA_DIR))

if not skin_files:
    print("\nNo 'skin' files found. Listing ALL extracted files:")
    for p in sorted(all_files)[:60]:
        if p.is_file():
            print(" ", p.relative_to(DATA_DIR))


# %% [code]
DATA_DIR = Path(DATA_DIR)

# Check whether files are directly in EXTRACT_ROOT or one level deeper
direct_files = [p for p in EXTRACT_ROOT.iterdir() if p.is_file()]
subdirs      = [p for p in EXTRACT_ROOT.iterdir() if p.is_dir()]

if direct_files:
    DATA_DIR = EXTRACT_ROOT
    print(f"Files are flat in {DATA_DIR}")
elif subdirs:
    # Common: tar extracts into a single subfolder named after the accession
    DATA_DIR = subdirs[0]
    print(f"Files are in subdirectory: {DATA_DIR}")
    print("  Contents:")
    for p in sorted(DATA_DIR.iterdir())[:20]:
        print("   ", p.name)
else:
    print("Extraction directory is empty — check TAR_PATH and re-run.")

print(f"\nDATA_DIR is now: {DATA_DIR}")


# %% [code]
from pathlib import Path

FIGDIR   = Path("./figures/SHARE-seq_skin")
MODEL_DIR = Path("./saved_models/SHARE-seq_skin")

for d in [DATA_DIR, FIGDIR, MODEL_DIR]:
    d.mkdir(parents=True, exist_ok=True)

# Discover what files are present — run this first to confirm filenames
print("Files in data directory:")
for f in sorted(DATA_DIR.iterdir()):
    print(" ", f.name)
    

# %% [code]
# ── List all skin files to identify exact naming convention ─────────────────
all_skin = sorted(DATA_DIR.glob("*skin*"))

# Split by suffix / modality tag
skin_rna_files  = [p for p in all_skin if "rna" in p.name.lower() and p.suffix in (".gz", ".txt")]
skin_atac_files = [p for p in all_skin if "atac" in p.name.lower() and p.suffix in (".gz", ".txt")]
skin_counts     = [p for p in all_skin if "counts" in p.name.lower() and p.suffix == ".gz"]
skin_peaks      = [p for p in all_skin if "peaks" in p.name.lower() or "peak" in p.name.lower()]
skin_frags      = [p for p in all_skin if "fragment" in p.name.lower() or "frag" in p.name.lower()]
skin_barcodes   = [p for p in all_skin if "barcode" in p.name.lower()]

print("=== All skin files ===")
for p in all_skin:
    print(" ", p.name)

print(f"\nRNA counts  ({len(skin_rna_files)}): ", [p.name for p in skin_rna_files])
print(f"ATAC counts ({len(skin_atac_files)}): ",  [p.name for p in skin_atac_files])
print(f"All counts  ({len(skin_counts)}): ",      [p.name for p in skin_counts])
print(f"Peaks       ({len(skin_peaks)}): ",       [p.name for p in skin_peaks])
print(f"Fragments   ({len(skin_frags)}): ",       [p.name for p in skin_frags])
print(f"Barcodes    ({len(skin_barcodes)}): ",    [p.name for p in skin_barcodes])


# %% [code]
# ── Assign RNA and ATAC paths based on the listing above ────────────────────
#
# SHARE-seq on GEO names the RNA counts with an explicit 'rna' tag but the ATAC
# counts file often omits the modality token (same convention as the cell-line
# samples in this archive, e.g. GSM4156590_GM12878.rep1.counts.txt.gz).
#
# The logic below handles both layouts:
#   Layout A: both files have modality tags  →  *skin*rna*counts*.txt.gz  /  *skin*atac*counts*.txt.gz
#   Layout B: ATAC has no modality tag       →  *skin*rna*counts*.txt.gz  /  *skin*counts*.txt.gz (non-RNA)
#
# If multiple stages exist (early.anagen, late.anagen, telogen) ALL are loaded
# below and concatenated before training.

def assign_paths(data_dir: Path):
    all_skin = sorted(data_dir.glob("*skin*"))
    count_gz = [p for p in all_skin
                if p.name.endswith(".txt.gz") and "counts" in p.name.lower()]

    rna_files  = [p for p in count_gz if "rna"  in p.name.lower()]
    atac_files = [p for p in count_gz if "atac" in p.name.lower()]

    # Layout B: no 'atac' tag → ATAC is every counts file that is NOT an RNA file
    if not atac_files:
        rna_stems = {p.name.replace("rna", "") for p in rna_files}
        atac_files = [p for p in count_gz
                      if "rna" not in p.name.lower()
                      and p.name not in rna_stems]
        if not atac_files:
            # Last resort: all counts files that are not RNA
            atac_files = [p for p in count_gz if "rna" not in p.name.lower()]

    return sorted(rna_files), sorted(atac_files)

rna_files, atac_files = assign_paths(DATA_DIR)

print("RNA  files detected:", [p.name for p in rna_files])
print("ATAC files detected:", [p.name for p in atac_files])

if not rna_files:
    raise FileNotFoundError("No skin RNA counts files found. Check DATA_DIR listing above.")
if not atac_files:
    raise FileNotFoundError("No skin ATAC counts files found. Check DATA_DIR listing above.")

# Use all stages if present (late.anagen, early.anagen, telogen) — they are
# concatenated in the loading cell below. If you want a single stage, set:
#   rna_files  = [rna_files[0]]
#   atac_files = [atac_files[0]]
print(f"\nWill load {len(rna_files)} RNA stage(s) and {len(atac_files)} ATAC stage(s).")

# For backwards compat with single-file references elsewhere in the notebook:
RNA_PATH  = rna_files[0]
ATAC_PATH = atac_files[0]

# Also look for barcode/metadata files
META_CANDIDATES = (
    sorted(DATA_DIR.glob("*skin*meta*.txt.gz"))   +
    sorted(DATA_DIR.glob("*skin*barcode*.txt.gz")) +
    sorted(DATA_DIR.glob("*skin*cell*.txt.gz"))    +
    sorted(DATA_DIR.glob("*skin*cluster*.csv"))
)
META_PATH = META_CANDIDATES[0] if META_CANDIDATES else None
print(f"Metadata file: {META_PATH}")
print(f"RNA  : {RNA_PATH}")
print(f"ATAC : {ATAC_PATH}")


# %% [markdown]
### Helper functions

# %% [code]
def _to_dense(X):
    if sp.issparse(X):
        return X.toarray()
    return np.asarray(X)

def _as_csr(X, dtype=np.float32):
    if sp.issparse(X):
        return X.tocsr().astype(dtype)
    return sp.csr_matrix(np.asarray(X), dtype=dtype)

def _row_sums_csr(X):
    return np.asarray(X.sum(axis=1)).ravel().astype(np.float32)

import scipy.io as sio, io

def load_shareseq_mtx(counts_path: Path, data_dir: Path):
    """
    Handles two formats found in GSE140203:
      - MatrixMarket (.mtx-style .txt.gz): ATAC counts — rows=features, cols=cells
      - TSV (.txt.gz with named rows/cols):  RNA counts — rows=genes, cols=cells
    """
    prefix = counts_path.name.split(".counts.txt.gz")[0]

    def find_companion(*suffixes):
        for s in suffixes:
            hits = sorted(data_dir.glob(f"{prefix}{s}"))
            if hits:
                return hits[0]
        return None

    # Peek at first line to detect format
    with gzip.open(counts_path, "rt") as fh:
        first_line = fh.readline()

    is_mtx = first_line.strip().startswith("%%MatrixMarket")

    print(f"  {counts_path.name}  →  {'MatrixMarket' if is_mtx else 'TSV'}")

    # ── MatrixMarket path (ATAC) ──────────────────────────────────────────────
    if is_mtx:
        barcodes_path = find_companion(".barcodes.txt.gz", ".barcodes.txt",
                                       ".cell_barcodes.txt.gz")
        peaks_path    = find_companion(".peaks.bed.gz", ".peaks.txt.gz",
                                       ".peaks.bed", ".features.txt.gz")
        print(f"    barcodes : {barcodes_path.name if barcodes_path else 'NOT FOUND — integer index'}")
        print(f"    peaks    : {peaks_path.name    if peaks_path    else 'NOT FOUND — integer index'}")

        # Decompress to BytesIO so mmread gets a clean seekable stream
        buf = io.BytesIO()
        with gzip.open(counts_path, "rb") as fh:
            buf.write(fh.read())
        buf.seek(0)
        mat = sio.mmread(buf).tocsr().astype(np.float32).T.tocsr()  # → cells x features
        print(f"    Shape (cells x peaks): {mat.shape}")

        def read_lines(p):
            opener = gzip.open if str(p).endswith(".gz") else open
            with opener(p, "rt") as fh:
                return [l.strip() for l in fh if l.strip()]

        barcodes = read_lines(barcodes_path) if barcodes_path \
                   else [str(i) for i in range(mat.shape[0])]

        if peaks_path:
            raw = read_lines(peaks_path)
            features = []
            for line in raw:
                parts = line.split("\t")
                features.append(f"{parts[0]}:{parts[1]}-{parts[2]}"
                                 if len(parts) >= 3 else line)
        else:
            features = [str(i) for i in range(mat.shape[1])]

    # ── TSV path (RNA) ────────────────────────────────────────────────────────
    else:
        sep = "\t" if "\t" in first_line else " "
        chunks = []
        for chunk in pd.read_csv(counts_path, sep=sep, index_col=0,
                                 compression="gzip", chunksize=2000):
            chunks.append(chunk)
        df = pd.concat(chunks)
        print(f"    Raw shape (rows x cols): {df.shape}")

        # Detect orientation: if index looks like gene names → rows are genes
        first_idx = str(df.index[0])
        if not re.match(r"^\d+$", first_idx):     # index is not purely numeric → features
            df = df.T
            print(f"    Transposed to cells x features: {df.shape}")

        mat      = sp.csr_matrix(df.values.astype(np.float32))
        barcodes = df.index.astype(str).tolist()
        features = df.columns.astype(str).tolist()

    assert mat.shape[0] == len(barcodes), \
        f"Barcodes ({len(barcodes)}) != rows ({mat.shape[0]})"
    assert mat.shape[1] == len(features), \
        f"Features ({len(features)}) != cols ({mat.shape[1]})"

    return ad.AnnData(
        X=mat,
        obs=pd.DataFrame(index=pd.Index(barcodes)),
        var=pd.DataFrame(index=pd.Index(features)),
    )


# %% [markdown]
### Read in data

# %% [code]
# ── Load RNA (all stages) ───────────────────────────────────────────────────
# If multiple stages exist (e.g. early.anagen + late.anagen + telogen),
# they are each loaded separately then concatenated. Barcodes are made
# globally unique by appending the stage name.

# ── Load RNA + ATAC (all stages, MTX format) ─────────────────────────────────

def load_stage(rna_path, atac_path, data_dir):
    stage = rna_path.name.split("_", 1)[1].split(".counts")[0]
    # Strip modality token so both get the same stage tag
    stage = re.sub(r"\.(rna|atac)$", "", stage)   # skin.late.anagen.rna → skin.late.anagen
    print(f"\n=== Stage: {stage} ===")
    rna  = load_shareseq_mtx(rna_path,  data_dir)
    atac = load_shareseq_mtx(atac_path, data_dir)

    # Normalize barcode separator: RNA uses commas, ATAC uses dots — unify to dots
    rna.obs_names  = pd.Index([bc.replace(",", ".") for bc in rna.obs_names])
    atac.obs_names = pd.Index([bc.replace(",", ".") for bc in atac.obs_names])

    rna.obs_names  = pd.Index([f"{bc}__{stage}" for bc in rna.obs_names])
    atac.obs_names = pd.Index([f"{bc}__{stage}" for bc in atac.obs_names])
    rna.obs["stage"]  = stage
    atac.obs["stage"] = stage
    return rna, atac

rna_parts, atac_parts = [], []
for rp, ap in zip(rna_files, atac_files):
    r, a = load_stage(rp, ap, DATA_DIR)
    rna_parts.append(r)
    atac_parts.append(a)

if len(rna_parts) == 1:
    skin_rna  = rna_parts[0]
    skin_atac = atac_parts[0]
else:
    skin_rna  = ad.concat(rna_parts,  join="inner", merge="same")
    skin_atac = ad.concat(atac_parts, join="inner", merge="same")

skin_rna.var_names_make_unique()
skin_atac.var_names_make_unique()
print(f"\nRNA  shape: {skin_rna.shape}")
print(f"ATAC shape: {skin_atac.shape}")
print("RNA obs  (first 3):", skin_rna.obs_names[:3].tolist())
print("ATAC obs (first 3):", skin_atac.obs_names[:3].tolist())


# %% [markdown]
### QC filtering

# %% [code]
# ---- RNA QC ----
sc.pp.calculate_qc_metrics(skin_rna, percent_top=None, log1p=False, inplace=True)

print("RNA QC summary:")
print(f"  Total genes: {skin_rna.n_vars}")
print(f"  Median UMI per cell: {np.median(skin_rna.obs['total_counts']):.0f}")
print(f"  Median genes detected: {np.median(skin_rna.obs['n_genes_by_counts']):.0f}")

# QC thresholds
MIN_COUNTS_RNA = 300
MAX_COUNTS_RNA = 30_000
MIN_GENES      = 200
MAX_PCT_MT     = 5.0

# Mito % - prefer existing column from upstream Seurat analysis, else recompute
if "percent.mt" in skin_rna.obs.columns:
    pct_mt = skin_rna.obs["percent.mt"].values.astype(np.float32)
    print(f"  Using existing 'percent.mt' column.")
elif "pct_counts_mt" in skin_rna.obs.columns:
    pct_mt = skin_rna.obs["pct_counts_mt"].values.astype(np.float32)
    print(f"  Using existing 'pct_counts_mt' column.")
else:
    # Mouse mito genes use lowercase 'mt-' prefix
    skin_rna.var["mt"] = skin_rna.var_names.str.startswith("mt-")
    if not skin_rna.var["mt"].any():
        # Fallback: human or differently-cased data
        skin_rna.var["mt"] = skin_rna.var_names.str.upper().str.startswith("MT-")
    sc.pp.calculate_qc_metrics(
        skin_rna, qc_vars=["mt"], percent_top=None, log1p=False, inplace=True
    )
    pct_mt = skin_rna.obs["pct_counts_mt"].values.astype(np.float32)
    print(f"  Computed pct_counts_mt from {int(skin_rna.var['mt'].sum())} mito genes.")

skin_rna.obs["pct_mt"] = pct_mt
print(f"  Median pct_mt: {np.median(pct_mt):.2f}%")

rna_pass = (
    (skin_rna.obs["total_counts"] >= MIN_COUNTS_RNA) &
    (skin_rna.obs["total_counts"] <= MAX_COUNTS_RNA) &
    (skin_rna.obs["n_genes_by_counts"] >= MIN_GENES) &
    (pct_mt <= MAX_PCT_MT)
)
n_pass = int(rna_pass.sum())
print(f"  Cells passing RNA QC: {n_pass} / {skin_rna.n_obs}")
print(f"    Removed by counts/genes: {int((~rna_pass & (pct_mt <= MAX_PCT_MT)).sum())}")
print(f"    Removed by pct_mt > {MAX_PCT_MT}%: {int((pct_mt > MAX_PCT_MT).sum())}")

skin_rna = skin_rna[rna_pass].copy()


# %% [code]
# ---- ATAC QC ----
sc.pp.calculate_qc_metrics(skin_atac, percent_top=None, log1p=False, inplace=True)

print("ATAC QC summary:")
print(f"  Total peaks: {skin_atac.n_vars}")
print(f"  Median fragments (peak counts) per cell: {np.median(skin_atac.obs['total_counts']):.0f}")

MIN_COUNTS_ATAC = 250
MAX_COUNTS_ATAC = 12_500

atac_pass = (
    (skin_atac.obs["total_counts"] >= MIN_COUNTS_ATAC) &
    (skin_atac.obs["total_counts"] <= MAX_COUNTS_ATAC)
)
print(f"  Cells passing ATAC QC: {int(atac_pass.sum())} / {skin_atac.n_obs}")
skin_atac = skin_atac[atac_pass].copy()


# %% [code]
# ---- ENCODE mm10 blacklist filter for ATAC peaks ----
# Removes peaks overlapping ENCODE blacklist regions (high-mappability satellite/repeat
# regions known to produce artifactual signal). Standard step in any rigorous ATAC pipeline.
# Download once with:
#   wget -O ./data/SHARE-seq_skin/mm10-blacklist.v2.bed.gz \
#     https://github.com/Boyle-Lab/Blacklist/raw/master/lists/mm10-blacklist.v2.bed.gz

BLACKLIST_PATH = Path("./data/SHARE-seq_skin/mm10-blacklist.v2.bed.gz")

if not BLACKLIST_PATH.exists():
    print(f"WARN: Blacklist file not found at {BLACKLIST_PATH}")
    print("To enable, download with:")
    print(f"  wget -O {BLACKLIST_PATH} \\")
    print(f"    https://github.com/Boyle-Lab/Blacklist/raw/master/lists/mm10-blacklist.v2.bed.gz")
    print("Skipping blacklist filter (results still valid, just less stringent).")
else:
    # Load blacklist intervals per chromosome
    bl_by_chrom = {}
    with gzip.open(BLACKLIST_PATH, "rt") as fh:
        for line in fh:
            parts = line.strip().split("\t")
            if len(parts) >= 3:
                chrom, start, end = parts[0], int(parts[1]), int(parts[2])
                bl_by_chrom.setdefault(chrom, []).append((start, end))

    n_bl_intervals = sum(len(v) for v in bl_by_chrom.values())
    print(f"Loaded {n_bl_intervals} blacklist intervals across {len(bl_by_chrom)} chromosomes.")

    # Parse peak coords from var_names (format "chrN:start-end")
    import re as _re
    peak_re = _re.compile(r"^(chr[\w]+):(\d+)-(\d+)$")

    peak_chroms = np.empty(skin_atac.n_vars, dtype=object)
    peak_starts = np.zeros(skin_atac.n_vars, dtype=np.int64)
    peak_ends   = np.zeros(skin_atac.n_vars, dtype=np.int64)
    n_parsed    = 0
    for i, name in enumerate(skin_atac.var_names):
        m = peak_re.match(str(name))
        if m:
            peak_chroms[i] = m.group(1)
            peak_starts[i] = int(m.group(2))
            peak_ends[i]   = int(m.group(3))
            n_parsed += 1
        else:
            peak_chroms[i] = "_UNPARSED_"

    if n_parsed < skin_atac.n_vars:
        print(f"WARN: only {n_parsed}/{skin_atac.n_vars} peak names matched 'chrN:start-end'.")

    # Vectorized overlap test per chromosome
    keep = np.ones(skin_atac.n_vars, dtype=bool)
    for chrom, intervals in bl_by_chrom.items():
        mask = peak_chroms == chrom
        if not mask.any():
            continue
        chrom_idx = np.where(mask)[0]
        ps = peak_starts[chrom_idx]
        pe = peak_ends[chrom_idx]
        bl_arr = np.array(intervals, dtype=np.int64)
        # Overlap: peak_start < bl_end AND bl_start < peak_end
        for bs, be in bl_arr:
            overlap = (ps < be) & (bs < pe)
            if overlap.any():
                keep[chrom_idx[overlap]] = False

    n_dropped = int((~keep).sum())
    print(f"Blacklist-overlapping peaks removed: {n_dropped} / {skin_atac.n_vars} "
          f"({n_dropped/max(skin_atac.n_vars,1):.1%})")
    skin_atac = skin_atac[:, keep].copy()
    print(f"ATAC peaks after blacklist filter: {skin_atac.n_vars}")
    

# %% [code]
# QC violin plots
fig, axes = plt.subplots(1, 3, figsize=(12, 4))
axes[0].violinplot(skin_rna.obs["total_counts"], showmedians=True)
axes[0].set_title("RNA total counts")
axes[1].violinplot(skin_rna.obs["n_genes_by_counts"], showmedians=True)
axes[1].set_title("RNA genes detected")
axes[2].violinplot(skin_atac.obs["total_counts"], showmedians=True)
axes[2].set_title("ATAC total counts")
plt.tight_layout()
plt.savefig(FIGDIR / "qc_violins.png", dpi=150)
plt.show()
plt.close()


# %% [markdown]
### Align paired barcodes (RNA ↔ ATAC)

SHARE-seq barcodes are combinatorial indices shared between RNA and ATAC; paired cells have identical barcodes. We take the intersection to build the strictly paired training set.

# %% [code]
# Intersect barcodes
common_barcodes = skin_rna.obs_names.intersection(skin_atac.obs_names)
print(f"Cells with both RNA and ATAC: {len(common_barcodes)}")
print(f"  RNA-only: {skin_rna.n_obs - len(common_barcodes)}")
print(f"  ATAC-only: {skin_atac.n_obs - len(common_barcodes)}")

skin_rna  = skin_rna[common_barcodes].copy()
skin_atac = skin_atac[common_barcodes].copy()

# Ensure identical order
skin_atac = skin_atac[skin_rna.obs_names].copy()
assert np.array_equal(skin_rna.obs_names.values, skin_atac.obs_names.values), \
    "obs_names must match exactly after intersection."

print(f"\nFinal paired dataset: {skin_rna.n_obs} cells")


# %% [markdown]
### Load cell type labels (if available)

Ma et al. 2020 provides cluster annotations (IFE, HF cycling, ORS, IRS, medulla, sebaceous, dermal papilla, fibroblast) in their supplemental tables. If the metadata file is absent, we derive provisional labels via Leiden clustering on the RNA for stratification purposes only — these are replaced by the publication labels once available.

# %% [code]
CELLTYPE_KEY = "cell_type"

celltype_candidates = sorted(DATA_DIR.glob("*skin*celltype*.txt.gz")) + \
                      sorted(DATA_DIR.glob("*skin*celltype*.txt"))
META_PATH = celltype_candidates[0] if celltype_candidates else None

if META_PATH is not None:
    with gzip.open(META_PATH, "rt") as fh:
        meta = pd.read_csv(fh, sep="\t", header=0)  # row 0 IS the header

    # Normalize: atac.bc and rna.bc both use dots already
    meta["atac.bc"] = meta["atac.bc"].str.replace(",", ".", regex=False)
    meta["rna.bc"]  = meta["rna.bc"].str.replace(",", ".", regex=False)

    print(f"  {meta.shape[0]} cells, columns: {meta.columns.tolist()}")
    print(meta["celltype"].value_counts().head(10))

    # Build lookup dicts: bare_barcode → celltype
    atac_to_ct = dict(zip(meta["atac.bc"], meta["celltype"]))
    rna_to_ct  = dict(zip(meta["rna.bc"],  meta["celltype"]))

    # Strip __stage suffix from obs_names to get bare barcodes
    def strip_stage(obs_names):
        return pd.Index([bc.split("__")[0] for bc in obs_names])

    rna_bare  = strip_stage(skin_rna.obs_names)
    atac_bare = strip_stage(skin_atac.obs_names)

    skin_rna.obs[CELLTYPE_KEY]  = [rna_to_ct.get(bc, "Unknown") for bc in rna_bare]
    #skin_atac.obs[CELLTYPE_KEY] = [atac_to_ct.get(bc, "Unknown") for bc in atac_bare]
    skin_atac.obs[CELLTYPE_KEY] = [rna_to_ct.get(bc, "Unknown") for bc in atac_bare]

    print(f"\n  RNA  matched: {(skin_rna.obs[CELLTYPE_KEY]  != 'Unknown').sum()} / {skin_rna.n_obs}")
    print(f"  ATAC matched: {(skin_atac.obs[CELLTYPE_KEY] != 'Unknown').sum()} / {skin_atac.n_obs}")
    print(skin_rna.obs[CELLTYPE_KEY].value_counts())

else:
    # Fallback Leiden — subsample to keep it fast on 40k+ cells
    print("No celltype file found — running Leiden on 10k subsample for stratification labels.")
    rng = np.random.default_rng(0)
    sub_idx = rng.choice(skin_rna.n_obs, size=min(10_000, skin_rna.n_obs), replace=False)
    tmp = skin_rna[sub_idx].copy()
    sc.pp.normalize_total(tmp, target_sum=1e4)
    sc.pp.log1p(tmp)
    sc.pp.highly_variable_genes(tmp, n_top_genes=2000)
    sc.pp.pca(tmp, use_highly_variable=True)
    sc.pp.neighbors(tmp, n_neighbors=30)
    sc.tl.leiden(tmp, resolution=0.4, key_added=CELLTYPE_KEY,
                 flavor="igraph", n_iterations=2, directed=False)
    skin_rna.obs[CELLTYPE_KEY]  = "Unknown"
    skin_rna.obs.iloc[sub_idx, skin_rna.obs.columns.get_loc(CELLTYPE_KEY)] = \
        tmp.obs[CELLTYPE_KEY].values
    skin_atac.obs[CELLTYPE_KEY] = skin_rna.obs[CELLTYPE_KEY].values.copy()
    print(skin_rna.obs[CELLTYPE_KEY].value_counts())


# %% [code]
# ---- Filter "Mix" cell type ----
# The "Mix" label in Ma et al. 2020 metadata is the original authors\' catch-all
# for cells they couldn\'t confidently assign — typically doublets, transitional
# states, and ambiguous cells. Most published benchmarks on this dataset exclude
# or treat them separately. Including them in training contaminates alignment metrics.

n_mix_rna  = int((skin_rna.obs[CELLTYPE_KEY]  == "Mix").sum())
n_mix_atac = int((skin_atac.obs[CELLTYPE_KEY] == "Mix").sum())

print(f"Cells labeled 'Mix' (RNA):  {n_mix_rna}")
print(f"Cells labeled 'Mix' (ATAC): {n_mix_atac}")

skin_rna  = skin_rna[skin_rna.obs[CELLTYPE_KEY]   != "Mix"].copy()
skin_atac = skin_atac[skin_atac.obs[CELLTYPE_KEY] != "Mix"].copy()

print(f"After Mix removal -- RNA: {skin_rna.n_obs}, ATAC: {skin_atac.n_obs}")
print(f"Remaining cell types: {skin_rna.obs[CELLTYPE_KEY].nunique()}")


# %% [markdown]
### Preprocessing

Identical pipelines to the Multiome PBMC analysis: RNA → log1p + HVG + *Z*-score; ATAC → TF-IDF → LSI → *Z*-score. Parameters are kept the same to ensure comparability.

# %% [code]
# ── Global hyperparameters ───────────────────────────────────────────────────
SPLIT_SEED     = 42
HVG_TOP_N      = 5000
RNA_TARGET_SUM = 1e4

# LSI components
ATAC_N_LSI     = 101

# Binarize ATAC counts before TF-IDF (Cusanovich/Signac/MUON convention).
ATAC_BINARIZE  = False

# TF-IDF method:
#   "log_tfidf" — log(TF * IDF * scale_factor + 1)  (Signac default; better
#                 discrimination on sparse data, more ideal for SHARE-seq)
#   "tfidf"     — plain TF * IDF                    (Cusanovich 2018 original)
ATAC_TFIDF_METHOD = "log_tfidf"
ATAC_TFIDF_SCALE  = 1e4   # only used for "log_tfidf"

BATCH_SIZE     = 128
NUM_WORKERS    = 0
PIN_MEMORY     = False


# %% [code]
# ── Preprocessing utility functions ─────────────────────────────────────────

EPS = 1e-8

def _to_dense(X, dtype=np.float32):
    return X.toarray().astype(dtype) if sp.issparse(X) else np.asarray(X, dtype=dtype)

def get_counts_matrix(adata, layer="counts"):
    """Return the count matrix from .layers[layer] if present, else .X."""
    if layer in adata.layers:
        return adata.layers[layer]
    return adata.X

# ── RNA ──────────────────────────────────────────────────────────────────────

def normalize_log1p(X, target_sum=1e4):
    if sp.issparse(X):
        X = X.tocsr().astype(np.float32)
        rs = np.maximum(_row_sums_csr(X), 1.0)
        Xn = sp.diags(target_sum / rs) @ X
        Xlog = Xn.copy()
        Xlog.data = np.log1p(Xlog.data)
        return Xlog
    X = np.asarray(X, dtype=np.float32)
    rs = np.maximum(X.sum(axis=1, keepdims=True), 1.0)
    return np.log1p((X / rs) * target_sum).astype(np.float32)


def hvg_exclusion_mask(var_names, exclude_prefixes=("mt-", "Mt-", "MT-", "Rps", "Rpl",
                                                    "Mrps", "Mrpl"),
                       exclude_suffixes=("Rik",),
                       exclude_contains=("-ps",),
                       cell_cycle_genes=None):
    """
    Return a boolean mask of var_names to *exclude* from HVG candidate pool.

    Removes mito, ribo (cytoplasmic + mitoribosomal), predicted/uncharacterized
    Riken genes, and pseudogenes. These are routinely high-variance for
    technical/stress reasons and don't carry cell-type identity — letting them
    occupy HVG slots crowds out actually-discriminative genes.

    Pass cell_cycle_genes as an iterable to also exclude S-phase/G2M markers.
    """
    var_names = pd.Index(var_names)
    excl = pd.Series(False, index=var_names)
    for p in exclude_prefixes:
        excl |= var_names.str.startswith(p)
    for s in exclude_suffixes:
        excl |= var_names.str.endswith(s)
    for c in exclude_contains:
        excl |= var_names.str.contains(c, regex=False)
    if cell_cycle_genes is not None:
        excl |= var_names.isin(list(cell_cycle_genes))
    return excl.values


def fit_hvg_seurat_v3(X_counts_train, var_names, top_n=5000,
                      exclude_mask=None, min_cells=10):
    """
    Seurat-v3 HVG selection on RAW counts (not log-transformed).

    Models mean-variance relationship via loess regression on log(var) vs
    log(mean), ranks genes by standardized variance. Robust on low-capture
    data where dispersion-based methods (var/mean ratio) get dominated by
    estimation noise — exactly your SHARE-seq case (median ~845 UMI/cell).

    Returns: (hvg_var_names, hvg_indices_into_original_var_names)
    """
    var_names = pd.Index(var_names)
    n_genes = len(var_names)

    # Drop genes detected in fewer than min_cells (seurat_v3 expects this)
    X = _as_csr(X_counts_train)
    n_cells_per_gene = np.asarray((X > 0).sum(axis=0)).ravel()
    candidate_mask = n_cells_per_gene >= int(min_cells)

    # Apply exclusion mask if provided
    if exclude_mask is not None:
        excl = np.asarray(exclude_mask).astype(bool)
        if excl.shape[0] != n_genes:
            raise ValueError(f"exclude_mask length {excl.shape[0]} != n_genes {n_genes}")
        candidate_mask &= ~excl
        n_excluded = int(excl.sum())
        print(f"  Excluded {n_excluded} genes from HVG pool by family mask "
              f"(mito/ribo/Rik/pseudogenes/cell-cycle).")

    print(f"  Candidate genes for HVG selection: {int(candidate_mask.sum())} / {n_genes}")

    # Run scanpy's seurat_v3 on the candidate gene set
    tmp = ad.AnnData(
        X=X[:, candidate_mask].copy(),
        var=pd.DataFrame(index=var_names[candidate_mask]),
    )
    sc.pp.highly_variable_genes(
        tmp, n_top_genes=int(top_n), flavor="seurat_v3", layer=None,
    )

    hvg_local = tmp.var["highly_variable"].values
    hvg_names = pd.Index(tmp.var_names[hvg_local])

    # Map selected gene names back to indices in the *original* var_names
    var_idx_lookup = pd.Series(np.arange(n_genes), index=var_names)
    hvg_idx = var_idx_lookup.loc[hvg_names].values.astype(np.int64)

    return hvg_names, hvg_idx


def fit_zscore_params(X, eps=EPS):
    if sp.issparse(X):
        X = X.tocsr().astype(np.float32)
        mean = np.asarray(X.mean(axis=0)).ravel().astype(np.float32)
        mean2 = np.asarray(X.power(2).mean(axis=0)).ravel().astype(np.float32)
        std = np.sqrt(np.maximum(mean2 - mean**2, 0.0) + eps).astype(np.float32)
    else:
        X = np.asarray(X, dtype=np.float32)
        mean = X.mean(axis=0).astype(np.float32)
        std = (X.std(axis=0) + eps).astype(np.float32)
    return mean, std

def apply_zscore(X, mean, std, clip=None):
    Xd = _to_dense(X)
    Z = (Xd - mean.reshape(1, -1)) / std.reshape(1, -1)
    if clip is not None:
        Z = np.clip(Z, -float(clip), float(clip))
    return Z.astype(np.float32)

# ── ATAC ─────────────────────────────────────────────────────────

def fit_tfidf_idf(X_counts):
    X = _as_csr(X_counts)
    n_cells = X.shape[0]
    df = X.getnnz(axis=0).astype(np.float32)
    return np.log1p(n_cells / (1.0 + df)).astype(np.float32)

def apply_tfidf(X_counts, idf, l2_norm=True, binarize=True,
                method="log_tfidf", scale_factor=1e4):
    X = _as_csr(X_counts)
    if binarize:
        X = X.copy()
        X.data = (X.data > 0).astype(np.float32)
    rs = np.maximum(_row_sums_csr(X), 1.0)
    tfidf = sp.diags(1.0 / rs) @ X
    tfidf = tfidf.multiply(idf).astype(np.float32)
    if method == "log_tfidf":
        tfidf = tfidf.multiply(float(scale_factor))
        tfidf.data = np.log1p(tfidf.data)
    elif method != "tfidf":
        raise ValueError(f"Unknown TF-IDF method: {method!r}.")
    tfidf.eliminate_zeros()
    if l2_norm:
        tfidf = normalize(tfidf, norm="l2", axis=1, copy=False)
    return tfidf

def fit_lsi(tfidf_train, n_components=101, seed=0):
    if not sp.issparse(tfidf_train):
        tfidf_train = _as_csr(tfidf_train)
    svd = TruncatedSVD(n_components=int(n_components), random_state=seed)
    Z = svd.fit_transform(tfidf_train).astype(np.float32)
    return svd, Z

def apply_lsi(tfidf, svd):
    if not sp.issparse(tfidf):
        tfidf = _as_csr(tfidf)
    return svd.transform(tfidf).astype(np.float32)
    

# %% [markdown]
### Train/val/test splits

# %% [code]
def stratified_split_indices(labels, train_frac=0.80, val_frac=0.10, test_frac=0.10,
                              seed=0, min_per_class=3):
    assert abs(train_frac + val_frac + test_frac - 1.0) < 1e-8
    rng = np.random.default_rng(seed)
    labels = pd.Series(labels).astype("category")
    idx_all = np.arange(labels.shape[0])
    train_idx, val_idx, test_idx = [], [], []
    for cat in labels.cat.categories:
        idx = idx_all[labels.values == cat]
        if idx.size < min_per_class:
            train_idx.extend(idx.tolist())
            continue
        rng.shuffle(idx)
        n = idx.size
        n_train = int(np.floor(train_frac * n))
        n_val   = int(np.floor(val_frac * n))
        train_idx.extend(idx[:n_train].tolist())
        val_idx.extend(idx[n_train:n_train+n_val].tolist())
        test_idx.extend(idx[n_train+n_val:].tolist())
    return {
        "train": np.array(train_idx, dtype=np.int64),
        "val":   np.array(val_idx,   dtype=np.int64),
        "test":  np.array(test_idx,  dtype=np.int64),
    }

splits = stratified_split_indices(
    skin_rna.obs[CELLTYPE_KEY].values, seed=SPLIT_SEED
)

skin_rna.obs["split"]  = "unsplit"
skin_atac.obs["split"] = "unsplit"
for k, idx in splits.items():
    skin_rna.obs.iloc[idx,  skin_rna.obs.columns.get_loc("split")]  = k
    skin_atac.obs.iloc[idx, skin_atac.obs.columns.get_loc("split")] = k

for split_name, idx in splits.items():
    print(f"  {split_name}: {len(idx)} cells")


# %% [markdown]
### Fit preprocessing on training cells and apply to all splits

Feature selectors (HVGs, IDF, SVD) are fit on training data only, then applied identically to val/test cells — same protocol as the Multiome PBMC analysis.

# %% [code]
# ── RNA ──────────────────────────────────────────────────────────────────────

# Store raw counts before any normalization (needed for reconstruction metrics)
skin_rna.layers["counts"] = skin_rna.X.copy()

train_idx = splits["train"]
rna_train_X = get_counts_matrix(skin_rna, "counts")[train_idx, :]

rna_train_log = normalize_log1p(rna_train_X, target_sum=RNA_TARGET_SUM)

excl = hvg_exclusion_mask(skin_rna.var_names)

print(f"\nHVG selection (seurat_v3 on raw counts, top {HVG_TOP_N}):")

hvg_genes, hvg_idx = fit_hvg_seurat_v3(
    X_counts_train=rna_train_X,           # RAW counts, not log!
    var_names=skin_rna.var_names,
    top_n=HVG_TOP_N,
    exclude_mask=excl,
    min_cells=10,
)
print(f"  HVGs selected: {len(hvg_genes)}")
print(f"  First 10: {hvg_genes[:10].tolist()}")

rna_train_log_hvg = (rna_train_log[:, hvg_idx] if sp.issparse(rna_train_log)
                     else rna_train_log[:, hvg_idx])
rna_mean, rna_std = fit_zscore_params(rna_train_log_hvg)

def apply_rna_pipeline(adata):
    Xc   = get_counts_matrix(adata, "counts")
    Xlog = normalize_log1p(Xc, target_sum=RNA_TARGET_SUM)
    hvg_loc = pd.Index(adata.var_names).get_indexer(hvg_genes)
    if np.any(hvg_loc < 0):
        raise ValueError(f"Missing HVGs: {list(hvg_genes[hvg_loc < 0][:5])}")
    Xlog_hvg = Xlog[:, hvg_loc] if sp.issparse(Xlog) else Xlog[:, hvg_loc]
    adata.obsm["X_rna_scaled_hvg"] = apply_zscore(Xlog_hvg, rna_mean, rna_std)
    # do NOT store log1p layer — not needed for training
    return adata

skin_rna = apply_rna_pipeline(skin_rna)
print("RNA pipeline done:", skin_rna.obsm["X_rna_scaled_hvg"].shape)


# %% [code]
# ── ATAC peak filtering + binarized TF-IDF + LSI ────────────────────────────

skin_atac.layers["counts"] = skin_atac.X.copy()

atac_train_X = get_counts_matrix(skin_atac, "counts")[train_idx, :]
n_train_cells = atac_train_X.shape[0]

peak_frac = np.asarray((atac_train_X > 0).sum(axis=0)).ravel() / n_train_cells

PEAK_MIN_FRAC = 0.0025   # loosened to preserve rare-population markers
PEAK_MAX_FRAC = 0.80
keep_peaks = (peak_frac > PEAK_MIN_FRAC) & (peak_frac < PEAK_MAX_FRAC)

print(f"Peak filter [{PEAK_MIN_FRAC:.0%}, {PEAK_MAX_FRAC:.0%}]: keeping "
      f"{keep_peaks.sum()} / {len(keep_peaks)} peaks ({keep_peaks.mean():.1%})")

skin_atac = skin_atac[:, keep_peaks].copy()
atac_train_X = get_counts_matrix(skin_atac, "counts")[train_idx, :]

# Fit TF-IDF + LSI on train cells
print(f"\nTF-IDF method: {ATAC_TFIDF_METHOD}, binarize: {ATAC_BINARIZE}")
idf            = fit_tfidf_idf(atac_train_X)
tfidf_train    = apply_tfidf(atac_train_X, idf=idf,
                              binarize=ATAC_BINARIZE,
                              method=ATAC_TFIDF_METHOD,
                              scale_factor=ATAC_TFIDF_SCALE)
svd, lsi_train = fit_lsi(tfidf_train, n_components=ATAC_N_LSI, seed=SPLIT_SEED)
lsi_mean       = lsi_train.mean(axis=0).astype(np.float32)
lsi_std        = (lsi_train.std(axis=0) + EPS).astype(np.float32)

del atac_train_X, tfidf_train, lsi_train
import gc; gc.collect()


# %% [code]
def apply_atac_pipeline(adata):
    """Apply the train-fit TF-IDF/LSI/z-score transform to all cells."""
    Xc         = get_counts_matrix(adata, "counts")
    tfidf      = apply_tfidf(Xc, idf=idf,
                              binarize=ATAC_BINARIZE,
                              method=ATAC_TFIDF_METHOD,
                              scale_factor=ATAC_TFIDF_SCALE)
    lsi        = apply_lsi(tfidf, svd)
    del tfidf
    lsi_scaled = (lsi - lsi_mean) / lsi_std
    del lsi
    adata.obsm["X_lsi_scaled"] = lsi_scaled.astype(np.float32)
    return adata

skin_atac = apply_atac_pipeline(skin_atac)
print("ATAC pipeline done:", skin_atac.obsm["X_lsi_scaled"].shape)

var_exp = svd.explained_variance_ratio_
sv      = svd.singular_values_

print(f"  LSI_0 variance explained:      {var_exp[0]:.4f}  (will be dropped)")
print(f"  LSI_1 variance explained:      {var_exp[1]:.4f}")
print(f"  LSI_1..{ATAC_N_LSI-1} cumulative:        {var_exp[1:].sum():.4f}")
print(f"  Singular value decay sv[0]/sv[-1]: {sv[0]/sv[-1]:.2f}x")
print(f"  Healthy multiomic ATAC LSI: SV decay >= 5x AND clear knee in spectrum")

fig, axes = plt.subplots(1, 2, figsize=(11, 3.5))
axes[0].plot(sv, marker="o", ms=3)
axes[0].set_yscale("log")
axes[0].set_xlabel("LSI component")
axes[0].set_ylabel("Singular value (log)")
axes[0].set_title("Singular value spectrum")
axes[0].axvline(0, color="red", ls="--", alpha=0.5, label="LSI_0 (depth, dropped)")
axes[0].legend()

axes[1].plot(np.cumsum(var_exp), marker="o", ms=3)
axes[1].set_xlabel("LSI component")
axes[1].set_ylabel("Cumulative variance explained")
axes[1].set_title("Cumulative explained variance")
axes[1].legend()

plt.tight_layout()
plt.savefig(FIGDIR / "atac_lsi_diagnostics.png", dpi=150)
plt.show()
plt.close()


# %% [code]
# ---- Pre-training diagnostic: UMAP on LSI to verify cell-type structure ----
# If cell types cluster well in LSI space alone, the encoder has good signal
# to learn from and alignment is plausibly achievable. If cell types are mixed
# in LSI space, the LSI is information-starved regardless of model tuning —
# fix preprocessing before tuning hyperparameters.

from umap import UMAP

# Subsample for speed
n_diag = min(8000, skin_atac.n_obs)
rng = np.random.default_rng(0)
diag_idx = rng.choice(skin_atac.n_obs, size=n_diag, replace=False)

X_lsi_diag = skin_atac.obsm["X_lsi_scaled"][diag_idx, 1:]   # drop LSI_0
ct_diag = skin_atac.obs[CELLTYPE_KEY].values[diag_idx]

print(f"Computing UMAP on {n_diag} cells x {X_lsi_diag.shape[1]} LSI dims...")
umap_lsi = UMAP(n_neighbors=15, min_dist=0.3, random_state=0,
                metric="euclidean").fit_transform(X_lsi_diag)

fig, ax = plt.subplots(figsize=(11, 7))
celltypes = sorted(pd.unique(ct_diag))
cmap = plt.cm.tab20(np.linspace(0, 1, len(celltypes)))
for ct, c in zip(celltypes, cmap):
    mask = ct_diag == ct
    ax.scatter(umap_lsi[mask, 0], umap_lsi[mask, 1], c=[c], s=3, alpha=0.7,
               label=f"{ct} (n={int(mask.sum())})")
ax.legend(bbox_to_anchor=(1.02, 1), loc="upper left", fontsize=7,
          markerscale=2.5, frameon=False)
ax.set_xlabel("UMAP1"); ax.set_ylabel("UMAP2")
ax.set_title("ATAC LSI space (pre-training): cell-type separation diagnostic")
plt.tight_layout()
plt.savefig(FIGDIR / "atac_lsi_celltype_umap.png", dpi=150, bbox_inches="tight")
plt.show()
plt.close()
    

# %% [code]
# ---- Pre-training diagnostic: UMAP on RNA z-scored HVGs ----
# Same goal as the ATAC LSI diagnostic: check whether the input representation
# already has clear cell-type structure before the encoder sees it.

from umap import UMAP
from sklearn.decomposition import PCA

# Subsample for speed
n_diag = min(8000, skin_rna.n_obs)
rng = np.random.default_rng(0)
diag_idx = rng.choice(skin_rna.n_obs, size=n_diag, replace=False)

X_rna_diag = skin_rna.obsm["X_rna_scaled_hvg"][diag_idx]
ct_diag = skin_rna.obs[CELLTYPE_KEY].values[diag_idx]

print(f"Computing PCA + UMAP on {n_diag} cells x {X_rna_diag.shape[1]} HVG dims...")
pca = PCA(n_components=50, random_state=0)
X_rna_pca = pca.fit_transform(X_rna_diag)
print(f"  PCA top-50 cumulative variance: {pca.explained_variance_ratio_.sum():.3f}")

umap_rna = UMAP(n_neighbors=15, min_dist=0.3, random_state=0,
                metric="euclidean").fit_transform(X_rna_pca)

fig, ax = plt.subplots(figsize=(11, 7))
celltypes = sorted(pd.unique(ct_diag))
cmap = plt.cm.tab20(np.linspace(0, 1, len(celltypes)))
for ct, c in zip(celltypes, cmap):
    mask = ct_diag == ct
    ax.scatter(umap_rna[mask, 0], umap_rna[mask, 1], c=[c], s=3, alpha=0.7,
               label=f"{ct} (n={int(mask.sum())})")
ax.legend(bbox_to_anchor=(1.02, 1), loc="upper left", fontsize=7,
          markerscale=2.5, frameon=False)
ax.set_xlabel("UMAP1"); ax.set_ylabel("UMAP2")
ax.set_title("RNA z-scored HVG space (pre-training): cell-type separation diagnostic")
plt.tight_layout()
plt.savefig(FIGDIR / "rna_celltype_umap.png", dpi=150, bbox_inches="tight")
plt.show()
plt.close()
    

# %% [code]
skin_rna_input = ad.AnnData(
    X=skin_rna.obsm["X_rna_scaled_hvg"],   # already dense float32 from apply_zscore
    obs=skin_rna.obs[["split", "cell_type", "stage"]].copy(),
    var=pd.DataFrame(index=hvg_genes),
)

skin_atac_input = ad.AnnData(
    X=skin_atac.obsm["X_lsi_scaled"][:, 1:],   # drop LSI_0 (depth) -> 50 dims
    obs=skin_atac.obs[["split", "cell_type", "stage"]].copy(),
    var=pd.DataFrame(index=[f"LSI_{i}" for i in range(1, ATAC_N_LSI)]),
)

print("UniVI input:")
print("  RNA  :", skin_rna_input.shape,  "(z-scored log1p HVGs)")
print("  ATAC :", skin_atac_input.shape, "(z-scored LSI, LSI_0 dropped)")


# %% [markdown]
### Build MultiModalDataset and DataLoaders

# %% [code]
def idx_from_split(adata, split_name):
    return np.where(adata.obs["split"].values == split_name)[0].astype(np.int32)

def make_subset_loader(dataset, idx, batch_size=128, shuffle=False,
                       num_workers=0, pin_memory=False):
    ds = Subset(dataset, idx)
    return DataLoader(ds, batch_size=batch_size, shuffle=shuffle,
                      num_workers=num_workers, pin_memory=pin_memory, drop_last=False)

skin_dict = align_paired_obs_names(
    {"rna": skin_rna_input, "atac": skin_atac_input})

skin_ds = MultiModalDataset(adata_dict=skin_dict, X_key="X", device=device)

train_idx = idx_from_split(skin_rna_input, "train")
val_idx   = idx_from_split(skin_rna_input, "val")
test_idx  = idx_from_split(skin_rna_input, "test")

train_loader = make_subset_loader(skin_ds, train_idx, batch_size=BATCH_SIZE, shuffle=True)
val_loader   = make_subset_loader(skin_ds, val_idx,   batch_size=BATCH_SIZE, shuffle=False)
test_loader  = make_subset_loader(skin_ds, test_idx,  batch_size=BATCH_SIZE, shuffle=False)

print(f"Train: {len(train_idx)} | Val: {len(val_idx)} | Test: {len(test_idx)}")


# %% [markdown]
### Build test AnnData objects for evaluation

Separate test-set RNA and ATAC AnnData objects are needed by `univi.evaluation` — same pattern as the Multiome PBMC analysis.

# %% [code]
rna_test_adata  = skin_rna_input[test_idx].copy()
atac_test_adata = skin_atac_input[test_idx].copy()

# Carry the raw log1p (all genes) for reconstruction evaluation
rna_test_adata_full = skin_rna_input[test_idx].copy()   # has .layers['log1p']

print("Test RNA :", rna_test_adata.shape)
print("Test ATAC:", atac_test_adata.shape)


# %% [markdown]
### Instantiate UniVI model (bimodal RNA–ATAC)

# %% [code]
# ── UniVI config — tuned for SHARE-seq skin (not just copied from PBMC) ─────

univi_cfg = UniVIConfig(
    latent_dim=30,
    beta=1.25,
    gamma=6.35,
    encoder_dropout=0.10,
    decoder_dropout=0.00,
    encoder_batchnorm=True,
    decoder_batchnorm=False,
    kl_anneal_start=50,
    kl_anneal_end=85,
    align_anneal_start=75,
    align_anneal_end=110,
    modalities=[
        ModalityConfig(
            name="rna",
            input_dim=skin_rna_input.n_vars,      # HVG_TOP_N
            encoder_hidden=[1024, 512, 256, 128],
            decoder_hidden=[128, 256, 512, 1024],
            likelihood="gaussian",
        ),
        ModalityConfig(
            name="atac",
            input_dim=skin_atac_input.n_vars,     # ATAC_N_LSI - 1 (LSI_0 dropped)
            encoder_hidden=[256, 128, 64],
            decoder_hidden=[64, 128, 256],
            likelihood="gaussian",
        ),
    ],
)

train_cfg = TrainingConfig(
    n_epochs=5000,
    batch_size=BATCH_SIZE,
    lr=1e-3,
    weight_decay=1e-4,
    device=device,
    log_every=20,
    grad_clip=5.0,
    num_workers=NUM_WORKERS,
    seed=42,
    early_stopping=True,
    patience=200,
    min_delta=0.0,
    best_epoch_warmup=0,
)

model = UniVIMultiModalVAE(
    univi_cfg,
    loss_mode="v1",
    v1_recon="avg",
    normalize_v1_terms=True,
).to(device)


# %% [markdown]
### Train model

# %% [code]
trainer = UniVITrainer(
    model=model,
    train_loader=train_loader,
    val_loader=val_loader,
    train_cfg=train_cfg,
    device=device,
)

history = trainer.fit()


# %% [code]
# Training curves
fig, axes = plt.subplots(1, 2, figsize=(11, 4))

axes[0].plot(history["train_loss"], label="train")
axes[0].plot(history["val_loss"],   label="val")
axes[0].set_xlabel("Epoch")
axes[0].set_ylabel("Loss")
axes[0].set_title("UniVI SHARE-seq skin: training curves")
axes[0].legend()

axes[1].plot(history["beta"],  label="beta (KL weight)")
axes[1].plot(history["gamma"], label="gamma (align weight)")
axes[1].set_xlabel("Epoch")
axes[1].set_ylabel("Annealing weight")
axes[1].set_title("KL / alignment annealing")
axes[1].legend()

plt.tight_layout()
plt.savefig(FIGDIR / "training_curves.png", dpi=150)
plt.show()
plt.close()


# %% [markdown]
### Save model checkpoint

# %% [code]
import hashlib, dataclasses

beta_used    = str(univi_cfg.beta).replace(".", "-")
gamma_used   = str(univi_cfg.gamma).replace(".", "-")
latent_used  = str(univi_cfg.latent_dim)

from datetime import date
today = date.today().strftime("%m-%d-%Y")

ckpt_name = (f"SHARE-seq_skin_RNA-ATAC_beta-{beta_used}_gamma-{gamma_used}"
             f"_latent-{latent_used}_{today}.pt")
ckpt_path = MODEL_DIR / ckpt_name

ckpt = {
    "model_state_dict": model.state_dict(),
    "univi_cfg":        dataclasses.asdict(univi_cfg),
    "train_cfg":        dataclasses.asdict(train_cfg),
    "history":          history,
    "preprocess": {
        "hvg_genes":    hvg_genes.tolist(),
        "rna_mean":     rna_mean,
        "rna_std":      rna_std,
        "idf":          idf,
        "svd_components_": svd.components_,
        "lsi_mean":     lsi_mean,
        "lsi_std":      lsi_std,
    },
    "split_idx":        splits,
    "n_cells_total":    skin_rna.n_obs,
    "dataset":          "Ma2020_SHARE-seq_skin",
    "geo_accession":    "GSE140203",
}

torch.save(ckpt, ckpt_path)
print(f"Checkpoint saved to: {ckpt_path}")


# %% [markdown]
### Evaluation

All metrics use the held-out test split. Metric definitions and code are identical to those used for the Multiome PBMC analysis to ensure direct comparability.

# %% [code]
# 1. Encode test cells into UniVI latent space
model.eval()

Z_rna  = trainer.encode_modality(rna_test_adata,  modality="rna",  batch_size=1024)
Z_atac = trainer.encode_modality(atac_test_adata, modality="atac", batch_size=1024)

rna_test_adata.obsm["X_univi"]  = Z_rna
atac_test_adata.obsm["X_univi"] = Z_atac

print("Latent shapes (test):")
print("  RNA :", Z_rna.shape)
print("  ATAC:", Z_atac.shape)


# %% [code]
# 2. FOSCTTM (paired correspondence, lower = better)

def foscttm_per_cell(z_src, z_tgt, metric="euclidean"):
    """Per-cell fraction of cells closer than the true match."""
    assert z_src.shape[0] == z_tgt.shape[0]
    dists = pairwise_distances(z_src, z_tgt, metric=metric)
    n = dists.shape[0]
    true_d = dists[np.arange(n), np.arange(n)]
    return (dists < true_d[:, None]).mean(axis=1)

fos_cells = foscttm_per_cell(Z_rna, Z_atac)
fos_mean  = fos_cells.mean()
fos_sem   = fos_cells.std(ddof=1) / np.sqrt(len(fos_cells))

print(f"FOSCTTM (RNA–ATAC): {fos_mean:.4f} ± {fos_sem:.4f} (mean ± SEM, lower is better)")


# %% [code]
# 3. Recall@K (cross-modal nearest-neighbor retrieval)

def topk_matching(z_src, z_tgt, pair_name, ks=(1, 5, 10, 25, 50, 100)):
    k_max = max(ks)
    nn = NearestNeighbors(n_neighbors=k_max, metric="euclidean")
    nn.fit(z_tgt)
    _, idx_knn = nn.kneighbors(z_src)
    true_idx = np.arange(z_src.shape[0])
    n_cells  = z_src.shape[0]
    results = {}
    for k in ks:
        hits = (idx_knn[:, :k] == true_idx[:, None]).any(axis=1)
        m, se = hits.mean(), np.sqrt(hits.mean() * (1 - hits.mean()) / n_cells)
        results[k] = (float(m), float(se))
        print(f"  {pair_name} Recall@{k:3d}: {m:.3f} ± {se:.3f}")
    return results

print("RNA → ATAC:")
recall_rna_atac = topk_matching(Z_rna, Z_atac, "RNA→ATAC")

print("\nATAC → RNA:")
recall_atac_rna = topk_matching(Z_atac, Z_rna, "ATAC→RNA")


# %% [code]
# 4. Modality mixing (stacked embeddings)

Z_joint = np.vstack([Z_rna, Z_atac])
modality_labels = np.array(["rna"] * Z_rna.shape[0] + ["atac"] * Z_atac.shape[0])

k_mix = 30

try:
    mixing_score = univi_eval.compute_modality_mixing(Z_joint, modality_labels, k_mix)
except TypeError:
    mixing_score = univi_eval.compute_modality_mixing(Z_joint, modality_labels)

nn_mix = NearestNeighbors(n_neighbors=k_mix + 1, metric="euclidean")
nn_mix.fit(Z_joint)
_, idx_mix = nn_mix.kneighbors(Z_joint)
neighbor_mods = modality_labels[idx_mix[:, 1:]]
same_mod_frac = (neighbor_mods == modality_labels[:, None]).mean()

sil_mod = silhouette_score(Z_joint, modality_labels, metric="euclidean")

print(f"Modality mixing score (k={k_mix}): {mixing_score:.4f}")
print(f"Same-modality neighbor fraction: {same_mod_frac:.4f}  (lower = better mixing)")
print(f"Silhouette (modality): {sil_mod:.4f}  (negative = good mixing)")


# %% [code]
# 5. UMAP on stacked latent (paired test cells)

rna_tmp  = rna_test_adata.copy()
atac_tmp = atac_test_adata.copy()
rna_tmp.obs["univi_source"]  = "rna"
atac_tmp.obs["univi_source"] = "atac"

combined = rna_tmp.concatenate(
    atac_tmp,
    join="outer",
    batch_key="univi_batch",
    batch_categories=["rna", "atac"],
    index_unique=None,
)
combined.obsm["X_univi"] = np.vstack([Z_rna, Z_atac])

sc.pp.neighbors(combined, use_rep="X_univi", n_neighbors=30)
sc.tl.umap(combined)
sc.tl.leiden(combined, key_added="univi_leiden", resolution=0.65)

# UMAP: colored by modality
sc.pl.umap(combined, color="univi_source", size=10, alpha=0.8, show=False,
           title="SHARE-seq skin — modality")
plt.savefig(FIGDIR / "umap_modality.png", bbox_inches="tight", dpi=200)
plt.show(); plt.close()

# UMAP: colored by cell type
sc.pl.umap(combined, color=CELLTYPE_KEY, size=10, alpha=0.8, show=False,
           title="SHARE-seq skin — cell type")
plt.savefig(FIGDIR / "umap_cell_type.png", bbox_inches="tight", dpi=200)
plt.show(); plt.close()

# UMAP: colored by Leiden
sc.pl.umap(combined, color="univi_leiden", size=10, alpha=0.8, show=False,
           title="SHARE-seq skin — Leiden (UniVI)")
plt.savefig(FIGDIR / "umap_leiden.png", bbox_inches="tight", dpi=200)
plt.show(); plt.close()


# %% [code]
# 6. kNN label transfer (RNA → ATAC and ATAC → RNA)
# Uses training RNA labels to annotate ATAC cells via kNN in the shared latent space.

labels_test = rna_test_adata.obs[CELLTYPE_KEY].values
known_mask  = labels_test != "Unknown"

if known_mask.sum() > 0:
    k_knn = 15
    nn_lt = NearestNeighbors(n_neighbors=k_knn, metric="euclidean")
    nn_lt.fit(Z_rna[known_mask])
    _, idx_lt = nn_lt.kneighbors(Z_atac[known_mask])

    labels_known = labels_test[known_mask]
    pred_labels  = np.array([pd.Series(labels_known[idx_lt[i]]).mode()[0]
                              for i in range(len(idx_lt))])

    acc = accuracy_score(labels_known, pred_labels)
    f1  = f1_score(labels_known, pred_labels, average="macro", zero_division=0)
    print(f"Label transfer ATAC→RNA cell type (kNN k={k_knn}):")
    print(f"  Accuracy: {acc:.3f}")
    print(f"  Macro F1: {f1:.3f}")
else:
    print("No annotated cells available for label transfer (all labels = Unknown).")
    print("Assign published cluster labels to skin_rna.obs[CELLTYPE_KEY] and re-run.")
    acc, f1 = np.nan, np.nan
    

# %% [code]
# 7. Cross-modal reconstruction: RNA → ATAC and ATAC → RNA
# Reports per-feature Pearson r and MSE in the log-normalized space.

def cross_modal_metrics(model, src_adata, tgt_adata, src_mod, tgt_mod, name_prefix):
    Xhat = univi_eval.cross_modal_predict(
        model, adata_src=src_adata, src_mod=src_mod, tgt_mod=tgt_mod,
        device=device, batch_size=512,
    )
    X_tgt   = _to_dense(tgt_adata.X)
    mse_f   = univi_eval.mse_per_feature(X_tgt, Xhat)
    corr_f  = univi_eval.pearson_corr_per_feature(X_tgt, Xhat)

    print(f"Cross-modal reconstruction: {src_mod} → {tgt_mod}")
    print(f"  Mean feature MSE:          {mse_f.mean():.4f}")
    print(f"  Mean feature Pearson r:    {corr_f.mean():.3f}")

    fig, axes = plt.subplots(1, 2, figsize=(9, 3))
    axes[0].hist(corr_f, bins=40)
    axes[0].set_xlabel("Per-feature Pearson r"); axes[0].set_title(f"{src_mod}→{tgt_mod} Pearson r")
    axes[1].hist(mse_f,  bins=40)
    axes[1].set_xlabel("Per-feature MSE");       axes[1].set_title(f"{src_mod}→{tgt_mod} MSE")
    plt.tight_layout()
    plt.savefig(FIGDIR / f"{name_prefix}_recon_metrics.png", dpi=150)
    plt.show(); plt.close()

    return float(mse_f.mean()), float(corr_f.mean())

mse_rna_atac, corr_rna_atac = cross_modal_metrics(
    model, rna_test_adata, atac_test_adata, "rna", "atac", "RNA_to_ATAC"
)
mse_atac_rna, corr_atac_rna = cross_modal_metrics(
    model, atac_test_adata, rna_test_adata, "atac", "rna", "ATAC_to_RNA"
)


# %% [code]
# 8. kNN distance distribution: same-modality vs. cross-modality

nn_dist = NearestNeighbors(n_neighbors=k_mix + 1, metric="euclidean")
nn_dist.fit(Z_joint)
dists_all, idx_all = nn_dist.kneighbors(Z_joint)
dists_neighbors = dists_all[:, 1:]
flat_dists = dists_neighbors.reshape(-1)

center_mods_flat   = np.repeat(modality_labels, k_mix)
neighbor_mods_flat = neighbor_mods.reshape(-1)
same_edge = neighbor_mods_flat == center_mods_flat

plt.figure(figsize=(6, 4))
sns.kdeplot(flat_dists[same_edge],  label="same modality",  fill=True, alpha=0.5)
sns.kdeplot(flat_dists[~same_edge], label="cross modality", fill=True, alpha=0.5)
plt.xlabel("Euclidean distance in UniVI latent")
plt.ylabel("Density")
plt.title(f"kNN distance: same vs. cross modality (k={k_mix})")
plt.legend()
plt.tight_layout()
plt.savefig(FIGDIR / "knn_distance_same_vs_cross.png", dpi=200)
plt.show()
plt.close()


# %% [markdown]
### Summarize and save benchmark metrics

These metrics feed into **Supplemental Fig. S[X]** and the cross-tissue robustness comparison against the Multiome PBMC results.

# %% [code]
skin_metrics = {
    # Dataset
    "dataset":                  "Ma2020_SHARE-seq_skin",
    "geo_accession":            "GSE140203",
    "n_cells_total":            int(skin_rna.n_obs),
    "n_cells_test":             int(rna_test_adata.n_obs),
    "n_genes_rna_hvg":          int(HVG_TOP_N),
    "n_lsi_components_atac":    int(ATAC_N_LSI),
    "n_peaks_after_filter":     int(skin_atac.n_vars),
    "latent_dim":               int(univi_cfg.latent_dim),

    # Correspondence
    "foscttm_rna_atac_mean":    float(fos_mean),
    "foscttm_rna_atac_sem":     float(fos_sem),
    "recall_at_1_rna_atac":     recall_rna_atac[1][0],
    "recall_at_5_rna_atac":     recall_rna_atac[5][0],
    "recall_at_10_rna_atac":    recall_rna_atac[10][0],
    "recall_at_1_atac_rna":     recall_atac_rna[1][0],
    "recall_at_5_atac_rna":     recall_atac_rna[5][0],
    "recall_at_10_atac_rna":    recall_atac_rna[10][0],

    # Modality mixing
    f"modality_mixing_k{k_mix}": float(mixing_score),
    "same_mod_neighbor_frac":   float(same_mod_frac),
    "silhouette_modality":      float(sil_mod),

    # Label transfer
    "label_transfer_accuracy":  float(acc) if not np.isnan(acc) else None,
    "label_transfer_macro_f1":  float(f1)  if not np.isnan(f1)  else None,

    # Cross-modal reconstruction
    "recon_rna_to_atac_mean_pearson": float(corr_rna_atac),
    "recon_rna_to_atac_mean_mse":     float(mse_rna_atac),
    "recon_atac_to_rna_mean_pearson": float(corr_atac_rna),
    "recon_atac_to_rna_mean_mse":     float(mse_atac_rna),
}

metrics_path = FIGDIR / "skin_univi_metrics.json"
with open(metrics_path, "w") as f:
    json.dump(skin_metrics, f, indent=2)

print(f"Benchmark metrics saved to: {metrics_path}")
print("\n=== SHARE-seq Skin Benchmark Summary ===")
for k, v in skin_metrics.items():
    print(f"  {k}: {v}")
    

# %% [code]


# %% [code]


# %% [code]


# %% [code]


# %% [code]


# %% [code]


# %% [code]


# %% [code]


# %% [code]


# %% [code]


# %% [code]


# %% [code]


# %% [code]


# %% [code]


# %% [code]


# %% [code]


# %% [code]


# %% [code]


# %% [code]


# %% [code]


# %% [code]


# %% [code]


# %% [code]


# %% [code]
````
