## Supplemental Code for "Unifying multimodal single-cell data with a mixture-of-experts *β*-variational autoencoder framework": Supp_scNMT_mouse_gastrulation.ipynb

```
# %% [code]
# Import non-UniVI modules
from __future__ import annotations

import os
import sys
from pathlib import Path

import numpy as np
import pandas as pd
import scanpy as sc
import anndata as ad
import re

import torch
from torch.utils.data import DataLoader, Subset

import scipy.sparse as sp

import io
import gc
import tarfile
import gzip
import warnings

from pathlib import Path
from collections import defaultdict, Counter
from typing import Dict, List, Optional, Tuple, Any

import numpy as np
import pandas as pd
import scipy.sparse as sp

from tqdm.auto import tqdm


# %% [code]
from sklearn.feature_extraction.text import TfidfTransformer
from sklearn.decomposition import TruncatedSVD
from sklearn.preprocessing import StandardScaler, normalize

from sklearn.neighbors import NearestNeighbors
from sklearn.metrics import confusion_matrix, accuracy_score, f1_score

import matplotlib.pyplot as plt


# %% [code]
# Import required UniVI modules
from univi import (
    ModalityConfig,
    UniVIConfig,
    TrainingConfig,
    UniVIMultiModalVAE,
    matching,
    UniVITrainer,
    write_univi_latent,
    MultiModalDataset,
)

import univi as uv
import univi.evaluation as ue
import univi.plotting as up

# Double check UniVI module version
print("Installed version is univi v" + str(uv.__version__))


# %% [code]
print("torch:", torch.__version__)
print("torch.cuda.is_available():", torch.cuda.is_available())
device = "cuda" if torch.cuda.is_available() else ("mps" if torch.mps.is_available() else "cpu")
print("Using device:", device)


# %% [code]
# =========================================================
# 1) Paths + manual member picks (recommended)
# =========================================================
DATA_DIR = Path("./data/scNMT-seq_data")  # edit if needed
DATA_DIR.mkdir(parents=True, exist_ok=True)

RAW_TAR_PATH = DATA_DIR / "scnmt_gastrulation.tar.gz"  # parsed EBI bundle
assert RAW_TAR_PATH.exists(), f"Missing file: {RAW_TAR_PATH}"

# Recommended first-pass members from parsed gastrulation tar
RNA_MEMBER = "rna/counts.txt.gz"
CPG_MEMBER = "met/feature_level/genebody.tsv.gz"
GPC_MEMBER = "acc/feature_level/genebody.tsv.gz"
METADATA_MEMBER = "sample_metadata.txt"

print("Using:")
print(" ", RNA_MEMBER)
print(" ", CPG_MEMBER)
print(" ", GPC_MEMBER)
print(" ", METADATA_MEMBER)


# %% [code]
# =========================================================
# 2) Tar helpers + preview utilities
# =========================================================
def _open_tar_member_stream(tar: tarfile.TarFile, member_name: str):
    """
    Return a readable file-like stream for a tar member.
    Handles .gz members transparently.
    """
    fobj = tar.extractfile(member_name)
    if fobj is None:
        raise FileNotFoundError(f"Member not found in tar: {member_name}")
    if member_name.endswith(".gz"):
        return gzip.GzipFile(fileobj=fobj)
    return fobj


def list_tar_members(tar_path: Path, n_preview: int = 80) -> pd.DataFrame:
    with tarfile.open(tar_path, "r:*") as tar:
        members = [m for m in tar.getmembers() if m.isfile()]

    names = [m.name for m in members]
    manifest = pd.DataFrame({
        "member_name": names,
        "size_bytes": [m.size for m in members],
    })

    print(f"Manifest files: {len(manifest)}")
    print("Preview:")
    for n in manifest["member_name"].head(n_preview):
        print(" ", n)

    return manifest


def preview_tar_member_df(tar_path: Path, member_name: str, nrows: int = 8, header="infer"):
    with tarfile.open(tar_path, "r:*") as tar:
        with _open_tar_member_stream(tar, member_name) as fh:
            df = pd.read_csv(fh, sep="\t", nrows=nrows, header=header)
    return df


def preview_tar_members(tar_path: Path, member_names: List[str], nrows: int = 8):
    previews = {}
    for name in member_names:
        try:
            header = None if ("feature_level/" in name and name.endswith(".tsv.gz")) else "infer"
            df = preview_tar_member_df(tar_path, name, nrows=nrows, header=header)
            previews[name] = {"ok": True, "df": df}
        except Exception as e:
            previews[name] = {"ok": False, "error": repr(e)}
    return previews


def display_previews(previews: Dict[str, Dict[str, Any]], max_rows: int = 8):
    for name, info in previews.items():
        print("\n" + "=" * 100)
        print(name)
        if not info["ok"]:
            print("ERROR:", info["error"])
            continue
        df = info["df"]
        print("shape:", df.shape)
        display(df.head(max_rows))
        

# %% [code]
# =========================================================
# 3) Optional sanity preview (recommended - might take a sec due to large file size)
# =========================================================
#to_preview = [RNA_MEMBER, CPG_MEMBER, GPC_MEMBER, METADATA_MEMBER]
#previews = preview_tar_members(RAW_TAR_PATH, to_preview, nrows=8)
#display_previews(previews)


# %% [code]
# =========================================================
# 4) RNA parser (wide matrix: genes x cells -> cells x genes)
# =========================================================
def read_rna_counts_from_tar(
    tar_path: Path,
    member_name: str,
    verbose: bool = True,
) -> pd.DataFrame:
    """
    Reads rna/counts.txt.gz from the scnmt_gastrulation parsed tar.
    Returns DataFrame with shape (cells x genes), index=id_rna strings, columns=ens_id.
    """
    if verbose:
        print(f"\nReading RNA matrix: {member_name}")

    with tarfile.open(tar_path, "r:*") as tar:
        with _open_tar_member_stream(tar, member_name) as fh:
            df = pd.read_csv(fh, sep="\t")

    if verbose:
        print("Raw shape:", df.shape)
        display(df.head())

    # First column is gene ID
    gene_col = df.columns[0]
    if str(gene_col).lower() not in {"ens_id", "gene", "gene_id", "feature"}:
        warnings.warn(f"Unexpected first RNA column: {gene_col}")

    df = df.set_index(gene_col)

    # rows=genes, cols=cells -> transpose to cells x genes
    df = df.T
    df.index = df.index.astype(str)
    df.columns = df.columns.astype(str)

    if verbose:
        print(f"Assumed rows=features, cols=cells -> transposed to cells x features: {df.shape}")

    return df
    

# %% [code]
# =========================================================
# 5) Long-table parser for met/acc feature_level genebody files
# =========================================================
def read_feature_level_long_from_tar(
    tar_path: Path,
    member_name: str,
    modality: str,                     # "cpg" or "gpc"
    feature_subset: Optional[List[str]] = None,
    cell_subset: Optional[List[str]] = None,
    min_coverage: int = 1,
    chunksize: int = 500_000,
    verbose: bool = True,
) -> Dict[str, Any]:
    """
    Read long-format feature-level table from scnmt_gastrulation parsed tar.

    Expected columns (headerless):
      0: cell_id
      1: feature_id
      2: feature_class (e.g. genebody)
      3: successes
      4: coverage
      5: percent (0-100)

    Returns dict with:
      - frac_df: cells x features (fractions)
      - successes_df: cells x features
      - coverage_df: cells x features
      - meta: summary
    """
    if verbose:
        print(f"\nReading feature-level long table: {member_name} ({modality})")

    feature_subset_set = set(map(str, feature_subset)) if feature_subset is not None else None
    cell_subset_set = set(map(str, cell_subset)) if cell_subset is not None else None

    chunk_results = []
    total_rows = 0
    kept_rows = 0

    with tarfile.open(tar_path, "r:*") as tar:
        with _open_tar_member_stream(tar, member_name) as fh:
            reader = pd.read_csv(
                fh,
                sep="\t",
                header=None,
                names=["cell_id", "feature_id", "feature_class", "successes", "coverage", "percent"],
                chunksize=chunksize,
                dtype={
                    "cell_id": "string",
                    "feature_id": "string",
                    "feature_class": "string",
                    "successes": "float32",
                    "coverage": "float32",
                    "percent": "float32",
                },
            )

            for i, chunk in enumerate(reader, start=1):
                total_rows += len(chunk)

                # basic cleaning
                chunk = chunk.dropna(subset=["cell_id", "feature_id", "successes", "coverage"])
                chunk["cell_id"] = chunk["cell_id"].astype(str)
                chunk["feature_id"] = chunk["feature_id"].astype(str)

                # filter by coverage
                chunk = chunk[chunk["coverage"] >= min_coverage]

                # optional subsets
                if cell_subset_set is not None:
                    chunk = chunk[chunk["cell_id"].isin(cell_subset_set)]
                if feature_subset_set is not None:
                    chunk = chunk[chunk["feature_id"].isin(feature_subset_set)]

                if len(chunk) == 0:
                    continue

                # fraction = successes / coverage
                cov = chunk["coverage"].to_numpy(dtype=np.float32)
                suc = chunk["successes"].to_numpy(dtype=np.float32)
                frac = np.divide(
                    suc,
                    cov,
                    out=np.zeros(len(chunk), dtype=np.float32),
                    where=(cov > 0),
                )

                chunk = chunk.copy()
                chunk["fraction"] = frac

                chunk_results.append(chunk[["cell_id", "feature_id", "successes", "coverage", "fraction"]])
                kept_rows += len(chunk)

                if verbose and (i <= 3 or i % 25 == 0):
                    print(f"  chunk {i}: input_rows={len(chunk):,}, retained_total={kept_rows:,}")

    if len(chunk_results) == 0:
        raise ValueError(f"No rows retained from {member_name}. Check subsets/min_coverage.")

    long_df = pd.concat(chunk_results, axis=0, ignore_index=True)

    if verbose:
        print(f"Total rows read: {total_rows:,}")
        print(f"Rows retained:   {kept_rows:,}")
        print("Retained long_df shape:", long_df.shape)
        display(long_df.head())

    # Aggregate duplicates if present
    dup_mask = long_df.duplicated(subset=["cell_id", "feature_id"], keep=False)
    if dup_mask.any():
        if verbose:
            print(f"Found duplicated (cell_id, feature_id) rows -> aggregating")
        long_df = (
            long_df.groupby(["cell_id", "feature_id"], as_index=False)[["successes", "coverage"]]
            .sum()
        )
        cov = long_df["coverage"].to_numpy(dtype=np.float32)
        suc = long_df["successes"].to_numpy(dtype=np.float32)
        long_df["fraction"] = np.divide(suc, cov, out=np.zeros(len(long_df), dtype=np.float32), where=(cov > 0))

    # Pivot wide: cells x features
    successes_df = long_df.pivot(index="cell_id", columns="feature_id", values="successes")
    coverage_df  = long_df.pivot(index="cell_id", columns="feature_id", values="coverage")
    frac_df      = long_df.pivot(index="cell_id", columns="feature_id", values="fraction")

    # Align and fill
    cells = successes_df.index.astype(str)
    feats = successes_df.columns.astype(str)

    successes_df = successes_df.reindex(index=cells, columns=feats).fillna(0).astype(np.float32)
    coverage_df  = coverage_df.reindex(index=cells, columns=feats).fillna(0).astype(np.float32)
    frac_df      = frac_df.reindex(index=cells, columns=feats).fillna(0).astype(np.float32)

    if verbose:
        print(f"Wide matrices built (cells x features): {frac_df.shape}")
        nnz_cov = int((coverage_df.to_numpy() > 0).sum())
        print(f"Coverage nonzero entries: {nnz_cov:,}")

    return {
        "frac_df": frac_df,
        "successes_df": successes_df,
        "coverage_df": coverage_df,
        "meta": {
            "member_name": member_name,
            "modality": modality,
            "n_cells": int(frac_df.shape[0]),
            "n_features": int(frac_df.shape[1]),
            "total_rows_read": int(total_rows),
            "rows_retained": int(kept_rows),
            "min_coverage": int(min_coverage),
        },
    }
    

# %% [code]
# =========================================================
# 6) Metadata reader
# =========================================================
def read_metadata_from_tar(
    tar_path: Path,
    member_name: str = "sample_metadata.txt",
    verbose: bool = True,
) -> pd.DataFrame:
    with tarfile.open(tar_path, "r:*") as tar:
        with _open_tar_member_stream(tar, member_name) as fh:
            meta = pd.read_csv(fh, sep="\t")

    # normalize common string columns
    for c in ["sample", "id_rna", "id_met", "id_acc", "embryo", "plate", "stage", "lineage10x", "lineage10x_2"]:
        if c in meta.columns:
            meta[c] = meta[c].astype("string")

    if verbose:
        print("Metadata shape:", meta.shape)
        print("Metadata columns:", list(meta.columns))
        display(meta.head())

    return meta
    

# %% [code]
# =========================================================
# 7) AnnData builders
# =========================================================
def _df_to_anndata_dense_or_sparse(
    df: pd.DataFrame,
    modality: str,
    feature_type: str,
    copy_to_counts: bool = False,
) -> ad.AnnData:
    x = df.copy()
    x.index = x.index.astype(str)
    x.columns = x.columns.astype(str)
    x = x.fillna(0.0)

    arr = x.to_numpy(dtype=np.float32)
    zero_frac = float((arr == 0).mean()) if arr.size else 0.0
    X = sp.csr_matrix(arr) if zero_frac > 0.5 else arr

    a = ad.AnnData(X=X)
    a.obs_names = pd.Index(x.index.astype(str), name="cell_id")
    a.var_names = pd.Index(x.columns.astype(str), name="feature")

    a.obs["modality"] = modality
    a.var["feature_type"] = feature_type

    if copy_to_counts:
        a.layers["counts"] = a.X.copy()

    return a


def build_methyl_like_anndata_from_long_feature(
    parsed: Dict[str, Any],
    modality: str,               # "cpg" or "gpc"
    successes_layer: str,
    coverage_layer: str,
    feature_type: str = "genebody",
) -> ad.AnnData:
    frac_df = parsed["frac_df"]
    suc_df = parsed["successes_df"]
    cov_df = parsed["coverage_df"]

    # ensure same order
    cells = frac_df.index.astype(str)
    feats = frac_df.columns.astype(str)
    suc_df = suc_df.reindex(index=cells, columns=feats).fillna(0).astype(np.float32)
    cov_df = cov_df.reindex(index=cells, columns=feats).fillna(0).astype(np.float32)

    a = _df_to_anndata_dense_or_sparse(
        frac_df,
        modality=modality,
        feature_type=feature_type,
        copy_to_counts=False,
    )

    suc_arr = suc_df.to_numpy(dtype=np.float32)
    cov_arr = cov_df.to_numpy(dtype=np.float32)

    a.layers[successes_layer] = sp.csr_matrix(suc_arr) if (suc_arr == 0).mean() > 0.5 else suc_arr
    a.layers[coverage_layer]  = sp.csr_matrix(cov_arr) if (cov_arr == 0).mean() > 0.5 else cov_arr

    # QC summaries from coverage
    cov_mat = a.layers[coverage_layer]
    if sp.issparse(cov_mat):
        cov_sum_cell = np.asarray(cov_mat.sum(axis=1)).ravel()
        cov_nnz_cell = np.asarray((cov_mat > 0).sum(axis=1)).ravel()
        cov_sum_feat = np.asarray(cov_mat.sum(axis=0)).ravel()
    else:
        cov_sum_cell = cov_mat.sum(axis=1)
        cov_nnz_cell = (cov_mat > 0).sum(axis=1)
        cov_sum_feat = cov_mat.sum(axis=0)

    a.obs["coverage_total"] = cov_sum_cell.astype(np.float32)
    a.obs["n_features_covered"] = cov_nnz_cell.astype(np.int32)
    a.var["coverage_total"] = cov_sum_feat.astype(np.float32)

    return a
    

# %% [code]
# =========================================================
# 8) Attach metadata + align paired cells
# =========================================================
def attach_metadata_and_align_modalities(
    rna: Optional[ad.AnnData],
    cpg: Optional[ad.AnnData],
    gpc: Optional[ad.AnnData],
    meta: pd.DataFrame,
    require_qc: bool = True,
    verbose: bool = True,
) -> Dict[str, ad.AnnData]:
    """
    - RNA obs_names are id_rna strings -> map to metadata['sample'] via metadata['id_rna']
    - cpg/gpc obs_names are sample IDs already (for parsed genebody files)
    - attach metadata to each modality
    - intersect common cells across available modalities
    """
    meta = meta.copy()

    if "sample" not in meta.columns:
        raise ValueError("metadata missing 'sample' column")

    # joint QC filter if requested
    if require_qc:
        mask = pd.Series(True, index=meta.index)
        for qc_col in ["pass_rnaQC", "pass_metQC", "pass_accQC"]:
            if qc_col in meta.columns:
                mask &= meta[qc_col].fillna(False).astype(bool)
        meta_qc = meta.loc[mask].copy()
        if verbose:
            print(f"QC filter retained {meta_qc.shape[0]} / {meta.shape[0]} metadata rows")
    else:
        meta_qc = meta

    meta_by_sample = meta_qc.drop_duplicates(subset=["sample"]).set_index("sample")
    meta_by_sample.index = meta_by_sample.index.astype(str)

    # RNA: map id_rna -> sample
    if rna is not None:
        if "id_rna" not in meta_qc.columns:
            raise ValueError("metadata missing 'id_rna' for RNA mapping")

        rna_map = (
            meta_qc[["sample", "id_rna"]]
            .dropna(subset=["sample", "id_rna"])
            .drop_duplicates(subset=["id_rna"])
            .set_index("id_rna")["sample"]
            .astype(str)
        )
        rna_map.index = rna_map.index.astype(str)

        keep_rna = [ix for ix in rna.obs_names.astype(str) if ix in rna_map.index]
        rna = rna[keep_rna].copy()

        rna.obs["sample"] = rna_map.loc[rna.obs_names.astype(str)].to_numpy().astype(str)
        rna.obs_names = pd.Index(rna.obs["sample"].astype(str), name="cell_id")

        common = rna.obs_names.intersection(meta_by_sample.index)
        rna = rna[common].copy()
        rna.obs = rna.obs.join(meta_by_sample, how="left")

    def _attach_sample_metadata(a: Optional[ad.AnnData]) -> Optional[ad.AnnData]:
        if a is None:
            return None
        a = a.copy()
        a.obs_names = pd.Index(a.obs_names.astype(str), name="cell_id")
        a.obs["sample"] = a.obs_names.astype(str)
        common = a.obs_names.intersection(meta_by_sample.index)
        a = a[common].copy()
        a.obs = a.obs.join(meta_by_sample, how="left")
        return a

    cpg = _attach_sample_metadata(cpg)
    gpc = _attach_sample_metadata(gpc)

    available = {k: v for k, v in {"rna": rna, "cpg": cpg, "gpc": gpc}.items() if v is not None}
    if len(available) == 0:
        raise ValueError("No modalities available after parsing and metadata attachment.")

    common_cells = None
    for k, a in available.items():
        idx = pd.Index(a.obs_names.astype(str))
        common_cells = idx if common_cells is None else common_cells.intersection(idx)

    common_cells = pd.Index(common_cells.astype(str))

    if verbose:
        print(f"Common paired cells across {list(available.keys())}: {len(common_cells)}")

    out = {}
    for k, a in available.items():
        out[k] = a[common_cells].copy()
        if verbose:
            print(k, out[k])

    return out
    

# %% [code]
# =========================================================
# 9) Optional feature filtering helpers (recommended)
# =========================================================
def filter_rna_genes_basic(
    rna: ad.AnnData,
    min_cells: int = 10,
    min_counts: int = 20,
    verbose: bool = True,
) -> ad.AnnData:
    r = rna.copy()

    X = r.layers["counts"] if "counts" in r.layers else r.X
    if sp.issparse(X):
        n_cells_by_gene = np.asarray((X > 0).sum(axis=0)).ravel()
        total_counts_by_gene = np.asarray(X.sum(axis=0)).ravel()
    else:
        n_cells_by_gene = (X > 0).sum(axis=0)
        total_counts_by_gene = X.sum(axis=0)

    keep = (n_cells_by_gene >= min_cells) & (total_counts_by_gene >= min_counts)
    if verbose:
        print(f"RNA features before: {r.n_vars}")
        print(f"RNA features kept:   {int(keep.sum())}")

    r = r[:, keep].copy()
    return r


def filter_methyl_features_by_coverage(
    a: ad.AnnData,
    coverage_layer: str,
    min_cells_covered: int = 10,
    min_total_coverage: float = 50.0,
    verbose: bool = True,
) -> ad.AnnData:
    b = a.copy()
    cov = b.layers[coverage_layer]

    if sp.issparse(cov):
        cells_covered = np.asarray((cov > 0).sum(axis=0)).ravel()
        total_cov = np.asarray(cov.sum(axis=0)).ravel()
    else:
        cells_covered = (cov > 0).sum(axis=0)
        total_cov = cov.sum(axis=0)

    keep = (cells_covered >= min_cells_covered) & (total_cov >= min_total_coverage)
    if verbose:
        print(f"{b.obs['modality'].iloc[0]} features before: {b.n_vars}")
        print(f"{b.obs['modality'].iloc[0]} features kept:   {int(keep.sum())}")

    b = b[:, keep].copy()
    return b
    

# %% [code]
# =========================================================
# 10) End-to-end loader for parsed gastrulation genebody triplet
# =========================================================
def load_scnmt_gastrulation_genebody_triplet(
    tar_path: Path,
    rna_member: str = "rna/counts.txt.gz",
    cpg_member: str = "met/feature_level/genebody.tsv.gz",
    gpc_member: str = "acc/feature_level/genebody.tsv.gz",
    metadata_member: str = "sample_metadata.txt",
    require_qc: bool = True,
    min_cov_cpg: int = 1,
    min_cov_gpc: int = 1,
    filter_features: bool = True,
    rna_min_cells: int = 10,
    rna_min_counts: int = 20,
    meth_min_cells_covered: int = 10,
    meth_min_total_coverage: float = 50.0,
    verbose: bool = True,
) -> Dict[str, Any]:
    # metadata first (lets us subset long tables to relevant cells)
    meta = read_metadata_from_tar(tar_path, metadata_member, verbose=verbose)

    meta_work = meta.copy()
    if require_qc:
        mask = pd.Series(True, index=meta_work.index)
        for qc_col in ["pass_rnaQC", "pass_metQC", "pass_accQC"]:
            if qc_col in meta_work.columns:
                mask &= meta_work[qc_col].fillna(False).astype(bool)
        meta_work = meta_work.loc[mask].copy()
        if verbose:
            print(f"Metadata rows after joint QC filter: {meta_work.shape[0]}")

    rna_keep_ids = meta_work["id_rna"].dropna().astype(str).unique().tolist() if "id_rna" in meta_work.columns else None
    sample_keep_ids = meta_work["sample"].dropna().astype(str).unique().tolist()

    # RNA
    rna_df = read_rna_counts_from_tar(tar_path, rna_member, verbose=verbose)
    if rna_keep_ids is not None:
        rna_df = rna_df.loc[rna_df.index.intersection(pd.Index(rna_keep_ids))].copy()
        if verbose:
            print("RNA after metadata/QC subset:", rna_df.shape)

    rna = _df_to_anndata_dense_or_sparse(
        rna_df,
        modality="rna",
        feature_type="gene",
        copy_to_counts=True,
    )

    # CpG (long -> wide)
    cpg_parsed = read_feature_level_long_from_tar(
        tar_path=tar_path,
        member_name=cpg_member,
        modality="cpg",
        cell_subset=sample_keep_ids,
        min_coverage=min_cov_cpg,
        chunksize=500_000,
        verbose=verbose,
    )
    cpg = build_methyl_like_anndata_from_long_feature(
        cpg_parsed,
        modality="cpg",
        successes_layer="meth_successes",
        coverage_layer="meth_total_count",
        feature_type="genebody",
    )

    # GpC (long -> wide)
    gpc_parsed = read_feature_level_long_from_tar(
        tar_path=tar_path,
        member_name=gpc_member,
        modality="gpc",
        cell_subset=sample_keep_ids,
        min_coverage=min_cov_gpc,
        chunksize=500_000,
        verbose=verbose,
    )
    gpc = build_methyl_like_anndata_from_long_feature(
        gpc_parsed,
        modality="gpc",
        successes_layer="acc_successes",
        coverage_layer="acc_total_count",
        feature_type="genebody",
    )

    # Attach metadata + intersect common paired cells
    aligned = attach_metadata_and_align_modalities(
        rna=rna,
        cpg=cpg,
        gpc=gpc,
        meta=meta,
        require_qc=require_qc,
        verbose=verbose,
    )
    rna = aligned["rna"]
    cpg = aligned["cpg"]
    gpc = aligned["gpc"]

    # Optional feature filtering (highly recommended for speed)
    if filter_features:
        if verbose:
            print("\nApplying feature filtering...")

        rna = filter_rna_genes_basic(
            rna, min_cells=rna_min_cells, min_counts=rna_min_counts, verbose=verbose
        )
        cpg = filter_methyl_features_by_coverage(
            cpg,
            coverage_layer="meth_total_count",
            min_cells_covered=meth_min_cells_covered,
            min_total_coverage=meth_min_total_coverage,
            verbose=verbose,
        )
        gpc = filter_methyl_features_by_coverage(
            gpc,
            coverage_layer="acc_total_count",
            min_cells_covered=meth_min_cells_covered,
            min_total_coverage=meth_min_total_coverage,
            verbose=verbose,
        )

        # Re-intersect cells just in case (vars changed only, but safe)
        common_cells = rna.obs_names.intersection(cpg.obs_names).intersection(gpc.obs_names)
        rna = rna[common_cells].copy()
        cpg = cpg[common_cells].copy()
        gpc = gpc[common_cells].copy()

    return {
        "rna": rna,
        "cpg": cpg,
        "gpc": gpc,
        "meta": meta,
        "cpg_parse_meta": cpg_parsed["meta"],
        "gpc_parse_meta": gpc_parsed["meta"],
    }
    

# %% [code]
# =========================================================
# 11) Load parsed gastrulation triplet into AnnData
# =========================================================
adapter_out = load_scnmt_gastrulation_genebody_triplet(
    tar_path=RAW_TAR_PATH,
    rna_member=RNA_MEMBER,
    cpg_member=CPG_MEMBER,
    gpc_member=GPC_MEMBER,
    metadata_member=METADATA_MEMBER,

    require_qc=False,
    min_cov_cpg=1,
    min_cov_gpc=1,

    filter_features=False,
    rna_min_cells=10,
    rna_min_counts=20,
    meth_min_cells_covered=10,
    meth_min_total_coverage=50.0,

    verbose=True,
)

rna = adapter_out["rna"]
cpg = adapter_out["cpg"]
gpc = adapter_out["gpc"]
meta = adapter_out["meta"]

print("\nFinal aligned objects:")
print("rna:", rna)
print("cpg:", cpg)
print("gpc:", gpc)

print("\nobs columns (rna):", list(rna.obs.columns))
display(rna.obs.head())
display(cpg.obs.head())
display(gpc.obs.head())


# %% [code]
# =========================================================
# 12) Build UniVI inputs from adapter output
# =========================================================
def build_univi_inputs_from_scnmt_triplet(
    rna: ad.AnnData,
    cpg: ad.AnnData,
    gpc: ad.AnnData,
    rna_target_sum: float = 1e4,
    log1p_rna: bool = True,
    copy: bool = True,
) -> Dict[str, Any]:
    """
    Returns:
      - adata_dict (paired, same obs_names expected after alignment)
      - recon_targets_spec (for beta_binomial on cpg/gpc)
    """
    r = rna.copy() if copy else rna
    m = cpg.copy() if copy else cpg
    a = gpc.copy() if copy else gpc

    # RNA preprocessing for model input in .X
    # Keep raw counts in .layers["counts"]
    sc.pp.normalize_total(r, target_sum=rna_target_sum)
    if log1p_rna:
        sc.pp.log1p(r)

    # cpg/gpc already have fractions in .X and successes/coverage in layers
    adata_dict = {"rna": r, "cpg": m, "gpc": a}

    recon_targets_spec = {
        "cpg": {
            "successes_layer": "meth_successes",
            "total_count_layer": "meth_total_count",
        },
        "gpc": {
            "successes_layer": "acc_successes",
            "total_count_layer": "acc_total_count",
        },
    }

    return {
        "adata_dict": adata_dict,
        "recon_targets_spec": recon_targets_spec,
    }


u = build_univi_inputs_from_scnmt_triplet(rna, cpg, gpc, copy=True)

adata_dict = u["adata_dict"]
recon_targets_spec = u["recon_targets_spec"]

print("adata_dict keys:", list(adata_dict.keys()))
for k, a in adata_dict.items():
    print(k, a)
print("recon_targets_spec:", recon_targets_spec)


# %% [code]
# =========================================================
# 13) UniVI dataset + train/val/test splits + loaders
# =========================================================
from univi.data import MultiModalDataset, align_paired_obs_names, collate_multimodal_xy_recon

# Ensure exact obs alignment/order
adata_dict = align_paired_obs_names(adata_dict)

n = next(iter(adata_dict.values())).n_obs
idx = np.arange(n)
rng = np.random.default_rng(0)
rng.shuffle(idx)

n_train = int(0.85 * n)
n_val = int(0.05 * n)

train_idx = idx[:n_train]
val_idx = idx[n_train:n_train + n_val]
test_idx = idx[n_train + n_val:]

np.savez(
    DATA_DIR / "splits_scnmt_gastrulation_genebody_seed0.npz",
    train_idx=train_idx,
    val_idx=val_idx,
    test_idx=test_idx,
)

dataset = MultiModalDataset(
    adata_dict=adata_dict,
    X_key="X",
    device=None,  # CPU tensors; trainer moves to device
    recon_targets_spec=recon_targets_spec,
)

train_loader = DataLoader(
    Subset(dataset, train_idx),
    batch_size=24,
    shuffle=True,
    num_workers=0,
    collate_fn=collate_multimodal_xy_recon,
)

val_loader = DataLoader(
    Subset(dataset, val_idx),
    batch_size=24,
    shuffle=False,
    num_workers=0,
    collate_fn=collate_multimodal_xy_recon,
)

test_loader = DataLoader(
    Subset(dataset, test_idx),
    batch_size=24,
    shuffle=False,
    num_workers=0,
    collate_fn=collate_multimodal_xy_recon,
)

print("n cells:", n)
print("train/val/test:", len(train_idx), len(val_idx), len(test_idx))


# %% [code]
# =========================================================
# 14) Batch debug (tuple/dict-safe)
# =========================================================
batch = next(iter(train_loader))
print("batch type:", type(batch))

if isinstance(batch, tuple):
    print("tuple len:", len(batch))
    for i, item in enumerate(batch):
        print(f"[{i}] type={type(item)}")
        if isinstance(item, dict):
            print("    keys:", list(item.keys()))
            for k, v in item.items():
                print("   ", k, type(v), getattr(v, "shape", None))
elif isinstance(batch, dict):
    print("dict keys:", list(batch.keys()))
    for k, v in batch.items():
        print(k, type(v), getattr(v, "shape", None))
        

# %% [code]
# =========================================================
# 15) Define and train UniVI (RNA + CpG + GpC)
# =========================================================
from univi import UniVIMultiModalVAE, ModalityConfig, UniVIConfig, TrainingConfig
from univi.trainer import UniVITrainer

device = "cuda" if torch.cuda.is_available() else ("mps" if torch.backends.mps.is_available() else "cpu")
print("device:", device)

univi_cfg = UniVIConfig(
    latent_dim=20,
    beta=1.35,
    gamma=6.25,
    encoder_dropout=0.010,
    decoder_dropout=0.005,
    encoder_batchnorm=True,
    decoder_batchnorm=False,
    kl_anneal_start=30,
    kl_anneal_end=60,
    align_anneal_start=40,
    align_anneal_end=70,
    modalities=[
        ModalityConfig(
            name="rna",
            input_dim=adata_dict["rna"].n_vars,
            encoder_hidden=[512, 256, 128, 64],
            decoder_hidden=[64, 128, 256, 512],
            likelihood="gaussian",      # because RNA .X is normalized/log1p
        ),
        ModalityConfig(
            name="cpg",
            input_dim=adata_dict["cpg"].n_vars,
            encoder_hidden=[256, 128, 64],
            decoder_hidden=[64, 128, 256],
            likelihood="beta_binomial", # uses meth_successes + meth_total_count
        ),
        ModalityConfig(
            name="gpc",
            input_dim=adata_dict["gpc"].n_vars,
            encoder_hidden=[256, 128, 64],
            decoder_hidden=[64, 128, 256],
            likelihood="beta_binomial", # uses acc_successes + acc_total_count
        ),
    ],
)

train_cfg = TrainingConfig(
    n_epochs=5000,
    batch_size=16,
    lr=1e-3,
    weight_decay=1e-4,
    device=device,
    early_stopping=True,
    best_epoch_warmup=30,
    patience=300,
    log_every=20,
)

model = UniVIMultiModalVAE(
    univi_cfg,
    loss_mode="v1",
    v1_recon="avg",
    normalize_v1_terms=True,
).to(device)


# %% [code]
trainer = UniVITrainer(
    model=model,
    train_loader=train_loader,
    val_loader=val_loader,
    train_cfg=train_cfg,
    device=device,
)

history = trainer.fit()

print("Best epoch:", getattr(trainer, "best_epoch", None))


# %% [code]
# =========================================================
# 16) Save checkpoint
# =========================================================
CKPT_DIR = DATA_DIR / "checkpoints"
CKPT_DIR.mkdir(exist_ok=True, parents=True)

ckpt_path = CKPT_DIR / "univi_scnmt_gastrulation_genebody_rna_cpg_gpc_beta_binom.pt"

ckpt = {
    "model_state_dict": model.state_dict(),
    "model_config": univi_cfg,
    "train_cfg": train_cfg,
    "history": history,
    "best_epoch": getattr(trainer, "best_epoch", None),
    "train_idx": train_idx,
    "val_idx": val_idx,
    "test_idx": test_idx,
}
torch.save(ckpt, ckpt_path)

print("Saved:", ckpt_path)
print("Best epoch:", ckpt.get("best_epoch"))


# %% [code]
# =========================================================
# 17) Encode fused latent + UMAP (tri-modal)
# =========================================================
from univi.evaluation import encode_fused_adata_pair, encode_adata
from univi.plotting import set_style, umap

set_style(font_scale=0.6, dpi=300)

# Subset to test cells for inductive eval if desired
rna_test = adata_dict["rna"][test_idx].copy()
cpg_test = adata_dict["cpg"][test_idx].copy()
gpc_test = adata_dict["gpc"][test_idx].copy()

#rna_test = adata_dict["rna"].copy()
#cpg_test = adata_dict["cpg"].copy()
#gpc_test = adata_dict["gpc"].copy()

assert np.array_equal(rna_test.obs_names, cpg_test.obs_names)
assert np.array_equal(rna_test.obs_names, gpc_test.obs_names)

# Fused encoding across all 3 modalities
fused = encode_fused_adata_pair(
    model,
    adata_by_mod={"rna": rna_test, "cpg": cpg_test, "gpc": gpc_test},
    device=device,
    batch_size=48,
    use_mean=True,
    return_gates=True,
    return_gate_logits=True,
    write_to_adatas=True,
    fused_obsm_key="X_univi_fused",
    gate_prefix="gate",
)

print("fused keys:", list(fused.keys()))
print("Z_fused shape:", None if "Z_fused" not in fused else fused["Z_fused"].shape)

# Plot on one modality object (same cells/order)
color_cols = [c for c in ["stage", "lineage10x", "lineage10x_2"] if c in rna_test.obs.columns]
if len(color_cols) == 0:
    color_cols = None

umap(
    rna_test,
    obsm_key="X_univi_fused",
    color=color_cols,
    legend="outside",
    show=True,
)


# %% [code]
# =========================================================
# 18) Optional cross-modal prediction examples
# =========================================================
from univi.evaluation import cross_modal_predict, evaluate_cross_reconstruction, reconstruction_metrics

# RNA -> CpG
cpg_hat_from_rna = cross_modal_predict(
    model,
    adata_src=rna_test,
    src_mod="rna",
    tgt_mod="cpg",
    device=device,
    layer=None,
    X_key="X",
    batch_size=48,
    use_moe=True,
)
cpg_test.layers["imputed_from_rna"] = cpg_hat_from_rna

# RNA -> GpC
gpc_hat_from_rna = cross_modal_predict(
    model,
    adata_src=rna_test,
    src_mod="rna",
    tgt_mod="gpc",
    device=device,
    layer=None,
    X_key="X",
    batch_size=48,
    use_moe=True,
)
gpc_test.layers["imputed_from_rna"] = gpc_hat_from_rna

print("Stored imputed layers:")
print("cpg:", cpg_test.layers.keys())
print("gpc:", gpc_test.layers.keys())


# %% [code]
# =========================================================
# 19) Quick evaluation helpers
# =========================================================
def to_dense(X):
    return X.toarray() if sp.issparse(X) else np.asarray(X)

# Example: compare predicted CpG fractions vs observed CpG fractions on test cells
true_cpg = to_dense(cpg_test.X)
pred_cpg = to_dense(cpg_test.layers["imputed_from_rna"])

# If dimensions match:
if true_cpg.shape == pred_cpg.shape:
    from univi.evaluation import reconstruction_metrics
    rep = reconstruction_metrics(true_cpg, pred_cpg)
    print("CpG (RNA->CpG) MSE mean:", rep.get("mse_mean"))
    print("CpG (RNA->CpG) Pearson mean:", rep.get("pearson_mean"))
else:
    print("Shape mismatch:", true_cpg.shape, pred_cpg.shape)
    

# %% [markdown]
NOTE: Since this is a small dataset, we can either use the test set to evaluate and visualize, or we can just use the whole dataset and treat it as transductive.

# %% [code]
from univi.evaluation import encode_adata, cross_modal_predict, evaluate_cross_reconstruction

#rna_test = adata_dict["rna"][test_idx].copy()
#cpg_test = adata_dict["cpg"][test_idx].copy()
#gpc_test = adata_dict["gpc"][test_idx].copy()
#rna_test = adata_dict["rna"].copy()
#cpg_test = adata_dict["cpg"].copy()
#gpc_test = adata_dict["gpc"].copy()

# Encode single-modality latents
rna_test.obsm["X_univi"] = encode_adata(
    model, rna_test, modality="rna", device=device, batch_size=48, latent="moe_mean"
)
cpg_test.obsm["X_univi"] = encode_adata(
    model, cpg_test, modality="cpg", device=device, batch_size=48, latent="moe_mean"
)
gpc_test.obsm["X_univi"] = encode_adata(
    model, gpc_test, modality="gpc", device=device, batch_size=48, latent="moe_mean"
)

print("Latent shapes:")
print("  RNA:", rna_test.obsm["X_univi"].shape)
print("  CpG:", cpg_test.obsm["X_univi"].shape)
print("  GpC:", gpc_test.obsm["X_univi"].shape)

# Cross-modal prediction: RNA -> CpG
cpg_hat_from_rna = cross_modal_predict(
    model,
    adata_src=rna_test,
    src_mod="rna",
    tgt_mod="cpg",
    device=device,
    batch_size=48,
    use_moe=True,
)
cpg_test.layers["imputed_from_rna"] = cpg_hat_from_rna

rep_rna_to_cpg = evaluate_cross_reconstruction(
    model,
    adata_src=rna_test,
    adata_tgt=cpg_test,
    src_mod="rna",
    tgt_mod="cpg",
    device=device,
    batch_size=48,
)
print("RNA -> CpG summary:")
print(rep_rna_to_cpg["summary"])


# %% [code]
print(rna_test)


# %% [code]
from univi.plotting import set_style, umap

set_style(font_scale=0.8, dpi=300)

# make UMAP on RNA latent
sc.pp.neighbors(rna_test, use_rep="X_univi")
sc.tl.umap(rna_test)

umap(
    rna_test,
    obsm_key="X_univi",
    color="lineage10x",
    legend="outside",
    show=True,
)


# %% [code]
print(rna_test)


# %% [code]
umap(
    rna_test,
    obsm_key="X_univi",
    color=["lineage10x"],
    legend="outside",
    legend_subset_topk=50,
    #savepath="umap_rna_univi.png",
    show=True,
)


# %% [code]
umap(
    cpg_test,
    obsm_key="X_univi",
    color=["lineage10x"],
    legend="outside",
    legend_subset_topk=50,
    #savepath="umap_rna_univi.png",
    show=True,
)


# %% [code]
umap(
    gpc_test,
    obsm_key="X_univi",
    color=["lineage10x"],
    legend="outside",
    legend_subset_topk=50,
    #savepath="umap_rna_univi.png",
    show=True,
)


# %% [code]
from univi.evaluation import encode_fused_adata_pair

# despite the name, it accepts adata_by_mod with multiple modalities in your README examples
fused = encode_fused_adata_pair(
    model,
    adata_by_mod={"rna": rna_test, "cpg": cpg_test, "gpc": gpc_test},
    device=device,
    batch_size=48,
    use_mean=True,
    return_gates=True,
    return_gate_logits=True,
    write_to_adatas=True,
    fused_obsm_key="X_univi_fused",
    gate_prefix="gate",
)

print(fused.keys())
print("Z_fused shape:", fused["Z_fused"].shape)


# %% [code]
from univi.evaluation import evaluate_alignment

# RNA vs CpG alignment in latent
metrics_rna_cpg = evaluate_alignment(
    Z1=rna_test.obsm["X_univi"],
    Z2=cpg_test.obsm["X_univi"],
    metric="euclidean",
    recall_ks=(1, 5, 10, 25, 50, 100),
    k_mixing=8,
    k_entropy=8,
    json_safe=True,
)
print(metrics_rna_cpg)


# %% [code]
from univi.evaluation import cross_modal_predict

cpg_hat_from_rna = cross_modal_predict(
    model,
    adata_src=rna_test,
    src_mod="rna",
    tgt_mod="cpg",
    device=device,
    X_key="X",
    batch_size=48,
    use_moe=True,
)

# Store as an imputed layer (mean-like output from decoder utility)
cpg_test.layers["imputed_from_rna"] = cpg_hat_from_rna
print("cpg_hat_from_rna shape:", np.asarray(cpg_hat_from_rna).shape)


# %% [code]


# %% [code]
import numpy as np
import scipy.sparse as sp
import torch

from univi.evaluation import (
    encode_adata,
    encode_fused_adata_pair,
    cross_modal_predict,
    denoise_adata,
    denoise_from_multimodal,
    evaluate_alignment,
    reconstruction_metrics,
    # NEW (generation + recon error workflows)
    generate_from_latent,
    fit_label_latent_gaussians,
    sample_latent_by_label,
    evaluate_cross_reconstruction,
)
from univi.plotting import (
    set_style,
    umap,
    umap_by_modality,
    compare_raw_vs_denoised_umap_features,
    plot_confusion_matrix,
    write_gates_to_obs,
    plot_moe_gate_summary,
    # NEW (reconstruction error plots)
    plot_reconstruction_error_summary,
    plot_featurewise_reconstruction_scatter,
)

set_style(font_scale=0.6, dpi=300)
device = "cuda" if torch.cuda.is_available() else ("mps" if torch.backends.mps.is_available() else "cpu")


# %% [code]
def to_dense(X):
    return X.toarray() if sp.issparse(X) else np.asarray(X)


# %% [code]
fused = encode_fused_adata_pair(
    model,
    adata_by_mod={"rna": rna_test, "cpg": cpg_test, "gpc": gpc_test},
    device=device,
    batch_size=48,
    use_mean=True,
    return_gates=True,
    return_gate_logits=True,
    write_to_adatas=True,
    fused_obsm_key="X_univi_fused",
    gate_prefix="gate",
)

# fused["Z_fused"] -> (n_cells, latent_dim)
# fused["gates"]  -> (n_cells, n_modalities) or None (if fused transformer posterior is used)


# %% [code]
print(rna_test)


# %% [code]
import anndata as ad
import scanpy as sc

# Stack once, label modality, compute UMAP once
stacked = ad.concat(
    {"rna": rna_test, "cpg": cpg_test, "gpc": gpc_test},
    label="univi_modality",
    join="outer",
    index_unique="-",   # avoids the obs_names duplicate warning
)
stacked = ad.concat(
    {"rna": rna_test, "cpg": cpg_test, "gpc": gpc_test},
    label="univi_modality",
    join="outer",
    index_unique="-",
)
# X_univi_fused is already on stacked.obsm at this point — no manual assignment

sc.pp.neighbors(stacked, use_rep="X_univi_fused", random_state=0)
sc.tl.umap(stacked, random_state=0)

for c in ["univi_modality", "lineage10x", "embryo", "stage"]:
    sc.pl.umap(stacked, color=c, size=20, legend_loc="right margin")
    

# %% [code]
umap_by_modality(
    {"rna": rna_test, "cpg": cpg_test, "gpc": gpc_test},
    obsm_key="X_univi_fused",
    color=["univi_modality"],
    legend="outside",
    size=20,
    #savepath="umap_fused_both_modalities.png",
    show=True,
)

umap_by_modality(
    {"rna": rna_test, "cpg": cpg_test, "gpc": gpc_test},
    obsm_key="X_univi_fused",
    color=["lineage10x"],
    legend="outside",
    size=20,
    #savepath="umap_fused_both_modalities.png",
    show=True,
)


# %% [code]
umap_by_modality(
    {"rna": rna_test, "cpg": cpg_test, "gpc": gpc_test},
    obsm_key="X_univi_fused",
    color=["stage"],
    legend="outside",
    size=20,
    #savepath="umap_fused_both_modalities.png",
    show=True,
)

umap_by_modality(
    {"rna": rna_test, "cpg": cpg_test, "gpc": gpc_test},
    obsm_key="X_univi_fused",
    color=["embryo"],
    legend="outside",
    size=20,
    #savepath="umap_fused_both_modalities.png",
    show=True,
)


# %% [code]
cpg_hat_from_rna = cross_modal_predict(
    model,
    adata_src=rna_test,
    src_mod="rna",
    tgt_mod="cpg",
    device=device,
    layer=None,
    X_key="X",
    batch_size=8,
    use_moe=True,
)

cpg_test.layers["imputed_from_rna"] = cpg_hat_from_rna


# %% [code]
gpc_hat_from_cpg = cross_modal_predict(
    model,
    adata_src=cpg_test,
    src_mod="cpg",
    tgt_mod="gpc",
    device=device,
    layer=None,
    X_key="X",
    batch_size=48,
    use_moe=True,
)

gpc_test.layers["imputed_from_cpg"] = gpc_hat_from_cpg


# %% [code]
rna_hat_from_cpg = cross_modal_predict(
    model,
    adata_src=cpg_test,
    src_mod="cpg",
    tgt_mod="rna",
    device=device,
    layer=None,
    X_key="X",
    batch_size=48,
    use_moe=True,
)

rna_test.layers["imputed_from_cpg"] = rna_hat_from_cpg


# %% [code]
true = to_dense(cpg_test.X)
pred = cpg_test.layers["imputed_from_rna"]

m = reconstruction_metrics(true, pred)
print("MSE mean:", m["mse_mean"])
print("Pearson mean:", m["pearson_mean"])


# %% [code]
rep = evaluate_cross_reconstruction(
    model,
    adata_src=rna_test,
    adata_tgt=cpg_test,
    src_mod="rna",
    tgt_mod="cpg",
    device=device,
    src_layer=None,
    tgt_layer=None,
    batch_size=48,
    # optionally restrict to a feature subset (e.g., top markers)
    feature_names=None,
)
print(rep["summary"])   # mse_mean/median, pearson_mean/median, etc.


# %% [code]
plot_reconstruction_error_summary(
    rep,
    title="RNA → CPG imputation error",
    #savepath="recon_error_summary.png",
    show=True,
)


# %% [code]
from univi.evaluation import to_dense, encode_moe_gates_from_tensors
from univi.plotting import write_gates_to_obs, plot_moe_gate_summary

gate = encode_moe_gates_from_tensors(
    model,
    x_dict={"rna": to_dense(rna_test.X), "cpg": to_dense(cpg_test.X), "gpc": to_dense(gpc_test.X)},
    device=device,
    batch_size=48,
    modality_order=["rna", "cpg", "gpc"],
    kind="router_x_precision",  # falls back to "effective_precision" if router logits are unavailable
    return_logits=True,
)

W    = gate["weights"]         # (n_cells, n_modalities), rows sum to 1
mods = gate["modality_order"]  # e.g. ["rna", "adt"]

print("Requested kind:", gate.get("requested_kind"))
print("Effective kind:", gate.get("kind"))
print("Per-modality mean:", gate.get("per_modality_mean"))
print("Has logits:", gate.get("logits") is not None)


# %% [code]
write_gates_to_obs(
    rna_test,
    gates=W,
    modality_names=mods,
    gate_prefix="moe_gate",          # creates obs cols: moe_gate_{mod}
    gate_logits=gate.get("logits"),  # optional; may be None
)


# %% [code]
print(rna_test)


# %% [code]
plot_moe_gate_summary(
    rna_test,
    gate_prefix="moe_gate",
    groupby="lineage10x",
    agg="mean",
    #savepath="moe_gates_by_celltype.png",
    show=True,
)


# %% [markdown]
Evaluate using all data rather than just the test set:

# %% [code]
# =========================================================
# 17) Encode fused latent + UMAP (tri-modal)
# =========================================================
from univi.evaluation import encode_fused_adata_pair, encode_adata
from univi.plotting import set_style, umap

set_style(font_scale=0.6, dpi=300)

# Subset to test cells for inductive eval if desired
#rna_test = adata_dict["rna"][test_idx].copy()
#cpg_test = adata_dict["cpg"][test_idx].copy()
#gpc_test = adata_dict["gpc"][test_idx].copy()

rna_test = adata_dict["rna"].copy()
cpg_test = adata_dict["cpg"].copy()
gpc_test = adata_dict["gpc"].copy()

assert np.array_equal(rna_test.obs_names, cpg_test.obs_names)
assert np.array_equal(rna_test.obs_names, gpc_test.obs_names)

# Fused encoding across all 3 modalities
fused = encode_fused_adata_pair(
    model,
    adata_by_mod={"rna": rna_test, "cpg": cpg_test, "gpc": gpc_test},
    device=device,
    batch_size=48,
    use_mean=True,
    return_gates=True,
    return_gate_logits=True,
    write_to_adatas=True,
    fused_obsm_key="X_univi_fused",
    gate_prefix="gate",
)

print("fused keys:", list(fused.keys()))
print("Z_fused shape:", None if "Z_fused" not in fused else fused["Z_fused"].shape)

# Plot on one modality object (same cells/order)
color_cols = [c for c in ["stage", "lineage10x", "lineage10x_2"] if c in rna_test.obs.columns]
if len(color_cols) == 0:
    color_cols = None

umap(
    rna_test,
    obsm_key="X_univi_fused",
    color=color_cols,
    legend="outside",
    show=True,
)


# %% [code]
# =========================================================
# 18) Optional cross-modal prediction examples
# =========================================================
from univi.evaluation import cross_modal_predict, evaluate_cross_reconstruction, reconstruction_metrics

# RNA -> CpG
cpg_hat_from_rna = cross_modal_predict(
    model,
    adata_src=rna_test,
    src_mod="rna",
    tgt_mod="cpg",
    device=device,
    layer=None,
    X_key="X",
    batch_size=48,
    use_moe=True,
)
cpg_test.layers["imputed_from_rna"] = cpg_hat_from_rna

# RNA -> GpC
gpc_hat_from_rna = cross_modal_predict(
    model,
    adata_src=rna_test,
    src_mod="rna",
    tgt_mod="gpc",
    device=device,
    layer=None,
    X_key="X",
    batch_size=48,
    use_moe=True,
)
gpc_test.layers["imputed_from_rna"] = gpc_hat_from_rna

print("Stored imputed layers:")
print("cpg:", cpg_test.layers.keys())
print("gpc:", gpc_test.layers.keys())


# =========================================================
# 19) Quick evaluation helpers
# =========================================================
def to_dense(X):
    return X.toarray() if sp.issparse(X) else np.asarray(X)

# Example: compare predicted CpG fractions vs observed CpG fractions on test cells
true_cpg = to_dense(cpg_test.X)
pred_cpg = to_dense(cpg_test.layers["imputed_from_rna"])

# If dimensions match:
if true_cpg.shape == pred_cpg.shape:
    from univi.evaluation import reconstruction_metrics
    rep = reconstruction_metrics(true_cpg, pred_cpg)
    print("CpG (RNA->CpG) MSE mean:", rep.get("mse_mean"))
    print("CpG (RNA->CpG) Pearson mean:", rep.get("pearson_mean"))
else:
    print("Shape mismatch:", true_cpg.shape, pred_cpg.shape)
    

from univi.evaluation import encode_adata, cross_modal_predict, evaluate_cross_reconstruction

#rna_test = adata_dict["rna"][test_idx].copy()
#cpg_test = adata_dict["cpg"][test_idx].copy()
#gpc_test = adata_dict["gpc"][test_idx].copy()
#rna_test = adata_dict["rna"].copy()
#cpg_test = adata_dict["cpg"].copy()
#gpc_test = adata_dict["gpc"].copy()

# Encode single-modality latents
rna_test.obsm["X_univi"] = encode_adata(
    model, rna_test, modality="rna", device=device, batch_size=48, latent="moe_mean"
)
cpg_test.obsm["X_univi"] = encode_adata(
    model, cpg_test, modality="cpg", device=device, batch_size=48, latent="moe_mean"
)
gpc_test.obsm["X_univi"] = encode_adata(
    model, gpc_test, modality="gpc", device=device, batch_size=48, latent="moe_mean"
)

print("Latent shapes:")
print("  RNA:", rna_test.obsm["X_univi"].shape)
print("  CpG:", cpg_test.obsm["X_univi"].shape)
print("  GpC:", gpc_test.obsm["X_univi"].shape)

# Cross-modal prediction: RNA -> CpG
cpg_hat_from_rna = cross_modal_predict(
    model,
    adata_src=rna_test,
    src_mod="rna",
    tgt_mod="cpg",
    device=device,
    batch_size=48,
    use_moe=True,
)
cpg_test.layers["imputed_from_rna"] = cpg_hat_from_rna

rep_rna_to_cpg = evaluate_cross_reconstruction(
    model,
    adata_src=rna_test,
    adata_tgt=cpg_test,
    src_mod="rna",
    tgt_mod="cpg",
    device=device,
    batch_size=48,
)
print("RNA -> CpG summary:")
print(rep_rna_to_cpg["summary"])


from univi.plotting import set_style, umap

set_style(font_scale=0.8, dpi=300)

# make UMAP on RNA latent
sc.pp.neighbors(rna_test, use_rep="X_univi")
sc.tl.umap(rna_test)


umap(
    rna_test,
    obsm_key="X_univi",
    color=["lineage10x"],
    legend="outside",
    legend_subset_topk=25,
    #savepath="umap_rna_univi.png",
    show=True,
)


umap(
    cpg_test,
    obsm_key="X_univi",
    color=["lineage10x"],
    legend="outside",
    legend_subset_topk=25,
    #savepath="umap_rna_univi.png",
    show=True,
)


umap(
    gpc_test,
    obsm_key="X_univi",
    color=["lineage10x"],
    legend="outside",
    legend_subset_topk=25,
    #savepath="umap_rna_univi.png",
    show=True,
)


from univi.evaluation import encode_fused_adata_pair

# despite the name, it accepts adata_by_mod with multiple modalities in your README examples
fused = encode_fused_adata_pair(
    model,
    adata_by_mod={"rna": rna_test, "cpg": cpg_test, "gpc": gpc_test},
    device=device,
    batch_size=48,
    use_mean=True,
    return_gates=True,
    return_gate_logits=True,
    write_to_adatas=True,
    fused_obsm_key="X_univi_fused",
    gate_prefix="gate",
)

print(fused.keys())
print("Z_fused shape:", fused["Z_fused"].shape)


from univi.evaluation import evaluate_alignment

# RNA vs CpG alignment in latent
metrics_rna_cpg = evaluate_alignment(
    Z1=rna_test.obsm["X_univi"],
    Z2=cpg_test.obsm["X_univi"],
    metric="euclidean",
    recall_ks=(1, 5, 10, 25, 50, 100),
    k_mixing=8,
    k_entropy=8,
    json_safe=True,
)
print(metrics_rna_cpg)


from univi.evaluation import cross_modal_predict

cpg_hat_from_rna = cross_modal_predict(
    model,
    adata_src=rna_test,
    src_mod="rna",
    tgt_mod="cpg",
    device=device,
    X_key="X",
    batch_size=128,
    use_moe=True,
)

# Store as an imputed layer (mean-like output from decoder utility)
cpg_test.layers["imputed_from_rna"] = cpg_hat_from_rna
print("cpg_hat_from_rna shape:", np.asarray(cpg_hat_from_rna).shape)


import numpy as np
import scipy.sparse as sp
import torch

from univi.evaluation import (
    encode_adata,
    encode_fused_adata_pair,
    cross_modal_predict,
    denoise_adata,
    denoise_from_multimodal,
    evaluate_alignment,
    reconstruction_metrics,
    # NEW (generation + recon error workflows)
    generate_from_latent,
    fit_label_latent_gaussians,
    sample_latent_by_label,
    evaluate_cross_reconstruction,
)
from univi.plotting import (
    set_style,
    umap,
    umap_by_modality,
    compare_raw_vs_denoised_umap_features,
    plot_confusion_matrix,
    write_gates_to_obs,
    plot_moe_gate_summary,
    # NEW (reconstruction error plots)
    plot_reconstruction_error_summary,
    plot_featurewise_reconstruction_scatter,
)

set_style(font_scale=0.8, dpi=300)
device = "cuda" if torch.cuda.is_available() else ("mps" if torch.backends.mps.is_available() else "cpu")


fused = encode_fused_adata_pair(
    model,
    adata_by_mod={"rna": rna_test, "cpg": cpg_test, "gpc": gpc_test},
    device=device,
    batch_size=8,
    use_mean=True,
    return_gates=True,
    return_gate_logits=True,
    write_to_adatas=True,
    fused_obsm_key="X_univi_fused",
    gate_prefix="gate",
)

# fused["Z_fused"] -> (n_cells, latent_dim)
# fused["gates"]  -> (n_cells, n_modalities) or None (if fused transformer posterior is used)


umap_by_modality(
    {"rna": rna_test, "cpg": cpg_test, "gpc": gpc_test},
    obsm_key="X_univi_fused",
    color=["univi_modality"],
    legend="outside",
    size=8,
    #savepath="umap_fused_both_modalities.png",
    show=True,
)

umap_by_modality(
    {"rna": rna_test, "cpg": cpg_test, "gpc": gpc_test},
    obsm_key="X_univi_fused",
    color=["lineage10x"],
    legend="outside",
    size=8,
    #savepath="umap_fused_both_modalities.png",
    show=True,
)

umap_by_modality(
    {"rna": rna_test, "cpg": cpg_test, "gpc": gpc_test},
    obsm_key="X_univi_fused",
    color=["embryo"],
    legend="outside",
    size=8,
    #savepath="umap_fused_both_modalities.png",
    show=True,
)

umap_by_modality(
    {"rna": rna_test, "cpg": cpg_test, "gpc": gpc_test},
    obsm_key="X_univi_fused",
    color=["stage"],
    legend="outside",
    size=8,
    #savepath="umap_fused_both_modalities.png",
    show=True,
)

cpg_hat_from_rna = cross_modal_predict(
    model,
    adata_src=rna_test,
    src_mod="rna",
    tgt_mod="cpg",
    device=device,
    layer=None,
    X_key="X",
    batch_size=8,
    use_moe=True,
)

cpg_test.layers["imputed_from_rna"] = cpg_hat_from_rna


gpc_hat_from_cpg = cross_modal_predict(
    model,
    adata_src=cpg_test,
    src_mod="cpg",
    tgt_mod="gpc",
    device=device,
    layer=None,
    X_key="X",
    batch_size=8,
    use_moe=True,
)

gpc_test.layers["imputed_from_cpg"] = gpc_hat_from_cpg


rna_hat_from_cpg = cross_modal_predict(
    model,
    adata_src=cpg_test,
    src_mod="cpg",
    tgt_mod="rna",
    device=device,
    layer=None,
    X_key="X",
    batch_size=8,
    use_moe=True,
)

rna_test.layers["imputed_from_cpg"] = rna_hat_from_cpg


true = to_dense(cpg_test.X)
pred = cpg_test.layers["imputed_from_rna"]

m = reconstruction_metrics(true, pred)
print("MSE mean:", m["mse_mean"])
print("Pearson mean:", m["pearson_mean"])


rep = evaluate_cross_reconstruction(
    model,
    adata_src=rna_test,
    adata_tgt=cpg_test,
    src_mod="rna",
    tgt_mod="cpg",
    device=device,
    src_layer=None,
    tgt_layer=None,
    batch_size=8,
    # optionally restrict to a feature subset (e.g., top markers)
    feature_names=None,
)
print(rep["summary"])   # mse_mean/median, pearson_mean/median, etc.


plot_reconstruction_error_summary(
    rep,
    title="RNA → CPG imputation error",
    #savepath="recon_error_summary.png",
    show=True,
)


from univi.evaluation import to_dense, encode_moe_gates_from_tensors
from univi.plotting import write_gates_to_obs, plot_moe_gate_summary

gate = encode_moe_gates_from_tensors(
    model,
    x_dict={"rna": to_dense(rna_test.X), "cpg": to_dense(cpg_test.X), "gpc": to_dense(gpc_test.X)},
    device=device,
    batch_size=8,
    modality_order=["rna", "cpg", "gpc"],
    kind="router_x_precision",  # falls back to "effective_precision" if router logits are unavailable
    return_logits=True,
)

W    = gate["weights"]         # (n_cells, n_modalities), rows sum to 1
mods = gate["modality_order"]  # e.g. ["rna", "adt"]

print("Requested kind:", gate.get("requested_kind"))
print("Effective kind:", gate.get("kind"))
print("Per-modality mean:", gate.get("per_modality_mean"))
print("Has logits:", gate.get("logits") is not None)


write_gates_to_obs(
    rna_test,
    gates=W,
    modality_names=mods,
    gate_prefix="moe_gate",          # creates obs cols: moe_gate_{mod}
    gate_logits=gate.get("logits"),  # optional; may be None
)


plot_moe_gate_summary(
    rna_test,
    gate_prefix="moe_gate",
    groupby="lineage10x",
    agg="mean",
    #savepath="moe_gates_by_celltype.png",
    show=True,
)


# %% [code]
import anndata as ad
import scanpy as sc

# Stack once, label modality, compute UMAP once
stacked = ad.concat(
    {"rna": rna_test, "cpg": cpg_test, "gpc": gpc_test},
    label="univi_modality",
    join="outer",
    index_unique="-",   # avoids the obs_names duplicate warning
)
stacked = ad.concat(
    {"rna": rna_test, "cpg": cpg_test, "gpc": gpc_test},
    label="univi_modality",
    join="outer",
    index_unique="-",
)
# X_univi_fused is already on stacked.obsm at this point — no manual assignment

sc.pp.neighbors(stacked, use_rep="X_univi_fused", random_state=0)
sc.tl.umap(stacked, random_state=0)

for c in ["univi_modality", "lineage10x", "embryo", "stage"]:
    sc.pl.umap(stacked, color=c, size=20, legend_loc="right margin")
    

# %% [code]


# %% [code]


# %% [code]


# %% [code]


# %% [code]


# %% [code]


# %% [code]


# %% [code]


# %% [code]


# %% [code]
```
