## Supplemental Code for "Unifying multimodal single-cell data with a mixture-of-experts *β*-variational autoencoder framework": Supplemental_Notebook_S1.ipynb

```
# %% [code]
# Import non-UniVI modules
import os
import sys
from pathlib import Path

import numpy as np
import pandas as pd
import scanpy as sc
import anndata as ad
import re

import torch
from torch.utils.data import DataLoader, Subset

import scipy.sparse as sp


# %% [code]
from sklearn.feature_extraction.text import TfidfTransformer
from sklearn.decomposition import TruncatedSVD
from sklearn.preprocessing import StandardScaler, normalize

from sklearn.neighbors import NearestNeighbors
from sklearn.metrics import confusion_matrix, accuracy_score, f1_score

import matplotlib.pyplot as plt


# %% [code]
# Import required UniVI modules
from univi import (
    ModalityConfig,
    UniVIConfig,
    TrainingConfig,
    UniVIMultiModalVAE,
    matching,
    UniVITrainer,
    write_univi_latent,
    MultiModalDataset,
)

import univi as uv
import univi.evaluation as ue
import univi.plotting as up

# Double check UniVI module version
print("Installed version is univi v" + str(uv.__version__))


# %% [code]
print("torch:", torch.__version__)
print("torch.cuda.is_available():", torch.cuda.is_available())
device = "cuda" if torch.cuda.is_available() else ("mps" if torch.mps.is_available() else "cpu")
print("Using device:", device)


# %% [code]
MULTIOME_DATA_ROOT = Path("./data/10x_Genomics_Multiome_data")

MULTIOME_RNA_PATH = MULTIOME_DATA_ROOT / "10x-Multiome-Pbmc10k-RNA.h5ad"
MULTIOME_ATAC_PATH = MULTIOME_DATA_ROOT / "10x-Multiome-Pbmc10k-ATAC.h5ad"

print("Multiome RNA file:", MULTIOME_RNA_PATH)
print("Multiome ATAC file:", MULTIOME_ATAC_PATH)


# %% [code]
multiome_rna = sc.read_h5ad(MULTIOME_RNA_PATH)
multiome_atac = sc.read_h5ad(MULTIOME_ATAC_PATH)

print(multiome_rna)
print(multiome_atac)

print("Multiome RNA obs names head:", multiome_rna.obs_names[:5].tolist())
print("Multiome ATAC obs names head:", multiome_atac.obs_names[:5].tolist())


# %% [code]
from univi.data import align_paired_obs_names

adata_dict = {'rna': multiome_rna, 'atac': multiome_atac}

align_paired_obs_names(adata_dict)


# %% [code]
multiome_rna = adata_dict['rna'].copy()
multiome_atac = adata_dict['atac'].copy()


# %% [code]
rna_raw  = multiome_rna.copy()
atac_raw = multiome_atac.copy()


# %% [code]
import numpy as np
import pandas as pd
import scipy.sparse as sp
import anndata as ad

# ----------------------------
# helpers
# ----------------------------
def _to_csr(X):
    return X if sp.issparse(X) else sp.csr_matrix(X)

def ensure_counts_in_layer(adata: ad.AnnData, counts_layer="counts") -> ad.AnnData:
    """Make sure raw counts exist in adata.layers[counts_layer]."""
    a = adata.copy()
    if counts_layer not in a.layers:
        a.layers[counts_layer] = a.X.copy()
    # keep counts sparse if possible
    a.layers[counts_layer] = _to_csr(a.layers[counts_layer])
    return a

def set_X_to_layer(adata: ad.AnnData, layer: str) -> ad.AnnData:
    """Set .X to a given layer (e.g. raw counts)."""
    a = adata.copy()
    if layer not in a.layers:
        raise KeyError(f"Layer {layer!r} not found.")
    a.X = a.layers[layer]
    return a

def stratified_split_indices(labels, train_frac=0.80, val_frac=0.10, seed=0, min_per_class=1):
    """
    Stratified train/val/test split.
    test_frac is implied as 1 - train_frac - val_frac.
    """
    assert 0 < train_frac < 1 and 0 <= val_frac < 1
    assert train_frac + val_frac < 1
    rng = np.random.default_rng(seed)

    y = pd.Series(labels).astype("category")
    idx_all = np.arange(y.shape[0])

    train_idx, val_idx, test_idx = [], [], []
    for ct in y.cat.categories:
        idx_ct = idx_all[(y.values == ct)]
        if idx_ct.size < min_per_class:
            # dump tiny classes into train (or handle however you prefer)
            train_idx.extend(idx_ct.tolist())
            continue

        rng.shuffle(idx_ct)
        n = idx_ct.size
        n_tr = int(np.floor(train_frac * n))
        n_va = int(np.floor(val_frac * n))

        train_idx.extend(idx_ct[:n_tr].tolist())
        val_idx.extend(idx_ct[n_tr:n_tr+n_va].tolist())
        test_idx.extend(idx_ct[n_tr+n_va:].tolist())

    train_idx = np.asarray(train_idx, dtype=int)
    val_idx   = np.asarray(val_idx, dtype=int)
    test_idx  = np.asarray(test_idx, dtype=int)

    rng.shuffle(train_idx); rng.shuffle(val_idx); rng.shuffle(test_idx)
    return train_idx, val_idx, test_idx

# ----------------------------
# raw-count "preprocess"
# ----------------------------
def preprocess_splits_raw_counts(
    rna: ad.AnnData,
    atac: ad.AnnData,
    train_idx: np.ndarray,
    val_idx: np.ndarray,
    test_idx: np.ndarray,
    *,
    counts_layer: str = "counts",
    set_X: bool = True,
):
    """
    Returns train/val/test AnnData for RNA and ATAC using raw counts only.
    If set_X=True, each returned AnnData will have .X == raw counts.
    """
    rna = ensure_counts_in_layer(rna, counts_layer=counts_layer)
    atac = ensure_counts_in_layer(atac, counts_layer=counts_layer)

    rna_tr, rna_va, rna_te = rna[train_idx].copy(), rna[val_idx].copy(), rna[test_idx].copy()
    atac_tr, atac_va, atac_te = atac[train_idx].copy(), atac[val_idx].copy(), atac[test_idx].copy()

    if set_X:
        rna_tr  = set_X_to_layer(rna_tr,  counts_layer)
        rna_va  = set_X_to_layer(rna_va,  counts_layer)
        rna_te  = set_X_to_layer(rna_te,  counts_layer)
        atac_tr = set_X_to_layer(atac_tr, counts_layer)
        atac_va = set_X_to_layer(atac_va, counts_layer)
        atac_te = set_X_to_layer(atac_te, counts_layer)

    return rna_tr, atac_tr, rna_va, atac_va, rna_te, atac_te


# %% [code]
# To make peaks binary and use a Bernoulli decoder model - uncomment the following two code chunks and run them in sequence:

'''
import numpy as np
import scipy.sparse as sp

def binarize_layer(adata, layer="counts", out_layer="bin"):
    a = adata.copy()
    X = a.layers[layer] if layer in a.layers else a.X
    if sp.issparse(X):
        Xb = X.copy()
        Xb.data = np.ones_like(Xb.data, dtype=np.float32)
    else:
        Xb = (X > 0).astype(np.float32)
    a.layers[out_layer] = Xb
    a.X = a.layers[out_layer]  # if your dataset reads from .X
    return a

atac_bin = binarize_layer(atac_raw, layer="counts", out_layer="bin")
'''

# %% [code]
#atac_raw_orig = atac_raw.copy()
#atac_raw = atac_bin.copy()


# %% [code]
CELLTYPE_KEY = "cell_type"
SEED = 0
COUNTS_LAYER = "counts"

assert CELLTYPE_KEY in rna_raw.obs.columns
assert CELLTYPE_KEY in atac_raw.obs.columns

# important: labels should correspond to the same ordering as rna_raw
y = rna_raw.obs[CELLTYPE_KEY].to_numpy()

train_idx, val_idx, test_idx = stratified_split_indices(
    y, train_frac=0.80, val_frac=0.10, seed=SEED, min_per_class=1
)

rna_tr, atac_tr, rna_va, atac_va, rna_te, atac_te = preprocess_splits_raw_counts(
    rna_raw, atac_raw,
    train_idx, val_idx, test_idx,
    counts_layer=COUNTS_LAYER,
    set_X=True,   # .X will be counts
)

print("train/val/test:", rna_tr.n_obs, rna_va.n_obs, rna_te.n_obs)


# %% [code]
adata_dict = {
    "rna":  rna_tr.concatenate(rna_va, rna_te, batch_key=None),
    "atac": atac_tr.concatenate(atac_va, atac_te, batch_key=None),
}

adata_dict = align_paired_obs_names(adata_dict)
dataset = MultiModalDataset(adata_dict=adata_dict, X_key="X", device=None)

n_tr, n_va, n_te = rna_tr.n_obs, rna_va.n_obs, rna_te.n_obs
TRAIN_IDX = np.arange(0, n_tr)
VAL_IDX   = np.arange(n_tr, n_tr + n_va)
TEST_IDX  = np.arange(n_tr + n_va, n_tr + n_va + n_te)


# %% [code]
def inspect_counts(adata, name):
    X = adata.X
    if sp.issparse(X):
        d = X.data
        mn = d.min() if d.size else 0.0
        mx = d.max() if d.size else 0.0
        has_nan = np.isnan(d).any() if d.size else False
        has_inf = np.isinf(d).any() if d.size else False
        sample = d[:200000] if d.size > 200000 else d
    else:
        arr = np.asarray(X)
        mn = np.nanmin(arr)
        mx = np.nanmax(arr)
        has_nan = np.isnan(arr).any()
        has_inf = np.isinf(arr).any()
        sample = arr.ravel()[:200000]

    integer_like = np.all(np.abs(sample - np.round(sample)) < 1e-6) if sample.size else True
    print(f"{name}: min={mn}, max={mx}, nan={has_nan}, inf={has_inf}, integer_like={integer_like}")

inspect_counts(rna_tr, "rna_tr")
inspect_counts(atac_tr, "atac_tr")


# %% [code]
#pin_memory = (device == "mps")
pin_memory = False

batch_size = 256
num_workers = 0


train_dict = {"rna": rna_tr, "atac": atac_tr}
val_dict   = {"rna": rna_va, "atac": atac_va}
test_dict  = {"rna": rna_te, "atac": atac_te}


# Build datasets (CPU tensors; trainer/model moves to GPU)
train_ds = MultiModalDataset(adata_dict=train_dict, X_key="X", device=None)
val_ds   = MultiModalDataset(adata_dict=val_dict,   X_key="X", device=None)
test_ds  = MultiModalDataset(adata_dict=test_dict,  X_key="X", device=None)


train_loader = DataLoader(
    train_ds,
    batch_size=batch_size,
    shuffle=True,
    num_workers=num_workers,
    pin_memory=pin_memory,
    drop_last=True,
)

val_loader = DataLoader(
    val_ds,
    batch_size=batch_size,
    shuffle=False,
    num_workers=num_workers,
    pin_memory=pin_memory,
    drop_last=False,
)

test_loader = DataLoader(
    test_ds,
    batch_size=batch_size,
    shuffle=False,
    num_workers=num_workers,
    pin_memory=pin_memory,
    drop_last=False,
)

print("n_train / n_val / n_test:", train_ds.n_cells, val_ds.n_cells, test_ds.n_cells)
print("batches:", len(train_loader), len(val_loader), len(test_loader))

# sanity check one batch
x = next(iter(train_loader))
print({k: v.shape for k, v in x.items()})


# %% [code]
'''
FOSCTTM mean:          0.07973345957826079
FOSCTTM sem:           0.005462308340337411
Recall@1:              0.07574206755373593
Recall@5:              0.23439099283520984
Recall@10:             0.3510747185261003
Recall@25:             0.5394063459570113
Recall@50:             0.7072671443193449
Recall@100:            0.8413510747185261
Modality mixing mean:  0.4307830110421078
Modality entropy mean: 0.9305325635349494
Label transfer ACC:    0.8300921187308086

# Stats from the below model run using binarized peak features
'''
'''
univi_cfg = UniVIConfig(
    latent_dim=60,
    beta=1.15,
    gamma=3.25,
    encoder_dropout=0.10,
    decoder_dropout=0.05,
    encoder_batchnorm=True,
    decoder_batchnorm=False,
    kl_anneal_start=75,
    kl_anneal_end=150,
    align_anneal_start=100,
    align_anneal_end=175,

    # ---- MoE gating (NEW) ----
    #use_moe_gating=True,
    #moe_gating_type="per_modality",   # or "shared"
    #moe_gating_hidden=[64],           # tiny MLP; you can do [32] too
    #moe_gating_dropout=0.0,
    #moe_gating_batchnorm=False,
    #moe_gating_activation="relu",
    #moe_gate_eps=1e-6,

    modalities=[
        ModalityConfig(
            name="rna",
            input_dim=rna_tr.n_vars,
            #encoder_hidden=[512, 256, 128],
            #decoder_hidden=[128, 256, 512],
            encoder_hidden=[2048, 1024, 512, 256, 128],
            decoder_hidden=[128, 256, 512, 1024, 2048],
            #likelihood="gaussian",
            likelihood="nb",
        ),
        ModalityConfig(
            name="atac",
            input_dim=atac_tr.n_vars,
            #encoder_hidden=[128, 64],
            #decoder_hidden=[64, 128],
            encoder_hidden=[2048, 1024, 512, 256, 128],
            decoder_hidden=[128, 256, 512, 1024, 2048],
            #likelihood="gaussian",
            likelihood="bernoulli",
        ),
    ],
)
'''


# %% [code]
univi_cfg = UniVIConfig(
    latent_dim=40,
    beta=0.85,
    gamma=1.75,
    encoder_dropout=0.075,
    decoder_dropout=0.035,
    encoder_batchnorm=False,
    decoder_batchnorm=False,
    kl_anneal_start=0,
    kl_anneal_end=100,
    align_anneal_start=30,
    align_anneal_end=130,

    # ---- MoE gating (NEW) ----
    #use_moe_gating=True,
    #moe_gating_type="per_modality",   # or "shared"
    #moe_gating_hidden=[64],           # tiny MLP; you can do [32] too
    #moe_gating_dropout=0.0,
    #moe_gating_batchnorm=False,
    #moe_gating_activation="relu",
    #moe_gate_eps=1e-6,

    modalities=[
        ModalityConfig(
            name="rna",
            input_dim=rna_tr.n_vars,
            #encoder_hidden=[512, 256, 128],
            #decoder_hidden=[128, 256, 512],
            encoder_hidden=[1024, 512, 256, 128],
            decoder_hidden=[128, 256, 512, 1024],
            #likelihood="gaussian",
            likelihood="nb",
        ),
        ModalityConfig(
            name="atac",
            input_dim=atac_tr.n_vars,
            #encoder_hidden=[128, 64],
            #decoder_hidden=[64, 128],
            encoder_hidden=[2048, 1024, 512, 256, 128],
            decoder_hidden=[128, 256, 512, 1024, 2048],
            #likelihood="gaussian",
            #likelihood="bernoulli",
            likelihood="poisson",
        ),
    ],
)


# %% [code]
model = UniVIMultiModalVAE(
    univi_cfg,
    loss_mode="v1",
    #loss_mode="lite"
    #v1_recon="cross",
    v1_recon="avg",
    #v1_recon_mix=0.5,
    normalize_v1_terms=True,
).to(device)


# %% [code]
print("use_moe_gating:", model.use_moe_gating)
print("moe_gating_type:", model.moe_gating_type)
print("shared_gate_net:", model.shared_gate_net is not None)
print("gate_nets:", list(model.gate_nets.keys())[:5], "… n=", len(model.gate_nets))


# %% [code]
train_cfg = TrainingConfig(
    n_epochs=5000,
    batch_size=batch_size,
    lr=1e-3,
    weight_decay=1e-4,
    device=device,
    log_every=25,
    grad_clip=5.0,
    num_workers=0,
    seed=0,
    early_stopping=True,
    best_epoch_warmup=10,
    patience=200,
    min_delta=0.0,
)


# %% [code]
trainer = UniVITrainer(
    model=model,
    train_loader=train_loader,
    val_loader=val_loader,
    train_cfg=train_cfg,
    device=device,
)


# %% [code]
history = trainer.fit()

# history is a dict with keys like "train_loss", "val_loss", "beta", "gamma"
print("Training finished.")
print("Best val loss:", np.min(history["val_loss"]))


# %% [code]
plt.figure()
plt.plot(history["train_loss"], label="train")
plt.plot(history["val_loss"], label="val")
plt.xlabel("Epoch")
plt.ylabel("Loss")
plt.legend()
plt.title("UniVI training (Multiome)")
plt.show()


# %% [code]
import torch

ckpt = {
    "model_state_dict": model.state_dict(),
    "model_config": univi_cfg,          # if picklable
    "train_cfg": train_cfg,             # optional, if picklable
    "history": getattr(trainer, "history", None),
    # optional fields if trainer exposes them:
    "best_epoch": getattr(trainer, "best_epoch", None),
    "best_metric": getattr(trainer, "best_metric", None),
}
torch.save(ckpt, "./saved_models/saved_models_2-27-2026/Multiome_raw_all_features_nb-poisson/univi_model_state.pt")


# %% [code]

#ckpt = torch.load("./saved_models/saved_models_2-22-2026/Multiome_raw_all_features_nb-burnoulli/univi_model_state.pt", map_location=device)
ckpt = torch.load("./saved_models/saved_models_2-27-2026/Multiome_raw_all_features_nb-poisson/univi_model_state.pt", map_location=device, weights_only=False)

model = UniVIMultiModalVAE(ckpt["model_config"]).to(device)  # or UniVI(univi_cfg) if you prefer
model.load_state_dict(ckpt["model_state_dict"])
model.eval()

print("Best epoch:", ckpt.get("best_epoch"))


# %% [code]
'''
import sys
import platform
import numpy as np
import torch

from univi.utils.io import save_checkpoint

def _maybe_to_dict(x):
    if x is None:
        return None
    if hasattr(x, "to_dict"):
        return x.to_dict()
    if hasattr(x, "__dict__"):
        # shallow best-effort; replace with dataclasses.asdict if needed
        return {k: v for k, v in vars(x).items()}
    return x

trainer_state = {
    "epoch": getattr(trainer, "epoch", None),
    "best_epoch": getattr(trainer, "best_epoch", None),
    "best_metric": getattr(trainer, "best_metric", None),
    "early_stopping_counter": getattr(trainer, "early_stopping_counter", None),
    # include scheduler state if trainer has one
    "scheduler_state": (
        trainer.scheduler.state_dict()
        if hasattr(trainer, "scheduler") and getattr(trainer, "scheduler", None) is not None
        else None
    ),
}

config_payload = {
    "model_config": _maybe_to_dict(univi_cfg),
    "train_config": _maybe_to_dict(train_cfg),
    "preprocess": {
        "counts_layer": "counts",
        "rna_input": "raw_counts",
        "atac_input": "raw_counts",
        "rna_likelihood": "nb",
        "atac_likelihood": "poisson",
        "celltype_key": "cell_type",
        "split_seed": 0,
    },
}

extra_payload = {
    "history": getattr(trainer, "history", None),
    "best_epoch": getattr(trainer, "best_epoch", None),
    "best_metric": getattr(trainer, "best_metric", None),
    "run_name": "Multiome_raw_all_features_nb-poisson",
    "device": str(device),
    "splits": {
        "train_idx": np.asarray(train_idx, dtype=np.int64),
        "val_idx": np.asarray(val_idx, dtype=np.int64),
        "test_idx": np.asarray(test_idx, dtype=np.int64),
    },
    "data_meta": {
        "rna_n_obs": int(rna_tr.n_obs),
        "rna_n_vars": int(rna_tr.n_vars),
        "atac_n_obs": int(atac_tr.n_obs),
        "atac_n_vars": int(atac_tr.n_vars),
    },
    "env": {
        "python": sys.version,
        "platform": platform.platform(),
        "torch": torch.__version__,
        "device": str(device),
        "mps_available": bool(getattr(torch.backends, "mps", None) and torch.backends.mps.is_available()),
    },
    # Optional but useful
    "rng_state": {
        "torch": torch.get_rng_state(),
        "numpy": np.random.get_state(),
    },
}

save_checkpoint(
    path="./saved_models/saved_models_2-22-2026/Multiome_raw_all_features_nb-poisson/univi_model_state.pt",
    model=model,
    optimizer=getattr(trainer, "optimizer", None),
    trainer_state=trainer_state,
    scaler_state=(trainer.scaler.state_dict() if hasattr(trainer, "scaler") and trainer.scaler is not None else None),
    config=config_payload,
    extra=extra_payload,
)
'''

# %% [code]
'''
from univi.utils.io import load_checkpoint, restore_checkpoint
from univi.config import UniVIConfig, ModalityConfig, ClassHeadConfig

#payload = load_checkpoint(
#    "./saved_models/saved_models_2-22-2026/Multiome_raw_all_features_nb-poisson/univi_model_state.pt",
#    map_location=device,
#)
payload = torch.load(
    "./saved_models/saved_models_2-22-2026/Multiome_raw_all_features_nb-poisson/univi_model_state.pt",
    map_location=device,
    weights_only=False,   # <-- important
)

# Rebuild model from payload["config"]["model_config"] (dict -> UniVIConfig as needed)
cfgd = payload["config"]["model_config"]

# rebuild modalities
modalities = [ModalityConfig(**m) if isinstance(m, dict) else m for m in cfgd["modalities"]]

# rebuild optional class heads (if present)
class_heads = None
if cfgd.get("class_heads") is not None:
    class_heads = [ClassHeadConfig(**h) if isinstance(h, dict) else h for h in cfgd["class_heads"]]

# rebuild top-level config
cfg_kwargs = dict(cfgd)
cfg_kwargs["modalities"] = modalities
if class_heads is not None:
    cfg_kwargs["class_heads"] = class_heads

univi_cfg = UniVIConfig(**cfg_kwargs)
model = UniVIMultiModalVAE(univi_cfg).to(device)

# If you already have the model instance constructed:
restore_checkpoint(
    payload,
    model=model,
    optimizer=getattr(trainer, "optimizer", None),  # optional
    scaler=getattr(trainer, "scaler", None),        # optional
    map_location=device,
    strict=False,
)

model.eval()

print("best_epoch:", payload.get("trainer_state", {}).get("best_epoch"))
print("best_metric:", payload.get("trainer_state", {}).get("best_metric"))
print("run_name:", payload.get("extra", {}).get("run_name"))
print("config keys:", list(payload.get("config", {}).keys()))
'''

# %% [code]
import numpy as np
import scipy.sparse as sp

from univi.evaluation import (
    encode_adata,
    encode_fused_adata_pair,
    cross_modal_predict,
    denoise_adata,
    denoise_from_multimodal,
    evaluate_alignment,
    reconstruction_metrics,
    # NEW (generation + recon error workflows)
    generate_from_latent,
    fit_label_latent_gaussians,
    sample_latent_by_label,
    evaluate_cross_reconstruction,
)
from univi.plotting import (
    set_style,
    umap,
    umap_by_modality,
    compare_raw_vs_denoised_umap_features,
    plot_confusion_matrix,
    write_gates_to_obs,
    plot_moe_gate_summary,
    # NEW (reconstruction error plots)
    plot_reconstruction_error_summary,
    plot_featurewise_reconstruction_scatter,
)

set_style(font_scale=1.2, dpi=150)
device = "mps"  # or "mps" (Mac), or "cpu"


# %% [code]
def to_dense(X):
    return X.toarray() if sp.issparse(X) else np.asarray(X)


# %% [code]
#rna = rna_te
#atac = atac_te
rna = rna_raw
atac = atac_raw


# %% [code]
Z_rna = encode_adata(
    model,
    adata=rna,
    modality="rna",
    device=device,
    layer=None,          # uses adata.X by default
    X_key="X",
    batch_size=1024,
    latent="moe_mean",   # {"moe_mean","moe_sample","modality_mean","modality_sample"}
    random_state=0,
)
rna.obsm["X_univi"] = Z_rna


# %% [code]
Z_atac = encode_adata(
    model,
    adata=atac,
    modality="atac",
    device=device,
    layer=None,          # uses adata.X by default
    X_key="X",
    batch_size=1024,
    latent="moe_mean",   # {"moe_mean","moe_sample","modality_mean","modality_sample"}
    random_state=0,
)
atac.obsm["X_univi"] = Z_atac


# %% [code]
umap(
    rna,
    obsm_key="X_univi",
    color=["cell_type"],
    legend="outside",
    legend_subset_topk=25,
    #savepath="umap_rna_univi.png",
    savepath=None,
    size=8,
    show=True,
)


# %% [code]
umap(
    atac,
    obsm_key="X_univi",
    color=["cell_type"],
    legend="outside",
    legend_subset_topk=25,
    #savepath="umap_rna_univi.png",
    savepath=None,
    size=8,
    show=True,
)


# %% [code]
fused = encode_fused_adata_pair(
    model,
    adata_by_mod={"rna": rna, "atac": atac},   # same obs_names, same order
    device=device,
    batch_size=1024,
    use_mean=True,
    return_gates=True,
    return_gate_logits=True,
    write_to_adatas=True,                   # writes obsm + gate columns
    fused_obsm_key="X_univi_fused",
    gate_prefix="gate",
)

# fused["Z_fused"] -> (n_cells, latent_dim)
# fused["gates"]  -> (n_cells, n_modalities) or None (if fused transformer posterior is used)


# %% [code]
umap(
    rna,
    obsm_key="X_univi_fused",
    color=["cell_type"],
    legend="outside",
    #savepath="umap_fused.png",
    savepath=None,
    size=8,
    show=True,
)


# %% [code]
from univi.plotting import umap_by_modality

# make sure both adatas have the fused embedding (encode_fused_adata_pair with write_to_adatas=True does this)
# rna.obsm["X_univi_fused"]
# adt.obsm["X_univi_fused"]

umap_by_modality(
    {"rna": rna, "atac": atac},
    obsm_key="X_univi_fused",
    color=["univi_modality", "cell_type"],
    legend="outside",
    size=8,
    show=True,
    savepath=None,
)


# %% [code]
atac_hat_from_rna = cross_modal_predict(
    model,
    adata_src=rna,
    src_mod="rna",
    tgt_mod="atac",
    device=device,
    layer=None,
    X_key="X",
    batch_size=512,
    use_moe=True,
)
atac.layers["imputed_from_rna"] = atac_hat_from_rna


# %% [code]
rna_hat_from_atac = cross_modal_predict(
    model,
    adata_src=atac,
    src_mod="atac",
    tgt_mod="rna",
    device=device,
    layer=None,
    X_key="X",
    batch_size=512,
    use_moe=True,
)
rna.layers["imputed_from_atac"] = rna_hat_from_atac


# %% [code]
denoise_adata(
    model,
    adata=rna,                         # output written here
    modality="rna",
    device=device,
    out_layer="denoised_fused",
    overwrite_X=False,
    batch_size=512,
    adata_by_mod={"rna": rna, "atac": atac},
    layer_by_mod={"rna": None, "atac": None},  # None -> use .X
    X_key_by_mod={"rna": "X", "atac": "X"},
    use_mean=True,
)


# %% [code]
compare_raw_vs_denoised_umap_features(
    rna,
    #obsm_key="X_univi",
    obsm_key="X_univi_fused",
    features=["MS4A1", "CD3D", "NKG7", "CD8A", "CD4"],
    raw_layer=None,
    denoised_layer="denoised_fused",
    #savepath="umap_raw_vs_denoised.png",
    show=True,
)


# %% [code]
denoise_adata(
    model,
    adata=atac,                         # output written here
    modality="atac",
    device=device,
    out_layer="denoised_fused",
    overwrite_X=False,
    batch_size=512,
    adata_by_mod={"rna": rna, "atac": atac},
    layer_by_mod={"rna": None, "atac": None},  # None -> use .X
    X_key_by_mod={"rna": "X", "atac": "X"},
    use_mean=True,
)


# %% [code]
print(atac.var_names[40000:40100])

# %% [code]
compare_raw_vs_denoised_umap_features(
    atac,
    obsm_key="X_univi",
    features=['chr1:1380624-1381248', 'chr1:1134636-1138947', 'chr1:983870-984728', 'chr11:121366986-121368317', 'chr17:3688002-3689045'],
    raw_layer=None,
    denoised_layer="denoised_fused",
    #savepath="umap_raw_vs_denoised.png",
    savepath=None,
    show=True,
)


# %% [code]
from univi.evaluation import to_dense, encode_moe_gates_from_tensors
from univi.plotting import write_gates_to_obs, plot_moe_gate_summary

gate = encode_moe_gates_from_tensors(
    model,
    x_dict={"rna": to_dense(rna.X), "atac": to_dense(atac.X)},
    device=device,
    batch_size=1024,
    modality_order=["rna", "atac"],
    kind="router_x_precision",  # will fall back to "effective_precision" if router logits are unavailable
    return_logits=True,
)

W    = gate["weights"]         # (n_cells, n_modalities), rows sum to 1
mods = gate["modality_order"]  # e.g. ["rna", "adt"]

print("Requested kind:", gate.get("requested_kind"))
print("Effective kind:", gate.get("kind"))
print("Per-modality mean:", gate.get("per_modality_mean"))
print("Has logits:", gate.get("logits") is not None)


# %% [code]
write_gates_to_obs(
    rna,
    gates=W,
    modality_names=mods,
    gate_prefix="moe_gate",          # creates obs cols: moe_gate_{mod}
    gate_logits=gate.get("logits"),  # optional; may be None
)


# %% [code]
plot_moe_gate_summary(
    rna,
    gate_prefix="moe_gate",
    groupby="cell_type",           # or "celltype.l2", "batch", etc.
    agg="mean",
    #savepath="moe_gates_by_celltype.png",
    show=False,
)


# %% [code]
print(rna)
print(atac)


# %% [code]
metrics = evaluate_alignment(
    Z1=rna.obsm["X_univi"],
    Z2=atac.obsm["X_univi"],
    metric="euclidean",
    recall_ks=(1, 5, 10, 25, 50, 100),
    k_mixing=20,
    k_entropy=30,
    labels_source=rna.obs["cell_type"].to_numpy(),
    labels_target=atac.obs["cell_type"].to_numpy(),
    compute_bidirectional_transfer=True,
    k_transfer=3,
    json_safe=True,
)

# %% [code]
plot_confusion_matrix(
    np.asarray(metrics["label_transfer_cm"]),
    labels=np.asarray(metrics["label_transfer_label_order"]),
    title="Label transfer (RNA → ATAC)",
    normalize=None,
    #savepath="label_transfer_confusion.png",
    show=False,
)


# %% [code]
plot_confusion_matrix(
    np.asarray(metrics["label_transfer_cm"]),
    labels=np.asarray(metrics["label_transfer_label_order"]),
    title="Label transfer (RNA → ATAC)",
    normalize="true",
    #savepath="label_transfer_confusion.png",
    show=False,
)


# %% [code]
#print(metrics)
print(f"FOSCTTM mean:          {metrics['foscttm_mean']}")
print(f"FOSCTTM sem:           {metrics['foscttm_sem']}")

print(f"Recall@1:              {metrics['recall_at_k']['1']['mean']}")
print(f"Recall@5:              {metrics['recall_at_k']['5']['mean']}")
print(f"Recall@10:             {metrics['recall_at_k']['10']['mean']}")
print(f"Recall@25:             {metrics['recall_at_k']['25']['mean']}")
print(f"Recall@50:             {metrics['recall_at_k']['50']['mean']}")
print(f"Recall@100:            {metrics['recall_at_k']['100']['mean']}")

print(f"Modality mixing mean:  {metrics['modality_mixing_mean']}")
print(f"Modality entropy mean: {metrics['modality_entropy_mean']}")
print(f"Label transfer ACC:    {metrics['label_transfer_acc']}")


# %% [code]


# %% [code]
import re
import gzip
from pathlib import Path
from typing import Optional, Iterable, Dict, List, Tuple, Union

import numpy as np
import pandas as pd
import scipy.sparse as sp
import anndata as ad
import matplotlib.pyplot as plt


# %% [code]
# -----------------------------
# Basic matrix helpers
# -----------------------------
def _to_dense(X):
    return X.toarray() if sp.issparse(X) else np.asarray(X)

def _row_to_dense(X, i: int):
    x = X[i]
    return x.toarray().ravel() if sp.issparse(x) else np.asarray(x).ravel()

def _ensure_float32(x):
    return np.asarray(x, dtype=np.float32)

def _copy_matrix_like(X):
    return X.copy() if sp.issparse(X) else np.array(X, copy=True)


# %% [code]
# -----------------------------
# Genomic coordinate parsing helpers
# -----------------------------
_coord_pat = re.compile(r"^(chr[^:]+):(\d+)-(\d+)$")

def parse_peak_name(s: str):
    """
    Parse 'chr:start-end' peak names. Returns (chrom, start, end) or None.
    """
    m = _coord_pat.match(str(s))
    if m is None:
        return None
    chrom, st, en = m.group(1), int(m.group(2)), int(m.group(3))
    return chrom, st, en

def ensure_atac_peak_coords(atac: ad.AnnData,
                            chrom_col: str = "chrom",
                            start_col: str = "chromStart",
                            end_col: str = "chromEnd") -> ad.AnnData:
    """
    Ensure atac.var has chrom/start/end columns.
    If missing, try parsing atac.var_names like 'chr1:100-200'.
    """
    v = atac.var.copy()

    have_cols = all(c in v.columns for c in [chrom_col, start_col, end_col])
    if have_cols:
        # normalize dtypes
        v[chrom_col] = v[chrom_col].astype(str)
        v[start_col] = v[start_col].astype(int)
        v[end_col] = v[end_col].astype(int)
        atac.var = v
        return atac

    parsed = [parse_peak_name(x) for x in atac.var_names.astype(str)]
    ok = np.array([p is not None for p in parsed], dtype=bool)
    if not ok.all():
        bad_n = (~ok).sum()
        raise ValueError(
            f"Could not parse {bad_n} ATAC peak names from atac.var_names. "
            f"Need columns [{chrom_col}, {start_col}, {end_col}] or var_names like chr:start-end."
        )

    v[chrom_col] = [p[0] for p in parsed]
    v[start_col] = [p[1] for p in parsed]
    v[end_col]   = [p[2] for p in parsed]
    atac.var = v
    return atac


# %% [code]
# -----------------------------
# GTF parser (gene entries) -> gene table with TSS (robust)
# -----------------------------
def _parse_gtf_attributes(attr_str: str) -> Dict[str, str]:
    out = {}
    for field in attr_str.strip().split(";"):
        field = field.strip()
        if not field:
            continue
        if " " not in field:
            continue
        k, v = field.split(" ", 1)
        out[k] = v.strip().strip('"')
    return out

def read_gtf_gene_table(gtf_path, feature_type: str = "gene", keep_biotypes=None) -> pd.DataFrame:
    """
    Parse GTF gene rows and return columns:
      chrom, start, end, strand, gene_id, gene_name, gene_biotype, tss
    Robust to missing gene_name (falls back to gene_id).
    """
    gtf_path = str(gtf_path)
    opener = gzip.open if gtf_path.endswith(".gz") else open

    rows = []
    with opener(gtf_path, "rt") as f:
        for line in f:
            if not line or line.startswith("#"):
                continue
            parts = line.rstrip("\n").split("\t")
            if len(parts) != 9:
                continue

            chrom, source, feat, start, end, score, strand, frame, attrs = parts
            if feat != feature_type:
                continue

            a = _parse_gtf_attributes(attrs)

            # Common alternatives across GTF/GFF-ish annotations
            gene_id = (
                a.get("gene_id")
                or a.get("ID")
                or a.get("gene")
                or a.get("Name")
            )
            if gene_id is None:
                continue

            gene_name = (
                a.get("gene_name")
                or a.get("gene_symbol")
                or a.get("Name")
                or a.get("gene")
                or gene_id
            )

            gene_biotype = (
                a.get("gene_biotype")
                or a.get("gene_type")
                or a.get("biotype")
                or None
            )

            start_i = int(start)
            end_i = int(end)
            tss = start_i if strand == "+" else end_i

            rows.append({
                "chrom": str(chrom),
                "start": start_i,
                "end": end_i,
                "strand": strand,
                "gene_id": str(gene_id),
                "gene_name": str(gene_name),  # always exists now
                "gene_biotype": gene_biotype,
                "tss": int(tss),
            })

    genes = pd.DataFrame(rows)
    if genes.empty:
        raise ValueError(f"No '{feature_type}' entries parsed from {gtf_path}")

    # Clean gene_id versions (optional but usually helpful)
    genes["gene_id"] = genes["gene_id"].astype(str).str.replace(r"\.\d+$", "", regex=True)
    genes["gene_name"] = genes["gene_name"].astype(str)

    if keep_biotypes is not None and "gene_biotype" in genes.columns:
        keep_biotypes = set(keep_biotypes)
        genes = genes[genes["gene_biotype"].isin(keep_biotypes)].copy()

    # Keep one row per gene_id (first occurrence)
    genes = genes.drop_duplicates(subset=["gene_id"]).reset_index(drop=True)
    return genes


# %% [code]
# -----------------------------
# Map RNA features to GTF genes
# -----------------------------
def annotate_rna_var_with_gtf(
    rna: ad.AnnData,
    genes_gtf: pd.DataFrame,
    rna_gene_id_col: Optional[str] = "gene_id",
    rna_gene_name_col: Optional[str] = None,  # if None, use rna.var_names
) -> ad.AnnData:
    """
    Add/overwrite RNA var columns:
      gene_id, gene_name, chrom, start, end, strand, tss
    using GTF matching by gene_id first, then gene_name.
    Robust to missing gene_name in genes_gtf (falls back to gene_id).
    """
    v = rna.var.copy()

    # RNA-side queries
    if rna_gene_id_col is not None and rna_gene_id_col in v.columns:
        v["_gene_id_query"] = v[rna_gene_id_col].astype(str).str.replace(r"\.\d+$", "", regex=True)
    else:
        v["_gene_id_query"] = pd.Series([None] * v.shape[0], index=v.index, dtype="object")

    if rna_gene_name_col is not None and rna_gene_name_col in v.columns:
        v["_gene_name_query"] = v[rna_gene_name_col].astype(str)
    else:
        v["_gene_name_query"] = v.index.astype(str)  # fallback to var_names

    g = genes_gtf.copy()

    # Ensure required columns exist in GTF table
    if "gene_id" not in g.columns:
        raise KeyError("genes_gtf is missing required column 'gene_id'")
    if "gene_name" not in g.columns:
        g["gene_name"] = g["gene_id"].astype(str)  # fallback
    for c in ["chrom", "start", "end", "strand", "tss"]:
        if c not in g.columns:
            raise KeyError(f"genes_gtf is missing required column '{c}'")

    g["gene_id"] = g["gene_id"].astype(str).str.replace(r"\.\d+$", "", regex=True)
    g["gene_name"] = g["gene_name"].astype(str)

    g_by_id = g.drop_duplicates("gene_id").set_index("gene_id")
    g_by_name = g.drop_duplicates("gene_name").set_index("gene_name")

    matched = []
    for _, row in v.iterrows():
        rec = None
        gid = row["_gene_id_query"]
        gname = row["_gene_name_query"]

        if gid is not None and gid in g_by_id.index:
            rec = g_by_id.loc[gid]
        elif gname in g_by_name.index:
            rec = g_by_name.loc[gname]

        if rec is None:
            matched.append({
                "gene_id": gid if gid is not None else None,
                "gene_name": gname if gname is not None else None,
                "chrom": None,
                "start": np.nan,
                "end": np.nan,
                "strand": None,
                "tss": np.nan,
            })
        else:
            if isinstance(rec, pd.DataFrame):
                rec = rec.iloc[0]
            rec_gene_id = str(rec["gene_id"]) if "gene_id" in rec.index else (str(gid) if gid is not None else None)
            rec_gene_name = str(rec["gene_name"]) if "gene_name" in rec.index else rec_gene_id

            matched.append({
                "gene_id": rec_gene_id,
                "gene_name": rec_gene_name,
                "chrom": str(rec["chrom"]) if pd.notna(rec["chrom"]) else None,
                "start": int(rec["start"]) if pd.notna(rec["start"]) else np.nan,
                "end": int(rec["end"]) if pd.notna(rec["end"]) else np.nan,
                "strand": str(rec["strand"]) if pd.notna(rec["strand"]) else None,
                "tss": int(rec["tss"]) if pd.notna(rec["tss"]) else np.nan,
            })

    ann = pd.DataFrame(matched, index=v.index)

    for col in ann.columns:
        v[col] = ann[col].values

    v = v.drop(columns=[c for c in ["_gene_id_query", "_gene_name_query"] if c in v.columns])
    rna.var = v
    return rna


# %% [code]
# REQUIRED: point this to your annotation
GTF_PATH = "./data/Homo_sapiens_GRCh38.p13.gencode.annotation.gtf"

# Example filter: keep protein-coding + lncRNA (customize)
genes_gtf = read_gtf_gene_table(
    GTF_PATH,
    feature_type="gene",
    keep_biotypes=None,
)

print(genes_gtf.columns.tolist())
display(genes_gtf.head())


# %% [code]
# normalize RNA-side gene_id once
if "gene_id" not in rna.var.columns:
    rna.var["gene_id"] = rna.var["gene_ids"].astype(str)
rna.var["gene_id"] = rna.var["gene_id"].astype(str).str.replace(r"\.\d+$", "", regex=True)

rna = annotate_rna_var_with_gtf(
    rna,
    genes_gtf,
    rna_gene_id_col="gene_id",      # now normalized
    rna_gene_name_col="gene_name",  # you do have this
)

print("RNA gene coords mapped:", rna.var["tss"].notna().sum(), "/", rna.n_vars)
display(rna.var[["gene_id","gene_name","chrom","start","end","strand","tss"]].head())


# %% [code]
# -----------------------------
# Peak <-> gene promoter mapping
# -----------------------------
def build_promoter_peak_gene_links(
    atac: ad.AnnData,
    rna: ad.AnnData,
    promoter_upstream: int = 2000,
    promoter_downstream: int = 500,
    peak_chrom_col: str = "chrom",
    peak_start_col: str = "chromStart",
    peak_end_col: str = "chromEnd",
) -> pd.DataFrame:
    """
    Returns long-form table of promoter-linked peak-gene pairs:
      peak_idx, peak_name, gene_idx, gene_name, gene_id, link_type, distance_to_tss
    Overlap criterion: peak overlaps [TSS-upstream, TSS+downstream] (strand-aware upstream/downstream)
    """
    vA = atac.var.copy()
    vR = rna.var.copy()

    need_rna_cols = ["chrom", "tss", "strand"]
    for c in need_rna_cols:
        if c not in vR.columns:
            raise ValueError(f"rna.var missing required column '{c}'. Run GTF annotation first.")

    # valid RNA genes with coords
    gene_mask = vR["chrom"].notna() & vR["tss"].notna() & vR["strand"].notna()
    vR2 = vR.loc[gene_mask].copy()

    # peak arrays
    p_ch = vA[peak_chrom_col].astype(str).values
    p_st = vA[peak_start_col].astype(int).values
    p_en = vA[peak_end_col].astype(int).values
    p_nm = atac.var_names.astype(str).values

    rows = []
    # group genes by chromosome to reduce work
    for chrom, gsub in vR2.groupby(vR2["chrom"].astype(str)):
        p_idx_chr = np.where(p_ch == str(chrom))[0]
        if p_idx_chr.size == 0:
            continue

        pst = p_st[p_idx_chr]
        pen = p_en[p_idx_chr]

        for g_idx, g in gsub.iterrows():
            tss = int(g["tss"])
            strand = str(g["strand"])

            if strand == "+":
                prom_start = tss - promoter_upstream
                prom_end   = tss + promoter_downstream
            else:
                # upstream/downstream reversed on negative strand
                prom_start = tss - promoter_downstream
                prom_end   = tss + promoter_upstream

            # overlap peak with promoter interval
            hit = (pen >= prom_start) & (pst <= prom_end)
            hit_idx_local = np.where(hit)[0]
            if hit_idx_local.size == 0:
                continue

            g_idx_pos = rna.var_names.get_loc(g_idx) if g_idx in rna.var_names else np.where(rna.var_names == g_idx)[0][0]

            for jloc in hit_idx_local:
                pi = int(p_idx_chr[jloc])
                # distance of peak midpoint to TSS
                pmid = 0.5 * (p_st[pi] + p_en[pi])
                d_tss = int(round(pmid - tss))
                rows.append({
                    "peak_idx": pi,
                    "peak_name": p_nm[pi],
                    "gene_idx": int(g_idx_pos),
                    "gene_var_name": str(g_idx),
                    "gene_id": str(g.get("gene_id", "")) if pd.notna(g.get("gene_id", None)) else None,
                    "gene_name": str(g.get("gene_name", g_idx)),
                    "chrom": str(chrom),
                    "link_type": "promoter",
                    "distance_to_tss": d_tss,
                    "weight": 1.0,
                })

    links = pd.DataFrame(rows)
    if links.empty:
        return links

    # de-dup if any
    links = links.drop_duplicates(subset=["peak_idx", "gene_idx", "link_type"]).reset_index(drop=True)
    return links


# %% [code]
# -----------------------------
# Optional: enhancer <-> gene mapping from external table
# -----------------------------
def build_enhancer_peak_gene_links_from_table(
    atac: ad.AnnData,
    rna: ad.AnnData,
    enhancer_gene_df: pd.DataFrame,
    enh_chrom_col: str = "chrom",
    enh_start_col: str = "start",
    enh_end_col: str = "end",
    gene_name_col: Optional[str] = "gene_name",
    gene_id_col: Optional[str] = None,
    score_col: Optional[str] = None,
    peak_chrom_col: str = "chrom",
    peak_start_col: str = "chromStart",
    peak_end_col: str = "chromEnd",
) -> pd.DataFrame:
    """
    Overlap ATAC peaks with enhancer intervals and map to RNA genes by gene_name or gene_id.
    Returns same schema as promoter links with link_type='enhancer'.
    """
    vA = atac.var.copy()
    vR = rna.var.copy()

    # peak arrays
    p_ch = vA[peak_chrom_col].astype(str).values
    p_st = vA[peak_start_col].astype(int).values
    p_en = vA[peak_end_col].astype(int).values
    p_nm = atac.var_names.astype(str).values

    # gene lookup
    gene_name_to_idx = {}
    gene_id_to_idx = {}
    for i, (vn, row) in enumerate(vR.iterrows()):
        gname = str(row["gene_name"]) if "gene_name" in vR.columns and pd.notna(row["gene_name"]) else str(vn)
        gid = str(row["gene_id"]) if "gene_id" in vR.columns and pd.notna(row["gene_id"]) else None
        gene_name_to_idx.setdefault(gname, i)
        if gid is not None:
            gene_id_to_idx.setdefault(gid, i)

    rows = []
    df = enhancer_gene_df.copy()
    for c in [enh_chrom_col, enh_start_col, enh_end_col]:
        if c not in df.columns:
            raise ValueError(f"enhancer_gene_df missing column '{c}'")

    for _, r in df.iterrows():
        chrom = str(r[enh_chrom_col])
        st = int(r[enh_start_col])
        en = int(r[enh_end_col])

        # resolve target gene
        g_idx = None
        gene_name = None
        gene_id = None
        if gene_id_col is not None and gene_id_col in df.columns and pd.notna(r.get(gene_id_col, None)):
            gene_id = str(r[gene_id_col])
            g_idx = gene_id_to_idx.get(gene_id, None)
        if g_idx is None and gene_name_col is not None and gene_name_col in df.columns and pd.notna(r.get(gene_name_col, None)):
            gene_name = str(r[gene_name_col])
            g_idx = gene_name_to_idx.get(gene_name, None)

        if g_idx is None:
            continue

        if gene_name is None:
            rv = vR.iloc[g_idx]
            gene_name = str(rv["gene_name"]) if "gene_name" in vR.columns and pd.notna(rv.get("gene_name", None)) else str(vR.index[g_idx])
        if gene_id is None:
            rv = vR.iloc[g_idx]
            gene_id = str(rv["gene_id"]) if "gene_id" in vR.columns and pd.notna(rv.get("gene_id", None)) else None

        # overlap peaks
        mask_chr = (p_ch == chrom)
        if not np.any(mask_chr):
            continue
        idx_chr = np.where(mask_chr)[0]
        hit = (p_en[idx_chr] >= st) & (p_st[idx_chr] <= en)
        hit_idx = idx_chr[np.where(hit)[0]]

        if hit_idx.size == 0:
            continue

        w = float(r[score_col]) if (score_col is not None and score_col in df.columns and pd.notna(r.get(score_col, None))) else 1.0

        # optional TSS distance if available
        tss = None
        if "tss" in vR.columns and pd.notna(vR.iloc[g_idx].get("tss", np.nan)):
            tss = int(vR.iloc[g_idx]["tss"])

        for pi in hit_idx:
            pmid = 0.5 * (p_st[pi] + p_en[pi])
            d_tss = int(round(pmid - tss)) if tss is not None else np.nan
            rows.append({
                "peak_idx": int(pi),
                "peak_name": p_nm[pi],
                "gene_idx": int(g_idx),
                "gene_var_name": str(vR.index[g_idx]),
                "gene_id": gene_id,
                "gene_name": gene_name,
                "chrom": chrom,
                "link_type": "enhancer",
                "distance_to_tss": d_tss,
                "weight": w,
            })

    links = pd.DataFrame(rows)
    if links.empty:
        return links

    links = links.drop_duplicates(subset=["peak_idx", "gene_idx", "link_type"]).reset_index(drop=True)
    return links


# %% [code]
# Build promoter links
prom_links = build_promoter_peak_gene_links(
    atac, rna,
    promoter_upstream=10000,
    promoter_downstream=2000,
)

print("Promoter links:", prom_links.shape)
display(prom_links.head())


# %% [code]
# OPTIONAL enhancer links (only if you have an enhancer-gene map)
# enhancer_df = pd.read_csv("/path/to/enhancer_gene_links.tsv", sep="\t")
# enh_links = build_enhancer_peak_gene_links_from_table(
#     atac, rna, enhancer_df,
#     enh_chrom_col="chrom", enh_start_col="start", enh_end_col="end",
#     gene_name_col="gene_name",  # or gene_id_col="gene_id"
#     score_col=None,
# )
# print("Enhancer links:", enh_links.shape)
# display(enh_links.head())

# If no enhancer file, use empty table for now
enh_links = pd.DataFrame(columns=prom_links.columns if prom_links is not None and not prom_links.empty else [
    "peak_idx","peak_name","gene_idx","gene_var_name","gene_id","gene_name","chrom","link_type","distance_to_tss","weight"
])

# Combine
peak_gene_links = pd.concat([prom_links, enh_links], axis=0, ignore_index=True)
if not peak_gene_links.empty:
    # keep max weight per (peak,gene,link_type)
    peak_gene_links = (
        peak_gene_links.sort_values(["peak_idx","gene_idx","link_type","weight"], ascending=[True, True, True, False])
        .drop_duplicates(subset=["peak_idx","gene_idx","link_type"])
        .reset_index(drop=True)
    )

print("Combined peak-gene links:", peak_gene_links.shape)
display(peak_gene_links.head(20))


# %% [code]
# -----------------------------
# Gene set -> peak selection
# -----------------------------
def _normalize_gene_list(gene_list: Iterable[str]) -> List[str]:
    return [str(g).strip() for g in gene_list if str(g).strip()]

def select_peaks_for_gene_set(
    peak_gene_links: pd.DataFrame,
    gene_set: Iterable[str],
    match_on: str = "gene_name",     # "gene_name" or "gene_id"
    include_link_types: Tuple[str, ...] = ("promoter", "enhancer"),
    dedup: bool = True,
) -> Dict[str, Union[np.ndarray, pd.DataFrame, List[str]]]:
    """
    Returns peak indices and metadata for peaks linked to genes in gene_set.
    """
    if peak_gene_links is None or peak_gene_links.empty:
        return {"peak_idx": np.array([], dtype=int), "links": pd.DataFrame(), "genes_found": [], "genes_missing": list(_normalize_gene_list(gene_set))}

    genes = _normalize_gene_list(gene_set)
    if match_on not in ("gene_name", "gene_id"):
        raise ValueError("match_on must be 'gene_name' or 'gene_id'")

    df = peak_gene_links.copy()
    df = df[df["link_type"].isin(include_link_types)].copy()

    # exact matching
    q = set(genes)
    mask = df[match_on].astype(str).isin(q)
    sub = df.loc[mask].copy()

    genes_found = sorted(sub[match_on].dropna().astype(str).unique().tolist())
    genes_missing = sorted(list(q - set(genes_found)))

    if sub.empty:
        return {"peak_idx": np.array([], dtype=int), "links": sub, "genes_found": genes_found, "genes_missing": genes_missing}

    if dedup:
        peak_idx = np.sort(sub["peak_idx"].astype(int).unique())
    else:
        peak_idx = sub["peak_idx"].astype(int).values

    return {
        "peak_idx": peak_idx,
        "links": sub.reset_index(drop=True),
        "genes_found": genes_found,
        "genes_missing": genes_missing,
    }


# %% [code]
# Example gene set (replace with your set)
gene_set = ["IL7R", "LTB", "MALAT1"]  # just example symbols

sel = select_peaks_for_gene_set(
    peak_gene_links,
    gene_set,
    match_on="gene_name",
    include_link_types=("promoter", "enhancer"),  # change to ("promoter",) if you want strict promoter-only
)

print("Genes found:", sel["genes_found"])
print("Genes missing:", sel["genes_missing"])
print("Selected peaks:", len(sel["peak_idx"]))
display(sel["links"].head(30))


# %% [code]
# -----------------------------
# Create perturbed ATAC AnnData from existing cells
# -----------------------------
def _get_atac_input_matrix(atac: ad.AnnData, layer: Optional[str] = None):
    if layer is None or layer == "X":
        return atac.X
    if layer not in atac.layers:
        raise KeyError(f"Layer '{layer}' not found in atac.layers")
    return atac.layers[layer]

def _set_atac_input_matrix(atac_new: ad.AnnData, X_new, layer: Optional[str] = None):
    if layer is None or layer == "X":
        atac_new.X = X_new
    else:
        atac_new.layers[layer] = X_new

def subset_cells_to_adata(atac: ad.AnnData, cell_idx: Union[int, np.ndarray, List[int]]) -> ad.AnnData:
    if np.isscalar(cell_idx):
        return atac[[int(cell_idx)]].copy()
    idx = np.asarray(cell_idx, dtype=int)
    return atac[idx].copy()

def perturb_atac_peaks(
    atac_src: ad.AnnData,
    peak_idx: Union[np.ndarray, List[int]],
    *,
    cell_idx: Optional[Union[int, List[int], np.ndarray]] = None,
    input_layer: Optional[str] = None,  # the layer used for ATAC->RNA prediction
    mode: str = "set",                  # "set", "off", "on", "scale", "add"
    value: float = 0.0,                 # used for set/scale/add
    clip_min: Optional[float] = 0.0,
    clip_max: Optional[float] = 1.0,
    keep_other_layers: bool = True,
) -> Tuple[ad.AnnData, ad.AnnData]:
    """
    Returns (baseline_adata, perturbed_adata), same cells/features, perturbation applied to selected peaks only.

    mode:
      - 'off'   -> set selected peaks to 0
      - 'on'    -> set selected peaks to 1
      - 'set'   -> set selected peaks to `value`
      - 'scale' -> selected peaks *= value
      - 'add'   -> selected peaks += value
    """
    peak_idx = np.unique(np.asarray(peak_idx, dtype=int))
    if peak_idx.size == 0:
        raise ValueError("peak_idx is empty; no peaks selected for perturbation")

    if cell_idx is None:
        base = atac_src.copy()
    else:
        base = subset_cells_to_adata(atac_src, cell_idx)

    pert = base.copy()

    Xb = _get_atac_input_matrix(base, layer=input_layer)
    Xp = _get_atac_input_matrix(pert, layer=input_layer)

    # Convert to editable format
    if sp.issparse(Xp):
        # LIL is convenient for column assignment in subsets; convert back after
        Xp_edit = Xp.tolil(copy=True)
        Xb_view = Xb  # baseline not edited
        is_sparse = True
    else:
        Xp_edit = np.array(Xp, copy=True)
        Xb_view = Xb
        is_sparse = False

    # Apply perturbation
    if mode == "off":
        set_val = 0.0
        if is_sparse:
            Xp_edit[:, peak_idx] = set_val
        else:
            Xp_edit[:, peak_idx] = set_val

    elif mode == "on":
        set_val = 1.0
        if is_sparse:
            Xp_edit[:, peak_idx] = set_val
        else:
            Xp_edit[:, peak_idx] = set_val

    elif mode == "set":
        set_val = float(value)
        if is_sparse:
            Xp_edit[:, peak_idx] = set_val
        else:
            Xp_edit[:, peak_idx] = set_val

    elif mode == "scale":
        if is_sparse:
            # operate row-by-row for sparse LIL; easiest is dense slice if small, else CSR multiply
            Xtmp = Xp_edit.tocsr(copy=True)
            Xtmp[:, peak_idx] = Xtmp[:, peak_idx].multiply(float(value))
            Xp_edit = Xtmp.tolil(copy=True)
        else:
            Xp_edit[:, peak_idx] *= float(value)

    elif mode == "add":
        if is_sparse:
            Xtmp = Xp_edit.tocsr(copy=True)
            Xtmp[:, peak_idx] = Xtmp[:, peak_idx] + float(value)
            Xp_edit = Xtmp.tolil(copy=True)
        else:
            Xp_edit[:, peak_idx] += float(value)
    else:
        raise ValueError("mode must be one of {'set','off','on','scale','add'}")

    # Clip if desired
    if clip_min is not None or clip_max is not None:
        if is_sparse:
            Xtmp = Xp_edit.tocsr(copy=True)
            if Xtmp.nnz > 0:
                data = Xtmp.data
                if clip_min is not None:
                    data = np.maximum(data, clip_min)
                if clip_max is not None:
                    data = np.minimum(data, clip_max)
                Xtmp.data = data
                Xtmp.eliminate_zeros()
            Xp_new = Xtmp
        else:
            Xp_new = Xp_edit
            if clip_min is not None:
                Xp_new = np.maximum(Xp_new, clip_min)
            if clip_max is not None:
                Xp_new = np.minimum(Xp_new, clip_max)
    else:
        Xp_new = Xp_edit.tocsr(copy=True) if is_sparse else Xp_edit

    # Write back
    _set_atac_input_matrix(pert, Xp_new, layer=input_layer)

    return base, pert


# %% [code]
# -----------------------------
# Run ATAC->RNA predictions before/after perturbation
# -----------------------------
def predict_rna_from_atac_adata(
    model,
    atac_in: ad.AnnData,
    *,
    device=None,
    atac_input_layer: Optional[str] = None,
    batch_size: int = 256,
    use_moe: bool = True,
):
    """
    Thin wrapper around your existing cross_modal_predict function.
    Returns dense ndarray [n_cells, n_genes] (or whatever cross_modal_predict returns converted to array).
    """
    y = cross_modal_predict(
        model,
        adata_src=atac_in,
        src_mod="atac",
        tgt_mod="rna",
        device=device,
        layer=atac_input_layer,
        X_key="X",
        batch_size=batch_size,
        use_moe=use_moe,
    )
    return y

def run_atac_ablation_to_rna(
    model,
    atac: ad.AnnData,
    rna: ad.AnnData,
    peak_idx: Union[np.ndarray, List[int]],
    *,
    cell_idx: Optional[Union[int, List[int], np.ndarray]] = None,
    atac_input_layer: Optional[str] = None,
    mode: str = "off",
    value: float = 0.0,
    clip_min: Optional[float] = 0.0,
    clip_max: Optional[float] = 1.0,
    device=None,
    batch_size: int = 256,
    use_moe: bool = True,
    return_observed_rna: bool = True,
    rna_obs_layer: Optional[str] = None,
):
    base_atac, pert_atac = perturb_atac_peaks(
        atac,
        peak_idx,
        cell_idx=cell_idx,
        input_layer=atac_input_layer,
        mode=mode,
        value=value,
        clip_min=clip_min,
        clip_max=clip_max,
    )

    pred_base = np.asarray(
        predict_rna_from_atac_adata(
            model, base_atac,
            device=device,
            atac_input_layer=atac_input_layer,
            batch_size=batch_size,
            use_moe=use_moe
        )
    )
    pred_pert = np.asarray(
        predict_rna_from_atac_adata(
            model, pert_atac,
            device=device,
            atac_input_layer=atac_input_layer,
            batch_size=batch_size,
            use_moe=use_moe
        )
    )

    out = {
        "base_atac": base_atac,
        "pert_atac": pert_atac,
        "pred_base": pred_base,
        "pred_pert": pred_pert,
        "delta_pred": pred_pert - pred_base,
        "peak_idx": np.unique(np.asarray(peak_idx, dtype=int)),
        "mode": mode,
        "value": value,
    }

    if return_observed_rna:
        # align observed RNA rows to selected cells if needed
        if cell_idx is None:
            rna_sub = rna
        elif np.isscalar(cell_idx):
            rna_sub = rna[[int(cell_idx)]].copy()
        else:
            rna_sub = rna[np.asarray(cell_idx, dtype=int)].copy()

        Xo = rna_sub.X if (rna_obs_layer is None or rna_obs_layer == "X") else rna_sub.layers[rna_obs_layer]
        out["obs_rna"] = _to_dense(Xo)

    return out


# %% [code]
# -----------------------------
# Gene-level delta summaries
# -----------------------------
def _gene_name_series(rna: ad.AnnData) -> pd.Series:
    if "gene_name" in rna.var.columns:
        return rna.var["gene_name"].astype(str)
    return pd.Series(rna.var_names.astype(str), index=rna.var_names)

def gene_indices_from_list(
    rna: ad.AnnData,
    genes: Iterable[str],
    match_on: str = "gene_name",  # "gene_name" or "gene_id" or "var_names"
) -> np.ndarray:
    genes = [str(g) for g in genes]
    if match_on == "gene_name":
        if "gene_name" in rna.var.columns:
            s = rna.var["gene_name"].astype(str).values
        else:
            s = rna.var_names.astype(str).values
    elif match_on == "gene_id":
        if "gene_id" not in rna.var.columns:
            raise KeyError("rna.var['gene_id'] not found")
        s = rna.var["gene_id"].astype(str).values
    elif match_on == "var_names":
        s = rna.var_names.astype(str).values
    else:
        raise ValueError("match_on must be 'gene_name', 'gene_id', or 'var_names'")

    lookup = {}
    for i, x in enumerate(s):
        lookup.setdefault(x, i)

    idx = [lookup[g] for g in genes if g in lookup]
    return np.unique(np.asarray(idx, dtype=int))

def summarize_delta_by_gene(
    rna: ad.AnnData,
    pred_base: np.ndarray,
    pred_pert: np.ndarray,
    *,
    obs_rna: Optional[np.ndarray] = None,
    sort_by: str = "mean_delta_abs",  # or "mean_delta"
) -> pd.DataFrame:
    """
    pred_base/pred_pert shape [n_cells, n_genes]
    """
    pb = np.asarray(pred_base)
    pp = np.asarray(pred_pert)
    d = pp - pb

    gene_name = rna.var["gene_name"].astype(str).values if "gene_name" in rna.var.columns else rna.var_names.astype(str).values
    gene_id = rna.var["gene_id"].astype(str).values if "gene_id" in rna.var.columns else np.array([None] * rna.n_vars, dtype=object)

    df = pd.DataFrame({
        "gene_idx": np.arange(rna.n_vars, dtype=int),
        "gene_var_name": rna.var_names.astype(str).values,
        "gene_name": gene_name,
        "gene_id": gene_id,
        "mean_pred_base": pb.mean(axis=0),
        "mean_pred_pert": pp.mean(axis=0),
        "mean_delta": d.mean(axis=0),
        "mean_delta_abs": np.abs(d).mean(axis=0),
        "median_delta": np.median(d, axis=0),
        "frac_pos_delta": (d > 0).mean(axis=0),
        "frac_neg_delta": (d < 0).mean(axis=0),
    })

    if obs_rna is not None:
        o = np.asarray(obs_rna)
        if o.ndim == 2 and o.shape[1] == rna.n_vars:
            df["mean_obs"] = o.mean(axis=0)

    ascending = False
    if sort_by not in df.columns:
        sort_by = "mean_delta_abs"
    df = df.sort_values(sort_by, ascending=ascending).reset_index(drop=True)
    return df

def summarize_target_gene_set_response(
    rna: ad.AnnData,
    delta_pred: np.ndarray,
    gene_set: Iterable[str],
    match_on: str = "gene_name"
) -> pd.DataFrame:
    idx = gene_indices_from_list(rna, gene_set, match_on=match_on)
    if idx.size == 0:
        return pd.DataFrame(columns=["gene_idx","gene_name","gene_id","mean_delta","median_delta","mean_abs_delta"])

    d = np.asarray(delta_pred)[:, idx]
    sub = rna.var.iloc[idx].copy()

    out = pd.DataFrame({
        "gene_idx": idx,
        "gene_var_name": rna.var_names[idx].astype(str),
        "gene_name": sub["gene_name"].astype(str).values if "gene_name" in sub.columns else rna.var_names[idx].astype(str),
        "gene_id": sub["gene_id"].astype(str).values if "gene_id" in sub.columns else [None] * idx.size,
        "mean_delta": d.mean(axis=0),
        "median_delta": np.median(d, axis=0),
        "mean_abs_delta": np.abs(d).mean(axis=0),
        "frac_pos_delta": (d > 0).mean(axis=0),
        "frac_neg_delta": (d < 0).mean(axis=0),
    }).sort_values("mean_abs_delta", ascending=False).reset_index(drop=True)

    return out


# %% [code]
# -----------------------------
# Null peak sets matched by chromosome/width (optional but very useful)
# -----------------------------
def sample_matched_null_peak_sets(
    atac: ad.AnnData,
    target_peak_idx: np.ndarray,
    n_sets: int = 100,
    *,
    peak_chrom_col: str = "chrom",
    peak_start_col: str = "chromStart",
    peak_end_col: str = "chromEnd",
    match_chrom: bool = True,
    match_width: bool = True,
    random_state: int = 0,
) -> List[np.ndarray]:
    rng = np.random.default_rng(random_state)
    v = atac.var.copy()

    p_ch = v[peak_chrom_col].astype(str).values
    p_st = v[peak_start_col].astype(int).values
    p_en = v[peak_end_col].astype(int).values
    p_w = (p_en - p_st).astype(int)

    tgt = np.unique(np.asarray(target_peak_idx, dtype=int))
    all_idx = np.arange(atac.n_vars, dtype=int)

    out = []
    for _ in range(n_sets):
        chosen = []
        used = set(tgt.tolist())  # avoid picking actual target peaks
        for pi in tgt:
            cand = all_idx.copy()
            if match_chrom:
                cand = cand[p_ch[cand] == p_ch[pi]]
            if match_width:
                # exact width if possible, else nearest width bucket
                exact = cand[p_w[cand] == p_w[pi]]
                if exact.size > 0:
                    cand = exact
                else:
                    # width tolerance fallback
                    tol = max(25, int(round(0.1 * p_w[pi])))
                    near = cand[np.abs(p_w[cand] - p_w[pi]) <= tol]
                    if near.size > 0:
                        cand = near

            # remove target/used
            cand = np.array([x for x in cand if x not in used], dtype=int)
            if cand.size == 0:
                # fallback to anything not used/target
                cand = np.array([x for x in all_idx if x not in used], dtype=int)
            pick = int(rng.choice(cand))
            chosen.append(pick)
            used.add(pick)
        out.append(np.array(chosen, dtype=int))
    return out


# %% [code]
def null_test_target_gene_set_response(
    model,
    atac: ad.AnnData,
    rna: ad.AnnData,
    target_peak_idx: np.ndarray,
    target_gene_set: Iterable[str],
    *,
    cell_idx: Optional[Union[int, List[int], np.ndarray]] = None,
    atac_input_layer: Optional[str] = None,
    mode: str = "off",
    value: float = 0.0,
    n_null: int = 50,
    device=None,
    batch_size: int = 256,
    use_moe: bool = True,
    random_state: int = 0,
):
    # observed target perturbation
    res_target = run_atac_ablation_to_rna(
        model, atac, rna, target_peak_idx,
        cell_idx=cell_idx,
        atac_input_layer=atac_input_layer,
        mode=mode, value=value,
        device=device, batch_size=batch_size, use_moe=use_moe,
        return_observed_rna=False
    )
    tgt_df = summarize_target_gene_set_response(rna, res_target["delta_pred"], target_gene_set, match_on="gene_name")
    tgt_score = float(tgt_df["mean_abs_delta"].mean()) if not tgt_df.empty else np.nan

    # nulls
    null_sets = sample_matched_null_peak_sets(
        atac, np.asarray(target_peak_idx, dtype=int),
        n_sets=n_null, random_state=random_state
    )

    null_scores = []
    for ns in null_sets:
        res_null = run_atac_ablation_to_rna(
            model, atac, rna, ns,
            cell_idx=cell_idx,
            atac_input_layer=atac_input_layer,
            mode=mode, value=value,
            device=device, batch_size=batch_size, use_moe=use_moe,
            return_observed_rna=False
        )
        null_df = summarize_target_gene_set_response(rna, res_null["delta_pred"], target_gene_set, match_on="gene_name")
        score = float(null_df["mean_abs_delta"].mean()) if not null_df.empty else np.nan
        null_scores.append(score)

    null_scores = np.array(null_scores, dtype=float)
    p_emp = np.mean(null_scores >= tgt_score) if np.isfinite(tgt_score) else np.nan

    return {
        "target_result": res_target,
        "target_gene_summary": tgt_df,
        "target_score": tgt_score,
        "null_scores": null_scores,
        "empirical_p_ge": p_emp,
    }


# %% [code]
# -----------------------------
# Visualization helpers
# -----------------------------
def plot_top_rna_deltas(
    delta_gene_df: pd.DataFrame,
    top_n: int = 30,
    metric: str = "mean_delta_abs",
    title: Optional[str] = None,
):
    if delta_gene_df.empty:
        print("No gene delta summary to plot.")
        return
    d = delta_gene_df.head(top_n).copy()
    ylab = d["gene_name"].astype(str).values if "gene_name" in d.columns else d["gene_var_name"].astype(str).values
    x = d[metric].values

    plt.figure(figsize=(8, max(4, 0.25 * len(d))))
    plt.barh(np.arange(len(d)), x)
    plt.yticks(np.arange(len(d)), ylab, fontsize=8)
    plt.gca().invert_yaxis()
    plt.xlabel(metric)
    if title:
        plt.title(title)
    plt.tight_layout()
    plt.show()

def plot_null_distribution(null_scores: np.ndarray, target_score: float, title: str = "Null test"):
    x = np.asarray(null_scores, dtype=float)
    x = x[np.isfinite(x)]
    plt.figure(figsize=(6, 4))
    plt.hist(x, bins=20, alpha=0.8)
    plt.axvline(target_score, linewidth=2)
    plt.xlabel("Mean abs ΔRNA in target gene set")
    plt.ylabel("Count")
    plt.title(title)
    plt.tight_layout()
    plt.show()


# %% [code]
# ============================================================
# EXAMPLE: gene-set-guided ATAC ablation/activation -> RNA
# ============================================================

# 1) Define the gene set you want to perturb through chromatin
gene_set = ["IL7R", "LTB", "MALAT1"]  # replace with your real pathway / signature genes

# 2) Select linked peaks (promoter only or promoter+enhancer)
sel = select_peaks_for_gene_set(
    peak_gene_links,
    gene_set,
    match_on="gene_name",
    include_link_types=("promoter", "enhancer"),  # or ("promoter",)
)
target_peak_idx = sel["peak_idx"]

print(f"Genes found: {sel['genes_found']}")
print(f"Genes missing: {sel['genes_missing']}")
print(f"Target peaks selected: {len(target_peak_idx)}")
display(sel["links"].sort_values(["gene_name", "link_type"]).head(50))


# %% [code]
# 3) Choose cells to perturb
#    - single cell: cell_idx = 0
#    - subset by cell type: cell_idx = np.where(atac.obs['celltype.l2'] == 'Treg')[0]
#    - all cells: cell_idx = None
cell_idx = 23  # or an integer / array of row indices


# %% [code]
# 4) Run perturbation
#    If your ATAC->RNA model expects atac.layers["bin"], set atac_input_layer="bin"
#    If it expects atac.X, set atac_input_layer=None
res = run_atac_ablation_to_rna(
    model=model,
    atac=atac,
    rna=rna,
    peak_idx=target_peak_idx,
    cell_idx=cell_idx,
    #atac_input_layer="bin",   # <-- change if needed
    atac_input_layer=None,
    mode="on",                 # "off", "on", "set", "scale", "add"
    value=100.0,                # only used for set/scale/add
    clip_min=0.0,
    clip_max=10000.0,
    device=device,             # if you have `device`
    batch_size=256,
    use_moe=True,
    return_observed_rna=True,
    rna_obs_layer=None,        # or "counts" depending on what you want to compare against
)

print("pred_base shape:", res["pred_base"].shape)
print("pred_pert shape:", res["pred_pert"].shape)
print("delta_pred shape:", res["delta_pred"].shape)


# %% [code]
# 5) Summarize all genes and target genes
df_delta_all = summarize_delta_by_gene(
    rna,
    pred_base=res["pred_base"],
    pred_pert=res["pred_pert"],
    obs_rna=res.get("obs_rna", None),
    sort_by="mean_delta_abs",
)

df_delta_targets = summarize_target_gene_set_response(
    rna,
    delta_pred=res["delta_pred"],
    gene_set=gene_set,
    match_on="gene_name",
)

display(df_delta_targets.head(50))
display(df_delta_all.head(50))


# %% [code]
# 6) Plot top affected genes
plot_top_rna_deltas(
    df_delta_all,
    top_n=30,
    metric="mean_delta_abs",
    title=f"ATAC perturbation ({res['mode']}) -> top RNA changes"
)


# %% [code]
# 7) (Optional but recommended) Null test for specificity
null_out = null_test_target_gene_set_response(
    model=model,
    atac=atac,
    rna=rna,
    target_peak_idx=target_peak_idx,
    target_gene_set=gene_set,
    cell_idx=cell_idx,
    #atac_input_layer="bin",   # <-- match training input
    atac_input_layer=None,
    mode="on",
    value=100.0,
    n_null=25,                # start small for speed
    device=device,
    batch_size=256,
    use_moe=True,
    random_state=0,
)

print("Target score (mean abs ΔRNA over target genes):", null_out["target_score"])
print("Empirical p-value (null >= target):", null_out["empirical_p_ge"])
display(null_out["target_gene_summary"])

plot_null_distribution(
    null_out["null_scores"],
    null_out["target_score"],
    title="Matched null peak sets vs target perturbation"
)


# %% [code]


# %% [code]
# ============================================================
# Comprehensive gene complex/pathway presets (human symbols)
# Immune-heavy, useful for single-cell perturbation experiments
# ============================================================
from typing import Sequence, Mapping, Any, Iterable, Dict, List

def _dedup_keep_order(xs):
    seen = set()
    out = []
    for x in xs:
        x = str(x)
        if x not in seen:
            seen.add(x)
            out.append(x)
    return out

def _merge_presets(*lists):
    out = []
    for lst in lists:
        out.extend(list(lst))
    return _dedup_keep_order(out)


GENE_COMPLEX_PRESETS = {
    # ------------------------------------------------------------------
    # HLA / MHC / Antigen presentation
    # ------------------------------------------------------------------
    "hla_class_i_core": [
        "HLA-A", "HLA-B", "HLA-C", "B2M"
    ],

    "hla_class_i_extended": [
        "HLA-A", "HLA-B", "HLA-C", "B2M",
        "HLA-E", "HLA-F", "HLA-G"
    ],

    "hla_class_i_antigen_processing_core": [
        "HLA-A", "HLA-B", "HLA-C", "B2M",
        "TAP1", "TAP2", "TAPBP",
        "PSMB8", "PSMB9", "PSMB10",
        "ERAP1", "ERAP2",
        "NLRC5"
    ],

    "hla_class_i_antigen_processing_broad": [
        # MHC-I heavy chain + beta2m
        "HLA-A", "HLA-B", "HLA-C", "B2M",
        # non-classical
        "HLA-E", "HLA-F", "HLA-G",
        # peptide loading / trimming
        "TAP1", "TAP2", "TAPBP", "ERAP1", "ERAP2",
        "CALR", "CANX", "PDIA3",
        # immunoproteasome / processing
        "PSMB8", "PSMB9", "PSMB10", "PSME1", "PSME2",
        # regulators
        "NLRC5", "STAT1", "IRF1"
    ],

    "hla_class_ii_core": [
        "HLA-DRA", "HLA-DRB1",
        "HLA-DPA1", "HLA-DPB1",
        "HLA-DQA1", "HLA-DQB1",
        "CD74", "CIITA"
    ],

    "hla_class_ii_extended": [
        "HLA-DRA", "HLA-DRB1", "HLA-DRB5",
        "HLA-DPA1", "HLA-DPB1",
        "HLA-DQA1", "HLA-DQA2",
        "HLA-DQB1", "HLA-DQB2",
        "HLA-DMA", "HLA-DMB",
        "HLA-DOA", "HLA-DOB",
        "CD74", "CIITA", "RFX5", "RFXAP", "RFXANK"
    ],

    "mhc_antigen_presentation_broad": [
        # class I
        "HLA-A", "HLA-B", "HLA-C", "B2M",
        "HLA-E", "HLA-F", "HLA-G",
        "TAP1", "TAP2", "TAPBP",
        "ERAP1", "ERAP2",
        "PSMB8", "PSMB9", "PSMB10", "PSME1", "PSME2",
        "CALR", "CANX", "PDIA3",
        "NLRC5",
        # class II
        "HLA-DRA", "HLA-DRB1", "HLA-DRB5",
        "HLA-DPA1", "HLA-DPB1",
        "HLA-DQA1", "HLA-DQB1",
        "HLA-DMA", "HLA-DMB", "HLA-DOA", "HLA-DOB",
        "CD74", "CIITA", "RFX5", "RFXAP", "RFXANK"
    ],

    # MHC locus-focused subset (chr6-heavy genes; useful when range-restricting to MHC locus)
    "mhc_chr6_locus_class_i_processing": [
        "HLA-A", "HLA-B", "HLA-C",
        "TAP1", "TAP2", "TAPBP",
        "PSMB8", "PSMB9", "PSMB10"
    ],

    "mhc_chr6_locus_class_ii": [
        "HLA-DRA", "HLA-DRB1", "HLA-DRB5",
        "HLA-DPA1", "HLA-DPB1",
        "HLA-DQA1", "HLA-DQB1",
        "HLA-DMA", "HLA-DMB", "HLA-DOA", "HLA-DOB",
        "CD74"
    ],

    # ------------------------------------------------------------------
    # Interferon / inflammatory signaling (strongly tied to HLA programs)
    # ------------------------------------------------------------------
    "ifn_gamma_response_core": [
        "IFNGR1", "IFNGR2",
        "JAK1", "JAK2", "STAT1",
        "IRF1",
        "NLRC5", "CIITA"
    ],

    "ifn_gamma_response_extended": [
        "IFNGR1", "IFNGR2",
        "JAK1", "JAK2", "STAT1", "STAT2",
        "IRF1", "IRF8", "NFKB1", "RELA",
        "NLRC5", "CIITA",
        "GBP1", "GBP2", "GBP4", "GBP5",
        "CXCL9", "CXCL10", "CXCL11",
        "IDO1",
        "B2M", "TAP1", "TAP2", "PSMB8", "PSMB9", "PSMB10"
    ],

    "type_i_ifn_response_core": [
        "IFNAR1", "IFNAR2",
        "JAK1", "TYK2", "STAT1", "STAT2", "IRF9"
    ],

    "type_i_ifn_isg_broad": [
        "IFNAR1", "IFNAR2", "JAK1", "TYK2", "STAT1", "STAT2", "IRF9",
        "ISG15", "IFI6", "IFI27", "IFI35", "IFI44", "IFI44L",
        "IFIT1", "IFIT2", "IFIT3", "IFIT5",
        "IFITM1", "IFITM2", "IFITM3",
        "MX1", "MX2", "OAS1", "OAS2", "OAS3", "OASL",
        "RSAD2", "BST2", "CMPK2", "EPSTI1",
        "XAF1", "USP18", "DDX58", "IFIH1", "TRIM22", "TRIM25",
        "HERC5", "HERC6", "SAMD9", "SAMD9L",
        "IRF7", "IRF1", "STAT1"
    ],

    "tnf_nfkb_response_core": [
        "TNFRSF1A", "TNFRSF1B",
        "TRAF2", "RIPK1", "IKBKB",
        "NFKB1", "RELA", "RELB"
    ],

    "tnf_nfkb_response_extended": [
        "TNF", "TNFRSF1A", "TNFRSF1B",
        "TRAF2", "TRAF5", "RIPK1", "IKBKB", "CHUK", "IKBKG",
        "NFKB1", "NFKB2", "RELA", "RELB", "BCL3",
        "NFKBIA", "NFKBIZ", "TNFAIP3",
        "ICAM1", "VCAM1", "CCL2", "CCL5", "CXCL8", "IL6"
    ],

    # ------------------------------------------------------------------
    # Immune checkpoints / coinhibitory / costimulatory
    # ------------------------------------------------------------------
    "immune_checkpoints_tcell_inhibitory": [
        "PDCD1", "CTLA4", "LAG3", "TIGIT", "HAVCR2", "TOX", "TOX2"
    ],

    "immune_checkpoints_tcell_extended": [
        "PDCD1", "CTLA4", "LAG3", "TIGIT", "HAVCR2", "BTLA", "CD244", "CD160",
        "VSIR", "ENTPD1", "LAYN", "TOX", "TOX2", "EOMES", "PRDM1"
    ],

    "checkpoint_ligands_apc_tumor": [
        "CD274", "PDCD1LG2", "CD80", "CD86",
        "LGALS9", "PVR", "NECTIN2", "TNFSF9", "TNFSF4", "ICOSLG"
    ],

    "costimulation_tcell_apc": [
        "CD28", "ICOS", "TNFRSF9", "TNFRSF4", "TNFRSF18",
        "CD80", "CD86", "ICOSLG", "TNFSF9", "TNFSF4", "TNFSF18"
    ],

    # ------------------------------------------------------------------
    # Cytotoxic / NK / effector programs
    # ------------------------------------------------------------------
    "cytotoxic_effector_core": [
        "NKG7", "PRF1", "GZMB", "GNLY"
    ],

    "cytotoxic_effector_extended": [
        "NKG7", "PRF1", "GZMB", "GZMH", "GZMK", "GNLY",
        "CTSW", "CST7", "KLRD1", "KLRK1", "FCGR3A", "TYROBP", "TRAC"
    ],

    "nk_receptors_activating_inhibitory": [
        "KLRK1", "KLRC1", "KLRC2", "KLRC3", "KLRD1",
        "NCR1", "NCR2", "NCR3",
        "CD244", "FCGR3A", "TYROBP", "HCST", "SH2D1B"
    ],

    # ------------------------------------------------------------------
    # T-cell identity / activation / exhaustion / Treg
    # ------------------------------------------------------------------
    "tcell_pan_identity": [
        "CD3D", "CD3E", "CD3G", "TRAC", "TRBC1", "TRBC2", "LTB", "IL7R"
    ],

    "tcell_activation_early": [
        "IL2RA", "CD69", "TNFRSF9", "ICOS", "CD40LG", "FOS", "JUN", "JUNB", "DUSP1"
    ],

    "tcell_memory_naive_like": [
        "IL7R", "LTB", "MALAT1", "CCR7", "LTB", "MALAT1", "TCF7", "LEF1", "MALAT1", "MALAT1"
    ],

    "tcell_memory_naive_refined": [
        "IL7R", "CCR7", "LTB", "MALAT1", "TCF7", "LEF1", "MALAT1", "MALAT1",
        "MALAT1", "SELL", "MALAT1", "LTB"
    ],

    # cleaner version (use this one in practice)
    "tcell_naive_memory_clean": [
        "IL7R", "CCR7", "TCF7", "LEF1", "LTB", "SELL", "MALAT1", "MALAT1"
    ],

    "tcell_exhaustion_like": [
        "PDCD1", "LAG3", "TIGIT", "HAVCR2", "TOX", "CTLA4", "ENTPD1", "CXCL13"
    ],

    "treg_core": [
        "FOXP3", "IL2RA", "IKZF2", "TIGIT", "CTLA4", "LTB", "IL7R"
    ],

    "treg_extended": [
        "FOXP3", "IL2RA", "IKZF2", "TIGIT", "CTLA4", "TNFRSF18", "TNFRSF4",
        "BATF", "LAYN", "ENTPD1", "ICOS", "CCR8", "IL7R"
    ],

    "th1_like": [
        "TBX21", "IFNG", "CXCR3", "STAT1", "IL12RB2"
    ],

    "th17_like": [
        "RORC", "IL7R", "CCR6", "KLRB1", "IL23R", "IL17A", "IL17F"
    ],

    "tfh_like": [
        "CXCR5", "PDCD1", "ICOS", "IL21", "BCL6", "MAF", "TOX"
    ],

    # ------------------------------------------------------------------
    # B cell / plasma cell modules
    # ------------------------------------------------------------------
    "bcell_pan_identity": [
        "MS4A1", "CD79A", "CD79B", "HLA-DRA", "CD74", "CD37", "BANK1", "BLNK"
    ],

    "bcell_naive_like": [
        "MS4A1", "CD79A", "CD79B", "HVCN1", "TCL1A", "IL4R", "FCER2"
    ],

    "bcell_memory_like": [
        "MS4A1", "CD79A", "CD79B", "CD27", "TNFRSF13B", "AIM2", "BANK1"
    ],

    "plasma_cell_core": [
        "SDC1", "MZB1", "XBP1", "JCHAIN", "PRDM1", "IRF4"
    ],

    "plasma_cell_extended": [
        "SDC1", "MZB1", "XBP1", "JCHAIN", "PRDM1", "IRF4", "DERL3", "SSR4", "TXNDC5", "FKBP11"
    ],

    # ------------------------------------------------------------------
    # Myeloid / monocyte / DC / macrophage modules
    # ------------------------------------------------------------------
    "monocyte_classical_cd14": [
        "LYZ", "S100A8", "S100A9", "CTSS", "FCN1", "SAT1", "LILRB1", "TYMP"
    ],

    "monocyte_nonclassical_cd16": [
        "FCGR3A", "LST1", "IFITM3", "TYMP", "SAT1", "MS4A7", "AIF1", "CTD"
    ],

    "monocyte_ifn_inflam": [
        "IFITM3", "ISG15", "IFI6", "IFI44L", "MX1", "STAT1", "CXCL10", "GBP1", "BST2"
    ],

    "dc_cdc_identity": [
        "FCER1A", "CST3", "HLA-DRA", "CD74", "CLEC10A", "CD1C"
    ],

    "dc_pdc_identity": [
        "GZMB", "IRF7", "TCF4", "JCHAIN", "PLD4", "IFITM1", "IL3RA", "CLEC4C"
    ],

    "apc_mhcii_program": [
        "HLA-DRA", "HLA-DRB1", "HLA-DPA1", "HLA-DPB1", "HLA-DQA1", "HLA-DQB1",
        "CD74", "CIITA", "HLA-DMA", "HLA-DMB"
    ],

    "macrophage_antigen_presentation_inflam": [
        "HLA-DRA", "CD74", "CIITA", "FCER1G", "TYROBP", "CTSB", "CTSD",
        "IL1B", "TNF", "NFKBIA", "CXCL8"
    ],

    # ------------------------------------------------------------------
    # Tumor immune evasion / interferon-antigen axis (useful in cancer)
    # ------------------------------------------------------------------
    "tumor_mhc_i_evasion_axis": [
        "B2M", "HLA-A", "HLA-B", "HLA-C", "TAP1", "TAP2", "TAPBP", "NLRC5",
        "JAK1", "JAK2", "STAT1", "IRF1", "IFNGR1", "IFNGR2"
    ],

    "tumor_ifn_response_antigen_presentation": [
        "IFNGR1", "IFNGR2", "JAK1", "JAK2", "STAT1", "IRF1",
        "B2M", "HLA-A", "HLA-B", "HLA-C",
        "TAP1", "TAP2", "TAPBP",
        "PSMB8", "PSMB9", "PSMB10",
        "NLRC5", "CXCL9", "CXCL10", "CXCL11"
    ],

    "pd_l1_axis": [
        "CD274", "PDCD1LG2", "JAK1", "JAK2", "STAT1", "IRF1", "IFNGR1", "IFNGR2"
    ],

    # ------------------------------------------------------------------
    # Cell stress / apoptosis / proliferation (handy controls)
    # ------------------------------------------------------------------
    "apoptosis_intrinsic_core": [
        "BAX", "BAK1", "BCL2", "BCL2L1", "MCL1", "PMAIP1", "BBC3", "CASP3", "CASP9"
    ],

    "apoptosis_extrinsic_core": [
        "FAS", "FASLG", "TNFRSF10A", "TNFRSF10B", "CASP8", "CFLAR"
    ],

    "cell_cycle_g1s_core": [
        "MKI67", "PCNA", "MCM2", "MCM3", "MCM4", "MCM5", "MCM6", "MCM7", "TYMS", "RRM2"
    ],

    "cell_cycle_g2m_core": [
        "TOP2A", "UBE2C", "BIRC5", "AURKA", "AURKB", "CCNB1", "CCNB2", "CDC20", "CENPF"
    ],

    "hypoxia_core": [
        "HIF1A", "VEGFA", "SLC2A1", "LDHA", "PGK1", "ENO1", "BNIP3"
    ],

    "emt_like_core": [
        "VIM", "ZEB1", "ZEB2", "SNAI1", "SNAI2", "TWIST1", "FN1", "ITGA5"
    ],

    # ------------------------------------------------------------------
    # Signaling pathways (compact but useful)
    # ------------------------------------------------------------------
    "jak_stat_core": [
        "JAK1", "JAK2", "JAK3", "TYK2", "STAT1", "STAT2", "STAT3", "STAT5A", "STAT5B", "STAT6"
    ],

    "mapk_ap1_core": [
        "MAPK1", "MAPK3", "MAPK14", "JUN", "JUNB", "JUND", "FOS", "FOSB", "ATF3", "DUSP1"
    ],

    "pi3k_akt_mtor_core": [
        "PIK3CA", "PIK3CB", "PIK3CD", "AKT1", "AKT2", "AKT3", "MTOR", "RPTOR", "RICTOR", "EIF4EBP1"
    ],
}

# Clean up duplicates across all presets (and duplicate entries within a preset)
GENE_COMPLEX_PRESETS = {k: _dedup_keep_order(v) for k, v in GENE_COMPLEX_PRESETS.items()}


# %% [code]
# ============================================================
# Optional aliases and compositional presets
# ============================================================
GENE_COMPLEX_ALIASES = {
    "mhc_i": "hla_class_i_core",
    "mhc_i_proc": "hla_class_i_antigen_processing_core",
    "mhc_ii": "hla_class_ii_core",
    "mhc_broad": "mhc_antigen_presentation_broad",
    "ifng": "ifn_gamma_response_extended",
    "isg": "type_i_ifn_isg_broad",
    "treg": "treg_core",
    "cytotoxic": "cytotoxic_effector_extended",
}

def resolve_complex_preset_name(name: str) -> str:
    name = str(name)
    return GENE_COMPLEX_ALIASES.get(name, name)

def add_composite_preset(name: str, preset_names: Sequence[str], overwrite: bool = False):
    """
    Create a new preset by merging existing presets.
    """
    global GENE_COMPLEX_PRESETS
    if (name in GENE_COMPLEX_PRESETS) and (not overwrite):
        raise KeyError(f"Preset '{name}' already exists. Use overwrite=True to replace.")
    merged = []
    for p in preset_names:
        p = resolve_complex_preset_name(p)
        if p not in GENE_COMPLEX_PRESETS:
            raise KeyError(f"Unknown preset '{p}'")
        merged.extend(GENE_COMPLEX_PRESETS[p])
    GENE_COMPLEX_PRESETS[name] = _dedup_keep_order(merged)


# %% [code]
add_composite_preset(
    "ifng_plus_mhc_i_axis",
    ["ifn_gamma_response_extended", "hla_class_i_antigen_processing_broad"],
    overwrite=True
)


# %% [code]
def list_gene_complex_presets(pattern: str = None, min_size: int = 0, max_size: int = None):
    """
    Return DataFrame of preset names and sizes; optionally filter by substring pattern.
    """
    rows = []
    for k, v in GENE_COMPLEX_PRESETS.items():
        n = len(v)
        if n < min_size:
            continue
        if max_size is not None and n > max_size:
            continue
        if pattern is not None and pattern.lower() not in k.lower():
            continue
        rows.append({"preset": k, "n_genes": n, "genes_preview": ", ".join(v[:8]) + ("..." if len(v) > 8 else "")})
    return pd.DataFrame(rows).sort_values(["n_genes", "preset"], ascending=[False, True]).reset_index(drop=True)

def show_preset(name: str):
    name = resolve_complex_preset_name(name)
    if name not in GENE_COMPLEX_PRESETS:
        raise KeyError(f"Unknown preset '{name}'")
    genes = GENE_COMPLEX_PRESETS[name]
    print(f"{name} ({len(genes)} genes)")
    print(", ".join(genes))
    return genes


# %% [code]
display(list_gene_complex_presets(pattern="mhc"))
show_preset("mhc_i_proc")


# %% [code]
# ============================================================
# Genomic range presets (human hg38) for locus perturbation/plotting
# ============================================================

# NOTE:
# - Coordinates here are intended as practical plotting/perturbation windows,
#   not exact gene body definitions.
# - Many are "buffered" windows to capture local promoters/enhancers.
# - Use tighter windows for clean mechanistic tests, broader windows for exploration.

GENOMIC_RANGE_PRESETS = {
    # ------------------------------------------------------------------
    # MHC / HLA locus (chr6p21; hg38 approximate practical windows)
    # ------------------------------------------------------------------
    "mhc_chr6_ultrabroad_hg38": [("chr6", 24_000_000, 36_000_000)],
    "mhc_chr6_broad_hg38":      [("chr6", 25_000_000, 35_000_000)],
    "mhc_chr6_core_hg38":       [("chr6", 28_000_000, 34_000_000)],

    # Class I-heavy portion (includes HLA-A/B/C and antigen processing genes nearby)
    "mhc_class_i_region_hg38":  [("chr6", 29_000_000, 33_500_000)],

    # Class II-heavy portion (roughly DRA/DRB/DP/DQ region)
    "mhc_class_ii_region_hg38": [("chr6", 32_300_000, 33_600_000)],

    # More focused windows often useful for plotting
    "hla_abc_focus_hg38":       [("chr6", 29_600_000, 31_500_000)],   # HLA-A/B/C neighborhood
    "tap_psmb_focus_hg38":      [("chr6", 32_700_000, 33_100_000)],   # TAP1/2, PSMB8/9/10-ish
    "hla_dra_drb_focus_hg38":   [("chr6", 32_400_000, 32_700_000)],
    "hla_dpq_focus_hg38":       [("chr6", 33_000_000, 33_500_000)],

    # ------------------------------------------------------------------
    # Immune receptor loci (TCR/BCR)
    # ------------------------------------------------------------------
    # TCR alpha/delta locus (TRA/TRD)
    "tcrad_chr14_broad_hg38":   [("chr14", 21_000_000, 24_500_000)],
    "tcrad_chr14_core_hg38":    [("chr14", 21_500_000, 23_500_000)],

    # TCR beta locus
    "tcrb_chr7_broad_hg38":     [("chr7", 141_000_000, 143_000_000)],
    "tcrb_chr7_core_hg38":      [("chr7", 141_800_000, 142_900_000)],

    # TCR gamma locus
    "tcrg_chr7_broad_hg38":     [("chr7", 38_000_000, 39_500_000)],

    # Immunoglobulin heavy chain (IGH)
    "igh_chr14_broad_hg38":     [("chr14", 104_000_000, 107_500_000)],
    "igh_chr14_core_hg38":      [("chr14", 105_000_000, 107_000_000)],

    # Immunoglobulin kappa (IGK)
    "igk_chr2_broad_hg38":      [("chr2", 88_000_000, 91_500_000)],

    # Immunoglobulin lambda (IGL)
    "igl_chr22_broad_hg38":     [("chr22", 22_000_000, 24_500_000)],

    # ------------------------------------------------------------------
    # Cytokine / chemokine clusters
    # ------------------------------------------------------------------
    # IFNG
    "ifng_locus_chr12_hg38":    [("chr12", 67_900_000, 68_600_000)],

    # IL2 / IL21 region (nearby on chr4 but not same exact function; useful neighborhood)
    "il2_locus_chr4_hg38":      [("chr4", 122_300_000, 123_100_000)],
    "il21_locus_chr4_hg38":     [("chr4", 122_600_000, 123_600_000)],

    # IL10 family-ish focus (IL10 on chr1)
    "il10_locus_chr1_hg38":     [("chr1", 206_700_000, 207_200_000)],

    # TNF/LTA/LTB cluster in MHC
    "tnf_lta_ltb_mhc_focus_hg38": [("chr6", 31_400_000, 31_800_000)],

    # CXCL9/10/11 chemokine cluster (IFN-responsive)
    "cxcl9_10_11_chr4_hg38":    [("chr4", 76_800_000, 77_400_000)],

    # CCL3/CCL4 cluster
    "ccl3_ccl4_chr17_hg38":     [("chr17", 36_000_000, 36_400_000)],

    # ------------------------------------------------------------------
    # NK receptor / leukocyte receptor clusters
    # ------------------------------------------------------------------
    # KLR cluster (NKG2 family, etc.) on chr12
    "klr_cluster_chr12_hg38":   [("chr12", 10_000_000, 11_500_000)],

    # LRC (Leukocyte receptor complex) on chr19 (LILR, KIR region vicinity is broad/complex)
    "lrc_chr19_broad_hg38":     [("chr19", 54_000_000, 56_500_000)],

    # KIR cluster on chr19
    "kir_cluster_chr19_hg38":   [("chr19", 54_700_000, 55_400_000)],

    # ------------------------------------------------------------------
    # Interferon / innate sensing / antigen processing related single loci
    # ------------------------------------------------------------------
    "b2m_locus_chr15_hg38":     [("chr15", 44_700_000, 45_100_000)],
    "nlrc5_locus_chr16_hg38":   [("chr16", 56_700_000, 57_600_000)],
    "ciita_locus_chr16_hg38":   [("chr16", 10_700_000, 11_300_000)],
    "tapbp_locus_chr6_hg38":    [("chr6", 33_250_000, 33_380_000)],
    "psmb8_9_10_tap1_2_chr6_hg38": [("chr6", 32_780_000, 33_020_000)],

    # IFN receptor / signaling loci
    "ifngr1_locus_chr6_hg38":   [("chr6", 137_100_000, 137_500_000)],
    "ifngr2_locus_chr21_hg38":  [("chr21", 33_300_000, 33_800_000)],
    "jak1_locus_chr1_hg38":     [("chr1", 64_700_000, 65_400_000)],
    "jak2_locus_chr9_hg38":     [("chr9", 4_800_000, 5_600_000)],
    "stat1_locus_chr2_hg38":    [("chr2", 190_800_000, 191_300_000)],
    "irf1_locus_chr5_hg38":     [("chr5", 132_450_000, 132_850_000)],

    # Type I IFN / ISG examples
    "ifnar1_ifnar2_chr21_hg38": [("chr21", 33_900_000, 35_200_000)],
    "oas_cluster_chr12_hg38":   [("chr12", 112_700_000, 113_500_000)],
    "ifit_cluster_chr10_hg38":  [("chr10", 89_800_000, 90_600_000)],
    "mx1_mx2_chr21_hg38":       [("chr21", 41_100_000, 41_800_000)],

    # ------------------------------------------------------------------
    # Immune checkpoints / ligands
    # ------------------------------------------------------------------
    "pdcd1_locus_chr2_hg38":    [("chr2", 241_650_000, 242_050_000)],  # PD-1
    "ctla4_cd28_icos_chr2_hg38":[("chr2", 203_700_000, 205_200_000)],
    "lag3_locus_chr12_hg38":    [("chr12", 6_700_000, 7_500_000)],
    "tig_it_cd226_pvrig_axis_chr3_hg38": [("chr3", 108_000_000, 112_500_000)],  # broad neighborhood
    "havcr2_locus_chr5_hg38":   [("chr5", 156_550_000, 157_050_000)],  # TIM-3
    "cd274_pdcd1lg2_chr9_hg38": [("chr9", 5_300_000, 5_900_000)],      # PD-L1 / PD-L2

    # Costimulation
    "cd80_locus_chr3_hg38":     [("chr3", 119_000_000, 120_000_000)],
    "cd86_locus_chr3_hg38":     [("chr3", 122_500_000, 123_500_000)],
    "icoslg_locus_chr21_hg38":  [("chr21", 44_200_000, 44_900_000)],

    # ------------------------------------------------------------------
    # Treg / lineage factor loci
    # ------------------------------------------------------------------
    "foxp3_locus_chrX_hg38":    [("chrX", 49_000_000, 49_600_000)],
    "il2ra_locus_chr10_hg38":   [("chr10", 6_000_000, 6_700_000)],
    "ikzf2_locus_chr2_hg38":    [("chr2", 213_000_000, 214_000_000)],
    "tbx21_locus_chr17_hg38":   [("chr17", 46_500_000, 47_100_000)],
    "rorc_locus_chr1_hg38":     [("chr1", 151_700_000, 152_300_000)],
    "bcl6_locus_chr3_hg38":     [("chr3", 186_800_000, 188_000_000)],

    # ------------------------------------------------------------------
    # Cytotoxic effector loci
    # ------------------------------------------------------------------
    "prf1_locus_chr10_hg38":    [("chr10", 70_200_000, 70_800_000)],
    "gzmb_locus_chr14_hg38":    [("chr14", 25_000_000, 25_500_000)],
    "gzmh_locus_chr14_hg38":    [("chr14", 24_800_000, 25_200_000)],
    "nkg7_locus_chr19_hg38":    [("chr19", 51_300_000, 51_700_000)],
    "gnly_locus_chr2_hg38":     [("chr2", 85_500_000, 86_200_000)],

    # ------------------------------------------------------------------
    # B cell / plasma loci
    # ------------------------------------------------------------------
    "ms4a1_cd20_locus_chr11_hg38": [("chr11", 60_000_000, 60_600_000)],
    "cd79a_locus_chr19_hg38":      [("chr19", 42_000_000, 42_500_000)],
    "cd79b_locus_chr17_hg38":      [("chr17", 62_000_000, 62_600_000)],
    "jchain_locus_chr4_hg38":      [("chr4", 71_100_000, 71_700_000)],
    "mzb1_locus_chr5_hg38":        [("chr5", 66_000_000, 66_500_000)],
    "prdm1_locus_chr6_hg38":       [("chr6", 105_600_000, 106_200_000)],
    "xbp1_locus_chr22_hg38":       [("chr22", 28_800_000, 29_500_000)],

    # ------------------------------------------------------------------
    # Cancer / IO-relevant loci (NSCLC-friendly starter set)
    # ------------------------------------------------------------------
    "kras_locus_chr12_hg38":    [("chr12", 25_000_000, 25_600_000)],
    "egfr_locus_chr7_hg38":     [("chr7", 54_700_000, 55_800_000)],
    "tp53_locus_chr17_hg38":    [("chr17", 7_400_000, 8_000_000)],
    "myc_locus_chr8_hg38":      [("chr8", 127_400_000, 129_000_000)],
    "cdkn2a_b_locus_chr9_hg38": [("chr9", 21_600_000, 22_400_000)],
    "stk11_lkb1_locus_chr19_hg38": [("chr19", 1_100_000, 1_800_000)],
    "keap1_locus_chr19_hg38":   [("chr19", 10_400_000, 10_900_000)],
    "nfe2l2_locus_chr2_hg38":   [("chr2", 177_100_000, 177_900_000)],
    "smarca4_locus_chr19_hg38": [("chr19", 10_700_000, 11_800_000)],
    "pten_locus_chr10_hg38":    [("chr10", 87_700_000, 88_300_000)],

    # EMT / TGFb axis useful loci
    "tgfb1_locus_chr19_hg38":   [("chr19", 41_200_000, 41_900_000)],
    "tgfbr2_locus_chr3_hg38":   [("chr3", 30_400_000, 31_300_000)],
    "smad4_locus_chr18_hg38":   [("chr18", 50_500_000, 51_500_000)],
    "vim_locus_chr10_hg38":     [("chr10", 17_000_000, 17_600_000)],
    "zeb1_locus_chr10_hg38":    [("chr10", 31_300_000, 32_400_000)],

    # ------------------------------------------------------------------
    # Neutral-ish controls / housekeeping-ish loci for sanity checks
    # ------------------------------------------------------------------
    "actb_locus_chr7_hg38":     [("chr7", 5_400_000, 5_900_000)],
    "gapdh_locus_chr12_hg38":   [("chr12", 6_500_000, 7_100_000)],
    "rplp0_locus_chr12_hg38":   [("chr12", 119_000_000, 119_500_000)],
}

GENOMIC_RANGE_PRESETS["cytotoxic_effector_multilocus_hg38"] = [
    ("chr14", 24_800_000, 25_500_000),  # GZMH/GZMB neighborhood
    ("chr10", 70_200_000, 70_800_000),  # PRF1
    ("chr19", 51_300_000, 51_700_000),  # NKG7
    ("chr2", 85_500_000, 86_200_000),   # GNLY
]

# Optional aliases (short names)
GENOMIC_RANGE_ALIASES = {
    "mhc": "mhc_chr6_core_hg38",
    "mhc_broad": "mhc_chr6_broad_hg38",
    "mhc_i": "mhc_class_i_region_hg38",
    "mhc_ii": "mhc_class_ii_region_hg38",
    "hla_abc": "hla_abc_focus_hg38",
    "tap_psmb": "tap_psmb_focus_hg38",
    "pdl1": "cd274_pdcd1lg2_chr9_hg38",
    "kras": "kras_locus_chr12_hg38",
    "egfr": "egfr_locus_chr7_hg38",
    "tp53": "tp53_locus_chr17_hg38",
}


# %% [code]
import re

def _parse_region_string(region_str: str):
    """
    Parse strings like:
      'chr6:28000000-34000000'
      'chr6:28,000,000-34,000,000'
    Returns (chrom, start, end)
    """
    s = str(region_str).strip()
    m = re.match(r"^([^:]+):([\d,]+)-([\d,]+)$", s)
    if not m:
        raise ValueError(f"Could not parse region string: {region_str}")
    chrom = m.group(1)
    st = int(m.group(2).replace(",", ""))
    en = int(m.group(3).replace(",", ""))
    if en < st:
        st, en = en, st
    return (chrom, st, en)

def resolve_range_preset_name(name: str) -> str:
    name = str(name)
    return GENOMIC_RANGE_ALIASES.get(name, name)

def normalize_ranges(ranges):
    """
    Normalize ranges into list of (chrom, start, end).

    Accepts:
      - tuple ('chr6', 28000000, 34000000)
      - list of tuples
      - preset string key in GENOMIC_RANGE_PRESETS
      - alias string in GENOMIC_RANGE_ALIASES
      - coordinate string 'chr6:28000000-34000000'
    """
    if ranges is None:
        return None

    # String input: alias, preset, or explicit coordinate string
    if isinstance(ranges, str):
        rname = resolve_range_preset_name(ranges)
        if rname in GENOMIC_RANGE_PRESETS:
            ranges = GENOMIC_RANGE_PRESETS[rname]
        else:
            # try parse literal coordinate string
            ranges = [_parse_region_string(ranges)]

    if isinstance(ranges, tuple):
        ranges = [ranges]

    if not isinstance(ranges, (list, tuple)):
        raise TypeError(f"ranges must be None, str, tuple, or list of tuples; got {type(ranges)}")

    out = []
    for r in ranges:
        if len(r) != 3:
            raise ValueError(f"Each range must be (chrom,start,end), got: {r}")
        chrom, st, en = r
        chrom = str(chrom)
        st = int(st)
        en = int(en)
        if en < st:
            st, en = en, st
        out.append((chrom, st, en))
    return out


# %% [code]
def list_genomic_range_presets(pattern: str = None):
    rows = []
    for k, v in GENOMIC_RANGE_PRESETS.items():
        n_intervals = len(v)
        total_bp = int(sum(int(en) - int(st) for _, st, en in v))
        preview = "; ".join([f"{c}:{st:,}-{en:,}" for c, st, en in v[:3]])
        if len(v) > 3:
            preview += " ..."
        if pattern and pattern.lower() not in k.lower():
            continue
        rows.append({
            "preset": k,
            "n_intervals": n_intervals,
            "total_bp": total_bp,
            "preview": preview
        })
    return pd.DataFrame(rows).sort_values(["preset"]).reset_index(drop=True)

def add_composite_range_preset(name: str, preset_names, overwrite: bool = False):
    global GENOMIC_RANGE_PRESETS
    if (name in GENOMIC_RANGE_PRESETS) and (not overwrite):
        raise KeyError(f"Range preset '{name}' already exists. Use overwrite=True.")
    intervals = []
    for p in preset_names:
        p2 = resolve_range_preset_name(p)
        if p2 not in GENOMIC_RANGE_PRESETS:
            raise KeyError(f"Unknown range preset '{p}'")
        intervals.extend(GENOMIC_RANGE_PRESETS[p2])
    GENOMIC_RANGE_PRESETS[name] = normalize_ranges(intervals)

def show_range_preset(name: str):
    name2 = resolve_range_preset_name(name)
    if name2 not in GENOMIC_RANGE_PRESETS:
        raise KeyError(f"Unknown range preset '{name}'")
    rr = GENOMIC_RANGE_PRESETS[name2]
    print(f"{name2} ({len(rr)} intervals)")
    for chrom, st, en in rr:
        print(f"  - {chrom}:{st:,}-{en:,} ({en-st:,} bp)")
    return rr


# %% [code]
display(list_genomic_range_presets(pattern="mhc"))
show_range_preset("mhc")
#show_range_preset("chr6:28,000,000-34,000,000")

# Make a combined MHC + B2M + NLRC5 perturbation region set
add_composite_range_preset(
    "mhc_plus_b2m_nlrc5_hg38",
    ["mhc_chr6_core_hg38", "b2m_locus_chr15_hg38", "nlrc5_locus_chr16_hg38"],
    overwrite=True
)
show_range_preset("mhc_plus_b2m_nlrc5_hg38")


# %% [code]
# ============================================================
# Select target peaks for named complex/pathway
# ============================================================
from typing import Sequence, Mapping, Any, Optional, Tuple, Dict, List
import numpy as np
import pandas as pd
import scipy.sparse as sp
import anndata as ad

def filter_links_by_peak_ranges(
    peak_gene_links: pd.DataFrame,
    atac: ad.AnnData,
    ranges=None,
    peak_chrom_col: str = "chrom",
    peak_start_col: str = "chromStart",
    peak_end_col: str = "chromEnd",
) -> pd.DataFrame:
    """
    Restrict peak_gene_links to peaks overlapping any provided genomic range(s).
    """
    if peak_gene_links is None or peak_gene_links.empty:
        return peak_gene_links.copy()

    rr = normalize_ranges(ranges)
    if rr is None:
        return peak_gene_links.copy()

    vA = atac.var.copy()
    for c in [peak_chrom_col, peak_start_col, peak_end_col]:
        if c not in vA.columns:
            raise KeyError(f"atac.var missing required column '{c}' for genomic range filtering")

    pchr = vA[peak_chrom_col].astype(str).values
    pst = vA[peak_start_col].astype(int).values
    pen = vA[peak_end_col].astype(int).values

    keep_peak = np.zeros(atac.n_vars, dtype=bool)
    for chrom, st, en in rr:
        m = (pchr == str(chrom)) & (pen >= int(st)) & (pst <= int(en))
        keep_peak |= m

    keep_idx = set(np.where(keep_peak)[0].tolist())
    out = peak_gene_links[peak_gene_links["peak_idx"].astype(int).isin(keep_idx)].copy()
    return out.reset_index(drop=True)

def get_gene_set_from_complex(
    complex_name_or_genes,
    custom_gene_sets: Optional[Mapping[str, Sequence[str]]] = None,
) -> List[str]:
    """
    Accepts either:
      - preset name (str), looked up in GENE_COMPLEX_PRESETS (and optional custom dict)
      - explicit list/tuple of gene symbols
    """
    if isinstance(complex_name_or_genes, str):
        # optional alias support if you added resolve_complex_preset_name()
        cname = resolve_complex_preset_name(complex_name_or_genes) if "resolve_complex_preset_name" in globals() else complex_name_or_genes

        if custom_gene_sets is not None and cname in custom_gene_sets:
            return [str(x) for x in custom_gene_sets[cname]]
        if cname in GENE_COMPLEX_PRESETS:
            return [str(x) for x in GENE_COMPLEX_PRESETS[cname]]

        avail = sorted(list(GENE_COMPLEX_PRESETS.keys()) + (list(custom_gene_sets.keys()) if custom_gene_sets else []))
        raise KeyError(f"Unknown complex preset '{complex_name_or_genes}'. Available: {avail}")

    return [str(x) for x in complex_name_or_genes]

def select_peaks_for_complex(
    peak_gene_links: pd.DataFrame,
    atac: ad.AnnData,
    complex_name_or_genes,
    *,
    match_on: str = "gene_name",
    include_link_types: Tuple[str, ...] = ("promoter", "enhancer"),
    restrict_peak_ranges=None,   # e.g. "mhc_chr6_core_hg38" or [("chr6", ...)]
    custom_gene_sets: Optional[Mapping[str, Sequence[str]]] = None,
    dedup: bool = True,
) -> Dict[str, Any]:
    """
    Returns same style output as select_peaks_for_gene_set(...) plus complex metadata.
    """
    genes = get_gene_set_from_complex(complex_name_or_genes, custom_gene_sets=custom_gene_sets)

    links_use = peak_gene_links.copy()
    if restrict_peak_ranges is not None:
        links_use = filter_links_by_peak_ranges(links_use, atac, ranges=restrict_peak_ranges)

    sel = select_peaks_for_gene_set(
        links_use,
        genes,
        match_on=match_on,
        include_link_types=include_link_types,
        dedup=dedup,
    )

    sel["complex_name"] = complex_name_or_genes if isinstance(complex_name_or_genes, str) else "custom"
    sel["target_gene_set"] = genes
    sel["restrict_peak_ranges"] = normalize_ranges(restrict_peak_ranges) if restrict_peak_ranges is not None else None
    return sel


# %% [code]
# ============================================================
# Pseudobulk helpers for ATAC and RNA + input semantics checks
# ============================================================

def _matrix_from_adata(adata: ad.AnnData, layer: Optional[str] = None):
    if layer is None or layer == "X":
        return adata.X
    if layer not in adata.layers:
        raise KeyError(f"Layer '{layer}' not found in adata.layers")
    return adata.layers[layer]

def _sample_array_from_matrix(X, max_n: int = 200_000, random_state: int = 0) -> np.ndarray:
    """
    Return a 1D float array sampled from matrix values (dense or sparse) for diagnostics.
    """
    if sp.issparse(X):
        arr = X.data
    else:
        arr = np.asarray(X).ravel()

    arr = np.asarray(arr, dtype=float)
    if arr.size == 0:
        return arr

    if arr.size > max_n:
        rng = np.random.default_rng(random_state)
        idx = rng.choice(arr.size, size=max_n, replace=False)
        arr = arr[idx]
    return arr

def summarize_matrix_values(X, max_n: int = 200_000, random_state: int = 0) -> Dict[str, Any]:
    """
    Lightweight numeric summary of a matrix for semantics checks/debugging.
    """
    arr = _sample_array_from_matrix(X, max_n=max_n, random_state=random_state)
    if arr.size == 0:
        return {
            "n_sampled": 0,
            "min": np.nan,
            "max": np.nan,
            "mean": np.nan,
            "std": np.nan,
            "frac_zero": np.nan,
            "frac_one": np.nan,
            "is_binary_like": False,
            "in_unit_interval": False,
        }

    uniq_probe = np.unique(arr[: min(arr.size, 50_000)])
    is_binary_like = np.all(np.isin(uniq_probe, [0.0, 1.0]))
    in_unit_interval = (np.nanmin(arr) >= -1e-6) and (np.nanmax(arr) <= 1.0 + 1e-6)

    return {
        "n_sampled": int(arr.size),
        "min": float(np.nanmin(arr)),
        "max": float(np.nanmax(arr)),
        "mean": float(np.nanmean(arr)),
        "std": float(np.nanstd(arr)),
        "frac_zero": float(np.mean(arr == 0.0)),
        "frac_one": float(np.mean(arr == 1.0)),
        "is_binary_like": bool(is_binary_like),
        "in_unit_interval": bool(in_unit_interval),
    }

def classify_atac_input_semantics(X, source_layer: Optional[str] = None) -> Dict[str, Any]:
    """
    Heuristic classification of ATAC input value semantics for perturbation mode safety.
    """
    s = summarize_matrix_values(X)
    layer_name = "X" if source_layer is None else str(source_layer)

    if s["n_sampled"] == 0:
        label = "empty"
        safe_onoff = False
        reason = "Matrix has no sampled values."
    elif s["is_binary_like"]:
        label = "binary_01"
        safe_onoff = True
        reason = "Values look binary (0/1)."
    elif s["in_unit_interval"]:
        label = "unit_interval_fraction_or_prob"
        safe_onoff = True
        reason = "Values are within [0,1]; likely probabilities/fractions (e.g., pseudobulk of bin)."
    else:
        # common for logits or transformed ATAC representations
        label = "transformed_or_logit_like"
        safe_onoff = False
        reason = "Values extend outside [0,1]; on/off semantics are not safe (use add/scale instead)."

    out = {
        "source_layer": layer_name,
        "semantics_label": label,
        "safe_for_on_off_set": bool(safe_onoff),
        "reason": reason,
        **s,
    }
    return out

def pseudobulk_adata_by_group(
    adata: ad.AnnData,
    groupby: str,
    *,
    layer: Optional[str] = None,
    agg: str = "mean",   # "mean" or "sum"
    min_cells: int = 5,
    groups: Optional[Sequence[str]] = None,
    out_X_dtype=np.float32,
) -> ad.AnnData:
    """
    Build pseudobulk profiles by adata.obs[groupby].
    Returns new AnnData with one row per group.
    """
    if groupby not in adata.obs.columns:
        raise KeyError(f"'{groupby}' not in adata.obs")

    if agg not in {"mean", "sum"}:
        raise ValueError("agg must be 'mean' or 'sum'")

    obs = adata.obs.copy()
    grp_series = obs[groupby].astype(str)

    if groups is None:
        group_levels = grp_series.value_counts().index.tolist()
    else:
        group_levels = [str(g) for g in groups]

    X = _matrix_from_adata(adata, layer=layer)

    rows = []
    out_obs = []
    for g in group_levels:
        idx = np.where(grp_series.values == g)[0]
        n = idx.size
        if n < min_cells:
            continue

        Xi = X[idx]
        if sp.issparse(Xi):
            xb = np.asarray(Xi.mean(axis=0)).ravel() if agg == "mean" else np.asarray(Xi.sum(axis=0)).ravel()
        else:
            Xi = np.asarray(Xi)
            xb = Xi.mean(axis=0) if agg == "mean" else Xi.sum(axis=0)

        rows.append(np.asarray(xb, dtype=out_X_dtype))
        out_obs.append({
            groupby: g,
            "n_cells": int(n),
            "pseudobulk_agg": str(agg),
            "source_layer": str(layer if layer is not None else "X"),
        })

    if len(rows) == 0:
        raise ValueError(f"No groups passed min_cells={min_cells} for groupby='{groupby}'")

    Xpb = np.vstack(rows).astype(out_X_dtype)
    pb = ad.AnnData(
        X=Xpb,
        obs=pd.DataFrame(out_obs).set_index(groupby),
        var=adata.var.copy()
    )

    # attach compact matrix summary for easy debugging
    pb.uns["pseudobulk_source"] = {
        "groupby": groupby,
        "agg": agg,
        "source_layer": str(layer if layer is not None else "X"),
        "value_summary": summarize_matrix_values(pb.X),
    }
    return pb

def aligned_pseudobulk_pair(
    atac: ad.AnnData,
    rna: ad.AnnData,
    groupby: str = "cell_type",
    *,
    atac_layer: Optional[str] = None,
    rna_layer: Optional[str] = None,
    agg: str = "mean",
    min_cells: int = 5,
) -> Tuple[ad.AnnData, ad.AnnData]:
    """
    Build pseudobulk ATAC and RNA with same groups (intersection of labels).
    Assumes atac.obs[groupby] and rna.obs[groupby] exist and use same labels.
    """
    if groupby not in atac.obs.columns or groupby not in rna.obs.columns:
        raise KeyError(f"groupby='{groupby}' must exist in both atac.obs and rna.obs")

    a_groups = set(atac.obs[groupby].astype(str).unique())
    r_groups = set(rna.obs[groupby].astype(str).unique())
    common = sorted(list(a_groups & r_groups))
    if len(common) == 0:
        raise ValueError(f"No overlapping groups for groupby='{groupby}' between atac and rna")

    atac_pb = pseudobulk_adata_by_group(
        atac, groupby, layer=atac_layer, agg=agg, min_cells=min_cells, groups=common
    )
    rna_pb = pseudobulk_adata_by_group(
        rna, groupby, layer=rna_layer, agg=agg, min_cells=min_cells, groups=common
    )

    common2 = sorted(list(set(atac_pb.obs_names.astype(str)) & set(rna_pb.obs_names.astype(str))))
    if len(common2) == 0:
        raise ValueError("No overlapping pseudobulk groups after min_cells filtering")

    atac_pb = atac_pb[common2].copy()
    rna_pb  = rna_pb[common2].copy()

    # annotate alignment metadata
    atac_pb.uns.setdefault("pseudobulk_pair", {})
    rna_pb.uns.setdefault("pseudobulk_pair", {})
    atac_pb.uns["pseudobulk_pair"]["aligned_groups"] = common2
    rna_pb.uns["pseudobulk_pair"]["aligned_groups"] = common2

    return atac_pb, rna_pb


# %% [code]
# ============================================================
# Group-level delta summaries
# ============================================================
def summarize_groupwise_target_gene_deltas(
    rna: ad.AnnData,
    delta_pred: np.ndarray,     # [n_groups, n_genes]
    group_names: Sequence[str],
    target_genes: Sequence[str],
    match_on: str = "gene_name",
) -> pd.DataFrame:
    """
    Returns long-form table: group x target_gene with delta summaries.
    """
    idx = gene_indices_from_list(rna, target_genes, match_on=match_on)
    if idx.size == 0:
        return pd.DataFrame(columns=["group", "gene_idx", "gene_name", "gene_id", "delta", "delta_abs"])

    d = np.asarray(delta_pred)[:, idx]  # [n_groups, n_targets]
    sub = rna.var.iloc[idx].copy()

    gnames = sub["gene_name"].astype(str).values if "gene_name" in sub.columns else rna.var_names[idx].astype(str)
    gids = sub["gene_id"].astype(str).values if "gene_id" in sub.columns else np.array([None] * idx.size, dtype=object)

    rows = []
    for i, grp in enumerate(group_names):
        for j, gi in enumerate(idx):
            rows.append({
                "group": str(grp),
                "gene_idx": int(gi),
                "gene_name": str(gnames[j]),
                "gene_id": str(gids[j]) if gids[j] is not None else None,
                "delta": float(d[i, j]),
                "delta_abs": float(abs(d[i, j])),
            })
    return pd.DataFrame(rows)

def summarize_group_scores(group_gene_delta_df: pd.DataFrame) -> pd.DataFrame:
    """
    Collapse long-form group x gene deltas into one row per group.
    """
    if group_gene_delta_df.empty:
        return pd.DataFrame(columns=["group", "mean_delta", "mean_abs_delta", "median_delta", "n_target_genes"])

    out = (
        group_gene_delta_df.groupby("group", as_index=False)
        .agg(
            mean_delta=("delta", "mean"),
            mean_abs_delta=("delta_abs", "mean"),
            median_delta=("delta", "median"),
            n_target_genes=("gene_name", "nunique"),
        )
        .sort_values("mean_abs_delta", ascending=False)
        .reset_index(drop=True)
    )
    return out


# %% [code]
# ============================================================
# Complex perturbation on pseudo-bulk groups (with input semantics guards)
# ============================================================
def run_complex_perturbation_pseudobulk(
    model,
    atac: ad.AnnData,
    rna: ad.AnnData,
    peak_gene_links: pd.DataFrame,
    complex_name_or_genes,
    *,
    groupby: str = "cell_type",
    atac_input_layer: Optional[str] = None,   # ATAC representation used as model input
    rna_obs_layer: Optional[str] = None,      # observed RNA layer for summaries (optional)
    pseudobulk_agg: str = "mean",
    min_cells: int = 10,
    include_link_types: Tuple[str, ...] = ("promoter", "enhancer"),
    restrict_peak_ranges=None,                # e.g. "mhc_chr6_core_hg38"
    mode: str = "off",                        # off/on/set/scale/add
    value: float = 0.0,
    clip_min: Optional[float] = 0.0,
    clip_max: Optional[float] = 1.0,
    match_on: str = "gene_name",
    device=None,
    batch_size: int = 256,
    use_moe: bool = True,
    custom_gene_sets: Optional[Mapping[str, Sequence[str]]] = None,

    # NEW safety / diagnostics controls
    strict_input_semantics: bool = True,
    verbose: bool = True,
):
    """
    Run complex/pathway-targeted ATAC perturbation on pseudobulked groups and summarize predicted RNA deltas.

    Important semantics:
      - mode in {'on','off','set'} assumes the ATAC input values are interpretable in [0,1]
        (binary peaks or pseudobulk fractions/probabilities).
      - If ATAC pseudobulk input is outside [0,1] (e.g., logits/transformed ATAC), use mode='add' or 'scale'
        OR set strict_input_semantics=False if you intentionally want to proceed.
    """
    mode = str(mode).lower()
    valid_modes = {"off", "on", "set", "scale", "add"}
    if mode not in valid_modes:
        raise ValueError(f"mode must be one of {valid_modes}, got {mode!r}")

    # 1) Choose peaks for complex
    sel = select_peaks_for_complex(
        peak_gene_links=peak_gene_links,
        atac=atac,
        complex_name_or_genes=complex_name_or_genes,
        match_on=match_on,
        include_link_types=include_link_types,
        restrict_peak_ranges=restrict_peak_ranges,
        custom_gene_sets=custom_gene_sets,
    )
    peak_idx = np.asarray(sel["peak_idx"], dtype=int)
    target_genes = sel["target_gene_set"]

    if peak_idx.size == 0:
        raise ValueError(
            f"No peaks selected for complex={complex_name_or_genes}, "
            f"link_types={include_link_types}, ranges={restrict_peak_ranges}. "
            f"Genes found={sel.get('genes_found', [])}, missing={sel.get('genes_missing', [])}"
        )

    # 2) Pseudobulk ATAC + RNA by group
    atac_pb, rna_pb = aligned_pseudobulk_pair(
        atac, rna,
        groupby=groupby,
        atac_layer=atac_input_layer,  # pseudobulk from same ATAC representation used for prediction input
        rna_layer=rna_obs_layer,
        agg=pseudobulk_agg,
        min_cells=min_cells,
    )

    # 2b) Diagnose pseudobulk ATAC input semantics before perturbation
    atac_input_info = classify_atac_input_semantics(atac_pb.X, source_layer=atac_input_layer)
    target_peak_summary_pre = summarize_matrix_values(atac_pb.X[:, peak_idx]) if peak_idx.size > 0 else None

    if verbose:
        print(f"[INFO] Pseudobulk groups retained: {atac_pb.n_obs}")
        print(f"[INFO] ATAC pseudobulk source layer: {atac_input_layer if atac_input_layer is not None else 'X'}")
        print(f"[INFO] ATAC input semantics: {atac_input_info['semantics_label']}")
        print(f"[INFO] ATAC value range: {atac_input_info['min']:.4g} .. {atac_input_info['max']:.4g}")
        print(f"[INFO] Selected target peaks: {peak_idx.size}")
        if target_peak_summary_pre is not None and target_peak_summary_pre["n_sampled"] > 0:
            print(f"[INFO] Target-peak baseline summary (pre-perturb): "
                  f"min={target_peak_summary_pre['min']:.4g}, max={target_peak_summary_pre['max']:.4g}, "
                  f"mean={target_peak_summary_pre['mean']:.4g}")

    # Guardrail: on/off/set only makes sense on [0,1] values
    if mode in {"on", "off", "set"} and not atac_input_info["safe_for_on_off_set"]:
        msg = (
            f"Unsafe perturbation mode '{mode}' for ATAC input semantics '{atac_input_info['semantics_label']}'. "
            f"ATAC pseudobulk values are outside [0,1] "
            f"(min={atac_input_info['min']:.4g}, max={atac_input_info['max']:.4g}). "
            f"This usually means logits/transformed ATAC (e.g., decoder outputs), where on/off semantics are invalid. "
            f"Use atac_input_layer='bin' (or a [0,1] probability/fraction layer), or use mode='add'/'scale'."
        )
        if strict_input_semantics:
            raise ValueError(msg)
        else:
            print("[WARN] " + msg)

    # Optional warning: 'sum' pseudobulk plus on/off often not what user intends
    if mode in {"on", "off", "set"} and str(pseudobulk_agg).lower() == "sum":
        print("[WARN] pseudobulk_agg='sum' with mode on/off/set may be hard to interpret "
              "(setting counts-like sums to 0/1). Consider agg='mean' for fraction-accessibility semantics.")

    # 2c) Run perturbation in pseudobulk input space
    # IMPORTANT: atac_pb.X already contains the chosen representation, so atac_input_layer=None downstream
    res = run_atac_ablation_to_rna(
        model=model,
        atac=atac_pb,
        rna=rna_pb,
        peak_idx=peak_idx,
        cell_idx=None,
        atac_input_layer=None,     # pseudobulk input is in X
        mode=mode,
        value=value,
        clip_min=clip_min,
        clip_max=clip_max,
        device=device,
        batch_size=batch_size,
        use_moe=use_moe,
        return_observed_rna=True,
        rna_obs_layer=None,        # pseudobulk RNA is in X
    )

    # 2d) Post-perturbation sanity diagnostics (ATAC level + delta sign)
    target_peak_summary_base = summarize_matrix_values(res["base_atac"].X[:, peak_idx]) if peak_idx.size > 0 else None
    target_peak_summary_pert = summarize_matrix_values(res["pert_atac"].X[:, peak_idx]) if peak_idx.size > 0 else None

    # robust target peak mean delta (exact, dense small matrix)
    base_tp = np.asarray(res["base_atac"].X[:, peak_idx], dtype=float)
    pert_tp = np.asarray(res["pert_atac"].X[:, peak_idx], dtype=float)
    target_peak_atac_delta_mean = float((pert_tp - base_tp).mean()) if peak_idx.size > 0 else np.nan

    # Check delta sign convention
    delta_pred_matches_pert_minus_base = bool(
        np.allclose(
            np.asarray(res["delta_pred"]),
            np.asarray(res["pred_pert"]) - np.asarray(res["pred_base"]),
            atol=1e-6,
            equal_nan=True
        )
    )

    if verbose:
        print(f"[INFO] Target-peak mean ΔATAC (pert-base): {target_peak_atac_delta_mean:.6f}")
        print(f"[INFO] delta_pred == pred_pert - pred_base ? {delta_pred_matches_pert_minus_base}")

    # 3) Summaries (RNA deltas)
    df_delta_all = summarize_delta_by_gene(
        rna_pb,
        pred_base=res["pred_base"],
        pred_pert=res["pred_pert"],
        obs_rna=res.get("obs_rna", None),
        sort_by="mean_delta_abs",
    )

    group_names = list(rna_pb.obs_names.astype(str))
    df_group_gene = summarize_groupwise_target_gene_deltas(
        rna=rna_pb,
        delta_pred=res["delta_pred"],
        group_names=group_names,
        target_genes=target_genes,
        match_on=match_on,
    )
    df_group_scores = summarize_group_scores(df_group_gene)

    # 4) Package output + diagnostics
    out = {
        "selection": sel,
        "peak_idx": peak_idx,
        "target_genes": target_genes,

        "atac_pseudobulk": atac_pb,
        "rna_pseudobulk": rna_pb,
        "result": res,

        "delta_all_genes": df_delta_all,
        "group_target_gene_deltas": df_group_gene,
        "group_scores": df_group_scores,

        # New diagnostics (super helpful for debugging semantics)
        "input_diagnostics": {
            "groupby": groupby,
            "pseudobulk_agg": pseudobulk_agg,
            "atac_input_layer": atac_input_layer if atac_input_layer is not None else "X",
            "rna_obs_layer": rna_obs_layer if rna_obs_layer is not None else "X",
            "atac_input_semantics": atac_input_info,
            "mode": mode,
            "value": float(value),
            "clip_min": None if clip_min is None else float(clip_min),
            "clip_max": None if clip_max is None else float(clip_max),
            "target_peak_summary_pre": target_peak_summary_pre,
            "target_peak_summary_base": target_peak_summary_base,
            "target_peak_summary_pert": target_peak_summary_pert,
            "target_peak_atac_delta_mean": target_peak_atac_delta_mean,
            "delta_pred_matches_pert_minus_base": delta_pred_matches_pert_minus_base,
        }
    }
    return out


# %% [code]
def print_complex_perturbation_diagnostics(out: dict):
    d = out.get("input_diagnostics", {})
    if not d:
        print("No input_diagnostics found.")
        return

    s = d.get("atac_input_semantics", {})
    print("=== Complex perturbation diagnostics ===")
    print(f"ATAC input layer      : {d.get('atac_input_layer')}")
    print(f"Pseudobulk agg        : {d.get('pseudobulk_agg')}")
    print(f"Mode / value          : {d.get('mode')} / {d.get('value')}")
    print(f"ATAC semantics        : {s.get('semantics_label')}")
    print(f"Safe on/off/set       : {s.get('safe_for_on_off_set')}")
    print(f"ATAC range            : {s.get('min')} .. {s.get('max')}")
    print(f"Target peak mean ΔATAC: {d.get('target_peak_atac_delta_mean')}")
    print(f"delta_pred sign check : {d.get('delta_pred_matches_pert_minus_base')}")


# %% [code]
# ============================================================
# Plot helpers for pseudobulk complex perturbation
# ============================================================
def plot_group_scores(df_group_scores: pd.DataFrame, metric: str = "mean_abs_delta", title: Optional[str] = None):
    if df_group_scores.empty:
        print("No group scores to plot.")
        return
    d = df_group_scores.sort_values(metric, ascending=True).copy()
    plt.figure(figsize=(7, max(4, 0.3 * len(d))))
    plt.barh(np.arange(len(d)), d[metric].values)
    plt.yticks(np.arange(len(d)), d["group"].astype(str).values, fontsize=8)
    plt.xlabel(metric)
    if title:
        plt.title(title)
    plt.tight_layout()
    plt.show()

def plot_group_target_heatmap(group_gene_delta_df: pd.DataFrame, value_col: str = "delta"):
    """
    Simple matplotlib heatmap for group x target_gene deltas.
    """
    if group_gene_delta_df.empty:
        print("No group x target-gene delta table to plot.")
        return
    pivot = group_gene_delta_df.pivot_table(index="group", columns="gene_name", values=value_col, aggfunc="mean")
    arr = pivot.values

    plt.figure(figsize=(max(6, 0.4 * pivot.shape[1]), max(4, 0.35 * pivot.shape[0])))
    im = plt.imshow(arr, aspect="auto")
    plt.colorbar(im, fraction=0.046, pad=0.04, label=value_col)
    plt.xticks(np.arange(pivot.shape[1]), pivot.columns.astype(str), rotation=90, fontsize=8)
    plt.yticks(np.arange(pivot.shape[0]), pivot.index.astype(str), fontsize=8)
    plt.xlabel("Target genes")
    plt.ylabel("Pseudo-bulk group")
    plt.tight_layout()
    plt.show()


# %% [code]
# ============================================================
# EXAMPLE 1: GNLY promoter region expression major increase (+1000 peak counts) perturbation
# ============================================================

# Sanity check available group labels
print(atac.obs["cell_type"].value_counts().head(20))
print(rna.obs["cell_type"].value_counts().head(20))

hla_on = run_complex_perturbation_pseudobulk(
    model=model,
    atac=atac,
    rna=rna,
    peak_gene_links=peak_gene_links,
    #complex_name_or_genes="hla_class_i_antigen_processing_core",
    #complex_name_or_genes=["B2M"], 
    #complex_name_or_genes="cytotoxic_effector_extended",
    complex_name_or_genes=["GNLY"],
    #complex_name_or_genes=["PRF1"],
    groupby="cell_type",
    #atac_input_layer="bin",          # <- important
    atac_input_layer=None,
    rna_obs_layer=None,
    pseudobulk_agg="mean",           # <- fraction accessibility
    min_cells=10,
    include_link_types=("promoter",),
    #restrict_peak_ranges="mhc_class_i_region_hg38",
    #restrict_peak_ranges="b2m_locus_chr15_hg38",
    restrict_peak_ranges="cytotoxic_effector_multilocus_hg38",
    #mode="on",
    mode="add",
    #value=1.00,
    #value=50.00,
    value=1000.00,
    #clip_min=0.0,
    #clip_min=-100.0,
    #clip_max=1.0,
    clip_max=1000.0,
    clip_min=0.0,
    #strict_input_semantics=True,
    strict_input_semantics=False,
    verbose=True,
    device=device,
)

print_complex_perturbation_diagnostics(hla_on)


# %% [code]
plot_group_scores(
    hla_on["group_scores"],
    metric="mean_abs_delta",
    title="GNLY promoter region peak perturbation effect by pseudo-bulk cell_type"
)

plot_group_target_heatmap(
    hla_on["group_target_gene_deltas"],
    value_col="delta"
)


# %% [code]
import numpy as np
import pandas as pd
import scipy.sparse as sp
import matplotlib.pyplot as plt

def _to_dense1(x):
    if sp.issparse(x):
        return x.toarray().ravel()
    return np.asarray(x).ravel()

def _get_X(adata, layer=None):
    if layer is None or layer == "X":
        return adata.X
    return adata.layers[layer]

def build_locus_peak_plot_df(
    perturb_out: dict,
    atac_ref: ad.AnnData,
    *,
    region=("chr6", 28_000_000, 34_000_000),
    group: str = None,          # pseudo-bulk group name (e.g. "T cell"), or None = average across groups
    groupby: str = "cell_type", # only used for labels
    peak_chrom_col: str = "chrom",
    peak_start_col: str = "chromStart",
    peak_end_col: str = "chromEnd",
):
    """
    Build peak-level plotting table for a genomic locus from run_complex_perturbation_pseudobulk() output.
    Uses pseudo-bulk ATAC before/after perturbation stored in perturb_out["result"].
    """
    chrom, reg_start, reg_end = region

    # Pseudobulk ATAC before/after from the perturbation result
    base_atac_pb = perturb_out["result"]["base_atac"]   # AnnData, groups x peaks
    pert_atac_pb = perturb_out["result"]["pert_atac"]   # AnnData, groups x peaks
    target_peak_idx = set(np.asarray(perturb_out["peak_idx"], dtype=int).tolist())

    # Coordinates from original ATAC var (same features/order expected)
    v = atac_ref.var.copy()
    pchr = v[peak_chrom_col].astype(str).values
    pst = v[peak_start_col].astype(int).values
    pen = v[peak_end_col].astype(int).values
    pnames = atac_ref.var_names.astype(str).values

    # Region mask
    m_reg = (pchr == str(chrom)) & (pen >= int(reg_start)) & (pst <= int(reg_end))
    peak_idx_reg = np.where(m_reg)[0]
    if peak_idx_reg.size == 0:
        raise ValueError(f"No peaks overlap region {region}")

    # Group selection in pseudobulk rows
    if group is None:
        row_idx = None
        base_vec = _to_dense1(base_atac_pb.X[:, peak_idx_reg].mean(axis=0))
        pert_vec = _to_dense1(pert_atac_pb.X[:, peak_idx_reg].mean(axis=0))
        group_label = "ALL_PSEUDOBULK_GROUPS_MEAN"
    else:
        pb_groups = base_atac_pb.obs_names.astype(str).tolist()
        if str(group) not in pb_groups:
            raise KeyError(f"group '{group}' not found in pseudobulk groups: {pb_groups}")
        row_idx = pb_groups.index(str(group))
        base_vec = _to_dense1(base_atac_pb.X[row_idx, peak_idx_reg])
        pert_vec = _to_dense1(pert_atac_pb.X[row_idx, peak_idx_reg])
        group_label = str(group)

    mid = ((pst[peak_idx_reg] + pen[peak_idx_reg]) / 2.0).astype(float)

    df = pd.DataFrame({
        "peak_idx": peak_idx_reg.astype(int),
        "peak_name": pnames[peak_idx_reg],
        "chrom": pchr[peak_idx_reg],
        "start": pst[peak_idx_reg].astype(int),
        "end": pen[peak_idx_reg].astype(int),
        "mid": mid,
        "width": (pen[peak_idx_reg] - pst[peak_idx_reg]).astype(int),
        "atac_base": np.asarray(base_vec, dtype=float),
        "atac_pert": np.asarray(pert_vec, dtype=float),
    })
    df["atac_delta"] = df["atac_pert"] - df["atac_base"]
    df["targeted"] = df["peak_idx"].isin(target_peak_idx)

    # Optional: attach link metadata if available
    if "selection" in perturb_out and "links" in perturb_out["selection"] and perturb_out["selection"]["links"] is not None:
        links = perturb_out["selection"]["links"].copy()
        if not links.empty:
            agg = (
                links.groupby("peak_idx")
                .agg(
                    linked_genes=("gene_name", lambda x: ",".join(sorted(set(map(str, x))))),
                    link_types=("link_type", lambda x: ",".join(sorted(set(map(str, x))))),
                    n_links=("gene_name", "size"),
                )
                .reset_index()
            )
            df = df.merge(agg, on="peak_idx", how="left")
        else:
            df["linked_genes"] = None
            df["link_types"] = None
            df["n_links"] = 0

    df = df.sort_values(["start", "end"]).reset_index(drop=True)
    df.attrs["region"] = region
    df.attrs["group"] = group_label
    return df


# %% [code]
def plot_locus_peaks_along_genome(
    locus_df: pd.DataFrame,
    *,
    mode: str = "delta",   # "delta", "base_pert", "stacked"
    title: str = None,
    show_targeted: bool = True,
    label_top_n: int = 0,  # label top |delta| peaks
    figsize=(14, 4),
):
    """
    Plot peak-level ATAC signal across a locus.
    x-axis = genomic midpoint
    """
    df = locus_df.copy()
    region = locus_df.attrs.get("region", None)
    group_label = locus_df.attrs.get("group", "group")

    x = df["mid"].values
    widths = np.maximum(df["width"].values, 1)

    if title is None:
        if region is not None:
            chrom, st, en = region
            title = f"{group_label} | {chrom}:{st:,}-{en:,}"
        else:
            title = f"{group_label} locus"

    if mode == "delta":
        y = df["atac_delta"].values
        plt.figure(figsize=figsize)
        # all peaks
        plt.vlines(x, 0, y, linewidth=1)
        # emphasize targeted
        if show_targeted and "targeted" in df.columns:
            dt = df[df["targeted"]]
            if not dt.empty:
                plt.vlines(dt["mid"].values, 0, dt["atac_delta"].values, linewidth=2)
                plt.scatter(dt["mid"].values, dt["atac_delta"].values, s=15)
        plt.axhline(0, linewidth=1)
        plt.xlabel("Genomic position")
        plt.ylabel("ΔATAC (pert - base)")
        plt.title(title + " | peak delta")
        plt.tight_layout()
        plt.show()

    elif mode == "base_pert":
        plt.figure(figsize=figsize)
        plt.plot(x, df["atac_base"].values, marker="o", linestyle="-", markersize=2, linewidth=1, label="base")
        plt.plot(x, df["atac_pert"].values, marker="o", linestyle="-", markersize=2, linewidth=1, label="pert")
        if show_targeted and "targeted" in df.columns:
            dt = df[df["targeted"]]
            if not dt.empty:
                ymax = float(np.nanmax(np.r_[df["atac_base"].values, df["atac_pert"].values]))
                for xm in dt["mid"].values:
                    plt.axvline(xm, linewidth=0.5, alpha=0.4)
        plt.xlabel("Genomic position")
        plt.ylabel("ATAC signal")
        plt.title(title + " | baseline vs perturbed")
        plt.legend()
        plt.tight_layout()
        plt.show()

    elif mode == "stacked":
        fig, ax = plt.subplots(2, 1, figsize=(figsize[0], figsize[1] * 1.8), sharex=True)
        ax[0].plot(x, df["atac_base"].values, marker="o", linestyle="-", markersize=2, linewidth=1, label="base")
        ax[0].plot(x, df["atac_pert"].values, marker="o", linestyle="-", markersize=2, linewidth=1, label="pert")
        ax[0].set_ylabel("ATAC")
        ax[0].legend()
        ax[0].set_title(title)

        ax[1].vlines(x, 0, df["atac_delta"].values, linewidth=1)
        if show_targeted and "targeted" in df.columns:
            dt = df[df["targeted"]]
            if not dt.empty:
                ax[1].vlines(dt["mid"].values, 0, dt["atac_delta"].values, linewidth=2)
                ax[1].scatter(dt["mid"].values, dt["atac_delta"].values, s=12)
        ax[1].axhline(0, linewidth=1)
        ax[1].set_ylabel("ΔATAC")
        ax[1].set_xlabel("Genomic position")
        plt.tight_layout()
        plt.show()

    else:
        raise ValueError("mode must be one of {'delta','base_pert','stacked'}")

    # Optional labels for top peaks
    if label_top_n > 0:
        dfl = df.reindex(df["atac_delta"].abs().sort_values(ascending=False).index).head(label_top_n)
        print(dfl[["peak_name", "start", "end", "atac_base", "atac_pert", "atac_delta", "targeted", "linked_genes"]]
              if "linked_genes" in dfl.columns else
              dfl[["peak_name", "start", "end", "atac_base", "atac_pert", "atac_delta", "targeted"]])


# %% [code]
def plot_locus_with_genes(
    locus_df: pd.DataFrame,
    rna: ad.AnnData,
    *,
    region=None,
    target_genes: list = None,      # optional: highlight these
    value_col: str = "atac_delta",  # "atac_delta", "atac_base", "atac_pert"
    figsize=(14, 6),
    max_gene_labels: int = 40,
):
    """
    Two-panel plot:
      top: peak signal along locus
      bottom: simple gene models/TSS markers from rna.var annotation
    Requires rna.var columns: chrom, start, end, strand, gene_name, tss
    """
    df = locus_df.copy()
    if region is None:
        region = locus_df.attrs.get("region", None)
    if region is None:
        raise ValueError("region not provided and not found in locus_df.attrs['region']")
    chrom, reg_start, reg_end = region

    # Select genes in region
    rv = rna.var.copy()
    needed = ["chrom", "start", "end", "strand", "gene_name", "tss"]
    for c in needed:
        if c not in rv.columns:
            raise KeyError(f"rna.var missing '{c}' (needed for gene track plotting)")

    gm = (
        rv["chrom"].astype(str).eq(str(chrom))
        & rv["start"].notna() & rv["end"].notna() & rv["tss"].notna()
        & (rv["end"].astype(float) >= reg_start)
        & (rv["start"].astype(float) <= reg_end)
    )
    gdf = rv.loc[gm, ["chrom", "start", "end", "strand", "gene_name", "tss"]].copy()
    gdf["start"] = gdf["start"].astype(int)
    gdf["end"] = gdf["end"].astype(int)
    gdf["tss"] = gdf["tss"].astype(int)
    gdf["gene_name"] = gdf["gene_name"].astype(str)

    # Dedup by gene_name for plotting cleanliness
    gdf = gdf.drop_duplicates(subset=["gene_name"]).sort_values(["start", "end"]).reset_index(drop=True)

    # Simple row packing to reduce label overlap
    rows_end = []
    gene_rows = []
    for _, row in gdf.iterrows():
        placed = False
        for ridx in range(len(rows_end)):
            if row["start"] > rows_end[ridx]:
                gene_rows.append(ridx)
                rows_end[ridx] = row["end"]
                placed = True
                break
        if not placed:
            rows_end.append(row["end"])
            gene_rows.append(len(rows_end) - 1)
    gdf["track_row"] = gene_rows

    # target highlight
    target_set = set([str(x) for x in target_genes]) if target_genes is not None else set()
    gdf["is_target_gene"] = gdf["gene_name"].isin(target_set)

    fig, (ax1, ax2) = plt.subplots(
        2, 1, figsize=figsize, sharex=True,
        gridspec_kw={"height_ratios": [2.2, 1.2]}
    )

    # --- top panel: peak values ---
    x = df["mid"].values
    y = df[value_col].values
    ax1.vlines(x, 0, y, linewidth=1)
    if "targeted" in df.columns:
        dt = df[df["targeted"]]
        if not dt.empty:
            ax1.vlines(dt["mid"].values, 0, dt[value_col].values, linewidth=2)
            ax1.scatter(dt["mid"].values, dt[value_col].values, s=10)
    if value_col == "atac_delta":
        ax1.axhline(0, linewidth=1)
    ax1.set_ylabel(value_col)
    reg_label = f"{chrom}:{reg_start:,}-{reg_end:,}"
    group_label = locus_df.attrs.get("group", "group")
    ax1.set_title(f"{group_label} | {reg_label}")

    # --- bottom panel: gene track ---
    nrows = max(1, (gdf["track_row"].max() + 1) if not gdf.empty else 1)
    for _, row in gdf.iterrows():
        y0 = nrows - 1 - row["track_row"]  # invert so first row on top
        # gene body
        ax2.hlines(y0, row["start"], row["end"], linewidth=2)
        # TSS marker
        ax2.vlines(row["tss"], y0 - 0.2, y0 + 0.2, linewidth=1.5)

        # strand arrow cue (very simple)
        if row["strand"] == "+":
            ax2.annotate("", xy=(row["end"], y0), xytext=(max(row["start"], row["end"] - (reg_end-reg_start)*0.01), y0),
                         arrowprops=dict(arrowstyle="->", lw=1))
        elif row["strand"] == "-":
            ax2.annotate("", xy=(row["start"], y0), xytext=(min(row["end"], row["start"] + (reg_end-reg_start)*0.01), y0),
                         arrowprops=dict(arrowstyle="->", lw=1))

    # label genes (all or subset if too many)
    if not gdf.empty:
        if len(gdf) > max_gene_labels and len(target_set) > 0:
            label_df = gdf[gdf["is_target_gene"]].copy()
        else:
            label_df = gdf.copy()

        for _, row in label_df.iterrows():
            y0 = nrows - 1 - row["track_row"]
            txt = row["gene_name"]
            ax2.text(row["tss"], y0 + 0.28, txt, fontsize=8, rotation=45, ha="left", va="bottom")

    ax2.set_ylim(-0.8, nrows - 0.2)
    ax2.set_yticks([])
    ax2.set_xlabel("Genomic position")
    ax2.set_ylabel("Genes")
    plt.tight_layout()
    plt.show()


# %% [code]
'''
# GZMB (not present)
region=("chr14", 24_800_000, 25_500_000)
# PRF1
region=("chr10", 70_200_000, 70_800_000)
# NKG7
region=("chr19", 51_300_000, 51_700_000)
# GNLY
region=("chr2", 85_500_000, 86_200_000)
'''

region=("chr2", 85_500_000, 86_200_000)
#region=("chr10", 70_200_000, 70_800_000)
#region=("chr10", 68_000_000, 72_000_000)
#region = ("chr6", 29_000_000, 33_500_000)


# %% [code]
# Inspect available pseudobulk groups
print(hla_on["atac_pseudobulk"].obs_names.tolist())

# Pick one
group_to_plot = "NK"
#group_to_plot = "CD8 TEM_1"
#group_to_plot = "CD14 Mono"

locus_df = build_locus_peak_plot_df(
    perturb_out=hla_on,
    atac_ref=atac,
    #region=("chr6", 28_000_000, 34_000_000),
    #region=("chr15", 44_700_000, 45_100_000),
    region=region,
    group=group_to_plot,
    groupby="cell_type",
)

display(locus_df.head())


# %% [code]
plot_locus_peaks_along_genome(
    locus_df,
    mode="stacked",     # try "delta", "base_pert", "stacked"
    #mode="base_pert",
    #mode="delta",
    label_top_n=20
)


# %% [code]
plot_locus_with_genes(
    locus_df,
    rna=rna,  # or rna_only if you made an RNA-only object with GTF coords
    #region=("chr6", 28_000_000, 34_000_000),
    #region=("chr15", 44_700_000, 45_100_000),    
    #region=("chr14", 24_800_000, 25_500_000),
    region=region,
    #region=("chr2", 0, 30_000_000),
    target_genes=hla_on["target_genes"],   # highlights HLA-related genes
    value_col="atac_delta",                 # or "atac_base"
)


# %% [code]
locus_df_all = build_locus_peak_plot_df(
    perturb_out=hla_on,
    atac_ref=atac,
    #region=("chr6", 28_000_000, 34_000_000),
    #region=("chr15", 44_700_000, 45_100_000),  
    #region=("chr14", 24_800_000, 25_500_000),
    region=region,
    group=None,  # average across pseudobulk groups
)

plot_locus_peaks_along_genome(locus_df_all, mode="delta", label_top_n=15)
plot_locus_with_genes(
    locus_df_all,
    rna=rna,
    #region=("chr6", 28_000_000, 34_000_000),
    #region=("chr15", 44_700_000, 45_100_000),  
    #region=("chr14", 24_800_000, 25_500_000),
    region=region,
    target_genes=hla_on["target_genes"],
    value_col="atac_delta",
)


# %% [code]
def plot_rna_delta_genes_in_locus(
    perturb_out: dict,
    rna_plot: ad.AnnData,
    *,
    region=("chr6", 28_000_000, 34_000_000),
    top_n=None,
    match_target_genes_only=False,
    figsize=(12, 4),
):
    """
    Plot gene-level predicted RNA delta (mean across pseudobulk groups) for genes in locus.
    Uses perturb_out["delta_all_genes"] from run_complex_perturbation_pseudobulk().
    """
    chrom, reg_start, reg_end = region
    dfg = perturb_out["delta_all_genes"].copy()
    rv = rna_plot.var.copy()

    for c in ["chrom", "start", "end", "gene_name"]:
        if c not in rv.columns:
            raise KeyError(f"rna.var missing '{c}' for locus RNA plotting")

    ann = rv[["chrom", "start", "end", "gene_name"]].copy()
    ann["gene_var_name"] = ann.index.astype(str)

    # Merge on gene_var_name
    dfg2 = dfg.merge(ann.reset_index().rename(columns={"index": "gene_var_name"}),
                     on="gene_var_name", how="left", suffixes=("", "_ann"))

    m = (
        dfg2["chrom"].astype(str).eq(str(chrom))
        & dfg2["start"].notna() & dfg2["end"].notna()
        & (dfg2["end"].astype(float) >= reg_start)
        & (dfg2["start"].astype(float) <= reg_end)
    )
    dfl = dfg2.loc[m].copy()
    if match_target_genes_only and "target_genes" in perturb_out:
        tgt = set(map(str, perturb_out["target_genes"]))
        dfl = dfl[dfl["gene_name"].astype(str).isin(tgt)].copy()

    if dfl.empty:
        print("No RNA genes found in region.")
        return

    dfl["mid"] = (dfl["start"].astype(float) + dfl["end"].astype(float)) / 2.0
    dfl = dfl.sort_values("mid")

    if top_n is not None and len(dfl) > top_n:
        dfl = dfl.reindex(dfl["mean_delta_abs"].sort_values(ascending=False).index).head(top_n).sort_values("mid")

    plt.figure(figsize=figsize)
    plt.vlines(dfl["mid"].values, 0, dfl["mean_delta"].values, linewidth=1.5)
    plt.scatter(dfl["mid"].values, dfl["mean_delta"].values, s=18)
    plt.axhline(0, linewidth=1)
    for _, r in dfl.iterrows():
        plt.text(r["mid"], r["mean_delta"], str(r["gene_name"]), fontsize=8, rotation=45, ha="left", va="bottom")
    plt.xlabel("Genomic position")
    plt.ylabel("Mean predicted RNA Δ")
    plt.title(f"RNA delta in locus {chrom}:{reg_start:,}-{reg_end:,}")
    plt.tight_layout()
    plt.show()


# %% [code]
#print(hla_on)


# %% [code]
plot_rna_delta_genes_in_locus(
    hla_on,
    rna_plot=hla_on["rna_pseudobulk"],   # pseudobulk RNA with same var annotation
    #region=("chr6", 28_000_000, 34_000_000),
    #region=("chr15", 44_700_000, 45_100_000), 
    #region=("chr14", 24_800_000, 25_500_000),
    region=region,
    #region = ("chr2", 1, 300_000_000),
    match_target_genes_only=False
)


# %% [code]
plot_rna_delta_genes_in_locus(
    hla_on,
    rna_plot=hla_on["rna_pseudobulk"],   # pseudobulk RNA with same var annotation
    #region=("chr6", 28_000_000, 34_000_000),
    #region=("chr15", 44_700_000, 45_100_000), 
    #region=("chr14", 24_800_000, 25_500_000),
    #region=region,
    #region = ("chr2", 70_000_000, 120_000_000),
    region = ("chr2", 0, 300_000_000),
    match_target_genes_only=False
)


# %% [code]
hla_logit_shift = run_complex_perturbation_pseudobulk(
    model=model,
    atac=atac,
    rna=rna,
    peak_gene_links=peak_gene_links,
    #complex_name_or_genes="hla_class_i_antigen_processing_core",
    #complex_name_or_genes=["B2M"],
    complex_name_or_genes="cytotoxic_effector_extended",
    groupby="cell_type",
    atac_input_layer="imputed_from_rna",    # logits/transformed-like
    pseudobulk_agg="mean",
    mode="add",                             # <- not on/off
    value=50.0,                             # add to logits/transformed values
    clip_min=None,
    clip_max=None,
    strict_input_semantics=True,            # safe because mode='add'
    verbose=True,
    device=device,
)


# %% [code]
import numpy as np
import scipy.sparse as sp

# 1) Confirm only 0/1 in bin layer
'''
Xb = atac_bin.layers["bin"]
if sp.issparse(Xb):
    print("unique nonzero values:", np.unique(Xb.data))
    print("min/max nonzero:", Xb.data.min(), Xb.data.max())
else:
    print("unique values:", np.unique(Xb))

# 2) Confirm sparsity pattern preserved vs counts
Xc = atac_raw.layers["counts"] if "counts" in atac_raw.layers else atac_raw.X
if sp.issparse(Xc) and sp.issparse(Xb):
    same_pattern = (Xc.indptr.shape == Xb.indptr.shape and
                    np.array_equal(Xc.indptr, Xb.indptr) and
                    np.array_equal(Xc.indices, Xb.indices))
    print("same sparse pattern:", same_pattern)

# 3) Confirm nnz unchanged
print("counts nnz:", Xc.nnz if sp.issparse(Xc) else np.count_nonzero(Xc))
print("bin nnz   :", Xb.nnz if sp.issparse(Xb) else np.count_nonzero(Xb))

# 4) Fraction accessible per peak / per cell (should be plausible)
# per-cell number of accessible peaks
cell_access = np.asarray(Xb.sum(axis=1)).ravel()
print("accessible peaks per cell (min/median/max):",
      cell_access.min(), np.median(cell_access), cell_access.max())

# per-peak fraction of cells accessible
peak_frac = np.asarray(Xb.mean(axis=0)).ravel() if sp.issparse(Xb) else Xb.mean(axis=0)
print("peak accessibility fraction (min/median/max):",
      peak_frac.min(), np.median(peak_frac), peak_frac.max())
'''


# %% [code]
import numpy as np
import pandas as pd

def summarize_delta_for_genes(
    perturb_out,
    rna,
    genes_of_interest,
    group=None,                  # e.g. "NK"
    groupby="cell_type",
):
    """
    Compare ΔRNA (pert - base) for selected genes to the distribution of ΔRNA across all genes.
    Supports perturb_out with nested structure:
      perturb_out["result"]["pred_base"], perturb_out["result"]["pred_pert"]
    """
    # --- get predictions (supports nested result dict) ---
    if "result" in perturb_out and isinstance(perturb_out["result"], dict):
        res = perturb_out["result"]
        if "pred_base" in res and "pred_pert" in res:
            pred_base = res["pred_base"]
            pred_pert = res["pred_pert"]
        else:
            raise KeyError("perturb_out['result'] is present but missing 'pred_base'/'pred_pert'")
    else:
        # fallback if already flat
        if "pred_base" in perturb_out and "pred_pert" in perturb_out:
            pred_base = perturb_out["pred_base"]
            pred_pert = perturb_out["pred_pert"]
        else:
            raise KeyError("Could not find predictions. Expected ['result']['pred_base'/'pred_pert'] or top-level 'pred_base'/'pred_pert'.")

    pred_base = np.asarray(pred_base)
    pred_pert = np.asarray(pred_pert)
    delta = pred_pert - pred_base  # (n_groups, n_genes)

    # --- determine group index ---
    group_names = None
    if "atac_pseudobulk" in perturb_out:
        group_names = list(perturb_out["atac_pseudobulk"].obs_names)
    elif "rna_pseudobulk" in perturb_out:
        group_names = list(perturb_out["rna_pseudobulk"].obs_names)
    elif "result" in perturb_out and isinstance(perturb_out["result"], dict):
        # if nested anndata is there instead
        if "base_atac" in perturb_out["result"]:
            group_names = list(perturb_out["result"]["base_atac"].obs_names)

    if group_names is None:
        if delta.shape[0] != 1 and group is None:
            raise ValueError("Could not infer group names; please provide group.")
        group_idx = 0 if group is None else int(group)
        group_label = str(group if group is not None else "group_0")
    else:
        if group is None:
            if len(group_names) != 1:
                raise ValueError(f"Please specify group. Available groups: {group_names}")
            group_idx = 0
        else:
            group_idx = group_names.index(group)
        group_label = group_names[group_idx]

    # --- gene indexing (prefer gene_name if available; fallback var_names) ---
    if "gene_name" in rna.var.columns:
        gene_name_series = rna.var["gene_name"].astype(str)
        # keep first occurrence if duplicates
        gene_name_to_idx = pd.Series(np.arange(rna.n_vars), index=gene_name_series).groupby(level=0).first()
        goi = [g for g in genes_of_interest if g in gene_name_to_idx.index]
        missing = [g for g in genes_of_interest if g not in gene_name_to_idx.index]
        gene_idx = gene_name_to_idx.loc[goi].values.astype(int)
        gene_labels = goi
    else:
        var_names = pd.Index(rna.var_names.astype(str))
        goi = [g for g in genes_of_interest if g in var_names]
        missing = [g for g in genes_of_interest if g not in var_names]
        gene_idx = var_names.get_indexer(goi)
        gene_labels = goi

    if len(gene_labels) == 0:
        raise ValueError(f"None of the genes_of_interest were found. Missing: {missing}")

    # --- per-group delta stats across all genes ---
    d = delta[group_idx, :]
    abs_d = np.abs(d)

    mu = float(np.mean(d))
    sd = float(np.std(d, ddof=1)) if d.size > 1 else 0.0
    abs_mu = float(np.mean(abs_d))
    abs_sd = float(np.std(abs_d, ddof=1)) if d.size > 1 else 0.0

    # --- selected genes summary ---
    rows = []
    for g, j in zip(gene_labels, gene_idx):
        db = float(pred_base[group_idx, j])
        dp = float(pred_pert[group_idx, j])
        dd = float(d[j])
        z = (dd - mu) / sd if sd > 0 else np.nan
        z_abs = (abs(dd) - abs_mu) / abs_sd if abs_sd > 0 else np.nan
        pct_of_baseline = (dd / db * 100.0) if db != 0 else np.nan
        abs_pct = (abs(dd) / abs(db) * 100.0) if db != 0 else np.nan
        percentile_abs = float((abs_d <= abs(dd)).mean())

        rows.append({
            "group": group_label,
            "gene": g,
            "pred_base": db,
            "pred_pert": dp,
            "delta": dd,
            "abs_delta": abs(dd),
            "delta_pct_of_base": pct_of_baseline,
            "abs_delta_pct_of_base": abs_pct,
            "delta_zscore_vs_all_genes": z,
            "abs_delta_zscore_vs_all_genes": z_abs,
            "abs_delta_percentile_vs_all_genes": percentile_abs,
        })

    out_df = pd.DataFrame(rows).sort_values(
        ["abs_delta_zscore_vs_all_genes", "abs_delta"],
        ascending=False
    )

    summary = {
        "group": group_label,
        "n_genes_total": int(d.size),
        "mean_delta_all_genes": mu,
        "sd_delta_all_genes": sd,
        "mean_abs_delta_all_genes": abs_mu,
        "sd_abs_delta_all_genes": abs_sd,
        "q95_abs_delta": float(np.quantile(abs_d, 0.95)),
        "q99_abs_delta": float(np.quantile(abs_d, 0.99)),
        "missing_genes": missing,
    }

    return out_df, summary


# %% [code]
#print(hla_on)


# %% [code]
df_b2m, summary_b2m = summarize_delta_for_genes(
    perturb_out=hla_on,          # or b2m_on / cyto_off etc.
    rna=rna,
    genes_of_interest=["GNLY"],
    #genes_of_interest=["PRF1"],
    group="NK",                  # change to your group of interest
    #group="CD8 TEM_1",
)

print(pd.Series(summary_b2m))
display(df_b2m)


# %% [code]
genes = ["B2M", "HLA-A", "HLA-B", "HLA-C", "TAP1", "TAP2", "TAPBP"]
df_prog, summary_prog = summarize_delta_for_genes(
    perturb_out=hla_on,
    rna=rna,
    genes_of_interest=genes,
    #group="CD14 Mono",
    group="CD8 TEM_1",
)
display(df_prog.sort_values("delta", ascending=False))
print(pd.Series(summary_prog))


# %% [code]


# %% [code]


# %% [code]


# %% [code]


# %% [code]


# %% [code]


# %% [code]


# %% [code]


# %% [code]


# %% [code]


# %% [code]


# %% [code]


# %% [code]
```
