## Supplemental Code for "Unifying multimodal single-cell data with a mixture-of-experts *β*-variational autoencoder framework": pyproject.toml

```
[build-system]
requires = ["setuptools>=68", "wheel"]
build-backend = "setuptools.build_meta"

[project]
name = "univi"
version = "0.4.7"
description = "UniVI: a scalable multi-modal variational autoencoder toolkit for seamless integration and analysis of multimodal single-cell data."
readme = "README.md"
requires-python = ">=3.10"
license = { file = "LICENSE" }
authors = [
  { name = "Andrew J. Ashford", email = "" }
]

dependencies = [
  "numpy>=1.26",
  "scipy>=1.11",
  "pandas>=2.1",
  "anndata>=0.10",
  "scanpy>=1.11",
  "torch>=2.2",
  "scikit-learn>=1.3",
  "h5py>=3.10",
  "pyyaml>=6.0",
  "matplotlib>=3.8",
  "seaborn>=0.13",
  "igraph>=0.11",
  "leidenalg>=0.10",
  "tqdm>=4.66",
  "openpyxl>=3.1",
]

classifiers = [
  "License :: OSI Approved :: MIT License",
  "Programming Language :: Python :: 3",
  "Programming Language :: Python :: 3.10",
  "Programming Language :: Python :: 3 :: Only",
]

[project.urls]
Homepage = "https://github.com/Ashford-A/UniVI"
Repository = "https://github.com/Ashford-A/UniVI"
"Bug Tracker" = "https://github.com/Ashford-A/UniVI/issues"

[tool.setuptools.packages.find]
include = ["univi*"]

[project.optional-dependencies]
bench = ["harmonypy>=0.0.9"]

[project.scripts]
univi = "univi.cli:main"
```
