## Supplemental Code for "Unifying multimodal single-cell data with a mixture-of-experts *β*-variational autoencoder framework": README.md

# UniVI

[![PyPI version](https://img.shields.io/pypi/v/univi?v=0.4.7)](https://pypi.org/project/univi/)
[![pypi downloads](https://img.shields.io/pepy/dt/univi?label=pypi%20downloads)](https://pepy.tech/project/univi)
[![Conda version](https://img.shields.io/conda/vn/conda-forge/univi?cacheSeconds=300)](https://anaconda.org/conda-forge/univi)
[![conda-forge downloads](https://img.shields.io/conda/dn/conda-forge/univi?label=conda-forge%20downloads&cacheSeconds=300)](https://anaconda.org/conda-forge/univi)
[![PyPI - Python Version](https://img.shields.io/pypi/pyversions/univi.svg?v=0.4.7)](https://pypi.org/project/univi/)

<picture>
  <source media="(prefers-color-scheme: dark)"
          srcset="https://raw.githubusercontent.com/Ashford-A/UniVI/v0.4.7/assets/figures/univi_overview_dark.png">
  <img src="https://raw.githubusercontent.com/Ashford-A/UniVI/v0.4.7/assets/figures/univi_overview_light.png"
       alt="UniVI overview and evaluation roadmap"
       width="100%">
</picture>

**UniVI** is a **multi-modal variational autoencoder (VAE)** toolkit for aligning and integrating single-cell modalities such as **RNA**, **ADT (CITE-seq)**, **ATAC**, and **coverage-aware / proportion-like assays** (e.g., **single-cell methylome** features).

Common use cases:

- **Joint embedding** of paired multimodal data (CITE-seq, Multiome, TEA-seq)
- **Bridge mapping / projection** of unimodal cohorts into a paired latent
- **Cross-modal imputation** (RNA→ADT, ATAC→RNA, RNA→methylome, …)
- **Denoising / reconstruction** with likelihood-aware decoders
- **Generating biologically-relevant samples** due to the generative nature of VAEs
- **Evaluation** (FOSCTTM, Recall@k, mixing/entropy, label transfer, clustering, basic MoE gating diagnostics)

Advanced/experimental use cases (all optional, model can be run entirely without these):

- **Supervised heads** (either a decoder classification head or a whole categorical encoder/decoder model VAE, treated as a modality)
- **Expanded MoE gating diagnostics** (setting a simple gating network during training)
- **Transformer encoders** (experimental, added for exploratory analysis)
- **Fused transformer latent space** (even more experimental, added for exploratory analysis/future model expansion)

---

## Preprint

If you use UniVI in your work, please cite:

> Ashford AJ, Enright T, Somers J, Nikolova O, Demir E.  
> **Unifying multimodal single-cell data with a mixture-of-experts β-variational autoencoder framework.**  
> *bioRxiv* (2025; updated 2026). doi: [10.1101/2025.02.28.640429](https://doi.org/10.1101/2025.02.28.640429)

```bibtex
@article{Ashford2025UniVI,
  title   = {Unifying multimodal single-cell data with a mixture-of-experts β-variational autoencoder framework},
  author  = {Ashford, A. J. and Enright, T. and Somers, J. and Nikolova, O. and Demir, E.},
  journal = {bioRxiv},
  date    = {2025},
  doi     = {10.1101/2025.02.28.640429},
  url     = {https://www.biorxiv.org/content/10.1101/2025.02.28.640429},
  note    = {Preprint (updated 2026)}
}
````

---

## Installation

### PyPI

```bash
pip install univi
```

> UniVI requires PyTorch. If `import torch` fails, install PyTorch for your platform/CUDA from [PyTorch's official install instructions](https://pytorch.org/get-started/locally/).

### Conda / mamba

```bash
conda install -c conda-forge univi
# or
mamba install -c conda-forge univi
```

### Development install (from source)

```bash
git clone https://github.com/Ashford-A/UniVI.git
cd UniVI

conda env create -f envs/univi_env.yml
conda activate univi_env

pip install -e .
```

---

## Data expectations

UniVI expects **per-modality AnnData** objects.

* Each modality is an `AnnData`
* For paired settings, modalities share the same cells (`obs_names`, same order)
* Raw counts often live in `.layers["counts"]`
* Model inputs typically live in `.X` (or `.obsm["X_*"]` for ATAC LSI)
* Model input is a dictionary of these `AnnData` objects with the dictionary key specifying the modality (e.g. `rna`, `adt`, `atac`). These keys are used later for evaluation functions (cross-reconstruction etc.).

Recommended convention:

* `.layers["counts"]` = raw counts / raw signal
* `.X` / `.obsm["X_*"]` = model input space (log1p RNA, CLR ADT, LSI ATAC, methyl fractions, etc.)
* `.layers["denoised_*"]` / `.layers["imputed_*"]` = UniVI outputs

---

## Quickstart (Python / Jupyter)

Minimal "notebook path": load paired AnnData → preprocess → train → encode/evaluate → plot.

The sections below walk through a complete CITE-seq (RNA + ADT) example.
All patterns generalize to Multiome (RNA + ATAC), TEA-seq (RNA + ADT + ATAC),
and any other paired combination supported by UniVI.

---

### 0) Imports

```python
import numpy as np
import scanpy as sc
import torch
from torch.utils.data import DataLoader, Subset

from univi import UniVIMultiModalVAE, ModalityConfig, UniVIConfig, TrainingConfig
from univi.data import MultiModalDataset, align_paired_obs_names, collate_multimodal_xy_recon
from univi.trainer import UniVITrainer
```

> **`collate_multimodal_xy_recon`** is the required collate function for `DataLoader` when
> using `MultiModalDataset`. It correctly handles the `(x, recon_targets)` batch format
> expected by the trainer, including coverage-aware modalities such as beta-binomial methylome.
> Always pass it as `collate_fn=collate_multimodal_xy_recon` when constructing your loaders.

---

### 1) Load paired AnnData

For CITE-seq data:

```python
rna = sc.read_h5ad("path/to/rna_citeseq.h5ad")
adt = sc.read_h5ad("path/to/adt_citeseq.h5ad")
```

For Multiome / SHARE-seq (RNA + ATAC):

```python
rna  = sc.read_h5ad("path/to/rna_multiome.h5ad")
atac = sc.read_h5ad("path/to/atac_multiome.h5ad")
```

For tri-modal TEA-seq / DOGMA-seq / ASAP-seq:

```python
rna  = sc.read_h5ad("path/to/rna.h5ad")
adt  = sc.read_h5ad("path/to/adt.h5ad")
atac = sc.read_h5ad("path/to/atac.h5ad")
```

---

### 2) Preprocess each modality

> After preprocessing, set `.X` to the model input space and keep raw counts in
> `.layers["counts"]`. Match the `likelihood` in `ModalityConfig` to your `.X` space
> (see the likelihood guidance table in step 4).

**RNA** — log-normalize, select HVGs, scale:

```python
rna.layers["counts"] = rna.X.copy()

rna.var["mt"] = rna.var_names.str.upper().str.startswith("MT-")
sc.pp.calculate_qc_metrics(rna, qc_vars=["mt"], percent_top=None, log1p=False, inplace=True)

sc.pp.normalize_total(rna, target_sum=1e4)
sc.pp.log1p(rna)
rna.raw = rna  # snapshot log-space for plotting/DE

sc.pp.highly_variable_genes(rna, flavor="seurat_v3", n_top_genes=2000, subset=True)
sc.pp.scale(rna, max_value=10)
```

**ADT** — CLR per cell, scale per protein:

```python
adt.layers["counts"] = adt.X.copy()

def clr_per_cell(X):
    X = X.toarray() if hasattr(X, "toarray") else np.asarray(X)
    logX = np.log1p(X)
    return logX - logX.mean(axis=1, keepdims=True)

adt.X = clr_per_cell(adt.layers["counts"])
sc.pp.scale(adt, zero_center=True, max_value=10)
```

**ATAC** — TF-IDF → LSI, drop first component:

```python
atac.layers["counts"] = atac.X.copy()

def tfidf(X):
    X = X.tocsr() if hasattr(X, "tocsr") else X
    cell_sum = np.asarray(X.sum(axis=1)).ravel()
    cell_sum[cell_sum == 0] = 1.0
    tf = X.multiply(1.0 / cell_sum[:, None])
    df = np.asarray((X > 0).sum(axis=0)).ravel()
    idf = np.log1p(X.shape[0] / (1.0 + df))
    return tf.multiply(idf)

X_tfidf = tfidf(atac.layers["counts"])

from sklearn.decomposition import TruncatedSVD
svd = TruncatedSVD(n_components=101, random_state=0)
X_lsi = svd.fit_transform(X_tfidf)
atac.obsm["X_lsi"] = X_lsi[:, 1:]  # drop first component (depth correlated)
```

**Post-preprocessing: assemble `adata_dict`**

```python
# Sanity check (CITE-seq)
assert rna.n_obs == adt.n_obs and np.all(rna.obs_names == adt.obs_names)

# CITE-seq
adata_dict = {"rna": rna, "adt": adt}

# Multiome/SHARE-seq
# adata_dict = {"rna": rna, "atac": atac}

# Tri-modal
# adata_dict = {"rna": rna, "adt": adt, "atac": atac}

# Unimodal VAE
# adata_dict = {"atac": atac}

align_paired_obs_names(adata_dict)  # ensures matching obs_names and order
```

> **Avoiding data leakage:** if you want to run UniVI inductively, apply feature selection,
> scaling, and any learned transforms (e.g., PCA/LSI) on the training set only, then apply
> the training-set-derived parameters to validation and test sets.

---

### 3) Dataset + DataLoaders

**Device detection** (CUDA → MPS → XPU → CPU):

```python
device = (
    "cuda" if torch.cuda.is_available() else
    ("mps" if getattr(torch.backends, "mps", None) is not None
               and torch.backends.mps.is_available() else
     ("xpu" if hasattr(torch, "xpu") and torch.xpu.is_available() else
      "cpu"))
)
```

**Build dataset:**

```python
dataset = MultiModalDataset(
    adata_dict=adata_dict,
    device=None,                  # dataset yields CPU tensors; model handles GPU transfer
    X_key_by_mod={
        "rna" : "X",              # uses rna.X
        "adt" : "X",              # uses adt.X
        # "atac": "obsm:X_lsi",  # uses atac.obsm["X_lsi"]
    },
)
```

**Train / val / test split (80 / 10 / 10):**

```python
n = rna.n_obs
idx = np.arange(n)
rng = np.random.default_rng(0)
rng.shuffle(idx)

n_train = int(0.8 * n)
n_val   = int(0.1 * n)

train_idx = idx[:n_train]
val_idx   = idx[n_train : n_train + n_val]
test_idx  = idx[n_train + n_val :]

# Save split indices for reproducibility
np.savez("splits_seed0.npz", train_idx=train_idx, val_idx=val_idx, test_idx=test_idx)
```

**Construct loaders** (always pass `collate_fn=collate_multimodal_xy_recon`):

```python
train_loader = DataLoader(
    Subset(dataset, train_idx),
    batch_size=256,
    shuffle=True,
    num_workers=0,
    collate_fn=collate_multimodal_xy_recon,
)
val_loader = DataLoader(
    Subset(dataset, val_idx),
    batch_size=256,
    shuffle=False,
    num_workers=0,
    collate_fn=collate_multimodal_xy_recon,
)
test_loader = DataLoader(
    Subset(dataset, test_idx),
    batch_size=256,
    shuffle=False,
    num_workers=0,
    collate_fn=collate_multimodal_xy_recon,
)
```

---

### 4) Model config + train

```python
univi_cfg = UniVIConfig(
    latent_dim=30,
    beta=1.15,
    gamma=3.25,
    encoder_dropout=0.10,
    decoder_dropout=0.05,
    encoder_batchnorm=True,
    decoder_batchnorm=False,
    kl_anneal_start=50,
    kl_anneal_end=100,
    align_anneal_start=75,
    align_anneal_end=125,
    modalities=[
        ModalityConfig(
            name="rna",
            input_dim=rna.n_vars,
            encoder_hidden=[1024, 512, 256, 128],
            decoder_hidden=[128, 256, 512, 1024],
            likelihood="gaussian",
        ),
        ModalityConfig(
            name="adt",
            input_dim=adt.n_vars,
            encoder_hidden=[256, 128, 64],
            decoder_hidden=[64, 128, 256],
            likelihood="gaussian",
        ),
    ],
)
```

#### `ModalityConfig` likelihood guidance

| Modality | `.X` / input space | Recommended `likelihood` |
|---|---|---|
| RNA | raw counts | `"nb"` or `"zinb"` (many zeros) |
| RNA | `normalize_total` + `log1p` (+ scale) | `"gaussian"` |
| ADT | CLR (+ scale) | `"gaussian"` |
| ADT | raw counts | `"nb"` (or `"gaussian"` post-CLR) |
| ATAC | binarized peaks | `"bernoulli"` |
| ATAC | raw peak counts | `"poisson"` |
| ATAC | LSI / reduced | `"gaussian"` |
| Methylome | fractions / beta values (0–1) | `"beta"` |
| Methylome | successes + coverage | `"beta_binomial"` (see methylome quickstart) |
| Methylome | PCA / LSI reduced | `"gaussian"` |

> **Manuscript-style tip:** Gaussian decoders on normalized feature spaces often produce
> the most cell-to-cell aligned latent spaces for integration-focused use cases.

**Training config and model construction:**

```python
train_cfg = TrainingConfig(
    n_epochs=3000,
    batch_size=256,
    lr=1e-3,
    weight_decay=1e-4,
    device=device,
    early_stopping=True,
    best_epoch_warmup=50,
    patience=50,
    log_every=25,
)

model = UniVIMultiModalVAE(
    univi_cfg,
    loss_mode="v1",       # "v1" (manuscript default) or "v2" (lite/fused)
    v1_recon="avg",
    normalize_v1_terms=True,
).to(device)
```

> **`loss_mode` note:**
> - `"v1"` (recommended): per-modality posteriors + cross/self/avg reconstruction + posterior alignment. Used in the manuscript.
> - `"v2"` ("lite"): fused posterior (MoE/PoE or fused transformer) + per-modality recon + β·KL + γ·L2 alignment. Useful for more loosely-paired/"pseudopaired" settings, but weaker overall cell-to-cell correspondence.

**Train:**

```python
trainer = UniVITrainer(
    model=model,
    train_loader=train_loader,
    val_loader=val_loader,
    train_cfg=train_cfg,
    device=device,
)

history = trainer.fit()
print("Best epoch:", getattr(trainer, "best_epoch", None))
```

---

### 5) Saving and loading trained models

**Save:**

```python
ckpt = {
    "model_state_dict": model.state_dict(),
    "model_config":     univi_cfg,
    "train_cfg":        train_cfg,
    "history":          getattr(trainer, "history", None),
    "best_epoch":       getattr(trainer, "best_epoch", None),
}
torch.save(ckpt, "./saved_models/univi_model_state.pt")
```

**Load:**

```python
import torch
from univi import UniVIMultiModalVAE

ckpt  = torch.load("./saved_models/univi_model_state.pt", map_location=device, weights_only=False)
model = UniVIMultiModalVAE(ckpt["model_config"]).to(device)
model.load_state_dict(ckpt["model_state_dict"])
model.eval()
print("Best epoch:", ckpt.get("best_epoch"))
```

---

### 6) Reproducibility helpers

UniVI provides utilities to lock down randomness across Python, NumPy, and PyTorch
(including CUDA), important for benchmarking and manuscript-level reproducibility.

```python
from univi.utils.seed import set_seed

set_seed(0)  # seeds Python, NumPy, torch, and torch.cuda
```

Call `set_seed` before dataset construction, model initialization, and training to ensure
results are deterministic across runs (given the same hardware and PyTorch version).

You can also snapshot environment + hyperparameter + dataset metadata to a supplemental table:

```python
from univi.diagnostics import export_supplemental_table_s1

export_supplemental_table_s1(
    model=model,
    train_cfg=train_cfg,
    adata_dict=adata_dict,
    outpath="./tables/Supplemental_Table_S1.xlsx",
)
```

---

## After training: encoding, evaluation, and plotting

### 0a) Post-training imports

```python
import numpy as np
import scipy.sparse as sp
import torch

from univi.evaluation import (
    encode_adata,
    encode_fused_adata_pair,
    cross_modal_predict,
    denoise_adata,
    denoise_from_multimodal,
    evaluate_alignment,
    reconstruction_metrics,
    generate_from_latent,
    fit_label_latent_gaussians,
    sample_latent_by_label,
    evaluate_cross_reconstruction,
)
from univi.plotting import (
    set_style,
    umap,
    umap_by_modality,
    compare_raw_vs_denoised_umap_features,
    plot_confusion_matrix,
    write_gates_to_obs,
    plot_moe_gate_summary,
    plot_reconstruction_error_summary,
    plot_featurewise_reconstruction_scatter,
)

set_style(font_scale=1.2, dpi=150)

def to_dense(X):
    return X.toarray() if sp.issparse(X) else np.asarray(X)
```

### 0b) Subset to test set before evaluating

```python
# Load the saved splits
splits = np.load("splits_seed0.npz")
test_idx = splits["test_idx"]

rna_test = rna[test_idx].copy()
adt_test = adt[test_idx].copy()

assert rna_test.n_obs == adt_test.n_obs
assert np.array_equal(rna_test.obs_names, adt_test.obs_names)
```

---

### 1) Encode a modality into latent space

Use when you have one observed modality at a time (RNA-only, ADT-only, etc.):

```python
Z_rna = encode_adata(
    model,
    adata=rna_test,
    modality="rna",
    device=device,
    layer=None,           # uses .X by default
    X_key="X",
    batch_size=1024,
    latent="moe_mean",    # options: "moe_mean", "moe_sample", "modality_mean", "modality_sample"
    random_state=0,
)
rna_test.obsm["X_univi"] = Z_rna

umap(
    rna_test,
    obsm_key="X_univi",
    color=["celltype.l2", "batch"],
    legend="outside",
    legend_subset_topk=25,
    savepath="umap_rna_univi.png",
    show=False,
)
```

---

### 2) Encode a fused multimodal latent

When you have multiple observed modalities for the same cells:

```python
fused = encode_fused_adata_pair(
    model,
    adata_by_mod={"rna": rna_test, "adt": adt_test},
    device=device,
    batch_size=1024,
    use_mean=True,
    return_gates=True,
    return_gate_logits=True,
    write_to_adatas=True,
    fused_obsm_key="X_univi_fused",
    gate_prefix="gate",
)
# fused["Z_fused"] → (n_cells, latent_dim)
# fused["gates"]   → (n_cells, n_modalities)

umap(
    rna_test,
    obsm_key="X_univi_fused",
    color=["celltype.l2", "batch"],
    legend="outside",
    savepath="umap_fused.png",
    show=False,
)

umap_by_modality(
    {"rna": rna_test, "adt": adt_test},
    obsm_key="X_univi_fused",
    color=["univi_modality", "celltype.l2"],
    legend="outside",
    size=8,
    savepath="umap_fused_both_modalities.png",
    show=False,
)
```

---

### 3) Cross-modal prediction (imputation)

Encode a source modality and decode into a target modality.
UniVI automatically handles decoder output types (Gaussian tensor, NB/ZINB/Poisson dicts, Beta/Beta-Binomial dicts)
and returns a mean-like prediction matrix ready for evaluation or storage.

```python
# RNA → ADT
adt_hat_from_rna = cross_modal_predict(
    model,
    adata_src=rna_test,
    src_mod="rna",
    tgt_mod="adt",
    device=device,
    layer=None,
    X_key="X",
    batch_size=512,
    use_moe=True,
)
adt_test.layers["imputed_from_rna"] = adt_hat_from_rna
```

---

### 4) Denoising

**Option A — self-denoise a single modality:**

```python
denoise_adata(
    model,
    adata=rna_test,
    modality="rna",
    device=device,
    out_layer="denoised_self",
    overwrite_X=False,
    batch_size=512,
)
```

**Option B — fused multimodal denoising:**

```python
denoise_adata(
    model,
    adata=rna_test,
    modality="rna",
    device=device,
    out_layer="denoised_fused",
    overwrite_X=False,
    batch_size=512,
    adata_by_mod={"rna": rna_test, "adt": adt_test},
    layer_by_mod={"rna": None, "adt": None},
    X_key_by_mod={"rna": "X", "adt": "X"},
    use_mean=True,
)

compare_raw_vs_denoised_umap_features(
    rna_test,
    obsm_key="X_univi",
    features=["MS4A1", "CD3D", "NKG7"],
    raw_layer=None,
    denoised_layer="denoised_fused",
    savepath="umap_raw_vs_denoised.png",
    show=False,
)
```

---

### 5) Reconstruction and imputation error

**Basic metrics on two matrices:**

```python
true = to_dense(adt_test.X)
pred = adt_test.layers["imputed_from_rna"]

m = reconstruction_metrics(true, pred)
print("MSE mean:", m["mse_mean"])
print("Pearson mean:", m["pearson_mean"])
```

**One-call cross-reconstruction evaluation:**

```python
rep = evaluate_cross_reconstruction(
    model,
    adata_src=rna_test,
    adata_tgt=adt_test,
    src_mod="rna",
    tgt_mod="adt",
    device=device,
    src_layer=None,
    tgt_layer=None,
    batch_size=512,
    feature_names=None,   # optionally restrict to a feature subset
)
print(rep["summary"])

plot_reconstruction_error_summary(
    rep,
    title="RNA → ADT imputation error",
    savepath="recon_error_summary.png",
    show=False,
)

plot_featurewise_reconstruction_scatter(
    rep,
    features=["CD3", "CD4", "MS4A1"],
    savepath="recon_scatter_selected_features.png",
    show=False,
)
```

---

### 6) Alignment evaluation

```python
metrics = evaluate_alignment(
    Z1=rna_test.obsm["X_univi"],
    Z2=adt_test.obsm["X_univi"],
    metric="euclidean",
    recall_ks=(1, 5, 10),
    k_mixing=20,
    k_entropy=30,
    labels_source=rna_test.obs["celltype.l2"].to_numpy(),
    labels_target=adt_test.obs["celltype.l2"].to_numpy(),
    compute_bidirectional_transfer=True,
    k_transfer=15,
    json_safe=True,
)

plot_confusion_matrix(
    np.asarray(metrics["label_transfer_cm"]),
    labels=np.asarray(metrics["label_transfer_label_order"]),
    title="Label transfer (RNA → ADT)",
    normalize="true",
    savepath="label_transfer_confusion.png",
    show=False,
)
```

---

### 7) Generate new data from the latent space

UniVI decoders define a per-modality likelihood. Generation works by:
1. picking latent samples `z` (from the prior or a conditional distribution),
2. decoding with the target modality's decoder,
3. returning a mean-like reconstruction or optionally sampling from the likelihood.

**Unconditional generation from standard normal prior:**

```python
Xgen = generate_from_latent(
    model,
    n=5000,
    target_mod="rna",
    device=device,
    z_source="prior",
    return_mean=True,
    sample_likelihood=False,
)
# Xgen shape: (5000, n_genes)
```

**Cell-type–conditioned generation via empirical latent neighborhoods:**

```python
Z      = rna_test.obsm["X_univi"]
labels = rna_test.obs["celltype.l2"].to_numpy()

# Fit a per-label Gaussian in latent space
label_gauss = fit_label_latent_gaussians(Z, labels)

# Sample latent points for a chosen label
z_B = sample_latent_by_label(label_gauss, label="B cell", n=2000, random_state=0)

# Decode to RNA space
X_B = generate_from_latent(
    model,
    z=z_B,
    target_mod="rna",
    device=device,
    return_mean=True,
)
```

> If you do not have cell-type annotations, you can cluster `Z` (e.g., k-means),
> fit cluster-specific Gaussians, and sample by cluster ID instead.

---

### 8) MoE gating diagnostics

UniVI can report per-cell modality contribution weights for the analytic fusion path.
Two notions of "who contributed" are available:

- **Precision-only** (always available): derived from each modality's posterior uncertainty.
- **Router × precision** (when a learnable gating network is present): combines router
  probabilities with precision to produce contribution weights.

> This section applies to **analytic fusion** (Gaussian experts).
> If you use the fused transformer posterior, gates may be unavailable or not meaningful.

**Compute per-cell contribution weights:**

```python
from univi.evaluation import encode_moe_gates_from_tensors

gate = encode_moe_gates_from_tensors(
    model,
    x_dict={"rna": to_dense(rna_test.X), "adt": to_dense(adt_test.X)},
    device=device,
    batch_size=1024,
    modality_order=["rna", "adt"],
    kind="router_x_precision",  # falls back to "effective_precision" if no router
    return_logits=True,
)

W    = gate["weights"]          # (n_cells, n_modalities), rows sum to 1
mods = gate["modality_order"]

print("Requested kind:", gate.get("requested_kind"))
print("Effective kind:", gate.get("kind"))
print("Per-modality mean:", gate.get("per_modality_mean"))
```

**Write to `.obs` and plot:**

```python
write_gates_to_obs(
    rna_test,
    gates=W,
    modality_names=mods,
    gate_prefix="moe_gate",
    gate_logits=gate.get("logits"),
)

plot_moe_gate_summary(
    rna_test,
    gate_prefix="moe_gate",
    groupby="celltype.l3",
    agg="mean",
    savepath="moe_gates_by_celltype.png",
    show=False,
)
```

**Optionally log gates alongside alignment metrics:**

```python
metrics["moe_gates"] = {
    "kind":              gate.get("kind"),
    "requested_kind":    gate.get("requested_kind"),
    "modality_order":    mods,
    "per_modality_mean": gate.get("per_modality_mean"),
}
```

---

## Quickstart: RNA + methylome (beta-binomial with recon_targets)

For coverage-aware methylome data where you have both methylated counts and total coverage:

```python
rna  = sc.read_h5ad("path/to/rna.h5ad")
meth = sc.read_h5ad("path/to/meth.h5ad")
# meth.X                         → model input (fractions or embeddings)
# meth.layers["meth_successes"]  → methylated counts
# meth.layers["meth_total_count"]→ coverage / trials

adata_dict = align_paired_obs_names({"rna": rna, "meth": meth})

recon_targets_spec = {
    "meth": {
        "successes_layer":   "meth_successes",
        "total_count_layer": "meth_total_count",
    }
}

dataset = MultiModalDataset(
    adata_dict=adata_dict,
    X_key="X",
    device=None,
    recon_targets_spec=recon_targets_spec,
)

train_loader = DataLoader(
    dataset,
    batch_size=256,
    shuffle=True,
    num_workers=0,
    collate_fn=collate_multimodal_xy_recon,
)

univi_cfg = UniVIConfig(
    latent_dim=30,
    beta=1.15,
    gamma=3.25,
    modalities=[
        ModalityConfig("rna",  rna.n_vars,  [1024, 512, 256, 128], [128, 256, 512, 1024], likelihood="gaussian"),
        ModalityConfig("meth", meth.n_vars, [512, 256, 128],       [128, 256, 512],       likelihood="beta_binomial"),
    ],
)

train_cfg = TrainingConfig(n_epochs=2000, batch_size=256, lr=1e-3, weight_decay=1e-4,
                           device=device, log_every=25)

model = UniVIMultiModalVAE(univi_cfg, loss_mode="v1", v1_recon="avg",
                           normalize_v1_terms=True).to(device)
trainer = UniVITrainer(model=model, train_loader=train_loader, val_loader=None,
                       train_cfg=train_cfg, device=device)
history = trainer.fit()
```

> When `recon_targets` are present in the batch, `UniVITrainer` automatically forwards them
> into `model(..., recon_targets=...)`.

---

## Bridge mapping / zero-shot projection (inductive use case)

UniVI can project unimodal cells that were not seen during training into the shared latent
space ("bridge mapping"), without retraining the model. This is useful for:

- projecting an external RNA-only cohort into a RNA+ADT latent trained on a reference,
- mapping a new patient's ATAC into an existing RNA+ATAC integration,
- any scenario where only a subset of modalities is available at inference time.

```python
# Load a held-out RNA-only cohort (no ADT available)
rna_query = sc.read_h5ad("path/to/rna_query.h5ad")

# Apply the same preprocessing as the reference RNA (HVG subset, scale params, etc.)
# ...

# Encode using only the RNA encoder (modality-specific posterior)
Z_query = encode_adata(
    model,
    adata=rna_query,
    modality="rna",
    device=device,
    layer=None,
    X_key="X",
    batch_size=1024,
    latent="modality_mean",   # use the RNA-specific posterior mean
    random_state=0,
)
rna_query.obsm["X_univi"] = Z_query
```

> **Fine-tuning (optional):** If you have a small amount of paired data in the query cohort,
> you can fine-tune the model on it (typically with a lower learning rate, e.g., `lr=1e-4`,
> and fewer epochs) to adapt the latent to query-specific batch effects. See
> `notebooks/GR/Fig07_AML_bridge_finetuning.ipynb`
> for an end-to-end example.

---

## CLI training and evaluation (script-based workflow)

For cluster/HPC use or for fully reproducible manuscript runs, UniVI exposes CLI entry points
that accept JSON parameter files.

**Train from a config JSON:**

```bash
python scripts/train_univi.py \
    --config parameter_files/defaults_cite_seq_scaled_gaussian_v1.json \
    --outdir runs/citeseq_v1_run1 \
    --data-root /path/to/your/data
```

**Evaluate a saved checkpoint:**

```bash
python scripts/evaluate_univi.py \
    --config parameter_files/defaults_cite_seq_scaled_gaussian_v1.json \
    --model-checkpoint runs/citeseq_v1_run1/checkpoints/univi_checkpoint.pt \
    --outdir runs/citeseq_v1_run1/eval
```

**Reproduce all manuscript figures and supplemental tables in one call:**

```bash
bash scripts/revision_reproduce_all.sh
```

**Exported run directory structure** (typical):

```
runs/<run_name>/
├── checkpoints/
│   ├── univi_checkpoint.pt   # model + optimizer + config
│   └── best.pt               # best-val checkpoint (if early stopping enabled)
├── eval/
│   ├── metrics.json          # machine-readable metrics summary
│   ├── metrics.csv           # flat table for comparisons
│   └── plots/                # UMAPs, heatmaps, benchmark figures
├── embeddings/
│   ├── mu_z.npy              # fused mean embeddings (cells × latent_dim)
│   ├── modality_mu/          # per-modality embeddings
│   │   ├── rna.npy
│   │   └── adt.npy
│   └── obs_names.txt         # row order for safe joins
├── reconstructions/
│   ├── rna_from_rna.npy
│   ├── adt_from_rna.npy      # cross-modal imputation
│   └── ...
├── tables/
│   └── Supplemental_Table_S1.xlsx
└── logs/
    ├── train.log
    └── history.csv
```

You can also invoke the package directly as a module:

```bash
python -m univi --help
```

---

## Advanced: Transformer encoders (per-modality)

By default, UniVI uses MLP encoders. You can switch any modality to a transformer encoder
by setting `encoder_type="transformer"` and providing a `TokenizerConfig` and `TransformerConfig`.

```python
from univi.config import TransformerConfig, TokenizerConfig

univi_cfg = UniVIConfig(
    latent_dim=40,
    beta=1.0,
    gamma=1.25,
    modalities=[
        ModalityConfig(
            name="rna",
            input_dim=rna.n_vars,
            encoder_hidden=[512, 256, 128],   # kept for compatibility; ignored by transformer encoder
            decoder_hidden=[128, 256, 512],
            likelihood="gaussian",
            encoder_type="transformer",
            tokenizer=TokenizerConfig(
                mode="topk_channels",
                n_tokens=512,
                channels=("value", "rank", "dropout"),
            ),
            transformer=TransformerConfig(
                d_model=256, num_heads=8, num_layers=4,
                dim_feedforward=1024, dropout=0.1, attn_dropout=0.1,
                activation="gelu", pooling="mean",
            ),
        ),
        ModalityConfig(
            name="adt",
            input_dim=adt.n_vars,
            encoder_hidden=[128, 64],
            decoder_hidden=[64, 128],
            likelihood="gaussian",
            encoder_type="mlp",
            tokenizer=TokenizerConfig(mode="topk_scalar", n_tokens=min(32, adt.n_vars)),
        ),
    ],
)
```

---

## Advanced: Distance attention bias for ATAC (transformer only)

For ATAC tokenizers, you can optionally incorporate genomic coordinate embeddings and
distance-aware attention bias to encourage local regulatory context:

```python
# Tokenizer config with coordinate embeddings enabled
TokenizerConfig(
    mode="topk_channels",
    n_tokens=512,
    channels=("value", "rank", "dropout"),
    use_coord_embedding=True,
    n_chroms=<num_chromosomes>,
    coord_scale=1e-6,
)

# Provide peak coordinates to the trainer
feature_coords = {
    "atac": {
        "chrom_ids": chrom_ids_long,   # (F,) integer chromosome IDs
        "start":     start_bp,          # (F,) start positions in bp
        "end":       end_bp,            # (F,) end positions in bp
    }
}

attn_bias_cfg = {
    "atac": {
        "type":            "distance",
        "lengthscale_bp":  50_000.0,
        "same_chrom_only": True,
    }
}

trainer = UniVITrainer(
    model,
    train_loader,
    val_loader=val_loader,
    train_cfg=TrainingConfig(...),
    device=device,
    feature_coords=feature_coords,
    attn_bias_cfg=attn_bias_cfg,
)
trainer.fit()
```

---

## Advanced: Fused multimodal transformer posterior

Instead of analytic MoE/PoE fusion of per-modality posteriors, you can use a single
transformer that sees concatenated tokens from all modalities and outputs a single
fused posterior `q(z | all modalities)`:

```python
from univi.config import TransformerConfig

univi_cfg = UniVIConfig(
    latent_dim=40,
    beta=1.0,
    gamma=1.25,
    modalities=[...],
    fused_encoder_type="multimodal_transformer",
    fused_modalities=("rna", "adt", "atac"),
    fused_require_all_modalities=True,   # fall back to MoE if any modality is missing
    fused_transformer=TransformerConfig(
        d_model=256, num_heads=8, num_layers=4,
        dim_feedforward=1024, dropout=0.1, attn_dropout=0.1,
        activation="gelu", pooling="cls",
    ),
)
```

Encoding with the fused transformer posterior:

```python
mu, logvar, z = model.encode_fused(
    {"rna": X_rna, "adt": X_adt, "atac": X_atac},
    use_mean=True,
)
```

> When `fused_require_all_modalities=True`, the model falls back to MoE fusion
> automatically for cells missing one or more modalities.

---

## Advanced: Learnable MoE gating

By default, fusion uses pure precision weighting. You can add a learnable gate network
that produces data-driven per-cell modality trust scores:

```python
univi_cfg = UniVIConfig(
    ...,
    use_moe_gating=True,
    moe_gating_type="per_modality",   # or "shared"
    moe_gating_hidden=[128, 64],
    moe_gating_dropout=0.1,
    moe_gating_batchnorm=True,
    moe_gate_eps=1e-6,
)
```

Retrieve gates after encoding:

```python
mu, logvar, z, gates, gate_logits = model.encode_fused(
    x_dict,
    use_mean=True,
    return_gates=True,
    return_gate_logits=True,
)
# gates: (n_cells, n_modalities)
```

---

## Advanced: Supervised classification heads

UniVI supports any number of supervised heads trained jointly with the VAE objective.
Heads can be categorical, binary, or adversarial (domain confusion via gradient reversal).

```python
from univi.config import ClassHeadConfig

univi_cfg = UniVIConfig(
    ...,
    class_heads=[
        ClassHeadConfig(
            name="celltype",
            n_classes=n_celltypes,
            loss_weight=1.0,
            ignore_index=-1,
            from_mu=True,
            warmup=0,
        ),
        ClassHeadConfig(
            name="batch",
            n_classes=n_batches,
            loss_weight=0.2,
            ignore_index=-1,
            from_mu=True,
            warmup=10,
            adversarial=True,
            adv_lambda=1.0,
        ),
        ClassHeadConfig(
            name="TP53_mut",
            n_classes=2,
            loss_weight=0.5,
            ignore_index=-1,
            from_mu=True,
        ),
    ],
)

model.set_head_label_names("celltype", list(rna.obs["celltype"].astype("category").cat.categories))
```

Pass labels as a dict during training:

```python
y = {
    "celltype": torch.tensor(celltype_codes[batch_idx], device=device),
    "batch":    torch.tensor(batch_codes[batch_idx], device=device),
    "TP53_mut": torch.tensor(tp53_flags[batch_idx], device=device),
}
out = model(x_dict, epoch=epoch, y=y)
loss = out["loss"]
loss.backward()
```

Predict heads after training:

```python
model.eval()
with torch.no_grad():
    probs = model.predict_heads(x_dict, return_probs=True)
for head_name, P in probs.items():
    print(head_name, P.shape)

# Inspect head config metadata
meta = model.get_classification_meta()
print(meta)
```

---

## Advanced: Categorical variables as a full modality

Rather than a prediction head, you can treat a categorical variable as a generative modality
with its own encoder `q(z|y)` and decoder `p(y|z)`, participating fully in fusion and
posterior alignment.

```python
univi_cfg = UniVIConfig(
    ...,
    modalities=[
        ModalityConfig("rna", rna.n_vars, [512, 256, 128], [128, 256, 512], likelihood="nb"),
        ModalityConfig("adt", adt.n_vars, [128, 64], [64, 128], likelihood="nb"),
        ModalityConfig(
            name="celltype_mod",
            input_dim=n_celltypes,    # number of classes
            encoder_hidden=[128, 64],
            decoder_hidden=[64, 128],
            likelihood="categorical",
        ),
    ],
)
```

Pass integer labels (or one-hot) via `x_dict["celltype_mod"]` during training.
Unlabeled cells should use `-1` (or your configured `ignore_index`). This is distinct
from classification heads: the categorical modality is part of the generative model, not
an auxiliary output.

---

## Advanced: Label expert injection into the fused posterior

Semi-supervised alignment where label information is fused as an additional Gaussian expert:

```python
univi_cfg = UniVIConfig(
    ...,
    use_label_encoder=True,
    n_label_classes=n_celltypes,
    label_encoder_warmup=50,     # epochs before label injection starts
    label_moe_weight=1.0,
    unlabeled_logvar=10.0,       # large → label contributes little when missing
)

# At encode time:
mu, logvar, z = model.encode_fused(
    x_dict,
    epoch=current_epoch,
    y={"label": y_ids},
    inject_label_expert=True,
)
```

---

## Advanced: Reconstruction loss balancing across modalities

When modalities have very different feature dimensionalities (RNA: 2k–20k; ADT: 30–200;
ATAC-LSI: ~50–500), reconstruction losses summed over features can cause high-dimensional
modalities to dominate gradients. UniVI supports per-modality normalization:

```python
model = UniVIMultiModalVAE(
    univi_cfg,
    loss_mode="v1",
    v1_recon="avg",
    normalize_v1_terms=True,
    recon_normalize_by_dim=True,   # divide recon loss by D^power
    recon_dim_power=0.5,           # 0.5 = soft correction; 1.0 = full per-feature mean
).to(device)
```

Alternatively, set per-modality weights directly in `ModalityConfig`:

```python
ModalityConfig(
    name="rna",
    ...,
    recon_weight=1.0,
)
ModalityConfig(
    name="adt",
    ...,
    recon_weight=2.0,   # upweight ADT relative to RNA
)
```

---

## Advanced: Hyperparameter optimization

UniVI ships with modality-specific hyperparameter search scripts:

```python
from univi.hyperparam_optimization import (
    run_citeseq_hparam_search,
    run_multiome_hparam_search,
    run_teaseq_hparam_search,
    run_rna_hparam_search,
    run_atac_hparam_search,
    run_adt_hparam_search,
)
```

Or via the CLI:

```bash
python scripts/run_multiome_hparam_search.py \
    --config parameter_files/defaults_multiome_v1.json \
    --outdir runs/hparam_search_multiome \
    --n-trials 50
```

See `univi/hyperparam_optimization/` and the grid-sweep notebooks in
`notebooks/GR/` for end-to-end examples with result aggregation.

---

## Repository structure

```text
UniVI/
├── README.md                              # Project overview, installation, quickstart
├── LICENSE                                # MIT license text file
├── pyproject.toml                         # Python packaging config (pip / PyPI)
├── assets/                                # Static assets used by README/docs
│   └── figures/                           # Schematic figure(s) for repository front page
├── conda.recipe/                          # Conda build recipe (for conda-build)
│   └── meta.yaml
├── envs/                                  # Conda/mamba environment examples you can use
│   ├── UniVI_working_environment.yml
│   ├── UniVI_working_environment_v2_full.yml
│   ├── UniVI_working_environment_v2_minimal.yml
│   ├── univi_env.yml                      
│   └── univi_v0.4.7.yml                   # Most recent working environment (v0.4.7) - recommended
├── data/                                  # Small example data notes (datasets are typically external)
│   └── README.md                          # Notes on data sources / formats
├── notebooks/                             # End-to-end Jupyter Notebook analyses and examples
│   ├── GR/     # Reproduce figures from revised manuscript (in progress for Genome Research; bioRxiv manuscript v2)
│   │   ├── Fig02_CITE_paired.ipynb
│   │   ├── Fig03_CITE_biological_latent.ipynb
│   │   ├── Fig04_Multiome_paired.ipynb
│   │   ├── Fig05_Multiome_bridge_finetuning.ipynb
│   │   ├── Fig06_TEAseq_trimodal.ipynb
│   │   ├── Fig07_AML_bridge_finetuning.ipynb
│   │   ├── Fig08_benchmark_PyTorch.ipynb
│   │   ├── Fig08_benchmark_R.ipynb
│   │   ├── Fig08_benchmark_merge_plot.ipynb
│   │   ├── Fig09_scaling_ablation.ipynb
│   │   ├── Fig09_scaling_compile.ipynb
│   │   ├── Fig10_population_ablation_MoE.ipynb
│   │   ├── Fig10_population_compile.ipynb
│   │   ├── Supp_mouse_skin_SHAREseq.ipynb
│   │   ├── Supp_scNMT_mouse_gastrulation.ipynb
│   │   ├── Supp_grid_sweep.ipynb
│   │   ├── Supp_grid_sweep_compile.ipynb
│   │   └── Supplemental_Notebook_S1.ipynb
│   └── examples/         # Additional examples of UniVI workflow functionality
│       └── Additional_AML_atlas_integration.ipynb
├── parameter_files/                       # JSON configs for model + training + data selectors
│   ├── defaults_*.json                    # Default configs (per experiment)
│   └── params_*.json                      # Example “named” configs (RNA, ADT, ATAC, methylome, etc.)
├── scripts/                               # Reproducible entry points (revision-friendly)
│   ├── train_univi.py                     # Train UniVI from a parameter JSON
│   ├── evaluate_univi.py                  # Evaluate trained models (FOSCTTM, label transfer, etc.)
│   ├── benchmark_univi_citeseq.py         # CITE-seq-specific benchmarking script
│   ├── run_multiome_hparam_search.py
│   ├── run_frequency_robustness.py        # Composition/frequency mismatch robustness
│   ├── run_do_not_integrate_detection.py  # “Do-not-integrate” unmatched population demo
│   ├── run_benchmarks.py                  # Unified wrapper (includes optional Harmony baseline)
│   └── revision_reproduce_all.sh          # One-click: reproduces figures + supplemental tables
└── univi/                                 # UniVI Python package (importable as `import univi`)
    ├── __init__.py                        # Package exports and __version__
    ├── __main__.py                        # Enables: `python -m univi ...`
    ├── cli.py                             # Minimal CLI (e.g., export-s1, encode)
    ├── pipeline.py                        # Config-driven model+data loading; latent encoding helpers
    ├── diagnostics.py                     # Exports Supplemental_Table_S1.xlsx (env + hparams + dataset stats)
    ├── config.py                          # Config dataclasses (UniVIConfig, ModalityConfig, TrainingConfig)
    ├── data.py                            # Dataset wrappers + matrix selectors (layer/X_key, obsm support)
    ├── evaluation.py                      # Metrics (FOSCTTM, mixing, label transfer, feature recovery)
    ├── matching.py                        # Modality matching / alignment helpers
    ├── objectives.py                      # Losses (ELBO variants, KL/alignment annealing, etc.)
    ├── plotting.py                        # Plotting helpers + consistent style defaults
    ├── trainer.py                         # UniVITrainer: training loop, logging, checkpointing
    ├── interpretability.py                # Helper scripts for transformer token weight interpretability
    ├── figures/                           # Package-internal figure assets (placeholder)
    │   └── .gitkeep
    ├── models/                            # VAE architectures + building blocks
    │   ├── __init__.py
    │   ├── mlp.py                         # Shared MLP building blocks
    │   ├── encoders.py                    # Modality encoders (MLP + transformer + fused transformer)
    │   ├── decoders.py                    # Likelihood-specific decoders (NB, ZINB, Gaussian, Beta, Binomial/Beta-Binomial, etc.)
    │   ├── transformer.py                 # Transformer blocks + encoder (+ optional attn bias support)
    │   ├── tokenizer.py                   # Tokenization configs/helpers (top-k / patch)
    │   └── univi.py                       # Core UniVI multi-modal VAE
    ├── hyperparam_optimization/           # Hyperparameter search scripts
    │   ├── __init__.py
    │   ├── common.py
    │   ├── run_adt_hparam_search.py
    │   ├── run_atac_hparam_search.py
    │   ├── run_citeseq_hparam_search.py
    │   ├── run_multiome_hparam_search.py
    │   ├── run_rna_hparam_search.py
    │   └── run_teaseq_hparam_search.py
    └── utils/                             # General utilities
        ├── __init__.py
        ├── io.py                          # I/O helpers (AnnData, configs, checkpoints)
        ├── logging.py                     # Logging configuration / progress reporting
        ├── seed.py                        # Reproducibility helpers (seeding RNGs)
        ├── stats.py                       # Small statistical helpers / transforms
        └── torch_utils.py                 # PyTorch utilities (device, tensor helpers)
```

---

## License

MIT License — see `LICENSE`.

---

## Contact, questions, and bug reports

* Questions / comments: open a GitHub Issue with the `question` label (or use Discussions)
* Bug reports: include:

  * UniVI version: `python -c "import univi; print(univi.__version__)"`
  * minimal notebook/code snippet
  * stack trace + OS/CUDA/PyTorch versions


## Compact supplemental archive note

To reduce journal-upload size and avoid Windows path-length errors, stored Jupyter outputs and execution counts were removed from all notebooks, and notebook filenames were shortened. The source code, markdown cells, notebook order, and parameters are unchanged. See `NOTEBOOK_FILE_MAP.tsv` for the original-to-compact filename mapping.
