## Supplemental Code for "Unifying multimodal single-cell data with a mixture-of-experts *β*-variational autoencoder framework": benchmark_univi_citeseq.py

```
#!/usr/bin/env python

"""
Benchmarking script for UniVI on paired CITE-seq (RNA + ADT) data.

What it does:
  1) Load trained UniVI model + trainer.
  2) Load RNA & ADT AnnData (same cells, paired).
  3) Encode into UniVI latent space.
  4) Compute:
       - FOSCTTM (RNA vs ADT)
       - Modality mixing score
       - kNN label transfer accuracy (ADT→RNA, RNA→ADT)
       - Silhouette scores (by cell type, by modality)
  5) Run UMAP on latent and make:
       - UMAP colored by celltype
       - UMAP colored by modality
  6) Save metrics.json + plots in output_dir and also show plots.

Assumptions:
  - RNA and ADT AnnData are aligned: same obs_names, same ordering.
  - Cell type column exists in obs (e.g. "celltype.l2").
  - UniVI training code saved a checkpoint + config that this script can use.

Usage (example):
  python benchmark_univi_citeseq.py \
      --rna-h5ad path/to/rna_test.h5ad \
      --adt-h5ad path/to/adt_test.h5ad \
      --celltype-key celltype.l2 \
      --checkpoint path/to/univi_checkpoint.pt \
      --config-json path/to/univi_config.json \
      --output-dir benchmarks/hao_level2_univi \
      --loss_mode v1

"""

import os
import json
import argparse
from pathlib import Path

import numpy as np
import pandas as pd
import matplotlib.pyplot as plt
import seaborn as sns

import scanpy as sc
import anndata as ad

from sklearn.neighbors import NearestNeighbors
from sklearn.metrics import silhouette_score

# UniVI imports – adjust if your module paths differ
from univi.config import UniVIConfig, TrainingConfig
from univi.models.univi import UniVIMultiModalVAE
from univi.trainer import UniVITrainer
from univi import evaluation as univi_eval
# from univi import plotting as univi_plot  # optional

sns.set(style="whitegrid")


# ----------------------------------------------------------------------
# Helper: label transfer via kNN
# ----------------------------------------------------------------------
def knn_label_transfer(
    Z_src,
    Z_tgt,
    labels_src,
    labels_tgt,
    n_neighbors=10,
):
    """
    Simple kNN-based label transfer accuracy.

    Parameters
    ----------
    Z_src : np.ndarray, shape (n_src, d)
        Latent embedding of source cells.
    Z_tgt : np.ndarray, shape (n_tgt, d)
        Latent embedding of target cells.
    labels_src : array-like, shape (n_src,)
        True labels for source cells.
    labels_tgt : array-like, shape (n_tgt,)
        True labels for target cells (used as ground truth).
    n_neighbors : int
        Number of neighbors for kNN voting.

    Returns
    -------
    acc : float
        Overall label transfer accuracy.
    preds : np.ndarray, shape (n_tgt,)
        Predicted labels for target cells.
    """
    labels_src = np.asarray(labels_src)
    labels_tgt = np.asarray(labels_tgt)

    nn = NearestNeighbors(n_neighbors=n_neighbors, metric="euclidean")
    nn.fit(Z_src)
    dists, idx = nn.kneighbors(Z_tgt)  # idx: (n_tgt, n_neighbors)

    preds = []
    for neigh_idx in idx:
        neigh_labels = labels_src[neigh_idx]
        # majority vote
        (values, counts) = np.unique(neigh_labels, return_counts=True)
        pred = values[np.argmax(counts)]
        preds.append(pred)

    preds = np.array(preds)
    acc = (preds == labels_tgt).mean()

    return acc, preds


# ----------------------------------------------------------------------
# Helper: make UMAP embedding & plots
# ----------------------------------------------------------------------
def run_umap_and_plot(
    adata_joint,
    latent_key="X_univi",
    celltype_key="celltype.l2",
    modality_key="modality",
    output_dir=None,
):
    """
    Assumes `adata_joint.obsm[latent_key]` already contains the UniVI latent.

    Computes neighbors + UMAP on that latent, and makes:
      - UMAP colored by celltype
      - UMAP colored by modality
    """
    sc.pp.neighbors(adata_joint, use_rep=latent_key, n_neighbors=30)
    sc.tl.umap(adata_joint)

    # UMAP by cell type
    plt.figure(figsize=(6, 5))
    sc.pl.umap(
        adata_joint,
        color=celltype_key,
        size=8,
        legend_loc="on data",
        title="UniVI latent UMAP – Hao level2 cell types",
        show=False,
    )
    if output_dir is not None:
        plt.savefig(os.path.join(output_dir, "umap_celltype.png"), dpi=200, bbox_inches="tight")
    plt.show()

    # UMAP by modality
    plt.figure(figsize=(6, 5))
    sc.pl.umap(
        adata_joint,
        color=modality_key,
        size=8,
        title="UniVI latent UMAP – modality",
        show=False,
    )
    if output_dir is not None:
        plt.savefig(os.path.join(output_dir, "umap_modality.png"), dpi=200, bbox_inches="tight")
    plt.show()


# ----------------------------------------------------------------------
# Main benchmarking function
# ----------------------------------------------------------------------
def benchmark_univi_citeseq(
    rna_h5ad,
    adt_h5ad,
    celltype_key,
    checkpoint_path,
    config_json,
    output_dir,
    n_neighbors_label=10,
):
    os.makedirs(output_dir, exist_ok=True)
    output_dir = str(Path(output_dir).resolve())
    print(f"[UniVI Benchmark] Output directory: {output_dir}")

    # -----------------------------
    # 1. Load data
    # -----------------------------
    print("[UniVI Benchmark] Loading RNA and ADT AnnData...")
    rna_adata = sc.read_h5ad(rna_h5ad)
    adt_adata = sc.read_h5ad(adt_h5ad)

    assert rna_adata.n_obs == adt_adata.n_obs, "RNA and ADT n_obs mismatch!"
    assert np.array_equal(rna_adata.obs_names, adt_adata.obs_names), (
        "RNA and ADT obs_names are not aligned – fix before benchmarking."
    )

    print(f"  RNA shape: {rna_adata.shape}")
    print(f"  ADT shape: {adt_adata.shape}")
    print(f"  Celltype key: {celltype_key}")

    if celltype_key not in rna_adata.obs.columns:
        raise ValueError(f"celltype_key '{celltype_key}' not found in rna_adata.obs")

    # -----------------------------
    # 2. Load UniVI config + model + trainer
    # -----------------------------
    print("[UniVI Benchmark] Loading UniVI config and model...")
    with open(config_json, "r") as f:
        cfg_dict = json.load(f)

    # Adjust this if your UniVIConfig.from_dict / from_json API is different
    univi_cfg = UniVIConfig.from_dict(cfg_dict["univi"])
    train_cfg = TrainingConfig.from_dict(cfg_dict["training"])

    #model = UniVIMultiModalVAE(univi_cfg)
    model = UniVIMultiModalVAE(univi_cfg, loss_mode=args.loss_mode)
    trainer = UniVITrainer(
        model=model,
        train_loader=None,
        val_loader=None,
        config=train_cfg,
    )

    print(f"[UniVI Benchmark] Loading checkpoint from: {checkpoint_path}")
    trainer.load_checkpoint(checkpoint_path, map_location="cpu")
    model.eval()

    # -----------------------------
    # 3. Encode UniVI latents
    # -----------------------------
    print("[UniVI Benchmark] Encoding latents for RNA and ADT...")
    # Assumes trainer.encode_modality exists; adjust if your API differs
    z_rna = trainer.encode_modality(rna_adata, modality="rna")
    z_adt = trainer.encode_modality(adt_adata, modality="adt")

    rna_adata.obsm["X_univi"] = z_rna
    adt_adata.obsm["X_univi"] = z_adt

    # -----------------------------
    # 4. Global metrics (FOSCTTM, mixing)
    # -----------------------------
    print("[UniVI Benchmark] Computing FOSCTTM...")
    foscttm = univi_eval.compute_foscttm(z_rna, z_adt)
    print(f"  FOSCTTM (rna vs adt): {foscttm:.4f}")

    print("[UniVI Benchmark] Computing modality mixing score...")
    Z_joint = np.concatenate([z_rna, z_adt], axis=0)
    modality_labels = np.array(["rna"] * z_rna.shape[0] + ["adt"] * z_adt.shape[0])

    mixing_score = univi_eval.compute_modality_mixing(
        Z_joint,
        modality_labels,
        k=20,
    )
    print(f"  Modality mixing score (k=20): {mixing_score:.4f}")

    # -----------------------------
    # 5. Label transfer (ADT→RNA and RNA→ADT)
    # -----------------------------
    print("[UniVI Benchmark] Computing label transfer accuracy...")

    ct_rna = rna_adata.obs[celltype_key].astype(str).values
    ct_adt = adt_adata.obs[celltype_key].astype(str).values

    # ADT -> RNA
    acc_adt2rna, preds_adt2rna = knn_label_transfer(
        Z_src=z_adt,
        Z_tgt=z_rna,
        labels_src=ct_adt,
        labels_tgt=ct_rna,
        n_neighbors=n_neighbors_label,
    )
    print(f"  Label transfer (ADT→RNA, k={n_neighbors_label}): {acc_adt2rna:.3f}")

    # RNA -> ADT
    acc_rna2adt, preds_rna2adt = knn_label_transfer(
        Z_src=z_rna,
        Z_tgt=z_adt,
        labels_src=ct_rna,
        labels_tgt=ct_adt,
        n_neighbors=n_neighbors_label,
    )
    print(f"  Label transfer (RNA→ADT, k={n_neighbors_label}): {acc_rna2adt:.3f}")

    # -----------------------------
    # 6. Silhouette scores
    # -----------------------------
    print("[UniVI Benchmark] Computing silhouette scores...")

    sil_by_celltype = silhouette_score(Z_joint, np.concatenate([ct_rna, ct_adt], axis=0))
    sil_by_modality = silhouette_score(Z_joint, modality_labels)

    print(f"  Silhouette (cell type): {sil_by_celltype:.3f}")
    print(f"  Silhouette (modality): {sil_by_modality:.3f}")

    # -----------------------------
    # 7. Join adatas for UMAP & plotting
    # -----------------------------
    print("[UniVI Benchmark] Building joint AnnData for UMAP...")
    # Stack RNA and ADT as two "views" in one AnnData, using UniVI latent
    joint = ad.AnnData(
        X=Z_joint,  # just to have a matrix; we'll use obsm for UMAP
    )
    joint.obs_names = list(rna_adata.obs_names) + list(adt_adata.obs_names)
    joint.obs[celltype_key] = np.concatenate([ct_rna, ct_adt], axis=0)
    joint.obs["modality"] = modality_labels
    joint.obsm["X_univi"] = Z_joint

    run_umap_and_plot(
        adata_joint=joint,
        latent_key="X_univi",
        celltype_key=celltype_key,
        modality_key="modality",
        output_dir=output_dir,
    )

    # -----------------------------
    # 8. Optional: FOSCTTM distribution plot
    # -----------------------------
    print("[UniVI Benchmark] Plotting FOSCTTM distribution (per cell)...")
    foscttm_per_cell = univi_eval.foscttm_per_cell(z_rna, z_adt)  # if you have such a helper
    # If not, comment above and implement a per-cell version, or just skip.

    plt.figure(figsize=(6, 4))
    sns.histplot(foscttm_per_cell, bins=50, kde=True)
    plt.xlabel("FOSCTTM per cell")
    plt.ylabel("Count")
    plt.title("Distribution of FOSCTTM across cells (RNA vs ADT)")
    plt.tight_layout()
    plt.savefig(os.path.join(output_dir, "foscttm_hist.png"), dpi=200, bbox_inches="tight")
    plt.show()

    # -----------------------------
    # 9. Save metrics as JSON
    # -----------------------------
    metrics = {
        "foscttm_rna_vs_adt": float(foscttm),
        "modality_mixing_k20": float(mixing_score),
        "label_transfer_acc_adt_to_rna": float(acc_adt2rna),
        "label_transfer_acc_rna_to_adt": float(acc_rna2adt),
        "silhouette_celltype": float(sil_by_celltype),
        "silhouette_modality": float(sil_by_modality),
        "n_cells": int(rna_adata.n_obs),
        "n_genes_rna": int(rna_adata.n_vars),
        "n_features_adt": int(adt_adata.n_vars),
        "celltype_key": celltype_key,
        "n_neighbors_label_transfer": int(n_neighbors_label),
    }

    metrics_path = os.path.join(output_dir, "metrics.json")
    with open(metrics_path, "w") as f:
        json.dump(metrics, f, indent=2)

    print(f"[UniVI Benchmark] Saved metrics to: {metrics_path}")
    print("[UniVI Benchmark] Done.")


# ----------------------------------------------------------------------
# CLI
# ----------------------------------------------------------------------
def parse_args():
    p = argparse.ArgumentParser(description="Benchmark UniVI on paired CITE-seq (RNA + ADT).")
    p.add_argument("--rna-h5ad", required=True, help="Path to RNA AnnData (.h5ad).")
    p.add_argument("--adt-h5ad", required=True, help="Path to ADT AnnData (.h5ad).")
    p.add_argument("--loss_mode", type=str, default="lite", help="Training objective: v1 or lite")
    p.add_argument("--celltype-key", default="celltype.l2", help="obs key for cell type labels.")
    p.add_argument("--checkpoint", required=True, help="Path to UniVI checkpoint (.pt).")
    p.add_argument("--config-json", required=True, help="Path to UniVI config JSON.")
    p.add_argument("--output-dir", required=True, help="Directory to write metrics & plots.")
    p.add_argument(
        "--n-neighbors-label",
        type=int,
        default=10,
        help="k for kNN label transfer.",
    )
    return p.parse_args()


if __name__ == "__main__":
    args = parse_args()
    benchmark_univi_citeseq(
        rna_h5ad=args.rna_h5ad,
        adt_h5ad=args.adt_h5ad,
        celltype_key=args.celltype_key,
        checkpoint_path=args.checkpoint,
        config_json=args.config_json,
        output_dir=args.output_dir,
        n_neighbors_label=args.n_neighbors_label,
    )
```
