## Supplemental Code for "Unifying multimodal single-cell data with a mixture-of-experts *β*-variational autoencoder framework": evaluate_univi.py

```
#!/usr/bin/env python3
"""
evaluate_univi.py
-----------------
CLI evaluation of a saved UniVI checkpoint on held-out test cells.

Runs: per-modality encode, fused encode, cross-modal imputation,
alignment metrics (FOSCTTM, Recall@k, mixing, label transfer),
MoE gating, self-reconstruction error, UMAPs, embedding export.

Usage
-----
python scripts/evaluate_univi.py \
    --config    parameter_files/params_citeseq_pbmc_GR_fig2_3.json \
    --checkpoint runs/citeseq_v1_fig2_3/checkpoints/univi_checkpoint.pt \
    --splits    runs/citeseq_v1_fig2_3/splits.npz \
    --outdir    runs/citeseq_v1_fig2_3/eval \
    --data-root /path/to/data \
    [--device cuda] [--transductive] [--skip-plots]
"""

import argparse, csv, json, warnings
from pathlib import Path

import numpy as np
import scipy.sparse as sp
import scanpy as sc
import torch

from univi import UniVIMultiModalVAE
from univi.data import align_paired_obs_names
from univi.evaluation import (
    encode_adata, encode_fused_adata_pair, cross_modal_predict,
    denoise_adata, evaluate_alignment, reconstruction_metrics,
    evaluate_cross_reconstruction, encode_moe_gates_from_tensors,
)
from univi.plotting import (
    set_style, umap, umap_by_modality, plot_confusion_matrix,
    write_gates_to_obs, plot_moe_gate_summary,
    plot_reconstruction_error_summary,
)
from univi.utils.logging import get_logger

logger = get_logger(__name__)
warnings.filterwarnings("ignore", category=FutureWarning)


def parse_args():
    p = argparse.ArgumentParser(description="Evaluate a saved UniVI checkpoint.")
    p.add_argument("--config",       required=True)
    p.add_argument("--checkpoint",   required=True)
    p.add_argument("--splits",       required=True)
    p.add_argument("--outdir",       required=True)
    p.add_argument("--data-root",    default=".")
    p.add_argument("--device",       default=None)
    p.add_argument("--transductive", action="store_true",
                   help="Encode all cells rather than test set only.")
    p.add_argument("--skip-plots",   action="store_true")
    return p.parse_args()


def resolve_device(arg):
    if arg: return arg
    if torch.cuda.is_available(): return "cuda"
    if getattr(torch.backends, "mps", None) and torch.backends.mps.is_available(): return "mps"
    return "cpu"


def to_dense(X):
    return X.toarray() if sp.issparse(X) else np.asarray(X)


def preprocess_rna(adata, cfg):
    adata.layers["counts"] = adata.X.copy()
    sc.pp.normalize_total(adata, target_sum=cfg.get("normalize_total", 1e4))
    sc.pp.log1p(adata); adata.raw = adata
    sc.pp.highly_variable_genes(adata, flavor=cfg.get("hvg_flavor","seurat_v3"),
                                  n_top_genes=cfg.get("n_hvg",2000), subset=True)
    sc.pp.scale(adata, max_value=cfg.get("scale_max_value", 10))
    return adata


def preprocess_adt(adata, cfg):
    adata.layers["counts"] = adata.X.copy()
    X = to_dense(adata.X); logX = np.log1p(X)
    adata.X = logX - logX.mean(axis=1, keepdims=True)
    if cfg.get("scale", True):
        sc.pp.scale(adata, zero_center=True, max_value=cfg.get("scale_max_value", 10))
    if cfg.get("clip"):
        lo, hi = cfg["clip"]; adata.X = np.clip(to_dense(adata.X), lo, hi)
    return adata


def preprocess_atac(adata, cfg):
    from sklearn.decomposition import TruncatedSVD
    adata.layers["counts"] = adata.X.copy()
    X = adata.X.tocsr() if hasattr(adata.X, "tocsr") else adata.X
    cs = np.asarray(X.sum(axis=1)).ravel(); cs[cs==0] = 1.0
    tf = X.multiply(1.0 / cs[:,None])
    df = np.asarray((X>0).sum(axis=0)).ravel()
    idf = np.log1p(X.shape[0] / (1.0+df))
    X_tfidf = tf.multiply(idf)
    n = cfg.get("lsi_n_components", 100); drop = cfg.get("drop_lsi1", False)
    svd = TruncatedSVD(n_components=n+1, random_state=0)
    X_lsi = svd.fit_transform(X_tfidf)
    start = 1 if drop else 0
    adata.obsm[cfg.get("store_lsi_obsm_key","X_lsi")] = X_lsi[:, start:start+n]
    return adata


def main():
    args   = parse_args()
    device = resolve_device(args.device)
    set_style(font_scale=1.2, dpi=150)

    outdir   = Path(args.outdir)
    plot_dir = outdir / "plots"
    plot_dir.mkdir(parents=True, exist_ok=True)

    with open(args.config) as f:
        full_cfg = json.load(f)
    data_cfg   = full_cfg["data"]
    preproc    = data_cfg.get("preprocessing", {})
    modalities = data_cfg["modalities"]
    X_key_by_mod = data_cfg.get("X_key_by_mod", {m: "X" for m in modalities})
    eval_labels  = data_cfg.get("eval_labels", [])

    data_root = Path(args.data_root)
    adata_dict = {}
    for mod in modalities:
        adata_dict[mod] = sc.read_h5ad(data_root / data_cfg[f"{mod}_filename"])

    if "rna"  in adata_dict: adata_dict["rna"]  = preprocess_rna(adata_dict["rna"],   preproc.get("rna",  {}))
    if "adt"  in adata_dict: adata_dict["adt"]  = preprocess_adt(adata_dict["adt"],   preproc.get("adt",  {}))
    if "atac" in adata_dict: adata_dict["atac"] = preprocess_atac(adata_dict["atac"], preproc.get("atac", {}))
    align_paired_obs_names(adata_dict)

    splits   = np.load(args.splits, allow_pickle=True)
    test_idx = splits["test_idx"] if not args.transductive \
               else np.arange(list(adata_dict.values())[0].n_obs)
    adatas   = {mod: adata_dict[mod][test_idx].copy() for mod in modalities}
    logger.info(f"Evaluating {len(test_idx)} cells ({'transductive' if args.transductive else 'test set'})")

    ckpt  = torch.load(args.checkpoint, map_location=device, weights_only=False)
    model = UniVIMultiModalVAE(ckpt["model_config"]).to(device)
    model.load_state_dict(ckpt["model_state_dict"]); model.eval()
    logger.info(f"Loaded model  |  best_epoch={ckpt.get('best_epoch')}")

    metrics = {"best_epoch": ckpt.get("best_epoch"), "n_test": len(test_idx)}

    # 1. Per-modality encode
    for mod in modalities:
        xkey = X_key_by_mod.get(mod, "X")
        Z = encode_adata(model, adata=adatas[mod], modality=mod, device=device,
                         X_key=xkey.replace("obsm:","") if xkey.startswith("obsm:") else xkey,
                         layer=None, batch_size=1024, latent="moe_mean", random_state=0)
        adatas[mod].obsm["X_univi"] = Z

    # 2. Fused encode
    if len(modalities) >= 2:
        encode_fused_adata_pair(model, adata_by_mod=adatas, device=device,
                                batch_size=1024, use_mean=True,
                                return_gates=True, return_gate_logits=True,
                                write_to_adatas=True, fused_obsm_key="X_univi_fused",
                                gate_prefix="gate")

    # 3. Alignment metrics
    if len(modalities) >= 2:
        m1, m2  = modalities[0], modalities[1]
        lbl_col = next((l for l in eval_labels
                        if l in adatas[m1].obs.columns and l in adatas[m2].obs.columns), None)
        aln = evaluate_alignment(
            Z1=adatas[m1].obsm["X_univi"], Z2=adatas[m2].obsm["X_univi"],
            metric="euclidean", recall_ks=(1,5,10), k_mixing=20, k_entropy=30,
            labels_source=adatas[m1].obs[lbl_col].to_numpy() if lbl_col else None,
            labels_target=adatas[m2].obs[lbl_col].to_numpy() if lbl_col else None,
            compute_bidirectional_transfer=True, k_transfer=15, json_safe=True,
        )
        metrics["alignment"] = aln
        foscttm = aln.get('foscttm_mean', 'N/A')
        recall10 = aln.get('recall_sym_10', 'N/A')
        foscttm_str  = f"{foscttm:.4f}"  if isinstance(foscttm,  float) else str(foscttm)
        recall10_str = f"{recall10:.4f}" if isinstance(recall10, float) else str(recall10)
        logger.info(f"FOSCTTM={foscttm_str}  Recall@10={recall10_str}")

        if lbl_col and not args.skip_plots:
            plot_confusion_matrix(
                np.asarray(aln["label_transfer_cm"]),
                labels=np.asarray(aln["label_transfer_label_order"]),
                title=f"Label transfer ({m1}→{m2})", normalize="true",
                savepath=str(plot_dir/f"label_transfer_confusion_{m1}_to_{m2}.png"), show=False)
            if f"label_transfer_cm_{m2}_to_{m1}" in aln:
                plot_confusion_matrix(
                    np.asarray(aln[f"label_transfer_cm_{m2}_to_{m1}"]),
                    labels=np.asarray(aln[f"label_transfer_label_order_{m2}_to_{m1}"]),
                    title=f"Label transfer ({m2}→{m1})", normalize="true",
                    savepath=str(plot_dir/f"label_transfer_confusion_{m2}_to_{m1}.png"), show=False)

    # 4. Cross-modal imputation + error
    if len(modalities) >= 2:
        for src, tgt in [(modalities[0], modalities[1]), (modalities[1], modalities[0])]:
            rep = evaluate_cross_reconstruction(
                model, adata_src=adatas[src], adata_tgt=adatas[tgt],
                src_mod=src, tgt_mod=tgt, device=device, batch_size=512)
            metrics[f"cross_recon_{src}_to_{tgt}"] = rep["summary"]
            logger.info(f"{src}→{tgt}: {rep['summary']}")
            if not args.skip_plots:
                plot_reconstruction_error_summary(
                    rep, title=f"{src}→{tgt} imputation error",
                    savepath=str(plot_dir/f"recon_error_{src}_to_{tgt}.png"), show=False)

    # 5. Self-denoising
    for mod in modalities:
        denoise_adata(model, adata=adatas[mod], modality=mod, device=device,
                      out_layer="denoised_self", overwrite_X=False, batch_size=512)
        m = reconstruction_metrics(to_dense(adatas[mod].X), adatas[mod].layers["denoised_self"])
        metrics[f"self_recon_{mod}"] = m
        logger.info(f"Self-recon {mod}: MSE={m['mse_mean']:.4f}  Pearson={m['pearson_mean']:.4f}")

    # 6. MoE gating
    if len(modalities) >= 2:
        try:
            gate = encode_moe_gates_from_tensors(
                model, x_dict={mod: to_dense(adatas[mod].X) for mod in modalities},
                device=device, batch_size=1024, modality_order=modalities,
                kind="router_x_precision", return_logits=True)
            W    = gate["weights"]; mods = gate["modality_order"]
            metrics["moe_gates"] = {k: gate.get(k) for k in
                                    ("kind","requested_kind","modality_order","per_modality_mean")}
            write_gates_to_obs(adatas[modalities[0]], gates=W, modality_names=mods,
                               gate_prefix="moe_gate", gate_logits=gate.get("logits"))
            if not args.skip_plots:
                lbl = next((l for l in eval_labels if l in adatas[modalities[0]].obs.columns), None)
                if lbl:
                    plot_moe_gate_summary(adatas[modalities[0]], gate_prefix="moe_gate",
                                         groupby=lbl, agg="mean",
                                         savepath=str(plot_dir/"moe_gates_by_celltype.png"),
                                         show=False)
        except Exception as e:
            logger.warning(f"MoE gating skipped: {e}")

    # 7. UMAPs
    if not args.skip_plots:
        ref = modalities[0]
        colors = [c for c in eval_labels if c in adatas[ref].obs.columns]
        umap(adatas[ref], obsm_key="X_univi", color=colors or [adatas[ref].obs.columns[0]],
             legend="outside", legend_subset_topk=25,
             savepath=str(plot_dir/f"umap_{ref}_univi.png"), show=False)
        if "X_univi_fused" in adatas[ref].obsm:
            umap_by_modality(adatas, obsm_key="X_univi_fused",
                             color=["univi_modality"] + colors,
                             legend="outside",
                             savepath=str(plot_dir/"umap_fused_by_modality.png"), show=False)

    # 8. Export embeddings
    embed_dir = outdir / "embeddings"; embed_dir.mkdir(exist_ok=True)
    ref = modalities[0]
    if "X_univi_fused" in adatas[ref].obsm:
        np.save(embed_dir/"mu_z_fused.npy", adatas[ref].obsm["X_univi_fused"])
    mod_mu = embed_dir/"modality_mu"; mod_mu.mkdir(exist_ok=True)
    for mod in modalities:
        np.save(mod_mu/f"{mod}.npy", adatas[mod].obsm["X_univi"])
    np.savetxt(embed_dir/"obs_names.txt", adatas[ref].obs_names.to_numpy(), fmt="%s")

    # 9. Metrics export
    with open(outdir/"metrics.json","w") as f:
        json.dump(metrics, f, indent=2, default=str)
    flat = []
    for sec, vals in metrics.items():
        if isinstance(vals, dict):
            for k, v in vals.items():
                if isinstance(v, (int,float,str)): flat.append({"section":sec,"metric":k,"value":v})
        elif isinstance(vals,(int,float)):
            flat.append({"section":"","metric":sec,"value":vals})
    with open(outdir/"metrics.csv","w",newline="") as f:
        w = csv.DictWriter(f, fieldnames=["section","metric","value"])
        w.writeheader(); w.writerows(flat)
    logger.info(f"Metrics -> {outdir/'metrics.json'}")
    logger.info("Evaluation done.")


if __name__ == "__main__":
    main()
```
