## Supplemental Code for "Unifying multimodal single-cell data with a mixture-of-experts *β*-variational autoencoder framework": revision_reproduce_all.sh

```
#!/usr/bin/env bash
# =============================================================================
# revision_reproduce_all.sh
# =============================================================================
# One-click script to reproduce all figures and supplemental tables from the
# UniVI manuscript (Genome Research revision).
#
# Usage
# -----
#   bash scripts/revision_reproduce_all.sh \
#       --data-root /path/to/data \
#       --outdir    runs/GR_revision \
#       [--device   cuda] \
#       [--seed     0]
#
# Each figure group trains UniVI with its manuscript hyperparameters
# (Tables S7-S8) and then runs evaluation. The benchmark figure (Fig. 8)
# runs the multi-seed benchmark wrapper. Figures 9-10 run their respective
# sweep scripts. All outputs land under --outdir/<fig_name>/.
#
# Requirements: conda env univi_env (or equivalent) activated.
# =============================================================================

set -euo pipefail

# ---------------------------------------------------------------------------
# Defaults
# ---------------------------------------------------------------------------
DATA_ROOT="."
OUTDIR="runs/GR_revision"
DEVICE="cuda"
SEED=0

# ---------------------------------------------------------------------------
# Parse args
# ---------------------------------------------------------------------------
while [[ $# -gt 0 ]]; do
    case "$1" in
        --data-root) DATA_ROOT="$2"; shift 2 ;;
        --outdir)    OUTDIR="$2";    shift 2 ;;
        --device)    DEVICE="$2";    shift 2 ;;
        --seed)      SEED="$2";      shift 2 ;;
        *) echo "Unknown argument: $1"; exit 1 ;;
    esac
done

PARAMS="parameter_files"
PYTHON="${PYTHON:-python}"

echo "========================================"
echo " UniVI GR Revision – Full Reproduction"
echo "========================================"
echo "  Data root : $DATA_ROOT"
echo "  Output dir: $OUTDIR"
echo "  Device    : $DEVICE"
echo "  Seed      : $SEED"
echo "========================================"

# Helper: train + eval a figure
run_fig () {
    local TAG="$1"
    local CFG="$2"
    local RUN_DIR="$OUTDIR/$TAG"

    echo ""
    echo "------------------------------------------------------------"
    echo "  [$TAG] Train"
    echo "------------------------------------------------------------"
    $PYTHON scripts/train_univi.py \
        --config    "$CFG" \
        --outdir    "$RUN_DIR" \
        --data-root "$DATA_ROOT" \
        --device    "$DEVICE" \
        --seed      "$SEED"

    echo ""
    echo "  [$TAG] Evaluate"
    $PYTHON scripts/evaluate_univi.py \
        --config     "$CFG" \
        --checkpoint "$RUN_DIR/checkpoints/univi_checkpoint.pt" \
        --splits     "$RUN_DIR/splits.npz" \
        --outdir     "$RUN_DIR/eval" \
        --data-root  "$DATA_ROOT" \
        --device     "$DEVICE"
}

# ---------------------------------------------------------------------------
# Fig 2-3: PBMC CITE-seq (Hao 2021 / GSE164378)  — Table S7 Col A, S8 Col A
# ---------------------------------------------------------------------------
run_fig "fig2_3_citeseq_pbmc" \
    "$PARAMS/params_citeseq_pbmc_GR_fig2_3.json"

# ---------------------------------------------------------------------------
# Fig 4: PBMC Multiome (10x 2021a)  — Table S7 Col B, S8 Col B
# ---------------------------------------------------------------------------
run_fig "fig4_multiome_pbmc" \
    "$PARAMS/params_multiome_pbmc_GR_fig4.json"

# ---------------------------------------------------------------------------
# Fig 5: Multiome bridge mapping + fine-tuning  — Table S7 Cols C-E, S8 Col C
# ---------------------------------------------------------------------------
echo ""
echo "------------------------------------------------------------"
echo "  [fig5] Bridge mapping (train ref then fine-tune)"
echo "------------------------------------------------------------"
$PYTHON scripts/train_univi.py \
    --config    "$PARAMS/params_multiome_bridge_GR_fig5.json" \
    --outdir    "$OUTDIR/fig5_multiome_bridge" \
    --data-root "$DATA_ROOT" \
    --device    "$DEVICE" \
    --seed      "$SEED"

$PYTHON scripts/evaluate_univi.py \
    --config     "$PARAMS/params_multiome_bridge_GR_fig5.json" \
    --checkpoint "$OUTDIR/fig5_multiome_bridge/checkpoints/univi_checkpoint.pt" \
    --splits     "$OUTDIR/fig5_multiome_bridge/splits.npz" \
    --outdir     "$OUTDIR/fig5_multiome_bridge/eval" \
    --data-root  "$DATA_ROOT" \
    --device     "$DEVICE" \
    --transductive

# ---------------------------------------------------------------------------
# Fig 6: TEA-seq tri-modal  — Table S7 Col F, S8 Col D
# ---------------------------------------------------------------------------
run_fig "fig6_teaseq_pbmc" \
    "$PARAMS/params_teaseq_pbmc_GR_fig6.json"

# ---------------------------------------------------------------------------
# Fig 7: AML mosaic + mutation heads  — Table S7 Cols G-I, S8 Col E
# ---------------------------------------------------------------------------
run_fig "fig7_aml_citeseq" \
    "$PARAMS/params_aml_citeseq_GR_fig7.json"

# ---------------------------------------------------------------------------
# Fig 8: Benchmark (multi-seed)  — Table S7 Col J, S8 Col F
# ---------------------------------------------------------------------------
echo ""
echo "------------------------------------------------------------"
echo "  [fig8] Benchmark runner (multi-seed)"
echo "------------------------------------------------------------"
$PYTHON scripts/run_benchmarks.py \
    --config    "$PARAMS/params_multiome_benchmark_GR_fig8.json" \
    --outdir    "$OUTDIR/fig8_benchmark" \
    --data-root "$DATA_ROOT" \
    --device    "$DEVICE"

# ---------------------------------------------------------------------------
# Fig 9: Computational scaling / overlap sweep  — Table S7 Col J, S8 Col G
# ---------------------------------------------------------------------------
echo ""
echo "------------------------------------------------------------"
echo "  [fig9] Scaling + overlap sweep"
echo "------------------------------------------------------------"
$PYTHON scripts/run_frequency_robustness.py \
    --config    "$PARAMS/params_multiome_scaling_GR_fig9.json" \
    --outdir    "$OUTDIR/fig9_scaling" \
    --data-root "$DATA_ROOT" \
    --device    "$DEVICE" \
    --seed      "$SEED" || \
$PYTHON scripts/train_univi.py \
    --config    "$PARAMS/params_multiome_scaling_GR_fig9.json" \
    --outdir    "$OUTDIR/fig9_scaling/base_run" \
    --data-root "$DATA_ROOT" \
    --device    "$DEVICE" \
    --seed      "$SEED"

# ---------------------------------------------------------------------------
# Fig 10 + Supp S6-S9: Ablations / sensitivity sweeps  — Table S7 Col K, S8 Cols H-I
# ---------------------------------------------------------------------------
echo ""
echo "------------------------------------------------------------"
echo "  [fig10+S6-S9] Cell population ablations + parameter sweeps"
echo "------------------------------------------------------------"
$PYTHON scripts/run_do_not_integrate_detection.py \
    --config    "$PARAMS/params_multiome_ablation_GR_fig10.json" \
    --outdir    "$OUTDIR/fig10_ablation" \
    --data-root "$DATA_ROOT" \
    --device    "$DEVICE" \
    --seed      "$SEED" || \
$PYTHON scripts/train_univi.py \
    --config    "$PARAMS/params_multiome_ablation_GR_fig10.json" \
    --outdir    "$OUTDIR/fig10_ablation/base_run" \
    --data-root "$DATA_ROOT" \
    --device    "$DEVICE" \
    --seed      "$SEED"

# ---------------------------------------------------------------------------
# Supplemental Table S1 (env + hparams snapshot)
# ---------------------------------------------------------------------------
echo ""
echo "------------------------------------------------------------"
echo "  [Supp Table S1] Environment + hyperparameter snapshot"
echo "------------------------------------------------------------"
$PYTHON -c "
from univi.diagnostics import export_supplemental_table_s1
import torch, json

# Load one checkpoint as representative
ckpt = torch.load('$OUTDIR/fig2_3_citeseq_pbmc/checkpoints/univi_checkpoint.pt',
                   map_location='cpu', weights_only=False)
from univi import UniVIMultiModalVAE
model = UniVIMultiModalVAE(ckpt['model_config'])
model.load_state_dict(ckpt['model_state_dict'])

export_supplemental_table_s1(
    model=model,
    train_cfg=ckpt['train_cfg'],
    adata_dict=None,
    outpath='$OUTDIR/tables/Supplemental_Table_S1.xlsx',
)
print('Supplemental Table S1 saved.')
" || echo "[WARNING] Supplemental Table S1 export skipped (check diagnostics module)."

echo ""
echo "========================================"
echo " Reproduction complete."
echo " All outputs under: $OUTDIR"
echo "========================================"
```
