## Supplemental Code for "Unifying multimodal single-cell data with a mixture-of-experts *β*-variational autoencoder framework": run_benchmarks.py

```
#!/usr/bin/env python3
"""
run_benchmarks.py
-----------------
Wrapper that trains UniVI under the benchmark config and exports results
in a format suitable for comparison against other methods (Table S6).

Runs multi-seed training, evaluates on the held-out test set, and writes
a consolidated results.csv + results plots.

Usage
-----
python scripts/run_benchmarks.py \
    --config  parameter_files/params_multiome_benchmark_GR_fig8.json \
    --outdir  runs/benchmark_fig8 \
    --data-root /path/to/data \
    [--seeds 0 1 2] [--device cuda]
"""

import argparse
import json
import subprocess
import sys
import csv
from pathlib import Path

import numpy as np

from univi.utils.logging import get_logger

logger = get_logger(__name__)


def parse_args():
    p = argparse.ArgumentParser(description="UniVI benchmark runner (multi-seed).")
    p.add_argument("--config",    required=True)
    p.add_argument("--outdir",    required=True)
    p.add_argument("--data-root", default=".")
    p.add_argument("--seeds",     nargs="+", type=int, default=None,
                   help="Seeds to run (overrides config). Default: uses config data.multi_seed seeds or [0].")
    p.add_argument("--device",    default=None)
    return p.parse_args()


def run_seed(config_path, outdir_seed, data_root, device, seed):
    cmd = [
        sys.executable, "scripts/train_univi.py",
        "--config",    str(config_path),
        "--outdir",    str(outdir_seed),
        "--data-root", str(data_root),
        "--seed",      str(seed),
    ]
    if device:
        cmd += ["--device", device]
    logger.info(f"Running seed {seed}: {' '.join(cmd)}")
    result = subprocess.run(cmd, check=True)
    return result.returncode


def eval_seed(config_path, outdir_seed, data_root, device):
    cmd = [
        sys.executable, "scripts/evaluate_univi.py",
        "--config",     str(config_path),
        "--checkpoint", str(outdir_seed / "checkpoints" / "univi_checkpoint.pt"),
        "--splits",     str(outdir_seed / "splits.npz"),
        "--outdir",     str(outdir_seed / "eval"),
        "--data-root",  str(data_root),
        "--skip-plots",
    ]
    if device:
        cmd += ["--device", device]
    logger.info(f"Evaluating seed: {outdir_seed}")
    subprocess.run(cmd, check=True)


def collect_results(outdir, seeds):
    """Aggregate metrics.csv across seeds into a summary table."""
    rows = []
    for seed in seeds:
        metrics_csv = outdir / f"seed_{seed}" / "eval" / "metrics.csv"
        if not metrics_csv.exists():
            logger.warning(f"Missing: {metrics_csv}")
            continue
        with open(metrics_csv) as f:
            reader = csv.DictReader(f)
            for row in reader:
                row["seed"] = seed
                rows.append(row)
    return rows


def main():
    args = parse_args()

    with open(args.config) as f:
        full_cfg = json.load(f)
    data_cfg = full_cfg.get("data", {})

    # Resolve seeds
    if args.seeds is not None:
        seeds = args.seeds
    elif data_cfg.get("multi_seed") and data_cfg.get("seeds"):
        seeds = data_cfg["seeds"]
    else:
        seeds = [0]

    outdir    = Path(args.outdir)
    data_root = Path(args.data_root)
    outdir.mkdir(parents=True, exist_ok=True)

    # Train + eval each seed
    for seed in seeds:
        seed_dir = outdir / f"seed_{seed}"
        run_seed(args.config, seed_dir, data_root, args.device, seed)
        eval_seed(args.config, seed_dir, data_root, args.device)

    # Aggregate
    all_rows = collect_results(outdir, seeds)
    if all_rows:
        agg_path = outdir / "results.csv"
        with open(agg_path, "w", newline="") as f:
            writer = csv.DictWriter(f, fieldnames=list(all_rows[0].keys()))
            writer.writeheader()
            writer.writerows(all_rows)
        logger.info(f"Aggregated results -> {agg_path}")

        # Summary statistics across seeds
        import collections
        by_metric = collections.defaultdict(list)
        for row in all_rows:
            try:
                by_metric[f"{row['section']}/{row['metric']}"].append(float(row["value"]))
            except (ValueError, KeyError):
                pass
        summary_path = outdir / "results_summary.csv"
        with open(summary_path, "w", newline="") as f:
            writer = csv.writer(f)
            writer.writerow(["metric", "mean", "std", "n_seeds"])
            for metric_key, vals in sorted(by_metric.items()):
                writer.writerow([metric_key,
                                  f"{np.mean(vals):.4f}",
                                  f"{np.std(vals):.4f}",
                                  len(vals)])
        logger.info(f"Summary stats -> {summary_path}")

    logger.info("Benchmark run complete.")


if __name__ == "__main__":
    main()
```
