## Supplemental Code for "Unifying multimodal single-cell data with a mixture-of-experts *β*-variational autoencoder framework": run_do_not_integrate_detection.py

```
#!/usr/bin/env python
"""Do-not-integrate population detection (Reviewer 1).

Simulates a population only present in one modality (e.g., an RNA-only shift)
and demonstrates that UniVI can flag it as 'non-integrating' based on latent distances.

Strategy:
- Start from paired CITE-seq (RNA/ADT) dataset.
- Choose (or user-specify) a held-out label L.
- Remove all cells of label L from modality-2 (ADT) *only* (conceptually unpaired).
- Compute nearest-neighbor distances from RNA(L) cells into ADT (without L).
- Compare to control distances from RNA(non-L) into ADT.
- Flag as non-integrating if distances exceed a high percentile threshold.

Outputs:
  runs/robustness/do_not_integrate_summary.csv
  runs/robustness/do_not_integrate_plot.(png|pdf)
"""

from __future__ import annotations

import argparse
import os
from typing import Dict, Any, Optional, Tuple, List

import numpy as np
import pandas as pd
import matplotlib.pyplot as plt

from sklearn.neighbors import NearestNeighbors

from univi.pipeline import load_model_and_data, encode_latents_paired


def _save_both(fig, outbase: str):
    os.makedirs(os.path.dirname(outbase) or ".", exist_ok=True)
    fig.savefig(outbase + ".png", bbox_inches="tight")
    fig.savefig(outbase + ".pdf", bbox_inches="tight")


def main():
    ap = argparse.ArgumentParser()
    ap.add_argument("--config", required=True)
    ap.add_argument("--checkpoint", required=True)
    ap.add_argument("--outdir", required=True)
    ap.add_argument("--data-root", default=None)
    ap.add_argument("--device", default="cpu")
    ap.add_argument("--batch-size", type=int, default=512)

    ap.add_argument("--m1", default="rna")
    ap.add_argument("--m2", default="adt")
    ap.add_argument("--label-key", default="cell_type")
    ap.add_argument("--heldout-label", default=None, help="Label to treat as RNA-only (if omitted: choose rarest).")
    ap.add_argument("--threshold-quantile", type=float, default=0.95)
    ap.add_argument("--control-subsample", type=int, default=2000)
    args = ap.parse_args()

    os.makedirs(args.outdir, exist_ok=True)

    cfg, adata_dict, model, layer_by, xkey_by = load_model_and_data(
        args.config,
        checkpoint_path=args.checkpoint,
        data_root=args.data_root,
        device=args.device,
        align_obs=True,
    )

    m1, m2 = args.m1, args.m2
    ad1 = adata_dict[m1]
    ad2 = adata_dict[m2]

    if args.label_key not in ad1.obs:
        raise KeyError(f"label_key={args.label_key!r} missing from {m1}.obs")
    y = np.asarray(ad1.obs[args.label_key].values).astype(str)

    # choose held-out label
    if args.heldout_label is None:
        uniq, counts = np.unique(y, return_counts=True)
        held = str(uniq[np.argmin(counts)])
    else:
        held = str(args.heldout_label)

    held_mask = (y == held)
    if held_mask.sum() < 5:
        raise ValueError(f"Held-out label {held!r} has too few cells ({held_mask.sum()}). Choose another.")
    print(f"[Do-not-integrate] heldout_label = {held} (n={held_mask.sum()})")

    # encode paired latents once
    Z = encode_latents_paired(
        model,
        {m1: ad1, m2: ad2},
        layer_by={m1: layer_by.get(m1), m2: layer_by.get(m2)},
        xkey_by={m1: xkey_by.get(m1, "X"), m2: xkey_by.get(m2, "X")},
        batch_size=args.batch_size,
        device=args.device,
        fused=False,
    )
    Z1 = Z[m1]
    Z2 = Z[m2]

    # Remove held-out label from modality-2 (simulate missing population)
    Z2_noheld = Z2[~held_mask]

    # Nearest neighbor distances from held-out RNA cells to ADT(noheld)
    nn = NearestNeighbors(n_neighbors=1, metric="euclidean")
    nn.fit(Z2_noheld)

    dist_held, _ = nn.kneighbors(Z1[held_mask], return_distance=True)
    dist_held = dist_held.reshape(-1)

    # control distances: sample from non-held cells
    rng = np.random.default_rng(0)
    non_idx = np.where(~held_mask)[0]
    if non_idx.size > args.control_subsample:
        non_idx = rng.choice(non_idx, size=args.control_subsample, replace=False)
    dist_ctrl, _ = nn.kneighbors(Z1[non_idx], return_distance=True)
    dist_ctrl = dist_ctrl.reshape(-1)

    thr = float(np.quantile(dist_ctrl, args.threshold_quantile))
    frac_flagged = float(np.mean(dist_held > thr))

    # centroid distance matrix (RNA labels vs ADT labels excluding held)
    labels = np.unique(y)
    labels_nonheld = [l for l in labels if l != held]
    cent_rna = {l: Z1[y == l].mean(axis=0) for l in labels}
    cent_adt = {l: Z2[y == l].mean(axis=0) for l in labels_nonheld}  # ADT centroids only if available post-drop
    mat = np.zeros((len(labels), len(labels_nonheld)), dtype=float)
    for i,l1 in enumerate(labels):
        for j,l2 in enumerate(labels_nonheld):
            mat[i,j] = float(np.linalg.norm(cent_rna[l1] - cent_adt[l2]))

    summary = pd.DataFrame([{
        "heldout_label": held,
        "n_heldout": int(held_mask.sum()),
        "threshold_quantile": float(args.threshold_quantile),
        "threshold_value": thr,
        "frac_flagged_nonintegrating": frac_flagged,
        "median_nn_dist_heldout": float(np.median(dist_held)),
        "median_nn_dist_control": float(np.median(dist_ctrl)),
    }])
    out_csv = os.path.join(args.outdir, "do_not_integrate_summary.csv")
    summary.to_csv(out_csv, index=False)

    # Plot
    fig, axes = plt.subplots(1, 2, figsize=(10.5, 4.2))
    ax = axes[0]
    ax.hist(dist_ctrl, bins=40, alpha=0.7, label="control (RNA non-held → ADT)")
    ax.hist(dist_held, bins=40, alpha=0.7, label=f"held-out (RNA {held} → ADT)")
    ax.axvline(thr, linestyle="--", linewidth=2, label=f"{args.threshold_quantile:.2f} quantile threshold")
    ax.set_title("Nearest-neighbor distance to modality-2")
    ax.set_xlabel("Euclidean distance in latent space")
    ax.set_ylabel("Count")
    ax.legend(frameon=False)

    ax2 = axes[1]
    im = ax2.imshow(mat, aspect="auto")
    ax2.set_title("Centroid distances (RNA types vs ADT types)")
    ax2.set_xlabel("ADT label (held-out removed)")
    ax2.set_ylabel("RNA label")
    ax2.set_xticks(np.arange(len(labels_nonheld)))
    ax2.set_xticklabels(labels_nonheld, rotation=90)
    ax2.set_yticks(np.arange(len(labels)))
    ax2.set_yticklabels(labels)
    fig.colorbar(im, ax=ax2, fraction=0.046, pad=0.04)

    fig.suptitle(f"Do-not-integrate detection: held-out label={held} | flagged={frac_flagged:.2f}", y=1.02)
    fig.tight_layout()

    outbase = os.path.join(args.outdir, "do_not_integrate_plot")
    _save_both(fig, outbase)
    plt.close(fig)

    print(f"[OK] Wrote: {out_csv}")
    print(f"[OK] Wrote: {outbase}.png/.pdf")


if __name__ == "__main__":
    main()
```
