## Supplemental Code for "Unifying multimodal single-cell data with a mixture-of-experts *β*-variational autoencoder framework": run_frequency_robustness.py

```
#!/usr/bin/env python
"""Frequency perturbation robustness (Reviewer 1).

Goal:
Quantify how UniVI alignment degrades as (i) shared cell-type *composition* overlap
and (ii) per-type *frequency* agreement degrade between modalities.

This script assumes paired CITE-seq-style data (RNA/ADT) but perturbs the observed
composition independently per modality, then evaluates on the *intersection* of cells
that remain shared after perturbation.

Outputs:
  runs/robustness/frequency_perturbation_results.csv
  runs/robustness/frequency_perturbation_plot.(png|pdf)
"""

from __future__ import annotations

import argparse
import os
import math
from typing import Dict, Any, List, Tuple, Optional

import numpy as np
import pandas as pd
import matplotlib.pyplot as plt

from univi.pipeline import load_model_and_data, encode_latents_paired
from univi.evaluation import compute_foscttm, label_transfer_knn


def _save_both(fig, outbase: str):
    os.makedirs(os.path.dirname(outbase) or ".", exist_ok=True)
    fig.savefig(outbase + ".png", bbox_inches="tight")
    fig.savefig(outbase + ".pdf", bbox_inches="tight")


def _sample_subset(
    obs_names: np.ndarray,
    labels: np.ndarray,
    *,
    keep_frac: float,
    drop_type_frac: float,
    rng: np.random.Generator,
) -> Tuple[np.ndarray, np.ndarray, List[str]]:
    """Return subset obs_names and labels, after dropping random types and sampling cells per type."""
    labels = np.asarray(labels)
    obs_names = np.asarray(obs_names)

    types = np.unique(labels)
    n_types = len(types)
    n_drop = int(round(drop_type_frac * n_types))
    drop_types = set(rng.choice(types, size=n_drop, replace=False).tolist()) if n_drop > 0 else set()

    keep_mask_type = np.array([lab not in drop_types for lab in labels], dtype=bool)
    obs_names2 = obs_names[keep_mask_type]
    labels2 = labels[keep_mask_type]

    # independent per-type sampling (binomial per type)
    keep_idx = []
    kept_types = []
    for t in np.unique(labels2):
        idx = np.where(labels2 == t)[0]
        if idx.size == 0:
            continue
        # per-type retention with mild noise around keep_frac
        # (helps simulate composition mismatch beyond global fraction)
        p = float(np.clip(rng.normal(loc=keep_frac, scale=max(0.02, 0.15 * keep_frac)), 0.0, 1.0))
        k = rng.binomial(idx.size, p)
        if k <= 0:
            continue
        sel = rng.choice(idx, size=k, replace=False)
        keep_idx.append(sel)
        kept_types.append(str(t))
    if len(keep_idx) == 0:
        return obs_names2[:0], labels2[:0], kept_types
    keep_idx = np.concatenate(keep_idx)
    keep_idx.sort()
    return obs_names2[keep_idx], labels2[keep_idx], kept_types


def main():
    ap = argparse.ArgumentParser()
    ap.add_argument("--config", required=True)
    ap.add_argument("--checkpoint", required=True)
    ap.add_argument("--outdir", required=True)
    ap.add_argument("--data-root", default=None)
    ap.add_argument("--device", default="cpu")
    ap.add_argument("--batch-size", type=int, default=512)

    ap.add_argument("--m1", default="rna")
    ap.add_argument("--m2", default="adt")
    ap.add_argument("--label-key", default="cell_type")

    ap.add_argument("--keep-fracs", nargs="+", type=float, default=[1.0, 0.8, 0.6, 0.4, 0.25])
    ap.add_argument("--drop-type-fracs", nargs="+", type=float, default=[0.0, 0.1, 0.25, 0.5])
    ap.add_argument("--repeats", type=int, default=5)
    ap.add_argument("--seed", type=int, default=0)
    ap.add_argument("--metric", default="euclidean")
    args = ap.parse_args()

    os.makedirs(args.outdir, exist_ok=True)

    cfg, adata_dict, model, layer_by, xkey_by = load_model_and_data(
        args.config,
        checkpoint_path=args.checkpoint,
        data_root=args.data_root,
        device=args.device,
        align_obs=True,
    )

    m1, m2 = args.m1, args.m2
    if m1 not in adata_dict or m2 not in adata_dict:
        raise KeyError(f"Modalities not found in config data.modalities: {list(adata_dict.keys())}")

    ad1 = adata_dict[m1]
    ad2 = adata_dict[m2]

    if args.label_key not in ad1.obs:
        raise KeyError(f"label_key={args.label_key!r} missing from {m1}.obs")
    y1_full = np.asarray(ad1.obs[args.label_key].values)

    # best-effort for modality 2 labels
    y2_full = np.asarray(ad2.obs[args.label_key].values) if args.label_key in ad2.obs else y1_full

    obs_names = np.asarray(ad1.obs_names.values)

    # encode once (paired order)
    Z = encode_latents_paired(
        model,
        {m1: ad1, m2: ad2},
        layer_by={m1: layer_by.get(m1), m2: layer_by.get(m2)},
        xkey_by={m1: xkey_by.get(m1, "X"), m2: xkey_by.get(m2, "X")},
        batch_size=args.batch_size,
        device=args.device,
        fused=False,
    )
    Z1_full = Z[m1]
    Z2_full = Z[m2]

    # map obs_name -> index
    name_to_i = {n: i for i, n in enumerate(obs_names.tolist())}

    rows: List[Dict[str, Any]] = []
    rng0 = np.random.default_rng(args.seed)

    for rep in range(args.repeats):
        rng = np.random.default_rng(rng0.integers(1, 2**32 - 1))
        for keep_frac in args.keep_fracs:
            for drop_frac in args.drop_type_fracs:
                # independent perturbation per modality
                n1, l1, kept1 = _sample_subset(obs_names, y1_full, keep_frac=keep_frac, drop_type_frac=drop_frac, rng=rng)
                n2, l2, kept2 = _sample_subset(obs_names, y2_full, keep_frac=keep_frac, drop_type_frac=drop_frac, rng=rng)

                set1 = set(n1.tolist())
                set2 = set(n2.tolist())
                overlap = sorted(set1.intersection(set2))

                types1 = set(np.unique(l1).astype(str).tolist())
                types2 = set(np.unique(l2).astype(str).tolist())
                jacc = (len(types1 & types2) / len(types1 | types2)) if (types1 | types2) else 0.0

                overlap_cells_frac = len(overlap) / max(len(obs_names), 1)

                entry: Dict[str, Any] = {
                    "repeat": rep,
                    "seed": int(args.seed),
                    "keep_frac": float(keep_frac),
                    "drop_type_frac": float(drop_frac),
                    "n_overlap_cells": int(len(overlap)),
                    "overlap_cells_frac": float(overlap_cells_frac),
                    "overlap_types_jaccard": float(jacc),
                }

                if len(overlap) < 10:
                    entry.update({"foscttm": np.nan, "lt_acc_%s_to_%s" % (m1,m2): np.nan, "lt_acc_%s_to_%s" % (m2,m1): np.nan})
                    rows.append(entry)
                    continue

                idx = np.array([name_to_i[n] for n in overlap], dtype=int)

                Z1 = Z1_full[idx]
                Z2 = Z2_full[idx]
                y1 = y1_full[idx]
                y2 = y2_full[idx]

                fos = compute_foscttm(Z1, Z2, metric=args.metric)
                _, acc12, _ = label_transfer_knn(Z1, y1, Z2, y2, k=15, metric=args.metric)
                _, acc21, _ = label_transfer_knn(Z2, y2, Z1, y1, k=15, metric=args.metric)

                entry["foscttm"] = float(fos)
                entry["lt_acc_%s_to_%s" % (m1, m2)] = float(acc12) if acc12 is not None else np.nan
                entry["lt_acc_%s_to_%s" % (m2, m1)] = float(acc21) if acc21 is not None else np.nan
                rows.append(entry)

    df = pd.DataFrame(rows)
    out_csv = os.path.join(args.outdir, "frequency_perturbation_results.csv")
    df.to_csv(out_csv, index=False)

    # Plot: overlap vs metrics (means)
    # bin by overlap_cells_frac rounded to 2dp for a clean curve
    df2 = df.dropna(subset=["foscttm"]).copy()
    if len(df2) > 0:
        df2["overlap_bin"] = df2["overlap_cells_frac"].round(2)
        g = df2.groupby("overlap_bin", as_index=False).agg(
            foscttm_mean=("foscttm", "mean"),
            foscttm_std=("foscttm", "std"),
            acc_mean=("lt_acc_%s_to_%s" % (m1, m2), "mean"),
            acc_std=("lt_acc_%s_to_%s" % (m1, m2), "std"),
            n=("foscttm", "size"),
        )
        g = g.sort_values("overlap_bin")
    else:
        g = pd.DataFrame({"overlap_bin": [], "foscttm_mean": [], "acc_mean": []})

    fig, ax1 = plt.subplots(figsize=(7, 4.5))
    ax1.set_title("UniVI robustness to composition mismatch (frequency perturbation)")
    ax1.set_xlabel("Shared cell overlap fraction (after independent perturbation)")
    ax1.set_ylabel("FOSCTTM (lower=better)")

    if len(g) > 0:
        ax1.plot(g["overlap_bin"], g["foscttm_mean"], marker="o", linewidth=2)
        if np.isfinite(g["foscttm_std"]).any():
            ax1.fill_between(
                g["overlap_bin"],
                g["foscttm_mean"] - g["foscttm_std"].fillna(0),
                g["foscttm_mean"] + g["foscttm_std"].fillna(0),
                alpha=0.2,
            )

    ax2 = ax1.twinx()
    ax2.set_ylabel(f"Label transfer accuracy ({m1}→{m2})")
    if len(g) > 0:
        ax2.plot(g["overlap_bin"], g["acc_mean"], marker="s", linewidth=2)

    ax1.grid(True, alpha=0.3)
    fig.tight_layout()

    outbase = os.path.join(args.outdir, "frequency_perturbation_plot")
    _save_both(fig, outbase)
    plt.close(fig)

    print(f"[OK] Wrote: {out_csv}")
    print(f"[OK] Wrote: {outbase}.png/.pdf")


if __name__ == "__main__":
    main()
```
