## Supplemental Code for "Unifying multimodal single-cell data with a mixture-of-experts *β*-variational autoencoder framework": run_multiome_hparam_search.py

```
# run_multiome_hparam_search.py

import json
import time
from copy import deepcopy
from typing import Dict, Any, List, Tuple

import numpy as np
import scanpy as sc
from torch.utils.data import DataLoader, Subset

from univi.config import UniVIConfig, ModalityConfig, TrainingConfig
from univi.data import MultiModalDataset
from univi.trainer import UniVITrainer
from univi.models.univi import UniVIMultiModalVAE
from univi import evaluation as univi_eval


def _prepare_multiome_views(rna, atac):
    """
    Given aligned RNA + ATAC AnnData objects, precompute multiple 'views'
    per modality corresponding to different decoder distributions.
    """
    # -------- RNA views --------
    rna_nb = rna.copy()
    if "counts" in rna_nb.layers:
        rna_nb.X = rna_nb.layers["counts"].copy()
    else:
        rna_nb.layers["counts"] = rna_nb.X.copy()

    rna_gauss = rna.copy()
    if "log1p" in rna_gauss.layers:
        rna_gauss.X = rna_gauss.layers["log1p"].copy()
    else:
        rna_gauss.layers["counts"] = rna_gauss.X.copy()
        sc.pp.normalize_total(rna_gauss, target_sum=1e4)
        sc.pp.log1p(rna_gauss)
        rna_gauss.layers["log1p"] = rna_gauss.X.copy()
    sc.pp.scale(rna_gauss, max_value=10)

    rna_views = {
        "rna_nb": rna_nb,
        "rna_gauss": rna_gauss,
    }

    rna_decoder_options = [
        {"name": "rna_nb_counts", "view_key": "rna_nb", "likelihood": "nb"},
        {"name": "rna_zinb_counts", "view_key": "rna_nb", "likelihood": "zinb"},
        {"name": "rna_gauss_log1p_scaled", "view_key": "rna_gauss", "likelihood": "gaussian"},
    ]

    # -------- ATAC views --------
    atac_nb = atac.copy()
    if "counts" in atac_nb.layers:
        atac_nb.X = atac_nb.layers["counts"].copy()
    else:
        atac_nb.layers["counts"] = atac_nb.X.copy()

    atac_gauss = atac.copy()
    if "counts" in atac_gauss.layers:
        atac_gauss.X = atac_gauss.layers["counts"].copy()
    else:
        atac_gauss.layers["counts"] = atac_gauss.X.copy()
    sc.pp.normalize_total(atac_gauss, target_sum=1e4)
    sc.pp.log1p(atac_gauss)
    sc.pp.scale(atac_gauss, max_value=10)

    atac_views = {
        "atac_nb": atac_nb,
        "atac_gauss": atac_gauss,
    }

    atac_decoder_options = [
        {"name": "atac_nb_counts", "view_key": "atac_nb", "likelihood": "nb"},
        {"name": "atac_poisson_counts", "view_key": "atac_nb", "likelihood": "poisson"},
        {"name": "atac_zinb_counts", "view_key": "atac_nb", "likelihood": "zinb"},
        {"name": "atac_gauss_lognorm_scaled", "view_key": "atac_gauss", "likelihood": "gaussian"},
    ]

    return rna_views, atac_views, rna_decoder_options, atac_decoder_options


def _iter_hparam_configs(search_space: Dict[str, List[Any]], max_configs: int, seed: int = 0):
    rng = np.random.default_rng(seed)
    keys = list(search_space.keys())
    for _ in range(max_configs):
        hp = {}
        for k in keys:
            options = search_space[k]
            idx = rng.integers(len(options))
            hp[k] = options[idx]
        yield hp


def run_multiome_hparam_search(
    rna,
    atac,
    device: str = "cuda",
    max_configs: int = 50,
    seed: int = 0,
    out_json: str | None = None,
) -> Tuple[Dict[str, Any], List[Dict[str, Any]]]:
    """
    Hyperparameter search for 2-modal multiome (RNA + ATAC) with
    different architectures and decoder distributions.

    Parameters
    ----------
    rna, atac : AnnData
        Aligned multiome RNA and ATAC AnnData objects.
    device : str
        "cuda" or "cpu".
    max_configs : int
        Number of random configurations to evaluate.
    seed : int
        Random seed for hparam sampling and splits.
    out_json : str or None
        If not None, write all results (list of dicts) to this JSON file.

    Returns
    -------
    best_config : dict
        Record for the best config (contains hparams, metrics, score).
    all_results : list of dict
        All configuration records.
    """

    # Precompute views + decoder options
    rna_views, atac_views, rna_decoder_options, atac_decoder_options = _prepare_multiome_views(rna, atac)

    # Arch options
    rna_arch_options = [
        {"name": "rna_med2",  "enc": [512, 256],         "dec": [256, 512]},
        {"name": "rna_wide2", "enc": [1024, 512],        "dec": [512, 1024]},
        {"name": "rna_wide3", "enc": [1024, 512, 256],   "dec": [256, 512, 1024]},
    ]

    atac_arch_options = [
        {"name": "atac_med2",  "enc": [512, 256],         "dec": [256, 512]},
        {"name": "atac_wide2", "enc": [1024, 512],        "dec": [512, 1024]},
        {"name": "atac_wide3", "enc": [2048, 1024, 512],  "dec": [512, 1024, 2048]},
    ]

    search_space = {
        "latent_dim":       [10, 20, 32, 40, 50, 64, 82, 120, 160, 200],
        "beta":             [0.0, 10.0, 20.0, 30.0, 40.0, 50.0, 60.0, 80.0, 100.0, 160.0, 200.0],
        "gamma":            [0.0, 10.0, 20.0, 30.0, 40.0, 50.0, 60.0, 80.0, 100.0, 160.0, 200.0],
        "lr":               [1e-3, 5e-4],
        "weight_decay":     [1e-4, 1e-5],
        "encoder_dropout":  [0.0, 0.1],
        "decoder_batchnorm":[False, True],
        "rna_arch":         rna_arch_options,
        "atac_arch":        atac_arch_options,
        "rna_input":        rna_decoder_options,
        "atac_input":       atac_decoder_options,
    }

    # fixed train/val/test split
    n_cells = rna.n_obs
    indices = np.arange(n_cells)
    rng = np.random.default_rng(seed)
    rng.shuffle(indices)

    frac_train = 0.8
    frac_val   = 0.1
    n_train = int(frac_train * n_cells)
    n_val   = int(frac_val   * n_cells)

    train_idx = indices[:n_train]
    val_idx   = indices[n_train:n_train + n_val]
    test_idx  = indices[n_train + n_val:]

    results: List[Dict[str, Any]] = []
    best_score = np.inf
    best_config: Dict[str, Any] | None = None

    config_id = 0

    for hp in _iter_hparam_configs(search_space, max_configs=max_configs, seed=seed):
        config_id += 1
        print("="*80)
        print(f"[Config {config_id}] Hyperparameters:")
        print(json.dumps({
            "latent_dim":   hp["latent_dim"],
            "beta":         hp["beta"],
            "gamma":        hp["gamma"],
            "lr":           hp["lr"],
            "weight_decay": hp["weight_decay"],
            "encoder_dropout": hp["encoder_dropout"],
            "decoder_batchnorm": hp["decoder_batchnorm"],
            "rna_arch":     hp["rna_arch"]["name"],
            "atac_arch":    hp["atac_arch"]["name"],
            "rna_input":    hp["rna_input"]["name"],
            "atac_input":   hp["atac_input"]["name"],
        }, indent=2))
        print("="*80)

        # select views + likelihoods
        rna_view_key   = hp["rna_input"]["view_key"]
        rna_likelihood = hp["rna_input"]["likelihood"]

        atac_view_key   = hp["atac_input"]["view_key"]
        atac_likelihood = hp["atac_input"]["likelihood"]

        rna_view  = rna_views[rna_view_key]
        atac_view = atac_views[atac_view_key]

        adata_dict = {"rna": rna_view, "atac": atac_view}

        univi_cfg = UniVIConfig(
            latent_dim=hp["latent_dim"],
            beta=hp["beta"],
            gamma=hp["gamma"],
            encoder_dropout=hp["encoder_dropout"],
            decoder_dropout=0.0,
            encoder_batchnorm=True,
            decoder_batchnorm=hp["decoder_batchnorm"],
            kl_anneal_start=0,
            kl_anneal_end=0,
            align_anneal_start=0,
            align_anneal_end=0,
            modalities=[
                ModalityConfig(
                    name="rna",
                    input_dim=rna_view.n_vars,
                    encoder_hidden=hp["rna_arch"]["enc"],
                    decoder_hidden=hp["rna_arch"]["dec"],
                    likelihood=rna_likelihood,
                ),
                ModalityConfig(
                    name="atac",
                    input_dim=atac_view.n_vars,
                    encoder_hidden=hp["atac_arch"]["enc"],
                    decoder_hidden=hp["atac_arch"]["dec"],
                    likelihood=atac_likelihood,
                ),
            ],
        )

        train_cfg = TrainingConfig(
            n_epochs=80,
            batch_size=256,
            lr=hp["lr"],
            weight_decay=hp["weight_decay"],
            device=device,
            log_every=5,
            grad_clip=5.0,
            num_workers=0,
            seed=seed,
            early_stopping=True,
            patience=15,
            min_delta=0.0,
        )

        dataset = MultiModalDataset(
            adata_dict=adata_dict,
            X_key="X",
            device=train_cfg.device,
        )
        train_ds = Subset(dataset, train_idx)
        val_ds   = Subset(dataset, val_idx)

        train_loader = DataLoader(
            train_ds,
            batch_size=train_cfg.batch_size,
            shuffle=True,
            num_workers=train_cfg.num_workers,
        )
        val_loader = DataLoader(
            val_ds,
            batch_size=train_cfg.batch_size,
            shuffle=False,
            num_workers=train_cfg.num_workers,
        )
        
        loss_mode = cfg.get("model", {}).get("loss_mode", "lite")

        model = UniVIMultiModalVAE(univi_cfg, loss_mode=loss_mode).to(train_cfg.device)
        trainer = UniVITrainer(
            model=model,
            train_loader=train_loader,
            val_loader=val_loader,
            config=train_cfg,
        )

        start_time = time.time()
        best_val_loss = trainer.train()
        elapsed = (time.time() - start_time) / 60.0

        print(f"[Config {config_id}] Done in {elapsed:.1f} min")
        print(f"  best_val_loss              = {best_val_loss:.3f}")

        # Validation FOSCTTM
        rna_val  = rna_view[val_idx].copy()
        atac_val = atac_view[val_idx].copy()

        z_rna_val  = trainer.encode_modality(rna_val, modality="rna")
        z_atac_val = trainer.encode_modality(atac_val, modality="atac")

        foscttm = univi_eval.compute_foscttm(z_rna_val, z_atac_val)
        print(f"  FOSCTTM (RNA vs ATAC, val) = {foscttm:.4f}")

        # composite score (tune this as you like)
        score = best_val_loss + 1e4 * foscttm

        print(f"  Composite score            = {score:.3f}")

        record = {
            "config_id": config_id,
            "hparams": hp,
            "best_val_loss": float(best_val_loss),
            "foscttm": float(foscttm),
            "score": float(score),
        }
        results.append(record)

        if score < best_score:
            best_score = score
            best_config = record
            print(f"--> New best config (id={config_id}) with score={score:.3f}")

    if out_json is not None:
        with open(out_json, "w") as f:
            json.dump(results, f, indent=2)

    print("\nBest config:")
    if best_config is not None:
        print(json.dumps(best_config["hparams"], indent=2))

    return best_config, results


if __name__ == "__main__":
    print(
        "This module is intended to be imported and run from a notebook.\n"
        "Example:\n"
        "  from univi.hparam_search_multiome import run_multiome_hparam_search\n"
        "  best_cfg, all_results = run_multiome_hparam_search(rna, atac, device='cuda', max_configs=50)\n"
    )
```
