## Supplemental Code for "Unifying multimodal single-cell data with a mixture-of-experts *β*-variational autoencoder framework": train_univi.py

```
#!/usr/bin/env python3
"""
train_univi.py
--------------
CLI entry point for training a UniVI model from a JSON parameter file.

Usage
-----
python scripts/train_univi.py \\
    --config  parameter_files/params_citeseq_pbmc_GR_fig2_3.json \\
    --outdir  runs/citeseq_v1_fig2_3 \\
    --data-root /path/to/data \\
    [--device cuda] [--seed 0] [--resume runs/.../checkpoints/univi_checkpoint.pt]
"""

import argparse
import csv
import json
import shutil
import time
import warnings
from pathlib import Path

import numpy as np
import scanpy as sc
import torch
from torch.utils.data import DataLoader, Subset

from univi import UniVIMultiModalVAE, ModalityConfig, UniVIConfig, TrainingConfig
from univi.data import (
    MultiModalDataset,
    align_paired_obs_names,
    collate_multimodal_xy_recon,
)
from univi.trainer import UniVITrainer
from univi.utils.seed import set_seed
from univi.utils.logging import get_logger

logger = get_logger(__name__)
warnings.filterwarnings("ignore", category=FutureWarning)


# ---------------------------------------------------------------------------
# Argument parsing
# ---------------------------------------------------------------------------

def parse_args():
    p = argparse.ArgumentParser(
        description="Train a UniVI model from a JSON parameter file."
    )
    p.add_argument(
        "--config", required=True,
        help="Path to JSON parameter file.",
    )
    p.add_argument(
        "--outdir", required=True,
        help="Output directory for checkpoints, embeddings, logs.",
    )
    p.add_argument(
        "--data-root", default=".",
        help="Root directory prepended to data filenames in the config.",
    )
    p.add_argument(
        "--device", default=None,
        help="Device override: 'cuda', 'mps', 'cpu'. Auto-detects if None.",
    )
    p.add_argument(
        "--seed", type=int, default=None,
        help="Random seed override (default: from config split.seed, or 0).",
    )
    p.add_argument(
        "--resume", default=None,
        help="Path to an existing checkpoint to resume training from.",
    )
    p.add_argument(
        "--no-save-splits", action="store_true",
        help="Skip saving train/val/test index arrays to disk.",
    )
    return p.parse_args()


# ---------------------------------------------------------------------------
# Device detection
# ---------------------------------------------------------------------------

def resolve_device(device_arg, config_device=None):
    if device_arg is not None:
        return device_arg
    if config_device is not None:
        return config_device
    if torch.cuda.is_available():
        return "cuda"
    if getattr(torch.backends, "mps", None) is not None and torch.backends.mps.is_available():
        return "mps"
    if hasattr(torch, "xpu") and torch.xpu.is_available():
        return "xpu"
    return "cpu"


# ---------------------------------------------------------------------------
# Preprocessing helpers
# ---------------------------------------------------------------------------

def preprocess_rna(adata, cfg: dict):
    adata.layers["counts"] = adata.X.copy()
    sc.pp.normalize_total(adata, target_sum=cfg.get("normalize_total", 1e4))
    sc.pp.log1p(adata)
    adata.raw = adata
    sc.pp.highly_variable_genes(
        adata,
        flavor=cfg.get("hvg_flavor", "seurat_v3"),
        n_top_genes=cfg.get("n_hvg", 2000),
        subset=True,
    )
    sc.pp.scale(adata, max_value=cfg.get("scale_max_value", 10))
    return adata


def preprocess_rna_apply(adata, ref_adata, cfg: dict):
    """Apply RNA transforms fitted on ref_adata to a new (query) adata."""
    adata.layers["counts"] = adata.X.copy()
    sc.pp.normalize_total(adata, target_sum=cfg.get("normalize_total", 1e4))
    sc.pp.log1p(adata)
    if "highly_variable" in ref_adata.var.columns:
        shared = adata.var_names.intersection(ref_adata.var_names[ref_adata.var["highly_variable"]])
        adata = adata[:, shared].copy()
    sc.pp.scale(adata, max_value=cfg.get("scale_max_value", 10))
    return adata


def preprocess_adt(adata, cfg: dict):
    adata.layers["counts"] = adata.X.copy()
    X = adata.X.toarray() if hasattr(adata.X, "toarray") else np.asarray(adata.X)
    logX = np.log1p(X)
    adata.X = logX - logX.mean(axis=1, keepdims=True)
    if cfg.get("scale", True):
        sc.pp.scale(adata, zero_center=True, max_value=cfg.get("scale_max_value", 10))
    if cfg.get("clip") is not None:
        lo, hi = cfg["clip"]
        X_arr = adata.X.toarray() if hasattr(adata.X, "toarray") else np.asarray(adata.X)
        adata.X = np.clip(X_arr, lo, hi)
    return adata


def preprocess_atac(adata, cfg: dict):
    from sklearn.decomposition import TruncatedSVD

    adata.layers["counts"] = adata.X.copy()
    X = adata.X.tocsr() if hasattr(adata.X, "tocsr") else adata.X
    cell_sum = np.asarray(X.sum(axis=1)).ravel()
    cell_sum[cell_sum == 0] = 1.0
    tf = X.multiply(1.0 / cell_sum[:, None])
    df = np.asarray((X > 0).sum(axis=0)).ravel()
    idf = np.log1p(X.shape[0] / (1.0 + df))
    X_tfidf = tf.multiply(idf)

    n_components = cfg.get("lsi_n_components", 100)
    drop_lsi1 = cfg.get("drop_lsi1", False)
    svd = TruncatedSVD(n_components=n_components + 1, random_state=0)
    X_lsi = svd.fit_transform(X_tfidf)
    adata.uns["_svd_model"] = svd

    start_col = 1 if drop_lsi1 else 0
    obsm_key = cfg.get("store_lsi_obsm_key", "X_lsi")
    adata.obsm[obsm_key] = X_lsi[:, start_col : start_col + n_components]
    return adata


# ---------------------------------------------------------------------------
# Split utilities
# ---------------------------------------------------------------------------

def build_split_indices(adata, cfg: dict):
    """Stratified (with optional per-class cap) or random train/val/test split."""
    n = adata.n_obs
    seed = cfg.get("seed", 0)
    rng = np.random.default_rng(seed)
    train_frac = cfg.get("train_frac", 0.80)
    val_frac   = cfg.get("val_frac",   0.10)
    stratify_by = cfg.get("stratify_by")
    cap = cfg.get("cap_per_class", {})

    if stratify_by is not None and stratify_by in adata.obs.columns:
        labels = adata.obs[stratify_by].astype(str).values
        unique_labels = np.unique(labels)
        train_idx, val_idx, test_idx = [], [], []
        for lbl in unique_labels:
            idxs = np.where(labels == lbl)[0]
            rng.shuffle(idxs)
            n_tr = min(len(idxs), cap.get("train", int(np.ceil(len(idxs) * train_frac))))
            n_va = min(len(idxs) - n_tr, cap.get("val", int(np.ceil(len(idxs) * val_frac))))
            train_idx.extend(idxs[:n_tr].tolist())
            val_idx.extend(idxs[n_tr : n_tr + n_va].tolist())
            test_idx.extend(idxs[n_tr + n_va:].tolist())
        return np.array(train_idx), np.array(val_idx), np.array(test_idx)
    else:
        idx = np.arange(n)
        rng.shuffle(idx)
        n_tr = int(train_frac * n)
        n_va = int(val_frac * n)
        return idx[:n_tr], idx[n_tr : n_tr + n_va], idx[n_tr + n_va:]


# ---------------------------------------------------------------------------
# Config -> UniVI dataclasses
# ---------------------------------------------------------------------------

def build_univi_config(model_cfg: dict, data_cfg: dict, adata_dict: dict) -> UniVIConfig:
    X_key_by_mod = data_cfg.get("X_key_by_mod", {})
    mod_cfgs = []
    for m in model_cfg["modalities"]:
        name = m["name"]
        xkey = X_key_by_mod.get(name, "X")
        if xkey.startswith("obsm:"):
            obsm_key = xkey.replace("obsm:", "")
            input_dim = adata_dict[name].obsm[obsm_key].shape[1]
        else:
            input_dim = adata_dict[name].n_vars
        mod_cfgs.append(
            ModalityConfig(
                name=name,
                input_dim=input_dim,
                encoder_hidden=m["encoder_hidden"],
                decoder_hidden=m["decoder_hidden"],
                likelihood=m.get("likelihood", "gaussian"),
            )
        )
    return UniVIConfig(
        latent_dim=model_cfg.get("latent_dim", 30),
        beta=model_cfg.get("beta", 1.0),
        gamma=model_cfg.get("gamma", 1.0),
        encoder_dropout=model_cfg.get("encoder_dropout", 0.10),
        decoder_dropout=model_cfg.get("decoder_dropout", 0.05),
        encoder_batchnorm=model_cfg.get("encoder_batchnorm", True),
        decoder_batchnorm=model_cfg.get("decoder_batchnorm", False),
        kl_anneal_start=model_cfg.get("kl_anneal_start", 0),
        kl_anneal_end=model_cfg.get("kl_anneal_end", 0),
        align_anneal_start=model_cfg.get("align_anneal_start", 0),
        align_anneal_end=model_cfg.get("align_anneal_end", 0),
        modalities=mod_cfgs,
    )


def build_training_config(train_dict: dict, device: str) -> TrainingConfig:
    return TrainingConfig(
        n_epochs=train_dict.get("n_epochs", 1000),
        batch_size=train_dict.get("batch_size", 256),
        lr=train_dict.get("lr", 1e-3),
        weight_decay=train_dict.get("weight_decay", 1e-4),
        device=device,
        early_stopping=train_dict.get("early_stopping", True),
        patience=train_dict.get("patience", 50),
        best_epoch_warmup=train_dict.get("best_epoch_warmup", 50),
        log_every=train_dict.get("log_every", 25),
    )


# ---------------------------------------------------------------------------
# Main
# ---------------------------------------------------------------------------

def main():
    args = parse_args()

    with open(args.config) as f:
        full_cfg = json.load(f)

    exp_cfg        = full_cfg.get("experiment", {})
    data_cfg       = full_cfg["data"]
    model_cfg      = full_cfg["model"]
    train_cfg_dict = full_cfg["training"]

    seed   = args.seed if args.seed is not None else data_cfg.get("split", {}).get("seed", 0)
    set_seed(seed)

    device = resolve_device(args.device, train_cfg_dict.get("device"))
    logger.info(f"Experiment : {exp_cfg.get('name', Path(args.config).stem)}")
    logger.info(f"Device     : {device}  |  Seed: {seed}")

    # Output directories
    outdir   = Path(args.outdir)
    ckpt_dir = outdir / "checkpoints"
    log_dir  = outdir / "logs"
    for d in [ckpt_dir, log_dir]:
        d.mkdir(parents=True, exist_ok=True)

    data_root  = Path(args.data_root)
    modalities = data_cfg["modalities"]
    preproc    = data_cfg.get("preprocessing", {})

    # Load AnnData
    adata_dict = {}
    for mod in modalities:
        fpath = data_root / data_cfg[f"{mod}_filename"]
        logger.info(f"Loading {mod}: {fpath}")
        adata_dict[mod] = sc.read_h5ad(fpath)

    # Preprocess (train-fit transforms)
    if "rna"  in adata_dict: adata_dict["rna"]  = preprocess_rna(adata_dict["rna"],   preproc.get("rna",  {}))
    if "adt"  in adata_dict: adata_dict["adt"]  = preprocess_adt(adata_dict["adt"],   preproc.get("adt",  {}))
    if "atac" in adata_dict: adata_dict["atac"] = preprocess_atac(adata_dict["atac"], preproc.get("atac", {}))

    align_paired_obs_names(adata_dict)

    # Split
    split_cfg = data_cfg.get("split", {})
    ref_mod   = modalities[0]
    train_idx, val_idx, test_idx = build_split_indices(adata_dict[ref_mod], split_cfg)
    logger.info(f"Split  -> train: {len(train_idx)}  val: {len(val_idx)}  test: {len(test_idx)}")

    if not args.no_save_splits:
        np.savez(outdir / "splits.npz",
                 train_idx=train_idx, val_idx=val_idx, test_idx=test_idx, seed=seed)

    # Dataset + loaders
    X_key_by_mod = data_cfg.get("X_key_by_mod", {m: "X" for m in modalities})
    dataset = MultiModalDataset(
        adata_dict=adata_dict,
        device=None,
        X_key=X_key_by_mod,
    )
    bs = train_cfg_dict.get("batch_size", 256)
    train_loader = DataLoader(Subset(dataset, train_idx), batch_size=bs, shuffle=True,
                              num_workers=0, collate_fn=collate_multimodal_xy_recon)
    val_loader   = DataLoader(Subset(dataset, val_idx),   batch_size=bs, shuffle=False,
                              num_workers=0, collate_fn=collate_multimodal_xy_recon) \
                   if len(val_idx) > 0 else None

    # Build model
    univi_cfg = build_univi_config(model_cfg, data_cfg, adata_dict)
    train_cfg = build_training_config(train_cfg_dict, device)
    model = UniVIMultiModalVAE(
        univi_cfg,
        loss_mode=model_cfg.get("loss_mode", "v1"),
        v1_recon=model_cfg.get("v1_recon", "avg"),
        normalize_v1_terms=model_cfg.get("normalize_v1_terms", True),
    ).to(device)

    # Optionally resume
    if args.resume:
        logger.info(f"Resuming from: {args.resume}")
        ckpt_r = torch.load(args.resume, map_location=device, weights_only=False)
        model.load_state_dict(ckpt_r["model_state_dict"])

    # Train
    t0 = time.time()
    trainer = UniVITrainer(
        model=model, train_loader=train_loader, val_loader=val_loader,
        train_cfg=train_cfg, device=device,
    )
    history   = trainer.fit()
    elapsed   = time.time() - t0
    best_epoch = getattr(trainer, "best_epoch", None)
    logger.info(f"Training complete in {elapsed:.1f}s  |  Best epoch: {best_epoch}")

    # Save checkpoint
    ckpt = {
        "model_state_dict": model.state_dict(),
        "model_config":     univi_cfg,
        "train_cfg":        train_cfg,
        "history":          history,
        "best_epoch":       best_epoch,
        "elapsed_seconds":  elapsed,
        "seed":             seed,
        "param_file":       str(args.config),
        "experiment":       exp_cfg,
    }
    ckpt_path = ckpt_dir / "univi_checkpoint.pt"
    torch.save(ckpt, ckpt_path)
    logger.info(f"Checkpoint saved -> {ckpt_path}")

    shutil.copy(args.config, outdir / "params_used.json")

    # Save history CSV
    if history:
        history_path = log_dir / "history.csv"
        if isinstance(history, dict):
            rows = [{"epoch": k, **v} if isinstance(v, dict) else {"epoch": k, "loss": v} for k, v in history.items()]
        else:
            rows = history if isinstance(history[0], dict) else [{"epoch": i, "loss": v} for i, v in enumerate(history)]
        with open(history_path, "w", newline="") as fh:
            writer = csv.DictWriter(fh, fieldnames=list(rows[0].keys()))
            writer.writeheader()
            writer.writerows(rows)
        logger.info(f"History CSV -> {history_path}")

    logger.info("Done.")


if __name__ == "__main__":
    main()
```
