## Supplemental Code for "Unifying multimodal single-cell data with a mixture-of-experts *β*-variational autoencoder framework": __init__.py

```
# univi/__init__.py

from __future__ import annotations

from typing import Any, List

__version__ = "0.4.7"

# Eager (fast/light) public API
from .config import ModalityConfig, UniVIConfig, TrainingConfig
from .models import UniVIMultiModalVAE
from . import matching

__all__ = [
    "__version__",
    # configs
    "ModalityConfig",
    "UniVIConfig",
    "TrainingConfig",
    # model
    "UniVIMultiModalVAE",
    # lightweight module
    "matching",
    # model state
    "save_checkpoint",
    "load_checkpoint",
    "restore_checkpoint",
    # lazy exports
    "UniVITrainer",
    "write_univi_latent",
    "MultiModalDataset",
    "pipeline",
    "diagnostics",
    # modules
    "evaluation",
    "plotting",
    # eval convenience (optional)
    "encode_adata",
    "evaluate_alignment",
    # interpretability
    "interpretability",
    "fused_encode_with_meta_and_attn",
    "feature_importance_for_head",
    "top_cross_modal_feature_pairs_from_attn",
]


def __getattr__(name: str) -> Any:
    """
    Lazy exports keep `import univi` fast/light and avoid heavy deps unless needed.
    """
    # ---- training ----
    if name == "UniVITrainer":
        from .trainer import UniVITrainer
        return UniVITrainer

    # ---- IO ----
    if name == "write_univi_latent":
        from .utils.io import write_univi_latent
        return write_univi_latent

    # ---- data ----
    if name == "MultiModalDataset":
        from .data import MultiModalDataset
        return MultiModalDataset

    # ---- model state ----
    if name in {"save_checkpoint", "load_checkpoint", "restore_checkpoint"}:
        from .utils.io import save_checkpoint, load_checkpoint, restore_checkpoint
        return {"save_checkpoint": save_checkpoint, "load_checkpoint": load_checkpoint, "restore_checkpoint": restore_checkpoint}[name]

    # ---- modules (return module objects) ----
    if name == "pipeline":
        from . import pipeline as _pipeline
        return _pipeline

    if name == "diagnostics":
        from . import diagnostics as _diagnostics
        return _diagnostics

    if name == "evaluation":
        from . import evaluation as _evaluation
        return _evaluation

    if name == "plotting":
        from . import plotting as _plotting
        return _plotting

    # ---- interpretability ----
    if name == "interpretability":
        from . import interpretability as _interpretability
        return _interpretability

    if name in {
        "fused_encode_with_meta_and_attn",
        "feature_importance_for_head",
        "top_cross_modal_feature_pairs_from_attn",
    }:
        from .interpretability import (
            fused_encode_with_meta_and_attn,
            feature_importance_for_head,
            top_cross_modal_feature_pairs_from_attn,
        )
        return {
            "fused_encode_with_meta_and_attn": fused_encode_with_meta_and_attn,
            "feature_importance_for_head": feature_importance_for_head,
            "top_cross_modal_feature_pairs_from_attn": top_cross_modal_feature_pairs_from_attn,
        }[name]

    # ---- eval convenience functions (re-export) ----
    if name in {"encode_adata", "evaluate_alignment"}:
        from .evaluation import encode_adata, evaluate_alignment
        return {"encode_adata": encode_adata, "evaluate_alignment": evaluate_alignment}[name]

    raise AttributeError(f"module {__name__!r} has no attribute {name!r}")


def __dir__() -> List[str]:
    return sorted(list(globals().keys()) + __all__)
```
