## Supplemental Code for "Unifying multimodal single-cell data with a mixture-of-experts *β*-variational autoencoder framework": cli.py

```
# univi/cli.py
from __future__ import annotations

import argparse
import os
import sys

from .pipeline import load_model_and_data, encode_latents_paired
from .diagnostics import export_supplemental_table_s1


def main(argv=None):
    ap = argparse.ArgumentParser(prog="univi")
    sub = ap.add_subparsers(dest="cmd", required=True)

    ap_s1 = sub.add_parser("export-s1", help="Export Supplemental_Table_S1.xlsx (env + hparams + dataset stats).")
    ap_s1.add_argument("--config", required=True)
    ap_s1.add_argument("--checkpoint", default=None)
    ap_s1.add_argument("--data-root", default=None)
    ap_s1.add_argument("--out", required=True)

    ap_encode = sub.add_parser("encode", help="Encode paired latents and save as .npz")
    ap_encode.add_argument("--config", required=True)
    ap_encode.add_argument("--checkpoint", required=True)
    ap_encode.add_argument("--data-root", default=None)
    ap_encode.add_argument("--out", required=True)
    ap_encode.add_argument("--device", default="cpu")
    ap_encode.add_argument("--batch-size", type=int, default=512)

    args = ap.parse_args(argv)

    if args.cmd == "export-s1":
        cfg, adata_dict, model, layer_by, xkey_by = load_model_and_data(
            args.config, checkpoint_path=args.checkpoint, data_root=args.data_root, device="cpu"
        )
        export_supplemental_table_s1(
            args.config,
            adata_dict,
            out_xlsx=args.out,
            layer_by=layer_by,
            xkey_by=xkey_by,
            extra_metrics=None,
        )
        return 0

    if args.cmd == "encode":
        cfg, adata_dict, model, layer_by, xkey_by = load_model_and_data(
            args.config, checkpoint_path=args.checkpoint, data_root=args.data_root, device=args.device
        )
        Z = encode_latents_paired(model, adata_dict, layer_by=layer_by, xkey_by=xkey_by, batch_size=args.batch_size, device=args.device, fused=True)
        os.makedirs(os.path.dirname(args.out) or ".", exist_ok=True)
        import numpy as np
        np.savez_compressed(args.out, **Z)
        return 0

    return 1


if __name__ == "__main__":
    raise SystemExit(main())
```
