## Supplemental Code for "Unifying multimodal single-cell data with a mixture-of-experts *β*-variational autoencoder framework": config.py

```
# univi/config.py
from __future__ import annotations

from dataclasses import dataclass, field
from typing import Dict, List, Optional, Literal, Sequence, Any, Union, Mapping


# =============================================================================
# Transformer + tokenizer config
# =============================================================================

@dataclass
class TransformerConfig:
    d_model: int
    num_heads: int
    num_layers: int
    dim_feedforward: int = 4096
    dropout: float = 0.1
    attn_dropout: float = 0.1
    activation: Literal["relu", "gelu"] = "gelu"
    pooling: Literal["cls", "mean"] = "mean"
    max_tokens: Optional[int] = None

    use_relpos_bias: bool = False
    relpos_num_bins: int = 32
    relpos_max_dist: float = 1e6  # basepairs


@dataclass
class TokenizerConfig:
    mode: Literal["topk_scalar", "topk_channels", "patch", "topk_embed"] = "topk_scalar"

    n_tokens: int = 256
    channels: Sequence[Literal["value", "rank", "dropout"]] = ("value",)

    patch_size: int = 32
    patch_proj_dim: Optional[int] = None

    add_cls_token: bool = False

    n_features: Optional[int] = None
    d_model: Optional[int] = None
    value_mlp_hidden: int = 256

    use_coords: bool = False
    chrom_vocab_size: int = 0
    coord_scale: float = 1e6
    feature_info: Optional[Dict[str, Any]] = None


# =============================================================================
# Core UniVI configs
# =============================================================================

LikelihoodName = Literal[
    # existing / common
    "gaussian", "normal", "mse", "gaussian_diag",
    "bernoulli", "poisson",
    "nb", "negative_binomial",
    "zinb", "zero_inflated_negative_binomial",
    "categorical", "cat", "ce", "cross_entropy", "multinomial", "softmax",
    "logistic_normal",
    # likely additions (safe to configure now; model/decoder can ignore until implemented)
    "beta",
    "beta_binomial",
    "zinb_with_libsize",
    "nb_with_libsize",
    "lognormal",
    "gamma",
    "student_t",
    "hurdle_poisson",
    "hurdle_nb",
]


@dataclass
class ModalityConfig:
    """
    Configuration for a single modality.

    Backwards compatible with prior versions.

    Added decoder/loss-related optional knobs so the model can support more
    likelihoods without changing the config schema again.
    """
    name: str
    input_dim: int
    encoder_hidden: List[int]
    decoder_hidden: List[int]
    likelihood: str = "gaussian"

    # ---- per-modality reconstruction weighting ----
    recon_weight: float = 1.0

    # ---- Count model settings (NB/ZINB and relatives) ----
    dispersion: str = "gene"  # e.g. "global", "gene", "feature" (model decides support)
    init_log_theta: float = 0.0

    # Optional library-size handling (for count decoders that use offsets)
    use_library_size: bool = False
    library_key: Optional[str] = None          # e.g. obs key if data loader uses obs-derived features
    library_log1p: bool = True                 # whether supplied library is already log1p-transformed
    predict_library: bool = False              # if you later add a library-size head/decoder branch

    # ---- Zero-inflation / hurdle settings ----
    use_zero_inflation: Optional[bool] = None  # None => infer from likelihood
    hurdle: bool = False                       # for hurdle-* likelihoods if implemented

    # ---- Bounded / proportion data (e.g. methylation fractions in [0,1]) ----
    # If you preprocess methylation to fractions, beta-like decoders often want clipping.
    clip_targets_min: Optional[float] = None
    clip_targets_max: Optional[float] = None

    # For beta/binomial-style likelihoods if you add them later
    concentration_init: float = 0.0           # log-concentration init or analogous parameter
    total_count_key: Optional[str] = None     # obs key / side-input key for trials if needed
    successes_are_fraction: bool = False      # True if x is fraction instead of counts

    # ---- Positive continuous data (if you add gamma/lognormal decoders) ----
    positive_eps: float = 1e-8

    # ---- Categorical modality support ----
    ignore_index: int = -1
    input_kind: Literal["matrix", "obs"] = "matrix"
    obs_key: Optional[str] = None

    # ---- Optional masking / weighting hooks (future-proof) ----
    mask_key: Optional[str] = None            # if your dataset returns feature or sample masks
    sample_weight_key: Optional[str] = None   # if you later support weighted recon losses

    # ---- Encoder backend (per-modality only) ----
    encoder_type: Literal["mlp", "transformer"] = "mlp"
    transformer: Optional[TransformerConfig] = None
    tokenizer: Optional[TokenizerConfig] = None

    # ---- Arbitrary decoder/likelihood kwargs passthrough (future-proof) ----
    # Example:
    # decoder_kwargs={"dispersion":"gene-cell", "min_scale":1e-4}
    decoder_kwargs: Optional[Dict[str, Any]] = None


@dataclass
class ClassHeadConfig:
    """
    Supervised / adversarial head configuration.

    Backwards compatible defaults preserve the old behavior:
    - head_type defaults to categorical (even if n_classes==2)
    """
    name: str
    n_classes: int
    loss_weight: float = 1.0
    ignore_index: int = -1
    from_mu: bool = True
    warmup: int = 0

    adversarial: bool = False
    adv_lambda: float = 1.0

    # NEW (optional): allow passing what the model reads via getattr
    head_type: Optional[Literal["categorical", "binary"]] = None
    hidden_dims: Optional[List[int]] = None
    dropout: Optional[float] = None
    batchnorm: Optional[bool] = None
    activation: Optional[Literal["relu", "gelu", "elu", "leakyrelu", "silu", "tanh"]] = None
    pos_weight: float = 1.0  # binary only


@dataclass
class UniVIConfig:
    latent_dim: int
    modalities: List[ModalityConfig]

    beta: float = 1.0
    gamma: float = 1.0

    encoder_dropout: float = 0.0
    decoder_dropout: float = 0.0
    encoder_batchnorm: bool = True
    decoder_batchnorm: bool = False

    kl_anneal_start: int = 0
    kl_anneal_end: int = 0
    align_anneal_start: int = 0
    align_anneal_end: int = 0

    class_heads: Optional[List[ClassHeadConfig]] = None
    label_head_name: str = "label"

    fused_encoder_type: Literal["moe", "multimodal_transformer"] = "moe"
    fused_transformer: Optional[TransformerConfig] = None
    fused_modalities: Optional[Sequence[str]] = None
    fused_add_modality_embeddings: bool = True
    fused_require_all_modalities: bool = True

    # ------------------------------------------------------------------
    # MoE gating (learned per-cell modality weights for fusion)
    # ------------------------------------------------------------------
    use_moe_gating: bool = False
    moe_gating_type: Literal["per_modality", "shared"] = "per_modality"
    moe_gating_hidden: Optional[List[int]] = None
    moe_gating_dropout: float = 0.0
    moe_gating_batchnorm: bool = False
    moe_gating_activation: Literal["relu", "gelu", "elu", "leakyrelu", "silu", "tanh"] = "relu"
    moe_gate_eps: float = 1e-6

    # ------------------------------------------------------------------
    # Optional global reconstruction/likelihood defaults (future-proof)
    # These do NOT override per-modality fields unless your model chooses to.
    # ------------------------------------------------------------------
    default_dispersion: Optional[str] = None
    default_use_library_size: Optional[bool] = None
    default_positive_eps: Optional[float] = None

    # ------------------------------------------------------------------
    # Arbitrary model kwargs passthrough (future-proof)
    # ------------------------------------------------------------------
    model_kwargs: Optional[Dict[str, Any]] = None

    def validate(self) -> None:
        if int(self.latent_dim) <= 0:
            raise ValueError(f"latent_dim must be > 0, got {self.latent_dim}")

        names = [m.name for m in self.modalities]
        if len(set(names)) != len(names):
            dupes = sorted({n for n in names if names.count(n) > 1})
            raise ValueError(f"Duplicate modality names in cfg.modalities: {dupes}")

        mod_by_name: Dict[str, ModalityConfig] = {m.name: m for m in self.modalities}

        for m in self.modalities:
            if int(m.input_dim) <= 0:
                raise ValueError(f"Modality {m.name!r}: input_dim must be > 0, got {m.input_dim}")

            if float(getattr(m, "recon_weight", 1.0)) < 0.0:
                raise ValueError(f"Modality {m.name!r}: recon_weight must be >= 0, got {m.recon_weight}")

            lk = (m.likelihood or "").lower().strip()

            # ---- categorical checks ----
            if lk in ("categorical", "cat", "ce", "cross_entropy", "multinomial", "softmax"):
                if int(m.input_dim) < 2:
                    raise ValueError(f"Categorical modality {m.name!r}: input_dim must be n_classes >= 2.")
                if m.input_kind == "obs" and not m.obs_key:
                    raise ValueError(f"Categorical modality {m.name!r}: input_kind='obs' requires obs_key.")

            # ---- count-like checks ----
            if lk in (
                "nb", "negative_binomial",
                "zinb", "zero_inflated_negative_binomial",
                "nb_with_libsize", "zinb_with_libsize",
                "hurdle_nb", "hurdle_poisson",
            ):
                disp = str(getattr(m, "dispersion", "gene")).lower().strip()
                # keep permissive (model can support subset)
                allowed_disp = {"global", "gene", "feature", "gene_cell", "gene-cell"}
                if disp not in allowed_disp:
                    raise ValueError(
                        f"Modality {m.name!r}: unsupported dispersion={m.dispersion!r}. "
                        f"Expected one of {sorted(allowed_disp)}."
                    )

                if float(getattr(m, "positive_eps", 1e-8)) <= 0:
                    raise ValueError(f"Modality {m.name!r}: positive_eps must be > 0.")

                if bool(getattr(m, "use_library_size", False)) and m.library_key is None:
                    # not always required if library is computed from x on-the-fly, so only warn via ValueError? No:
                    # keep permissive for backward compatibility.
                    pass

            # ---- bounded / proportion checks ----
            if lk in ("beta", "beta_binomial"):
                cmin = getattr(m, "clip_targets_min", None)
                cmax = getattr(m, "clip_targets_max", None)
                if cmin is not None and cmax is not None and float(cmin) >= float(cmax):
                    raise ValueError(
                        f"Modality {m.name!r}: clip_targets_min must be < clip_targets_max "
                        f"(got {cmin} >= {cmax})."
                    )
                if float(getattr(m, "positive_eps", 1e-8)) <= 0:
                    raise ValueError(f"Modality {m.name!r}: positive_eps must be > 0.")

            # ---- positive continuous checks ----
            if lk in ("gamma", "lognormal", "student_t"):
                if float(getattr(m, "positive_eps", 1e-8)) <= 0:
                    raise ValueError(f"Modality {m.name!r}: positive_eps must be > 0.")

            # ---- decoder kwargs sanity ----
            if m.decoder_kwargs is not None and not isinstance(m.decoder_kwargs, dict):
                raise ValueError(f"Modality {m.name!r}: decoder_kwargs must be dict or None.")

            # ---- encoder backend checks ----
            enc_type = (m.encoder_type or "mlp").lower().strip()
            if enc_type not in ("mlp", "transformer"):
                raise ValueError(
                    f"Modality {m.name!r}: encoder_type must be 'mlp' or 'transformer', got {m.encoder_type!r}"
                )

            if enc_type == "transformer":
                if m.transformer is None:
                    raise ValueError(f"Modality {m.name!r}: encoder_type='transformer' requires transformer config.")
                if m.tokenizer is None:
                    raise ValueError(f"Modality {m.name!r}: encoder_type='transformer' requires tokenizer config.")
                _validate_tokenizer(m.name, m.tokenizer)

        # -------------------------
        # fused encoder checks
        # -------------------------
        fe = (self.fused_encoder_type or "moe").lower().strip()
        if fe not in ("moe", "multimodal_transformer"):
            raise ValueError(
                f"fused_encoder_type must be 'moe' or 'multimodal_transformer', got {self.fused_encoder_type!r}"
            )

        if fe == "multimodal_transformer":
            if self.fused_transformer is None:
                raise ValueError("fused_encoder_type='multimodal_transformer' requires UniVIConfig.fused_transformer.")

            fused_names = list(self.fused_modalities) if self.fused_modalities is not None else list(mod_by_name.keys())
            if not fused_names:
                raise ValueError("fused_modalities is empty; expected at least one modality name.")

            missing = [n for n in fused_names if n not in mod_by_name]
            if missing:
                raise ValueError(f"fused_modalities contains unknown modalities: {missing}. Known: {list(mod_by_name)}")

            for n in fused_names:
                tok = mod_by_name[n].tokenizer
                if tok is None:
                    raise ValueError(
                        f"Fused multimodal transformer requires ModalityConfig.tokenizer for modality {n!r}."
                    )
                _validate_tokenizer(n, tok)

        # -------------------------
        # MoE gating checks
        # -------------------------
        if bool(self.use_moe_gating):
            gt = (self.moe_gating_type or "per_modality").lower().strip()
            if gt not in ("per_modality", "shared"):
                raise ValueError(
                    f"moe_gating_type must be 'per_modality' or 'shared', got {self.moe_gating_type!r}"
                )

            h = self.moe_gating_hidden
            if h is not None:
                if not isinstance(h, (list, tuple)) or len(h) == 0:
                    raise ValueError("moe_gating_hidden must be a non-empty list[int] if provided.")
                bad = [x for x in h if int(x) <= 0]
                if bad:
                    raise ValueError(f"moe_gating_hidden must contain only positive ints; bad entries: {bad}")

            if float(self.moe_gating_dropout) < 0.0:
                raise ValueError(f"moe_gating_dropout must be >= 0, got {self.moe_gating_dropout}")
            if float(self.moe_gate_eps) < 0.0:
                raise ValueError(f"moe_gate_eps must be >= 0, got {self.moe_gate_eps}")

            act = (self.moe_gating_activation or "relu").lower().strip()
            if act not in ("relu", "gelu", "elu", "leakyrelu", "silu", "tanh"):
                raise ValueError(
                    "moe_gating_activation must be one of "
                    "['relu','gelu','elu','leakyrelu','silu','tanh'], "
                    f"got {self.moe_gating_activation!r}"
                )

        # -------------------------
        # class head checks
        # -------------------------
        if self.class_heads is not None:
            hn = [h.name for h in self.class_heads]
            if len(set(hn)) != len(hn):
                dupes = sorted({n for n in hn if hn.count(n) > 1})
                raise ValueError(f"Duplicate class head names in cfg.class_heads: {dupes}")

            for h in self.class_heads:
                ht = (h.head_type or "categorical")
                if int(h.n_classes) < 2 and ht != "binary":
                    raise ValueError(f"Class head {h.name!r}: n_classes must be >= 2 for categorical heads.")
                if float(h.loss_weight) < 0:
                    raise ValueError(f"Class head {h.name!r}: loss_weight must be >= 0.")
                if int(h.warmup) < 0:
                    raise ValueError(f"Class head {h.name!r}: warmup must be >= 0.")
                if float(getattr(h, "adv_lambda", 1.0)) < 0.0:
                    raise ValueError(f"Class head {h.name!r}: adv_lambda must be >= 0.")
                if float(getattr(h, "pos_weight", 1.0)) < 0.0:
                    raise ValueError(f"Class head {h.name!r}: pos_weight must be >= 0.")

                if h.hidden_dims is not None:
                    if not isinstance(h.hidden_dims, (list, tuple)) or len(h.hidden_dims) == 0:
                        raise ValueError(f"Class head {h.name!r}: hidden_dims must be non-empty if provided.")
                    bad = [x for x in h.hidden_dims if int(x) <= 0]
                    if bad:
                        raise ValueError(f"Class head {h.name!r}: hidden_dims must be positive ints; bad: {bad}")

        # -------------------------
        # annealing checks
        # -------------------------
        for k in ("kl_anneal_start", "kl_anneal_end", "align_anneal_start", "align_anneal_end"):
            v = int(getattr(self, k))
            if v < 0:
                raise ValueError(f"{k} must be >= 0, got {v}")

        # -------------------------
        # optional global defaults checks
        # -------------------------
        if self.default_positive_eps is not None and float(self.default_positive_eps) <= 0:
            raise ValueError("default_positive_eps must be > 0 if provided.")

        if self.model_kwargs is not None and not isinstance(self.model_kwargs, dict):
            raise ValueError("model_kwargs must be dict or None.")


def _validate_tokenizer(mod_name: str, tok: TokenizerConfig) -> None:
    mode = (tok.mode or "").lower().strip()
    if mode not in ("topk_scalar", "topk_channels", "patch", "topk_embed"):
        raise ValueError(
            f"Modality {mod_name!r}: tokenizer.mode must be one of "
            f"['topk_scalar','topk_channels','patch','topk_embed'], got {tok.mode!r}"
        )

    if mode in ("topk_scalar", "topk_channels", "topk_embed"):
        if int(tok.n_tokens) <= 0:
            raise ValueError(f"Modality {mod_name!r}: tokenizer.n_tokens must be > 0 for topk_*")

    if mode == "topk_channels":
        if not tok.channels:
            raise ValueError(f"Modality {mod_name!r}: tokenizer.channels must be non-empty for topk_channels")
        bad = [c for c in tok.channels if c not in ("value", "rank", "dropout")]
        if bad:
            raise ValueError(f"Modality {mod_name!r}: tokenizer.channels has invalid entries: {bad}")

    if mode == "topk_embed":
        if tok.n_features is None or int(tok.n_features) <= 0:
            raise ValueError(f"Modality {mod_name!r}: tokenizer.n_features must be set (>0) for topk_embed")
        if tok.d_model is None or int(tok.d_model) <= 0:
            raise ValueError(f"Modality {mod_name!r}: tokenizer.d_model must be set (>0) for topk_embed")
        if not tok.channels:
            raise ValueError(f"Modality {mod_name!r}: tokenizer.channels must be non-empty for topk_embed")
        bad = [c for c in tok.channels if c not in ("value", "rank", "dropout")]
        if bad:
            raise ValueError(f"Modality {mod_name!r}: tokenizer.channels has invalid entries: {bad}")

        if tok.use_coords:
            if int(tok.chrom_vocab_size) <= 0:
                raise ValueError(
                    f"Modality {mod_name!r}: tokenizer.chrom_vocab_size must be > 0 when use_coords=True"
                )
            if tok.feature_info is not None:
                for k in ("chrom", "start", "end"):
                    if k not in tok.feature_info:
                        raise ValueError(
                            f"Modality {mod_name!r}: tokenizer.feature_info missing key {k!r} (required for coords)"
                        )

    if mode == "patch":
        if int(tok.patch_size) <= 0:
            raise ValueError(f"Modality {mod_name!r}: tokenizer.patch_size must be > 0 for patch")
        if tok.patch_proj_dim is not None and int(tok.patch_proj_dim) <= 0:
            raise ValueError(f"Modality {mod_name!r}: tokenizer.patch_proj_dim must be > 0 if set")


@dataclass
class TrainingConfig:
    n_epochs: int = 200
    batch_size: int = 256
    lr: float = 1e-3
    weight_decay: float = 0.0
    device: str = "cpu"
    log_every: int = 10
    grad_clip: Optional[float] = None
    num_workers: int = 0
    seed: int = 0

    early_stopping: bool = False
    patience: int = 20
    min_delta: float = 0.0

    best_epoch_warmup: int = 0
```
