## Supplemental Code for "Unifying multimodal single-cell data with a mixture-of-experts *β*-variational autoencoder framework": data.py

```
# univi/data.py

from __future__ import annotations

from typing import Any, Dict, Mapping, Optional, Tuple, Union, List, Sequence

import os
import numpy as np
import scipy.sparse as sp
import pandas as pd
import anndata as ad
from anndata import AnnData

import torch
from torch.utils.data import Dataset

from .config import ModalityConfig

# -----------------------------------------------------------------------------
# Types
# -----------------------------------------------------------------------------
LayerSpec = Union[None, str, Mapping[str, Optional[str]]]
XKeySpec = Union[str, Mapping[str, str]]
LabelSpec = Union[
    np.ndarray,
    torch.Tensor,
    Sequence[int],
    Mapping[str, Union[np.ndarray, torch.Tensor, Sequence[int]]],
]

# For binomial / beta-binomial recon targets (e.g., methylome):
# recon_targets_spec = {
#   "meth": {"successes_layer": "meth_successes", "total_count_layer": "meth_total_count"}
# }
ReconTargetSpec = Mapping[str, Mapping[str, str]]


# -----------------------------------------------------------------------------
# Helpers
# -----------------------------------------------------------------------------
def _is_categorical_likelihood(lk: Optional[str]) -> bool:
    lk = (lk or "").lower().strip()
    return lk in ("categorical", "cat", "ce", "cross_entropy", "multinomial", "softmax")


def _get_matrix(adata_obj: AnnData, *, layer: Optional[str], X_key: str):
    if X_key != "X":
        if X_key not in adata_obj.obsm:
            raise KeyError("X_key=%r not found in adata.obsm. Keys=%s" % (X_key, list(adata_obj.obsm.keys())))
        return adata_obj.obsm[X_key]

    if layer is not None:
        if layer not in adata_obj.layers:
            raise KeyError("layer=%r not found in adata.layers. Keys=%s" % (layer, list(adata_obj.layers.keys())))
        return adata_obj.layers[layer]

    return adata_obj.X


def infer_input_dim(adata_obj: AnnData, *, layer: Optional[str], X_key: str) -> int:
    X = _get_matrix(adata_obj, layer=layer, X_key=X_key)
    if not hasattr(X, "shape") or len(X.shape) != 2:
        raise ValueError("Selected matrix for (layer=%r, X_key=%r) is not 2D." % (layer, X_key))
    return int(X.shape[1])


def align_paired_obs_names(
    adata_dict: Dict[str, AnnData],
    how: str = "intersection",
    require_nonempty: bool = True,
    sort: bool = True,
    copy: bool = True,
) -> Dict[str, AnnData]:
    if not adata_dict:
        raise ValueError("adata_dict is empty")
    if how != "intersection":
        raise ValueError("Unsupported how=%r. Only 'intersection' is supported." % how)

    names = list(adata_dict.keys())
    shared = None
    for nm in names:
        idx = adata_dict[nm].obs_names
        shared = idx if shared is None else shared.intersection(idx)

    if shared is None:
        shared = pd.Index([])

    if require_nonempty and len(shared) == 0:
        raise ValueError("No shared obs_names across modalities (intersection is empty).")

    if sort:
        shared = shared.sort_values()

    out: Dict[str, AnnData] = {}
    for nm in names:
        slc = adata_dict[nm][shared, :]
        out[nm] = slc.copy() if copy else slc
    return out


def _as_modality_map(
    spec: Union[str, None, Mapping[str, Any]],
    adata_dict: Dict[str, AnnData],
    kind: str,
) -> Dict[str, Any]:
    if isinstance(spec, Mapping):
        out = dict(spec)
    else:
        out = {k: spec for k in adata_dict.keys()}

    for k in adata_dict.keys():
        if k not in out:
            out[k] = None if kind == "layer" else "X"
    return out


# -----------------------------------------------------------------------------
# Dataset
# -----------------------------------------------------------------------------
class MultiModalDataset(Dataset):
    """
    Multi-modal AnnData-backed torch Dataset.

    Returns (depending on what you provide):
      - x_dict
      - (x_dict, y)
      - (x_dict, recon_targets)
      - (x_dict, y, recon_targets)

    Where:
      - x_dict: Dict[modality -> FloatTensor] (1D per item; collate stacks to (B, D))
      - y:
          * LongTensor scalar (back-compat), OR
          * dict[str -> LongTensor scalar] (multi-head)
      - recon_targets:
          * Dict[modality -> {"successes": FloatTensor(1D), "total_count": FloatTensor(1D)}]
        (collate stacks to (B, D) per target)

    Categorical modality support:
      - If modality_cfgs marks a modality as categorical with input_kind="obs",
        x_dict[modality] is a (1,) float tensor holding an integer code.
    """

    def __init__(
        self,
        adata_dict: Dict[str, AnnData],
        layer: LayerSpec = None,
        X_key: XKeySpec = "X",
        paired: bool = True,
        device: Optional[torch.device] = None,
        labels: Optional[LabelSpec] = None,
        dtype: torch.dtype = torch.float32,
        modality_cfgs: Optional[List[ModalityConfig]] = None,
        recon_targets_spec: Optional[ReconTargetSpec] = None,
    ):
        if not adata_dict:
            raise ValueError("adata_dict is empty")

        self.adata_dict: Dict[str, AnnData] = adata_dict
        self.modalities: List[str] = list(adata_dict.keys())
        self.paired = bool(paired)
        self.device = device
        self.dtype = dtype

        self.layer_by_modality: Dict[str, Optional[str]] = _as_modality_map(layer, adata_dict, kind="layer")
        self.xkey_by_modality: Dict[str, str] = _as_modality_map(X_key, adata_dict, kind="xkey")

        self.mod_cfg_by_name: Dict[str, ModalityConfig] = {}
        if modality_cfgs is not None:
            self.mod_cfg_by_name = {m.name: m for m in modality_cfgs}

        first = next(iter(adata_dict.values()))
        self._n_cells: int = int(first.n_obs)
        self._obs_names = first.obs_names

        if self.paired:
            for nm, adata_obj in self.adata_dict.items():
                if int(adata_obj.n_obs) != self._n_cells:
                    raise ValueError(
                        f"Paired dataset requires matching n_obs across modalities. "
                        f"First={self._n_cells}, {nm}={adata_obj.n_obs}"
                    )
                if not np.array_equal(adata_obj.obs_names.values, self._obs_names.values):
                    raise ValueError(
                        "Paired dataset requires identical obs_names order; %r differs. "
                        "Tip: use dataset_from_anndata_dict(..., align_obs=True)." % nm
                    )

        # Labels (optional): either a single vector or a dict of vectors
        self.labels: Optional[Union[torch.Tensor, Dict[str, torch.Tensor]]] = None
        if labels is not None:
            if isinstance(labels, Mapping):
                yd: Dict[str, torch.Tensor] = {}
                for hk, hv in labels.items():
                    t = hv if torch.is_tensor(hv) else torch.as_tensor(hv)
                    if t.ndim != 1:
                        t = t.reshape(-1)
                    if int(t.shape[0]) != self._n_cells:
                        raise ValueError(
                            f"labels[{hk!r}] length ({int(t.shape[0])}) must equal n_cells ({self._n_cells})"
                        )
                    t = t.long()
                    if self.device is not None:
                        t = t.to(self.device)
                    yd[str(hk)] = t
                self.labels = yd
            else:
                y = labels if torch.is_tensor(labels) else torch.as_tensor(labels)
                if y.ndim != 1:
                    y = y.reshape(-1)
                if int(y.shape[0]) != self._n_cells:
                    raise ValueError(f"labels length ({int(y.shape[0])}) must equal n_cells ({self._n_cells})")
                y = y.long()
                if self.device is not None:
                    y = y.to(self.device)
                self.labels = y

        # Reconstruction targets (optional): for (beta_)binomial etc.
        # spec is per-modality:
        #   {"successes_layer": "...", "total_count_layer": "..."}
        self.recon_targets_spec: Optional[Dict[str, Dict[str, str]]] = None
        if recon_targets_spec is not None:
            rts: Dict[str, Dict[str, str]] = {}
            for mod, spec in recon_targets_spec.items():
                if mod not in self.adata_dict:
                    raise KeyError(
                        f"recon_targets_spec contains mod={mod!r} not in adata_dict keys {list(self.adata_dict.keys())}"
                    )
                if not isinstance(spec, Mapping):
                    raise TypeError(f"recon_targets_spec[{mod!r}] must be a mapping, got {type(spec)}")

                s_layer = spec.get("successes_layer", None)
                n_layer = spec.get("total_count_layer", None)
                if not s_layer or not n_layer:
                    raise ValueError(
                        f"recon_targets_spec[{mod!r}] must define both "
                        f"'successes_layer' and 'total_count_layer'. Got keys={list(spec.keys())}"
                    )

                a = self.adata_dict[mod]
                if s_layer not in a.layers:
                    raise KeyError(
                        f"recon_targets_spec[{mod!r}] successes_layer={s_layer!r} not in adata.layers keys={list(a.layers.keys())}"
                    )
                if n_layer not in a.layers:
                    raise KeyError(
                        f"recon_targets_spec[{mod!r}] total_count_layer={n_layer!r} not in adata.layers keys={list(a.layers.keys())}"
                    )

                rts[str(mod)] = {"successes_layer": str(s_layer), "total_count_layer": str(n_layer)}

            self.recon_targets_spec = rts

    @property
    def n_cells(self) -> int:
        return self._n_cells

    @property
    def obs_names(self):
        return self._obs_names

    def __len__(self) -> int:
        return self._n_cells

    def _get_X_row(self, adata_obj: AnnData, idx: int, layer: Optional[str], X_key: str) -> np.ndarray:
        X = _get_matrix(adata_obj, layer=layer, X_key=X_key)
        row = X[idx]
        if sp.issparse(row):
            row = row.toarray()
        return np.asarray(row).reshape(-1).astype(np.float32, copy=False)

    def _get_layer_row(self, adata_obj: AnnData, idx: int, layer: str) -> np.ndarray:
        if layer not in adata_obj.layers:
            raise KeyError(f"layer={layer!r} not found in adata.layers. Keys={list(adata_obj.layers.keys())}")
        row = adata_obj.layers[layer][idx]
        if sp.issparse(row):
            row = row.toarray()
        return np.asarray(row).reshape(-1).astype(np.float32, copy=False)

    def _get_obs_label_row(self, adata_obj: AnnData, idx: int, obs_key: str) -> np.ndarray:
        if obs_key not in adata_obj.obs:
            raise KeyError(f"obs_key={obs_key!r} not found in adata.obs columns.")

        col = adata_obj.obs[obs_key]

        if pd.api.types.is_categorical_dtype(col):
            v = int(col.cat.codes.iloc[idx])
            return np.asarray([v], dtype=np.float32)

        v = col.iloc[idx]
        if isinstance(v, (np.integer, int)):
            return np.asarray([int(v)], dtype=np.float32)
        if isinstance(v, (np.floating, float)):
            return np.asarray([float(v)], dtype=np.float32)

        raise TypeError(
            f"adata.obs[{obs_key!r}] must be numeric integer codes (or pandas Categorical). "
            f"Got type {type(v)} at row {idx}. Encode categories to int codes first."
        )

    def __getitem__(self, idx: int):
        x_dict: Dict[str, torch.Tensor] = {}

        for name, adata_obj in self.adata_dict.items():
            mcfg = self.mod_cfg_by_name.get(name, None)

            if (
                mcfg is not None
                and _is_categorical_likelihood(mcfg.likelihood)
                and (mcfg.input_kind or "matrix") == "obs"
            ):
                if not mcfg.obs_key:
                    raise ValueError(f"Modality {name!r}: input_kind='obs' requires obs_key.")
                row_np = self._get_obs_label_row(adata_obj, idx, obs_key=mcfg.obs_key)
            else:
                layer = self.layer_by_modality.get(name, None)
                xkey = self.xkey_by_modality.get(name, "X")
                row_np = self._get_X_row(adata_obj, idx, layer=layer, X_key=xkey)

            x = torch.from_numpy(row_np).to(dtype=self.dtype)
            if self.device is not None:
                x = x.to(self.device)
            x_dict[name] = x

        recon_targets = None
        if self.recon_targets_spec is not None:
            rt: Dict[str, Dict[str, torch.Tensor]] = {}
            for mod, spec in self.recon_targets_spec.items():
                a = self.adata_dict[mod]
                k_np = self._get_layer_row(a, idx, layer=spec["successes_layer"])
                n_np = self._get_layer_row(a, idx, layer=spec["total_count_layer"])

                k = torch.from_numpy(k_np).to(dtype=self.dtype)
                n = torch.from_numpy(n_np).to(dtype=self.dtype)
                if self.device is not None:
                    k = k.to(self.device)
                    n = n.to(self.device)

                rt[mod] = {"successes": k, "total_count": n}
            recon_targets = rt

        # Return patterns:
        #   - x_dict
        #   - (x_dict, y)
        #   - (x_dict, recon_targets)
        #   - (x_dict, y, recon_targets)
        if self.labels is None and recon_targets is None:
            return x_dict

        if self.labels is None and recon_targets is not None:
            return x_dict, recon_targets

        # labels exist
        if isinstance(self.labels, dict):
            y_out: Dict[str, torch.Tensor] = {k: v[idx] for k, v in self.labels.items()}
            if recon_targets is None:
                return x_dict, y_out
            return x_dict, y_out, recon_targets

        # single-label
        if recon_targets is None:
            return x_dict, self.labels[idx]
        return x_dict, self.labels[idx], recon_targets


# -----------------------------------------------------------------------------
# Factory helpers
# -----------------------------------------------------------------------------
def dataset_from_anndata_dict(
    adata_dict: Dict[str, AnnData],
    layer: LayerSpec = None,
    X_key: XKeySpec = "X",
    paired: bool = True,
    align_obs: bool = True,
    labels: Optional[LabelSpec] = None,
    device: Optional[torch.device] = None,
    dtype: torch.dtype = torch.float32,
    copy_aligned: bool = True,
    modality_cfgs: Optional[List[ModalityConfig]] = None,
    recon_targets_spec: Optional[ReconTargetSpec] = None,
) -> Tuple[MultiModalDataset, Dict[str, AnnData]]:
    if align_obs and paired:
        adata_dict = align_paired_obs_names(adata_dict, how="intersection", copy=copy_aligned)

    ds = MultiModalDataset(
        adata_dict=adata_dict,
        layer=layer,
        X_key=X_key,
        paired=paired,
        device=device,
        labels=labels,
        dtype=dtype,
        modality_cfgs=modality_cfgs,
        recon_targets_spec=recon_targets_spec,
    )
    return ds, adata_dict


def load_anndata_dict_from_config(
    modality_cfgs: List[Dict[str, Any]],
    data_root: Optional[str] = None,
) -> Dict[str, AnnData]:
    out: Dict[str, AnnData] = {}
    for m in modality_cfgs:
        if "name" not in m or "h5ad_path" not in m:
            raise KeyError("Each modality config must contain keys: 'name' and 'h5ad_path'.")

        name = m["name"]
        path = m["h5ad_path"]

        if data_root is not None and not os.path.isabs(path):
            path = os.path.join(data_root, path)

        out[name] = ad.read_h5ad(path)

    if not out:
        raise ValueError("No modalities loaded (empty modality_cfgs?)")

    return out


# -----------------------------------------------------------------------------
# Collate functions
# -----------------------------------------------------------------------------
def collate_multimodal_xy(batch):
    """
    Back-compat collate:
      - works for [x_dict, ...] or [(x_dict, y), ...]
      - stacks per-modality tensors into (B, D)
      - supports y as:
          * scalar tensor/int
          * dict[str -> scalar tensor/int]
    """
    if isinstance(batch[0], (tuple, list)) and len(batch[0]) == 2:
        xs, ys = zip(*batch)

        y0 = ys[0]
        if isinstance(y0, Mapping):
            y_out: Dict[str, torch.Tensor] = {}
            keys = list(y0.keys())
            for k in keys:
                y_out[str(k)] = torch.stack([torch.as_tensor(yy[k], dtype=torch.long) for yy in ys], dim=0)
            y = y_out
        else:
            y = torch.stack([torch.as_tensor(yy, dtype=torch.long) for yy in ys], dim=0)
    else:
        xs, y = batch, None

    keys = xs[0].keys()
    x = {k: torch.stack([d[k] for d in xs], dim=0) for k in keys}
    return x if y is None else (x, y)


def collate_multimodal_xy_recon(batch):
    """
    Collate that supports recon_targets.

    Accepts batch items in any of these shapes:
      - x_dict
      - (x_dict, y)
      - (x_dict, recon_targets)
      - (x_dict, y, recon_targets)

    Returns:
      - x
      - (x, y)
      - (x, recon_targets)
      - (x, y, recon_targets)

    recon_targets is:
      Dict[mod -> Dict["successes"->(B,D), "total_count"->(B,D)]]
    """
    x_items: List[Dict[str, torch.Tensor]] = []
    y_items: List[Optional[Union[torch.Tensor, Mapping[str, Any]]]] = []
    rt_items: List[Optional[Mapping[str, Mapping[str, Any]]]] = []

    def _looks_like_recon_targets(obj: Any) -> bool:
        if not isinstance(obj, Mapping) or len(obj) == 0:
            return False
        # expected: {mod: {"successes": ..., "total_count": ...}}
        for v in obj.values():
            if not isinstance(v, Mapping):
                return False
            if ("successes" not in v) or ("total_count" not in v):
                return False
        return True

    for item in batch:
        if isinstance(item, (tuple, list)):
            if len(item) == 2:
                a, b = item
                if _looks_like_recon_targets(b):
                    x_items.append(a)
                    y_items.append(None)
                    rt_items.append(b)
                else:
                    x_items.append(a)
                    y_items.append(b)
                    rt_items.append(None)
            elif len(item) == 3:
                a, b, c = item
                x_items.append(a)
                y_items.append(b)
                rt_items.append(c)
            else:
                raise ValueError(f"Unsupported batch item tuple length={len(item)}")
        else:
            x_items.append(item)
            y_items.append(None)
            rt_items.append(None)

    # Stack X
    keys = x_items[0].keys()
    x = {k: torch.stack([d[k] for d in x_items], dim=0) for k in keys}

    # Stack y if present
    y_present = any(v is not None for v in y_items)
    y_out = None
    if y_present:
        ys = [v for v in y_items if v is not None]
        y0 = ys[0]
        if isinstance(y0, Mapping):
            y_out = {
                str(k): torch.stack([torch.as_tensor(yy[k], dtype=torch.long) for yy in ys], dim=0)
                for k in y0.keys()
            }
        else:
            y_out = torch.stack([torch.as_tensor(yy, dtype=torch.long) for yy in ys], dim=0)

    # Stack recon_targets if present
    rt_present = any(v is not None for v in rt_items)
    rt_out = None
    if rt_present:
        rts = [v for v in rt_items if v is not None]
        rt_out = {}
        for mod in rts[0].keys():
            rt_out[mod] = {
                "successes": torch.stack(
                    [torch.as_tensor(rr[mod]["successes"], dtype=torch.float32) for rr in rts], dim=0
                ),
                "total_count": torch.stack(
                    [torch.as_tensor(rr[mod]["total_count"], dtype=torch.float32) for rr in rts], dim=0
                ),
            }

    # Return in a stable order
    if (y_out is None) and (rt_out is None):
        return x
    if (y_out is None) and (rt_out is not None):
        return x, rt_out
    if (y_out is not None) and (rt_out is None):
        return x, y_out
    return x, y_out, rt_out
```
