## Supplemental Code for "Unifying multimodal single-cell data with a mixture-of-experts *β*-variational autoencoder framework": diagnostics.py

```
# univi/diagnostics.py
from __future__ import annotations

from typing import Any, Dict, Optional, List
import os
import platform
import importlib

import numpy as np
import pandas as pd
from anndata import AnnData

from .data import _get_matrix
from .utils.io import load_config


def _safe_version(pkg: str) -> str:
    try:
        mod = importlib.import_module(pkg)
        return getattr(mod, "__version__", "unknown")
    except Exception:
        return "not_installed"


def collect_environment_info() -> Dict[str, Any]:
    return {
        "python": platform.python_version(),
        "platform": platform.platform(),
        "numpy": _safe_version("numpy"),
        "scipy": _safe_version("scipy"),
        "pandas": _safe_version("pandas"),
        "anndata": _safe_version("anndata"),
        "scanpy": _safe_version("scanpy"),
        "torch": _safe_version("torch"),
        "sklearn": _safe_version("sklearn"),
        "h5py": _safe_version("h5py"),
        "matplotlib": _safe_version("matplotlib"),
        "seaborn": _safe_version("seaborn"),
    }


def dataset_stats_table(
    adata_dict: Dict[str, AnnData],
    *,
    layer_by: Optional[Dict[str, Optional[str]]] = None,
    xkey_by: Optional[Dict[str, str]] = None,
) -> pd.DataFrame:
    rows = []
    for nm, adata in adata_dict.items():
        layer = None if layer_by is None else layer_by.get(nm, None)
        xkey = "X" if xkey_by is None else xkey_by.get(nm, "X")
        X = _get_matrix(adata, layer=layer, X_key=xkey)
        rows.append(
            {
                "modality": nm,
                "n_cells": int(adata.n_obs),
                "n_features": int(X.shape[1]),
                "X_key": xkey,
                "layer": layer if layer is not None else "",
            }
        )
    return pd.DataFrame(rows)


def model_hparams_table(cfg: Dict[str, Any]) -> pd.DataFrame:
    model = cfg.get("model", {})
    training = cfg.get("training", {})
    rows = []
    # flatten a curated set of keys
    keys = [
        "loss_mode",
        "v1_recon",
        "v1_recon_mix",
        "normalize_v1_terms",
        "latent_dim",
        "beta",
        "gamma",
        "hidden_dims_default",
        "dropout",
        "encoder_dropout",
        "decoder_dropout",
        "batchnorm",
        "encoder_batchnorm",
        "decoder_batchnorm",
        "kl_anneal_start",
        "kl_anneal_end",
        "align_anneal_start",
        "align_anneal_end",
    ]
    for k in keys:
        if k in model:
            rows.append({"section": "model", "key": k, "value": str(model[k])})
    tkeys = ["n_epochs", "batch_size", "lr", "weight_decay", "device", "seed", "num_workers", "early_stopping", "patience", "min_delta"]
    for k in tkeys:
        if k in training:
            rows.append({"section": "training", "key": k, "value": str(training[k])})

    # per-modality entries
    for m in cfg.get("data", {}).get("modalities", []):
        name = m.get("name", "modality")
        for k in ["likelihood", "layer", "X_key", "hidden_dims", "encoder_hidden", "decoder_hidden"]:
            if k in m:
                rows.append({"section": f"data.{name}", "key": k, "value": str(m[k])})

    return pd.DataFrame(rows)


def export_supplemental_table_s1(
    config_path: str,
    adata_dict: Dict[str, AnnData],
    *,
    out_xlsx: str,
    layer_by: Optional[Dict[str, Optional[str]]] = None,
    xkey_by: Optional[Dict[str, str]] = None,
    extra_metrics: Optional[Dict[str, Any]] = None,
):
    """Write Supplemental_Table_S1.xlsx: environment + hparams + dataset stats (+ optional metrics)."""
    cfg = load_config(config_path)
    env = collect_environment_info()
    df_env = pd.DataFrame([env])
    df_hp = model_hparams_table(cfg)
    df_ds = dataset_stats_table(adata_dict, layer_by=layer_by, xkey_by=xkey_by)

    os.makedirs(os.path.dirname(out_xlsx) or ".", exist_ok=True)
    with pd.ExcelWriter(out_xlsx, engine="openpyxl") as w:
        df_env.to_excel(w, index=False, sheet_name="environment")
        df_hp.to_excel(w, index=False, sheet_name="hyperparameters")
        df_ds.to_excel(w, index=False, sheet_name="datasets")
        if extra_metrics:
            pd.DataFrame([extra_metrics]).to_excel(w, index=False, sheet_name="metrics")
```
