## Supplemental Code for "Unifying multimodal single-cell data with a mixture-of-experts *β*-variational autoencoder framework": evaluation.py

```
# univi/evaluation.py
from __future__ import annotations

from typing import Any, Dict, Optional, Tuple, Union, Sequence, List, Mapping

import numpy as np
import scipy.sparse as sp
import torch

from sklearn.neighbors import NearestNeighbors
from sklearn.metrics import (
    accuracy_score,
    confusion_matrix,
    f1_score,
    roc_auc_score,
)


# =============================================================================
# Helpers
# =============================================================================
def to_dense(X):
    return X.toarray() if sp.issparse(X) else np.asarray(X)


def _mean_sem(x: np.ndarray) -> Tuple[float, float]:
    x = np.asarray(x, dtype=float)
    if x.size == 0:
        return 0.0, 0.0
    if x.size == 1:
        return float(x.mean()), 0.0
    return float(x.mean()), float(x.std(ddof=1) / np.sqrt(x.size))


def _json_safe(obj: Any) -> Any:
    """Convert numpy/torch-ish objects into JSON-safe python types recursively."""
    if torch.is_tensor(obj):
        obj = obj.detach().cpu().numpy()

    if isinstance(obj, (np.floating, np.integer)):
        return obj.item()
    if isinstance(obj, np.ndarray):
        return obj.tolist()
    if isinstance(obj, dict):
        return {str(k): _json_safe(v) for k, v in obj.items()}
    if isinstance(obj, (list, tuple)):
        return [_json_safe(v) for v in obj]
    return obj


def _is_tensor(x: Any) -> bool:
    return torch.is_tensor(x)


def _sigmoid(x: torch.Tensor) -> torch.Tensor:
    return torch.sigmoid(x)


def _softmax(x: torch.Tensor, dim: int = -1) -> torch.Tensor:
    return torch.softmax(x, dim=dim)


def _extract_tensor_from_decoder_output(
    out: Any,
    *,
    prefer: str = "auto",
    apply_link: Optional[str] = None,
) -> torch.Tensor:
    """
    Turn a decoder output into a tensor suitable for returning/storing.

    out:
      - torch.Tensor OR a dict produced by a decoder.

    prefer:
      - "auto" : pick sensible default by keys
      - "mean" : mean-like for Gaussian
      - "mu"   : mu-like for NB/ZINB
      - "rate" : rate for Poisson
      - "logits": logits for Bernoulli/Categorical etc.
      - "probs": probs if available, otherwise compute if possible

    apply_link:
      - None
      - "sigmoid": logits -> probs
      - "softmax": logits -> probs
      - "softplus": log_rate -> rate
    """
    if _is_tensor(out):
        x = out
    elif isinstance(out, dict):
        p = str(prefer).lower().strip()

        if p in out and _is_tensor(out[p]):
            x = out[p]
        else:
            if p == "probs":
                key_order = ["probs", "logits", "mean", "mu", "rate", "log_rate"]
            elif p == "logits":
                key_order = ["logits", "probs", "mean", "mu", "rate", "log_rate"]
            elif p == "mean":
                key_order = ["mean", "mu", "rate", "probs", "logits", "log_rate"]
            elif p == "mu":
                key_order = ["mu", "mean", "rate", "probs", "logits", "log_rate"]
            elif p == "rate":
                key_order = ["rate", "log_rate", "mu", "mean", "probs", "logits"]
            else:
                key_order = ["mean", "mu", "rate", "probs", "logits", "log_rate", "logvar"]

            x = None
            for k in key_order:
                if k in out and _is_tensor(out[k]):
                    x = out[k]
                    break

            if x is None:
                if len(out) == 1:
                    x = next(iter(out.values()))
                    if not _is_tensor(x):
                        raise TypeError(f"Decoder output dict has non-tensor leaf: {type(x)}")
                else:
                    raise TypeError(f"Could not extract tensor from decoder output keys={list(out.keys())}")
    else:
        raise TypeError(f"Unsupported decoder output type: {type(out)}")

    if apply_link is not None:
        link = str(apply_link).lower().strip()
        if link == "sigmoid":
            x = _sigmoid(x)
        elif link == "softmax":
            x = _softmax(x, dim=-1)
        elif link == "softplus":
            x = torch.nn.functional.softplus(x)
        else:
            raise ValueError(f"Unknown apply_link={apply_link!r}")

    return x


def _to_numpy(x: torch.Tensor) -> np.ndarray:
    return x.detach().cpu().numpy()


def _infer_model_modality_order(model, fallback: Sequence[str]) -> List[str]:
    """
    Best-effort inference of the modality order used by the model (for gates).
    Falls back to the provided sequence.
    """
    # common: model.cfg.modalities is list[ModalityConfig(name=...)]
    cfg = getattr(model, "cfg", None)
    if cfg is not None and hasattr(cfg, "modalities"):
        mods = getattr(cfg, "modalities")
        try:
            names = [getattr(m, "name") for m in mods]
            if all(isinstance(x, str) and x for x in names):
                return list(names)
        except Exception:
            pass
    return list(map(str, list(fallback)))


# =============================================================================
# Manuscript helpers (LSC17 score)
# =============================================================================
LSC17_GENES = [
    "DNMT3B", "ZBTB46", "NYNRIN", "ARHGAP22", "LAPTM4B", "MMRN1", "DPYSL3", "FAM30A",
    "CDK6", "CPXM1", "SOCS2", "SMIM24", "EMP1", "BEX3", "CD34", "AKR1C3", "ADGRG1",
]


def add_lsc17_scores(
    adata,
    layer: Optional[str] = None,
    gene_key: Optional[str] = None,
    prefix: str = "LSC17",
) -> Any:
    var_names = np.asarray(adata.var_names if gene_key is None else adata.var[gene_key])
    gene_to_idx = {g: i for i, g in enumerate(var_names)}
    present = [g for g in LSC17_GENES if g in gene_to_idx]
    idx = [gene_to_idx[g] for g in present]

    X = adata.X if layer is None else (adata.layers[layer] if layer in adata.layers else adata.X)
    X = X[:, idx]
    X = X.toarray() if sp.issparse(X) else np.asarray(X, dtype=np.float32)

    score = X.mean(axis=1)
    mu = float(np.mean(score))
    sd = float(np.std(score, ddof=0))
    z = (score - mu) / (sd + 1e-8)

    adata.obs[f"{prefix}_score"] = np.asarray(score).ravel()
    adata.obs[f"{prefix}_z"] = np.asarray(z).ravel()
    adata.obs[f"{prefix}_n_genes"] = int(len(present))
    return adata


# =============================================================================
# FOSCTTM + Recall@k
# =============================================================================
def compute_foscttm(
    Z1: np.ndarray,
    Z2: np.ndarray,
    metric: str = "euclidean",
    block_size: int = 512,
    return_sem: bool = False,
    return_per_cell: bool = False,
) -> Union[float, Tuple[float, float], Tuple[float, np.ndarray], Tuple[float, float, np.ndarray]]:
    Z1 = np.asarray(Z1, dtype=np.float32)
    Z2 = np.asarray(Z2, dtype=np.float32)

    if Z1.shape != Z2.shape:
        raise ValueError(f"Z1/Z2 must have same shape for FOSCTTM. Got {Z1.shape} vs {Z2.shape}")

    n = int(Z1.shape[0])
    if n <= 1:
        fos = np.zeros(n, dtype=np.float32)
        m, s = _mean_sem(fos.astype(float))
        if return_sem and return_per_cell:
            return float(m), float(s), fos
        if return_sem:
            return float(m), float(s)
        if return_per_cell:
            return float(m), fos
        return float(m)

    metric = str(metric).lower().strip()
    if metric not in {"euclidean", "cosine"}:
        raise ValueError("compute_foscttm supports metric in {'euclidean','cosine'}.")

    fos = np.empty(n, dtype=np.float32)

    if metric == "euclidean":
        Z2_T = Z2.T
        n2 = np.sum(Z2 * Z2, axis=1)
        for i0 in range(0, n, int(block_size)):
            i1 = min(i0 + int(block_size), n)
            A = Z1[i0:i1]
            n1 = np.sum(A * A, axis=1)[:, None]
            d2 = n1 + n2[None, :] - 2.0 * (A @ Z2_T)
            true = d2[np.arange(i1 - i0), np.arange(i0, i1)]
            fos[i0:i1] = (d2 < true[:, None]).sum(axis=1) / (n - 1)
    else:
        Z2_T = Z2.T
        n2 = np.linalg.norm(Z2, axis=1) + 1e-8
        for i0 in range(0, n, int(block_size)):
            i1 = min(i0 + int(block_size), n)
            A = Z1[i0:i1]
            n1 = np.linalg.norm(A, axis=1) + 1e-8
            sim = (A @ Z2_T) / (n1[:, None] * n2[None, :])
            d = 1.0 - sim
            true = d[np.arange(i1 - i0), np.arange(i0, i1)]
            fos[i0:i1] = (d < true[:, None]).sum(axis=1) / (n - 1)

    m, s = _mean_sem(fos.astype(float))
    if return_sem and return_per_cell:
        return float(m), float(s), fos
    if return_sem:
        return float(m), float(s)
    if return_per_cell:
        return float(m), fos
    return float(m)


def compute_match_recall_at_k(
    Z1: np.ndarray,
    Z2: np.ndarray,
    k: int = 10,
    metric: str = "euclidean",
    block_size: int = 512,
    return_sem: bool = False,
    return_per_cell: bool = False,
) -> Union[float, Tuple[float, float], Tuple[float, np.ndarray], Tuple[float, float, np.ndarray]]:
    Z1 = np.asarray(Z1, dtype=np.float32)
    Z2 = np.asarray(Z2, dtype=np.float32)

    if Z1.shape != Z2.shape:
        raise ValueError(f"Z1/Z2 must have same shape. Got {Z1.shape} vs {Z2.shape}")

    n = int(Z1.shape[0])
    if n == 0:
        raise ValueError("Empty inputs.")
    if n == 1:
        hits = np.array([1.0], dtype=np.float32)
        if return_sem and return_per_cell:
            return 1.0, 0.0, hits
        if return_sem:
            return 1.0, 0.0
        if return_per_cell:
            return 1.0, hits
        return 1.0

    k = int(max(1, min(int(k), n)))
    metric = str(metric).lower().strip()
    if metric not in {"euclidean", "cosine"}:
        raise ValueError("compute_match_recall_at_k supports metric in {'euclidean','cosine'}.")

    hits = np.empty(n, dtype=np.float32)

    if metric == "euclidean":
        Z2_T = Z2.T
        n2 = np.sum(Z2 * Z2, axis=1)
        for i0 in range(0, n, int(block_size)):
            i1 = min(i0 + int(block_size), n)
            A = Z1[i0:i1]
            n1 = np.sum(A * A, axis=1)[:, None]
            d2 = n1 + n2[None, :] - 2.0 * (A @ Z2_T)
            topk = np.argpartition(d2, kth=k - 1, axis=1)[:, :k]
            for r in range(i1 - i0):
                hits[i0 + r] = 1.0 if (i0 + r) in topk[r] else 0.0
    else:
        Z2_T = Z2.T
        n2 = np.linalg.norm(Z2, axis=1) + 1e-8
        for i0 in range(0, n, int(block_size)):
            i1 = min(i0 + int(block_size), n)
            A = Z1[i0:i1]
            n1 = np.linalg.norm(A, axis=1) + 1e-8
            sim = (A @ Z2_T) / (n1[:, None] * n2[None, :])
            d = 1.0 - sim
            topk = np.argpartition(d, kth=k - 1, axis=1)[:, :k]
            for r in range(i1 - i0):
                hits[i0 + r] = 1.0 if (i0 + r) in topk[r] else 0.0

    m, s = _mean_sem(hits.astype(float))
    if return_sem and return_per_cell:
        return float(m), float(s), hits
    if return_sem:
        return float(m), float(s)
    if return_per_cell:
        return float(m), hits
    return float(m)


# =============================================================================
# Modality mixing / entropy
# =============================================================================
def compute_modality_mixing(
    Z: np.ndarray,
    modality_labels: np.ndarray,
    k: int = 20,
    metric: str = "euclidean",
    return_sem: bool = False,
    return_per_cell: bool = False,
) -> Union[float, Tuple[float, float], Tuple[float, np.ndarray], Tuple[float, float, np.ndarray]]:
    Z = np.asarray(Z, dtype=np.float32)
    modality_labels = np.asarray(modality_labels)
    if Z.shape[0] != modality_labels.shape[0]:
        raise ValueError("Z and modality_labels must align on n_cells.")

    n = int(Z.shape[0])
    if n <= 1:
        frac_other = np.zeros(n, dtype=np.float32)
        m, s = _mean_sem(frac_other.astype(float))
        if return_sem and return_per_cell:
            return float(m), float(s), frac_other
        if return_sem:
            return float(m), float(s)
        if return_per_cell:
            return float(m), frac_other
        return float(m)

    metric = str(metric).lower().strip()
    k_eff = int(min(max(int(k), 1), n - 1))

    nn = NearestNeighbors(n_neighbors=k_eff + 1, metric=metric)
    nn.fit(Z)
    neigh_idx = nn.kneighbors(Z, return_distance=False)[:, 1:]

    neigh_mods = modality_labels[neigh_idx]
    frac_other = (neigh_mods != modality_labels[:, None]).mean(axis=1).astype(np.float32)

    m, s = _mean_sem(frac_other.astype(float))
    if return_sem and return_per_cell:
        return float(m), float(s), frac_other
    if return_sem:
        return float(m), float(s)
    if return_per_cell:
        return float(m), frac_other
    return float(m)


def compute_modality_entropy(
    Z: np.ndarray,
    modality_labels: np.ndarray,
    k: int = 30,
    metric: str = "euclidean",
    return_sem: bool = False,
    return_per_cell: bool = False,
) -> Union[float, Tuple[float, float], Tuple[float, np.ndarray], Tuple[float, float, np.ndarray]]:
    Z = np.asarray(Z, dtype=np.float32)
    modality_labels = np.asarray(modality_labels)

    n = int(Z.shape[0])
    if n <= 1:
        ent = np.zeros(n, dtype=np.float32)
        m, s = _mean_sem(ent.astype(float))
        if return_sem and return_per_cell:
            return float(m), float(s), ent
        if return_sem:
            return float(m), float(s)
        if return_per_cell:
            return float(m), ent
        return float(m)

    uniq_mods = np.unique(modality_labels)
    mcount = int(uniq_mods.size)
    if mcount <= 1:
        ent = np.zeros(n, dtype=np.float32)
        m, s = _mean_sem(ent.astype(float))
        if return_sem and return_per_cell:
            return float(m), float(s), ent
        if return_sem:
            return float(m), float(s)
        if return_per_cell:
            return float(m), ent
        return float(m)

    k_eff = int(min(max(int(k), 1), n - 1))
    nn = NearestNeighbors(n_neighbors=k_eff + 1, metric=str(metric))
    nn.fit(Z)
    neigh_idx = nn.kneighbors(Z, return_distance=False)[:, 1:]

    mod_to_code = {m: i for i, m in enumerate(uniq_mods)}
    codes = np.vectorize(mod_to_code.get)(modality_labels)

    ent = np.empty(n, dtype=np.float32)
    denom = np.log(float(mcount) + 1e-12)

    for i in range(n):
        neigh_codes = codes[neigh_idx[i]]
        p = np.bincount(neigh_codes, minlength=mcount).astype(np.float32)
        p = p / (p.sum() + 1e-8)
        h = -(p * np.log(p + 1e-12)).sum()
        ent[i] = float(h / (denom + 1e-12))

    m, s = _mean_sem(ent.astype(float))
    if return_sem and return_per_cell:
        return float(m), float(s), ent
    if return_sem:
        return float(m), float(s)
    if return_per_cell:
        return float(m), ent
    return float(m)


# =============================================================================
# Label transfer (kNN)
# =============================================================================
def label_transfer_knn(
    Z_source: np.ndarray,
    labels_source: np.ndarray,
    Z_target: np.ndarray,
    labels_target: Optional[np.ndarray] = None,
    k: int = 15,
    metric: str = "euclidean",
    return_label_order: bool = False,
    return_f1: bool = False,
):
    Z_source = np.asarray(Z_source, dtype=np.float32)
    Z_target = np.asarray(Z_target, dtype=np.float32)
    labels_source = np.asarray(labels_source)
    if labels_target is not None:
        labels_target = np.asarray(labels_target)

    n_source = int(Z_source.shape[0])
    if n_source == 0:
        raise ValueError("Z_source is empty.")

    k_eff = int(min(max(int(k), 1), n_source))
    nn = NearestNeighbors(n_neighbors=k_eff, metric=metric)
    nn.fit(Z_source)
    neigh_idx = nn.kneighbors(Z_target, return_distance=False)

    uniq_src, src_codes = np.unique(labels_source, return_inverse=True)

    pred_codes = np.empty(Z_target.shape[0], dtype=np.int64)
    for i in range(Z_target.shape[0]):
        votes = src_codes[neigh_idx[i]]
        bc = np.bincount(votes, minlength=len(uniq_src))
        pred_codes[i] = int(bc.argmax())

    pred_labels = uniq_src[pred_codes]

    if labels_target is None:
        return pred_labels, None, np.array([])

    label_order = np.unique(np.concatenate([labels_target, pred_labels]))
    acc = float(accuracy_score(labels_target, pred_labels))
    cm = confusion_matrix(labels_target, pred_labels, labels=label_order)

    extras = []
    if return_label_order:
        extras.append(label_order)
    if return_f1:
        extras.append({
            "macro_f1": float(f1_score(labels_target, pred_labels, average="macro")),
            "weighted_f1": float(f1_score(labels_target, pred_labels, average="weighted")),
        })

    if not extras:
        return pred_labels, acc, cm
    return (pred_labels, acc, cm, *extras)


def summarize_bidirectional_transfer(
    Z_a: np.ndarray,
    y_a: np.ndarray,
    Z_b: np.ndarray,
    y_b: np.ndarray,
    k: int = 15,
    metric: str = "euclidean",
) -> Dict[str, Any]:
    pred_ab, acc_ab, cm_ab, order_ab, f1_ab = label_transfer_knn(
        Z_source=Z_a, labels_source=y_a,
        Z_target=Z_b, labels_target=y_b,
        k=k, metric=metric,
        return_label_order=True, return_f1=True,
    )
    pred_ba, acc_ba, cm_ba, order_ba, f1_ba = label_transfer_knn(
        Z_source=Z_b, labels_source=y_b,
        Z_target=Z_a, labels_target=y_a,
        k=k, metric=metric,
        return_label_order=True, return_f1=True,
    )

    return {
        "ab": {"acc": acc_ab, "f1": f1_ab, "cm": cm_ab, "order": order_ab, "pred": pred_ab},
        "ba": {"acc": acc_ba, "f1": f1_ba, "cm": cm_ba, "order": order_ba, "pred": pred_ba},
        "worst_direction_macro_f1": float(min(f1_ab["macro_f1"], f1_ba["macro_f1"])),
        "worst_direction_weighted_f1": float(min(f1_ab["weighted_f1"], f1_ba["weighted_f1"])),
        "k": int(k),
        "metric": str(metric),
    }


# =============================================================================
# Reconstruction metrics (continuous + binary)
# =============================================================================
def mse_per_feature(x_true: np.ndarray, x_pred: np.ndarray) -> np.ndarray:
    x_true = np.asarray(x_true)
    x_pred = np.asarray(x_pred)
    if x_true.shape != x_pred.shape:
        raise ValueError("x_true and x_pred must have same shape.")
    return np.mean((x_true - x_pred) ** 2, axis=0)


def pearson_corr_per_feature(x_true: np.ndarray, x_pred: np.ndarray) -> np.ndarray:
    x_true = np.asarray(x_true, dtype=np.float32)
    x_pred = np.asarray(x_pred, dtype=np.float32)
    if x_true.shape != x_pred.shape:
        raise ValueError("x_true and x_pred must have same shape.")

    x_true_c = x_true - x_true.mean(axis=0, keepdims=True)
    x_pred_c = x_pred - x_pred.mean(axis=0, keepdims=True)

    num = (x_true_c * x_pred_c).sum(axis=0)
    denom = np.sqrt((x_true_c ** 2).sum(axis=0) * (x_pred_c ** 2).sum(axis=0)) + 1e-8
    return num / denom


def auc_per_feature_binary(x_true01: np.ndarray, x_score: np.ndarray) -> np.ndarray:
    """
    Per-feature AUC for binary ground-truth.
    x_true01: 0/1
    x_score: probabilities or logits (both ok for AUC)
    """
    x_true01 = np.asarray(x_true01)
    x_score = np.asarray(x_score)
    if x_true01.shape != x_score.shape:
        raise ValueError("x_true01 and x_score must have same shape.")
    aucs = np.full(x_true01.shape[1], np.nan, dtype=np.float32)
    for j in range(x_true01.shape[1]):
        y = x_true01[:, j]
        if y.min() == y.max():
            continue
        aucs[j] = float(roc_auc_score(y, x_score[:, j]))
    return aucs


def reconstruction_metrics(
    x_true: np.ndarray,
    x_pred: np.ndarray,
    *,
    kind: str = "continuous",   # "continuous" | "binary"
) -> Dict[str, Any]:
    kind = str(kind).lower().strip()
    x_true = np.asarray(x_true)
    x_pred = np.asarray(x_pred)

    if kind == "binary":
        aucs = auc_per_feature_binary(x_true, x_pred)
        return {
            "auc_mean": float(np.nanmean(aucs)),
            "auc_median": float(np.nanmedian(aucs)),
            "auc_per_feature": aucs,
        }

    pf_mse = mse_per_feature(x_true, x_pred)
    pf_r = pearson_corr_per_feature(x_true, x_pred)
    return {
        "mse_mean": float(np.mean(pf_mse)),
        "mse_median": float(np.median(pf_mse)),
        "pearson_mean": float(np.mean(pf_r)),
        "pearson_median": float(np.median(pf_r)),
        "mse_per_feature": pf_mse,
        "pearson_per_feature": pf_r,
    }


# =============================================================================
# Encoding (single modality) + cross-modal prediction
# =============================================================================
def encode_adata(
    model,
    adata,
    modality: str,
    device: str = "cpu",
    layer: Optional[str] = None,
    X_key: str = "X",
    batch_size: int = 1024,
    latent: str = "moe_mean",
    random_state: int = 0,
) -> np.ndarray:
    """
    Encode a *single* modality into z.

    latent:
      - "moe_mean" / "moe_sample": fused MoE/PoE posterior using only provided experts
      - "modality_mean" / "modality_sample": that modality posterior only
    """
    from .data import _get_matrix

    latent = str(latent).lower().strip()
    valid = {"moe_mean", "moe_sample", "modality_mean", "modality_sample"}
    if latent not in valid:
        raise ValueError("latent must be one of %s; got %r" % (sorted(valid), latent))

    def _sample_gaussian(mu: torch.Tensor, logvar: torch.Tensor, gen: torch.Generator) -> torch.Tensor:
        eps = torch.randn(mu.shape, device=mu.device, generator=gen, dtype=mu.dtype)
        return mu + eps * torch.exp(0.5 * logvar)

    model.eval()
    X = _get_matrix(adata, layer=layer, X_key=X_key)
    X = X.toarray() if sp.issparse(X) else np.asarray(X)

    dev = torch.device(device)
    gen = torch.Generator(device=dev)
    gen.manual_seed(int(random_state))

    zs: List[np.ndarray] = []
    with torch.no_grad():
        for start in range(0, X.shape[0], int(batch_size)):
            end = min(start + int(batch_size), X.shape[0])
            xb = torch.as_tensor(np.asarray(X[start:end]), dtype=torch.float32, device=dev)

            mu_dict, logvar_dict = model.encode_modalities({modality: xb})

            if "modality" in latent:
                mu = mu_dict[modality]
                lv = logvar_dict[modality]
                z = mu if latent.endswith("_mean") else _sample_gaussian(mu, lv, gen)
            else:
                if hasattr(model, "mixture_of_experts"):
                    mu_z, logvar_z = model.mixture_of_experts(mu_dict, logvar_dict)
                elif hasattr(model, "fuse_posteriors"):
                    mu_z, logvar_z = model.fuse_posteriors(mu_dict, logvar_dict)
                else:
                    mu_z, logvar_z = mu_dict[modality], logvar_dict[modality]

                z = mu_z if latent.endswith("_mean") else _sample_gaussian(mu_z, logvar_z, gen)

            zs.append(_to_numpy(z))

    return np.vstack(zs) if zs else np.zeros((0, 0), dtype=np.float32)


def cross_modal_predict(
    model,
    adata_src,
    src_mod: str,
    tgt_mod: str,
    device: str = "cpu",
    layer: Optional[str] = None,
    X_key: str = "X",
    batch_size: int = 512,
    use_moe: bool = True,
    decoder_prefer: str = "auto",
    decoder_link: Optional[str] = None,
) -> np.ndarray:
    """
    Encode src_mod then decode tgt_mod.

    Robust to decoder outputs that are dicts (NB/ZINB/Poisson/Bernoulli/...).
    Returns a mean-like matrix for evaluation/plotting.
    """
    from .data import _get_matrix

    model.eval()
    X = _get_matrix(adata_src, layer=layer, X_key=X_key)
    X = X.toarray() if sp.issparse(X) else np.asarray(X)

    dev = torch.device(device)

    preds: List[np.ndarray] = []
    with torch.no_grad():
        for start in range(0, X.shape[0], int(batch_size)):
            end = min(start + int(batch_size), X.shape[0])
            xb = torch.as_tensor(np.asarray(X[start:end]), dtype=torch.float32, device=dev)

            mu_dict, logvar_dict = model.encode_modalities({src_mod: xb})

            if use_moe and hasattr(model, "mixture_of_experts"):
                mu_z, _ = model.mixture_of_experts(mu_dict, logvar_dict)
            elif hasattr(model, "fuse_posteriors"):
                mu_z, _ = model.fuse_posteriors(mu_dict, logvar_dict)
            else:
                mu_z = mu_dict[src_mod]

            xhat_dict = model.decode_modalities(mu_z)
            if tgt_mod not in xhat_dict:
                raise KeyError(f"Target modality {tgt_mod!r} not found. Available: {list(xhat_dict.keys())}")

            xhat_t = _extract_tensor_from_decoder_output(
                xhat_dict[tgt_mod],
                prefer=decoder_prefer,
                apply_link=decoder_link,
            )
            preds.append(_to_numpy(xhat_t))

    return np.vstack(preds) if preds else np.zeros((0, 0), dtype=np.float32)


# =============================================================================
# README: fused encoding + gate retrieval (paired / multi-observed)
# =============================================================================
def encode_moe_gates_from_tensors(
    model,
    x_dict: Mapping[str, Union[np.ndarray, "sp.spmatrix", torch.Tensor]],
    *,
    device: str = "cpu",
    batch_size: int = 1024,
    modality_order: Optional[Sequence[str]] = None,
    kind: str = "effective_precision",   # "effective_precision" | "router_x_precision" | "uniform"
    return_logits: bool = True,
    eps: float = 1e-8,
) -> Dict[str, Any]:
    """
    Compute per-cell modality *contribution weights* (aka "gates") for analytic fusion diagnostics.

    This helper is designed for the **analytic fusion** path (MoE/PoE-style), where each modality
    has a Gaussian posterior (mu_m, logvar_m) in latent space. We summarize how much each modality
    contributes to the fused posterior on a per-cell basis.

    Importantly, this function is intentionally **independent of cfg.use_moe_gating**:
      - If your model exposes router logits (best-effort via model.encode_fused(..., return_gate_logits=True)),
        we can optionally incorporate them.
      - If logits are unavailable, we still provide meaningful diagnostics using precision-only weights.

    Parameters
    ----------
    x_dict:
      dict {modality: X} with shape (n_cells, n_features) for each modality.
      Arrays may be numpy, scipy sparse, or torch tensors.
      All modalities must have the same n_cells.

    modality_order:
      Order to report/stack modalities in outputs. If None, uses model cfg order if possible,
      otherwise uses sorted(x_dict.keys()).

    kind:
      - "effective_precision":
          Precision-only weights derived from per-modality posterior uncertainty:
            s_m = sum_d exp(-logvar_m[d])
            w_m = s_m / sum_j s_j
          Always available (no router needed).
      - "router_x_precision":
          If router logits are available, combine router probabilities with effective precision:
            r = softmax(logits)
            w_m ∝ s_m * r_m
          If logits are unavailable, this silently falls back to "effective_precision".
      - "uniform":
          Equal weights across modalities (sanity check).

    return_logits:
      If True, attempt to retrieve router logits. If not available, "logits" will be None.

    Returns
    -------
    dict with:
      - weights: (n_cells, n_modalities) float32, rows sum to 1
      - logits: (n_cells, n_modalities) float32 or None
      - modality_order: list[str]
      - kind: str (the *effective* kind actually used; may downgrade router_x_precision -> effective_precision)
      - requested_kind: str
      - per_modality_mean: dict {modality: mean weight}
    """
    model.eval()
    dev = torch.device(device)

    def _normalize_rows(x: torch.Tensor) -> torch.Tensor:
        return x / (x.sum(dim=1, keepdim=True) + float(eps))

    # Decide output modality order
    mods_in = list(x_dict.keys())
    if len(mods_in) == 0:
        raise ValueError("x_dict is empty.")

    if modality_order is None:
        modality_order = _infer_model_modality_order(model, fallback=sorted(mods_in))
    else:
        modality_order = list(map(str, list(modality_order)))

    # Normalize inputs to numpy + validate n_cells match
    X_np: Dict[str, np.ndarray] = {}
    n: Optional[int] = None
    for m in modality_order:
        if m not in x_dict:
            raise KeyError(f"modality {m!r} not found in x_dict. Available: {list(x_dict.keys())}")

        Xm = x_dict[m]
        if torch.is_tensor(Xm):
            Xm = Xm.detach().cpu().numpy()
        Xm = Xm.toarray() if sp.issparse(Xm) else np.asarray(Xm)
        Xm = np.asarray(Xm)

        if n is None:
            n = int(Xm.shape[0])
        if int(Xm.shape[0]) != int(n):
            raise ValueError(f"All modalities must have same n_cells. {m} has {Xm.shape[0]} vs {n}.")

        X_np[m] = Xm

    n = int(n)

    requested_kind = str(kind).lower().strip()
    if requested_kind not in {"effective_precision", "router_x_precision", "uniform"}:
        raise ValueError("kind must be 'effective_precision', 'router_x_precision', or 'uniform'.")

    weights_chunks: List[np.ndarray] = []
    logits_chunks: List[np.ndarray] = []

    with torch.no_grad():
        for start in range(0, n, int(batch_size)):
            end = min(start + int(batch_size), n)

            xb_dict = {
                m: torch.as_tensor(np.asarray(X_np[m][start:end]), dtype=torch.float32, device=dev)
                for m in modality_order
            }

            # modality posteriors
            mu_dict, logvar_dict = model.encode_modalities(xb_dict)

            # ---- router logits (best-effort) ----
            logits = None
            if return_logits and hasattr(model, "encode_fused"):
                try:
                    out = model.encode_fused(
                        xb_dict,
                        use_mean=True,
                        return_gates=True,
                        return_gate_logits=True,
                    )
                    # expected: mu, logvar, z, gates, gate_logits
                    if isinstance(out, (tuple, list)) and len(out) >= 5 and out[4] is not None:
                        logits = out[4]
                except Exception:
                    logits = None

            if logits is not None:
                logits_chunks.append(_to_numpy(logits))

            # ---- base scalars: effective precision per modality per cell ----
            if requested_kind == "uniform":
                w = torch.ones((end - start, len(modality_order)), device=dev, dtype=torch.float32)
                w = _normalize_rows(w)

            else:
                s_list = []
                for m in modality_order:
                    lv = logvar_dict[m]          # (batch, latent_dim)
                    prec = torch.exp(-lv)        # (batch, latent_dim)
                    s = prec.sum(dim=1)          # (batch,)
                    s_list.append(s)
                S = torch.stack(s_list, dim=1)   # (batch, n_mods)

                if requested_kind == "router_x_precision" and (logits is not None):
                    R = _softmax(logits, dim=1)  # (batch, n_mods)
                    w = _normalize_rows(S * R)
                else:
                    w = _normalize_rows(S)

            weights_chunks.append(_to_numpy(w))

    W = np.vstack(weights_chunks).astype(np.float32) if weights_chunks else np.zeros((0, 0), dtype=np.float32)
    L = np.vstack(logits_chunks).astype(np.float32) if logits_chunks else None

    # What kind did we *actually* deliver?
    effective_kind = requested_kind
    if requested_kind == "router_x_precision" and L is None:
        effective_kind = "effective_precision"

    per_mod_mean = (
        {m: float(W[:, j].mean()) for j, m in enumerate(modality_order)} if W.size else {m: 0.0 for m in modality_order}
    )

    return {
        "weights": W,
        "logits": L,
        "modality_order": list(modality_order),
        "kind": effective_kind,
        "requested_kind": requested_kind,
        "per_modality_mean": per_mod_mean,
    }


def encode_fused_adata_pair(
    model,
    adata_by_mod: Mapping[str, Any],
    *,
    device: str = "cpu",
    batch_size: int = 1024,
    use_mean: bool = True,
    return_gates: bool = True,
    return_gate_logits: bool = True,
    write_to_adatas: bool = True,
    fused_obsm_key: str = "X_univi_fused",
    gate_prefix: str = "gate",
    layer_by_mod: Optional[Mapping[str, Optional[str]]] = None,
    X_key_by_mod: Optional[Mapping[str, str]] = None,
) -> Dict[str, Any]:
    """
    Encode a fused posterior for paired/multi-observed cells.

    Expected:
      - all AnnData share identical obs_names in the same order.

    Returns dict with keys:
      - "Z_fused" (n_cells, latent_dim)
      - "mu", "logvar"
      - "gates" (or None)
      - "gate_logits" (or None)
      - "modality_order" (the order corresponding to columns of gates)
    """
    from .data import _get_matrix

    mods_in = list(adata_by_mod.keys())
    if len(mods_in) == 0:
        raise ValueError("adata_by_mod is empty.")

    # validate paired obs
    first = adata_by_mod[mods_in[0]]
    n = int(first.n_obs)
    obs0 = list(first.obs_names)
    for m in mods_in[1:]:
        a = adata_by_mod[m]
        if int(a.n_obs) != n:
            raise ValueError(f"All modalities must have same n_obs. {mods_in[0]}={n}, {m}={a.n_obs}")
        if list(a.obs_names) != obs0:
            raise ValueError(f"All modalities must have identical obs_names order. Mismatch at {m!r}.")

    # stable ordering for inputs; gates order should follow model order if possible
    mods_sorted = sorted(mods_in)
    modality_order = _infer_model_modality_order(model, fallback=mods_sorted)

    # fetch matrices once (can be sparse); slice per batch
    X_by_mod = {}
    for m in mods_sorted:
        layer = None if layer_by_mod is None else layer_by_mod.get(m, None)
        xkey = "X" if X_key_by_mod is None else X_key_by_mod.get(m, "X")
        X_by_mod[m] = _get_matrix(adata_by_mod[m], layer=layer, X_key=xkey)

    dev = torch.device(device)
    model.eval()

    mu_chunks: List[np.ndarray] = []
    lv_chunks: List[np.ndarray] = []
    z_chunks: List[np.ndarray] = []
    gates_chunks: List[np.ndarray] = []
    glog_chunks: List[np.ndarray] = []

    with torch.no_grad():
        for start in range(0, n, int(batch_size)):
            end = min(start + int(batch_size), n)

            x_dict_t = {}
            for m in mods_sorted:
                Xm = X_by_mod[m][start:end]
                Xm = Xm.toarray() if sp.issparse(Xm) else np.asarray(Xm)
                x_dict_t[m] = torch.as_tensor(np.asarray(Xm), dtype=torch.float32, device=dev)

            out = model.encode_fused(
                x_dict_t,
                use_mean=bool(use_mean),
                return_gates=bool(return_gates),
                return_gate_logits=bool(return_gate_logits),
            )

            # model.encode_fused returns:
            #   mu, logvar, z  OR  mu, logvar, z, gates, gate_logits
            if isinstance(out, (tuple, list)) and len(out) >= 3:
                mu, logvar, z = out[0], out[1], out[2]
                gates = out[3] if len(out) > 3 else None
                gate_logits = out[4] if len(out) > 4 else None
            else:
                raise TypeError("encode_fused returned unexpected output type.")

            mu_chunks.append(_to_numpy(mu))
            lv_chunks.append(_to_numpy(logvar))
            z_chunks.append(_to_numpy(z))

            if gates is not None:
                gates_chunks.append(_to_numpy(gates))
            if gate_logits is not None:
                glog_chunks.append(_to_numpy(gate_logits))

    mu_all = np.vstack(mu_chunks) if mu_chunks else np.zeros((0, 0), dtype=np.float32)
    lv_all = np.vstack(lv_chunks) if lv_chunks else np.zeros((0, 0), dtype=np.float32)
    z_all = np.vstack(z_chunks) if z_chunks else np.zeros((0, 0), dtype=np.float32)
    gates_all = np.vstack(gates_chunks) if gates_chunks else None
    glog_all = np.vstack(glog_chunks) if glog_chunks else None

    rep: Dict[str, Any] = {
        "mu": mu_all,
        "logvar": lv_all,
        "Z_fused": z_all,
        "gates": gates_all,
        "gate_logits": glog_all,
        "modality_order": modality_order,
    }

    if write_to_adatas:
        for m in mods_in:
            adata_by_mod[m].obsm[str(fused_obsm_key)] = z_all

        # write gates into .obs columns if present
        if gates_all is not None:
            # gates columns correspond to model modality_order (best-effort)
            for j, name in enumerate(modality_order[: gates_all.shape[1]]):
                col = f"{gate_prefix}_{name}"
                for m in mods_in:
                    adata_by_mod[m].obs[col] = gates_all[:, j].astype(np.float32)

        if glog_all is not None:
            for j, name in enumerate(modality_order[: glog_all.shape[1]]):
                col = f"{gate_prefix}_logit_{name}"
                for m in mods_in:
                    adata_by_mod[m].obs[col] = glog_all[:, j].astype(np.float32)

    return rep


# =============================================================================
# README: denoising
# =============================================================================
def denoise_adata(
    model,
    adata,
    *,
    modality: str,
    device: str = "cpu",
    out_layer: str = "denoised_fused",
    overwrite_X: bool = False,
    batch_size: int = 512,
    # If provided -> true multimodal denoising via fused latent:
    adata_by_mod: Optional[Mapping[str, Any]] = None,
    layer_by_mod: Optional[Mapping[str, Optional[str]]] = None,
    X_key_by_mod: Optional[Mapping[str, str]] = None,
    use_mean: bool = True,
    # Decoder output handling:
    decoder_prefer: str = "auto",
    decoder_link: Optional[str] = None,
) -> Any:
    """
    Denoise (reconstruct) a modality.

    - If adata_by_mod is None: encodes only this modality, decodes this modality (self denoise).
    - If adata_by_mod is provided: encodes a fused posterior from adata_by_mod (paired/multi-observed),
      then decodes `modality` and writes into `adata.layers[out_layer]`.

    Returns the modified `adata` (for chaining).
    """
    from .data import _get_matrix

    dev = torch.device(device)
    model.eval()

    n = int(adata.n_obs)
    preds: List[np.ndarray] = []

    # fast path: fused denoise
    if adata_by_mod is not None:
        # ensure target modality exists in mapping (or at least decodable)
        if modality not in adata_by_mod:
            # still okay: you can decode a modality even if it wasn't observed,
            # but we need paired obs_names source for fused encoding.
            pass

        fused = encode_fused_adata_pair(
            model,
            adata_by_mod=adata_by_mod,
            device=device,
            batch_size=batch_size,
            use_mean=use_mean,
            return_gates=False,
            return_gate_logits=False,
            write_to_adatas=False,
            layer_by_mod=layer_by_mod,
            X_key_by_mod=X_key_by_mod,
        )
        Z = np.asarray(fused["Z_fused"], dtype=np.float32)

        with torch.no_grad():
            for start in range(0, n, int(batch_size)):
                end = min(start + int(batch_size), n)
                zb = torch.as_tensor(Z[start:end], dtype=torch.float32, device=dev)
                dec = model.decode_modalities(zb)
                if modality not in dec:
                    raise KeyError(f"Decoder did not return modality {modality!r}. Available: {list(dec.keys())}")
                x = _extract_tensor_from_decoder_output(dec[modality], prefer=decoder_prefer, apply_link=decoder_link)
                preds.append(_to_numpy(x))

    # self denoise: encode only this modality
    else:
        X = _get_matrix(adata, layer=None, X_key="X")
        X = X.toarray() if sp.issparse(X) else np.asarray(X)

        with torch.no_grad():
            for start in range(0, n, int(batch_size)):
                end = min(start + int(batch_size), n)
                xb = torch.as_tensor(np.asarray(X[start:end]), dtype=torch.float32, device=dev)
                mu_dict, logvar_dict = model.encode_modalities({modality: xb})

                # prefer fused/moe if model wants to do that with single expert? for self-denoise,
                # simplest/most intuitive is the modality posterior mean.
                mu = mu_dict[modality]
                dec = model.decode_modalities(mu)
                if modality not in dec:
                    raise KeyError(f"Decoder did not return modality {modality!r}. Available: {list(dec.keys())}")
                x = _extract_tensor_from_decoder_output(dec[modality], prefer=decoder_prefer, apply_link=decoder_link)
                preds.append(_to_numpy(x))

    Xhat = np.vstack(preds) if preds else np.zeros((0, 0), dtype=np.float32)

    if overwrite_X:
        adata.X = Xhat
    else:
        adata.layers[str(out_layer)] = Xhat

    return adata


def denoise_from_multimodal(
    model,
    adata,
    *,
    modality: str,
    adata_by_mod: Mapping[str, Any],
    device: str = "cpu",
    out_layer: str = "denoised_fused",
    overwrite_X: bool = False,
    batch_size: int = 512,
    layer_by_mod: Optional[Mapping[str, Optional[str]]] = None,
    X_key_by_mod: Optional[Mapping[str, str]] = None,
    use_mean: bool = True,
    decoder_prefer: str = "auto",
    decoder_link: Optional[str] = None,
) -> Any:
    """README alias: true multimodal denoising via fused latent."""
    return denoise_adata(
        model,
        adata=adata,
        modality=modality,
        device=device,
        out_layer=out_layer,
        overwrite_X=overwrite_X,
        batch_size=batch_size,
        adata_by_mod=adata_by_mod,
        layer_by_mod=layer_by_mod,
        X_key_by_mod=X_key_by_mod,
        use_mean=use_mean,
        decoder_prefer=decoder_prefer,
        decoder_link=decoder_link,
    )


# =============================================================================
# README: alignment evaluation (one-call)
# =============================================================================
def evaluate_alignment(
    *,
    Z1: np.ndarray,
    Z2: np.ndarray,
    metric: str = "euclidean",
    recall_ks: Sequence[int] = (1, 5, 10),
    k_mixing: int = 20,
    k_entropy: int = 30,
    labels_source: Optional[np.ndarray] = None,
    labels_target: Optional[np.ndarray] = None,
    compute_bidirectional_transfer: bool = True,
    k_transfer: int = 15,
    json_safe: bool = True,
    block_size: int = 512,
) -> Dict[str, Any]:
    """
    README-facing alignment evaluation.

    Computes:
      - FOSCTTM (mean + SEM)
      - Recall@k (mean + SEM) for each k
      - modality mixing + entropy in the *stacked* space
      - label transfer (source->target), and optionally bidirectional summary

    Returns a flat dict with stable keys used by README examples.
    """
    Z1 = np.asarray(Z1, dtype=np.float32)
    Z2 = np.asarray(Z2, dtype=np.float32)

    out: Dict[str, Any] = {
        "metric": str(metric),
        "n_cells": int(Z1.shape[0]),
    }

    fos_m, fos_s = compute_foscttm(Z1, Z2, metric=metric, block_size=block_size, return_sem=True)
    out["foscttm_mean"] = float(fos_m)
    out["foscttm_sem"] = float(fos_s)

    recalls = {}
    for k in recall_ks:
        rm, rs = compute_match_recall_at_k(Z1, Z2, k=int(k), metric=metric, block_size=block_size, return_sem=True)
        recalls[int(k)] = {"mean": float(rm), "sem": float(rs)}
    out["recall_at_k"] = recalls

    # modality mixing/entropy computed on stacked space
    Z_stack = np.vstack([Z1, Z2])
    mod_labels = np.array(["mod1"] * Z1.shape[0] + ["mod2"] * Z2.shape[0], dtype=object)

    mix_m, mix_s = compute_modality_mixing(Z_stack, mod_labels, k=int(k_mixing), metric=metric, return_sem=True)
    ent_m, ent_s = compute_modality_entropy(Z_stack, mod_labels, k=int(k_entropy), metric=metric, return_sem=True)
    out["modality_mixing_mean"] = float(mix_m)
    out["modality_mixing_sem"] = float(mix_s)
    out["modality_entropy_mean"] = float(ent_m)
    out["modality_entropy_sem"] = float(ent_s)
    out["k_mixing"] = int(k_mixing)
    out["k_entropy"] = int(k_entropy)

    # label transfer
    if labels_source is not None and labels_target is not None:
        pred, acc, cm, order, f1d = label_transfer_knn(
            Z_source=Z1,
            labels_source=np.asarray(labels_source),
            Z_target=Z2,
            labels_target=np.asarray(labels_target),
            k=int(k_transfer),
            metric=str(metric),
            return_label_order=True,
            return_f1=True,
        )
        out["label_transfer_acc"] = float(acc)
        out["label_transfer_cm"] = cm
        out["label_transfer_label_order"] = order
        out["label_transfer_macro_f1"] = float(f1d["macro_f1"])
        out["label_transfer_weighted_f1"] = float(f1d["weighted_f1"])
        out["label_transfer_pred"] = pred
        out["k_transfer"] = int(k_transfer)

        if compute_bidirectional_transfer:
            bi = summarize_bidirectional_transfer(Z1, np.asarray(labels_source), Z2, np.asarray(labels_target), k=int(k_transfer), metric=str(metric))
            out["bidirectional_transfer"] = bi
            out["worst_direction_macro_f1"] = float(bi["worst_direction_macro_f1"])
            out["worst_direction_weighted_f1"] = float(bi["worst_direction_weighted_f1"])

    return _json_safe(out) if json_safe else out


# =============================================================================
# README: latent sampling / generation utilities
# =============================================================================
def fit_latent_gaussians_by_label(
    Z: np.ndarray,
    labels: Sequence[Any],
    *,
    min_n: int = 20,
    shrink: float = 1e-3,
) -> Dict[str, Dict[str, Any]]:
    """
    Fit diagonal Gaussians per label:
      mean_c, var_c (per dim)

    shrink: add to var for stability.
    """
    Z = np.asarray(Z, dtype=np.float32)
    y = np.asarray(labels).astype(str)

    out: Dict[str, Dict[str, Any]] = {}
    for lab in np.unique(y):
        idx = np.where(y == lab)[0]
        if idx.size < int(min_n):
            continue
        Zc = Z[idx]
        mu = Zc.mean(axis=0)
        var = Zc.var(axis=0) + float(shrink)
        out[str(lab)] = {"n": int(idx.size), "mean": mu.astype(np.float32), "var": var.astype(np.float32)}
    return out


def sample_latent_from_label_gaussians(
    gauss: Dict[str, Dict[str, Any]],
    label: str,
    n: int,
    *,
    random_state: int = 0,
) -> np.ndarray:
    if label not in gauss:
        raise KeyError(f"label {label!r} not present. Available: {list(gauss.keys())[:20]}")
    rng = np.random.default_rng(int(random_state))
    mu = gauss[label]["mean"]
    var = gauss[label]["var"]
    eps = rng.standard_normal((int(n), mu.shape[0]), dtype=np.float32)
    return mu[None, :] + eps * np.sqrt(var[None, :]).astype(np.float32)


def decode_from_latent(
    model,
    Z: np.ndarray,
    *,
    device: str = "cpu",
    batch_size: int = 512,
    decoder_prefer: str = "auto",
    decoder_link: Optional[str] = None,
) -> Dict[str, np.ndarray]:
    """
    Decode latents into *all* modalities returned by model.decode_modalities.
    Returns modality -> numpy array
    """
    model.eval()
    dev = torch.device(device)
    Z = np.asarray(Z, dtype=np.float32)

    outs: Dict[str, List[np.ndarray]] = {}
    with torch.no_grad():
        for start in range(0, Z.shape[0], int(batch_size)):
            end = min(start + int(batch_size), Z.shape[0])
            zb = torch.as_tensor(Z[start:end], dtype=torch.float32, device=dev)

            dec = model.decode_modalities(zb)
            for mod, v in dec.items():
                x = _extract_tensor_from_decoder_output(v, prefer=decoder_prefer, apply_link=decoder_link)
                outs.setdefault(mod, []).append(_to_numpy(x))

    return {m: np.vstack(chunks) for m, chunks in outs.items()}


def generate_from_latent(
    model,
    n: Optional[int] = None,
    *,
    target_mod: Optional[str] = None,
    device: str = "cpu",
    z_source: str = "prior",
    z: Optional[np.ndarray] = None,
    batch_size: int = 512,
    return_mean: bool = True,        # kept for API parity
    sample_likelihood: bool = False, # accepted but not implemented
    decoder_prefer: str = "auto",
    decoder_link: Optional[str] = None,
    random_state: int = 0,
) -> Union[np.ndarray, Dict[str, np.ndarray]]:
    """
    README-compatible generator.

    Notes
    -----
    - If z is None, samples z ~ N(0, I) when z_source == "prior".
    - return_mean is kept for API parity (decoders return mean-like outputs).
    - sample_likelihood is accepted but not implemented here.
    """
    if not return_mean:
        # decoders here return mean-like outputs; we keep the knob but warn by behavior.
        pass
    if sample_likelihood:
        raise NotImplementedError("sample_likelihood=True is not implemented yet in generate_from_latent().")

    if z is None:
        if n is None:
            raise ValueError("Provide either `z` or `n`.")
        if str(z_source).lower().strip() != "prior":
            raise ValueError("Only z_source='prior' is supported when `z` is not provided.")

        latent_dim = getattr(getattr(model, "cfg", None), "latent_dim", None)
        if latent_dim is None:
            latent_dim = getattr(model, "latent_dim", None)
        if latent_dim is None:
            raise ValueError("Could not infer latent_dim from model; provide `z` explicitly.")

        rng = np.random.default_rng(int(random_state))
        z = rng.standard_normal((int(n), int(latent_dim)), dtype=np.float32)

    dec = decode_from_latent(
        model,
        np.asarray(z, dtype=np.float32),
        device=device,
        batch_size=batch_size,
        decoder_prefer=decoder_prefer,
        decoder_link=decoder_link,
    )

    if target_mod is None:
        return dec
    if target_mod not in dec:
        raise KeyError(f"target_mod {target_mod!r} not found. Available: {list(dec.keys())}")
    return dec[target_mod]


# =============================================================================
# README: Evaluate reconstruction / imputation error against ground truth
# =============================================================================
def evaluate_prediction_against_adata(
    adata_true,
    X_pred: np.ndarray,
    *,
    layer_true: Optional[str] = None,
    X_key_true: str = "X",
    kind: str = "continuous",     # "continuous" | "binary"
    feature_names: Optional[Sequence[str]] = None,
) -> Dict[str, Any]:
    """
    Compare X_pred (n_cells, n_features) to adata_true ground truth matrix.

    If feature_names is provided, it subsets BOTH true and pred to those features
    (by matching adata_true.var_names).
    """
    from .data import _get_matrix

    X_true = _get_matrix(adata_true, layer=layer_true, X_key=X_key_true)
    X_true = to_dense(X_true).astype(np.float32, copy=False)

    X_pred = np.asarray(X_pred, dtype=np.float32)
    if X_true.shape != X_pred.shape:
        raise ValueError(f"Shape mismatch: true {X_true.shape} vs pred {X_pred.shape}")

    feat_names_out: Optional[List[str]] = None
    if feature_names is not None:
        wanted = [str(x) for x in list(feature_names)]
        name_to_j = {str(n): j for j, n in enumerate(map(str, list(adata_true.var_names)))}
        cols = []
        for f in wanted:
            if f not in name_to_j:
                raise KeyError(f"Feature {f!r} not found in adata_true.var_names.")
            cols.append(int(name_to_j[f]))
        cols = np.asarray(cols, dtype=int)
        X_true = X_true[:, cols]
        X_pred = X_pred[:, cols]
        feat_names_out = wanted
    else:
        feat_names_out = [str(x) for x in list(map(str, list(adata_true.var_names)))]

    met = reconstruction_metrics(X_true, X_pred, kind=kind)

    per_cell = np.mean((X_true - X_pred) ** 2, axis=1)
    met["per_cell_mse"] = per_cell.astype(np.float32)
    met["per_cell_mean"] = float(np.mean(per_cell))

    # For plotting + downstream use
    met["X_true"] = X_true
    met["X_pred"] = X_pred
    met["feature_names"] = feat_names_out

    # README convenience: "summary" + "per_feature"
    if str(kind).lower().strip() == "binary":
        met["summary"] = {
            "auc_mean": float(met["auc_mean"]),
            "auc_median": float(met["auc_median"]),
            "per_cell_mean": float(met["per_cell_mean"]),
        }
        met["per_feature"] = {"auc": np.asarray(met["auc_per_feature"])}
    else:
        met["summary"] = {
            "mse_mean": float(met["mse_mean"]),
            "mse_median": float(met["mse_median"]),
            "pearson_mean": float(met["pearson_mean"]),
            "pearson_median": float(met["pearson_median"]),
            "per_cell_mean": float(met["per_cell_mean"]),
        }
        met["per_feature"] = {
            "mse": np.asarray(met["mse_per_feature"]),
            "pearson": np.asarray(met["pearson_per_feature"]),
        }

    return met


def evaluate_cross_reconstruction(
    model,
    adata_src,
    adata_tgt,
    *,
    src_mod: str,
    tgt_mod: str,
    device: str = "cpu",
    src_layer: Optional[str] = None,
    src_X_key: str = "X",
    tgt_layer: Optional[str] = None,
    tgt_X_key: str = "X",
    batch_size: int = 512,
    kind: str = "continuous",
    feature_names: Optional[Sequence[str]] = None,
    use_moe: bool = True,
    decoder_prefer: str = "auto",
    decoder_link: Optional[str] = None,
) -> Dict[str, Any]:
    """
    README-style one-call cross-reconstruction evaluation:

      1) X_pred = cross_modal_predict(model, adata_src, src_mod -> tgt_mod)
      2) compare X_pred against adata_tgt truth (layer/X_key)
      3) return dict with:
          - "summary"
          - "per_feature"
          - "X_true", "X_pred", "feature_names" (for plotting)
    """
    X_pred = cross_modal_predict(
        model,
        adata_src=adata_src,
        src_mod=src_mod,
        tgt_mod=tgt_mod,
        device=device,
        layer=src_layer,
        X_key=src_X_key,
        batch_size=batch_size,
        use_moe=use_moe,
        decoder_prefer=decoder_prefer,
        decoder_link=decoder_link,
    )

    rep = evaluate_prediction_against_adata(
        adata_tgt,
        X_pred,
        layer_true=tgt_layer,
        X_key_true=tgt_X_key,
        kind=kind,
        feature_names=feature_names,
    )

    rep["src_mod"] = str(src_mod)
    rep["tgt_mod"] = str(tgt_mod)
    rep["kind"] = str(kind)
    rep["device"] = str(device)
    rep["batch_size"] = int(batch_size)
    rep["decoder_prefer"] = str(decoder_prefer)
    rep["decoder_link"] = decoder_link

    return rep


# =============================================================================
# Backwards/README-compatible aliases
# =============================================================================
def fit_label_latent_gaussians(
    Z: np.ndarray,
    labels: Sequence[Any],
    *,
    min_n: int = 20,
    shrink: float = 1e-3,
) -> Dict[str, Dict[str, Any]]:
    return fit_latent_gaussians_by_label(Z, labels, min_n=min_n, shrink=shrink)


def sample_latent_by_label(
    gauss: Dict[str, Dict[str, Any]],
    label: str,
    n: int,
    *,
    random_state: int = 0,
) -> np.ndarray:
    return sample_latent_from_label_gaussians(gauss, label, n, random_state=random_state)
```
