## Supplemental Code for "Unifying multimodal single-cell data with a mixture-of-experts *β*-variational autoencoder framework": __init__.py

```
# univi/hyperparam_optimization/__init__.py

from .run_multiome_hparam_search import run_multiome_hparam_search
from .run_citeseq_hparam_search import run_citeseq_hparam_search
from .run_teaseq_hparam_search import run_teaseq_hparam_search
from .run_rna_hparam_search import run_rna_hparam_search
from .run_atac_hparam_search import run_atac_hparam_search
from .run_adt_hparam_search import run_adt_hparam_search

__all__ = [
    "run_multiome_hparam_search",
    "run_citeseq_hparam_search",
    "run_teaseq_hparam_search",
    "run_rna_hparam_search",
    "run_atac_hparam_search",
    "run_adt_hparam_search",
]
```
