## Supplemental Code for "Unifying multimodal single-cell data with a mixture-of-experts *β*-variational autoencoder framework": common.py

```
# univi/hyperparam_optimization/common.py

from __future__ import annotations
from dataclasses import dataclass
from typing import Dict, Any, List, Optional, Iterable, Tuple

import json
import time

import numpy as np
import pandas as pd
import torch
from torch.utils.data import DataLoader

from univi import UniVIMultiModalVAE, ModalityConfig, UniVIConfig, TrainingConfig
from univi.data import MultiModalDataset
from univi.trainer import UniVITrainer
from univi import evaluation as univi_eval


@dataclass
class SearchResult:
    config_id: int
    hparams: Dict[str, Any]
    best_val_loss: float
    metrics: Dict[str, float]
    runtime_min: float


def iter_hparam_configs(
    space_dict: Dict[str, List[Any]],
    max_configs: int,
    seed: int = 0,
) -> Iterable[Dict[str, Any]]:
    """
    Random sampler over a hyperparameter space dict.
    Each config independently samples a value for each key.
    """
    rng = np.random.default_rng(seed)
    keys = list(space_dict.keys())
    for _ in range(max_configs):
        hp = {}
        for k in keys:
            options = space_dict[k]
            idx = rng.integers(len(options))
            hp[k] = options[idx]
        yield hp


def build_modality_configs(
    arch_config_per_mod: Dict[str, Dict[str, Any]],
    likelihood_per_mod: Dict[str, str],
    input_dims: Dict[str, int],
) -> List[ModalityConfig]:
    """
    arch_config_per_mod: e.g. {"rna": {"enc": [...], "dec": [...]}, ...}
    likelihood_per_mod:  e.g. {"rna": "nb", "atac": "gaussian"}
    input_dims:          e.g. {"rna": rna.n_vars, "atac": atac.n_vars}
    """
    mod_cfgs: List[ModalityConfig] = []
    for mod_name, arch_cfg in arch_config_per_mod.items():
        enc = arch_cfg["enc"]
        dec = arch_cfg["dec"]
        lik = likelihood_per_mod.get(mod_name, "gaussian")
        in_dim = int(input_dims[mod_name])
        mod_cfgs.append(
            ModalityConfig(
                name=mod_name,
                input_dim=in_dim,
                encoder_hidden=list(enc),
                decoder_hidden=list(dec),
                likelihood=lik,
            )
        )
    return mod_cfgs


def make_dataloaders(
    adata_train: Dict[str, "AnnData"],
    adata_val: Dict[str, "AnnData"],
    layer: Optional[str],
    X_key: str,
    batch_size: int,
    num_workers: int,
    device: Optional[str] = None,
) -> Tuple[DataLoader, DataLoader]:
    """
    Thin wrapper to build MultiModalDataset and DataLoader for train/val.
    """
    device_obj = None
    if device is not None and device != "cpu":
        device_obj = torch.device(device)

    train_ds = MultiModalDataset(
        adata_dict=adata_train,
        layer=layer,
        X_key=X_key,
        paired=True,
        device=device_obj,
    )
    val_ds = MultiModalDataset(
        adata_dict=adata_val,
        layer=layer,
        X_key=X_key,
        paired=True,
        device=device_obj,
    )

    train_loader = DataLoader(
        train_ds,
        batch_size=batch_size,
        shuffle=True,
        num_workers=num_workers,
    )
    val_loader = DataLoader(
        val_ds,
        batch_size=batch_size,
        shuffle=False,
        num_workers=num_workers,
    )

    return train_loader, val_loader


def train_single_config(
    config_id: int,
    hparams: Dict[str, Any],
    mod_arch_space: Dict[str, List[Dict[str, Any]]],
    modalities: List[str],
    input_dims: Dict[str, int],
    likelihood_per_mod: Dict[str, List[str]],
    adata_train: Dict[str, "AnnData"],
    adata_val: Dict[str, "AnnData"],
    base_train_cfg: TrainingConfig,
    layer: Optional[str],
    X_key: str,
    celltype_key: Optional[str],
    device: str = "cuda",
    multimodal_eval: bool = True,
) -> SearchResult:
    """
    Train one UniVI model for a given hyperparam config and return metrics.

    multimodal_eval:
        - True: compute FOSCTTM, label transfer, modality mixing where possible
        - False: only val_loss (for unimodal).
    """
    start = time.time()

    # ----- pick architectures -----
    arch_cfg_per_mod: Dict[str, Dict[str, Any]] = {}
    like_cfg_per_mod: Dict[str, str] = {}
    for mod in modalities:
        arch_list = mod_arch_space[mod]
        arch_choice = hparams[f"{mod}_arch"]
        if isinstance(arch_choice, dict) and "name" in arch_choice:
            arch_cfg = arch_choice
        else:
            # if user passed just the dict, keep as-is
            arch_cfg = arch_choice
        arch_cfg_per_mod[mod] = arch_cfg

        like_options = likelihood_per_mod.get(mod, ["gaussian"])
        like_choice = hparams.get(f"{mod}_likelihood", like_options[0])
        like_cfg_per_mod[mod] = like_choice

    # ----- build modality configs -----
    mod_cfgs = build_modality_configs(
        arch_config_per_mod=arch_cfg_per_mod,
        likelihood_per_mod=like_cfg_per_mod,
        input_dims=input_dims,
    )

    # ----- UniVI config -----
    univi_cfg = UniVIConfig(
        latent_dim=hparams["latent_dim"],
        modalities=mod_cfgs,
        beta=hparams["beta"],
        gamma=hparams["gamma"],
        encoder_dropout=hparams["encoder_dropout"],
        decoder_dropout=0.0,
        encoder_batchnorm=True,
        decoder_batchnorm=hparams["decoder_batchnorm"],
        kl_anneal_start=0,
        kl_anneal_end=0,
        align_anneal_start=0,
        align_anneal_end=0,
    )

    model = UniVIMultiModalVAE(univi_cfg)

    # ----- dataloaders -----
    train_loader, val_loader = make_dataloaders(
        adata_train=adata_train,
        adata_val=adata_val,
        layer=layer,
        X_key=X_key,
        batch_size=base_train_cfg.batch_size,
        num_workers=base_train_cfg.num_workers,
        device=device,
    )

    # ----- training config -----
    train_cfg = TrainingConfig(
        n_epochs=base_train_cfg.n_epochs,
        batch_size=base_train_cfg.batch_size,
        lr=hparams["lr"],
        weight_decay=hparams["weight_decay"],
        device=device,
        log_every=base_train_cfg.log_every,
        grad_clip=base_train_cfg.grad_clip,
        num_workers=base_train_cfg.num_workers,
        seed=base_train_cfg.seed,
        early_stopping=base_train_cfg.early_stopping,
        patience=base_train_cfg.patience,
        min_delta=base_train_cfg.min_delta,
    )

    trainer = UniVITrainer(
        model=model,
        train_loader=train_loader,
        val_loader=val_loader,
        train_cfg=train_cfg,
        device=device,
    )

    print("=" * 80)
    print(f"[Config {config_id}] Hyperparameters:")
    print(json.dumps(hparams, indent=2))
    print("=" * 80, flush=True)

    history = trainer.fit()
    best_val_loss = trainer.best_val_loss

    metrics: Dict[str, float] = {"best_val_loss": float(best_val_loss)}

    # ----- evaluation metrics -----
    if multimodal_eval and len(modalities) >= 2:
        # encode all modalities for val set
        Z_val: Dict[str, np.ndarray] = {}
        for mod in modalities:
            # use same layer used in training
            Z = trainer.encode_modality(
                adata_val[mod],
                modality=mod,
                layer=layer,
                X_key=X_key,
                batch_size=1024,
            )
            Z_val[mod] = Z
            print(f"  Encoded modality {mod} into latent shape {Z.shape}")

        # pairwise FOSCTTM
        mods = modalities
        fos_vals = []
        for i in range(len(mods)):
            for j in range(i + 1, len(mods)):
                m1, m2 = mods[i], mods[j]
                fos = univi_eval.compute_foscttm(Z_val[m1], Z_val[m2])
                key = f"foscttm_{m1}_vs_{m2}"
                metrics[key] = float(fos)
                fos_vals.append(fos)
                print(f"  FOSCTTM ({m1} vs {m2}): {fos:.4f}")

        if fos_vals:
            metrics["foscttm_mean"] = float(np.mean(fos_vals))

        # label transfer: use first modality as "reference" if celltype_key is given
        if celltype_key is not None and celltype_key in adata_val[modalities[0]].obs:
            ref_mod = modalities[0]
            labels_ref = adata_val[ref_mod].obs[celltype_key].astype(str).values

            for tgt_mod in modalities[1:]:
                labels_tgt = (
                    adata_val[tgt_mod].obs[celltype_key].astype(str).values
                    if celltype_key in adata_val[tgt_mod].obs
                    else None
                )
                _, acc, _ = univi_eval.label_transfer_knn(
                    Z_source=Z_val[ref_mod],
                    labels_source=labels_ref,
                    Z_target=Z_val[tgt_mod],
                    labels_target=labels_tgt,
                    k=15,
                )
                if acc is not None:
                    key = f"label_acc_{tgt_mod}_from_{ref_mod}"
                    metrics[key] = float(acc)
                    print(f"  Label transfer ({ref_mod}→{tgt_mod}) accuracy: {acc:.3f}")

        # modality mixing
        Z_joint = np.concatenate(list(Z_val.values()), axis=0)
        modality_labels = np.concatenate(
            [[m] * Z_val[m].shape[0] for m in modalities]
        )
        mix_score = univi_eval.compute_modality_mixing(Z_joint, modality_labels)
        metrics["modality_mixing"] = float(mix_score)
        print(f"  Modality mixing score: {mix_score:.4f}")

        # composite score: simple weighted combo
        if "foscttm_mean" in metrics:
            comp = best_val_loss + 1000.0 * metrics["foscttm_mean"]
        else:
            comp = best_val_loss
        metrics["composite_score"] = float(comp)
        print(f"  Composite score: {comp:.3f}")
    else:
        # unimodal: composite == val loss
        metrics["composite_score"] = float(best_val_loss)

    runtime_min = (time.time() - start) / 60.0
    print(f"[Config {config_id}] Done in {runtime_min:.2f} min")
    print(
        f"  best_val_loss         = {best_val_loss:.3f}\n"
        f"  composite_score       = {metrics['composite_score']:.3f}"
    )

    return SearchResult(
        config_id=config_id,
        hparams=hparams,
        best_val_loss=float(best_val_loss),
        metrics=metrics,
        runtime_min=runtime_min,
    )


def results_to_dataframe(results: List[SearchResult]) -> pd.DataFrame:
    rows = []
    for r in results:
        row = {
            "config_id": r.config_id,
            "runtime_min": r.runtime_min,
        }
        row.update(r.hparams)
        row.update(r.metrics)
        rows.append(row)
    df = pd.DataFrame(rows)
    df = df.sort_values("composite_score", ascending=True).reset_index(drop=True)
    return df
```
