## Supplemental Code for "Unifying multimodal single-cell data with a mixture-of-experts *β*-variational autoencoder framework": run_atac_hparam_search.py

```
# univi/hyperparam_optimization/run_atac_hparam_search.py

from __future__ import annotations
from typing import List, Any, Optional

from anndata import AnnData

from univi.config import TrainingConfig
from .common import (
    iter_hparam_configs,
    train_single_config,
    results_to_dataframe,
)


def run_atac_hparam_search(
    atac_train: AnnData,
    atac_val: AnnData,
    device: str = "cuda",
    layer: Optional[str] = "counts",  # raw peaks; or change to TF-IDF/LSI etc.
    X_key: str = "X",
    max_configs: int = 50,
    seed: int = 0,
    base_train_cfg: Optional[TrainingConfig] = None,
):
    """
    Hyperparameter search for *unimodal* ATAC.
    """

    adata_train = {"atac": atac_train}
    adata_val = {"atac": atac_val}
    modalities = ["atac"]

    if base_train_cfg is None:
        base_train_cfg = TrainingConfig(
            n_epochs=80,
            batch_size=256,
            lr=1e-3,
            weight_decay=1e-5,
            device=device,
            log_every=5,
            grad_clip=5.0,
            num_workers=0,
            seed=42,
            early_stopping=True,
            patience=15,
            min_delta=0.0,
        )

    atac_arch_options = [
        {"name": "atac_med2", "enc": [512, 256], "dec": [256, 512]},
        {"name": "atac_wide2", "enc": [1024, 512], "dec": [512, 1024]},
    ]

    mod_arch_space = {"atac": atac_arch_options}
    likelihood_per_mod = {
        "atac": ["nb", "poisson", "zinb", "gaussian"],
    }

    search_space = {
        "latent_dim":       [10, 20, 32, 40, 50, 64, 82, 120, 160, 200],
        "beta":             [0.0, 1.0, 5.0, 10.0, 15.0, 20.0, 30.0, 40.0, 60.0, 80.0, 100.0, 160.0, 200.0, 300.0],
        "gamma":               [0.0],
        "lr":                  [1e-3, 5e-4],
        "weight_decay":        [1e-4, 1e-5],
        "encoder_dropout":     [0.0, 0.1],
        "decoder_batchnorm":   [False, True],
        "atac_arch":           atac_arch_options,
        "atac_likelihood":     likelihood_per_mod["atac"],
    }

    input_dims = {"atac": atac_train.n_vars}

    results: List[Any] = []
    best_score = float("inf")
    best_cfg = None
    best_model = None

    for cfg_id, hp in enumerate(
        iter_hparam_configs(search_space, max_configs=max_configs, seed=seed),
        start=1,
    ):
        res = train_single_config(
            config_id=cfg_id,
            hparams=hp,
            mod_arch_space=mod_arch_space,
            modalities=modalities,
            input_dims=input_dims,
            likelihood_per_mod={"atac": likelihood_per_mod["atac"]},
            adata_train=adata_train,
            adata_val=adata_val,
            base_train_cfg=base_train_cfg,
            layer=layer,
            X_key=X_key,
            celltype_key=None,
            device=device,
            multimodal_eval=False,
        )
        results.append(res)

        score = res.metrics["composite_score"]
        if score < best_score:
            best_score = score
            best_model = res
            best_cfg = hp
            print(f"--> New best ATAC-only config (id={cfg_id}) with score={score:.3f}")

    df = results_to_dataframe(results)
    return df, best_model, best_cfg
```
