## Supplemental Code for "Unifying multimodal single-cell data with a mixture-of-experts *β*-variational autoencoder framework": interpretability.py

```
# univi/interpretability.py
from __future__ import annotations

from dataclasses import dataclass
from typing import Dict, Any, Optional, Tuple, List, Sequence, Literal, Union

import numpy as np
import torch
from torch import nn
import torch.nn.functional as F


# -----------------------------
# Utilities: dense conversion
# -----------------------------
def _to_tensor(X, device: str, dtype=torch.float32) -> torch.Tensor:
    if torch.is_tensor(X):
        return X.to(device=device, dtype=dtype)
    X = np.asarray(X)
    return torch.as_tensor(X, device=device, dtype=dtype)

@dataclass
class TokenMap:
    """Mapping from fused token positions -> modality + feature identity."""
    modalities: List[str]
    # token slices in the concatenated sequence (excluding global CLS if present)
    slices: Dict[str, Tuple[int, int]]
    # modality-specific meta returned by tokenizer
    meta: Dict[str, Any]
    # whether a global CLS token was prepended
    has_global_cls: bool


def select_attn_matrix(
    attn_all: List[torch.Tensor],
    *,
    layer: int = -1,
    reduce_heads: Literal["mean", "none"] = "mean",
    reduce_batch: Literal["mean", "none"] = "mean",
) -> torch.Tensor:
    """
    attn_all[layer] is either (B,T,T) if heads averaged, or (B,H,T,T) if not.
    Returns:
      - (T,T) if reduce_batch="mean"
      - (B,T,T) if reduce_batch="none"
    """
    attn = attn_all[layer]

    if attn.dim() == 4:  # (B,H,T,T)
        if reduce_heads == "mean":
            attn = attn.mean(dim=1)  # (B,T,T)
        else:
            raise ValueError("reduce_heads='none' not supported here (pick a head upstream or add support).")

    if attn.dim() != 3:
        raise ValueError(f"Expected attn as (B,T,T) after head reduce; got {tuple(attn.shape)}")

    if reduce_batch == "mean":
        return attn.mean(dim=0)  # (T,T)
    return attn  # (B,T,T)

# -----------------------------
# Getting tokens + attention from your fused encoder
# -----------------------------
@torch.no_grad()
def fused_encode_with_meta_and_attn(
    model: nn.Module,
    x_dict: Dict[str, torch.Tensor],
    *,
    return_attn: bool = True,
    attn_average_heads: bool = True,
) -> Tuple[torch.Tensor, torch.Tensor, TokenMap, Optional[List[torch.Tensor]]]:
    if not hasattr(model, "fused_encoder") or model.fused_encoder is None:
        raise ValueError("Model has no fused_encoder. Set fused_encoder_type='multimodal_transformer'.")

    fused = model.fused_encoder

    # Fused encoder now returns one of:
    #   (mu, logvar)
    #   (mu, logvar, meta)
    #   (mu, logvar, attn_all)
    #   (mu, logvar, attn_all, meta)
    ret = fused(
        x_dict,
        return_token_meta=True,
        return_attn=return_attn,
        attn_average_heads=attn_average_heads,
    )

    if return_attn:
        mu, logvar, attn_all, meta = ret
    else:
        mu, logvar, meta = ret
        attn_all = None

    tokmap = TokenMap(
        modalities=list(meta["modalities"]),
        slices=dict(meta["slices"]),  # slices exclude CLS; you handle shift elsewhere
        meta={m: meta.get(m, {}) for m in meta["modalities"]},
        has_global_cls=bool(getattr(fused, "use_global_cls", False)),
    )
    return mu, logvar, tokmap, attn_all


# -----------------------------
# Map tokens back to feature names
# -----------------------------
def token_to_feature_names(
    tokmap: TokenMap,
    *,
    var_names_by_mod: Dict[str, Sequence[str]],
    tokenizer_mode_by_mod: Dict[str, str],
    patch_size_by_mod: Optional[Dict[str, int]] = None,
) -> Dict[str, List[str]]:
    """
    Returns for each modality: a list of length T_mod giving a human-readable
    feature name per token position (token order as produced by tokenizer).
    """
    out: Dict[str, List[str]] = {}

    for m in tokmap.modalities:
        mode = tokenizer_mode_by_mod[m]
        var_names = list(map(str, var_names_by_mod[m]))

        if mode in ("topk_scalar", "topk_channels"):
            topk_idx = tokmap.meta[m].get("topk_idx", None)
            if topk_idx is None:
                raise ValueError(f"Missing topk_idx for modality {m}. Ensure return_token_meta=True.")
            # topk_idx is (B,K). For naming tokens, pick a canonical naming:
            # we will name by index position; actual per-cell chosen features vary.
            # We'll return placeholders; per-cell mapping happens later.
            K = int(topk_idx.shape[1])
            out[m] = [f"{m}:topk_token_{i}" for i in range(K)]
        elif mode == "patch":
            if patch_size_by_mod is None or m not in patch_size_by_mod:
                raise ValueError(f"patch_size_by_mod is required for patch token naming for modality {m}")
            P = int(patch_size_by_mod[m])
            T = (len(var_names) + P - 1) // P
            names = []
            for t in range(T):
                a = t * P
                b = min((t + 1) * P, len(var_names))
                names.append(f"{m}:patch[{t}] {var_names[a]}..{var_names[b-1]}")
            out[m] = names
        else:
            raise ValueError(f"Unknown tokenizer mode {mode!r} for modality {m}")

    return out


# -----------------------------
# Feature-level attribution (Integrated Gradients or grad×input)
# -----------------------------
def integrated_gradients(
    f,                          # function mapping inputs -> scalar
    x: torch.Tensor,            # (B,F)
    baseline: Optional[torch.Tensor] = None,
    steps: int = 32,
) -> torch.Tensor:
    """
    Standard IG: returns attribution of same shape as x.
    """
    if baseline is None:
        baseline = torch.zeros_like(x)

    # interpolate
    alphas = torch.linspace(0.0, 1.0, steps=steps, device=x.device, dtype=x.dtype).view(steps, 1, 1)
    x0 = baseline.unsqueeze(0)
    x1 = x.unsqueeze(0)
    xs = x0 + alphas * (x1 - x0)  # (S,B,F)

    grads = []
    for s in range(steps):
        xi = xs[s].detach().clone().requires_grad_(True)
        y = f(xi)
        if y.dim() != 0:
            y = y.sum()
        g = torch.autograd.grad(y, xi, retain_graph=False, create_graph=False)[0]
        grads.append(g)

    grads = torch.stack(grads, dim=0)          # (S,B,F)
    avg_grads = grads.mean(dim=0)              # (B,F)
    return (x - baseline) * avg_grads          # (B,F)


def feature_importance_for_head(
    model: nn.Module,
    x_dict: Dict[str, torch.Tensor],
    *,
    head_name: str,
    class_index: int,
    method: Literal["grad_x_input", "ig"] = "ig",
    ig_steps: int = 32,
    per_cell_topk: int = 50,
    feature_names_by_mod: Optional[Dict[str, Sequence[str]]] = None,
) -> Dict[str, Any]:
    """
    Computes feature attributions for predicting `head_name` class `class_index`
    using the fused latent (best for multimodal interpretability).

    Returns:
      {modality: {names, scores, indices}} with top features.
    """
    device = next(model.parameters()).device
    model.eval()

    # Clone inputs and ensure grad
    x_in = {m: x_dict[m].detach().clone().to(device=device) for m in x_dict.keys()}
    for m in x_in:
        x_in[m].requires_grad_(True)

    # Define scalar function: logit(class_index) from the fused representation
    def scalar_from_inputs(x_mod: torch.Tensor, mod: str) -> torch.Tensor:
        # This wrapper is used for IG per modality, so we rebuild x_dict each time.
        xd = {k: (x_mod if k == mod else x_in[k]) for k in x_in.keys()}

        # You can either:
        # (A) call model.predict_heads on fused z (preferred), or
        # (B) call model(...) and read out head logits.
        #
        # Easiest robust path: call model(x_dict) and extract head logits.
        out = model(xd, epoch=0, y=None)
        if "head_logits" in out and head_name in out["head_logits"]:
            logits = out["head_logits"][head_name]  # (B,C)
        else:
            # fallback: use helper if available
            probs_or_logits = model.predict_heads(xd, return_probs=False)
            logits = probs_or_logits[head_name]

        return logits[:, int(class_index)].sum()

    results: Dict[str, Any] = {}

    for mod in x_in.keys():
        if method == "grad_x_input":
            out = model(x_in, epoch=0, y=None)
            if "head_logits" in out and head_name in out["head_logits"]:
                logits = out["head_logits"][head_name]
            else:
                logits = model.predict_heads(x_in, return_probs=False)[head_name]

            score = logits[:, int(class_index)].sum()
            grads = torch.autograd.grad(score, x_in[mod], retain_graph=True)[0]
            attr = grads * x_in[mod]  # (B,F)

        else:  # IG
            attr = integrated_gradients(
                lambda xm: scalar_from_inputs(xm, mod),
                x_in[mod],
                baseline=torch.zeros_like(x_in[mod]),
                steps=int(ig_steps),
            )

        # Aggregate across cells (mean abs is usually a good default)
        attr_agg = attr.detach().abs().mean(dim=0)  # (F,)

        k = min(int(per_cell_topk), int(attr_agg.numel()))
        vals, idx = torch.topk(attr_agg, k=k, largest=True, sorted=True)

        names = None
        if feature_names_by_mod is not None and mod in feature_names_by_mod:
            vn = list(map(str, feature_names_by_mod[mod]))
            names = [vn[i] for i in idx.detach().cpu().tolist()]

        results[mod] = {
            "indices": idx.detach().cpu().numpy(),
            "scores": vals.detach().cpu().numpy(),
            "names": names,
        }

    return results


# -----------------------------
# Cross-modal token interaction from attention
# -----------------------------
def top_cross_modal_attention_pairs(
    attn: torch.Tensor,
    tokmap: TokenMap,
    *,
    mod_a: str,
    mod_b: str,
    top_n: int = 50,
    reduce: Literal["mean", "max"] = "mean",
) -> List[Tuple[str, str, float]]:
    """
    Takes one attention matrix (T,T) and returns strongest A->B token pairs.
    `attn` should already correspond to the fused token sequence *including* global CLS if present.
    """
    if attn.dim() != 2:
        raise ValueError(f"Expected attn as (T,T), got {tuple(attn.shape)}")

    # compute slices in the attention matrix
    # If global CLS exists, token indices shift by +1 for everything else.
    shift = 1 if tokmap.has_global_cls else 0
    a0, a1 = tokmap.slices[mod_a]
    b0, b1 = tokmap.slices[mod_b]
    a0 += shift; a1 += shift
    b0 += shift; b1 += shift

    sub = attn[a0:a1, b0:b1]  # (Ta,Tb)

    # flatten and take top pairs
    flat = sub.reshape(-1)
    k = min(int(top_n), int(flat.numel()))
    vals, idx = torch.topk(flat, k=k, largest=True, sorted=True)

    Ta = a1 - a0
    Tb = b1 - b0

    pairs: List[Tuple[str, str, float]] = []
    for v, ii in zip(vals.detach().cpu().tolist(), idx.detach().cpu().tolist()):
        ia = ii // Tb
        ib = ii % Tb
        pairs.append((f"{mod_a}:token_{ia}", f"{mod_b}:token_{ib}", float(v)))

    return pairs

@torch.no_grad()
def top_cross_modal_feature_pairs_from_attn(
    attn_all: List[torch.Tensor],
    tokmap: TokenMap,
    *,
    mod_a: str,
    mod_b: str,
    var_names_by_mod: Dict[str, Sequence[str]],
    tokenizer_mode_by_mod: Dict[str, str],
    layer: int = -1,
    top_pairs_per_cell: int = 50,
    top_n: int = 100,
) -> List[Tuple[str, str, float]]:
    """
    Returns strongest (feature_a -> feature_b) pairs aggregated across the batch.

    Works best when tokenizer mode for both modalities is topk_* (because tokens map
    to feature indices via per-cell topk_idx).
    """
    mode_a = tokenizer_mode_by_mod[mod_a]
    mode_b = tokenizer_mode_by_mod[mod_b]
    if mode_a not in ("topk_scalar", "topk_channels") or mode_b not in ("topk_scalar", "topk_channels"):
        raise ValueError("This helper currently supports only topk_* tokenizers for both modalities.")

    A = attn_all[layer]
    if A.dim() == 4:               # (B,H,T,T)
        A = A.mean(dim=1)          # avg heads -> (B,T,T)
    if A.dim() != 3:
        raise ValueError(f"Expected attn as (B,T,T); got {tuple(A.shape)}")

    Bsz, Ttot, _ = A.shape

    shift = 1 if tokmap.has_global_cls else 0
    a0, a1 = tokmap.slices[mod_a]
    b0, b1 = tokmap.slices[mod_b]
    a0 += shift; a1 += shift
    b0 += shift; b1 += shift

    Ta = a1 - a0
    Tb = b1 - b0

    sub = A[:, a0:a1, b0:b1]                 # (B,Ta,Tb)
    flat = sub.reshape(Bsz, Ta * Tb)         # (B, Ta*Tb)

    k = min(int(top_pairs_per_cell), int(Ta * Tb))
    vals, idx = torch.topk(flat, k=k, dim=1, largest=True, sorted=True)  # (B,k)

    ia = idx // Tb  # (B,k) token index within A-slice
    ib = idx % Tb   # (B,k) token index within B-slice

    topk_a = tokmap.meta[mod_a].get("topk_idx", None)
    topk_b = tokmap.meta[mod_b].get("topk_idx", None)
    if topk_a is None or topk_b is None:
        raise ValueError("Missing topk_idx in tokmap.meta. Ensure return_token_meta=True.")

    # Map token positions -> feature indices (per-cell)
    fa = torch.gather(topk_a.to(device=ia.device), 1, ia)  # (B,k)
    fb = torch.gather(topk_b.to(device=ib.device), 1, ib)  # (B,k)

    # Aggregate into a python dict (sparse aggregation)
    scores: Dict[Tuple[int, int], float] = {}
    fa_np = fa.detach().cpu().numpy()
    fb_np = fb.detach().cpu().numpy()
    v_np  = vals.detach().cpu().numpy()

    for b in range(Bsz):
        for j in range(k):
            key = (int(fa_np[b, j]), int(fb_np[b, j]))
            scores[key] = scores.get(key, 0.0) + float(v_np[b, j])

    # Convert to top list with names
    a_names = list(map(str, var_names_by_mod[mod_a]))
    b_names = list(map(str, var_names_by_mod[mod_b]))

    items = sorted(scores.items(), key=lambda kv: kv[1], reverse=True)[: int(top_n)]
    out: List[Tuple[str, str, float]] = []
    for (ia_feat, ib_feat), s in items:
        out.append((a_names[ia_feat], b_names[ib_feat], float(s)))
    return out
```
