## Supplemental Code for "Unifying multimodal single-cell data with a mixture-of-experts *β*-variational autoencoder framework": matching.py

```
# univi/matching.py

import warnings
import numpy as np
from sklearn.metrics import pairwise_distances
from sklearn.neighbors import NearestNeighbors
from scipy.optimize import linear_sum_assignment
from typing import Optional, Dict


def _subsample_indices(n: int, max_cells: int, rng: np.random.Generator) -> np.ndarray:
    """
    Helper to subsample up to max_cells indices from range(n) without replacement.
    """
    idx_full = np.arange(n)
    if n <= max_cells:
        return idx_full
    return rng.choice(idx_full, size=max_cells, replace=False)


# ---------------------------------------------------------------------------
# 1. Basic bipartite matching (Hungarian) in a shared embedding
# ---------------------------------------------------------------------------

def bipartite_match_adata(
    adata_A,
    adata_B,
    emb_key: str = "X_pca",
    metric: str = "euclidean",
    max_cells: int = 20000,
    random_state: int = 0,
):
    """
    Bipartite matching between cells in adata_A and adata_B based on a shared embedding.

    This is the basic "Hungarian in latent space" matcher. It assumes that both
    adata_A and adata_B have a *comparable* embedding in .obsm[emb_key], e.g.:

        - both are in the same PCA space, or
        - both are in a shared latent space (CCA, UniVI encoder, etc.)
    """
    rng = np.random.default_rng(random_state)

    XA = np.asarray(adata_A.obsm[emb_key])
    XB = np.asarray(adata_B.obsm[emb_key])

    na, nb = XA.shape[0], XB.shape[0]
    n = min(na, nb, max_cells)

    idx_A = _subsample_indices(na, n, rng)
    idx_B = _subsample_indices(nb, n, rng)

    XA_sub = XA[idx_A]
    XB_sub = XB[idx_B]

    # cost matrix
    D = pairwise_distances(XA_sub, XB_sub, metric=metric)

    # Hungarian algorithm (min-cost)
    row_ind, col_ind = linear_sum_assignment(D)

    matched_A = idx_A[row_ind]
    matched_B = idx_B[col_ind]

    return matched_A, matched_B


# ---------------------------------------------------------------------------
# 2. Stratified bipartite matching (per group / cell type)
# ---------------------------------------------------------------------------

def stratified_bipartite_match_adata(
    adata_A,
    adata_B,
    group_key_A: str,
    group_key_B: Optional[str] = None,
    group_map: Optional[Dict] = None,
    emb_key: str = "X_pca",
    metric: str = "euclidean",
    max_cells_per_group: int = 20000,
    random_state: int = 0,
    shuffle: bool = True,
):
    """
    Per-group (e.g. per-celltype) bipartite matching in a shared embedding.

    This wraps `bipartite_match_adata` but runs it separately within each group,
    then concatenates the matches.
    """
    rng = np.random.default_rng(random_state)

    if group_key_B is None:
        group_key_B = group_key_A

    if group_key_A not in adata_A.obs.columns:
        raise KeyError(f"{group_key_A!r} not found in adata_A.obs")
    if group_key_B not in adata_B.obs.columns:
        raise KeyError(f"{group_key_B!r} not found in adata_B.obs")

    labels_A = adata_A.obs[group_key_A].astype(str).to_numpy()
    labels_B = adata_B.obs[group_key_B].astype(str).to_numpy()

    unique_A = np.unique(labels_A)

    all_matched_A = []
    all_matched_B = []
    group_counts: Dict[str, int] = {}

    for gA in unique_A:
        # Determine which label in B we should match to
        if group_map is not None:
            gB = group_map.get(gA, None)
            if gB is None:
                # group not mapped; skip
                continue
        else:
            gB = gA

        idx_A_g = np.where(labels_A == gA)[0]
        idx_B_g = np.where(labels_B == gB)[0]

        if (idx_A_g.size == 0) or (idx_B_g.size == 0):
            continue

        # Build small views
        adata_A_g = adata_A[idx_A_g]
        adata_B_g = adata_B[idx_B_g]

        # n per group for bipartite matching
        n_grp = min(idx_A_g.size, idx_B_g.size, max_cells_per_group)
        if n_grp == 0:
            continue

        mA_local, mB_local = bipartite_match_adata(
            adata_A_g,
            adata_B_g,
            emb_key=emb_key,
            metric=metric,
            max_cells=n_grp,
            random_state=random_state,
        )

        if mA_local.size == 0:
            continue

        mA = idx_A_g[mA_local]
        mB = idx_B_g[mB_local]

        all_matched_A.append(mA)
        all_matched_B.append(mB)
        group_counts[gA] = mA.size

    if not all_matched_A:
        raise RuntimeError("No stratified matches were found for any group.")

    matched_A = np.concatenate(all_matched_A)
    matched_B = np.concatenate(all_matched_B)

    if shuffle:
        perm = rng.permutation(matched_A.size)
        matched_A = matched_A[perm]
        matched_B = matched_B[perm]

    return matched_A, matched_B, group_counts


# ---------------------------------------------------------------------------
# 3. Mutual Nearest Neighbor (MNN) anchors
# ---------------------------------------------------------------------------

def mnn_anchors_adata(
    adata_A,
    adata_B,
    emb_key: str = "X_pca",
    k: int = 20,
    max_cells: int = 20000,
    random_state: int = 0,
):
    """
    Mutual Nearest Neighbor (MNN) anchors between adata_A and adata_B.
    """
    rng = np.random.default_rng(random_state)

    XA = np.asarray(adata_A.obsm[emb_key])
    XB = np.asarray(adata_B.obsm[emb_key])

    na, nb = XA.shape[0], XB.shape[0]

    idx_A = _subsample_indices(na, max_cells, rng)
    idx_B = _subsample_indices(nb, max_cells, rng)

    XA_sub = XA[idx_A]
    XB_sub = XB[idx_B]

    k_A = min(k, idx_B.size)
    k_B = min(k, idx_A.size)

    # A -> B neighbors
    nn_B = NearestNeighbors(n_neighbors=k_A)
    nn_B.fit(XB_sub)
    dist_A2B, ind_A2B = nn_B.kneighbors(XA_sub, return_distance=True)

    # B -> A neighbors
    nn_A = NearestNeighbors(n_neighbors=k_B)
    nn_A.fit(XA_sub)
    dist_B2A, ind_B2A = nn_A.kneighbors(XB_sub, return_distance=True)

    # Mutual neighbors
    anchors_A_list = []
    anchors_B_list = []

    # For quick membership tests: for each j, set of neighbors in A
    neighbors_B2A = [set(ind_B2A[j]) for j in range(idx_B.size)]

    for i in range(idx_A.size):
        for j_local in ind_A2B[i]:
            # Check mutuality
            if i in neighbors_B2A[j_local]:
                anchors_A_list.append(idx_A[i])
                anchors_B_list.append(idx_B[j_local])

    if not anchors_A_list:
        warnings.warn("mnn_anchors_adata: no mutual nearest neighbors found.")
        return np.array([], dtype=int), np.array([], dtype=int)

    anchors_A = np.array(anchors_A_list, dtype=int)
    anchors_B = np.array(anchors_B_list, dtype=int)

    return anchors_A, anchors_B


# ---------------------------------------------------------------------------
# 4. Cluster / cell-type centroid matching (for building group maps)
# ---------------------------------------------------------------------------

def cluster_centroid_matching_adata(
    adata_A,
    adata_B,
    group_key_A: str,
    group_key_B: Optional[str] = None,
    emb_key: str = "X_pca",
    metric: str = "euclidean",
):
    """
    Match cluster / cell-type centroids across datasets via Hungarian.
    """
    if group_key_B is None:
        group_key_B = group_key_A

    if group_key_A not in adata_A.obs.columns:
        raise KeyError(f"{group_key_A!r} not found in adata_A.obs")
    if group_key_B not in adata_B.obs.columns:
        raise KeyError(f"{group_key_B!r} not found in adata_B.obs")

    labels_A = adata_A.obs[group_key_A].astype(str).to_numpy()
    labels_B = adata_B.obs[group_key_B].astype(str).to_numpy()

    XA = np.asarray(adata_A.obsm[emb_key])
    XB = np.asarray(adata_B.obsm[emb_key])

    groups_A = np.unique(labels_A)
    groups_B = np.unique(labels_B)

    # Compute centroids
    centroids_A = []
    for g in groups_A:
        idx = np.where(labels_A == g)[0]
        centroids_A.append(XA[idx].mean(axis=0))
    centroids_A = np.vstack(centroids_A)

    centroids_B = []
    for g in groups_B:
        idx = np.where(labels_B == g)[0]
        centroids_B.append(XB[idx].mean(axis=0))
    centroids_B = np.vstack(centroids_B)

    # Cost between centroids
    D = pairwise_distances(centroids_A, centroids_B, metric=metric)
    row_ind, col_ind = linear_sum_assignment(D)

    group_map: Dict[str, str] = {}
    for i, j in zip(row_ind, col_ind):
        gA = groups_A[i]
        gB = groups_B[j]
        group_map[gA] = gB

    return group_map


# ---------------------------------------------------------------------------
# 5. Gromov–Wasserstein OT-based anchors (optional; requires POT)
# ---------------------------------------------------------------------------

def gw_ot_anchors_adata(
    adata_A,
    adata_B,
    emb_key: str = "X_pca",
    max_cells: int = 3000,
    random_state: int = 0,
    normalize_distances: bool = True,
):
    """
    Geometry-aware anchors via Gromov–Wasserstein optimal transport.
    """
    try:
        import ot  # type: ignore
    except ImportError as e:
        raise ImportError(
            "gw_ot_anchors_adata requires the 'pot' package. "
            "Install with: pip install pot"
        ) from e

    rng = np.random.default_rng(random_state)

    XA = np.asarray(adata_A.obsm[emb_key])
    XB = np.asarray(adata_B.obsm[emb_key])

    na, nb = XA.shape[0], XB.shape[0]
    idx_A = _subsample_indices(na, max_cells, rng)
    idx_B = _subsample_indices(nb, max_cells, rng)

    XA_sub = XA[idx_A]
    XB_sub = XB[idx_B]

    # Distance matrices within each dataset
    DA = pairwise_distances(XA_sub, XA_sub, metric="euclidean")
    DB = pairwise_distances(XB_sub, XB_sub, metric="euclidean")

    if normalize_distances:
        if DA.max() > 0:
            DA = DA / DA.max()
        if DB.max() > 0:
            DB = DB / DB.max()

    # Uniform weights
    p = np.ones(DA.shape[0]) / DA.shape[0]
    q = np.ones(DB.shape[0]) / DB.shape[0]

    # Compute GW coupling
    T = ot.gromov.gromov_wasserstein(
        DA, DB, p, q, loss_fun="square_loss", verbose=False
    )

    # For each i in A, pick the j in B with maximum coupling mass
    anchors_A_list = []
    anchors_B_list = []
    for i in range(T.shape[0]):
        j = int(np.argmax(T[i]))
        anchors_A_list.append(idx_A[i])
        anchors_B_list.append(idx_B[j])

    anchors_A = np.array(anchors_A_list, dtype=int)
    anchors_B = np.array(anchors_B_list, dtype=int)

    return anchors_A, anchors_B


# ---------------------------------------------------------------------------
# 6. Group latent statistics (for distribution-level alignment)
# ---------------------------------------------------------------------------

def group_latent_stats_adata(
    adata,
    group_key: str,
    emb_key: str = "X_pca",
):
    """
    Compute simple group-wise latent statistics (mean, covariance) to
    support distribution-level alignment strategies.
    """
    if group_key not in adata.obs.columns:
        raise KeyError(f"{group_key!r} not found in adata.obs")

    labels = adata.obs[group_key].astype(str).to_numpy()
    X = np.asarray(adata.obsm[emb_key])

    groups = np.unique(labels)
    stats = {}

    for g in groups:
        idx = np.where(labels == g)[0]
        if idx.size == 0:
            continue
        Xg = X[idx]
        mu = Xg.mean(axis=0)
        # rowvar=False so that columns are variables
        cov = np.cov(Xg, rowvar=False)
        stats[g] = {
            "mean": mu,
            "cov": cov,
            "n": idx.size,
        }

    return stats
```
