## Supplemental Code for "Unifying multimodal single-cell data with a mixture-of-experts *β*-variational autoencoder framework": __init__.py

```
# univi/models/__init__.py
from __future__ import annotations

from .univi import UniVIMultiModalVAE
from .transformer import TransformerEncoder
from .tokenizers import build_tokenizer

__all__ = ["UniVIMultiModalVAE", "TransformerEncoder", "build_tokenizer"]
```
