## Supplemental Code for "Unifying multimodal single-cell data with a mixture-of-experts *β*-variational autoencoder framework": decoders.py

```
from __future__ import annotations

from dataclasses import dataclass
from typing import Any, Dict, List

import torch
from torch import nn
import torch.nn.functional as F

from .mlp import build_mlp


@dataclass
class DecoderConfig:
    """Generic configuration for feed-forward decoders."""
    output_dim: int
    hidden_dims: List[int]
    dropout: float = 0.0
    batchnorm: bool = False


# ---------------------------------------------------------------------
# Continuous decoders
# ---------------------------------------------------------------------
class GaussianDecoder(nn.Module):
    """z -> mean reconstruction (use with MSE/Gaussian losses)."""

    def __init__(self, cfg: DecoderConfig, latent_dim: int):
        super().__init__()
        self.cfg = cfg
        self.net = build_mlp(
            in_dim=latent_dim,
            hidden_dims=cfg.hidden_dims,
            out_dim=cfg.output_dim,
            dropout=cfg.dropout,
            batchnorm=cfg.batchnorm,
        )

    def forward(self, z: torch.Tensor) -> torch.Tensor:
        return self.net(z)


class GaussianDiagDecoder(nn.Module):
    """z -> {'mean','logvar'} for full diagonal Gaussian likelihoods."""

    def __init__(self, cfg: DecoderConfig, latent_dim: int):
        super().__init__()
        self.cfg = cfg
        self.backbone = build_mlp(
            in_dim=latent_dim,
            hidden_dims=cfg.hidden_dims,
            out_dim=2 * cfg.output_dim,
            dropout=cfg.dropout,
            batchnorm=cfg.batchnorm,
        )

    def forward(self, z: torch.Tensor) -> Dict[str, torch.Tensor]:
        out = self.backbone(z)
        mean, logvar = out.chunk(2, dim=-1)
        return {"mean": mean, "logvar": logvar}


class BetaDecoder(nn.Module):
    """
    z -> parameters for Beta likelihoods on continuous targets in (0,1).

    Useful for fraction-valued features (e.g., methylation/accessibility fractions)
    when you do NOT explicitly model coverage counts.

    Returns:
      {
        'alpha': alpha,
        'beta': beta,
        'mu_logits': mu_logits,
        'log_conc': log_conc,
        'mu': mu,         # convenience
        'conc': conc,     # convenience
      }

    Parameterization:
      mu   = sigmoid(mu_logits) in (0,1)
      conc = softplus(log_conc_raw) + min_conc
      alpha = mu * conc
      beta  = (1-mu) * conc
    """

    def __init__(
        self,
        cfg: DecoderConfig,
        latent_dim: int,
        eps: float = 1e-8,
        min_conc: float = 1e-4,
    ):
        super().__init__()
        self.cfg = cfg
        self.eps = float(eps)
        self.min_conc = float(min_conc)

        self.backbone = build_mlp(
            in_dim=latent_dim,
            hidden_dims=cfg.hidden_dims,
            out_dim=2 * cfg.output_dim,
            dropout=cfg.dropout,
            batchnorm=cfg.batchnorm,
        )

    def forward(self, z: torch.Tensor) -> Dict[str, torch.Tensor]:
        out = self.backbone(z)
        mu_logits, log_conc_raw = out.chunk(2, dim=-1)

        mu = torch.sigmoid(mu_logits).clamp(self.eps, 1.0 - self.eps)
        conc = F.softplus(log_conc_raw) + self.min_conc

        alpha = (mu * conc).clamp_min(self.eps)
        beta = ((1.0 - mu) * conc).clamp_min(self.eps)

        return {
            "alpha": alpha,
            "beta": beta,
            "mu_logits": mu_logits,
            "log_conc": torch.log(conc + self.eps),
            "mu": mu,
            "conc": conc,
        }


# ---------------------------------------------------------------------
# Bernoulli / count / proportion-count decoders
# ---------------------------------------------------------------------
class BernoulliDecoder(nn.Module):
    """z -> {'logits'} for Bernoulli likelihoods."""

    def __init__(self, cfg: DecoderConfig, latent_dim: int):
        super().__init__()
        self.cfg = cfg
        self.net = build_mlp(
            in_dim=latent_dim,
            hidden_dims=cfg.hidden_dims,
            out_dim=cfg.output_dim,
            dropout=cfg.dropout,
            batchnorm=cfg.batchnorm,
        )

    def forward(self, z: torch.Tensor) -> Dict[str, torch.Tensor]:
        logits = self.net(z)
        return {"logits": logits}


class BinomialDecoder(nn.Module):
    """
    z -> {'logits'} for Binomial likelihoods.

    Intended for targets like:
      successes m ~ Binomial(total_count=n, p)
    where `n` (coverage / trials) is provided to the loss function, not the decoder.

    Decoder returns logits for p.
    """

    def __init__(self, cfg: DecoderConfig, latent_dim: int):
        super().__init__()
        self.cfg = cfg
        self.net = build_mlp(
            in_dim=latent_dim,
            hidden_dims=cfg.hidden_dims,
            out_dim=cfg.output_dim,
            dropout=cfg.dropout,
            batchnorm=cfg.batchnorm,
        )

    def forward(self, z: torch.Tensor) -> Dict[str, torch.Tensor]:
        logits = self.net(z)
        return {"logits": logits}


class BetaBinomialDecoder(nn.Module):
    """
    z -> parameters for Beta-Binomial likelihoods (overdispersed Binomial).

    Intended for count-with-coverage observations such as methylome or
    proportion-like assays with sampling noise and extra-binomial variance.

    Returns:
      {
        'mu_logits': mu_logits,
        'log_conc': log_conc,
        'mu': mu,         # convenience
        'conc': conc,     # convenience
        'alpha': alpha,   # convenience (for losses that prefer alpha/beta)
        'beta': beta,     # convenience
      }

    Parameterization:
      mu   = sigmoid(mu_logits) in (0,1)
      conc = softplus(log_conc_raw) + min_conc
      alpha = mu * conc
      beta  = (1-mu) * conc
    """

    def __init__(
        self,
        cfg: DecoderConfig,
        latent_dim: int,
        eps: float = 1e-8,
        min_conc: float = 1e-4,
    ):
        super().__init__()
        self.cfg = cfg
        self.eps = float(eps)
        self.min_conc = float(min_conc)

        self.backbone = build_mlp(
            in_dim=latent_dim,
            hidden_dims=cfg.hidden_dims,
            out_dim=2 * cfg.output_dim,
            dropout=cfg.dropout,
            batchnorm=cfg.batchnorm,
        )

    def forward(self, z: torch.Tensor) -> Dict[str, torch.Tensor]:
        out = self.backbone(z)
        mu_logits, log_conc_raw = out.chunk(2, dim=-1)

        mu = torch.sigmoid(mu_logits).clamp(self.eps, 1.0 - self.eps)
        conc = F.softplus(log_conc_raw) + self.min_conc

        alpha = (mu * conc).clamp_min(self.eps)
        beta = ((1.0 - mu) * conc).clamp_min(self.eps)

        return {
            "mu_logits": mu_logits,
            "log_conc": torch.log(conc + self.eps),
            "mu": mu,
            "conc": conc,
            "alpha": alpha,
            "beta": beta,
        }


class PoissonDecoder(nn.Module):
    """z -> {'log_rate','rate'} for Poisson likelihoods."""

    def __init__(self, cfg: DecoderConfig, latent_dim: int):
        super().__init__()
        self.cfg = cfg
        self.net = build_mlp(
            in_dim=latent_dim,
            hidden_dims=cfg.hidden_dims,
            out_dim=cfg.output_dim,
            dropout=cfg.dropout,
            batchnorm=cfg.batchnorm,
        )

    def forward(self, z: torch.Tensor) -> Dict[str, torch.Tensor]:
        log_rate = self.net(z)
        rate = F.softplus(log_rate)
        return {"log_rate": log_rate, "rate": rate}


class NegativeBinomialDecoder(nn.Module):
    """z -> {'mu','log_theta'} (theta can be global or gene-wise)."""

    def __init__(
        self,
        cfg: DecoderConfig,
        latent_dim: int,
        dispersion: str = "gene",
        init_log_theta: float = 0.0,
        eps: float = 1e-8,
    ):
        super().__init__()
        self.cfg = cfg
        self.dispersion = dispersion
        self.eps = float(eps)

        self.mu_net = build_mlp(
            in_dim=latent_dim,
            hidden_dims=cfg.hidden_dims,
            out_dim=cfg.output_dim,
            dropout=cfg.dropout,
            batchnorm=cfg.batchnorm,
        )

        if dispersion == "global":
            self.log_theta = nn.Parameter(torch.full((1,), float(init_log_theta)))
        elif dispersion == "gene":
            self.log_theta = nn.Parameter(torch.full((cfg.output_dim,), float(init_log_theta)))
        else:
            raise ValueError("Unknown dispersion mode: %r (expected 'global' or 'gene')" % dispersion)

    def forward(self, z: torch.Tensor) -> Dict[str, torch.Tensor]:
        mu = F.softplus(self.mu_net(z)) + self.eps
        return {"mu": mu, "log_theta": self.log_theta}


class ZeroInflatedNegativeBinomialDecoder(nn.Module):
    """z -> {'mu','log_theta','logit_pi'}."""

    def __init__(
        self,
        cfg: DecoderConfig,
        latent_dim: int,
        dispersion: str = "gene",
        init_log_theta: float = 0.0,
        eps: float = 1e-8,
    ):
        super().__init__()
        self.cfg = cfg
        self.dispersion = dispersion
        self.eps = float(eps)

        self.backbone = build_mlp(
            in_dim=latent_dim,
            hidden_dims=cfg.hidden_dims,
            out_dim=2 * cfg.output_dim,
            dropout=cfg.dropout,
            batchnorm=cfg.batchnorm,
        )

        if dispersion == "global":
            self.log_theta = nn.Parameter(torch.full((1,), float(init_log_theta)))
        elif dispersion == "gene":
            self.log_theta = nn.Parameter(torch.full((cfg.output_dim,), float(init_log_theta)))
        else:
            raise ValueError("Unknown dispersion mode: %r (expected 'global' or 'gene')" % dispersion)

    def forward(self, z: torch.Tensor) -> Dict[str, torch.Tensor]:
        out = self.backbone(z)
        mu_logits, logit_pi = out.chunk(2, dim=-1)
        mu = F.softplus(mu_logits) + self.eps
        return {"mu": mu, "log_theta": self.log_theta, "logit_pi": logit_pi}


# ---------------------------------------------------------------------
# Compositional / categorical decoders
# ---------------------------------------------------------------------
class LogisticNormalDecoder(nn.Module):
    """z -> {'logits','probs'} for compositions/toy probability vectors."""

    def __init__(self, cfg: DecoderConfig, latent_dim: int):
        super().__init__()
        self.cfg = cfg
        self.net = build_mlp(
            in_dim=latent_dim,
            hidden_dims=cfg.hidden_dims,
            out_dim=cfg.output_dim,
            dropout=cfg.dropout,
            batchnorm=cfg.batchnorm,
        )

    def forward(self, z: torch.Tensor) -> Dict[str, torch.Tensor]:
        logits = self.net(z)
        probs = F.softmax(logits, dim=-1)
        return {"logits": logits, "probs": probs}


class CategoricalDecoder(nn.Module):
    """z -> {'logits','probs'} for discrete labels."""

    def __init__(self, cfg: DecoderConfig, latent_dim: int):
        super().__init__()
        self.cfg = cfg
        self.net = build_mlp(
            in_dim=latent_dim,
            hidden_dims=cfg.hidden_dims,
            out_dim=cfg.output_dim,
            dropout=cfg.dropout,
            batchnorm=cfg.batchnorm,
        )

    def forward(self, z: torch.Tensor) -> Dict[str, torch.Tensor]:
        logits = self.net(z)
        probs = F.softmax(logits, dim=-1)
        return {"logits": logits, "probs": probs}


# ---------------------------------------------------------------------
# Registry / factory
# ---------------------------------------------------------------------
DECODER_REGISTRY = {
    # gaussian
    "gaussian": GaussianDecoder,
    "normal": GaussianDecoder,
    "gaussian_diag": GaussianDiagDecoder,

    # bounded continuous (fractions)
    "beta": BetaDecoder,

    # bernoulli / proportion-count models
    "bernoulli": BernoulliDecoder,
    "binomial": BinomialDecoder,
    "beta_binomial": BetaBinomialDecoder,
    "betabinomial": BetaBinomialDecoder,
    "bb": BetaBinomialDecoder,

    # count models
    "poisson": PoissonDecoder,
    "nb": NegativeBinomialDecoder,
    "negative_binomial": NegativeBinomialDecoder,
    "zinb": ZeroInflatedNegativeBinomialDecoder,
    "zero_inflated_negative_binomial": ZeroInflatedNegativeBinomialDecoder,

    # compositions / discrete
    "logistic_normal": LogisticNormalDecoder,
    "categorical": CategoricalDecoder,
    "cat": CategoricalDecoder,
    "ce": CategoricalDecoder,
    "cross_entropy": CategoricalDecoder,
}


def build_decoder(kind: str, cfg: DecoderConfig, latent_dim: int, **kwargs: Any) -> nn.Module:
    key = str(kind).lower()
    if key not in DECODER_REGISTRY:
        raise ValueError(
            "Unknown decoder kind: %r. Available: %s" % (kind, list(DECODER_REGISTRY.keys()))
        )
    cls = DECODER_REGISTRY[key]
    return cls(cfg=cfg, latent_dim=latent_dim, **kwargs)
```
