## Supplemental Code for "Unifying multimodal single-cell data with a mixture-of-experts *β*-variational autoencoder framework": encoders.py

```
# univi/models/encoders.py
from __future__ import annotations

from dataclasses import dataclass, replace
from typing import Dict, List, Optional, Sequence, Tuple, Union, Any, Mapping

import torch
from torch import nn
import torch.nn.functional as F

from ..config import UniVIConfig, ModalityConfig, TokenizerConfig, TransformerConfig as CFGTransformerConfig
from .mlp import build_mlp
from .transformer import TransformerEncoder, TransformerConfig as ModelTransformerConfig


# =============================================================================
# Small config helper
# =============================================================================

@dataclass
class EncoderConfig:
    input_dim: int
    hidden_dims: List[int]
    latent_dim: int
    dropout: float = 0.1
    batchnorm: bool = True


# =============================================================================
# Base encoders
# =============================================================================

class GaussianEncoder(nn.Module):
    """Base: x -> (mu, logvar) for a diagonal Gaussian."""
    def forward(self, x: torch.Tensor) -> Tuple[torch.Tensor, torch.Tensor]:
        raise NotImplementedError


class MLPGaussianEncoder(GaussianEncoder):
    """MLP encoder: x -> (mu, logvar) directly."""
    def __init__(self, cfg: EncoderConfig):
        super().__init__()
        self.cfg = cfg
        self.net = build_mlp(
            in_dim=int(cfg.input_dim),
            hidden_dims=list(cfg.hidden_dims),
            out_dim=2 * int(cfg.latent_dim),
            dropout=float(cfg.dropout),
            batchnorm=bool(cfg.batchnorm),
        )

    def forward(self, x: torch.Tensor) -> Tuple[torch.Tensor, torch.Tensor]:
        h = self.net(x)
        mu, logvar = torch.chunk(h, 2, dim=-1)
        return mu, logvar


# =============================================================================
# Compatibility: safe TransformerEncoder call
# =============================================================================

def _call_transformer_encoder(
    enc: nn.Module,
    tokens: torch.Tensor,
    *,
    key_padding_mask: Optional[torch.Tensor],
    return_attn: bool,
    attn_average_heads: bool = True,
    attn_bias: Optional[torch.Tensor] = None,
):
    """
    Call TransformerEncoder.forward in a backward-compatible way across
    versions that may or may not support:
      - attn_bias kwarg
      - attn_average_heads kwarg

    Returns
    -------
    If return_attn:
        (h, attn_all)
    else:
        h
    """
    # Build base kwargs
    kwargs: Dict[str, Any] = {
        "key_padding_mask": key_padding_mask,
        "return_attn": bool(return_attn),
    }

    # Only include attn_average_heads when returning attention
    if return_attn:
        kwargs["attn_average_heads"] = bool(attn_average_heads)

    # Only include attn_bias when provided (older encoders choke on the kwarg)
    if attn_bias is not None:
        kwargs["attn_bias"] = attn_bias

    # Try the "full" call first
    try:
        return enc(tokens, **kwargs)
    except TypeError:
        # Drop attn_bias first (most common incompatibility)
        if "attn_bias" in kwargs:
            kwargs.pop("attn_bias", None)
            try:
                return enc(tokens, **kwargs)
            except TypeError:
                pass

        # Then drop attn_average_heads (some versions don’t support it)
        if "attn_average_heads" in kwargs:
            kwargs.pop("attn_average_heads", None)
            try:
                return enc(tokens, **kwargs)
            except TypeError:
                pass

        # Final fallback: keep only the essentials
        kwargs_min: Dict[str, Any] = {
            "key_padding_mask": key_padding_mask,
            "return_attn": bool(return_attn),
        }
        return enc(tokens, **kwargs_min)


# =============================================================================
# Tokenization: vector -> tokens (+ optional embeddings / bias)
# =============================================================================

def _mlp(in_dim: int, out_dim: int, hidden: int = 128) -> nn.Module:
    return nn.Sequential(
        nn.LayerNorm(in_dim),
        nn.Linear(in_dim, hidden),
        nn.GELU(),
        nn.Linear(hidden, out_dim),
    )


class _VectorToTokens(nn.Module):
    """
    Turn a vector x (B, F) into tokens (B, T, D_in) and optional key_padding_mask.

    TokenizerConfig modes
    ---------------------
    - topk_scalar:
        Select top-k features per cell (by value), output tokens (B, K, 1)
    - topk_channels:
        Select top-k features per cell, output tokens (B, K, C) where C=len(channels)
        channels in {"value","rank","dropout"}:
          * value: raw x at selected indices
          * rank: rank01 among selected K tokens (0..1), per cell
          * dropout: indicator (value==0)
    - patch:
        Split contiguous features into patches of size P:
          tokens (B, T, P) or (B, T, patch_proj_dim) if patch_proj_dim is set

    add_cls_token:
        If True, prepend a learned CLS token embedding to tokens.

    NEW (optional)
    --------------
    - use_feature_embedding:
        Adds learned embedding for selected feature IDs (topk modes), or patch IDs (patch mode).
    - use_coord_embedding (ATAC):
        Adds chromosome embedding + coordinate MLP for selected feature coords (topk modes).
        Call set_feature_coords(chrom_ids, start, end) to attach coords.
    - token_proj_dim:
        If set (or implied by embeddings), project raw token channels/patches to token_proj_dim.

    Notes
    -----
    - Expects dense float input (B,F). If modality data is sparse, densify upstream.
    - key_padding_mask uses True = PAD/ignore (MultiheadAttention convention).
    """
    def __init__(self, *, input_dim: int, tok: TokenizerConfig):
        super().__init__()
        self.input_dim = int(input_dim)
        self.tok = tok

        mode = str(tok.mode).lower().strip()
        if mode not in ("topk_scalar", "topk_channels", "patch"):
            raise ValueError(f"Unknown tokenizer mode {tok.mode!r}")

        self.mode = mode
        self.add_cls_token = bool(getattr(tok, "add_cls_token", False))

        # ------------------------------------------------------------------
        # NEW options (all default to False/None -> no behavior change)
        # ------------------------------------------------------------------
        self.use_feature_emb = bool(getattr(tok, "use_feature_embedding", False))
        self.feature_emb_mode = str(getattr(tok, "feature_emb_mode", "add")).lower().strip()
        self.n_features = getattr(tok, "n_features", None)
        self.feature_emb_dim = getattr(tok, "feature_emb_dim", None)

        self.use_coord_emb = bool(getattr(tok, "use_coord_embedding", False))
        self.coord_mode = str(getattr(tok, "coord_mode", "midpoint")).lower().strip()
        self.coord_scale = float(getattr(tok, "coord_scale", 1e-6))
        self.coord_emb_dim = getattr(tok, "coord_emb_dim", None)
        self.n_chroms = getattr(tok, "n_chroms", None)
        self.coord_mlp_hidden = int(getattr(tok, "coord_mlp_hidden", 128))

        self.token_proj_dim = getattr(tok, "token_proj_dim", None)
        self._has_coords = False
        self.register_buffer("_chrom_ids", torch.empty(0, dtype=torch.long), persistent=False)
        self.register_buffer("_start", torch.empty(0), persistent=False)
        self.register_buffer("_end", torch.empty(0), persistent=False)

        # ------------------------------------------------------------------
        # Base token dims per mode (pre-projection)
        # ------------------------------------------------------------------
        if self.mode == "topk_scalar":
            self.n_tokens = int(tok.n_tokens)
            base_d = 1
            self.channels: Sequence[str] = ("value",)

        elif self.mode == "topk_channels":
            self.n_tokens = int(tok.n_tokens)
            ch = list(tok.channels)
            if not ch:
                raise ValueError("topk_channels requires tokenizer.channels non-empty")
            bad = [c for c in ch if c not in ("value", "rank", "dropout")]
            if bad:
                raise ValueError(f"topk_channels invalid channels: {bad}")
            self.channels = tuple(ch)
            base_d = len(self.channels)

        else:  # patch
            P = int(tok.patch_size)
            if P <= 0:
                raise ValueError("patch_size must be > 0")
            self.patch_size = P
            T = (self.input_dim + P - 1) // P
            self.n_tokens = int(T)

            proj_dim = tok.patch_proj_dim
            if proj_dim is None:
                self.patch_proj = None
                base_d = P
            else:
                proj_dim = int(proj_dim)
                if proj_dim <= 0:
                    raise ValueError("patch_proj_dim must be > 0 if set")
                self.patch_proj = nn.Linear(P, proj_dim)
                base_d = proj_dim

        # ------------------------------------------------------------------
        # Decide final token dim (d_in)
        # ------------------------------------------------------------------
        implied_proj: Optional[int] = None
        if self.token_proj_dim is not None:
            implied_proj = int(self.token_proj_dim)
        elif self.use_feature_emb or self.use_coord_emb:
            implied_proj = int(self.feature_emb_dim or self.coord_emb_dim or 64)

        self._d_in = int(implied_proj) if implied_proj is not None else int(base_d)

        # ------------------------------------------------------------------
        # Optional projection from base_d -> d_in
        # ------------------------------------------------------------------
        self.val_proj: Optional[nn.Module] = None
        if self._d_in != int(base_d):
            self.val_proj = _mlp(int(base_d), self._d_in, hidden=self.coord_mlp_hidden)

        # ------------------------------------------------------------------
        # NEW: feature ID embedding (topk) / patch ID embedding (patch)
        # ------------------------------------------------------------------
        self.id_emb: Optional[nn.Embedding] = None
        self.id_fuse: Optional[nn.Linear] = None
        if self.use_feature_emb:
            if self.mode in ("topk_scalar", "topk_channels"):
                if self.n_features is None or int(self.n_features) <= 0:
                    raise ValueError("use_feature_embedding=True for topk requires tok.n_features > 0")
                emb_dim = int(self.feature_emb_dim or self._d_in)
                self.id_emb = nn.Embedding(int(self.n_features), emb_dim)
                if self.feature_emb_mode == "concat":
                    self.id_fuse = nn.Linear(self._d_in + emb_dim, self._d_in)
            else:
                emb_dim = int(self.feature_emb_dim or self._d_in)
                self.id_emb = nn.Embedding(int(self.n_tokens), emb_dim)
                if self.feature_emb_mode == "concat":
                    self.id_fuse = nn.Linear(self._d_in + emb_dim, self._d_in)

        # ------------------------------------------------------------------
        # NEW: coord embeddings (topk only)
        # ------------------------------------------------------------------
        self.chrom_emb: Optional[nn.Embedding] = None
        self.coord_mlp: Optional[nn.Module] = None
        if self.use_coord_emb:
            if self.mode not in ("topk_scalar", "topk_channels"):
                raise ValueError("use_coord_embedding=True is only supported for topk_* tokenizers.")
            if self.n_chroms is None or int(self.n_chroms) <= 0:
                raise ValueError("use_coord_embedding=True requires tok.n_chroms > 0")
            self.chrom_emb = nn.Embedding(int(self.n_chroms), self._d_in)
            cd_in = 1 if self.coord_mode == "midpoint" else 2
            self.coord_mlp = _mlp(cd_in, self._d_in, hidden=self.coord_mlp_hidden)

        # ------------------------------------------------------------------
        # CLS token (learned, matches d_in)
        # ------------------------------------------------------------------
        self.cls_token: Optional[nn.Parameter] = None
        if self.add_cls_token:
            self.cls_token = nn.Parameter(torch.zeros(1, 1, self._d_in))
            nn.init.normal_(self.cls_token, mean=0.0, std=0.02)

    @property
    def d_in(self) -> int:
        return int(self._d_in)

    def set_feature_coords(self, chrom_ids: torch.Tensor, start: torch.Tensor, end: torch.Tensor) -> None:
        chrom_ids = chrom_ids.long().contiguous()
        start = start.to(dtype=torch.float32).contiguous()
        end = end.to(dtype=torch.float32).contiguous()
        if chrom_ids.ndim != 1 or start.ndim != 1 or end.ndim != 1:
            raise ValueError("set_feature_coords expects 1D tensors: chrom_ids/start/end.")
        if not (chrom_ids.shape[0] == start.shape[0] == end.shape[0] == self.input_dim):
            raise ValueError(
                f"set_feature_coords expects length F={self.input_dim}; got "
                f"{chrom_ids.shape[0]}, {start.shape[0]}, {end.shape[0]}"
            )
        self._chrom_ids = chrom_ids
        self._start = start
        self._end = end
        self._has_coords = True

    def _apply_id_emb(self, tokens: torch.Tensor, ids: torch.Tensor) -> torch.Tensor:
        if self.id_emb is None:
            return tokens
        idv = self.id_emb(ids)

        if self.feature_emb_mode == "add":
            if idv.shape[-1] != tokens.shape[-1]:
                raise ValueError(
                    f"feature_emb_mode='add' requires emb_dim==d_in; got emb_dim={idv.shape[-1]} vs d_in={tokens.shape[-1]}"
                )
            return tokens + idv

        if self.feature_emb_mode == "concat":
            if self.id_fuse is None:
                raise RuntimeError("id_fuse not initialized for concat mode.")
            return self.id_fuse(torch.cat([tokens, idv], dim=-1))

        raise ValueError(f"Unknown feature_emb_mode={self.feature_emb_mode!r}")

    def _apply_coord_emb(self, tokens: torch.Tensor, topk_idx: torch.Tensor) -> torch.Tensor:
        if not self.use_coord_emb:
            return tokens
        if not self._has_coords:
            return tokens
        if self.chrom_emb is None or self.coord_mlp is None:
            return tokens

        B, K = topk_idx.shape
        Fdim = self.input_dim
        if self._chrom_ids.numel() != Fdim:
            raise ValueError(f"Tokenizer coords are for F={self._chrom_ids.numel()} but input_dim={Fdim}")

        chrom = torch.gather(self._chrom_ids.view(1, Fdim).expand(B, Fdim), 1, topk_idx)
        start = torch.gather(self._start.view(1, Fdim).expand(B, Fdim), 1, topk_idx)
        end = torch.gather(self._end.view(1, Fdim).expand(B, Fdim), 1, topk_idx)

        chrom_e = self.chrom_emb(chrom)

        if self.coord_mode == "midpoint":
            mid = 0.5 * (start + end)
            pos = (mid * self.coord_scale).unsqueeze(-1)
        else:
            pos = torch.stack([start * self.coord_scale, end * self.coord_scale], dim=-1)

        pos_e = self.coord_mlp(pos)
        return tokens + chrom_e + pos_e

    def build_distance_attn_bias(
        self,
        topk_idx: torch.Tensor,
        *,
        lengthscale_bp: float = 50_000.0,
        same_chrom_only: bool = True,
        include_cls: bool = False,
        cls_is_zero: bool = True,
    ) -> torch.Tensor:
        if self.mode not in ("topk_scalar", "topk_channels"):
            raise ValueError("build_distance_attn_bias is only supported for topk_* modes.")
        if not self._has_coords:
            raise ValueError("build_distance_attn_bias requires feature coords; call set_feature_coords first.")
        if float(lengthscale_bp) <= 0:
            raise ValueError("lengthscale_bp must be > 0")

        B, K = topk_idx.shape
        Fdim = self.input_dim

        chrom = torch.gather(self._chrom_ids.view(1, Fdim).expand(B, Fdim), 1, topk_idx)
        start = torch.gather(self._start.view(1, Fdim).expand(B, Fdim), 1, topk_idx)
        end = torch.gather(self._end.view(1, Fdim).expand(B, Fdim), 1, topk_idx)
        mid = 0.5 * (start + end)

        dist = (mid.unsqueeze(2) - mid.unsqueeze(1)).abs()

        if same_chrom_only:
            same = (chrom.unsqueeze(2) == chrom.unsqueeze(1))
        else:
            same = torch.ones((B, K, K), device=dist.device, dtype=torch.bool)

        ls = float(lengthscale_bp)
        bias = -((dist / ls) ** 2)
        bias = torch.where(same, bias, torch.full_like(bias, -1e4))

        if include_cls:
            T = K + 1
            out = torch.zeros((B, T, T), device=bias.device, dtype=bias.dtype)
            out[:, 1:, 1:] = bias
            if not cls_is_zero:
                pass
            return out

        return bias

    def forward(
        self,
        x: torch.Tensor,
        *,
        return_indices: bool = False,
    ) -> Union[
        Tuple[torch.Tensor, Optional[torch.Tensor]],
        Tuple[torch.Tensor, Optional[torch.Tensor], Dict[str, Any]],
    ]:
        if x.dim() != 2:
            raise ValueError(f"_VectorToTokens expects x as (B,F); got shape {tuple(x.shape)}")
        B, Fdim = x.shape
        if Fdim != self.input_dim:
            raise ValueError(f"Expected input_dim={self.input_dim}, got F={Fdim}")

        key_padding_mask: Optional[torch.Tensor] = None
        meta: Dict[str, Any] = {}

        if self.mode == "topk_scalar":
            K = min(int(self.n_tokens), Fdim)
            vals, idx = torch.topk(x, k=K, dim=1, largest=True, sorted=False)
            tokens = vals.unsqueeze(-1)
            key_padding_mask = None
            if return_indices:
                meta["topk_idx"] = idx

            if self.val_proj is not None:
                tokens = self.val_proj(tokens)

            if self.use_feature_emb:
                tokens = self._apply_id_emb(tokens, idx)

            tokens = self._apply_coord_emb(tokens, idx)

        elif self.mode == "topk_channels":
            K = min(int(self.n_tokens), Fdim)
            vals, idx = torch.topk(x, k=K, dim=1, largest=True, sorted=True)
            feats = []
            for c in self.channels:
                if c == "value":
                    feats.append(vals)
                elif c == "rank":
                    if K <= 1:
                        rank01 = torch.zeros((B, K), device=x.device, dtype=torch.float32)
                    else:
                        rank01 = torch.linspace(0.0, 1.0, steps=K, device=x.device, dtype=torch.float32)
                        rank01 = rank01.unsqueeze(0).expand(B, K)
                    feats.append(rank01)
                elif c == "dropout":
                    feats.append((vals == 0).to(torch.float32))
                else:
                    raise RuntimeError(f"Unhandled channel: {c!r}")
            tokens = torch.stack(feats, dim=-1)
            key_padding_mask = None
            if return_indices:
                meta["topk_idx"] = idx

            if self.val_proj is not None:
                tokens = self.val_proj(tokens)

            if self.use_feature_emb:
                tokens = self._apply_id_emb(tokens, idx)

            tokens = self._apply_coord_emb(tokens, idx)

        else:  # patch
            P = int(self.patch_size)
            T = int(self.n_tokens)
            pad = T * P - Fdim
            if pad > 0:
                x_pad = F.pad(x, (0, pad), mode="constant", value=0.0)
            else:
                x_pad = x
            patches = x_pad.view(B, T, P)

            tokens = self.patch_proj(patches) if self.patch_proj is not None else patches

            if pad > 0:
                real_counts = torch.full((T,), P, device=x.device, dtype=torch.int64)
                last_real = Fdim - (T - 1) * P
                if last_real < P:
                    real_counts[-1] = max(int(last_real), 0)
                key_padding_mask = (real_counts == 0).unsqueeze(0).expand(B, T)
            else:
                key_padding_mask = None

            if return_indices:
                meta["patch_size"] = P
                meta["n_patches"] = T
                meta["pad"] = pad

            if self.use_feature_emb and self.id_emb is not None:
                pid = torch.arange(T, device=x.device).view(1, T).expand(B, T)
                tokens = self._apply_id_emb(tokens, pid)

        if self.add_cls_token:
            assert self.cls_token is not None
            cls = self.cls_token.expand(B, -1, -1)
            tokens = torch.cat([cls, tokens], dim=1)
            if key_padding_mask is not None:
                cls_mask = torch.zeros((B, 1), device=x.device, dtype=torch.bool)
                key_padding_mask = torch.cat([cls_mask, key_padding_mask], dim=1)

        if return_indices:
            return tokens, key_padding_mask, meta
        return tokens, key_padding_mask


# =============================================================================
# Per-modality transformer Gaussian encoder
# =============================================================================

def _cfg_to_model_tcfg(cfg: CFGTransformerConfig) -> ModelTransformerConfig:
    return ModelTransformerConfig(
        d_model=int(cfg.d_model),
        num_heads=int(cfg.num_heads),
        num_layers=int(cfg.num_layers),
        dim_feedforward=int(cfg.dim_feedforward),
        dropout=float(cfg.dropout),
        attn_dropout=float(cfg.attn_dropout),
        activation=str(cfg.activation),
        pooling=str(cfg.pooling),
        max_tokens=None if cfg.max_tokens is None else int(cfg.max_tokens),
    )


class TransformerGaussianEncoder(GaussianEncoder):
    """
    (B,F) -> tokens (B,T,D_in) -> TransformerEncoder -> (mu, logvar)

    - attach coords: self.vec2tok.set_feature_coords(...)
    - optional attn_bias passthrough (e.g., distance bias)
    """
    def __init__(
        self,
        *,
        input_dim: int,
        latent_dim: int,
        tokenizer: _VectorToTokens,
        tcfg: ModelTransformerConfig,
        use_positional_encoding: bool = True,
    ):
        super().__init__()
        self.vec2tok = tokenizer

        if use_positional_encoding and tcfg.max_tokens is None:
            tcfg.max_tokens = int(self.vec2tok.n_tokens + (1 if self.vec2tok.add_cls_token else 0))

        self.encoder = TransformerEncoder(
            cfg=tcfg,
            d_in=int(self.vec2tok.d_in),
            d_out=2 * int(latent_dim),
            use_positional_encoding=bool(use_positional_encoding),
        )

    def forward(
        self,
        x: torch.Tensor,
        *,
        return_attn: bool = False,
        attn_average_heads: bool = True,
        return_token_meta: bool = False,
        attn_bias: Optional[torch.Tensor] = None,
    ):
        if return_token_meta:
            tokens, key_padding_mask, meta = self.vec2tok(x, return_indices=True)
        else:
            tokens, key_padding_mask = self.vec2tok(x, return_indices=False)
            meta = None

        if return_attn:
            h, attn_all = _call_transformer_encoder(
                self.encoder,
                tokens,
                key_padding_mask=key_padding_mask,
                return_attn=True,
                attn_average_heads=attn_average_heads,
                attn_bias=attn_bias,
            )
        else:
            h = _call_transformer_encoder(
                self.encoder,
                tokens,
                key_padding_mask=key_padding_mask,
                return_attn=False,
                attn_average_heads=attn_average_heads,
                attn_bias=attn_bias,
            )
            attn_all = None

        mu, logvar = torch.chunk(h, 2, dim=-1)

        if return_attn and return_token_meta:
            return mu, logvar, attn_all, meta
        if return_attn:
            return mu, logvar, attn_all
        if return_token_meta:
            return mu, logvar, meta
        return mu, logvar


# =============================================================================
# Multimodal concatenated-token transformer Gaussian encoder (fused)
# =============================================================================

def _tokcfg_without_cls(tok_cfg_in: TokenizerConfig) -> TokenizerConfig:
    return replace(tok_cfg_in, add_cls_token=False)


class MultiModalTransformerGaussianEncoder(nn.Module):
    """
    Fused encoder over multiple modalities by concatenating tokens.

    Produces ONE fused posterior q(z | x_all). It does not replace per-modality q(z|x_m).
    """
    def __init__(
        self,
        *,
        modalities: Sequence[str],
        input_dims: Dict[str, int],
        tokenizers: Dict[str, TokenizerConfig],
        transformer_cfg: CFGTransformerConfig,
        latent_dim: int,
        add_modality_embeddings: bool = True,
        use_positional_encoding: bool = True,
    ):
        super().__init__()

        self.modalities = list(modalities)
        self.latent_dim = int(latent_dim)

        tcfg_model = _cfg_to_model_tcfg(transformer_cfg)
        d_model = int(tcfg_model.d_model)

        self.vec2tok = nn.ModuleDict()
        self.proj = nn.ModuleDict()
        self.mod_emb = nn.ParameterDict() if add_modality_embeddings else None

        total_tokens = 0
        for m in self.modalities:
            tok_cfg_in = tokenizers[m]
            tok_cfg = _tokcfg_without_cls(tok_cfg_in)

            tok = _VectorToTokens(input_dim=int(input_dims[m]), tok=tok_cfg)
            self.vec2tok[m] = tok
            self.proj[m] = nn.Linear(int(tok.d_in), d_model, bias=True)

            if self.mod_emb is not None:
                self.mod_emb[m] = nn.Parameter(torch.zeros(1, 1, d_model))
                nn.init.normal_(self.mod_emb[m], mean=0.0, std=0.02)

            total_tokens += int(tok.n_tokens)

        self.pooling = str(tcfg_model.pooling).lower().strip()
        self.use_global_cls = (self.pooling == "cls")
        self.cls_token = nn.Parameter(torch.zeros(1, 1, d_model)) if self.use_global_cls else None
        if self.cls_token is not None:
            nn.init.normal_(self.cls_token, mean=0.0, std=0.02)

        if use_positional_encoding and tcfg_model.max_tokens is None:
            tcfg_model.max_tokens = int(total_tokens + (1 if self.use_global_cls else 0))

        self.encoder = TransformerEncoder(
            cfg=tcfg_model,
            d_in=d_model,
            d_out=2 * int(latent_dim),
            use_positional_encoding=bool(use_positional_encoding),
        )

    def set_feature_coords(self, modality: str, chrom_ids: torch.Tensor, start: torch.Tensor, end: torch.Tensor) -> None:
        modality = str(modality)
        if modality not in self.vec2tok:
            raise KeyError(f"Unknown modality {modality!r}. Known: {list(self.vec2tok.keys())}")
        self.vec2tok[modality].set_feature_coords(chrom_ids, start, end)

    def forward(
        self,
        x_dict: Dict[str, torch.Tensor],
        *,
        return_token_meta: bool = False,
        return_attn: bool = False,
        attn_average_heads: bool = True,
        attn_bias: Optional[torch.Tensor] = None,
        attn_bias_fn: Optional[Any] = None,
    ):
        tokens_list: List[torch.Tensor] = []
        masks_list: List[Optional[torch.Tensor]] = []

        meta: Dict[str, Any] = {
            "modalities": self.modalities,
            "slices": {},
            "has_global_cls": bool(self.use_global_cls),
            "cls_index": 0 if self.use_global_cls else None,
        }
        t_cursor = 0

        for m in self.modalities:
            x = x_dict[m]

            if return_token_meta:
                tok, mask, mmeta = self.vec2tok[m](x, return_indices=True)
                meta[m] = mmeta
            else:
                tok, mask = self.vec2tok[m](x, return_indices=False)

            tok = self.proj[m](tok)
            if self.mod_emb is not None:
                tok = tok + self.mod_emb[m]

            Tm = tok.shape[1]
            meta["slices"][m] = (t_cursor, t_cursor + Tm)
            t_cursor += Tm

            tokens_list.append(tok)
            masks_list.append(mask)

        tokens = torch.cat(tokens_list, dim=1)

        key_padding_mask: Optional[torch.Tensor] = None
        if any(m is not None for m in masks_list):
            B = tokens.shape[0]
            built: List[torch.Tensor] = []
            for i, mname in enumerate(self.modalities):
                mask = masks_list[i]
                if mask is None:
                    Tm = tokens_list[i].shape[1]
                    built.append(torch.zeros((B, Tm), device=tokens.device, dtype=torch.bool))
                else:
                    built.append(mask.to(dtype=torch.bool))
            key_padding_mask = torch.cat(built, dim=1)

        if self.use_global_cls:
            assert self.cls_token is not None
            cls = self.cls_token.expand(tokens.shape[0], -1, -1)
            tokens = torch.cat([cls, tokens], dim=1)

            meta["slices_with_cls"] = {k: (a + 1, b + 1) for k, (a, b) in meta["slices"].items()}

            if key_padding_mask is not None:
                cls_mask = torch.zeros((tokens.shape[0], 1), device=tokens.device, dtype=torch.bool)
                key_padding_mask = torch.cat([cls_mask, key_padding_mask], dim=1)
        else:
            meta["slices_with_cls"] = dict(meta["slices"])

        if attn_bias is None and attn_bias_fn is not None:
            attn_bias = attn_bias_fn(meta)

        if return_attn:
            h, attn_all = _call_transformer_encoder(
                self.encoder,
                tokens,
                key_padding_mask=key_padding_mask,
                return_attn=True,
                attn_average_heads=attn_average_heads,
                attn_bias=attn_bias,
            )
        else:
            h = _call_transformer_encoder(
                self.encoder,
                tokens,
                key_padding_mask=key_padding_mask,
                return_attn=False,
                attn_average_heads=attn_average_heads,
                attn_bias=attn_bias,
            )
            attn_all = None

        mu, logvar = torch.chunk(h, 2, dim=-1)

        if return_attn and return_token_meta:
            return mu, logvar, attn_all, meta
        if return_attn:
            return mu, logvar, attn_all
        if return_token_meta:
            return mu, logvar, meta
        return mu, logvar


# =============================================================================
# Factories
# =============================================================================

def build_gaussian_encoder(*, uni_cfg: UniVIConfig, mod_cfg: ModalityConfig) -> GaussianEncoder:
    kind = (mod_cfg.encoder_type or "mlp").lower().strip()

    if kind == "mlp":
        return MLPGaussianEncoder(
            EncoderConfig(
                input_dim=int(mod_cfg.input_dim),
                hidden_dims=list(mod_cfg.encoder_hidden),
                latent_dim=int(uni_cfg.latent_dim),
                dropout=float(uni_cfg.encoder_dropout),
                batchnorm=bool(uni_cfg.encoder_batchnorm),
            )
        )

    if kind == "transformer":
        if mod_cfg.transformer is None:
            raise ValueError(f"Modality {mod_cfg.name!r}: encoder_type='transformer' requires mod_cfg.transformer.")
        if mod_cfg.tokenizer is None:
            raise ValueError(f"Modality {mod_cfg.name!r}: encoder_type='transformer' requires mod_cfg.tokenizer.")

        tokenizer = _VectorToTokens(
            input_dim=int(mod_cfg.input_dim),
            tok=mod_cfg.tokenizer,
        )

        tcfg = _cfg_to_model_tcfg(mod_cfg.transformer)
        if tcfg.max_tokens is None:
            tcfg.max_tokens = int(tokenizer.n_tokens + (1 if tokenizer.add_cls_token else 0))

        return TransformerGaussianEncoder(
            input_dim=int(mod_cfg.input_dim),
            latent_dim=int(uni_cfg.latent_dim),
            tokenizer=tokenizer,
            tcfg=tcfg,
            use_positional_encoding=True,
        )

    raise ValueError(f"Unknown encoder_type={kind!r} for modality {mod_cfg.name!r}")


def build_multimodal_transformer_encoder(
    *,
    uni_cfg: UniVIConfig,
    modalities: Sequence[ModalityConfig],
    fused_modalities: Optional[Sequence[str]] = None,
) -> MultiModalTransformerGaussianEncoder:
    if uni_cfg.fused_transformer is None:
        raise ValueError("UniVIConfig.fused_transformer must be set for fused_encoder_type='multimodal_transformer'.")

    mods = {m.name: m for m in modalities}
    use_names = list(fused_modalities) if fused_modalities is not None else list(mods.keys())

    input_dims = {n: int(mods[n].input_dim) for n in use_names}
    tokenizers: Dict[str, TokenizerConfig] = {}
    for n in use_names:
        if mods[n].tokenizer is None:
            raise ValueError(f"Fused multimodal encoder requires tokenizer for modality {n!r}")
        tokenizers[n] = mods[n].tokenizer

    return MultiModalTransformerGaussianEncoder(
        modalities=use_names,
        input_dims=input_dims,
        tokenizers=tokenizers,
        transformer_cfg=uni_cfg.fused_transformer,
        latent_dim=int(uni_cfg.latent_dim),
        add_modality_embeddings=bool(getattr(uni_cfg, "fused_add_modality_embeddings", True)),
        use_positional_encoding=True,
    )
```
