## Supplemental Code for "Unifying multimodal single-cell data with a mixture-of-experts *β*-variational autoencoder framework": mlp.py

```
# univi/models/mlp.py

from __future__ import annotations
from typing import List, Optional
from torch import nn


def build_mlp(
    in_dim: int,
    hidden_dims: List[int],
    out_dim: int,
    activation: Optional[nn.Module] = None,
    dropout: float = 0.0,
    batchnorm: bool = True,
) -> nn.Sequential:
    """
    Generic MLP builder: [Linear -> BN -> Act -> Dropout]* + final Linear.
    """
    if activation is None:
        activation = nn.ReLU()

    layers = []
    last_dim = in_dim
    for h in hidden_dims:
        layers.append(nn.Linear(last_dim, h))
        if batchnorm:
            layers.append(nn.BatchNorm1d(h))
        layers.append(activation.__class__() if isinstance(activation, nn.Module) else nn.ReLU())
        if dropout and dropout > 0:
            layers.append(nn.Dropout(float(dropout)))
        last_dim = h

    layers.append(nn.Linear(last_dim, out_dim))
    return nn.Sequential(*layers)
```
