## Supplemental Code for "Unifying multimodal single-cell data with a mixture-of-experts *β*-variational autoencoder framework": tokenizers.py

```
# univi/models/tokenizers.py
from __future__ import annotations

from typing import Optional, Tuple, Sequence, Literal, Dict, Any

import torch
from torch import nn

from ..config import TokenizerConfig


class Tokenizer(nn.Module):
    """
    Base tokenizer interface.

    Backwards-compatible:
      forward(x) -> (tokens, key_padding_mask)

    Extras:
      - self.last_meta is updated on each forward()
      - forward_with_meta(x) -> (tokens, key_padding_mask, meta)

    Conventions
    -----------
    - tokens: (B, T, D_in)
    - key_padding_mask: Optional[(B, T)] where True means "PAD / ignore"
    - meta: dict (optional), e.g. {"token_pos": (B, T) basepair positions}
    """
    def __init__(self):
        super().__init__()
        self.last_meta: Dict[str, Any] = {}

    @property
    def d_in(self) -> int:
        raise NotImplementedError

    def forward(self, x: torch.Tensor) -> Tuple[torch.Tensor, Optional[torch.Tensor]]:
        raise NotImplementedError

    def forward_with_meta(self, x: torch.Tensor) -> Tuple[torch.Tensor, Optional[torch.Tensor], Dict[str, Any]]:
        tokens, mask = self.forward(x)
        return tokens, mask, dict(self.last_meta)


class TopKScalarTokenizer(Tokenizer):
    """(B,F) -> (B,K,1) using top-k by absolute value per cell."""
    def __init__(self, n_tokens: int, add_cls_token: bool = False):
        super().__init__()
        self.n_tokens = int(n_tokens)
        self.add_cls_token = bool(add_cls_token)

    @property
    def d_in(self) -> int:
        return 1

    def forward(self, x: torch.Tensor):
        B, F = x.shape
        K = min(self.n_tokens, F)

        _, idx = torch.topk(x.abs(), k=K, dim=1, largest=True, sorted=True)
        vals = torch.gather(x, 1, idx)      # (B,K)
        tokens = vals.unsqueeze(-1)         # (B,K,1)

        key_padding_mask = None
        self.last_meta = {"feature_idx": idx}

        if self.add_cls_token:
            cls = torch.zeros((B, 1, 1), device=x.device, dtype=x.dtype)
            tokens = torch.cat([cls, tokens], dim=1)

        return tokens, key_padding_mask


class TopKChannelsTokenizer(Tokenizer):
    """
    (B,F) -> (B,K,C) multi-dim tokens, where channels can include:
      - value: raw x_i
      - rank: rank within selected K (0..1)
      - dropout: 1 if x_i == 0 else 0
    """
    def __init__(
        self,
        n_tokens: int,
        channels: Sequence[Literal["value", "rank", "dropout"]] = ("value", "rank", "dropout"),
        add_cls_token: bool = False,
    ):
        super().__init__()
        self.n_tokens = int(n_tokens)
        self.channels = tuple(channels)
        self.add_cls_token = bool(add_cls_token)

        if len(self.channels) == 0:
            raise ValueError("TopKChannelsTokenizer requires at least one channel.")
        for c in self.channels:
            if c not in ("value", "rank", "dropout"):
                raise ValueError(f"Unknown channel {c!r}. Allowed: value, rank, dropout")

    @property
    def d_in(self) -> int:
        return len(self.channels)

    def forward(self, x: torch.Tensor):
        B, F = x.shape
        K = min(self.n_tokens, F)

        _, idx = torch.topk(x.abs(), k=K, dim=1, largest=True, sorted=True)
        vals = torch.gather(x, 1, idx)  # (B,K)

        chans = []
        for c in self.channels:
            if c == "value":
                chans.append(vals)
            elif c == "dropout":
                chans.append((vals == 0).to(vals.dtype))
            elif c == "rank":
                r = torch.arange(K, device=x.device, dtype=vals.dtype).view(1, K).expand(B, K)
                chans.append(r / max(K - 1, 1))
            else:
                raise RuntimeError("unreachable")

        tokens = torch.stack(chans, dim=-1)  # (B,K,C)
        key_padding_mask = None
        self.last_meta = {"feature_idx": idx}

        if self.add_cls_token:
            cls = torch.zeros((B, 1, tokens.size(-1)), device=x.device, dtype=x.dtype)
            tokens = torch.cat([cls, tokens], dim=1)

        return tokens, key_padding_mask


class PatchTokenizer(Tokenizer):
    """
    Split features into patches:

      (B,F) -> (B,T,patch_size)  where T = ceil(F/patch_size)

    Optionally project:
      patch_vec (patch_size) -> patch_proj_dim
    """
    def __init__(
        self,
        patch_size: int,
        add_cls_token: bool = False,
        patch_proj_dim: Optional[int] = None,
    ):
        super().__init__()
        self.patch_size = int(patch_size)
        self.add_cls_token = bool(add_cls_token)
        self.patch_proj_dim = int(patch_proj_dim) if patch_proj_dim is not None else None

        if self.patch_size <= 0:
            raise ValueError("patch_size must be > 0")

        if self.patch_proj_dim is not None:
            self.proj = nn.Sequential(
                nn.LayerNorm(self.patch_size),
                nn.Linear(self.patch_size, self.patch_proj_dim),
                nn.GELU(),
                nn.Linear(self.patch_proj_dim, self.patch_proj_dim),
            )
        else:
            self.proj = None

    @property
    def d_in(self) -> int:
        return self.patch_proj_dim if self.patch_proj_dim is not None else self.patch_size

    def forward(self, x: torch.Tensor):
        B, F = x.shape
        P = self.patch_size
        T = (F + P - 1) // P
        pad = T * P - F

        if pad > 0:
            x_pad = torch.cat([x, torch.zeros((B, pad), device=x.device, dtype=x.dtype)], dim=1)
        else:
            x_pad = x

        patches = x_pad.view(B, T, P)  # (B,T,P)
        key_padding_mask = None

        if self.proj is not None:
            patches = self.proj(patches)  # (B,T,patch_proj_dim)

        if self.add_cls_token:
            cls = torch.zeros((B, 1, patches.size(-1)), device=x.device, dtype=x.dtype)
            patches = torch.cat([cls, patches], dim=1)

        self.last_meta = {}
        return patches, key_padding_mask


class TopKEmbeddedTokenizer(Tokenizer):
    """
    Top-k tokenizer with explicit feature identity embeddings:

      token = Emb(feature_id) + MLP(channels(value/rank/dropout))

    Optional ATAC coordinate embeddings:
      token += Emb(chrom_id) + MLP(midpoint_bp / coord_scale)

    Meta
    ----
    self.last_meta will include:
      - "feature_idx": (B,K) long
      - "token_pos":  (B,K) float basepairs (if use_coords=True)
    """
    def __init__(
        self,
        *,
        n_tokens: int,
        n_features: int,
        d_model: int,
        channels: Sequence[Literal["value", "rank", "dropout"]] = ("value", "rank", "dropout"),
        add_cls_token: bool = False,
        value_mlp_hidden: int = 256,
        # coordinate extras
        use_coords: bool = False,
        chrom_vocab_size: int = 0,
        feature_info: Optional[Dict[str, Any]] = None,
        coord_scale: float = 1e6,
    ):
        super().__init__()
        self.n_tokens = int(n_tokens)
        self.n_features = int(n_features)
        self._d_model = int(d_model)
        self.channels = tuple(channels)
        self.add_cls_token = bool(add_cls_token)
        self.use_coords = bool(use_coords)
        self.chrom_vocab_size = int(chrom_vocab_size)
        self.coord_scale = float(coord_scale)

        if len(self.channels) == 0:
            raise ValueError("TopKEmbeddedTokenizer requires at least one channel.")
        for c in self.channels:
            if c not in ("value", "rank", "dropout"):
                raise ValueError(f"Unknown channel {c!r}. Allowed: value, rank, dropout")

        self.id_embed = nn.Embedding(self.n_features, self._d_model)

        c_in = len(self.channels)
        self.val_proj = nn.Sequential(
            nn.LayerNorm(c_in),
            nn.Linear(c_in, int(value_mlp_hidden)),
            nn.GELU(),
            nn.Linear(int(value_mlp_hidden), self._d_model),
        )

        # Feature metadata buffers for coords
        self.feature_chrom: Optional[torch.Tensor] = None
        self.feature_start: Optional[torch.Tensor] = None
        self.feature_end: Optional[torch.Tensor] = None

        if self.use_coords:
            if self.chrom_vocab_size <= 0:
                raise ValueError("chrom_vocab_size must be > 0 when use_coords=True.")
            if feature_info is None:
                raise ValueError("feature_info must be provided when use_coords=True (keys: chrom,start,end).")
            for k in ("chrom", "start", "end"):
                if k not in feature_info:
                    raise ValueError(f"feature_info missing key {k!r} (required for coords).")

            chrom = torch.as_tensor(feature_info["chrom"], dtype=torch.long)
            start = torch.as_tensor(feature_info["start"], dtype=torch.float32)
            end = torch.as_tensor(feature_info["end"], dtype=torch.float32)

            if chrom.numel() != self.n_features or start.numel() != self.n_features or end.numel() != self.n_features:
                raise ValueError(
                    f"feature_info arrays must have length n_features={self.n_features}; "
                    f"got chrom={chrom.numel()}, start={start.numel()}, end={end.numel()}."
                )

            # register buffers so they follow .to(device)
            self.register_buffer("feature_chrom", chrom, persistent=False)
            self.register_buffer("feature_start", start, persistent=False)
            self.register_buffer("feature_end", end, persistent=False)

            self.chrom_embed = nn.Embedding(self.chrom_vocab_size, self._d_model)
            self.coord_mlp = nn.Sequential(
                nn.LayerNorm(1),
                nn.Linear(1, int(value_mlp_hidden)),
                nn.GELU(),
                nn.Linear(int(value_mlp_hidden), self._d_model),
            )

    @property
    def d_in(self) -> int:
        return self._d_model

    def forward(self, x: torch.Tensor):
        B, F = x.shape
        if F != self.n_features:
            raise ValueError(f"Expected F={self.n_features}, got {F}. Did you set TokenizerConfig.n_features correctly?")

        K = min(self.n_tokens, F)

        _, idx = torch.topk(x.abs(), k=K, dim=1, largest=True, sorted=True)  # (B,K)
        vals = torch.gather(x, 1, idx)                                       # (B,K)

        # channels -> (B,K,C)
        chans = []
        for c in self.channels:
            if c == "value":
                chans.append(vals)
            elif c == "dropout":
                chans.append((vals == 0).to(vals.dtype))
            elif c == "rank":
                r = torch.arange(K, device=x.device, dtype=vals.dtype).view(1, K).expand(B, K)
                chans.append(r / max(K - 1, 1))
            else:
                raise RuntimeError("unreachable")

        ch = torch.stack(chans, dim=-1)  # (B,K,C)

        id_emb = self.id_embed(idx)      # (B,K,D)
        val_emb = self.val_proj(ch)      # (B,K,D)
        tokens = id_emb + val_emb

        meta: Dict[str, Any] = {"feature_idx": idx}

        if self.use_coords:
            # buffers exist because we register_buffer above
            chrom = self.feature_chrom[idx]  # (B,K)
            mid = 0.5 * (self.feature_start[idx] + self.feature_end[idx])  # (B,K)
            mid_scaled = (mid / self.coord_scale).unsqueeze(-1)            # (B,K,1)

            tokens = tokens + self.chrom_embed(chrom) + self.coord_mlp(mid_scaled)
            meta["token_pos"] = mid  # basepairs

        if self.add_cls_token:
            cls = torch.zeros((B, 1, tokens.size(-1)), device=x.device, dtype=x.dtype)
            tokens = torch.cat([cls, tokens], dim=1)
            # keep meta aligned if present
            if "token_pos" in meta:
                cls_pos = torch.zeros((B, 1), device=x.device, dtype=meta["token_pos"].dtype)
                meta["token_pos"] = torch.cat([cls_pos, meta["token_pos"]], dim=1)

        self.last_meta = meta
        return tokens, None


def build_tokenizer(cfg: TokenizerConfig) -> Tokenizer:
    mode = (cfg.mode or "").lower().strip()

    if mode == "topk_scalar":
        return TopKScalarTokenizer(n_tokens=cfg.n_tokens, add_cls_token=cfg.add_cls_token)

    if mode == "topk_channels":
        return TopKChannelsTokenizer(n_tokens=cfg.n_tokens, channels=cfg.channels, add_cls_token=cfg.add_cls_token)

    if mode == "patch":
        return PatchTokenizer(
            patch_size=cfg.patch_size,
            add_cls_token=cfg.add_cls_token,
            patch_proj_dim=cfg.patch_proj_dim,
        )

    if mode == "topk_embed":
        if cfg.n_features is None or cfg.d_model is None:
            raise ValueError("TokenizerConfig.mode='topk_embed' requires n_features and d_model to be set.")
        return TopKEmbeddedTokenizer(
            n_tokens=cfg.n_tokens,
            n_features=int(cfg.n_features),
            d_model=int(cfg.d_model),
            channels=cfg.channels,
            add_cls_token=cfg.add_cls_token,
            value_mlp_hidden=int(cfg.value_mlp_hidden),
            use_coords=bool(cfg.use_coords),
            chrom_vocab_size=int(cfg.chrom_vocab_size),
            feature_info=cfg.feature_info,
            coord_scale=float(cfg.coord_scale),
        )

    raise ValueError(f"Unknown tokenizer mode {cfg.mode!r}")
```
