## Supplemental Code for "Unifying multimodal single-cell data with a mixture-of-experts *β*-variational autoencoder framework": transformer.py

```
# univi/models/transformer.py
from __future__ import annotations

from dataclasses import dataclass
from typing import Optional, Literal, List, Tuple, Union

import torch
from torch import nn
import torch.nn.functional as F


@dataclass
class TransformerConfig:
    d_model: int
    num_heads: int
    num_layers: int
    dim_feedforward: int = 4096
    dropout: float = 0.1
    attn_dropout: float = 0.1
    activation: Literal["relu", "gelu"] = "gelu"
    pooling: Literal["cls", "mean"] = "mean"
    max_tokens: Optional[int] = None

    # Optional: binned relative-position attention bias (e.g., genomic distance)
    use_relpos_bias: bool = False
    relpos_num_bins: int = 32
    relpos_max_dist: float = 1e6  # basepairs


def _act(name: str):
    name = str(name).lower().strip()
    if name == "relu":
        return F.relu
    if name == "gelu":
        return F.gelu
    raise ValueError(f"Unknown activation: {name!r}")


class GenomicRelPosBias(nn.Module):
    """
    Simple distance-binned relative attention bias.

    Given token positions pos (B,T) in basepairs, returns an additive bias
    (B, H, T, T). Intended for ATAC peak midpoints.

    Notes
    -----
    - Uses log1p compression to allocate more bins to shorter distances.
    - Bias table is learned: (H, num_bins).
    """
    def __init__(self, num_heads: int, num_bins: int = 32, max_dist: float = 1e6):
        super().__init__()
        self.num_heads = int(num_heads)
        self.num_bins = int(num_bins)
        self.max_dist = float(max_dist)
        self.bias = nn.Parameter(torch.zeros(self.num_heads, self.num_bins))

    def _bin(self, dist: torch.Tensor) -> torch.Tensor:
        # dist: (B,T,T) >= 0
        d = dist.clamp(min=0.0, max=self.max_dist)
        d = torch.log1p(d)
        dmax = torch.log1p(torch.tensor(self.max_dist, device=d.device, dtype=d.dtype))
        b = (d / dmax) * (self.num_bins - 1)
        return b.to(torch.long)

    def forward(self, pos: torch.Tensor) -> torch.Tensor:
        # pos: (B,T)
        dist = (pos[:, :, None] - pos[:, None, :]).abs()  # (B,T,T)
        bins = self._bin(dist)                             # (B,T,T)
        # bias[:, bins] -> (H,B,T,T) then permute -> (B,H,T,T)
        out = self.bias[:, bins]
        return out.permute(1, 0, 2, 3).contiguous()


def _as_mha_attn_mask(
    bias: torch.Tensor,
    *,
    num_heads: int,
    batch_size: int,
    seq_len: int,
    dtype: torch.dtype,
    device: torch.device,
) -> torch.Tensor:
    """
    Convert an additive attention bias into a shape accepted by nn.MultiheadAttention.

    Accepted input shapes:
      - (T, T)                     -> returned as-is (shared across batch/heads)
      - (B, T, T)                  -> expanded to (B*H, T, T)
      - (B, H, T, T)               -> reshaped to (B*H, T, T)
      - (B*H, T, T)                -> returned as-is
      - (1, T, T)                  -> treated like (B, T, T) and expanded
    """
    if bias.dim() == 2:
        # (T,T)
        if bias.shape != (seq_len, seq_len):
            raise ValueError(f"attn_bias (T,T) must be ({seq_len},{seq_len}); got {tuple(bias.shape)}")
        return bias.to(device=device, dtype=dtype)

    if bias.dim() == 3:
        # (B,T,T) or (1,T,T) or (B*H,T,T)
        b0, t1, t2 = bias.shape
        if (t1, t2) != (seq_len, seq_len):
            raise ValueError(f"attn_bias (...,T,T) must have T={seq_len}; got {tuple(bias.shape)}")

        if b0 == batch_size * num_heads:
            return bias.to(device=device, dtype=dtype)

        if b0 == 1:
            bias = bias.expand(batch_size, seq_len, seq_len)

        if b0 != batch_size:
            raise ValueError(
                f"attn_bias (B,T,T) must have B={batch_size} (or 1); got {b0} with shape {tuple(bias.shape)}"
            )

        # expand across heads -> (B*H,T,T)
        bias = bias.to(device=device, dtype=dtype)
        bias = bias.unsqueeze(1).expand(batch_size, num_heads, seq_len, seq_len)
        return bias.reshape(batch_size * num_heads, seq_len, seq_len).contiguous()

    if bias.dim() == 4:
        # (B,H,T,T)
        b0, h0, t1, t2 = bias.shape
        if (t1, t2) != (seq_len, seq_len):
            raise ValueError(f"attn_bias (B,H,T,T) must have T={seq_len}; got {tuple(bias.shape)}")
        if b0 != batch_size:
            raise ValueError(f"attn_bias (B,H,T,T) must have B={batch_size}; got {b0}")
        if h0 != num_heads:
            raise ValueError(f"attn_bias (B,H,T,T) must have H={num_heads}; got {h0}")
        bias = bias.to(device=device, dtype=dtype)
        return bias.reshape(batch_size * num_heads, seq_len, seq_len).contiguous()

    raise ValueError(f"attn_bias must be 2D/3D/4D; got ndim={bias.dim()} shape={tuple(bias.shape)}")


class TransformerBlock(nn.Module):
    """
    Single pre-norm style block:
      x -> MHA -> residual -> LN
        -> FFN -> residual -> LN

    Supports optional additive attention bias (e.g., genomic distance).
    """
    def __init__(self, cfg: TransformerConfig):
        super().__init__()
        self.cfg = cfg
        d_model = int(cfg.d_model)
        self.num_heads = int(cfg.num_heads)

        self.attn = nn.MultiheadAttention(
            embed_dim=d_model,
            num_heads=self.num_heads,
            dropout=float(cfg.attn_dropout),
            batch_first=True,
        )
        self.attn_drop = nn.Dropout(float(cfg.dropout))
        self.ln1 = nn.LayerNorm(d_model)

        self.ff = nn.Sequential(
            nn.Linear(d_model, int(cfg.dim_feedforward)),
            nn.GELU() if str(cfg.activation).lower().strip() == "gelu" else nn.ReLU(),
            nn.Dropout(float(cfg.dropout)),
            nn.Linear(int(cfg.dim_feedforward), d_model),
        )
        self.ff_drop = nn.Dropout(float(cfg.dropout))
        self.ln2 = nn.LayerNorm(d_model)

        self.relpos: Optional[GenomicRelPosBias] = None
        if bool(getattr(cfg, "use_relpos_bias", False)):
            self.relpos = GenomicRelPosBias(
                num_heads=self.num_heads,
                num_bins=int(getattr(cfg, "relpos_num_bins", 32)),
                max_dist=float(getattr(cfg, "relpos_max_dist", 1e6)),
            )

    def forward(
        self,
        x: torch.Tensor,
        *,
        key_padding_mask: Optional[torch.Tensor] = None,
        token_pos: Optional[torch.Tensor] = None,  # (B,T) basepairs or other coordinates
        attn_bias: Optional[torch.Tensor] = None,  # additive bias: (T,T) or (B,T,T) or (B,H,T,T) or (B*H,T,T)
        return_attn: bool = False,
        attn_average_heads: bool = True,
    ) -> Union[torch.Tensor, Tuple[torch.Tensor, torch.Tensor]]:
        need_weights = bool(return_attn)

        # Build additive attention mask for nn.MultiheadAttention
        attn_mask = None
        B, T, _ = x.shape
        H = self.num_heads

        # 1) relpos bias -> (B,H,T,T) -> (B*H,T,T)
        if self.relpos is not None and token_pos is not None:
            rel = self.relpos(token_pos).to(dtype=x.dtype)          # (B,H,T,T)
            rel = rel.reshape(B * H, T, T).contiguous()             # (B*H,T,T)
            attn_mask = rel

        # 2) external attn_bias -> normalize to MHA shape and add
        if attn_bias is not None:
            ext = _as_mha_attn_mask(
                attn_bias,
                num_heads=H,
                batch_size=B,
                seq_len=T,
                dtype=x.dtype,
                device=x.device,
            )
            attn_mask = ext if attn_mask is None else (attn_mask + ext)

        attn_out, attn_w = self.attn(
            x, x, x,
            key_padding_mask=key_padding_mask,        # (B, T) True = PAD
            attn_mask=attn_mask,                      # None or (T,T) or (B*H,T,T)
            need_weights=need_weights,
            average_attn_weights=bool(attn_average_heads),
        )

        x = self.ln1(x + self.attn_drop(attn_out))
        ff_out = self.ff(x)
        x = self.ln2(x + self.ff_drop(ff_out))

        if return_attn:
            if attn_w is None:
                raise RuntimeError("Expected attn_w when return_attn=True, got None.")
            return x, attn_w
        return x


class TransformerEncoder(nn.Module):
    """
    Generic encoder:
      tokens (B,T,D_in) -> proj -> blocks -> pool -> out_proj -> (B,d_out)

    Optional:
      - learned absolute positional embeddings (use_positional_encoding=True)
      - relative attention bias via token_pos (if cfg.use_relpos_bias=True)
      - external additive attention bias via attn_bias (passed through to blocks)
    """
    def __init__(
        self,
        *,
        cfg: TransformerConfig,
        d_in: int,
        d_out: int,
        use_positional_encoding: bool = True,
    ):
        super().__init__()
        self.cfg = cfg
        self.use_positional_encoding = bool(use_positional_encoding)

        d_model = int(cfg.d_model)
        self.input_proj = nn.Identity() if int(d_in) == d_model else nn.Linear(int(d_in), d_model, bias=True)
        self.blocks = nn.ModuleList([TransformerBlock(cfg) for _ in range(int(cfg.num_layers))])
        self.dropout = nn.Dropout(float(cfg.dropout))
        self.out_proj = nn.Linear(d_model, int(d_out), bias=True)

        self.pooling = str(cfg.pooling).lower().strip()
        if self.pooling not in ("cls", "mean"):
            raise ValueError(f"Unknown pooling={cfg.pooling!r}")

        # learned positional embeddings (optional)
        self.pos_emb: Optional[nn.Parameter] = None
        if self.use_positional_encoding:
            if cfg.max_tokens is None:
                raise ValueError("use_positional_encoding=True requires cfg.max_tokens to be set.")
            max_tokens = int(cfg.max_tokens)
            self.pos_emb = nn.Parameter(torch.zeros(1, max_tokens, d_model))
            nn.init.normal_(self.pos_emb, mean=0.0, std=0.02)

    def _pool(self, x: torch.Tensor, *, key_padding_mask: Optional[torch.Tensor]) -> torch.Tensor:
        if self.pooling == "cls":
            return x[:, 0, :]

        if key_padding_mask is None:
            return x.mean(dim=1)

        keep = (~key_padding_mask).to(dtype=x.dtype)  # (B, T)
        denom = keep.sum(dim=1, keepdim=True).clamp_min(1.0)
        return (x * keep.unsqueeze(-1)).sum(dim=1) / denom

    def forward(
        self,
        tokens: torch.Tensor,
        *,
        key_padding_mask: Optional[torch.Tensor] = None,
        token_pos: Optional[torch.Tensor] = None,  # (B,T) for relpos bias (optional)
        attn_bias: Optional[torch.Tensor] = None,  # (T,T) or (B,T,T) or (B,H,T,T) or (B*H,T,T)
        return_attn: bool = False,
        attn_average_heads: bool = True,
    ) -> Union[torch.Tensor, Tuple[torch.Tensor, List[torch.Tensor]]]:
        x = self.input_proj(tokens)

        if self.use_positional_encoding:
            assert self.pos_emb is not None
            T = x.shape[1]
            if T > self.pos_emb.shape[1]:
                raise ValueError(f"Sequence length T={T} exceeds max_tokens={self.pos_emb.shape[1]}.")
            x = x + self.pos_emb[:, :T, :]

        x = self.dropout(x)

        attn_all: List[torch.Tensor] = []
        for blk in self.blocks:
            if return_attn:
                x, aw = blk(
                    x,
                    key_padding_mask=key_padding_mask,
                    token_pos=token_pos,
                    attn_bias=attn_bias,
                    return_attn=True,
                    attn_average_heads=attn_average_heads,
                )
                attn_all.append(aw)
            else:
                x = blk(
                    x,
                    key_padding_mask=key_padding_mask,
                    token_pos=token_pos,
                    attn_bias=attn_bias,
                    return_attn=False,
                    attn_average_heads=attn_average_heads,
                )

        pooled = self._pool(x, key_padding_mask=key_padding_mask)
        out = self.out_proj(pooled)

        if return_attn:
            return out, attn_all
        return out
```
