## Supplemental Code for "Unifying multimodal single-cell data with a mixture-of-experts *β*-variational autoencoder framework": univi.py

```
# univi/models/univi.py
from __future__ import annotations

from typing import Dict, Tuple, Optional, Any, List, Union, Mapping, Callable, overload, Literal
import math

import torch
from torch import nn
import torch.nn.functional as F

from ..config import UniVIConfig, ModalityConfig, ClassHeadConfig
from .mlp import build_mlp
from .decoders import DecoderConfig, build_decoder
from .encoders import build_gaussian_encoder, build_multimodal_transformer_encoder


YType = Union[torch.Tensor, Dict[str, torch.Tensor]]


class _GradReverseFn(torch.autograd.Function):
    """Gradient reversal layer: forward identity, backward -lambda * grad."""
    @staticmethod
    def forward(ctx, x: torch.Tensor, lambd: float):
        ctx.lambd = float(lambd)
        return x.view_as(x)

    @staticmethod
    def backward(ctx, grad_output: torch.Tensor):
        return -ctx.lambd * grad_output, None


class _LogitsMLPHead(nn.Module):
    """
    Simple MLP head that returns logits in a dict for API compatibility with decoders.

    Used for binary heads (and can be used for categorical if desired),
    but we keep categorical default as build_decoder("categorical") to preserve behavior.
    """
    def __init__(
        self,
        in_dim: int,
        out_dim: int,
        *,
        hidden_dims: List[int],
        dropout: float = 0.0,
        batchnorm: bool = False,
        activation: Optional[nn.Module] = None,
    ):
        super().__init__()
        act = activation if activation is not None else nn.ReLU()
        self.net = build_mlp(
            in_dim=int(in_dim),
            hidden_dims=list(hidden_dims) if hidden_dims else [max(64, int(in_dim))],
            out_dim=int(out_dim),
            activation=act,
            dropout=float(dropout),
            batchnorm=bool(batchnorm),
        )

    def forward(self, z: torch.Tensor) -> Dict[str, torch.Tensor]:
        logits = self.net(z)
        return {"logits": logits}


def _as_list_int(x: Any) -> Optional[List[int]]:
    if x is None:
        return None
    if isinstance(x, (list, tuple)):
        out = []
        for v in x:
            if v is None:
                continue
            out.append(int(v))
        return out
    try:
        return [int(x)]
    except Exception:
        return None


def _parse_activation(act: Any) -> Optional[nn.Module]:
    if act is None:
        return None
    if isinstance(act, nn.Module):
        return act
    s = str(act).lower().strip()
    if s in ("relu",):
        return nn.ReLU()
    if s in ("gelu",):
        return nn.GELU()
    if s in ("elu",):
        return nn.ELU()
    if s in ("leakyrelu", "leaky_relu", "lrelu"):
        return nn.LeakyReLU(0.01)
    if s in ("silu", "swish"):
        return nn.SiLU()
    if s in ("tanh",):
        return nn.Tanh()
    return None


class UniVIMultiModalVAE(nn.Module):
    """
    Multi-modal β-VAE with per-modality encoders and decoders.
    """

    LOGVAR_MIN = -10.0
    LOGVAR_MAX = 10.0
    EPS = 1e-8

    def __init__(
        self,
        cfg: UniVIConfig,
        *,
        loss_mode: str = "v1",
        v1_recon: str = "avg",
        v1_recon_mix: float = 0.0,
        normalize_v1_terms: bool = True,

        recon_normalize_by_dim: Optional[bool] = None,
        recon_dim_power: Optional[float] = None,

        n_label_classes: int = 0,
        label_loss_weight: float = 1.0,
        use_label_encoder: bool = False,
        label_moe_weight: float = 1.0,
        unlabeled_logvar: float = 20.0,
        label_encoder_warmup: int = 0,
        label_ignore_index: int = -1,
        classify_from_mu: bool = True,
        label_head_name: Optional[str] = None,
    ):
        super().__init__()
        self.cfg = cfg
        self.cfg.validate()

        self.loss_mode = str(loss_mode).lower().strip()
        self.v1_recon = str(v1_recon).lower().strip()
        self.v1_recon_mix = float(v1_recon_mix)
        self.normalize_v1_terms = bool(normalize_v1_terms)

        mode = (self.loss_mode or "v2").lower()
        is_v1 = mode in ("v1", "paper", "cross")

        if recon_normalize_by_dim is None:
            recon_normalize_by_dim = False if is_v1 else True
        if recon_dim_power is None:
            recon_dim_power = 0.5 if is_v1 else 1.0

        self.recon_normalize_by_dim = bool(recon_normalize_by_dim)
        self.recon_dim_power = float(recon_dim_power)

        self.latent_dim = int(cfg.latent_dim)
        self.beta_max = float(cfg.beta)
        self.gamma_max = float(cfg.gamma)

        self.modality_names: List[str] = [m.name for m in cfg.modalities]
        self.mod_cfg_by_name: Dict[str, ModalityConfig] = {m.name: m for m in cfg.modalities}

        self.encoders = nn.ModuleDict()
        self.encoder_heads = nn.ModuleDict()
        self.decoders = nn.ModuleDict()

        for m in cfg.modalities:
            if not isinstance(m, ModalityConfig):
                raise TypeError(f"cfg.modalities must contain ModalityConfig, got {type(m)}")

            self.encoders[m.name] = build_gaussian_encoder(uni_cfg=cfg, mod_cfg=m)
            self.encoder_heads[m.name] = nn.Identity()

            dec_hidden = list(m.decoder_hidden) if m.decoder_hidden else [max(64, self.latent_dim)]
            dec_cfg = DecoderConfig(
                output_dim=int(m.input_dim),
                hidden_dims=dec_hidden,
                dropout=float(cfg.decoder_dropout),
                batchnorm=bool(cfg.decoder_batchnorm),
            )

            lk = (m.likelihood or "gaussian").lower().strip()
            decoder_kwargs: Dict[str, Any] = {}

            if lk in ("nb", "negative_binomial", "zinb", "zero_inflated_negative_binomial"):
                dispersion = getattr(m, "dispersion", "gene")
                init_log_theta = float(getattr(m, "init_log_theta", 0.0))
                decoder_kwargs = dict(
                    dispersion=dispersion,
                    init_log_theta=init_log_theta,
                    eps=self.EPS,
                )
            elif lk in ("beta",):
                decoder_kwargs = dict(
                    eps=self.EPS,
                    min_conc=float(getattr(m, "min_conc", 1e-4)),
                )
            elif lk in ("beta_binomial", "betabinomial", "bb"):
                decoder_kwargs = dict(
                    eps=self.EPS,
                    min_conc=float(getattr(m, "min_conc", 1e-4)),
                )
            elif lk in ("binomial",):
                decoder_kwargs = {}

            self.decoders[m.name] = build_decoder(
                lk,
                cfg=dec_cfg,
                latent_dim=self.latent_dim,
                **decoder_kwargs,
            )

        self.register_buffer("prior_mu", torch.zeros(self.latent_dim))
        self.register_buffer("prior_logvar", torch.zeros(self.latent_dim))

        self.fused_encoder_type = (getattr(cfg, "fused_encoder_type", "moe") or "moe").lower().strip()
        self.fused_require_all = bool(getattr(cfg, "fused_require_all_modalities", True))
        self.fused_modalities = list(getattr(cfg, "fused_modalities", None) or self.modality_names)

        if self.fused_encoder_type == "multimodal_transformer":
            self.fused_encoder = build_multimodal_transformer_encoder(
                uni_cfg=cfg,
                modalities=cfg.modalities,
                fused_modalities=self.fused_modalities,
            )
        else:
            self.fused_encoder = None

        self.n_label_classes = int(n_label_classes) if n_label_classes is not None else 0
        self.label_loss_weight = float(label_loss_weight)
        self.label_ignore_index = int(label_ignore_index)
        self.classify_from_mu = bool(classify_from_mu)
        self.label_head_name = str(label_head_name or getattr(cfg, "label_head_name", "label"))

        if self.n_label_classes > 0:
            dec_cfg = DecoderConfig(
                output_dim=self.n_label_classes,
                hidden_dims=[max(64, self.latent_dim)],
                dropout=float(cfg.decoder_dropout),
                batchnorm=bool(cfg.decoder_batchnorm),
            )
            self.label_decoder = build_decoder("categorical", cfg=dec_cfg, latent_dim=self.latent_dim)
        else:
            self.label_decoder = None

        self.label_names: Optional[List[str]] = None
        self.label_name_to_id: Optional[Dict[str, int]] = None

        self.use_label_encoder = bool(use_label_encoder)
        self.label_moe_weight = float(label_moe_weight)
        self.unlabeled_logvar = float(unlabeled_logvar)
        self.label_encoder_warmup = int(label_encoder_warmup)

        if self.n_label_classes > 0 and self.use_label_encoder:
            self.label_encoder = build_mlp(
                in_dim=self.n_label_classes,
                hidden_dims=[max(64, self.latent_dim)],
                out_dim=self.latent_dim * 2,
                activation=nn.ReLU(),
                dropout=float(cfg.encoder_dropout),
                batchnorm=bool(cfg.encoder_batchnorm),
            )
        else:
            self.label_encoder = None

        self.class_heads = nn.ModuleDict()
        self.class_heads_cfg: Dict[str, Dict[str, Any]] = {}
        self.head_label_names: Dict[str, List[str]] = {}
        self.head_label_name_to_id: Dict[str, Dict[str, int]] = {}

        if cfg.class_heads:
            for h in cfg.class_heads:
                if not isinstance(h, ClassHeadConfig):
                    raise TypeError(f"cfg.class_heads must contain ClassHeadConfig, got {type(h)}")

                name = str(h.name)
                n_classes = int(getattr(h, "n_classes", 0))

                head_type = (
                    getattr(h, "head_type", None)
                    or getattr(h, "type", None)
                    or getattr(h, "task", None)
                )
                head_type = (str(head_type).lower().strip() if head_type is not None else None)

                if head_type is None:
                    head_type = "categorical"

                if head_type in ("bin", "binary", "bce", "bernoulli", "logistic"):
                    head_type = "binary"
                elif head_type in ("cat", "categorical", "softmax", "multiclass", "ce"):
                    head_type = "categorical"
                else:
                    head_type = "categorical"

                hidden_dims = (
                    _as_list_int(getattr(h, "hidden_dims", None))
                    or _as_list_int(getattr(h, "head_hidden", None))
                    or _as_list_int(getattr(h, "mlp_hidden", None))
                    or _as_list_int(getattr(h, "layers", None))
                )
                if not hidden_dims:
                    hidden_dims = [max(64, self.latent_dim)]

                h_dropout = float(getattr(h, "dropout", cfg.decoder_dropout))
                h_batchnorm = bool(getattr(h, "batchnorm", cfg.decoder_batchnorm))
                h_act = _parse_activation(getattr(h, "activation", None))

                if head_type == "categorical":
                    dec_cfg = DecoderConfig(
                        output_dim=n_classes,
                        hidden_dims=list(hidden_dims),
                        dropout=h_dropout,
                        batchnorm=h_batchnorm,
                    )
                    self.class_heads[name] = build_decoder("categorical", cfg=dec_cfg, latent_dim=self.latent_dim)
                    out_dim = n_classes
                else:
                    self.class_heads[name] = _LogitsMLPHead(
                        in_dim=self.latent_dim,
                        out_dim=1,
                        hidden_dims=list(hidden_dims),
                        dropout=h_dropout,
                        batchnorm=h_batchnorm,
                        activation=h_act,
                    )
                    out_dim = 1

                self.class_heads_cfg[name] = {
                    "type": head_type,
                    "n_classes": n_classes,
                    "out_dim": out_dim,
                    "loss_weight": float(getattr(h, "loss_weight", 1.0)),
                    "ignore_index": int(getattr(h, "ignore_index", -1)),
                    "from_mu": bool(getattr(h, "from_mu", True)),
                    "warmup": int(getattr(h, "warmup", 0)),
                    "adversarial": bool(getattr(h, "adversarial", False)),
                    "adv_lambda": float(getattr(h, "adv_lambda", 1.0)),
                    "pos_weight": float(getattr(h, "pos_weight", 1.0)),
                }

        self.use_moe_gating = bool(getattr(cfg, "use_moe_gating", False))
        self.moe_gating_type = str(getattr(cfg, "moe_gating_type", "per_modality")).lower().strip()

        gate_hidden = getattr(cfg, "moe_gating_hidden", None)
        if gate_hidden is None:
            gate_hidden = [max(32, self.latent_dim // 2)]
        if isinstance(gate_hidden, (int, float)):
            gate_hidden = [int(gate_hidden)]
        self.moe_gating_hidden = [int(x) for x in gate_hidden]

        self.moe_gating_dropout = float(getattr(cfg, "moe_gating_dropout", 0.0))
        self.moe_gating_batchnorm = bool(getattr(cfg, "moe_gating_batchnorm", False))
        self.moe_gating_activation = _parse_activation(getattr(cfg, "moe_gating_activation", "relu")) or nn.ReLU()
        self.moe_gate_eps = float(getattr(cfg, "moe_gate_eps", 1e-6))

        self.gate_nets = nn.ModuleDict()
        self.shared_gate_net = None

        if self.use_moe_gating:
            if self.moe_gating_type == "shared":
                self.shared_gate_net = build_mlp(
                    in_dim=2 * self.latent_dim,
                    hidden_dims=list(self.moe_gating_hidden),
                    out_dim=1,
                    activation=self.moe_gating_activation,
                    dropout=self.moe_gating_dropout,
                    batchnorm=self.moe_gating_batchnorm,
                )
            else:
                for m in self.modality_names:
                    self.gate_nets[m] = build_mlp(
                        in_dim=2 * self.latent_dim,
                        hidden_dims=list(self.moe_gating_hidden),
                        out_dim=1,
                        activation=self.moe_gating_activation,
                        dropout=self.moe_gating_dropout,
                        batchnorm=self.moe_gating_batchnorm,
                    )

    # ----------------------------- label name utilities -----------------------------

    def set_label_names(self, label_names: List[str]) -> None:
        if self.n_label_classes <= 0:
            raise ValueError("n_label_classes=0; cannot set label names.")
        if len(label_names) != int(self.n_label_classes):
            raise ValueError(f"label_names length {len(label_names)} != n_label_classes {self.n_label_classes}")

        self.label_names = [str(x) for x in label_names]

        def norm(s: str) -> str:
            return " ".join(str(s).strip().lower().split())

        m: Dict[str, int] = {}
        for i, name in enumerate(self.label_names):
            m[name] = i
            m[norm(name)] = i
        self.label_name_to_id = m

    def set_head_label_names(self, head: str, label_names: List[str]) -> None:
        head = str(head)
        if head not in self.class_heads_cfg:
            raise KeyError(f"Unknown head {head!r}. Known heads: {list(self.class_heads_cfg)}")

        n = int(self.class_heads_cfg[head]["n_classes"])
        if len(label_names) != n:
            raise ValueError(f"Head {head!r}: label_names length {len(label_names)} != n_classes {n}")

        names = [str(x) for x in label_names]
        self.head_label_names[head] = names

        def norm(s: str) -> str:
            return " ".join(str(s).strip().lower().split())

        m: Dict[str, int] = {}
        for i, name in enumerate(names):
            m[name] = i
            m[norm(name)] = i
        self.head_label_name_to_id[head] = m

    # ----------------------------- helpers -----------------------------

    def _reparameterize(self, mu: torch.Tensor, logvar: torch.Tensor) -> torch.Tensor:
        std = torch.exp(0.5 * logvar)
        eps = torch.randn_like(std)
        return mu + eps * std

    def _kl_gaussian(
        self,
        mu_q: torch.Tensor,
        logvar_q: torch.Tensor,
        mu_p: torch.Tensor,
        logvar_p: torch.Tensor,
    ) -> torch.Tensor:
        var_q = torch.exp(logvar_q)
        var_p = torch.exp(logvar_p)
        kl = logvar_p - logvar_q + (var_q + (mu_q - mu_p) ** 2) / var_p - 1.0
        return 0.5 * kl.sum(dim=-1)

    def _alignment_loss_l2mu_pairwise(self, mu_per_mod: Dict[str, torch.Tensor]) -> torch.Tensor:
        names = list(mu_per_mod.keys())
        if len(names) < 2:
            if len(names) == 1:
                k = names[0]
                return torch.zeros(mu_per_mod[k].size(0), device=mu_per_mod[k].device)
            return torch.tensor(0.0, device=next(self.parameters()).device)

        losses = []
        for i in range(len(names)):
            for j in range(i + 1, len(names)):
                losses.append(((mu_per_mod[names[i]] - mu_per_mod[names[j]]) ** 2).sum(dim=-1))
        return torch.stack(losses, dim=0).mean(dim=0)

    def _alignment_loss_to_fused(self, mu_per_mod: Dict[str, torch.Tensor], mu_fused: torch.Tensor) -> torch.Tensor:
        if len(mu_per_mod) == 0:
            return torch.zeros(mu_fused.size(0), device=mu_fused.device)
        losses = [((mu - mu_fused) ** 2).sum(dim=-1) for mu in mu_per_mod.values()]
        return torch.stack(losses, dim=0).mean(dim=0)

    @staticmethod
    def _is_categorical_likelihood(likelihood: Optional[str]) -> bool:
        lk = (likelihood or "").lower().strip()
        return lk in ("categorical", "cat", "ce", "cross_entropy", "multinomial", "softmax")

    # -------------------------- recon scaling --------------------------

    def _recon_scale(self, mod_name: str, x: torch.Tensor, likelihood: Optional[str]) -> float:
        m_cfg = self.mod_cfg_by_name[mod_name]
        w = float(getattr(m_cfg, "recon_weight", 1.0))

        if not self.recon_normalize_by_dim:
            return w

        lk = (likelihood or "gaussian").lower().strip()
        if lk == "mse":
            return w

        D = 0
        if x is not None and torch.is_tensor(x) and x.dim() == 2:
            D = int(x.shape[1])
        if D <= 0:
            try:
                D = int(getattr(m_cfg, "input_dim", 0))
            except Exception:
                D = 0
        if D <= 0:
            D = 1

        denom = float(D) ** float(self.recon_dim_power)
        denom = max(denom, 1.0)
        return w / denom

    # ------------------------------ categorical utilities ------------------------------

    def _categorical_targets_and_mask(
        self,
        x: torch.Tensor,
        *,
        n_classes: int,
        ignore_index: int,
    ) -> Tuple[torch.Tensor, torch.Tensor]:
        if x.dim() == 2 and x.size(1) == 1:
            x = x[:, 0]

        if x.dim() == 2:
            mask = x.sum(dim=-1) > 0.5
            y = x.argmax(dim=-1).long()
            return y, mask

        y = x.view(-1)
        if y.dtype.is_floating_point:
            y = y.round()
        y = y.long()

        mask = (y != int(ignore_index))
        return y, mask

    def _encode_categorical_input_if_needed(self, mod_name: str, x: torch.Tensor) -> torch.Tensor:
        m_cfg = self.mod_cfg_by_name[mod_name]
        if not self._is_categorical_likelihood(m_cfg.likelihood):
            return x

        C = int(m_cfg.input_dim)
        ignore_index = int(getattr(m_cfg, "ignore_index", self.label_ignore_index))

        if x.dim() == 2 and x.size(1) == 1:
            x = x[:, 0]
        if x.dim() == 2:
            return x.float()

        y, mask = self._categorical_targets_and_mask(x, n_classes=C, ignore_index=ignore_index)
        B = y.shape[0]
        x_oh = torch.zeros((B, C), device=y.device, dtype=torch.float32)
        if mask.any():
            x_oh[mask] = F.one_hot(y[mask], num_classes=C).float()
        return x_oh

    # -------------------------- attention-bias helpers (optional, safe no-op) --------------------------

    def _build_distance_bias_for_permod_transformer(
        self,
        enc: nn.Module,
        x_in: torch.Tensor,
        cfg_m: Mapping[str, Any],
    ) -> Tuple[Optional[torch.Tensor], Optional[torch.Tensor], Optional[torch.Tensor]]:
        typ = str(cfg_m.get("type", "")).lower().strip()
        if typ != "distance":
            return None, None, None

        vec2tok = getattr(enc, "vec2tok", None)
        core = getattr(enc, "encoder", None)
        if vec2tok is None or core is None:
            return None, None, None

        if not hasattr(vec2tok, "build_distance_attn_bias"):
            return None, None, None

        try:
            tokens, key_padding_mask, meta = vec2tok(x_in, return_indices=True)
        except Exception:
            return None, None, None

        topk_idx = None if meta is None else meta.get("topk_idx", None)
        if topk_idx is None:
            return None, None, None

        lengthscale_bp = float(cfg_m.get("lengthscale_bp", 50_000.0))
        same_chrom_only = bool(cfg_m.get("same_chrom_only", True))
        include_cls = bool(getattr(vec2tok, "add_cls_token", False))

        try:
            attn_bias = vec2tok.build_distance_attn_bias(
                topk_idx,
                lengthscale_bp=lengthscale_bp,
                same_chrom_only=same_chrom_only,
                include_cls=include_cls,
            )
        except Exception:
            return None, None, None

        return tokens, key_padding_mask, attn_bias

    def _build_fused_attn_bias_fn(
        self,
        attn_bias_cfg: Mapping[str, Any],
        sub_x_dict: Dict[str, torch.Tensor],
    ) -> Optional[Callable[[Dict[str, Any]], Optional[torch.Tensor]]]:
        fused = self.fused_encoder
        if fused is None:
            return None

        vec2tok_map = getattr(fused, "vec2tok", None)
        if vec2tok_map is None:
            return None

        want_any = any(
            str(v.get("type", "")).lower().strip() == "distance"
            for v in attn_bias_cfg.values()
            if isinstance(v, Mapping)
        )
        if not want_any:
            return None

        def fn(meta: Dict[str, Any]) -> Optional[torch.Tensor]:
            if not meta:
                return None

            any_x = next(iter(sub_x_dict.values()))
            B = int(any_x.shape[0])

            slices = meta.get("slices_with_cls", meta.get("slices", {}))
            if not slices:
                return None

            T = 0
            for _, (a, b) in slices.items():
                T = max(T, int(b))
            if bool(meta.get("has_global_cls", False)):
                T = max(T, 1)

            bias_full = torch.zeros((B, T, T), device=any_x.device, dtype=torch.float32)

            for m, cfg_m in attn_bias_cfg.items():
                if not isinstance(cfg_m, Mapping):
                    continue
                if str(cfg_m.get("type", "")).lower().strip() != "distance":
                    continue
                if m not in slices:
                    continue
                if m not in vec2tok_map:
                    continue

                tok = vec2tok_map[m]
                if not hasattr(tok, "build_distance_attn_bias"):
                    continue

                mmeta = meta.get(m, {}) or {}
                topk_idx = mmeta.get("topk_idx", None)
                if topk_idx is None:
                    continue

                lengthscale_bp = float(cfg_m.get("lengthscale_bp", 50_000.0))
                same_chrom_only = bool(cfg_m.get("same_chrom_only", True))

                try:
                    local = tok.build_distance_attn_bias(
                        topk_idx,
                        lengthscale_bp=lengthscale_bp,
                        same_chrom_only=same_chrom_only,
                        include_cls=False,
                    )
                except Exception:
                    continue

                a, b = slices[m]
                a = int(a)
                b = int(b)
                if (b - a) != int(local.shape[1]):
                    continue

                bias_full[:, a:b, a:b] = local

            return bias_full

        return fn

    # -------------------------- encode/decode/fuse --------------------------

    def encode_modalities(
        self,
        x_dict: Dict[str, torch.Tensor],
        *,
        attn_bias_cfg: Optional[Mapping[str, Any]] = None,
    ) -> Tuple[Dict[str, torch.Tensor], Dict[str, torch.Tensor]]:
        mu_dict: Dict[str, torch.Tensor] = {}
        logvar_dict: Dict[str, torch.Tensor] = {}

        for m in self.modality_names:
            if m not in x_dict or x_dict[m] is None:
                continue

            x_in = self._encode_categorical_input_if_needed(m, x_dict[m])
            enc = self.encoders[m]

            tokens = key_padding_mask = attn_bias = None
            if attn_bias_cfg is not None and isinstance(attn_bias_cfg, Mapping):
                cfg_m = attn_bias_cfg.get(m, None)
                if isinstance(cfg_m, Mapping):
                    tokens, key_padding_mask, attn_bias = self._build_distance_bias_for_permod_transformer(enc, x_in, cfg_m)

            if tokens is not None and getattr(enc, "encoder", None) is not None:
                h = enc.encoder(
                    tokens,
                    key_padding_mask=key_padding_mask,
                    attn_bias=attn_bias,
                    return_attn=False,
                )
                mu, logvar = torch.chunk(h, 2, dim=-1)
            else:
                try:
                    mu, logvar = enc(x_in, attn_bias=attn_bias)
                except TypeError:
                    mu, logvar = enc(x_in)

            mu = self.encoder_heads[m](mu)
            logvar = torch.clamp(logvar, self.LOGVAR_MIN, self.LOGVAR_MAX)

            mu_dict[m] = mu
            logvar_dict[m] = logvar

        return mu_dict, logvar_dict

    @overload
    def mixture_of_experts(
        self,
        mu_dict: Dict[str, torch.Tensor],
        logvar_dict: Dict[str, torch.Tensor],
        *,
        return_weights: Literal[False] = False,
        return_logits: Literal[False] = False,
        modality_order: Optional[List[str]] = None,
    ) -> Tuple[torch.Tensor, torch.Tensor]:
        ...

    @overload
    def mixture_of_experts(
        self,
        mu_dict: Dict[str, torch.Tensor],
        logvar_dict: Dict[str, torch.Tensor],
        *,
        return_weights: Literal[True],
        return_logits: Literal[False] = False,
        modality_order: Optional[List[str]] = None,
    ) -> Tuple[torch.Tensor, torch.Tensor, torch.Tensor]:
        ...

    @overload
    def mixture_of_experts(
        self,
        mu_dict: Dict[str, torch.Tensor],
        logvar_dict: Dict[str, torch.Tensor],
        *,
        return_weights: Literal[True],
        return_logits: Literal[True],
        modality_order: Optional[List[str]] = None,
    ) -> Tuple[torch.Tensor, torch.Tensor, torch.Tensor, torch.Tensor]:
        ...

    def mixture_of_experts(
        self,
        mu_dict: Dict[str, torch.Tensor],
        logvar_dict: Dict[str, torch.Tensor],
        *,
        return_weights: bool = False,
        return_logits: bool = False,
        modality_order: Optional[List[str]] = None,
    ):
        if len(mu_dict) == 0:
            raise ValueError("mixture_of_experts requires at least one modality posterior.")

        if return_logits and not return_weights:
            raise ValueError("return_logits=True requires return_weights=True.")

        if modality_order is None:
            modality_order = [m for m in self.modality_names if (m in mu_dict and m in logvar_dict)]
            if not modality_order:
                modality_order = sorted(list(mu_dict.keys()))

        mus = [mu_dict[m] for m in modality_order]
        logvars = [logvar_dict[m] for m in modality_order]

        if len(mus) == 1:
            mu_comb = mus[0]
            logvar_comb = logvars[0]
            if not return_weights and not return_logits:
                return mu_comb, logvar_comb

            B = int(mu_comb.shape[0])
            w = torch.ones((B, 1), device=mu_comb.device, dtype=mu_comb.dtype)
            logits = torch.zeros((B, 1), device=mu_comb.device, dtype=mu_comb.dtype)

            if return_weights and return_logits:
                return mu_comb, logvar_comb, w, logits
            return mu_comb, logvar_comb, w

        precisions = [torch.exp(-lv) for lv in logvars]

        logits_mat = None
        weights = None

        if self.use_moe_gating:
            logit_list = []
            for m, mu_m, lv_m in zip(modality_order, mus, logvars):
                feat = torch.cat([mu_m, lv_m], dim=-1)
                if self.shared_gate_net is not None:
                    gl = self.shared_gate_net(feat)
                else:
                    gn = self.gate_nets[m] if (m in self.gate_nets) else None
                    if gn is None:
                        gl = torch.zeros((feat.shape[0], 1), device=feat.device, dtype=feat.dtype)
                    else:
                        gl = gn(feat)
                logit_list.append(gl.view(-1))

            logits_mat = torch.stack(logit_list, dim=1)
            weights = F.softmax(logits_mat, dim=1)

            w_list = [weights[:, i].unsqueeze(-1) for i in range(weights.shape[1])]
            precisions = [(w_i + self.moe_gate_eps) * p for w_i, p in zip(w_list, precisions)]

        precision_sum = torch.stack(precisions, dim=0).sum(dim=0).clamp_min(self.EPS)
        mu_weighted = torch.stack([m * p for m, p in zip(mus, precisions)], dim=0).sum(dim=0)
        mu_comb = mu_weighted / precision_sum
        var_comb = 1.0 / precision_sum
        logvar_comb = torch.log(var_comb.clamp_min(self.EPS))

        if not return_weights and not return_logits:
            return mu_comb, logvar_comb

        if weights is None:
            B = int(mu_comb.shape[0])
            M = len(mus)
            weights = torch.full((B, M), 1.0 / float(M), device=mu_comb.device, dtype=mu_comb.dtype)
        if logits_mat is None:
            logits_mat = torch.zeros_like(weights)

        if return_weights and return_logits:
            return mu_comb, logvar_comb, weights, logits_mat
        return mu_comb, logvar_comb, weights

    def decode_modalities(self, z: torch.Tensor) -> Dict[str, Any]:
        return {m: self.decoders[m](z) for m in self.modality_names}

    # -------------------- fused posterior --------------------

    def _present_fused_modalities(self, x_dict: Dict[str, torch.Tensor]) -> List[str]:
        return [m for m in self.fused_modalities if (m in x_dict) and (x_dict[m] is not None)]

    def _can_use_fused_encoder(self, x_dict: Dict[str, torch.Tensor]) -> bool:
        if self.fused_encoder is None:
            return False
        if not self.fused_modalities:
            return False

        present = self._present_fused_modalities(x_dict)
        if self.fused_require_all:
            return len(present) == len(self.fused_modalities)
        return len(present) >= 1

    def _compute_fused_posterior(
        self,
        x_dict: Dict[str, torch.Tensor],
        *,
        attn_bias_cfg: Optional[Mapping[str, Any]] = None,
    ) -> Tuple[torch.Tensor, torch.Tensor]:
        assert self.fused_encoder is not None
        present = self._present_fused_modalities(x_dict)
        sub = {m: x_dict[m] for m in present}

        attn_bias_fn = None
        want_meta = False
        if attn_bias_cfg is not None and isinstance(attn_bias_cfg, Mapping):
            attn_bias_fn = self._build_fused_attn_bias_fn(attn_bias_cfg, sub)
            want_meta = attn_bias_fn is not None

        out = self.fused_encoder(sub, return_token_meta=want_meta, attn_bias_fn=attn_bias_fn)
        if want_meta:
            mu_f, logvar_f, _meta = out  # type: ignore[misc]
        else:
            mu_f, logvar_f = out  # type: ignore[misc]

        logvar_f = torch.clamp(logvar_f, self.LOGVAR_MIN, self.LOGVAR_MAX)
        return mu_f, logvar_f

    # -------------------- label expert --------------------

    def _encode_labels_as_expert(self, y: torch.Tensor, B: int, device: torch.device) -> Tuple[torch.Tensor, torch.Tensor]:
        mu_y = torch.zeros(B, self.latent_dim, device=device)
        logvar_y = torch.full((B, self.latent_dim), float(self.unlabeled_logvar), device=device)

        if self.label_encoder is None or y is None:
            return mu_y, logvar_y

        y = y.long()
        mask = (y >= 0) & (y != self.label_ignore_index)
        if not mask.any():
            return mu_y, logvar_y

        y_oh = F.one_hot(y[mask], num_classes=self.n_label_classes).float()
        h = self.label_encoder(y_oh)
        mu_l, logvar_l = torch.chunk(h, 2, dim=-1)
        logvar_l = torch.clamp(logvar_l, self.LOGVAR_MIN, self.LOGVAR_MAX)

        w = float(self.label_moe_weight)
        if w != 1.0:
            logvar_l = logvar_l - math.log(max(w, 1e-8))

        mu_y[mask] = mu_l
        logvar_y[mask] = logvar_l
        return mu_y, logvar_y

    def _extract_legacy_y(self, y: Optional[YType]) -> Optional[torch.Tensor]:
        if y is None:
            return None
        if isinstance(y, Mapping):
            v = y.get(self.label_head_name, None)
            return v.long() if v is not None else None
        return y.long()

    def _grad_reverse(self, x: torch.Tensor, lambd: float = 1.0) -> torch.Tensor:
        return _GradReverseFn.apply(x, float(lambd))

    # -------------------- UPDATED multihead losses --------------------

    def _apply_multihead_losses(
        self,
        *,
        mu_z: torch.Tensor,
        z: torch.Tensor,
        y: Optional[YType],
        epoch: int,
        loss: torch.Tensor,
        loss_annealed: torch.Tensor,
        loss_fixed: torch.Tensor,
    ) -> Tuple[torch.Tensor, torch.Tensor, torch.Tensor, Dict[str, torch.Tensor], Dict[str, torch.Tensor]]:
        head_logits: Dict[str, torch.Tensor] = {}
        head_loss_means: Dict[str, torch.Tensor] = {}

        if y is None or not isinstance(y, Mapping) or len(self.class_heads) == 0:
            return loss, loss_annealed, loss_fixed, head_logits, head_loss_means

        B = int(mu_z.size(0))
        device = mu_z.device

        for name, head in self.class_heads.items():
            cfg_h = self.class_heads_cfg[name]
            if int(epoch) < int(cfg_h["warmup"]):
                continue

            y_h = y.get(name, None)
            if y_h is None:
                continue

            z_in = mu_z if bool(cfg_h["from_mu"]) else z
            if bool(cfg_h.get("adversarial", False)):
                z_in = self._grad_reverse(z_in, cfg_h.get("adv_lambda", 1.0))

            dec_out = head(z_in)
            logits = dec_out.get("logits", dec_out.get("logit", dec_out.get("scores", dec_out))) if isinstance(dec_out, dict) else dec_out

            head_logits[name] = logits

            ignore_index = int(cfg_h["ignore_index"])
            per_cell = torch.zeros(B, device=device)

            head_type = str(cfg_h.get("type", "categorical")).lower().strip()

            if head_type == "binary":
                yy = y_h.view(-1)
                yy_f = yy.float()
                mask = (yy != ignore_index) & (yy >= 0)
                if mask.any():
                    logit_1d = logits.view(-1)
                    pos_w = float(cfg_h.get("pos_weight", 1.0))
                    if pos_w != 1.0:
                        pos_weight = torch.tensor(pos_w, device=logit_1d.device, dtype=logit_1d.dtype)
                        bce = F.binary_cross_entropy_with_logits(
                            logit_1d[mask], yy_f[mask], reduction="none", pos_weight=pos_weight
                        )
                    else:
                        bce = F.binary_cross_entropy_with_logits(logit_1d[mask], yy_f[mask], reduction="none")
                    per_cell[mask] = bce
            else:
                yy = y_h.long().view(-1)
                mask = (yy != ignore_index) & (yy >= 0)
                if mask.any():
                    per_cell[mask] = F.cross_entropy(logits[mask], yy[mask], reduction="none")

            w = float(cfg_h["loss_weight"])
            if w != 0.0:
                loss = loss + w * per_cell
                loss_annealed = loss_annealed + w * per_cell
                loss_fixed = loss_fixed + w * per_cell

            head_loss_means[name] = per_cell.mean()

        return loss, loss_annealed, loss_fixed, head_logits, head_loss_means

    # ------------------------------ forward dispatcher ------------------------------

    def forward(
        self,
        x_dict: Dict[str, torch.Tensor],
        epoch: int = 0,
        y: Optional[YType] = None,
        attn_bias_cfg: Optional[Mapping[str, Any]] = None,
        recon_targets: Optional[Dict[str, Any]] = None,
        **kwargs: Any,
    ) -> Dict[str, torch.Tensor]:
        mode = (self.loss_mode or "v2").lower()
        if mode in ("v1", "paper", "cross"):
            return self._forward_v1(
                x_dict=x_dict,
                epoch=epoch,
                y=y,
                attn_bias_cfg=attn_bias_cfg,
                recon_targets=recon_targets,
            )
        if mode in ("v2", "lite", "light", "moe", "poe", "fused"):
            return self._forward_v2(
                x_dict=x_dict,
                epoch=epoch,
                y=y,
                attn_bias_cfg=attn_bias_cfg,
                recon_targets=recon_targets,
            )
        raise ValueError(f"Unknown loss_mode={self.loss_mode!r}.")

    # ------------------------------ v2 / lite ------------------------------

    def _forward_v2(
        self,
        x_dict: Dict[str, torch.Tensor],
        epoch: int = 0,
        y: Optional[YType] = None,
        attn_bias_cfg: Optional[Mapping[str, Any]] = None,
        recon_targets: Optional[Dict[str, Any]] = None,
    ) -> Dict[str, torch.Tensor]:
        mu_dict, logvar_dict = self.encode_modalities(x_dict, attn_bias_cfg=attn_bias_cfg)
        if len(mu_dict) == 0:
            raise ValueError("At least one modality must be present in x_dict.")

        use_fused = (self.fused_encoder_type == "multimodal_transformer") and self._can_use_fused_encoder(x_dict)
        mu_fused: Optional[torch.Tensor] = None
        logvar_fused: Optional[torch.Tensor] = None

        if use_fused:
            mu_fused, logvar_fused = self._compute_fused_posterior(x_dict, attn_bias_cfg=attn_bias_cfg)
            mu_z, logvar_z = mu_fused, logvar_fused
        else:
            mu_z, logvar_z = self.mixture_of_experts(mu_dict, logvar_dict)

        y_legacy = self._extract_legacy_y(y)
        if self.label_encoder is not None and y_legacy is not None and epoch >= self.label_encoder_warmup:
            B = mu_z.shape[0]
            device = mu_z.device
            mu_y, logvar_y = self._encode_labels_as_expert(y=y_legacy, B=B, device=device)

            base_mu_dict = {"__base__": mu_z, "__label__": mu_y}
            base_lv_dict = {"__base__": logvar_z, "__label__": logvar_y}
            mu_z, logvar_z = self.mixture_of_experts(base_mu_dict, base_lv_dict)

        z = self._reparameterize(mu_z, logvar_z)
        xhat_dict = self.decode_modalities(z)

        recon_total: Optional[torch.Tensor] = None
        recon_losses: Dict[str, torch.Tensor] = {}

        for name, m_cfg in self.mod_cfg_by_name.items():
            if name not in x_dict or x_dict[name] is None:
                continue

            x_recon = x_dict[name]
            if recon_targets is not None and name in recon_targets and recon_targets[name] is not None:
                x_recon = recon_targets[name]

            loss_m = self._recon_loss(
                x=x_recon,
                raw_dec_out=xhat_dict[name],
                likelihood=m_cfg.likelihood,
                mod_name=name,
            )

            s = self._recon_scale(name, x_dict[name], m_cfg.likelihood)
            if s != 1.0:
                loss_m = loss_m * float(s)

            recon_losses[name] = loss_m
            recon_total = loss_m if recon_total is None else (recon_total + loss_m)

        if recon_total is None:
            raise RuntimeError("No modalities in x_dict produced recon loss.")

        mu_p = self.prior_mu.expand_as(mu_z)
        logvar_p = self.prior_logvar.expand_as(logvar_z)
        kl = self._kl_gaussian(mu_z, logvar_z, mu_p, logvar_p)

        real_mu = {k: v for k, v in mu_dict.items() if not k.startswith("__")}
        if use_fused and (mu_fused is not None):
            align_loss = self._alignment_loss_to_fused(real_mu, mu_fused)
        else:
            align_loss = self._alignment_loss_l2mu_pairwise(real_mu)

        beta_t = self._anneal_weight(epoch, self.cfg.kl_anneal_start, self.cfg.kl_anneal_end, self.beta_max)
        gamma_t = self._anneal_weight(epoch, self.cfg.align_anneal_start, self.cfg.align_anneal_end, self.gamma_max)

        beta_used = beta_t if self.training else self.beta_max
        gamma_used = gamma_t if self.training else self.gamma_max

        loss_annealed = recon_total + beta_t * kl + gamma_t * align_loss
        loss_fixed = recon_total + self.beta_max * kl + self.gamma_max * align_loss
        loss = recon_total + beta_used * kl + gamma_used * align_loss

        class_loss = None
        class_logits = None
        if self.label_decoder is not None:
            z_for_cls = mu_z if self.classify_from_mu else z
            dec_out = self.label_decoder(z_for_cls)
            class_logits = dec_out["logits"] if isinstance(dec_out, dict) and "logits" in dec_out else dec_out

            if y_legacy is not None:
                B = loss.shape[0]
                yy = y_legacy.long()
                mask = (yy >= 0) & (yy != self.label_ignore_index)
                per_cell = torch.zeros(B, device=loss.device)
                if mask.any():
                    per_cell[mask] = F.cross_entropy(class_logits[mask], yy[mask], reduction="none")
                class_loss = per_cell

                loss = loss + self.label_loss_weight * class_loss
                loss_annealed = loss_annealed + self.label_loss_weight * class_loss
                loss_fixed = loss_fixed + self.label_loss_weight * class_loss

        loss, loss_annealed, loss_fixed, head_logits, head_loss_means = self._apply_multihead_losses(
            mu_z=mu_z,
            z=z,
            y=y,
            epoch=epoch,
            loss=loss,
            loss_annealed=loss_annealed,
            loss_fixed=loss_fixed,
        )

        out: Dict[str, Any] = {
            "loss": loss.mean(),
            "recon_total": recon_total.mean(),
            "kl": kl.mean(),
            "align": align_loss.mean() if torch.is_tensor(align_loss) else torch.tensor(float(align_loss), device=loss.device),
            "mu_z": mu_z,
            "logvar_z": logvar_z,
            "z": z,
            "xhat": xhat_dict,
            "mu_dict": mu_dict,
            "logvar_dict": logvar_dict,
            "recon_per_modality": {k: v.mean() for k, v in recon_losses.items()},
            "beta": torch.tensor(beta_t, device=loss.device),
            "gamma": torch.tensor(gamma_t, device=loss.device),
            "beta_used": torch.tensor(beta_used, device=loss.device),
            "gamma_used": torch.tensor(gamma_used, device=loss.device),
            "loss_annealed": loss_annealed.mean(),
            "loss_fixed": loss_fixed.mean(),
            "used_fused_encoder": torch.tensor(1.0 if use_fused else 0.0, device=loss.device),
        }
        if class_loss is not None:
            out["class_loss"] = class_loss.mean()
        if class_logits is not None:
            out["class_logits"] = class_logits
        if head_logits:
            out["head_logits"] = head_logits
        if head_loss_means:
            out["head_losses"] = head_loss_means
        return out

    # ------------------------------ v1 ------------------------------

    def _forward_v1(
        self,
        x_dict: Dict[str, torch.Tensor],
        epoch: int = 0,
        y: Optional[YType] = None,
        attn_bias_cfg: Optional[Mapping[str, Any]] = None,
        recon_targets: Optional[Dict[str, Any]] = None,
    ) -> Dict[str, torch.Tensor]:
        mu_dict, logvar_dict = self.encode_modalities(x_dict, attn_bias_cfg=attn_bias_cfg)
        if len(mu_dict) == 0:
            raise ValueError("At least one modality must be present in x_dict.")

        present = list(mu_dict.keys())
        K = len(present)

        use_fused = (self.fused_encoder_type == "multimodal_transformer") and self._can_use_fused_encoder(x_dict)
        if use_fused:
            mu_moe, logvar_moe = self._compute_fused_posterior(x_dict, attn_bias_cfg=attn_bias_cfg)
        else:
            mu_moe, logvar_moe = self.mixture_of_experts(mu_dict, logvar_dict)

        z_moe = self._reparameterize(mu_moe, logvar_moe)
        z_mod = {m: self._reparameterize(mu_dict[m], logvar_dict[m]) for m in present}

        def recon_from_z_to_target(z_src: torch.Tensor, target_mod: str) -> torch.Tensor:
            raw = self.decoders[target_mod](z_src)
            m_cfg = self.mod_cfg_by_name[target_mod]
            x_recon = x_dict[target_mod]
            if recon_targets is not None and target_mod in recon_targets and recon_targets[target_mod] is not None:
                x_recon = recon_targets[target_mod]
            loss_t = self._recon_loss(
                x=x_recon,
                raw_dec_out=raw,
                likelihood=m_cfg.likelihood,
                mod_name=target_mod,
            )
            s = self._recon_scale(target_mod, x_dict[target_mod], m_cfg.likelihood)
            if s != 1.0:
                loss_t = loss_t * float(s)
            return loss_t

        device = next(iter(mu_dict.values())).device
        B = next(iter(mu_dict.values())).shape[0]

        recon_per_target: Dict[str, torch.Tensor] = {m: torch.zeros(B, device=device) for m in present}
        recon_counts: Dict[str, int] = {m: 0 for m in present}

        recon_total: Optional[torch.Tensor] = None
        n_terms = 0

        def add_term(t: torch.Tensor, tgt: str):
            nonlocal recon_total, n_terms
            recon_per_target[tgt] = recon_per_target[tgt] + t
            recon_counts[tgt] += 1
            recon_total = t if recon_total is None else (recon_total + t)
            n_terms += 1

        v1_recon = self.v1_recon

        if v1_recon in ("avg", "both", "paper", "self+cross", "self_cross", "hybrid"):
            if K <= 1:
                tgt = present[0]
                add_term(recon_from_z_to_target(z_mod[tgt], tgt), tgt=tgt)
            else:
                w_self = 0.5 / float(K)
                w_cross = 0.5 / float(K * (K - 1))
                for tgt in present:
                    add_term(w_self * recon_from_z_to_target(z_mod[tgt], tgt), tgt=tgt)
                for src in present:
                    for tgt in present:
                        if src == tgt:
                            continue
                        add_term(w_cross * recon_from_z_to_target(z_mod[src], tgt), tgt=tgt)

        elif v1_recon.startswith("src:"):
            src_name = v1_recon.split("src:", 1)[1].strip()
            if src_name not in z_mod:
                raise ValueError(f"v1_recon={self.v1_recon!r} but '{src_name}' not present. Present={present}")
            for tgt in present:
                add_term(recon_from_z_to_target(z_mod[src_name], tgt), tgt=tgt)

        elif v1_recon == "self":
            for tgt in present:
                add_term(recon_from_z_to_target(z_mod[tgt], tgt), tgt=tgt)

        elif v1_recon in ("avg_z", "mean_z", "average_z"):
            z_avg = torch.stack([z_mod[m] for m in present], dim=0).mean(dim=0)
            for tgt in present:
                add_term(recon_from_z_to_target(z_avg, tgt), tgt=tgt)
            z_moe = z_avg

        elif v1_recon in ("moe", "poe", "fused"):
            for tgt in present:
                add_term(recon_from_z_to_target(z_moe, tgt), tgt=tgt)

        else:
            if K >= 2:
                for src in present:
                    for tgt in present:
                        if src == tgt:
                            continue
                        add_term(recon_from_z_to_target(z_mod[src], tgt), tgt=tgt)
            else:
                tgt = present[0]
                add_term(recon_from_z_to_target(z_mod[tgt], tgt), tgt=tgt)

            if self.v1_recon_mix > 0.0 and K >= 2:
                mix = float(self.v1_recon_mix)
                z_avg = torch.stack([z_mod[m] for m in present], dim=0).mean(dim=0)
                for tgt in present:
                    add_term(mix * recon_from_z_to_target(z_avg, tgt), tgt=tgt)

        if recon_total is None or n_terms == 0:
            raise RuntimeError("v1 reconstruction produced no loss terms.")

        weighted_mode = v1_recon in ("avg", "both", "paper", "self+cross", "self_cross", "hybrid")
        if self.normalize_v1_terms and (not weighted_mode):
            recon_total = recon_total / float(n_terms)

        recon_per_target_mean: Dict[str, torch.Tensor] = {}
        for tgt in present:
            ct = max(int(recon_counts[tgt]), 1)
            recon_per_target_mean[tgt] = recon_per_target[tgt] / float(ct)

        mu_p = self.prior_mu.expand_as(mu_dict[present[0]])
        logvar_p = self.prior_logvar.expand_as(logvar_dict[present[0]])

        kl_terms = [self._kl_gaussian(mu_dict[m], logvar_dict[m], mu_p, logvar_p) for m in present]
        kl = torch.stack(kl_terms, dim=0).sum(dim=0)
        if self.normalize_v1_terms:
            kl = kl / float(max(K, 1))

        if K < 2:
            cross_kl = torch.zeros_like(kl)
        else:
            cross_terms = []
            for i in range(K):
                for j in range(K):
                    if i == j:
                        continue
                    mi, lvi = mu_dict[present[i]], logvar_dict[present[i]]
                    mj, lvj = mu_dict[present[j]], logvar_dict[present[j]]
                    cross_terms.append(self._kl_gaussian(mi, lvi, mj, lvj))
            cross_kl = torch.stack(cross_terms, dim=0).sum(dim=0)
            if self.normalize_v1_terms:
                cross_kl = cross_kl / float(K * (K - 1))

        beta_t = self._anneal_weight(epoch, self.cfg.kl_anneal_start, self.cfg.kl_anneal_end, self.beta_max)
        gamma_t = self._anneal_weight(epoch, self.cfg.align_anneal_start, self.cfg.align_anneal_end, self.gamma_max)

        beta_used = beta_t if self.training else self.beta_max
        gamma_used = gamma_t if self.training else self.gamma_max

        loss_annealed = recon_total + beta_t * kl + gamma_t * cross_kl
        loss_fixed = recon_total + self.beta_max * kl + self.gamma_max * cross_kl
        loss = recon_total + beta_used * kl + gamma_used * cross_kl

        y_legacy = self._extract_legacy_y(y)
        class_loss = None
        class_logits = None
        if self.label_decoder is not None:
            z_for_cls = mu_moe if self.classify_from_mu else z_moe
            dec_out = self.label_decoder(z_for_cls)
            class_logits = dec_out["logits"] if isinstance(dec_out, dict) and "logits" in dec_out else dec_out

            if y_legacy is not None:
                yy = y_legacy.long()
                mask = (yy >= 0) & (yy != self.label_ignore_index)
                per_cell = torch.zeros(B, device=loss.device)
                if mask.any():
                    per_cell[mask] = F.cross_entropy(class_logits[mask], yy[mask], reduction="none")
                class_loss = per_cell

                loss = loss + self.label_loss_weight * class_loss
                loss_annealed = loss_annealed + self.label_loss_weight * class_loss
                loss_fixed = loss_fixed + self.label_loss_weight * class_loss

        loss, loss_annealed, loss_fixed, head_logits, head_loss_means = self._apply_multihead_losses(
            mu_z=mu_moe,
            z=z_moe,
            y=y,
            epoch=epoch,
            loss=loss,
            loss_annealed=loss_annealed,
            loss_fixed=loss_fixed,
        )

        xhat_dict = self.decode_modalities(z_moe)

        out: Dict[str, Any] = {
            "loss": loss.mean(),
            "recon_total": recon_total.mean(),
            "kl": kl.mean(),
            "align": cross_kl.mean(),
            "cross_kl": cross_kl.mean(),
            "mu_z": mu_moe,
            "logvar_z": logvar_moe,
            "z": z_moe,
            "xhat": xhat_dict,
            "mu_dict": mu_dict,
            "logvar_dict": logvar_dict,
            "recon_per_modality": {k: v.mean() for k, v in recon_per_target_mean.items()},
            "beta": torch.tensor(beta_t, device=loss.device),
            "gamma": torch.tensor(gamma_t, device=loss.device),
            "beta_used": torch.tensor(beta_used, device=loss.device),
            "gamma_used": torch.tensor(gamma_used, device=loss.device),
            "loss_annealed": loss_annealed.mean(),
            "loss_fixed": loss_fixed.mean(),
            "v1_recon_terms": torch.tensor(float(n_terms), device=loss.device),
            "used_fused_encoder": torch.tensor(1.0 if use_fused else 0.0, device=loss.device),
        }
        if class_loss is not None:
            out["class_loss"] = class_loss.mean()
        if class_logits is not None:
            out["class_logits"] = class_logits
        if head_logits:
            out["head_logits"] = head_logits
        if head_loss_means:
            out["head_losses"] = head_loss_means
        return out

    # ------------------------------ reconstruction loss ------------------------------

    def _unwrap_decoder_out(self, dec_out: Any) -> Any:
        if isinstance(dec_out, (tuple, list)):
            if len(dec_out) != 1:
                raise TypeError(f"Unsupported decoder output container of length {len(dec_out)}: {type(dec_out)!r}")
            dec_out = dec_out[0]
        if torch.is_tensor(dec_out):
            return dec_out
        if isinstance(dec_out, dict):
            out = dict(dec_out)
            if "mu" not in out and "mean" in out:
                out["mu"] = out["mean"]
            return out
        raise TypeError(f"Unsupported decoder output type: {type(dec_out)!r}")

    @staticmethod
    def _nb_nll(x: torch.Tensor, mu: torch.Tensor, theta: torch.Tensor, eps: float = 1e-8) -> torch.Tensor:
        mu = mu.clamp(min=eps)
        theta = theta.clamp(min=eps)
        t1 = torch.lgamma(x + theta) - torch.lgamma(theta) - torch.lgamma(x + 1.0)
        t2 = theta * (torch.log(theta) - torch.log(theta + mu))
        t3 = x * (torch.log(mu) - torch.log(theta + mu))
        return -(t1 + t2 + t3)

    @staticmethod
    def _zinb_nll(x: torch.Tensor, mu: torch.Tensor, theta: torch.Tensor, logit_pi: torch.Tensor, eps: float = 1e-8) -> torch.Tensor:
        mu = mu.clamp(min=eps)
        theta = theta.clamp(min=eps)
        pi = torch.sigmoid(logit_pi)
        t1 = torch.lgamma(x + theta) - torch.lgamma(theta) - torch.lgamma(x + 1.0)
        t2 = theta * (torch.log(theta) - torch.log(theta + mu))
        t3 = x * (torch.log(mu) - torch.log(theta + mu))
        log_nb = t1 + t2 + t3
        is_zero = (x < eps)
        log_prob_pos = torch.log1p(-pi + eps) + log_nb
        log_nb_zero = theta * (torch.log(theta) - torch.log(theta + mu))
        log_prob_zero = torch.log(pi + (1.0 - pi) * torch.exp(log_nb_zero) + eps)
        log_prob = torch.where(is_zero, log_prob_zero, log_prob_pos)
        return -log_prob

    @staticmethod
    def _beta_nll(
        x: torch.Tensor,
        alpha: torch.Tensor,
        beta: torch.Tensor,
        eps: float = 1e-8,
    ) -> torch.Tensor:
        x = x.clamp(min=eps, max=1.0 - eps)
        alpha = alpha.clamp(min=eps)
        beta = beta.clamp(min=eps)

        log_norm = torch.lgamma(alpha) + torch.lgamma(beta) - torch.lgamma(alpha + beta)
        log_prob = (alpha - 1.0) * torch.log(x) + (beta - 1.0) * torch.log(1.0 - x) - log_norm
        return -log_prob

    @staticmethod
    def _binomial_nll(
        successes: torch.Tensor,
        total_count: torch.Tensor,
        logits: torch.Tensor,
    ) -> torch.Tensor:
        log_prob = -F.binary_cross_entropy_with_logits(logits, successes / total_count.clamp_min(1.0), reduction="none") * total_count
        # The BCE-with-logits trick above gives the Bernoulli cross-entropy scaled by n,
        # but omits the combinatorial term log(n choose k). Add it back for true Binomial NLL:
        comb = (
            torch.lgamma(total_count + 1.0)
            - torch.lgamma(successes + 1.0)
            - torch.lgamma((total_count - successes) + 1.0)
        )
        return -(comb + log_prob)

    @staticmethod
    def _beta_binomial_nll(
        successes: torch.Tensor,
        total_count: torch.Tensor,
        alpha: torch.Tensor,
        beta: torch.Tensor,
        eps: float = 1e-8,
    ) -> torch.Tensor:
        successes = successes.clamp_min(0.0)
        total_count = total_count.clamp_min(0.0)
        alpha = alpha.clamp_min(eps)
        beta = beta.clamp_min(eps)

        failures = (total_count - successes).clamp_min(0.0)

        log_comb = (
            torch.lgamma(total_count + 1.0)
            - torch.lgamma(successes + 1.0)
            - torch.lgamma(failures + 1.0)
        )

        log_beta_term = (
            torch.lgamma(successes + alpha)
            + torch.lgamma(failures + beta)
            - torch.lgamma(total_count + alpha + beta)
            + torch.lgamma(alpha + beta)
            - torch.lgamma(alpha)
            - torch.lgamma(beta)
        )

        return -(log_comb + log_beta_term)

    def _extract_count_target(
        self,
        x: Any,
        *,
        mod_name: str,
        ref_device: torch.device,
        ref_dtype: torch.dtype,
    ) -> Tuple[torch.Tensor, torch.Tensor, torch.Tensor]:
        """
        Parse count-with-coverage targets for Binomial/Beta-Binomial losses.

        Accepts either:
          - dict with aliases for successes and totals, e.g.
              {'successes': m, 'total_count': n}
              {'m': m, 'n': n}
              {'counts': m, 'coverage': n}
          - tuple/list length 2: (successes, total_count)

        Returns:
          successes, total_count, valid_mask  (all shaped [B, D], mask bool)
        """
        successes = None
        total_count = None

        if isinstance(x, Mapping):
            for k in ("successes", "m", "count", "counts", "k"):
                if k in x and x[k] is not None:
                    successes = x[k]
                    break
            for k in ("total_count", "total_counts", "n", "coverage", "depth", "trials"):
                if k in x and x[k] is not None:
                    total_count = x[k]
                    break
        elif isinstance(x, (tuple, list)) and len(x) == 2:
            successes, total_count = x[0], x[1]

        if successes is None or total_count is None:
            raise ValueError(
                f"Modality {mod_name!r} with binomial/beta_binomial likelihood requires recon target "
                f"with successes + total_count (e.g., {{'successes': m, 'total_count': n}})."
            )

        if not torch.is_tensor(successes):
            successes = torch.as_tensor(successes, device=ref_device, dtype=ref_dtype)
        else:
            successes = successes.to(device=ref_device, dtype=ref_dtype)

        if not torch.is_tensor(total_count):
            total_count = torch.as_tensor(total_count, device=ref_device, dtype=ref_dtype)
        else:
            total_count = total_count.to(device=ref_device, dtype=ref_dtype)

        if successes.dim() == 1:
            successes = successes.unsqueeze(-1)
        if total_count.dim() == 1:
            total_count = total_count.unsqueeze(-1)

        if successes.shape != total_count.shape:
            raise ValueError(
                f"Modality {mod_name!r}: successes shape {tuple(successes.shape)} != total_count shape {tuple(total_count.shape)}"
            )

        successes = successes.clamp_min(0.0)
        total_count = total_count.clamp_min(0.0)
        successes = torch.minimum(successes, total_count)

        valid = total_count > 0.0
        return successes, total_count, valid

    def _recon_loss(self, x: Any, raw_dec_out: Any, likelihood: str, mod_name: str) -> torch.Tensor:
        likelihood = (likelihood or "gaussian").lower().strip()
        dec_out = self._unwrap_decoder_out(raw_dec_out)

        if self._is_categorical_likelihood(likelihood):
            m_cfg = self.mod_cfg_by_name[mod_name]
            C = int(m_cfg.input_dim)
            ignore_index = int(getattr(m_cfg, "ignore_index", self.label_ignore_index))

            logits = dec_out["logits"] if isinstance(dec_out, dict) else dec_out
            y, mask = self._categorical_targets_and_mask(x, n_classes=C, ignore_index=ignore_index)

            nll = torch.zeros(y.shape[0], device=logits.device)
            if mask.any():
                nll[mask] = F.cross_entropy(logits[mask], y[mask], reduction="none")
            return nll

        if likelihood in ("gaussian", "normal", "mse", "gaussian_diag"):
            if not torch.is_tensor(x):
                raise TypeError(f"Gaussian-like recon expects tensor target for modality {mod_name!r}; got {type(x)!r}")

            if isinstance(dec_out, dict) and ("mean" in dec_out) and ("logvar" in dec_out):
                mean = dec_out["mean"]
                logvar = dec_out["logvar"].clamp(self.LOGVAR_MIN, self.LOGVAR_MAX)
                var = torch.exp(logvar)
                nll = 0.5 * (logvar + (x - mean) ** 2 / (var + self.EPS))
                return nll.sum(dim=-1)

            pred = dec_out["mean"] if isinstance(dec_out, dict) and ("mean" in dec_out) else dec_out
            if likelihood == "mse":
                return ((x - pred) ** 2).mean(dim=-1)
            return ((x - pred) ** 2).sum(dim=-1)

        if likelihood == "beta":
            if not torch.is_tensor(x):
                raise TypeError(f"Beta recon expects tensor fraction target for modality {mod_name!r}; got {type(x)!r}")

            if not isinstance(dec_out, dict):
                raise ValueError("Beta recon expects dict decoder output with alpha/beta or mu/conc params.")

            if ("alpha" in dec_out) and ("beta" in dec_out):
                alpha = dec_out["alpha"]
                beta = dec_out["beta"]
            elif ("mu" in dec_out) and ("conc" in dec_out):
                mu = dec_out["mu"].clamp(self.EPS, 1.0 - self.EPS)
                conc = dec_out["conc"].clamp_min(self.EPS)
                alpha = (mu * conc).clamp_min(self.EPS)
                beta = ((1.0 - mu) * conc).clamp_min(self.EPS)
            elif ("mu_logits" in dec_out) and ("log_conc" in dec_out):
                mu = torch.sigmoid(dec_out["mu_logits"]).clamp(self.EPS, 1.0 - self.EPS)
                conc = torch.exp(dec_out["log_conc"]).clamp_min(self.EPS)
                alpha = (mu * conc).clamp_min(self.EPS)
                beta = ((1.0 - mu) * conc).clamp_min(self.EPS)
            else:
                raise ValueError("Beta recon expects alpha/beta or (mu,conc) or (mu_logits,log_conc).")

            return self._beta_nll(x, alpha, beta, eps=self.EPS).sum(dim=-1)

        if likelihood == "bernoulli":
            if not torch.is_tensor(x):
                raise TypeError(f"Bernoulli recon expects tensor target for modality {mod_name!r}; got {type(x)!r}")
            logits = dec_out["logits"] if isinstance(dec_out, dict) else dec_out
            nll = F.binary_cross_entropy_with_logits(logits, x, reduction="none")
            return nll.sum(dim=-1)

        if likelihood == "binomial":
            if not isinstance(dec_out, dict) or ("logits" not in dec_out):
                raise ValueError("Binomial recon expects dict decoder output with key 'logits'.")
            logits = dec_out["logits"]
            successes, total_count, valid = self._extract_count_target(
                x,
                mod_name=mod_name,
                ref_device=logits.device,
                ref_dtype=logits.dtype,
            )
            if successes.shape != logits.shape:
                raise ValueError(
                    f"Binomial recon shape mismatch for {mod_name!r}: target {tuple(successes.shape)} vs logits {tuple(logits.shape)}"
                )
            nll = self._binomial_nll(successes, total_count, logits)
            nll = torch.where(valid, nll, torch.zeros_like(nll))
            return nll.sum(dim=-1)

        if likelihood in ("beta_binomial", "betabinomial", "bb"):
            if not isinstance(dec_out, dict):
                raise ValueError("Beta-Binomial recon expects dict decoder output.")
            logits_ref = None
            for k in ("mu_logits", "alpha", "mu"):
                if k in dec_out and torch.is_tensor(dec_out[k]):
                    logits_ref = dec_out[k]
                    break
            if logits_ref is None:
                raise ValueError("Beta-Binomial decoder output missing expected parameter tensors.")

            successes, total_count, valid = self._extract_count_target(
                x,
                mod_name=mod_name,
                ref_device=logits_ref.device,
                ref_dtype=logits_ref.dtype,
            )

            if ("alpha" in dec_out) and ("beta" in dec_out):
                alpha = dec_out["alpha"]
                beta = dec_out["beta"]
            elif ("mu" in dec_out) and ("conc" in dec_out):
                mu = dec_out["mu"].clamp(self.EPS, 1.0 - self.EPS)
                conc = dec_out["conc"].clamp_min(self.EPS)
                alpha = (mu * conc).clamp_min(self.EPS)
                beta = ((1.0 - mu) * conc).clamp_min(self.EPS)
            elif ("mu_logits" in dec_out) and ("log_conc" in dec_out):
                mu = torch.sigmoid(dec_out["mu_logits"]).clamp(self.EPS, 1.0 - self.EPS)
                conc = torch.exp(dec_out["log_conc"]).clamp_min(self.EPS)
                alpha = (mu * conc).clamp_min(self.EPS)
                beta = ((1.0 - mu) * conc).clamp_min(self.EPS)
            else:
                raise ValueError("Beta-Binomial recon expects alpha/beta or (mu,conc) or (mu_logits,log_conc).")

            if successes.shape != alpha.shape:
                raise ValueError(
                    f"Beta-Binomial recon shape mismatch for {mod_name!r}: target {tuple(successes.shape)} vs params {tuple(alpha.shape)}"
                )

            nll = self._beta_binomial_nll(successes, total_count, alpha, beta, eps=self.EPS)
            nll = torch.where(valid, nll, torch.zeros_like(nll))
            return nll.sum(dim=-1)

        if likelihood == "poisson":
            if not torch.is_tensor(x):
                raise TypeError(f"Poisson recon expects tensor target for modality {mod_name!r}; got {type(x)!r}")
            log_rate = dec_out["log_rate"] if isinstance(dec_out, dict) and "log_rate" in dec_out else dec_out
            nll = F.poisson_nll_loss(log_rate, x, log_input=True, full=False, reduction="none")
            return nll.sum(dim=-1)

        if likelihood in ("nb", "negative_binomial"):
            if not torch.is_tensor(x):
                raise TypeError(f"NB recon expects tensor target for modality {mod_name!r}; got {type(x)!r}")
            if not isinstance(dec_out, dict) or ("mu" not in dec_out) or ("log_theta" not in dec_out):
                raise ValueError(f"NB recon expects dict with keys ('mu','log_theta'); got {type(dec_out)}")
            mu = dec_out["mu"]
            theta = torch.exp(dec_out["log_theta"])
            if theta.dim() == 1:
                theta = theta.unsqueeze(0).expand_as(mu)
            return self._nb_nll(x, mu, theta, eps=self.EPS).sum(dim=-1)

        if likelihood in ("zinb", "zero_inflated_negative_binomial"):
            if not torch.is_tensor(x):
                raise TypeError(f"ZINB recon expects tensor target for modality {mod_name!r}; got {type(x)!r}")
            if (
                not isinstance(dec_out, dict)
                or ("mu" not in dec_out)
                or ("log_theta" not in dec_out)
                or ("logit_pi" not in dec_out)
            ):
                raise ValueError("ZINB recon expects dict with keys ('mu','log_theta','logit_pi').")
            mu = dec_out["mu"]
            theta = torch.exp(dec_out["log_theta"])
            logit_pi = dec_out["logit_pi"]
            if theta.dim() == 1:
                theta = theta.unsqueeze(0).expand_as(mu)
            if logit_pi.dim() == 1:
                logit_pi = logit_pi.unsqueeze(0).expand_as(mu)
            return self._zinb_nll(x, mu, theta, logit_pi, eps=self.EPS).sum(dim=-1)

        if not torch.is_tensor(x):
            raise TypeError(f"Fallback recon expects tensor target for modality {mod_name!r}; got {type(x)!r}")
        pred = dec_out["mean"] if isinstance(dec_out, dict) and ("mean" in dec_out) else dec_out
        return ((x - pred) ** 2).sum(dim=-1)

    # ------------------------------ annealing ------------------------------

    def _anneal_weight(self, epoch: int, start: int, end: int, max_val: float) -> float:
        start = int(start)
        end = int(end)
        if end <= start:
            return float(max_val)
        if epoch <= start:
            return 0.0
        if epoch >= end:
            return float(max_val)
        frac = (epoch - start) / float(end - start)
        return float(max_val) * float(frac)

    # --------------------------- convenience API ---------------------------

    @property
    def device(self) -> torch.device:
        return next(self.parameters()).device

    @torch.no_grad()
    def encode_fused(
        self,
        x_dict: Dict[str, torch.Tensor],
        *,
        epoch: int = 0,
        y: Optional[YType] = None,
        use_mean: bool = True,
        inject_label_expert: bool = True,
        attn_bias_cfg: Optional[Mapping[str, Any]] = None,
        return_gates: bool = False,
        return_gate_logits: bool = False,
    ) -> Union[
        Tuple[torch.Tensor, torch.Tensor, torch.Tensor],
        Tuple[torch.Tensor, torch.Tensor, torch.Tensor, Optional[torch.Tensor], Optional[torch.Tensor]],
    ]:
        mu_dict, logvar_dict = self.encode_modalities(x_dict, attn_bias_cfg=attn_bias_cfg)
        if len(mu_dict) == 0:
            raise ValueError("At least one modality must be present in x_dict.")

        use_fused = (self.fused_encoder_type == "multimodal_transformer") and self._can_use_fused_encoder(x_dict)

        gates: Optional[torch.Tensor] = None
        gate_logits: Optional[torch.Tensor] = None

        if use_fused:
            mu_z, logvar_z = self._compute_fused_posterior(x_dict, attn_bias_cfg=attn_bias_cfg)
        else:
            if return_gates or return_gate_logits:
                if return_gate_logits:
                    mu_z, logvar_z, gates, gate_logits = self.mixture_of_experts(
                        mu_dict, logvar_dict,
                        return_weights=True,
                        return_logits=True,
                    )
                else:
                    mu_z, logvar_z, gates = self.mixture_of_experts(
                        mu_dict, logvar_dict,
                        return_weights=True,
                        return_logits=False,
                    )
                    gate_logits = None
            else:
                mu_z, logvar_z = self.mixture_of_experts(mu_dict, logvar_dict)

        y_legacy = self._extract_legacy_y(y)
        if (
            inject_label_expert
            and (self.label_encoder is not None)
            and (y_legacy is not None)
            and (epoch >= self.label_encoder_warmup)
        ):
            B = mu_z.shape[0]
            dev = mu_z.device
            mu_y, logvar_y = self._encode_labels_as_expert(y=y_legacy, B=B, device=dev)
            base_mu_dict = {"__base__": mu_z, "__label__": mu_y}
            base_lv_dict = {"__base__": logvar_z, "__label__": logvar_y}
            mu_z, logvar_z = self.mixture_of_experts(base_mu_dict, base_lv_dict)

        z = mu_z if use_mean else self._reparameterize(mu_z, logvar_z)

        if return_gates or return_gate_logits:
            return mu_z, logvar_z, z, gates, gate_logits

        return mu_z, logvar_z, z

    @torch.no_grad()
    def predict_heads(
        self,
        x_dict: Dict[str, torch.Tensor],
        *,
        epoch: int = 0,
        y: Optional[YType] = None,
        use_mean: bool = True,
        inject_label_expert: bool = True,
        return_probs: bool = True,
        attn_bias_cfg: Optional[Mapping[str, Any]] = None,
    ) -> Dict[str, torch.Tensor]:
        mu_z, logvar_z, z = self.encode_fused(
            x_dict,
            epoch=epoch,
            y=y,
            use_mean=use_mean,
            inject_label_expert=inject_label_expert,
            attn_bias_cfg=attn_bias_cfg,
        )
        out: Dict[str, torch.Tensor] = {}

        if self.label_decoder is not None:
            z_for_cls = mu_z if self.classify_from_mu else z
            dec_out = self.label_decoder(z_for_cls)
            logits = dec_out.get("logits", dec_out.get("logit", dec_out.get("scores", dec_out))) if isinstance(dec_out, dict) else dec_out
            out[self.label_head_name] = logits if not return_probs else F.softmax(logits, dim=-1)

        for name, head in self.class_heads.items():
            cfg_h = self.class_heads_cfg[name]
            z_in = mu_z if bool(cfg_h["from_mu"]) else z
            dec_out = head(z_in)
            logits = dec_out.get("logits", dec_out.get("logit", dec_out.get("scores", dec_out))) if isinstance(dec_out, dict) else dec_out

            head_type = str(cfg_h.get("type", "categorical")).lower().strip()
            if not return_probs:
                out[name] = logits
            else:
                if head_type == "binary":
                    out[name] = torch.sigmoid(logits)
                else:
                    out[name] = F.softmax(logits, dim=-1)

        return out

    def get_classification_meta(self) -> Dict[str, Any]:
        meta: Dict[str, Any] = {
            "label_head_name": self.label_head_name,
            "legacy": {
                "n_label_classes": int(self.n_label_classes),
                "label_ignore_index": int(self.label_ignore_index),
            },
            "multi": {
                "heads": {k: dict(v) for k, v in self.class_heads_cfg.items()},
            },
        }
        if self.label_names is not None:
            meta["legacy"]["label_names"] = list(self.label_names)
        if self.head_label_names:
            meta["multi"]["label_names"] = {k: list(v) for k, v in self.head_label_names.items()}
        return meta
```
