## Supplemental Code for "Unifying multimodal single-cell data with a mixture-of-experts *β*-variational autoencoder framework": objectives.py

```
# univi/objectives.py

from __future__ import annotations
from typing import Dict
import torch


def kl_diag_gaussians(
    mu_q: torch.Tensor,
    logvar_q: torch.Tensor,
    mu_p: torch.Tensor,
    logvar_p: torch.Tensor,
) -> torch.Tensor:
    """
    KL(q||p) for diagonal Gaussians, summed over latent dim, per sample.
    """
    var_q = torch.exp(logvar_q)
    var_p = torch.exp(logvar_p)
    kl = (
        logvar_p
        - logvar_q
        + (var_q + (mu_q - mu_p) ** 2) / var_p
        - 1.0
    )
    return 0.5 * kl.sum(dim=-1)


def symmetric_alignment_loss(
    mu_per_mod: Dict[str, torch.Tensor],
) -> torch.Tensor:
    """
    Simple symmetric cross-modal alignment: mean pairwise L2 distance
    between latent means across modalities.
    """
    names = list(mu_per_mod.keys())
    if len(names) < 2:
        return torch.zeros(mu_per_mod[names[0]].size(0), device=mu_per_mod[names[0]].device)

    losses = []
    for i in range(len(names)):
        for j in range(i + 1, len(names)):
            mu_i = mu_per_mod[names[i]]
            mu_j = mu_per_mod[names[j]]
            losses.append(((mu_i - mu_j) ** 2).sum(dim=-1))
    stacked = torch.stack(losses, dim=0)
    return stacked.mean(dim=0)
```
