## Supplemental Code for "Unifying multimodal single-cell data with a mixture-of-experts *β*-variational autoencoder framework": pipeline.py

```
# univi/pipeline.py
from __future__ import annotations

from dataclasses import asdict
from typing import Any, Dict, Optional, Tuple

import os
import numpy as np
import torch
import anndata as ad
from anndata import AnnData

from .config import UniVIConfig, ModalityConfig, TrainingConfig
from .models import UniVIMultiModalVAE
from .data import align_paired_obs_names, infer_input_dim, _get_matrix
from .utils.io import load_config, load_checkpoint
from .utils.seed import set_seed


def load_anndata_dict(cfg: Dict[str, Any], *, data_root: Optional[str] = None) -> Dict[str, AnnData]:
    adata_dict: Dict[str, AnnData] = {}
    for m in cfg["data"]["modalities"]:
        name = m["name"]
        path = m["h5ad_path"]
        if data_root is not None and not os.path.isabs(path):
            path = os.path.join(data_root, path)
        adata_dict[name] = ad.read_h5ad(path)
    return adata_dict


def build_univi_from_config(
    cfg: Dict[str, Any],
    adata_dict: Dict[str, AnnData],
) -> Tuple[UniVIMultiModalVAE, UniVIConfig, Dict[str, Optional[str]], Dict[str, str]]:
    """Build UniVI model + selectors from a loaded config and loaded AnnData dict."""
    mcfgs = cfg["data"]["modalities"]
    model_cfg = cfg.get("model", {})

    # selectors
    layer_by = {m["name"]: m.get("layer", None) for m in mcfgs}
    xkey_by = {m["name"]: m.get("X_key", "X") for m in mcfgs}

    modalities = []
    for m in mcfgs:
        name = m["name"]
        input_dim = infer_input_dim(adata_dict[name], layer=layer_by[name], X_key=xkey_by[name])

        hidden_default = model_cfg.get("hidden_dims_default", [256, 128])
        enc = m.get("encoder_hidden", m.get("hidden_dims", hidden_default))
        dec = m.get("decoder_hidden", m.get("decoder_hidden", list(enc)[::-1]))
        modalities.append(
            ModalityConfig(
                name=name,
                input_dim=int(input_dim),
                encoder_hidden=list(enc),
                decoder_hidden=list(dec),
                likelihood=m.get("likelihood", "gaussian"),
            )
        )

    univi_cfg = UniVIConfig(
        latent_dim=int(model_cfg.get("latent_dim", 32)),
        modalities=modalities,
        beta=float(model_cfg.get("beta", 1.0)),
        gamma=float(model_cfg.get("gamma", 1.0)),
        encoder_dropout=float(model_cfg.get("encoder_dropout", model_cfg.get("dropout", 0.0))),
        decoder_dropout=float(model_cfg.get("decoder_dropout", model_cfg.get("dropout", 0.0))),
        encoder_batchnorm=bool(model_cfg.get("encoder_batchnorm", model_cfg.get("batchnorm", True))),
        decoder_batchnorm=bool(model_cfg.get("decoder_batchnorm", False)),
        kl_anneal_start=int(model_cfg.get("kl_anneal_start", 0)),
        kl_anneal_end=int(model_cfg.get("kl_anneal_end", 0)),
        align_anneal_start=int(model_cfg.get("align_anneal_start", 0)),
        align_anneal_end=int(model_cfg.get("align_anneal_end", 0)),
    )

    model = UniVIMultiModalVAE(
        univi_cfg,
        loss_mode=model_cfg.get("loss_mode", "v2"),
        v1_recon=model_cfg.get("v1_recon", "cross"),
        v1_recon_mix=float(model_cfg.get("v1_recon_mix", 0.0)),
        normalize_v1_terms=bool(model_cfg.get("normalize_v1_terms", True)),
    )
    return model, univi_cfg, layer_by, xkey_by


@torch.no_grad()
def encode_latents_single_modality(
    model: UniVIMultiModalVAE,
    adata: AnnData,
    modality: str,
    *,
    layer: Optional[str] = None,
    X_key: str = "X",
    batch_size: int = 512,
    device: str = "cpu",
) -> np.ndarray:
    """Encode *one modality only* (for unimodal / independent dataset experiments)."""
    model.eval()
    model.to(device)

    X = _get_matrix(adata, layer=layer, X_key=X_key)
    # materialize sparse batches safely
    n = X.shape[0]
    zs = []
    for start in range(0, n, batch_size):
        end = min(n, start + batch_size)
        xb = X[start:end]
        if hasattr(xb, "A"):
            xb = xb.A
        xb = torch.as_tensor(np.asarray(xb), dtype=torch.float32, device=device)
        mu_dict, logvar_dict = model.encode_modalities({modality: xb})
        mu_z, logvar_z = model.mixture_of_experts(mu_dict, logvar_dict)
        zs.append(mu_z.detach().cpu().numpy())
    return np.vstack(zs)


@torch.no_grad()
def encode_latents_paired(
    model: UniVIMultiModalVAE,
    adata_dict: Dict[str, AnnData],
    *,
    layer_by: Optional[Dict[str, Optional[str]]] = None,
    xkey_by: Optional[Dict[str, str]] = None,
    batch_size: int = 512,
    device: str = "cpu",
    fused: bool = True,
) -> Dict[str, np.ndarray]:
    """Encode paired cells for each modality and optionally the fused MoE latent.

    Returns dict with keys for each modality (mu of that encoder) and optionally 'fused'.
    """
    model.eval()
    model.to(device)

    names = list(adata_dict.keys())
    n = adata_dict[names[0]].n_obs
    # require paired order
    for nm in names[1:]:
        if not np.array_equal(adata_dict[nm].obs_names.values, adata_dict[names[0]].obs_names.values):
            raise ValueError(f"obs_names mismatch between {names[0]} and {nm}")

    out = {nm: [] for nm in names}
    if fused:
        out["fused"] = []

    for start in range(0, n, batch_size):
        end = min(n, start + batch_size)
        x_dict = {}
        for nm, adata in adata_dict.items():
            layer = None if layer_by is None else layer_by.get(nm, None)
            xkey  = "X" if xkey_by is None else xkey_by.get(nm, "X")
            X = _get_matrix(adata, layer=layer, X_key=xkey)[start:end]
            if hasattr(X, "A"):
                X = X.A
            x_dict[nm] = torch.as_tensor(np.asarray(X), dtype=torch.float32, device=device)

        mu_dict, logvar_dict = model.encode_modalities(x_dict)
        # per-modality mus
        for nm in names:
            out[nm].append(mu_dict[nm].detach().cpu().numpy())
        if fused:
            mu_z, logvar_z = model.mixture_of_experts(mu_dict, logvar_dict)
            out["fused"].append(mu_z.detach().cpu().numpy())

    for k in list(out.keys()):
        out[k] = np.vstack(out[k])
    return out


def load_model_and_data(
    config_path: str,
    *,
    checkpoint_path: Optional[str] = None,
    data_root: Optional[str] = None,
    device: str = "cpu",
    align_obs: bool = True,
) -> Tuple[Dict[str, Any], Dict[str, AnnData], UniVIMultiModalVAE, Dict[str, Optional[str]], Dict[str, str]]:
    cfg = load_config(config_path)
    seed = int(cfg.get("training", {}).get("seed", 0))
    set_seed(seed, deterministic=False)

    adata_dict = load_anndata_dict(cfg, data_root=data_root)
    if align_obs:
        adata_dict = align_paired_obs_names(adata_dict)

    model, univi_cfg, layer_by, xkey_by = build_univi_from_config(cfg, adata_dict)
    if checkpoint_path is not None:
        ck = load_checkpoint(checkpoint_path)
        # common patterns
        state = ck.get("model_state", ck.get("state_dict", ck))
        model.load_state_dict(state, strict=False)

    model.to(device)
    return cfg, adata_dict, model, layer_by, xkey_by
```
