## Supplemental Code for "Unifying multimodal single-cell data with a mixture-of-experts *β*-variational autoencoder framework": plotting.py

```
# univi/plotting.py
from __future__ import annotations

from typing import Dict, Optional, Sequence, List, Union, Tuple, Any

import numpy as np
import matplotlib.pyplot as plt
import scanpy as sc
import scipy.sparse as sp
from anndata import AnnData

from .evaluation import to_dense


# =============================================================================
# Style / defaults
# =============================================================================
def set_style(
    font_scale: float = 1.1,
    dpi: int = 150,
    *,
    rc: Optional[Dict[str, Any]] = None,
) -> None:
    import matplotlib as mpl

    base = 10.0 * float(font_scale)
    defaults = {
        "figure.dpi": int(dpi),
        "savefig.dpi": 300,
        "font.size": base,
        "axes.titlesize": base * 1.15,
        "axes.labelsize": base * 1.05,
        "xtick.labelsize": base * 0.90,
        "ytick.labelsize": base * 0.90,
        "legend.fontsize": base * 0.90,
        "pdf.fonttype": 42,
        "ps.fonttype": 42,
    }
    if rc:
        defaults.update(rc)

    mpl.rcParams.update(defaults)
    sc.settings.set_figure_params(dpi=int(dpi), dpi_save=300, frameon=False)


# =============================================================================
# Display / lifecycle helpers
# =============================================================================
def _default_close(show: bool, savepath: Optional[str], return_fig: bool) -> bool:
    """
    Sensible default:
      - If the user wants a figure object back, do NOT close by default.
      - If show=True, do NOT close by default (so interactive use behaves normally).
      - If show=False AND not returning fig: close by default (prevents Jupyter double-render).
      - If saving without showing and not returning fig: close by default.
    """
    if return_fig:
        return False
    if show:
        return False
    # show=False and not returning fig => close to prevent stray display
    return True


def _finalize_figure(
    fig: plt.Figure,
    *,
    savepath: Optional[str],
    show: bool,
    close: bool,
    tight: bool = True,
    bbox_inches: str = "tight",
    pad_inches: float = 0.02,
) -> None:
    if tight:
        try:
            fig.tight_layout()
        except Exception:
            pass
    if savepath is not None:
        fig.savefig(savepath, dpi=300, bbox_inches=bbox_inches, pad_inches=float(pad_inches))
    if show:
        plt.show()
    if close:
        plt.close(fig)


# =============================================================================
# Scanpy legend / category helpers
# =============================================================================
def _is_categorical_obs(adata: AnnData, key: str) -> bool:
    if key not in adata.obs:
        return False
    s = adata.obs[key]
    return (str(s.dtype) == "category") or (s.dtype == object) or (str(s.dtype).startswith("string"))


def _ensure_scanpy_colors(adata: AnnData, key: str) -> None:
    if key not in adata.obs:
        return
    if not _is_categorical_obs(adata, key):
        return
    color_key = f"{key}_colors"
    if color_key in adata.uns and adata.uns[color_key] is not None:
        return
    try:
        sc.pl.umap(adata, color=key, show=False)
    except Exception:
        pass


def _get_categories_and_colors(
    adata: AnnData,
    key: str,
    *,
    subset_topk: Optional[int] = None,
) -> Tuple[List[str], List[Any]]:
    s = adata.obs[key].astype("category")
    cats_all = [str(c) for c in s.cat.categories]

    if subset_topk is not None and subset_topk > 0 and len(cats_all) > subset_topk:
        vc = s.value_counts()
        cats = [str(x) for x in vc.index[: int(subset_topk)]]
    else:
        cats = cats_all

    _ensure_scanpy_colors(adata, key)
    colors_all = adata.uns.get(f"{key}_colors", None)

    if colors_all is None:
        colors_all = [None] * len(cats_all)
    else:
        colors_all = list(colors_all)

    if len(colors_all) < len(cats_all):
        colors_all = (colors_all * (len(cats_all) // max(1, len(colors_all)) + 1))[: len(cats_all)]
    else:
        colors_all = colors_all[: len(cats_all)]

    if cats != cats_all:
        idx = {lab: i for i, lab in enumerate(cats_all)}
        colors = [colors_all[idx[lab]] for lab in cats]
    else:
        colors = colors_all[: len(cats)]

    return cats, colors


def _add_outside_legend(
    ax: plt.Axes,
    adata: AnnData,
    key: str,
    *,
    title: Optional[str] = None,
    ncols: int = 2,
    fontsize: Optional[float] = None,
    markerscale: float = 1.2,
    max_items: int = 40,
    subset_topk: Optional[int] = None,
    loc: str = "center left",
    bbox_to_anchor: Tuple[float, float] = (1.02, 0.5),
) -> None:
    if key not in adata.obs or not _is_categorical_obs(adata, key):
        return

    s = adata.obs[key].astype("category")
    n_cats = len(s.cat.categories)

    if n_cats > int(max_items) and (subset_topk is None or subset_topk <= 0):
        return

    cats_str, colors = _get_categories_and_colors(adata, key, subset_topk=subset_topk)
    if len(cats_str) > int(max_items):
        return

    handles = []
    for lab, col in zip(cats_str, colors):
        h = plt.Line2D(
            [0], [0],
            marker="o", linestyle="",
            markersize=6,
            markerfacecolor=col, markeredgecolor=col,
            label=lab,
        )
        handles.append(h)

    ax.legend(
        handles=handles,
        title=title or key,
        loc=loc,
        bbox_to_anchor=bbox_to_anchor,
        frameon=False,
        ncol=int(ncols),
        fontsize=fontsize,
        markerscale=float(markerscale),
        borderaxespad=0.0,
        handletextpad=0.35,
        columnspacing=0.9,
    )


# =============================================================================
# Embedding / UMAP helpers
# =============================================================================
def ensure_neighbors_umap(
    adata: AnnData,
    *,
    rep_key: str = "X_univi",
    n_neighbors: int = 30,
    random_state: int = 0,
    neighbors_key: Optional[str] = None,
    force: bool = False,
) -> None:
    if rep_key not in adata.obsm:
        raise KeyError(f"Missing obsm[{rep_key!r}]. Available: {list(adata.obsm.keys())}")

    if neighbors_key is None:
        needs_neighbors = force or ("neighbors" not in adata.uns)
        if needs_neighbors:
            sc.pp.neighbors(adata, use_rep=rep_key, n_neighbors=int(n_neighbors))
    else:
        needs_neighbors = force or (neighbors_key not in adata.uns)
        if needs_neighbors:
            sc.pp.neighbors(adata, use_rep=rep_key, n_neighbors=int(n_neighbors), key_added=str(neighbors_key))

    needs_umap = force or ("X_umap" not in adata.obsm)
    if needs_umap:
        if neighbors_key is None:
            sc.tl.umap(adata, random_state=int(random_state))
        else:
            sc.tl.umap(adata, random_state=int(random_state), neighbors_key=str(neighbors_key))


# =============================================================================
# UMAP plotting
# =============================================================================
def umap(
    adata: AnnData,
    *,
    obsm_key: str = "X_univi",
    color: Union[str, Sequence[str]] = (),
    layer: Optional[str] = None,
    title: Optional[Union[str, Sequence[str]]] = None,
    size: Optional[float] = None,
    n_neighbors: int = 30,
    random_state: int = 0,
    force_recompute: bool = False,
    ncols: int = 2,
    panel_size: Tuple[float, float] = (4.2, 3.8),
    wspace: float = 0.25,
    hspace: float = 0.25,
    legend: str = "outside",  # "outside", "right_margin", "on_data", "none"
    legend_ncols: int = 2,
    legend_fontsize: Optional[float] = None,
    legend_max_items: int = 40,
    legend_subset_topk: Optional[int] = None,
    legend_bbox_to_anchor: Tuple[float, float] = (1.02, 0.5),
    savepath: Optional[str] = None,
    show: bool = True,
    close: Optional[bool] = None,
    return_fig: bool = False,
    bbox_inches: str = "tight",
    pad_inches: float = 0.02,
    **scanpy_kwargs,
) -> Union[None, Tuple[plt.Figure, np.ndarray]]:
    ensure_neighbors_umap(
        adata,
        rep_key=obsm_key,
        n_neighbors=n_neighbors,
        random_state=random_state,
        force=force_recompute,
    )

    colors = [color] if isinstance(color, str) else list(color)
    n_panels = max(1, len(colors)) if len(colors) > 0 else 1

    if close is None:
        close = _default_close(show=show, savepath=savepath, return_fig=return_fig)

    ncols_eff = int(min(max(1, ncols), n_panels))
    nrows_eff = int(np.ceil(n_panels / ncols_eff))

    fig = plt.figure(figsize=(panel_size[0] * ncols_eff, panel_size[1] * nrows_eff))
    gs = fig.add_gridspec(nrows_eff, ncols_eff, wspace=wspace, hspace=hspace)

    axes = np.empty((nrows_eff, ncols_eff), dtype=object)
    for r in range(nrows_eff):
        for c in range(ncols_eff):
            axes[r, c] = fig.add_subplot(gs[r, c])

    if title is None:
        titles = [None] * n_panels
    elif isinstance(title, str):
        titles = [title] + [None] * (n_panels - 1)
    else:
        titles = list(title) + [None] * max(0, n_panels - len(title))

    layer_eff = None if (layer is None or layer == "X") else layer

    if len(colors) == 0:
        ax = axes[0, 0]
        sc.pl.umap(
            adata, color=None, ax=ax, show=False,
            title=titles[0] if titles[0] is not None else "",
            size=size, **scanpy_kwargs,
        )
        for j in range(1, nrows_eff * ncols_eff):
            r, c = divmod(j, ncols_eff)
            axes[r, c].axis("off")

        _finalize_figure(fig, savepath=savepath, show=show, close=bool(close),
                         tight=False, bbox_inches=bbox_inches, pad_inches=pad_inches)
        return (fig, axes) if return_fig else None

    for i, key in enumerate(colors):
        r, c = divmod(i, ncols_eff)
        ax = axes[r, c]
        is_cat = _is_categorical_obs(adata, str(key))

        if legend == "none":
            legend_loc = None
        elif legend == "on_data":
            legend_loc = "on data"
        elif legend == "right_margin":
            legend_loc = "right margin"
        elif legend == "outside":
            legend_loc = None
        else:
            legend_loc = "right margin" if is_cat else None

        sc.pl.umap(
            adata,
            color=str(key),
            layer=layer_eff,
            ax=ax,
            show=False,
            title=titles[i] if titles[i] is not None else str(key),
            size=size,
            legend_loc=legend_loc,
            **scanpy_kwargs,
        )

        if legend == "outside" and is_cat:
            _add_outside_legend(
                ax, adata, str(key),
                ncols=int(legend_ncols),
                fontsize=legend_fontsize,
                max_items=int(legend_max_items),
                subset_topk=legend_subset_topk,
                bbox_to_anchor=legend_bbox_to_anchor,
            )

    for j in range(len(colors), nrows_eff * ncols_eff):
        r, c = divmod(j, ncols_eff)
        axes[r, c].axis("off")

    _finalize_figure(fig, savepath=savepath, show=show, close=bool(close),
                     tight=False, bbox_inches=bbox_inches, pad_inches=pad_inches)
    return (fig, axes) if return_fig else None


def umap_by_modality(
    adata_dict: Dict[str, AnnData],
    *,
    obsm_key: str = "X_univi",
    color: Union[str, Sequence[str]] = ("cell_type", "dataset"),
    layer: Optional[str] = None,
    ncols: int = 2,
    panel_size: Tuple[float, float] = (4.2, 3.8),
    wspace: float = 0.25,
    hspace: float = 0.25,
    legend: str = "outside",
    legend_ncols: int = 2,
    legend_fontsize: Optional[float] = None,
    legend_max_items: int = 40,
    legend_subset_topk: Optional[int] = None,
    legend_bbox_to_anchor: Tuple[float, float] = (1.02, 0.5),
    n_neighbors: int = 30,
    random_state: int = 0,
    force_recompute: bool = False,
    savepath: Optional[str] = None,
    show: bool = True,
    close: Optional[bool] = None,
    return_fig: bool = False,
    bbox_inches: str = "tight",
    pad_inches: float = 0.02,
    **scanpy_kwargs,
) -> Union[None, Tuple[plt.Figure, np.ndarray]]:
    mods = list(adata_dict.keys())
    if len(mods) == 0:
        raise ValueError("adata_dict is empty.")

    Zs = []
    obs_frames = []
    for mod in mods:
        a = adata_dict[mod]
        if obsm_key not in a.obsm:
            raise KeyError(f"{mod!r} missing obsm[{obsm_key!r}]")

        Zs.append(np.asarray(a.obsm[obsm_key]))
        obs = a.obs.copy()
        obs["univi_modality"] = str(mod)
        obs_frames.append(obs)

    Z = np.vstack(Zs)
    if len(obs_frames) == 1:
        obs_all = obs_frames[0]
    else:
        import pandas as pd
        obs_all = pd.concat(obs_frames, axis=0)

    combined = AnnData(X=np.zeros((Z.shape[0], 0), dtype=np.float32), obs=obs_all)
    combined.obsm[obsm_key] = Z

    # NOTE: close/return_fig behavior is handled inside umap()
    return umap(
        combined,
        obsm_key=obsm_key,
        color=color,
        layer=layer,
        ncols=ncols,
        panel_size=panel_size,
        wspace=wspace,
        hspace=hspace,
        legend=legend,
        legend_ncols=legend_ncols,
        legend_fontsize=legend_fontsize,
        legend_max_items=legend_max_items,
        legend_subset_topk=legend_subset_topk,
        legend_bbox_to_anchor=legend_bbox_to_anchor,
        n_neighbors=n_neighbors,
        random_state=random_state,
        force_recompute=force_recompute,
        savepath=savepath,
        show=show,
        close=close,
        return_fig=return_fig,
        bbox_inches=bbox_inches,
        pad_inches=pad_inches,
        **scanpy_kwargs,
    )


# =============================================================================
# README: Confusion matrix plot
# =============================================================================
def plot_confusion_matrix(
    cm: np.ndarray,
    *,
    labels: Optional[Sequence[str]] = None,
    xlabels: Optional[Sequence[str]] = None,
    ylabels: Optional[Sequence[str]] = None,
    title: Optional[str] = None,
    normalize: Optional[str] = None,  # None | "true" | "pred" | "all"
    figsize: Tuple[float, float] = (6.0, 5.4),
    rotate_xticks: int = 90,
    show: bool = True,
    savepath: Optional[str] = None,
    close: Optional[bool] = None,
    return_fig: bool = False,
) -> Optional[plt.Figure]:
    """
    Simple matplotlib confusion matrix plot.

    normalize:
      - None: raw counts
      - "true": rows sum to 1
      - "pred": cols sum to 1
      - "all": total sum to 1
    """
    M = np.asarray(cm, dtype=float)
    if M.ndim != 2 or M.shape[0] != M.shape[1]:
        raise ValueError(f"cm must be square, got shape={M.shape}")

    if normalize is not None:
        norm = str(normalize).lower().strip()
        if norm == "true":
            denom = M.sum(axis=1, keepdims=True) + 1e-12
            M = M / denom
        elif norm == "pred":
            denom = M.sum(axis=0, keepdims=True) + 1e-12
            M = M / denom
        elif norm == "all":
            M = M / (M.sum() + 1e-12)
        else:
            raise ValueError("normalize must be one of: None, 'true', 'pred', 'all'.")

    n = M.shape[0]
    if labels is None:
        labels0 = [str(i) for i in range(n)]
    else:
        labels0 = [str(x) for x in labels]
        if len(labels0) != n:
            raise ValueError(f"labels length must match cm size ({n}).")

    if ylabels is None:
        ylabels = labels0
    else:
        ylabels = [str(x) for x in ylabels]
        if len(ylabels) != n:
            raise ValueError(f"ylabels length must match cm size ({n}).")

    if xlabels is None:
        xlabels = labels0
    else:
        xlabels = [str(x) for x in xlabels]
        if len(xlabels) != n:
            raise ValueError(f"xlabels length must match cm size ({n}).")

    if close is None:
        close = _default_close(show=show, savepath=savepath, return_fig=return_fig)

    fig = plt.figure(figsize=figsize)
    ax = plt.gca()
    im = ax.imshow(M, aspect="auto")
    plt.colorbar(im, ax=ax, fraction=0.046, pad=0.04)

    ax.set_xticks(np.arange(n))
    ax.set_yticks(np.arange(n))
    ax.set_xticklabels(xlabels, rotation=rotate_xticks, ha="right")
    ax.set_yticklabels(ylabels)

    ax.set_xlabel("Predicted")
    ax.set_ylabel("True")
    if title is not None:
        ax.set_title(title)

    _finalize_figure(fig, savepath=savepath, show=show, close=bool(close),
                     tight=True, bbox_inches="tight", pad_inches=0.02)

    return fig if return_fig else None


# =============================================================================
# README: MoE gate helpers
# =============================================================================
def write_gates_to_obs(
    adata: AnnData,
    gates: np.ndarray,
    *,
    modality_names: Sequence[str],
    gate_prefix: str = "gate",
    gate_logits: Optional[np.ndarray] = None,
    logits_prefix: Optional[str] = None,
) -> None:
    """
    Write gate weights (and optionally gate logits) into adata.obs columns.

    Columns:
      gate_prefix_{mod}
      (optional) {logits_prefix or gate_prefix+"_logit"}_{mod}
    """
    G = np.asarray(gates, dtype=np.float32)
    mods = [str(m) for m in modality_names]
    if G.ndim != 2:
        raise ValueError(f"gates must be 2D, got shape={G.shape}")
    if G.shape[0] != adata.n_obs:
        raise ValueError(f"gates n_cells ({G.shape[0]}) != adata.n_obs ({adata.n_obs})")
    if G.shape[1] != len(mods):
        raise ValueError(f"gates n_mods ({G.shape[1]}) != len(modality_names) ({len(mods)})")

    for j, m in enumerate(mods):
        adata.obs[f"{gate_prefix}_{m}"] = G[:, j]

    if gate_logits is not None:
        L = np.asarray(gate_logits, dtype=np.float32)
        if L.shape != G.shape:
            raise ValueError(f"gate_logits shape {L.shape} must match gates shape {G.shape}")
        lp = logits_prefix if logits_prefix is not None else f"{gate_prefix}_logit"
        for j, m in enumerate(mods):
            adata.obs[f"{lp}_{m}"] = L[:, j]


def plot_moe_gate_summary(
    adata: AnnData,
    *,
    gate_prefix: str = "gate",
    modality_names: Optional[Sequence[str]] = None,
    groupby: str = "celltype.l2",
    agg: str = "mean",  # "mean" | "median"
    figsize: Tuple[float, float] = (7.2, 4.2),
    title: Optional[str] = None,
    show: bool = True,
    savepath: Optional[str] = None,
    close: Optional[bool] = None,
    return_fig: bool = False,
) -> Optional[plt.Figure]:
    """
    Heatmap summary of MoE gate usage by group.

    Expects columns in adata.obs:
      {gate_prefix}_{modality}
    """
    if groupby not in adata.obs:
        raise KeyError(f"groupby={groupby!r} not in adata.obs")

    if modality_names is None:
        modality_names = []
        for c in adata.obs.columns:
            if c.startswith(f"{gate_prefix}_") and not c.startswith(f"{gate_prefix}_logit"):
                modality_names.append(c[len(gate_prefix) + 1 :])
        modality_names = sorted(set(modality_names))
    mods = [str(m) for m in modality_names]
    if len(mods) == 0:
        raise ValueError("Could not infer modality_names; pass modality_names explicitly.")

    cols = [f"{gate_prefix}_{m}" for m in mods]
    for c in cols:
        if c not in adata.obs:
            raise KeyError(f"Missing gate column {c!r} in adata.obs")

    import pandas as pd
    df = adata.obs[[groupby] + cols].copy()
    df[groupby] = df[groupby].astype("category")

    if str(agg).lower().strip() == "median":
        mat = df.groupby(groupby)[cols].median()
    else:
        mat = df.groupby(groupby)[cols].mean()

    sizes = df[groupby].value_counts()
    mat = mat.loc[sizes.index]

    M = mat.to_numpy(dtype=float)
    ylabels = [str(x) for x in mat.index]
    xlabels = [str(x) for x in mods]

    if close is None:
        close = _default_close(show=show, savepath=savepath, return_fig=return_fig)

    fig = plt.figure(figsize=figsize)
    ax = plt.gca()
    im = ax.imshow(M, aspect="auto")
    plt.colorbar(im, ax=ax, fraction=0.046, pad=0.04)

    ax.set_yticks(np.arange(len(ylabels)))
    ax.set_yticklabels(ylabels)
    ax.set_xticks(np.arange(len(xlabels)))
    ax.set_xticklabels(xlabels, rotation=45, ha="right")

    ax.set_xlabel("Modality")
    ax.set_ylabel(groupby)
    if title is None:
        title = f"MoE gates by {groupby} ({agg})"
    ax.set_title(title)

    _finalize_figure(fig, savepath=savepath, show=show, close=bool(close),
                     tight=True, bbox_inches="tight", pad_inches=0.02)

    return fig if return_fig else None


# =============================================================================
# Convenience helpers for logits/probs + error plots
# =============================================================================
def layer_sigmoid_inplace(adata: AnnData, src_layer: str, dst_layer: str) -> None:
    """
    Convert logits layer -> probs layer using sigmoid.
    Works for dense or sparse; for sparse, applies sigmoid to stored data only.
    """
    X = adata.layers[src_layer]
    if sp.issparse(X):
        X2 = X.copy()
        X2.data = 1.0 / (1.0 + np.exp(-X2.data))
        adata.layers[dst_layer] = X2
    else:
        adata.layers[dst_layer] = 1.0 / (1.0 + np.exp(-np.asarray(X)))


def compare_raw_vs_pred_umap_features(
    adata: AnnData,
    *,
    obsm_key: str = "X_univi",
    features: Sequence[str],
    pred_layer: str,
    raw_layer: Optional[str] = None,
    pred_is_logits: bool = False,
    n_neighbors: int = 30,
    random_state: int = 0,
    force_recompute: bool = False,
    panel_size: Tuple[float, float] = (4.0, 3.6),
    wspace: float = 0.15,
    hspace: float = 0.15,
    savepath: Optional[str] = None,
    show: bool = True,
    close: Optional[bool] = None,
    return_fig: bool = False,
    bbox_inches: str = "tight",
    pad_inches: float = 0.02,
    **scanpy_kwargs,
) -> Union[None, Tuple[plt.Figure, np.ndarray]]:
    """
    2-row grid:
      row 1: raw
      row 2: predicted

    pred_is_logits=True will plot sigmoid(pred_layer) by creating a temporary layer.
    """
    feats = list(features)
    if len(feats) == 0:
        raise ValueError("features must be non-empty.")

    ensure_neighbors_umap(
        adata,
        rep_key=obsm_key,
        n_neighbors=n_neighbors,
        random_state=random_state,
        force=force_recompute,
    )

    raw_layer_eff = None if (raw_layer is None or raw_layer == "X") else raw_layer
    if pred_layer not in adata.layers:
        raise KeyError(f"pred_layer {pred_layer!r} not found in adata.layers.")

    plot_pred_layer = pred_layer
    tmp_layer = None
    if pred_is_logits:
        tmp_layer = f"__tmp_sigmoid__{pred_layer}"
        layer_sigmoid_inplace(adata, pred_layer, tmp_layer)
        plot_pred_layer = tmp_layer

    if close is None:
        close = _default_close(show=show, savepath=savepath, return_fig=return_fig)

    ncol = len(feats)
    fig, axes = plt.subplots(
        nrows=2,
        ncols=ncol,
        figsize=(panel_size[0] * ncol, panel_size[1] * 2),
        squeeze=False,
        gridspec_kw={"wspace": wspace, "hspace": hspace},
    )

    for j, f in enumerate(feats):
        sc.pl.umap(
            adata, color=f, layer=raw_layer_eff, ax=axes[0, j], show=False,
            title=f"{f} (raw)", **scanpy_kwargs
        )
        sc.pl.umap(
            adata, color=f, layer=plot_pred_layer, ax=axes[1, j], show=False,
            title=f"{f} (pred)", **scanpy_kwargs
        )

    _finalize_figure(fig, savepath=savepath, show=show, close=bool(close),
                     tight=False, bbox_inches=bbox_inches, pad_inches=pad_inches)

    if tmp_layer is not None and tmp_layer in adata.layers:
        try:
            del adata.layers[tmp_layer]
        except Exception:
            pass

    return (fig, axes) if return_fig else None


def plot_feature_scatter_observed_vs_pred(
    adata: AnnData,
    feature: str,
    *,
    layer_obs: Optional[str] = None,
    layer_pred: str,
    pred_is_logits: bool = False,
    max_points: int = 20000,
    random_state: int = 0,
    s: float = 4.0,
    alpha: float = 0.3,
    title: Optional[str] = None,
    show: bool = True,
    savepath: Optional[str] = None,
    close: Optional[bool] = None,
    return_fig: bool = False,
) -> Optional[plt.Figure]:
    """
    Scatter: predicted vs observed for one feature (AnnData-based).

    IMPORTANT NOTE (Jupyter):
      If you call with show=False and you don't want any display, also ensure
      the function closes the figure (default behavior here does that unless return_fig=True).
    """
    j = adata.var_names.get_loc(feature)

    Xo = adata.X if layer_obs is None else adata.layers[layer_obs]
    Xp = adata.layers[layer_pred]

    yo = np.asarray(Xo[:, j].toarray()).ravel() if sp.issparse(Xo) else np.asarray(Xo[:, j]).ravel()
    yp = np.asarray(Xp[:, j].toarray()).ravel() if sp.issparse(Xp) else np.asarray(Xp[:, j]).ravel()

    if pred_is_logits:
        yp = 1.0 / (1.0 + np.exp(-yp))

    n = yo.shape[0]
    if n > int(max_points):
        rng = np.random.default_rng(int(random_state))
        idx = rng.choice(n, size=int(max_points), replace=False)
        yo, yp = yo[idx], yp[idx]

    if close is None:
        close = _default_close(show=show, savepath=savepath, return_fig=return_fig)

    fig = plt.figure(figsize=(4.8, 4.2))
    plt.scatter(yp, yo, s=float(s), alpha=float(alpha))
    plt.xlabel("pred")
    plt.ylabel("obs")
    plt.title(title or feature)

    _finalize_figure(fig, savepath=savepath, show=show, close=bool(close),
                     tight=True, bbox_inches="tight", pad_inches=0.02)

    return fig if return_fig else None


# =============================================================================
# README-compatible names (FULL implementations)
# =============================================================================
def compare_raw_vs_denoised_umap_features(
    adata: AnnData,
    *,
    obsm_key: str = "X_univi",
    features: Sequence[str],
    raw_layer: Optional[str] = None,
    denoised_layer: str = "denoised_fused",
    denoised_is_logits: bool = False,
    n_neighbors: int = 30,
    random_state: int = 0,
    force_recompute: bool = False,
    panel_size: Tuple[float, float] = (4.0, 3.6),
    wspace: float = 0.15,
    hspace: float = 0.15,
    savepath: Optional[str] = None,
    show: bool = True,
    close: Optional[bool] = None,
    return_fig: bool = False,
    bbox_inches: str = "tight",
    pad_inches: float = 0.02,
    **scanpy_kwargs,
) -> Union[None, Tuple[plt.Figure, np.ndarray]]:
    """
    README name -> plots raw vs denoised in a 2-row UMAP feature grid.
    """
    return compare_raw_vs_pred_umap_features(
        adata,
        obsm_key=obsm_key,
        features=features,
        pred_layer=denoised_layer,
        raw_layer=raw_layer,
        pred_is_logits=denoised_is_logits,
        n_neighbors=n_neighbors,
        random_state=random_state,
        force_recompute=force_recompute,
        panel_size=panel_size,
        wspace=wspace,
        hspace=hspace,
        savepath=savepath,
        show=show,
        close=close,
        return_fig=return_fig,
        bbox_inches=bbox_inches,
        pad_inches=pad_inches,
        **scanpy_kwargs,
    )


def plot_featurewise_reconstruction_scatter(
    rep: Dict[str, Any],
    *,
    features: Sequence[str],
    max_points: int = 20000,
    random_state: int = 0,
    s: float = 4.0,
    alpha: float = 0.3,
    title: Optional[str] = None,
    savepath: Optional[str] = None,
    show: bool = True,
    close: Optional[bool] = None,
    return_fig: bool = False,
) -> Union[None, plt.Figure, Dict[str, plt.Figure]]:
    """
    Scatter plots of TRUE vs PRED for selected features,
    using dict returned by univi.evaluation.evaluate_cross_reconstruction(...).

    If return_fig=False, returns None (prevents notebook auto-render).
    If len(features)==1 and return_fig=True, returns a Figure.
    Otherwise returns dict[str, Figure].
    """
    feats = list(features)
    if len(feats) == 0:
        raise ValueError("features must be non-empty.")
    if "X_true" not in rep or "X_pred" not in rep:
        raise KeyError("rep must contain keys 'X_true' and 'X_pred'.")

    X_true = np.asarray(rep["X_true"])
    X_pred = np.asarray(rep["X_pred"])
    if X_true.shape != X_pred.shape:
        raise ValueError(f"Shape mismatch: X_true {X_true.shape} vs X_pred {X_pred.shape}")

    feat_names = rep.get("feature_names", None)
    if feat_names is None:
        feat_names = [str(i) for i in range(X_true.shape[1])]
    else:
        feat_names = [str(x) for x in list(feat_names)]

    name_to_j = {n: j for j, n in enumerate(feat_names)}

    figs: Dict[str, plt.Figure] = {}
    for f in feats:
        if f not in name_to_j:
            raise KeyError(f"Feature {f!r} not found in rep['feature_names'] (n={len(feat_names)}).")

        j = name_to_j[f]
        yt = X_true[:, j].astype(float, copy=False)
        yp = X_pred[:, j].astype(float, copy=False)

        n = yt.shape[0]
        if n > int(max_points):
            rng = np.random.default_rng(int(random_state))
            idx = rng.choice(n, size=int(max_points), replace=False)
            yt = yt[idx]
            yp = yp[idx]

        # decide lifecycle per-figure
        if close is None:
            close_i = _default_close(show=show, savepath=savepath, return_fig=return_fig)
        else:
            close_i = bool(close)

        fig = plt.figure(figsize=(4.8, 4.2))
        plt.scatter(yp, yt, s=float(s), alpha=float(alpha))
        plt.xlabel("pred")
        plt.ylabel("true")
        plt.title(title or f)
        plt.tight_layout()

        if savepath is not None:
            if len(feats) == 1:
                out = savepath
            else:
                root, ext = (savepath.rsplit(".", 1) + ["png"])[:2]
                out = f"{root}__{f}.{ext}"
            fig.savefig(out, dpi=300, bbox_inches="tight", pad_inches=0.02)

        if show:
            plt.show()
        if close_i:
            plt.close(fig)

        figs[f] = fig

    if not return_fig:
        return None
    return figs[feats[0]] if len(feats) == 1 else figs


def plot_reconstruction_error_summary(
    rep: Dict[str, Any],
    *,
    top_k: int = 25,
    metric: str = "mse",  # "mse" | "pearson" | "auc"
    sort: str = "worst",  # "worst" | "best"
    title: Optional[str] = None,
    show: bool = True,
    savepath: Optional[str] = None,
    close: Optional[bool] = None,
    return_fig: bool = False,
) -> Optional[plt.Figure]:
    """
    README-friendly summary plot for reconstruction reports.

    If return_fig=False, returns None (prevents notebook auto-render).
    """
    if "per_feature" not in rep or "feature_names" not in rep:
        raise KeyError("rep must contain keys 'per_feature' and 'feature_names'.")

    metric = str(metric).lower().strip()
    sort = str(sort).lower().strip()
    if sort not in {"worst", "best"}:
        raise ValueError("sort must be 'worst' or 'best'.")

    feat_names = [str(x) for x in rep["feature_names"]]

    pf = rep["per_feature"]
    if metric == "mse":
        if "mse" not in pf:
            raise KeyError("rep['per_feature'] must contain 'mse' for metric='mse'.")
        vals = np.asarray(pf["mse"], dtype=float)
        order = np.argsort(vals)[::-1] if sort == "worst" else np.argsort(vals)
        xlabel = "MSE (higher = worse)"
    elif metric == "pearson":
        if "pearson" not in pf:
            raise KeyError("rep['per_feature'] must contain 'pearson' for metric='pearson'.")
        vals = np.asarray(pf["pearson"], dtype=float)
        order = np.argsort(vals) if sort == "worst" else np.argsort(vals)[::-1]
        xlabel = "Pearson r (higher = better)"
    elif metric == "auc":
        if "auc" not in pf:
            raise KeyError("rep['per_feature'] must contain 'auc' for metric='auc'.")
        vals = np.asarray(pf["auc"], dtype=float)
        order = np.argsort(vals) if sort == "worst" else np.argsort(vals)[::-1]
        xlabel = "AUC (higher = better)"
    else:
        raise ValueError("metric must be one of: 'mse', 'pearson', 'auc'.")

    nan_mask = np.isnan(vals)
    if nan_mask.any():
        bad = np.where(nan_mask)[0]
        order = np.concatenate([order[~nan_mask[order]], bad])

    top_k = int(max(1, min(int(top_k), len(feat_names))))
    idx = order[:top_k]

    names_top = [feat_names[i] for i in idx]
    vals_top = vals[idx]

    if close is None:
        close = _default_close(show=show, savepath=savepath, return_fig=return_fig)

    fig = plt.figure(figsize=(7.6, max(3.2, 0.22 * top_k + 1.8)))
    y = np.arange(top_k)[::-1]
    plt.barh(y, vals_top[::-1])
    plt.yticks(y, names_top[::-1], fontsize=9)

    if title is None:
        base = "Reconstruction summary"
        if "summary" in rep and isinstance(rep["summary"], dict):
            s = rep["summary"]
            if metric == "mse" and "mse_mean" in s:
                base += f" (mse_mean={s['mse_mean']:.4g}, r_mean={s.get('pearson_mean', float('nan')):.3g})"
            elif metric == "pearson" and "pearson_mean" in s:
                base += f" (r_mean={s['pearson_mean']:.3g})"
            elif metric == "auc" and "auc_mean" in s:
                base += f" (auc_mean={s['auc_mean']:.3g})"
        title = base

    plt.title(title)
    plt.xlabel(xlabel)

    _finalize_figure(fig, savepath=savepath, show=show, close=bool(close),
                     tight=True, bbox_inches="tight", pad_inches=0.02)

    return fig if return_fig else None


# =============================================================================
# Backwards-compatible aliases (older names)
# =============================================================================
def umap_single_adata(
    adata_obj: AnnData,
    *,
    obsm_key: str = "X_univi",
    color: Optional[Sequence[str]] = None,
    layer: Optional[str] = None,
    savepath: Optional[str] = None,
    n_neighbors: int = 30,
    random_state: int = 0,
    title: Optional[str] = None,
    size: Optional[float] = None,
    show: Optional[bool] = None,
    close: Optional[bool] = None,
    legend: str = "right_margin",
    legend_ncols: int = 2,
    legend_fontsize: Optional[float] = None,
    legend_max_items: int = 40,
    legend_subset_topk: Optional[int] = None,
    legend_bbox_to_anchor: Tuple[float, float] = (1.02, 0.5),
    ncols: int = 2,
    panel_size: Tuple[float, float] = (4.2, 3.8),
    wspace: float = 0.25,
    hspace: float = 0.25,
    force_recompute: bool = False,
    return_fig: bool = False,
    **scanpy_kwargs,
) -> Union[None, Tuple[plt.Figure, np.ndarray]]:
    # Preserve historical behavior defaults: show=False, close=True
    if show is None:
        show = False
    if close is None:
        close = True

    color_list = list(color) if color is not None else []
    return umap(
        adata_obj,
        obsm_key=obsm_key,
        color=color_list,
        layer=layer,
        title=title,
        size=size,
        n_neighbors=n_neighbors,
        random_state=random_state,
        force_recompute=force_recompute,
        ncols=ncols,
        panel_size=panel_size,
        wspace=wspace,
        hspace=hspace,
        legend=legend,
        legend_ncols=legend_ncols,
        legend_fontsize=legend_fontsize,
        legend_max_items=legend_max_items,
        legend_subset_topk=legend_subset_topk,
        legend_bbox_to_anchor=legend_bbox_to_anchor,
        savepath=savepath,
        show=bool(show),
        close=bool(close),
        return_fig=return_fig,
        **scanpy_kwargs,
    )


def umap_by_modality_old(
    adata_dict: Dict[str, AnnData],
    *,
    obsm_key: str = "X_univi",
    color: Union[str, Sequence[str]] = "celltype",
    layer: Optional[str] = None,
    savepath: Optional[str] = None,
    n_neighbors: int = 30,
    random_state: int = 0,
    title: Optional[str] = None,
    size: Optional[float] = None,
    show: Optional[bool] = None,
    close: Optional[bool] = None,
    **kwargs,
) -> None:
    # Preserve historical behavior defaults: show=False, close=True
    if show is None:
        show = False
    if close is None:
        close = True

    if isinstance(color, str):
        color_list = [color, "univi_modality"]
    else:
        color_list = list(color) + ["univi_modality"]

    umap_by_modality(
        adata_dict,
        obsm_key=obsm_key,
        color=color_list,
        layer=layer,
        n_neighbors=n_neighbors,
        random_state=random_state,
        force_recompute=kwargs.pop("force_recompute", False),
        savepath=savepath,
        show=bool(show),
        close=bool(close),
        **{k: v for k, v in kwargs.items() if k in {
            "ncols", "panel_size", "wspace", "hspace",
            "legend", "legend_ncols", "legend_fontsize", "legend_max_items",
            "legend_subset_topk", "legend_bbox_to_anchor",
            "return_fig", "bbox_inches", "pad_inches",
        }},
    )
```
