## Supplemental Code for "Unifying multimodal single-cell data with a mixture-of-experts *β*-variational autoencoder framework": trainer.py

```
# univi/trainer.py
from __future__ import annotations

from dataclasses import asdict
from typing import Any, Dict, List, Optional, Tuple, Union, Mapping

import contextlib

import anndata as ad
import numpy as np
import scipy.sparse as sp
import torch
from torch import nn, optim
from torch.utils.data import DataLoader
from tqdm.auto import tqdm

from .config import TrainingConfig
from .utils.io import restore_checkpoint, save_checkpoint
from .utils.logging import get_logger

YType = Union[torch.Tensor, Dict[str, torch.Tensor]]

# Batch can be:
#   x_dict
#   (x_dict, y)
#   (x_dict, recon_targets)
#   (x_dict, y, recon_targets)
#
# recon_targets is typically:
#   Dict[modality -> {"successes": Tensor(B,D), "total_count": Tensor(B,D)}]
# but trainer also supports generic nested target mappings for forward compatibility.
ReconTargetsType = Dict[str, Dict[str, torch.Tensor]]

BatchType = Union[
    Dict[str, torch.Tensor],
    Tuple[Dict[str, torch.Tensor], YType],
    Tuple[Dict[str, torch.Tensor], ReconTargetsType],
    Tuple[Dict[str, torch.Tensor], YType, ReconTargetsType],
]


class UniVITrainer:
    """
    Lightweight training loop for UniVI models.

    Supports:
      - mixed precision (AMP) via use_amp + amp_dtype
      - optional coordinate metadata and attention-bias configuration for tokenized ATAC
        (passed through to model forward when supported)
      - optional recon_targets (successes + total_count) for binomial / beta-binomial losses
      - checkpoint save/load via utils.io helpers

    Behavior notes:
      - If val_loader is provided, we track best validation epoch + weights
        after an optional warmup window (TrainingConfig.best_epoch_warmup).
      - Early stopping (patience) is optional and only controls stopping.
      - If best weights were tracked, we restore them at the end of fit().
    """

    def __init__(
        self,
        model: nn.Module,
        train_loader: DataLoader,
        val_loader: Optional[DataLoader] = None,
        train_cfg: Optional[TrainingConfig] = None,
        device: Optional[str] = None,
        use_amp: bool = False,
        amp_dtype: str = "fp16",  # "fp16" or "bf16"
        *,
        feature_coords: Optional[Dict[str, Dict[str, Any]]] = None,
        attn_bias_cfg: Optional[Dict[str, Dict[str, Any]]] = None,
    ):
        self.model = model
        self.train_loader = train_loader
        self.val_loader = val_loader

        self.cfg = train_cfg or TrainingConfig()
        self.device = device or self.cfg.device

        self.use_amp = bool(use_amp)
        self.amp_dtype = str(amp_dtype).lower().strip()
        if self.amp_dtype not in ("fp16", "bf16"):
            raise ValueError(f"amp_dtype must be 'fp16' or 'bf16', got {amp_dtype!r}")

        # Optional genomic context / transformer bias configuration (safe defaults).
        # These are *data-derived* and intentionally not serialized by default.
        self.feature_coords: Dict[str, Dict[str, Any]] = feature_coords or {}
        self.attn_bias_cfg: Dict[str, Dict[str, Any]] = attn_bias_cfg or {}

        self.model.to(self.device)

        self.optimizer = optim.Adam(
            self.model.parameters(),
            lr=float(self.cfg.lr),
            weight_decay=float(self.cfg.weight_decay),
        )

        self._scaler: Optional[torch.cuda.amp.GradScaler] = None
        if self.use_amp and torch.cuda.is_available() and str(self.device).startswith("cuda"):
            # GradScaler is only used for fp16. bf16 autocast generally does not need scaling.
            self._scaler = torch.cuda.amp.GradScaler(enabled=(self.amp_dtype == "fp16"))

        self.logger = get_logger("UniVITrainer")

        # Best tracking (independent of early stopping)
        self.best_val_loss = float("inf")
        self.best_state_dict: Optional[Dict[str, Any]] = None
        self.best_epoch: Optional[int] = None

        # Early stopping counters
        self.epochs_no_improve = 0

        self.history: Dict[str, List[float]] = {
            "train_loss": [],
            "val_loss": [],
            "beta": [],
            "gamma": [],
        }

        self._log_config()

    # ------------------------------ training ------------------------------

    def fit(self) -> Dict[str, List[float]]:
        epoch_iter = tqdm(range(1, int(self.cfg.n_epochs) + 1), desc="Training UniVI", leave=True)

        for epoch in epoch_iter:
            tr_loss, tr_beta, tr_gamma = self._run_one_epoch(epoch, train=True)
            self.history["train_loss"].append(tr_loss)
            self.history["beta"].append(tr_beta)
            self.history["gamma"].append(tr_gamma)

            if self.val_loader is not None:
                va_loss, _, _ = self._run_one_epoch(epoch, train=False)
                self.history["val_loss"].append(va_loss)

                epoch_iter.set_postfix(
                    train_loss="%.4f" % tr_loss,
                    val_loss="%.4f" % va_loss,
                    beta="%.3f" % tr_beta,
                    gamma="%.3f" % tr_gamma,
                )

                improved, counted = self._update_best(va_loss, epoch)

                # Only update patience/stop logic if early stopping is enabled.
                # Warmup epochs do NOT increment patience (counted=False).
                if self.cfg.early_stopping and counted:
                    self._update_patience(improved)

                if self._should_stop():
                    self.logger.info(
                        "Early stopping at epoch %d (best val loss=%.4f, best epoch=%s)"
                        % (epoch, self.best_val_loss, str(self.best_epoch))
                    )
                    break
            else:
                epoch_iter.set_postfix(
                    train_loss="%.4f" % tr_loss,
                    beta="%.3f" % tr_beta,
                    gamma="%.3f" % tr_gamma,
                )

        # Restore best model if we have it (tracked after warmup), otherwise leave as-is.
        if self.val_loader is not None:
            if self.best_state_dict is not None:
                self.model.load_state_dict(self.best_state_dict)
                if self.best_epoch is not None:
                    self.logger.info(
                        "Restored best model from epoch %d (val loss=%.4f)"
                        % (self.best_epoch, self.best_val_loss)
                    )
            else:
                self.logger.info("No best checkpoint was tracked (best_epoch_warmup may exceed training).")

        return self.history

    # ------------------------------ model state handling ------------------------------

    def state_dict(self) -> Dict[str, Any]:
        # Keep this small. Large dataset-derived items (coords) are not stored here.
        return {
            "history": self.history,
            "best_val_loss": float(self.best_val_loss),
            "best_epoch": self.best_epoch,
            "epochs_no_improve": int(self.epochs_no_improve),
            "use_amp": bool(self.use_amp),
            "amp_dtype": str(self.amp_dtype),
        }

    def save(self, path: str, *, extra: Optional[Dict[str, Any]] = None, save_best: bool = False) -> None:
        model_state = (
            self.best_state_dict
            if (save_best and self.best_state_dict is not None)
            else self.model.state_dict()
        )

        scaler_state = None
        if self._scaler is not None and self._scaler.is_enabled():
            try:
                scaler_state = self._scaler.state_dict()
            except Exception:
                scaler_state = None

        save_checkpoint(
            path,
            model_state=model_state,
            optimizer_state=self.optimizer.state_dict(),
            extra=extra,
            model=self.model,
            trainer_state=self.state_dict(),
            scaler_state=scaler_state,
        )

    def load(
        self,
        path: str,
        *,
        map_location: Union[str, torch.device, None] = "cpu",
        strict: bool = True,
        restore_label_names: bool = True,
        enforce_label_compat: bool = True,
    ) -> Dict[str, Any]:
        payload = restore_checkpoint(
            path,
            model=self.model,
            optimizer=self.optimizer,
            scaler=self._scaler,
            map_location=map_location,
            strict=strict,
            restore_label_names=restore_label_names,
            enforce_label_compat=enforce_label_compat,
        )

        ts = payload.get("trainer_state", None)
        if isinstance(ts, dict):
            self.history = ts.get("history", self.history)
            self.best_val_loss = float(ts.get("best_val_loss", self.best_val_loss))
            self.best_epoch = ts.get("best_epoch", self.best_epoch)
            self.epochs_no_improve = int(ts.get("epochs_no_improve", self.epochs_no_improve))

        self.model.to(self.device)
        return payload

    # ------------------------------ batch handling ------------------------------

    @staticmethod
    def _looks_like_recon_targets(obj: Any) -> bool:
        """
        Heuristic used only to disambiguate (x_dict, y) vs (x_dict, recon_targets)
        when batch is a 2-tuple.

        Accepts the common shape:
          {mod: {"successes": ..., "total_count": ...}}
        """
        if not isinstance(obj, Mapping) or len(obj) == 0:
            return False
        for v in obj.values():
            if not isinstance(v, Mapping):
                return False
            if ("successes" not in v) or ("total_count" not in v):
                return False
        return True

    def _split_batch(
        self, batch: BatchType
    ) -> Tuple[Dict[str, torch.Tensor], Optional[YType], Optional[ReconTargetsType]]:
        """
        Accepts any of:
          - x_dict
          - (x_dict, y)
          - (x_dict, recon_targets)
          - (x_dict, y, recon_targets)

        where:
          y can be:
            * LongTensor (B,)                    [back-compat]
            * dict[str -> LongTensor(B,)]        [multi-head]
          recon_targets is:
            * dict[modality -> dict[target_name -> tensor]]
              e.g. {"atac": {"successes": ..., "total_count": ...}}

        Moves tensors to device.
        - x tensors: preserves dtype unless non-tensor (casts to float32)
        - y tensors: coerced to long (classification labels)
        - recon target tensors: preserves dtype if already tensor; otherwise float32
        """
        x_dict = None
        y: Optional[YType] = None
        recon_targets: Optional[ReconTargetsType] = None

        # ---------- parse batch shape ----------
        if isinstance(batch, (tuple, list)):
            if len(batch) == 2:
                a0, a1 = batch
                x_dict = a0

                # Prefer explicit helper for common recon-target shape.
                if self._looks_like_recon_targets(a1):
                    recon_targets = a1  # type: ignore[assignment]
                # Fallback heuristic: any nested mapping is likely recon_targets.
                elif isinstance(a1, Mapping):
                    is_nested_mapping = any(isinstance(v, Mapping) for v in a1.values()) if len(a1) > 0 else False
                    if is_nested_mapping:
                        recon_targets = a1  # type: ignore[assignment]
                    else:
                        y = a1  # type: ignore[assignment]
                else:
                    y = a1  # type: ignore[assignment]

            elif len(batch) == 3:
                a0, a1, a2 = batch
                x_dict = a0
                y = a1  # type: ignore[assignment]
                recon_targets = a2  # type: ignore[assignment]

            else:
                raise TypeError(
                    f"Expected batch to be dict, (dict,y), (dict,recon_targets), or (dict,y,recon_targets). "
                    f"Got tuple/list length={len(batch)}"
                )
        else:
            x_dict = batch

        if not isinstance(x_dict, dict):
            raise TypeError(f"Expected batch to be dict or (dict, ...). Got {type(x_dict)!r}")

        # ---------- x_dict ----------
        x_out: Dict[str, torch.Tensor] = {}
        for k, v in x_dict.items():
            if v is None:
                x_out[k] = None  # type: ignore[assignment]
                continue
            if torch.is_tensor(v):
                x_out[k] = v.to(self.device, non_blocking=True)
            else:
                x_out[k] = torch.as_tensor(v, dtype=torch.float32, device=self.device)

        # ---------- y ----------
        y_out: Optional[YType] = None
        if y is not None:
            if isinstance(y, Mapping):
                yd: Dict[str, torch.Tensor] = {}
                for hk, hv in y.items():
                    if hv is None:
                        yd[str(hk)] = None  # type: ignore[assignment]
                        continue
                    if not torch.is_tensor(hv):
                        hv = torch.as_tensor(hv)
                    yd[str(hk)] = hv.long().to(self.device, non_blocking=True)
                y_out = yd
            else:
                if not torch.is_tensor(y):
                    y = torch.as_tensor(y)
                y_out = y.long().to(self.device, non_blocking=True)

        # ---------- recon_targets ----------
        rt_out: Optional[ReconTargetsType] = None
        if recon_targets is not None:
            if not isinstance(recon_targets, Mapping):
                raise TypeError(f"recon_targets must be a mapping, got {type(recon_targets)!r}")

            rt_cast: ReconTargetsType = {}
            for mod, target_dict in recon_targets.items():
                if target_dict is None:
                    continue
                if not isinstance(target_dict, Mapping):
                    raise TypeError(
                        f"recon_targets[{mod!r}] must be a mapping of target_name->tensor, "
                        f"got {type(target_dict)!r}"
                    )

                td_cast: Dict[str, torch.Tensor] = {}
                for tk, tv in target_dict.items():
                    if tv is None:
                        continue
                    if torch.is_tensor(tv):
                        td_cast[str(tk)] = tv.to(self.device, non_blocking=True)
                    else:
                        # keep float32 by default for count-like tensors too (works with torch distributions)
                        td_cast[str(tk)] = torch.as_tensor(tv, dtype=torch.float32, device=self.device)

                if len(td_cast) > 0:
                    rt_cast[str(mod)] = td_cast

            rt_out = rt_cast if len(rt_cast) > 0 else None

        return x_out, y_out, rt_out

    # ------------------------------ forward wrappers ------------------------------

    def _forward_model(
        self,
        x_dict: Dict[str, torch.Tensor],
        epoch: int,
        y: Optional[YType],
        recon_targets: Optional[ReconTargetsType] = None,
    ):
        """
        Best-effort forward dispatch for multiple model signatures.

        Preferred signature (newest):
          model(x_dict, epoch=..., y=..., feature_coords=..., attn_bias_cfg=..., recon_targets=...)

        Then gracefully degrades to older signatures.
        """
        fc = self.feature_coords if self.feature_coords else None
        ab = self.attn_bias_cfg if self.attn_bias_cfg else None

        # Most specific / newest
        try:
            return self.model(
                x_dict,
                epoch=epoch,
                y=y,
                recon_targets=recon_targets,
                feature_coords=fc,
                attn_bias_cfg=ab,
            )
        except TypeError:
            pass

        # Without feature coords / attn bias
        try:
            return self.model(
                x_dict,
                epoch=epoch,
                y=y,
                recon_targets=recon_targets,
            )
        except TypeError:
            pass

        # Without y (but with recon targets)
        try:
            return self.model(
                x_dict,
                epoch=epoch,
                recon_targets=recon_targets,
            )
        except TypeError:
            pass

        # Older signatures (no recon_targets support)
        try:
            return self.model(x_dict, epoch=epoch, y=y, feature_coords=fc, attn_bias_cfg=ab)
        except TypeError:
            pass

        try:
            return self.model(x_dict, epoch=epoch, y=y)
        except TypeError:
            pass

        # Epoch-only
        try:
            return self.model(x_dict, epoch=epoch, recon_targets=recon_targets)
        except TypeError:
            pass

        try:
            return self.model(x_dict, epoch=epoch)
        except TypeError:
            pass

        # y-only
        if y is not None:
            try:
                return self.model(x_dict, y=y, recon_targets=recon_targets)
            except TypeError:
                pass

            try:
                return self.model(x_dict, y=y)
            except TypeError:
                pass

        # recon_targets-only
        if recon_targets is not None:
            try:
                return self.model(x_dict, recon_targets=recon_targets)
            except TypeError:
                pass

        return self.model(x_dict)

    @staticmethod
    def _as_float(v: Any) -> float:
        if v is None:
            return 0.0
        if isinstance(v, (float, int)):
            return float(v)
        if torch.is_tensor(v):
            return float(v.detach().cpu().item())
        try:
            return float(v)
        except Exception:
            return 0.0

    def _amp_context(self):
        if not (self.use_amp and torch.cuda.is_available() and str(self.device).startswith("cuda")):
            return contextlib.nullcontext()
        dtype = torch.float16 if self.amp_dtype == "fp16" else torch.bfloat16
        return torch.autocast(device_type="cuda", dtype=dtype, enabled=True)

    # ------------------------------ epoch loop ------------------------------

    def _run_one_epoch(self, epoch: int, train: bool = True) -> Tuple[float, float, float]:
        if train:
            self.model.train()
            loader = self.train_loader
        else:
            self.model.eval()
            if self.val_loader is None:
                raise ValueError("val_loader is None but train=False")
            loader = self.val_loader

        total_loss = 0.0
        total_beta = 0.0
        total_gamma = 0.0
        n_batches = 0

        for batch in loader:
            x_dict, y, recon_targets = self._split_batch(batch)

            if train:
                self.optimizer.zero_grad(set_to_none=True)

            with torch.set_grad_enabled(train):
                with self._amp_context():
                    out = self._forward_model(
                        x_dict,
                        epoch=epoch,
                        y=y,
                        recon_targets=recon_targets,
                    )

                    # Back-compat: prefer loss_fixed for eval if present, else loss.
                    loss = out["loss"] if train else out.get("loss_fixed", out["loss"])

                if train:
                    if self._scaler is not None and self._scaler.is_enabled():
                        self._scaler.scale(loss).backward()
                        if self.cfg.grad_clip is not None and float(self.cfg.grad_clip) > 0:
                            self._scaler.unscale_(self.optimizer)
                            nn.utils.clip_grad_norm_(self.model.parameters(), float(self.cfg.grad_clip))
                        self._scaler.step(self.optimizer)
                        self._scaler.update()
                    else:
                        loss.backward()
                        if self.cfg.grad_clip is not None and float(self.cfg.grad_clip) > 0:
                            nn.utils.clip_grad_norm_(self.model.parameters(), float(self.cfg.grad_clip))
                        self.optimizer.step()

            total_loss += float(loss.detach().cpu().item())
            total_beta += self._as_float(out.get("beta_used", out.get("beta", 0.0)))
            total_gamma += self._as_float(out.get("gamma_used", out.get("gamma", 0.0)))
            n_batches += 1

        avg_loss = total_loss / max(1, n_batches)
        avg_beta = total_beta / max(1, n_batches)
        avg_gamma = total_gamma / max(1, n_batches)

        # Robust logging even if log_every <= 0
        log_every = int(getattr(self.cfg, "log_every", 10) or 0)
        if epoch == 1 or (log_every > 0 and epoch % log_every == 0):
            self.logger.info(
                "[Epoch %03d] %s loss=%.4f (beta=%.3f, gamma=%.3f)"
                % (epoch, "Train" if train else "Val", avg_loss, avg_beta, avg_gamma)
            )

        return avg_loss, avg_beta, avg_gamma

    # ------------------------------ best tracking + early stopping ------------------------------

    def _best_warmup(self) -> int:
        v = int(getattr(self.cfg, "best_epoch_warmup", 0) or 0)
        return max(0, v)

    def _update_best(self, val_loss: float, epoch: int) -> Tuple[bool, bool]:
        """
        Update best validation metrics + weights with warmup support.

        Warmup behavior:
          - if epoch < best_epoch_warmup:
              * do NOT update best (improved=False)
              * do NOT count toward patience (counted=False)

        Returns:
          improved: bool  (whether this epoch became a new best)
          counted:  bool  (whether this epoch should affect patience)
        """
        val_loss_f = float(val_loss)
        warmup = self._best_warmup()

        if int(epoch) < int(warmup):
            if epoch == warmup - 1:
                self.logger.info(
                    "[Epoch %03d] Best tracking warmup ends next epoch (best_epoch_warmup=%d)."
                    % (epoch, warmup)
                )
            return False, False

        counted = True

        if self.best_state_dict is None or self.best_epoch is None:
            self.best_val_loss = val_loss_f
            self.best_state_dict = {k: v.detach().cpu().clone() for k, v in self.model.state_dict().items()}
            self.best_epoch = int(epoch)
            self.logger.info("[Epoch %03d] New best val loss: %.4f" % (epoch, val_loss_f))
            return True, counted

        improved = (self.best_val_loss - val_loss_f) > float(self.cfg.min_delta)
        if improved:
            self.best_val_loss = val_loss_f
            self.best_state_dict = {k: v.detach().cpu().clone() for k, v in self.model.state_dict().items()}
            self.best_epoch = int(epoch)
            self.logger.info("[Epoch %03d] New best val loss: %.4f" % (epoch, val_loss_f))
            return True, counted

        return False, counted

    def _update_patience(self, improved: bool) -> None:
        if improved:
            self.epochs_no_improve = 0
        else:
            self.epochs_no_improve += 1

    def _should_stop(self) -> bool:
        if not self.cfg.early_stopping:
            return False
        return int(self.epochs_no_improve) >= int(self.cfg.patience)

    # ------------------------------ encoding utility ------------------------------

    def encode_modality(
        self,
        adata: ad.AnnData,
        modality: str,
        layer: Optional[str] = None,
        X_key: str = "X",
        obs_key: Optional[str] = None,
        batch_size: int = 512,
        *,
        use_moe: bool = True,
    ) -> np.ndarray:
        """
        Encode a single modality AnnData into latent means.

        By default, this returns the fused mean via MoE/PoE if available (use_moe=True),
        which is identical to the modality posterior when only one modality is provided.

        If you want strictly per-modality posterior mean, set use_moe=False.
        """
        names = getattr(self.model, "modality_names", None)
        if names is not None and modality not in names:
            raise ValueError("Unknown modality %r. Available: %s" % (modality, names))

        self.model.eval()

        if obs_key is not None:
            if obs_key not in adata.obs:
                raise KeyError(f"obs_key={obs_key!r} not found in adata.obs.")
            col = adata.obs[obs_key]
            if hasattr(col, "cat"):
                vals = col.cat.codes.to_numpy()
            else:
                vals = np.asarray(col.values)
            X = vals.astype(np.float32).reshape(-1, 1)
        else:
            if X_key != "X":
                if X_key not in adata.obsm:
                    raise KeyError(f"X_key={X_key!r} not in adata.obsm.")
                X = adata.obsm[X_key]
            else:
                X = adata.layers[layer] if layer is not None else adata.X

            if sp.issparse(X):
                X = X.toarray()
            X = np.asarray(X, dtype=np.float32)

        zs = []
        dev = torch.device(self.device)

        with torch.no_grad():
            for start in range(0, X.shape[0], int(batch_size)):
                end = min(start + int(batch_size), X.shape[0])
                xb = torch.as_tensor(X[start:end], dtype=torch.float32, device=dev)

                mu_dict, logvar_dict = self.model.encode_modalities({modality: xb})

                if use_moe:
                    if hasattr(self.model, "mixture_of_experts"):
                        mu_z, _ = self.model.mixture_of_experts(mu_dict, logvar_dict)
                    elif hasattr(self.model, "fuse_posteriors"):
                        mu_z, _ = self.model.fuse_posteriors(mu_dict, logvar_dict)
                    else:
                        mu_z = mu_dict[modality]
                else:
                    mu_z = mu_dict[modality]

                zs.append(mu_z.detach().cpu().numpy())

        return np.vstack(zs)

    # ------------------------------ logging ------------------------------

    def _log_config(self) -> None:
        cfg_dict = asdict(self.cfg)
        self.logger.info("TrainingConfig:")
        for k, v in cfg_dict.items():
            self.logger.info("  %s: %r" % (k, v))
```
