## Supplemental Code for "Unifying multimodal single-cell data with a mixture-of-experts *β*-variational autoencoder framework": io.py

```
# univi/utils/io.py
from __future__ import annotations

from typing import Any, Dict, Optional, Sequence, Union, Mapping, Literal
import os
import json

import numpy as np
import torch
import scipy.sparse as sp
import anndata as ad


SplitKey = Literal["train", "val", "test"]
SplitMap = Dict[SplitKey, Any]


# =============================================================================
# Checkpointing
# =============================================================================

def save_checkpoint(
    path: str,
    model_state: Optional[Dict[str, Any]] = None,
    optimizer_state: Optional[Dict[str, Any]] = None,
    extra: Optional[Dict[str, Any]] = None,
    *,
    model: Optional[torch.nn.Module] = None,
    optimizer: Optional[torch.optim.Optimizer] = None,
    trainer_state: Optional[Dict[str, Any]] = None,
    scaler_state: Optional[Dict[str, Any]] = None,
    config: Optional[Dict[str, Any]] = None,
    strict_label_compat: bool = True,
) -> None:
    if model_state is None and model is not None:
        model_state = model.state_dict()
    if optimizer_state is None and optimizer is not None:
        optimizer_state = optimizer.state_dict()

    if model_state is None:
        raise ValueError("save_checkpoint requires either model_state=... or model=...")

    payload: Dict[str, Any] = {
        "format_version": 3,
        "model_state": model_state,
    }
    if optimizer_state is not None:
        payload["optimizer_state"] = optimizer_state
    if trainer_state is not None:
        payload["trainer_state"] = dict(trainer_state)
    if scaler_state is not None:
        payload["scaler_state"] = dict(scaler_state)
    if config is not None:
        payload["config"] = dict(config)
    if extra is not None:
        payload["extra"] = dict(extra)

    # --- classification metadata (legacy + multi-head) ---
    if model is not None:
        meta: Dict[str, Any] = {}

        n_label_classes = getattr(model, "n_label_classes", None)
        label_names = getattr(model, "label_names", None)
        label_head_name = getattr(model, "label_head_name", None)

        if n_label_classes is not None:
            meta.setdefault("legacy", {})["n_label_classes"] = int(n_label_classes)
        if label_names is not None:
            meta.setdefault("legacy", {})["label_names"] = list(label_names)
        if label_head_name is not None:
            meta["label_head_name"] = str(label_head_name)

        class_heads_cfg = getattr(model, "class_heads_cfg", None)
        head_label_names = getattr(model, "head_label_names", None)

        if isinstance(class_heads_cfg, dict) and len(class_heads_cfg) > 0:
            meta.setdefault("multi", {})["heads"] = {k: dict(v) for k, v in class_heads_cfg.items()}
        if isinstance(head_label_names, dict) and len(head_label_names) > 0:
            meta.setdefault("multi", {})["label_names"] = {k: list(v) for k, v in head_label_names.items()}

        if hasattr(model, "get_classification_meta"):
            try:
                meta = dict(model.get_classification_meta())
            except Exception:
                pass

        if meta:
            payload["label_meta"] = meta
            payload["strict_label_compat"] = bool(strict_label_compat)

    os.makedirs(os.path.dirname(path) or ".", exist_ok=True)
    torch.save(payload, path)


def load_checkpoint(path: str, *, map_location: Union[str, torch.device, None] = "cpu") -> Dict[str, Any]:
    return torch.load(path, map_location=map_location)


def restore_checkpoint(
    payload_or_path: Union[str, Dict[str, Any]],
    *,
    model: torch.nn.Module,
    optimizer: Optional[torch.optim.Optimizer] = None,
    scaler: Optional[torch.cuda.amp.GradScaler] = None,
    map_location: Union[str, torch.device, None] = "cpu",
    strict: bool = True,
    restore_label_names: bool = True,
    enforce_label_compat: bool = True,
) -> Dict[str, Any]:
    payload = load_checkpoint(payload_or_path, map_location=map_location) if isinstance(payload_or_path, str) else payload_or_path

    if enforce_label_compat:
        meta = payload.get("label_meta", {}) or {}

        legacy = meta.get("legacy", {}) if isinstance(meta, dict) else {}
        ckpt_n = legacy.get("n_label_classes", None)
        model_n = getattr(model, "n_label_classes", None)
        if ckpt_n is not None and model_n is not None and int(ckpt_n) != int(model_n):
            raise ValueError(
                f"Checkpoint n_label_classes={ckpt_n} does not match model n_label_classes={model_n}. "
                "Rebuild the model with the same n_label_classes."
            )

        multi = meta.get("multi", {}) if isinstance(meta, dict) else {}
        ckpt_heads = multi.get("heads", None)
        model_heads = getattr(model, "class_heads_cfg", None)
        if isinstance(ckpt_heads, dict) and isinstance(model_heads, dict):
            for hk, hcfg in ckpt_heads.items():
                if hk not in model_heads:
                    raise ValueError(
                        f"Checkpoint contains head {hk!r} but model does not. "
                        f"Model heads: {list(model_heads.keys())}"
                    )
                ckpt_c = int(hcfg.get("n_classes", -1))
                model_c = int(model_heads[hk].get("n_classes", -1))
                if ckpt_c != model_c:
                    raise ValueError(f"Head {hk!r} n_classes mismatch: checkpoint={ckpt_c}, model={model_c}.")

    model.load_state_dict(payload["model_state"], strict=bool(strict))

    if optimizer is not None and "optimizer_state" in payload:
        optimizer.load_state_dict(payload["optimizer_state"])

    if scaler is not None and payload.get("scaler_state") is not None:
        try:
            scaler.load_state_dict(payload["scaler_state"])
        except Exception:
            pass

    if restore_label_names:
        meta = payload.get("label_meta", {}) or {}

        legacy = meta.get("legacy", {}) if isinstance(meta, dict) else {}
        label_names = legacy.get("label_names", None)
        if label_names is not None and hasattr(model, "set_label_names"):
            try:
                model.set_label_names(list(label_names))
            except Exception:
                pass

        multi = meta.get("multi", {}) if isinstance(meta, dict) else {}
        head_names = multi.get("label_names", None)
        if isinstance(head_names, dict) and hasattr(model, "set_head_label_names"):
            for hk, names in head_names.items():
                try:
                    model.set_head_label_names(str(hk), list(names))
                except Exception:
                    pass

    return payload


# =============================================================================
# JSON config helpers
# =============================================================================

def save_config_json(config: Dict[str, Any], path: str) -> None:
    os.makedirs(os.path.dirname(path) or ".", exist_ok=True)
    with open(path, "w") as f:
        json.dump(config, f, indent=2, sort_keys=True)


def load_config(config_path: str) -> Dict[str, Any]:
    with open(config_path) as f:
        return json.load(f)


# =============================================================================
# AnnData split helpers (SAFE + FLEXIBLE)
# =============================================================================

def _is_sequence_of_str(x: Any) -> bool:
    return isinstance(x, (list, tuple, np.ndarray)) and len(x) > 0 and isinstance(x[0], (str, np.str_))


def _is_sequence_of_int(x: Any) -> bool:
    return isinstance(x, (list, tuple, np.ndarray)) and len(x) > 0 and isinstance(x[0], (int, np.integer))


def _normalize_split_selector(
    adata: ad.AnnData,
    selector: Any,
    *,
    name: str,
) -> Union[np.ndarray, Sequence[str], Sequence[int]]:
    """
    Convert a selector into one of:
      - boolean mask (np.ndarray[bool] length n_obs)
      - list of obs_names (Sequence[str])
      - list of integer indices (Sequence[int])

    Rejects AnnData objects or other iterables that could trigger accidental iteration.
    """
    if selector is None:
        return np.zeros(adata.n_obs, dtype=bool)

    # HARD FAIL: AnnData passed where indices were expected
    if isinstance(selector, ad.AnnData):
        raise TypeError(
            f"{name}: expected indices/obs_names/mask, but got AnnData. "
            "Pass `splits={'train': adata_train, ...}` instead."
        )

    # bool mask
    if isinstance(selector, np.ndarray) and selector.dtype == bool:
        if selector.shape[0] != adata.n_obs:
            raise ValueError(f"{name}: boolean mask length {selector.shape[0]} != n_obs {adata.n_obs}.")
        return selector

    # pandas Series of bool
    try:
        import pandas as pd  # optional
        if isinstance(selector, pd.Series) and selector.dtype == bool:
            v = selector.to_numpy()
            if v.shape[0] != adata.n_obs:
                raise ValueError(f"{name}: boolean mask length {v.shape[0]} != n_obs {adata.n_obs}.")
            return v
    except Exception:
        pass

    # list/tuple/np array of strings: obs_names
    if _is_sequence_of_str(selector):
        return [str(s) for s in list(selector)]

    # list/tuple/np array of ints: indices
    if _is_sequence_of_int(selector):
        idx = [int(i) for i in list(selector)]
        if len(idx) > 0:
            mx = max(idx)
            mn = min(idx)
            if mn < 0 or mx >= adata.n_obs:
                raise IndexError(f"{name}: index out of bounds (min={mn}, max={mx}, n_obs={adata.n_obs}).")
        return idx

    # empty list/tuple
    if isinstance(selector, (list, tuple)) and len(selector) == 0:
        return np.zeros(adata.n_obs, dtype=bool)

    raise TypeError(
        f"{name}: unsupported selector type {type(selector)}. "
        "Use obs_names (list[str]), indices (list[int]), or boolean mask (np.ndarray[bool]). "
        "If you already have split AnnData objects, pass `splits=...`."
    )


def _subset_adata(
    adata: ad.AnnData,
    selector: Union[np.ndarray, Sequence[str], Sequence[int]],
    *,
    copy: bool,
) -> ad.AnnData:
    if isinstance(selector, np.ndarray) and selector.dtype == bool:
        out = adata[selector]
    elif _is_sequence_of_str(selector):
        out = adata[list(selector)]
    else:
        out = adata[list(selector), :]
    return out.copy() if copy else out


def _write_h5ad_safe(adata_obj: ad.AnnData, path: str, *, write_backed: bool = False) -> None:
    os.makedirs(os.path.dirname(path) or ".", exist_ok=True)
    # write_backed kept for API compatibility; anndata's backed writing patterns vary.
    adata_obj.write_h5ad(path)


def save_anndata_splits(
    adata: Optional[ad.AnnData] = None,
    outdir: str = ".",
    *,
    prefix: str = "dataset",
    # Mode A: split from adata.obs[split_key]
    split_key: Optional[str] = "split",
    train_label: str = "train",
    val_label: str = "val",
    test_label: str = "test",
    # Mode B: split from selectors (indices/obs_names/mask)
    split_map: Optional[Dict[str, Any]] = None,
    # Mode C: already split objects (no resplitting)
    splits: Optional[Dict[str, ad.AnnData]] = None,
    # Behavior
    copy: bool = False,
    write_backed: bool = False,
    save_h5ad: bool = True,
    save_split_map: bool = True,
    split_map_name: Optional[str] = None,
    require_disjoint: bool = True,
) -> Dict[str, ad.AnnData]:
    """
    Save train/val/test splits to {outdir}/{prefix}_{train|val|test}.h5ad and optionally a split map JSON.

    You can provide EXACTLY ONE of:
      1) splits={...}                      (already split AnnData objects)
      2) adata + split_map={...}           (selectors: obs_names, indices, or boolean masks)
      3) adata + split_key in adata.obs    (labels in .obs)

    Safety: passing AnnData objects inside split_map is rejected with a clear error.
    """
    os.makedirs(outdir, exist_ok=True)

    if splits is not None:
        if adata is not None and split_map is not None:
            raise ValueError("Provide only one of: splits=, (adata+split_map), or (adata+split_key).")
        missing = [k for k in ("train", "val", "test") if k not in splits]
        if missing:
            raise ValueError(f"splits is missing keys: {missing}. Expected train/val/test.")
        train = splits["train"]
        val = splits["val"]
        test = splits["test"]
        if require_disjoint:
            s1 = set(train.obs_names)
            s2 = set(val.obs_names)
            s3 = set(test.obs_names)
            if (s1 & s2) or (s1 & s3) or (s2 & s3):
                raise ValueError("splits are not disjoint by obs_names (overlap found).")
    else:
        if adata is None:
            raise ValueError("If splits is not provided, you must provide adata=...")

        if split_map is not None:
            sel_train = _normalize_split_selector(adata, split_map.get("train", None), name="split_map['train']")
            sel_val   = _normalize_split_selector(adata, split_map.get("val", None),   name="split_map['val']")
            sel_test  = _normalize_split_selector(adata, split_map.get("test", None),  name="split_map['test']")

            if require_disjoint:
                def to_set(sel):
                    if isinstance(sel, np.ndarray) and sel.dtype == bool:
                        return set(np.where(sel)[0].tolist())
                    if _is_sequence_of_str(sel):
                        return set(map(str, sel))
                    return set(map(int, sel))
                a = to_set(sel_train); b = to_set(sel_val); c = to_set(sel_test)
                if (a & b) or (a & c) or (b & c):
                    raise ValueError("split_map splits overlap (require_disjoint=True).")

            train = _subset_adata(adata, sel_train, copy=copy)
            val   = _subset_adata(adata, sel_val,   copy=copy)
            test  = _subset_adata(adata, sel_test,  copy=copy)

        else:
            if split_key is None or split_key not in adata.obs:
                raise ValueError(
                    f"Expected split labels in adata.obs[{split_key!r}], or provide split_map=..., or splits=...."
                )
            s = adata.obs[split_key].astype(str)
            train = adata[s == train_label].copy() if copy else adata[s == train_label]
            val   = adata[s == val_label].copy()   if copy else adata[s == val_label]
            test  = adata[s == test_label].copy()  if copy else adata[s == test_label]

            if require_disjoint:
                s1 = set(train.obs_names); s2 = set(val.obs_names); s3 = set(test.obs_names)
                if (s1 & s2) or (s1 & s3) or (s2 & s3):
                    raise ValueError("split_key-derived splits are not disjoint by obs_names.")

    paths = {
        "train": os.path.join(outdir, f"{prefix}_train.h5ad"),
        "val":   os.path.join(outdir, f"{prefix}_val.h5ad"),
        "test":  os.path.join(outdir, f"{prefix}_test.h5ad"),
    }

    if save_h5ad:
        _write_h5ad_safe(train, paths["train"], write_backed=write_backed)
        _write_h5ad_safe(val,   paths["val"],   write_backed=write_backed)
        _write_h5ad_safe(test,  paths["test"],  write_backed=write_backed)

    if save_split_map:
        sm = {
            "train": train.obs_names.tolist(),
            "val":   val.obs_names.tolist(),
            "test":  test.obs_names.tolist(),
            "prefix": prefix,
        }
        if splits is None and split_map is None and adata is not None:
            sm.update({
                "split_key": split_key,
                "train_label": train_label,
                "val_label": val_label,
                "test_label": test_label,
            })

        fn = split_map_name or f"{prefix}_split_map.json"
        with open(os.path.join(outdir, fn), "w") as f:
            json.dump(sm, f, indent=2)

    return {"train": train, "val": val, "test": test}


# =============================================================================
# Loading helpers (ADDED)
# =============================================================================

def load_split_map(outdir: str, prefix: str = "dataset", split_map_name: Optional[str] = None) -> Dict[str, Any]:
    fn = split_map_name or f"{prefix}_split_map.json"
    path = os.path.join(outdir, fn)
    with open(path) as f:
        return json.load(f)


def load_anndata_splits(
    outdir: str,
    prefix: str = "dataset",
    *,
    backed: Optional[Union[bool, str]] = None,
) -> Dict[str, ad.AnnData]:
    """
    Load {prefix}_{train|val|test}.h5ad from outdir.

    backed:
      - None: normal in-memory load
      - "r":  backed read-only (useful for huge files)
      - True: alias for "r"
    """
    paths = {
        "train": os.path.join(outdir, f"{prefix}_train.h5ad"),
        "val":   os.path.join(outdir, f"{prefix}_val.h5ad"),
        "test":  os.path.join(outdir, f"{prefix}_test.h5ad"),
    }
    missing = [k for k, p in paths.items() if not os.path.exists(p)]
    if missing:
        raise FileNotFoundError(f"Missing split files for prefix={prefix!r}: {missing}")

    if backed is True:
        backed = "r"

    return {k: ad.read_h5ad(p, backed=backed) for k, p in paths.items()}


def subset_anndata_from_split_map(
    adata: ad.AnnData,
    split_map: Dict[str, Any],
    *,
    copy: bool = False,
    require_all: bool = True,
) -> Dict[str, ad.AnnData]:
    """
    Recreate splits from a loaded split_map JSON (expects obs_names lists in split_map['train'/'val'/'test']).
    """
    keys = ("train", "val", "test")
    if require_all:
        for k in keys:
            if k not in split_map:
                raise KeyError(f"split_map missing key {k!r}")

    def get_names(k: str):
        v = split_map.get(k, [])
        if v is None:
            v = []
        if not isinstance(v, (list, tuple)):
            raise TypeError(f"split_map[{k!r}] must be a list of obs_names.")
        return [str(x) for x in v]

    train_names = get_names("train")
    val_names   = get_names("val")
    test_names  = get_names("test")

    train = adata[train_names]
    val   = adata[val_names]
    test  = adata[test_names]

    if copy:
        train = train.copy()
        val = val.copy()
        test = test.copy()

    return {"train": train, "val": val, "test": test}


def load_or_recreate_splits(
    outdir: str,
    prefix: str,
    *,
    adata: Optional[ad.AnnData] = None,
    backed: Optional[Union[bool, str]] = None,
    split_map_name: Optional[str] = None,
    copy: bool = False,
) -> Dict[str, ad.AnnData]:
    """
    Convenience:
      - if {prefix}_train/val/test.h5ad exist: load them
      - else if split map exists and adata is provided: recreate splits from adata
    """
    paths = {
        "train": os.path.join(outdir, f"{prefix}_train.h5ad"),
        "val":   os.path.join(outdir, f"{prefix}_val.h5ad"),
        "test":  os.path.join(outdir, f"{prefix}_test.h5ad"),
    }
    if all(os.path.exists(p) for p in paths.values()):
        return load_anndata_splits(outdir, prefix=prefix, backed=backed)

    # fallback to split map + adata
    sm = load_split_map(outdir, prefix=prefix, split_map_name=split_map_name)
    if adata is None:
        raise ValueError(
            f"Split .h5ad files not found for prefix={prefix!r}. "
            "Provide `adata=...` to recreate from split_map JSON."
        )
    return subset_anndata_from_split_map(adata, sm, copy=copy)


# =============================================================================
# Latent writer
# =============================================================================

def _select_X(adata_obj: ad.AnnData, layer: Optional[str], X_key: str):
    if X_key != "X":
        if X_key not in adata_obj.obsm:
            raise KeyError(f"X_key={X_key!r} not found in adata.obsm. Keys={list(adata_obj.obsm.keys())}")
        return adata_obj.obsm[X_key]
    if layer is not None:
        if layer not in adata_obj.layers:
            raise KeyError(f"layer={layer!r} not found in adata.layers. Keys={list(adata_obj.layers.keys())}")
        return adata_obj.layers[layer]
    return adata_obj.X


@torch.no_grad()
def write_univi_latent(
    model,
    adata_dict: Dict[str, ad.AnnData],
    *,
    obsm_key: str = "X_univi",
    batch_size: int = 512,
    device: Optional[Union[str, torch.device]] = None,
    use_mean: bool = False,
    epoch: int = 0,
    y: Optional[Union[np.ndarray, torch.Tensor, Dict[str, Union[np.ndarray, torch.Tensor]]]] = None,
    layer: Union[None, str, Mapping[str, Optional[str]]] = None,
    X_key: Union[str, Mapping[str, str]] = "X",
    require_paired: bool = True,
) -> np.ndarray:
    model.eval()

    names = list(adata_dict.keys())
    if len(names) == 0:
        raise ValueError("adata_dict is empty.")

    n = int(adata_dict[names[0]].n_obs)

    if require_paired:
        ref = adata_dict[names[0]].obs_names.values
        for nm in names[1:]:
            if adata_dict[nm].n_obs != n:
                raise ValueError(f"n_obs mismatch: {nm} has {adata_dict[nm].n_obs} vs {n}")
            if not np.array_equal(adata_dict[nm].obs_names.values, ref):
                raise ValueError(f"obs_names mismatch between {names[0]} and {nm}")

    if device is None:
        try:
            device = next(model.parameters()).device
        except StopIteration:
            device = "cpu"

    layer_by_mod = dict(layer) if isinstance(layer, dict) else {nm: layer for nm in names}
    xkey_by_mod = dict(X_key) if isinstance(X_key, dict) else {nm: X_key for nm in names}

    y_t = None
    if y is not None:
        if isinstance(y, dict):
            y_t = {str(k): (v if torch.is_tensor(v) else torch.as_tensor(v)).long() for k, v in y.items()}
        else:
            y_t = (y if torch.is_tensor(y) else torch.as_tensor(y)).long()

    zs = []
    bs = int(batch_size)
    if bs <= 0:
        raise ValueError("batch_size must be > 0")

    for start in range(0, n, bs):
        end = min(n, start + bs)
        x_dict = {}
        for nm in names:
            adata_obj = adata_dict[nm]
            X = _select_X(adata_obj, layer_by_mod.get(nm, None), xkey_by_mod.get(nm, "X"))
            xb = X[start:end]
            if sp.issparse(xb):
                xb = xb.toarray()
            xb = np.asarray(xb)
            x_dict[nm] = torch.as_tensor(xb, dtype=torch.float32, device=device)

        yb = None
        if isinstance(y_t, dict):
            yb = {k: v[start:end].to(device) for k, v in y_t.items()}
        elif torch.is_tensor(y_t):
            yb = y_t[start:end].to(device)

        if hasattr(model, "encode_fused"):
            mu_z, logvar_z, z = model.encode_fused(x_dict, epoch=int(epoch), y=yb, use_mean=bool(use_mean))
            z_use = z
        else:
            out = model(x_dict)
            z_use = out["mu_z"] if (use_mean and ("mu_z" in out)) else out["z"]

        zs.append(z_use.detach().cpu().numpy())

    Z = np.vstack(zs)

    for nm in names:
        adata_dict[nm].obsm[obsm_key] = Z

    return Z
```
