## Supplemental Code for "Unifying multimodal single-cell data with a mixture-of-experts *β*-variational autoencoder framework": logging.py

```
# univi/utils/logging.py

from __future__ import annotations
import logging
from typing import Optional


def get_logger(name: str = "univi", level: int = logging.INFO) -> logging.Logger:
    logger = logging.getLogger(name)
    if not logger.handlers:
        handler = logging.StreamHandler()
        fmt = "[%(asctime)s] [%(name)s] [%(levelname)s] %(message)s"
        handler.setFormatter(logging.Formatter(fmt))
        logger.addHandler(handler)
    logger.setLevel(level)
    return logger
```
