## Supplemental Code for "Unifying multimodal single-cell data with a mixture-of-experts *β*-variational autoencoder framework": seed.py

```
# univi/utils/seed.py

from __future__ import annotations
import random
import numpy as np
import torch


def set_seed(seed: int = 0, deterministic: bool = False):
    random.seed(seed)
    np.random.seed(seed)
    torch.manual_seed(seed)
    if torch.cuda.is_available():
        torch.cuda.manual_seed_all(seed)

    if deterministic:
        torch.backends.cudnn.deterministic = True
        torch.backends.cudnn.benchmark = False
```
