## Supplemental Code for "Unifying multimodal single-cell data with a mixture-of-experts *β*-variational autoencoder framework": stats.py

```
# univi/utils/stats.py

from __future__ import annotations
from typing import Dict
import numpy as np


def mean_dict(d: Dict[str, float]) -> float:
    """
    Simple helper: mean of dictionary values.
    """
    if not d:
        return float("nan")
    return float(np.mean(list(d.values())))


def zscore(x: np.ndarray, axis: int = 0, eps: float = 1e-8) -> np.ndarray:
    """
    Z-score normalization along given axis.
    """
    mu = x.mean(axis=axis, keepdims=True)
    std = x.std(axis=axis, keepdims=True) + eps
    return (x - mu) / std
```
