## Supplemental Code for "Unifying multimodal single-cell data with a mixture-of-experts *β*-variational autoencoder framework": torch_utils.py

```
# univi/utils/torch_utils.py

from __future__ import annotations
from typing import Dict, Any
import numpy as np
import torch


def get_device(prefer: str = "cuda") -> torch.device:
    if prefer == "cuda" and torch.cuda.is_available():
        return torch.device("cuda")
    return torch.device("cpu")


def to_device(batch: Dict[str, torch.Tensor], device: torch.device) -> Dict[str, torch.Tensor]:
    return {k: (v.to(device) if v is not None else None)
            for k, v in batch.items()}


def to_numpy(x: Any):
    if isinstance(x, torch.Tensor):
        return x.detach().cpu().numpy()
    return np.asarray(x)
```
