## Supplementary figures and images for "Unifying multimodal single-cell data with a mixture-of-experts *β*-variational autoencoder framework"

### univi_overview_dark.png

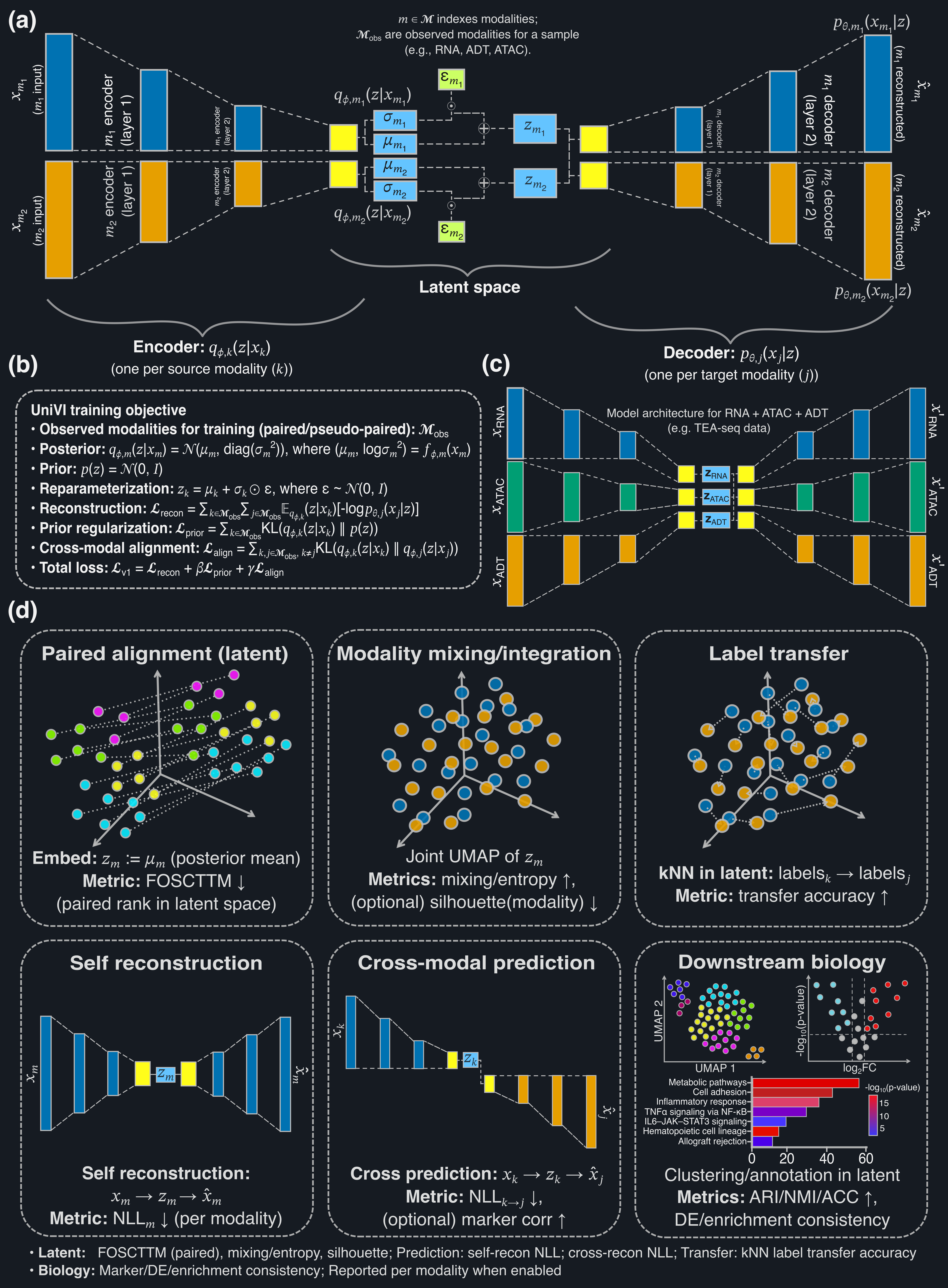

### univi_overview_light.png

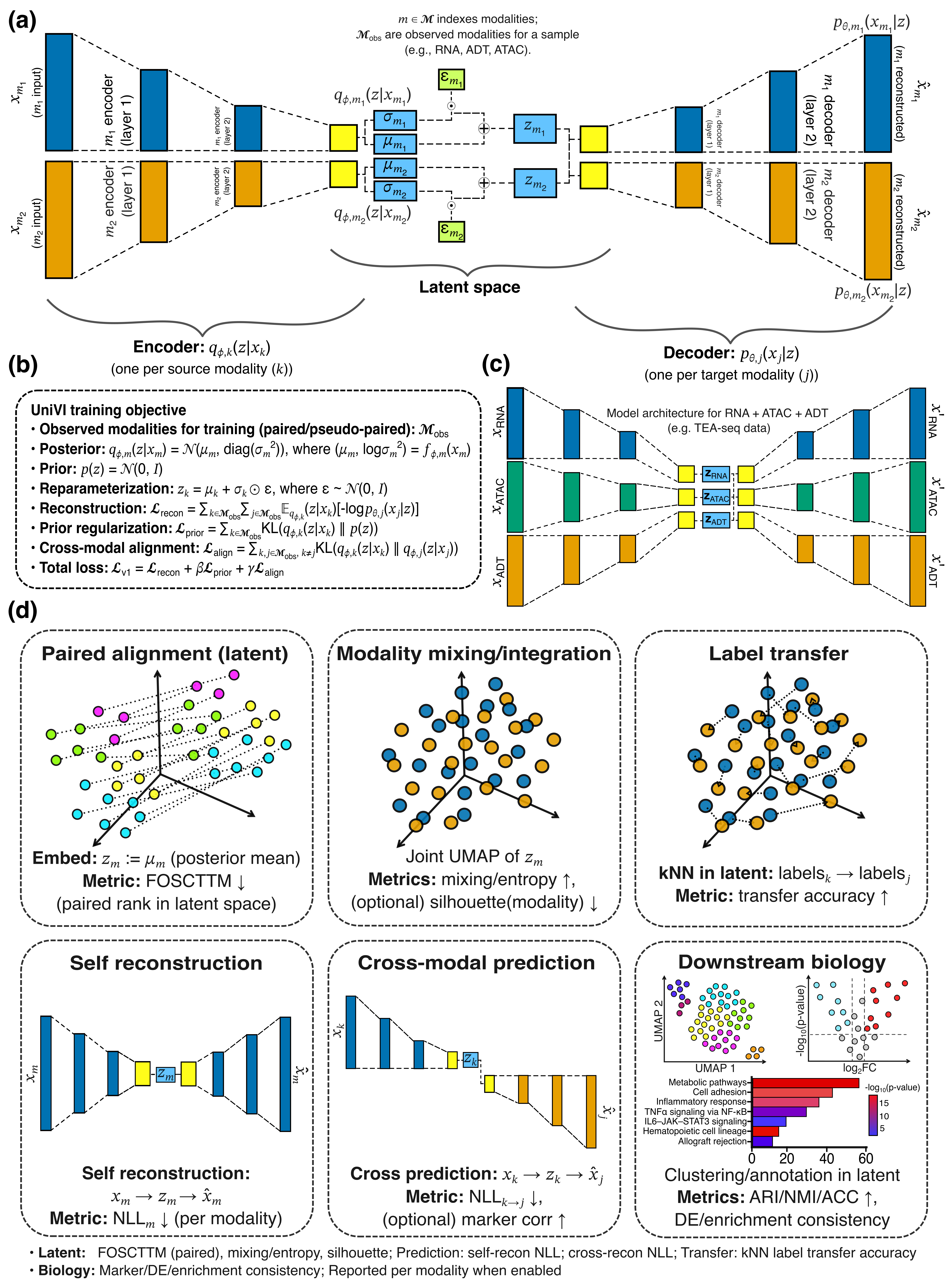
